# Phylogenetic Investigation of the 100 kDa Hexokinase Enzyme Family with the Topiary Ancestral Sequence Reconstruction Pipeline

**DOI:** 10.64898/2026.01.30.702642

**Authors:** Carolin Freye, A. Carl Whittington, Brian G. Miller

## Abstract

The 100 kDa hexokinase (HK) enzyme family represents an attractive model to investigate the molecular origins of allosteric regulation in multidomain enzymes. Extant HK homologs are subject to various allosteric phenomena, including activation and inhibition by both homotropic and heterotropic ligands. Here, we report the results of a phylogenetic investigation of this enzyme family using the recently developed *Topiary* ancestral sequence reconstruction pipeline. The results agree with prior studies that used a smaller number of sequences from individual HK domains and suggest that modern HK3 isozymes diverged first from a 100 kDa ancestor, followed by gene duplication and divergence of the HK2 isozymes. A subsequent gene duplication event led to divergence of HK1 and the hexokinase domain containing protein 1 (HKDC1). To probe the ability of *Topiary* to yield functional, allosterically regulated ancestral enzymes, we resurrected and biochemically characterized two HKs from early vertebrate evolution, Anc1 and Anc2. Both enzymes were functionally similar to extant HK1, and possessed a low activity, regulatory N-terminal domain that governs allosteric regulation of the C-terminal active site by two heterotropic effectors, glucose 6-phosphate and inorganic phosphate. Neither ancestor was subject to homotropic regulation by substrate glucose, a characteristic observed in several extant HK3 family members. Our phylogenetic analysis provides a foundation for investigating the evolution of allostery in this enzyme family. It also demonstrates the need to sequence and biochemically characterize additional full-length HKs, especially those from jawless vertebrates, to enable more robust inferences of ancestral regulatory traits.

## INTRODUCTION

Allosteric regulation of protein function is a common mechanism for controlling biological processes (Nussinov and Tsai, 2013). Evolutionary approaches offer the potential to reveal both the mechanistic basis of regulation and the historical molecular events that enabled an allosteric pathway to first emerge (Hochberg and Thornton, 2017). The vertebrate hexokinase (HK) enzyme family presents an excellent model system to investigate fundamental aspects of allosteric enzyme regulation. HKs are well represented across all five vertebrate classes, are subject to allosteric inhibition and activation by both homotropic and heterotropic effectors and share a common biochemical function and structural scaffold (Cárdenas et al., 1998; Wilson, 2003). Notably, however, the extent to which these homologs share a common allosteric regulatory mechanism remains unclear.

HKs catalyze the first step of glycolytic metabolism, the ATP-dependent phosphorylation of glucose to glucose 6-phosphate (G6P) (Wilson, 2003). The four 100 kDa HKs are comprised of two 50 kDa homologous N- and C-terminal domains linked by an α-helix (Aleshin et al., 1998; Irwin and Tan, 2008; Wilson, 2003). Despite this common structural arrangement, each of the four homologs displays distinct functional and regulatory features (Figure 1) (Cárdenas et al., 1998; Piwko et al., 2025; Wilson, 2003). HK1 has an inactive, regulatory N-terminal domain and a catalytically active C-terminal domain (Cárdenas et al., 1998; White and Wilson, 1989; Wilson, 2003). Human HK1 has a low glucose *K*_m_ value (60 μM) and is potently inhibited by G6P, with a *K*_i_ value of ∼25 μM (Magnani et al., 1988). G6P binding to the N-terminal domain stabilizes the C-terminal domain in an ATP-incompetent state, impeding catalysis (Aleshin et al., 2000). That inhibition is relieved by binding of P_i_, a function exclusive to HK1, to an overlapping pocket that allosterically converts the C-terminal active site to an ATP-competent state (Aleshin et al., 2000). These interdomain allosteric signals are relayed > 50 Å via a pathway involving coulombic interactions at an interdomain interface (Aleshin et al., 2000; Fang et al., 1998).

**Figure 1.**
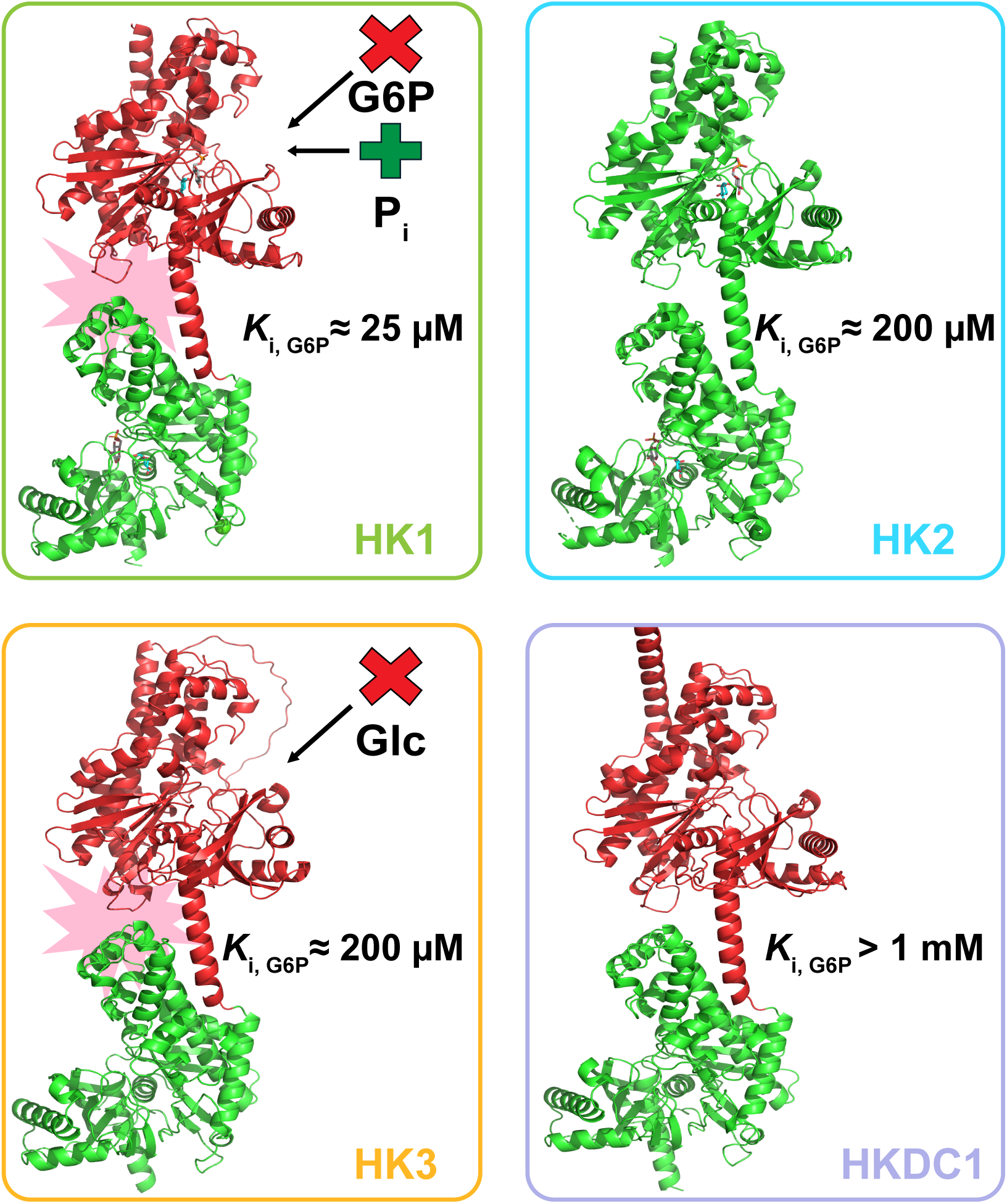
Vertebrate hexokinases share a common structural scaffold and display different functional and regulatory features (Wilson, 2003). The N-terminal domains of HK1 (top left; PDB: 1HKB) (Aleshin et al., 1998), HK3 (bottom left; AF-P52790), and HKDC1 (bottom right; AF-Q2TB90) lack catalytic activity (red) while the C-domains are catalytically active (green) (Jumper et al., 2021; Varadi et al., 2024, 2022). Both domains of HK2 (top right; PDB: 2NZT) are catalytically active (green) (Nawaz et al., 2018). Glucose 6-phosphate (G6P) potently inhibits HK1-HK3 (Ardehali et al., 1996; Magnani et al., 1988; Palma et al., 2002) while HKDC1 displays much lower sensitivity to G6P inhibition (Piwko et al., 2025). In HK1, G6P binding to the N-terminal domain allosterically inhibits C-terminal domain activity which is relieved by inorganic phosphate (P_i_) (Aleshin et al., 2000). In HK3, glucose (Glc) binding to the N-domain allosterically inhibits C-terminal domain activity (Freye and Miller, 2025). Allosteric regulation of HK1 and HK3 by their respective heterotropic and homotropic effectors is relayed, in part, through a similar interdomain interface (pink) (Freye and Miller, 2025).

HK2 is unique among the 100 kDa isozymes as both domains are catalytically active (Ahn et al., 2009; Ardehali et al., 1999, 1996; Ferreira et al., 2021; Nawaz et al., 2018; Tsai and Wilson, 1996). Crystal structures of human HK2 show glucose bound at the active site of each domain (Nawaz et al., 2018), suggesting catalytic independence. Specific activity of the full-length human enzyme matches the sum of the activities of the isolated domains (Ahn et al., 2009; Ardehali et al., 1996). Interestingly, the specific activity of the full-length, rat enzyme is <u>∼6</u>0% lower than the activity of the isolated N-domain (Ardehali et al., 1996). In addition, rat HK2 reportedly displays a 1:1 stoichiometry with *N*-bromoacetylglucosamine, an irreversible active-site targeted inhibitor (Connolly and Trayer, 1979). Together these observations suggest the possibility of half-of-sites activity in some HK2s. Human HK2 has a relatively low *K*_m_ value for glucose (∼300 μM) and is potently inhibited by G6P, with a *K*_i_ value of ∼200 μM (Ardehali et al., 1996). The *K*_i_ value for G6P inhibition of the isolated N-terminal domain resembles the full-length enzyme, whereas the *K*_i_ value for the isolated C-domain is significantly higher (Ahn et al., 2009; Ardehali et al., 1999, 1996).

The functional organization of HK3 resembles HK1. The N-terminal domain serves a regulatory function while the C-terminal domain is catalytically active (Freye and Miller, 2025; Palma et al., 1996; Tsai and Wilson, 1997). Human HK3 displays a low *K*_m_ value for glucose (∼30 μM), but is inhibited by G6P with a *K*_i_ value (∼200 μM) that is 10-fold higher than that observed with HK1 (Palma et al., 2002). The extent to which G6P inhibition is direct or allosteric in nature is unknown. A unique feature of HK3 regulation is its susceptibility to substrate inhibition by high, but physiologically relevant, glucose concentrations (Cárdenas et al., 1998; Wilson, 2003). Biochemical studies support an allosteric model for the homotropic regulation of HK3, wherein glucose binding to the N-terminal domain impedes catalysis at the C-domain (Freye and Miller, 2025; Palma et al., 1996; Tsai and Wilson, 1997). This homotropic signal involves similar coulombic interdomain interactions as in HK1, suggesting that HK1 and HK3 may share a common mechanism for interdomain communication (Freye and Miller, 2025).

HKDC1 likely diverged from HK1 upon a tandem gene duplication event (Irwin and Tan, 2008) and displays a similar functional organization as HK1 (Piwko et al., 2025). In contrast to HK1, HKDC1 displays lower overall hexokinase activity, a higher glucose *K*_m_ value (∼0.5 mM), and is largely insensitive to physiological concentrations of G6P (Piwko et al., 2025). Despite these functional differences between HKDC1 and HK1, a sequence comparison of the two human isozymes demonstrates that nearly all residues that contact G6P in HK1 (Aleshin et al., 2000, 1998; Rosano et al., 1999) are conserved in HKDC1, suggesting divergent regulatory properties despite a conserved structural arrangement.

The current understanding of HK evolution posits that an ancestral 100 kDa HK likely evolved via duplication and subsequent fusion of a gene encoding an ancestral 50 kDa hexokinase during early vertebrate evolution (Bork et al., 1993; Cárdenas et al., 1998; González-Alvarez et al., 2009; Griffin et al., 1991; Irwin and Tan, 2008; Li et al., 2014; Mochizuki, 1981). Successive gene duplication events during the evolution of jawless and jawed vertebrates, led to the divergence of the present-day HKs (Bork et al., 1993; Cárdenas et al., 1998; González-Alvarez et al., 2009; Irwin and Tan, 2008; Li et al., 2014; Mochizuki, 1981). Importantly, all past phylogenetic investigations of HKs divided the protein sequence into separate N- and C-terminal domains prior to analysis. Several investigators speculated about the traits of early 100 kDa HKs (Ardehali et al., 1996; Cárdenas et al., 1998; Griffin et al., 1991; Irwin and Tan, 2008; Li et al., 2014; Tsai and Wilson, 1996; White and Wilson, 1989), however, ancestors have not been resurrected in the lab to probe these hypotheses. As a first step in that direction, we utilized the recently developed python framework *Topiary,* a pipeline specifically developed for phylogenetic “non-experts” (Orlandi et al., 2023), to perform the first phylogenetic analysis of this enzyme family using full-length sequences. We successfully resurrected the two oldest ancestors from our resulting HK *Topiary* tree and performed steady-state kinetic and inhibition assays to determine the functional and regulatory features of each enzyme. Both ancestors were found to possess a catalytically active C-terminal domain and a low activity, regulatory N-terminal domain to which G6P can bind and allosterically inhibit catalysis at the C-terminal domain. Our results provide a starting point for investigating the evolutionary history of the diverse functional and regulatory strategies present in the vertebrate HK enzyme family.

## RESULTS & DISCUSSION

### Phylogenetic analysis of the 100 kDa HK enzyme family by *Topiary* agrees with literature precedent

We used the *Topiary* python framework (Orlandi et al., 2023) to perform a phylogenetic analysis of the HK enzyme family. *Topiary* combines the individual steps of ancestral sequence reconstruction (ASR) into one pipeline and was developed to expand accessibility of ASR to the broader scientific community by facilitating handling of large datasets, eliminating data manipulation for the transfer between various software, and reducing the need for informed decision making by phylogenetic experts (Orlandi et al., 2023). Past phylogenetic studies involving the HK enzyme family divided the 100 kDa HK sequences into separate N- and C-terminal domains prior to analysis (Bork et al., 1993; Cárdenas et al., 1998; González-Alvarez et al., 2009; Griffin et al., 1991; Irwin and Tan, 2008; Li et al., 2014). Allosteric regulation in 100 kDa HKs requires communication between both domains. Thus, we chose to perform analyses on the full-length HK sequences.

From our eight sequence seed-dataset, ∼5000 HK sequences were automatically collected (Table SI). A combination of automated and manual curation (Materials and Methods) reduced the number of sequences used for the final multiple sequence alignment (MSA) to 231 (Supplementary phylogenetic material S1). *Topiary* selected an HK1-like protein from the lamprey *Lethenteron reissneri* (accession number: XP_061404494.1), a jawless vertebrate, as the outgroup. Analysis of the *L. reissneri* genome yielded three *hk*-like genes, one annotated as *hk1*-like and two adjacent genes that are annotated as *hk2*-like. The synteny for each *hk*-like gene differs from the respective human isozymes likely due to taxon-specific traits, e.g. genome rearrangements and losses that occur during lamprey embryogenesis (Smith et al., 2009).

Using the final MSA and the lamprey HK1-like protein as an outgroup, *Topiary* generated a reconciled maximum likelihood (ML) phylogenetic tree for the 100 kDa HK enzyme family and inferred full-length ancestral HK sequences at individual nodes (Figure 2, Supplementary phylogenetic material S2-3). The phylogenetic tree supports a model in which HK3s diverged first from the common ancestor of all 100 kDa jawed vertebrate HKs (Anc1) following a gene duplication event. A subsequent gene duplication event led to the divergence of HK2. Finally, following a third gene duplication event, HK1 and HKDC1 diverged from one another. Notably, HK3s from bony fish are underrepresented in our phylogenetic tree due to lineage specific genetic events including duplications and potential exon loss (Yates et al., 2026), which can complicate phylogenetic inference by obscuring orthology/paralogy and reducing the number of informative sites in the MSA, respectively.

**Figure 2.**
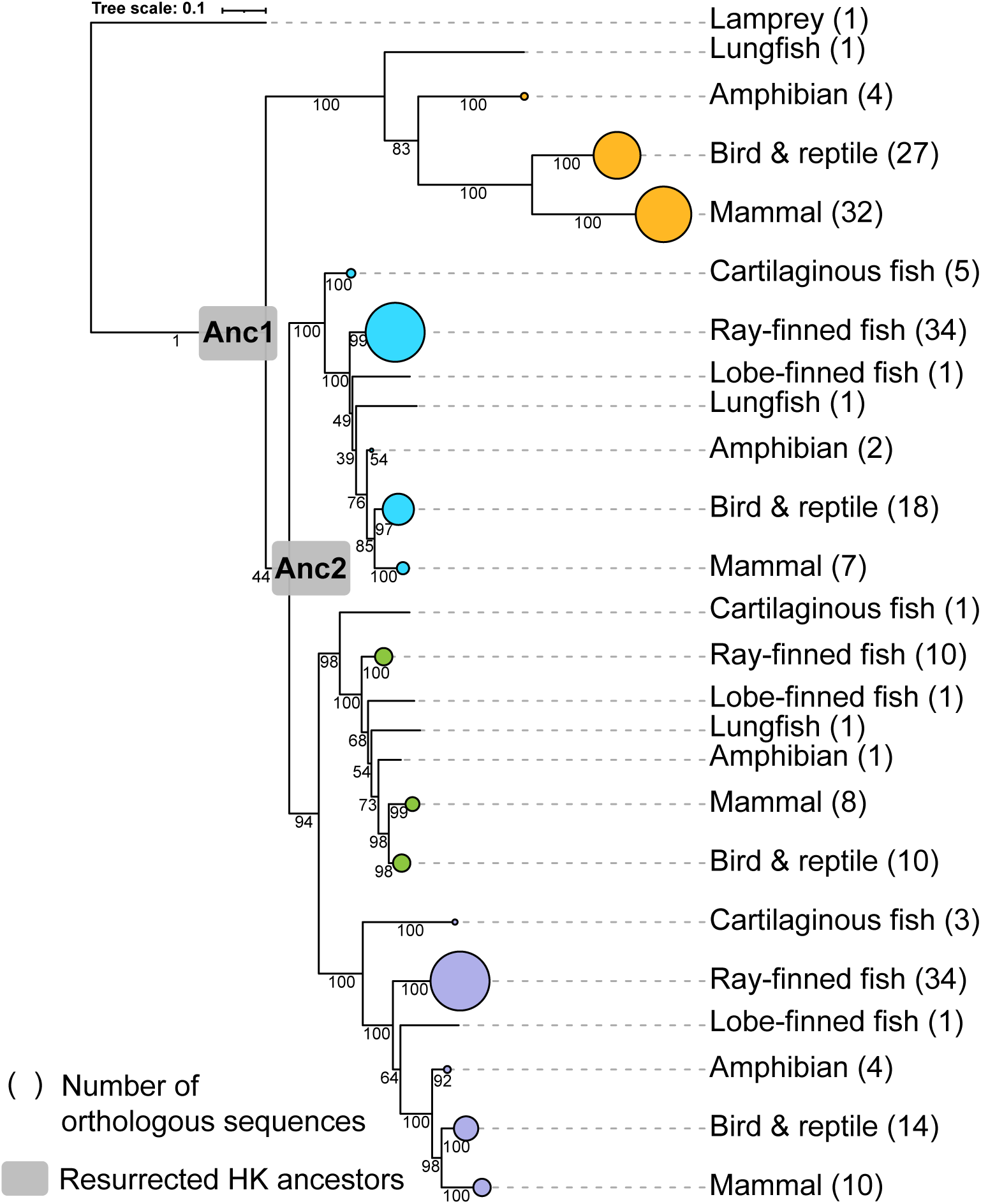
Phylogenetic tree of the hexokinase enzyme family estimated with *Topiary*. The sizes of the circles located at the tips of the tree correspond to the number of extant HK sequences used for our phylogenetic analysis from each class. Each color represents the distinct HK paralogs. HK1 orthologs are marked in green, HK2 orthologs are blue, HK3 orthologs are orange and HKDC1 orthologs are purple. Branch supports are indicated next to each node on a scale from 0 to 100.

*Topiary* utilizes the non-parametric bootstrapping method of RAxML-NG to determine branch support values as a measure of confidence in the tree topology (Kozlov et al., 2019; Orlandi et al., 2023). The branch support values for our ancestors Anc1 (1/100) and Anc2 (44/100) are low (Figure 2). *Topiary* selected a HK1-like sequence from lamprey as the outgroup, which is expected to group with other HK1 orthologs. This mismatch may be one source of low branch support for Anc1. Low branch support values were also observed by Li and coworkers during their most recent phylogenetic analysis of the HK domains (Li et al., 2014). Additionally, poor branch support may stem from constrained sequence sampling. We postulate that a lack of extant full-length sequences from invertebrates and jawless vertebrates, and the back-to-back duplication of the ancestral *hk* gene during early vertebrate evolution, are detrimental for the resolution at early nodes. Despite the low branch supports of the two ancestral nodes, and the uncertainty of the outgroup, the topology of our *Topiary* tree generally agrees with other sources including the *Ensembl* HK tree (Yates et al., 2026) and trees previously inferred using small datasets with HKs split into isolated N- and C-terminal domains (Bork et al., 1993; Cárdenas et al., 1998; González-Alvarez et al., 2009; Irwin and Tan, 2008; Li et al., 2014). This observation provides additional support that the tree presented in Figure 2 provides a useful model of HK evolution.

### Anc1 and Anc2 are catalytically competent HKs

The ML sequence consists of the amino acid with the highest posterior probability (PP) at each position of the alignment with gaps reconstructed via maximum parsimony within *Topiary* (Ishikawa et al., 2019; Orlandi et al., 2023). We resurrected the two most ancient ML ancestors from our *Topiary* reconstruction, Anc1 and Anc2 (Figure 2), and found that Anc1 and Anc2 display *k*_cat_ values of 35 ± 10 s^−1^ and 56 ± 12 s^−1^, and *K*_m_ values for glucose of 75 ± 8 μM and 68 ± 5 μM, respectively (Table I, Figures 3 and S1-2). The *K*_m_ values for ATP of Anc1 and Anc2 are 1.4 ± 0.1 mM and 1.8 ± 0.1 mM, respectively (Figures S3-4). Overall, these values are comparable to the characteristics of extant HKs (Cárdenas et al., 1998; Ureta, 1982; Wilson, 1995, 2003).

**Figure 3.**
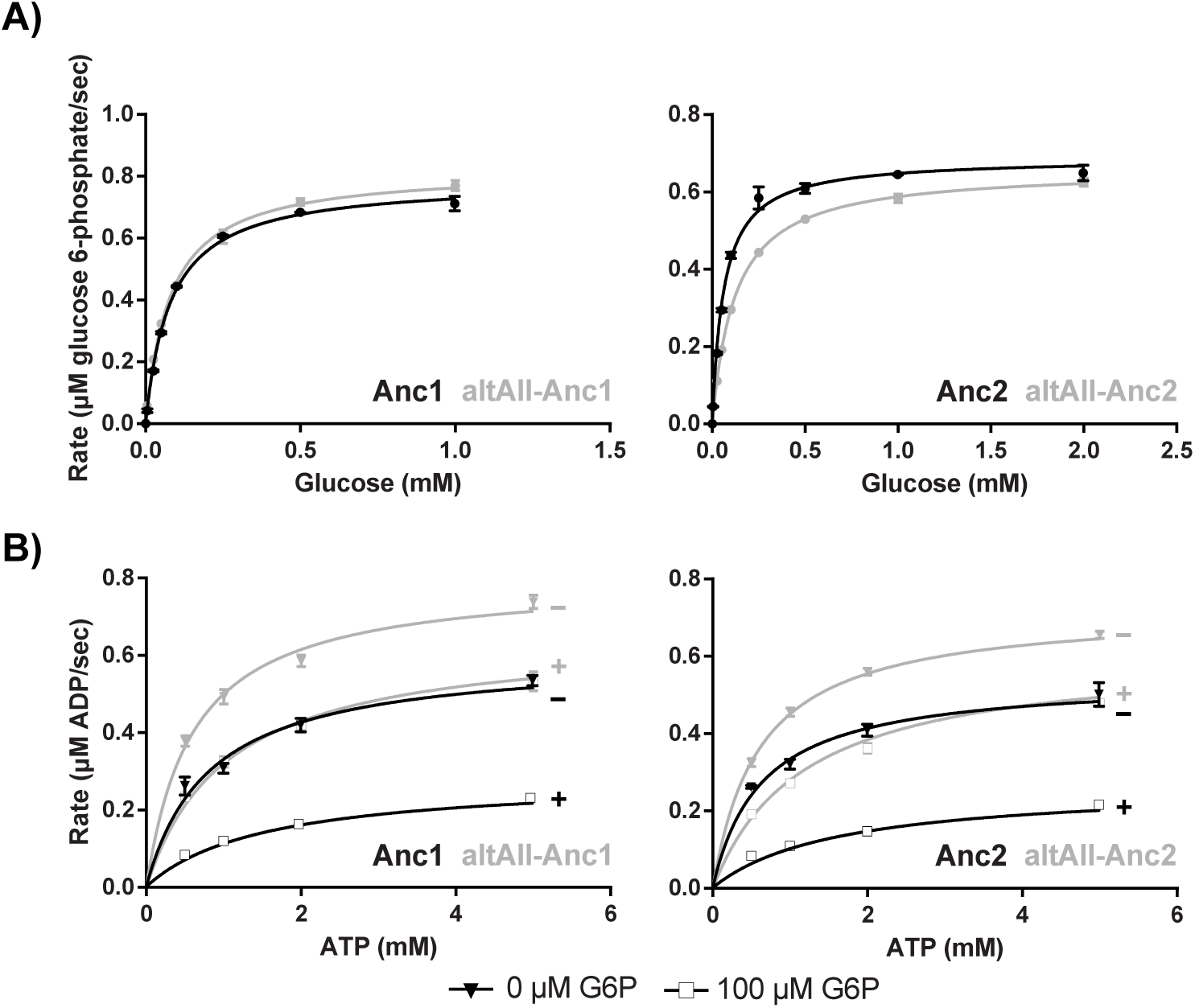
Comparison of steady-state kinetics for ML and altAll versions of resurrected hexokinase ancestors. A) Enzymatic activities of ML and altAll Anc1 (left), and ML and altAll Anc2 (right), at varying glucose concentrations and saturating ATP concentrations. B) Activities of ML and altAll Anc1 (left), and ML and altAll Anc2 (right), in the absence (−) and presence of G6P (+, 100 μM) at saturating glucose concentrations and sub-saturating ATP concentrations.

**Table I.** Apparent steady-state kinetic parameters of resurrected ML and altAll HK ancestors and variants thereof.

| | $K_m$, glucose ($\mu\text{M}$) | $K_m$, ATP (mM) | $K_i$, glucose 6-phosphate<br>($\mu\text{M}$) | $\alpha^*K_i$, glucose 6-phosphate<br>($\mu\text{M}$) | $k_{\text{cat}}$ ($\text{s}^{-1}$) |
| --- | --- | --- | --- | --- | --- |
| Anc1 | $75 \pm 8$ | $1.4 \pm 0.1$ | $49 \pm 21^a$ | $68 \pm 30^a$ | $35 \pm 10$ |
| Anc1C | $80 \pm 1$ | $2.0 \pm 0.1$ | $85 \pm 15^a$ | $116 \pm 51^a$ | $27 \pm 3$ |
| Anc1N | nd | nd | nd | nd | $0.10 \pm 0.01$ |
| Anc1 D643A | nd | nd | nd | nd | $0.36 \pm 0.02$ |
| altAll-Anc1 | $80 \pm 1$ | $1.0 \pm 0.1$ | $78 \pm 16^a$ | $508 \pm 91^a$ | $74 \pm 13$ |
| Anc2 | $68 \pm 5$ | $1.8 \pm 0.1$ | $24 \pm 1^a$ | $104 \pm 12^a$ | $56 \pm 12$ |
| Anc2C | $72 \pm 4$ | $2.8 \pm 0.1$ | $55 \pm 13^a$ | $175 \pm 56^a$ | $51 \pm 5$ |
| Anc2N | nd | nd | nd | nd | $0.08 \pm 0.01$ |
| altAll-Anc2 | $124 \pm 1$ | $1.8 \pm 0.4$ | $96 \pm 28^a$ | $542 \pm 4^a$ | $52 \pm 7$ |
| Anc1 D518A | $17 \pm 3$ | $8 \pm 1$ | $34 \pm 7^b$ | - | $1.1 \pm 0.1$ |
| Anc2 D518A | $10 \pm 2$ | $6.7 \pm 0.4$ | $74 \pm 16^b$ | - | $0.42 \pm 0.02$ |

Kinetic parameters are reported as the average ± standard deviation of two biological replicates with three technical replicates each. The $k_{\text{cat}}$ values of Anc1N, Anc2N, and Anc1 D643A were determined at 1 mM glucose and 15 mM ATP. All other $k_{\text{cat}}$ values were calculated from $V_{\text{max}}$ following data fitting to the Michaelis-Menten equation and are reported as the average ± standard deviation of four $k_{\text{cat}}$ values, two from each biological replicate.
<sup>a</sup> Inhibition constants were determined by fitting data to equation 1.
<sup>b</sup> Inhibition constants were determined by fitting data to equation 2.
nd: not determined

To evaluate the phenotypic robustness of our ancestors, we also characterized the altAll versions (Table I, Figures 3A and S5-8) where any position with a PP lower than 0.75 is replaced with the second most probable amino acid from our reconstruction. The ML sequence of Anc1 has an average PP across all residues of 0.955 and, among the human HK paralogs, Anc1 shares the highest sequence identity with HK2 (84%). The altAll sequence for Anc1 has an average PP of 0.944 and shares 93% sequence identity with the ML sequence. The ML sequence of Anc2 has an average PP of 0.975 and shares a sequence identity of 99% and 85% with Anc1 and human HK2, respectively. The altAll sequence of Anc2 has an average PP of 0.967 and shares a sequence identity of 96% with the ML sequence. Both altAll-Anc1 and altAll-Anc2 were catalytically active and displayed functional characteristics very similar to the ML versions (Table I, Figure 3A), demonstrating that the kinetic parameters are robust to variation in sequence and to model uncertainty, which is generally seen in ASR studies (Eick et al., 2016).

### Functional organization of Anc1 and Anc2

To investigate the functional organization of individual domains of Anc1 and Anc2, we characterized the isolated N- and C-domains of each ML ancestor. The kinetic parameters obtained for the isolated C-domains of Anc1 (Anc1C) and Anc2 (Anc2C) resemble those of the full-length enzymes (Table I, Figures S9-12). In contrast, we found that the isolated N-terminal domains (Anc1N and Anc2N) display only marginal catalytic activity, with apparent *k*_cat_ values of 0.10 ± 0.01 s^−1^ and 0.08 ± 0.01 s^−1^, respectively (Table I). To verify that the lack of activity of the N-terminal domain is not caused by the truncation of the C-terminus, we prepared a full-length variant of Anc1 in which the C-terminal catalytic aspartate (Asp643) was replaced with alanine. The D643A substitution reduced the activity from 35 ± 10 s^−1^ to 0.36 ± 0.02 s^−1^ (Table I). We also collected CD spectra for both the isolated N- and C-terminal domains of Anc1 and found them to be highly comparable (Figures S13-14), consistent with a similar, folded structure in both domains. Together, these results demonstrate that Anc1 and Anc2 share a similar functional organization with extant HK1, HK3, and HKDC1.

Among the human HK isozymes, HK2 is unique in that its N-terminal domain displays robust catalytic activity (Ahn et al., 2009; Ardehali et al., 1999, 1996; Ferreira et al., 2021; Nawaz et al., 2018; Tsai and Wilson, 1996). Of the two ancestors, Anc2 shares highest sequence identity with human HK2; they differ at 136 positions, 83 of which are located at the N-domain while 53 are at the C-domain. We analyzed our MSA and identified only one residue that is highly conserved in extant HK2s (Arg470 in the interdomain α-helix) but differs in Anc2 (Ile456) (Figure 4). We also identified one HK2 N-terminal active site residue (Thr88) that was ambiguously reconstructed in our ASR and corresponds to Ser74 in Anc2. To determine whether Anc2N activity can be enhanced by changes at either of these positions, we characterized the Anc2N I456R and S74T variants. The I456R variant had an unimproved *k*_cat_ value of 0.07 ± 0.01 s^−1^, whereas the S74T variant displayed a 6-fold increase in *k*_cat_ value (0.55 ± 0.01 s^−1^) compared to Anc2N. Together, these data suggests that lack of activity in Anc2N is not solely caused by amino acid differences at these two positions.

**Figure 4.**
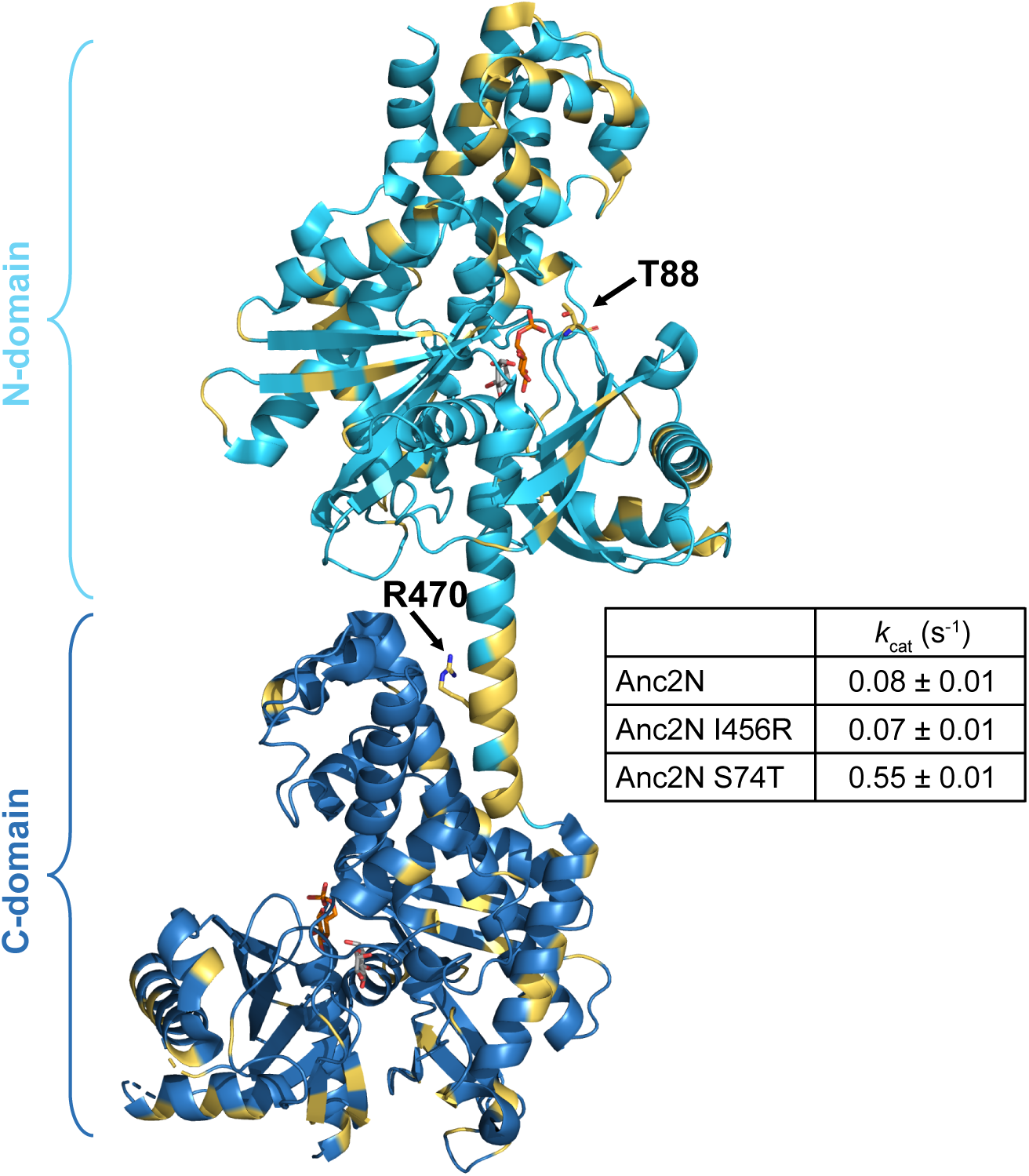
Crystal structure of human HK2 (PDB: 2NZT) bound to glucose (gray) and glucose 6-phosphate (orange) (Nawaz et al., 2018). Anc2 residues that differ from human HK2 are yellow. Arg470 (human HK2 numbering), is an isoleucine in Anc2 (Ile456) and lies at the end of the interdomain α-helix. Thr88 (human HK2 numbering) lies in the active site cavity of HK2 and is a serine in Anc2 (Ser74). Mutagenesis of neither residue in Anc2 to the respective amino acid found in extant human HK2 introduces robust catalytic activity.

### Anc1 and Anc2 display heterotropic allosteric regulation

HK1, HK2, and HK3 are subject to potent heterotropic inhibition by G6P (Wilson, 2003). We performed kinetic assays of Anc1 and Anc2 in the presence of G6P and observed potent inhibition of both ancestors (Table I, Figures 3B and S15-16). Both altAll-Anc1 and altAll-Anc2 displayed similar levels of inhibition (Table I, Figure 3B and S17-18), suggesting phenotypic robustness. To characterize the pattern of inhibition, we globally fit our kinetic data to several models and found that both mixed and pure non-competitive inhibition models provided reasonable fits (Table SII). Both models are consistent with the presence of multiple binding sites for G6P.

Human HK1 has G6P binding sites in both the N-terminal and C-terminal domains (Aleshin et al., 2000, 1998; Rosano et al., 1999). To determine which binding site is responsible for the observed inhibition of human HK1, Liu *et al*. demonstrated that site-directed mutagenesis of G6P binding residues at the C-terminus impairs inhibition of the isolated domain (Liu et al., 1999) while potent inhibition is retained in the full-length variant with a G6P *K*_i_ value unchanged from wild-type (Liu et al., 1999; Zeng et al., 1996). This supports a model in which inhibition is allosteric in nature and originates from the N-terminal site (Liu et al., 1999). Subsequent structural studies supported this allosteric mechanism by revealing conformational changes in the full-length enzyme that link the N-terminal domain G6P binding site to the C-terminal active site (Aleshin et al., 2000).

To test whether the heterotropic G6P inhibition of Anc1 and Anc2 involves an allosteric mechanism that resembles HK1, or if it acts under an orthosteric mechanism, we undertook a similar experimental approach. We prepared and characterized the analogous C-terminal G6P binding site variants in our ancestors (Anc1 D518A and Anc2 D518A, Figures S19-22). Upon fitting the obtained kinetic data to the competitive inhibition equation, we observed that the *K*_i_ value for G6P remained unaltered in both variants relative to the full-length enzymes and the isolated C-domains (Table I and Table SII, Figures S23-26). We also tested whether P_i_ relieves G6P inhibition, as has been observed in human HK1 with a G6P analog (Fang et al., 1998; Sui and Wilson, 2002; Tsai, 2007). We found that [P_i_] ≤ 10 mM enhance the activities of Anc1 and Anc2 in the presence of inhibitory G6P concentrations (Figures S27-28). The magnitude of inhibition relief reaches a maximum of ∼10%. By comparison, we observed a larger maximum level of relief of G6P inhibition by phosphate in human HK1 that reached ∼25% (Figures S29). Together, these data are consistent with a model in which both Anc1 and Anc2 are subject to heterotropic allosteric regulation involving the N-terminal domain, like extant human HK1.

Several extant amphibian and mammalian HK3s display homotropic allosteric regulation, whereby glucose binds at the catalytically inactive N-terminal domain and inhibits catalysis at the C-terminal domain (Balinsky and Fromm, 1978; Cayanis and Balinsky, 1975; Freye and Miller, 2025; Heumann et al., 1974; Magnani, 1983; Palma et al., 2002, 1996; Radojković and Ureta T, 1987; Stocchi et al., 1983; Tsai and Wilson, 1997; Ureta, 1976). We did not observe inhibition of Anc1 and Anc2 activity at glucose concentrations up to 100 mM in our assays (Table SIII). The putative N-domain glucose binding residues of human HK3, inferred from X-ray structural data of other HK paralogs (Aleshin et al., 2000, 1998, 1998; Nawaz et al., 2018; Rosano et al., 1999) due to the lack of structural data for full-length human HK3, appear to be fully conserved in Anc1. We analyzed residues in our MSA that are fully conserved in all extant HK3 sequences but not present in Anc1. The only residue to which this criterion applied is position 83, which is a valine in Anc1 and a leucine in the extant HK3s. The V83L Anc1 variant displayed similar kinetic parameters to Anc1 with a *k*_cat_ value of 55 ± 2 s^−1^ and a *K*_m_ value for ATP and glucose of 2.3 ± 0.1 mM and 96 ± 1 μM, respectively (Figures S30-31). However, it lacked inhibition by glucose concentrations up to 100 mM (Table SIII). Together, these results demonstrate that introducing a leucine at position 83 is not sufficient to install substrate inhibition in Anc1.

### Evaluation of *Topiary* as an ancestral sequence reconstruction tool for non-experts

Biochemists generally receive little to no exposure to molecular evolution while pursuing their degrees. *Topiary* makes ASR more accessible to biochemists by combining individual steps of the process into one pipeline, requiring little user input, yet having detailed explanations of each step in the pipeline (Orlandi et al., 2023). *Topiary* collects large datasets from only a handful of input sequences, curates and transfers these datasets between several ASR software, and generates a species-aware reconciled tree (Mctavish et al., 2021; Orlandi et al., 2023). Reconciliation of the gene and species trees may enhance the accuracy of the inferred ancestral sequences by rooting the phylogenetic tree and resolving ambiguous relationships within the gene tree (Orlandi et al., 2023). Tree reconciliation also assigns speciation, duplication and gene loss events to the tree’s nodes (Orlandi et al., 2023).

Our results demonstrate the usefulness of *Topiary*. A non-expert was able to successfully execute an ASR analysis on the HK family of enzymes. We did, however, encounter several unexpected challenges, some of which are addressed in methods and materials. *Topiary* requires cross compatibility between specific software versions, and we needed to downgrade to older versions at the time of installation. We also experienced issues connecting the graphical application of RAxML-NG to the display. *Topiary* uses the proteomes of each species listed in the seed-dataset, which serves as the input file for the pipeline, to collect sequences via reciprocal BLAST from NCBI. *Topiary* did not automatically download these proteomes which required user intervention at this stage of the analysis. We had to manually set-up the messenger passing interference (MPI) to parallelize the quantification of branch support, as it was originally not compatible with our institutional high performance computing cluster. This step of the pipeline was also significantly slower than expected, which revealed problems with the --restart flag. We found that the command did not restart the partially completed job from the last saved checkpoint but rather restarted the entire bootstrapping process. Despite these issues, *Topiary* provides a useful tool for non-experts who don’t have formal training in phylogenetics and evolutionary biochemistry. We appreciated that the protocol provides guidance on the sequence selection for the seed-data set and recommendations for MSA edits. The fast collection of large sequence datasets and the determination of the taxonomic scope from which sequences are available is one of *Topiary’s* most significant benefits as it eliminates hours of tedious manual work.

### *Topiary* informs future directions

The present study represents the first attempt to perform a phylogenetic analysis using a large number of full-length 100 kDa HKs protein sequences. Our results, while demonstrating the ability to reconstruct functional, allosteric-regulated ancestors also identified several key areas for improvement. Future studies would benefit from the incorporation of full-length HK3 sequences from cartilaginous and ray-finned fish. We could not identify such full-length sequences in currently available databases, suggesting the need to sequence additional fish genomes. Alternatively, gene truncation (Irwin and Tan, 2008) or loss may have occurred in those lineages, in which case future studies may benefit from the incorporation of more lobe-finned fish and amphibian sequences. To the extent possible, the inclusion of more HK sequences corresponding to the base of our tree, e.g. sequences from jawless vertebrates and potentially non-craniate chordates, would also be useful. Access to such data should improve the resolution at early nodes of the tree providing a more robust reconstruction with the potential to reveal the functional and regulatory features of HKs from basal vertebrates. However, addressing these information gaps may be challenging due to the limited number of modern species from which to draw sequences.

The biochemical characterization of extant HKs from a broad taxonomic scope offers another route to expand our understanding of the evolutionary distribution of allosteric properties in this enzyme family. Apart from amphibian HK3s, comprehensive biochemical investigations have been limited to mammalian HKs. Biochemical data of HKs from species that diverged during early vertebrate evolution, such as jawless vertebrates or fish, may provide insights into the degree to which the distinct features found in the HK isozymes are conserved across classes. In addition, kinetic studies of enzymes from these organisms may aid the approximation of functional or regulatory transitions along the evolutionary trajectories leading to modern HKs. Mochizuki isolated a low *K*_m_, 90 kDa HK from the lamprey *Lethenteron camtschaticum* and found it displays low sensitivity to G6P (Mochizuki, 1981). Unfortunately, the functional organization of the enzyme nor the G6P *K*_i_ value were reported. Characterization of the lamprey HK1-like enzyme used as the outgroup in this study, or the enzyme encoded by its *hk2*-like gene, could reveal the extent to which features of the respective mammalian orthologs (i.e. heterotropic regulation in HK1 and robust N-domain activity in HK2) are conserved in HKs from jawless vertebrates. Similarly, characterization of fish HK3s may provide further insights into the evolutionary origins of homotropic regulation. In conclusion, our phylogenetic investigation of the HK enzyme family using *Topiary* revealed gaps in both sequencing and biochemical data from early vertebrate species. Filling these gaps will likely improve the resolution at early nodes of our phylogenetic tree and reveal the functional and regulatory features of HKs from basal vertebrates.

## MATERIALS & METHODS

### Phylogenetic analysis and reconstruction of hexokinase ancestors

We performed phylogenetic analysis and ancestral sequence reconstruction in the python framework *Topiary* using the following software packages: NCBI blast+ 2.16.0, MUSCLE 5.1.linux64, GeneRax 2.0.4, RaxML-NG 1.1.0, and Open MPI 4.1.6 (Camacho et al., 2009; Edgar, 2022; Kozlov et al., 2019; Morel et al., 2020). For a detailed description of the workflow, the *Topiary* documentation can be consulted. Briefly, a seed-dataset with a total of eight extant HK sequences served as the input file for the *Topiary* pipeline (Table SI). This included the four human 100kDa HKs (HK1-3, and HKDC1) and a respective ortholog for each HK paralog to define the taxonomic scope. Orthologs were verified via reciprocal BLAST and synteny analysis.

*Topiary* initially collected ∼5000 sequences from the NCBI nr database, automatically culled these sequences using default parameters, and generated a MSA with a total of 544 HK sequences. Manual refinement of the MSA as recommended by *Topiary* included the removal of: 1) Amino acid residues preceding the conserved Leu26 (human HK1 numbering) due to high variability in N-terminal lengths, 2) Low-quality sequences containing unknown/unassigned/ambiguous amino acid residues, 3) Sequences with long indels yielding a total sequence length below 847 or above 898 residues, 4) lower quality sequence(s) of lineage specific duplicates, if present. In addition, sequences that were annotated as more than one paralog or mis/unassigned were removed. This resulted in a MSA containing 368 sequences. Initial attempts to construct a reconciled phylogenetic tree with this dataset yielded a tree in which a cluster of 20 bird HK2 sequences grouped with HK3s. In addition, we observed spurious duplication assignments arising from discordances between the gene and species trees with unknown origins. These ambiguities might stem from incomplete lineage sorting; however, this lies outside the scope of GeneRax/this study. To resolve the excessive gene duplications, we removed the 20 bird HK2 sequences and any sequences at points of tree discordance, yielding a final MSA with a total of 231 HK sequences (Supplementary phylogenetic material S1). This MSA was used to construct the reconciled phylogenetic tree shown in Figure 2, and to infer ancestral HK sequences at each node of the tree (Supplementary phylogenetic material S2-3). *Topiary* selected JTT+G8+FO+IO as the best fit evolutionary model for phylogenetic analysis. The resulting reconstructed sequences for the ML and altAll ancestors, as well as their posterior probabilities, are provided in the Supplementary phylogenetic material S3.

### Protein production and mutagenesis

Both extant and resurrected HKs were expressed and purified as recently described, with minor changes (Freye and Miller 2025). Briefly, N-terminal His^6^-tagged HKs were expressed from pET-22b(+) constructs (GenScript) in BL21(DE3) cells grown in Luria-Bertani broth with induction at 20°C and 0.75 mM IPTG. HKs were purified via HisTrap FF affinity column (Cytiva Life Sciences) and Superdex 200 10/300 GL size-exclusion chromatography (Cytiva Life Sciences). Mutagenesis was performed using QuikChange (Agilent) or Q5 (NEB) according to manufacturer protocols. DNA sequences were verified via whole plasmid sequencing (Plasmidsaurus).

### Enzyme activity and inhibition assays

Hexokinase activity was assayed at 25°C as described previously (Freye and Miller 2025). Inhibition assays were conducted by coupling the production of ADP to the oxidation of NADH via an enzyme-coupled spectrophotometric assay using rabbit muscle pyruvate kinase (MP Biomedicals) and rabbit muscle or porcine heart lactate dehydrogenase (MilliporeSigma). Reaction progress was followed at 340 nm at 25 °C (Agilent Cary UV-Vis spectrophotometer). Assays contained Tris (0.1 M, pH 8.0). KCl (50 mM), dithiothreitol (10 mM), pyruvate kinase (5 - 10 units), lactate dehydrogenase (5 units), NADH (0.5 mM), phosphoenolpyruvate (5 mM), glucose (0.25 - 2 mM), and variable ATP (0.5 - 16 mM), MgCl_2_ (1.5 - 17 mM), and G6P (10 - 500 μM) depending on the enzyme under investigation. MgCl_2_ concentrations were kept in 1 mM excess relative to ATP concentrations. Assays were initiated with HK and data were fit to the mixed non-competitive (eq. 1), competitive (eq. 2), pure non-competitive (eq. 3), or uncompetitive (eq. 4) inhibition equation in GraphPad Prism. GraphPad Prism defines the mechanism of inhibition by α. When α is very large the model approaches competitive inhibition. When α = 1 the model approaches pure non-competitive inhibition and when α is very small it cannot be fit separately from *K*_i_, hence the combined product, *K*_i_’, is reported for uncompetitive inhibition.

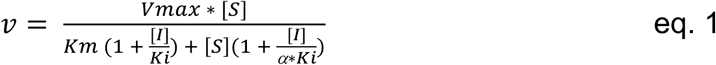

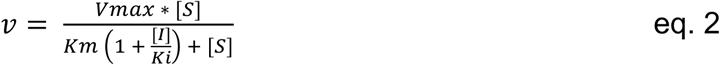

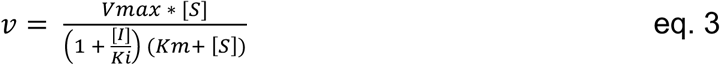

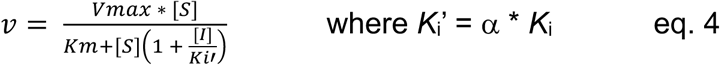

Relief of G6P inhibition by inorganic phosphate (1 - 20 mM) was investigated using the pyruvate kinase/lactate dehydrogenase assay as described above, while maintaining constant concentrations of glucose (1 mM), ATP (1.5 mM), MgCl_2_ (2.5 mM), and G6P (15-70 μM, depending on the enzyme investigated).

### Circular dichroism spectroscopy

A truncated version of Anc1N (stop codon inserted after Ala446) and Anc1C were extensively dialyzed against a HEPES buffer (10 mM, pH 8.0) at 4°C. Circular dichroism spectra of each construct (final concentration = 9 μM) were obtained from 200-280 nm using a Chirascan VX spectrophotometer (AppliedPhotophysics) with a path length of 0.5 mm. The baseline was obtained by averaging three scans of the HEPES buffer (without protein). The baseline was subtracted from the average of three scans of protein sample to correct the spectra obtained for each isolated domain. Data was processed using the Chirascan system (pro-data viewer).

## Supporting information

Supplementary phylogenetic material

## SUPPLEMENTARY MATERIAL DESCRIPTION

The Supplementary material.pdf file includes the sequences of all proteins characterized in this study, the seed-dataset used as input for *Topiary*, a comparison of different kinetic models for the G6P inhibition of HKs, circular dichroism spectra, and all steady-state kinetics conducted in this study. In addition, the final MSA of all extant HK sequences and their respective accession codes, reconciled phylogenetic tree(s) in newick format, and the ML and altAll protein sequences of all reconstructed HK ancestors are provided in a separate file (Supplementary phylogenetic material.docx).

## ACKNOWLEDGEMENTS

This work was supported by the National Institutes of Health Grants GM133843 and GM157172 (B.G.M.) and the Pfeiffer Professorship for Cancer Research (B.G.M). We acknowledge Dr. Diego Zorio for assistance with Q5 mutagenesis, Dr. S. Alexander Townsend and Nodin Weddington for computational support, and Dr. Peter Randolph for assistance with circular dichroism.

## AUTHOR CONTRIBUTIONS

C.F.: conceptualization; formal analysis; investigation; methodology; visualization; writing – original draft; writing – review and editing. A.C.W.: supervision writing – review and editing. B.G.M.: conceptualization; funding acquisition; methodology; project administration; supervision; writing – review and editing.

