## Supplementary material for "Phylogenetic Investigation of the 100 kDa Hexokinase Enzyme Family with the Topiary Ancestral Sequence Reconstruction Pipeline"

### TABLE OF CONTENTS


**Protein sequences**

1. Protein sequences of the resurrected HK ancestors, respective isolated N- and C-domain variants, and human HK1 .......... 1-3

**Tables**

SI. Input seed-dataset for *Topiary* .......... 4-7

SII. Comparison of inhibition models .......... 8

SIII. Enzymatic turnover of Anc1, Anc2, and Anc1 V83L at saturating and super-saturating glucose concentrations .......... 9

**Supporting figures**

S1. Steady-state kinetic assays of Anc1 with varying glucose concentrations fit to the Michaelis-Menten equation .......... 10

S2. Steady-state kinetic assays of Anc2 with varying glucose concentrations fit to the Michaelis-Menten equation .......... 11

S3. Steady-state kinetic assays of Anc1 with varying ATP concentrations fit to the Michaelis-Menten equation .......... 12

S4. Steady-state kinetic assays of Anc2 with varying ATP concentrations fit to the Michaelis-Menten equation .......... 13

S5. Steady-state kinetic assays of altAll-Anc1 with varying glucose concentrations fit to the Michaelis-Menten equation .......... 14

S6. Steady-state kinetic assays of altAll-Anc2 with varying glucose concentrations fit to the Michaelis-Menten equation .......... 15

S7. Steady-state kinetic assays of altAll-Anc1 with varying ATP concentrations fit to the Michaelis-Menten equation .......... 16

S8. Steady-state kinetic assays of alt-All Anc2 with varying ATP concentrations fit to the Michaelis-Menten equation .......... 17

S9. Steady-state kinetic assays of Anc1C with varying glucose concentrations fit to the Michaelis-Menten equation .......... 18

S10. Steady-state kinetic assays of Anc1C with varying ATP concentrations fit to the Michaelis-Menten equation .......... 19

S11. Steady-state kinetic assays of Anc2C with varying glucose concentrations fit to the Michaelis-Menten equation .......... 20

S12. Steady-state kinetic assays of Anc2C with varying ATP concentrations fit to the Michaelis-Menten equation .......... 21

S13. Circular dichroism spectrum of Anc1N .......... 22

S14. Circular dichroism spectrum of Anc1C .......... 23

S15. Steady-state kinetic inhibition assays of Anc1 with varying glucose 6-phosphate and ATP concentrations fit to the mixed non-competitive inhibition equation .......... 24

S16. Steady-state kinetic inhibition assays of Anc2 with varying glucose 6-phosphate and ATP concentrations fit to the mixed non-competitive inhibition equation .......... 25

S17. Steady-state kinetic inhibition assays of altAll-Anc1 with varying glucose 6-phosphate and ATP concentrations fit to the mixed non-competitive inhibition equation .......... 26

S18. Steady-state kinetic inhibition assays of altAll-Anc2 with varying glucose 6-phosphate and ATP concentrations fit to the mixed non-competitive inhibition equation .......... 27

S19. Steady-state kinetic assays of Anc1 D518A with varying glucose concentrations fit to the Michaelis-Menten equation .......... 28

S20. Steady-state kinetic assays of Anc1 D518A with varying ATP concentrations fit to the Michaelis-Menten equation .......... 29

S21. Steady-state kinetic assays of Anc2 D518A with varying glucose concentrations fit to the Michaelis-Menten equation .......... 30

S22. Steady-state kinetic assays of Anc2 D518A with varying ATP concentrations fit to the Michaelis-Menten equation .......... 31

S23. Steady-state kinetic inhibition assays of Anc1 D518A with varying glucose 6-phosphate and ATP concentrations fit to the mixed non-competitive inhibition equation .......... 32

S24. Steady-state kinetic inhibition assays of Anc2 D518A with varying glucose 6-phosphate and ATP concentrations fit to the mixed non-competitive inhibition equation .......... 33

S25. Steady-state kinetic inhibition assays of Anc1C with varying glucose 6-phosphate and ATP concentrations fit to the mixed non-competitive inhibition equation .......... 34

S26. Steady-state kinetic inhibition assays of Anc2C with varying glucose 6-phosphate and ATP concentrations fit to the mixed non-competitive inhibition equation .......... 35

S27. Glucose 6-phosphate inhibition relief of Anc1 by inorganic phosphate .......... 36

S28. Glucose 6-phosphate inhibition relief of Anc2 by inorganic phosphate .......... 37

S29. Glucose 6-phosphate inhibition relief of human HK1 by inorganic phosphate .......... 38

S30. Steady-state kinetic assays of Anc1 V83L with varying glucose concentrations fit to the Michaelis-Menten equation .......... 39

S31. Steady-state kinetic assays of Anc1 V83L with varying ATP concentrations fit to the Michaelis-Menten equation .......... 40

#### Protein sequences

*ML-Anc1 (Topiary numbering: anc229)*
MHHHHHHGSGSLYHMRLSDETLQDIMNRMRTEMEKGLGRDTNPTATVKMLPTFVRSTPDGTEKGDFLALD
LGGSNFRVLRVKVSDDGKQKVEMESEIYAIPEDIMRGSGTQLFDHVAECLGNFMEKLQIKDKKLPLGFTF
SFPCRQTKLDESVLITWTKGFKASGVEGRDVVKLLRKAIQRRGDFDIDIVAVVNDTVGTMMTCGYDDHRC
EIGLIVGTGTNACYMEEMRNIDLVEGDEGRMCINMEWGAFGDDGSLDDIRTEFDLEIDRGSLNPGKQLFE
KMISGMYMGELVRLILVKMAKQGLLFGGKITPELLTKGHFETKYVSAIEKDKEGLSKAKEILTKLGLEPS
EEDCVAVQRICTIVSTRSANLCAATLAAVVTRIKENKGVKRLRTTVGVDGTVYKTHPQFARRLHKTVRRL
APDCDVRFLLSEDGSGKGAAMVTAVAYRLASQRKQIDETLAPFKLSHEQLLEVKKRMRTEMERGLKKETH
SSATVKMLPTYVRSTPDGTEKGDFLALDLGGTNFRVLLVKIRSGKRRGVEMHNKIYSIPQEVMQGTGEEL
FDHIVHCISDFLDYMGMKGARLPLGFTFSFPCRQTSLDQGILLNWTKGFKATGCEGEDVVNLLREAIKRR
EEFDLDVVAVVNDTVGTMMTCAYEDPNCEIGLIVGTGSNACYMEEMRNVEMVEGDEGRMCINMEWGAFGD
NGCLDDIRTEFDRAVDELSLNPGKQRYEKMISGMYLGEIVRNILIDFTKRGLLFRGRISERLKTRGIFET
KFLSQIESDRLALLQVRAILQQLGLESTCDDSIIVKEVCGVVSRRAAQLCGAGMAAVVDKIRENRGLDHL
KITVGVDGTLYKLHPHFSKIMHETVKELAPKCDVTFLQSEDGSGKGAALITAVACRLREAGQH

*ML-Anc1N*
MHHHHHHGSGSLYHMRLSDETLQDIMNRMRTEMEKGLGRDTNPTATVKMLPTFVRSTPDGTEKGDFLALD
LGGSNFRVLRVKVSDDGKQKVEMESEIYAIPEDIMRGSGTQLFDHVAECLGNFMEKLQIKDKKLPLGFTF
SFPCRQTKLDESVLITWTKGFKASGVEGRDVVKLLRKAIQRRGDFDIDIVAVVNDTVGTMMTCGYDDHRC
EIGLIVGTGTNACYMEEMRNIDLVEGDEGRMCINMEWGAFGDDGSLDDIRTEFDLEIDRGSLNPGKQLFE
KMISGMYMGELVRLILVKMAKQGLLFGGKITPELLTKGHFETKYVSAIEKDKEGLSKAKEILTKLGLEPS
EEDCVAVQRICTIVSTRSANLCAATLAAVVTRIKENKGVKRLRTTVGVDGTVYKTHPQFARRLHKTVRRL
APDCDVRFLLSEDGSGKGAAMVTAVAYRLASQRKQIDETLAPFKL

*ML-Anc1C*
MHHHHHHGSGSYRLASQRKQIDETLAPFKLSHEQLLEVKKRMRTEMERGLKKETHSSATVKMLPTYVRST
PDGTEKGDFLALDLGGTNFRVLLVKIRSGKRRGVEMHNKIYSIPQEVMQGTGEELFDHIVHCISDFLDYM
GMKGARLPLGFTFSFPCRQTSLDQGILLNWTKGFKATGCEGEDVVNLLREAIKRREEFDLDVVAVVNDTV
GTMMTCAYEDPNCEIGLIVGTGSNACYMEEMRNVEMVEGDEGRMCINMEWGAFGDNGCLDDIRTEFDRAV
DELSLNPGKQRYEKMISGMYLGEIVRNILIDFTKRGLLFRGRISERLKTRGIFETKFLSQIESDRLALLQ
VRAILQQLGLESTCDDSIIVKEVCGVVSRRAAQLCGAGMAAVVDKIRENRGLDHLKITVGVDGTLYKLHP
HFSKIMHETVKELAPKCDVTFLQSEDGSGKGAALITAVACRLREAGQH

*altAll-Anc1 (Topiary numbering: anc229_altAll)*
MHHHHHHGSGSLYHMRLSDETLQDIMNRMRTEMEKGLGKETNPTATVKMLPTFVRSTPDGSEKGDFLALD
LGGTNFRVLRVKVSDDGKQTVEMENQIYEIPEDIMHGTGTQLFDHVAECLGSFMEKQQIKDKKLPLGFTF
SFPCRQTKLDESILITWTKGFKASGVEGQDVVKLLREAIKKRGDYDVDIVAVVNDTVGTMMTCGYDDHRC
EIGLIVGTGTNACYMEEMRHIEQVEGDEGRMCINMEWGAFGDDGALDDIRTEFDREVDRGSLNPGKQLFE
KMISGMYMGELVRLILVKMATEGLLFGGKTTPELLTKGHFETKYVSAIEKDKEGLSKAKEILTKLGLEPS
EEDCIAVQQICTIVSTRSANLCAAALAAVVTRLRENKGVQRLRTTVGVDGTVYKKHPQYSKRLQQTVRRL
VPDCDVQFLLSEDGSGKGAAMVTAVAYRLAAQRKQIDETLAPFKLSHEQLLEVKRRMRTEMERGLKKETH
STATVKMLPTYVRSTPDGTEKGDFLALDLGGTNFRVLLVKVRSGKKRSVEMHNKIYAIPQEVMQGTGEEL
FDHIVDCISDFLEYMGMKGARLPLGFTFSFPCKQTSLDQGILVNWTKGFKASGCEGEDVVNLLREAIKRR
EEFELDVVAIVNDTVGTMMTCAYEDPNCEIGLIVGTGSNACYMEEMRNVEMVEGDEGRMCINMEWGAFGD
NGCLDDIRTEYDRAVDELSLNPGKQRYEKMISGMYLGEIVRNILIDFTKRGLLFRGQISERLKTRGIFET
KFLSQIESDRLALLQVRAILQELGLDSTCDDSIIVKEVCGVVSRRAAQLCGAGMAAVVDKIRENRGLDHL
KVTVGVDGTLYKLHPHFSRIMHETVKELAPKCEVTFLQSEDGSGKGAALITAVACRLREAGQH

*ML-Anc2 (Topiary numbering: anc165)*
MHHHHHHGSGSLYHMRLSDETLQDIMNRFRTEMEKGLGRDTNPTATVKMLPTFVRSTPDGTEKGDFLALD
LGGSNFRVLRVKVSDDGKQKVEMESQIYAIPEDIMRGSGTQLFDHVAECLGNFMEKLQIKDKKLPLGFTF
SFPCRQTKLDESILITWTKGFKASGVEGRDVVKLLRKAIKKRGDFDIDIVAVVNDTVGTMMTCGYDDHRC
EIGLIVGTGTNACYMEEMRHIDLVEGDEGRMCINMEWGAFGDDGSLDDIRTEFDREIDRGSLNPGKQLFE
KMISGMYMGELVRLILVKMAKEGLLFGGKITPELLTKGHFETKYVSAIEKDKEGLSKAKEILTKLGLEPS
EEDCIAVQRICTIVSTRSANLCAATLAAVLTRIKENKGVKRLRTTVGVDGTVYKTHPQFARRLHKTVRRL
VPDCDVRFLLSEDGSGKGAAMVTAVAYRLASQRKQIDETLAPFKLSHEQLLEVKKRMRTEMERGLKKETH
SSATVKMLPTYVRSTPDGTEKGDFLALDLGGTNFRVLLVKIRSGKRRGVEMHNKIYAIPQEVMQGTGEEL
FDHIVHCISDFLDYMGMKGARLPLGFTFSFPCRQTSLDQGILLNWTKGFKATGCEGEDVVNLLREAIKRR
EEFDLDVVAVVNDTVGTMMTCAYEDPNCEIGLIVGTGSNACYMEEMRNVEMVEGDEGRMCINMEWGAFGD
NGCLDDIRTEFDRAVDELSLNPGKQRYEKMISGMYLGEIVRNILIDFTKRGLLFRGRISERLKTRGIFET
KFLSQIESDRLALLQVRAILQQLGLESTCDDSIIVKEVCGVVSRRAAQLCGAGMAAVVDKIRENRGLDHL
KITVGVDGTLYKLHPHFSKIMHETVKELAPKCDVTFLQSEDGSGKGAALITAVACRLREAGQH

*ML-Anc2N*
MHHHHHHGSGSLYHMRLSDETLQDIMNRFRTEMEKGLGRDTNPTATVKMLPTFVRSTPDGTEKGDFLALD
LGGSNFRVLRVKVSDDGKQKVEMESQIYAIPEDIMRGSGTQLFDHVAECLGNFMEKLQIKDKKLPLGFTF
SFPCRQTKLDESILITWTKGFKASGVEGRDVVKLLRKAIKKRGDFDIDIVAVVNDTVGTMMTCGYDDHRC
EIGLIVGTGTNACYMEEMRHIDLVEGDEGRMCINMEWGAFGDDGSLDDIRTEFDREIDRGSLNPGKQLFE
KMISGMYMGELVRLILVKMAKEGLLFGGKITPELLTKGHFETKYVSAIEKDKEGLSKAKEILTKLGLEPS
EEDCIAVQRICTIVSTRSANLCAATLAAVLTRIKENKGVKRLRTTVGVDGTVYKTHPQFARRLHKTVRRL
VPDCDVRFLLSEDGSGKGAAMVTAVAYRLASQRKQIDETLAPFKL

*ML-Anc2C*
MHHHHHHGSGSYRLASQRKQIDETLAPFKLSHEQLLEVKKRMRTEMERGLKKETHSSATVKMLPTYVRST
PDGTEKGDFLALDLGGTNFRVLLVKIRSGKRRGVEMHNKIYAIPQEVMQGTGEELFDHIVHCISDFLDYM
GMKGARLPLGFTFSFPCRQTSLDQGILLNWTKGFKATGCEGEDVVNLLREAIKRREEFDLDVVAVVNDTV
GTMMTCAYEDPNCEIGLIVGTGSNACYMEEMRNVEMVEGDEGRMCINMEWGAFGDNGCLDDIRTEFDRAV
DELSLNPGKQRYEKMISGMYLGEIVRNILIDFTKRGLLFRGRISERLKTRGIFETKFLSQIESDRLALLQ
VRAILQQLGLESTCDDSIIVKEVCGVVSRRAAQLCGAGMAAVVDKIRENRGLDHLKITVGVDGTLYKLHP
HFSKIMHETVKELAPKCDVTFLQSEDGSGKGAALITAVACRLREAGQH

*altAll-Anc2 (Topiary numbering: anc165_altAll)*
MHHHHHHGSGSLYHMRLSDETLQDIMNRFRTEMEKGLGRDTNPTATVKMLPTFVRSTPDGSEKGDFLALD
LGGTNFRVLRVKVSDDGKQKVEMENQIYAIPEDIIHGSGTQLFDHVAECLGDFMEKQQIKDKKLPLGFTF
SFPCRQTKLDESVLITWTKGFKASGVEGRDVVSLLRKAIQKRGDYDVDIVAVVNDTVGTMMTCGYDDHNC
EIGLIVGTGTNACYMEEMRHIDLVEGDEGRMCINMEWGAFGDDGALDDIRTEFDREIDRGSLNPGKQLFE
KMISGMYMGELVRLILVKMAKEGLLFGGKITPELLTKGHFETKYVSAIEKDKEGLSKAKEILTKLGLEPS
EEDCVAVQHICTIVSTRSANLCAAALAAVLTRLRENKGVQRLRTTVGVDGTVYKKHPQYARRLHKTVRRL
VPDCDVRFLLSEDGSGKGAAMVTAVAYRLASQRKQIDETLAPFKLSHEQLLEVKRRMRTEMERGLKKETH
NTATVKMLPTYVRSTPDGTEKGDFLALDLGGTNFRVLLVKVRSGKRRSVEMHNKIYSIPQEVMQGTGEEL
FDHIVHCISDFLEYMGMKGVRLPLGFTFSFPCKQTSLDQGILLNWTKGFKATGCEGEDVVNLLREAIKRR
EEFELDVVAIVNDTVGTMMTCAYEDPNCEVGLIVGTGSNACYMEEMRNVEMVEGDEGRMCINMEWGAFGD
NGCLDDIRTEFDRAVDELSLNPGKQRYEKMISGMYLGEIVRNILIDFTKRGLLFRGQISERLKTRGIFET
KFLSQIESDRLALLQVRAILQQLGLDSTCDDSIIVKEVCGVVSRRAAQLCGAGMAAVVDKIRENRGLDHL
KVTVGVDGTLYKLHPHFSRIMHETVKELAPKCEVTFLQSEDGSGKGAALITAVACRIREAGQH

*Human HK1 (Δ10)*

MHHHHHHGSGSELNYFQSMFTELKDDQVKKIDKYLYAMRLSDETLIDIMTRFRKEMKNGLSRDFNPTATV
KMLPTFVRSIPDGSEKGDFIALDLGGSSFRILRVQVNHEKNQNVHMESEVYDTPENIVHGSGSQLFDHVA
ECLGDFMEKRKIKDKKLPVGFTFSFPCQQSKIDEAILITWTKRFKASGVEGADVVKLLNKAIKKRGDYDA
NIVAVVNDTVGTMMTCGYDDQHCEVGLIIGTGTNACYMEELRHIDLVEGDEGRMCINTEWGAFGDDGSLE
DIRTEFDREIDRGSLNPGKQLFEKMVSGMYLGELVRLILVKMAKEGLLFEGRITPELLTRGKFNTSDVSA
IEKNKEGLHNAKEILTRLGVEPSDDDCVSVQHVCTIVSFRSANLVAATLGAILNRLRDNKGTPRLRTTVG
VDGSLYKTHPQYSRRFHKTLRRLVPDSDVRFLLSESGSGKGAAMVTAVAYRLAEQHRQIEETLAHFHLTK
DMLLEVKKRMRAEMELGLRKQTHNNAVVKMLPSFVRRTPDGTENGDFLALDLGGTNFRVLLVKIRSGKKR
TVEMHNKIYAIPIEIMQGTGEELFDHIVSCISDFLDYMGIKGPRMPLGFTFSFPCQQTSLDAGILITWTK
GFKATDCVGHDVVTLLRDAIKRREEFDLDVVAVVNDTVGTMMTCAYEEPTCEVGLIVGTGSNACYMEEMK
NVEMVEGDQGQMCINMEWGAFGDNGCLDDIRTHYDRLVDEYSLNAGKQRYEKMISGMYLGEIVRNILIDF
TKKGFLFRGQISETLKTRGIFETKFLSQIESDRLALLQVRAILQQLGLNSTCDDSILVKTVCGVVSRRAA
QLCGAGMAAVVDKIRENRGLDRLNVTVGVDGTLYKLHPHFSRIMHQTVKELSPKCNVSFLLSEDGSGKGA
ALITAVGVRLRTEASS

#### Tables

**SI.** Input seed-dataset for *Topiary*.

| Species | Name | Sequence | Aliases |
|---|---|---|---|
| Homo sapiens<br>(UniProt: P19367-1<br>NCBI: P19367) | HK1 | MIAAQLLAYYFTELKDDQVKKIDKYLYAMRLSDETLIDIMT<br>RFRKEMKNGLSRDFNPTATVKMLPTFVRSIPDGSEKGDFIA<br>LDLGGSSFRILRVQVNHEKNQNVHMESEVYDTPENIVHGSG<br>SQLFDHVAECLGDFMEKRKIKDKKLPVGFTFSFPCQQSKID<br>EAILITWTKRFKASGVEGADVVKLLNKAIKKRGDYDANIVA<br>VVNDTVGTMMTCGYDDQHCEVGLIIGTGTNACYMEELRHID<br>LVEGDEGRMCINTEWGAFGDDGSLEDIRTEFDREIDRGSLN<br>PGKQLFEKMVSGMYLGELVRLILVKMAKEGLLFEGRITPEL<br>LTRGKFNTSDVSAIEKNKEGLHNAKEILTRLGVEPSDDDCV<br>SVQHVCTIVSFRSANLVAATLGAILNRLRDNKGTPRLRTTV<br>GVDGSLYKTHPQYSRRFHKTLRRLVPDSDVRFLLSESGSGK<br>GAAMVTAVAYRLAEQHRQIEETLAHFHLTKDMLLEVKKRMR<br>AEMELGLRKQTHNNAVVKMLPSFVRRTPDGTENGDFLALDL<br>GGTNFRVLLVKIRSGKKRTVEMHNKIYAIPIEIMQGTGEEL<br>FDHIVSCISDFLDYMGIKGPRMPLGFTFSFPCQQTSLDAGI<br>LITWTKGFKATDCVGHDVVTLLRDAIKRREEFDLDVVAVVN<br>DTVGTMMTCAYEEPTCEVGLIVGTGSNACYMEEMKNVEMVE<br>GDQGQMCINMEWGAFGDNGCLDDIRTHYDRLVDEYSLNAGK<br>QRYEKMISGMYLGEIVRNILIDFTKKGFLFRGQISETLKTR<br>GIFETKFLSQIESDRLALLQVRAILQQLGLNSTCDDSILVK<br>TVCGVVSRRAAQLCGAGMAAVVDKIRENRGLDRLNVTVGVD<br>GTLYKLHPHFSRIMHQTVKELSPKCNVSFLLSEDGSGKGAA<br>LITAVGVRLRTEASS | hexokinase-1; brain form hexokinase; HKI; hexokinase-A; HK1; HXK1; HK1-ta; HK1-tb; HK1-tc |
| Danio rerio<br>(NCBI: NP_998417) | HK1 | MIAAQLLAYYFTELKDDQVKKIDKYLYAMRFSDETLRDIMA<br>RFRREMENGLARDTNPTATAKMLPTFVRSIPDGSEKGDFIA<br>LDLGGSNFRILRVKVSHEKKQTVQMESQIYETPEDIIHGSR<br>SRLFDHVAECLGDFMEKQKIKDKKLPVGFTFSFPCSQSKLD<br>EAVLLTWTKRFKVNGVEGMDVVKLLNKAIKKRGDYEADIMA<br>VVNDTVGTMMTCGFDDQRCEVGIIIGTGTNACYMEELRHMD<br>MVEGDEGRMCINTEWGAFGDDGTLEDIRTEFDREIDRGSLN<br>PGKQLFEKMVSGMYMGELVRLILVKMAKEGLLFEGRITPEL<br>LTKGKIETKHVSAIEKSKEGLTKAKEILTRLGVEPSEDDCI<br>AVQHVCAIVSFRSANLIAATLGAILTRLKDNKNTPRLRTTV<br>GIDGSLYKMHPQYARRLHKTVRRLVPESDVRFLLSESGSGK<br>GAALVTAWAYRLADQERQIAETLEEFRLTKDQLLEVKKRMR<br>TEIQNGLSKSTQNTATVKMLPTYVRSTPDGSENGDFLALDL<br>GGTNFRVLLVKIRSGKRRTVEMHNKIYAIPIEVMQGTGEEL<br>FDHIVYCISDFLDYMGMKNARLPLGFTFSFPCRQTSLDAGL<br>LVNWTKGFKATDCEGEDVVGLLREGIKRREEFDLDVVAIVN<br>DTVGTMMTCAYEEPTCEVGLIAGTGSNACYMEEMRNIETVS<br>GEEGRMCVNMEWGAFGDNGCLDDIRTKYDDAVDDLSLNAGK<br>QKYEKMCSGMYLGEIVRNILIDLTKRGFLFRGQISETLKTR<br>GIFETKFLSQIESDRLALLQVRSILQHLGLDSTCDDSIIVK<br>EVCGAVSRRAAQLCGAGMAAVVDKIRENRGLDHLDITVGVD<br>GTLYKLHPHFSRIMHQTVKELAPKCNVTFLLSEDGSGKGAA<br>LITAVGCRLRQQEQKS | hexokinase-1 |

| | | | |
|---|---|---|---|
| Homo sapiens (UniProt and NCBI: P52789) | HK2 | MIASHLLAYFFTELNHDQVQKVDQYLYHMRLSDETLLEIS<br>KRFRKEMEKGLGATTHPTAAVKMLPTFVRSTPDGTEHGEF<br>LALDLGGTNFRVLWVKVTDNGLQKVEMENQIYAIPEDIMR<br>GSGTQLFDHIAECLANFMDKLQIKDKKLPLGFTFSFPCHQ<br>TKLDESFLVSWTKGFKSSGVEGRDVVALIRKAIQRRGDFD<br>IDIVAVVNDTVGTMMTCGYDDHNCEIGLIVGTGSNACYME<br>EMRHIDMVEGDEGRMCINMEWGAFGDDGSLNDIRTEFDQE<br>IDMGSLNPGKQLFEKMISGMYMGELVRLILVKMAKEELLF<br>GGKLSPELLNTGRFETKDISDIEGEKDGIRKAREVLMRLG<br>LDPTQEDCVATHRICQIVSTRSASLCAATLAAVLQRIKEN<br>KGEERLRSTIGVDGSVYKKHPHFAKRLHKTVRRLVPGCDV<br>RFLRSEDGSGKGAAMVTAVAYRLADQHRARQKTLEHLQLS<br>HDQLLEVKRRMKVEMERGLSKETHASAPVKMLPTYVCATP<br>DGTEKGDFLALDLGGTNFRVLLVRVRNGKWGGVEMHNKIY<br>AIPQEVMHGTGDELFDHIVQCIADFLEYMGMKGVSLPLGF<br>TFSFPCQQNSLDESILLKWTKGFKASGCEGEDVVTLLKEA<br>IHRREEFDLDVVAVVNDTVGTMMTCGFEDPHCEVGLIVGT<br>GSNACYMEEMRNVELVEGEEGRMCVNMEWGAFGDNGCLDD<br>FRTEFDVAVDELSLNPGKQRFEKMISGMYLGEIVRNILID<br>FTKRGLLFRGRISERLKTRGIFETKFLSQIESDCLALLQV<br>RAILQHLGLESTCDDSIIVKEVCTVVARRAAQLCGAGMAA<br>VVDRIRENRGLDALKVTVGVDGTLYKLHPHFAKVMHETVK<br>DLAPKCDVSFLQSEDGSGKGAALITAVACRIREAGQR | hexokinase-2; HK II, hexokinase-B; muscle form hexokinase; HK2; HXK2; hexokinase-2, muscle |
| Scyliorhinus canicular (NCBI: XP_038641198) | HK2 | MIASHLLAYYFTELKHDQVQKVDKYLYHMRLSDEALHDIS<br>NRFLIEMEQGLRKDTNPTSSLKMLPTFVRSTPDGTESGDF<br>LALDLGGTNFRVLRVKVSDNGKQNVQIENRICAIPEDIMH<br>GTGTQLFDHIANCLGDVMEELDIKDKKLPLGFTFSFPCQQ<br>SKLDESLLLHWTKGFKASGVEGRDVVNLLRQAIQRRGDFD<br>IDVVAVVNDTVGTMMSCGYDDHNCEIGLIVGTGTNACYME<br>DMQHIDLVEGDEGQMCINMEWGAFGDLGELHAVRTEFDRE<br>IDRGSINPGKQLFEKMISGMYMGELVRIILVKMAKDGLLF<br>GGIISPDLLIKGHFETKYVSAIERDRDGLQKACEILTRLG<br>LQPTPDDCAATQRVCTIVSTRSANLCAATLAAVLRRIKEN<br>KGVQRLRTTIGVDGSVYKKHPQFARRLHKAVRRLVPTCEV<br>RFLPSEDGSGKGAAMVTAVAYRLAKQQKVREEILATLMLS<br>DAQLLEVKRRMREEMERGLKKETHCSAAVKMLATYVRSTP<br>DGTEKGDFLALDLGGTNFRVLLVKVRSGKRRAVEMHNKIY<br>TIPMDAMEGTGEELFDHIVHCILDFLEYMGMKDCVLPLGF<br>TFSFPCEQRNLDQGILLKWTKGFKATGCEGEDVVDLLREA<br>IKRREEFELEVVAIVNDTVGTMMSCGYDDPFCEVGLIVGT<br>GCNACYMEEMRNVEMVEGDEGRMCVNMEWGAFGDNDCLDD<br>IRTEFDREVDLLSTNPGKQRFEKLISGMYLGEIVRNILIN<br>FTKRGLLFRGKISERLKTRGIFQTKFLSHIESDRLALLQV<br>RAILQQLGLQSSCDDSIIVKEVCGVVSRRAAQLCGAGMAA<br>VVDKIRENRHLDYLKVTVGVDGTLYKLHPHFSAIMHETVG<br>LLAPKCKVTFVQSEDGSGKGAALITAVACRIREAGQH | hexokinase-2 |
| Homo sapiens (UniProt and NCBI: P52790) | HK3 | MDSIGSSGLRQGEETLSCSEEGLPGPSDSSELVQECLQQF<br>KVTRAQLQQIQASLLGSMEQALRGQASPAPAVRMLPTYVG<br>STPHGTEQGDFVVLELGATGASLRVLWVTLTGIEGHRVEP<br>RSQEFVIPQEVMLGAGQQLFDFAAHCLSEFLDAQPVNKQG<br>LQLGFSFSFPCHQTGLDRSTLISWTKGFRCSGVEGQDVVQ<br>LLRDAIRRQGAYNIDVVAVVNDTVGTMMGCEPGVRPCEVG<br>LVVDTGTNACYMEEARHVAVLDEDRGRVCVSVEWGSFSDD<br>GALGPVLTTFDHTLDHESLNPGAQRFEKMIGGLYLGELVR | hexokinase-3; HK III; hexokinase-C; HK3; HXK3; hexokinase 3 (white cell) |

| | | | |
|---|---|---|---|
| | | LVLAHLARCGVLFGGCTSPALLSQGSILLEHVAEMEDPSTG<br>AARVHAILQDLGLSPGASDVELVQHVCAAVCTRAAQLCAAA<br>LAAVLSCLQHSREQQTLQVAVATGGRVCERHPRFCSVLQGT<br>VMLLAPECDVSLIPSVDGGGRGVAMVTAVAARLAAHRRLLE<br>ETLAPFRLNHDQLAAVQAQMRKAMAKGLRGEASSLRMLPTF<br>VRATPDGSERGDFLALDLGGTNFRVLLVRVTTGVQITSEIY<br>SIPETVAQGSGQQLFDHIVDCIVDFQQKQGLSGQSLPLGFT<br>FSFPCRQLGLDQGILLNWTKGFKASDCEGQDVVSLLREAIT<br>RRQAVELNVVAIVNDTVGTMMSCGYEDPRCEIGLIVGTGTN<br>ACYMEELRNVAGVPGDSGRMCINMEWGAFGDDGSLAMLSTR<br>FDASVDQASINPGKQRFEKMISGMYLGEIVRHILLHLTSLG<br>VLFRGQQIQRLQTRDIFKTKFLSEIESDSLALRQVRAILED<br>LGLPLTSDDALMVLEVCQAVSQRAAQLCGAGVAAVVEKIRE<br>NRGLEELAVSVGVDGTLYKLHPRFSSLVAATVRELAPRCVV<br>TFLQSEDGSGKGAALVTAVACRLAQLTRV | |
| Protopterus annectens (NCBI: XP_043925231) | HK3 | MNSHSEQVNSILSLFVLSDEKLQKIVSLMLENMEKGLRKQK<br>HTTSTLKMLPTFVHSTLDGTERGDFLALDLRGTKFRVLQVK<br>LTDDDSYKVQSESFEIPEDIMTGTGIQLFDYFAKHLSSFLD<br>KQQLKDKKLPLGFTFPFPCKQIDLDKSVLLTWTKGFKCSGV<br>EGEDVVQMLRDAIHRQLDCDVNVIAVVNDTVGTMMSCGYKD<br>RACEIGLVVGTGTNACYMEEMGNIEQIEGDEGRMCVIMEWG<br>AFGDDGCLRDIQTEFDLELDRHSRNPGRQTFEKKISGMYMG<br>EIVRLIVVKLASLHLLFNGVTSPALLKSGTFETNYISEIED<br>EKVGLTKAKEILLSFNLNPSEQDCVIVQKICNNVSTRSANL<br>CAAGLAAVVTRIQRNRHLKHLQTTVGVDGTVYRKHPKFSGR<br>LQATLKQLTSNCDVQFMLSEDGTGKGTAMVTAVARRLFAQR<br>QQINETMAPFCLSRKQLVQVQQRLRLEMLRGLEKETQSTAT<br>VKMLPTYVRGTPDGTERGHFLALDLGGTNFRVLLVNVNSRE<br>EGGVKMVNQIYSIPESLMQGSGTKLFDHIVDCIIDFVKKQG<br>KMGCRLPLGFTFSFPCRQTGLDKGILITWTKGFNATECEGK<br>DVVMLLREAIKRKQGVELDVVAIVNDTVGTMMSCAYNDSNC<br>EIGLIVGTGSNACYMEEMSNIKNVEGDIGQMCINMEWGAFG<br>DNGCLDVFVTQYDQIVDKASINPGKQRYEKMMSGMYLGEIV<br>RNILIDLTKQGIVFRGQISERLNTKGIFETKFLSEIESDSL<br>VLLQVRAILQKLGLDVTCDDTLIVKEVCTIVSRRAAQLCGA<br>GIAAVVEKIRENRRLNHLAITVGVDGTLYKLHPHFSKIVHE<br>TVAELAPCCDVTFHQSEDGSGKGAALITAVACRLRETGQH | hexokinase-3 |
| Homo sapiens (UniProt: Q2TB90-1 NCBI: Q2TB90) | HKDC1 | MFAVHLMAFYFSKLKEDQIKKVDRFLYHMRLSDDTLLDIMR<br>RFRAEMEKGLAKDTNPTAAVKMLPTFVRAIPDGSENGEFLS<br>LDLGGSKFRVLKVQVAEEGKRHVQMESQFYPTPNEIIRGNG<br>TELFEYVADCLADFMKTKDLKHKKLPLGLTFSFPCRQTKLE<br>EGVLLSWTKKFKARGVQDTDVVSRLTKAMRRHKDMDVDILA<br>LVNDTVGTMMTCAYDDPYCEVGVIIGTGTNACYMEDMSNID<br>LVEGDEGRMCINTEWGAFGDDGALEDIRTEFDRELDLGSLN<br>PGKQLFEKMISGLYLGELVRLILLKMAKAGLLFGGEKSSAL<br>HTKGKIETRHVAAMEKYKEGLANTREILVDLGLEPSEADCI<br>AVQHVCTIVSFRSANLCAAALAAILTRLRENKKVERLRTTV<br>GMDGTLYKIHPQYPKRLHKVVRKLVPSCDVRFLLSESGSTK<br>G | hexokinase hkdc1; hexokinase domain-containing protein 1; hkdc1; hexokinase domain containing 1; putative hexokinase hkdc1 |

| | | | |
|---|---|---|---|
| | | AAMVTAVASRVQAQRKQIDRVLALFQLTREQLVDVQAKMRA<br>ELEYGLKKKSHGLATVRMLPTYVCGLPDGTEKGKFLALDLG<br>GTNFRVLLVKIRSGRRSVRMYNKIFAIPLEIMQGTGEELFD<br>HIVQCIADFLDYMGLKGASLPLGFTFSFPCRQMSIDKGTLI<br>GWTKGFKATDCEGEDVVDMLREAIKRRNEFDLDIVAVVNDT<br>VGTMMTCGYEDPNCEIGLIAGTGSNMCYMEDMRNIEMVEGG<br>EGKMCINTEWGGFGDNGCIDDIWTRYDTEVDEGSLNPGKQR<br>YEKMTSGMYLGEIVRQILIDLTKQGLLFRGQISERLRTRGI<br>FETKFLSQIESDRLALLQVRRILQQLGLDSTCEDSIVVKEV<br>CGAVSRRAAQLCGAGLAAIVEKRREDQGLEHLRITVGVDGT<br>LYKLHPHFSRILQETVKELAPRCDVTFMLSEDGSGKGAALI<br>TAVAKRLQQAQKEN | |
| Hypanus sabinus (NCBI: XP_059803278) | HKDC1 | MFSIHLLAFYLAKLQEDQAKKIDRFLYNMQLNPEKLQDIM<br>SRFGNEMRKGLSINETSTVKMLPTFVQSTPDRTEKGDFLA<br>LDLGGSRFEVLRVRVSEDGQQVIQTERQLYPMPDEMKQSN<br>ASEFFNYVAECLGDFMEKREIKNRKFPLGFTFSFPCRQTK<br>LDEGFLISWGKGYKISGSKGANVISLLRKGIEQQGDFDVD<br>VLALVNDTVGTLMSGGFDDPYCEVGVVIGTGTNACYMEEM<br>SNIESVEGNIGRMCINTEWGAFGDDGSLNDIRTEFDLELD<br>ANSLNPGKQLFEKMISAMYLGELVRLILVRLTKSHTLFSG<br>TASAKLLTKGSFGIQHLTAVVNSNSGMENSMQILMDLDLQ<br>PSEANCKALQHICFIVMSRSISLCAAALATILIHLKNTRK<br>IKRLRTTVGIDGTVYRTVPQYPKNLHRLVRQLAPGCDVRF<br>LLVDTGTGKGAAMVTAVAYRLVNQRKYQDKTLAPLQLSRE<br>QLLKIMQIMKNEMKLGLKTETHSTATLKMLPTFIQKFPDG<br>TERGKFLALDLGGTNFRVLLLVLRNRPRMISQTYQKSYSV<br>PMEVMQGIGEELFDWIVECIVEFLDYMGMKCAPLPLGFTF<br>SFPCIQHRLDQGVLVNWTKGFKATDVEGDDVVFLLKEAIK<br>RQGELNLDIVALLNDTVGTMMTCAYENPNCEVGLIAGTGT<br>NACYMEEMKNIEIVDGVEGRMCINTEWGAFGDNGCLDDIR<br>TTFDRLVDINSLNPGKQKYEKMISGMYLGEIVRNILINFT<br>KHGLLFRGFVSPVLKTSGIFETSYMSQIESDRLGLLQVHS<br>ILQNLGLDCTCDDSIIVKEVCSVVSKRSAQLCGAGMAAVV<br>DKIRENRGLNHLEITVGVDGTLYKLHPHFSTIMQETVKDL<br>APSCDVKFVLSEDGSGKGAALVTAAVKQSN | hexokinase HKDC1-like isoform X1 |

**SII.** Comparison of inhibition models.

| | Competitive | Uncompetitive | Pure non-competitive | Mixed non-competitive | Probability that model is correct |
|---|---|---|---|---|---|
| Anc1 (1) | $K_i$ = 14.02<br>$R^2$ = 0.9074 | $K_i$' = 29.83<br>$R^2$ = 0.9569 | **$K_i$ = 51.52**<br>**$R^2$ = 0.9721** | $K_i$ = 63.58<br>$\alpha$ = 0.7304<br>$R^2$ = 0.9726 | > 88.51 |
| Anc1 (1) | $K_i$ = 12.85<br>$R^2$ = 0.9457 | $K_i$' = 40.99<br>$R^2$ = 0.9412 | $K_i$ = 62.2<br>$R^2$ = 0.9795 | **$K_i$ = 33.49**<br>**$\alpha$ = 2.668**<br>**$R^2$ = 0.9853** | > 61.82 |
| Anc1C (1) | $K_i$ = 35.48<br>$R^2$ = 0.9534 | $K_i$' = 72.26<br>$R^2$ = 0.9508 | **$K_i$ = 126.4**<br>**$R^2$ = 0.9889** | $K_i$ = 95.25<br>$\alpha$ = 1.602<br>$R^2$ = 0.9902 | > 76.01 |
| Anc1C (2) | $K_i$ = 20.90<br>$R^2$ = 0.9142 | $K_i$' = 45.37<br>$R^2$ = 0.9418 | **$K_i$ = 78.14**<br>**$R^2$ = 0.9668** | $K_i$ = 74.28<br>$\alpha$ = 1.082<br>$R^2$ = 0.9668 | > 89.79 |
| AltAll-Anc1 (1) | $K_i$ = 41.46<br>$R^2$ = 0.9660 | $K_i$' = 164.5<br>$R^2$ = 0.9306 | $K_i$ = 237.5<br>$R^2$ = 0.9788 | **$K_i$ = 89.52**<br>**$\alpha$ = 4.953**<br>**$R^2$ = 0.9946** | > 99.99 |
| altAll-Anc1 (2) | $K_i$ = 40.98<br>$R^2$ = 0.9847 | $K_i$' = 144.3<br>$R^2$ = 0.9285 | $K_i$ = 215.6<br>$R^2$ = 0.9743 | **$K_i$ = 67.47**<br>**$\alpha$ = 8.478**<br>**$R^2$ = 0.9945** | > 99.77 |
| Anc2 (1) | $K_i$ = 9.900<br>$R^2$ = 0.9476 | $K_i$' = 39.11<br>$R^2$ = 0.9255 | $K_i$ = 56.48<br>$R^2$ = 0.9696 | **$K_i$ = 23.46**<br>**$\alpha$ = 4.068**<br>**$R^2$ = 0.9819** | > 87.7 |
| Anc2 (2) | $K_i$ = 12.39<br>$R^2$ = 0.9585 | $K_i$' = 39.54<br>$R^2$ = 0.9195 | $K_i$ = 60.12<br>$R^2$ = 0.9685 | **$K_i$ = 25.10**<br>**$\alpha$ = 4.482**<br>**$R^2$ = 0.9817** | > 89.57 |
| Anc2C (1) | $K_i$ = 26.31<br>$R^2$ = 0.9614 | $K_i$' = 69.65<br>$R^2$ = 0.9283 | $K_i$ = 106.2<br>$R^2$ = 0.9635 | **$K_i$ = 45.91**<br>**$\alpha$a = 4.675**<br>**$R^2$ = 0.9734** | > 59.04 |
| Anc2C (2) | $K_i$ = 28.93<br>$R^2$ = 0.9769 | $K_i$' = 59.66<br>$R^2$ = 0.9673 | $K_i$ = 97.98<br>$R^2$ = 0.9930 | **$K_i$ = 63.64**<br>**$\alpha$ = 2.123**<br>**$R^2$ = 0.9953** | > 72.17 |
| altAll-Anc2 (1) | $K_i$ = 41.85<br>$R^2$ = 0.9778 | $K_i$' = 161.4<br>$R^2$ = 0.9176 | $K_i$ = 236.1<br>$R^2$ = 0.9725 | **$K_i$ = 76.15**<br>**$\alpha$ = 7.157**<br>**$R^2$ = 0.9954** | > 99.99 |
| altAll-Anc2 (2) | $K_i$ = 60.56<br>$R^2$ = 0.9795 | $K_i$' = 181.4<br>$R^2$ = 0.9449 | $K_i$ = 277.7<br>$R^2$ = 0.9840 | **$K_i$ = 116.1**<br>**$\alpha$ = 4.642**<br>**$R^2$ = 0.9949** | > 99.90 |
| Anc1 D518A (1) | **$K_i$ = 28.56**<br>**$R^2$ = 0.9960** | $K_i$' = 41.46<br>$R^2$ = 0.9453 | $K_i$ = 79.23<br>$R^2$ = 0.9780 | $K_i$ = 28.56<br>$\alpha$ = nd<br>$R^2$ = 0.9960 | > 89.86 |
| Anc1 D518A (2) | **$K_i$ = 38.96**<br>**$R^2$ = 0.9745** | $K_i$' = 0.1833<br>$R^2$ = 0.9490 | $K_i$ = 48.50<br>$R^2$ = 0.9728 | $K_i$ = 38.96<br>$\alpha$ = nd<br>$R^2$ = 0.9745 | > 62.6 |
| Anc2 D518A (1) | **$K_i$ = 85.29**<br>**$R^2$ = 0.9935** | $K_i$' = 172.2<br>$R^2$ = 0.9209 | $K_i$ = 293.1<br>$R^2$ = 0.9626 | $K_i$ = 85.29<br>$\alpha$ = nd<br>$R^2$ = 0.9935 | > 89.86 |
| Anc2 D518A (2) | **$K_i$ = 60.57**<br>**$R^2$ = 0.9973** | $K_i$' = 94.49<br>$R^2$ = 0.9499 | $K_i$ = 174.2<br>$R^2$ = 0.9803 | $K_i$ = 60.57<br>$\alpha$ = nd<br>$R^2$ = 0.9973 | > 89.86 |

Kinetic data for each biological replicate were fit to equations 1-4 (provided in the main manuscript) using GraphPad Prism. Parameters for the preferred model of inhibition are bold.

**SIII.** Enzymatic turnover of Anc1, Anc2, and Anc1 V83L at saturating and super-saturating glucose concentrations.

| | $k_{cat}$ ($s^{-1}$) at 1 mM glucose | $k_{cat}$ ($s^{-1}$) at 100 mM glucose |
|---|---|---|
| Anc1 | 36 ± 13 | 38 ± 15 |
| Anc2 | 49 ± 4 | 50 ± 4 |
| Anc1 V83L | 49 ± 2 | 53 ± 3 |

$k_{cat}$ values were determined as described in materials and methods using saturating ATP concentrations (15 mM).

**Supporting figures**

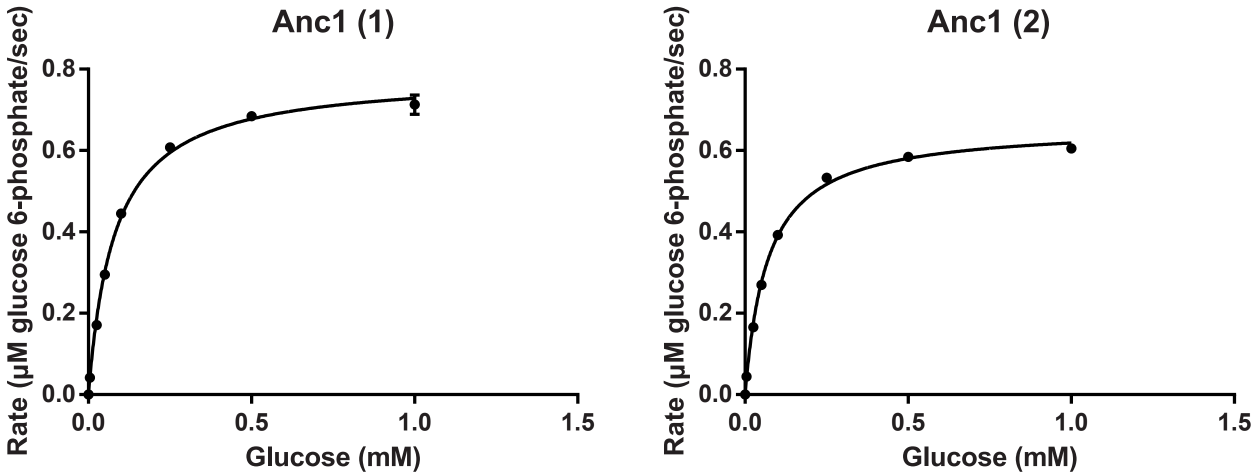


**S1.** Two biological replicates of steady-state kinetic assays of ML Anc1 with varying glucose concentrations at 15 mM ATP. Each data point represents the average of three technical replicates ± standard deviation. Data were fit to the Michaelis-Menten equation in GraphPad Prism.

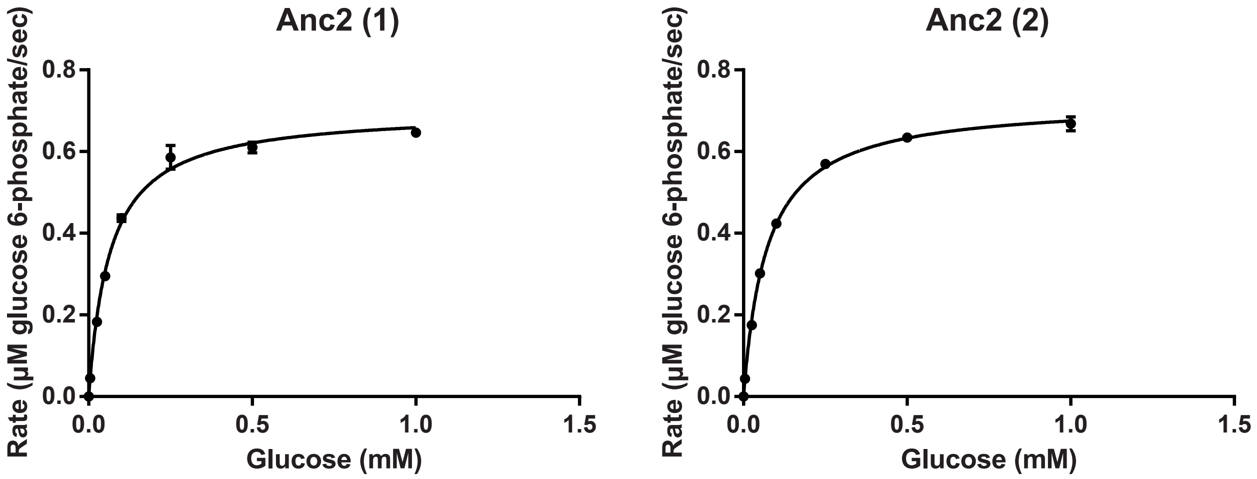


**S2.** Two biological replicates of steady-state kinetic assays of ML Anc2 with varying glucose concentrations at 15 mM ATP. Each data point represents the average of three technical replicates ± standard deviation. Data were fit to the Michaelis-Menten equation in GraphPad Prism.

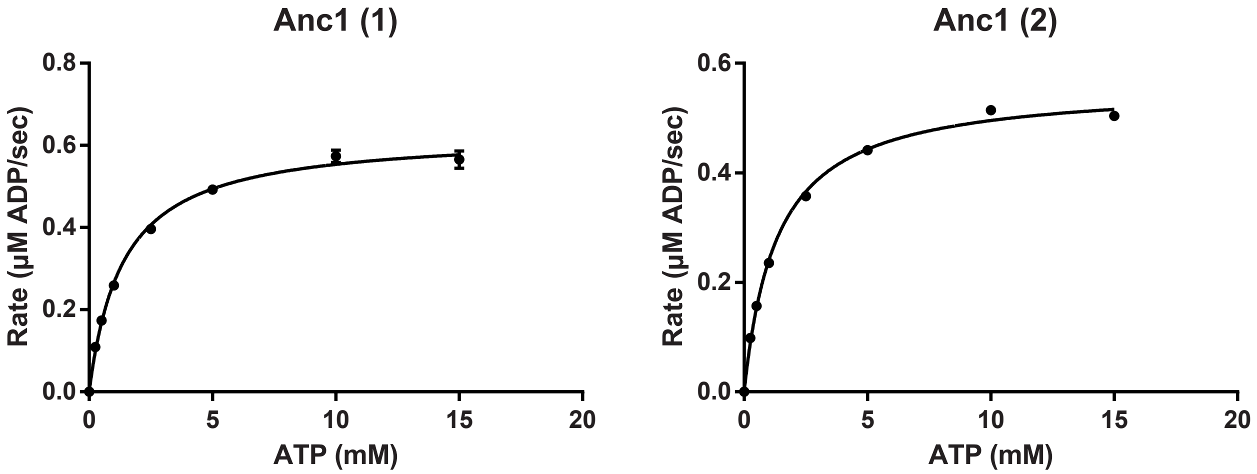


**S3.** Two biological replicates of steady-state kinetic assays of ML Anc1 with varying ATP concentrations at 1 mM glucose. Each data point represents the average of three technical replicates ± standard deviation. Data were fit to the Michaelis-Menten equation in GraphPad Prism.

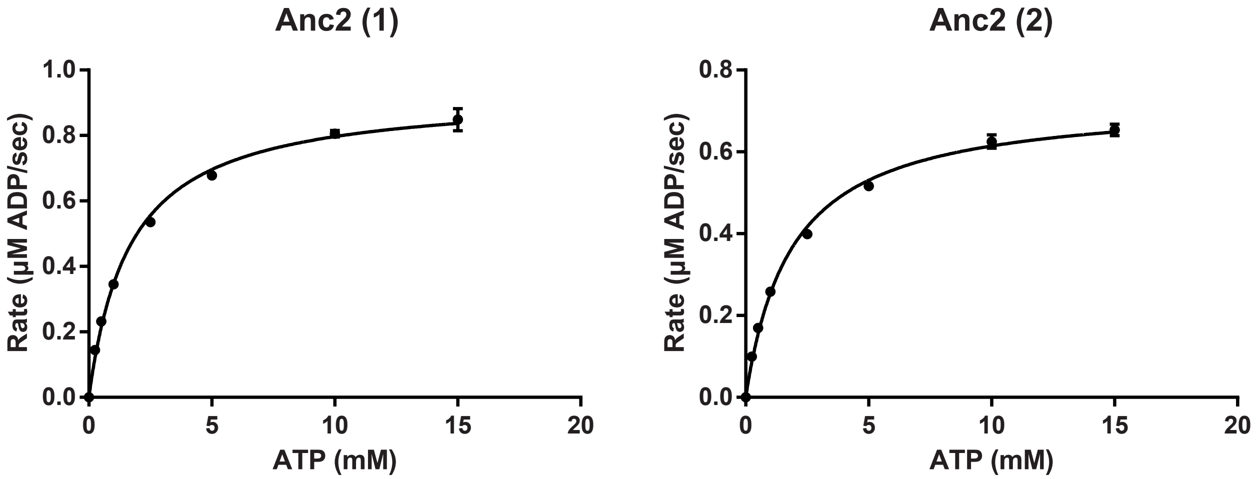


**S4.** Two biological replicates of steady-state kinetic assays of ML Anc2 with varying ATP concentrations at 1 mM glucose. Each data point represents the average of three technical replicates ± standard deviation. Data were fit to the Michaelis-Menten equation in GraphPad Prism.

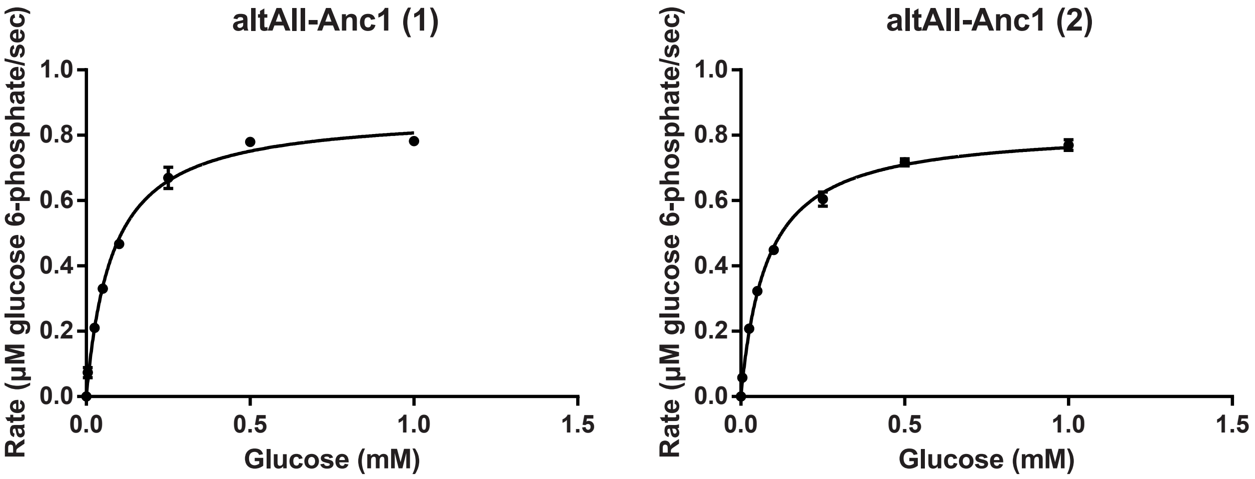


**S5.** Two biological replicates of steady-state kinetic assays of altAll-Anc1 with varying glucose concentrations at 15 mM ATP. Each data point represents the average of three technical replicates ± standard deviation. Data were fit to the Michaelis-Menten equation in GraphPad Prism.

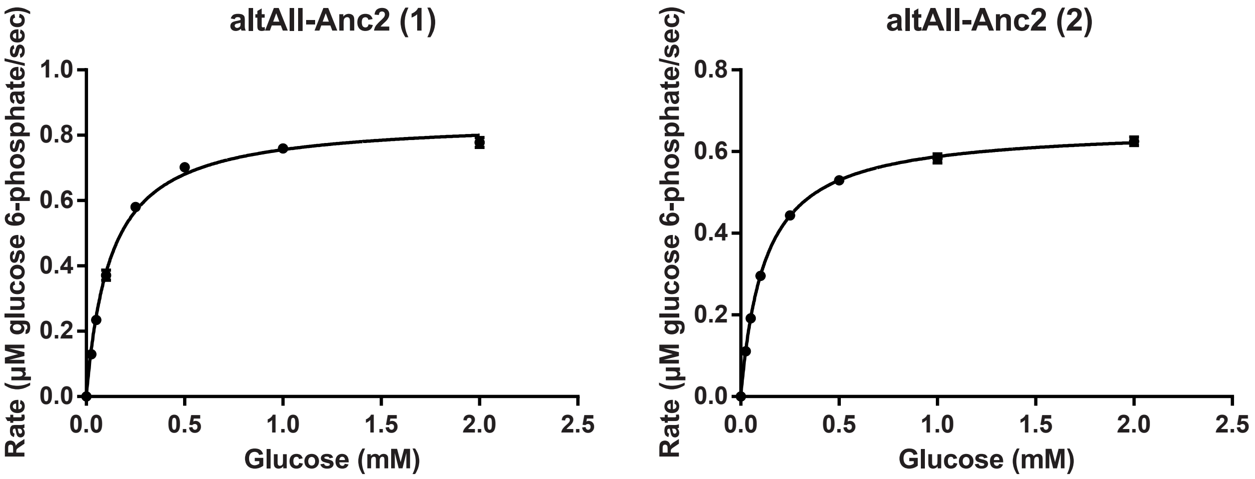


**S6.** Two biological replicates of steady-state kinetic assays of altAII-Anc2 with varying glucose concentrations at 15 mM ATP. Each data point represents the average of three technical replicates ± standard deviation. Data were fit to the Michaelis-Menten equation in GraphPad Prism.

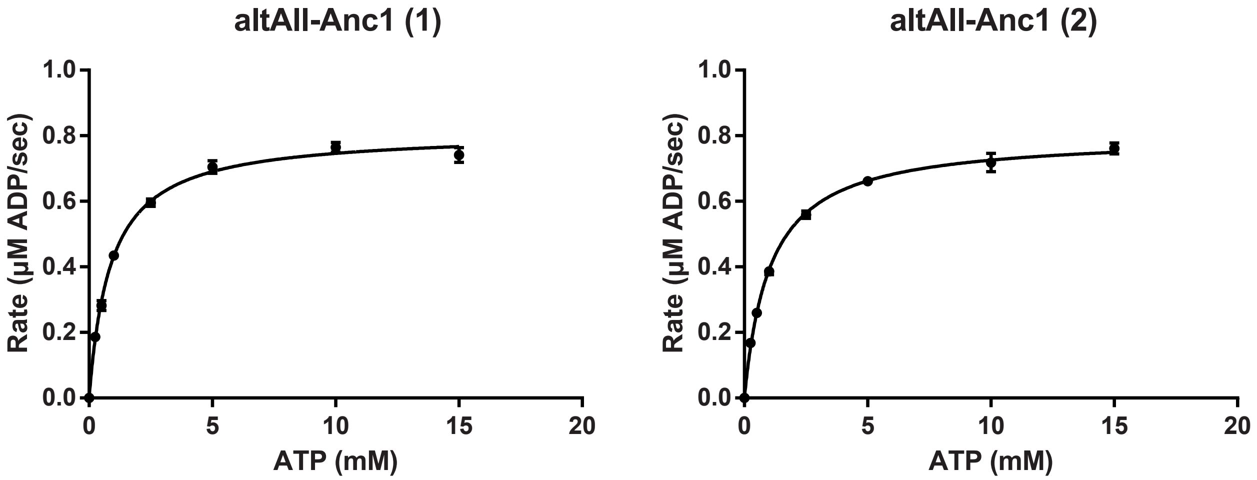


**S7.** Two biological replicates of steady-state kinetic assays of altAII-Anc1 with varying ATP concentrations at 1 mM glucose. Each data point represents the average of three technical replicates ± standard deviation. Data were fit to the Michaelis-Menten equation in GraphPad Prism.

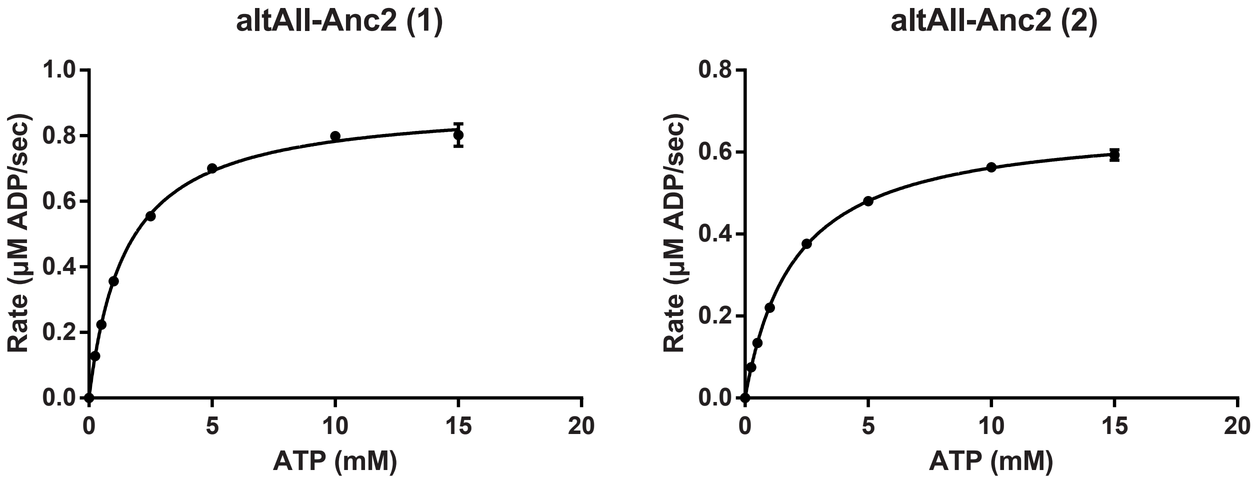


**S8.** Two biological replicates of steady-state kinetic assays of altAll-Anc2 with varying ATP concentrations at 2 mM glucose. Each data point represents the average of three technical replicates ± standard deviation. Data were fit to the Michaelis-Menten equation in GraphPad Prism.

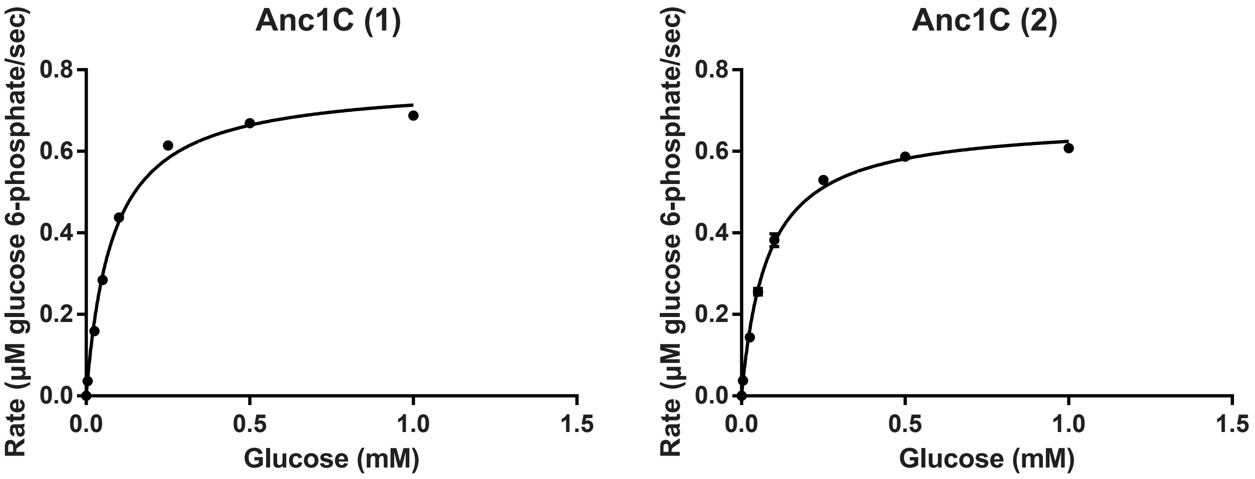


**S9.** Two biological replicates of steady-state kinetic assays of ML Anc1C with varying glucose concentrations at 15 mM ATP. Each data point represents the average of three technical replicates ± standard deviation. Data were fit to the Michaelis-Menten equation in GraphPad Prism.

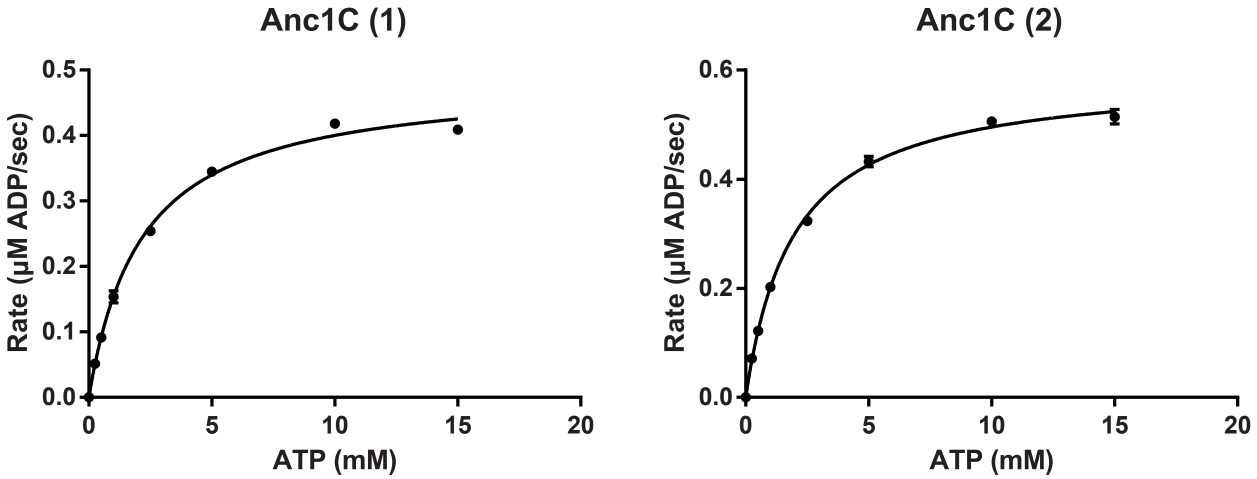


**S10.** Two biological replicates of steady-state kinetic assays of ML Anc1C with varying ATP concentrations at 1 mM glucose. Each data point represents the average of three technical replicates ± standard deviation. Data were fit to the Michaelis-Menten equation in GraphPad Prism.

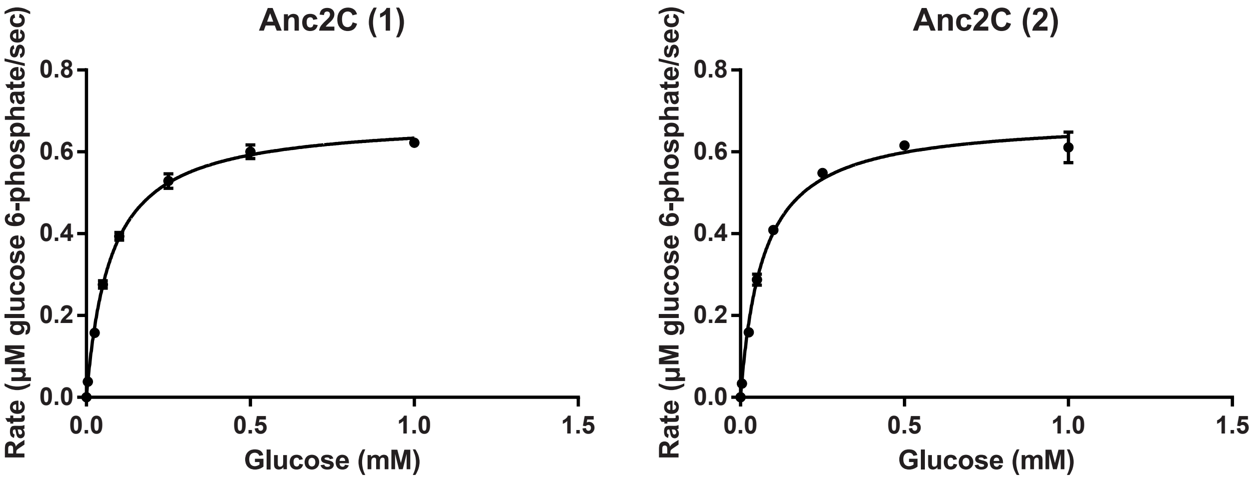


**S11.** Two biological replicates of steady-state kinetic assays of ML Anc2C with varying glucose concentrations at 15 mM ATP. Each data point represents the average of three technical replicates ± standard deviation. Data were fit to the Michaelis-Menten equation in GraphPad Prism.

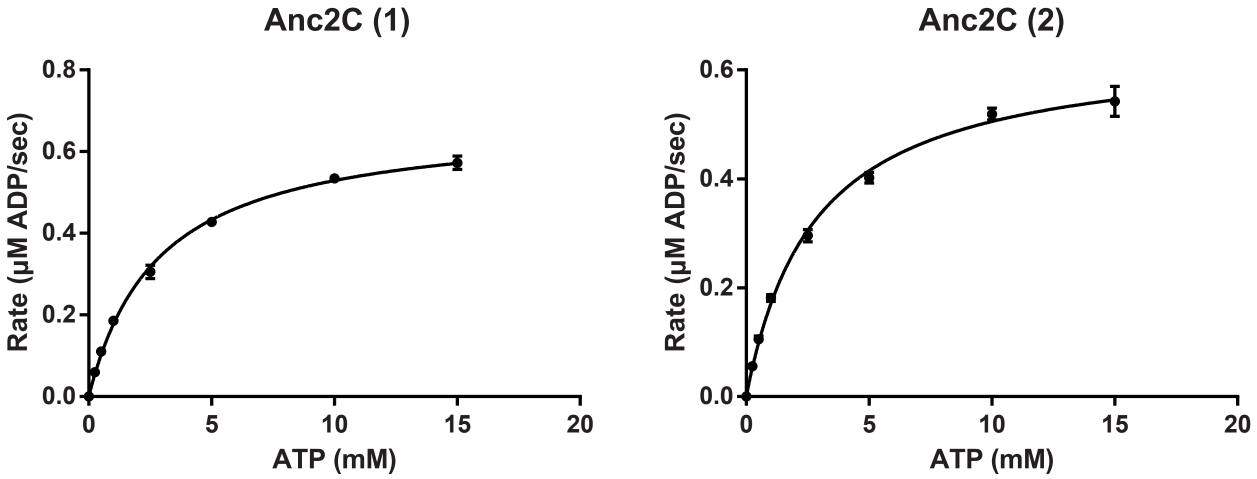


**S12.** Two biological replicates of steady-state kinetic assays of ML Anc2C with varying ATP concentrations at 1 mM glucose. Each data point represents the average of three technical replicates ± standard deviation. Data were fit to the Michaelis-Menten equation in GraphPad Prism 6.

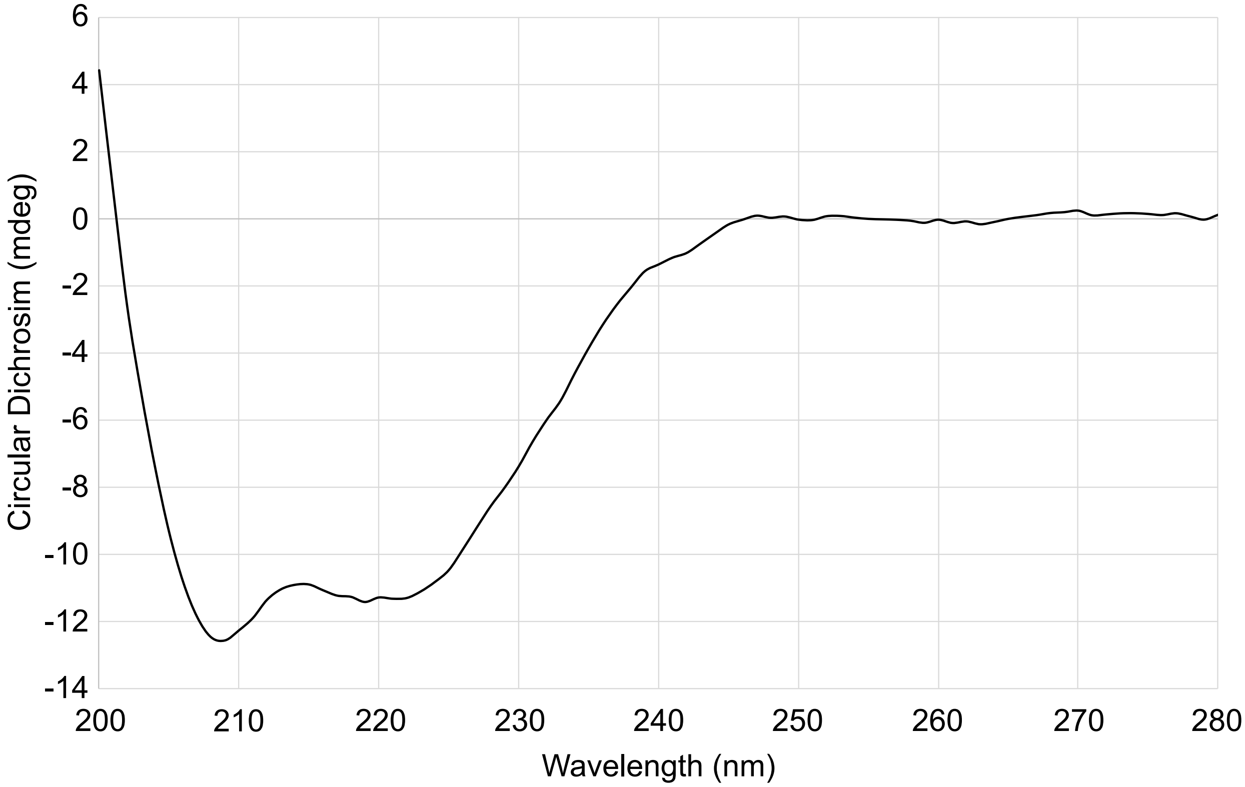


**S13.** Circular dichroism spectrum of truncated ML Anc1N.

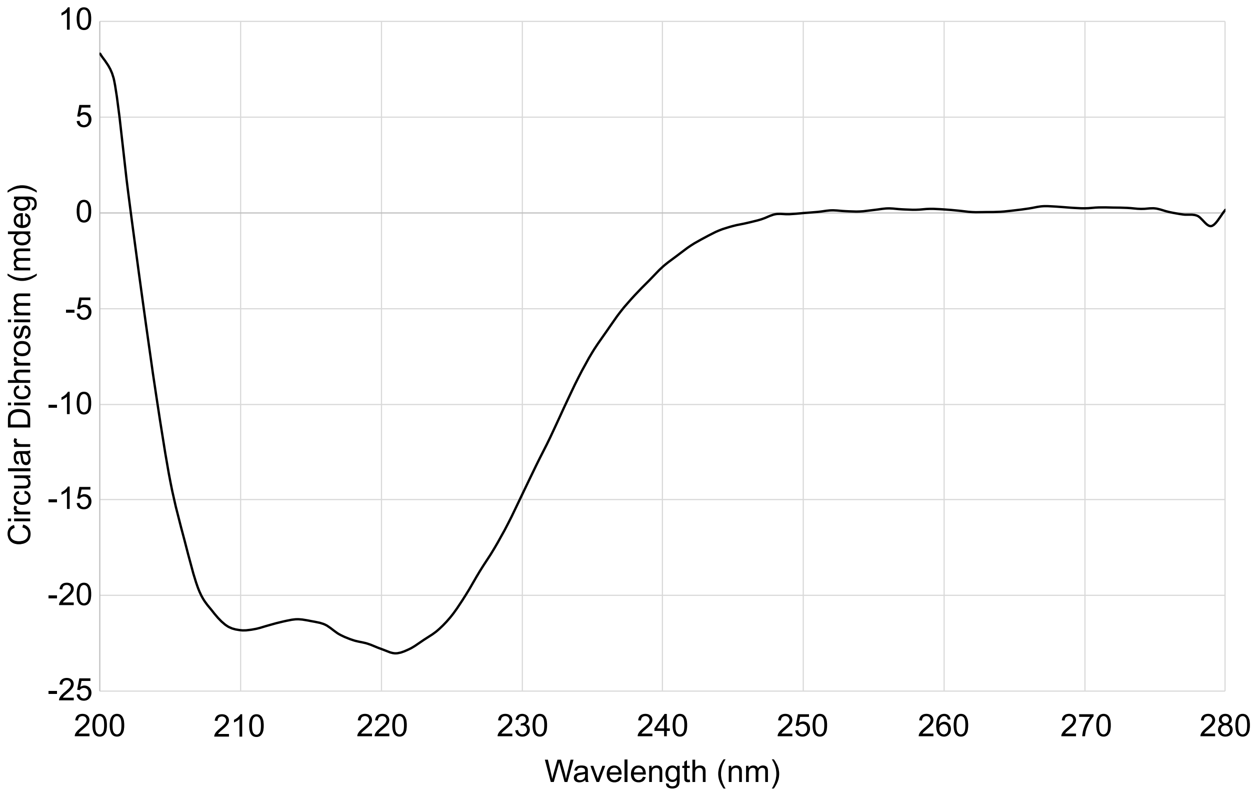


**S14.** Circular dichroism spectrum of ML Anc1C.

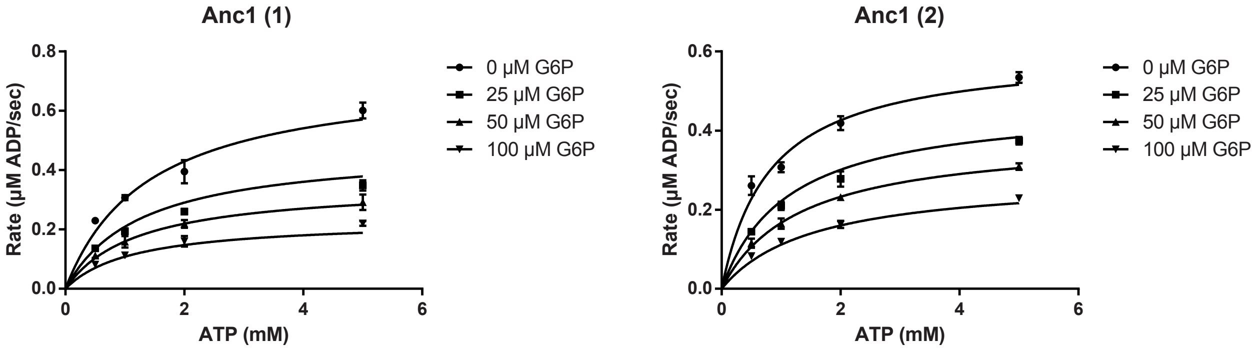


**S15.** Two biological replicates of steady-state kinetic assays of ML Anc1 with varying ATP and glucose 6-phosphate concentrations at 1 mM glucose. Each data point represents the average of three technical replicates ± standard deviation. Data were fit to the mixed non-competitive inhibition equation in GraphPad Prism.

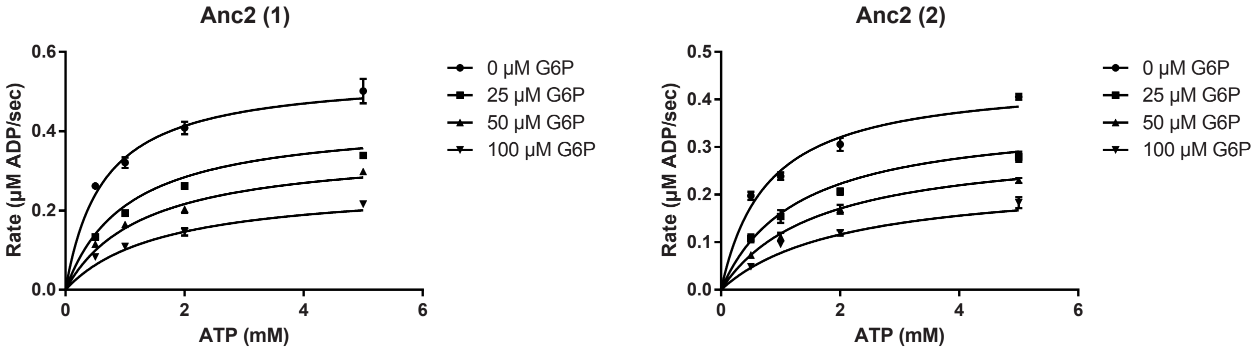


**S16.** Two biological replicates of steady-state kinetic assays of ML Anc2 with varying ATP and glucose 6-phosphate concentrations at 1 mM glucose. Each data point represents the average of three technical replicates ± standard deviation. Data were fit to the mixed non-competitive inhibition equation in GraphPad Prism.

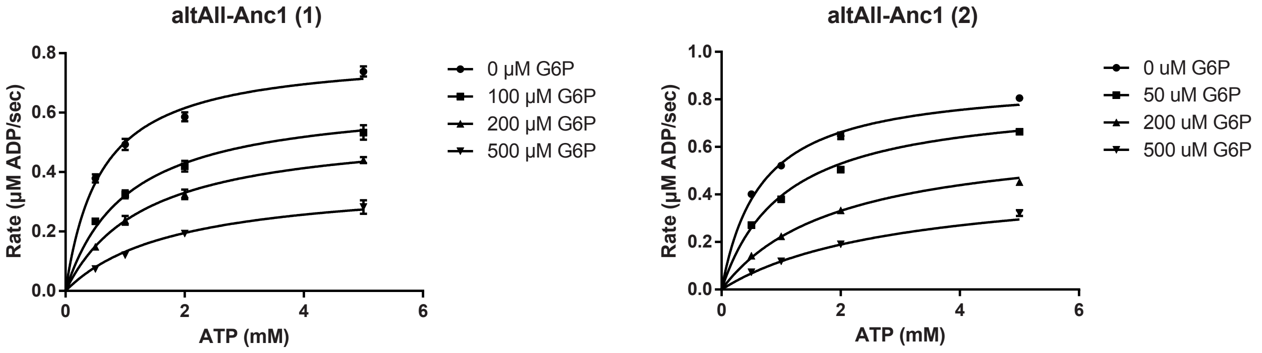


**S17.** Two biological replicates of steady-state kinetic assays of altAII-Anc1 with varying ATP and glucose 6-phosphate concentrations at 1 mM glucose. Each data point represents the average of three technical replicates ± standard deviation. Data were fit to the mixed non-competitive inhibition equation in GraphPad Prism.

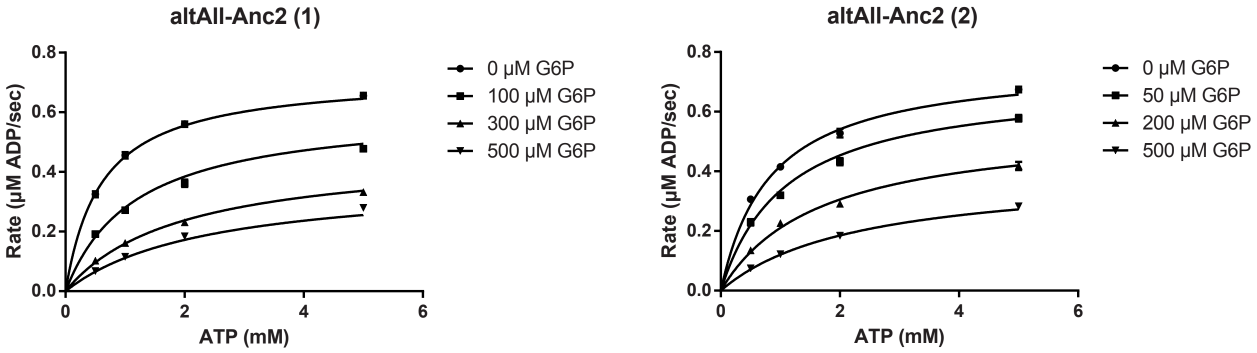


**S18.** Two biological replicates of steady-state kinetic assays of altAII-Anc2 with varying ATP and glucose 6-phosphate concentrations at 2 mM glucose. Each data point represents the average of three technical replicates ± standard deviation. Data were fit to the mixed non-competitive inhibition equation in GraphPad Prism.

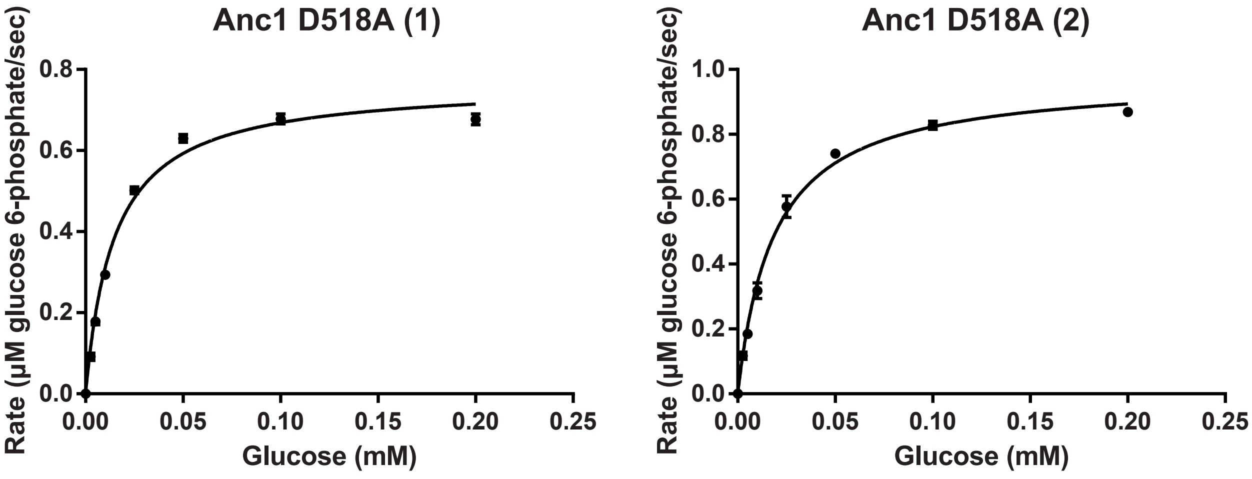


**S19.** Two biological replicates of steady-state kinetic assays of the ML Anc1 D518A variant with varying glucose concentrations at 50 mM ATP. Each data point represents the average of three technical replicates ± standard deviation. Data were fit to the Michaelis-Menten equation in GraphPad Prism.

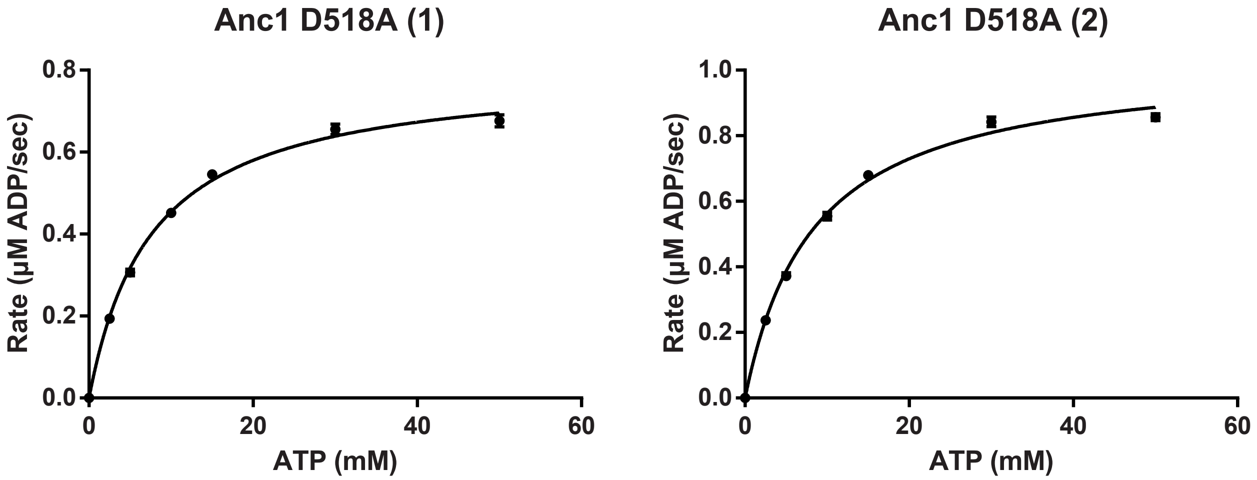


**S20.** Two biological replicates of steady-state kinetic assays of the ML Anc1 D518A variant with varying ATP concentrations at 0.2 mM glucose. Each data point represents the average of three technical replicates ± standard deviation. Data were fit to the Michaelis-Menten equation in GraphPad Prism.

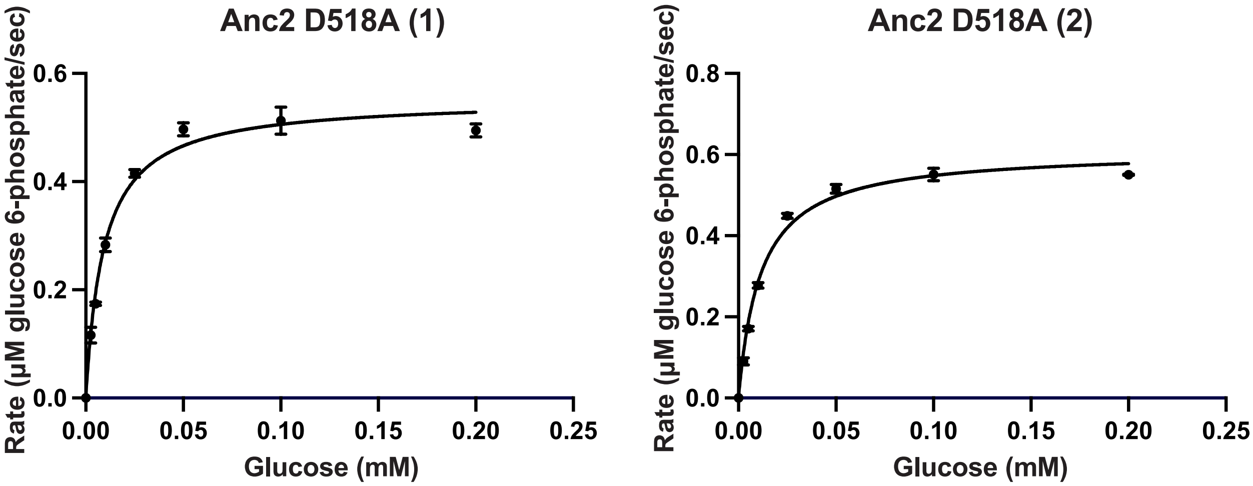


**S21.** Two biological replicates of steady-state kinetic assays of the ML Anc2 D518A variant with varying glucose concentrations at 50 mM ATP. Each data point represents the average of three technical replicates ± standard deviation. Data were fit to the Michaelis-Menten equation in GraphPad Prism.

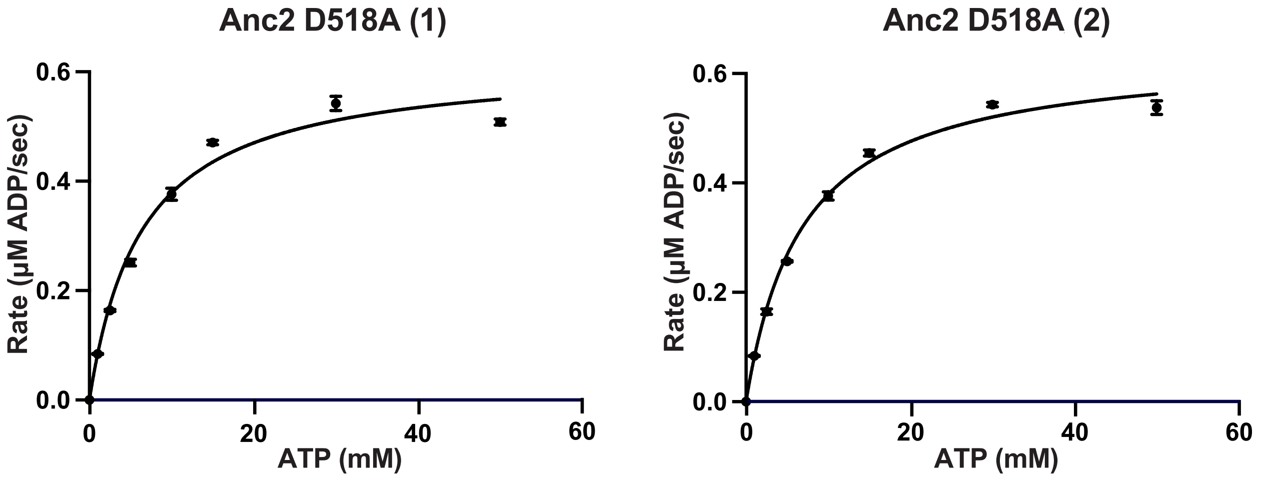


**S22.** Two biological replicates of steady-state kinetic assays of the ML Anc2 D518A variant with varying ATP concentrations at 0.2 mM glucose. Each data point represents the average of three technical replicates ± standard deviation. Data were fit to the Michaelis-Menten equation in GraphPad Prism.

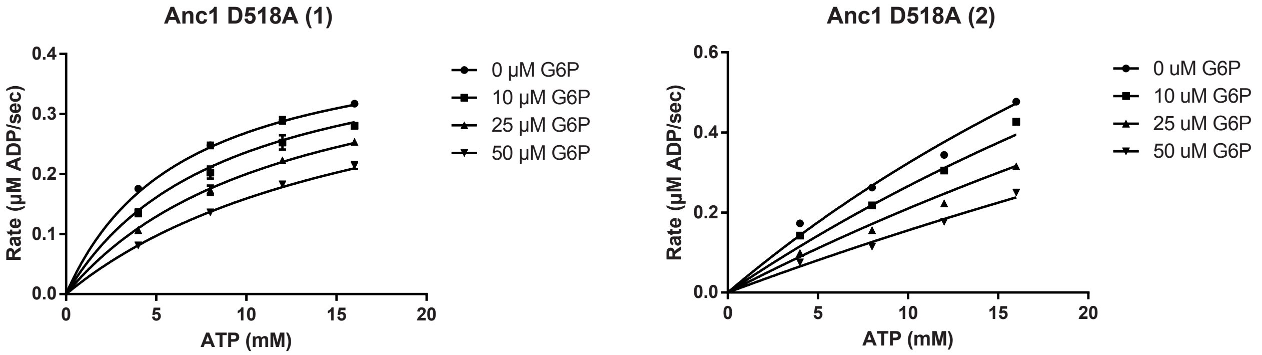


**S23.** Two biological replicates of steady-state kinetic assays of ML Anc1 D518A with varying ATP and glucose 6-phosphate concentrations at 0.25 mM glucose. Each data point represents the average of three technical replicates ± standard deviation. Data were fit to the competitive inhibition equation in GraphPad Prism.

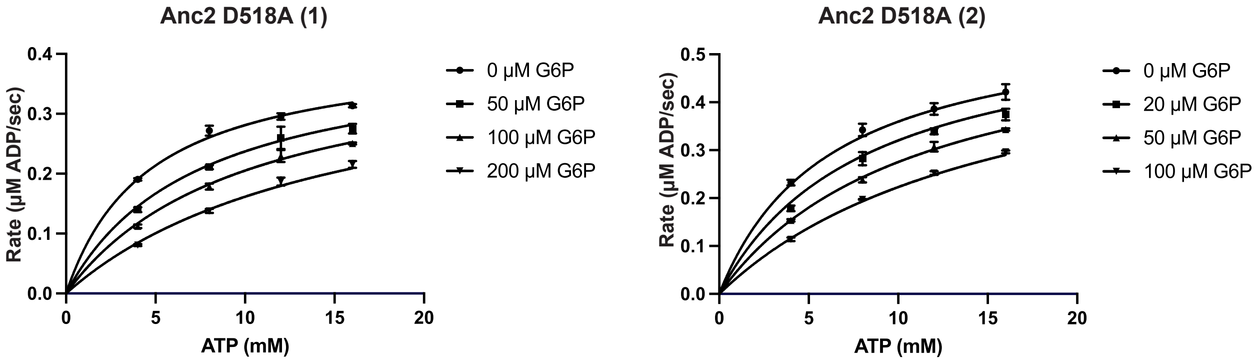


**S24.** Two biological replicates of steady-state kinetic assays of ML Anc2 D518A with varying ATP and glucose 6-phosphate concentrations at 0.25 mM glucose. Each data point represents the average of three technical replicates ± standard deviation. Data were fit to the competitive equation in GraphPad Prism.

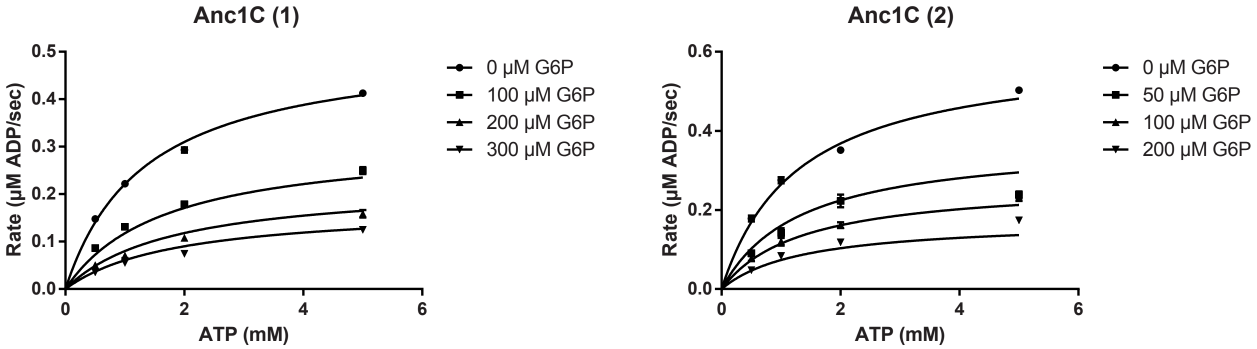


**S25.** Two biological replicates of steady-state kinetic assays of ML Anc1C with varying ATP and glucose 6-phosphate concentrations at 1 mM glucose. Each data point represents the average of three technical replicates ± standard deviation. Data were fit to the mixed non-competitive inhibition equation in GraphPad Prism.

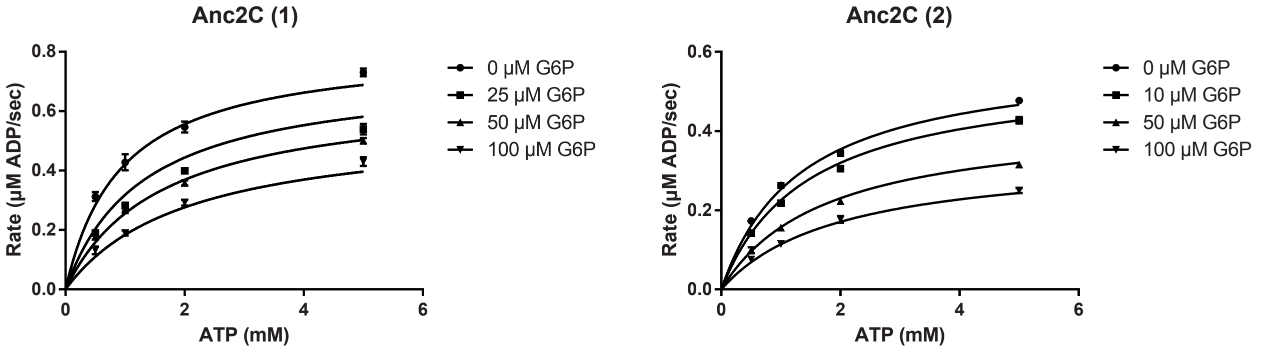


**S26.** Two biological replicates of steady-state kinetic assays of ML Anc2C with varying ATP and glucose 6-phosphate concentrations at 1 mM glucose. Each data point represents the average of three technical replicates ± standard deviation. Data were fit to the mixed non-competitive inhibition equation in GraphPad Prism.

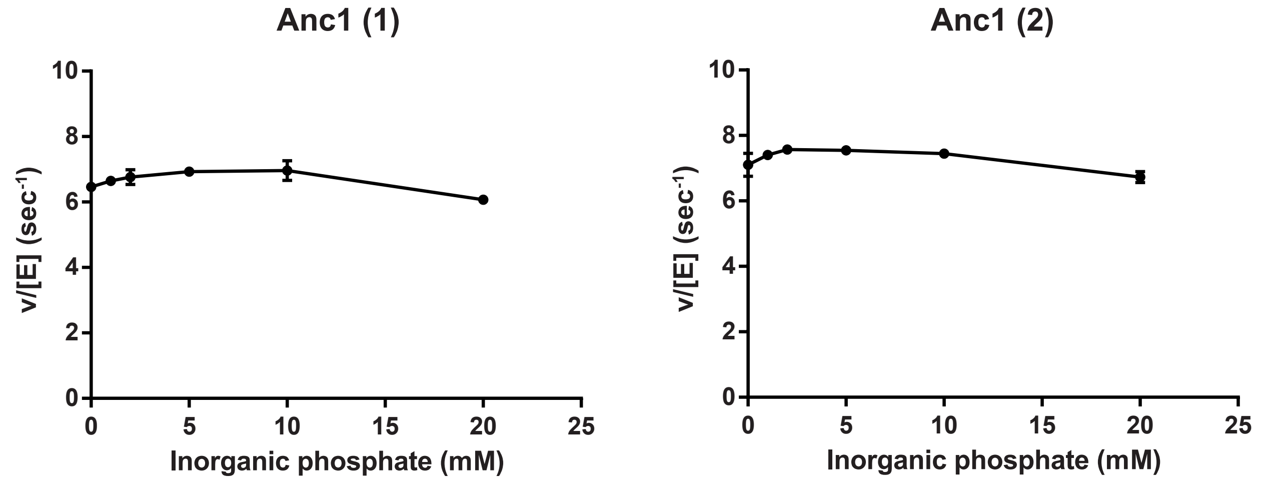


**S27.** Two biological replicates of ML Anc1 activity at varying concentrations of inorganic phosphate. Glucose, glucose 6-phosphate and ATP concentrations were held constant at 1 mM, 70 μM and 1.5 mM, respectively. Each data point represents the average of three technical replicates ± standard deviation.

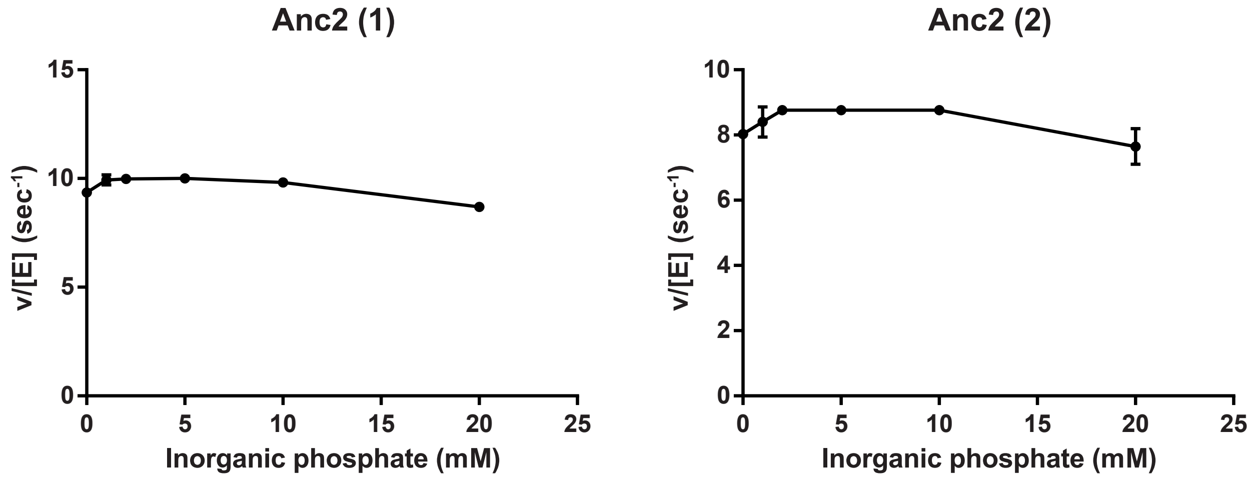


**S28.** Two biological replicates of ML Anc2 activity at varying concentrations of inorganic phosphate. Glucose, glucose 6-phosphate and ATP concentrations were held constant at 1 mM, 25 $\mu$M and 1.5 mM, respectively. Each data point represents the average of three technical replicates ± standard deviation.

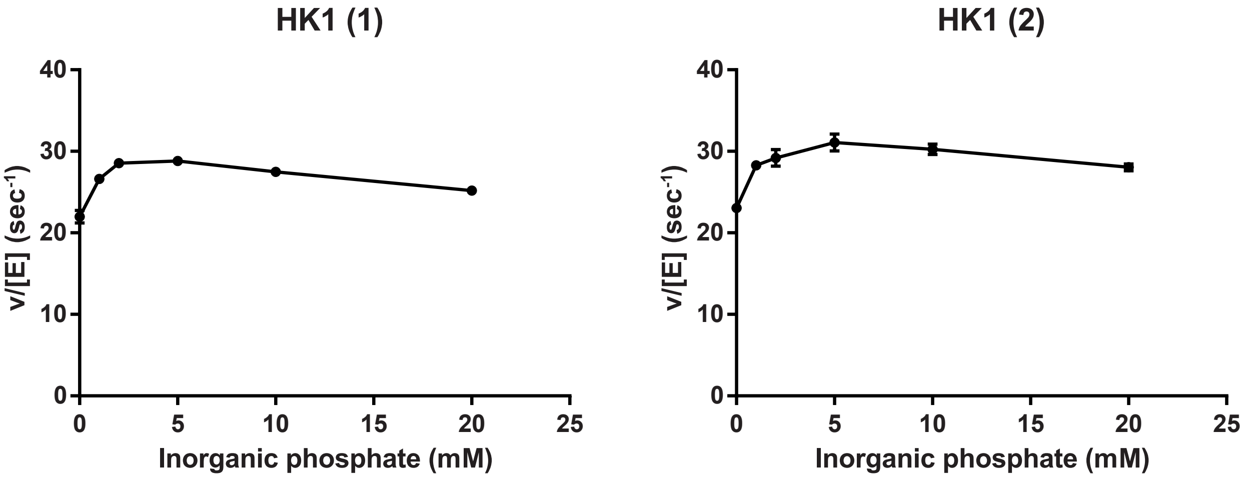


**S29.** Two biological replicates of human HK1 activity at varying concentrations of inorganic phosphate. Glucose, glucose 6-phosphate and ATP concentrations were held constant at 1 mM, 15 μM and 1.5 mM, respectively. Each data point represents the average of three technical replicates ± standard deviation.

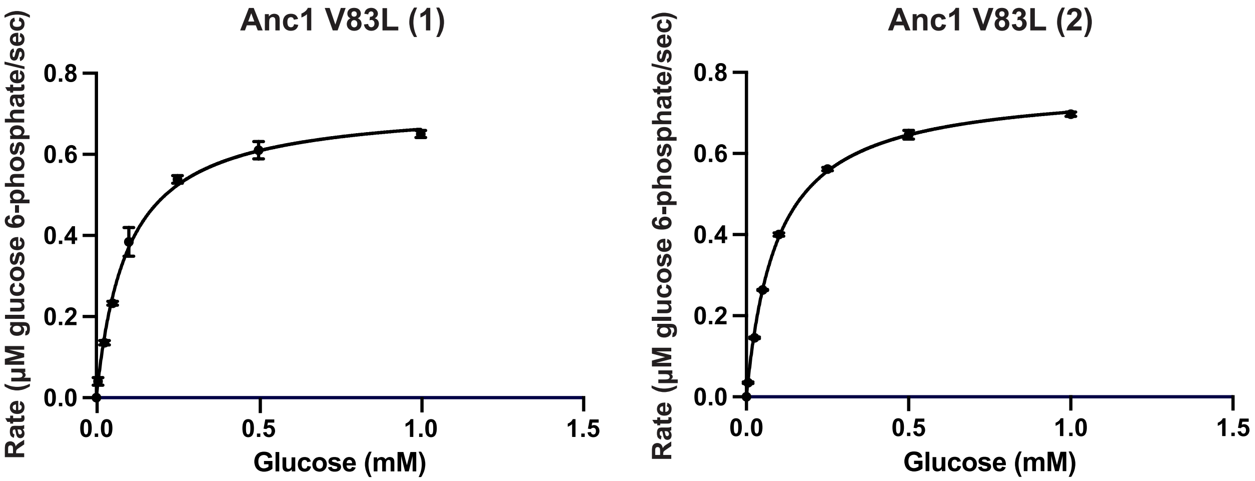


**S30.** Two biological replicates of steady-state kinetic assays of ML Anc1 V83L with varying glucose concentrations at 15 mM ATP. Each data point represents the average of three technical replicates ± standard deviation. Data were fit to the Michaelis-Menten equation in GraphPad Prism.

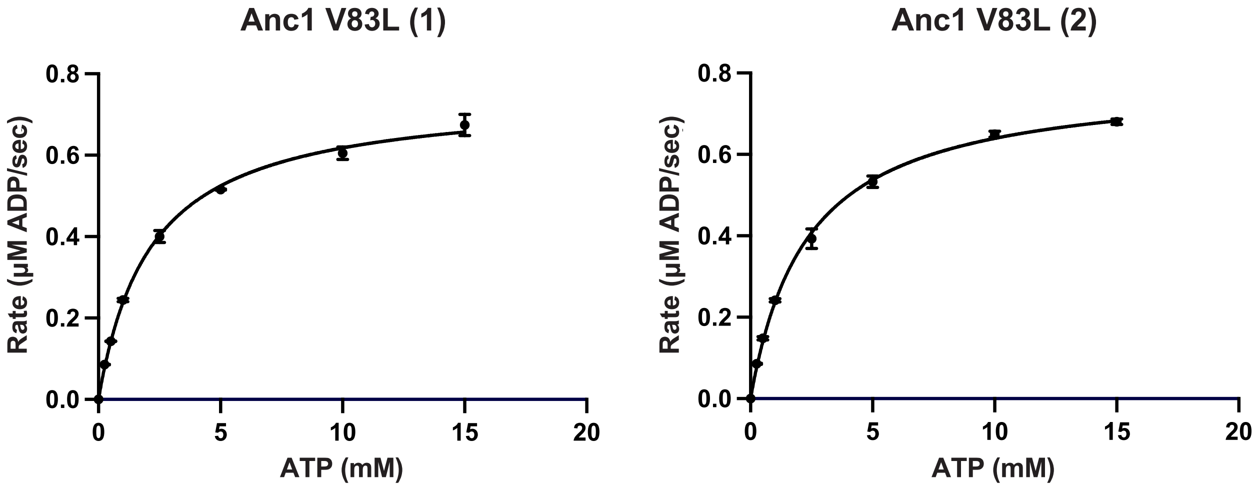


**S31.** Two biological replicates of steady-state kinetic assays of ML Anc1 V83L with varying ATP concentrations at 1 mM glucose. Each data point represents the average of three technical replicates ± standard deviation. Data were fit to the Michaelis-Menten equation in GraphPad Prism.
