## Supplementary phylogenetic material for "Phylogenetic Investigation of the 100 kDa Hexokinase Enzyme Family with the Topiary Ancestral Sequence Reconstruction Pipeline"

### TABLE OF CONTENTS

#### Supporting phylogenetic files

S1. Multiple sequence alignment of 100 kDa extant HKs used for ancestral sequence reconstruction ........ 1-85

S2. Phylogenetic files for the reconciled 100 kDa HK tree ........ 86-101

S3. Reconstructed maximum-likelihood and altAll sequences of HK ancestors ........ 102-226

**Supporting phylogenetic files**

**S1.** Multiple sequence alignment of 100 kDa extant HKs used for ancestral sequence reconstruction.

```
>XP_015676026|Protobothrops mucrosquamatus|HK3
ILPFYVTLEQLQSVKKLMVAALEKGL-KEKDKATSY--LPMLPTYVSALPDGSERGDFLGLELT--FDQV
KVLLLQLTDN---VD-EP--MVVKKKTFSFPEKILSGNGQQ---------------VFEFMAQCLSTFLG
GL-----NLS---KKYFPLGFSFPFSCDHIALN------------------------Q------------
-----------CKLIRWTKDFRWPGVEGKDVTKLLQTAINEV---------CK-----------------
--------------NYQIEVLAVVNDAVGDLMSCSSNVRPCQMGVMI-----------------------
-----D-----AGTNCCYMEDTQCIPGNVQETR--GYMCINTEWGSFGNNGELNGMLTEFDTLVDKESMD
QGKCRFEKMTSSYYVRETVRIILANLSENAELFSGVLTPTLLIKGKLALQDIVEVTD-------------
--DK--VGLAKTKNFLCR-LGLVASNQDCFHVQEICQAVYTRSAKLCAAGVAGVLTHIHKQQ-QLPEMKT
CVVMSGDMYKSH-SVYSEMLQQTLKELVPEYSLTFIPSDDGSTLGAAIVTAVEVKLRNHREGLAHVLAPL
KLSIVDLERLRNGIRQEMENGLSKAASKKS-CLHMLPTFVCYLPDGS-----------------------
---ERGDFLALDLGGTNFRVLMVQVKSQEEGGVHIINETYTIPPEITQSTGIQ-------LFDHIVNCIV
DFHIKHGMSEH-ILPLGFTFSFPCNQITLDKGILLRWTKGFSATDCVGEDVVNLLRDAVQRQK----TID
VDVIA-----LVNDTVGTMMSCAYLDHNCEVGLIVGTGCNACYMEEM---RNVGSME-------------
--SN------DG---------------------RMCINMEWGAFGDDGC-LDHYITPYDDIVDKESLNPN
QQR-FEKMISGMYLGEIVRYVLLELASRKILFRDHKLKGLQNKGIFPTKLLSDIEN--------------
------------DSESRQHVYHILEELGL-PVSPEDAKVVKEVCHTVTSRAAMMCAAGVAAVVEKIRENK
RMTKLEVTVGVDGTVYKMN-----------PYFAKKLQDTVALLAPNCNVKF---------LLSEDG---
----------SGKGAALIAAVACRTKI-------S-----------------
>XP_062979419|Elgaria multicarinata webbii|HK3
LQPFYLSKEQLLQVKNSMLAAMERGL-KQKVKETGS--LPMLNTYVCTLPNGTEEGDFLGLELS--QTQV
KVLLLQLTDD---ID-EP--EVVKRRTFPLTEKILIGSGQE---------------VFEFMAQCLATFLS
GL-----NIN---KKYFSLGLSFPFPCENSSLT------------------------Q------------
-----------CKLIRFTKDCRWPGVEGCDVTQMLQLAINQH---------CK-----------------
--------------NYQIEVLAVVNDTVGILLSCSSTKQPCEIGLLI-----------------------
-----D-----SGTNCCYMEEKQRI-AGMEESE--GHMCVNLEWGTLGNAGELDDLLTEFDLLMDKQTMD
PGKCRFEKFISSMYFCETVRVILASLAEKAELFNGVLTPPLLTKGKLEIQDILEIVH-------------
--EK--VGLAKAKSFLFR-LGLVASNQDCFHVQQICQAVFVRSANLCAAGLAAILTHIRATQ-GLSELKT
FVAVDGELYRSE-TQYGEILQQTLTSLIPECTIKFVLTEGGATLGAAQLAAVAVRLRNQRQGVSQALAPF
RLSMVDLERLRNLMRQEMEKGLSKETNCKA-SVRMLPTYVCNLPDGT-----------------------
---EKGDFLALDLGGTNFRVLVVQVQSKQEGGVHMISESYIIPPEIAQSTAEK-------LFDHIVSCIV
DFQTKNNMLGS-MLPLGFTFSFPCNQRGLDKGILLRWTKGFTASGAVGQDVVNLLREAVQRQQ----HVD
VDVVA-----LVNDTVGTMMACAYIDPKCEVGLIVGTGTNACYMEEM---RNIGTVE-------------
--GD------EG---------------------RMCINMEWGAFGDDGC-LDHYITSFDTIVDKNSLNPG
QQR-YEKLISGMYLGEIVRHILLDLISRKILFKGRESFVLQTPNIFPTKFLTQIED--------------
------------SSQGNRYVCDFLEDIGL-SVIPEDAEVVKDVCHAVSSRAAKVCAAGIAAVVEKIRENK
RMTHLEVTVGVDGTLYKMH-----------PYFSRKLQDTVPLLAPGCSVKF---------LLSEDG---
----------SGKGAALIAAVANRTK--------------------------
>XP_053573022|Bombina bombina|HK3
LQPLCLSQTQLAQLRDMIMDNIERGL-KKDPDRPST--LKMLPTYVRSTPDGTERGHFLVLDLG--GAKF
RVLQVKLDKD------RP--VEMESEIYAIPEDIMTGTGRQ---------------LFDHVAECLGRFLR
QR-----HLQ---QKALPLGFTFSFPCVQSQLD------------------------K------------
-----------SVLISWTKGFHCSGVEGEDVVQLLRDAIHRR----------D-----------------
--------------DCNVDVIAIVNDTVGTMMSCGYHDHSCEVGFIV-----------------------
-----G-----TGTNACYMEEMGNV-ETVEGDE--GKMCVNMEWGALGDDGELRPFQTDYDHQVDQNSFA
```

```
PGKQCFDKMISGMYLGEIVRLVLVRLASQGLLFGGAKTSALLTKGSFATKHVSAIED-------------
--DH--SGLAAAKQILTL-LGLKPSEQDCIQVQRVCSTVSTRSANLCAAGIAAVVTRIQRNR-HDNHSCV
TVGVDGSVYKLH-PKFSERLHYALKFLAPQCQIKFIVSEDGSGKGTAIVTAVAQRLAAQRQLINKVLAPL
ILSNEKLREVQKRMCVEMEMGLRKETQPKA-TVKMLPTYVRATPDGT-----------------------
---ERGDFLALDLGGTNFRVLYVHVGLKEEGGVQIKSKTFSLSEEVITGTGEE-------LFDHIVDCIT
DFQNETHLRGK-RLPLGFTFSFPCKQTSLDQGILISWTKGFSASGCEGEDVVRLLREAAVRKN----NSD
LFVVA-----LVNDTVGTMMSCGYDDPSCEIGLIVGTGSNACYMEDM---KNVEVLE-------------
--GD------EG---------------------QMCINMEWGAFGDNGC-LDDITTSFDHNVDQFSMNPR
KQK-YEKMISGMYLGEIVRQILIVLTENGLLFGGKISQRLQTRNIFPTKFLSDIEK--------------
-------------DTLGLIQVRSILTDLGL-RSTCDDAVVVKEVCTTVSRRAQFCAAGIAAVVEKIRANR
GLDHLDVTVGVDGTLYKLH-----------PHFAGVVQETVKALAPQCEVTF---------LRSDDG---
----------SGKGAALITAVACRIAG--SEA--RAA---------------
> XP_063784001|Pseudophryne corroboree|HK3
LEPFLLSQAQLEAVRDLMSVNMERGL-KAKKKRPST--LKMLPMYVRSTPDGTERGVFLALDLG--GANI
RIIKVKLDGS------RG--VEMDSEVYDVPEAVVTGAGKQ---------------LFDHVAECLGLFLE
KQ-----NLK---DEVLPLGFTFSFPCKQTELD------------------------Q------------
-----------SVLISWTKGFRCSGVEGRDVVQLLKDAIQRC----------G-----------------
--------------DYKVDVLAIVNDTVGTMMSCGYEDHSCEVGFII-----------------------
-----D-----SGTNVCYMEEMGNV-ESVEGDE--GTMCISMEWGALGDDGALT-IETSYDAQVDQASRL
PGKQRFEKMTSGMYMGEIVRYVLLELTNKGLLFGGFKTSQLDTEGSVGTKEVLEIEN-------------
--DP--FGLDKAKSMLTS-LGLNPSDQDCRAVKSVCNAVLMRSAKLCAAGLAAVANRIKNNR-QDPVGRI
TVGVDGYVYKTN-PKFGAYLNKALEELAPSCQIKFLVSEDGSGKGTAIVTAVAQRRVEQRQCINEILAPF
IMSESQLKEVQRLMRKEMDFGLEGGAR--A-TVKMLPTYVSATPDGT-----------------------
---ERGEFLALDLGGTNFRVLYTRVGIKNEGGIQMVSKTFTLSTQVIQGTGEE-------LFDHIVDCIT
EFQKENGLKGK-RLPLGFTFSFPCQQTSLDKGILISWTKGFSASGCVGEDVIKLLREAAIRKK----NPD
FHVVA-----LVNDTVGTMMSCGYDDAACEIGLIVGTGTNACYMEEM---RNVKVLE-------------
--GD------EG---------------------QMCINMEWGAFGDNGC-LHDILTSYDTDVDQFSINSG
RQR-FEKMISGMYLGEIVRQVLIVLTQRGILFGGKISEQLNTRNLFPTKFLSDIER--------------
-------------DTLGLLEVRAILTGLGL-ESTCDDTVVVKEVCIKVSRRAQLCAAGIAAVVEKIRENR
GLDHLQITVGVDGTLYKQH-----------PHFAGEVQKAVKMLAPQCDVTF---------HISEDG---
----------SGKGAALITAVACRVAG--AEA--RAA---------------
> XP_018407907|Nanorana parkeri|HK3
LQPFLLSIGQLEKVRDLTLDNMERGL-KAKPRTQST--LKMLPTYVRSTPDGTELGDFLVLDLG--GAKF
RILKVKLSCA---SG-----VEMDSEVFDISEDIMTGTGEQ---------------LFDHVAECLGHFLQ
HR-----NLS---KEVFQLGFTFSFPCEQSQLD------------------------K------------
-----------SVLISWTKGFRCSGVEGKDVVQLLKDAIQRR----------G-----------------
--------------DYKVDVVAIVNDTVGTMMSCGYKDHSCEVGFIV-----------------------
-----G-----TGTNVCYMEEMGNV-EAVEGDE--GTMCINLEWGSLGDDGALNDILTQYDLQVDQTSTV
PGKQRFDKMISGLYMGEIVRHVLAKLTNRGLLFGGVSTQQLLDKGTFGTKDVSDIEN-------------
--DN--FPFEKTKNVLTA-LGLKPSEKDCVQVNCVCSAVSMRSARLCAAGLAAVANRIKNNR-RHQLGSI
TVGVDGSVYKTN-PRFGEHLNQALKELAPGCQIKFLVSEDGSGKGTAIVTAVAQRLASQRKHIDKILAPF
LMSHEKLIEVQSRMRNEMEIGLHKQTQPEA-TVKMLPTYVRATPDGT-----------------------
---EEGEFIALDLGGTNFRVLCINVGLKNESGVQMKSKTFTLPTEVIQGTGEA-------LFDHIVNCIT
EFQKENGLQGK-KLPLGFTFSFPCKQTSLDQGILISWTKGFSASGCVGEDVVRLLREAANRNN----NHD
FRVVA-----LVNDTVGTMMSCGYDDTACEIGLIVGTGTNACYMEEM---RNVEVLE-------------
--GH------EG---------------------KMCINMEWGAFGDNGY-LDDITTSFDSNVDKFSINPG
KQR-YEKMISGMYLGEIVRQVLIELTKDRILFRGKISEQLNTRDLFPTKYLSQIES--------------
------------DSLGLLQVRSILTKLGL-NSTCDDTVIVKEVCTIVSKRAAQLCAAGIAAVVEKMRGSR
GLDHLKITVGVDGTLYKLH-----------PHFAGVLQKTVKTLAPQCDVSF---------HLSEDG---
----------SGKGAGLITAVACRVAD--AEP--RAA---------------
> XP_040200337|Rana temporaria|HK3
LQPFSLSIGQLEKVRDLMLGNMERGL-QVNPEKQST--LKMLPTYVRSTPDGTECGDFMVLDLG--GSKF
```

RILKVKLSRA---SG-----VEMDNEEFDISEEIRTGTGER---------------LFDHVADCLGRFLQ
HR-----NLS---ESVFPLGFTFSFPCVQSQLD-----------------------K------------
-----------SVLISWTKGFCCSGVEGKDVVQLLKDAIQRR----------G-----------------
--------------DYKVDVIAIVNDTVGTMMSCGYKDHSCEVGFIV-----------------------
-----G-----TGTNVCYMEEMGNV-EALEGDE--GTMCINIEWGGFGDDGTLNDIVTEYDSQVDQTSRV
PGRQRFGKMIGGMYMGEIVRHVLVKLTNQGLLF-GVSALKLLNKATFGMKDVSDIEN------------
--DN--FPFEKTKSVLAV-FGLKPNEEDCVRVKCVCCAVSLRSARLCAAGLSAVANRIKNNR-PHHLGRV
TVGVAGSVYKTN-PKFGEHLNEALKELAPGCQIKFLVSEDGSGKGTAIVTAVAQRLATQRKHINEILAPF
LMSLAKLKEVQSRMRNEMEIGLHKQTQPGA-TVKMLPTYVRATPDGT-----------------------
---EVGEFIALDLGGTNFRVLCVNVGLKNEGGVQMKSKTFTLPTEVIQGTGEG-------LFDHIVDCIT
EFQKENGLLGK-KLPLGFTFSFPCKQTSLDQGILISWTKGFSASGCVGEDVVRLLREAANRKN----NHD
FRVVA-----LVNDTVGTMMSCGYDDTACEIGLIVGTGTNACYMEEM---RNVEVLE------------
--GG------EG---------------------RMCINMEWGAFGDNGC-LDDITTSFDNDVDKFSINPG
KQR-YEKMISGMYLGEIVRQVLIELTKDRILFRGKISEQLNTRGLFPTKYLSQIES-------------
------------DSLGLLQVRSILTKLGL-NSTCDDTLIVKEVCTVVAKRAAQVCASGIAAVVEKMRENR
GLDHLKITVGVDGTLYKLH-----------PHFAGVLQKTVKLLAPQCDVSF---------HQSEDG---
----------SGKGAGLITAVACRVSD--AEP--RAA---------------
>XP_043925231|Protopterus annectens|HK3
LSLFVLSDEKLQKIVSLMLENMEKGL-RKQKHTTST--LKMLPTFVHSTLDGTERGDFLALDLR--GTKF
RVLQVKLTDD------DS--YKVQSESFEIPEDIMTGTGIQ---------------LFDYFAKHLSSFLD
KQ-----QLK---DKKLPLGFTFPFPCKQIDLD-----------------------K------------
-----------SVLLTWTKGFKCSGVEGEDVVQMLRDAIHRQ----------L-----------------
--------------DCDVNVIAVVNDTVGTMMSCGYKDRACEIGLVV-----------------------
-----G-----TGTNACYMEEMGNI-EQIEGDE--GRMCVIMEWGAFGDDGCLRDIQTEFDLELDRHSRN
PGRQTFEKKISGMYMGEIVRLIVVKLASLHLLFNGVTSPALLKSGTFETNYISEIED------------
--EK--VGLTKAKEILLS-FNLNPSEQDCVIVQKICNNVSTRSANLCAAGLAAVVTRIQRNR-HLKHLQT
TVGVDGTVYRKH-PKFSGRLQATLKQLTSNCDVQFMLSEDGTGKGTAMVTAVARRLFAQRQQINETMAPF
CLSRKQLVQVQQRLRLEMLRGLEKETQSTA-TVKMLPTYVRGTPDGT-----------------------
---ERGHFLALDLGGTNFRVLLVNVNSREEGGVKMVNQIYSIPESLMQGSGTK-------LFDHIVDCII
DFVKKQGKMGC-RLPLGFTFSFPCRQTGLDKGILITWTKGFNATECEGKDVVMLLREAIKRKQ----GVE
LDVVA-----IVNDTVGTMMSCAYNDSNCEIGLIVGTGSNACYMEEM---SNIKNVE------------
--GD------IG---------------------QMCINMEWGAFGDNGC-LDVFVTQYDQIVDKASINPG
KQR-YEKMMSGMYLGEIVRNILIDLTKQGIVFRGQISERLNTKGIFETKFLSEIES-------------
------------DSLVLLQVRAILQKLGL-DVTCDDTLIVKEVCTIVSRRAAQLCGAGIAAVVEKIRENR
RLNHLAITVGVDGTLYKLH-----------PHFSKIVHETVAELAPCCDVTF---------HQSEDG---
----------SGKGAALITAVACRLRE--TGQ--H-----------------
>XP_048407507|Stegostoma fasciatum|HKDC1
---MQLNKDQIEDIMVRFKAEMLKGL---DFPSSSC--LKMLPTHVRSTPDGSEKGEFLALDIG--GSMF
RVLRVKVSED---GR-QM--VHISSNVYPMTEDMKHENSRQ---------------FFDYIAECLADFIE
KQ-----QLK---DRKLPLGFTFSFPCRQSKLD-----------------------E------------
-----------GILISWSKGFFVKGVQGVDVVSLLREAISRR----------R-----------------
--------------DYEVDVLALVNDTVGTMMTCAFDDPYCEVGVVI-----------------------
-----G-----TGTNACYMEELRHI-QLVEGDE--GRMCINTEWGGFGDDGSLEDIRTVFDREVDDGSIN
PGKQLFEKMASAMYMGELVRIILVRIVRNGILFKGKTSSKLLTKGSIDIEDIIAIQN------------
--YS--TGLESAIKMLKN-LELEPTEEDGIAVKQICKIVSVRSATLCSATLAVILQHIKKNK-KIKRLRT
TVGITGTLYRQN-TQYAKYFHKMVRSLVPDCDVRFFLAEYGSGKGAAMVTAVAQRLVHQRNYIDKTLAPL
RLNCDQLLDVMDKMRSNMEQGLKTDTHNTA-TVKMLPTFICKFPTGN-----------------------
---ESGKFIALDLGGTNLRILLVEMQSGMRKSLRMYSKIFPIPLEIMQGTGEE-------LFDHIVECIA
NFLEYMGVKGT-RLSLGFTFSFPCVQTRLDQGILINWTKGFSATGVEGQDVVSLLKEAITRNG----ELK
LDIVA-----LLNDTVGTMMSCAYDHPNCEIGMIAGTGCNLCYMEEM---RNIETVK------------
--GK------EG---------------------RMCINTEWGAFGDDGC-LDNIRTTFDKLVDINSLNMG
HQK-FEKMISGMYLGEIVRNILINFTKQGLLFRGLMSQSLMTNGIFQTKYLSQIES-------------

------------DSLALLQIRTILQKLGL-DSSCDDSIIVKEVCGVVSTRAAKICGAGLAAVAEKIRQSR
HLDYLEVTVGVDGTLYKLH-----------PHFSTILENTVKDLAPNCDMTF---------VLSEDG---
----------SGKGAALVTAAAIQRDK--DSN--AQSNR-------------
> XP_059803278|Dasyatis sabina|HKDC1
LYNMQLNPEKLQDIMSRFGNEMRKGL---SINETST--VKMLPTFVQSTPDRTEKGDFLALDLG--GSRF
EVLRVRVSED---GQ-QV--IQTERQLYPMPDEMKQSNASE---------------FFNYVAECLGDFME
KR-----EIK---NRKFPLGFTFSFPCRQTKLD------------------------E------------
-----------GFLISWGKGYKISGSKGANVISLLRKGIEQQ----------G-----------------
--------------DFDVDVLALVNDTVGTLMSGGFDDPYCEVGVVI-----------------------
-----G-----TGTNACYMEEMSNI-ESVEGNI--GRMCINTEWGAFGDDGSLNDIRTEFDLELDANSLN
PGKQLFEKMISAMYLGELVRLILVRLTKSHTLFSGTASAKLLTKGSFGIQHLTAVVN-------------
--SN--SGMENSMQILMD-LDLQPSEANCKALQHICFIVMSRSISLCAAALATILIHLKNTR-KIKRLRT
TVGIDGTVYRTV-PQYPKNLHRLVRQLAPGCDVRFLLVDTGTGKGAAMVTAVAYRLVNQRKYQDKTLAPL
QLSREQLLKIMQIMKNEMKLGLKTETHSTA-TLKMLPTFIQKFPDGT-----------------------
---ERGKFLALDLGGTNFRVLLLVLRNRPRMISQTYQKSYSVPMEVMQGIGEE-------LFDWIVECIV
EFLDYMGMKCA-PLPLGFTFSFPCIQHRLDQGVLVNWTKGFKATDVEGDDVVFLLKEAIKRQG----ELN
LDIVA-----LLNDTVGTMMTCAYENPNCEVGLIAGTGTNACYMEEM---KNIEIVD-------------
--GV------EG---------------------RMCINTEWGAFGDNGC-LDDIRTTFDRLVDINSLNPG
KQK-YEKMISGMYLGEIVRNILINFTKHGLLFRGFVSPVLKTSGIFETSYMSQIES--------------
------------DRLGLLQVHSILQNLGL-DCTCDDSIIVKEVCSVVSKRSAQLCGAGMAAVVDKIRENR
GLNHLEITVGVDGTLYKLH-----------PHFSTIMQETVKDLAPSCDVKF---------VLSEDG---
----------SGKGAALVTAAVKQSN--------------------------
> XP_051883560|Pristis pectinata|HKDC1
LYNMHLNSEQLQDIMAQFDNEMRKGL---STIETST--VKMLPTFVRSTPDGTEKGDFLALDLG--GSRF
EVLRVKVSED---GR-QV--IHTERQTYPMSDEMKQGNAPE---------------FFNYVAECLGDFME
KQ-----EIK---GKKLPLGFTFSFPCRQTRLD------------------------E------------
-----------AFLISWGKGFKIKGTKGANVISLLRKGIEQQ----------G-----------------
--------------DFDVDVLALVNDTVGTMMSGGFDDPYCEVGVIV-----------------------
-----G-----TGTNACYMEELSHI-ESVEGDI--GRMCINTEWGAFGDDGSLEDIRTEFDRELDSGSLN
PGKQLFEKMISAMYLGEIVRLILVKIAQNGMLFKGNTSLKLLTERSFGIQHVSAIMN-------------
--HK--SGLENAMQILTD-LDLNPSEEDCLAVQHICTIVLSRSVSLCAATLATILIRLKRNK-KVKRLRT
TVGIDGTVYRTV-PQYSKNLHRMVRRLAPECDVRFILVETGTGKGAAMVTAVAYRLVHQRKYQEKTLAPL
KLTREQLLKVMKMMRSEMEQGLKKETHSTA-TVKMLPTFIQKFPDGT-----------------------
---ERGKFLALDLGGTNLRVLLLVLQRRPRRSTQVYQKSYRIPMEIMQGTGEE-------LFDYIAECIV
EFLDYMGMKCA-QLPLGFTFSFPCVQHQLDQGFLVNWTKGFKATDVEGNDVVTLLKEAIKRHG----ELD
LEIVA-----LLNDTVGTMMTCAYENPNCEVGMIAGTGSNACYMEEM---KNIEIVE-------------
--GD------EG---------------------RMCINTEWGAFGDNGC-LDDIRTTFDRLVDINSLNPG
KQK-YEKMISGMYLGEIVRNILINFTKHGLLFRGLVSPVLKTVGIFETSYISQIES--------------
------------DRLALLQVRTILQNLGL-DSTCDDSIIVKEVCGVVSERAAQLCGAGMAAVVDKIRENR
GLDHLEITVAVDGTLYKLH-----------PHFATILKETVKDLAPNCDVTF---------MLSEDG---
----------SGKGAALVTAAVKRPD--------------------------
> XP_061404494|Lethenteron reissneri|HK1
LYEMKLSDERLRAIMEQMKLEMERGL-DKEARSRAP--VKMLPTFVRSTPDGSEKGDFLALDLG--GSSF
RILRVRV------ET-QT--IDMEDEVYEVPDTVIRGTGPQ---------------LFDHVAACLSSFME
KL-----KIK---DKKLPLGFTFSFPCRHNVNG------------------------E------------
-----------SVLLRWTKRFHVSGVEGQNVVKMLRDAIQRR----------G-----------------
--------------DFEIDIVAVVNDTVGTMMTCGYDDHRCEIGLIV-----------------------
-----G-----TGSNCCYMEDLRNI-EQVEGDE--GRMCINMEWGAFGDDGSLDDIRTEYDLEVDAGSLN
PGQQIFEKMISGLYLGELVRLILVKMTTQALLFNGKATPELLTRGHFQTQFITAIDTR------------
--NK--EGVLKARELLSDHFSLHPSEEDCAALQQICAIVSTRSAYLVAAALGAVVSRLRLNK-AVKKLQT
TVGVDGTVYKTH-PQYSKRLQLMMRRLAPESRVQFLLSECGSSKGSAMVTAVAYRLAEQRKLIDATLAPL
KLSLTQLLDVKRRMRQEMDRGLARDSHATA-SVKMLPSYVRSTPDGS-----------------------

---ENGDFLSLDLGGTNFRVLLVKIRSSKKRSVEMHNKIYPIPAEIMQGSAED-------LFDHIVGCIS
DFLDYMGMAGW-RLPLGFTFSFPCRQTSLDVGFLVNWTKGFKASGCEGKDVVMLLREAIQRRN----EFE
LDVVA-----VVNDTVGTMMTCAYEDPKCEIGLIVGTGSNCCYMEEM---RNVELVE-------------
--GD------SG---------------------RMCINMEWGAFGDSGV-IDDIRTEYDQLVDAGSLNPG
KQR-FEKLMSGMYLGEIVRNILMDFTKHGLLFRGKISERLKTRGIFETRFLSQIES--------------
------------DRLALLQVRNILHELGL-ESTCDDSIIVKEVCGVVSRRAAQMCGAGMAAIVDKIRENR
GLEHLKVTVGVDGSMYKLH-----------PHFARFLHDTVKQLSPKCEVKF---------LQSEDG---
----------SGKGAALITAVACRFRN--TSE--S-----------------
>XP_040218119|Rana temporaria|HKDC1
LYHMRLTDETLLDIMARFQVELEKGL-KKDTNSTAS--LKMLPTYVRSTPDGSERGDFLALDLG--GSMF
RVLKVKVSED---GK-QK--VHMDSQFYPTPSDIATGNKTE---------------LFEYVVDCLGDFIK
KR-----QIT----ETLPLGFTFSFPCQQTKLD------------------------E------------
-----------GVLINWTKEFKVKGMQGMDVVKTLRKTLKRK----------K-----------------
--------------IPNVEVMAIVNDTVGTMMTCGFEDQLCEVGVIV-----------------------
-----G-----TGTNACYMENLSSI-DLIEGDE--GRMCINTEWGAFGDDGSLEDIRTEFDKEIDLGSVD
PGKQLFEKMISGLYMGELVRLILLEMAKKGLLFGGKISTALRTKGAIETSHVASMEN-------------
--NQ---GLYNTKQVLLD-LGLEPSDDDCIAVQHVATIVSFRSANLCAAALAAILTRLRQSK-KLDRMRT
TVGMDGTVYKTH-PQYAKRLHKVVRRLVPNCDVRFLLSECGSGRGAAMVTAVACRLLTQRKEIDEVLDVF
RASDEKLADIKQRMLTELKCGLQKETHDKA-TVKMLPTYVYATPDGT-----------------------
---ENGNFLALDLGGTNFRVLLVKIRSGRRRYVKMYNKIFAIPLEIMQGTGEE-------LFDHIVECIA
EFLDYMGIKGA-PLPLGFTFSFPCKQDRIDKGTLVGWTKGFKATDCEGQDVVEMLREAIKRRN----EFD
LDIVA-----IVNDTVGTMMTCGYEDPHCEIGLIAGTGSNLCYMEEM---KNIEFLE-------------
--GD------EG---------------------RMCINTEWGGFGDNDC-LADIQTQYDRHVDEKSLNPG
KQR-YEKMTSGMYLGEIVRQILIDLTKKGLLFRGQISERLRTKGIFETKFLSQIES--------------
------------DRLALLQVRKILQQLGL-DSTCDDSIIVKEVCGVVSTRAAKLCGAGLAAVVDKIRENR
GLDHLKITVGVDGTLYKLH-----------PHFSRILQETVKKLSPECDVSI---------LLSEDG---
----------SGKGAALITAVAKRFHS--QGA--ENKNSIGLNI--------
>XP_002936904|Xenopus tropicalis|HKDC1
LYHMRLSDETLLDIMGRFRNEMEKGL-KKDTNPTAS--LKMLSTHVRSTPDGSEKGDFLALDLG--GSNF
RVLRVKVSEN---GK-QK--VQMDSQYYPMPNETMTGSGAE---------------LFDYVADCLEDFIK
KR-----QIK----ENLPLGFTFSFPCKQTKLD------------------------E------------
-----------GVLISWTKEFKARGVQGSDVVNVLRSTLQKK----------Q-----------------
--------------CANVDVLALVNDTVGTMMTCGYDDQLCEVGVIV-----------------------
-----G-----TGTNACYMENLSNI-DFVEGDE--GRMCINTEWGAFGDDGLLDDIRTEFDEEIDLGSLN
PGKQLFEKMISGLYMGELVRLVLLKMAKKGLLFGGKISNALRTKGSIATSHVAAIEK-------------
--YK--EGLYNTNQILTE-LELEPAEEDCIAVQHVCTIVSFRSANLCAAALAAILERLRENK-KLERMRT
TVGMDGTVYKTH-PQYAKRLHKVVRRLVPNCDVRFLLSESGSGKGAAMVTAVACRLLSQHKQIDEILALL
QLSNENLFDIKERMMIELKNGLMKETHGSA-TVKMLPTFVYGTPDGT-----------------------
---EWGKFLALDLGGTNFRVLLVKIRSGRRKYVRMYNKIFAIPLEIMQGTGEE-------LFDHIVECIA
EFLDYMGIKGA-PLPLGFTFSFPCKQERIDKGTLVGWTKGFKATDCEGQDVVDMLREAIKRRN----EFD
LDIVA-----IVNDTVGTMMTCGYEDPNCEIGLIAGTGSNVCYMEEM---KNIEIVD-------------
--GD------EG---------------------KMCINTEWGGFGDNGC-LADIQTKYDIDVDEKSLNPG
KQR-YEKMTSGMYLGEIVRQILIDLTKCGLLFRGQISERLRTKGIFETKFLSQIES--------------
------------DRLALLQVRKILQQLGL-DSTCDDSIIVKEVCGAVSTRAAKLCGAGLAAVVDKIRENR
DLEHLKVTVGVDGTLYKLH-----------PHFSRILQETVKKLSPQCDVTF---------MLSEDG---
----------SGKGAALITAVAKRFHS--RGE--EETK--------------
>XP_040293980|Bufo bufo|HKDC1
LYHMRLSDETLMNIMARFQAEMDKGL-KKDTNSTAS--VKMLPTYVRSTPDGSEKGDFLALDLG--GSKF
KVLRVKVSED---GT-QN--VQMDSQFYPTPSEIINGSKAE---------------LFDYVANCLGDFMK
KR-----QIT----EKLPLGFTFSFPCKQTRLD------------------------E------------
-----------GVLISWTKDYKVRGVHQTDVVNTLRKTLKKK----------K-----------------
--------------YTNVDVLALVNDTVGTMMTCGYDDQLCEVGVIV-----------------------

-----G-----SGTNACYMENLSNI-DLVEGDE--GRMCINTEWGAFGDDGALDDIRTGFDNEIDLGSVN
PGKQLFEKMISGLYMGELVRLILLKMAKKGLLFGGKVSNALRTKGKIETSQVASMEK------------
--YN--EGLYNTRQILLD-LGMEPSEDDCIAVQHVATIVSFRSANLCAAALAAILTRLRENK-KLDRMRI
TVGMDGTVYRTH-PQYAKRLHKVVRRLVPNCDVRFLLSESGSGKGAAMVTAVACRLFSQHKQIDEVLDLL
HISDEKLADIKLRMITELKYGLKKETHDSA-TVKMLPTYVYGTPDGT----------------------
---EHGKFLALDLGGTNFRVLLVKIRSGRRRYVRMYNKIFAIPLEIMQGTGEE-------LFDHIVECIA
EFLDYMGIKGA-PLPLGFTFSFPCKQDRIDKGTLVGWTKGFKATDCEGQDVVDMLREAIKRRN----EFD
LDIVA-----IVNDTVGTMMTCGYEDPDCEIGLIAGTGSNVCYMEEM---KNIEFID------------
--GD------EE--------------------KMCINTEWGGFGDNDC-LSDIQTQYDIRVDEKSLNSG
KQR-YEKMTSGMYLGEIVRQILIDLTKRGLLFRGQISERLRTKGIFETKFISQIES-------------
------------DRLALLQVRKILQQLGL-DSTCDDSIIVKEVCGAVSTRAAKLCGAGLAAVVDKIRENR
GLDYLKTTVGVDGTLYKLH-----------PHFSRILQETVKKLSPQCDVTF---------ILSEDG---
----------SGKGAALITAVAKRFHS--QEE--PGKN--------------
>XP_005990008|Latimeria chalumnae|HKDC1
LYHMRLSDDTLLDVMTRFRVEMEKGL-AKATNPTAT--VKMLPTFVRSMPDGSEKGDFLALDLG--GSNF
RVLRVKVSED---GK-QK--VEMESQFYPTPEEVIHGTGKQ---------------LFDYVAECVVNFME
KK-----KIK---DKKLPLGFTFSFPCKQAKLD-----------------------E------------
-----------GILISWTKNFKARGVQGTDVVSSLRKAMLKY----------Q----------------
--------------DVDVDVLALVNDTVGSMMTCGFEDKYCEAGVII----------------------
-----G-----SGTNACYMEELRHI-DLVEGDE--GRMCINTEWGAFGDDGSLEDIRTEFDREIDLGSIN
PGKQLFEKMISGPHMGELVRLILVKMAKKGLLFKGNLSAALRTKGKFEMKHVAAIEK------------
--YK--EGLRNAREILSN-LGLDPTEEDCIAVQHVCTIVSFRAANLCATTLAAILARLRENK-KLRRLRT
TVGMDGTVYKTH-PQFAKRLHKVVRRLAPECDVRFLLSESGSGKGAAMVTAVAYRLVSQRKQVDEALAPF
KLTKDQLCEVKKRMRTELEHGLKKETHHTA-SVKMLPTYVYGTPDGT----------------------
---ERGKFLALDLGGTNFRVLLVKIRSGLRRSVRMYNKIYAIPLWIMQGTGEE-------LFDHIVQCIS
DFLDYMGMKGA-QLPLGFTFSFPCRQAGIDKGTLVGWTKGFKATDCEGEDVVDLLREAIKRRN----EFD
LDIVA-----VVNDTVGTMMTCGYEDPKCEIGLIAGTGSNVCYMEEM---RNIELME------------
--GD------EG--------------------RMCINTEWGAFGDNGC-IDDLRTQFDREVDEGSLNPG
KQR-YEKMTSGMYLGEIVRQILIHLTKQGLLFRGQVSERLKTRGIFETKFLSQIES-------------
------------DGLALLQVRGILQQLGL-DSTCDDSIIVKEVCGVISRRAAQLCGAGLAAIVDKIRENR
NLDCLEVTVGVDGTLYKLH-----------PHFSRVLQETVKEQAPTCGVTF---------MLSEDG---
----------SGKGAALITAVARRFHS--NRQ--H-----------------
>XP_029465520|Rhinatrema bivittatum|HKDC1
LYQMRLSDDDLLDIMTRFQKEMDKGL-KKDTNPTAM--VKMLPTHVRSTPDGSEKGDFLALDLG--GSKF
RVLNVRVSEN---GK-QS--VQMESQIYPAPKEITHGNGEE---------------LFEYVADCLVDFME
TK-----NLK---HKKLPLGATFSFPCKQTKLE-----------------------E------------
-----------GVLLSWTKNFKARRVQGTDVVSSLHNAIKKH----------K----------------
--------------DIDVDVLALVNDTVGTMMACGYDDQHCETAVII----------------------
-----G-----SGTNACYMEEMSNV-DLVEGDE--GRMCINTEWGAFGDDGLLDDIRTEFDKEIDLGSIN
PGMQLFEKMIGGLYLGELVRLVLLKMAKKGLLFKGKISTALCTKDKIETEHVSAMEK------------
--YK--EGLSNTKQILVD-LDLDPSEEDCIAVQHVCTIVSFRSANLCAAALAAILTRIRENR-KLRRLRT
TVGMDGTVYKTH-PQYAKRLHKVTRRLVPNCDVRFLVSESGSGKGAAMVTAVALRLLEQRKQIDEILAKF
KLTKEHLIDVKMKMMVEMKKGLKKETQESA-TVKMLPTYVYGTPDGT----------------------
---EHGKFLALDLGGTNFRVLLVKIRSGRRKSVKMYNKIFAIPLEIMQGTGEE-------LFDHIVECIA
EFLDYMGLKGA-PLPLGFTFSFPCRQTSIDKGSLVEWTKGFKATDCEGEEVVDMLREAIKRRN----EFD
LDIVA-----LVNDTVGTMMTCGYEDPNCKIGLIVGTGSNVCYMEEM---KNIETVN------------
--GN------EG--------------------KMCINTEWGGFGDNGC-LDDIRTNYDKEVDEGSLNSG
KQR-FEKMISGMYLGEIVRQILIDLTKQGLLFRGQISERLRTKGIFETKFISQIES-------------
------------DQLALLQVRRILQELGL-DSTCDDSVIVKEVCGTISTRAAQLCGAGMAAIVEKIWENS
GLDHLDITVGVDGTLYKLH-----------PHFSRILKETMADLAPECNVTF---------MQSEDG---
----------SGKGAALITAVAKRMHN--NLK--TGGNDAPPPECA------
>XP_026651632|Zonotrichia albicollis|HKDC1

```
LHQLRLSDDVLQDVAARFQAEMLKGL-ARDTNPTAA--VKMLPSFVRSLPDGSEKGEFLAVDLG--GSQL
RAHQVRVSDD---GK-QS--SQLESKFYPTPKEVTQGNGAE---------------LFDYIADCLSDFMD
TR-----NLK---QKKLPLGFTFSFPCKQTKLD------------------------E------------
-----------GVLLEWTKHFRVRGVQGTDVVSSLHRALQKH-QASPLMAIFK-----------------
--------------DIDVDVLALVNDTVGTMMTCGYDDPRCEVGLII-----------------------
-----G-----TGTNACYMEEMRHI-SLVEGDE--GRMCINTEWGAFGDDGALDDLRTEFDRELDLGSLN
PGKQLFEKMISSLYLGELVRLILLKMTKEGLLFNGKVSAALLTKGKIEMKHVSAMEK-------------
--YK--EGLSNTREILTE-LKLFPSEEDCVAVQHVCTIVSFRSANLCAAALAAILTRLRDNK-KLLRMRT
TVGIDGGLYKTH-PRYAQRLHKVLRRLVPTCDVRFVLSRSGSGRGAAMVTAVALRLAAQRRRLDAALAPF
LLPPSTLQQVRDSMRAELEYGLKREAQGQA-TVKMLPTYVCGMPDGT-----------------------
---EKGKFLALDLGGTNFRVLLVKIKSGRRRSVQMYNKIFAIPLEIMQGTGEE-------LFDHIVQCIA
DFLDYMGIKGA-RLPLGFTFSFPCRQTSIDKGTLVGWTKGFKATDCEGEDVVDMLREAIRRRN----EFD
LDIVA-----VVNDTVGTMMTCGYEDPNCEIGLIAGTGTNVCYMEDM---KNIETVE-------------
--GN------EG---------------------KMCINTEWGAFGDNGC-IDHIRTKYDREVDEGSLNPG
KQR-YEKMTSGMYLGEIVREILIDLTKQGLLFRGHISESLRKRGIFETKFLSQIES--------------
------------DRLALLQVRRVLQHLGL-DSTCEDSIIVKEVCAAVSTRAARLCGAGLAAVVEKRRQNR
GVERLQITVGVDGTLYKLH-----------PHFSRILQETVKELAPQCDVTF---------MLSEDG---
----------SGKGAALITAVAKRLHN--VGQ--K-----------------
>XP_009077222|Acanthisitta chloris|HKDC1
LYHLRLSDDVLLDVLARFQAEMVKGL-GRDTNPTAA--VKMLPSFVRSLPDGSEKGDFLAVDLG--GSQF
RAHQVKVFDD---GK-QS--SQLESKFYPTPKEVTQGNGAE---------------LFDYVADCLLDFMA
TK-----NLK---HKKLPLGFTFSFPCKQTKLE------------------------E------------
-----------GVLLAWTKHFKVRGVQDTDVVSSLHKALQKH----------K-----------------
--------------DVDVEVLALVNDTVGTMMTCGYDDPRCEVGLII-----------------------
-----G-----TGTNACYMEEMRHI-ELVEGDE--GRMCINTEWGAFGDDGALDDLRTEFDRELDLGSLN
PGKQLFEKMISSLYLGELVRLILLKMAKEGLLFNGKVSTALLTKGKIEMKHVSAMEK-------------
--YK--EGLSNAKEILTE-LSLFPSEEDCIAVQHVCTIVSFRSANLCAAALAAILTRLRENK-KLLRMRT
TVGIDGGLYKTH-PQYAKRLHKVVRRLVPNCDVRFLLSVSGSGKGAAMVTAVAHRLVTQRKQIDATLAPF
LLSLKTLREVKDKMRTELEYGLKRETQANA-TVKMLPTYVCGTPDGT-----------------------
---EKGKFLALDLGGTNFRVLLVKIRSGRRRSVQMYNKIFAIPLEIMQGTGEE-------LFDHIVQCIA
NFLDYMGIKGA-RLPLGFTFSFPCRQASIDKGTLVGWTKGFKATGCEGEDVVDMLREAIRRRN----EFD
LDIVA-----VVNDTVGTMMTCGYEDPNCEIGLIAGTGSNVCYMEDM---KNIEIVE-------------
--GN------EG---------------------KMCINTEWGAFGDNGC-IDNIRTKYDKEVDEGSLNPG
KQR-YEKMTSGMYLGEIVRQILIDLTKQGLLFRGHISESLRKRGIFETKFLSQIES--------------
------------DRLALLQVRQILRQLGL-DSKCDDSIIVKEVCGVVSRRAAQLCGAGLAAVVEKKRENQ
GVEHLQITVGVDGTLYKLH-----------PHFSRILQETVKQLAPQCDVTF---------MLSEDG---
----------SGKGAALITAVAKRLHS--VGQ--K-----------------
>XP_008495878|Calypte anna|HKDC1
LYHMRLSDDVLLDVMARFQAEMVKGL-GRDTNPTAT--VKMLPSFVRSLPDGSEKGEFLAVDVG--GSQF
RAHRVKVFDD---GK-QS--SELETKFYPSPKEIIQGNGAE---------------FFDYIADCLSDFME
AE-----NLK---HKKLPLGFTFSFPCKQTKLD------------------------E------------
-----------GVLLSWTKHFKVRGVQDTDVVSSLCKALEKH----------K-----------------
--------------ELDLDVLALVNDTVGTMMTCGYDDQRCEVGLII-----------------------
-----G-----TGTNACYMEDMRHI-DLVEGDE--GRMCINTEWGAFGDDGALDDLRTEFDLELDQGSLN
PGKQLFEKMISSLHLGELVRLILLKMTKDGLLFNGKVSTALLTKGKIEMEHVSAMEK-------------
--HK--EGLSNTKKILTE-LNLFPSEEDCIAVQHVCTIVSFRSANLCAAALAAILTRLRENK-KLSRMRT
TVGIDGGLYKTH-PQYAKRLHKVVRRLVPNCDVRFFLSLSGSGKGAAMVTAVGYRLAAQRKKIDAALAPF
LLSLDTLREVKNKMRTELEYGLKRETQASA-TVKMLPTYVCGTPDGT-----------------------
---EKGKFLALDLGGTNFRVLLVKIRSGRRKSVQMYNKIFAIPLEIMQGTGEE-------LFDHIVQCIA
DFLEYMGIKGA-RLPLGFTFSFPCRQASIDKGTLVGWTKGFKATDCEGEDVVDMLREAIKRRN----EFD
LDIVA-----VVNDTVGTMMTCGYEDPKCEIGLIAGTGSNVCYMEDM---KNIEIVE-------------
--GN------EG---------------------KMCINTEWGGFGDNGC-IDNIRTKYDKEVDEGSLNPG
```

KQR-YEKMTSGMYLGEIVRQILIDLTKQGLLFRGQISESLRKRGIFETKFLSQIES-------------
------------DRLALLQVRQILQHLGL-DSTCEDSIIVKEVCGAVSKRAAQLCGAGLAAIVEKKRENQ
GVERLQITVGVDGTLYKLH-----------PHFSRVLQETVKELAPQCDVTF---------MLSEDG---
----------SGKGAALITAVAKRLHN--VGQ--K-----------------
>XP_015721892|Coturnix japonica|HKDC1
LYHMRLSDDVLLDVMARFQAEMVKGL-GRDTNPTAT--VKMLPSFVRSLPDGSESGDFLAVDLG--GSQF
RALEVKVFND---GK-QS--SQLESKFYPTPKEVIQGSGTE---------------LFSYVADCLSDFME
SR-----NLK---HKKLPLGFTFSFPCKQTKLE-----------------------E-----------
-----------GVLLSWTKHFKARGVQGTDVVSSLHKALQKH----------K-----------------
--------------DIDVDVLAMVNDTVGTMMTCGYDDPRCEVGLII----------------------
-----G-----TGTNACYMEEMRHI-DLVEGDE--GRMCINTEWGAFGDDGALDDLRTEFDRELDLGSLN
PGKQLFEKMISSLYLGELVRLILLKMTKEGLLFNGKVSAALLTKGKIEMKHVSAMEK------------
--YK--EGLSNTKEILTE-LNLFPSEDDCIAVQHVCTIVSFRSANLCAAALAAILTRLRENK-KLLRLRT
TVGIDGGLYKTH-PQYPKRLHKVVRRLVPNCDVRFLLSQSGSAKGAAMVTAVAYRLAAQRKQIDAVLAPF
LLSLETLRDVMNKMRTELEYGLKRETQDRA-TVKMLPTYVCGTPDGT----------------------
---EKGKYLALDLGGTNFRVLLVKIRSGRRRSVRMYNKIFAIPLEIMQGTGEE-------LFDHIVQCIA
DFLEYMGIKGA-RLPLGFTFSFPCRQASIDKGTLVGWTKGFKATDCEGEDVVDMLREAIRRRN----EFD
LDIVA-----VVNDTVGTMMTCGYEDPNCEIGLIAGTGSNVCYMEDM---KNIEIVE------------
--GN------EG---------------------KMCINTEWGGFGDNGC-IDNIRTKYDKEVDEGSLNAG
KQR-YEKMTSGMYLGEIVRQILIDLTKQGLLFRGQISESLRKRGIFETKFLSQIES-------------
------------DRLALLQVRRILQQLGL-DSTCEDSIIVKEVCGAVSRRAAQLCGAGLAAIVEKKRENR
GVEHLQITVGVDGTLYKLH-----------PHFSRVLQETVRELAPRCDVTF---------MLSEDG---
----------SGKGAALITAVAKRLHN--VGQ--K-----------------
>XP_057229735|Malurus melanocephalus|HKDC1
LYPLRLSDDVLLDVMARFQAEMVKSL-GRDTNPTAT--VKMLPSFVRSLPDGSEKGDFLAVDLG--GSQF
RAHQVKVFDD---GK-QS--SQLESKFYPTPKEVTQGNGAE---------------LFDYIADCLLDFME
TK-----NLK---HKKLPLGFTFSFPCRQTKLE-----------------------E-----------
-----------GVLLAWTKHFKVRGVQDTDVVSFLRKALQKH----------K-----------------
--------------DIDVDVLALVNDTVGTMMTCGYDDQRCEVGLII----------------------
-----G-----TGTNACYMEEMRHI-DLVEGDE--GRMCINTEWGAFGDDGALDDLRTEFDRELDLGSLN
PGKQLFEKMISSLYLGELVRLILLKMTKEGLLFNGKVSTALLTKGKIETKHVSAMEK------------
--YK--EGLSNTKEILTE-LNLFPSEEDCVAVQHVCTIVSFRSANLCAAALAAILTRLRENK-KLLRMRT
TVGIDGGLYKTH-PQYAKRLHKVVRRLVPTCDVRFLLSVSGSGRGAAMVTAVARRLAAQRQQINATLAPF
LLPLGTLREVKDKMRAELEYGLKRETQGNA-TVKMLPTYVCGTPDGT----------------------
---EKGKFLALDLGGTNFRVLLVKIKSGRRRSVQMYNKIFAIPLEIMQGTGEE-------LFDHIVQCIA
DFLEYMGIKGA-RLPLGFTFSFPCRQASIDKGTLVGWTKGFKATDCEGEDVVDMLREAIRRRN----EFD
LDIVA-----VVNDTVGTMMTCGYEDPNCEIGLIAGTGSNVCYMEDM---KNIEIVE------------
--GN------EG---------------------KMCINTEWGAFGDNGC-IDNIRTKYDKEVDEGSLNPG
KQR-YEKMTSGMYLGEIVRQILIDLTKQGLLFRGHISESLQKRGIFETKFLSQIES-------------
------------DRLALLQVRRILQQLGL-DSTCDDSIIVKEVCGVVSRRAAQLCGAGLAAIVEKKRENR
GVERLQITVGVDGTLYKLH-----------PHFSRVLQETVKELAPQCDVTF---------MLSEDG---
----------SGKGAALITAVAKRLHN--VGQ--K-----------------
>XP_061229926|Neopsephotus bourkii|HKDC1
LYHMRLSDDILLDVMSRFQAEMVKGL-GRDTNPTAT--VKMLPSFVRSLPDGSEKGDFLAVDVG--GSLF
RAHQVKVFDD---GK-QS--SQLENKFYPTPKEITQGNGAE---------------LFDYIADCLLDFME
TK-----NLK---HKKLPLGFTFSFPCKQTKLE-----------------------E-----------
-----------GVLLAWTKHFKVRGVQDTDVVSSLRKALQKH----------K-----------------
--------------DIDVDVLALVNDTVGTMMTCGYDDQHCEVGLII----------------------
-----G-----TGTNACYMEEMRHI-DLVEGDE--GRMCINTEWGAFGDDGALDDLRTEFDRELDLGSLN
PGKQLFEKMISSLYLGELVRLILLKMTKEGLLFNGKVSTALLTKGKIEMKHVSAMEK------------
--YK--EGLNNTKEILTE-LNLFPSEEDCIAVQHVCTIVSFRSANLCAAALAAILTRLRENK-KVLRMRT
TVGIDGGLYKTH-PQYAKRLHKVVRRLVPNCDVRFLLSVSGSGKGAALVTAVAYRLAAQRKRIDAALAPF

LLSLETLREVKNKMRTELEYGLKRETQASA-TVKMLPTYVCGTPDGT-----------------------
---EKGKFLALDLGGTNFRVLLVKIRSGRRRSVQMYNKIFAIPLEIMQGTGEE-------LFDHIVQCIA
DFLEYMGIKGA-RLPLGFTFSFPCRQASIDKGTLVGWTKGFKATDCEGEDVVDMLREAIRRRN----EFD
LDIVA-----VVNDTVGTMMTCGYEDPNCEIGLIAGTGSNVCYMEDM---KNIEIVE-------------
--GN------EG---------------------KMCINTEWGAFGDNGC-IDNIRTKYDKEVDEGSLNPG
KQR-YEKMTSGMYLGEIVRQILIDLTKQGLLFRGQISESLRKRGIFETKFLSQIES--------------
------------DRLALLQVRRILQQLGL-DSTCEDSIIVKEVCGAVSKRAAQLCGAGLAAIVEKKRENR
GVERLQITVGVDGTLYKLH-----------PHFSRVLQETVKQLAPQCDVTF---------MLSEDG---
----------SGKGAALITAVAKRLHN--VGQ--K-----------------
> XP_054688625|Grus americana|HKDC1
LYHMRLSDDVLLDVMTRFQAEMVKGL-GKDTNPTAT--VKMLPTFVRSLPDGSEKGDFLAVDLG--GSQF
RAHQVKVFDD---GK-QN--SQLESKFYPTPKEVIQGNGAE---------------LFDYVADCLLDFME
TK-----NLK---NKKLPLGFTFSFPCKQTKLE------------------------E------------
-----------GVLLAWTKHFKVRGVQDTDVVNSLRKALQKH----------K-----------------
--------------DIDVDVLALVNDTVGTMMTCGYDDQRCEVGVII-----------------------
-----G-----TGTNACYMEEMRHI-DLVEGDE--GRMCINTEWGAFGDDGALDDLRTEFDRELDLGSLN
PGKQLFEKMISSLYLGELVRLILLKMTKEGLLFNGKVSTALLTKGKIEMKHVSAMEK-------------
--YK--EGLSNTKEILTE-LNLFPSEEDCIAVQHVCTIVSFRSANLCAAALAAILTRLRENK-KLLRMRT
TVGIDGGLYKTH-PQYAKRLHKVVRKLVPNCDVRFLLSVSGSGKGAAMVTAVAYRLAAQRKQIDAALAPF
LLSLETLREVKNKMRTELEYGLKRETQASA-TVKMLPTYVCGTPDGT-----------------------
---EKGKFLALDLGGTNFRVLLVKIRSGRRRSVQMYNKIFAIPLEIMQGTGEE-------LFDHIVQCIA
DFLEYMGIKGA-RLPLGFTFSFPCRQASIDKGTLVGWTKGFKATDCEGEDVVDMLREAIRRRN----EFD
LDIVA-----VVNDTVGTMMTCGYEDPNCEIGLIAGTGSNVCYMEDM---KNIEIVE-------------
--GN------EG---------------------KMCINTEWGGFGDNGC-IDNIRTKYDKEVDEGSLNPG
KQR-YEKMTSGMYLGEIVRQILIDLTKQGLLFRGQISESLRKRGIFETKFLSQIES--------------
------------DRLALLQVRRILQQLGL-DSTCEDSIIVKEVCGAVSKRAAQLCGAGLAAIVEKKRENR
GVEHLQITVGVDGTLYKLH-----------PHFSRVLRETVKELAPQCDVTF---------MLSEDG---
----------SGKGAALITAVAKRLHN--VGQ--K-----------------
> XP_026707802|Athene cunicularia|HKDC1
LYHMRLSDDVLLDVMARFQAEMVKGL-GRDTNPTAT--VKMLPSFVRSLPDGSEKGDFLAVDLG--GSQF
RAHQVKVFDD---GK-QS--SQLESKFYPTPKEVIQGNGAE---------------LFDYIAECLLDFTE
TK-----NLK---HKKLPLGFTFSFPCKQTKLD------------------------E------------
-----------GVLLAWTKHFKVRGVQDTDVVSSLRKALQKH----------K-----------------
--------------DIDVDVLALVNDTVGTMMTCGYDDQRCEVGLII-----------------------
-----G-----TGTNACYMEEMRHI-DLVEGDE--GRMCINTEWGAFGDDGALDDLRTEFDRELDLGSLN
PGRQLFEKMISSLYLGELVRLILLKMTKEGLLFNGKVSTALLTKGKIEMKHVSAMEK-------------
--YK--EGLSNTKEILTE-LNLFPSEEDCIAVQHVCTIVSFRSANLCAAALAAILTRLRENK-KLLRMRT
TVGIDGGLYKTH-PQYAKRLHKVVRRLVPNCDVRFLLSVSGSGKGAAMVTAVAYRLAAQRKRIDAALAPF
LLSLETLQEVKNKMRTELEYGLKRETQASA-SVKMLPTYVCGTPDGT-----------------------
---EKGKFLALDLGGTNFRVLLVKIRSGRRRSVQMYNKIFAIPLEIMQGTGEE-------LFDHIVQCIA
DFLEYMGIKGA-RLPLGFTFSFPCRQASIDKGTLVGWTKGFKATDCEGEDVVDMLREAIRRRN----EFD
LDIVA-----VVNDTVGTMMTCGYEDPNCEIGLIAGTGSNVCYMEDM---KNIEIVE-------------
--GN------EG---------------------KMCINTEWGGFGDNGC-IDNIRTKYDKEVDEGSLNPG
KQR-YEKMTSGMYLGEIVRQILIDLTKQGLLFRGQISESLRKRGIFETKFLSQIES--------------
------------DRLALLQVRRILQQLGL-DSTCEDSIIVKEVCGAVSKRAAQLCGAGLAAIVEKKRENQ
GVEHLQITVGVDGTLYKLH-----------PHFSRVLRETVKELAPQCDVTF---------MLSEDG---
----------SGKGAALITAVAKRLHN--VGQ--K-----------------
> XP_009814831|Gavia stellata|HKDC1
LYHMRLSDDVLLDVMARFQAEMVKGL-GRDTNPTAT--VKMLPSFVRSLPDGSEKGDFLAVDLG--GSQF
RAHQVKVFDD---GK-HS--SQLESKFYPTPKEVIQGNGAE---------------LFDYIADCLLDFME
TK-----NLK---HKKLPLGFTFSFPCKQTKLD------------------------E------------
-----------GVLLAWTKHFKVRGVQDTDVVSSLRKALQKH----------K-----------------

--------------DIDVDILALVNDTVGTMMTCGYDDPRCEVGLII-----------------------
-----G-----TGTNACYMEEMRHI-DLVEGDE--GRMCVNTEWGAFGDDGALDDLRTEFDRELDLGSLN
PGKQLFEKMISSLYLGELVRLILLKMTKEGLLFNGKVSTALLTKGKIEMKHVSAMEK-------------
--YK--EGLSNTREILTE-LNLFPSEEDCIAVQHVCTIVSFRSANLCAAALAAILTRLRENK-KLLRMRT
TVGIDGGLYKTH-PQYAKRLHKVVRRLVPNCDVRFLLSVSGSGKGAAMVTAVAYRLAAQRKQIDAALAPF
LLSLETLREVKNKMRTELEYGLKRETQASA-TVKMLPTYVCGTPDGT-----------------------
---EKGKFLALDLGGTNFRVLLVKIRSGRRRSVQMYNKIFAIPLEIMQGTGEE-------LFDHIVQCIA
DFLEYMGIKGA-RLPLGFTFSFPCRQASIDKGTLVGWTKGFKATDCEGEDVVDMLREAIRRRN----EFD
LDIVA-----VVNDTVGTMMTCGYEDPNCEIGLIAGTGSNVCYMEDM---KNIEIVE-------------
--GN------EG---------------------KMCINTEWGGFGDNGC-IDNIRTKYDKVVDEGSLNPG
KQR-YEKMTSGMYLGEIVRQILIDLTKQGLLFRGQISESLRKRGIFETKFLSQIES--------------
------------DRLALLQVRRILQQLGL-DSTCEDSIIVKEVCGVVSKRAAQLCGAGLAAIVEKKRENR
GVERLQITVGVDGTLYKLH-----------PHFSRVLRETVKELAPQCDVTF---------MLSEDG---
----------SGKGAALITAVAKRLHN--VGQ--K-----------------
>XP_007427559|Python bivittatus|HKDC1
LYQMRLSDDILHDVMTRFQAEMVKGL-GRDTSPTAT--VKMLPTFVRSLPDGSEKGEFLALDLG--GSKF
KVLRVKVAEE---GK-QH--VQMESQFYPTPKEIIQGSGTD---------------LFEYVANCLADFME
TK-----DIK---YKKLPLGFTFSFPCRQNKLE------------------------E------------
-----------GVLLAWTKHFKVRGVQQTDVVSSLCKALRKH----------E-----------------
--------------DIDVEVLALVNDTVGTMMTCGYDDQRCEVGVII-----------------------
-----G-----TGTNACYMEEMSNI-DLVEGDE--GRMCINTEWGAFGDDASLEDIRTEFDKEIDLGSMN
PGKQLFEKMISGLYLGELVRLILLKMSRKGLLFSGKLSTALCTKGKIETKHVAAMEK-------------
--YK--EGLSNTREILRE-LGLNPSEEDCVAVQHVCTIVSFRSANLCATALAAILTRLRENK-KVARLRT
TVGIDGTLYKTH-PQYAKRLHKVVRRLVPNCDVRFLLSESGSGKGAAMVTAVAHRLLSQRKQIDEVLAPF
KLTKENLIEVRNKMRTELEYGLKKETHPTA-TVKMLPTFVHGTPDGT-----------------------
---EKGKFLALDLGGTNFRVLLVKIRSGRNKSVRIYNKIFAIPLEIMQGTGEE-------LFDHIVQCIA
DFLEYMGIKGA-RLPLGFTFSFPCRQTSIDKGALVRWTKGFKATNCEGEDVVDLLREAIKRRN----EFD
LDIVA-----VVNDTVGTMMTCGYEDPHCEIGLIAGTGSNVCYMEEM---KHIEIIG-------------
--GA------EG---------------------RMCINTEWGGFGDNGC-IDNFRTKYDKEVDSWSLNPG
NQK-YEKMTGGMYLGEIVRQILIDLTKRGLLFRGQISESLRRRRIFETKYLSQIES--------------
------------DRLALLQVRRILQQLGL-DGTCDDSIIVKEVCGCVSKRAAQLCGAGLAAVVEKIRDNR
KLEHLKTTVGVDGTLYKLH-----------PHFSGILQETVKELAPQCDVTF---------MLSEDG---
----------SGKGAALITAVAKRMHQ--VAE--N-----------------
>XP_028585286|Podarcis muralis|HKDC1
LYHMRLSDDVLLDLMARFQAEMVKGL-ARDTNPTAT--LKMLPTFVRSIPDGTEKGEFLALDLG--GSRF
RVLKVKVSEN---GK-QN--VQMESQIYPTPKEIIHGSGTD---------------LFQYVADCLADFME
TK-----NIK---HKKFPLGFTLSFPYIQRKLE------------------------E------------
-----------GILISWTKHFKARGVQQTDVVKSLQKALRKH----------Q-----------------
--------------DIDVDILALVNDTVGTMMTCGYDDQRCEVGVII-----------------------
-----G-----TGTNACYMEEMSNI-DLVEGDE--GRMCINTEWGAFGDDASLDDIRTEFDKEIDLGSFN
PGKQLFEKMISGLYLGELVRIILLKMTREGLLFNGKLSTALCTKGKIETKHIAAMEK-------------
--YK--EGLSNTKEILME-LGLAPSEDDCIAVQHVCTVVSFRSANLCAAALAAILTRLRENK-KVARLRT
TVGVDGTLYKIH-PQYAKRLHKVVRRLVPNCDVRFLLSESGSGKGAAMVTAVACRLLSQRKQINEVLSLF
KITKENLIGVKNRMRAELEYGLKKETHPVA-TVKMLPTFVYGTPDGT-----------------------
---ENGKFLALDLGGTNFRVLLVKIRSGRSKSVRMYNKIFAIPLEIMQGTGEE-------LFDHIVQCIA
DFLEYMGIKGA-RLPLGFTFSFPCRQACIDKGTLVGWTKGFKATDCEGEDVVDMLREAIKRRN----EFD
LDIVA-----VVNDTVGTMMTCGYEDSNCEIGLIAGTGSNVCYMEDM---RNIEIVE-------------
--GD------EG---------------------KMCINTEWGGFGDNGC-IDNLRTKYDKEVDEGSLNPG
EQR-YEKMTSGMYLGEIVRQILIDLTKQGLLFRGQISERLRTRGIFETKFLSQIES--------------
------------DRLVLLQVRRILQQLGL-DSTCDDSIIVKEVCGSVSKRAAQLCGAGLAAIVEKIRDNR
KLEHLKTTVGVDGTLYKLH-----------PHFSRILQETVKELAPQCDVTF---------MLSEDG---
----------SGKGAALITAVAKRVHN--VAE--N-----------------

>XP_056705300|Euleptes europaea|HKDC1
LYHMRLSDDVLLDVMKRFQAEMVKGL-GRDTNPTAT--LKMLPTFVRSIPDGSEKGDFLALDLG--GSRF
RVLKVKVSED---GK-QN--VQMESQFYPTPKEIIHGNATD---------------LFKYVAECLTDFME
TK-----GIK---HKKLPLGFTFSFPCRQNKLE------------------------E------------
-----------GVLLSWTKHFKVRGVQQTDVVSSLRKALGKH----------Q-----------------
--------------DIDVDVLALVNDTVGTMMTCGYDDQRCEVGVII-----------------------
-----G-----TGTNACYMEEMSNI-ELVEGDE--GQMCINTEWGAFGDDGSLESIRTEFDREIDLGSIN
PGKQLFEKMISGLYLGELVRLILLKMTREGLLFNGKLSTDLCTKGKIETRHVAAMEK-------------
--YK--EGLNNTKEMLKE-LGLAPSEEDCIAVQHVCTIVSFRSANLCASALAAILTRLRENK-KLIRLRT
TVGMDGTLYKTH-PQYAKRLHKVVRRLVPNCDVRFLLSESGSGKGAAMVTAVAYRLLSQRKQIDEVLAYF
RLTEEDLIEVKNKMRMELESGLKKESQPTA-TVRMLPTFVCGTPDGT-----------------------
---EKGKFLALDLGGTNFRVLLVKIRSGRSKSVRMYNKIFAIPLEIMQGTGEE-------LFDHIVQCIA
EFLEYMGIKGA-QLPLGFTFSFPCRQASIDKGALIGWTKGFKATDCEGEDVVDMLREAIKRRN----EFD
LDIVA-----IVNDTVGTMMTCGYEDPNCEIGLIAGTGSNVCYMEDM---KNIEIVD-------------
--GD------EG---------------------RMCINTEWGGFGDNGC-INNLRTKYDKEVDEGSLNPG
KQR-YEKMTSGMYLGEIVRQILIDLTKQGLLFRGQISESLRTRGIFETKFLSQIES--------------
------------DRLALLQVRKILQQLGL-DSTCDDSIIVKEVCGCVSKRAAQLCGAGLAAVVEKIRENR
NLEHLKTTVGVDGTLYKLH-----------PHFSRILQETVKELAPRCDVTF---------LLSEDG---
----------SGKGAALITAVAKRMHS--AFTEN------------------
>XP_065264132|Emys orbicularis|HKDC1
LYHMHLSDDVLLDVMTRFQAEMVKGL-ERDAHPTAA--VKMLPTFVRSIPDGSEKGDFLALDLG--GSKF
RVLKVKVSED---GK-QN--VQMESQFYPTPKEITHGNGTE---------------LFDYVADCLADFME
TK-----NIK---HKKFPLGFTFSFPCIQTKLD------------------------E------------
-----------GVLLSWTKYFKVRGVQETDVISSLRHVLRKH----------K-----------------
--------------DVDVDVLALVNDTVGTMMTCGYDDQRCEVGVII-----------------------
-----G-----TGTNACYMEEMSNI-DLVEGDE--GRMCINTEWGAFGDDGSLDDLRTEFDREIDLGSIN
PGKQLFEKMISGLYLGELVRLILLKMAKKGLLFNGKISTALRTKDKIETKHVAAMEK-------------
--HK--KGLQNTKEILME-LGLTPSEEDCIAVQHVCTIVSFRSANLCAAALAAILTRLRENK-KLIRLRT
TVGIDGTLYKTH-PQYPKRLHKVVRRLVPNCDVRFLLSESGSAKGAAMVTAVACRLVSQRKQIDEILASF
ELSKENLIEVKNKMREELEYGLKKETHATA-TVKMLPTYVCGTPDGT-----------------------
---EKGKFLALDLGGTNFRVLLVKIRSGRRKSVRMYNKIFAIPLEIMQGTGEE-------LFDHIVQCIA
DFLEYMGIKGA-RLPLGFTFSFPCRQASIDKGALVEWTKGFKATDCEGEDVVDMLREAIKRRN----EFD
LDIVA-----VVNDTVGTMMTCGYEDPNCEIGFIAGTGSNVCYMEEM---KNIEIVE-------------
--GT------EG---------------------KMCINTEWGGFGDNGC-IDSIRTKYDKEVDEGSLNPG
KQR-YEKMTSGMYLGEIVRQILIDLTKHGLLFRGQISESLRTRGIFETKFLSHIES--------------
------------DRLALLQVRRILQQLGL-DSTCDDSIIVKEVCGAVSKRAAQLCGAGLAAIVEKKRENR
NLEHLKITVGVDGTLYKLH-----------PHFSGILRETVKEMAPQCDVTF---------MLSEDG---
----------SGKGAALITAVAKRLHN--VGE--N-----------------
>XP_006273539|Alligator mississippiensis|HKDC1
LYHMRLSDDVLLDLMTRFQTEMVKGL-GRDTNPTAA--VKMLPTFVRSIPDGSEKGDFLALDLG--GSKF
RVLKVKVSED---GK-QN--VQMESQLYPTPKEIIHGNATE---------------LFDYVADCLEDFME
NK-----NIK---HKKLPVGFTFSFPCKQAKLD------------------------E------------
-----------GVLLSWTKHFKARGVQETDVVSSLRNALRKH----------K-----------------
--------------DVHADVLALVNDTVGTMMTCGYDDQRCEVGVII-----------------------
-----G-----TGTNACYMEEMSHI-DLVEGDE--GRMCINTEWGAFGDDGSLEDIRTEFDREIDLGSIN
PGKQLFEKMISGLYLGELVRLILLKMAKEGLLFNGEISTDLRTKGKIETKHVAAMEK-------------
--YK--EGLSNTKEILME-LGLFPSEEDCVAVQHVSTIVSFRSANLCAAALAAILTRLRENK-KLMRLRT
TVGMDGTLYKTH-PQYAKRLHKVVRRLVPNCDVRFLLSESGSGKGAAMVTAVAYRLASQRKQIDEILASF
KLSKENLREVKTKMRAELEYGLKKETQPNA-TVKMLPTYVCGTPDGT-----------------------
---EKGKFLSLDLGGTNFRVLLVKIRSGRRKAVRMYNKIFAIPLEIMQGTGEE-------LFDHIVQCIA
DFLEYMGIKGA-RLPLGFTFSFPCRQASIDKGTLVGWTKGFKATDCEGEDVVDMLREAIKRRN----EFD
LDIVA-----VVNDTVGTMMTCGYEDPNCEIGLIAGTGSNVCYMEDM---KNIEVVE-------------

--GN------EG---------------------KMCINTEWGGFGDNGC-IDNIRTKYDKEVDEGSLNPG
KQR-YEKMTSGMYLGEIVRQILIDLTKHGLLFRGQISETLRTRGIFETKFLSQIES--------------
------------DRLALLQVRRILQQLGL-DSTCDDSIIVKEVCGAVSRRAAQLCGAGLAAVVEKKRENR
GVEHLKITVGVDGTLYKLH-----------PHFSRILREMVKELAPQCDVTF---------MLSEDG---
----------SGKGAALITAVAKRLHS--IGE--N-----------------
>XP_004701445|Echinops telfairi|HKDC1
LYHMRLSDETLLEIMARFQAEMEKGL-RKDTNPTAA--VKMLPTFVRAIPDGSEHGEFLSLDLG--GSKF
RVSKVQVSEE---GK-HK--VHMESQFYPVTGEITRGNGSQ---------------LFEYVADCLADFME
AK-----DLK---QKELPLGFTFSFPYWQTKLD------------------------E------------
-----------GVLLSWTKKFKARGVQDTDVVSHLRRALQKH----------P-----------------
--------------DLHVDILALVSDTVGTMMTCAYDDPYCEVGVII-----------------------
-----G-----TGSNACYMEDMSNI-DLVEGDE--GRMCINTEWGAFGDDGVLEDIRTEFDRELDLGSLH
PGKQLFEKMISGLYLGELVRLILLKMTRAGLLFGGKASCALHTKDKIETRHVAAMEK-------------
--YK--EGLANTKEILTN-LGLEPSEADCAAVQQVCTIVSFRSANLCAAALAAILTRLRENK-KLVRLRT
TVGVDGTLYKTH-PQYPKRLHKVVRKLAPNCDVRFLLSESGSTKGAAMVTAVASRVQAQRKQIDEVLALF
QLTRDNLVDVKNRMRAELEYGLKKETHPMA-TVKMLPTYVCGLPDGT-----------------------
---EKGKFLALDLGGTNFRVLLVKFRKG-RSSVRMYNKIFAIPLEIMQGTGEE-------LFDHIVQCIA
DFLDYMGLKGA-QLPLGFTFSFPCRQTSIDKGILVEWTKGFKATDCEGEDVVDLLREAIKRRN----EFD
LDIVA-----LVNDTVGTMMTCAHEDPKCEIGLIAGS---------------------------------
--TE------RG-----------------------VVRM-----------APELESPMPGPLMPVLLQLG
DPQ-YEKMTSGMYLGEIVRQILIDLTKQGLLFRGHISECLRTRGIFETKFLSQIES--------------
------------DRLALLQVRAILQQLGL-DSTCDDSIVVKEVCGAVSRRAAQLCGAGLAAIVQKRREDQ
GLKHLKITVGVDGTLYKLH-----------PHFSQILQETVKELAPECDVTF---------MLSEDG---
----------SGKGAALITVVAQRFRQ--LQK-EN-----------------
>XP_028915043|Ornithorhynchus anatinus|HKDC1
LYHMRLSDDSLLDMMARFQAEMEKGL-GRDTNPTAT--VKMLPTFVRAIPDGSEKGEFLALDLG--GSKF
RVLKVKVSED---GK-QT--VQMESQFYPTPEEIIQGNGPE---------------LFEYVADCLVDFVK
AK-----DMQ---HKKIPLGFTFSFPCKQTKLE------------------------E------------
-----------GVLLSWTKKFKVRGVQGTDVVSSLTTALKKR----------K-----------------
--------------DIDVDVLALVNDTVGTMMTCAYDDPRCEVGVII-----------------------
-----G-----TGTNACYMEDMSNI-DLVEGDE--GRMCINTEWGAFGDDGSLEDIRTEFDRELDLGSIN
PGKQLFEKMISGLYLGELVRLILLKMAKEGLLFGGRTSTALLTKGKIETQHVAAMEK-------------
--HK--EGLGNTKEILRE-LGLEPTEADCVAVQHVCTVVSFRSANLCAATLAAILTRLRENK-KVARLKT
TVGMDGTLYKTH-PQFPKRLHKVVRRLAPNCDVRFLLSESGSAKGAAMVTAVASRVQAQRKQIDQVLAML
KLSRGHLVSVRNKMRAELEYGLKKETQPLA-TVKMLPTYVCGLPDGT-----------------------
---EKGKYLALDLGGTNFRVLLVKIRSG-RKSVRMFNKIFAIPQEIMQGTGEE-------LFDHIVQCIA
DFLDYMGLKGA-RLPLGFTFSFPCRQTSIDKGILIGWTKGFKATDCEGEDVVDLLRDAIKRRN----EFE
LDIVA-----VVNDTVGTMMTCAYEDPECEIGLIAGTGSNVCYMEEM---RNIELVE-------------
--GD------TG---------------------KMCINTEWGAFGDKGC-LDEIRTQYDREVDEGSLYPG
KQR-YEKMTSGMYLGEIVRQILIDLTKSGLLFRGQISERLRTKGIFETKFLSQIES--------------
------------DRLALLQVRRVLQQLGL-DSTCEDSIVVKEVCGAVSRRAAQLCGAGLAAVVDKRREDR
KLEHLKITVGVDGTLYKLH-----------PHFSRILQETVKELAPRCDVTF---------MLSEDG---
----------SGKGAALITAVAQRMQQ--EKE--N-----------------
>XP_043840961|Dromiciops gliroides|HKDC1
LYDMRLSDDTLLDIMTRFRAEMEKGL-GKDTNPTSC--VKMLPTFVRAIPDGSEKGEFLALDLG--GSKF
RVLTVQVSEE---GK-PN--VHMESQFYPTPMEIIRGNASE---------------LFDYVADCVADFMK
TR-----NIE---HKKLPLGLTFSFPCQQRKLE------------------------E------------
-----------GILLSWTKKFKVRGVQDTDVVSVLTKALKRH----------K-----------------
--------------DIDVDVLALVNDTVATMMTCAYDDPHCEVGVII-----------------------
-----G-----TGTNACYMEDMSNI-DLVEGDE--GRMCINTEWGAFGDDGCLEDIRTDFDREIDLGSIN
PGKQLFEKMISGLYIGELVRLILLKMTKAGLLFRGEKSAALYTKGKIETQHVAAMEK-------------
--YK--EGLQNTREILME-LGLEPSEGDCVAVQHVCTIVSFRSANLCAAALAAILTRLRENK-KLVRLRT

TVGMDGTLYKTH-PQYPKRLHKVVRKLVPNCDVRFLLSESGSAKGAAMVTAVAARVQAQRKQIDEVLAMF
QLTREHLVSVKTKMRAEMDYGLKKETQPLA-TVKMLPTYVCAMPDGT-----------------------
---EKGKYLALDLGGTNFRVLLVKIRSG-RRSVRMYNKIFAIPLEIMQGTGEE-------LFDHIVQCIA
DFLDYMGIKGANSLPLGFTFSFPCQQTNLDKGMLVEWTKGFKATDCKGEDVVDMLREAVKRRN----EFE
LDIVA-----IVNDTVGTMMTCGYEDPDCEIGLIAGTGSNVCYMEEM---RNIELVE-------------
--GN------EG---------------------KMCINTEWGGFGDNGC-LDKIRTQYDAEVDEGSLNPG
KQR-YEKMTSGMYLGEIVRQILIDLTKHGLLFRGQISERLRTRGIFETKFLSQIES--------------
------------DRLALLQVRRILQQLGL-DSTCEDSIVVKEVCEAVSRRAAQLCGAGLAAIVEKRREDR
NLEHLKITVGVDGTLYKLH-----------PHFSRILRETVKELAPQCDVTF---------MLSEDG---
----------SGKGAALITAVAKRLHL--KKD--H-----------------
>XP_006835906|Chrysochloris asiatica|HKDC1
LYHMRLSDETLLDIMKKFHAEMKKGL-GQDTNPTAS--VKMLPTFVRAIPDGSENGEFLSLDLG--GSKF
RIQKVQVSGE---GK-RN--VHMESQFYPMTDEITRGNGSE---------------LFEFIAECLHDFMK
AK-----DLK---NKKLPLGFTFSFPCRQTKLD------------------------E------------
-----------GFLLSWTKKFKKQGVLNTDVVSRLSKAMKKY----------P-----------------
--------------DMDVDIVTVVNDTVATMMTCAYDDPYCEVGVII-----------------------
-----G-----TGSNACYMEDMSNI-DLVEGDE--GRMCISTEWGAFGDDGALEDIRTEFDRELDLGSLN
PGKQLFEKMISGLYLGELVRLILLKMARAGLLFGGKTSSALLTKDMIETQHVAAMEK------------
--YK--EGLANTKEILTD-LGLEPSEADCIAVQHVCTIVSFRSANLCAAALAAILTRLRENK-KLERLRT
TVGVDGTLYKIH-PQYPKRLHKVVRKLVPNCDVRFLLSENGSSKGAAMVTAVALRVQAQRKQIDKLLALF
RLTHEHLVGVRDKMRAELEYGLKKTTHPMA-TVKMLPTYVYGIPDGT-----------------------
---ENGKFLALDLGGTNFRVLLVKIKRG-RKSVQMYSKIFAIPLEIMQGTGEE-------LFDHIVQCIS
DFLDYMGLKGA-HLPLGFTFSFPCRQMSIDKGVLIKWTKGFKATDCENEDVVEMLREAIKRRN----EFD
LDIVA-----LINDTVGTMMTCAYEDPSCEIGLIAGTGSNACYMEEM---RNIELMK-------------
--EN------KG---------------------KMCINTEWGGFGDNGC-IDDIQTQYDREVDEGSLNQG
KQR-YEKMTSGMYLGEIVRQILIDLTRQGLLFHGKVSERLQTRGIFETKFLSQIES--------------
------------DRLALLQVRGILQQLGL-DSTCEDSIVVKEVCGAVSRRAAQLCGAGLAAIVEKRREDQ
GLESLKITVGVDGTLYKLH-----------PHFSRVLQETVKELAPQCNVTF---------MLSEDG---
----------SGKGAALITAVAKRLQQ--SQK-EN-----------------
>XP_049620508|Suncus etruscus|HKDC1
LYHMRLSDETLLDLMARFQAEMRKGL-DKDTNPTAA--VKMLPTFVRAIPDGSEKGEFLSLDLG--GSKF
RVLKVQVSEE---GK-HE--VQMESQFYPTPNEIIQGNGTK---------------LFDYVADCLEDFMK
AK-----GLQ---HKKLPLGLTFSFPCRQTKLD------------------------E------------
-----------GILLSWTKNFKAWGIQDTDVVSVLKKAMKKH----------K-----------------
------------LPDLDVDVLALVNDTVGTMMTCAYDDPHCEVGVII-----------------------
-----G-----TGTNACYMEDLSNI-DLVEGDE--GRMCINTEWGAFGDDGALEDIRTEFDRELDLGSLN
PGKQLFEKMISSLYLGELVRIILLKMAKAGLLFGGKVSSALLTKGKLETRHVAAMEQ------------
--YK--EGLANTREILTD-LGLEPSEADCVAVQHVCTVISFRSANLCAAALAAILKRLQENK-KVARLRT
TVGMDGTLYKIH-PQYPKRLHKVVRKLVPTCDVRFLLSESGSTKGAAMVTAVACRLQAQRKRIDQVLAQF
RLTRQQLEGVRDKMRAELEYGLKRDTHKQA-TVKMLPTYVYGVPDGT-----------------------
---ENGKFLALDLGGTNFRVLLVKIRSG-RRSVRMYNKIFAIPLEIMQGTGEE-------LFDHIVQCIA
DFLDYMGLKGA-QLPLGFTFSFPCSQTSIDKGILIGWTKGFKATDCEGEDVVDMLREAIKRRN----EFD
LDIVA-----VVNDTVGTMMTCGYEDPNCEIGLIAGTGSNMCYMEEM---RNIETVE-------------
--GD------EG---------------------RMCINSEWGGFGDNGC-LEDIRTQYDKEVDQGSLNSG
KQR-YEKMTSGMYLGEIVRQILIDLTKQGLLFRGQISERLRTRGIFETKFLSQIES--------------
------------DRLALLQVRSILQQLGL-DSTCEDSIVVKEVCGAVSRRAAQLCGAGLAAVVEKRRQDQ
GLEHLRVTVGVDGTLYKLH-----------PHFSRILQETVQELAPKCTVTF---------LLSEDG---
----------SGKGAALITAVAKRMQQ--RQK-EP-----------------
>EPQ14288|Myotis brandtii|HKDC1
LYHMRLSDETLLDIMARFQAEMQRGL-GKDTNPTAS--VKMLPTFVRAIPDRSENGEFLSLDLG--GSKF
RVLKVQVSEE---GK-QK--VQMESQFYPTPNEIIRGNGAE---------------LFEYVADCLADFMK
TK-----GLM---HKKLPLGMTFSFPCWQTKLE------------------------E------------

-----------GVLLSWTKKFKARGVQDTDVVSRLTKAMKNH----------K-----------------
--------------DIDVDILALVNDTVGTMMTCAYDDPSCEVGVII-----------------------
-----G-----TGTNACYMEDMSNI-DLVEGDE--GRMCINTEWGAFGDDGALEDIRTEFDRDLDLGSLN
PGKQLFEKMISGLYLGELVRLILVKMTKAGLLFGGKKSSSLHTKGKIETRHVAAMEK-------------
--YK--EGLANTREILTN-LGLEPSEADCIAVQHVCTIVSFRSANLCAAALAAILTRLRENK-KLVRLRT
TVGVDGTVYKIH-PQYPKRLHKVVRRLVPNCDVRFFLSESGSTKGAAMVTAVAARVQAQRKLVDQVLALF
RLTREQLVGVRDKMRAELEYGLKRDTHLLA-TVKMLPSYVCGMPDGT-----------------------
---ERGNFLALDLGGTNFRVLLVKIRSG-RRSVRMYNKIFAIPLEIMQGTGEE-------LFDHIVQCIS
DFLDYMGLKGV-PLPLGFTFSFPCRQTGIDKGILIEWTKGFKATDCEGEDMVDMLREAIKRRN----EFD
LDIVA-----VVNDTVGTMMTCGYEDPNCEIGLIAGTGSNMCYMEEM---RNIEVVE-------------
--GD------EG---------------------KMCINTEWGGFGDNGC-IDDIRTQYDKEVDEGSLNPG
KQR-YEKMTSGMYLGEIVRQILIDLTKQGLLFRGQISERLRTRGIFETKFLSQIES--------------
------------DRLALLQVRRILQQLGL-DSTCEDSIVVKEVCGAVSRRAAQLCGAGMAAIVEKRRQDQ
GLEHLKITVGVDGTLYKLH-----------PHFSWILQETVKELAPKCDVTF---------MLSEDG---
----------SGKGAALITAVAKRLQQ--AQK-EN-----------------
>XP_007938860|Orycteropus afer afer|HKDC1
LYHMRLSDETLLDIKARFQAEMEKGL-RKDTNPTAS--VKMLPTFVRAIPDGSENGEFLSLDLG--GSKF
RVLKVQVSEE---GK-RN--VQMESQFYPTPNEIIRGNGSE---------------LFDYVVDCLADFMK
TK-----DLK---HKKFPLGLTFSFPCKQTKLE------------------------E------------
-----------GVLLSWTKKFKARGVQDTDVVGHLTKAMKKH----------P-----------------
--------------DMDVDILALVNDTVGTMMTCAYDDPYCEVGVII-----------------------
-----G-----TGTNACYMEDMSNI-DLVEGDE--GRMCINTEWGAFGDDGALEDIRTEFDRELDLGSLN
PGKQLFEKMISGLYLGELVRLILLKMARAGLLFGSKKSSALHTKGKIETRHVAAMEK-------------
--YK--EGLANTREILTD-LGLEPSEADCIAVQHVCTIVSFRSANLCAAALAAILTRLRENK-KLVRLRT
TVGMDGTLYKIH-PQYPKRLHKVVRKLVPNCDVRFLLSECGSAKGAAMVTAVASRVQAQRKRIDEVLALF
RLTHEHLVDMQEKMRVELEYGLKRKTHPMA-TVKMLPTYVYGMPDGT-----------------------
---EKGKFLALDLGGTNFRVLLVKIRRG-RKSVRMYNKIFAIPLEIMQGTGEE-------LFDHIVQCIA
DFLDYMGLKGA-QLPLGFTFSFPCRQTSIDKGILIEWTKGFKATDCEGEDVVDMLREAIKRRN----EFD
LDIVA-----VVNDTVGTMMTCAYEDPNCEIGLIAGTGSNVCYMEEM---RNIELVD-------------
--ED------EG---------------------RMCINTEWGGFGDNGC-MDDIRTQYDKEVDEGSLNPG
KQR-YEKMTSGMYLGEIVRQILIDLTKQGLLFRGQVSERLRTRGIFETKFLSQIES--------------
------------DRLALLQVRGILQQLGL-DSTCEDSIVVKEVCGAVSRRAAQLCGSGLAAIVEKRRQDQ
GLEQLRITVGVDGTLYKLH-----------PHFSRILQETVKELAPQCDVTF---------MLSEDG---
----------SGKGAALITAVAKRLQQ--PWR-EN-----------------
>XP_060052799|Erinaceus europaeus|HKDC1
LYHMRLSDEILLDIMARFQAEMQKGL-DKDTNPTAS--VKMLPTFVRAIPDGSEHGEFLSLDLG--GSKF
RVLKVQVSEE---GK-RN--VQMESQFYPTPNEIIRGNSSE---------------LFEYVADCLADFMK
TK-----GLE---SKKLPLGLTFSFPCLQNKLE------------------------E------------
-----------GVLLSWTKKFKARGVQHTDVVNSLKKAIQKH----------K-----------------
-------------QDIDVDILALVNDTVGTMMTCAYDDPYCEVGVII-----------------------
-----G-----TGTNACYMEDMSNI-DLVEGDE--GRMCINTEWGAFGDDGALEDIRTEFDRELDLGSLN
PGKQLFEKMISGLYMGELVRLILLKMAKAGLLFGGKKSSALHTKGKIETRHVASMEK-------------
--YK--EGLANTREILTD-LGLEPSEADCIAVQHVCTIVSFRSANLCAAALAAILTRLQENK-KLERLRT
TVGMDGTVYKIH-PQYPKRLHKVVRRLVPNCDVRFLLSESGSTKGAAMVTAVACRLQAQRKQIDRVLALF
QLTQEQLVGIQDKMRAEFEYGLKRDTHPLA-TVKMLPTYVYGMPDGT-----------------------
---EKGKFLALDLGGTNFRVLLVKMRSG-RRSVRMYNKIFAIPLEIMQGTGEE-------LFDHIVQCIA
DFLDYMGLKGA-QLPLGFTFSFPCRQKNIDKGILIEWTKGFKATDCEGEDVVDMLREAIKRRN----EFD
LDIVA-----VVNDTVGTMMTCGYENPNCEIGLIAGTGSNMCYMEDM---KNIEMVD-------------
--GD------EG---------------------KMCINTEWGGFGDNSC-IDDIRTQYDKEVDEGSLNAG
KQR-YEKMTSGMYLGEIVRQILIDLTKQGLLFRGQISERLRTRGIFETKFLSQIES--------------
------------DRLALLQVRSILQQLGL-DSTCEDSIVVKEVCGAVSRRAAQLCGAGLAAIVEKRREDQ
GLEHLKITVGVDGTLYKLH-----------PHFSRILKETVKELAPQCDVAF---------MLSEDG---

----------SGKGAALITAVAKRLQQ--AQE--N-----------------
>XP_045883199|Meles meles|HKDC1
LYHMRLSDETLLDIMARFRAEMQKGL-GKDTNPTAS--VKMLPTFVRAIPDGSENGEFLSLDLG--GSKF
RVLKVQVSEA---GK-RK--VQMESQFYATPNEIIRGNGSE---------------LFDYIADCLADFMK
TR-----GLK---HKKLPLGLTFSFPCRQTKLE-----------------------E------------
-----------GILLSWTKKFKARGVQGTDVVSCLTKAIRKH----------K-----------------
--------------DIDVDILALVNDTVGTMMTCAYDDPYCEVGVII----------------------
-----G-----TGTNACYMEDMNNI-DLVEGDE--GRMCISTEWGAFGDDGALEDIRTEFDRELDLGSLN
PGKQLFEKMISGLYLGELVRIILLKMAKAGLLFGGKRSSALHTKGKIETRHVAAMEK------------
--YK--EGLANTREILTD-LGLEPSEADCIAVQHVCTIVSFRSANLCAAALAAILTRLRENK-KVARLRT
TVGVDGTLYKIH-PQYPKRLHKVVRRLVPNCDVRFLLSESGSTKGAAMVTAVASRVQAQRKQIDHVLALF
RLTREQLKDVQGKMRTEFENGLRKDTHLMA-TVKMLPTYVCGMPDGT----------------------
---EKGKFLALDLGGTNFRVLLVKIRSG-RRSVRMYNKIFAIPLEIMQGTGEE-------LFDHIVQCIA
DFLDYMGLKGA-RLPLGFTFSFPCRQTSIDKATLIEWTKGFKATDCEGEDMVDMLREAIKRRN----EFD
LDIVA-----VVNDTVGTMMTCGHEDPNCEIGLIAGTGSNMCYMEEM---RNIELLE------------
--GG------EG---------------------KMCINTEWGGFGDNGC-LDDIRTQYDKEVDEGSLNPG
KQR-YEKMTSGMYLGEIVRQILIDLTKQGLLFRGQISERLRTRGIFETKFLSQIES-------------
------------DRLALLQVRRILQQLGL-DSTCEDSIVVKEVCGAVSRRAAQLYGAGLAAIVEKRREDQ
GLEYLNITVGVDGTLYKLH-----------PHFSRILQETVKELAPRCDVTL---------ILSEDG---
----------SGKGAALITAVAKRLQQ--AQE-EN-----------------
>Q2TB90|Homo sapiens|HKDC1
LYHMRLSDDTLLDIMRRFRAEMEKGL-AKDTNPTAA--VKMLPTFVRAIPDGSENGEFLSLDLG--GSKF
RVLKVQVAEE---GK-RH--VQMESQFYPTPNEIIRGNGTE---------------LFEYVADCLADFMK
TK-----DLK---HKKLPLGLTFSFPCRQTKLE-----------------------E------------
-----------GVLLSWTKKFKARGVQDTDVVSRLTKAMRRH----------K-----------------
--------------DMDVDILALVNDTVGTMMTCAYDDPYCEVGVII----------------------
-----G-----TGTNACYMEDMSNI-DLVEGDE--GRMCINTEWGAFGDDGALEDIRTEFDRELDLGSLN
PGKQLFEKMISGLYLGELVRLILLKMAKAGLLFGGEKSSALHTKGKIETRHVAAMEK------------
--YK--EGLANTREILVD-LGLEPSEADCIAVQHVCTIVSFRSANLCAAALAAILTRLRENK-KVERLRT
TVGMDGTLYKIH-PQYPKRLHKVVRKLVPSCDVRFLLSESGSTKGAAMVTAVASRVQAQRKQIDRVLALF
QLTREQLVDVQAKMRAELEYGLKKKSHGLA-TVRMLPTYVCGLPDGT----------------------
---EKGKFLALDLGGTNFRVLLVKIRSG-RRSVRMYNKIFAIPLEIMQGTGEE-------LFDHIVQCIA
DFLDYMGLKGA-SLPLGFTFSFPCRQMSIDKGTLIGWTKGFKATDCEGEDVVDMLREAIKRRN----EFD
LDIVA-----VVNDTVGTMMTCGYEDPNCEIGLIAGTGSNMCYMEDM---RNIEMVE------------
--GG------EG---------------------KMCINTEWGGFGDNGC-IDDIWTRYDTEVDEGSLNPG
KQR-YEKMTSGMYLGEIVRQILIDLTKQGLLFRGQISERLRTRGIFETKFLSQIES-------------
------------DRLALLQVRRILQQLGL-DSTCEDSIVVKEVCGAVSRRAAQLCGAGLAAIVEKRREDQ
GLEHLRITVGVDGTLYKLH-----------PHFSRILQETVKELAPRCDVTF---------MLSEDG---
----------SGKGAALITAVAKRLQQ--AQK-EN-----------------
>XP_061084721|Conger conger|HKDC1
LYAMRLSDKQLQDISGRFRVEMEKGL-SAKSNATAT--VKMLPTHVLSTPDGSEKGEFLALDLG--GSSL
EVLRVKLSAD---RK-RN--VEMKSHTYSIPDAVINGRGTD---------------LFNHVARSLKDFME
RN-----HIS---NMKIPLGFTFSFPCRQSKLD-----------------------E------------
-----------GVLLSWSKNYNTRGVQGMDVVESLRKAINRT----------G-----------------
--------------GMDVEVLALVNDTVGTMMSCGYDDQCCEVGVTV----------------------
-----G-----TGTNACYMEELRHI-DQVEGDD--GRMCISTEWGAFGDDGVLNNFITDFDRENDAASSN
PGNQIFEKMVSGLYMGELVRIILLKMTKKGLLFNGQVSDTLRTKGKFQTKQIALIEE------------
--NE--DGLRNTKEILVD-LDLNPSEEDCIAVQHVCTIVSFRSANLCAAALAAILTRIRQNR-KLKSLRT
TVGMDGTVYRTH-PQYSRNLHEAVRRLVPDCRVEFMLSES-SGKGVAMVTAVAQRQVMQRQQISETLAAF
KLSPEQLQEVRDKMKIEMERGLRKDSHSTA-SVKMLPTYVYNMPDGT----------------------
---ETGKYLALDLGGTNFQVHVVKIRSG--ESVHTDNKVYVLPLEVMQGNQEE-------LFDHIVQCIS
DFLDCMGMKNT-SLPLGFTFSFPCRQTGIDKGNLVCWTKGFKAMDCEGNDVVDMLREAIKRRN----EFE

```
LDIVA-----VMNDTVGTMMTCAYDDSKCQIGLIAGTGSNVCYMEDM---QNIEMVE-------------
--GD------EG---------------------RMCVNTEWGGFGENGC-IDDIQTQFDREVDEGSLNIG
KQR-FEKMIGGMYLGEIVRKILIDLTRRDLLFRGEITEKLKTSGIFETKFLSQIES--------------
------------DRLALLQVRSILQHLGL-DSTCDDSIIVKQVCGTVSRRAAQLCGAGMAAIVEKIRENR
GLDHLDITVGVDGTLYKLH-----------PHFSRILNETVQELAPKCDVTF---------VLSEDG---
----------SGKGAALITDVAQRRYA---EM--------------------
> XP_035255802|Anguilla anguilla|HKDC1
LDAMRLSDEQLQDISAHFQAEMEKGL-SAKSNETAT--VKMLPTHVLSTPDGSEKGDFLALKLG--SSSL
KVLQVKVSED---GK-RK--VEMNSHTYPIPYGIINGRGTD---------------LFRHVAESLKDFME
KN-----HIN---NRKTPLGFTFSFPCRQSKLD------------------------E------------
-----------GVLLSWSKNYKARGVQGVDVVESLRQAIKQT----------G-----------------
--------------GMDVEVLALVNDTVGTMMSCGYDDQHCEVGVTV-----------------------
-----G-----TGTNACYMEELRHI-DLVEGDE--GRMCVSTEWGAFGDDGVLNDFITDLDREIDAPSSN
PGKQLFEKMVSGLYMGELVRIVLLKMIKKGLLFEGRASDALRTKGKFQTKHIALIEE-------------
--DK--EGLKNTKEILVD-LDLSPSEEDCIAVQRVCTIVSFRSANLCAAALAAILTRIRHNH-KLKTLRT
TVGVGGTVYRTH-PQYPKRLQEVVQRLVPDCHVRFMLSKSGSGKGVAMVTAVAQKLAIQRQQISETLAAF
KLSLAQLQEVRDKMRIEMERGLRKDTHSTA-SVKMLPTYIYSIPNGT-----------------------
---ETGKYLALDLGGTNFRVLVVNIRSGLHKSVHMDNKIYTVPQEIMQGTREE-------LIDHIVQCIS
DFLDYMGMKNT-CLPLGFTFSFPCRQNEIDKGSLVCWTKGFKATDCEGNDVVDMLKEAIKRRN----EFN
LDIVA-----VMNDTVGTMMTCAYEDHECEIGLIAGTGSNVCYMEDM---RNIETVE-------------
--GD------NG---------------------QMCVNTEWGGFGENGC-IDDIWTKFDKEVDEESLNVG
KQR-FEKMMSGMYLGEIVRHILIDLTRHGLLFRGHITDLLKTRGIFETKFLSQIES--------------
------------DRLALLQVRSILRRLGL-DSTCDDSIIVKEVCGTVSQRAAQLCGAGMAAVVDKIRENR
GLDHLDISVGVDGTLYKLH-----------PHFSRILRKTVRDLAPRCDVTF---------VLSEDG---
----------SGKGAALITTVAQQRYS---QM--------------------
> XP_046884627|Hypomesus transpacificus|HKDC1
LYAMRLSDEQLKDISARFMVEMEKGL-SNESNATST--VKMLPTHVRCTPDGSEKGEFLALDLG--GSKF
KVLQVRVCED---GRGGK--VEMDSETYPVPAELRSGRGSE---------------FFDHVAESLKDFMQ
KK-----NIM---QKKQPLGLTFSFPCLQSTVD------------------------Q------------
-----------GVLLSWSKNFKARGVQGTNVMESLRQAIQRA----------G-----------------
--------------VIDVDVLALVNDTVGAMMTCGYDDQRCEVGVIV-----------------------
-----G-----TGTNACYMEELRHI-DLVEGDE--GRMCINTEWGAFGDDGRLDDFITKFDREIDAASVN
PGKQLFEKMVSGLYLGELVRLILLKMAKEGLLFGGGLSDALRTKGKLETRHICMMEE-------------
--YK--DGLRNTREVLVD-LGLSPSDDDCLAVQHVCTLVSFRSSNLCAATLAAILTRIRTNR-KLKTLRI
TVGMDGTVYKTH-PQYARRLHKVVRRLVPDCHVRFLLSESGSGKGAAMVTAVAQRLAEQHAQMEATLEPF
RLTPEQLGLVRDRMRASLEQGLKRESKDSVKPVKMLPSYVFKTPDGT-----------------------
---EGGKFLALDLGGTNFRVLVVKIRSGIRRSVCMYNKIYAIPQEIMQGTGEE-------LFDHIVHCVS
DFLDYMGMKNT-RLPLGFTFSFPCQQAGIDKGTLVSWTKGFKATDCEGYDVVDMLREAIKRRN----EFD
LDIVA-----VVNDTVGTMMTCAYEDTRCEVGLIAGTGSNVCYMEEM---RNIEMVE-------------
--GD------EG---------------------QMCINTEWGGFGEDGC-IDDIRTRFDGEVDKGSLNIG
KQQ-FEKMTSGMYLGEIVRQILIDLTRRGLLFRGHVTETLNTRGIFETKFLSQIES--------------
------------DRLALLQVRSILQHLGL-DSTCDDSIIVKEVCGVVSHRAAQLCGAGMAAVVDKIRENR
GLATMDITVGVDGTLYKLH-----------PHFSRVLQETVKALAPQCEVNF---------VLSEDG---
----------SGKGAALITAVARQSCR---ET--------------------
> XP_048863144|Brienomyrus brachyistius|HKDC1
LHAMRLSDEQLLDISTRFRAQMEKGL-SAESNATAS--VRMLPTHVRSTPDGSEKGEFLALDLG--GSKF
KVLLVKVSED---GR-GK--VEMESQSYPLPEEVLGGRGTD---------------LFDHVAESLRDFME
RN-----KLDLSTHERRSLGFTFSFPCRQTKVD------------------------E------------
-----------GVLLSWSKNFKARGVLGTDVVESLRQAVDRV----------G-----------------
--------------DLKVNVLALVNDTVGTMMTCGFDDRNCEVGVII-----------------------
-----G-----TGTNACYMEEMRHI-DLVEGDE--GRMCINTEWGAFGDDGTLDDFITEFDREIDAASGH
PGKQTFEKMVSGMYLGELVRIVMLKMARRGLLFGGRVSDALCTKGKLQTAHVALMEM-------------
```

--YK--EGLRNTWNILME-LGLQPSEDDCVAAQHVCTIVSFRSANLCAAVLAAILTRIRENK-RLKSLRT
TVGMDGTVYRTH-PQYAKRLHKVVRRLVPECHVRFVLSESGSSKGAAMVTAVAGRLHSQRLQVRHILAPF
SLSAGQLQQVRDRMRVDLERGLREDTHTGT-PVKMLPSFVCTLPSGN-----------------------
---ECGKYLALDLGGTNFRVMVVKIRSGLRRSVRIYNKIYAIPLEIMQGTGEE-------LFDHIVQCIA
DFLDYMGMKNT-RLPLGFTFSFPCRQTGIDKGTLVSWTKGFKATDCEGHDVVDMLREAIKRKN----EFD
LDIVA-----VVNDTVGTMMTCAYEDPKCEIGLIAGTGSNVCYMEEM---RNIECME-------------
--GN------EG---------------------RMCVNTEWGGFGEDGA-LQDICTKFDQEVDEGSLNMG
KQR-FEKMTSGMYLGEIVRQILIELTSQGLLFRGRITERLKTPGIFETRFLSQIES--------------
------------DRLALLQVRAILQQLGL-DSTCDDSIIVKEVCGVVSRRAAQLCGAGLAAVVDKIRENR
GLDHLNITVGVDGTLYKLH-----------PHFSTILKETVNALAPLCDVTF---------VLSEDG---
----------SGKGAALITAVACSLCR---GV--------------------
> XP_065152912|Paramisgurnus dabryanus|HKDC1
LYAMRLSNDQLVDISARFRLEMEKGL-SSESTAAAT--VKMLPTHVYSTPDGSERGEFLALDLG--GSKF
KVLQVKVSDN---GK-RS--VQMESETYPIPEEIMNGRGSE---------------LFEHVAESLKAFLL
KH-----HVN---QNRKPLGLTFSFPCAQSKIN------------------------E------------
-----------GVLLSWSKNFKARGVQGSNVVQSLKDAMNEV----------G-----------------
--------------GVNVDVLAIVNDTVGTMMTCGYDDHNCEVGVII-----------------------
-----G-----TGTNACYMEEMRYI-DMVEGDE--GRMCINTEWGAFGDDGALDDFITDFDREIDAASIN
PGKQLFEKMISGMYMGELVRLILLKMAKKGLLFNGQVTDDLRTQGKFHTSYICSIEQ-------------
--YK--SGLENTRDILKQ-LGLNPSDDDCIAVQHVCTIVSFRSANLVAAALAAILTRIRENK-KLKTLRT
TVGVDGTVYRTH-PQYPKRLHKVVRYLVPDCHVRFVLSESGSAKGAAMVTAVAQRLAAQRQKVDDTLAAF
ALDFTQLQEIKEKMHVELERGLKKETHSTA-SVKMLPTYVYRTPDGT-----------------------
---ERGKFLALDLGGTNFRVLVVKLRTGIHNSVRMYNKIYAIPLEIMQGTGEE-------LFDHIVQCIS
DFLDYMGMKNT-CLPLGFTFSFPCSQSGIDKGSLVSWTKGFKATDCEGTDVVDLLREAIKRRN----EFD
LDIVA-----LVNDTVGTMMTCAYEDPKCQIGLIAGTGSNVCYMEEM---KNIEFLD-------------
--GD------EG---------------------QMCINTEWGGFGENEN-IEDIRTRYDREVDEGSLNVG
KQR-FEKMTSGMYLGEIVRQILIDLTKQGLLFRGRITERLKTKGIFETKFLSQIES--------------
------------DRLALLQVRSILQHLGL-DSTCDDSIIVKEVCGAVSRRAAQLCGAGMAAVVDKIRENR
GLDHLEITVGVDGTLYKLH-----------PHFSTILKDTVRQLAPKCKVDF---------ILSEDG---
----------SGKGAALITAVARQHKG--DKQ--DVR---------------
> XP_041125226|Polyodon spathula|HKDC1
LYHMRLSDEQLRDIMMRFRAEMEKGL-SRATNPTAA--VKMLPTHVRSTPDGSEKGEFLALDLG--GSKF
KVLRVKVSDN---GN-QR--VEMESQHYPTPEEITSGKGTG---------------LFDHVAECLKDFME
RN-----HVN---TKRHPLGFTFSFPCSQTKLD------------------------E------------
-----------GVLLSWSKHFKARGVQGTDVVDSLRKAIDKT----------G-----------------
--------------GMDVDVLALVNDTVGTMMTCGYDDKYCEVGAII-----------------------
-----G-----TGTNACYMEEMRHI-DLVEGDE--GRMCINTEWGSFGDDGSLEDIRTKFDRDIDLGSIN
PGKQLFEKLISGMYMGELVRIILVKMARRGLLFQGRMSDALRTKGKFQTQHVAEIEN-------------
--YK--EGLSNAKTILAG-LGLSPSEDDCIAVQHVCTIVSFRSANLCAAALAAILTRLQENK-KLKRLRT
TVGMDGTVYRTH-PQYAKRLHKVVRRLVPDCNVRFLLSESGSGKGAAMVTAVAYRLVSQRKQIDETLAPF
KLSRQHLQDVKDKMRVELERGLKKDTHHSA-SVKMLPSYVCGTPDGT-----------------------
---ERGKFLALDLGGTNFRVLLVKIYTGLRKSIRMYNKIYAIPLEIMQGTGEE-------LFDHIVHCIS
DFLDYMGMKNA-RLPLGFTFSFPCRQAGIDKGSLVSWTKGFKATDCEGQDVVDMLREAIKRRN----EFD
LDIVA-----VVNDTVGTMMTCAYEDPNCEIGLIAGTGSNVCYMEEM---RNIETVE-------------
--GE------EG---------------------RMCVNTEWGGFGDNDC-IEFIRTKFDREVDEGSLNQG
KQR-YEKMTSGMYLGEIVRQILIDLTKRGLLFRGQISERLRTRGIFETKFLSQIES--------------
------------DRLALLQVRSILQQLGL-DSTCDDSIIVKEVCGAVSRRAAQICGAGMAAVADKIRENR
GLEQLDITVGVDGTLYKLH-----------PHFSKVLKETVKDLAPQCDVTF---------VLSEDG---
----------SGKGAALITAVAQKLHS---HS--------------------
> XP_018604952|Scleropages formosus|HKDC1
LYAMRLSDDQLLDISARFRAEMVKGL-SAESNATAT--VRMLPTHVRSIPDGSERGEFLALDLG--GSKF
KVLRVKVSEN---GK-RK--VDMESETYPIPEEVAHGRGTD---------------LFNHVAESLRDFMQ

```
KI-----DVN---QKKMPLGFTFSFPCRQSRLD------------------------E------------
-----------GVLLSWSKNFKARGVQGTDVVESLREAIDRV----------G-----------------
--------------DIDVNVLALVNDTVGTMMTCGFDDQRCEVGLIV-----------------------
-----G-----TGTNACYMEEMRHI-DLVEGDE--GRMCINTEWGAFGEDGALDDFITSFDQEIDAASLN
PGKQLFEKMISGMYMGELVRIILLKMARSGLLFGGRVSDALCTKGKLETRHVVLMEK-------------
--YK--EGLQNTKNILTD-LDLTPTEDDCVAVQHVCTIVSFRSANLCAAALAAILTRMRESR-KLQTLRT
TVGVDGTVYRTH-PQYAKRLHKVVRRLVPECHVRFVLSESGSGKGAAMVTAVAQRLLAQRREVDQTLAPF
VLMAKQLYQVRDRMRRELENGLRLETHATA-SVKMLPSYVYNTPSGR-----------------------
---EHGNYLALDLGGTNFRVLVVKVRSSRRRSVRMYNKIYAIPLEIMQGTGEE-------LFDHIVQCIS
DFLDYMGMKNT-CLPLGFTFSFPCRQTGIDKGTLVSWTKGFKATDCEGNDMVEMLREAIKRRN----EFD
LDIVA-----LVNDTVGTMMTCAYEDPKCEIGLIVGTGCNVCYMEEM---TNIETAE-------------
--GD------EG---------------------HMCINTEWGGFGENGE-LDDIRTRFDREVDEGSLNVG
KQT-FEKMTSGMYLGEIVRQILIDLTRRGLLFRGRITERLKTPGIFETKFLSQIES--------------
------------DRLALLQVRSILQHLGL-DSTCDDSIIVKEVCGTVSRRAAQLCGAGMAAVVEKIRENR
GLDRLEITVGVDGTLYKLH-----------PHFSSILKETVGELAPKCNVSF---------VLSEDG---
----------SGKGAALITAVARSLGD--RPV--TL----------------
>XP_030627404|Chanos chanos|HKDC1
LYAMRLSDDQLKDISARFRMEMEKGL-SRESNATAA--VKMLPTHVRSIPEGSEKGEFLALDLG--GSKF
KVLRVDVSAD---GK-KK--VHMESETYPIPGEILNGRGAD---------------LFNHVAMSLKAFME
KH-----KIS---ERKTPLGFTFSFPCAQSKID------------------------E------------
-----------GVLLSWSKHYKARGVVGVNVVQSLRQAIRKQ----------R-----------------
--------------DVDVDVLALVNDTVGAMMTCGYDDQQCEVGVII-----------------------
-----G-----TGTNACYMEEMRHI-DLVEGDE--GRMCINTEWGAFGDDGALDDFITDFDREIDAASIN
PGKQLFEKMVSGMYLGELVRLILLKMAKKGLLFQGRITNALRTQGKFLTKHVCLIEE-------------
--YK--DGLKNTRQILRD-LGLHPTADDCIAVQHVCTIVSFRSANLCAAALAAILNRIRENK-KLKSLRT
TVGVDGTVYRTH-PQYAKRLHKVVRRLVPDCYVRFVLSESGSGKGAAMVTAVAQRRASQRKEIDDTLASF
RLSLEQLQKVRDKMKVELERGLRKDSHGSA-SVKMLPTYVYRTPDGT-----------------------
---ESGNYLALDLGGTNFRVLLVKIRSGLRKSVRMYNKIYAIPLEIMQGTGEE-------LFDHIVQCIS
DFLDYMGMKNT-CFPLGFTFSFPCRQTGIDKGSLVNWTKGFKATDCVGHDVVDMLREAIKRRN----DFD
LDIVA-----VVNDTVGTMMTCAYEDPNCEIGLIAGTGSNVCYMEEM---CNIETVE-------------
--GD------EG---------------------QMCVNTEWGGFGGNDN-IEDIRTKYDHEVDTGSLNVG
QQR-FEKMTSGMYLGEIVRQILIDLTKRGLLFRGRITEPLKTRGIFETKFLSQIES--------------
------------DRLALLQVRAILQHLGL-DSTCDDSIIVKEVCGAVSRRAAQLCGAGMAAVVDKIRENR
RLDHLHITVGVDGTLYKLH-----------PHFSGILKETVKELAPDCTVDF---------VLSEDG---
----------SGKGAALITAVARSSQN---KT--------------------
>MBN3295896|Amia calva|HKDC1
---MRLSDDQLLDIAARFQAEMEKGL-SAESNPRAA--LKMLPTHVRSTPDTSEKGEFLALDLG--GSKF
KVLQVKVSED---GK-KR--VEMESKMYPNPEEILSGKGTD---------------LFDYVAACLKDFMQ
SK-----HIN---QKKQPLGFTFSFPCRQSTVD------------------------E------------
-----------GVLLSWSKHFKARGVQGKDVVNLLRKAIDRA----------G-----------------
--------------GIDVDVLALVNDTVGTMMTCGYDDQHCEVGVII-----------------------
-----G-----TGTNACYMEELRHI-DLVEGDE--GRMCINTEWGAFGDDGSLDDFITDFDREIDAASIN
PGKQLFEKMISGLYLGELVRLILLKMTKKGLLFKGRASDALRTKGKFETRHISAIEK-------------
--YK--EGLNNTREILTA-LGLQPSEDDCIAVQHVCTIVSFRSANLCAAALSAILARIRENK-KVSRLRT
TVGVDGTVYRTH-PQYAKRLHKVVRRLVPDCHVRFVLSESGSGKGAAMVTAVAQRLVSQRKQIDQTLAPF
LLSRQQLQQVRDKMRVELERGLKKDTHHTA-SVKMLPSYVYRTPDGT-----------------------
---EGGKFLALDLGGTNFRVLVVKIRSGLRRSVRMYNKIYAIPLEIMQGTGEE-------LFDHIVHCIS
DFLDYMGMKNT-TLPLGFTFSFPCRQTGIDKGTLVNWTKGFKATDCEGHDVVDMLREAIKRRN----EFD
LDIVA-----VVNDTVGTMMTCAYEDPQCEIGLIAGTGSNVCYMEEM---RNIETVE-------------
--GD------EG---------------------RMCVNTEWGGFGENGC-IEDIRTKFDREVDEGSLNIG
KQR-FEKMTSGMYLGEIVRQILIDLTRRGLLFRGRVTERLKTRGIFETKFLSQIES--------------
------------DRLALLQVRAILQQLGL-DGTCDDSIIVKEVCSTVSRRAAQLCGAGMAAVVDKIRENR
```

GLDKLDITVGVDGTLYKLH-----------PHFSSILKETVQDLAPQCEVNF---------VLSEDG---
----------SGKGAALITAVARKYKE--PLR--PAV---------------
>MBN3326727|Atractosteus spatula|HKDC1
LYAMRLSDDQLLDMAARFRVEMEKGL-SSESCATAA--VKMLPTHVRSTPDGSEKGEFLALDVG--GSKF
KVLQVKVSDD---GK-RT--VEMDSITYPTPEEIRSGKGTD---------------LFDHVAESLKDFMQ
RK-----HIN---QKKQPLGITFSFPCRQSKLD------------------------E------------
-----------GVLLSWSKNFKARGVRGTDVVELLREAIDRL----------G-----------------
--------------GIDVDVLALVNDTVGTMMTCGYDDQNCEVGVIV-----------------------
-----G-----TGTNACYMEEMRHI-DLVEGDE--GRMCINTEWGAFGDDGSLNDFITDFDREIDAASIN
PGQQLFEKMISGLYMGELVRLILLKMTKKGLLFNGQASDALRTKGKFETKYVCAIEK-------------
--YK--EGLSNTKEILTS-LGLDPSEQDCIAVQHVCTIVSFRSANLCAAALAAILTRIRENK-RLKVLRT
TVGVDGTVYRTH-PQYAKRLHKVARRLVPDCHVRFLLSESGSSKGAAMVTAVAQRLVSQHRQIAETLAPL
RLSHQQLRDVRDKMRVELERGLRRDTHHTA-SVKMLPTYVYRTPNGT-----------------------
---ERGKFLALDLGGTNFRVLVVKIRSGLRRSVRMYNKIYAIPLEIMQGTGEE-------LFDHIVQCIS
DFLDYMGMKNS-RLPLGFTFSFPCRQAGIDKGTLVSWTKGFKATYCEGNDVVDMLREAVKRRN----EFD
LDIVA-----VVNDTVGTMMTCAYEDPNCEIGLIAGTGSNVCYMEEM---KNIEVVD-------------
--GD------EG---------------------RMCVNTEWGGFGENGC-IEDIRTKFDREVDEGSLNVG
KQK-FEKMTSGMYLGEVVRQILIDLTRRGLLFRGRITERLKTRGIFETKFLSQIES--------------
------------DRLALLQVRAILQQLGL-DSTCDDSIIVKEVCGAVSRRAAQICGAGMAAVVDKIRENR
GLDKLDITVGVDGTLYKLH-----------PHFSTSLRETVQELAPQCSVAF---------VLSEDG---
----------SGKGAALITAVARGLHT--QA---------------------
>XP_017319568|Ictalurus punctatus|HKDC1
LYAMRLSDEQLLNIKSRFCAEMEKGL-SKESNVAAT--VKMLPTHVRSTPDGSEKGEFLALDLG--GSKF
KVLQVKVSED---GK-RK--VQMESETFPVPEEILNGRGTD---------------LFDHVAESLKTFMQ
KN-----NIN---QKKKPLGFTFSFPCLQSKLD------------------------E------------
-----------GMLLSWSKNYKARGVKGTNVVQLLRQAINNV----------G-----------------
--------------GIDVDVLALVNDTVGTMMTCGYDDQHCEVGVII-----------------------
-----G-----TGTNACYMEEMRHI-DLVEGDE--GRMCINTEWGAFGDDGALDDFITSFDQEIDAASIN
PGKQLFEKMVSGMHMGELVRLILLKMAKRGLLFKGKVSDALRTKGRFQTKHVCMIEE-------------
--YK--DGLKNTKDILTE-LGLSPTEDDCIAVQHVCTIVSFRSANLCAATLAAILTRIRENK-KLKTLRT
TVGVDGTVYRIH-PQYAKRLHKVVRRLVPECHVRFVLSESGSGKGAAMVTAVAQRLASQRKEIDNTLAEF
TLSHTQLQQVKDKMRVELERGLRKDSHGSA-SVKMLPSYVYRTPDGT-----------------------
---ERGKFLALDLGGTNFRVLVVKIRTGIRKSVRMYNKIYAIPLEIMQGTGEE-------LFDHIVHCIS
DFLDYMGMKNT-CLPLGFTFSFPCRQNGIDKGILVSWTKGFKATHCEGNDVVDMLREAIKRRN----EFD
LDIVA-----VVNDTVGTMMTCAYEDPNCEIGLIAGTGSNVCYMEHM---KNIETVD-------------
--GD------EG---------------------HMCVNTEWGGFGDNDN-IEDIRTRFDREVDSGSLNVG
KQR-FEKMTSGMYLGEIVRQILIDLTKRGLLFRGHITECLKTRGIFETKFLSQIES--------------
------------DRLALLQVRTILQNLGL-NSTCNDSIIVKEVCGVVSHRAAQLCGAGMAAIVDKIRENR
GLDHLDITVGVDGTLYKLH-----------PHFSRILKDTVKELAPECEVDF---------VLSEDG---
----------SGKGAALITAVARQYCK---KT--------------------
>XP_026856323|Electrophorus electricus|HKDC1
LYAMRLSDEQLCDISSRFHVEMEKGL-CGESNATAT--VKMLPTHVCSTPDGSEQGEFLALDLG--GSRF
KVLQVKVSED---GK-RK--VQMESETFPVPEEVLNGRGTD---------------LFGHVAESLKAFMQ
KN-----DIN---HKRKPLGFTFSFPCLQSKLD------------------------E------------
-----------GVLLSWSKNYKARGVQGTDVVQLLRQAINNV----------G-----------------
--------------GTDVDVLALVNDTVGTMMTCGYDDQHCEVGVII-----------------------
-----G-----TGTNACYMEELRHI-DLVEGDE--GRMCINTEWGAFGDDGALDDFITSFDREIDAASIN
PGKQLFEKLVSGMYMGEIVRLVLLKMAKAGLLFKGRMSDALRTKGKFHTEHVCLIEE-------------
--YK--DGLKNTRSILTE-LGLSASEEDCIAVQHVCTIVSFRSANLCAATLAAILTRMRQNK-KLKSLRT
TVGVDGTVYKTH-PQYAKRLHKVVRRLVPECNVRFVLSESGSGKGAAMVTAVAQRLASRRRDVDDTLAGF
ALSTAQLQQVKEKMRRELELGLRKESHGSA-SVKMLPSYVYRTPDGT-----------------------
---ERGKFLALDLGGTNFRVLVVKIRSGIRKSVRMYNKIYAIPLEIMQGTGEE-------LFDHIVQCIS

DFLDYMGMRNT-CLPLGFTFSFPCRQTGIDKGILVNWTKGFKATDCEGNDVVDMLREAIKRRN----EFD
LDIVA-----VVNDTVGTMMTCAYEDPKCEIGLIAGTGSNVCYMEEM---QNIESVE------------
--GD------QG---------------------RMCVNTEWGGFGDNAN-IEDIRTRFDREVDEGSFNVG
KQR-FEKMTSGMYLGEIVRQILIDLTRRGLLFRGRITEPLKTRGIFETKFLSQIES-------------
------------DRLALLQVRSILQHLGL-DSTCDDSIIVKEVCGTVSRRAARICGAGMAAVVDKIRENR
GLDHLDITVGVDGTLYKLH-----------PHFSRILKETVKELAPKCNVDF---------MLSEDG---
----------SGKGAALITAVARQHFR---NT--------------------
>XP_028844788|Denticeps clupeoides|HKDC1
LYAMRLSDDQLVDVSTRFRVEMEKGL-SCESNATAA--VKMLPTHVRSTPDGSEKGEFLALDLG--GSRF
KVLQVKVSEN---GK-RK--VEMESETYPIPDEIFNGRGTD---------------LFSHVAECLKDFME
KK-----NMN---QKKKPLGFTFSFPCAQSKLD------------------------E------------
-----------GVLLSWSKNFKSRGVQGTNVVHSLRQAIRRV----------G-----------------
--------------DIDVDVLALVNDTVGTMMTCGYDDHHCEVGVIV-----------------------
-----G-----TGTNACYMEEMRHI-DLVEGDE--GRMCINTEWGAFGDDGTLDDFVTSFDKEIDAASIN
PGKQLFEKMVSGMYMGELVRLILLKMAKKGLLFGGMVSDALRTKGKLQTKHICSIEE------------
--YK--NGLSNTKGILTE-LGLRPSDDDCIAVQHVCTIVSFRSANLCAAALAAILTRIRENR-KVKTLRT
TVGVDGTVYRTH-PQYSKRLHKVVRRLVPDCHVRFVLSESGSGKGAAMVTAVAQRLLSQRKQVDDTLAHF
SLNRDQLRVVRDKMRVELERGLRKDSHSSA-SVKMLPAFVFQTPDGT----------------------
---ERGKYLALDLGGTNFRVLVVKIRSGIRKSVRMYNKIYAIPHEIMQGTGEE-------LFDHIVHCIS
DFLDYMGMKNT-RLPLGFTFSFPCRQTGIDKGSLVSWTKGFKATDCEDHDVVDMLRDAIKRRN----EFD
LDIVA-----VVNDTVGTMMTCAYEDSKCEIGLIAGTGSNVCYMEDM---RNIETVE------------
--GD------EG---------------------QMCVNTEWGGFGENGC-IEDIRTKFDREVDEGSLNSG
KQR-FEKMTSGMYLGEIVRQILIDLTKRGLLFRGRITEPLKTRGIFETKFLSQIES-------------
------------DRLALLQVRCILQQLGL-DSTCDDSIIVKEVCGAVSHRAAQLCGAGMAAIVEKIRENR
GLEHLEITVGVDGTLYKLH-----------PHFSRVLKDTVRDLAPKCTVSF---------VLSEDG---
----------SGKGAALITAVARRYCG--GAQ--QQVN--------------
>XP_036402269|Megalops cyprinoides|HKDC1
LYAMRLSDDQLLDISARFRAEMEKGL-NGESNPTAA--VKMLPTHVLSTPDGSEKGEFLALDLG--GSRF
KVLQVKVSED---GK-QK--VEMESQTYPIPEEVVNGRGAD--------VSFFYTFLFNHVAESLKDFMQ
KN-----HIN---QKKKPLGFTFSFPCRQSKLD------------------------E------------
-----------GVLLSWSKNYKARGVQGTNVVESLRQAIARA----------G-----------------
--------------GMDVEVLALVNDTVGTMMTCGYDDQHCEVGVIV-----------------------
-----G-----TGTNACYMEELRHI-DLVEGDE--GRMCINTEWGAFGDDGSLDDFITDFDREIDAASIN
PGKQLFEKMVSGMYMGELVRLILLKMSKQGLLFEGRASDALRTKGKFQTKHVALIEK------------
--YK--EGLKNTRDILQD-LGLNPSEDDCVAVQHVCTIVSFRSANLCAAALAAILTRLRENR-KQKTLRT
TVGVDGTVYRTH-PQYPKRLHKVVRRLVPECHVRFVLSESGSGKGAAMVTAVAQRLVAQRREVDDTLAPF
ALTRQQLQQVRDRMREGLERGLKKDTHSTA-SVKMLPSYVYRTPDGT----------------------
---ETGKYLALDLGGTNFRVLVVKIRTGLRKSVRMYNKIYAIPLEIMQGTGEE-------LFDHIVQCIS
DFLDYMGMKNT-CLPLGFTFSFPCRQTGIDKGSLVSWTKGFKATDCEGNDVVDMLREAIKRRN----EFE
LDIVA-----VVNDTVGTMMTCAYEDPECEIGLIAGTGSNVCYMEEM---RNIETVE------------
--GD------EG---------------------RMCVNTEWGGLGENGC-LEDIRTKYDREVDEGSLNVG
KQR-FEKMTSGMYLGEIVRQILTDLTKRGLLFRGHITECLKTRGIFETKFLSQIES-------------
------------DRLALLQVRSILKNLGL-DSTCDDSIIVKEVCGAVSRRAAQLCGAGMAAVVEKIRENR
GLDHLNVTVGVDGTLYKLH-----------PHFSRILKDTVRELAPKCDVTF---------VLSEDG---
----------SGKGAALITAVAR------GET--------------------
>XP_031435294|Clupea harengus|HKDC1
LYAMRLSDEQLKDISARFRAEMEKGL-SGESNATAA--VRMLPTHVCSTPDGSEKGEFLALDLG--GSRF
KVLQVKVSEN---GK-RK--VQMESETYPIPEEVLNGRGTE---------------LFDHVAESLKAFMQ
RN-----GIN---QKKKPLGFTFSFPCAQSRLN------------------------E------------
-----------GVLLSWTKHFKARGVQGTNVAQSLRQAINRV----------K-----------------
--------------DIDVDVLALVNDTVGTMMTCGYDDQRCEVGVIV-----------------------
-----G-----TGTNACYMEELRHI-ELVEGDE--GRMCINTEWGAFGDDGALDDYITGFDREIDAASSN

AEKQLFEKMVSGLYMGELVRLILLKMAKKGLLFGGVISDSLRTKGKFQTKHVCQIEQ-------------
--YK--DGLKNTKQILTE-LGLQPSTDDCVAVQQVCTIVSFRSANLCAAALAAILTRIRENR-KLKSLRT
TVGMDGTVYRMH-PQYAKRLHKVVRRLVPDCHVRFVLSESGSGKGAAMVTAVAQRLMSQRQEIDDTLATF
RLDQEQLQQVRDRMRAELERGLKKDSHSTA-SVKMLPSYVCRTPDGS-----------------------
---ETGKYLALDLGGTNFRVLVVKIRTGLRKSVRMYNKIYAIPQEIMQGTGEE-------LFDHIVQCIS
DFLDYMGMKNT-RLPLGFTFSFPCRQTGIDKGTLVSWTKGFKATDCVGNDVVEMLREAIKRRN----EFD
LDIVA-----VVNDTVGTMMTCAYEDPKCQIGLIAGTGSNVCYMEEM---RNIETAE-------------
--GD------EG---------------------QMCVNTEWGGFGENLEDIEGIRTKYDQEVDEGSLNNG
KQR-FEKMTSGMYLGEIVRQILIDLTSRGLLFRGHITEPLKTRGIFETKFLSQIES--------------
------------DRLALLQVRCILQQLGL-DSTCDDSIIVKEVCGAVSHRAAQLCGAGMAAVADKIRENR
GLDHLDVTVGVDGTLYKLH-----------PHFSRILKDTVKELAPKCDVEF---------VLSEDG---
----------SGKGAALITAVAKSFNG---ET--------------------
>XP_030231928|Gadus morhua|HKDC1
LYSMRLNDDQLRDVSARLLQEMQDVL-SSQSSAAES--IKMLPTHVCSTPDGSENGDFLALDMG--GSRF
KVLRVKVRQQ---VG-KKGGVEMEEKTYSIPKELLNGKGSE---------------LFDHVSKSLKDFAD
TN-----NID--LEKKHPLGFTFSFPCHQNTLN------------------------Q------------
-----------GVLLNWTKNYRARGVQGKDVTRALMESIQRT----------G-----------------
--------------GMDVELLALVNDTVGTMMTCAFDDPYCEVGLII-----------------------
-----G-----TGTNACYMEELRHI-NLVEGDE--GRMCVNTEWGAFGDDGALKDYITDFDRDIDAASAT
PGKQIFEKLISGLYLGELVRLVVLKMAKQGLLFEGHAPDALRTKGKITTAHVASMEQ-------------
--YK--DGLRNTRAILED-LGLAPSDDDCVAVQHVGTVVSFRSSNLVAAGLAAILTRVRQNR-KLKTLRI
TVGVDGTVYRTH-TQYAKRLHKVVRRLVPDCHVRFVLSESGSSKGAAMVTAVAQRLASQRRQVDEALSSL
KLTQEHLGYVKSRMRAGLEAGLKSQGPS---PVKMLPAYVYRTPDGT-----------------------
---EHGNYLALDLGGTNFRVLLVKIRSGLSRSVRLYNKIYAIPLEVMQGTGEE-------LFDHIVQCIS
DFLDYMGMKNT-RLPLGFTFSFPCYQKAIDTGTLVSWTKGFKATGCQGNDVVDMLREAIKRRN----EFD
LDIVA-----IVNDTVGTMMSCAYEDPQCEIGLIAGTGSNVCYMEEL---SNVEKLHVPQNGKEGRNGGK
VEEECTEAQAKDVIGREAGEEEEEEEEQSGPNVRMCINTEWGGLGEDGC-LDDIITPYDSAVDRDSLNQG
RQR-FEKMTSGLYLGEIVRQVLLDLTRRGLLFRGHVTEALKTPGIFHTKYLSQIES--------------
------------DRLALLQVRSILQQLGL-ESTINDSIVVKEVCGAVSHRAAQICGAGMAAVVDKIRENR
GLDHLDVTVGVDGTLYKLH-----------PHFAGVMQQTVNKLAPQCNVNF---------LLSEDG---
----------SGKGAALITAVGCRLRE--ELK--SQ----------------
>XP_056135122|Lampris incognitus|HKDC1
LYSMRLHDDQLRDISARFLGEMTKGL-SSESNATAT--VKMLPTHVRSTPDGSEKGEFLALDLG--GSKF
KVLQVKVREEAGKNS-RR--VEMEEQTYPMPKELLFGKGVE---------------LFDHVSECLKDFLH
QK-----NIS--LDKKHPLGFTFSFPCQQTALN------------------------Q------------
-----------GILINWSKNFRARGLQGKDVVQALRQAINRT----------G-----------------
--------------DMDVDVLALVNDTVGTMMTCGFDDQHCEVGLII-----------------------
-----G-----TGTNACYMEQLCHI-GLVEGDE--GRMCVNTEWGAFGDDGALDDYITEFDREIDAASTN
PGKQLFEKLIGGLYLGELVRLIVLKMTKLELLFNGNASSALKVKGKITTAHVAAMEE-------------
--YK--NGLRNTRDILLN-LGLTPSHDDCIAVQQVSTVVSFRSSNLVAAGLAAILNRIRQNR-KLKTLKI
TVGVDGTLYRTH-PQYAKRLHKVVRRLVPDCHVRFILSESGSSKGAAVVTAVAQRLASQRRQVEETLSPF
RLSQEQLGLVKVRMRAGLDAGLRNVSA----PVKMLPAFVYRTPDGT-----------------------
---EHGTYLALDLGGTNFRVLLVKIYSGLRRSVRLYNKIYSIPLEIMQGTGEE-------LFDHIVQCIS
DFLDYMGMKNV-HLPLGFTFSFPCKQSGINTGTLVSWTKGFKATGCEGNDVIDMLRDAIKRRN----EFD
MDIEA-----IINDTVGTMMSCAYEDPQCEIGLIAGTGSNVCYMEEL---RNIKKIA----TEEGSMEGG
EKKEQEEWRAEGVEKKEDKGMEGEGKQGNGERKRMCVNTEWGGLGEDGC-LDDIITHYDIEVDQASLNHG
KQR-FEKLTSGLYLGEIVRHVLLDLTRRGLLFRGYVTEALKTPGIFETKYLSYIES--------------
------------DHLALLQVRAILQKLGL-DSTCDDSIIVKEVCETVSHRAAQLCGAGLAAVVDKIRENR
GLEHLDITVGVDGTLYKLH-----------PHFARVLQATVRELAPQCCVTF---------VLSEEG---
----------SGKGAALITAVAIQQK---VEI--------------------
>XP_029904144|Myripristis murdjan|HKDC1
LYAMRLHDDQLRDISARFQAEMKKGL-SAESNAAAT--LKMLPTHVCSTPDGSENGEFLAVDLG--GSKF

KVLQVKVRKEAGRVR-RS--VEMQEQTYPLPKELLIGRGVE---------------LFDHVSECLKDFLL
QK-----KIS--LEKKHPLGFTFSFPCQQTALN------------------------Q------------
-----------GLLLNWSKNYWARGLQGKDMVNTLRQAISRA----------G-----------------
--------------GMDVDVLALVNDTVGTMMTCGFDDQHCEVGLII-----------------------
-----G-----TGTNACYMEELRHI-DLVEGDE--GRMCVNTEWGAFGDDGALDDYLTEFDRDIDAASIN
PGKQCLEKMVSGLYLGELVRLVVLKMTKLGLLFDGRVTDALRTKGKITTAHVAAMEQ-------------
--YK--DGLKNTRDILLD-LSLNPSPDDCIAVQHVCTVVSFRSSNLVAAGLAAILTRIRQNR-KLKTLRI
TVGVDGTVYKTH-PQYAKRLHKVVRRLVPDCHVRFVLSESGSSKGAAMVTAVAQRVTARRHQVDESLSSF
RLSRDQLRSVKARMRAGLEAGLKSNSHS---PVKMLPSYVFRTPDGT-----------------------
---ECGKYLALDLGGTNFRVLLVKIRSGLRRSVRLYNKLYPIPLEIMQGTGEE-------LFDHIVHCIS
DFLDYVGMKKA-RLPLGFTFSFPCLQAAINVGTLVSWTKGFKATDCEGNDVVDMLREAIKRRN----EFD
VDIVA-----IVNDTVGTMMSCAYEDPQCEIGLIAGTGSNVCYMEEL---RNIEKVK-MEEGEPERKQDS
DGED-----------KTDDGMDREAKNGLGEMQRMCINTEWGGLGEDGC-LDDIITEYDNAVDRESINHG
RQR-FEKLTSGLYLGEVVRQVLLDLTRRGLLFKGRVTEALKTPGIFETKYLSQIES--------------
------------DRLALLQVRSILQHLGL-DSTCDDSIIVKEVCGAVSRRAAQLCGAGMAAVVDKIRENR
GLDQLNITVGVDGTLYKLH-----------PHFSTVLQETVVALAPQCYVTY---------LLSEEG---
----------SGRGAALITAVARRDQ---VEM--------------------
>XP_029950878|Salarias fasciatus|HKDC1
LHKMRLNDEQLNDISARFQAEMKKGL-SAESNAAAA--VKMLPTHVRSTPDGTEKGQFLALDLG--GSKF
KVLQIKVREDQGIRK-GG--VEIEEKTYPIPNEVLNGRGTE---------------LFDHVSESLKDFMH
EK-----NIN--PKKKHPLAFTFSFPCEQPTLD------------------------Q------------
-----------GFLLEWTKDFSVRGLQGKDVVDSLRQAIKRT----------G-----------------
--------------GMDMEILAMVNDTVATMMTCGFDDKNCEVGVII-----------------------
-----G-----SGTNACYMEQLNHV-DLVEGDE--GQMCVSTEWGALGDDGTLDDYITEFDREVDACSIN
PGKQIFEKMVSGMYLGELVRLVVLKMAKLGLLFDGRVTNALRTKGTITTAHVAAMEE-------------
--YK--NCLGHTREILES-LNLSPSADDCIAVQHISTIVSFRSSNLVAAGLAAILTRIQKNR-NTRELRI
TIGVDGTVYKTH-PQYPKRLHKVVRRLLPECHVRFVLSESGSSKGAALVSAVAQRLASRKRQVDETLNSF
KLSWEQLQLVKARMRAALQAGLKTDGPS---EVKMLPSFVYRTPDGS-----------------------
---EHGKYLALDLGGTNFRAMLVNFKKGLQQNTKLYHKIYKIPLEIMQGTGEE-------LFDYLAQCVG
DFLDYMGMKNT-RLPAGFTFSFPCEQTALDTATLLSWTKGFKATNCEGHDVVDMLREAIKRRD----EFD
LDIVA-----LVNDTVGTLMSCAYEDPQCQVALIAGTGSNVCYMEEV---KNIEKSK-EYMRKPNGVQDL
RDGD------EK---KVKGG---NGQHSDADFSKMCINTEWGGLGDDGT-LDDIMTSDDTEVDQNSINPG
KQR-FEKLTSGMYLGEIVRRVLLNLTIRGLLFKGYVSEALKTPGIFETKYLSQIES--------------
------------DRLALLQVRSILQHMGL-EGTCVDIITVKEVCGAISRRAAQLCGAGMAAVVDKIRENR
GLDRLNITVGVDGTLYKLH-----------PHFSRILQETTRALAPQCDVTF---------LPSEEG---
----------SGKGAALIAALAKHMQ--------------------------
>XP_011490494|Oryzias latipes|HKDC1
LHAMRLNDDQICEISARFLAEMRKGL-SSESNAAAA--VKMLPTHVRSTPDGSEKGQFLALDLG--GSKF
KVLQVKVREGVGVRK-GE--VEMEEKIYPMPTELRTGTGAE---------------LFDHVSECLKEFLH
EK-----KID--TKKKHPLAFTFSFPCEQPSLD------------------------Q------------
-----------GFLLSWCKNYRVRGLQGKDVVRTLRDAISRA----------G-----------------
--------------EMDMEVLAMVNDTVATMMTCGFDDPYCEVGLII-----------------------
-----G-----TGTNACYMEDMRHV-DLIEGDE--GRMCINTEWAGFGDDGALNDFTTEFDKEVDAASSN
PGKQIFEKMVSGMYLGELVRLVVLKMAKLGKLFDGHVSDALRTKGKITTADVAAMEE-------------
--YR--NGLKNTKDILMH-LGLSPSHDDCIAVQHVSTIVSFRSSNLVAAGLSAILTRIRQNR-NVRVLRV
TVGVDGTVYKTH-PQYPKRLHKVVRRLLPECHVRFVLSDSGSSKGAALVSAVAQRLTSQSRKVDEILSNF
KLSEEHQQLVKDRMRAGLEAGLRSQGPF---TIKMLPSFIFHKPNGK-----------------------
---EHGKFLALDLGGTNFRALLVRFK---KQKSRIYHKIYTIPLEIMQGTGEE-------LFDHLAQCVC
DFLDYMGVKNA-RLPTGFTFSFPCEQTAIDTGKLVTWTKGFNATDCEGHDVVGLLREAIKRRN----DFD
LDIVA-----LINDTVGTMMSCAYEDPRCEIGLIAGTGTNLCYMEEL---KNVKKLH-QKSGTAEKDQET
PEVE------NG---KETDGETRR--------SNMCINTEWGGLGDDGS-LDNIITAFDTEVDQNSVNPG
KQR-FEKLTSGMYLGEIVRQVLLDLTRRGLLFRGHISECLKKPGIFETRYLSQIES--------------

------------DHLALLQVRSVLQKLGL-DSTCHDSIIVKQVCGAVSRRAAQLCGAGMAAVVDKIRQNR
GLDHLNTTVGVDGALYKLH-----------PHFSQILQETVKVLAPQCDVTF---------FPSEEG---
----------SGKGAALVAAVARQNQE--MLT--------------------
>XP_028298855|Gouania willdenowi|HKDC1
LYAMRLNDGQLTDISARFQAEMKKGL-LRESNAAAA--VKMLPTHVRSTPDGSENGRFLALDLG--GSKF
KVLQVKVREGRGLRK-GQ--VEMEEKIYPVPVELLSGRGAE---------------LFDHVAESLKDFLQ
EK-----KIS--LDQKHPLAFTFSFPCELPSLH------------------------Q------------
-----------GFLLSWSKNYRVRGFQGKDVVHALRQAIYRT----------G-----------------
--------------GVDVEVLAMVNDTVATMMTCGFDDQYCEVGLII-----------------------
-----G-----TGTNACYMEEMRHV-DLVEGDE--GRMCVNTEWGAFGDDGALKDFITEFDEDVDAASVN
PGKQIFEKMVSGLYLGELVRLVVVKMAKLGKLFDGCVSEALRTKGKITTAHVAAMEE-------------
--YK--TGLKNTREILMG-LGLSPSPDDCIAVQHVSTIVSFRSSNLVAAGLVAILNRIKKSR-GVRGLRI
TIGVDGTVYKTH-PQYPKRLHKVVRRLLPECNVRFVLSESGSSKGAALVAAVAQRLASRKRQVDQRLSPF
RLSEEHLQLVKARMRVGLDAGLKATGPS---AVKMLPSFVYHTPDGT-----------------------
---EHGKYLALDLGGTNFRALLVKFKNGRQHNSRLYHKIYTIPLEIMQGTGEE-------LFDYLAHCVG
DFLDYMGMKNV-RLPTGFTFSFPCEQSAIDTGKLVSWTKGFKATDCEGHDVVNMLREAIKRRN----DYD
LDIVS-----IVNDTVGTMMSCAYEDPECEVALIAGTGSNVCYMEEL---RNIKKVK--QSTHKTEDVNT
DGGE-----EKG---------TEKTTNHTADISRMCVNTEWGGLGDDGS-LDDIITSYDTAVDQNSINPG
KQR-FEKLTSGMYLGETVRHVLLDLTREGLLFRGNVTETLKTPDIFETKYLSQIES--------------
------------DRLALLQVRSILQKLGL-DSTCDDGIIVKEVCGAISQRAARLCGAGMAAVVDKIRENR
GLDRLTVTAGVDGALYKLH-----------PHFSQVLKETTRVLAPQCNVTF---------LPSEEG---
----------SGKGAALITAVAKKRQL-------------------------
>XP_038150506|Cyprinodon tularosa|HKDC1
LHAMRLHDDRLSDISKRFLAEMKKGL-SAESTAAAA--VKMLPTQVRSTPDGSEKGEFLALDLG--GSNF
KVLRVKVREDVGIKK-GG--VEMEEKTYPIPNELHAGKAAE---------------LFDHVSESLKDFLH
EK-----NIS--PSNKHLLAFTFSFPCEHPSLD------------------------Q------------
-----------GLSVSWGKNYRVRGLQGKDVVQALKEAIGRT----------G-----------------
--------------DMDVEVLAMVNDTVATMMTCGFDDQCCEVGLII-----------------------
-----G-----TGTNVCYMEELRHI-DLVEGDE--GRMCVNTEWGGFGDDGALDDFITEFDRDVDAASLN
PGQQTFEKMVSGMYLGELVRLVVLKMAKLGLLFDGRVSDALRTKGNITTADVAAMEE-------------
--NR--NGLKNTKDILTR-LHLSPSPDDCIAVQHVSTIVSFRSSNLVAAGLAAILTRIRQNR-NLRGLKI
TVGVDGTLYKTH-PQYPKRLHKVVRRLLPDCQVRFALSESGSSKGAALVAAVAQRLTSQRRRVDEALSLF
RLNQQQLQLVKSRMREGLEAGLRGKGC----TVKMLPSFVYHTPDGT-----------------------
---ENGNYLALDLGGTNFRAMLVRFK---RHKSQLYHKIYTIPLEIMQGTGEE-------LFDHLAQCVC
DFLDYMGLKKA-CLPAGFTFSFPCEQSGINKGTLVSWTKGFKATDCVGQDVVFMLRDAIKRRN----EFD
LDIVA-----LVNDTVGTMMSCAYEDPQCEIGMIAGTGSNLCYMEEV---KNIEKLK-QGHGTAKKDK--
---E------QE---E-NHAMKPEKSNVEADESKMCINTEWGGLGDDHC-LDDIINPYDAEVDIHSVNPG
KQR-FEKMTGGMYLGEIVRQVLLDLTRRGLLFKGHVTEPLKTPGIFETKYLSQIES--------------
------------DRLALLQVRSILQRLGL-RGTSGDSIIVKEVCGAVSRRAAQLCGAGMAAVVDKIREDR
RLERLSITVGVDGALYKLH-----------PHFSQVVKETVRVMAPQCDVTF---------LPSEEG---
----------SGKGAALIAAVVQSINN-------I-----------------
>XP_021163457|Fundulus heteroclitus|HKDC1
LHAMRLQDDQLIDISKRFLAEMKKGL-SAESTAAAA--VKMLPTHVRSTPDGSEKGQFLALDLG--GSKF
KVLRVKVREDVGIRK-GG--VEMEEKIYPIPKELHVGRAAE---------------LFDHVSESLKDFLH
EK-----NIG--LEKKHPLAFTFSFPCEHPSLD------------------------Q------------
-----------GLLVSWCKNYKIRGLQGKDVVQALRQSIGRA----------G-----------------
--------------DMDVEILAMVGDTVATMMTCGFDDQCCEVGLII-----------------------
-----G-----TGTNACYMEELRHI-DLVEGDE--GRMCVNTEWGAFGDDGALDDHITEFDRDVDAASLN
PGKQIFEKMISGMYLGELVRLVVLKMAKLGLLFDGCVSDALRTKGKITTADVAAMEE-------------
--YR--NGLNNTRNILTH-LHLDPSPDDCIAVQHVSTIVSFRSSNLVAAGLAAILTRIGQNR-NVRGLRI
TVGVDGTVYKTH-PQYPKRLHKVVRRLLPECQVRFVLSESGSSKGAALVAAVAQRLTSQRRKVDEALSLF
TLNQQQLQLVKSRMREGLEAGLKSKGC----SVKMLPSFVYHTPDGT-----------------------

---EYGKYLALDLGGTNFRAMLVRFK---RQKSRLYHKIYTIPLEIMQGTGEE-------LFDHLAQCVC
DFLDYMGLKKA-RLPAGFTFSFPCEQSGVDTGTLVSWTKGFKATDCEGQDVVSMLREAIKRRN----EFD
LDIVA-----LLNDTVGTMMSCAYEDPQCEIGMIAGTGSNLCYMEQL---KNIEKIK-EGGGITRTDTEA
LEGE------ED----EDVVTEAEKSVLEAGKSKMCINTEWGGLGDDRC-LDDIITPYDAEVDIHSVNPG
KQR-FEKLTSGMYLGEIVRQVLLDLTRRGLLFKGNVTETLKTPGIFETKYLSQIES--------------
------------DRLALLQVRSILQHLGL-RGTCDDSIVVKEVCGAVSRRAAQLCGAGMAAVVDKIREDR
RLERLSITVGVDGALYKLH-----------PHFSQVLKETVKVMAPQCDVTF---------LPSEEG---
----------SGKGAALIAAVVRQKQE-------M-----------------
> XP_007549861|Poecilia formosa|HKDC1
LHAMRLQDEQLVDISSRFLAEMKKGL-SAESLAAAA--VKMLPTHVRSTPDGSEKGQFVALDLG--GSKF
KVLRVKVREDEETNK-GG--VEMEEKTYPMPKELYEGRAVE---------------LFDHVSESLKDFLH
EK-----NMS--PEKKHPLAFTFSFPCEHPGLD------------------------Q------------
-----------GLLLSWCKNYKVRGLQGKDVVQFLRQAIDRT----------G-----------------
--------------DMDIDVLAMVNDTVATMMTCGFDDQFCEVGLII-----------------------
-----G-----TGTNACYMEELRHI-DLVQGDE--GRMCVNTEWGAFGDDGALDDFITDFDRDVDAASLN
PGKQIFEKMVSGLYLGELVRLVVLKMAKLGLLFDGHVSDALRTKGKITTGDVAAMEE-------------
--YR--NGLKNTKEILTR-LHLTPSPDDCVAVQHVSTIVSFRSSNLVAAGLVAVLTRIRQNR-NLRGLRI
TVGVDGTVYKTH-PQYPKRLHKVVRRLLPDCQVRFVLSESGSSKGAALVAAVAQRLISQRRKVDEALSLF
TLNQEKLQLVKSRMREGLEAGLKSKDC----TIKMLPSFVYHTPNGT-----------------------
---ERGKYLALDLGGTNFRAMLVRFK---KQKTRLYHKIYTIPLEIMQGTGEE-------LFDHLAQCVC
DFLDYMGLKKV-RLPAGFTFSFPCQQSDIDTGTLVSWTKGFKATDCEGEDVVSMLREAIKRRN----DFD
LDIVA-----LVNDTVGTMMSCAYEDPKCEVGMIAGTGSNLCYMEEL---RNIKKLD-QSNSILKTNKQT
SEGE------EG-------------EAAVKSGLKMCVNTEWGGLGNDRC-LEDIITPYDAEVDIHSVNPG
KQR-FEKLTSGMYLGEIVRQVLLDLTRRGLLFKGHVTETLKTPGIFETKYLSQIES--------------
------------DRLALLQVRSILQQLGL-HGTCDDSIIVKEVCGAVSRRAAQMCGAGMAAVVDKIREDR
GLDRLSITVGVDGALYKLH-----------PHFSQVLKETVGLMAPQCDVTF---------LPSEEG---
----------SGKGAALIAAVVQQKQE----M--------------------
> XP_054588160|Nothobranchius furzeri|HKDC1
LHAMRLNDDQLNELSARFQAEMKKGL-SAESNAAAA--VKMLPTHVCSTPDGSEKGQFLALDLG--GSKF
KVLQVKVRECVGIKK-GD--VEMEEKIYPVPDELLTGRATE---------------LFDHVSESLKDFMH
EK-----NIS--LEKKHPLAFTFSFPCDQSRLD------------------------Q------------
-----------GTLLNWSKNYKARGLQGKDVVQALREAVERT----------E-----------------
--------------GLDVEILAMVNDTVATMMTCGFDDSYCEVGIIV-----------------------
-----G-----TGTNACYMEELRHI-DLVEGDE--GRMCVNTEWGSFGDDGALKDFLTEFDREVDAASSN
PGKQIFEKMVSGMHLGELVRLVVLKMAKLGKLFDGRVSDALRTKGKITTADVAAMDE-------------
--YR--NGLKNTKDVLDE-LDLNPSDDDCIAVQHVSTIVSFRSSNLVAAGLAAILTRIRQNR-NMGALRI
TVGVDGTVYKTH-PQYPKRLHKVVRRLLPECQVRFVLSESGSSKGAALVAAVAQRLTSLRRQVSETLSLF
RLKEEQLQLVKARMRSGLDAGLRSKSSS---TIKMLPSFVYLTPDGT-----------------------
---ERGKYLALDLGGTNFRALLVRFK---KQTSRLYHKIYTIPLEIMQGTGEE-------LFDHLAQCVC
DFLDYMGLKNT-RLPAGFTFSFPCEQSAIDTGTLVAWTKGFKATDSEGHDVVSMLREAIKRRN----DID
LDIVA-----LINDTVGTMMSCAHEDQRCEIGMIAGTGTNLCYMEEL---KNIEKIE---QRATKTEEKT
QKGE------KN------ENAGAEKNKKDGEMQKMCINTEWGGLGDDGC-LDDIITLYDTEVDQNSINQG
KQR-FEKMTSGMYLGEVVRQVLLDLTRRGLLFKGHVSETLKTPGIFETKYLSQIES--------------
------------DRLALLQVRSILQQLGL-HGTCHDSIIVKHVCLVVSRRAAQLCGAGMAAVVDKIRENR
GLDHLDIAVGVDGSLYKLH-----------PHFSQVLKETVRVMAPQCDVTF---------QPSEEG---
----------SGKGAALIAAVARRKQD-------V-----------------
> XP_061600571|Cololabis saira|HKDC1
LHAMRLQDDQLSDISARFQAEMKKGL-SIESNAAAA--VKMLPTYVRSTPDGSEKGQFLALDVG--GSKF
KVLQVKVREGVGIKK-GE--VEMAEKIYPLPKELQTGKGVE---------------LFDHVSESLKDFLH
EK-----NIS--LEKKHPLAFTFSFPCEHRCLD------------------------Q------------
-----------GILLSWCKNYRARGLQGKDVVQVLRDAISRA----------G-----------------
--------------DMDMEVLAMVNDTVATMMTCGFDDEYCEVGLII-----------------------

-----G-----TGTNACYMEERRHV-DLVAGDE--GRMCVNTEWGAFGDDGALDDFITEFDRDVDAASAN
PGKQIFEKMVSGMYLGELVRLIVLKMAKLGLLFDGNISDALRTKGKITTAHVAAMEE-------------
--YR--HGLKNTIDLLMQ-LDLTPSADDCIAVQHVSTIVSFRSSNLVAAGLAAVLTRIKQNR-NIRDLRI
TVGVDGTVYKTH-PQYPKRLHKVVRQLLPECQVRFVLSDGGSSKGAALVTAVAQRLAAQSRKVKETLSAF
TLSHDQLQMVKARMRAGLEAGLKSQGPF---TIKMLPSYVYHTPDGT-----------------------
---QRGKYLALDLGGTNFRALLVTFKGQ---TSRLYHKIYTIPLVIMQGTGEE-------LFDHLADCVC
DFLDYMGMKNV-RLPAGFTFSFPCKQTAVDTGTLVSWTKGYKATDCEGHDVVKMLREAIKRRK----EFD
LDIVA-----LVNDTVGTMMSCAYEDPQCEIGLIAGTGSNLCYMEDL---KNIEKIE-QRRGTTETDEGT
LDGQ------ED-----EDCCQAEKTEEDAEIPKMCINTEWGGLGDDGS-LNDVITAFDKEVDQHSVNPG
KQR-FEKMTSGMYLGEVVRHVLLDLTKRGLLFRGHVTETLKRHGKFETKYLSQIES--------------
------------DNLALLEVRSILQKLGL-DSTCDDSIIVKEVCGAVSRRAAQLCGAGLAAVVDKIRENR
ELDHLNITVGVDGALYKLH-----------PHFSQILQNTVKVMAPQCDVTF---------LPSEEG---
----------SGKGAALIAAVTQTTAG-DVTE--------------------
> XP_063738053|Eleginops maclovinus|HKDC1
LYTMHLHDDQLSDISACFQAEMKKGL-SAESNAAAA--VKMLPTHVHSTPDGSEKGQFLVLDLG--GSKF
KVLQVKVREGTGIGR-GG--VDIEEKIYPIPEELLTGSGKE---------------LFDHVSESLKDFLH
DK-----NIS--VENKLPLAFTFSFPCEQSTLN------------------------Q------------
-----------GLLLHWSKNFRARGVQGKDVVQALRGAVDRT----------G-----------------
--------------GMDVEVLALVNDTVATMMTCGFDDRYCEVGLIV-----------------------
-----G-----SGTSACYMEELRHI-DLVEGDE--GRMCVNTEWGAFGDDGALNDYITEFDREVDAASNN
PGKQILEKMVSGMYLGELVRLVVLKMAKLGLLFEGSVSDALRTKGKITTAHVAAMEE-------------
--YK--NGLKNTRNILVD-LDLSPTPDDCIAVQHVSTTVSFRSSNLVAAGLAAILTRIRQNR-HLRTLRI
TVGVDGTVYKTH-PRYANRLHKVVQRLVPDCHVRFVLSDGGSSKGAALVTAVAERLASQRQQVDETLSPL
RLSREKLLLVKKRMRGALEAGLKSKGPF---AVKMLPSYVYQTPDGT-----------------------
---EHGKYLALGLGRTNFRALLVDLNKGLQQNSRLNHKIYPIPVEITQGSGEE-------LFDHIAQCVG
DFLDYMGMKGA-RLPAGFTFSFPCKQTGIDTGTLVSWTKGFKATDCEGHDVVSMLREAIKRRN----DFD
LDIVS-----VVNDAVGTMMSCAYEDPQCEIGLIVGTGTNVCYMEEL---KNIEKVE-QEKGKEER----
-EGE------EE---KQDDGAERQKKEEDAGIPKMCINTEWGGLGDDGS-LDDIITHYDMEVDQNSLNPG
KQR-FEKLISGMYLGEVVRQVLLDLTSRGLLFRGQVTETLKTPGIFQTKYLSQIES--------------
------------DRLALLQVRSVLQQLGL-DSTCDDSIIVKEVCGAVSHRAAQLCGAGLAAVVDKIRENR
GLDHLDIPVGVAGSLYKLH-----------PHFSGFLQETVKMLALQCNVTF---------LPSEGG---
----------SGTGAALTAAVARKTLE-------M-----------------
> XP_056269604|Pseudoliparis swirei|HKDC1
LSAMRLHDDQLSDISARFQAEMKKGL-SEDSNAAAA--MKMLPTHVRSTPDGSEKGEFLVLDLG--GSKF
KVLQVKVREGTGITR-GG--VEMEEKTYPIPKELHIGSGAE---------------LFDHVSESLKDFLH
EK-----NIS--LEKKHPLAFTFSFPCEQTTVN------------------------Q------------
-----------GLLLNWSKNFRARGLQGTDVVQALRGSIDRT----------G-----------------
--------------GMDVEVLAVVNDTVATMMTCGVDDQYCEVGLIV-----------------------
-----G-----TGTNACYMEELRHV-GLVEGDE--GRMCVNTEWGAFGDDGALEGYITEFDRDVDAASNN
PGKQILEKMVSGMYLGELVRLVVLKMAKLRLLFHGHVSDALRTKGTITTEHVAAMEE-------------
--DK--TGLKNTRNILVD-LDLTPSSDDCVAVRHVGTVVSFRSSNLVAAGLAAILTRIKQNR-NLRTFRI
TVGVDGSVYKRH-PQFPRRLHEVVRRLLPECDVRFVLSDSGSSKGAALVAAVAQRLASRRLQVDETLSPF
RLSQEQLQVMKTRMRMGLEAGLKSKGPS---ALKMLPSFVYRTPDGT-----------------------
---EHGKYLSLDLGARNFRAMLVNFKQGLQPNSPLYHKMYPVPPEVMQGSGEE-------LFDLIAQCVS
DFLDYTGMKNA-RLPACFIFSFPCEQTAIDTGSLVSWTKGFHATGCEGRDVVSMLREAIERRN----EFD
LDIAA-----VVNDTVATMMSCACEDALCEIGLIAGSGTNVCYMEEL---KNIEKSE-DKTGQMER----
--GE------ED---K-DDGRERERT----EIPKMCVNTEWGGLGDDGS-LDDIITPYDIEVDRNSLHPE
KQR-FEKLTSGMYLGEVVRRVLLDLTRGGLLFRGHVTEALKTPGMFQTKYLSQIES--------------
------------GRLALLQVRSILQQLGL-DCTCDDSIIVKEVCGAVSRRAAQLCGAGMAAVVDKIRENR
GLDHLNITVGVDGTLYKLH-----------PHFSGILQETARVLAPRCDMTF---------LPSE-G---
----------RWKGAALIAAMARQQLE-------M-----------------
> XP_054464542|Anoplopoma fimbria|HKDC1

LYAMRLHDDQLSDISARFQAEMKRGL-SVGSNVTAA--VKMLPTHVRSTPDGSEKGQFLALDLG--GSRF
KVLQVKVREG--MGN-KRGGVEIVQKTYPIPKELHVGKATE---------------LFDHVSESLKDFLH
EK-----NIS--LEKKHPLAFTFSFPCEQSTLN------------------------Q------------
-----------GFLMNWSKNFRARGLQGKDVVQALRDAIDRT----------G-----------------
--------------GMDVEVLAMVNDTVATMMTCGFDDQYCEVGLII-----------------------
-----G-----TGTNACYMEELRHV-DLVEGDE--GRMCMNTEWGGFGDDGALNDYITEFDRDVDAASNN
PGKQILEKMVSGMYLGELVRLAVLKMAKLRLLFNGHVSDALRTRGTITTEHVAAMEE------------
--YK--NGLKNTRNLLMD-LDLSPTSDDCVAVQHISTIVSFRSSNLVAAGLAAILTRIKQNR-NLRRLRI
TVGVDGSVYKTH-PQYPKRLHKVVQRLLPECHVRFVLSDSGSSKGAALVSAVAQRLASQRRKVDETLSPF
RLSQEQLQLVKTRMRAGLEAGLKSKGPS---AIKMLPSFVYHTPDGT-----------------------
---ECGKHLVLDLGGTNFRALLVNCNQGLQQNSQLYHKIYTIPLEIMQGSGEE-------LFDHIVQCVS
DFLDYIGMKNT-RLPASFIFSFPCEQTAIDTGTLVSWTKGFKATDCEGHDVVSMLREAIKRRN----EFD
LDIAA-----VVNGTVATMMSCAYENPQCEIGLIAGTGTNVCYMEEL---KNIEKTE-------QKTEKN
ERGE---GTPEGEKDEQDDGREIERQEGHAEIPKMCINTEWGGLGDDGS-LDDIITPCDIEVDQNSLNPG
KQR-FEKLMSGMYLGEVVRQVLLDLTRGGLLFRGHVTEALKTPGMFQTKYLSQIES--------------
------------DRQALLQVRSILQQLGL-DSTCDDSIIVKEVCGVVSRRAAQLCGAGMAAVVDKIRENR
GLDHLNITVGVDGKLYKLH-----------PHFSGILKETARVLAPQCNVTF---------LPSEEG---
----------SGKGAALITAAARQKLE-------M-----------------
>XP_049421211|Epinephelus fuscoguttatus|HKDC1
LHAMRLHDDQLSDISARFQAEMKKGL-SAESNAAAA--VKMLPTHVRSTPDGSEKGQFLVLDLG--GSRF
KVLQVKVREGMGIRR-GG--VEMEEKTYPIPKELLNGKGTE---------------LFDHVSESLKDFMH
EK-----NIS--LEKKHPLALTFSFPCEQSALN------------------------Q------------
-----------GLLLDWSKNYQARGLQGKDVVQALRGAIERT----------G-----------------
--------------GMDVEVLAMVNDTVATMMTCGFDDQYCEVGLIV-----------------------
-----G-----TGTNACYMEEMRHV-DLVEGDE--GRMCVNTEWGGFGDDGALNDYITEFDREVDAASNN
PGKQIMEKMVSGMYLGELMRLVVLKMAKLGLLFDGHLSEALKTKGTITTAHVAAMEE------------
--YK--TGLKNTINILTD-LDLNPSSDDCAAVQHISTIVSFRSSNLVAAGLAAILTRIKQNR-KLRKFRI
TVGVDGTVYKTH-PQYAKRLHKVVRRLLPDCQVRFVLSDNGSSKGAALVTAVAQRLASRRRQVDETLSPF
KLTPEQLQLVKSRMRAGLEAGLKSKGPS---AIKMLPSFVYRTPDGT-----------------------
---EHGKYLALDLGGTNFRALLVNLTKGQQPNTQLYHKIYTIPLEIMQGTGEE-------LFDHLAQCVS
DFLDYMGMKNA-HLPAGFTFSFPCEQSALDTGTLVSWTKGFKATDCEGQDVVSMLREAIKRRN----EFD
LDIVA-----VVNDTVGTMMSCAYEDPQCEIGLIAGTGTNVCYMEEL---KNIEKTE-QKKGRLDREEGT
PEGE---DWEDG--------TEREGEEGDAEISKMCVNTEWGGLGDDGS-LDDIITPYDIEVDKNSVNPG
KQR-FEKLTSGMYLGEVVRQVLLDLTSGGLLFRGHVTEALKTPGIFETKYLSQIES--------------
------------DRLALLQVRAILQQLGL-DSNCDDSIIVKEVCGAVSRRAAQMCGAGMAAVVEKIRENR
GLDHLDITVGVDGALYKLH-----------PHFSIVLQETARVMAPQCNITF---------LPSEEG---
----------SGKGAALITAVAKQKPE-------M-----------------
>XP_044202873|Thunnus albacares|HKDC1
LYAMRIHDDQLTDISARFQAEMKKGL-SAESNAAAA--VKMLPTHVRSTPDGSEKGQFLALDLG--GSKF
KVLQVKVREGMGIRR-GG--VEMEEKTYSIPKELLIGKGTE---------------LFDHVSESLKDFLH
EK-----NIS--LEKKHPLAFTFSFPCEQSTLN------------------------Q------------
-----------GLLLNWSKNYRARGLQGKDVVQVLREAVDRT----------G-----------------
--------------GMDVEVLAMVNDTVATMMTCGFDDRYCEVGLII-----------------------
-----G-----TGTNACYMEELRHV-DLVEGDE--GRMCVNTDWGAFGDDGALKDFITEFDRDVDAASNN
PGKQLFEKMVSGMYLGELVRLVVLKMAKLGLLFDGHVSNALKTKGKITTAHVAAMEE------------
--YK--NGLKNTRDILTE-LDLTPSPDDCIAVQHVSTIVSFRSSNLVAAGLAAILTRIKQNR-NLRKLRI
TVGMDGTLYKTH-PQYPKRLHKVVRRLLPECHVRFVLSDSGSSKGAALVAAVAQRLASQRRKVDETLSPF
RLNQEQLQLVKARMRAGLEAGLKSKISS---GIKMLPSFVYHTPDGT-----------------------
---ERGKYLALDLGGTNFRTLLVDFKKGLQQNSRLYHKIYTIPLEIMQGTGEE-------LFDHIVQCIS
DFLDYMGMKNA-HLPAGFTFSFPCEQTAIDTGTLVSWTKGYKATDCEGHDVVGMLREAIKRHN----EFD
LDIVA-----VVNDTVGTMMSCAYEDPQCEIGLIAGTGSNLCYMEEL---KNIEKIK------------
--QE------EGKMEREEGTPERQGDMQDDEIRRMCINTEWGGLGDDGS-LDDIITPYDVEVDQNSVNAG

KQR-FEKLTSGMYLGEVVRQVLLDLTRGGLLFRGHVTEPLKTPGIFQTKYLSQIES-------------
------------DRLALLQVRSILQQLGL-DSTCDDSIIVKEVCGAVSRRAAQLCGAGMAAVVDKMRENR
GLDHLNITVGVDGALYKLH-----------PHFSKVLQETTSVLAPQCDVTF---------VPSEEG---
----------SGKGAALITAVARQKQD-------M-----------------
>XP_029314653|Cottoperca gobio|HKDC1
LYAMRLHDDQLSDISACFQAEMKKGL-SAESNAAAA--VKMLPTHVRSTPDGSENGQFLALDLG--GSRF
KVLQVKVREGTGIRK-GG--VEMEENTYPIPEELLTGRGTE---------------LFDHVSESLKDFLH
EK-----NIS--LEKKHPLAFTFSFPCEQSALN-----------------------Q-----------
-----------GLLLNWSKNCRARGLQGKDVVQALRGAIDRT----------G-----------------
--------------GMDVEVLAMVNDTVATMMTCGFDDQHCEVGLIV----------------------
-----G-----TGTNACYMEELRHV-DLVEGDE--GRMCVNTEWGAFGDDGALNDYITEFDRDVDAASNN
PGKQILEKMVSGLYLGELVRLVVLKMAKLGLLFDGHVSDALRTKGKITTAHVAAMEE------------
--YK--DGLKNTRNILVD-LDLTPTSDDCVAVQHVSTIVSFRSSNLVAAGLAAILTRIKQNR-NLRTLRI
TVGVDGTVYKTH-PRYPNRLHEVVRRLLPECDVRFVLSDSGSSKGAALVTAVAQRLASRRRKVDETLSPF
RLSQEQLQLVKTRMRAGLEAGLKSKGPS---AIKMLPSFVYSTPDGT----------------------
---EHGKYLALGLGGTNFRALLVNFKKGLQQNSQLHHKIYTIPLEIKQGSGEE-------LFDHIAQCVS
DFLDYMGMKNA-RLPAGFTFSFPCEQTAIDRGTLVNWTKGFKATDCEGHDVVSMLREAIKRRN----DFD
VDIVA-----VVNDTVGTMMSCAYEDPQCEIGLIAGTGTNVCYMEEL---KNIEKTE----QKKEEEEEM
SEGE------EG----KDDGTERERVEGDAKTPKMCINTEWGGLGDDGA-LEDFITLYDIEVDRNSLNPG
KQR-FEKLTSGMHLGEVVRQVLLDLTRGGLLFRGHVTEALKTPGIFQTKYLSQIES-------------
------------DRLALLQVRSILQQLGL-DSTCDDSIIVKEVCGAVSHRAAQLCGAGMAAVVDKIRENR
GVDHLNITVGVDGTLYKQH-----------PHFSGVLQETAKVLSPRCNVTF---------LPSEGG---
----------SGKGAALITA---RQKM---EL--------------------
>XP_059202062|Centropristis striata|HKDC1
LYAMRLHDDQLSDISARFQAEMKKGL-SAESNVAAA--VKMLPTHVRATPDGSEKGQFLALDLG--GSRF
KVLQVKVREGMGIRR-GG--VQMEEKVYPIPKELLNGRATE---------------LFDHVSESLKNFLQ
EK-----KIS--LEKKHPLAFTFSFPCEQPTLN-----------------------Q-----------
-----------GLLSNWSKDYRARGLQGKDVVQTLESAIGRT----------G-----------------
--------------GMDVEVIALVNDTVATMMTCGFDDQYCEVGLII----------------------
-----G-----TGTNACYMEELRHV-DLVEGDE--GRMCVNTEWGAFGDDGALKDYITEFDRDVDAASNN
PGKQIFEKMVSGMYLGELVRLVVLKMAKLGLLFDGQVSDALKTKGRITTANVAAMEE------------
--YK--NGLKNTEKILLD-LDLKPTCDDCTAVQHVSTIVSFRSSNLVAAGLAAILTRIRQNR-NLRTFRI
TVGVDGTVYKTH-PQYPKRLHKVVRRLLPECHVRFVLSDSGSSKGAALVTAVAQRLASRRRQVDETLAPF
RVSQEQLQLVKTRMRAGLEAGLKRKGSS---AIKMLPSFVYRTPDGT----------------------
---EHGKYLALDLGGTNFRALLVNFKKGPQQNTRLYHKIYTIPEEIMQGSGEE-------LFDHIAQCVS
DFLDYMGMKNA-RLPAGFTFSFPCEQTSIDTGTLVSWTKGFKATNCEGNDVVSMLREAIKRA-----EFD
LDIVA-----VVNDTVGTMMSCAYEDPECEIGLIAGTGTNVCYMEEL---KNIKKTE-QKTEQVEREEGT
PDGE------ED---KHDAWTEREKKEGDAEIPKMCINTEWGGLGDDGS-LDDIITPYDTEVNQNSINPG
KQR-FEKLTSGMYLGEVVRQVLLDLTRGGLLFRGHVTEALKTPGIFQTQYLSQIES-------------
------------DRLALLQVRSILQHLGL-DSTCDDSIIVKEVCGAVSRRAAQLCGAGMAAVVDKIRENR
GLDRLNITVGVDGALYKLH-----------PHFAGVLQETARALAPQCNVNF---------LPSEEG---
----------SGKGAALITAVARQNPE-------M-----------------
>XP_037650464|Sebastes umbrosus|HKDC1
LYAMRLHDDQLSDISARFQAEMKKGL-SRESNAAAA--VKMLPTHVHSTPDGSEKGQFLALDLG--GSKF
KVLQVKVREGMGIRR-GG--VEMVEKTYPIPKELLIGRGAE---------------LFDHVSESLKDFLH
EK-----KIS--LEKKHPLAFTFSFPCEQSTLN-----------------------Q-----------
-----------GLLLNWSKNYRARGLQGKDVVQALRGAIDRT----------G-----------------
--------------GMDVEVLAMVNDTVATMMTCGFDDQYCEVGLII----------------------
-----G-----TGTNACYMEELRHV-DLVEGDE--GRMCVNTEWGAFGDDGALDDYITEFDRDVDAASNN
PGKQILEKMVSGMYLGELVRLVVLKMAKLGLLFDGHVSNALKTKGRITTANVAAMEE------------
--YK--NGLDNTRKILMG-LDLTPTDDDCVAVQHVSTVVSFRSSNLVAAGLAAILTRIKQNR-HLRTFRI
TVGVDGTVYKIH-PQYAKRLHKVVRRLHPECQVRFVLSDSGSSKGAALVTAVAQRLASRRQQVDETLSPF

RLSRETLQLVKTRMRAGLEAGLKSKGSS---AIKMLPSFVYHTPDGT-----------------------
---EHGKYLALDLGGSNFRALLVNFKKGLQQNSKLYHKIYTIPLEIMQGSGEE-------LFDHIAQCVG
DFLDYMGMKNA-RLPAGFTFSFPCEQTSIDTGTLLSWTKGFKATNCEGHDVVGMLRDAIKRRN----EFD
LDVMA-----VVNDTVGTMMSCAYEDPQCEIGLIAGTGTNVCYMEEL---RNIEKTE-----QKTGNVER
EEGT-----PEGAQVKQDDGTERERKKGEAEIPRMCINTEWGGLGDDGS-LDDIITPYDVEVDQHSLNPG
KQR-FEKLTSGMYLGEVVRQVLLDLTRGGLLFRGRVTEALKTPGIFQTKYLSQIES--------------
------------DRLALLQVRSILQQLGL-DSTCDDSIIVKEVCGAVSRRAAQLCGAGMAAVVDKIRENR
GLDHLDITVGVDGTLYKLH-----------PHFSPVLQETARLLAPQCNATF---------LPSEEG---
----------SGKGAALITAVSRQKME-------M-----------------
>XP_043839845|Dromiciops gliroides|HK1
LYSMRLSDEMLNEVTNRFKKAMQNGL-SRNYHVTAS--VKMLPTFVRSIPDGSEKGDFIALDLG--GSSF
RILRVQVSHE---KN-QT--VQMESEVYETPEGIMNGSGSM---------------LFDHVAECLGDFME
KK-----NIK---DKNLPVGFTFSFPCRQTKID------------------------E------------
-----------AVLITWTKRFKASGVEGSDVVQLLKKAIKKR----------G-----------------
--------------DYDANIVAVVNDTVGTMMTCGFDDQECEVGLII-----------------------
-----G-----TGTNACYMEEMRHI-DMVEGDE--GRMCINTEWGAFGDDGCIQEIRTEFDKEIDRGSLN
PGKQLFEKMVSGMYMGELVRLIIVKMAKGGLLFEGRITPELLTKGKFETKNVSAIEK-------------
--TK--EGLQRAQEILTN-LGLEPSPEDCISVQHVCAIVSHRSANLVAAMLAGILHRLKENK-GVTRLRT
TVGVDGTLYKFH-PQYSRRFHKTLRRLVPDCDVRFILSESGSGKGAAMVTAVAYRLAEQHKQIEEMLAPF
KLSTEQLTQVKNRMRVEMDLGLQKKTNESA-KVKMLPSFVCGTPDGT-----------------------
---ENGDFLALDLGGTNFRVLLVKIRSGKKRSVEMHNKIYAIPEEVMQGTGEE-------LFDHIVSCIS
DFLDYMGIKGA-RLPLGFTFSFPCRQTSLDGGILLTWTKGFNASNCEGQDVPTLLREAIKRRE----EFD
LDVVA-----LVNDTVGTMMTCAYEEPTCEIGLIVGTGSNACYMEEM---KNIEMVE-------------
--GD------EG---------------------RMCINMEWGAFGDNGC-LEDIRTEYDRIVDDYSLNAN
KQR-FEKMISGMYLGEIVRNLLIDFTRKGFLFRGQISETLKTRGIFKTKFLSQIES--------------
------------DRLALLQVRTILQELGL-NSTCDDSILVKMVCGIVSKRAAQLCGAGMAAVVEKIKENR
GLKTLNVTVGVDGTLYKLH-----------PHFSRIMHQTVSELAPNCNVKF---------LLSEDG---
----------SGKGAALISAVGVKLRD--AKA---KAS--------------
>XP_007505933|Monodelphis domestica|HK1
LYAMRLSDDVLHEVTNRFKKAMSNGL-SRNYNVTAS--VKMLPTFVRSIPDGSEKGDFIALDLG--GSSF
RILRVQVSHD---KN-QH--VEMDSEVYETPDRIMRGSGPE---------------LFDHVAECLGNFME
KK-----KIK---DKNLPIGFTFSFPCRQTKIN------------------------E------------
-----------AVLITWTKRFKVSGVEGSDVVQLLNKAIKKR----------G-----------------
--------------EYEANIMAVVNDTVGTMMTCAFDDPECEVGLIV-----------------------
-----G-----TGTNACYMEELRHI-DMVEGDE--GRMCINTEWGSFGDDGCINEIRTEFDKEIDRGSLN
PGKQLFEKMVSGMYMGELVRLIIVKMAKGGLLFDGRLTPELLTKGKFETKNVSAIEK-------------
--TK--EGLDRAKDILTN-LGLEPSPEDCVSVQHVCAIVSHRSANLVAAMLAGILFRLKENK-NLPRLRT
TVGVDGTLYKFH-PQYSRRLQKTLRRLFPDSDVRFILSESGSGKGAAMVTAVAYRLAEQQKHMAEMLAPL
KLTSQQLMQVKNRLRVEMDLGLKKSSHDSA-KVKMWPSYVTGTLDGS-----------------------
---ESGDFLALDLGGTNFRVLLVKIRSGKKKSVEMHNKIYAIPESVMQGTGEE-------LFDHIVSCIS
DFLDYMGMKTA-RLPLGFTFSFPCHQTSLDGGTLVTWTKGFNASNCVGQDVPTLLRDAIKRRG----EFD
LDVVA-----LVNDTVGTMMTCAYEERTCEVGLIVGTGSNACYMEEM---KNIELVE-------------
--GN------EG---------------------RMCINMEWGAFGDNGC-LEDLRTEYDRIVDDFSLNAG
KQR-YEKMISGMYLGEIVRNILIDFTKNGFLFRGHISEALKTRGIFKTKFLSQIES--------------
------------DRLALLQVRVILQELGL-NSTCDDSILVKMVCGVVSKRAAQLCGAGVAAVVDKIRENR
NLETLKVTVGVDGTLYKLH-----------PHFSRIMNQTVQDLAPKCNVKF---------LLSEDG---
----------SGKGAALISAVGVRLRE--GKG---KAS--------------
>XP_055503280|Leucoraja erinacea|HK1
LYAARLSDDTLRDVMTRFRTEMEKGL-ARDTNPTAT--VKMLPTFVRSPPDGSEKGDFLALDLG--GSNF
RILRVKVSHE---RQ-QK--VVMESQIYSTPEDIIHGSGSR---------------LFEHVAECLGDFME
KQ-----EIK---DKKLPLGFTFSFPCRQSKVD------------------------E------------
-----------SILITWTKGFKASGVEGSDVVKLLRKAIDKK----------A-----------------

--------------EYDVDIMAVVNDTVGTMMTCGFDDQRCEVGLII-----------------------
-----G-----TGTNACYMEELRHI-DLVEGDE--GRMCINTEWGAFGDDGSLEDIRTEFDRELDRGSLN
PGKQLFEKLISGMYMGELVRLILVKMAKEGLLFEGNITPELLTKGKLETKHVSAIEK-------------
--SK--GGLSKADEILTR-LGLEPSEEDCIAVQHVCAIVSFRSANLSAAALGAILTQLRHNK-GVPRLRT
TVAMDGSLYKMH-PQFARRFQKTVRRLVPDCEVRFLLSESGSGKGAAMVTAVAERLVSQRKEMDKTLATF
TLTREQLFEVKGRLRKEMEMGLQKKTHEAA-TVKMLPTFVRSTPDGT-----------------------
---ENGDFVALDLGGTNFRVLLVKIRSGKRRSVEMHNKIYAIPLEVMQGTGEE-------LFDHIVHCIS
DFLDYMGMKGA-RLPLGFTFSFPCEQKSLDEGILLKWTKGFKATDCEGEDVVTMLRDAIKRRD----EFE
LDIVS-----IVNDTVGTMMTCAYEDPLCEVGMIVGTGSNTCYMEEA---RNTETVP-------------
--GD------EG---------------------LMCVNMEWGAFGDNGC-LDDIRTEYDNAVDDLSLNHG
KAKVYEKMISGMYLGEIVRNVVIDFTRRGFLFRGQISERLKTRSIFETKFLSQIES--------------
------------DRLALLQVRAILQNLGL-DSTCDDSIIVKEVCGAVSRRAGQLCGAGMAAVVDKIRENR
GLDRLAITVGVDGTLYKLH-----------PHFSKIMHETVRELAPNCDVTF---------LLSEDG---
----------SGKGAALITAVACRLRE--SAA--------------------
>KAF6303823|Myotis myotis|HK1
LYAMRLSDETLIDIMTRFKKEMRHGL-SRDFNPTAT--VKMLPTFVRSTPDGSEKGDFIALDLG--GSYF
RILRVQVSLE---KN-QN--VHMESEVYDTPENIMHGSGSQ---------------LFDHVAECLGDFME
KK-----KIK---DKKLPVGLTFSFPCRQSRID------------------------E------------
-----------GILITWTKRFKASGVEGADVVKLLNKAIKKR----------G-----------------
--------------DYDANIVAVVNDTVGTMMTCGYDDQHCEVGLII-----------------------
-----G-----TGTNACYMEELRHI-DLVEGDE--GRMCINTEWGAFGDDGALEDIRTEFDREIDRGSLN
PGKQLFEKMISGMYLGELVRLILVKMAKEGLLFEGRITPELLTRGKFTTSDVSAIER-------------
--NK--EGLSNAKDILTR-LGVEPSDVDCVAVQHVCTIVSFRSANLVAAALGAILNRLRDNK-GIPRLRT
TVGVDGSLYKMH-PQYSRRFHKTLRRLVPDCDVRFLLSESGSGKGAAMVTAVAYRLAEQHRQIEETLAHF
RLTREMLLEVKKRMRTEMEMGLKKKTNDKA-VVKMLPSFVRSTPDGT-----------------------
---ENGDFLALDLGGTNFRVLLVKIRSGKKRTVEMHNKIYAIPIEIMQGTGEE-------LFDHIVSCIS
DFLDYMGIKGP-RMPLGFTFSFPCKQMSLDAGILITWTKGFKATDCVGHDVATLLREAVKRRE----EFD
LDVVA-----VVNDTVGTMMTCAYEEPTCEIGLIVGTGSNACYMEEM---KNVEMLE-------------
--GN------VG---------------------RMCINMEWGAFGDNGC-LDDIRTEYDVLVDENSLNAG
KQR-YEKMISGMYLGEIVRNILLDFTKKGFLFRGKISEPLKTQGIFQTKYLSQIES--------------
------------DRLALLQVRAILQQLGL-NSTCDDSILVKTVCGVVSRRAAQLCGAGMAAVVDKIRENR
GLDHLDVTVGVDGTLYKLH-----------PHFSRIMHQTVKELSPKCNVSF---------LLSEDG---
----------SGKGAALITAVGVRLRN-ETGS--------------------
>CAH6789752|Phodopus roborovskii|HK1
LYAMRLSDEILIDILTRFKKEMKNGL-SRDYNPTAS--VKMLPTFVRSIPDGSEKGDFIALDLG--GSSF
RILRVQVNHE---KN-QN--VHMESEAYDTPENIVHGSGSQ---------------LFDHVAECLGDFME
KR-----KIK---DKKLPVGFTFSFPCRQSKID------------------------E------------
-----------AILITWTKRFKASGVEGADVVTLLNKAIKKR----------G-----------------
--------------DYDANIVAVVNDTVGTMMTCGYDDQQCEVGLII-----------------------
-----G-----TGTNACYMEELRHI-DLVEGDE--GRMCVNTEWGAFGDDGALEDIRTEFDRELDRGSLN
PGKQLFEKMVSGMYMGELVRLILVKMAKEGLLFEGRITPELLTRGKFNTSDVSAIENTVYYRWFSCVVSK
TVNK--EGIQNAKEILTR-LGVEPSDDDCVSVQHVCTIVSFRSANLVAATLGAILNRLRDNK-GTARLRT
TVGVDGSLYKTH-PQYSRRFHKTLRRLVPDSDVRFLLSESGSGKGAAMVTAVAYRLAEQHRQIEETLAHF
RLSKQTLMEVKKRMRTEMETGLKKETHSRA-TVKMLPSFVRSIPDGT-----------------------
---ENGDFLALDLGGTNFRVLLVKIRSGKKRTVEMHNKIYSIPVEIMQGTGEE-------LFDHIVSCIS
DFLDYMGIKGP-RMPLGFTFSFPCNQTSLDCGILITWTKGFKATDCVGHDVVTLLKDAVKRRE----EFD
LDVVA-----VVNDTVGTMMTCAYEEPTCEIGLIVGTGSNACYMEEM---RNVEMVE-------------
--GN------QG---------------------QMCINMEWGAFGDNGC-LDDIRTDFDRVVDEYSLNAG
KQR-FEKMISGMYLGEIVRNILIDFTKKGFLFRGQISEPLKTRGIFETKFLSQIES--------------
------------DRLALLQVRAILQQLGL-NSTCDDSILVKTVCGVVSKRAAQLCGAGMAAVVEKIRENR
GLDHLNVTVGVDGTLYKLH-----------PHFSRIMHQTVKELSPKCTVSF---------LLSEDG---
----------SGKGAALITAVGVRLRG--DVM--TA----------------

> XP_007938858|Orycteropus afer afer|HK1
LYTMRLSDETLLDIMARFRKEMKNGL-SRDFNPTAT--IKMLPTFVRSIPDGSEKGDFIALDLG--GSSF
RILRVQVNHD---QH-QN--VHMESEVYDTPENIIHGSGSQ---------------LFDHVAECLGDFME
KK-----SIK---DKKLPVGFTFSFPCRQSKID------------------------E------------
-----------AVLITWTKRFKASGVEGADVVKLLDKAIKKR----------G-----------------
--------------DYEANIVAVVNDTVGTMMTCGYDDQHCEVGLII-----------------------
-----G-----TGTNACYMEELRHI-DLVEGDE--GRMCINTEWGAFGDNGSLEDIRTEFDREIDRGSLN
PGKQLFEKMVSGMYMGELVRLILVKMAKEGLLFEGRITPELLTKGKFNTSDVSAIEK------------
--NK--EGLHNAKEILTR-LGVEPSHDDCISVQHVCTIVSFRSANLVAATLGAILNRLRDNK-GTPRLRT
TVGVDGSLYKTH-PQYSRRFHKTLRRLVPDSDVRFLLSESGSGKGAAMVTAVAYRLAEQHRQIEETLAYF
RLTKEMLMEVKKRMRNEMEFGLKKATHDAA-TVKMLPSFVRSTPDGT-----------------------
---ENGDFLALDLGGTNFRVLLVKIRSGKRRTVEMHNKIYAIPIEIMQGTGEE-------LFDHIVSCIS
DFLDYMGIKGP-RMPLGFTFSFPCKQTSLDAGVLITWTKGFKATDCVGHDVATLLRDAIKRRE----EFD
LDVVA-----VVNDTVGTMMTCAYEEPTCEVGLIVGTGSNACYMEEM---RNVETLE-------------
--GT------NG---------------------EMCINMEWGAFGDNGC-LDDIRTNYDKLVDEYSLNAG
KQR-FEKMISGMYLGEIVRNILIDFTKKGFLFRGQISETLKTRGIFETKYLSQIES--------------
------------DRLALLQVRAILQQLGL-NSTCDDSILVKTVCGVVSKRAAQLCGAGMAAVVDKIRENR
GLDHLNVTVGVDGTLFKLH-----------PHFSRIMHQTVKELAPQCNVSF---------LLSEDG---
----------SGKGAALITAVGVRLRE--MSP--SS----------------
> XP_059981502|Lagenorhynchus albirostris|HK1
LYAMRLSDETLLDIMARFKKEMKNGL-SRDFNPTAT--VKMLPTFVRSIPDGSEKGDFIALDLG--GSSF
RILLVQVNHE---QK-QT--VHMESEVYDTPESIMHGSGSQ---------------LFGHVAECLGDFME
KK-----KIK---DKKLPVGFTFSFPCRQSKID------------------------E------------
-----------AILITWTKRFKASGVEGTDVVKLLEKAIKKH----------G-----------------
--------------DYDANVVAVVNDTVGTMMTCGYDDQQCEVGLII-----------------------
-----G-----TGTNACYMEELRHI-DLVEGDE--GRMCINTEWGAFGDDGALEDIRTEFDREIDRGSLN
PGKQLFEKMVSGMYMGELVRLILVKMAKEGLLFEGRITPELLTRGKFNTSDVSAIEK------------
--NK--EGLHNAREILTR-LGVEPSDDDCISVQHVCTIVSFRSANLVAATLGAILNRLRDNK-STPRLRT
TVGVDGSLYKTH-PQYSRRFHKTLRRLVPDCDVRFLLSESGSGKGAAMVTAVAYRLAEQHRQIEETLAHF
SLTKDTLLEVKRRMRAEMELGLGKQTHDKA-VVKMLPSFVRSTPDGT-----------------------
---EHGDFLALDLGGTNFRVLLVKIRSGKKRTVEMHNKIYAIPTEIMQGTGEE-------LFDHIVSCIS
DFLDYMGIKGP-RMPLGFTFSFPCKQTSLDAGILITWTKGFKATDCVGQDVATLLRDAVKRRE----EFD
LDVVA-----VVNDTVGTMMTCAYEEPTCEVGLIVGTGSNACYMEEM---KNVETLA-------------
--GN------EG---------------------QMCINMEWGAFGDNGC-LDDIRTIYDKLVDEYSLNAG
KQR-YEKMISGMYLGEIVRNILIDFTKRGFLFRGQISEPLKTRGIFQTKYLSQIES--------------
------------DRLALLQVRAILQQLGL-NSTCDDSILVKTVCGVVSKRAAQLCGAGMAAVVDKIRENR
GLDHLNVTVGVDGTLYKLH-----------PHFSRIMHQTVKELSPKCNVSF---------LLSEDG---
----------SGKGAALITAVGVRLRQ--ETS--S-----------------
> P19367|Homo sapiens|HK1
LYAMRLSDETLIDIMTRFRKEMKNGL-SRDFNPTAT--VKMLPTFVRSIPDGSEKGDFIALDLG--GSSF
RILRVQVNHE---KN-QN--VHMESEVYDTPENIVHGSGSQ---------------LFDHVAECLGDFME
KR-----KIK---DKKLPVGFTFSFPCQQSKID------------------------E------------
-----------AILITWTKRFKASGVEGADVVKLLNKAIKKR----------G-----------------
--------------DYDANIVAVVNDTVGTMMTCGYDDQHCEVGLII-----------------------
-----G-----TGTNACYMEELRHI-DLVEGDE--GRMCINTEWGAFGDDGSLEDIRTEFDREIDRGSLN
PGKQLFEKMVSGMYLGELVRLILVKMAKEGLLFEGRITPELLTRGKFNTSDVSAIEK------------
--NK--EGLHNAKEILTR-LGVEPSDDDCVSVQHVCTIVSFRSANLVAATLGAILNRLRDNK-GTPRLRT
TVGVDGSLYKTH-PQYSRRFHKTLRRLVPDSDVRFLLSESGSGKGAAMVTAVAYRLAEQHRQIEETLAHF
HLTKDMLLEVKKRMRAEMELGLRKQTHNNA-VVKMLPSFVRRTPDGT-----------------------
---ENGDFLALDLGGTNFRVLLVKIRSGKKRTVEMHNKIYAIPIEIMQGTGEE-------LFDHIVSCIS
DFLDYMGIKGP-RMPLGFTFSFPCQQTSLDAGILITWTKGFKATDCVGHDVVTLLRDAIKRRE----EFD
LDVVA-----VVNDTVGTMMTCAYEEPTCEVGLIVGTGSNACYMEEM---KNVEMVE------------

--GD------QG---------------------QMCINMEWGAFGDNGC-LDDIRTHYDRLVDEYSLNAG
KQR-YEKMISGMYLGEIVRNILIDFTKKGFLFRGQISETLKTRGIFETKFLSQIES--------------
------------DRLALLQVRAILQQLGL-NSTCDDSILVKTVCGVVSRRAAQLCGAGMAAVVDKIRENR
GLDRLNVTVGVDGTLYKLH-----------PHFSRIMHQTVKELSPKCNVSF---------LLSEDG---
----------SGKGAALITAVGVRLRT--EAS--S-----------------
>XP_043912214|Protopterus annectens|HK1
LYAMRLSDETLLDITGRFRREMENGL-SRDFNPTCS--VKMLPTFVRSIPDGTERGDFIALDLG--GSFF
RILRVKVSHD---KK-QM--VQMESQVYDTPEDIIHGTGIR---------------LFDHVAECLGDFMA
KQ-----EIK---DKKLPVGFTFSFPCRQTKLD------------------------E------------
-----------AVLITWTKKFKASGVEGADVVKLLNKAIKKR----------G-----------------
--------------DYDADIIAVVNDTVGTMMTCGFDDQRCEVGIII-----------------------
-----G-----TGTNACYMEELRHI-DLVEGDE--GRMCINTEWGAFGDDGELEDIRTEFDKEIDRGSLN
PGKQLFEKLTSGMYMGELVRLILVKMAKEGLLFEGRITPELLTKGKLTTKDVSAMEK-------------
--SK--DGLSKAKDILTR-LGVEPSQEDCVAVQHVCTIVSFRSANLVAACLGAILTRLRDNK-GVPRLRT
TVGVDGSLYKMH-PQYCRRLQKTVRRLNPDCDVRFLLSESGSGKGAAMVTAVAHRLAEQRRQIDATLAQF
ELSNEQLMEVKKRMRIEIDAGLRKKTHENA-TVKMLPTFVRSTPDGT-----------------------
---ENGDFLALDLGGTNFRVLLVAIRSGKRRSVEMHNKIYAIPVEVMQGTGEE-------LFDHIVHCIS
DFLDYMGMKGA-RLPLGFTFSFPCKQTSLDAGILLKWTKGFKATDCEGEDVVGLLREAIKRRE----EFE
LDIVA-----VVNDTVGTMMTCAYEDSNCEIGLIVGTGSNACYMEEM---RNIEMVD-------------
--GD------EG---------------------RMCVNMEWGAFGDNGC-LDDIRTTSDKAVDELSLNSG
KQR-YEKMISGMYLGEIVRNILIDFTKRGFLFRGQISEALKTRSIFETKFLSQIES--------------
------------DRLALLQVRAILQQLGL-NSTCDDSIIVKEVCGAVSRRAAKLCGAGMAAVVDKIRENR
GMEHLDVTVGVDGTLYKLH-----------PHFSKIMKHTVKELAPKCDVTF---------LLSEDG---
----------SGKGAALITAVGCRLRQ--MEQ--K-----------------
>XP_059903079|Gadus macrocephalus|HK1
LYSMRFSDETVRDIMTRFRREMQNGL-GRDTNATATVKVKMLPTFVRSIPDGSEKGDFLALDLG--GSNF
RILRVKVTRD---KK-QP--VQMESQVYETPDDIIHGSSTR---------------LFDHVADCLGDFME
KQ-----NIK---DKNLPVGFTFSFPCAQTKLN------------------------E------------
-----------AVLVTWTKKFKVSGVEGMDVVKLLNKAIKKR----------G-----------------
--------------DYEADIMAVVNDTTGTMMTCGFDDQRCEVGIII-----------------------
-----G-----TGTNACYMEELRHM-DLVEGDE--GRMCINTEWGAFGDDGSLEDIRTEFDREIDRGSLN
PGKQLFEKMASGMYMGELVRLILVKMAKESLLFEGRITPELLTKGTIETKHVSAIEK-------------
--SK--EGLKKCMEILTR-LGVEPSDEDCLAVQHVCTIVSFRSANLVAATLGGILLRLKENR-GGMRLRT
TVGIDGSLYKMH-PQYARRLHKTVRRLVPETDVRFLLSESGSAKGAAMVTAVAYRLSDQARQIQQTLAEF
RLTKAQLLEVKKRMRTEIERGLKKESHEEA-TVKMLPTFVRSTPDGT-----------------------
---ENGDFLALDLGGTNFRVLLVKIRSGKRRTVEMHNKIYAIPIEVMQGTGEE-------LFDHIVFCIS
DFLDYMGMKSA-RLPLGFTFSFPCHQTSLDAGRLVNWTKGFKATDCVGEDVVELLREGIKRKEMEDPEFE
LDVVA-----IVNDTVGTMMTCAYEEPTCEVGLIAGTGSNACYMEEM---RNIESVE-------------
--GN------EG---------------------RMCVNMEWGAFGDNGC-LDDIRTQYDRAVDENSLNEG
KQR-YEKMCSGMYLGEIVRNILIDLTRRGFLFRGQISETLKTRGIFETKFLSQIES--------------
------------DRLALLQVRAILQGLGL-DSTCDDSIIVKEVCGSVSRRAAQICGAGMAAVVDKIRENR
GLDHLDITVGVDGTLYKLH-----------PHFSRIFHQTVKDLAPKCDVNF---------LLSEDG---
----------SGKGAALITAVGCRQREQEAQQ--H-----------------
>XP_056266310|Pseudoliparis swirei|HK1
LYSMRFSDDTVKDIMNRFRREMENGL-GRDTGPTAT--VKMLPTFVRAIPDGSEKGDFIALDLG--GSNF
RILRVKVTED---KK-QP--VQMESQVYETPDDIIHGSGTR---------------LFSHVADCLGDFME
KQ-----KIK---DKKLPVGFTFSFPCAQTKLD------------------------E------------
-----------AVLLTWTKKFRASGVEGTDVVSLLNRAIKKR----------G-----------------
--------------DYEADIMAVVNDTVGTMMTCGFDDQRCEVGVII-----------------------
-----G-----TGTNACYMEELRHI-DQVEGDE--GRMCINTEWGAFGDDGSLEDVRTEFDREIDRGSIN
PGRQLFEKMASGMYMGELVRLILVKMAKEGLLFEGRITPELLTKGKVETKHVSAIEK-------------
--SK--EGLKKCMETLTR-LGVEPSAEDCLAVQHVCTIVSFRSANLVAATLGAILARLKDNR-GAARLRT

TVGIDGSLYKMH-PQYARRLHKTVRRLVPDSDVRFLLSESGSAKGAAMVTAVAYRLTEQARQIQQTMAEF
RLSKAQLLEVKKRMRQEMERGLKKTSHHEA-TVKMLPTFVRRTPDGT-----------------------
---ENGDFLALDLGGTNFRVLLVKIRSGKKRSVEMHNKIYAIPVEVMQGTGEE-------LFDHIVHCIS
DFLDYMGMKSA-RLPLGFTFSFPCHQTSLDAGILVNWTKGFKATDCEGENVVELLREAIKRKE----EFE
LDVVA-----IVNDTVGTMMTCAYEEPSCEVGLIAGTGSNCCYMEEA---GNVELVE------------
--GA------AG------------------AGGGMCVNMEWGAFGDNGC-LDDIRTQYDQAVDENSLNDG
KQR-YEKMCSGMYLGEIVRQVLIDLTRRGFLFRGQISETLKTRGIFETKFLSQIES-------------
------------DRLALLQVRAILQQLGL-DSTCDDSIIVKEVCGTVSRRAAQMCGAGMAAVVDKIRENR
GLDHLDTTVGVDGTLYKLH-----------PHFSRIFQQTVKELAPKCDVQF---------LLSEDG---
----------SGKGAALITAVGCRQRE--LDA----QH--------------
>XP_034043790|Thalassophryne amazonica|HK1
LYSMRFSDETLKDIKNRFRRDMENGL-GRDTNPTAT--VKMLPTFVRSIPDGSEKGDFIALDLG--GSNF
RILRVKVTED---KK-QP--VQMESQVYETPDDIIHGSGTQ---------------LFDHVAECLGDFME
KQ-----KVK---DKNLPVGFTFSFPCAQTKLN------------------------E------------
-----------AVLLTWTKKFKASGVEGADVVKLLSKAIMRR----------G-----------------
--------------DYEADIMAVVNDTVGTMMTCGFDDQRCEVGIII-----------------------
-----G-----TGTNACYMEELRHI-DLVEGDE--GRMCINTEWGAFGDNGSLEDIRTEFDREIDRGSIN
PGKQLFEKMASGMYMGELVRLILVKMAKEGLLFEGRITPELLTKGKIKTKHVSAIEK------------
--TK--EGLKKCMEILTR-LGVEPSDEDCLAVQHVCTIVSFRSANLISASLGAILSRLKDNK-GVARLRT
TVGIDGSLYKMH-PQYARRLHKTVRRLVPDSDVRFLLSESGSAKGAAMVTAVAYRLTEQARQIQQTLAEF
RLSKAQLLEVKKRMRVEMERGLKKDTHKDA-SVKMLPSFVRSTPDGT-----------------------
---ENGDFLALDLGGTNFRVLLVKIRSGKKRRTVEMHNKIYAIPLEVMQGTGEE-------LFDHIVHCIS
DFLDYMGMKSA-RLPLGFTFSFPCHQTSLDAGILLTWTKGFKATDCEGENVVELLREAIKRKE----EFD
LDVVA-----IVNDTVGTMMTCAYEEPTCEIGLIAGTGSNACYMEET---RNIEMVD------------
--GN------EG---------------------RMCVNMEWGAFGDNGC-LDDIRTIYDKAVDENSLNEG
KQR-YEKMCSGMYLGEIVRNILIDLTKRGFLFRGHISETLKTTGIFETKFLSQIES-------------
------------DRLALLQVRAILQQLGL-NSTCDDSIIVKEVCGTVSRRAAQICGAGMAAVVDKIRENR
GLDHLDVTVGVDGTLYKLH-----------PHFSRIFHQTVKELAPKCDVNF---------LLSEDG---
----------SGKGAALITAVGCRQREQEAQH--H-----------------
>XP_056135455|Lampris incognitus|HK1
LYSMRFSDETLKDIMSRYRREMENGL-GRDTSPTAT--VKMLPTFVRSIPDGSEKGDFIALDLG--GSNF
RILRVKVTQD---KK-QP--VQMESQVYETPDDITHGSGTQ---------------LFDHVADCLGDFME
KQ-----KIK---NKKLPVGFTFSFPCAQSKLN------------------------E------------
-----------AVLVTWTKKFKASGVEGMDVVKLLNKAIKKR----------G-----------------
--------------DYDADIMAVVNDTVGTMMTCGFDDQRCEVGIIV-----------------------
-----G-----TGTNACYMEELRHI-DLVEGDE--GRMCINTEWGAFGDDGLLEDIRTEFDREIDRGSLN
PGKQLFEKMASGMYMGELVRLILVKMAKEGLLFEGRITPELLTKGKIETKHVSAIEK------------
--SK--EGLKKCMEILTR-LGVEPSDEDCLAVQHVCTIVSFRSANLIAATLGGILARLKENK-GVARLRT
TVGIDGSLYKMH-PQYARRLHKTVRRLVPDSDVRFLLSESGSAKGAAMVTAVAYRLADQARQIQQTLAEF
RLSKAQLLEVKKKMRTEIERGLQKNTHNDA-TVKMLPTFVRRTPDGT-----------------------
---ENGDFLALDLGGTNFRVLLVKIRSGKKRRTVEMHNKIYAIPIEVMQGTGEE-------LFDHIVHCIS
DFLDYMGMKSA-RLPLGFTFSFPCHQTSLDAGILVNWTKGFKATDCEGEDVVELLREGIKRKE----EFE
LDVVA-----IVNDTVGTMMTCAYEEPTCEVGLIAGTGSNACYMEEM---KNIEMVD------------
--GN------EG---------------------RMCVNMEWGAFGDNGC-LDDIRTQYDCAVDENSLNEG
KQR-YEKMCSGMYLGEIVRNILIDLTKRGFLFRGQISETLKTRGIFETKFLSQIES-------------
------------DRLALLQVRAILQQLGL-DSTCDDSIIVKEVCGTVSRRAAQICGAGMAAVVDKIRENR
GLDHLDITVGVDGTLYKLH-----------PHFSRIFHQTVKELAPKCDVNF---------LLSEDG---
----------SGKGAALITAVGCRQRE--QEA----QH--------------
>XP_059188067|Centropristis striata|HK1
LYSMRFSDETLRDIMNRFRREMENGL-GRDTSPTAT--VKMLPTFVRSIPDGSEKGDFIALDLG--GSNF
RILRVKVTQD---KK-QP--VQMESQVYETPDDIIHGNGTR---------------LFGHVADCLGDFME
KQ-----KIK---DKKLPVGFTFSFPCAQTKLN------------------------E------------

```
-----------AVLVTWTKKFKVSGVEGMDVVSLLNRAIKKR----------G-----------------
--------------DYEADIMAVVNDTVGTMMTCGFDDQRCEVGIII-----------------------
-----G-----TGTNACYMEELRHI-DQVEGDE--GRMCINTEWGAFGDDGSLEDIRTEFDREIDRGSIN
PGKQLFEKMASGMYMGELVRLILVKMAKEGLLFEGRITPELLTKGKIETKHVSAIEK-------------
--SK--EGLKKCMEILTR-LGVEPSDEDCLAVQHVCTIVSFRSANLIASTLGGILSRLKENK-GVARLRT
TVGIDGSLYKMH-PQYARRLHKTVRRLVPDSDVRFLLSESGSAKGAAMVTAVAYRLTEQARQIQQTMAEF
RLSKAQLLEVKKRMRVEIERGLKKTSHEDA-TVKMLPTFVRTTPDGT-----------------------
---ENGDFLALDLGGTNFRVLLVKIRSGKKRSVEMHNKIYAIPIEVMQGTGEE-------LFDHIVHCIS
DFLDYMGMKSA-RLPLGFTFSFPCHQTSLDAGILVTWTKGFKATDCEGEDVVELLREAIKRKE----EFE
LDVVA-----IVNDTVGTMMTCAYEEPSCEVGLIAGTGSNACYMEEA---KNIEIVE-------------
--GS------EG---------------------RMCVNMEWGAFGDNGC-LDDIRTKYDQAVDENSLNEG
KQR-YEKMCSGMYLGEIVRQILIDLTRRGFLFRGQISETLKTRGIFETKFLSQIES--------------
------------DRLALLQVRAILQQLGL-DSTCDDSIIVKEVCGTVSRRAAQICGAGMAAVVDKIRENR
GLDHLDITVGVDGTLYKLH-----------PHFSRIFQQTVKELAPKCDVNF---------LLSEDG---
----------SGKGAALITAVGCRQRE--LEA--QQQH--------------
> XP_038567794|Micropterus salmoides|HK1
LYSMRFSDETLKDIMNRFRREMENGL-GRDTNPTAT--LKMLPTFVRSIPDGSEKGDFIALDLG--GSNF
RILRVKVTQD---KK-QP--VQMESQVYDTPDDIIHGSGTQ---------------LFDHVADCLGDFME
KQ-----KIK---DKKLPVGFTFSFPCTQTKLD------------------------E------------
-----------AVLVTWTKKFKASGVEGMDVVKLLNKAIKKR----------G-----------------
--------------DYEADIMAVVNDTVGTMMTCGFDDQRCQVGIII-----------------------
-----G-----TGTNACYMEEMRHI-DLVEGDE--GRMCINTEWGAFGDDGSLEDIRTEYDREIDRGSIN
PGKQLFEKMVSGMYLGELVRLILVKMAKEGLLFEGRITPELLTKGKIETKHVSAIEK-------------
--TK--EGLKKCMEILTR-LGVEPSDEDCLAVQHVCTIVSFRSANLIASTLGGILSRLKENK-GPVRLRT
TVGIDGSLYKMH-PQYARRLHKTVRRLVPDSDVRFLLSESGSAKGAAMVTAVAYRLTEQARQIQETMAEF
RLSKAQLLEVKKRMRMEIERGLKKDSHKEA-TVKMLPTFVRSTPDGT-----------------------
---ENGDFLALDLGGTNFRVLLVKIRSGKRRSVEMHNKIYAIPIEVMQGTGEE-------LFDHIVHCIS
DFLDYMGMKSA-RLPLGFTFSFPCHQTSLDAGILVTWTKGFKATDCEGEDVVELLREAIKRKEMENPEFE
LDVVA-----IVNDTVGTMMTCAYEEPTCEVGLIAGTGSNACYMEET---KNIEIVE-------------
--GN------EG---------------------RMCVNMEWGAFGDNGC-LDDIRTKYDQAVDDNSLNEG
KQR-YEKMCSGMYLGEIVRQILIDLTRRGFLFRGQISETLKTRGIFETKFLSQIES--------------
------------DRLALLQVRAILQQLGL-DSTCDDSIIVKEVCGTVSRRAAQICGAGMAAVVDKIRENR
GLDHLDITVGVDGTLYKLH-----------PHFSRIFQQTVKDLAPKCDVTF---------LLSEDG---
----------SGKGAALITAVGCRQRELEAQQ--H-----------------
> XP_063331828|Pelmatolapia mariae|HK1
LYSMRFSDETLKDIMNRFRREMENGL-ERDTNPTAT--VKMLPTFVRSIPDGSEKGDFIALDLG--GSNF
RILRVKVTQD---KK-QP--VQMESQIYETPDDIIHGSGAQ---------------LFDHVADCLGDFME
KQ-----KIK---DKKLPVGFTFSFPCAQTKLD------------------------E------------
-----------AVLLTWTKKFKASGVEGMDVVKLLNKAIKKR----------G-----------------
--------------DYQADIMAVVNDTTGTMMTCGFDDQRCEVGIII-----------------------
-----G-----TGTNACYMEELRHI-DLVEGDE--GRMCINTEWGAFGDNGSLEDIRTEFDREIDRGSIN
PGKQLFEKMASGMYMGELVRLILVKMAKEGLLFEGRITPELLTRGKIETKHVSAIEK-------------
--TK--EGLKKCMEILTR-LGVEPSEEDCLAVQHVCTIVSFRSANLISATLGAILCRLKENK-GVVRLRT
TVGIDGSLYKMH-PQYARRLHKTVRRLVPDSDVRFLLSESGSGKGAAMVTAVAYRLTEQARQIQQTLAEF
RLSKNQLLEVKKRMKVEIERGLKKDSHKDA-TVKMLPTFVRSTPDGT-----------------------
---ENGDFLALDLGGTNFRVLLVKIRSGKRRSVEMHNKIYAIPIEVMQGTGEE-------LFDHIVHCIS
DFLDYMGMKSA-RLPLGFTFSFPCHQTSLDAGILVTWTKGFKATDCEGEDVVELLREAIKRKE----EFE
LDVVA-----IVNDTVGTMMTCAYEEPTCEVGLIAGTGSNACYMEEM---KNIEIVP-------------
--GT------EG---------------------RMCVNMEWGAFGDNGC-LDDIRTTYDQAVDENSLNEG
KQR-YEKMCSGMYLGEIVRQILIDLTKRGFLFRGQISETLKTRGIFETKFLSQIES--------------
------------DRLALLQVRAILQQLGL-DSTCDDSIIVKEVCGTVSRRAAQLCGAGMAAVVDKIRENR
RLDHLDVTVGVDGTLYKLH-----------PHFSRIFQQTVKELAPKCDVNF---------LLSEDG---
```

----------SGKGAALITAVGCRQRELDAQQ--H-----------------
> XP_057902432|Doryrhamphus excisus|HK1
LYSMRFSDETLKDIMNRFRREMENGL-GHDTNPTAT--VKMLPTFVRSIPDGSEKGDFIALDLG--GSYF
RILRVKVTQD---KK-QP--VQMESQSYETPDNIIHGSGTQ---------------LFDHVAECLGDFME
KQ-----KIK---DKKLPVGFTFSFPCAQTKLD------------------------E------------
-----------AVLVTWTKKFKASGVEGMDVVKLLNKAIKKR----------G-----------------
--------------DYEADIMAVVNDTVGTMMTCGFDDQRCEVGIII-----------------------
-----G-----TGTNACYMEEMRHI-DQVEGDE--GRMCINTEWGAFGDDGSLEDIRTEFDREIDRGSIN
PGKQLFEKMASGMYMGELVRLILVKMAKEGLLFEGRITPELLTKGKIETKHVSAIEK-------------
--TK--EGLKKCMEILTR-LGVEPSDEDCLAVQHVCTIVSFRSANLISATLGGILSRLKENK-GVARLRT
TVGIDGSLYKMH-PQYARRLHKTVRRLVPDSDVRFLLSESGSAKGAAMVTAVAYRLTEQARQIQQTLAEF
RLSKAQLLEVKKRMRVEIERGLKKDTHEEA-TVKMLPTYVRSTPDGT-----------------------
---ENGDFLALDLGGTNFRVLLVKIRSGKRRSVEMHNKIYAIPIEVMQGTGEE-------LFDHIVHCIS
DFLDYMGMKSA-RLPLGFTFSFPCHQTSLDAGILVTWTKGFKATDCEGEDVVELLREAIKRKE----EFE
LDVVA-----IVNDTVGTMMTCAYEEPTCEVGLIAGTGSNACYMEEL---RNIEIVD-------------
--GN------EG---------------------RMCVNMEWGAFGDNGC-LDDIRTKYDQAVDENSLNEG
KQR-YEKMCSGMYLGEIVRQILIDLTKRGFLFRGQISETLKTRGIFETKFLSQIES--------------
------------DRLALLQVRAILQQLGL-DSTCDDSIIVKEVCGTVSRRAAQICGAGMAAVVDKIRENR
KLDHLDVTVGVDGTLYKLH-----------PHFSRIFHQTVKELAPKCDVTF---------LLSEDG---
----------SGKGAALITAVGCRQREMEAQQ--H-----------------
> XP_022070933|Acanthochromis polyacanthus|HK1
LYSMRFSDETLKDIMNRFRREMENGL-GNDTNPTAT--VKMLPTFVRSIPDGSEKGDFIALDLG--GSNF
RILRVKVTQD---KK-QP--VQMESQVYETPDDIIHGSGTR---------------LFDHVAECLGDFME
KQ-----KIK---DKKLPVGFTFSFPCAQTKLD------------------------E------------
-----------AVLVTWTKKFKASGVEGVDVVKLLNKAIKKR----------G-----------------
--------------DYEADIMAVVNDTVGTMMTCGFDDQRCEVGIII-----------------------
-----G-----TGTNACYMEELRHI-DLVEGDE--GRMCINTEWGAFGDDGSLEDIRTEFDREIDRGSIN
PGKQLFEKMASGMYMGELVRLILVKMAKEGLLFEGRITPELLTKGKIETKHVSAIEK-------------
--TK--EGLKKCMEILTR-LGVEPSDEDCLAVQHVCTIVSFRSANLISATLGAILARLKENK-GVARLRT
TVGIDGSLYKMH-PQYARRLHKTVRRLVPDSDVRFLLSESGSAKGAAMVTAVAYRLTEQARQIQQTLAEF
RLSKTQLLEVKKRMREEIERGLKKSSHQEA-TVKMLPTFVRSTPDGT-----------------------
---ENGDFLALDLGGTNFRVLLVKIRSGKRRTVEMHNKIYAIPIEVMQGTGEE-------LFDHIVHCIS
DFLDYMGMKSA-RLPLGFTFSFPCHQTSLDAGILVTWTKGFKATDCEGEDVVELLREAIKRKE----EFE
LDVVA-----IVNDTVGTMMTCAYEEPTCEVGLIAGTGSNACYMEEM---KNVEIVE-------------
--GN------EG---------------------RMCVNMEWGAFGDNGC-LDDIRTIYDQAVDENSLNEG
KQR-YEKMCSGMYLGEIVRQILIDLTKRGFLFRGQISETLKTRGIFETKFLSQIES--------------
------------DRLALLQVRAILQQLGL-DSTCDDSIIVKEVCGTVSRRAAQICGAGMAAVVDKIRENR
GLDHLDVTVGVDGTLYKLH-----------PHFSRIFQQTVKELAPKCDVTF---------LLSEDG---
----------SGKGAALITAVGCRQRE-LDAQ--QH----------------
> XP_005990009|Latimeria chalumnae|HK1
LFSMRLSDETLLDIMTRFRSEMKRGL-SSDFNPTAT--IKMLPTFVRSTPDGSEKGDFIALDLG--GSFF
RILRVKVSHE---KK-QT--VQMESKIYDTPEDIMSGSGTR---------------LFDHVAECLGDFME
KQ-----QIK---DKKLPVGFTFSFPCVQTKLN------------------------E------------
-----------GVLITWTKRFKASGVEGTDVVTLLNKAIKKR----------G-----------------
--------------DYDADIMAVVNDTVGTMMTCGFDDQRCEVGLII-----------------------
-----G-----TGTNACYMEELRHI-DLVEGDE--GRMCINTEWGAFGDDGSLEDIRTEFDREIDRGSLN
PGKQLFEKMASGLYLGELVRLILVKMAKEELLFEGRITPELLTKGKFETKHVSAIEK-------------
--NK--EGLTKAKEILTR-LGVEPSREDCVAVQHVCTIVSFRSANLIAAVLGGILTHMKENK-GVTRLRT
TVGVDGSLYKMH-PQYARRLHKTVRRLVPDCDVRFLLSESGSGKGAAMVTAVAYRLAEQHRQIDETLESF
NLTHKQLMEIKKRMRTEMEAGLRKKTHEEA-KVKMLPTFVRSTPDGT-----------------------
---EHGDFLALDLGGTNFRVLLVKIRSGKRRTVEMHNKIYAIPIEVMQGTGEE-------LFDHIVNCIS
DFLDYMGMKGA-CLPLSFTFSFPCQQTSLDAGILLTWTKGFKATDCEGEDMVNMLREAIKRRE----EFE

```
LDVVA-----IVNDTVGTMMTCAYEDQNCEIGLIVGTGSNACYMEEV---KNIEMVE-------------
--GE------QG---------------------RMCVNMEWGAFGDNGC-LDDIRTVYDEAVDDLSLNPG
KQR-YEKMISGMYLGEIVRNILIDLTKRGFLFRGQISETLKTRGIFETKFLSQIES--------------
------------DRLALLQVRAILQQLGL-DSTCEDSIIVKAVCGAISKRAAQVCGAGMAAVVDKIRENR
GLNHLEITVGVDGTLYKLH-----------PHFSKIMHQTVKELAPKCDVTF---------LLSEDG---
----------SGKGAALITAVACRLRE---LE--Q-KLN-------------
> XP_028651798|Erpetoichthys calabaricus|HK1
LYSMRFSDETLTDIMHRFRREMENGL-GRDTNPTAT--VKMLPTFVRSIPDGSEKGDFIALDLG--GSNF
RILRVKVSHE---KR-QT--VQMESQIYETPEDIIHGSGTQ---------------LFDHVAECLGDFME
KR-----KIK---DKKLPVGFTFSFPCRQSKLD------------------------E------------
-----------GILITWTKRFKASGVEGADVVKLLNKAIKKR----------G-----------------
--------------DYDADIMAVVNDTVGTMMTCGFDDQRCEVGIII-----------------------
-----G-----TGTNACYMEELRHI-DLVEGDE--GRMCINTEWGAFGDDGSLEDIRTEFDREIDRGSLN
PGKQLFEKMVSGMYMGELVRLILVKMAKEGLLFEGRITPELLTKGKFDTKHVSAIEK-------------
--SK--EGLNKAKEILTR-LGVEPSHEDCIAVQHVCTIVSFRSANLIAATLCGILSRLKDNK-GVPRLRT
TVGLDGSLYKMH-PQYARRLHKTVRRLVPDSDVRFLLSESGSGKGAAMVTAVAYRLLEQHRQIEETLDEF
KLSHVQLLEVKKIMRTEMENGLRKKTHESA-TVKMLPTFVRSTPDGT-----------------------
---ENGDFLALDLGGTNFRVLLVKIRSGKKRSVEMHNKIYAIPIEVMQGTGEE-------LFDHIVHCIS
DFLDYMGMKNA-RLPLGFTFSFPCKQTSLDAGILINWTKGFKATDCEGEDVVNLLREGIKRRE----EFD
LDVVA-----IVNDTVGTMMTCAYEEPTCEVGLIAGTGSNACYMEEM---RNIETVS-------------
--GD------EG---------------------RMCVNMEWGAFGDSGC-LDDFRTQYDRAVDDLSLNPG
KQR-YEKMCSGMYLGEIVRNILIDLTKRGFLFRGQISEALKTRGIFETKFLSQIES--------------
------------DRLALLQVRAILQGLGL-DSTCDDSIIVKEVCGVVSRRGAQLCGAGMAAVVDKIRENQ
GLDHLEITVGVDGTLYKLH-----------PHFSKIMHQTVKELAPKCDVTF---------LLSEDG---
----------SGKGAALITAVGCRLRE--QEQ--QN----------------
> XP_029465522|Rhinatrema bivittatum|HK1
LYSMRLSDETLLDIQARFRREMENGL-SRDFNPTAT--VKMLPTFVRSIPDGTEKGDFIALDLG--GSYF
RILRVKVSHE---KK-QT--VQMESEIYDTPEDIMHGSGAR---------------LFDHVAECLGDFME
KK-----QIK---DKKLPVGFTFSFPCRQSKID------------------------E------------
-----------GFLITWTKRYKASGVEGADVVKLLNKAIKKR----------G-----------------
--------------DYDADIMAVVNDTVGTMMTCGFDDQRCEVGIII-----------------------
-----G-----TGTNACYMEELRHI-DLVEGDE--GRMCINTEWGAFGDDGLLEDIRTEFDREIDRGSLN
PGKQLFEKMVSGMYLGELVRLILVKMAKEGLLFEGRITPELLTKGKFDTKHVSAIEK-------------
--SK--EGLTKAKEILTR-LGVEPSHEDCTAVQHVCTIVSFRSANLIAATLGAILTRLRDNK-GTPRLRT
TVAVDGSLYKMH-PQYARRLHKTVRRLVPDSDVRFLLSESGSGKGAAMVTAVAYRLAEQHRQIDETLEEF
QLSQEQLLEVKKRMRMEIEAGLRKKTHQSA-KVKMLPTFVLSTPDGS-----------------------
---ENGDFLALDLGGTNFRVLLVKIRSGKRRTVEMHNKIYAIPEDVMQGTGEE-------LFDHIVHCIS
DFLDYMGIKGA-RLPLGFTFSFPCSQTSLDAGILLTWTKGFKATDCEGEDVVQLLREGIKRRE----EFE
LDVVA-----VVNDTVGTMMTCAYEDPNCEIGLIVGTGSNACYMEEM---KNIEMVE-------------
--GD------KG---------------------RMCVNMEWGAFGDNGC-LDDIRTIYDKAVDELSLNSG
KQR-YEKMISGMYLGEIVRNILIDFTKRGFLFRGQISEALKTRGIFETKFLSQIES--------------
------------DRLALLQVRAILQQLGL-DGTCDDSIIVKEVCGAVSRRAAQVCGAGMAAVVDKIRENR
GLDHLDITVGVDGTLYKLH-----------PHFSRIMHQTVKTLAPKCDVTF---------LLSEDG---
----------SGKGAALITAVACRLRE--AEQ--S-----------------
> XP_038599558|Tachyglossus aculeatus|HK1
LYAMRLSDETLLDVMDRFSKEMKNGL-SRDFNPTAT--VKMLPTFVRSIPDGSEKGDFIALDIG--GSCF
RILRVKVSHE---KK-QT--VQMETEIYDTPEDIMHGSGSQ---------------LFDHVAECLGDFME
KK-----QIK---DKKLPVGFTFSFPCRQSRID------------------------E------------
-----------GVLITWTKRFKASGVEGADVVKLLNKAIKKR----------G-----------------
--------------DYDADIVAVVNDTVGTMMTCGFDDQRCEVGLII-----------------------
-----G-----TGTNACYMEELRHI-DSVEGDE--GRMCINTEWGAFGDDGSLEDIRTEFDREIDRGSLN
PGKQLFEKMVSGMYMGELVRLILVKMAKEGLLFEGRITPELLTKGKFETKHVSAIEK-------------
```

--SK--EGLNKAKEVLTR-LGVEPSHEDCIAVQHVCAIVSFRSANLVATTLGAILTRLRDNK-GVARLRT
TVGVDGSLYKMH-PQYSRRFHKTLRRLVPESDVRFILSESGSGKGAAMVTAVAYRLAEQHRLIEETLAEF
KLTHEQLLQVKRRMRVEIEAGLQKKSHESA-KVKMLPTFVRSTPDGT-----------------------
---ENGDFLALDLGGTNFRVLLVKIRSGKRRTVEMHNKIYAIPIEVMQGTGEE-------LFDHIVHCIS
DFLDYMGIKGA-RLPLGFTFSFPCKQTSLDAGILLNWTKGFKATDCEGQDVANLLREGIKRRE----EFD
LDVVA-----VVNDTVGTMMTCAYEEPTCEVGLIVGTGSNACYMEEM---KNIEMVE-------------
--GS------EG---------------------RMCVNMEWGAFGDNGC-LDDIRTIYDKAVDELSLNAG
KQR-YEKMISGMYLGEIVRNILIDFTKRGFLFRGQISETLKTRGIFETKFLSQIES--------------
------------DRLALLQVRAILQQLGL-NGTCDDSILVKTVCGTVSRRAAQMCGAGMAAVVDKIRENR
GLDHLEVTVGVDGTLYKLH-----------PHFSRIMHQTVKDLAPKCNVTF---------LLSEDG---
----------SGKGAALITAVGVRLRD--GGQ--S-----------------
>XP_020663615|Pogona vitticeps|HK1
LYAMRLSDETLLDIMGRFRSEMKNGL-SRDFNPTAS--VKMLPTFVRSIPDGSEKGDFIALDLG--GSFF
RILRVKVSHE---KK-QT--VQMETEIYETPEDIMHGSGTR---------------LFDHVAECLGDFME
KQ-----EIK---DKKLPVGFTFSFPCRQSKLD------------------------E------------
-----------GILITWTKRFKASGVEGADVVKLLNKAIKKR----------G-----------------
--------------DYDADIMAVVNDTVGTMMTCGFDDQRCEVGLII-----------------------
-----G-----TGTNACYMEEMRHI-DLVEGDE--GRMCINTEWGAFGDDGLLEDIRTEFDREIDRGSLN
PGKQLFEKMVSGMYMGELVRLILVKMAKEGLIFEGRITPELLTKGKLETKHVSAIEK-------------
--SK--EGLQKAKEILTR-LGVEPSHEDCIAVHHVCTIVSFRSANLVAATLGAILNQLRENK-GVGRLRT
TVGVDGSLYKMH-PQYSRRLHKTVRRLVPDSDVRFLLSESGSGKGAAMVTAVAYRLAEQHRLINETLAEF
KLTHEQLLQVKKRMRIEIEAGLRKKSHDHA-KVKMLPTFVRSTPDGT-----------------------
---ENGDFLALDLGGTNFRVLLVKIRSGKRRMVEMHNKIYAIPIEVMQGTGEE-------LFDHIVSCIS
DFLDYMGIKGA-RLPLGFTFSFPCKQTSLDAGILLNWTKGFKATDCEGEDVVNLLREGIKRRE----EFD
LDVVA-----VVNDTVGTMMTCAYEDPNCEIGLIVGTGSNACYMEEM---RNIEMVE-------------
--GE------QG---------------------RMCVNMEWGAFGDNGC-LDDIRTIYDKTVDEYSLNAG
KQR-YEKMISGMYLGEIVRNILIDFTKRGFLFRGQISETLKTRSIFETKFLSQIES--------------
------------DRLALLQVRAILQQLGL-NGTCDDSIIVKTVCGAVSRRAAQLCGAGMAAVVDKIRENR
GLDHLDITVGVDGTLYKLH-----------PHFSRIMHQTVKDLAPKCDVTF---------LLSEDG---
----------SGKGAALITAVGCRLRD--AET--HN----------------
>XP_006121588|Pelodiscus sinensis|HK1
LYSMRLSDETLLDIMARFKREMQNGL-SRDFNPTAT--VKMLPTFVRSIPDGSEKGDFIALDLG--GSYF
RILRVKVSHE---KK-QT--VQMETEIYDTPEDIMHGSGTR---------------LFDHVAECLGDFME
KQ-----QIK---DKKLPVGFTFSFPCRQSKLD------------------------E------------
-----------GILITWTKRFKASGVEGADVVKLLNKAIKKR----------G-----------------
--------------DYDADIMAVVNDTVGTMMTCGFDDQRCEVGLII-----------------------
-----G-----TGTNACYMEEMRHI-DLVEGDE--GRMCINTEWGAFGDDGSLEDIRTEFDREIDRGSLN
PGKQLFEKMVSGMYMGELVRLILVKMAKEGLLFEGRITPELLTKGKFETKHVSAIEK-------------
--SK--EGLNKAKEILTR-LGVEPSHEDCIAVQHVCTIVTFRSANLVASTLGAILNQLRDNK-GVTRLRT
TVGVDGSLYKMH-PQYSRRLHKTTRRLVPDSDVRFLLSESGSGKGAAMVTAVAYRLSEQHRLIDETLAEF
KLTHEQLLQVKKRMRTEIEAGLRKKTHETA-KVKMLPTFVRSTPDGT-----------------------
---ENGDFLALDLGGTNFRVLLVKIRSGKKRTVEMHNKIYAIPIEVMLGTGEE-------LFDHIVTCIS
DFLDYMGIKGA-RLPLGFTFSFPCKQTSLDAGILLNWTKGFKATECEGEDVVYLLREGIKRRE----EFD
LDVVA-----VVNDTVGTMMTCAYEDPNCEIGLIVGTGSNACYMEEM---RNIEMVE-------------
--GD------QG---------------------RMCVNMEWGAFGDNGC-LDDIRTTYDKAVDDYSLNAG
KQR-YEKMISGMYLGEIVRNILIDFTKRGFLFRGQISETLKTRGIFETKFLSQIES--------------
------------DRLALLQVRAILQQLGL-NGTCDDSIIVKMVCGAVSRRAAQVCGAGMAAVVDKIRENR
GVDHLEVTVGVDGTLYKLH-----------PHFSRIMHQTVKDLAPKCNVTF---------LLSEDG---
----------SGKGAALITAVGCRLRD--AEQ--N-----------------
>XP_006023198|Alligator sinensis|HK1
LYAMRLSDETLLDIMARFRREMKNGL-SRDFNPTAT--IKMLPTFVRSIPDGSEKGDFIALDLG--GSYF
RILRVKVSHE---KK-QT--VQMETEIYNTPEDIMHGTGTR---------------LFDHVAECLGDFME

KQ-----KIK---DKKLPVGFTFSFPCRQSKLD------------------------E------------
-----------GILVTWTKRYKASGVEGADVVKLLNKAIKKR----------G-----------------
--------------DYDADIMAVVNDTVGTMMTCGFDDQRCEVGLII-----------------------
-----G-----TGTNACYMEEMRHI-DLVEGDE--GRMCINTEWGAFGDDGSLEDIRTEFDREIDRGSLN
PGKQLFEKMVSGMYMGELVRLILVKMAKEGLLFEGRITPELLTKGKFETKHVSAIEK-------------
--SK--EGLNKAKEILTR-LGVEPSHEDCIAVHHVCTIVSFRSANLVACTLGAILNQLRDNK-GVPRLRT
TVGVDGSLYKMH-PQYARRLHKTTRRLVPDSDVRFLLSESGSGKGAAMVTAVAYRLAEQHRLIAETLAEF
KLTHEQLLQVKKRMRMEMEAGMKKKTHDSA-KVKMLPTFVRSTPDGT-----------------------
---ENGDFLALDLGGTNFRVLLVKIRSGKRRTVEMHNKIYAIPVEVMLGTGEE-------LFDHIVACIS
DFLDYMGIKGA-RLPLGFTFSFPCKQTSLDAGILLNWTKGFKATDCEGEDVVYLLREGIKRRE----EFD
LDVVA-----VVNDTVGTMMTCAYEDPNCEIGLIVGTGSNACYMEEM---RNIEMVE-------------
--GD------QG---------------------RMCVNMEWGAFGDNGC-LDDIRTIYDKAVDDYSLNAG
KQR-YEKMISGMYLGEIVRNILIDFTKRGFLFRGQISETLKTRSIFETKFLSQIES--------------
------------DRLALLQVRAILQQLGL-NSTCDDSIIVKTVCGAVSRRAAQLCGAGMAAVVDKIRENR
GLEHLEITVGVDGTLYKLH-----------PHFSKVMHQTVKDLAPKCDVTF---------LLSEDG---
----------SGKGAALITAVGCRLRD--AEQ--N-----------------
>XP_025962176|Dromaius novaehollandiae|HK1
LYAMRLSDETLIDIMARFRREMKNGL-SRDFNPTAT--VKMLPTFVRSIPDGSEKGDFIALDLG--GSYF
RILRVKVSHE---KK-QT--VQMETEIYNTPEDIMHGSGTR---------------LFDHVAECLGDFME
KQ-----QIK---DKKLPVGFTFSFPCRQSKLD------------------------E------------
-----------GILITWTKRFKASGVEGADVVKLLNRAIKKR----------G-----------------
--------------DYDADIMAVVNDTVGTMMTCGFDDQRCEVGLII-----------------------
-----G-----TGTNACYMEEMRHI-DLVEGDE--GRMCINTEWGAFGDDGSLEDIRTEFDREIDRGSLN
PGKQLFEKMVSGLYMGELVRLILVKMAKEGLLFEGRITPELLTKGKFETKHVSAIEK-------------
--SK--EGLSKAKDILTR-LGVEPSHEDCVAVQHVCTIVSFRSANLVAATLGAILCQLRDNK-GVGRLRT
TVGVDGSLYKMH-PQYARRLHKTTRRLVPDSDVRFLLSESGSGKGAAMVTAVAYRLSEQHRLIDETLAEF
KLTHEQLLQVKKRMRTEMEAGLRKKTHETA-KVKMLPTFVRSTPDGT-----------------------
---ENGDFLALDLGGTNFRVLLVKIRSGKRRTVEMHNKIYAIPIEVMQGTGEE-------LFDHIVTCIS
DFLDYMGIKGA-RLPLGFTFSFPCKQTSLDAGILLNWTKGFKATDCEGEDVVYLLREGIKRRE----EFD
LDVVA-----VVNDTVGTMMTCAYEDPNCEIGLIVGTGSNACYMEEM---RNVEMVD-------------
--GE------QG---------------------RMCVNMEWGAFGDNGC-LDDIRTIYDKAVDDYSLNAG
KQR-YEKMISGMYLGEIVRNILIDFTKRGFLFRGQISETLKTRHIFETKFLSQIES--------------
------------DRLALLQVRTILQQLGL-NSTCDDSIIVKTVCGAVSRRAAQLCGAGMAAVVDKIRENR
GLEHLEITVGVDGTLYKLH-----------PHFSRIMHQTVKDLAPKCDVTF---------LLSEDG---
----------SGKGAALITAVGCRLRD--AEQ--N-----------------
>XP_039409749|Corvus cornix cornix|HK1
LYAMRLSDETLLDVMARFRREMKNGL-SRDFNPTAV--VKMLPTFVRSIPDGSEKGDFIALDLG--GSYF
RILRVKVSHE---KK-QT--VQMESEIYNTPEDIMHGSGTR---------------LFDHVAECLGDFME
KQ-----QIK---DKKLPVGFTFSFPCRQSKLD------------------------E------------
-----------GILITWTKRFKASGVEGADVVRLLNKAIKKR----------G-----------------
--------------DYDADIMAVVNDTVGTMMTCGFDDQRCEVGLII-----------------------
-----G-----TGTNACYMEEMRHI-DLVEGDE--GRMCINTEWGAFGDDGSLEDIRTEFDREIDRGSLN
PGKQLFEKMVSGLYMGELVRLILVKMAKEGLLFEGRITPELLTKGKFETKHVSAIEK-------------
--SK--EGLNKAKEILTR-LGVEPSHEDCIAVQHVCTIVSFRSANLVASTLGAILNQLRDNK-GVGRLRT
TVGVDGSLYKMH-PQYARRLHKTTRRLVPDSEVRFLLSESGSGKGAAMVTAVAYRLSEQHRLIDETLAEF
KLTHEQLLQVKKRMRAEMEAGLKKKTHETA-KVKMLPTFVRSTPDGT-----------------------
---ENGDFLALDLGGTNFRVLLVKIRSGKRRTVEMHNKIYAIPIEVMQGTGEE-------LFDHIVTCIS
DFLDYMGIKGA-RLPLGFTFSFPCKQTSLDAGILLNWTKGFKATDCEGEDVVYLLREGIKRRE----EFD
LDVVA-----VVNDTVGTMMTCAYEDPNCEIGLIVGTGSNACYMEEM---RNIEMVD-------------
--GE------QG---------------------RMCVNTEWGAFGDNGC-LDDIRTIYDKAVDDFSLNAG
KQR-YEKMISGMYLGEIVRNILIDFTKRGFLFRGQISETLKTRHIFETKFLSQIES--------------
------------DRLALLQVRTILQQLGL-NSTCDDSIIVKTVCGAVSRRAAQLCGAGMAAIVDKIRENR

GLEHLEITVGVDGTLYKLH-----------PHFSRIMHQTVKDLAPNCDVTF---------LLSEDG---
----------SGKGAALITAVGCRLRE--AEQ--N-----------------
>XP_050161742|Myiozetetes cayanensis|HK1
LYAMRLSDETLLDVMARFRREMKNGL-SRDFNPTAA--VKMLPTFVRSIPDGSEKGDFIALDLG--GSYF
RILRVKVSHE---KK-QT--VQMESEIYNTPEDIMHGSGTR---------------LFDHVAECLGDFME
KQ-----QIK---DKKLPVGFTFSFPCRQSKLD------------------------E------------
-----------GILITWTKRFKASGVEGADVVRLLNKAIKKR----------G-----------------
--------------DYDADIMAVVNDTVGTMMTCGFDDQRCEVGLII-----------------------
-----G-----TGTNACYMEEMRHI-DLVEGDE--GRMCINTEWGAFGDDGSLEDIRTEFDREIDRGSLN
PGKQLFEKMVSGLYMGELVRLILVKMAKEGLLFEGRITPELLTKGKFETKHVSAIEK-------------
--SK--EGLNKAKEILTR-LGVEPSHEDCIAVQHVCTIVSFRSANLVASTLGAILNQLRDNK-GVGRLRT
TVGVDGSLYKMH-PQYARRLHKTTRRLVPDSEVRFLLSESGSGKGAAMVTAVAYRLSEQHRLIDETLAEF
KLTHEQLLQVKKRMRAEMEAGLKKKTHETA-KVKMLPTFVRSTPDGT-----------------------
---ENGDFLALDLGGTNFRVLLVKIRSGKRRTVEMHNKIYAIPIEVMQGTGEE-------LFDHIVTCIS
DFLDYMGIKGA-RLPLGFTFSFPCKQTSLDAGILLNWTKGFKATDCEGEDVVYLLREGIKRRE----EFD
LDVVA-----VVNDTVGTMMTCAYEDPNCEIGLIVGTGSNACYMEEM---RNIEMVD-------------
--GE------QG---------------------RMCVNTEWGAFGDNGC-LDDIRTIYDKAVDDYSLNAG
KQR-YEKMISGMYLGEIVRNILIDFTKRGFLFRGQISETLKTRHIFETKFLSQIES--------------
------------DRLALLQVRTILQQLGL-NSTCDDSIIVKTVCGAVSRRAAQLCGAGMAAIVDKIRENR
GLERLEITVGVDGTLYKLH-----------PHFSRIMHQTVKDLAPNCDVTF---------LLSEDG---
----------SGKGAALITAVGCRLRE--AEQ--N-----------------
>XP_057229717|Malurus melanocephalus|HK1
LYAMRLSDETLLDVMARFRREMKNGL-SRDFNPTAA--VKMLPTFVRSIPDGSEKGDFIALDLG--GSYF
RILRVKVSHE---KK-QT--VQMESEIYNTPEDIMHGSGTR---------------LFDHVAECLGDFME
KQ-----QIK---DKKLPVGFTFSFPCRQSKLD------------------------E------------
-----------GILITWTKRFKASGVEGADVVRLLNKAIKKR----------G-----------------
--------------DYDADIMAVVNDTVGTMMTCGFDDQRCEVGLII-----------------------
-----G-----TGTNACYMEEMRHI-DLVEGDE--GRMCINTEWGAFGDDGSLEDIRTEFDREIDRGSLN
PGKQLFEKMVSGLYMGELVRLILVKMAKEGLLFEGRITPELLTKGKFETKHVSAIEK-------------
--SK--EGLNKAKEILTR-LGVEPSHEDCIAVQHVCTIVSFRSANLVASTLGAILNQLRDNK-GVGRLRT
TVGVDGSLYKMH-PQYARRLHKTTRRLVPDSEVRFLLSESGSGKGAAMVTAVAYRLSEQHRLIDETLAEF
KLTHEQLLQVKKRMRAEMEAGLKKKTHETA-KVKMLPTFVRSTPDGT-----------------------
---ENGDFLALDLGGTNFRVLLVKIRSGKRRTVEMHNKIYAIPIEVMQGTGEE-------LFDHIVTCIS
DFLDYMGIKGA-RLPLGFTFSFPCKQTSLDAGILLNWTKGFKATDCEGEDVVYLLREGIKRRE----EFD
LDVVA-----VVNDTVGTMMTCAYEDPNCEIGLIVGTGSNACYMEEM---RNIEMVD-------------
--GE------QG---------------------RMCVNTEWGAFGDNGC-LDDIRTIYDKAVDDFSLNAG
KQR-YEKMISGMYLGEIVRNILIDFTKRGFLFRGQISETLKTRHIFETKFLSQIES--------------
------------DRLALLQVRTILQQLGL-NSTCDDSIIVKTVCGAVSRRAAQLCGAGMAAIVDKIRENR
GLERLEITVGVDGTLYKLH-----------PHFSRIMHQTVKDLAPNCDVTF---------LLSEDG---
----------SGKGAALITAVGCRLRE--AEQ--N-----------------
>XP_005153748|Melopsittacus undulatus|HK1
LYAMRLSDETLIDVMARFRREMKNGL-SRDFNPTAA--VKMLPTFVRSIPDGSEKGDFIALDLG--GSYF
RILRVKVSHE---KK-QT--VQMESEIYNTPEDIMHGSGTR---------------LFDHVAECLGDFME
KQ-----QIK---DKKLPVGFTFSFPCRQSKLD------------------------E------------
-----------GILITWTKRFKASGVEGADVVRLLNRAIKKR----------G-----------------
--------------DYDADIMAVVNDTVGTMMTCGFDDQRCEVGLII-----------------------
-----G-----TGTNACYMEEMRHI-DLVEGDE--GRMCINTEWGAFGDDGSLEDIRTEFDREIDRGSLN
PGKQLFEKMVSGLYMGELVRLILVKMAKEGLLFEGRITPELLTKGKFETKHVSAIEK-------------
--SK--EGLNKAKEILTR-LGVEPSHEDCIAVQHVCTIVSFRSANLVASTLGAILNQLRDNK-GVGRLRT
TVGVDGSLYKMH-PQYARRLHKTTRRLVPDSEVRFLLSESGSGKGAAMVTAVAYRLSEQHRLIDETLAEF
KLTHEQLLQVKKRMRAEMEAGLKKKTHETA-KVKMLPTFVRSTPDGT-----------------------
---ENGDFLALDLGGTNFRVLLVKIRSGKRRTVEMHNKIYAIPIEVMQGTGEE-------LFDHIVTCIS

DFLDYMGIKGA-RLPLGFTFSFPCKQTSLDAGILLNWTKGFKATDCEGEDVVYLLREGIKRRE----EFD
LDVVA-----VVNDTVGTMMTCAYEDPNCEIGLIVGTGSNACYMEEM---RNIEMVD-------------
--GE------QG---------------------RMCVNTEWGAFGDNGC-LDDIRTIYDKAVDDYSLNAG
KQR-YEKMISGMYLGEIVRNILIDFTKRGFLFRGQISETLKTRHIFETKFLSQIES--------------
------------DRLALLQVRAILQQLGL-NSTCDDSIIVKTVCGAVSRRAAQLCGAGMAAMVDKIRENR
GLEHLEITVGVDGTLYKLH-----------PHFSRIMHQTVKDLAPNCDVTF---------LLSEDG---
----------SGKGAALITAVGCRLRD--AEQ--N-----------------
>XP_009942739|Opisthocomus hoazin|HK1
LYAMRLSDETLIDIMARFFREMKNGL-SRDFNPTAT--VKMLPTFVRSIPDGSEKGDFIALDLG--GSYF
RILRVKVSHE---KK-QT--VQMESEIYNTPEDIMHGSGTR---------------LFDHVAECLGDFME
KQ-----QIK---DKKLPVGFTFSFPCRQSKLD------------------------E------------
-----------GILITWTKRFKASGVEGADVVRLLNKAIKKR----------G-----------------
--------------DYDADIMAVVNDTVGTMMTCGFDDQRCEVGIII-----------------------
-----G-----TGTNACYMEEMRHI-DLVEGDE--GRMCINTEWGAFGDDGSLEDIRTEFDREIDRGSLN
PGKQLFEKMVSGLYMGELVRLILVKMAKEGLLFEGRITPELLTKGKFETKHVSAIEK-------------
--SK--EGLNKAKEILTR-LGVEPSHEDCIAVQHVCTIVSFRSANLVASTLGAILNQLRDNK-GVGRLRT
TVGVDGSLYKMH-PQYARRLHKTTRRLVPDSEVRFLLSESGSGKGAAMVTAVAYRLSEQHRLIDETLAEF
KLTHEQLLQVKKRMRAEMEAGLKKKTHETA-KVKMLPTFVRSTPDGT-----------------------
---ENGDFLALDLGGTNFRVLLVKIRSGKRRTVEMHNKIYAIPIEVMQGTGEE-------LFDHIVTCIS
DFLDYMGIKGA-RLPLGFTFSFPCKQTSLDAGILLNWTKGFKATDCEGEDVVYLLREGIKRRE----EFD
LDVVA-----VVNDTVGTMMTCAYEDPNCEIGLIVGTGSNACYMEEM---RNIEMVD-------------
--GE------QG---------------------RMCVNTEWGAFGDNGC-LDDIRTIYDKAVDDYSLNAG
KQR-YEKMISGMYLGEIVRNILIDFTKRGFLFRGQISETLKTRHIFETKFLSQIES--------------
------------DRLALLQVRAILQQLGL-NSTCDDSIIVKTVCGAVSRRAAQLCGAGMAAVVDKIRENR
GLEHLEITVGVDGTLYKLH-----------PHFSRIMHQTVKDLAPNCDVTF---------LLSEDG---
----------SGKGAALITAVGCRLRD--AEQ--N-----------------
>XP_026707605|Athene cunicularia|HK1
LYAMRLSDETLIDIMARFFREMKNGL-SRDFNPTAT--VKMLPTFVRSIPDGSEKGDFIALDLG--GSYF
RILRVKVSHE---KK-QT--VQMESEIYNTPEDIMHGSGTR---------------LFDHVAECLGDFME
KQ-----QIK---DKKLPVGFTFSFPCRQSKLD------------------------E------------
-----------GILITWTKRFKASGVEGADVVRLLNKAIKKR----------G-----------------
--------------DYDADIMAVVNDTVGTMMTCGFDDQRCEVGLII-----------------------
-----G-----TGTNACYMEEMRHI-DLVEGDE--GRMCINTEWGAFGDDGSLEDIRTEFDREIDRGSLN
PGKQLFEKMVSGLYMGELVRLILVKMAKEGLLFEGRITPELLTKGKFETKHVSAIEK-------------
--SK--EGLNKAKEILTR-LGVEPSHEDCIAVQHVCTIVSFRSANLVASTLGAILNQLRDNK-GVGRLRT
TVGVDGSLYKMH-PQYARRLHKTTRRLVPDSEVRFLLSESGSGKGAAMVTAVAYRLSEQHRLIDETLAEF
KLTHEQLLQVKKRMRAEMEAGLKKKTHETA-KVKMLPTFVRSTPDGT-----------------------
---ENGDFLALDLGGTNFRVLLVKIRSGKRRTVEMHNKIYAIPIEVMQGTGEE-------LFDHIVTCIS
DFLDYMGIKGA-RLPLGFTFSFPCKQTSLDAGILLNWTKGFKATDCEGEDVVYLLREGIKRRE----EFD
LDVVA-----VVNDTVGTMMTCAYEDPNCEIGLIVGTGSNACYMEEM---RNIEMVD-------------
--GE------QG---------------------RMCVNTEWGAFGDNGC-LDDIRTIYDKAVDDYSLNAG
KQR-YEKMISGMYLGEIVRNILIDFTKRGFLFRGQISETLKTRHIFETKFLSQIES--------------
------------DRLALLQVRAILQQLGL-NSTCDDSIIVKTVCGAVSRRAAQLCGAGMAAVVDKIRENR
GLEHLEITVGVDGTLYKLH-----------PHFSRIMHQTVKELAPNCDVTF---------LLSEDG---
----------SGKGAALITAVGCRLRD--AEQ--N-----------------
>XP_038603089|Tachyglossus aculeatus|HK2
LYHMRLSDENLQEISERFRKEMEKGL-AADTHPTAS--VKMLPTFVRSTPDGTEDGDFLALDLG--GTNF
RVLRVRVTDN---GL-QK--VDMENQIYAIPEDLMRGSGTQ---------------LFDHIAECLANFME
KL-----QIK---NKKLPLGFTFSFPCHQTKLD------------------------E------------
-----------SFLVTWTKGFKSSGVEGRDVVALLRDAIRKR----------G-----------------
--------------DFDLDIVAVVNDTVGTMMTCGYDDQNCEIGLIV-----------------------
-----G-----TGSNACYMEELRHV-ELVEGDE--GRMCVNMEWGAFGDNGELGDFRTDFDREIDAASLN

PGKQLFEKMISGLYLGELVRLVLVRMAKEEQLFGGKLTPELLTAGCLETRDVSAIEG------------
--EK--DGVREAHVVLSR-LGLEPTQEDCVAAHRVCQIVTTRSAGLCAATLAAVLRRLKENK-GEERLRS
TIGVDGSVYKKH-PHFARRLQKTLRRLVPDCDVRFLRSEDGSGKGAAMVTAVAHRLANQQRARQETLETL
RLSPDQLLDVKRRLRLEMDNGLARDTHAAA-SVKMLPSYVRATPDGT----------------------
---EKGDFLALDLGGTNFRVLLVRVRAGLRRGVEMHSKIYSIPQEITQGTGDE-------LFDHIVHCIA
DFLEYMGMKGV-SLPLGFTFSFPCHQTNLDEGILLKWTKGFKATGCEGEDVVALLKEAIHRRE----EFD
LDVVA-----IVNDTVGTMMACGHEDPRCEVGLIVGTGSNVCYMEEL---RHVELVE------------
--GD------EG---------------------RMCINMEWGAFGDNGE-LDHLRTDFDVAVDESSLNVG
RQR-FEKMISGLYLGEIVRTVLLHFTERGLLFRGRISERLKTRGIFETKFLSQIES--------------
------------DGLALLQVRAVLQHLGL-ESTCDDSIIVKEVCTVVARRAAQLCGAGVAAVVDKIRDGR
RLDSLQVTVGVDGTLFKLH-----------PHFSTIMMETVKQLAPNCEVSF---------LPSEDG---
----------SGKGAALIAAVACRIRE--AGQ--R-----------------
> XP_026571063|Pseudonaja textilis|HK2
LYHLRLSDDALRDISKRFRKEMEKGL-RADTNPTAS--VKMLPTFVRSIPDGTEEGDFLALDLG--GTNF
RVLRVKVADN---GS-KK--VEMENRIYDIPEDLMRASGTQ---------------LFDYIADCLANFME
QL-----QIK---ERKLPLGFTFSFPCRQTKLD-----------------------E------------
-----------SFLDTWTKGFKCSGVEGKDVVSLLREAIKKR----------G-----------------
--------------DFEIDVVAVVNDTVGTMMTCGYDDHDCEVGLIV----------------------
-----G-----TGSNACYMEEMRHI-EMVEGDE--GRMCINMEWGAFGDDGSLKDIRTEFDQEIDMGSLN
PGKQLFEKMISGLYMGELVRLILVKMAKEKLLFSGKVPEDLRKKGCFETRYVSAIEK------------
--EK--EGLSKARAVLEQ-LGLEPSHEDCVATQRVCEIVSTRSAHLCGAALAAVLHRLKENR-GLDWLRT
TVGVDGTVYKKH-PHFARRLHKVVRRLLSGHEVRFVRSEDGSGKGAAMVTAVAYRLAAQDEVRQQILRPL
RLSQQQLLEVKGRMQQEMAWGLARQTHAKA-TVKMLPTYVCATPDGT----------------------
---EKGDFLALDLGGTNFRVLLVRVRNGMRRSVEMHNKIYTISEDIMQGTGEE-------LFDHIVSCIS
DFLEYMGMKGA-SLPLGFTFSFPCQQKRLDEGILLQWTKGFSASGCEGENVVGLLKEAILRRK----EFD
LDVVA-----LVNDTVGTMMSCGYEDPLCEVGLIVGTGTNACYMEEL---RNVELVP------------
--GD------QG---------------------RMCVNMEWGAFGDNGC-LEDFWTEFDATVDENSLNPG
KQR-FEKMISGMYLGEIVRNILIDFTKRGLLFRGRISERLKTKGIFKTKFLSQIES--------------
------------DCLALLQVRHILQDLGL-KSTCDDSIIVKEVCTVVARRAAQLCGAGVAAVVDKIRENR
DLDFLKVTVGVDGTLYKLH-----------PHFSKEMDKTVKQLSPKCEVTF---------LQSEDG---
----------SGKGAALITAVACRLRE--AGQ--H-----------------
> XP_043939013|Protopterus annectens|HK2
LYHMRLSDEVLEDISKRFRREMEKGL-GEGTNATAS--VKMLPTFVRSTPDGTENGEFLALDLG--GTNF
RVLRVKVSDN---GR-QK--VEMENQIYAIPEDLMRGSGTQ---------------LFDHIAECLANFME
QL-----NIK---DKKLPLGFTFSFPCQQSKLD-----------------------E------------
-----------SILVNWTKGFKACGVEGRDVVQLLRNSIKRR----------G-----------------
--------------DFDIDIVAVVNDTVGTMMTCGYDDHNCEIGLIV----------------------
-----G-----TGSNACYMEEMRHI-ETVEGDE--GRMCINMEWGAFGDDGVLNGIRTEFDREIDRGSIN
PGKQLFEKMISGMYMGELVRLILVKMAKEGLLFEGRLTPDLLTTGHFETRFVSAIEN------------
--EK--EGIETALELLTK-LGLEPSEDDAIATQRICQIVSTRSSNLCAATLAAVLKRIKENK-GVERLRT
TVGVDGTVYKKH-PQFARRLHKAVRRLVPECDVRFVRSEDGSGKGAAMVTAVASRLISQQIAREKTLEPL
KLTNDQLLKVKQRMFVEMERGLKKETHNSA-TVKMLPTYVRSTPDGT----------------------
---ERGDFLALDLGGTNFRVLLVRVRNGMRRGVEMHNKIYTIPQEVMQGTGEE-------LFDHIVHCIA
DFLEYMGMKGL-ALPLGFTFSFPCHQNNLDQSILLKWTKGFKATGCEGEDVVHLLKEAIHRRE----EFD
LDIVA-----VVNDTVGTMMTCGYEDSCCEVGLIVGTGSNACYMEEM---KNVELVE------------
--GK------EG---------------------RMCINMEWGAFGDNGC-LDDFRTEFDMAVDNLSLNPG
KQI-YEKMISGMYLGEIVRNILIDFTKRGLLFRGRISERLKTRGIFETKFLSQIES--------------
------------DRLALLQVRAVLQHLGL-QSTCDDSIIVKEVCTVVAKRAAQLCGAGMAAVVDKIRLNR
GLTSLKVTVGVDGTLYKLH-----------PHFSVIMQDTVKILSPNCEVTF---------IESEDG---
----------SGKGAALITAVACRVRE--AGQ--R-----------------
> MBN3276225|Polyodon spathula|HK2
LYHLRLSEETLLDVSKRFLKEMKKGL-GRDTNATAT--IKMLPTFVRSTPDGTENGEFLALDLG--GTNF

RVLWVKVLDN---GK-QK--VEMENQIYAIPEDLMRGSGTQ---------------LFDHIAECLANFLE
KL-----KIK---DKKLPLGFTFSFPCQQSKLD------------------------E------------
-----------SILVSWTKGFKSSGVEGRDVVTLLRKAISKR----------G-----------------
--------------DFDIDIVAVINDTVGTMMTCGYDDHDCEIGLIV-----------------------
-----G-----TGTNACYMEEMRHI-DLVDGDE--GRMCVNMEWGAFGDDGVLDDIRTEFDREIDMGSLN
PGKQLFEKMISGMYMGELVRLILVKMAKEGQLFGGRITPDLLTTGHFETTFVSAIEK-------------
--EK--EGLSKAEEILSQ-LGLEPSKEDCVATQRICKIVSERAANLCAATLAAVLGQIKENK-GVEQLKS
TVGVDGSVYKKH-PQFARHLQKMVRRLAPGCVVRFLRSEDGSGKGAAMVTAVAYRLISQKEERRATLDAF
RLSREQLARVKQSMREEMERGLRKETHDAA-TVKMLPTYVRSTPDGT-----------------------
---EKGDFLALDLGGTNFRVLLVRVRNG-KRTVEMHNKIYSIPLEAMQGTGEE-------LFDHIVHCIA
DFLEYMGMKGL-ALPMGFTFSFPCQQKSLDEGILLKWTKGFKATGCEGEDVVRLLKDAIHRRE----EFD
LDVVA-----VVNDTVGTMMTCGYEDPCCEVGLIVGTGSNACYMEEM---CNVELVD-------------
--GD------EG---------------------HMCVNMEWGAFGDNGC-LDDVRTPFDCAVDELSLNPG
KQK-FEKMISGMYLGEIVRNILIDFTKRGLLFRGRISERLKTRGIFETKFLSQIES--------------
------------DRLALRQVRSILQHLGL-ESTCDDSIIVKEVCTAVACRGAQLCGAGLAAVVDKIRENR
GLDSLRVTVGVDGTLYKLH-----------PHFSNVLQETVRDLAPKCTVSF---------LQSEDG---
----------SGKGAALITAVACRIRE--AGQ--H-----------------
>XP_039624365|Polypterus senegalus|HK2
LYHMRLSDETLLEVSNRFRKEMEKGL-GRDTNPTAS--VKMLPTFVCSTPDGTEKGDFLALDLG--GTNF
RVLYVKVSDN---GK-QK--VEMQNQIYAIPENLMRGSGSQ---------------LFDHIAECLANFLE
KL-----EIK---DKKLPLGFTFSFPCQQNKLD------------------------E------------
-----------SILLTWTKGFKASGVEGRDVVGLLRKAIRKR----------G-----------------
--------------DFDIDVVAVVNDTVGTMMTCGYDDHNCEVGLII-----------------------
-----G-----TGTNACYMEEMRHI-DLVEGDE--GRMCINMEWGAFGDDGVLDDVRTEFDREIDMGSLN
PGKQLFEKMISGMYMGELVRLIMVKMAKEDLLFGGKITPNLLTTGHFETRFVSAIEK-------------
--DK--EGLEKANEVLSK-LGLDPSHEDCLATQRICQIVSERAAKLCAATLAGVLRQIKENK-AVDRLRT
TVGVDGSVYKKH-PQFARRLHKMVRRLVPDCDVRFLRSEDGSGKGAAMVTAVAYRLASQQAERQTTLDMF
RLNKEQLLEIKRRMQVEMERGLKKETHSSA-TVKMLPTYVRSTPDGT-----------------------
---EKGDFLALDLGGTNFRVLLVRVRNGMRRGVEMHNKIYSIPAEVMQGTGEE-------LFDHIVHCIS
DFLEYMGMKGV-SLPMGFTFSFPCHQNNLDEGILLKWTKGFKATGCEGEDVVTLLKEAIHRRE----EFD
LDVVA-----VVNDTVGTMMTCGYEDPYCEVGLIVGTGSNACYMEEM---RNVDLVS-------------
--GE------EG---------------------RMCINMEWGAFGDNGC-LDDFRTPFDKSVDDLSLNPG
KQR-YEKMISGMYLGEIVRNILIDFTKRGLLFRGRISERLKTRSIFETKFLSQIES--------------
------------DRLALRQVRSILQHLGL-DSTCDDSIIVKEVCTAVARRAAQLCGAGLAAVVDRIRENR
GLDFLKITVGVDGTLYKLH-----------PHFATVLHQTVQDLAPNCSVTF---------LQSEDG---
----------SGKGAALITAVACRIRE--AGQ--R-----------------
>XP_005988265|Latimeria chalumnae|HK2
LYHMRLSDEQLQDISNRFMKEMEKGL-GRATNPTAS--VKMLPTFVRSTPNGTENGEFLALDLG--GTNF
RVLRVKVSDN---GS-QQ--VEMENQIYAIPEHIMRGSGTQ---------------LFDHIAECLANFMD
KL-----NIK---DKRLPLGFTFSFPCHQTKLD------------------------E------------
-----------SILVTWTKGFKSSGVEGKDIVSLLRKAIQKR----------G-----------------
--------------DFETDIVAVINDTVGTMMSCGFDDHNCEVGLIV-----------------------
-----G-----TGTNACYMEEMQHI-DLVEGDE--GRMCINMEWGAFGDDGVLDDIRTEFDREIDMGSLN
PGKQLFEKMISGMYMGELVRLIMVKMAKEGLLFGGKSTPDLLTTGHFETRFVSAIEK-------------
--DK--EGVQKAHEILTN-LGLEPSKEDCVATHRICQIVSTRSSNLCAATLAGVLKRIKNNK-GTERLRS
SVGVDGTVYKKH-PQFARRLHKAVRRLVPDCDIRFIRSEDGSGKGAAMVTAVAYRLLSQHKTRQKILDDF
RLSHAQLLEVKSRMRIEMEKGLKRETRNSA-AVKMLPTYVRSTPDGT-----------------------
---EKGDFLALDLGGTNFRVLLVRVRSGMRRGVEMHNKIYSIPQEVMQGTGEE-------LFDHIVHCIA
DFLEYMGMKGV-SLPLGFTFSFPCQQNNLDEGILLKWTKGFNATGCEGKNVVALLKEAIHRIE----DFD
LDVVA-----VVNDTVGTMMTCGYDHPRCEVGLIVGTGSNACYMEEM---KNVELVS-------------
--GE------EG---------------------QMCVNMEWGAFGDNCC-LDDIRTEFDRSVDEFSLNPG
KQI-YEKMISGMYLGEIVRNILIDFTKRGLLFRGRISERLKTKGIFETKFLSQIES--------------

------------DRLALLQVRSILQHLGL-KSTCDDSIIVKEVCTAVARRAAQLCGAGIAAVVDKIRENR
DLDLLKVTVGVDGTLYKLH-----------PHFSAVMQETVKELAPNCEVTF---------VQSEDG---
----------SGKGAALITAVACRIRE--AGQ--R-----------------
>XP_029460023|Rhinatrema bivittatum|HK2
LYHLRLSDEILHDVSNRFRKEMEKGL-RADTNPTAS--VKMLPTFVRSTPDGTEDGDFLALDLG--GTNF
RVLRVKVSNN---GL-KK--VEMENQIYAIPEDLMRASGTQ---------------LFDHIAECLANFME
TL-----HIK---DKKLPLGFTFSFPCHQTKLD-----------------------E-----------
-----------SILVTWTKGFKASGVEGKDVVSLLRKAIKRR----------G----------------
--------------DFDIDVVAVVNDTVGTMMTCGYDDHNCEIGLIV----------------------
-----G-----TGTNSCYMEEMRHI-DLVEGDE--GRMCINMEWGAFGDDGSLNDIRTEFDHEIDMGSLN
PGKQLFEKMISGMYMGELVRLILVKMAKEDLLFMGRITPELLTTGHFKTGFVSAIEQ------------
--DK--EGIQKAYDILTQ-LGLEPSHEDCVATQRICQIVSTRSANLCAATLAAVLHRIKENK-GVERLRS
SIGVDGSVYKKH-PHFAKRLHKTVRRLAPGCDTRFIRSEDGSGKGAAMITAVAYRLAAQQETRQETLETL
KLNFEQLLAVKKKLKMEMERGLGKESHSSA-TVKMLPTYVRATPDGT----------------------
---EKGDFLALDLGGTNFRVLLVRVRNGMRRGVEMHHKIYSISQDIMQGTGEE-------LFDHIVHCIA
DFLEYMGMKGV-SLPLGFTFSFPCHQSNLDEGILLKWTKGFQATGCEGEDVVNLLKEAIHRRE----EFD
LDVVA-----VVNDTVGTMMTCGYEDPYCEVGLIVGTGSNACYMEEM---HNVELVE------------
--GE------EG---------------------KMCVNMEWGAFGDNGC-LEEFRTDFDRTVDELSLNPG
KQR-FEKMISGMYLGEIVRNILIDFTKRGLLFRGRISERLKTRGIFETKFLSQIES-------------
------------DCLALLQVRAILQHLGL-ESTCDDSIIVKEVCTIVAKRAAQLCGAGIAAVVDKIRENR
RLDFLKVTVGVDGTLYKLH-----------PHFSAVMHKTVKELSPKCEVTF---------LLSEDG---
----------SGKGAALITAVACRVRE--AGQ--R-----------------
>XP_027720948|Vombatus ursinus|HK2
LYHMRLSDETLQEVSKRFRKEMEKGL-GADSHPTAS--VKMLPTFVRSTPDGTEDGDFLALDLG--GTNF
RVLRVRVTDN---GL-QK--VEMENQIYAIPEDIMRGSGTQ---------------LFDHIAECLANFMD
KL-----QIK---NKKLPLGFTFSFPCHQTKLD-----------------------E-----------
-----------SYLVSWTKGFKSSGVEGRDVVSLIRKAIQRR----------G----------------
--------------DFDIDIVAVVNDTVGTMMTCGYDDQNCEIGLIV----------------------
-----G-----TGTNACYMEEMRYI-DTVEGDE--GRMCINMEWGAFGDDGVLDDIRTEFDREIDMGSLN
PGKQLFEKMISGMYMGELVRLILVKMAKEEMLFDGKLSPELLTTGHFETKDVSDIEG------------
--EK--DGIKKALEVLTR-LGLQPSQEDCVATHRICQIVSTRSANLCAATLAAVLRRLKENK-GEDRLRS
TIGVDGSVYKKH-PHFAKRLHKAVRRLVPDCDVRFLRSEDGSGKGAAMVTAVAYRLADQHRARQKTLEAL
KLSHEQLLEVKKNMKKEMERGLGKETHAGA-SVKMLPTYVCATPDGT----------------------
---EKGDFLALDLGGTNFRVLLVRVRNGMRRGVEMHNKIYAIPQEVMHGTGDE-------LFDHIVQCIA
DFLEYMGMKGV-SLPLGFTFSFPCQQNSLDESILLKWTKGFKATGCEGEDVVSLLKEAIHRRE----EFD
LDVVA-----VVNDTVGTMMTCGYEDPHCEVGLIVGTGSNACYMEEM---RNVELVE------------
--GD------EG---------------------RMCINMEWGAFGDNGC-LDDIRTEFDLAVDELSLNPG
KQR-FEKMISGMYLGEIVRNILINFTKRGLLFRGRISERLKTRGIFETKFLSQIES-------------
------------DCLALLQVRTILRHLGL-ESTCDDSIIVKEVCTVVARRAAQLCGAGMAAVVDKIRENR
QLDSLKITVGVDGTLYKLH-----------PHFAKVMHETVKELAPKCDVSF---------LESEDG---
----------SGKGAALITAVACRIRD--AGQ--R-----------------
>XP_004704430|Echinops telfairi|HK2
LYHMRLSDETLQEISERFRKEMEKGL-GATTHPTAS--VKMLPTFVRSTPDGTEHGEFLALDLG--GTNF
RVLRVRVMDN---GL-RR--VEMENQIYAIPEDIMRGSGTQ---------------LFDHIAECLANFMG
QL-----QIK---DKKLPLGFTFSFPCHQTKLD-----------------------E-----------
-----------SFLVTWTKGFKSSGVEGQDVVRLIRKAIQRR----------G----------------
--------------DFDIDIVAVVNDTVGTMMTCGYDDQNCEIGLIV----------------------
-----G-----TGSNACYMEEMRHI-DTVEGDE--GRMCINMEWGAFGDDGTLNDIRTEFDQEVDMGSLN
PGKQLFEKMISGMYMGELVRLILVKMTKEELLFGGKLSSELLTTGLFETKDVSDIEG------------
--EK--DGVRKAHEILVR-LGLQPTHEDCVATHRICQVVSTRSANLCAATLAAVLRRLKENKGGEERLRS
TIGVDGSVYKKH-PHFAKRLHKTVRRLVPDCDVRFLRSEDGSGKGAAMVTAVAYRLADQHRARQKTLESL
KLSQEQLLEVKRRMKVEMEQGLRKETHASA-PVKMLPTYVCATPDGT----------------------

```
---EKGDFLALDLGGTNFRVLLVRVRNGKRRGVEMHNKIYSIPQEVMHGTGDE-------LFDHIVQCIA
DFLEYMGMKGV-SLPLGFTFSFPCQQNRLDESILLKWTKGFKASGCEGEDVVNLLKEAIHRRE----EFD
LDVVA-----VVNDTVGTMMTCGYEDPHCEVGLIVGTGSNACYMEEM---RNVELVE-------------
--GE------EG---------------------RMCVNMEWGAFGDNGC-LDDFRTEFDVAVDELSLNAG
KQR-FEKMISGMYLGEIVRNILIDFTKRGLLFRGRISERLKTRGIFETKFLSQIES--------------
------------DCLALLQVRAILQHLGL-ESTCDDSIIVKEVCAVVARRAAQLCGAGMAAVVDKIRENR
GLDTLKVTVGVDGTLYKLH-----------PHFAKIMHETVQDLAPRCDVSF---------LESEDG---
----------SGKGAALITAVACRIRE--AGQ--R-----------------
>KAG8518722|Galemys pyrenaicus|HK2
LYHMRLSDEELQDISKRFRKEMEKGL-GVTTHPTAS--VKMLPTFVRSTPDGTEHGEFLALDLG--GTNF
RVLRVRVTDN---GL-QK--VEMDNQIYAIPEDIMRGSGTQ---------------LFDHIAECLGNFME
KL-----QIK---DKKLPLGFTFSFPCHQTKLD------------------------E------------
-----------SVLVSWTKGFKCSGVEGRDVVTLIRKAIQRR----------G-----------------
--------------DFDIDIVAVVNDTVGTMMTCGYDDQNCEIALIV-----------------------
-----G-----TGSNACYMEEMRHI-DTVEGDE--GRMCINMEWGAFGDDGALDDFRTEFDREIDMGSLN
PGKQLFEKMISGMYMGELVRLILVKMAKEELLFGGKLSSELLATGRFETKDVSDIEG-------------
--EK--DNIRKAREVLMR-LGMDPTPEDCVATHRICQIVSTRSASLCAATLAAVLRRIKENK-GEERLRS
TIGVDGSVYKKH-PHFAKRLHKAVRRLVPDCDVRFLRSEDGSGKGAAMVTAVAHRLADQHRARQKTLEFL
KLSHEQLLEVKRRMKVEMERGLRKETHAIA-PVKMLPTYVCATPDGT-----------------------
---EKGDFLALDLGGTNFRVLLVRVRNGKRRGVEMHNKIYSIPQEVMHGTGDECPNFSLQLFDHIVQCIA
DFLEYMGMKGV-SLPLGFTFSFPCQQNSLDESILLKWTKGFKASGCEGEDVVTLLKEAIHRRE----EFD
LDVVA-----VVNDTVGTMMTCGYEDPHCEVGLIVGTGSNACYMEEM---RNIELVE-------------
--GE------EG---------------------QMCINMEWGAFGDNGC-LDDLRTEFDVAVDEHSLNPG
KQR-FEKMISGMYLGEIVRNILMDFTKRGLLFRGRISERLKTRGIFETKFLSQIES--------------
------------DCLALLQVRAILHHLGL-ESTCDDSIIVKEVCAVVARRAAQLCGAGMAAVVDKIRENR
GLDTLKVTVGVDGTLYKLH-----------PHFAKVMHETVKDLAPKCEVSF---------LQSEDG---
----------SGKGAALITAVACRIRE--AGQ--R-----------------
> P52789|Homo sapiens|HK2
LYHMRLSDETLLEISKRFRKEMEKGL-GATTHPTAA--VKMLPTFVRSTPDGTEHGEFLALDLG--GTNF
RVLWVKVTDN---GL-QK--VEMENQIYAIPEDIMRGSGTQ---------------LFDHIAECLANFMD
KL-----QIK---DKKLPLGFTFSFPCHQTKLD------------------------E------------
-----------SFLVSWTKGFKSSGVEGRDVVALIRKAIQRR----------G-----------------
--------------DFDIDIVAVVNDTVGTMMTCGYDDHNCEIGLIV-----------------------
-----G-----TGSNACYMEEMRHI-DMVEGDE--GRMCINMEWGAFGDDGSLNDIRTEFDQEIDMGSLN
PGKQLFEKMISGMYMGELVRLILVKMAKEELLFGGKLSPELLNTGRFETKDISDIEG-------------
--EK--DGIRKAREVLMR-LGLDPTQEDCVATHRICQIVSTRSASLCAATLAAVLQRIKENK-GEERLRS
TIGVDGSVYKKH-PHFAKRLHKTVRRLVPGCDVRFLRSEDGSGKGAAMVTAVAYRLADQHRARQKTLEHL
QLSHDQLLEVKRRMKVEMERGLSKETHASA-PVKMLPTYVCATPDGT-----------------------
---EKGDFLALDLGGTNFRVLLVRVRNGKWGGVEMHNKIYAIPQEVMHGTGDE-------LFDHIVQCIA
DFLEYMGMKGV-SLPLGFTFSFPCQQNSLDESILLKWTKGFKASGCEGEDVVTLLKEAIHRRE----EFD
LDVVA-----VVNDTVGTMMTCGFEDPHCEVGLIVGTGSNACYMEEM---RNVELVE-------------
--GE------EG---------------------RMCVNMEWGAFGDNGC-LDDFRTEFDVAVDELSLNPG
KQR-FEKMISGMYLGEIVRNILIDFTKRGLLFRGRISERLKTRGIFETKFLSQIES--------------
------------DCLALLQVRAILQHLGL-ESTCDDSIIVKEVCTVVARRAAQLCGAGMAAVVDRIRENR
GLDALKVTVGVDGTLYKLH-----------PHFAKVMHETVKDLAPKCDVSF---------LQSEDG---
----------SGKGAALITAVACRIRE--AGQ--R-----------------
>XP_052503623|Budorcas taxicolor|HK2
LYHMRLSDETLLEIAKRFRKEMEKGL-GATTHPTAS--VKMLPTFVRSTPDGTEHGEFLALDLG--GTNF
RVLWVRVTDN---GL-QK--VEMENQIYAIPEDIMRGSGTQ---------------LFDHIAECLANFMD
KL-----QIK---DKKLPLGFTFSFPCLQTKLD------------------------E------------
-----------SFLVSWTKGFKSSGVEGKDVVTLIRKAIQRR----------G-----------------
--------------DFDIDIVAVVNDTVGTMMTCGYDDQNCEIGLIV-----------------------
```

-----G-----TGSNACYMEEMRHI-DTVEGDE--GRMCINMEWGAFGDDGALDDIRTEFDQEIDMGSLN
PGKQLFEKMISGMYMGELVRLILVKMAKEELLFGGKLSPELLATGHFETKDVSDIEG-------------
--EK--DGIRKAREILVR-LGLNPTDEDCVATHRVCQIVSTRSASLCAATLAAVLRRIKENK-GVDRLRS
TVGVDGSVYKKH-PQFAKRLHKTVRRLVPDCDVRFLRSEDGSGKGAAMVTAVAYRLADQHRARQKTLESL
KLSREQLLEVKKRMKIEMERGLSKETHAIA-PVKMLPTYVCATPDGT-----------------------
---EKGDFLALDLGGTNFRVLLVRVRNGKRRGVEMHNKIYSIPQEVMHGTGDE-------LFDHIVQCIA
DFLEYMGMKGV-SLPLGFTFSFPCQQNSLDESILLKWTKGFKASGCEGEDVVMLLKEAIHRRE----EFD
LDVVA-----VVNDTVGTMMTCGYEDPHCEVGLIVGTGSNACYMEEM---RNVELVE-------------
--GE------EG---------------------RMCVNMEWGAFGDNGC-LDDLRTEFDAAVDELSLNPG
RQR-FEKMISGMYLGEIVRNILIDFTKRGLLFRGRISERLKTRGIFETKFLSQIES--------------
------------DCLALLQVRAILHHLGL-ESTCDDSIIVKEVCTVVARRAAQLCGAGMAAVVDKIRENR
GLDTLKVTVGVDGTLYKLH-----------PHFAKVMHETVKDLAPKCDVSF---------LESEDG---
----------SGKGAALITAVACRIRE--AGQ--R-----------------
>XP_039096060|Hyaena hyaena|HK2
LYHMRLSDENLLEISQRFRKEMEKGL-GATTHPTAS--VKMLPTFVRSTPDGTEHGEFLALDLG--GTNF
RVLRVRVTDN---GL-QK--VEMENQIYAIPEDLMRGSGTQ---------------LFDHIAECLANFMD
KL-----QIK---DKKLPLGFTFSFPCVQTKLD------------------------E------------
-----------SFLVSWTKGFKSSGVEGKDVVTLIRKAIQRR----------G-----------------
--------------DFDIDIVAVVNDTVGTMMTCGYDDQNCEIGLIV-----------------------
-----G-----TGSNACYMEEMRHI-DMVEGDE--GRMCINMEWGAFGDDGALDDFRTEFDQEIDMGSLN
PGKQLFEKMISGMYMGELVRLILVKMAKEELLFGGKVSPGLLATGQFETKDVSDIEG-------------
--EK--DGIRKAREVLVR-LGIDPTQEDCVATHRVCQIVSTRSASLCAATLAAVLRRIKENK-GEERLRS
TIGVDGSVYKKH-PHFAKRLHKTVRRLVPDCDVRFLRSEDGSGKGAAMVTAVAYRLADQHRARQNTLESL
KLSREQLLEVKRRMKVEMERGLSKETHAIA-PVKMLPTYVCATPDGT-----------------------
---EKGDFLALDLGGTNFRVLLVRVRNGKRRGVEMHNKIYSIPQEVMHGTGDE-------LFDHIVQCIA
DFLEYMGMKGV-SLPLGFTFSFPCQQNSLDESILLKWTKGFKASGCEGEDVVTLLKEAIHRRE----EFD
LDVVA-----VVNDTVGTMMTCGYEDPHCEIGLIVGTGSNACYMEEM---RNVELVE-------------
--GE------EG---------------------RMCVNMEWGAFGDNGC-LDDFRTEFDVAVDELSLNPG
KQR-FEKMISGMYLGEIVRNILIDFTKRGLLFRGRISERLKTRGIFETKFLSQIES--------------
------------DCLALLQVRAILHHLGL-ESTCDDSIIVKEVCTVVARRAAQLCGAGMAAVVDKIRENR
GLDTLKVTVGVDGTLYKLH-----------PHFAKVMHETVKDLAPKCDVSF---------LESEDG---
----------SGKGAALITAVACRIRE--AGQ--R-----------------
>XP_062995434|Elgaria multicarinata webbii|HK2
---MRLPDDTLQEISNRFRKDMEKGL-AEDTNPTAA--VKMLPTFVRSTPDGTEEGDFLALDLG--GTNF
RVLRVKVADN---GS-KK--VEMENQIYAIPEDLMRSSGVQ---------------LFDHIAECLANFME
KL-----EMK---DKKLPLGFTFSFPCYQTKLD------------------------E------------
-----------SVLVTWTKGFKSSGVEGKDVVSLLRQAIKKR----------G-----------------
--------------DFDIDVVAVVNDTVGTMMTCGYDDPNCEVGLIV-----------------------
-----G-----TGSNACYMEEMRHI-ELVEGDE--GRMCINMEWGAFGDDGALNDIRTEFDRDIDMGSLN
PGKQLFEKMISGMYMGELVRLILVKMAKQGLLFHGRPTPELLTMGCFETRYVSAIEK-------------
--EN--EGLLKAEEILVK-LGLEPSHEDCVATRRVCEIVSTRSASLCGAALAAVLRRIKENK-GVDQMRS
TVGVDGTVYKKH-PYFARRLHKTVRRLLPDCDIRFIRSEDGSGKGAAMVTAVAYRLAAQHKARQEILDPL
RLSHDQLLEVKKRMRQEMARGLARESHGQA-TVKMLPTYVCSTPDGK-----------------------
---ENGDFLALDLGGTNFRVLLVRIRNGMRRTVEMHNKIYSIPQDIMQGTGDE-------LFDHIVHCIS
DFLEYMGMKGV-SLPLGFTFSFPCHQKNLDEGILLKWTKGFSATGCEGEDVVGLLKDAIHRRE----EFD
LDVVA-----LVNDTVGTMMTCGYEDPLCEVGLIVGTGSNACYMEEM---QNVELVE-------------
--GD------QG---------------------RMCINMEWGAFGDSGC-LDDFRTAFDAAVDENSLNPG
KQR-FEKMISGMYLGEIVRTILIDFTRRGLLFRRRISERLKTRGIFETKFLSQIES--------------
------------DRLALLQVRSILQHLGL-DSTCDDSIIVKEVCTVIARRAAQLCGAGIAAVVDKIRENR
GLDSLKLTVGVDGTLYKLH-----------PHFSTVMAETVKQLSPKCEVTF---------LQSEDG---
----------SGKGAALITAVACRIRE--AGQ--H-----------------
>XP_053321222|Spea bombifrons|HK2

LYHMRLSDETLLEVSSRFGKQMEKGL-SADTNPTAC--VKMLPTFVRSTPDGTENGDFLALDLG--GTNF
RVLRVKVCDN---GM-KK--VEMENQIYAIPEDLMRGSGMQ---------------LFDHIAECLANFME
KL-----HIK---DKKLPLGFTFSFPCHQSKLD------------------------E------------
-----------SILVNWTKGFKASGVEGKDVVTLLRKAIKKR----------G-----------------
--------------DFDIDIVAVVNDTVGTMMTCGYDDHNCQIGLIV-----------------------
-----G-----TGTNACYMEDMRHI-DLVEGDE--GRMCINMEWGAFGDDGCLDDIRTEFDREIDRGSLN
PGKQLFEKMISGMYMGELVRLILVKMAKEDLLFEGRITADLLTTGHFETRFVSAIEK-------------
--DK--EGIQKAHEILSK-LGLEPSQEDCVATHRICQIVSSRSASLCAATLAAVLQRIRENK-GVERLKS
TVGVDGSVYKKH-PHFARLLHKTVRRLVPECDIRFVRSEDGSGKGAAMVTAVAYRLADQQHEREKTLEAL
NLSTEQLLEIKKIMRIEMERGLREETHDSA-TVKMLPTYVRSTPDGT-----------------------
---EKGDFLALDLGGTNFRVLLVRVRNGMRRGVEMHNKIYSIPQDIMQGTGEE-------LFDHIVHCIA
DFLEYMGMKGV-SLPLGFTFSFPCHQNSLDEGILLKWTKGFKATGCEGEDVVNLLKEAIHRRE----EFD
LDVVA-----IVNDTVGTMMTCGYEDPYCEMGLIVGTGSNACYMEEM---RNVELVE-------------
--GQ------EG---------------------RMCINMEWGAFGDNDC-LEDFRTEFDRAVDELSLNPG
KQR-YEKMISGMYLGEIVRNILIDFTKRGLLFRGKISERLKTRGIFETKFLSQIES--------------
------------DRLALLQVRSILQHLGL-ESTCDDSIIVKEVCTVVARRAAQLCGAGVAAVVDKIRENR
GLDFLKVTVGVDGTLYKLH-----------PHFSTIMRQTVKELSPKCQVTF---------LQSEDG---
----------SGKGAALITAVACRIRE--AGQ--R-----------------
> XP_054853439|Eublepharis macularius|HK2
LYHMRLSEETLQEVSSRFRKEMEKGL-GIDTNPTAS--VKMLPTFVRSTPDGTEEGDFLALDLG--GTNF
RVLRVKVSDN---GS-KK--VEMENQIYAIPEDLMRGSGVQ---------------LFDHIAECLANFME
QL-----QMK---DKKLPLGFTFSFPCYQTKLD------------------------E------------
-----------SILVTWTKGFKSSGVEGKDVVTLLRKAIKKR----------G-----------------
--------------DFDIDIVAVVNDTVGTMMTCGYDDHNCEVGLII-----------------------
-----G-----TGSNACYMEEMRHI-DSVEGDE--GRMCINMEWGAFGDDGVLNDIRTEFDRDIDMGSLN
PGKQLFEKMISGMYMGELVRLILVKMAKEHLLFGGKLTSDLLTTGHFETRYVSAIEK-------------
--EK--EGLQKAYEVLSK-LGLDPTHEDCVATRRICEIVSTRSANLCGAALAAVLRRIKENK-GVDRLRS
TVGVDGTVYKKH-PHFARRLHKMVRCLLPDCDIRFVRSEDGSGKGAAMVTAVAYRLAAQHKTRQKILEPL
RLSHEQLLEVKERMKEEMERGLARATHNEA-TVKMLPTYVCATPDGT-----------------------
---EKGDFLALDLGGTNFRVLLVRIRNGMRRSVEMHNKIYSIPLDIMQGTGEE-------LFDHIVHCIS
DFLEYMGMKGV-SLPLGFTFSFPCHQTRLDEGILLNWTKGFKATGCEGEDVVGLLKDAIHRRE----EFD
LDVVA-----VVNDTVGTMMTCGYEDPYCEVGLIVGTGSNACYMEEM---KNVELVE-------------
--GD------EG---------------------RMCVNMEWGAFGDNGC-LDDFRTEFDVAVDEHSLNPG
KQR-FEKMISGMYLGEIVRNILICFTKRGLLFRGRISQRLKTRGIFETKFLSQIES--------------
------------DRLALLQVRSILQHLGL-ESTCDDSIIVKEVCTAIARRAAQLCGAGMAAVVDKIRENR
GLDFLKVTVGVDGTLYKLH-----------PHFSSVMYETVKQLSPKCEVTF---------LQSEDG---
----------SGKGAALITAVACRIRE--AGQ--H-----------------
> XP_024059147|Terrapene carolina triunguis|HK2
LYHMRLSDETLQEVSERFRKEMEKGL-GVDTNPTSS--VKMLPSFVRSTPDGTENGDFLALDLG--GTNF
RVLRVKVSDN---GL-RK--VEMENQIYAIPEDLMRGSGVQ---------------LFDHIAECLANFME
KL-----NMK---DKKLPLGFTFSFPCHQTKLD------------------------E------------
-----------SILVTWTKGFKSSGVEGKDVVAMIRKAIKKR----------G-----------------
--------------DFDIDLVAVVNDTVGTMMTCGYDDHNCEVGLIV-----------------------
-----G-----TGTNACYMEEMRHI-DLVEGDE--GRMCINMEWGAFGDDGVLNDIRTEFDREIDMGSLN
PGKQLFEKMISGMYMGELVRLILVKMAKEELLFGGRLTPSLLTTGHFETGYVSAIEK-------------
--EK--EGLQKAHEILTK-LGLDPSAEDCIATQHICQIVSTRSANLCGATLAAVLRRIKENK-GVDRLRS
TVGVDGSVYKKH-PHFARRLQKTVRKLLPDCDIRFVRSEDGSGKGAAMVTAVAYRLAAQHKARQKTLEAL
KLSHEQLLEVKNRLRIEMERGLSKASHLDA-TVKMLPTYVCATPDGT-----------------------
---EKGDFLALDLGGTNFRVLLVRVRNGIRRGVEMHNKIYSIPQEIMQGTGEE-------LFDHIVHCIS
DFLEYMGMKGV-SLPMGFTFSFPCQQNNLDEGILLKWTKGFKATGCEGEDVVNLLKEAIHRRE----EFD
LDVVA-----VVNDTVGTMMTCGYEDPYCEVGLIVGTGSNACYMEEM---RNVELVE-------------
--GE------EG---------------------RMCINMEWGAFGDNGC-LDDFRTEFDVAVDELSLNAG

KQR-FEKMISGMYLGEIVRNILIDFTKRGLLFGRISERLKTRGIFETKFLSQIES---------------
------------DCLALLQVRSILQHLGL-ESTCDDSIIVKEVCTVVARRAAQLCGAGMAAVVDKIRENR
KLDFLKVTVGVDGTLYKLH-----------PHFSSIMHETVKQLTPKCEVTF---------LQSEDG---
----------SGKGAALITAVACRIRE--AGQ--R-----------------
>XP_025047675|Alligator sinensis|HK2
LYHMRLSDETIQEVAERFRKEMEKGL-GADTNPTAS--VKMLPTFVRSTPDGTEDGDFLALDLG--GTNF
RVLRVKVSDN---GL-QK--VEMENQIYAIPEDLMRGSGAQ---------------LFDHIAECLANFMD
KL-----GIK---DKKLPLGFTFSFPCHQTKLD------------------------E------------
-----------SFLVTWTKGFKSSGVEGKDVVSMLRKAIKKR----------G-----------------
--------------DFDIDIVAVVNDTVGTMMTCGYDDHNCEIGLIV-----------------------
-----G-----TGTNACYMEEMRHI-DLVEGDE--GRMCINMEWGAFGDDGVLNDIRTEFDREIDMGSLN
PGKQLFEKMISGMYMGELVRLILVKMAKEELLFGGRLTPDLLTTGHFETRYVSAIEK-------------
--EK--EGLQKAHEVLTK-LGLEPSHEDCIATHRICQIVSTRSANLCAATLAAVLRRIKENK-GVERLRS
TVGVDGTVYKKH-PQFARRLHKMVRKLLPDCDIRFIRSEDGSGKGAAMVTAVAYRLAAQHKARQKTLEAL
KLSQEQLLEVKKRMRVEMERGLRKETHPEA-TVKMLPTYVCSTPDGT-----------------------
---EKGDFLALDLGGTNFRVLLVRVRNGMRRGVEMHNKIYSIPQEIMQGTGEE-------LFDHIVHCIS
DFLEYMGMKGV-SLPLGFT----GKELDLLWGILLKWTKGFNASGCEGEDVVNLLKEAIHRRE----EFD
LDVVA-----VVNDTVGTMMTCGYEDPYCEVGLIVGTGSNACYMEEM---RNVELVE-------------
--GE------EG---------------------RMCVNMEWGAFGDNGC-LDDFWTEFDVAVDEHSLNPG
KQR-FEKMISGMYLGEIVRNILIDFTKRGLLFGRISERLKTRGIFETKFLSQIES---------------
------------DCLALLQVRSILQHLGL-ESTCDDSIIVKEVCTVVARRAAQLCGAGMAAVVDKIRENR
QLDFLKVTVGVDGTLYKLH-----------PHFASIMHETVKQLSPKCEVTF---------LQSEDG---
----------SGKGAALITAVACRIRE--AGQ--R-----------------
>XP_064353791|Dromaius novaehollandiae|HK2
LYHMRLSEETLQDVAQRFRKEMEKGL-GADTNPTAS--VKMLPTFVRSTPDGTEDGDFLALDLG--GTNF
RVLRVRVSDN---GL-QK--VEMENQIYAIPEELMRGSGVQ---------------LFDHIAECLANFLD
KL-----SIK---DKKLPLGFTFSFPCRQSKLD------------------------E------------
-----------SILVTWTKGFKCSGVEGKDVVSLLRKSIKKR----------G-----------------
--------------DFDIDIVAVVNDTVGTMMTCGYDDHHCEVGLIV-----------------------
-----G-----TGSNACYMEEMRHI-DLVEGDE--GRMCINMEWGAFGDDGVLNDIRTEFDHEIDMGSLN
PGKQLFEKMISGMYMGELVRLILVKMAKEGLLFGGRLTPDLLTTGHFETRYVSTIEK-------------
--DK--EGLQKAHEILTK-LGLEPSHEDCVATHRICQIVSTRSASLCGATLAAVLQRIKESK-GGDRLRS
TVGVDGSVYKKH-PHFARRLHKTVRKLLPDCDIRFVRSEDGSGKGAAMVTAVAYRLAAQHKARQETLEAL
RLSREQLLEVKQRMRLEMEKGLGKETHAEA-TVKMLPTYVCSTPDGT-----------------------
---EKGDFLALDLGGTNFRVLLVRVRNGMRRGVEMHNKIYSIPLEIMQGTGEE-------LFDHIVHCIS
DFLEYMGMKGV-SLPLGFTFSFPCQQTNLDEGILLKWTKGFKASGCEGEDVVSLLKEAIHRRE----EFD
LDVVA-----VVNDTVGTMMTCGYEDPYCEVGLIVGTGSNACYMEEM---RNVELVE-------------
--GE------EG---------------------RMCVNMEWGAFGDNGC-LDNVWTEFDLAVDELSLNPG
KQR-FEKMISGMYLGEIVRNILIDFTKRGLLFGRISERLKTRGIFETKFLSQIES---------------
------------DRLALLQVRSILQHLGL-ESTCDDSIIVKEVCAVVARRAAQLCGAGMAAVVDKIRENR
GLDFLKVTVGVDGTLYKLH-----------PHFSTVMHETVKQLSPKCEVTF---------LQSEDG---
----------SGKGAALITAVACRIRE--AGQ--R-----------------
>XP_065515161|Lathamus discolor|HK2
LYHMRLSEETLQEVSQRFGKEMEKGL-GADTNPTAS--VKMLPTFVRSTPDGTEDGDFLALDLG--GTNF
RVLRVKVSDN---GL-QK--VEMENQIYAIPEELMRGSGVQ---------------LFDHIAECLANFMD
KL-----KIK---DKKLPLGFTFSFPCHQTKLD------------------------E------------
-----------SILVTWTKGFKCSSVEGKDVVSMLRKSIKKR----------G-----------------
--------------DFDIDIVAVVNDTVGTMMTCGYDDHNCEVGLIV-----------------------
-----G-----TGTNACYMEEMRHI-DLVEGDE--GRMCINMEWGAFGDDGVLKDIRTEFDHEIDMGSLN
PGKQLFEKMISGMYMGELVRLILVKMAKEGLLFGGRLTPDLLTTGHFETRYISAIEK-------------
--EK--EGLQKAHEILSK-LGLEPSHEDCVATLRICQIVSTRSANLCGATLAAVLRRIKDNK-GVERLRS
TVGVDGSVYKKH-PHFARRLHKTVRKLLPDCEIRFVRSEDGSGKGAAMVTAVAYRLAAQHKARQKILEAL

KLSHEQLLEVKQRMRMEMEKGLGKETHAEA-AVKMLPTYVCSTPDGT-----------------------
---EKGDFLALDLGGTNFRVLLVRVRNGMRRGVEMHNKIYSIPVEIMQGTGEE-------LFDHIVHCIS
DFLEYMGMKGV-SLPLGFTFSFPCQQTSLDEGILLKWTKGFKATGCEGEDVVNLLKEAIHRRE----EFD
LDVVA-----VVNDTVGTMMTCGYEDPYCEVGLIVGTGSNACYMEEM---RNVELVE-------------
--GE------EG---------------------RMCINMEWGAFGDNGC-LDDIRTEFDLAVDELSLNPG
KQR-FEKMISGMYLGEIVRNILMDFTKRGLLFRGRISERLKTRGIFETKFLSQIES--------------
------------DCLALLQVRSILQHLGL-ESTCDDSIIVKEVCAVVARRAAQLCGAGMAAVVDKIRENR
GLDFLKVTVGVDGTLYKLH-----------PHFSTVMHETVKQLSPKCEVTF---------LQSEDG---
----------SGKGAALITAVACRIRE--AGQ--R-----------------
>XP_063214541|Chroicocephalus ridibundus|HK2
LYHMRLSEETLQEVSQRFGKEMEKGL-GADTNPTAA--VKMLPTFVRSTPDGTEDGDFLALDLG--GTNF
RVLRVKVSDN---GL-QK--VEMENQIYAIPEELMRGSGVQ---------------LFDHIAECLANFME
KL-----KIK---DKKLPLGFTFSFPCHQTKLD------------------------E------------
-----------SILVTWTKGFKCSSVEGKDVVSLLRKSIKKR----------G-----------------
--------------DFDIDIVAVVNDTVGTMMTCGYDDHNCEVGLIV-----------------------
-----G-----TGTNACYMEEMRHI-DLVEGDE--GRMCINMEWGAFGDDGVLNDIRTEFDREIDMGSLN
PGKQLFEKMISGMYMGELVRLILVKMAKEGLLFGGKLTPDLLTTGHFETRFVSAIEK-------------
--EK--EGLQKAHEILTK-LGLEPSHEDCIATHRICQIVSTRSANLCGATLAAVLRRIKENK-GVDRLRS
TVGVDGSVYKKH-PHFARRLHKTVRKLLPDCEIRFVRSEDGSGKGAAMVTAVAYRLAAQHKARQKILEAL
KLSHDQLLEVKQRMKIEMEKGLGKETHAEA-TVKMLPTYVCSTPDGT-----------------------
---EKGDFLALDLGGTNFRVLLVRVRNGMRRGVEMHNKIYSIPVEIMQGTGEE-------LFDHIVHCIS
DFLEYMGMKGV-SLPLGFTFSFPCQQTNLDEGILLKWTKGFKATGCEGEDVVNLLKEAIHRRE----EFD
LDVVA-----VVNDTVGTMMTCGYEDPYCEVGLIVGTGSNACYMEEM---RNVELVE-------------
--GE------EG---------------------RMCVNMEWGAFGDNGC-LDDIRTEFDLAVDELSLNPG
KQR-FEKMISGMYLGEIVRNILMDFTKRGLLFRGRISERLKTRGIFETKFLSQIES--------------
------------DCLALLQVRSILQHLGL-ESTCDDSIIVKEVCAVVARRAAQLCGAGMAAVVDKIRENR
GLDFLKITVGVDGTLYKLH-----------PHFSTVMHETVKQLSPKCEVTF---------LQSEDG---
----------SGKGAALITAVACRIRE--AGQ--R-----------------
>XP_064897188|Columba livia|HK2
LYDMRLSEETLQEVSQRFGKEMEKGL-GADTNPTAS--VKMLPTFVRSTPDGTEDGDFLALDLG--GTNF
RVLRVKVSDN---GL-QK--VEMENQIYAIPEELMRGSGVQ---------------LFDHIAECLANFME
KL-----KIK---DKKLPLGFTFSFPCHQTKLD------------------------E------------
-----------SILVTWTKGFKCSSVEGKDVVSMLRKSIKKR----------G-----------------
--------------DFDIDIVAVVNDTVGTMMTCGYDDHNCEVGLIV-----------------------
-----G-----TGTNACYMEEMRHI-DLVEGDE--GRMCINMEWGAFGDDGVLNDIRTEFDREIDMGSLN
PGKQLFEKMISGMYMGELVRLILVKMAKEGLLFGGRLTPDLLTTGHFETRYVSAIEK-------------
--EK--EGLQKAHEILSK-LGLEPSHEDCVATHRICQIVSTRSANLCGATLAAVLRRIKENK-GLDRLRS
TVGVDGSVYKKH-PHFARRLHKTVRKLLPDCEIRFVRSEDGSGKGAAMVTAVAYRLAAQHQARQKILEAL
KLSHEQLLEVKQRMRTEMEKGLGKETHAEA-TVKMLPTYVCSTPDGT-----------------------
---EKGDFLALDLGGTNFRVLLVRVRNGMRRGVEMHNKIYSIPVEIMQGTGEE-------LFDHIVHCIS
DFLEYMGMKGV-SLPLGFTFSFPCQQTNLDEGILLKWTKGFKATGCEGEDVVNLLKEAIHRRE----EFD
LDVVA-----VVNDTVGTMMTCGYEDPYCEVGLIVGTGSNACYMEEM---RNVELVE-------------
--GE------EG---------------------RMCVNMEWGAFGDNGC-LDDIRTEFDMAVDELSLNPG
KQR-FEKMISGMYLGEIVRNILMDFTKRGLLFRGRISERLKTRGIFETKFLSQIES--------------
------------DCLALLQVRSILQHLGL-ESTCDDSIIVKEVCTVVARRAAQLCGAGMAAVVDKIRENR
GLDFLKVTVGVDGTLYKLH-----------PHFSTVMHETVKQLSPKCEVTF---------LQSEDG---
----------SGKGAALITAVACRIRE--AGQ--R-----------------
>XP_049687159|Accipiter gentilis|HK2
LYHMRLSEDTLQEVSQRFGKEMEKGL-GADTNPTAS--VKMLPTFVRSTPDGTEDGDFLALDLG--GTNF
RVLRVKVSDN---GL-QK--VEMENQIYAIPEELMRGSGVQ---------------LFDHIAECLANFME
KL-----KIK---DKKLPLGFTFSFPCHQTKLD------------------------E------------
-----------SILVTWTKGFKCSSVEGKDVVSMLRKSIKKR----------G-----------------

--------------DFDIDIVAVVNDTVGTMMTCGYDDHNCEVGLIV-----------------------
-----G-----TGTNACYMEEMRHI-DLVEGDE--GRMCINMEWGAFGDDGVLNDIRTEFDREIDMGSLN
PGKQLFEKMISGMYMGELVRLILVKMAKEGLLFGGRLTPDLLTTGHFETRYVSAIEK-------------
--EK--EGLQKAHEILTK-LGLEPSHEDCVATHRICQIVSTRSANLCGATLAAVLRRIKENK-GVDRLRS
TVGVDGSVYKKH-PHFARRLHKTVRKLLPDCEIRFVRSEDGSGKGAAMVTAVAYRLAAQHKARQKILEAL
KLSHEQLLEVKQRMRLEMEKGLGKETHEEA-AVKMLPTYVCSTPDGT-----------------------
---EKGDFLALDLGGTNFRVLLVRVRNGMRRGVEMHNKIYSIPVEIMQGTGEE-------LFDHIVHCIS
DFLEYMGMKGV-SLPLGFTFSFPCQQNNLDEGILLKWTKGFKATGCEGEDVVNLLKEAIHRRE----EFD
LDVVA-----VVNDTVGTMMTCGYEDPYCEVGLIVGTGSNACYMEEM---RNVELVE-------------
--GE------EG---------------------RMCVNMEWGAFGDNGC-LDDIRTEFDMAVDELSLNPG
KQR-FEKMISGMYLGEIVRNILMDFTKRGLLFRGRISERLKTRGIFETKFLSQIES--------------
------------DCLALLQVRSILQHLGL-ETTCDDSIIVKEVCTVVARRAAQLCGAGMAAVVDKIRENR
GLDFLKVTVGVDGTLYKLH-----------PHFSTVMHETVKQLSPKCEVTF---------LQSEDG---
----------SGKGAALITAVACRIRE--AGQ--R-----------------
>XP_059687487|Gavia stellata|HK2
LYHMRLSEETLQEVSQRFGKEMEKGL-GADTNPTAS--VKMLPTFVRSTPDGTEDGDFLALDLG--GTNF
RVLRVKVSDN---GL-QK--VEMENQIYAIPEELMRGSGVQ---------------LFDHIAECLANFME
KL-----KIK---DKKLPLGFTFSFPCHQTKLD------------------------E------------
-----------SILVTWTKGFKCSSVEGKDVVSMLRKSIKKR----------G-----------------
--------------DFDIDIVAVVNDTVGTMMTCGYDDHNCEVGLIV-----------------------
-----G-----TGTNACYMEEMRHI-DLVEGDE--GRMCINMEWGAFGDDGVLNDIRTEFDREIDMGSLN
PGKQLFEKMISGMYMGELVRLILVKMAKEGLLFGGKLTPDLLTTGHFETRYVSAIEK-------------
--EK--EGLQKAHEILTK-LGLEPSHEDCVATHRICQIVSTRSANLCGATLAAVLRRIKENK-GVDRLRS
TVGVDGSVYKKH-PHFARRLHKTVRKLLPDCEIRFIRSEDGSGKGAAMVTAVAYRLAAQHKARQKILEAL
KLSHEQLLEVKQRMRIEMEKGLGKETHAEA-TVKMLPTYVCSTPDGT-----------------------
---EKGDFLALDLGGTNFRVLLVRVRNGMRRGVEMHNKIYSIPVEIMQGTGEE-------LFDHIVHCIS
DFLEYMGMKGV-SLPLGFTFSFPCQQTNLDEGILLKWTKGFKATGCEGEDVVNLLKEAIHRRE----EFD
LDVVA-----VVNDTVGTMMTCGYEDPYCEVGLIVGTGSNACYMEEM---RNVELVE-------------
--GE------EG---------------------RMCVNMEWGAFGDSGC-LDDIRTEFDLAVDELSLNPG
KQR-FEKMISGMYLGEIVRNILMDFTKRGLLFRGRISERLKTRGIFETKFLSQIES--------------
------------DCLALLQVRSILQHLGL-ETTCDDSIIVKEVCTVVARRAAQLCGAGMAAVVDKIRENR
GLDFLKVTVGVDGTLYKLH-----------PHFSTVMHETVKQLSPKCEVTF---------LQSEDG---
----------SGKGAALITAVACRIRE--AGQ--R-----------------
>NXV88722|Calonectris borealis|HK2
LYHMRLSEETLQEVSQRFGKEMEKGL-GADTNPTAS--VKMLPTFVRSTPDGTEDGDFLALDLG--GTNF
RVLRVKVSDN---GL-QK--VEMENQIYAIPEELMRGSGVQ---------------LFDHIAECLANFME
KL-----KIK---DKKLPLGFTFSFPCHQTKLD------------------------E------------
-----------SILVTWTKGFKCSSVEGKDVVSMLRKSIKKR----------G-----------------
--------------DFDIDIVAVVNDTVGTMMTCGYDDHNCEVGLIV-----------------------
-----G-----TGTNACYMEEMRHI-DLVEGDE--GRMCINMEWGAFGDDGVLNDIRTEFDREIDMGSLN
PGKQLFEKMISGMYMGELVRLILVKMAKEGLLFGGKLTPDLLTTGHFETRYVSAIEK-------------
--EK--EGLQKAHEILTK-LGLEPSHEDCVAIHRICQIVSTRSANLCGATLAAVLRRIKENK-GVDRLRS
TVGVDGSVYKKH-PHFARRLHKTVRKLLPDCEIRFVRSEDGSGKGAAMVTAVAYRLAAQHKARQKILEAL
KLSHEQLLEVKQRMRIEMEKGLGKETHAEA-TVKMLPTYVCSTPDGT-----------------------
---EKGDFLALDLGGTNFRVLLVRVRNGMRRGVEMHNKIYSIPVEIMQGTGEE-------LFDHIVHCIS
DFLEYMGMKGV-SLPLGFTFSFPCQQTNLDEGILLKWTKGFKATGCEGEDVVNLLKEAIHRRE----EFD
LDVVA-----VVNDTVGTMMTCGYEDPYCEVGLIVGTGSNACYMEEM---RNVELVE-------------
--GE------EG---------------------RMCVNMEWGAFGDNGC-LDDIRTEFDLAVDELSLNPG
KQR-FEKMISGMYLGEIVRNILMDFTKRGLLFRGRISERLKTRGIFETKFLSQIES--------------
------------DCLALLQVRSILQHLGL-ETTCDDSIIVKEVCTVVARRAAQLCGAGMAAVVDKIRENR
GLDFLKVTVGVDGTLYKLH-----------PHFSTVMRETVKQLSPKCEVTF---------LQSEDG---
----------SGKGAALITAVACRIRE--AGQ--R-----------------

>XP_027524830|Corapipo altera|HK2
LYHMRLSDETLLEVSQRFEKEMEKGL-GAETNPTAS--VKMLPTFVRSTPDGTEDGDFLALDLG--GTNF
RVLRVKVSDN---GL-QK--VEMENQIYAIPEELMRGSGVQ---------------LFDHITECLANFLD
KL-----NIK---DKKLPLGFTFSFPCHQTKLD------------------------E------------
-----------SILVTWTKGFKCSSVEGKDVVSMLRKSIKKR----------G-----------------
--------------DFDIDIVAVVNDTVGTMMTCGYDDHNCEVGLIV-----------------------
-----G-----TGTNACYMEEMRHI-DLVEGDE--GRMCINMEWGAFGDDGVLNDIRTEFDREIDMGSLN
PGKQLFEKMISGMYMGELVRLILVKMTKEGLLFGGRLTPDLLTTGHFETRYVSAIEK------------
--EK--EGLQKAHEILSK-LGLEPSHEDCVATLRICQIVSTRSASLCGATLAAVLRRIKDNK-GVDRLRS
TVGVDGSVYKKH-PHFARRLHKTVRKLLPDCEIRFIRSEDGSGKGAAMVTAVAYRLDAQHKARQKILEPF
KLSHEQLLEVKERMRKEMEKGLGKETHEEA-TVKMLPTYVCSTPDGT-----------------------
---EKGDFLALDLGGTNFRVLLVRVRNGKRRGVEMHNKIYSIPVEIMQGTGEE-------LFDHIVHCIS
DFLEYMGMKGV-SLPLGFTFSFPCQQNNLDEGILLKWTKGFKATGCEGEDVVGLLKEAIHRRE----EFD
LDVVA-----VVNDTVGTMMTCGYEDPLCEVGLIVGTGSNACYMEEM---RNVELVE-------------
--GD------EG---------------------RMCVNMEWGAFGDSGC-LDDVRTEFDVAVDELSLNPG
KQR-FEKMISGMYLGEIVRNILMAFTKRGLLFRGRISERLKTRGIFETKFLSQIES--------------
------------DCLALLQVRSILQHLGL-ESTCDDSIIVKEVCTVVARRAAQLCGAGMAAVVDKIRENR
GLDFLKVTVGVDGTLYKLH-----------PHFSTVMHQTVKELSPKCEVTF---------LQSEDG---
----------SGKGAALITAVACRIRE--AGQ--R-----------------
>NXT64159|Chaetops frenatus|HK2
LYHMRLSDETLQDVSQRFEKEMEKGL-GAETNPTAC--VKMLPTFVRSTPDGTEDGDFLALDLG--GTNF
RVLRVKVSDN---GL-QK--VEMENQIYAIPEELMRGSGVQ---------------LFDHIAECLANFME
KL-----QIK---DKKLPLGFTFSFPCHQTKLD------------------------E------------
-----------SILVNWTKGFKCSSVEGRDVVSLLRKSIKKR----------G-----------------
--------------DFDIDIVAVVNDTVGTMMTCGYDDHNCEVGLIV-----------------------
-----G-----TGTNACYMEEMRNV-ELVEGDE--GRMCVNMEWGAFGDDGVLNDIRTEFDREIDMGSLN
PGKQLFEKMISGMYMGELVRLILVKMTKEGLLFGGRLTPDLLTTGHFETRFVSAIEK------------
--EK--EGLQKAHEILSK-LGLEPSHEDCVAALRICQVVSTRSASLCGAALAAVLRRIRHNK-GLQRLRS
SVGVDGSVYKKH-PHFARRLHKTVRKLLPDCEIRFIRSEDGSGKGAAMVTAVAYRLAAQHKARQKILEPL
KLSHEQLLEVKARMRMEMEKGLGKETHAEA-TVKMLPTYVCSTPDGT-----------------------
---EKGDFLALDLGGTNFRVLLVRVRNGMRRGVEMHNKIYSIPVEIMQGTGEE-------LFDHIVHCIS
DFLEYMGMKGV-SLPLGFTFSFPCQQNNLDEGILLKWTKGFKATGCEGEDVVGLLKEAIHRRE----EFD
LDVVA-----VVNDTVGTMMTCGYEDPYCEVGLIVGTGSNACYMEEM---RNVELVE-------------
--GD------EG---------------------RMCVNMEWGAFGDSGC-LDDIRTEFDLAVDELSLNPG
KQR-FEKMISGMYLGEIVRNILMDFTKRGLLFRGRISERLKTRGIFETKFLSQIES--------------
------------DCLALLQVRSILQHLGL-ESTCDDSIIVKEVCTVVARRAAQLCGAGMAAVVDKIRENR
GLDFLKITVGVDGTLYKLH-----------PHFSTVMHETVKQLSPKCEVTF---------LQSEDG---
----------SGKGAALITAVACRIRE--AGQ--R-----------------
>NXC06527|Orthonyx spaldingii|HK2
LYHMRLSDETLLEVSQRFEKEMEKGL-GADTNPTAS--VKMLPTFVRSTPDGTEDGDFLALDLG--GTNF
RVLRVKVSDN---GL-QK--VEMENQIYAIPEELMRGSGVQ---------------LFDHIAECLANFME
KL-----QIK---DKKLPLGFTFSFPCHQTKLD------------------------E------------
-----------SILVNWTKGFKCSSVEGRDVVSMLRKSIKKR----------G-----------------
--------------DFDIDIVAVVNDTVGTMMTCGYDDHNCEVGLIV-----------------------
-----G-----TGTNACYMEEMRHI-DLVEGDE--GRMCINMEWGAFGDDGVLNDIRTEFDREIDMGSLN
PGKQLFEKMISGMYMGELVRLILVKMAKEGLLFGGRLTPDLLTTGHFETRFVSAIEK------------
--EK--EGLQKAHEILSK-LGLEPSHEDCVATLRICQIVSTRSASLCGATLAAVLRRIKDNK-GVERLRS
TVGVDGSVYKKH-PHFARRLHKTVRKLLPDCEIRFIRSEDGSGKGAAMVTAVAYRLAAQHKARQKILEPL
KLSHEQLLEVKERMRAEMEKGLGKETHAEA-TVKMLPTYVCSTPDGT-----------------------
---EKGDFLALDLGGTNFRVLLVRVRNGMRRGVEMHNKIYSIPVEIMQGTGEE-------LFDHIVHCIS
DFLEYMGMKGV-SLPLGFTFSFPCQQNNLDEGILLKWTKGFKATGCEGEDVVGLLKEAIHRRE----EFD
LDVVA-----VVNDTVGTMMTCGYEDPYCEVGLIVGTGSNACYMEEM---RNVELVE-------------

--GD------EG---------------------RMCVNMEWGAFGDSGC-LDDIRTEFDVAVDELSLNPG
KQR-FEKMISGMYLGEIVRNILMDFTKRGLLFRGRISERLKTRGIFETKFLSQIES--------------
------------DCLALLQVRSILQHLGL-ESTCDDSIIVKEVCTVVARRAAQLCGAGMAAVVDKIRENR
GLDFLKITVGVDGTLYKLH-----------PHFSTVMHETVKELSPKCEVTF---------LQSEDG---
----------SGKGAALITAVACRIRE--AGQ--R-----------------
>NXU36883|Drymodes brunneopygia|HK2
LYHMRLSDETLQEVSQRFEKEMEKGL-GAETNPTAS--VKMLPTFVRSTPDGTEDGDFLALDLG--GTNF
RVLRVKVSDN---GL-QK--VEMENQIYAIPEELMRGSGVQ---------------LFDHIAECLANFMD
KL-----QIK---DKKLPLGFTFSFPCHQTKLD------------------------E------------
-----------SILVNWTKGFKCSSVEGRDVVSLLRKSIKKR----------G-----------------
--------------DFDIDIVAVVNDTVGTMMTCGYDDHNCEVGLIV-----------------------
-----G-----TGTNACYMEEMRHI-DLVEGDE--GRMCINMEWGAFGDDGVLNDIRTEFDREIDMGSLN
PGKQLFEKMISGMYMGELVRLILVKMTKEGLLFGGRLTPDLLTTGHFETRFVSAIEK-------------
--EK--EGLQKAHEILSK-LGLEPSHEDCLATLRICQIVSTRSASLCGATLAAVLRRIKDNK-GVDRLRS
TVGVDGSVYKKH-PHFARRLHKTVRKLLPDCEIRFIRSEDGSGKGAAMVTAVAYRLAAQHKARQKILEPL
KLSHEQLLEVKERMRIEMEKGLGKETHAEA-TVKMLPTYVCSTPDGT-----------------------
---EKGDFLALDLGGTNFRVLLVRVRNGMRRGVEMHNKIYSIPVEIMQGTGEE-------LFDHIVHCIS
DFLEYMGMKGV-SLPLGFTFSFPCQQNNLDEGILLKWTKGFKATGCEGEDVVGLLKEAIHRRE----EFD
LDVVA-----VVNDTVGTMMTCGYEDPYCEVGLIVGTGSNACYMEEM---RNVELVE-------------
--GD------EG---------------------RMCVNMEWGAFGDSGC-LDDIRTEFDLAVDELSLNPG
KQR-FEKMISGMYLGEIVRNILMDFTKRGLLFRGRISERLKTRGIFETKFLSQIES--------------
------------DCLALLQVRSILQHLGL-ESTCDDSIIVKEVCTVVARRAAQLCGAGMAAVVDKIRENR
GLDFLKITVGVDGTLYKLH-----------PHFSTVMHETVKQLSPKCEVTF---------LQSEDG---
----------SGKGAALITAVACRIRE--AGQ--R-----------------
>NWV05970|Ptilonorhynchus violaceus|HK2
LYHMRLSDETLLEVSQRFEKEMEKGL-GADTNPTAS--VKMLPTFVRSTPDGTEDGDFLALDLG--GTNF
RVLRVKVSDN---GL-QK--VEMENQIYAIPEELMRGSGVQ---------------LFDHIAECLANFMD
KL-----QIK---DKKLPLGFTFSFPCHQTKLD------------------------E------------
-----------SILVTWTKGFKCSSVEGRDVVSMLRKSIKKR----------G-----------------
--------------DFDIDIVAVVNDTVGTMMTCGYDDHNCEVGLIV-----------------------
-----G-----TGTNACYMEEMRHI-DLVEGDE--GRMCINMEWGAFGDDGVLNDIRTEFDREIDMGSLN
PGKQLFEKMISGMYMGELVRLILVKMAKEGLLFGGKLSPDLLATGHFETRFVSAIEK-------------
--EK--EGLQKAHEILSK-LGLEPSHEDCLATLRICQIVSTRSASLCGATLAAVLRRIKDNK-GVERLRS
TVGVDGSVYKKH-PHFARRLHKTVRKLLPDCEIRFIRSEDGSGKGAAMVTAVAYRLAAQHKARQKILEPL
KLSHEQLLEVKERMRIEMEKGLGKETHAEA-TVKMLPTYVCSTPDGT-----------------------
---EKGDFLALDLGGTNFRVLLVRVRNGMRRGVEMHNKIYSIPVEIMQGTGEE-------LFDHIVHCIS
DFLEYMGMKGV-SLPLGFTFSFPCQQNNLDEGILLKWTKGFKATGCEGEDVVGLLKEAIHRRE----EFD
LDVVA-----VVNDTVGTMMTCGYEDPCCEVGLIVGTGSNACYMEEM---RNVELVE-------------
--GD------EG---------------------RMCVNMEWGAFGDSGC-LDDIRTEFDLVVDELSLNPG
KQR-FEKMISGMYLGEIVRNILMDFTKRGLLFRGRISERLKTRGIFETKFLSQIES--------------
------------DCLALLQVRSILQHLGL-ESTCDDSIIVKEVCTVVARRAAQLCGAGMAAVVDKIRENR
GLDFLKVTVGVDGTLYKLH-----------PHFSTVMHETVKQLSPKCEVTF---------LQSEDG---
----------SGKGAALITAVACRIRE--AGQ--R-----------------
>NXY14364|Atrichornis clamosus|HK2
LYHMRLSDETLLEVSQRFEKEMEKGL-GAETNPTAS--VKMLPTFVRSTPDGTEDGDFLALDLG--GTNF
RVLRVKVSDN---GL-QK--VEMENQIYAIPEELMRGSGVQ---------------LFDHIAECLANFMD
KL-----QIK---DKKLPLGFTFSFPCHQTKLD------------------------E------------
-----------SILVTWTKGFKCSSVEGKDVVSMLRKSIKKR----------G-----------------
--------------DFDIDIVAVVNDTVGTMMTCGYDDHNCEVGLIV-----------------------
-----G-----TGTNACYMEEMRHI-DLVEGDE--GRMCINMEWGAFGDDGVLNDIRTEFDREIDMGSLN
PGKQLFEKMISGMYMGELVRLILVKMAKEGLLFGGRLTPDLLTTGHFETRFVSAIEK-------------
--EK--EGLQKAHEILSK-LGLEPSHEDCLATLRICQIVSTRSASLCGATLAAVLRRIKDNK-GVERLRS

TVGVDGSVYKKH-PHFARRLHKTVRKLLPDCEIRFIRSEDGSGKGAAMVTAVAYRLAAQHKARQKILEAL
KLSHEQLLEVKERMRIEMEKGLGKETHADA-TVKMLPTYVCSTPDGT-----------------------
---EKGDFLALDLGGTNFRVLLVRVRNGMRRGVEMHNKIYSIPVEIMQGTGEE-------LFDHIVHCIS
DFLEYMGMKGV-SLPLGFTFSFPCQQNNLDEGILLKWTKGFKATGCEGEDVVGLLKEAIHRRE----EFD
LDVVA-----VVNDTVGTMMTCGYEDPYCEVGLIVGTGSNACYMEEM---RNVELVE-------------
--GD------EG---------------------RMCVNMEWGAFGDSGC-LDDIRTEFDLAVDELSLNPG
KQR-FEKMISGMYLGEIVRNILMDFTKRGLLFRGRISERLKTRGIFETKFLSQIES--------------
------------DCLALLQVRSILQHLGL-ESTCDDSIIVKEVCTVVARRAAQLCGAGMAAVVDKIRENR
GLDFLKVTVGVDGTLYKLH-----------PHFSTVMHETVKQLSPKCEVTF---------LQSEDG---
----------SGKGAALITAVACRIRE--AGQ--R-----------------
> XP_032870023|Amblyraja radiata|HK2
LYHMRLSDEVLHDISERFLKEMKRGL-GKDTNPTSS--VKMLPSFVRSTPDGTESGDFLALDLG--GTNF
RVLRVKVSDN---GK-QK--VVIENRICSIPEEVMHGTGTQ---------------LFDHIAKCLADVMR
EL-----DIK---DTKLPLGFTFSFPCQQSKLD------------------------E------------
-----------SILLTWTKGFKASGVEGQDVVGLLHKAIDRR----------G-----------------
--------------DFDIDIVAVVNDTVGTMMSCGYEDHNCEIGLIV-----------------------
-----G-----TGTNACYMEEMRHI-DLVEGDE--GRMCINMEWGAFGDQGELADVRTEFDTEIDSGSIN
PGKQLFEKMISGMYMGELVRLILVKMTKDGLLFRGKLTPQLLVKDNFETQYVSAIEK-------------
--DS--DGLQRAHGVLTK-LGLEPTSDDCISLRRICSIVSTRSSDLCAAALAAVLRRIQENK-GLQRMQT
TIGVDGSVYKMH-PQFARRLNKTVKRLVPSCEVRFLLSEDGSGKGAAMVTAVAHRLDKQQKMRDECLSTF
KLSEKQLLEVKSRMRLEMERGLKKATHSIA-TVKMLATYVRATPDGT-----------------------
---EKGDYLALDLGGTNFRVLLVKIRDGNRRAVELHNKIYTIPKHAIEGTGEE-------LFDHIVQCIA
DFLKYMGIQGS-RLPLGFTFSFPCEQRTLDQGILLNWTKGFKATGCEGKDVVNLLREAIKRRE----EFD
LDVVA-----IVNDTVGTMMSCGYDDPLCEVGLIVGTGSNACYMEEM---KNVEMVD-------------
--GD------EG---------------------RMCVNMEWGAFGDNGC-LDDIRTEFDLEVDENSLYAG
KQR-FEKMISGMYLGEIVRNILIKFTKQGLLFRGKISERLKTQGIFETKFLSQIES--------------
------------DRLALLQVRAILQQLGL-ESTCDDSIIVKEVCGVVSRRAAQLCGIGLAAIVDKIRVNR
NLDNLKVTVGIDGTLYKLH-----------PHFSTIMQETVQRMSPKCKVLF---------VQSEDG---
----------SGKGAALITAVACRLRE--AGQ--H-----------------
> XP_042200526|Callorhinchus milii|HK2
LYHMRLSDETLQDISKRFLVEMERGL-GRDTNPTSS--VKMLPTFVRSTPDGTESGDFLALDLG--GTNF
RVLRVKVSDN---SR-QR--VEMENQIFSIPEDLMRGSGSQ---------------LFDHIADCLGSFME
KL-----GIK---DQKLPLGFTFSFPCQQTKLD------------------------E------------
-----------SVLVNWTKGFKSSGVEGRDVVTLLRQAIQKR----------G-----------------
--------------DFDIDIVAVINDTVGTMMTCGYDDHNCEVGLIV-----------------------
-----G-----TGTNACYMEEMRHI-DLVEGDE--GRMCINMEWGGFGDQNALDDVRTEFDREIDRGSLN
PGKQLYEKMISGMYMGELVRLILVKMTKEGLLFGGKISPELMVHGHFETKYVSAIEK-------------
--DG--DGLQKAHEILSK-LGLEPSQEDCIATQRICTIVSTRSANLCAATLAAVLRRIKENK-GLQRLRS
TIGVDGTVYKKH-PQFARRLHKAVRRLVPDCDVRFLRSEDGSGKGAAMVTAVAYRLVAQRKAREETLAAF
ELSHQQLLKVKARMRQEMEQGLQRETHTSA-TVKMLASYVRSTPDGT-----------------------
---EKGDFLALDLGGTNFRVLLVKVRNGMRRGVEMQNKIYAIPTEAMEGTGEE-------LFDHIVHCIG
DFLEYMGMKGV-MLPLGFTFSFPCEQRNLDQGILLKWTKGFKASGCEGEDVVDLLREAIRRRE----EFE
LDVVA-----VVNDTVGTMMSCGYDDPHCEVGLIVGTGSNACYMEEM---RNVEMVE-------------
--GD------EG---------------------RMCINMEWGAFGDNGC-LEDMRTPFDEEVDKFSMNPG
KQR-YEKMISGMYLGEIVRNILLDFTKRGLLFRGRISERLKTRGIFETKFLSHIES--------------
------------DRLALLQVRAILQQLGL-ESTCDDSIIVKEVCGVVARRAAMLCGAGMAAVVDKIRENR
NLEHLRITVGVDGTLYKLH-----------PHFSTIMHDTVAKLAPKCKVSF---------VQSEDG---
----------SGKGAALITAVACRIRE--AGQ--H-----------------
> XP_060712556|Hemiscyllium ocellatum|HK2
LYHMRLSDDVLRDISDRFLLGMKKGL-RKDTNPTSS--VKMLPTFVRSTPDGTESGDFLALDLG--GTNF
RVLRVKVSDN---GK-QK--VEIENRICAVPEDLMHGTGTQ---------------LFDHIANCLASVME
EL-----DIK---NEKLPLGFTFSFPCQQTKLD------------------------E------------

-----------SILVHWTKGFKASGVEGRDVVTLLRQAITRR----------G----------------
--------------DFDIDVVSVVNDTVGTMMTCGYDDHNCEIGLIV----------------------
-----G-----TGTNACYMEEMQYI-DMVEGDE--GRMCINMEWGAFGDFGELDDIRTEFDREIDRGSIN
PGKQLFEKLISGMYMGELVRIILVQMAKDGLLFEGKLSPELLIKGHFETRYVSAIEK------------
--DS--DGLQKAQEILAK-LGIEASADDCAATQRVCTVVSSRSSDLCAATLAAVLRRLKENK-GLERLRT
TIGVDGTVYKKH-PQFARRLQKSVRRLAPWCDVRFLPSEDGSGKGAAMVTAVASRLAQQQELREATLRAL
TLDEEQLLQVKQRMRAEMERGLTKDGHATA-PVKMLPTYVRSTPDGT----------------------
---EKGDFLALDLGGTNFRVLLVKLRSGKRRVAEMHNKIYAIPVDAMEGTGEE-------LFDHIVHCIS
EFMEYMGIKQC-VLPLGFTFSFPCEQRGLDQAILLKWMKGFKTTGCEGEDIVDLLREAIKRRE----EFD
LDVVA-----IVNDTVGTMMSCGYDDPFCEVGLIVGTGCNACYMEEM---KNVETVE------------
--GD------EG---------------------RMCINTEWGAFGDNGC-LDDICTEFDREVDRLSMNPG
KQT-YEKLISGMYLGEIVRNILIDFTKRGLLFRGRISERLKTRGIFQTRFLSQIES-------------
------------DRLALLQVRAILQKLGL-QSTCDDSIIVKEVCGVVSRRAAQLCGAGLAAVVDKIRENR
NLERLRVTVGMDGTLYKLH-----------PHFSSIMHETVARLAPRCEVSY---------VQSEDG---
----------SGKGAALVTAVACRIRE--AGQ--H-----------------
>XP_038641198|Scyliorhinus canicula|HK2
LYHMRLSDEALHDISNRFLIEMEQGL-RKDTNPTSS--LKMLPTFVRSTPDGTESGDFLALDLG--GTNF
RVLRVKVSDN---GK-QN--VQIENRICAIPEDIMHGTGTQ---------------LFDHIANCLGDVME
EL-----DIK---DKKLPLGFTFSFPCQQSKLD-----------------------E------------
-----------SLLLHWTKGFKASGVEGRDVVNLLRQAIQRR----------G----------------
--------------DFDIDVVAVVNDTVGTMMSCGYDDHNCEIGLIV----------------------
-----G-----TGTNACYMEDMQHI-DLVEGDE--GQMCINMEWGAFGDLGELHAVRTEFDREIDRGSIN
PGKQLFEKMISGMYMGELVRIILVKMAKDGLLFGGIISPDLLIKGHFETKYVSAIER------------
--DR--DGLQKACEILTR-LGLQPTPDDCAATQRVCTIVSTRSANLCAATLAAVLRRIKENK-GVQRLRT
TIGVDGSVYKKH-PQFARRLHKAVRRLVPTCEVRFLPSEDGSGKGAAMVTAVAYRLAKQQKVREEILATL
MLSDAQLLEVKRRMREEMERGLKKETHCSA-AVKMLATYVRSTPDGT----------------------
---EKGDFLALDLGGTNFRVLLVKVRSGKRRAVEMHNKIYTIPMDAMEGTGEE-------LFDHIVHCIL
DFLEYMGMKDC-VLPLGFTFSFPCEQRNLDQGILLKWTKGFKATGCEGEDVVDLLREAIKRRE----EFE
LEVVA-----IVNDTVGTMMSCGYDDPFCEVGLIVGTGCNACYMEEM---RNVEMVE------------
--GD------EG---------------------RMCVNMEWGAFGDNDC-LDDIRTEFDREVDLLSTNPG
KQR-FEKLISGMYLGEIVRNILINFTKRGLLFRGKISERLKTRGIFQTKFLSHIES-------------
------------DRLALLQVRAILQQLGL-QSSCDDSIIVKEVCGVVSRRAAQLCGAGMAAVVDKIRENR
HLDYLKVTVGVDGTLYKLH-----------PHFSAIMHETVGLLAPKCKVTF---------VQSEDG---
----------SGKGAALITAVACRIRE--AGQ--H-----------------
> XP_059820589|Dasyatis sabina|HK2
LYHMQLSDEQLHEISKRFLTEMKKGL-RKDTNLTSS--LKMLPSFVRSTPDGTESGNFLALDLG--GTNF
RVLRVKVSDN---GK-QN--VEIENRICAIPEDIMHGTGSQ---------------LFDHIANCLGDVME
EL-----NIK---DEKLPLGFTFSFPCKQSKLD-----------------------E------------
-----------SILVNWTKGFKASGVEGRNVVGLLRTAIQKR----------G----------------
--------------DFDIDVVAVVNDTVGTMMSCGYGDHNCEIGLIV----------------------
-----G-----TGTNACYMEEMQHI-DLVEGDE--GRMCVNMEWGAFGDLGELDDIRTDFDKEIDRGSIN
PGKQLFEKMISGMYMGELVRVILVKMAKDGLLFRGKITPELLIKGHFETKYVSAIEK------------
--DS--DGLQKAKEILTK-LGLEPTADDCAATKRVCSIVSTRSSNLCAATLAAVLRRIKENK-DVERLRT
TIGVDGTVYKKH-PQFARRLQKAVRRLVPNCEVRFLPSEDGSGKGAAMVTAVAYRLAKQQKMREEILSAF
NLSDDQLLEVKRRMIVEMERGLKKETHSTA-TVKMLATYVRATPDGT----------------------
---ENGDFLALDLGGTNFRVLLVKVRGGKRRNVEMHNKIYTIPLDAMEGTGEE-------LFDHIVHCIG
DFLEYMGMKGC-RLPLGFTFSFPCEQRNLDQGILLKWTKGFKATGCEGEDIVDLLREAIRRRE----EFD
LDVVA-----IVNDTVGTMMSCGYDDPQCEVGLIVGTGCNACYMEEM---RNVEMVE------------
--GD------EG---------------------RMCVNTEWGAFGDNGC-LDDIRTEFDMEVDKLSLNPG
RQS-YEKMISGMYLGEIVRNILIKFTKEGLLFRGRISERLKTRGIFQTKFLSQIES-------------
------------DRLALLQVRTILQQLGL-QSTCDDSIIVKEVCGVVSRRAAQLCGAGLAAIVDKIRENR
SLDTLKVTVGLDGTLYKLH-----------PHFSTIMHETVERLSPKCQVTF---------LQSEDG---

----------SGKGAALITAVACRIRE--AGQ--H-----------------
>XP_034027133|Thalassophryne amazonica|HK2
LQNLQISDEKLLELSVRFRREMDKGL-CSDTNPTAT--VKMLPTFVRSTPDGTEKGEFLALDLG--GSSF
RVLLVKVMSN---GN-QT--VEMESQIYPMPNELMQACSSELLNGTELMKDKKLVWLFDHIADCLANFLE
KL-----EIK---DKKLPLGFTFSFPCQQTKLD-----------------------E------------
-----------SILLSWTKGFKSSGVEGQDVVQLLRAAIKKR----------G-----------------
--------------DFDIDLVTVINDTVGAMMTCGYEDRNCEVGLIV----------------------
-----G-----TGTNVCYMEELRNM-DLVEGDE--GRMCVNVEWGAFGDDGALEDLRTDVDREMDEGSLH
PGKQLFEKMISGMYMGELVRLILVRMTKEQLLFQGKMTAQLLTTQHIRSSYIYEIEC-------------
--NNDIEGLLNAEKVLKG-LDLDPSLEDCIATRRVCQIVSTRAAHLCAASLVAVLRQIRDNK-AAEKLRA
TIGVDGSVYKNH-PEFGKKLQKMVRRLMPDCDPHFLLSEDGSAKGSAMVTAVASRLAVQHAQRQRVLDTL
HLSREQLLEVKKRMSEGMVLGLSKQTHKQT-SIKMLPTYVRSIPDGT----------------------
---EQGNFLALDFGGSTFRVLLVQLHSGMRRRAEIHSKIYSIPQETMQGTGEE-------LFDHIVDCLA
RFLEYKGMKGA-SLPLGFTFSFPCQQSKLDEGILLNWTKGFKASDCVGKDVVTLLKDAVHRRK----EFE
LNFVA-----VVNDTVGTMMTCGYADPKCEVGLIVGTGTNICYMEEV---ENIDLVE------------
--GY------EG---------------------RMCVNVEWGAFGELGE-LDEFYTQFDLTVDNCSNNPG
KQR-YEKMISGMYLGEIVRNVLLDFTTKGLLYGGKPSERLKTRGIFETKFLSQIET--------------
------------DRLALRQVRAILQHLGLTSSTCDDSVLVKEVCCVVARRAAQLCGAGLAAVVDKIRQNR
HLQQLCITVGVDGTLYKIH-----------PHFSSVMQETLQDLAPQCQVTF---------LRSEDG---
----------SGKGAALITAVACCENG--EQQ--C-----------------
>XP_053725971|Synchiropus splendidus|HK2
LHRFQLSDELLMDISLRFRREMDKGL-CRDTNPTAA--VKMLPTFVLSTPDGTEQGEFLALDLG--GSNF
RVLRVKVISN---GK-Q---VEMENEIFEIPEHIMRGSAFE---------------LFDHVADCLAKFME
KL-----GIK---DQKLPLGFTFSFPCQQNKLD-----------------------E------------
-----------AVLVSWTKRFRTRDVEGKDVVKLLKQAIKKR----------G-----------------
--------------DFEVDIMAVINDTVGAMMTCGYDDPLCEVGLIV----------------------
-----G-----TGTNACYMEHMRNL-EVLDGDE--GRMCVNTEWGAFGDDGVLEDVRNDFDREIDAGSLN
PGKQLFEKMISGMYMGELVRLIMVKMTKEKLLFHGQTSKQLLTTGCFNTKFIYAIES------------
--EKDDEGLARAEEVLKN-LALQPSADDCLATRRICQIVSTRAAHLCAAALVAVLRQIRDNK-AAEKLRT
TIGVDGSVYKNH-PEFPRRLHKMVSRLVPDCDVRFLKSEVGSGKGAAMVTAVAYRLAAQNAERQRILDTL
RLSREQLLEVKERMSYEMMRGLSKQTHEVA-TVKMLPSFVRSLPDGT----------------------
---EQGDFLALDLGGSSFRVLMVQVRSGNKRNVDMRQKVYGIPPELMQGSAEE-------LFTYIVECMA
NFLENMGMTGA-SLPLGFTFSFPCQQSKLDEGILLNWTKCFAASGCVGYDVVKLLKDAVRRRG----EFH
LNFVA-----VVNDTVGTMMTCAYEDPKCEVGLIVGTGTNACYMEEM---ENIEVLE------------
--GN------EG---------------------RMCVNIEWGAFGDNGE-LDDFFTCFDRLVDESSNNPG
KQR-YEKMIGGLYLGEIVRNVLLDFTAKGLLFRGKVSERLKTFGIFETKFLARIES--------------
------------DRLAMRQIRSILQHLGLTFSTCDDSVLVKEVCAVVSRRAAQLCGAGLAAVVDRIRQNR
NLQRLEVTVGVDGTLYKTH-----------PHFSRVMQEVLQDLAPQCSVTF---------HKSEDG---
----------SGKGAALITAVACRVNN--EGQ--Y-----------------
>XP_061907577|Entelurus aequoreus|HK2
LHQLHLSDTTVMDISMRFRREMDKGL-CRDTNPTAA--VKMLPTYVLSTPDGSEQGDFLALDLG--GSTF
RVLLVTVMAN---GE-QK--VDMENQIYAIPEYLMRGSGSE---------------LFDHIAECLANFME
KL-----GIK---DKRLPLGFTFSFPCQQSKLD-----------------------E------------
-----------SVLISWTKAFKASGVEGNNVVSLLQRSIKKR----------G-----------------
--------------DINVDIMAVINDTVGAMMTCGYDDHRCEIGLIV----------------------
-----G-----TGTNACYMEQMRNI-DVLDGDE--GQMCINTEWGAFGDDGALEDLRTDFDREIDAGSLN
PGKQLFEKMISGMYMGELVRLILVKMAKDRLLFNGKTTSQLNTTGKFTTKCIYSIES------------
--DNIEEGLKSAQSVLQA-LGLDPTLEDCIATLRVCQIVSIRAAHLCAATLVAVLRQIRDNK-AAEKLRT
TIGVDGSVYKNH-PEFPRRLHKMVGRLVPDCDVRFLRSEVGSGKGAAMVTAVALRLAAQNADRQRILNTL
RLNREQLLRVKTLVSEGMVRGLSKPEHDQA-SIRMLPTFVGSIPDGT----------------------
---EHGDFLVLDLGGSSFRVLLVHVRSGKRHNVDTHQKIYNIPQETMQGTGEE-------LFTHIVDCMA
NFLENMGMSGA-SLPLGFTFSFPCHQSKLDQGILLKWTKGFKASGCEGQDVVKLLKDAVRRQG----VFN

```
LTFVA-----VVNDTVGTMMTCGYEDPKCEVGLIVGTGTNVCYMEEM---QNIEVVD-------------
--GD------DG---------------------QMCVNMEWGAFGENGE-LDDFITEFDRVVDASSNHPG
KQR-YEKMISGMYLGEVVRHVLMEFTAKGLLFRGKLSEQLKTRGIFATRFLSQIES--------------
------------DRLAMRQVRCILQHLGLVTSTYDDSVLVKEVCSVVSRRAAQLCGAGLAAVVDKIRENR
NMQQLTITVGVDGTLYKMH-----------PHFSRIMQETLQDLAPQCNVTF---------HKSEDG---
----------SGKGAALITAVACKSNS--SGQ--NYSG--------------
>XP_061528924|Phycodurus eques|HK2
LHYLQLPDETLMDISMRFRREMDKGL-CRDTNPTAV--VKMLPTYVRSTPDGTEQGEFLALDLG--GSTF
RVLLVTVMAN---GE-PK--VDMENQIYTIPEHLKRGCGSE---------------LFDHIAECLADFLE
KL-----GIK---DKKLPLGFTFSFPCQQSKLD------------------------E------------
-----------SILISWTKAFRANGVEGKDVVNLLQRSVNKR----------G-----------------
--------------DISVDIMAVINDTVGAMMTCGYDDRNCEIGLIV-----------------------
-----G-----TGTNACYMEQMRNL-ESLDGDE--GQMCVNTEWGAFGDDGTLEDLRTDFDREIDAGSLN
PGKQLFEKMIGGMYIGELVRLILVRMTRDQLLFQGKTTSQLLMTGKFPTNYIYSIDS-------------
--DNREEGLMSAQNVLRS-LGLDPTLEDCIATQRVCQIVSTRAAHLCAATLVAVLRQIRDNK-ATEKLRS
TIGVDGSVYKNH-QEFSRRLHKMVRRLVPDCDVRFLRSEVGSGKGAAMVTAVALRLAAQNAERQHILDTL
RLNREQLMEVKRLISEDMVRGLSRQAHNQA-SIKMLPTFVGSTPDGS-----------------------
---EHGDFLALDLGGSSFRVLLVRVRSG-KTHVDMHHKNYSIPQETMQGTGEE-------LFSHIVDCIA
KFLENMGMRGA-SLPLGFTFSFPCHQSKLDQGILLKWTKGFKASGCEGHDVVTLLKDAVRCRG----EFD
LSFVA-----VVNDTVGTMMTCGYEDPKCEVGLIVGTGTNACYMEEM---QNIEMVD-------------
--GD------DG---------------------RMCVNMEWGAFGENGE-LDDFLTNFDRVVDASSSNPG
KQR-YEKLISGMYLGEIVRHVLMEFTAKGLLFRGKLSERLKTWGIFETKFLSQIES--------------
------------DRLAMRQIRFILQHLGLTSSTCDDSVLVKEVCSVVSRRAAQLCGAGLAAVVDKIRQNR
NMQQLSVTVGVDGTLYKMH-----------PHFAGVVQETLQDLAPQCKVTF---------HKSEDG---
----------SGKGAALITAVACRIRS--NTL--ANSPN-------------
>KAK0134095|Merluccius polli|HK2
LYHLRLSDENLMDVSLRFRREMDKGL-GRDTNPTAA--LKMLPTFVRSTPDGTEKGDFLALDLG--GTHF
RVLLVKVSSN---GK-QK--VEMENQIYAIPENIMKGRASE---------------LFDHIADCLANFLD
KL-----GIK---EKKLPLGFTFSFPCQQTKLD------------------------E------------
-----------SILVSWTKGFRVSGVEGKDVVALLRRAIQKR----------G-----------------
--------------DFDIDIVAVINDTVGTMMTCGYDDHHCEIGLII-----------------------
-----G-----TGTNACYMEEMRNM-ELVEGDE--GRMCVNMEWGAFGDDGVLDDLRTAFDREIDAGSLN
PGKQLFEKMISGFYMGELVRLILVKMAKDQMIFQGQVTANLLTTGHFKTSQVSAIET-------------
--DRDNGGLLAAKTVLQG-LGLTPSTEDCVATQRVCQVVSTRAAHLCAATLVSVLRQIRDNK-AAEKLRT
TIGVDGSIYKCH-PQFARRLHKMVRRLAPDCDVRFLQSEDGSGKGAAMVTAVAYRLAMQHAERQCILNTL
RLSREQLLEVKRRLEKAMEQGLAKETHPQS-SVKMLPTYVRATPDGT-----------------------
---EHGDFLALDLGGTNFRVLLVRVRSGKRRSVEMHNKIYSIPQDTMQGTGEE-------LFDHIVHCIA
DFLEYMGMKGA-FLPLGFTFSFPCQQRQLDQGILLKWTKGFKATGCEGEDVVTLLKDAIHRRE----EFD
LDVVA-----VVNDTVGTMMTCGYEDPLCEVGLIVGTGTNACYMEEM---TNVELVE-------------
--GD------EG---------------------RMCVNMEWGAFGDGGE-LDDLCAEFDRIVDENSNNQG
KQR-YEKMISGMYLGEIVRNVLLDFTAKGLLFRGKLSERLKTRGIFETKFLSQIES--------------
------------DRLALRQVRSILRHLGLTSSTCDDSILVKEVCGVVGRRAAQLCGAGLAAVVDKVRRNR
GLEHLRITVGVDGTLYKLH-----------PHFSGIMRETVQDLAPCCEVSF---------LQSEDG---
----------SGKGAALITAVACRMRD--AAL--H-----------------
>XP_038868897|Salvelinus namaycush|HK2
LYHLRLSDENLIDVSERFRREMDKGL-GRDSNPTAA--VKMLPTFVRSTPDGTEKGDFLALDLG--GTNF
RVLLVKVSDN---GK-QK--VEMENQIYAIPEELMRGSGEE---------------LFDHIAECLANFLE
KL-----GIK---NQKLPLGFTFSFPCQQTKLD------------------------E------------
-----------SILVSWTKGFKSHGVEGRDVVSFLRKAIIKR----------G-----------------
--------------DFDIDIVSVINDTVGTMMTCGYDDHHCEIGLIV-----------------------
-----G-----TGTNACYMEEMRHL-ELVEGDE--GRMCVNMEWGAFGDDGTLDDLRTDFDQEIDAGSLN
PGKQLFEKMISGMYMGELVRLILVKMAKEDMVFQGHTTPDLVTNGQLQTSFVSAIEN-------------
```

--DKDKEGLVSAEKMLRG-LGLDPSVEDCVATRRVCQVVSTRAAHLCAATLAAVLRQIRDNK-AAERLRT
TIGVDGSVYKNH-PQFARRLHKMVRRLVPDCNVRFLRSEDGSGKGAAMVTAVAFRLAIQHAGRQRILDAL
RLSQEQLLDVKRRMGEEMNRGLAKESHDQA-TVKMLPTFVRSIPDGT-----------------------
---ESGEFLALDLGGTNFRVLLVRVRCGKRRSVEMHNKIYSIPQEAMQGTGEE-------LFDHIVHCIA
DFLEYMGMKGA-SLPLGFTFSFPCHQSKLDQGILLKWTKGFKATGCEGEDVVTLLKDAIYHRE----EFD
LDVVA-----VVNDTVGTMMTCGYEDPLCEVGLIVGTGTNVCYMEEM---QNMELVD-------------
--GS------EG---------------------KMCVNMEWGAFGDHGE-LDNFSTDFDKAVDEHSANPG
KQT-YEKMISGMYLGEIVRNVLLEFTTKGLLFRGKLSERLKTRGIFETKFLSQIES--------------
------------DRLALRQVRSILQHLGLTSSTCDDSILVKEVCSVVACRAAQLCGAGLAAVVDKIRQNR
NLPELKITVGVDGTLYKLH-----------PRFSNFMHETVRDLAPQCKVTF---------IQSEDG---
----------SGKGAALITAVACRIRD--AGQ--R-----------------
>XP_063051015|Engraulis encrasicolus|HK2
LSPLRLSDQSIMDVSLRFRREMDRGL-CRDTNPTAA--IKMLPTFVRSTPDGTEQGDFLALDLG--GTNF
RVLLVKVSAN---GK-QK--VEMENQIYAIPEDLMRGDGSE---------------LFDHIAECLANFLS
KL-----GIK---DKKLPLGFTFSFPCQQTKLD------------------------E------------
-----------SILVSWTKGFRARGVEGNDVVGMLRAAIKKR----------G-----------------
--------------DFDIDIVAVINDTVGTMMTCGYDDHHCEIGLIV-----------------------
-----G-----TGTNACYMEEMRNL-QTVEGDE--GRMCVNLEWGAFGDDGALDDLRTSYDQEIDAGSLN
PGKQLFEKMISGMYMGELVRLILVKMAKEELLFQGTTTPQLLNSGQFQTTYVSAIEN-------------
--EKDNEGLLRAEEVLLG-LGLQPSAEDCMATQRVCQIVSTRAAHLCAATLSAVLRQIRDNK-AADRLRT
TVGVDGSVYKNH-PQFSRHLHKMVRALVPDCDVRFLRSEDGSGKGAAMVTAVAERLANQHAARQAVLDTL
RLSRAQLAEVKHRITEGMELGLARSTHANA-TIKMLPTYVRSTPNGT-----------------------
---EQGDFLALDLGGTNFRVLLVKVRSGSKKRVEMHNKIYAIPQETMQGTGEQ-------LFDHIVQCIA
DFLEYKGMKGA-TLPLGFTFSFPCQQNRLDQGILLKWTKGFRASGCEGRDVVTLLKDAVHRHE----GFD
LDVVA-----VVNDTVGTMMTCGYEDPHCEVGLIVGTGTNACYMEEL---ENVELVE-------------
--GQ------GG---------------------RMCVNMEWGAFGEHGE-LDDFCTTFDHTVDQSSAHPG
KQR-YEKMISGMYLGEIVRNVLLEFTAKGLLFRGRLSERLKTRGIFETKFLSQIES--------------
------------DRLALRQVRSILQHLGLTSSTCDDSILVKEVCAVVSRRAAQLCGAGLAAVVDKIRRNR
HLEQLRITVGVDGTLYKLH-----------PHFSTIMRETLRELAPQCEVKL---------LQSEDG---
----------SGKGAALITAVACRLRD--GEQ--------------------
>XP_041956986|Alosa sapidissima|HK2
LYHLRLSEENLMDVSLRFRREMDRGL-GRDTSPTAA--VKMLPTFVRSTPDGTEQGDFLALDLG--GTNF
RVLLVRVSSN---SK-KQ-EVEMENQIYAIPEDLMRGSGSE---------------LFDHIVECLAIFLE
KL-----GIK---DKKLPLGFTFSFPCQQTKLD------------------------E------------
-----------SILVSWTKGFKASGVEGKDVVSMLRAAIKKR----------G-----------------
--------------DFDIDIVAVINDTVGTMMTCGYDDHHCEIGLIV-----------------------
-----G-----TGTNACYMEEMRNL-ETVEGDE--GRMCVNMEWGAFGDDGALDDLRTPFDVEIDAGSLN
PGKQLFEKMISGLYMGELVRLILVKMAKEELLFQGKTTPELLTTGHFQTSFISAIEN-------------
--EKNNQGLLNAEQILQG-LGLTPSAEDCVATQRVCQIVSTRASQLCAATLAAVLRQIRDNK-AAERLRT
TVGVDGSVYKNH-PQFARRLHKMVRRLVPDCDVRFLRSEDGSGKGAAMVTAVAYRLANQHAERQRILDKL
RLSHDQLLEVKRRMAEAMERGLSRDTHGNA-TVKMLPTYVRSTPNGT-----------------------
---ERGDFLALDLGGTNFRVLLVRVRSGKKRSVEMHNKIYAIPQDLMQGTGDE-------LFDHIVHCIA
DFMEYMGMKGA-SLPLGFTFSFPCQQNRLDEGILLKWTKGFRATGCEGQDVVTLLKDAIHRHE----EFD
LDVVA-----VVNDTVGTMMTCGYEDPYCEVGLIVGTGTNACYMEEM---ENVELVE-------------
--GN------EG---------------------RMCINMEWGAFGDHGE-LNDFCTTFDHAVDERSANPG
KQQ-YEKMISGMYLGEVVRNVLLDFTSKGLLFRGRLSERLKTRGIFETKFLSQIES--------------
------------DRLALRQVRAILQHLGLTSSTCDDSILVKEVCTVVARRAAQLCAAGLAAVVDKIRKNR
QLEHLRVTVGVDGTLYKLH-----------PHFSTILRQTLRELAPQCEVSI---------MQSEDG---
----------SGKGAALITAVACRMRN--EAK--------------------
>XP_051957323|Xyrauchen texanus|HK2
LYHMRLSEENLMDISKRFRTEMDKGL-GRDTNLTAA--VKMLPTFVRSTPDGTETGNFLALDLG--GTNF
RVLLVKVCSN---GK-QTVEVEMENQIYAIPENIMRGSGTE---------------LFDHIAECLANFLE

KL-----GIK---EKKLPLGFTFSFPCLQTKLD------------------------E------------
-----------SFLVSWTKGFKASGVEGRDVVDLLRKAIKKR----------G-----------------
--------------DFDIDIVAVINDTVGTMMTCGYDDHHCEIGLIV-----------------------
-----G-----TGTNACYMEEMRHL-ELVDGDE--GRMCVNMEWGAFGDDGALDDFRTDFDREIDAGSLN
PGKQLFEKMISGMYMGELVRLILVKMAKDGLLFQGHTTPDLLTTGHFQTAFVSAIEN-------------
--EKNNEGMVSAEQVLRG-LGLNPSSEDCEATQRVCQIVSTRAAHLCAASLAAVLRQIRDNK-AERLRT
TIGVDGSVYKNH-PLFARRMQKMVRWLVPDCDVRFLRSEDGSGKGAAMVTAVAYRLAKQHVERQRILDTL
RLSRDQLLEVKKRMTEEMNRGLAKQTHSTA-SVKMLPTFVRSTPDGT-----------------------
---ERGDFLALDLGGTNFRVLLVRVRSGKRRSAEMHNKIYTIPEDITQGTGEE-------LFDHIVHCIA
DFLEYMGMKGA-SLPLGFTFSFPCHQTRLDQGVLIKWTKGFKASGCEGEDVATLLKDAIHRSE----EFD
LEVVA-----VVNDTVGTMMTCGYEDPQCEVGLIVGTGTNACYMEDM---SNVELVD-------------
--GD------EG---------------------RMCVNMEWGAFGDLGE-LDDICTEFDRAVDDQSANSG
KQR-YEKMISGMYLGEVVRNVLLDFTDKGLLFRGKLSERLKTRGIFETKFLSQIES--------------
------------DRLALRQVRSILQHLGLTSSTCDDSILVKEVCSVVSKRAAQLCGAGLAAVVDKIRLNR
GLDQLKITVGVDGTLYKLH-----------PHFSTIMRETLKDLAPNCEVTV---------IQSEDG---
----------SGKGAALITAVACRLRG--GGK--------------------
>XP_028850948|Denticeps clupeoides|HK2
LYHLRLPEETLMDVSKRFRQEMDRGL-GRDTNPTAA--VKMLPTFVRSTPDGTESGDFLALDLG--GTNF
RVLLVKVSAN---GK-QR--VEMENQIYAIPESLMRGSGTE---------------LFDHIAECLANFLE
KL-----GIK---DKKLPLGFTFSFPCQQTKLD------------------------E------------
-----------SILVSWTKGFKSSGVEGQDVVSLLRNAIRRRRREWEVRYQMQ-----------------
--------------DFDIDIVAVINDTVGTMMTCGYDDHHCEIGLIV-----------------------
-----G-----TGTNACYMEEMRHL-ELVEGDE--GRMCVNMEWGAFGDDGALDDLRTDFDREIDAGSLN
PGKQLFEKMISGMYMGELVRLILVKMAREGIVFQGHTTPDLLATGHFQTSFVSAIEN-------------
--EKDNEGLLSAERILKD-LGLNPSPEDCVTTQRVCQIVSTRAAHLCAASLAAVLRQIRDNK-AERLRT
TIGVDGSVYKNH-PQFARRLHKMVRRLVPDCDVRFLRSEDGSGKGAAMVTAVAYRLAKQQAERQSILDTL
RLGREQLVAVKRRLLEEMTRGLARETHSSA-AVKMLPTYVRSTPDGT-----------------------
---ERGDFLALDLGGTNFRVLLVRVRQGRRRGVEMHNKIYAIPQDLMQGTGEE-------LFDHIVHCIA
DFLEYMGMRGV-SLPLGFTFSFPCHQTRLNQGILLKWTKGFRASGCEGEDVVTLLKDAIHRHE----EFD
LDVVA-----VVNDTVGTMMTCGYDDPRCEVGLIVGTGTNACYMEEM---RNVELVE-------------
--GN------AG---------------------LMCINMEWGAFGDQGE-LDEFCTEFDRLVDERSSNPG
KQR-YEKMISGMYLGEIVRNVLLDFTAKGLLFRGKLSERLKTRGIFETKFLSQIES--------------
------------DRLALRQVRTILQHLGLTSSTCEDSILVKEVCSVVSRRAAQLCGAGLAAVVDKIRQNR
NLNHLQVTVGVDGTLYKLH-----------PHFSTVLRETLRDLAPQCEVTL---------VQSEDG---
----------SGKGAALITAVACRLRE--AGL--------------------
>XP_062874325|Trichomycterus rosablanca|HK2
LYHLRLSDENLMDISKRFRREMDKGL-GKDTNLTAS--VKMLPTYVRSTPDGTEKGHFLALDLG--GTNF
RVLLVKVSSN---GK-QT--VEMENQIYAIPETLMRGSGTE---------------LFDHIAECLANFLD
KM-----GIK---EKKLPLGFTFSFPCQQNKLD------------------------E------------
-----------SILVSWTKGFKASGVEGRDVVSLLRKAIKKR----------G-----------------
--------------DFDIDIVAVINDTVGTMMTCGFDDHHCEIGLII-----------------------
-----G-----TGTNACYMEEMRNL-ELVEGDE--GRMCVNMEWGAFGDDGALDDIRTDFDREIDAGSLN
PGKQLFEKMISGMYMGELVRIILVKMTKQGLLFQGHTTPDLLTTGHFKTSFVSAIEN-------------
--DKSKEGLVCVEEILRS-LGLEPSNEDCVAVHHVCKVVSTRAAHLCAAALVAVLRQIRDNK-AERLRT
TIGVDGSVYKNH-PQFARRLHKMVRRLVPDCDVRFMRSEDGSGKGAAMVTAVAYRLAKQHAERQQVLDSL
RLDREQLMVVKRRMEEQMRRGLAKDTHSSA-SVKMLPSYVRATPDGT-----------------------
---ERGDFLALDLGGTNFRVLLVRLRSGKRRSADMQNKIYAIPQEVMQGTGEE-------LFDHIVHCIS
DFMEYMGIKGA-SLPLGFTFSFPCHQTALDQGVLMKWTKGFKATGCEGHDVAVLLKDAIHRME----EFD
LDVVA-----VVNDTVGTMMSCAFEDPQCEVGLIVGTGTNACYMEEM---SNVELVD-------------
--GD------EG---------------------QMCVNMEWGAFGDDGE-LDDICTPFDRTVDENSTNPG
KQR-FEKMISGMYLGEIVRNVLLDFTSKGLLFRGKLSERLKTHGIFETKFLSQIES--------------
------------DRLALRQVRSILQHLGLTSSTCDDSILVKEVCTVVAQRAAQLCGAALAAVVDKIRQNR

```
NLNTLSVTVGVDGTLYKLH-----------PHFSSVMRDTLQDLAPQCDVSL---------VQSEDG---
----------SGKGAALITAVACRING--AKK--------------------
>KAI5088099|Silurus meridionalis|HK2
LFHLRLSPENLMDVSQRFRREMDKGL-GRDTNLTAS--VKMLPTFVRSTPDGTEKGDFLALDLG--GTNF
RVLRVKVSSN---GK-QT--VEMENQIYAIPETLMRGSGTE---------------LFDHIAECLANFLE
KL-----GIK---EKKLPLGFTFSFPCQQTKLD------------------------E------------
-----------SILVSWTKGFKASGVEGRDVVGLLRKAIKKR----------G-----------------
--------------DFDIDIVAVINDTVGTMMTCGFDDHHCEIGLIV-----------------------
-----GESSFRTGTNACYMEEMRNM-ELVEGDE--GRMCVNMEWGAFGDDGSLDDIRTAFDREIDAGSLN
PGKQLFEKMISGMYMGELVRIILVKMTKEGLLFQGRTTPNLLVNGCFKTSYVSAIEN-------------
--DKSKEGLVMVEQVLRD-LGFEPSQEDCVAVQRVCQIVSTRAAHLCAASLAAVLRQIRDNK-AVDRLRT
TIGVDGSVYKNH-PQFARRLHKMVRCLVPDCDVRFLRSEDGSGKGAAMVTAVAYRLAKQHAERQKVLDTL
RLSREQLMEVKRRMEVQMKRGLAQNTHDSA-SVKMLPTYVRATPDGT-----------------------
---EHGDFLALDLGGTNFRVLLVRVRSG-RRSVEMHNKIYTISQDIMQGTGEE-------LFDHIVHCIA
DFLEYMGMKGA-SLPLGFTFSFPCHQAGLNKGILLRWTKGFKASGCEGEDVAMLLKDAIHRME----EFD
LDVVA-----VVNDTVGTMMSCGYEDPQCEVGLIVGTGTNACYMEEM---RNVELVE-------------
--GD------EG---------------------RMCVNMEWGAFGDDSE-LDNVSTPFDLAVDENSTNPG
KQR-FEKMISGMYLGEIVRNVLLDFTAKGLLFRGKLSERLKTRGIFETKFLSQIES--------------
------------DRLALRQVRSILQHLGLTSSTCDDSILVKEVCSVVSQRAAQLCAAALAAVVDKIRQNR
NLNSLSVTVGVDGTLYKLH-----------PHFSCILQETLRDLAPQCDVTL---------LQSEDG---
----------SGKGAALITAVACRIRD-------V-----------------
>KAG9262061|Astyanax mexicanus|HK2
LYHMRLSEENLMDVSRRFRREMDKGL-GRDTNLTAS--VKMLPTFVRSTPDGTESGDFLALDLG--GTNF
RVLLVKVSAN---GK-QK--VEMENQIYAIPETLMRGSGCE---------------LFDHIAECLANFLD
KL-----GIK---DKKLPLGFTFSFPCRQTKLD------------------------E------------
-----------SFLVSWTKGFKSSGVEGKDVVGLLRKAIKKR----------G-----------------
--------------DFDIDIVAVINDTVGTMMTCGYDDHHCELGLIV-----------------------
-----G-----TGTNACYMEEMRNL-ELVEGDE--GRMCVNMEWGAFGDDGALDDLRTEFDREIDAGSLN
PGKQLFEKMISGMYMGELVRIILVKMAREGLVFQGHTTPDLLITGHFKTSYVSAIEN-------------
--QKEKEGLLYAEQVLRG-LGLDPSAEDCVAVQRVCHTVSTRAAHLCAATLAAVLRQIRDNK-AAERLRT
TIGVDGSVYKNH-PQFARRLHKMVRRLVPDCDVRFLRSEDGSGKGAAMVTAVAYRLAEQHAERQRVLDTL
RLSREQLMEVKRRMEEGMKRGLARDTHDSA-SVKMLPTYVRSTPDGT-----------------------
---ERGDFLALDLGGTNFRVLLVRVRSGKRRSVEMHNKIYTISQDIMQGTGEE-------LFDHIVHCIG
DFLEYMGLKGA-SLPLGFTFSFPCHQTRLDQGILLKWTKGFKATGCEGEDVAALLKDAIHRME----EFD
LDVVA-----VVNDTVGTMMTCGYEDQRCEVGLIVGTGTNACYMEEM---ENVELVE-------------
--GD------DG---------------------LMCVNMEWGAFGDRGE-LDEFCTQFDAAVDERSANPG
KQR-FEKMISGMYLGEIVRNVLLDFTTRGLLFRGKLSERLKTRGIFETKFLSQIES--------------
------------DRLALRQVRSILQHLGLVSSTCEDSILVKEVCSVVSRRAAQLCGAGLAAVVDKIRQNR
NLEQLTVTVGVDGTLYKLH-----------PHFSTIMRETLKDLAPQCDVSL---------VQSEDG---
----------SGKGAALITAVACRLRD--AKK--------------------
>XP_056889203|Takifugu flavidus|HK2
LQRFQLSYGDLMEISNRFRREMDKGL-CRDTNPTAV--VKMLPTFVHSTPDRTEHGEFLALDLG--GSTF
RVLLVKVMAD---GE-QK--VEMENQIYVIPEHLMKGNGVK---------------LFDHIADCLANFME
KI-----GMK---EKKLPLGFTFSFPCQQTKLD------------------------E------------
-----------SILLSWTKAFKSSGVEGKDVVSLLRAAIKKR----------G-----------------
--------------DFDVDVMAVINDTVGAMMTCGYDDQLCEIGLIV-----------------------
-----G-----TGTNACYMEQMRNI-EVLDGDE--GRMCINTEWGAFGDDGTLEDHRTDFDREIDAGSLN
PGNQLFEKMVSGMYMGELVRVILVKMTKEQLLFKGKTTVGLLTTGSFKTSYIYTIDN-------------
--DNDEEGLASAEKVLRG-LNLNPTSEDCLAVRRVCQIVSTRSAHLCAATLVAVLRQIRDNK-AAEKLRT
TIGADGSVYKNH-QEFSRRLHKMVRQLMPDSDVRFLTSQCGSGKGAAMVTAVAYRYAAQQAERQAVLNTL
RLSRQQLLEVKRRMREKMLQGLSKQTHEES-SVKMLPTYVISTPNGT-----------------------
---EQGDFLVVDLGGSKFRVLLAQIQNGKKRNETLHQKIYNIPQETMQGTGKE-------LFDHIVDCIA
```

DFLEYMGMSGA-SLPLGFTFSFPCHQSRLDQGILLKWTKGFKASGCEGQDVIVLLKEAVCRRK----EFD
LNFVA-----VVNDTVGTMMTCGYEDPRCEVGLIVGTGTNACYMEEV---QNIDLVD-------------
--GD------EG---------------------RMCVNMEWGAFGDAGE-LDDFCTRFDAFVDESSTNPG
KQR-YEKMVSGMYLGEIVRNILMHFTDRGLLFRGKVSEKLKMHGIFATKFLSQIES--------------
------------DRLAMRQVRAILQHLGLTNSTCDDSVLVKEVCGVVSRRAAQLCGAGLAAVVDKMRENR
NLNQLSVTVGVDGTLYKTH-----------PHFSRIMQETLQDLAPQCQVTF---------HKSEDG---
----------SGKGAALITAVACRLRR--EEE--Q-----------------
>XP_056150125|Lampris incognitus|HK2
LCRLRLSDENLMDVSVRFRREMDKGL-CRDTNPTAA--VKMLPTFVRSTPDGTEEGDFLALDLG--GTNF
RVLLVRVSAN---GK-QK--VEMENQIYAIPEHLMRGSGSE---------------LFDHIADCLANFLD
KL-----GMK---DKKLPLGFTFSFPCQQAKLD------------------------E------------
-----------SILVSWTKGFKVSEVEGKDVVSLLRKAIKKR----------G-----------------
----------LCLQDFDIDIVAVINDTVGTMMTCGYDDHHCEIGLIV-----------------------
-----G-----TGTNACYMEEMRNL-DLIEGDE--GRMCVNMEWGAFGDDNSLEDLRTEFDREIDAASLN
PGKQLFEKMISGLYMGELVRQILVKMTKEQLLFHGRGTPDLLTTGCFHTSYIFSIEN-------------
--EK--EGRMNAEKVLKG-LGLDPSPEDCIATQRVCEIVSTRAAHLCAATLVAVLRQIRDNK-AAERLRT
TIGVDGSIYKNH-PQFARKLHKMVRRLVPDCDVRFLLSEDGSGKGAAMVTAVAYRLATQHAQRQQILDTL
RLNREQLLEVKRRMAEEMVRGLTKQSHEQA-SVKMLPTYVRSTPDGT-----------------------
---EQGDFLALDLGGSNFRVLLVRVQSGSRRRIEMHNKIYNIPQEFMQGTGEE-------LFDHVVCCMA
DFLEYMSMKGA-TLPLGFTFSFPCHQSKLDQGILLKWTKGFKASGCEGEDVVRLLKDAVHRRQ----EFD
LDVVA-----VVNDTVGTMMTCGYKDPLCEVGLIVGTGTNACYMEEM---QNVDLVE-------------
--GD------LG---------------------RMCVNMEWGGFGEHGE-LDDFCTEFDYTVDDQSSTPG
KQR-YEKMISGMYLGEIVRNVLLEFTSKGLLFRGKLSERLKTRGIFETKFLTQIES--------------
------------DRLALRQVRSILQHLGLTSSTCEDGILVKEVCSVVARRAAQLCGAGLAAVVDKIRQNR
NLEQLCITVGVDGTLYKMH-----------PHFSSIMRETVQDLAPQCEVTF---------LQSEDG---
----------SGKGAALITAVACRIRE--SEQ--H-----------------
>XP_028312572|Gouania willdenowi|HK2
MHQFQLSDQTIMDISLRFRREMDKGL-CRDTNPTAV--VKMLPTYVHSTPDGTEQGEFLALDLG--GSNF
RVLLVRVMGN---GE-QK--VEMENQIYDIPEHIMRGSGSK---------------LFDHVADCLANFLE
KK-----GMK---NKKLPLGFTFSFPCQQTKLD------------------------E------------
-----------SFLLSWTKGFKSSGVEGKDVVSLLRKSIEKR----------G-----------------
--------------DFEIDIVAVINDTVGTMMTCGFDDRFCEIGLIV-----------------------
-----G-----TGTNACYMEHMRNI-ELLDGDE--GRMCINTEWGAFGDDGALEDIRTEIDRELDAGSLN
SGNQLFEKMVSGMYQGELVRLILVKMAREHLMFKGKTTPGLLTTGHFLTSYIYDIDT-------------
--EKEEEGLASTEKVLRG-LDLDPTAEDCKATRRVCHIISKRAAHLCAATLVALLRQIRDNK-AAEKLRT
TIGVDGSVYKNH-NAFSRRLNKMVRRLVPDCDVRFLLSQHGSSKGAAMVTAVAFRFAAQQAERQRILDTL
RLSRQQLLEVKRRMSEQMLRGLSKQTHEQS-SLKMLPTYVRSTPDGT-----------------------
---EHGDFLALDLGGSSFRVLLVRIESGERNNVDMHHKIYSIPQEFMQGTGDE-------LFSHIVTCIA
DFLEYMGMRGA-SLPLGFTFSFPCHQSKLDEGILLQWTKGFKASGCEGQDVVTLLKEATRCKG----EFD
LSFVA-----VVNDTVGTMMTCGYEDCQCEVGLIVGTGTNICYMEEM---QNVELVE-------------
--GD------DG---------------------RMCVNMEWGAFGENGE-LDDFCTKFDQEVDEFSNYPG
KQR-YEKMISGMYLGEIVRNVLIEFTAKGLLFRGKDPERLRTRGIFETKFLSQIES--------------
------------DRLAMRQIRSILQHLGLTGSTCDDSVLVKEVCAVVSRRAAQLCGAGLAAVVEKIRQSR
NLNQLSVTVGVDGTLYKRH-----------PHFSSIMQETLQDLAPQCQVTF---------HKSEDG---
----------SGKGAALITAVACRMDR--CGQ--------------------
>KAF6726383|Oryzias melastigma|HK2
LQQFMVSDQAVMDLSLRFRREMDKGL-CRDTNPTAA--VKMLPTFVRSTPDGTEQGEFLSLDLG--GSNF
RVLLVKVKAD---GE-QK--VEMENKIYAIPEHVMRGCGSG---------------LFDHVVDCLANFME
MK-----GIK---DKKLPLGFTFSFPCLQTKLD------------------------E------------
-----------SVLLSWTKGFKLSGVEGKDVVSLLRTAIKKR----------G-----------------
--------------DFDIDVVAVVNDTVGTMMTCGYEDQYCEIGLIV-----------------------
-----G-----TGTNACYMEQMRNL-DLLDGDE--GRMCVNTEWGAFGDDGALEDLRTNIDQEIDSGSLN

PGKQLFEKMVSGMYQGELVRLILVKMARQRQLFQGRTTPELLTTGRFSTSYIFTIDS------------
--DKEAESLSSTEKVLRD-LGLEPSLEDCICTRRVCQIVSTRAARLCAATLVAILRQIRDNK-AAEKLRT
TIGVDGSVYKNH-PEFSRRLNKMVRRLVPDCDVRFLQSQDGSGKGAAMVTAVAYRLAAQHAERQRILDTL
RLSREQLLEVKRRMTEEMMRGLAKQTHEQT-SVKMLPAFVRHTPDGT-----------------------
---EHGDFLTLDLGGSSFRVLLVRLQSGGHPKVDMHQKIYSVPQEIMQGTGEE-------LFDHIVQCIA
DFLEYMGMRGA-SLPLAFAFSFPCHQNKLDEGILLKWTKGFKASGCEGRDVVTLLKEAVRRKR----EFG
LSFVA-----VVNDTVGTMMTCGYEDPKCEVGLIVGTGTNACYMEEM---HNIEVVE------------
--GN------DG---------------------QMCVNVEWGAFGDNGE-LDDFRTRFDRAVDEGSNYPG
KQS-YEKMISGMYLGEIVRNVLLDFTERGLLFRGKMSERLKTRGIFATKFLAQIES--------------
------------DRLAMRQVRSILQHLGLTGSTCDDSVLVKEVCSVVARRAAQLCGAGLAAVVDKMRQNR
NLNQLSVTVGVDGTLYKTH-----------PHFSSIMQETLQDLAPQCQVTF---------LKSEDG---
----------SGKGAALITAVACRNNS-------------------------

>XP_017270188|Kryptolebias marmoratus|HK2
LEKLKLSDEAVMDLSVRFRREMDKGL-CRDTNPTAT--VKMLPTFVRSTPDGTEQGEFLALDLG--GTFF
RILLVKVKGN---GE-QK--VEMADQIYDIPELVMRGTGTE---------------LFDHVADCLANFLE
KM-----GIK---HKKLPLGFTFSFPCQQTKLD-----------------------E------------
-----------SVLIYWTKGFRATGVEGKEVVSLLRKSIEKR----------G-----------------
--------------DFDVDILAVINDTVGTMMTCGYDDHLCEIGLIV----------------------
-----G-----TGTNACYMEQMRNL-ELMDGDE--GRMCVNTEWGAFGDDGALEYLRTDIDREIDAGSLN
PGKQLFEKMVSGMYLGELVRLILVKMARDQLLFQGKTTSELLTSGSISTSYIYTIET------------
--DKDEEGQANAEKVLRG-LGLDPSAEDCIATRRVCQIVSTRAAHLCAATLAAVLRQIRDNK-AAEKLRT
TIGVDGSVYKNH-PEFPRRLHKMVRRLVPDCDPRFLQSQCGSGKGAAMVTAVAYRVAAQQAERQRILDTL
RLSREQLLEVKRRMTEGMVLGLSKQTHDQT-SVKMLPTYVRSTPDGT-----------------------
---EQGDFLALDLGGSSFRVLLVRLRSEKRQKVDMHQKIYSIPQETMQGTGEE-------LFDHIVHCIA
DFLEYMGMRGA-SLPLGFTFSFPCHQSKLDEGILLKWTKGFKASDCEGKDVVMLLKEAVRRKQ----EFD
LNFVA-----IVNDTVGTMMTCGYEDPKCEVGLIVGTGTNACYMEEM---HNIEMVE------------
--GN------EG---------------------RMCVNVEWGAFGEKGE-LDDFCTQFDHAVDDCSNFPG
KQR-YEKMISGMYLGEIVRNVLLDFTTKGLLFRGKLSERLKTRGIFETKFLSQIES--------------
------------DRLAMRQVRSILQHLGLTVSTCDDSVLVKEVCSVVARRAAQLCGAGLAALVDRIRQSR
NLNQLSITVGVDGTLYKTH-----------PHFSSIMQETLQDLAPQCQVTF---------LKSEDG---
----------SGKGAALITAVACRVES--EGR--Q-----------------

>XP_054591383|Nothobranchius furzeri|HK2
LQKLMVSDEILMDLSVRFRREMDKGL-CRDTNPTAA--VKMLPTYVRSTPDGTEQGEFLALDLG--GSNF
RVLLVKVKGN---GE-QK--VEMENQIYDIPEHVMRGSGSE---------------LFDHIAECLANFLE
KM-----GIK---DKKLPLGFTFSFPCQQTKLD-----------------------E------------
-----------SVLIYWTKGFRLSGVEGKEVVSLLRNAIRKR----------G-----------------
--------------DFDVDILAVINDTVGTMMTCGYDDHHCEVGLIV----------------------
-----G-----TGTNACYMEQMRNL-DVLDGDE--GRMCVNTEWGAFGDDGALEDLRTNIDREIDAGSLN
PGKQLFEKMISGMYMGELVRLILVKLARDQMVFQGKTTPELLTTGRFSTSYIYTIET------------
--DKDEEGLANADKVLRD-LGLNPSAEDCKATRRVCQIVSTRAARLCAAALAAVLRQIRDNK-AAERLRI
TIGADGSVYKTH-PGFSRRLQKMVRRLVPDCDVRFLQSPCGSGKGAAMVTAVAYRLAAQQAERQRILDTL
RLSREQLLEVKRRMTDGIVLGLSKQTHEQT-SVKMLPTYVRSTPDGT-----------------------
---ENGDFLALDLGGSSFRVLLVRLRSGKRHKVDMHQKIYTIPQETMQGTGEE-------LFDHIVQCIA
DFLEYMGMRGA-SLPLGFTFSFPCNQSKLDEGILLKWTKGFKASDCEGKDVVMLLKEAIQRKQ----EFD
LNFVA-----VVNDTVGTMMTCGYEDPKCEVGLIVGTGTNACYMEEM---HNVELVE------------
--GN------EG---------------------LMCVNVEWGAFGDFGE-LDDFCTQFDRSVDDCSNYPG
KQR-YEKMISGMYLGEIVRNVLIDFTSKGLLFRGKMSERLKSRGIFETKFLAQIES--------------
------------DRLAMRQVRSILQHLGLTGSTCDDSVLVKEVCSVVARRAAQLCGAGLAAVVDRMRQNR
NQNQLDITVGVDGTLYKTH-----------PHFSSIMQETLQDLAPQCRVTF---------LKSEDG---
----------SGKGAALITAVACRGNN--QGQ--H-----------------

>XP_061586344|Cololabis saira|HK2
LHQFKLSDRTLMDLSKRFRREMDKGL-CRDTNATAS--VKMLPTFVTSTPDGTEQGEFLALDLG--GSNF

RVLLVKVKAN---GE-QR--VEMKSQIFAIPENIMRGSGSE---------------LFDHIVDCLANFMD
MM-----GIK---DQKLPLGFTFSFPCQQTKLD------------------------E------------
-----------SILLSWTKGFKISGVEGKDVVGLLRESIRKR----------G-----------------
--------------DFDIDVVAVVNDTVGTMMTCGYDDHHCEIGLIV-----------------------
-----G-----TGTNACYMEQMRNL-DLLDGDE--GRMCVNTEWGAFGDDGSLEDVRTDTDREIDAGSLN
PGTQLFEKMISGMYLGELVRLIVVKMAREKLLFQGKTTSELLTTGRFSTSHLYTIEN-------------
--DSDEEGLASAEKVLRG-LGLEPSMEDCIAIRKVCQIVSTRAAHLCAATLAALLRQIRDNK-AAEKLRT
TIGVDGSVYKNH-PEFPRRLNKMLRRLVPDCDVRFQLSQDGSGKGSAMVTAVAHRLAAQHVERQRILDAL
RLSREQLLEVKRRMAEGMARGLSKNSREQT-SVKMLPTYVRSIPDGT-----------------------
---EDGNFLSLDLGGSSFRVLLVGLHSGTKHKVDMHQKIYSIPQETMQGTGDE-------LFSHIVDCIA
DFLEYMGMRGA-SLPLGFTFSFPCHQSKLDEGILLKWTKGFKASGCEGRDVVTLLKEAVRRKQ----EFD
LNFVA-----VVNDTVGTMMTCAYGDPRCEVGLIVGTGTNVCYMEEM---CNIDTVE-------------
--GN------DG---------------------QMCVNVEWGAFGDSGE-LDDFVTQFDCIVDDCSNHPG
KQR-YEKMISGMYLGEIVRNVLLDLTSKGLLFRGKGSERLKTRGIFETKFLSQIER--------------
------------DRLAMRQVRSILQHLGLTGSTSDDSVLVKEVCSVVARRAAQLCGAGLAAVVDKIRQNR
NLNQLSITVGVDGTLYKTH-----------PHFSSMMQETLQDLAPQCQVTF---------LKSEDG---
----------SGKGAALITAVACRING--GES--------------------
> XP_029960666|Salarias fasciatus|HK2
LHNFQLSDQVVMDVSMRFRREMDKGL-CRDTNPTAA--VKMLPTFVRSTPDGTEQGEFLALDLG--GSNF
RVLLVKVLGN---GK-QK--VEMENQIYEIPENIMRGCGSE---------------LFDHVADCLADFLE
KL-----GIK---DRKLPLGFTFSFPCLQTKLD------------------------E------------
-----------SVLISWTKAFRSSGVEGKDVVGLLRESIKKR----------G-----------------
--------------DFDVDILAVINDTVGTMMTCGFDDHHCEIGLIV-----------------------
-----G-----TGTNACYMEHMRNL-ELLDGDE--GRMCVNTEWGAFGDDGALEDLRTDIDREIDAGSLN
PGKQLFEKMISGMYMGELVRLILVKMTRERLLFQGKVTSALLTTGRFNTSYIYSIDN-------------
--DKEDEGLASAEKALRG-LGLDPSAEDCAATRRVCQIVSARAAHLCAATLASLLRQIRDNK-AAEKLRT
TIGVDGSVYKNH-AEFSRRLNKMLRRLVPDCDVRFLLSQDGSSKGAAMVTAVAFRVAAQQAERQRILDTL
RLSREQLLEVKRRMGEEMLRGLSRQSHDLT-SVKMLPTYVRSVADGT-----------------------
---EHGDFLALDLGGSSFRVLLVRVRSGKKRSVDMNHKIYNIPQETMQGTGEE-------LFSHIVNCIA
DFLEYMGMRGA-SLPLGFTFSFPCHQTKLDQGILLKWTKGFKASGCEGKDVVTLLKEAVRRKQ----EFD
LNFVA-----IVNDTVGTMMTCSYEDPRCEVGLIVGTGTNACYMEEM---HNIELVD-------------
--GN------DG---------------------RMCVNVEWGAFGDNGE-LDDFCTEFDRAVDDCSNFPG
KQR-YEKMISGMYLGEIVRNVLMDFTAKGLLFRGKVSERLKTRGIFETKFLSQIES--------------
------------DRLAMRQVRSILQHLGLTGSTCDDSVLVKEVCAVVARRAAQLCGAGLAALVDRIRQNR
NLNQLSVTVGVDGTLYKTH-----------PHFSSIMQETLQDLAPQCQVTF---------HKSEDG---
----------SGKGAALITAVACRMKD--EQQ--R-----------------
> CAJ1068369|Xyrichtys novacula|HK2
LLHFQLSVETIMDVSVRFRREMDKGL-CRDTNPAAT--VKMLPTFVLSTPDGTEQGEFLALDLG--GSTF
RVLLVKVRAN---GK-QE--VEMEDQIYDIPENLMRGSGSE---------------LFDHIADCLADFME
KM-----GIK---DKKLPLGFTFSFPCQQTKLD------------------------E------------
-----------GVLLSWTKNFKASGVEGKDVVSLLKKSIKKR----------G-----------------
--------------DFDIDILAVINDTVGTMMTCGYDDHLCEIGLIV-----------------------
-----G-----TGTNACYMEQMRNL-ELLDGDE--GRMCVNTEWGAFGDDGALDDLRTDVDREIDAGSLN
PGKQLFEKMISGMYMGEMVRLILVKMTKEKQLFQGNATAQLLTTGSFKTSYIYSIDT-------------
--DKDEEGLQKAEVVLRG-LGLDPSVDDCFATQRVCQTVSTRAAHLCAASLVAVLRQIRDNK-AAEKLRA
TIGADGSVYKNH-TEFSRRLHKMVRRLVPDCDVRFLQSQCGSGKGAAMVTAVAYRLAAQNAERQRILDTL
RLSREQLLEVKNRMSEAMVRGLSRQTHEQS-SVKMLPTYVRSTPDGT-----------------------
---EHGDFLALDLGGSSFRVLLVQLRSGTKQTVDMHQKIYGIPQETMQSTGEE-------LFNHIVDCIA
DFLDYMGMRGA-SLPLGFTFSFPCHQKKLDQGILLKWTKGFKASGCEGQDVIMLLKDAVRRRR----EFD
LNFVA-----VVNDTVGTMMTCGYEDCKCEVGLIVGTGTNVCYMEEM---RNIDLVE-------------
--GD------NG---------------------LMCVNMEWGAFGENGE-LDDFCTQYDRIVDESSNYPG
KQR-YEKMISGMYLGEIVRNILMDYTAKGLLFRGKLSERLKTRGIFETKFLSQIER--------------

------------DRLAMRQVRSILQHLGLTSSTCDDSFLVKEVCSVVARRAAQLCGAGLAAVVDKIRENR
NLNQLSITVGVDGTLYKTH-----------PHFSSIMQETLQDLAPQCQVTF---------HKSEDG---
----------SGKGAALITAVACRIQN--EEQ--N-----------------
> XP_060893251|Labrus mixtus|HK2
LPHLQLSDQTIMDVSVRFRREMDKGL-CRDTNPAAA--VKMLPTFVLSTPDGTEQGEFLALDLG--GSNF
RVLLVKVSAN---DK-QE--VEMENQIYAIPENLMRGSGSE---------------LFDHIADCLADFME
KL-----GIK---DKKLPLGFTFSFPCQQTKLD-----------------------E------------
-----------SVLLSWTKAFKAGGVEGKDVVGLLRRSIKKR----------G-----------------
--------------DLNIDILAVINDTVGTMMTCGFDDPLCEIGLIV-----------------------
-----G-----TGTNACYMEQMRNL-ELLDGDE--GRMCVNTEWGAFGDDGALEDLRTDIDREIDAGSLN
PGKQLFEKMISGMYMGEMVRLVLVRMTKEKLLFQGKTTAQLLTTGRFKTNYIYTIDT-------------
--DKDEEGLLSAEEVLRG-LGLDPSVKDCIATQRVCQIVSTRAAHLCAASLVALLRQIRDNK-AAEKLRA
TIGVDGSVYKNH-AEFSRRLHKMVRRLVPDCDVRFLTSQCGSGKGAAMVTAVAYRLAAQNAERQSILDTL
RLSHEQLLEVKKRLSAEMVRGLLRQMRDQT-SLKMLPTYVRSTPDGT-----------------------
---EHGDFLALDLGGSSFRVLLVHLKSGKKHTVDMQQKIYSIPQETMQSTGEE-------LFDHIVSCIA
DFLEYMGMKGA-SLPLGFTFSFPCHQSKLDQGILLKWTKGFKASGCEGQDVIVLLKDAVRRRQ----EFD
LNFVA-----VVNDTVGTMMTCSYEDPQCEVGLIVGTGTNACYMEEM---RNVEVVE-------------
--GD------DG---------------------RMCVNMEWGAFGEKGE-LDDFCTEFDRIVDESSNNPG
KQR-YEKMISGMYLGEIVRNVLIDYTAKGLLFRGKLSKRLETRGIFETKFLSQIER--------------
------------DRLAMRQVRSILQHLGLTSSTCDDSVLVKEVCSVVARRAAQLCGAGLAAVVDKIRENR
NLNQLSVTVGVDGTLYKTH-----------PHFSSIMQETLQELAPQCQVTF---------HKSEDG---
----------SGKGAALITAVACRIQN--GGQ--L-----------------
> XP_024660229|Maylandia zebra|HK2
LHNLQLSDKTLMDVSLRFRREMDKGL-CRDTNPTAA--VKMLPTFVRSTPDGTEQGQFLALDLG--GSNF
RVLLVKVMGN---GE-QK--VEMESQIYDIPEHIMRGSGSE---------------FFDHIAACLANYLD
KM-----GMK---DKKLPLGFTFSFPCQQTKLD-----------------------E------------
-----------AVLMSWTKGFRSSGVEGQDVVSLLRKSIKKR----------G-----------------
--------------DFDIDIVSVVNDTVGTMMTCGFDDRHCEVGLIV-----------------------
-----G-----TGTNACYMEQMRNI-GVLDGDE--GRMCVNTEWGAFGDDGALEDLRTDIDRELDAGSFN
PGKQLFEKMISGMYMGELVRLILVKMARSQLLFQGKTTSELLTTGHFSTSHIYAIEN-------------
--DKDEEGIASAEKILRG-LGLDPSVEDCIATRRVCQIVSTRAAHLCASSLASVLRQIRDNK-AAEKLRV
TIGVDGSVYKNH-PEFSRRLNKMVRRLVPDCDVRFLQSQDGSGKGAAMVTAVAFRLANQNADRQHILDTL
GLSREQLLEVKRRFSEEMTRGLSKQTHKQA-SIKMLPTYVRSTPDGS-----------------------
---EHGDFLALDLGGSSFRVLLVRVRSGTKRSVDMQQKIYSIPQEIMQGTGEE-------LFNHIVDCIA
DFLEYMGMKGA-SLPLGFTFSFPCDQTKLDEGILLKWTKGFKASGCEGKDVVALLKEAVRSRG----EFD
LNFVA-----VVNDTVGTMMTCGYEDPKCEVGLIVGTGTNACYMEEM---HNIELVE-------------
--GD------NG---------------------RMCVNVEWGAFGENGE-LEEFCTEFDRLVDACSNYPG
KQR-YEKMISGMYLGEIVRNVLLDFTAKGLLFRGKVSERLKTRGIFETKFLSQIES--------------
------------DRLAMRQVRSILQHLGLTGSTCDDSVLVKEVCSVVARRAAQLSGAGLAAVVDKIRQNR
NLNQLSITVGVDGTLYKTH-----------PHFSAIMQETLRDLAPQCEVTF---------LKSEDG---
----------SGKGAALITAVACRVKN--EGQ--Q-----------------
> XP_040051867|Gasterosteus aculeatus aculeatus|HK2
LDHFQLSDKTIMDLSIRFRREMDKGL-CRDTNPTAA--VKMLPTFVRSTPDGTEQGEFLALDLG--GSNF
RVLLVKVMAN---GK-QE--VEMENQIYAIPEHLMRGSGSQ---------------LFDHIADCLANFLE
KL-----GIK---EKKLPLGFTFSFPCQQTKLD-----------------------E------------
-----------SVLVSWTKSFKSSGVEGKDVVSLLRKSIKKR----------G-----------------
--------------DFDVDILAVINDTVGTMMTCGYDDHHCEIGLIV-----------------------
-----G-----TGTNACYMEHMRNL-DLLDGDE--GRMCVNTEWGAFGDDGSLEDLRTGIDREIDAGSLN
PGKQLFEKMVSGLYMGELVRLILLRMTKEQLLFQGKTTPQLLTTGSFNTSHIFAIEN-------------
--DKDEEGLASAQQVLQG-LGLDPSVEDCRTTRRVCQIVSTRAAHLCAATLVAVLRQIRDNK-AAEKLRT
TIGVDGSVYKNH-PEFSRRLHKMVRRLVPDCDVRFLQSQCGSGKGAAMVTAVAYRLAAQHAERQRILDTL
RLSREKLLEVKRRMSEEMARGLSKLTHEEA-SVKMLPTFVRSTPDGT-----------------------

---EHGDYLALDLGGSTFRVLLVRLRSEKGHNVDMHHKIYSIPQEIMQGTGDE-------LFGHIVCCIG
DFLEYMGMKGA-SIPMGFTFSFPCHQSKLDQGILLNWTKGFKASGCEGQDVLTLLKDAVRRRQ----DFD
LNFVA-----LVNDTVGTMMTCSYEDPQCEVGLIVGTGTNACFMEEM---QNIELVE-------------
--GD------HG---------------------RMCVNMEWGAFGDKGE-LDDFCTEFDRVVDNSSNNPG
KQR-YEKMISGMYLGEIVRNVLIDFTNKGLLFRGKLSERLKTRCIFETKFLAQIES--------------
------------DRLAMRQIRSILQHLGLTTSTCDDSVLVKEVCRVVSSRAAQLCGAGLAAVVDKIRENR
NLNQLSITVGVDGTLYKTH-----------PHFSSIMQETLQDLAPQCQVTF---------NKSEDG---
----------SGKGAALITAVACRMKS--EGQ--P-----------------
> XP_030271525|Sparus aurata|HK2
LQTLQLSDETLMDVSIRFRREMDKGL-CRDTNPTAA--VRMLPTFVLSTPDGTEQGEFLALDLG--GANF
RVLLVKVMAD---GE-QK--VEMENQIYDIPEHLMRGSGTE---------------LFDHIADCLANFLE
KL-----GIK---DKKLPLGFTFSFPCQQTKLD------------------------E------------
-----------SVLLSWTKAFKSSGVEGKDVVSLLRKSIKKR----------G-----------------
--------------DFDVDILAVINDTVGTMMTCGYDDHLCEIGLIV-----------------------
-----G-----TGTNACYMEQMRNI-ELLDADE--GRMCVNTEWGAFGDDGTLEDLRTDIDREIDAGSLN
PGKQLFEKMISGMYMGELVRLILVRMTREQLLFQAKTTAELLTTGSFKTSYIYGIDN-------------
--ENNEEGLASAEKVLRG-LNLDPSVEDCIATQRVCQIVSTRAAHLCAATLVAVLRQIRDNK-AAEKLRT
TIGVDGSVYKNH-PEFSRRLHKMVRRLVPDCDVRFLQSQCGSGKGAAMVTAVAYRLAAQNAERQRILDTM
CLSREQLLEVKRRMSEEMVRGLSKQTHEQA-SVKMLPAYVRSTPDGT-----------------------
---EHGDFLALDLGGSSFRVLLVQVQSGQRHKAKMQQKIYSIPQETMQGTGEE-------LFDHIVDCIA
AFLEYMGMKGA-SLPLGFTFSFPCHQSRLDQGILLKWTKGFKASGCEGHDVILLLKDAVRRRQ----EFD
LNFVA-----VVNDTVGTMMTCGYEDPKCEVGLIVGTGTNACYMEEM---SNIELVE-------------
--GN------EG---------------------RMSINMEWGAFGDNGE-LDDFCTRFDHIVDDSSNNPG
KQR-YEKMISGMYLGEIVRNVLIDFTSKGLLFRGKLSERLKTWGIFATKFLSQIES--------------
------------DRLAMRQIRSILQHLGLTSSTIDDSVLVKEVCSVVARRAAQLCGAGLAAVVDRMRENR
NLNQFSVTVGVDGTLYKTH-----------PHFSSIMQETLQDLAPQCQVTF---------QKSEDG---
----------SGKGAALITAVACRIKS--EGQ--H-----------------
> XP_063753384|Eleginops maclovinus|HK2
LHHFQLSDKTLMDLSIRFRREMDKGL-CRDTNPTAA--VKMLPTFVRSTPDGTEQGEFLALDLG--GSNF
RVLLVKVMDN---GK-QE--VEMENQIYDIPEHLMRGCGSE---------------LFDHIADCLANFLE
KL-----GIK---DKKLPLGFTFSFPCQQTKLD------------------------E------------
-----------SVLVHWTKSFKAYGVEGKDVVSLLRKAIKKR----------G-----------------
--------------DFDVDILAVINDTVGTMMTCGYDDHHCEIGLIV-----------------------
-----G-----TGTNACYMEQMRNL-DLLDGDE--GRMCVNTEWGAFGDDGALEDLRTNVDRDIDAGSLN
PGNQLFEKMISGLYMGEVVRLILVKMTKEKLLFQGKTSAKLLTTGSFSTSFIYGIDT-------------
--DKDEEGVERAEEVLRG-LGLNPSVEDCIATQRVCEIVSTRAAHLCASALVAVLRQIRDNK-AAEKLRT
TIGVDGSVYKNH-SEFSRRLHKMVRRLVPDCDVRFLQSQCGSGKGAAMVTAVAYRLAAQHAERQRTLDTL
RVSREQLLEVKKRMSEEMERGLSKQTHEQS-SVKMLPSYVRDTPDGT-----------------------
---EHGDFLALDLGGSSFRVLLVRLRGGKGHNVDMHHKIYSIPQETMQGTGEE-------LFSHIVDCIA
DFLEYMGMSGA-SLPLGFTFSFPCFQSKIDQGILLKWTKGFKASGCEGQDVITLLKDAVRRRR----EFD
LNFVA-----VVNDTVGTMMTCSYEDPQCEVGLIVGTGTNACYMEEM---QNIGRVE-------------
--GN------DG---------------------RMCVNMEWGAFGDNGE-LDDFCTAFDHMVDDSSNYPG
KQR-YEKMISGMYLGEIVRNVLIDFTAKGLLFRGKLSERLKTRGIFETKFLAQIES--------------
------------DRLAMRQVRSILQHLGLTISTCDDSVLVKEVCSVVSRRAAQLCGAGLAAVVDKIRQNR
NLNQLSITVGVDGTLYKTH-----------PHFSSIMQETLQDLAPQCQVTF---------HKSEDG---
----------SGKGAALITAVACRVKN--EVQ--KQ----------------
> XP_029365634|Echeneis naucrates|HK2
LKRLQLSDETLMDLSLRFRREMDKGL-CRDTNPTAA--VKMLPTFVCSTPDGTEQGEFLALDLG--GCNF
RVLLVKVMAD---GE-RK--VEMENQIYAIPENLMRGCGSE---------------LFDHIADCLSNFME
KK-----GIK---EKKLPLGFTFSFPCQQTKLD------------------------E------------
-----------SVLLSWTKRFKASGVEGKDVVSLLRKAIQKR----------G-----------------
--------------DFDIDIVTVVNDTVGTMMTCGYDDQHCEIGLIV-----------------------

-----G-----TGTNACYMEQMRNL-ELLDGDE--GRMCVNTEWGAFGDDGSLEDLRTDIDRELDAGSLN
PGKQLFEKMIGEMYMGELVRLILLRMAKEQLLFQGKTTAELLTTGGFKTTDIYAIEN-------------
--DKAEEGLASAEKVLRG-LHLEPSKEDCIATQRVCQIVSTRAAHLCSATLAAVLRQIRDNK-AAEKLRA
TIGVDGSVYKYH-PEFSRRLNKMVRRLVPDCDVRFLLSQDGSGKGSAMVTAVAYRFAAQQAERQRILDTL
RLSRNQLLEVKERMNEEMARGLSKQTHEQA-SIKMLPTYVRSTPDGT-----------------------
---EHGDFLALDLGGSSFRVLLVQVRSGTKSNVDMHHKIYSIPQETMQGTAEE-------LFDHIVDCIA
DFLKYRDMRGV-SLPLGFTFSFPCHQSKLDQGILLRWTKGFKASGCEGHDVVSLLKDAVRRKQ----EFD
LNFVA-----VVNDTVGTMMTCSYEDPKCEVGLIVGTGTNACYMEEM---HNIELVE-------------
--GD------SG---------------------LMCVNMEWGAFGEGGE-LDDFCTQFDRVVDACSSNPG
KQR-YEKMISGMYLGEIVRNVLLDFTAKGLLFRGKLSERLKTHGIFETKFLTQIEK--------------
------------DRLAMRQIRSILQHLGLTSSTCDDSVVVKEVCSVVARRAAQLCGAGLAAVIDKIRQNR
NLNQLSITVGVDGTLYKTH-----------PHFSSIMQETLQDLAPQCQVTF---------HKSEDG---
----------SGKGAALITAVACRIMT--QDN--NCEV--------------
>XP_028269324|Parambassis ranga|HK2
LQCLQLSDETLMDVSMRFRREMDKGL-CRDTNPAAA--VKMLPTFVRSTPDGTEQGEFLALDLG--GSNF
RVLLVKVMAD---GM-QK--VEMESQIYAIPEHIMRGSGSE---------------LFDHVVDCLANFVD
KM-----GIK---DKKLPLGFTFSFPCQQTKLD------------------------E------------
-----------SVLLSWTKGFRSSGVEGKDVVSLLRKSIKKR----------G-----------------
--------------DFDIDIVAVVNDTVGTMMTCGYDDQHCEIGLIV-----------------------
-----G-----TGTNACYMEQMRNI-ELLDGDE--GRMCVNTEWGAFGDDGALEDLRTDIDREIDAGSLN
PGKQLFEKMISGMYLGELVRLILVKMAREQLLFQGKTTPDLLTTGRFSTSYIYSIEN-------------
--DNNEEGLTSAEKVLRG-LSLDPSVEDCIATRRVCQIVSTRAAHLCAASLAAVLRQIRDNK-AAEKLRS
TIGVDGSVYKNH-PEFPRRLNKMLRRLVPDCDVRFLLSQDGSGKGSAMVTAVAYRLAAQHAERQRILDTL
RLSREQLLEVKKRMSEEMVRGLSKQTHEQA-SVKMLPTYVRSTPDGT-----------------------
---EHGDFLALDLGGSSFRVLLVQVRSGKRHNVDMHHKIYSIPQETMQGTGEE-------LFNHIVYCIT
DFLEYMGMRGA-SLPLGFTFSFPCHQSRLDQGILLKWTKGFKASGCEGQDVVMLLKEAVQRKK----AFD
LNYVA-----VVNDTVGTMMTCGYEDPKCEVGLIVGTGTNICYMEEM---HNIETVE-------------
--GD------DG---------------------RMCINIEWGAFGENGE-LDDFRTQFDCIVDDCSNYPG
KQR-YEKMISGMYLGEIVRNVLLDFTAKGLLFRGKVSERLKTRGIFETKFLSQIER--------------
------------DRLAMRQVRSILQHLGLTGSTCDDSVLVKEVCSIVARRAAQLCGAGLAAVVDKIRENR
NLNELYITVGVDGTLYKTH-----------PHFSSIMQETLQDLAPQCHVTF---------HKSEDG---
----------SGKGAALITAVACRIKS--GGQ--H-----------------
>XP_042276480|Thunnus maccoyii|HK2
LHHLQLSDETLMDVSIRFRREMDKGL-CRDTNPTAA--VKMLPTFVLSTPDGTEQGEFLALDLG--GTNF
RVLLVKVMAD---GK-QK--VEMEDQIYAIPEHIMRGSGSE---------------LFDHIADCLANFLE
KL-----GMK---NKKLPLGFTFSFPCKQTKLD------------------------E------------
-----------SVLVSWTKGFRISGVEGQDVVSLLRKSIKKR----------G-----------------
--------------DFDVDIMAVINDTVGAMMTCGYDDRHCEIGLIV-----------------------
-----G-----TGTNACYMEQMRNL-ELLDGDE--GRMCVNTEWGAFGDDGALEDLRTDIDREIDAGSLN
PGKQLFEKMISGMYMGELVRLILVRMAKAQLLFQGKTTAELLTTGSFNTSYIYAIEN-------------
--DKDEEGVASAEKVLRG-LGLDPSLEDCIATRRVCQIVSTRAAHLCAATLVALLRQIRDNK-AAEKLRA
TIGVDGSVYKNH-PEFHRRLNKMVRRLVPDCDVRFLRSEVGSGKGAAMVTAVAYRLAAQHAERQGILDTL
RLSREQLLEVKKRMSEEMVRGLSKQTYEQA-TVKMLPTYVRSTPDGT-----------------------
---EHGDFLALDLGGSTFRVLLVRVRSGKRHKVDMHHKIYSIPQETMQGTGEE-------LFSHIVYCIS
DFLEYMGMRGA-SLPLGFTFSFPCQQSKLDQGILLNWTKGFKASGCVGEDVVTLLKDAVRCRG----EFD
LNFVA-----VVNDTVGTMMTCGYEDPKCEVGLIVGTGTNVCYMEEM---QNIEVVE-------------
--GD------DG---------------------RMCVNMEWGAFGDNGE-LDDFFTQFDRFVDDCSNNPG
KQR-YEKMISGMYLGEIVRNVLMDFTAKGLLFRGKLSDRLKTRGIFETKFLSQIES--------------
------------DRLAMRQVRSILQHLGLTSSTCDDSVLVKEVCSVVSRRAAQLCGAGLAAVVDKIRQNR
NLQQLSITVGVDGTLYKTH-----------PHFSSIMQETLQDLAPQCKVTF---------HKSEDG---
----------SGKGAALITAVACRIKS--GGQ--H-----------------
>XP_059192483|Centropristis striata|HK2

LHHFQLSDKTLMDLSIRFRREMDKGL-CRDTNPTAA--VKMLPTFVLSTPDGTEEGEFLALDLG--GSNF
RVLLVKVMAD---GR-QE--VEMEDQIYDIPEHLMRGCGSE---------------LFDHIADCLANFME
KL-----DIK---DKKLPLGFTFSFPCLQTNLD------------------------E------------
-----------SVLVSWTKGFKSSGVEGKDVVNLLRKSIKKR----------G-----------------
--------------DFDIDIVAVTNDTVGTMMTCGYDDHLCEIGLIV-----------------------
-----G-----TGTNACYMEQMRNL-ELLDGDE--GRMCVNTEWGAFGDDGALEDLRTDIDREIDAGSLN
PGKQLFEKMISGLYMGELVRLILVRMTKEKLLFQGKTTAQLLTTGSFNTSYIYAIEN-------------
--DKDEEGLASAEKVLRG-LGLDPLVEDCIATQRVCQIISTRAAHLCAATLVAVLRQIRDNK-AAEKLRT
TIGVDGSVYKNH-PEFSRRLHKMVRRLVPDCDVRFLQSQCGSGKGAAMVTAVAYRLAAQHAERQGILDTL
RLSHEQLLEVKEQMSKEMVRGLSKQTHEQA-SVKMLPTFVRATPDGT-----------------------
---EHGDFLALDLGGSSFRVLLVRVRSGKKHNVDMHHKIYSIPQETMQGTGQE-------LFSHIVDCIA
DFLEYMGMRGA-SLPLGFTFSFPCHQSKLDQGILLKWTKGFKATGCEGQDVITLLKDAVRRRQ----EFD
LNFVA-----VVNDTVGTMMTCSYEDPQCEVGLIVGTGTNACYMEEM---KNIEVVE-------------
--GD------EG---------------------LMCVNMEWGAFGDNGE-LDDFCTQFDHTVDECSNNPG
KQR-YEKMVSGMYLGEIVRNVLIDFTAKGLLFRGKMSERLKTRCIFETKFLSQIET--------------
------------DRLAMRQIRSILQHLGLTTSTCDDSVLVKEVCSVVARRAAQLCGAGLAAVVDKMRQSR
NLNQLSVTVGVDGTLYKTH-----------PHFSSIMQETLQALAPQCQVTF---------HKSEDG---
----------SGKGAALITAVACRVKS--EEQ--H-----------------

> XP_022607332|Seriola dumerili|HK2

LHRLQLSDETLMDLSIRFRREMDKGL-CRDTNPTAA--VKMLPTFVRSTPDGTEQGEFLALDLG--GSNF
RVLLVKVMAD---GE-QK--VEMENQIYAIPEHLMRGSGSE---------------LFDHIADCLSSFLE
KM-----GIK---DKKLPLGFTFSFPCQQTKLD------------------------E------------
-----------SVLVSWTKGFKSSGVEGKDVVSLLRKSIKKR----------G-----------------
--------------DFDIDIVAVVNDTVGTMMTCGFDDQHCEIGLIV-----------------------
-----G-----TGTNACYMEQMRNL-ELLDGDE--GRMCVNTEWGAFGDDGALEDLRTDIDQELDAGSLN
PGKQLFEKMISGLYMGELVRLILVRMTKEQLLFQGKTTAQLLTTGSFSTSYIYAIEN-------------
--DKGEEGLTSAEKVLRD-LGLEPSVEDCISTQRVCQIVSTRAAHLCAATLVAVLRQIRDNK-AAERLRT
TIGVDGSVYKNH-PEFSRRLHKMVRRLVPDCDVRFLLSQDGSSKGSAMVTAVAYRLAAQHAERQRVLDTL
RLSREQLLEVKNRMSEEMGRGLSKQAHEQA-SVKMLPTYVRSTPDGT-----------------------
---EHGDFLALDLGGSSFRVLLVRVRSGKRHNVDMHHKIYSIPQEIMQGTGEE-------LFNHIVDCIA
DFLEYMGMRGT-SLPLGFTFSFPCHQSKLDQGILLKWTKGFKASGCEGQDVVTLLKDAVRRRQ----EFD
LNFVA-----VVNDTVGTMMTCGYEDPNCEVGLIVGTGTNACYMEEM---HNIELVE-------------
--GN------DG---------------------CMCVNMEWGAFGDNGE-LDDFCTQFDRTVDDCSNNPG
KQR-YEKMISGMYLGEIVRNVLMDFTSKGLLFRGKLSERLKTHGIFETKFLSQIEK--------------
------------DRLAMRQIRSILQHLGLTSSTCDDSVLVKEVCSVVARRAAQLCGAGLAAVVDKIRQNR
NLNQLSITVGVDGTLYKTH-----------PHFSSIMQETLQDLAPQCPVTF---------HKSEDG---
----------SGKGAALITAVACRIKS--GGQ--H-----------------

> XP_038627012|Tachyglossus aculeatus|HK3

LQVYDLSEEQLRVIQAHILEDMERALSRRQTKSTSS--IQMLPTYVRSTPHGNEKGDFLVLELGGTDATL
RVLWVTLTGD---AH-HK--IEPKSQVFSIPSDVMLGSGLQ---------------LFDFVAKCLTNFLD
QQ-----QVG-GDKQPIKMGFSFSFPCHQTGLD------------------------R------------
-----------STLISWTKGFKCSDVEGKDVVQLLRDAIKRQ----------E-----------------
--------------KYRIDVVAVVNDTVGTMMSCGSEELPCEIGLIV-----------------------
-----D-----TGTNSCYMEEARHV-AGLEEEQEGGKICVNTQWGSFGDDGSLK--QTFFDVQLDLFSLN
PGKQRFEKMVSGMYLGELVRLTLVHLAEKGALFGGNVSEALKKHNAILIQHVVKMED-------------
--PS--AGLAQTLALLKE-LGLAASEQDCRLVQQACKAVSTRAARLCAAALAATLNRLQSNR-ELARLPV
AVATAGAVCQQH-PSFSRILQETVKALAPACDVSFVPSEDGGGRGVAVVTAVASRQATHRRLLAETLAPL
RLKGHQLQALQAQMQEAMERGLCGEA---S-SLRMLPTFVRATPDGS-----------------------
---ERGDFLALDLGGTNFRVLLVQLRARVESGISITSQVYSLPQDVIQGTGEQ-------LFDHIVNCIV
DFQKKNGLAGR-VLPLGFTFSFPCQQLGLDQGILLSWTKGFNASNCEGKDVVTLLRQAIARRK----DVQ
LNVVA-----IVNDTVGTMMSCAYDDLRCEVGLIVGTGTNACYMEEM---KHVMTLP-------------
--GE------EG---------------------RMCINMEWGAFGDDGG-LDHLITDFDCMVDKNSINPG

KQK-FEKLISGMFLGEIVRSILLHLTKREVLFRGREVNRLQTKDIFKTKFLSDIER--------------
------------DSLALRQVRAILLDLGL-QLTCDDALVVREVCQAVSRRAAQLCGAGVAAIVEKIRRNR
GLDQLSITVGVDGTLYKLH-----------PHFSSHVQEVVRELAPHCTVTF---------LQSEDG---
----------SGKGAALITAVACRVAN-QTGP--------------------
> XP_051834965|Antechinus flavipes|HK3
LSQFDVTEEQLQQIQAGILRDMERAL-GRQAGSSPS--VQMLPTYVRSTPRGTEQGDFLVLELGASGATL
RVLWVTLTGI---SG-PR--KEPVSQEFTIPTEITLGTGQQ---------------LFDFAAGCLVQFLD
QL-----QVG---SQPIQLGFNFSFPCHQTGLN------------------------Q------------
-----------STLISWTKGFRCSGVEGQDVVQLLRQAIQRQ----------G-----------------
--------------AYSIEVVAVVNDTVGTMMSCDPSPQACEVGLVV-----------------------
-----D-----TGTNACYMEEVRHV-AELREDW--GRICVNVQWGSFGDGGALEPVQTPFDLQLDQESLN
PGTQRFEKMVGGLYMGELVRLVLIHLAGQGALFGGVPSPALLTHGSVHVQHVMEIED-------------
--PS--HGRARAQAVLQG-LGLHPSAQDCAQVQRVCGAVSSRAARLCAAALAAVLARVRSNP-ERSPLGV
SVATGGRVYEQH-PRFSALLQQTVRLLAPDADVTFVPSEDGGGRGVAVVTAVASRLALQRRQLEETLAPF
RLHSQQLKMVQAQMRESMARGLREEG---S-SVRMLPTYVCALPDGT-----------------------
---ERGDFLALDLGGTNFRVLWVKVGSRSEGGVKIINRVFSLPESVTQGTGQQ-------LFDHIVDCIV
DFQKENNLSGQ-TLPLGFTFSFPCKQLDLDQGILLNWTKGFKASGCEGQDVVSLLRAAIRRRQ----GVE
LNVVA-----IVNDTVGTMMSCGYEDPHCEVGLIVGTGTNACYMEEL---GNVSAVP-------------
--GD------KG---------------------RMCINMEWGAFGDDGLALQEFYTPFDISVDQASINPG
KQR-YEKLISGMYLGEIVRQVLLHLTRLGVLFGGQVFQSLQTKDLIKTKFLSEIES--------------
------------D-LERWQVRAILEKLDL-SANCDDALLVREVCQAVSQRAAKLCGAGVAAVVEKIRENR
GLDQLRVTVGVDGTLYKLH-----------PHFSEQVKDTVRQLAPQCSVRF---------LLSEDG---
----------SGKGAALVTAVACRLAG--ATQ--S-----------------
> XP_007474181|Monodelphis domestica|HK3
LKQFYVTEEQLQQIQAGILRDMEKAL-SRQTDPSPT--VQMLPTYVRSTPHGTEQGNFLVLELGASGATL
RVLWVTLTGT---SG-PR--KEPISREFTIPPEIMLGTGQQ---------------LFDFAASCLVEFLD
QL-----KVG---SQHIQLGFNFSFPCHQTGLD------------------------R------------
-----------STLISWTKGFQCSGVEGQDVVQLLREAIQRQ----------G-----------------
---------PLLPQGYSIDVVAVVNDTVGTMMSCDPRPQACEIGLVV-----------------------
-----D-----TGTNACYMEEARHV-GELGEDQ--GRVCVNVQWGSFGDSGALEPVQTPFDLLLDRESLN
PGTQRFEKMVGGLYLGELIRLVLIHLAGQRALFDGATSPALLTHGSVPIEHVMEIED-------------
--PS--QGRARAQAVLQG-LGLNPSSQDCQWVQQVCGAVCARAARLCAAALVAILARVEHNQ-ERSQPGI
AVATGGRICEQH-PRFSTILQQTVKLLAPDSDVTFVPSEDGRGRGVAVVTAVASRLALHRRQLEETLAPF
RLNSQQLKAVQVQMQEAMVRGLRGEA---S-SLPMLPTYVCSTPNGT-----------------------
---ERGDFLALDLGGTNFRVLLVKVGSKSEGGVKITNQVFSIPESVTQGTGQQ-------LFDHIVDCIV
DFQKEHGLVGQ-ILPLGFTFSFPCKQLGLDQGILLNWTKGFKASDCEGQDVVALLRAAIRRRK----GVE
LNVVA-----IVNDTVGTMMSCGYEDPRCEVGLIVGTGTNACYMEEL---CNVSSVP-------------
--GD------QG---------------------RMCINMEWGAFGDDGL-LKELCTPFDINVDKSTINPG
KQR-FEKLISGMYLGEIVRQVLLQLTHLGVLFGGEVSQRLQTKDIFKTKFLSEIES--------------
------------EGLERRQVRAILGALGL-RVNCDDALMVREVCQAVSQRAAGLCGAGVAAVVEKIRENR
GLEELNVAVGVDGTLYKLH-----------PHFSEQVKDTVQRLAPRCSVTF---------LQSEDG---
----------SGKGAALVTAVACRLAA--GAL--------------------
> XP_043837583|Dromiciops gliroides|HK3
LKQFNVTEKQLRQIQEGILQDMEQAL-SRQTDPSPT--VRMLPTYVRSTPHGTEQGDFLVLELGASGATL
RVLWVTLTGI---TG-PR--KEPMSREFTIPTEITLGPGQQ---------------LFDFAAGCLVEFLD
QL-----KVG---SQPIQLGFNFSFPCHQTGLD------------------------Q------------
-----------STLISWTKGFRCSGVEGQDVVQLLREAIQRQ----------G-----------------
--------------AYSIDVVAVVNDTVGTMMSCDPRSQACEIGLVV-----------------------
-----D-----TGTNACYMEEVRHV-VELGEDQ--GRVCVNVQWGSFGDRGALEPVQTPFDLQLDQESLN
PRTQRFEKMVGGLYLGELVRLVMIHLAGQRALFGGVTSPALLTRGSVHIEHVMEIED-------------
--PS--HGRVRAQAVLQR-LGLNPSTQDCELVQQICAAVSARAAHLCAAALAAILARVKSNR-ERVQPGT
AVATGGHVYEQH-PRFSSILQQTVRLLAPDSDITFVPSEDGGGRGVAVVTAVASRLALHRQQLEETLAPF

RLNSEQLKAVQAQMREAMVRGLRGEQ---S-SLPMLPTYVCATPDGT-----------------------
---ERGDFLALDLGGTNFRVLWVKVGSRSEGGVKITNQVFSIPESVTQGTGQQ-------LFDHIVACIV
DFQKRHGLSGQ-TLPLGFTFSFPCRQLGLDQGVLLNWTKGFKASDCEGQDVVSLLRAAIRRRQ----GVE
LNVVA-----IVNDTVGTMMSCGYEDPQCEVGLIVGTGTNACYMEEL---CNVSAVP-------------
--GD------RG---------------------RMCINMEWGAFGDEGL-LKELCTPFDISVDKSSINPC
KQR-FEKLISGMYLGEIVRQILLHLTRLGVLFRGQVFQCLQTKDIFKTKFLSEIES--------------
------------NDLERRQVRAILEELGL-PVSYDDALMVREVCQAVSQRAAKLCGAGVAAVVEKIRENR
GLEELSVAVGVDGTLYKLH-----------PHFSDQVKDTVRHLAPRCSVRF---------LQSEDG---
----------SGKGAALVTAVACRLAE--SPQ--A-----------------
>XP_036607607|Trichosurus vulpecula|HK3
LKQFDVMEKQLQEIQAGILQDMEQAL-CRQTDPTPT--VRMLPTYVRSTPQGTEQGNFLVLELGASGATL
RVLWVTLGGV---TG-QR--TQPISWEFTIPTEIMLGTGQQ---------------LFDFAAGCLVEFLD
QL-----KVG---SQSIQLGFNFSFPCHQTGLD------------------------Q------------
-----------STLISWTKGFRCSGVEGQDVVQLLRDAIQRQ----------G-----------------
--------------AYSIEVVAVVNDTVGTMMSCDPRPQACEIGLVV-----------------------
-----D-----TGTNACYMEEARHV-RELGEDK--GRICVNVQWGSFGDSGALEPVQTPFDLQLDQESLN
PGTQRFEKMVGGLYLGELVRLVLIHLAGQRALFGGDTSPALLTRGSVHIEHVMEIEH-------------
--PC--HGRARAQAVLQG-LGLNPSTQDCEQVQQVCGAVCARAARLCAAALTAILVRVKSNQ-ERSQPGI
AVATGGHVYEQH-PRFRTILQQTVRLLAPDSDITFVPSEDGGGWGVAVVTAVASRLALHRRQLEQTLAPF
RLDSEQLKAVQAQMREAMVQGLQGGD---S-SLRMLPTYVCATPDGT-----------------------
---ERGDFLALDLGGTNFRVLWVKVDSRNEGGIKIINQVFSLPQSVTQGTGQQ-------LFDHIVDCIV
DFQKQHDLSGQ-TLPLGFTFSFPCKQLGLDQGILLNWTKGFKASDCEGQDIVSLLRAAISRRE----GIE
LNVVA-----IVNDTVGTMMSCGYEDPHCEVGLIVGTGTNACYMEEL---CNVSAVP-------------
--GD------QG---------------------RMCINMEWGAFGDDGL-LKEFCTPFDINVDKCSINPG
KQR-FEKLISGMYLGEIVRQILLLLSRLGVLFGGQVFQCLKTKDIFKTKFLSEIER--------------
------------DNLERWQVQAILEKLGL-TVSCDDALMVREVCQAVSQRAAKLCGAGVAAIVEKIRENR
GLEELSVTVGVDGTLYKLH-----------PHFSNQVKDTVQHLAPRCSVRF---------LQSEDG---
----------SGKGAALVTAVACRLAK--STQ--A-----------------
>XP_049631651|Suncus etruscus|HK3
LQQFLVTEAQLRQIQASLRSSMDQAL-KGPASPCPA--VRMLPTYVGSIPHGTEKGDFVVLELGAKGAFL
RVLWVTLMGT---EG-PR--MTSRSQEFIIPQEVMQGPGQQ---------------LFDFAARCLLEFLG
PL-----PEM---EYKLQLGFSFSFPCQQTALD------------------------Q------------
-----------STLISWTKGFRCSDVEGRDVVQLLREALERQDLVSGKGSVHG-----------------
--------------SMNVDVVAVVNDTVGTMMGCEPGSGPCEIGLVV-----------------------
-----D-----TGTNACYMEEAGHV-AMLDEDR--GRVCISVEWGSFGQDGVLGPVLTPFDRTLDLESLN
PGAQRFEKMIGGLYLGELVRLALVHLAQAGVLFGGCTSSALLTPGSFRLEDVAEMEA-------------
--SW--ALEPHTCAVLQR-LGLSVGAADTMWVQHLCVVVCARAARLCAAALAAVLSRLQHSR-EQPGLQL
SVAIGGRVFEQL-PRYDGVLKGTVGFLVPECEVSFHRSLDGSGQGVAMVTAVAARLAAHQRLLASTLAPF
RLSREQLASVQEQMREAMARGLRGEA---S-SLRMLPTYVRATPDGS-----------------------
---ERGDFLALDLGGTHFRVLLVHLAAG---SVQIAQQVYSVPEHVAQGSGQE-------LFDHIVDCIV
DFQQRQGLSGQ-SLPLGFTFSFPCRQMGLDQGILLNWTKGFKASDCEGHDVVRLLREAIERKQ----GVE
LNVVA-----IVNDTVGTMMSCGYEDPQCEVGLIVGTGTNACYMEEL---QNMAEFQ---------NVAG
AAGD------SG---------------------HMCINMEWGAFGDDGS-LDTLSTRFDANVDQASINPY
KQR-FEKMISGMYLGEIVRHVLLHLTNLGVLFRGQESQRLQVQDIFKTQFLSEIES--------------
------------DSLALRQVRTILEDLGL-VLTSDDAQMVLEVCQAVSRRAAQLCGAGVAAVVEKIRENR
GLDKLAVSVGVDGTLYKMH-----------PHFSSLVAATVHELAPHCKVTF---------LQSEDG---
----------SGKGAALVTAVACRLAS--VTR--V-----------------
>XP_023582186|Trichechus manatus latirostris|HK3
LQQFTVTVAQLQQIRTSLLGSMEQAL-MGQNDPAPA--VRMLPTYVGSTPHGTEQGDFVVLELGATGASV
RVLWVTLTGT---EG-HR--VEPRSQEFVIPPEVMLGPGKQ---------------LFDFAACCLTKFLE
AL-----PVS---NQCLPLGFSFSFPCHQTGLD------------------------R------------
-----------SILISWTKGFRCSDVEGQDVVQLLRDAIERQ----------G-----------------

--------------APSLNVVAVVNDTVGTMMGCDPELGPCEIGLLV-----------------------
-----D-----LGTKACYMEETKHV-AVLGEDL--GSVCVSIEWGSFCDDEALGPVLTSFDHALDQESLD
PGAQRFEKMIGGLHLGELVRLVLVDLAQRGLLFGGCTSPALLSQGSVALEHVAEMED-------------
--PS--AGTARVLAVLQN-LGLSPEASDVELVQRVCAAVTTRAARLCAAALAAILGRLQHSR-EEHALQI
AVATGGRVFERH-PSFLGLLKETVMLLAPECGVSFIPSVDGGGRGVAVVTAVVTRLATNRRLLEETLVPF
RLDREQLAAVQVQMQEAMAKGLRGEA---S-SLRMLPTYVRAVPDGS-----------------------
---ERGDFLALDLGGTNFRVLLVRVAAT---GVQITSEVYTIPELMAQGPGQQ-------LFDHIVDCIV
DFQQKKGLVGQ-SLPLGFTFSFPCRQLGLDQGILLNWTKGFSASDCVGQDVVALLREAIRRRQ----AVE
LNVVA-----IVNDTVGTMMSCGYKDPRCEVGLIVGTGTNACYMEEL---QNVAGVD-------------
--GD------SG---------------------HMCINMEWGAFGDDGT-LGVLSTHFDASVDGASINPG
RQR-FEKLISGMYLGEIVCHILLHLTRLGVLFQGQQTQQLQARGIFKTKFLSQIES--------------
------------DSPALRKVRDILQGLGL-SLTSDDALIVREVCQAVSRRAAQLCGAGVAAVVEKIRENR
GLEELTVTVGVDGTLYKLH-----------PHFSRLVEATVRDLAPRCKVTF---------RQSEDG---
----------SGKGAALVTAVACRLAQ--TPR--I-----------------
>XP_006884497|Elephantulus edwardii|HK3
LQQFMVTEAQLQQIQASLLGSMEQEL-MGRSSPAPT--VRMLPTYVGSTPHGTEKGDFVVLELGATGASL
RVLCVTLTGT---EG-QR--VEPRSREFVIPSEVMLGPGQQ---------------LFDFAARCLAEFLE
AL-----PMG---DRHLPLGFNFSFPCHQTGLD-----------------------K------------
-----------STLISWTKGFRCSGVEGQDVVQLLREAIERQ----------G-----------------
--------------IPNLDVVAVVNDTVGTMMGCDPRLGPCEVGLVV-----------------------
-----D-----TGTNACYMEEVRHV-AVLDEDR--GRICISVEWGSFGDDEALGPVLTPFDKTLDQESLK
PGAQRFEKMIGGLYLGELVRLALAHLAQCGLLFGGSTSPALMSQGSVTLEHMAAMED-------------
--PS--AG--RALAILES-LGLSPRASDVELVQRVCEAVSTRAARLCAASLAAVLGRLQHSR-EQHTLQV
SVATGGRVFEQR-PRFLSILQETVKLLTPGCDVSFVPSVDGGGQGVAMVTAVAARLAAHRRLLAETLAPF
QLDRAQLAAVQAQMREAMARGLRGEA---S-SLRMLPTYVRATPDGS-----------------------
---ERGDFLALDLGGTNFRVLLVRVASD---GVTISSQVYAIPEQVTQGSGQQ-------LFDHIVDCIV
DFQQRQGLSGQ-SLPLGFTFSFPCRQLGLDQGILLNWTKGFKASDCEGQDVVTLLREAIARRQ----AVE
LNVVA-----IVNDTVGTMMSCGYEDHRCEVGLIVGTGTNACYMEEL---QNVAAVP-------------
--GE------SG---------------------HMCINMEWGAFGDDGV-LDVPRTHFDALVDEASINPG
KQR-FEKLISGMYLGEIVRHILLHLNRLGVLFRGQQTQCLQTRDIFKTKFLSEIES--------------
------------DSLALRQVRAILEDLGL-PLTSDDALVVREVCQVVSRRAAQLCGAGVAAVVEKIRDNR
GLKEFTVTVGVDGTLYKLH-----------PHFASLVAATVRELAPCCRVTF---------LQSEDG---
----------SGKGAALVTAVACRLAQ--TTR--V-----------------
>XP_004697201|Echinops telfairi|HK3
LKQFEVTEAQLRQIQASLLGSMEQAL-LGQNGPAPA--VRMLPAYVGSTPHGTEQGDFVVLELGATGASL
RVLWVTLTGT---EG-SR--VEPRSREFVIPPDVMLGPGQQ---------------LFDFAARCLAEFLE
AL-----PVH---SRTLPLGFNFSFPCHQTGLD-----------------------Q------------
-----------STLISWTKGFKCSDVEGQDVVRLLREALKRQ----------G-----------------
--------------VTSLSVVAVVNDTVGTMMGCDPGLGPCEVGLVV-----------------------
-----D-----TGTNACYMEEARHV-AALDEDR--GRVCVSVEWGSFGDDRALGLVLTPFDHALDLESLN
PGAQRFEKLIGGLYLGELVRLVLAHLAQHGLLFGGCTPPALLSRGTLGLEHVAELQD-------------
--PS--AGTARVLTVLQS-LGLSPGASDVGLVQSVCMAVSTRAARLCATALAAVLGRLQHSR-EQQALQL
SVATGGRVLARL-PRFRSLLQETVMLLAPECSVTFIPSEDGGGRGVAVVTAVAARLAAHRCLLEETLAPF
RLDQEQLVVVQAQMREAMAQGLRGEA---S-SLRMLPTYVRATPDGS-----------------------
---ERGDFLALDLGGTHFRVLLVRVAAG---GVQITSQVYAIPEQVAQGTGRQ-------LFDHIVDCIV
DFQQRQGLSGQ-SLPLGFTFSFPCRQLGLDQGILLTWTKGFNASDCEGQDVVTLLREAIGRRQ----AVE
LNVVA-----IVNDTVGTMMSCGYEDHRCEVGLIVGTGTNACYMEEL---QNVAGVP-------------
--GD------SG---------------------RMCINMEWGAFGDDGR-LSVPSTCFDASVDRASINPG
KQR-FEKLISGMYLGEIVRHILLHLNSLGVLFRGQQTQRLQSRDIFKTKFLSEIES--------------
------------DSLALRQVRAILEDLGL-PLTSDDALVVREVCQAVSRRAAQLCGAGVAAVVEKIRENR
GLEELAVTVGVDGTLYKLH-----------PHFSSLVADTVRALAPRCAVTF---------LQSEDG---
----------SGKGAALVTAVACRLAQ--MAR--V-----------------

>XP_006873553|Chrysochloris asiatica|HK3
LQQFMVTEAQLQQIRASLLSSMEQGL--MGQNPASA--VRMLPTYVGSTPHGTEQGDFVVLELGATGASM
RVLWVTLTGI---EG-CR--VDPRSREFVIPPEVMLGPGQQ---------------LFDFAARCLAEFLE
AF-----PVK---NRCLPLGFNFSFPCHQTGLD------------------------R------------
-----------STLISWTKGFRCSGVEGRDVVQLLREAIERQ----------G-----------------
--------------ASSLAVVAVVNDTVGTMMDCDPGLGPCEVGLVV-----------------------
-----D-----TGTNACYMEEARHV-ASLDEDR--GRVCVSVEWGSFSDDKALGPVLTPFDKALDQGSLN
PGAQRFEKMIGGLYLGELVRLVLAHLARRGLLFGGCTSPALLNQGSVGLEHVAEMED-------------
--PS--SGEARVLAVLQN-LGLSPGASDVKLVQLVCEAVSTRAARLCAAALAAVLGRLQHSR-EQHALQI
AVATGGQVFKRH-PRFHSLLQKTVILLAPGCDISFIPSVDGGGRGVAVVTAVAARLADHRRLLEETLAPF
RLDREQLAAVQVQMREAMAKGLQGEA---S-SLRMLPTYVRATPDGS-----------------------
---ERGDFLALDLGGTNFRVLLVRVTTG---GVQITSQVYAIPERVTQGSGQQ-------LFDHIVDCIV
DFQQRQGLGGQ-SLPLGFTFSFPCRQLGLDQGILLNWTKGFNASDCEGQDVVALLREAIGRRQ----AVE
LNVVA-----IVNDTVGTMMSCGYEDHHCEVGLIVGTGTNACYMEEL---QNVAAVA-------------
--GD------SG---------------------HMCINMEWGAFGDDGT-LSVPSTCFDASVDRASINPG
KQR-FEKLISGMYLGEIVRYILLHLNSLGVLFRGQQTQCLQTRDIFKTKFLSEIES--------------
------------DSLALRQVRAILEDLGL-PLTSDDALIVREVCQAVSRRAAQLCGAGVAAVVEKIRENR
GLEELMVTVGVDGTLYKLH-----------PHFSSLVAATVKDLAPRCTVTF---------LQSEDG---
----------SGKGAALVTAVACRLAQ--TTR--V-----------------
>XP_007951171|Orycteropus afer afer|HK3
LQQFMVTKAQLQQIQDSLLDSMEQAL-MGHSGPMPA--VRMLPTYVGSTPHGTEQGDFVVLELGATGASM
RVLWVTLTGT---KG-RR--VQPRSQEFVIPPEVMLGPGQQ---------------LFDFAAHCLAEFLE
AL-----PVG---NQCLPLGFNFSFPCHQTGLD------------------------R------------
-----------STLISWTKGFRCSDVEGQDVVQLLREAIKRQ----------G-----------------
--------------TSNLDVVAMVNDTVGTMMGCDPELGPCEIGLVV-----------------------
-----D-----TGTNACYMEEARHV-AALDEDR--GRVCVSIEWGSFGDDEVLGPLLTPFDRALDQESLN
PGAQRFEKMIGGLYLGELVRLVLAHLARRGILFGGCTTPALLSQGSVTLEHVAEMED-------------
--PS--TGAARVLAVLQN-LGLRPGPSDVELIQHVCAAVSTRSARLCAAALAAVLSRLQHSR-EQHTLQI
AVATGGQVFERH-PRFHNVLQETVMLLAPECDISFIPSVDGGGQGVAMVTAVAARLAAHRHLVEETLAPF
QLDQERLASVQVQMREAMAKGLRGEA---S-SLSMLPSYVRATPDGS-----------------------
---ERGDFLALDLGGTNFRVLLVRVAAA---GVKITSQVYAIPECVAQGSGQQ-------LFDHIVDCIM
DFQQRQGLGGQ-SLPLGFTFSFPCKQVGLDQGILLNWTKGFNASDCEGHDVVALLREAIGRRQ----AVE
LKVVA-----IVNDTVGTMMSCGYEDHRCEVGLIVGTGTNACYMEEL---QNVAGVP-------------
--GD------SG---------------------HMCINMEWGAFGDDGM-LDVPITCFDASVDRESINPG
KQR-FEKLISGMYLGEIVRHILLHLNSLGVLFRGQQTQCLQTRDIFKTKFLSEIES--------------
------------DSLALRQVRAILEDLGL-PLTSDDALVVREVCQAVSRRAAQLCGVGVAAVVEKIRENR
SLEELTVTVGVDGTLYKLH-----------PHFSSLVAATVRELAPRCAVTF---------LQSEDG---
----------SGKGAALVTAVACRLAQ--AAQ--V-----------------
>XP_040819904|Ochotona curzoniae|HK3
LQQFQVTNAQLQHIQASLLGCMEQAL-SNQAGPAPA--VRMLPTYVASTPHGTEKGDFLVLELGATGASL
RVLWVTLTGM---DG-HR--VEPRSQEFVIPKEVMLGTGQE---------------LFDFAARCLSEFLD
AN-----PVG---KHKMQLGFNFPFPCRQTSLD------------------------K------------
-----------STLISWTKGFQCSGVEGEDVVQLLRNAIQRQ----------G-----------------
--------------TYNIEVIAVVNDTVGTMMGCEPGDRPCEVGLVV-----------------------
-----G-----MGTNACYMEEARHV-AVLDEDR--GRMCVSVEWGSFNDNEALGPVLTTFDHTLDQESLN
PGAQRFEKMFGGLYLGELVRLVLVHLTQQGVLFGGCTSPALQSPDSIRAGHVADMED-------------
--AS--SGVARAFSILKD-LGLSPAASDAELVHCVCAAVSMRAARLCAAALAAILSRLQRSR-QQQTLQV
TVATGGQVFEQH-SRFRRILQDTIVPLVPGCEVSFVPSVDGGGRGVAVVTAVAARLAAHRHLLKETLMPF
RLEHKQLVQVQTQMREAMVRGLRGEA---S-SLRMLPTYVRATPDGN-----------------------
---ERGDFLALDLGGTNFRVLLVHVGDG---GVRIINQIYQVPEHVAQGSGQQ-------LFDHIVDCIV
DFQRKQGLSGR-SLPLGFTFSFPCRQLGLDQGILLNWTKGFNASDCEGKDVVCLLREAIRRRQ----GVE
LNVVA-----IVNDTVGTMMSCGYEDPRCEMGLIVGTGTNACYMEEL---QNVASVA-------------

```
--GD------SG---------------------RMCINMEWGAFGDDGS-LNSLNTCFDRSVDQESINPN
KQR-FEKMISGMYLGEIVRHILLYLTSRGVLFRGKQTQRLQTKDIFKTKFLSDIES--------------
------------DSLALRQVRTILEDLGL-PVTSDDTLMVLEVCQAVSRRAAQLCGAGVAAVVEKIRENR
NLQELTISVGVDGTLYKLH-----------PRFSSLVAATVQQLAPRCKVTF---------LESEDG---
----------SGKGAALVTAVACRLAE--NTC--V-----------------
>XP_057643111|Chionomys nivalis|HK3
LNQFKVTRTQLQQIQASLLCSMEQAL-KRQDSPAPS--VRMLPTYIGSMPNGTEQGDFLVLELGTKGASL
RVLWVTLTDT---KE-QR--VEPKSQEFVIPQEVMRGTGRQ---------------LFDFAAHCLSEFLD
AH-----TLK---NQDLKLGFNFPFPCHQTGLD------------------------R------------
-----------STLISWTRGFRCSGVEGQDVVQLLREAIQRQ----------K-----------------
--------------AHSIDVVAMVNDTMGIMMDCKCGSKSCEVGLIV-----------------------
-----D-----TGTNACYMEEARHV-KALDENR--GRTCISIEWGSFCDEEALGPVLTTFDHALDCESLA
PGAQRFEKMIGGLYLGELVRLVLVHFAQHGILFGGCTSPALLSQGSILLEHVIKMED-------------
--PA--TGKAQILAVLQD-LGLSPQVSDVEHVQHVCAAVCTRAAQLCAAALAAVLSHLQRSR-EQQTLHV
AVATGGRVLEKH-PRFHHTLKEMVMLLVPECDISFIPSVDDGGRGVAMVTAVAARLAAHKRILEDTLAPF
QLSFEQLKMVQAQMRTAMIKGLQGEA---S-SLRMLPTYVRATPDGS-----------------------
---ERGDFLALDLGGTNFRVLLVRVVEG---SVQITNQVYSIPEHTAQGSGEQ-------LFDHIVDCIV
DFQQSQGLSGH-SLPLGFTFSFPCKQLGLDQAILLNWTKGFSASGCEGQDVVNLLREAIRRKQ----GVE
LNVVA-----IVNDTVGTMMSCGYDDPRCEIGLIVGTGTNACYMEEL---QNVASVD-------------
--GN------SG---------------------RMCINMEWGAFGDDGT-LGMFLTRFDDSVDQASINPG
KQR-FEKMISGMYLGEIVRHILLHLTSLGALFRGQMTQSLQTRDIFKTKFLSEIES--------------
------------DGLALRQVRAILEDLGL-PLTSDDALMVLEVCQAVSRRAAQLCGAGVAAVVEKMRENR
GLQELTVSVGVDGTLYKLH-----------PHFSKLVSATVQKLAPRCKVTF---------LQSEDG---
----------SGKGAALVTAVACRLAQ--KAS--V-----------------
>XP_060053196|Erinaceus europaeus|HK3
LQLFTVTDAQLQQIQAALLASMEQAL-QDPEGPASA--VRMLPTYVGSIPHGTEQGDFVVLELGATGASL
RVLWVTLTGT---EG-PR--LEPRSQEFVIPQEVMLGPGQQ---------------LFDFAARCLCEFLD
TL-----PLC---QQPLQLGFSFSFPCHQTGLD------------------------K------------
-----------STLISWTKGFWCSGMEGQDVVQMLRDAIRKQ----------G-----------------
--------------AYSIDVVAVVNDTVGTMMGGQLGPGPCEVGLVV-----------------------
-----D-----TGTNACYMEEARHV-AVLDEDR--GRVCVSVEWGSFHDAGALGPLLTTFDHVLDHGSLN
PGAQRFEKMIGGLYLGELVRLVLAHLTERRALFGGCPSPALLCPGSILLEHLAAMED-------------
--PS--SGEAGAHVLLQD-LGLSTGMVDAKLVQQVGWAVSTRAARLCAAALSAVLARLQRGR-ELQTLRV
AVATGGQVFERL-PRFLSVLRETVQLLAPQCEVSFTPSEDGGGRGVAVVTAVAARLAAHRRLLEETLAPF
RLGHRELEAVQAQMRGAMSRGLHGEA---S-SLRMLPTYVRATPDGS-----------------------
---ERGDFLALDLGGTNFRVLLVRVAAQ---GVRITNQVYSIPEGVTQGSGQQ-------LFDHIVDCIV
DFQRKQGLSGQ-SLPLGFTFSFPCQQLGLDQGILLNWTKGFKASDCEGQDVVKLLREAIHRRQ----AAK
LNVVA-----IVNDTVGTMMSCGYEDSRCEVGLIVGTGTNACYMEEL---RNVAAVA-------------
--GD------SG---------------------HMCINMEWGAFGDDGS-LDALSTSFDASVDQASINPG
KQR-FEKMISGMYLGEIVRHILLHLTQLGVLFRGQQTQSLQARDIFKTKFLSEIES--------------
------------DSLALRQVRAILEDLGL-PLTSDDALMVLEVCQAVSQRAARLCGAGVAAVVEKIRENR
GLEELSVSVGVDGTLYKLH-----------PHFSSLVAATVRELAPRCTVTF---------LQSEDG---
----------SGKGAALVTAVACRLAQ--STR--V-----------------
>XP_035146538|Callithrix jacchus|HK3
LQQFKVTVAQLQQIQASLLGSMEQAL-RGQASPFPV--VRMLPTYVGSTPHGTEQGDFVVLELGATGASL
RVLWVTLTGI---EG-HR--VEPRSQEFVIPQDVMLGAGQQ---------------LFDFAAHCLSEFLD
AQ-----PVS---KQCLQLGFSFSFPCHQTGLD------------------------R------------
-----------STLISWTKGFRCSGVEGQDVVQLLRDAIQRQ----------G-----------------
--------------AYNIDVVAVVNDTVGTMMGCELGARPCEVGLVV-----------------------
-----D-----TGTNVCYMEEARHV-AVLDEDR--GRVCVSVEWGSFSDDGALGPVLTTFDHTLDHESLN
PGAQRFEKMIGGLYLGELVRLVLAHLTRRGVLFRGCPSPALLSQGSILLEHVAEMED-------------
--PS--AGAARVHAILQD-LGLSPEASDVELVQRVCAAVCMRAAQLCAAALAAVLSRLQHSR-EQQTLQV
```

TVATGGRVCERH-PRFCSILQGTVMLLAPECDVSFIPSADMGGRGVAMVTAVAARLAAHWRLLEETLAPF
RLNHDQLAAVQAQMREAMAKGLRGEA---S-SLRMLPTYVRATPDGS-----------------------
---ERGDFLALDLGGTNFRVLLVRVTT----CVQITSQIYSIPESVAQGSGQQ-------LFDHIVDCIV
DFQQKQGLSGQ-SLPLGFTFSFPCRQLGLDQGILLNWTKGFNASDCEGQDVVSLLREAIMRRQ----ELA
PMLATWRSSGMWQECLGTQATCASTWSGVPLGMMALWTRSAPALTQVWTRRPSTLAS-------------
----------RG--------------------------VGWAQMVVAGC--DQCWGQWPVGPFAMCAHTC
SRR-FEKMISGMYLGEIVRHILLHLTSLGVLFRGQQTQCLQTRDIFKTKFLSEIES--------------
------------DSLALRQVRAILEDLGL-PLTSDDALMVLEVCQAVSQRAAQLCGAGVAAVVEKIRENR
GLEELAVSVGVDGTLYKLH-----------PHFSSLVAATVRELAPRCAVTF---------LQSEDG---
----------SGKGAALVTAVACRLAQ--STH--I-----------------
>XP_063086899|Cavia porcellus|HK3
LQQFKVTKTQLQQIQASLLGSMEQAL-TGGASPASG--VRMLPTYVGSTPHGNEQGNFVVLELGASGASL
RVLWVTLTGI---KG-QR--VEPKSQEFVIPQEVVLGTGQQ---------------LFDFAARCLSEFLD
AH-----SVG---SQGLQLGFNFSFPCHQTSLD------------------------R------------
-----------STLISWTKGFRCSGVEGQDVVQLLREAIQRQ----------G-----------------
--------------GYSIEVVAVVNDTVGTMMGCEPAVGPCEIGLVV-----------------------
-----D-----TGTNACYMEEARHV-AVLDEDR--GRTCISIEWGSFNDDRALGPVLTTFDHALDHASLN
PGAQRFEKMIGGLYLGELVRLVLVHLAQNRVLFGGYTSPALLSQGSILLEHVTEMED------------
--PS--TGVARVHAILQD-LGLSPETSDAELVQRVCAAVCSRAAQLCAAALVAILSHLQHSR-EQQTLQV
AVATGGRVLERH-PRFQSILQETVTLLASECDVSFVPSVDGGGRGVAMVTAVAARLAYHRRLLEKTLAPF
RLRCEQLAAVRTQMRDAMAKGLRGEP---S-SLRMLPTYIRATPDGS-----------------------
---ERGDFLALDLGGTNFRVLLVRVTEG---GVQIINQIYSIPESVAQGSGQK-------LFDHIVDCIV
DFQEKQGLRGQ-SLPLGFTFSFPCKQLGLDQGILLNWTKGFNASGCEGQDVVYLLREAIRRKQ----AME
LNVVA-----IVNDTVGTMMSCGYEDPRCEMGLIVGTGTNACYMEEL---RNVAGVP------------
--GD------SG---------------------HMCINMEWGAFGDDGS-LGMLSTCFDEMVDQASINPG
KQR-FEKMISGMYLGEIVRHILLHLTSLGVLFRGQQTQRLQTRDIFKTKFLSEIES--------------
------------DSLALRQVRAILEDLGL-SLTSDDALIVLEVCQAVSRRAAQLCGAGVAAVVEKIRENR
GLEELTVSVGVDGTLYKLCQSCGCHSTGAGPSLCGHLLAVRRWVWERCSFGHCCCLSPCPCLRKSPGITV
SQPPPCWTRAGPRYPGLLKQPLCDLSV--AIC--PTRWFSPREVAQGLAIYI
>XP_053769214|Desmodus rotundus|HK3
LDQFKLTEAQLRQIQARILGSMELAL-RGEASPVPA--VRMLPAYVGSIPHGTEQGDFLVLELGATGASL
RVLWVTLTGT---EE-PQ--IEPRSQEFLIPQELMLGPGQK---------------LFDFAAHCLSEFLD
AC-----PVG---DRTLQLGFNFSFPCHQTGLD------------------------R------------
-----------STLISWTKGFRCSGVEGQDVVQLLREAIERQ----------G-----------------
--------------AYSINVAAVVNDTVGTMMGCGPGSAPCEVGLVV-----------------------
-----D-----TGTNACYMEEARHV-AVLDEDR--GRVCVSIEWGSFGDDGALGPVLTTFDRILDQESLN
PGAQRFEKMIGGLYLGELVRLVLAHLAQHGVLFGGRTSPALLSQGGVLLEHVAEMQD------------
--PS--AGTARVQAVLQG-LGLSPGTTDAELVQQVCAAVCTRAARLCAAALAAVLTRLQHGR-EQQTLQI
AVATGGRVFEQH-PRFLSVLRETVKLLAPKCDVSFTPSVDGGGRGVAMVTAVAARVAAHRRMLEETLAPF
RLNREQLAQVQMQMREAMAKGLRGEA---S-SLRMLPTYVRATPDGS-----------------------
---ERGDFLALDLGGTNFRVLLVRVAAG---GVQITSQIYSIPERVAQGPGQQ-------LFDHIVDCIL
DFQQKQGLSGQ-SLPLGFTFSFPCRQVGLDQGILLTWTKGFNASDCEGQDVVGLLRQAIGRRQ----AVK
LNVVA-----IVNDTVGTMMSCGYEDPRCEVGLIIGTGTNACYMEEL---RNVAGVA------------
--GD------TG---------------------HMCINMEWGAFGDDGS-LDMLTTCFDKLVDQASINPG
RQR-FEKMISGMYLGEIVRHILLHLTSLGVLFRGQKTQRLQTRDIFKTKFLSEIES--------------
------------DSLALRQVRAILEDLGL-PLTSDDALTVLEVCQAVSQRAAQLCGAGVAAVVEKIRENR
GLDELTVSVGVDGTLYKLH-----------PHFSGLVAATVRQLAPRCAVTF---------LQSEDG---
----------SGKGAALVTAVACRLAQ--PAS--V-----------------
>XP_042523024|Dipodomys spectabilis|HK3
LQQFKVTRTQLQQIQASLLGSMEQAL-RGQAGPAPS--VRMLPTYVGSTPHGTEQGDFLVLELGATGASL
RVLWVTLTGT---EG-HR--VEPRTQEFAIPQEVVLGPGQQ---------------LFDFAARCLSEFLD
TQ-----PVG---SQRLQLGFSFSFPCHQTGLD------------------------R------------

-----------SILISWTKGFRCSGVEGQDVVQLLRDAIRRQ----------G-----------------
--------------TYSIDVVAVVNDTVGTMMGCELGAGPCEIGLIV-----------------------
-----D-----TGTNACYMEEARHV-AVLDEDR--GRSCVSVEWGSFDDHAALGPVLTTFDLTLDRESLN
PGAQRFEKMIGGLYLGELVRLVLLHLAQRGVLFGGSTSPALLTPGSIDLEHVACMED-------------
--PS--TGAACAQDILKG-LGLSPRARDAELVQRVCTAVSTRAAQLCAAALAAVLVRLQRSR-EQQELRV
AVAVGGRVFEQH-SRFHGILRDMVVLLAPECDVSFVPSVDGGGRGVAMVTAVAARLAAQRHLLEETLAPF
RLSREQLVAVQAQMREAMARGLKGEA---S-SLRMLPTYVRATPDGS-----------------------
---ERGDFLALDLGGTNFRVLLVRVTEA---GVQITSHIYSIPECVAQGSGQQ-------LFDHIVDCIV
DFQQQQGLSGR-SLPLGFTFSFPCKQLGLDQGILLNWTKGFNASDCEGRDVVCLLREAIGRRQ----AVE
LNVVA-----IVNDTVGTMMSCGYEDPRCEMGLIVGTGTNACYMEEL---QNVPGVP-------------
--GD------QG---------------------RMCINMEWGAFGDDGS-LGMLGTCFDERVDQASINPG
KQR-FEKMISGMYLGEIVRHILLHLTSLGVLFRGQVTQCLQTRDIFKTKFLSEIES--------------
------------DSLALRQVRAILEALGL-PLTSDDALMVLEVCQAVSRRAAQLCGAGVAAVVEKIRENR
GLDELAVSVGVDGTLYKLH-----------PHFSSLVAATVRELAPRCAVTF---------LQSEDG---
----------SGKGAALVTAVACRLAQ--EAQ--V-----------------
>XP_004665026|Jaculus jaculus|HK3
LKQFKVTRSQLQQIQVSLLGCMERAL-KGKADSAPA--VRMLPTYVGSVPHGTEKGDFLVLELGATGASL
RVLWVTLTGT---EG-YK--VEPRSQEFVIPQDVVVGAGQQ---------------LFDFAAHCLSEFLD
AQ-----PVG---NQPLQLGFSFSFPCHQTSLD------------------------R------------
-----------STLISWTKGFKCSGVEGQDVVQLLRDAIQRK----------G-----------------
--------------TYNIDVVAVVNDTVGTMMGCELGARPCEVGLIV-----------------------
-----D-----TGTNACYMEEARHV-AALDEDR--GRTCVSVEWGSFSDDGALAPVLTTFDHVLDQGSLN
PGAQRFEKMIGGLYLGELVRLVLAHLAQHGALFGGHTSPALLSPGSLLLEHVAEMED-------------
--AC--NGAAHTHTVLQG-LGVSPDASDAELVQRVCAAVCSRAARLCAAALAAVLARLQHSR-EQRALQV
AVATGGRVFERH-PRFHNILKETVMHLAPECEVSFIPSVDGGGRGVAMVTAVAARLAAHRQLLEETLAPF
QLNHEQLRAVQAQMREAMVRGLRGEA---S-SLRMLPTYVRATPDGS-----------------------
---ERGDFLALDLGGTNFRVLVVHVTEG---GVQITNQVYSIPEGVAQGSGQQ-------LFDHIVDCIV
DFQQRQGLSGQ-SLPLGFTFSFPCKQLGLDQGILLNWTKGFNASDCEGQDVVCLLREAIRRRQ----AVE
LNVVA-----IVNDTVGTMMSCGYEDPRCEMGLIVGTGTNACYMEEL---RNVTSVA-------------
--GD------SG---------------------HMCINMEWGAFGDDGS-LGLLSTCFDAIVDQASINPG
KQR-FEKMISGMYLGEIVRHILLHLTSCGVLFRGRQAQCLQTKDIFKTKFLSEIES--------------
------------DSLALRQVRAILEDLGL-LLTSDDALMVREVCQAVSRRAARLCGAGVAAVVEKIRENR
GLEELTVSVGVDGTLYKMH-----------PHFSSLVATTVRELAPRCAVSF---------LQSEDG---
----------SGKGAALVTAVACRLAQ--TAQ--V-----------------
>XP_037355781|Talpa occidentalis|HK3
LQKFKVTEAQLRQIQASLLASMEQAL-KGQASSASA--VRMLPTFVGSTPHGTEQGDFVVLELGATGASL
RVLWVTLMGT---AG-PR--MEPQSQEFVIPQEVMLGPGEQ---------------LFDFASRCLSEFLD
ML-----PLD---KQSLQLGFNFPFPCHQTGLD------------------------K------------
-----------STLISWTKGFKCSGVEGQDVVQMLREAIQRQ----------G-----------------
--------------AYSIDVVAVVNDTVGTMMGCEPGVGPCEIGLVV-----------------------
-----D-----TGTNACYMEEARHV-AVLDEDR--GRVCVSVEWGSFSDDKALGPVLTTFDHALDHQSLN
PGAQRFEKMIGGLYLGELVRLVLVHLVERGVLFGGCVSSSLLSQGSILLQHVAEMES-------------
--FS--AGAARVQAVLQD-LGLSVGASDAKLVQQVCVAVCTRAARLCAAALAAVLTRLQHSR-EQQTLQI
AVATGGGVFERH-PRFLSILQDTVMLLAPECDVSFIPTVDGGGRGVAMVTAVAARLAAHRRLLEETLAPF
RLSQQQLAAVQTQMREAMAKGLRGEV---S-SLRMLPTYVRATPDGS-----------------------
---ERGDFLALDLGGTNFRVLLVRVATG---GVQITNQVYSIPESVAQGSGQQ-------LFDHIVDCIV
DFQQKQGLSGQ-SLPLGFTFSFPCKQIGLDQGILLNWTKGFKASDCEGQDVVCLLREAIGRKQ----GVE
LNVVA-----IVNDTVGTMMSCGYEDPCCEVGLIVGTGTNACYMEEL---QNVASVP-------------
--GD------SG---------------------NMCINMEWGAFGDDGS-LDMLSTCFDARVDQASINPG
KQR-FEKMISGMYLGEIVRHVLLHLTSLGVLFRGQQTQHLQTRDIFKTKFLSEIES--------------
------------DSLALRQVRAILEDLGL-PLNTDDALMVLEVCQAVSQRAAQLCGAGVAAVVEKIRENR
GLEELTVSVGVDGTLYKLH-----------PHFSTLVAATVQELAPRCVVTF---------LQSEDG---

----------SGKGAALVTAVACRLAQ--KTH--V-----------------
>XP_016052623|Miniopterus natalensis|HK3
LNQFKVTGTQLRQIQASLLGSMEQAL-RGQVSPAPV--VRMLPTYVGSIPHGTEQGDFLVLELGATEASL
RVLWVTLTGI---EG-HR--IEPRSQEFLIPQEVMLGPGQQ---------------LFDFAARCLSEFLD
AL-----PVG---NKGLQLGFNFSFPCHQTGLD------------------------R------------
-----------STLISWTKGFKCSGVEGQDVVQLLRDAIKRQ----------G-----------------
--------------TYSINVVAVVNDTVGTMMGCEPGTEPCEVGLIV-----------------------
-----D-----TGTNACYMEEARHV-AALDEDR--GRVCVSIEWGSFGDDAALGPVLTTFDHILDQESLN
PGAQRFEKMIGGLYLGELVRQVLAHLAQQGILFGGCTSPALLSRGCILLEHVAQMED-------------
--PS--AGTAHVHAILQD-LGLNPGASDAELVQHVCAAVCARAARLCAAALAAVLTRLQHSR-EQQTLHV
AVAIGGRVFERH-PRFLSILQETVKLLVPECDVSFIPSVDGGGRGVAMVTAVAARLAAHRCLLEETLAPF
RLNREQLAMVQAQMREAMAKGLRGEA---S-SLRMLPTYVRATPDGS-----------------------
---ERGNFLALDLGGTNFRVLLVHVATE---DVQITSQIYSIPECVAQGSGQQ-------LFDHIVDCIL
DFQQKQGLSGQ-SLPLGFTFSFPCRQLGLDQGILLNWTKGFSASGCEGQDVVRLLREALGRRQ----AME
LNVVA-----IVNDTVGTMMSCGYKDPHCEVGLIVGTGTNACYMEEL---RNVEGVA-------------
--GD------TG---------------------HMCINMEWGAFGDDGS-LDMLSTCFDASVDQATINPG
KQR-FEKMISGMYLGEIVRHVLLHLTSLGVLFRGQLSQCLQTRDIFKTKFLSEIES--------------
------------DSLALRQVRAILEDLGL-SLTSDDALMVLEVCQAVSQRAAQLCGAGVAAVVEKIRENR
GLEELTVSVGVDGTLYKLH-----------PHFSRLVAATVQELAPRCVVTF---------LQSEDG---
----------SGKGAALVTAVACRLAQ--LAH--V-----------------
>XP_033617541|Fukomys damarensis|HK3
LQRFKVTRTQLQHIQASLLDSMEQAL-TGKASSAPG--VQMLPTYVDSTPHGNEQGNFVVLELGASGASL
RVLWVTLTGI---KG-QR--VEPKSQEFVIPQEVVLGTGQQ---------------LFDFAARCLFEFLD
AH-----SVR---NQGLQLGFSFSFPCHQTGLD------------------------R------------
-----------STLISWTKGFRCSGVEGQDVVQLLRDAIQRQ----------G-----------------
--------------AYNIDVVAVVNDTVGTMMSCEPGVRPCEVGLIV-----------------------
-----D-----TGTNACYMEEARHV-AVLDEDR--GRTCVSVEWGSFSDDGALGPVLTTFDHALDHASLN
PGAQRFEKMIGGLYLGELVRLVLAHLAEHRVLFGGYTSSALLSQGSILLEHVAEMED-------------
--PS--TGAARVHAILQD-LGLSPEASDAELVQRVCAAVCSRAAQLCAAALAAILSRLQRSR-EQQILQV
AVATGGRVLEQH-PRFYSILQETVKLLASECDVSFIPSVDGGGRGVAMVTAVAARLAAHRRLLEETLAPF
RLSREQLAAVQTQMREAMAKGLRREA---S-SLRMLPTYVRATPDGS-----------------------
---ERGDFLALDLGGTNFRVLLVRVAEG---GVRIVNQVYAIPECVAQGSGQQ-------LFDHIVDCIM
DFQEKQGLRGQ-SLPLGFTFSFPCKQLGLDQGILLNWTKGFNASGCEGQDVVYLLREAIKRKQ----AVE
LNVVA-----IVNDTVGTMMSCGYEDPRCEMGLIVGTGTNACYMEEL---QNVAGVA-------------
--GD------SG---------------------HMCINMEWGAFGDDGS-LGMLSTCFDVRVDQVSINPG
KQR-FEKMISGMYLGEIVRHILLHLTSLGVLFQGQQTQRLQTRDIFKTKFLSEIES--------------
------------DSLALRQVRAILEDLGL-PLTSDDALIVLEVCQAVSRRAAQLCGAGVAAVVEKIRENR
GLEELTVSVGVDGTLYKVH-----------PHFASLVAATVRELAPRCMVTF---------LQSEDG---
----------SGKGAALVTAVACRLAQ--MAH--V-----------------
>XP_008253672|Oryctolagus cuniculus|HK3
LQQFQVTRAQLQQIQAGLLDSMEQAL-RGQANPAPA--VRMLPTYVGSTPHGTEQGDFLVLELGATGASL
RVLWVTLTGV---DG-HR--VEPRSQEFVIPQEVMLGTGQQ---------------LFDFAACCLSEFLD
AH-----PVG---NQGLQLGFSFSFPCHQTGLA------------------------R------------
-----------STLISWTKGFRCSGVEGQDVVQLLRDAIQRR----------G-----------------
--------------ASNIDVIAVVNDTVGTMMGCEPGVRPCEVGLVV-----------------------
-----D-----TGTNACYMEEARHV-AALDEDR--GRVCVSVEWGSFGDDGALGPLLTAFDHALDLESLN
PHAQRFEKMVGGLYLGELVRLVLSHLAQRGVLFGGSSSPALQSQGSILPEDVAAMED-------------
--PS--VGAAHVLTILQG-LGLSPEVSDVELVRSVCAAVSTRAARLCAAALAAVLARLQRSR-EQQMLQV
TVATGGRVFEWH-RRFHSILQETVALLVPECDVSFIPSVDGGGRGVAVVTAVAARLAAHRRLLEETLMPF
RLNCEQLAGVQAQMREAMAKGLRGEA---S-SLRMLPTYVRATPDGS-----------------------
---ERGDFLALDLGGTNFRVLLVRVTEG---GVQITNQVYSVPEHVAQGSGQQ-------LFDHIVDCIV
DFQRAQGLSGQ-SLPLGFTFSFPCKQLGLDQGILLNWTKGFNALDCEGRDVVCLLREAIRRRQ----AVE

LNVVA-----IVNDTVGTMMSCGYEDPRCEMGLIVGTGTNACYMEEL---RNVAGVA-------------
--GD------SG---------------------QMCINMEWGAFGDDGS-LGSLSTCFDDSVDQASINPG
KQR-FEKMISGMYLGEIVRHILLHLTSLGVLFRGQQTQRLQTRDIFKTKFLSDIES--------------
------------DSLALRQVRAILEDLGL-PVTSDDALMVLEVCQAVSRRAAQLCGAGVAAVVEKIRENR
GLEELTVSVGVDGTLYKLH-----------PHFSSLVAATVRQLAPRCTVTF---------LESEDG---
----------SGKGAALVTAVACRLAQ--TTR--V-----------------
> XP_055451751|Psammomys obesus|HK3
LQQFAVTRTQLQQIQASLLCSMEQAL-KGQDGPAPS--VRMLPTYVGSTPHGTEQGDFLVLELGATGASL
RVLWVTLTGT---QE-YH--VEPRSQEFEIPQEVMLGPGQQ---------------LFDFAARCLSEFLD
AH-----PVK---NQDLKLGFNFSFPCHQTGLD------------------------KVRWGHTGPARAK
RATLTLILYPQSTLISWTKGFRCSDVEGQDVVQLLRDAIQRQ----------G-----------------
--------------TYSIDVLAIVNDTVGTMMGCELGAGPCEVGLIV-----------------------
-----D-----TGTNACYMEEARHV-AALDEDR--GRTCVSVEWGSFCDEQALGPALTTFDRALDHESLT
PGAQRFEKMIGGLYLGELVRLVLVYLAQHGLLFGGCTSPALLSQGSILLEHVAELED-------------
--PT--TGTARAHTILQD-LGLSPQASDAQLVRRVCAAVCARAAQLCAAALAAVLSRLQHGR-EQQALRV
AVATGGRVLERQ-PRFQRTLKETVAFLAPECDVSFIPSVDGGGRGVAMVTAVAARLAAHRHILEETLAPF
RLTLEQLAVVQAQMREAMVRGLRGEA---S-SLRMLPTYVRSTPDGT-----------------------
---ERGDFLALDLGGTNFRVLLVRVAEG---SVQITNQVYSIPECVAQGSGQQ-------LFDHIVDCIV
DFQQRQGLSGQ-SLPLGFTFSFPCKQLGLDQGILLNWTKGFKASGCEGQDVVHLLREAIRRRQ----AME
LNVVA-----IVNDTVGTMMSCGYDDPRCEMGLIVGTGTNACYMEEL---QNVASVA-------------
--GH------SG---------------------LMCINMEWGAFGDHGS-LGTLSTCFDARVDQASINPG
KQR-FEKMISGMYLGEIVRHILLHLTSLGVLFRGRKSRCLQTRGIFKTKFLSEIES--------------
------------DSLALRQVRAILEDLGL-TLTSDDALMVLEVCQVVSRRAAQLCGAGVAAVVEKIRENR
GLQELTVSVGVDGTLYKLH-----------PHFSRLVSATVEKLAPHCTVTF---------LQSEDG---
----------SGKGAALVTAVACRLTP--MAV--SEGISKSGGLSLA-----
> XP_008060946|Carlito syrichta|HK3
LQPFKVTETQLQRIQAGLLGSMEQAL-RGQPSPAPA--VRMLPTYVGSTPNGTEQGDFVVLELGATGASL
RVLWVTLTGA---EG-QR--VEPRSQEFAIPQEVMLGAGPQ---------------LFDFAARCLSEFLE
AH-----PVS---SRSLQLGFSFSFPCHQTGLD------------------------R------------
-----------STLISWTKGFRCSGVEGQDVVQLLREAMRRQ----------G-----------------
--------------VHNIDVVAVVNDTVGTMMGCEPGARPCEVGLVV-----------------------
-----D-----TGTNACYMEEARHV-AVLDEDR--GRVCVSVEWGSFSDDGALGPVLTAFDRALDRASLS
PGAQRFEKMIGGLYLGELVRLVLAHLAAHGALFGGRTSPALLSAGSIHLEHVAEMEH-------------
--PS--TGAARVRAILQG-LGLSPEASDTELVQHVCAAVCTRAARLCAAALAAVLARLQCSR-EQRALQV
AVATGGRVCDRH-PRFCSVLRETVLFLAPECDVSFVPSVDGGGRGVAMVTAVAARLAAHRRLLEETLAPF
RLSREQLAVVQAQMREAMARGLRGEA---S-SLRMLPTYVRATPDGS-----------------------
---ERGDFLALDLGGTNFRVLLVRVAAG---GVQITSQVYSIPECVAQGSGQQ-------LFDHIVDCIV
DFQQKQGLSGQ-SLPLGFTFSFPCRQLGLDQGVLLNWTKGFNASGCEGQDVVSLLREAIARRQ----AVE
LNVVA-----IVNDTVGTMMSCGYEDPRCEMGLIVGTGTNACYMEEL---RNVAGVA-------------
--GD------TG---------------------HMCINMEWGAFGDDGS-LGVFSTRFDENVDRASINPG
KQR-FEKMISGMYLGEIVRHVLLHLTSLGVLFRGQQSRRLQTRDIFKTKFLSEIES--------------
------------DSLALRQVRAILEDLGL-PLTSDDALMVLEVCQAVSQRAARLCGAGVAAVVEQIRENR
GLAELTVSVGVDGTLYKLH-----------PHFSSLVAATVQELAPCCAVTF---------LQSEDG---
----------SGKGAALVTAVACRLSR--PTR--I-----------------
> XP_012642744|Microcebus murinus|HK3
LQQFKVTEAQLQQIQASLLVSMEQAL-RGEASPAPA--VRMLPTYVGSIPHGTEQGDFLVLELGAKGASL
RVLWVTLTGI---EG-HR--VEPRSQEFVIPQEVMLGAGQQ---------------LFDFAAHCLSEFLD
AH-----PVS---RQGLQLGLSFSFPCHQTGLD------------------------R------------
-----------STLISWTKGFRCSGVEGQDVVQLLRDAIQRQ----------G-----------------
--------------AHSVEVVAVVNDTVGTMMGCAPGLRPCEVGLVV-----------------------
-----D-----TGTNACYMEEARHV-AALDDDR--GRMCVSVEWGSFRDDAALGPLLTAFDRTLDRESLN
PGAQRFEKMIGGLYLGELVRLVLVHLARRGVLFGGCTSPALLSRGSVLLEHVAEMED-------------

--PS--AGAARVHAILQG-LGLSPDSSDAELVRHVCAAVCTRAAQLCAAALASVLSRMQRGR-EHRALRV
AVATGGRVCEQH-PRFCRILRETVKVLAPECDVSFVPSVDGGGRGVAMVTAVAARLASHRRLLEETLAPF
RLSREQLAVVQAQMREAMAKGLRGEA---S-SLRMLPTYVRATPDGS-----------------------
---ERGDFLALDLGGTNFRVLLVRVAAG---GVQITSQVYSIPECVAQGSGQQ-------LFDHIVDCIV
DFQQKQGLSGQ-SLPLGFTFSFPCRQLGLDQGILLNWTKGFNASDCEGQDIVRLLRAAIGRRQ----AVE
LNVVA-----IVNDTVGTMMSCGYEDPRCEIGLIVGTGTNACYMEEL---RNVAGLT-------------
--GG------PG---------------------RMCVNMEWGAFGDDGS-LDALSTCFDASVDRASINPG
KQR-FEKMISGMYLGEIVRHVLLRLTSLGVLFRGQQMGRLQTRDIFKTKFLSEIES--------------
------------DSLALRQVRAILEDLGL-PLTSDDALVVLEVCQAVSQRAARLCGAGVAAVAERMRESR
GLEELAVSVGVDGTLYKLH-----------PHFSSLVAATVRELAPRCAVTF---------LQSEDG---
----------SGKGAALVTAVACRLAQ--KTR--V-----------------
>XP_028627988|Grammomys surdaster|HK3
LQQFKVTRTQLQQIQASLLCSMEQAL-KGQDSPAPS--VRMLPTYVRSTPHGTEQGDFLVLELGATGASL
RVLWVTLTGT---KE-HS--MEPRSREFVIPQEVILGAGQQ---------------LFDFAACCLSEFLD
AY-----PVE---NQGLKLGFNFSFPCHQTGLD------------------------R------------
-----------STLISWTKGFRCSGVEGQDVVQLLRDAIQRQ----------G-----------------
--------------TYSIDVVAMVNDTVGTMMGCELGARPCEVGLIV-----------------------
-----D-----TGTNACYMEEARHV-AALDEDR--GRTCVSIEWGSFYDEEALGPVLTTFDYALDHESLI
PGAQRFEKMIGGLYLGELVRLVLVHLSQHGVLFGGCTSPALLSQGSILLEHVAEMED-------------
--PA--TGTAHVHTILQA-LGLSLQVSDGELVQCVCVAVCTRAAQLCAAALAAVLSRLQRSR-EQQTLRV
AVATGGRVFDQH-PRFLCILKETVKLLAPECDVSFIRSVDGGGQGVAMVTAVAARLAAHRRILEETLAPF
QLTLEQLTVVQAQMREAMIRGLQGDA---S-SLRMLPTYVRATPDGS-----------------------
---ERGDFLALDLGGTNFRVLLVRVAEG---SVRIINQIYSIPECIAQGSGQK-------LFDHIVDCIV
DFQKRQGLSGQ-SLPLGFTFSFPCKQLGLDQGILLNWTKGFNASGCEGQDVVYLLREAIRRRQ----AVE
LNVVA-----IVNDTVGTMMSCGYDDPRCEMGLIVGTGTNACYMEEL---RNVASVP-------------
--GD------SG---------------------RMCINMEWGAFGDDGS-LSMLSTCFDASVDQASINPG
KQR-FEKMISGMYLGEIVRHILLQLTNLGVLFRGQKTQCLQTRDIFKTKFLSEIES--------------
------------DSLALRQVRAILEDLGL-TLTSDDALMVLEVCQAVSRRAAHLCGAGVAAVVEKIRENR
GLRELTVSVGVDGTLYKLH-----------PHFSRLVSATVRKLAPQCTVTF---------LQSEDG---
----------SGKGAALVTAVACRLTQ--MAH--I-----------------
>XP_008845528|Nannospalax galili|HK3
LQQFKVTRKQLQQIQVSLLRSMEQAL-MEQNNPAPA--VRMLPTYVGSTPHGTEQGDFLVLELGATGASL
RVLWVTLTGT---EE-HR--VEPRSQEFVIPQEVVLGSGQQ---------------LFDFAACCLSEFLD
AH-----PVG---NQDLKLGFSFSFPCHQTGLD------------------------K------------
-----------STLISWTKGFKCSGVEGHDVVQLLRDAIQRQ----------G-----------------
--------------TYNIDVVAVVNDTVGTMMGCELGAGPCEVGLIV-----------------------
-----D-----TGTNACYMEEARHV-AALDEDR--GRTCVSIEWGSFNDDGALGPVMTIFDHALDYESLN
PGAQRFEKMIGGLYLGELVRLVLAHLAQHGVLFGGCTSPALLSRGSILLEHVVEMED-------------
--PF--TGAAHVHAVLQD-LDLSPQASDAELVQHVCAAVCTRAAQLCAAALAAVLSRLQHSR-EQQTLQV
AVATGGQVFEVH-PRFRSILKEMVMLLVPECDVSFIPAVDGGGQGVAMVTAVAARLAAHRHLLEETLAPF
QLNCQQLIVVQAQMREAMIRGLQGEA---S-SLRMLPTYVRATPDGS-----------------------
---ERGDFLALDLGGTNFRVLLVRVAKG---SVQITNQVYSIPECVAQGSGQE-------LFDHIVDCIV
DFQQRQGLSGQ-SLPLGFTFSFPCKQLGLDQGILLNWTKGFNASGCEGQDVVCLLREAIGRRQ----AVE
LNVVA-----IVNDTVGTMMSCGYDDPHCEMGLIVGTGTNACYMEEL---RNVASVA-------------
--GD------SG---------------------YMCINMEWGAFGDDGS-LSMLSTCFDASVDQASINPG
KQR-FEKMISGMYLGEIVRHILLHLTSLGVLFQGQQTQCLQTRDIFKTKFLSEIES--------------
------------DSLALRQVRAILEDLGL-TLTSDDALMVLEVCQAVSRRAAQLCGAGVAAVVEKIRENR
GLEELTVSVGVDGTLYKLH-----------PHFSSLVSATVRKLAPHCTVTF---------LQSEDG---
----------SGKGAALVTAVACRLAQ--TTR--V-----------------
>XP_015344007|Marmota marmota marmota|HK3
LQQFKVTRTQLQQIQASLLDSMEQAL-KGQASPAPA--VRMLPTYVGSTPHGTEQGDFLVLELGATGASL
RVLWVTLTGI---EG-HR--VEPRSQEFMIPEEVVLGTGQQ---------------LFDFAACCLSEFLD

AY-----PVG---NQGLQLGFSFSFPCHQTGLD------------------------K------------
-----------STLISWTKGFRCSGVEGQDVVQLLRDAIQRQ----------G-----------------
--------------AYSIDVVAVVNDTVGTMMGCEPGTRPCEIGLIV-----------------------
-----D-----TGTNACYMEEARHV-AVLDEDR--GRTCVSIEWGSFNDHGALGPVMTTFDHALDQESLN
PGAQRFEKMIGGLYLGELVRLALAHLARRGVLFGGCTSPALLSQGSIRLEHVAEMED------------
--PS--TGAARVHTILQD-LGLSLEASDAELVQRVCAAVCTRAAQLCAAALAAVLSRLQHSR-EQQTLQV
AVATGGQVFEQH-LRFHSILQETVMVLVPECDVSFIPSVDGGGRGVAMVTAVAARLAAHQRLLEETLAPF
RLNLEQLKAVQAQMREAMAKGLRGEA---S-SLRMLPTYVRATPDGS-----------------------
---ERGDFLALDLGGTNFRVLLVRLTEG---GVQIINQVYSIPECVAQGSGQQ-------LFDHIVDCIV
DFQQRQGMSGQ-SLPLGFTFSFPCKQVGLDQGILLNWTKGFNASDCEGHDVVHLLREAIKRRQ----AVE
LNVVA-----IVNDTVGTMMSCGYEDPRCEIGLIVGTGTNACYMEEL---PNVVDTA-------------
--GE------SG---------------------QMCINMEWGAFGDDGS-LGMLSTCFDESVDQASINPG
KQR-FEKMISGMYLGEIVRHILLHLTSLGVLFRGQQTQCLQTRDIFKTKFLSEIES--------------
------------DSLALRQVRAILEDLGL-PLTSDDALMVLEVCQAVSQRAAQLCGAGVAAVVEKIRENR
GLEELTISVGVDGTLYKLH-----------PHFSSLVAATVRELAPCCKVTF---------LQSEDG---
----------SGKGAALVTAVACRLAQ--KTR--V-----------------

>XP_003781951|Otolemur garnettii|HK3

LQQFKVTGAQLQQIQASLLASMEQAL-KERASPVPA--VQMLPTYVRSTPHGTEQGDFIVLELGAKGASL
RVLWVTLTGT---EG-HR--VVPRSQEFVIPQEVMLGPGQQ---------------LFDFAAHCLAEFLD
TH-----PVS---SQSLQLGFSFSFPCHQTGLD------------------------R------------
-----------STLISWTKGFRCSGVEGQDVVQLLRDAIQRL----------G-----------------
--------------AYNIDVVAVVNDTVGTMMGCEPGVRPCEVGLVV-----------------------
-----D-----TGTNACYMEEAQHV-AVLDNDR--GRMCVSVEWGSFSDNGALGPVLTTFDHALDHESLN
PGDQRFEKMIGGLYLGELVRLVLVHLAHHGVLFGGYTSPALLSQGSILLEHVAEMED------------
--PS--AGVARVHAILQD-LGLSPEASDIELVQHVCEAVCTRAAQLCGAALAAVLSRLQRSR-EHQTLQV
AVATGGQVFEQH-PRFSSILQDTVMLLAPQCDVSFVPSVDGGGQGVAMVTAVAARLAAHQLLLEETLAPF
RLNRKQLAAVQAQMQEAMARGLRGEA---S-SLRMLPTYVRATPNGS-----------------------
---ERGDFLALDLGGTNFRVLLVRVATG---GVQITSQVYSIPECVAQGSGQQ-------LFDHIVDCIV
DFQDKQGLSGQ-SLPLGFTFSFPCRQIGLDQGILLNWTKGFNASDCEGQDVVYLLREAIRRRQ----AVE
LKVVA-----IVNDTVGTMMSCGYEDPHCEIGLIVGTGTNACYMEEL---RNVAGVA-------------
--GD------SG---------------------HMCINMEWGAFGDDGS-LGMLSTRFDASVDQMSINPG
KQR-FEKMISGMYLGEIVRHILLHLTSLGVLFRGQQIQCLQTRDIFKTKFLSEIES--------------
------------DSLALRQVRAILEDLGL-PLTSDDAQMVLEVCQAVSQRAAQLCGAGVAAVVERIRENR
GLEELTVSVGVDGTLYKLH-----------PHFSSLVAATVRELAPRCAVTF---------LQSEDG---
----------SGKGAALVTAVACRLAQ--KIR--V-----------------

>XP_029798653|Suricata suricatta|HK3

LQQFKVTRTQLQQIQANLLGSMEQAL-AGQVSPAPA--VRMLPTYVGSTPHGTEQGDFLVLELGATGASL
RVLWVTLMGT---EG-HR--MEPRSQEFVIPPEVMLGPGQQ---------------LFDFAAHCLSEFLD
AL-----PVS---TQDLQLGFSFSFPCHQTGLD------------------------R------------
-----------STLISWTKGFRCSGVEGHDVVQLLRDAIKRQ----------G-----------------
--------------AYNIDVVAVVNDTVGTMMGCEPGVGPCEVGLVV-----------------------
-----D-----TGTNACYMEEARHV-AVLDEDR--GRVCVSVEWGSFGDDEALGPVLTTFDHSLDQESLN
PGAQRFEKMIGGLYLGELVRLVLAHLAQRGVLFGGRTSPALLSRGSILLEHVAEMED------------
--PL--AGAARVHAILQD-LGLSVGASEAELVQYVCAAILTRAARLCAAALAAVLSHLQHSR-EQQMLQI
AVATGGRVFERY-PSFLSILQETVKLLVPECDVSFIPSVDGGGRGVAMVTAVAARMAAHRRLLEETLAPF
RLNHEQLAAVQAQMQEAMAKGLRGEA---S-SLRMLPTYVRATPDGS-----------------------
---ERGDFLALDLGGTNFRVLLVRVTAG---GVQITNQVYSIPECVAQGSGQQ-------LFDHIVDCIV
DFQQKQGLSGQ-SLPLGFTFSFPCRQLGLDQGILLNWTKGFSASDCEGQDVVCLLREAIRRRQ----AVK
LNVVA-----IVNDTVGTMMSCGYEDPRCEVGLIVGTGTNACYMEEL---RNVAGVA-------------
--GD------YG---------------------HMCINMEWGAFGDDGS-LGLLRTSFDANVDQASINPG
KQS-FEKMISGMYLGEIVRHILLHLTSLGVLFRGQQTQRLQTRDIFKTKFLSEIES--------------
------------DSLALRQVRAILEDLGL-PLTSDDALMVLEVCQAVSQRAAQLCGAGVAAVVEKIRENR

GLDELTISVGVDGTLYKLH-----------PHFSGLVAATVRELAPRCAVTF---------LQSEDG---
----------SGKGAALVTAVACRLAQ--MDH--V-----------------
> P52790|Homo sapiens|HK3
LQQFKVTRAQLQQIQASLLGSMEQAL-RGQASPAPA--VRMLPTYVGSTPHGTEQGDFVVLELGATGASL
RVLWVTLTGI---EG-HR--VEPRSQEFVIPQEVMLGAGQQ---------------LFDFAAHCLSEFLD
AQ-----PVN---KQGLQLGFSFSFPCHQTGLD-----------------------R------------
-----------STLISWTKGFRCSGVEGQDVVQLLRDAIRRQ----------G-----------------
--------------AYNIDVVAVVNDTVGTMMGCEPGVRPCEVGLVV-----------------------
-----D-----TGTNACYMEEARHV-AVLDEDR--GRVCVSVEWGSFSDDGALGPVLTTFDHTLDHESLN
PGAQRFEKMIGGLYLGELVRLVLAHLARCGVLFGGCTSPALLSQGSILLEHVAEMED------------
--PS--TGAARVHAILQD-LGLSPGASDVELVQHVCAAVCTRAAQLCAAALAAVLSCLQHSR-EQQTLQV
AVATGGRVCERH-PRFCSVLQGTVMLLAPECDVSLIPSVDGGGRGVAMVTAVAARLAAHRRLLEETLAPF
RLNHDQLAAVQAQMRKAMAKGLRGEA---S-SLRMLPTFVRATPDGS----------------------
---ERGDFLALDLGGTNFRVLLVRVTTG----VQITSEIYSIPETVAQGSGQQ-------LFDHIVDCIV
DFQQKQGLSGQ-SLPLGFTFSFPCRQLGLDQGILLNWTKGFKASDCEGQDVVSLLREAITRRQ----AVE
LNVVA-----IVNDTVGTMMSCGYEDPRCEIGLIVGTGTNACYMEEL---RNVAGVP------------
--GD------SG---------------------RMCINMEWGAFGDDGS-LAMLSTRFDASVDQASINPG
KQR-FEKMISGMYLGEIVRHILLHLTSLGVLFRGQQIQRLQTRDIFKTKFLSEIES-------------
------------DSLALRQVRAILEDLGL-PLTSDDALMVLEVCQAVSQRAAQLCGAGVAAVVEKIRENR
GLEELAVSVGVDGTLYKLH-----------PRFSSLVAATVRELAPRCVVTF---------LQSEDG---
----------SGKGAALVTAVACRLAQ--LTR--V-----------------
> XP_043376493|Dermochelys coriacea|HK3
LQPLCLSLEKLHRVKDSMLEAMDRGL-SRRTHAQAT--VRMLPNFIRSTPDGTERGEFLVLELR--DTDL
RVLLVELSGD---TR-QM--VTVKQQLFPVSDAMAQGPGEE---------------LFAYMATCLCEFLD
RL-----RLP---RRAFQLGFSFPFHCRQTRLD-----------------------Q------------
-----------SYLISWTKSFKCPGVEGRDVVELLQNAINKQ----------D-----------------
--------------RYPITVIAVVNDTTGTMMSCEASAEPCEVGLVV-----------------------
-----D-----AGTNSCFMAEARHV-EMVEESE--GRMGVVTEWGSFGDDGTLNDMLSPYDRLVDEASLN
PGKSRFEKMISVLYLGEIVRHIL---ATQGGLFSERTTPALLTKGSVQTRDILEITD------------
--EQ--VGLVRAQCVLRQ-LGLEASDQDCFSMQQVCSTVCIRAASLCAAGLAAILSHMCRNR-ELTHLKT
SVAVDGAMFRDH-AKFSQTLQQLVKQLAPECDVTLVQAEDGSGKGIAMVAAVASRQVAQRLQVNQTLALL
QLSPDALGKVQALLRREMELGLKKETHAMS-SVRMLPTYICATPDGS----------------------
---ERGNFLALDLGGTNFRVLLVQVGSREQGGVQIVSEVYSIPPYMAQGPGSQ-------LFDHIVECIM
DFQAKQGMMGQ-TLPLGFTFSFPCQHTGLDQGVLLSWTKGFNASDCVGQDVVKLLREAVKRKN----HFA
LDVVA-----IVNDTVGTMMSCGYDDPSCEVGLIVGTGTNACYMEEL---RNVGTLA------------
--GG------EG---------------------RMCVNMEWGAFGDNGC-LDAFVTPFDRQVDEASLNPG
RQR-FEKLISGMYLGEIVRRILLELTAKGLLFRGRPSPKLQTRDIFQTKFLSSIES-------------
------------DGLALRQVCAILEGLGL-HVTCEDSVLVREVCQVVARRAAQLCGAGVAAVVEKIRENR
GLEQLSVTVGVDGTLYKLH-----------PHFAQEVQKTVQLLAPRCSVTF---------LRSEDG---
----------SGKGAALIAAVACRVAR--HQQ--A-----------------
> NWI97254|Pitta sordida|HK3
LCAFTIPLERLQMMKGHMMQAMHTGL-SCQTHAQAT--VQMLPTYVCSVPDGTEEGDFLVVELC--QSQV
RTLMVTLSGD---GN-LS--PQMVYRIFDMPEDIMHGVGEA---------------LFDFMAQCLSKFLA
DT-----SSI---QERLPLGFVFPFSCHKTKLD-----------------------K------------
-----------AVLISWSKGFSCSGVEGKDVVQLLQEAIYKQ----------E-----------------
--------------LYHVDVVALMNDTVGTMMSCISSGRPCEISLIA-----------------------
-----D-----KGSNCCFMAEAHLV-EMAEETS--GRMCVNTEWGCFGDDGMLSDILTPYDESMDNESSS
PGVKRFEKLVGTLYLGEIVRHVLITLITQKALFNRSDIAVLKKKGVLKTQQVLEIIN------------
--NE--EGTAAAKRILES-LGLEPNEQDCSRVQQICQAVVTRAATLHAAALAAVLSFMCRSR-EMEELLV
NVAVDGELYHSH-TRFKEILQNVIKLLAPECTAIPLPSTDGTGRGAAMVTAVAVRLEARRREVNEVLAPL
RLSRADLAKVQALMRQEMELGLGRETSATA-CVRMLPTYVCHTPDGT----------------------
---ERGDFLALDLGGTNFRVLVVHVSQE---GIQMASEIYVIPTAIMQGTGEQ-------LFDHIMDCII

DFQLKQNLMEQ-VLPLGFTFSFPCQQVGLDKAFLLSWTKGFCASGCVGQDVVQLLWEAARRKN----HSA
LNVVA-----LVNDTVGTMMSCGYDDPKCEIGLIVGTGTNACYMEEM---QNVGTVE------------
--GE------QG--------------------QMCINMEWGAFGNNGC-LDHIFTSFDTVVDEQTINPG
KQR-FEKLISGMYLGEIVRHILLAMTERRLLFKGNPSPKLLTKDIFPTKFLSTIEL-------------
------------HGLALRHVQAIVQDLEL-NATFEDCVLMREVCQTVSLRAAQLCAAGLAAVVEKMRENR
GLDQLTVTVGVDGTLYKLH-----------PNFSRNVQKTLQDLAPNCIVTF---------MPSEDG---
----------SGKGAALVAAVASRTTS-------P-----------------
>NXF12746|Smithornis capensis|HK3
LRMLTIPLERLQMMKVHMMAAMCKGL-SRETHTQAT--VQMLPTYICSTPDGTEKGDFLVVELC--QSQV
RILLVTLFGD---GN-LS--PQMIYKIFDMPENIMHGEGEA---------------LFDFIAQRLSEFLG
GT-----SSS---QERIPLGFVFPFSCRQTKLD-----------------------K-----------
-----------AELISWSKGFSCSDVEGKDVVQLLQWAIYKQ----------E----------------
--------------LSHVDVVALMNDTVGTMMTCSTEGRPCEIALIA----------------------
-----D-----KGSNCCFMAEAHLV-ETAEETS--GRMCVNTEWGCFGDDGMLSDILTPYDESIDDESSN
PGLKRFEKIVGTLYLGEIVRHVLITLVAQKALCSRSNVAVLKQKGVLKTQQVLEIIN------------
--NG--EGMAAAKRVLEA-LGLESSEQDCSRMQQICQAVVTRAATLHAAGLAAVLSFMCQSR-DMDQLLV
NVGVDGELYHGC-ARFREVLENVVKLLAPDCTATLLPSTDGSGRGAAMVTAVAVRLEAQRREVDEVLGPL
RLSRADLERVQALLRREMDLGLCKETNATS-SVRMLPTYVCGTPDGT---------------------
---EQGDFLALDLGGTNFRVLVVRVSQE---GIRMASEIYVIPTEITQGTGEQ-------LFDHIVDCIM
DFQLKQNLMKQ-VLPLGFTFSFPCQQVGLDKAFLLTWTKGFSASGCVGQDVVHLLREAAQRKN-----VG
VEVVA-----LVNDTVGTMMSCGYDDPKCEIGLIVGTGTNACYMEEM---KNVGTVE------------
--GE------QG--------------------RMCINMEWGAFGDNGC-LEHIFTSFDQVVDKDTINPG
KQK-FEKLISGMYLGEIVRHILLAMVDRQLLFKGNPTPKLQTKNIFQTKFLSTIEL-------------
------------NGLALRQIRTILQDLEL-DASFEDSVLMRDVCHTVSQRAAKLCAAGLAAVVEKMRENR
SLDQLSVTVGVDGTLYKLH-----------PTFSHVVQETLRDLAPKCNVTF---------MPSEDG---
----------SGKGAALVAAVAC-----------------------------
>NWI59033|Calyptomena viridis|HK3
LRALTIPLERLQMMKGHMMQDMRKGL-SRQTHAQAT--VQMLPTYICSTPDGTEKGDFLVVELC--QSQV
RILLVTLFGD---GN-LS--PQMVYKIFDMPEDIVHGEGEA---------------LFDFIAQRLSEFLG
DT-----SSS---QERIPLGFVFPFSCHQTKLD-----------------------K-----------
-----------AELLSWSKGFSCGDVEGKDVVQLLQWAIYKQ----------E----------------
--------------LCHVDVVALMNDTVGTMMTCSTEGRPCEIALIAGEEWASNLPEWWVMGALHQCSPP
SLPP-D-----KGSNCCFMAEAYLV-EITEETS--GRMCINTEWGSFGDDGMLSDILTVYDESMDDESSN
PGLKRFEKLVGTLYLGEIVRHVLIVLAAQKALFTGSDVAILKEKDVIKTQQVLEIIN------------
--NG--ENTAAVKRILEG-LGLESSERDCSRVQQICQAVVTRAATLYAAGLAAVLSYMCQCR-DMDQLLV
NVGVDGELYHTC-TRFKEILENVVKLLAPECTATLLPSTDGSGRGAAMVTAVAVRLEAQRREVDELLAPL
RLSRADLERVQALMRQEMELGLGQETNPTA-CVRMLPTYVCATPDGT---------------------
---EQGKFLALDLGGTNFRVLVVHVSEG---GIRMASEIYVIPTGVMQGTGDQ-------LFDHIVDCIM
DFQLKQNMMEE-VLPLGFTFSFPCQQVGLDKALLLTWTKGFSASGCVGQDVVQMLRDAAQRKN-----VG
LQVVA-----VVNDTVGTMMSCGYDDPKCEIGLIVGTGTNACYMEEM---QNVGTVE------------
--GE------QG--------------------RMCINMEWGAFGDSGC-LDNILTSFDTVVDKDTINPG
KQR-YEKLISGMYLGEIVRHILLAMVDRELLFKGNPSPKLQTKNIFQTKFLSTIEL-------------
------------NGLALRQIRAILQDLEL-NASFEDSVLMREVCHVVSERAAKLCAAGLAAVVEKMRENR
GLDQLSVTVGVDGTLYKLH-----------PTFSHILQETLQDLAPKCTVTF---------MPSEDG---
----------SGKGAALVAAVACNISS--RGK-QP----------------
>NXG25478|Grallaria varia|HK3
LRAFTIPLERLQLMKSHMMQAMRRGL-SRHTHREAT--VQMLPTYICATPDGTEKGDFLVVELC--QSQV
RTLLVTLFGD---GN-LR--PQIMHKTFDMPEEIMQSRGEV---------------LFDFIAQCVSQFLA
DT-----GSS---QERLPLGFVFPFSCRQTQLD-----------------------K-----------
-----------AELISWSKGFSCSDVKGRDVGQLLQSAIHKQ----------E----------------
--------------LSHVEVVALMNDTVGTMMTCSTEGTPCEVALVM----------------------
-----D-----KGSNCCFMAEAHLV-EIAKETS--GRMCVNTEWGCFGDNGIVNDIVTPYDECVDNESSS

PGEKRFEKMVGSLYLGETVRHVLITLAAEKALFTGRSAAVLKEKGALKTQQVLEIFD-------------
--KE--EGMAVTKGILEA-LGLKPNERDCSRVQQICRAVVSRAATLNAAGLAAVLSYMCQTR-DLDQLVV
NVGVDGELYHGH-PRFEEIVQSVLRLLAPECVAIFLPSRDGSGRGAAMVTAVALRLEAQRREVAEILAPL
RLSYADLVQVQALMRQEMELGLGQETNANA-SVRMLPTYVCNTPDGT-----------------------
---EQGEFLALDLGGTNFRVLVVRVSQE---GIRMASEVYVIPTAIMQGTGEA-------LFDHIIDCII
NFQRKQDLMQH-ALPLGFTFSFPCQQVGLDKALLLTWTKGFSASDCVGRDVVQLLREAAQRKH----HSA
LNVVA-----LVNDTVGTMMSCGYDDPKCEIGLIVGTGTNACYMEEM---RNVGTVK-------------
--GD------QG---------------------RMCINMEWGAFGDNGC-LDHIFTSFDRVVDNQTINPG
KQR-FEKLISGMYLGEIVRHILVKMADKALLFHGKPSPKLFCRDIFKTKFLSTIEL--------------
------------NGLALRQIRIILQDLEL-NASYEDSVLMREVCEVVSLRAAQLCAAGLAAVVEKMRENR
GLDQLSVTVGVDGTLYKLH-----------PNFSHNLQKTLQGLAPNCTVTF---------MPSEDG---
----------SGKGAALVAAVACR----------------------------
>NXP27561|Scytalopus superciliaris|HK3
LRALTIPLERLQVMKNHMMQAMCKGL-SRQTHAQAT--LQMLPTYICSTPDGTEKGNFLVVELC--QSQV
RTLLVTLLGD---GN-LS--PQMMYKTFDMPEEIMKGEGEA---------------LFDFIAQCLSQFLA
DI-----GSF---QERLPLGFVFPFSCQQTQLN------------------------K------------
-----------AELISWSKGFSCSNVEGKDVVQLLQSAINKQ----------E-----------------
--------------LSHVEVVALMNDTVGTMMTCSTERAPCEVALVM-----------------------
-----D-----KGSNCCFMAEAHLV-EMAEETS--GRMCVNTEWGCFGDNGSLNDIVTPYDTCVDGESSS
PGEKRFEKMVGSLYLGEIVRHVLIALAAEKVLFTGKNVAVLKEKGVLKTQQVLDIIK-------------
--HE--EGMAATKNALEA-LGLQPNERDCGRVQQICLAVVSRAASLYAAGLAAVLTYMCHTR-DMDQLLV
NVGVDGELYHGH-TRFEEVLHTVTRLLAPECVATFIPTTDGSGRGAAMVTAVALRLEARRREVNEVLAPL
RLSRADLEQVQALMRQEMERGLGRETHDNA-SVRMLPSYVCSTPDGT-----------------------
---EQGQFLALDLGGTNFRVLVVHVTEE---GIRMASEIYVIPTAIMQGSGEA-------LFDHIIDCIL
DFQMKQNLMNH-VLPLGFTFSFPCKQVGLDKALLLTWTKGFSASNCVGQDVVQMLREAAQRKQ----HSG
LNVVA-----LVNDTVGTMMSCGYDDPKCEIGLIVGTGTNACYMEEM---RNVGTVE-------------
--GD------EG---------------------RMCINMEWGAFGDNGC-LDHIFSNFDKVVDDQTINPG
KQR-FEKLISGMYLGEIVRHILLAMADRKLLFQGKPTPKLYCKDIFKTKFLSTIEL--------------
------------DELALRQVRAILQELEL-NASHEDSVLMLEVCRTVSLRAAQLCAAGLAAVVEKMRENQ
GLNQLSVTVGVDGTLYKLH-----------PNFSKNLQKTLQDLAPNCTVTF---------MPSEDG---
----------SGKGAALVAAVACRINS-------P-----------------
>NXU93189|Xiphorhynchus elegans|HK3
LRALTIPLERLQVMKDHMMQAMQKGL-SRSTHSQAP--VQMLPTYICSTPDGTEKGDFLVVELC--QSQV
RTLMVTLLGD---GN-LS--PQMMYKTFDMPEEIMKGSGEA---------------LFDFIAQCVSQFLA
DT-----SSS---QERLPLGFVFPFSCRQIQLD------------------------K------------
-----------AELISWSKDFSCSDVEGKDVGQLLQSAINKQ----------E-----------------
--------------LSHVELVAVMNDTVGTMMTCSTEGRPCEVALVM-----------------------
-----D-----KGSNCCFMAEAHLV-EIAKETS--GRMCVNTEWGCFGDNGSLNDIVTPYDECVDNESSS
PGEKRFEKMVGSLYLGEIVRHVLITLAAEKVLFKGSNVAILKEKGVVKTQQVLEIIN-------------
--HE--EGLAVTKRMLEA-LGLPPNERDCSRVQQICLAVVSRAASLYAAGLAAVLTYMCQTR-EMDQLLV
NVGVDGELYQGH-TRFEEILQSVTRLLAPECVATFLPSTDGSGRGAAMVTAVALRLEAQHREVNEVLAPL
RLSLADLEQVQALMRQEMELGLGQETNANA-SVRMLPTFVCNTPDGT-----------------------
---EQGEFLALDLGGTNFRVLVVRVTEE---GIRMASEIYVIPTAIMQGTGEA-------LFDHIIECII
DFQTKQDLMNH-VLPLGFTFSFPCKQVGLDKALLLTWTKGFSATDCVGQDVVQLLREAAQRKH----QTG
LNVVA-----LVNDTVGTMISCGYDDPKCEVGLIVGTGTNACYMEEM---RNVGTVE-------------
--GD------EG---------------------RMCINMEWGAFGDNGC-LDHIFTSFDKLVDDQTINPG
KQR-FEKLISGMYLGEIVRHILLAMADRQLLFHGKPGPKLYCKDIFKTKFLSTIEL--------------
------------NGLALRQIRAILQDLDL-DVSYEDSVLLREVCEAVSLRAAQLCAAGLAAVVEKMRESR
GLDQLTTTVGVDGTLYKLH-----------PTFSENVQKTLQDLAPKCSVTF---------MPSEDG---
----------SGKGAALVAAVACRITS--SGQ--------------------
>NWW71186|Climacteris rufus|HK3
LRSLSVPLDRLHIMKGHMMQDMCRGL-SRYTHTQAK--VRMLPTYICSTPNGTEKGNFLVVELC--QSQV

RTLLVTLYGD---GN-LS--PQMMYKIFDMPEGIMQGEGEA---------------LFDFIAQCISQFLA
ETISSDSSSS---EERLPLGFVFPFSCRQTQLD-----------------------K------------
-----------AELISWSKGFSCSGVVGKDVVQLLQSAIDKQ----------E-----VGSGGAGGRWLS
PWRSWKLSPFAPGQLSRVNVVALMNDTVGTMMTCSTEGRPCEVAVVA----------------------
-----D-----KGSNCCFMAEAHLV-ETTEETS--GRMCVNTEWGCFGDDGILNDIITPYDECVDQESSN
PGEKRFEKLVGSLYLGEIVRHALLGLTAEKAIFTGTDAAVLKQKGVFTTQQVLEIIN------------
--NE--DSTTEVKRVLEV-LGLQPSERDCGRVQQICRAVVGRAATLYATGLAAILSYMCRSR-EMEQLMV
NVGVEGELYQGY-TRFEEILQSVSKLLAPECLATLLPSRDGSGRGAAMVTAVALRLEAQRREVNEVLAPL
RLNHADLEKVQALMRQEMELGLGRDTNATA-SVRMLPTFVCGTPDGT----------------------
---ERGDFLALDLGGTNFRVLLVRVTEE---GISMASEIYVIPTTIMQGTGEE-------LFDHIIECIM
DFQMKQKLAMD-NLPLGFTFSFPCQQVGLDKALLLNWTKGFSASDCVGQDVVHLLRNAAQRKQ----HVG
LQVVA-----LVNDTVGTMMSCGYDDPRCEIGLIVGTGTNACYMEEM---KNVGTVE------------
--GE------QG---------------------RMCINMEWGAFGDNGC-LDHIFTHFDRVVDDSTINPG
KQR-FEKLISGMYLGEIVRHILLEMTERELLFQGRPCPKLQTKNIFQTKFLSTIEL-------------
-----------SNGLALRQIRAILQELEL-DASFEDSVLMREVCQTVSLRAAQLCAAGLAAVVEKMRENR
GLDQLSVTVGVDGTLYKLH-----------PCFSYNLQKTLKDLAPNCNVSF---------LLSEDG---
----------SGKGAALVAAVASRAA-------------------------
> NXU35529|Drymodes brunneopygia|HK3
LRSLSIPLERLHIMKGHMMQDMCKGL-CRQTHTQAK--VRMLPTYICSTPNGTEKGNFLVVELC--QNLV
RTLLVTLYGD---GN-MS--PHMVYKIFDMPEGIMQGEGEA---------------LFDFIAQCVTQFLA
ETTNPDTNTS---GENLPLGFVFPFSCRQTRLD-----------------------K------------
-----------AELLSWSKGFSCSGVVGKDVVELLQSAINKQ----------E-----AGGQGGGGNWLS
SWRGRKPSQPAAGQFRHVEVVALMNDTVGTMMTCSMEGRPCEVAVVA----------------------
-----D-----KGSNCCFMAEAFLV-EITDETS--GRMCVNTEWGCFGDDGTLNDIFTPYDVCVDEESSD
PGEKRFEKLVGSLYLGEIIRHALIALTAEKAIFTGTDVGVLKQKGVFTIQHVLEIIN------------
--NE--EGTTEVKRILEA-LGLQPSERDCGRVQQICRAVVGRAATLYATGLAAILSYMCQTR-EMETLMV
NVGMEGELYTGY-PRFEEILQSVSRLLAPECMATLLPSRDGSGRGAAMVTAVALRLAAQRREVNEVLAPL
RLSREDLEGVQALMRQEMERGLNKETHDTA-SVRMLPSYVSHTPDGT----------------------
---ERGNFLALDLGGTNFRVLVVRVTEE---GINIASEIYVIPASIMQGTGEG-------LFDHIIDCIL
DFQTKQNLMTN-VLPLGFTFSFPCKQVGLDKALLLTWTKGFSASGCVGQDVVQLLREAAVRKQ----HSG
LRVVA-----LVNDTVGTMMSCGYEDPNCEIGLIVGTGTNACYMEEM---RNVGTVE------------
--GD------EG---------------------RMCINMEWGAFGDNGC-IDHLFTHFDKVVDETTINPG
KQR-FEKLISGMYLGEIVRQILMVMAEKQLLFQGRPCPKLQTKDIFQTKFLSTIEL-------------
-----------SNGLALRQIRDILNELEL-DASFDDSVLMREVCQTVSLRAAQLCAAGLAAVVEKMRENR
GLDQLSVTVGVDGTLYKLH-----------PCFSDNLQKTLKELAPKCNVTF---------LLSEDG---
----------SGKGAALVAAVANRA----AGQ--------------------
> XP_064244153|Passer domesticus|HK3
LRSFTIPLERLHVMKGHMVQDMCKGL-SRETHAQAK--VRMLPTYICSTPNGTEKGSFLVVELC--QNQV
RTLLVTLYGD---GN-MS--PQMMYKIYELPEDIMQGEGEA---------------LFDFIAHCLTQFLA
ETISPDTNIS---EQHLPLGFVFPFTCQQTRLD-----------------------K-----------
-----------AELLSWSKGFSCSGVVGKDVVQLLQSAIDKQ----------AKPDKQDGEGNGSGNWLS
SWKGRKPSQSAPSLPYNVEVVALMNDTVGTMMTCSMEGRACEVAMVA----------------------
-----D-----KGSNCCFMAEAYLV-ETTEETS--GRMCVNTEWGCFGDDGTLNDILTPYDQSVDVESSN
PGEKRFEKLVGTLYLGELVRHTLVALTAEKAVFTGTNAAVLKEKGVFTMQHVLDIAN------------
--NE--DGITEVRRILEV-LGLQPSERDCGRVQQICRAVVARAVTLHATGLAAILSYMCQTR-DLESLMV
NVGVEGELYAGY-PRFEEVLLSVSRLLSPECMATILPSRDGSGRGAAMVTAVALRLASQRRVVDEVLAPL
RLTRDDLVKVQALMRQEMDQGLDKETNPTA-SVRMLPTYVNHTPDGT----------------------
---EKGDFLALDLGGTNFRVLVVRITEE---GITMASEIYVIPVSIMQGSGEE-------LFDHIIDCII
DFQTKQNLVTQ-MLPLGFTFSFPCKQVGLDKALLLTWTKGFSASGCVGQDVVQLLRDAAKRKR----HLG
MQVVA-----LVNDTVGTMMACGYDDPKCEIGLIVGTGTNACYMEEM---KNVGTVE------------
--GD------EG---------------------RMCINMEWGAFGDNGC-LDHLFTHFDKVVDETTINPG
KQR-FEKLISGMYLGEIVRQILLVLTEKELLFQGKPCPKLQTKDIFKTKFLSTIEL-------------

------------NGLALRQIRNILNELEL-DASFEDSVLMREVCQTVSLRAAQLCAAGLAAVVEKMRENR
GVDQLSVTVGVDGTLYKLH-----------PCFSSNLQKTLKDLAPNCNVTF---------LLSEDG---
----------SGKGAALVAAVADRAAN--AME--------------------
>NWR50242|Regulus satrapa|HK3
LRSLSIPLERLHIMKGHMIQDMCKGL-SRQTHAQAK--VRMLPTYICSTPNGTEKGNFLVVELC--QNQV
RILLVTLYGD---GN-MS--PQMMYKIFDMPEGIMQGEGEA---------------LFDFIAQCVSQFLA
ETISPEARGS---EEHLPLGFVFPFSCRQTQLD------------------------K------------
-----------AELLSWSKGFTCSGVVGKDVVQLLQSAIDKQELQEVGDKGTG------------GRWLS
LWKGQKSSKSTPGQLCHVDVVALMNDTVGTMMTCSMEGRPCEVAIVA-----------------------
-----D-----KGSNCCFMAEAYLV-ETAEETS--GRMCVNTEWGCFGDDGSLDDVFTPYDDCVDEESLN
PGVKRFEKMVGTLYLGEIVRHALVGLTAEKAVFTGTNADVLKEKGVFTMQHVLNIIN-------------
--NE--ESITEVRRILEV-LGLQPSERDCGRVQQICRAVVGRAATLHATGLSAILSYMCQTR-DLEALMV
NVGVEGELYTGY-PRFEEILLSVSRLLSPECMATLLPSRDGSGRGAAMVTAVALRLAAQRREVNEVLAPL
RLSLADLEKVQALMRDEMEHGLCKETNPSA-SVRMLPTYVSHTPDGT-----------------------
---ERGDFLALDLGGTNFRVLVVRVTEE---GISMASEIYVIPAAIMQGTGEG-------LFDHIIDCIV
DFQTKQNLVAQ-KLPLGFTFSFPCKQVGLDKALLLTWTKGFSASDCVGQDVVQLLRDAAQRKQ----HLG
MHVVA-----LVNDTVGTMMACGYDDPKCEIGLIVGTGTNACYMEEM---RNVGTVE-------------
--GD------QG---------------------RMCINMEWGAFGDNGC-LDDIFTHFDRVVDESTINPG
KQR-FEKLISGMYLGEIVRQILLVMTDKQLLFQGKSSTKLQTKDIFKTKFLSTIEL--------------
-----------SNGLALRHIRAVLNELEI-DASFEDCVLMREVCQTVSLRAAQLCAAGLAAVVEKMRENQ
GVEQLSVTVGVDGTLYKLH-----------PCFSRNLQETLKILAPNCDVTF---------LLSEDG---
----------SGKGAALVAAVANRATE--AME--------------------
>NWZ72822|Acrocephalus arundinaceus|HK3
LRSLSIPLERLHIMKGHMIQDMCKGL-SRQTHAQAK--VRMLPTYLCSTPNGTEKGSFLVVELC--QSQV
RILLVTLFGD---GN-MS--PHMVYKIFDLPEGITQGEGEA---------------LFDFIAQCVSQFLA
ETFSSDTSKP---EERLPLGFVFPFTCQQTQLD------------------------K------------
-----------AELLSWSKGFNCSGVVGEDVVKLLQTAINRQ----------Q-----------------
--------------LCHVDVVALMNDTVGTMMTCSTEGRPCEVAIVA-----------------------
-----D-----KGSNCCFMAEAYLV-ETTDETS--GRMCVNTEWGCFGDDGTLNDVFTPYDESVDVESYN
PGEKRFEKMVGTLYLGEIVRHALIALAAEKAIFVGVNIAVLKEKGVFTMQHIMDIIN-------------
--NE--DGISDVKRILEV-LGLQPTERDCGRVQQICRAVVGRASTLHATGLAAILSYMCQTR-DLETLMV
NVGVEGELYTGY-PRFEEILMSVSRLLSPECMATLLPSKDGSGRGAAMVTAVALRLAALRREVNEVLSPL
RLSRSDLEKVQALMRQEMDQGLGKETNANA-SVRMLPTYVSHTPDGT-----------------------
---ERGDFLALDLGGTNFRVLVVHVTEE---GISMASEIYVIPAAIMQGTGEA-------LFDHIIDCII
DFQTKQNLVTD-TLPLGFTFSFPCKQMGLDKALLLTWTKGFSASGCVGQDVVQLLREAARRKQ----HLG
MQVVA-----LVNDTVGTMMACGYDDPKCEIGLIVGTGTNACYMEEM---KNVGTVE-------------
--GD------QG---------------------RMCINMEWGAFGDNGC-LDHIFTHFDKVVDETTINPG
KQR-FEKLISGMYLGEIVRQVLLVMTEKQLLFQGKLTPKLQTRDIFKTKFLSTIEV--------------
-----------SNGLALRQIRAILSELEI-DASFEDSVLLREVCQTVSLRAAQLCAAGLAAVVEKMRENQ
GVDQLSVTVGVDGTLYKLH-----------PRFSDNLQKTLKDLAPNCNVSF---------LLSEDG---
----------SGKGAALVAAVASRAAN--AME--------------------
>NWY47829|Sylvia atricapilla|HK3
LRSLSVPLERLHIMKGHMVEDMCRGL-CRQTHAQAK--VRMLPTYICSTPNGTEQGNFLVVELC--QSQI
RTVLVTLYGD---GN-MS--PNMVYKIYDLPEGITQGKGEA---------------LFDFIAQCVSQFLA
ETIRSDPSKS---VERLPLGFVFPFTCRQTQLD------------------------K------------
-----------AELLSWSKGFSCNDVIGKDVVELLQSAINKQ----------E-----------------
--------------LCHVDVVAVMNDTVGTMMTCSTEGRPCEVAMVA-----------------------
-----D-----KGSNCCFMAEAYLV-ETADETS--GRMCVNTEWGCFGDDGTLNDIITPYDECVDVESSN
PGEKRFEKMVGSLYLGEIVRHALIGLTAEKAIFTGTDIGVLRQKGVFTMQHILDIVN-------------
--NE--EGTTEVKRILDV-LGLQSSERDCSRVQQICRAVVGRAATFHATGLAAILSYMCQTR-DMETLMV
NVGMDGELYKGY-PRFQEILLSVSRLLSPECMATLLPSRDGSGRGAAMVTAVALRLAALRREVNEVLAPL
RLSRADLEKVQALMREEMDRGLGKETNAIA-SVRMLPTYVSHTPDGT-----------------------

---ERGDFLALDLGGTNFRVLVVHVTEE---GITMASEIYVIPVAIMQGTGEA-------LFDHIIDCII
DFQMKQNLVTD-MLPLGFTFSFPCKQLGLDKALLLTWTKGFSASGCVGQDVVQLLREAARRKQ----HLG
MQVVA-----LVNDTVGTMMACGYDDPKCEIGLIVGTGTNACYMEEM---KNVGTVE-------------
--GD------QG---------------------RMCINMEWGAFGDNGC-LDHIFTHFDQVVDQSTINPG
KQR-FEKLISGMYLGEIVRQILMVMTEKQLLFQGKLSSKLQTKNIFQTKFLSTIEV--------------
------------KGLALRQIQAILNELEI-EASFEDSVLLREVCQTVSLRAAQLCAAGLAAVVEKMRENR
GLDQLSVTVGVDGTLYKLH-----------PCFSTNLQNTLKDLAPNCNVSF---------MLSEDG---
----------SGKGAALVAAVASREAN--AME--------------------
>NXU52420|Turnix velox|HK3
LVALTIPLELLHEVKRRMMAAMQQGL-SRLTHAEAP--VKMLPTFICSTPNGMEKGYMLVVELC--QIHV
RTLFMELLGD---GN-KT--PFMKNQIFNMPEDRMHGTGEE---------------LFDFIAQCVRQFLT
EH-----RLT---QQHLSLGFVFPFSCRQTQLD------------------------K------------
-----------AELISWSKGFSCSGVEGKDVVQLLQLALNKQ----------E-----------------
--------------LSNVKVVALLNDTVGTMMSCSTEEIPCEVGLVV-----------------------
-----D-----KGTNSCFMAEAQYI-EMVEETS--GRMCVNTEWGSFGDDDALNDILTPYDQHMDQESSN
PGKKRFEKLVGSLYLGEIVRHTLMTLAAEKAVFVGSNVAFLGTKDALKTQQVLEILD-------------
--NE--HGMTKARAVLEA-VGLRPSERDCCRVQQICRAVVSRSASLCAAGLAAILTHMRLSR-EMSELIV
NVAVGGELYRAH-RRYGEILQNVVALLAPECMVNFVPSEDGTGRGSAIVTAVSLRLTEKHHEVGQVLAPL
RLSRADLQGIQELMTREMELGLNKETNPSS-SIRMLPTYVRGTPDGT-----------------------
---ERGHFLALDLGGTNFRVLVVRIAEN---SIHMDSKVYLIPPSIMQGTGEA-------LFDHIISCIM
DFRVEQNLMEE-ILPLGFTFSFPCQQLGLDKAVLLTWTKGFNASGCVGEDVVQLLREAAQRKQ----HLA
LKVVA-----VVNDTVGTMMSCGYNDPKCEIGLIVGTGTNACYMEEM---KNVGTVE-------------
--GD------QG---------------------RMCINMEWGAFGDNGC-LDHIITDFDRMVDERTINPG
KQR-FEKLISGMYLGEIVRHILLALVEKRVLFHGNSCTTLETRDIFKTSFLSTIEI--------------
-TYDDTLGVWGSDGLALLQVRDILMDLEL-DASFEDSTLVREVCQTVSLRAAQLCAAGLAAVVEKMRKSR
GLAHLDVTVGVDGSLYKMH-----------PHFSQYIQQMLQDLAPNCTVTL---------MQSEDG---
----------SGKGAALVAAVASREAE-SQEQ--P-----------------
>NXG31793|Dromaius novaehollandiae|HK3
LLELTLSLETLQAVKGHMLEAMAKGL-SRQTHAQAT--VRMLPTYICSTPDGTEKGDFLVVELR--RRHV
RTLWVTLPGD---GN-QS--PQLKERLFDMPEDITQGKGEA---------------LFDFIARCLCQFLE
GV-----CPS-EPQQRLPLGFVFPFSCRQMRLD------------------------Q------------
-----------GELITWSKGFHCSEVEGHDVVQLLQAAIDKQ----------E-----------------
--------------RCQVDVVALMNNTVGTMMSCSMEGMPCEVGLVI-----------------------
-----D-----VGTNSCFMAEGWHV-ETVEETS--GRMCVNTEWGFFGDDGALSNVLTPYDQCVDQESAK
PGEKRFEKLIGGLYLGEIVRHVLAKLAAEGALFGGDSIAILETKDTFKTQQVLEIID-------------
--DE--AGLAMARSTLQA-LGLAPSEQDCCHVQQVCRAVVTRAAGLCAAGLAAILSHMCQSR-ELEQLKA
NVGVDGAVYRGH-ARFGEILQRITKLLAPECMATLVPSADGGGRGAAVVTAVALRLAAQRREVDRLLAPL
RLGPADLQRVQALMRWEMELGLNQDTNAQS-SVRMLPTYVRSTPDGT-----------------------
---ERGDFLALDLGGTNFRVLVVRVGQD---GVHMASEIYAIPLAVMQGTGEA-------LFDHIIDCIV
DFQQKQGLTGQ-VLPLGFTFSFPCQQLGLDKAVLLSWTKGFSASGCVGKDVVQMLREAAQRKQ----HFG
LDVVA-----VVNDTVGTMMSCGYDDPSCEIGLIVGTGTNACYMEEM---QNVGTAE-------------
--GE------QG---------------------RMCINMEWGAFGDNGC-LDDIFTDFDRLVDEKTINPG
KQR-FEKLISGMYLGEIVRYILLALVEKHILFRGKPSPKLQTRDIFQTKFLSTIEM--------------
-----GPRCAGSDGLALRQVRGVLHDLEL-DASFEDGVLVREVCQAVSLRAAQLCAAGLAAVVEKMRESR
GLAQLAVTVGVDGTLYKMH-----------PRFSQHLQRTLQDLAPNCTVTF---------LQSEDG---
----------SGKGAALVAAVHCRVAQ-AQGP--P-----------------
>NWJ02243|Crypturellus undulatus|HK3
LLELTVSLETLHEVKGHMLEAMAKGL-SQQTHEEAT--VHMLPTYVCSTPDGTEKGEFLVVELR--RRHV
RTLWVTLPGD---GN-QN--LQLKDQLFDMPEDIMQGKGEA---------------LFDFIAQCLCQFLD
GA-----GLS-NPQQRLPLGFVFPFSCRQTRLDQARHSQQGARPVQHRVPTARPCSPQ------------
-----------GELIAWSKDVRCSDVEGHDVVQLLQAAINKQ----------E-----------------
--------------RYQVDVVALLNDTVGTMMSCSTEGMPCEVALLA-----------------------

-----D-----TGTNSCFMAEGCHV-EIVEETS--GRMCVNTEWGSFGDDGALSSVLTPYDQSVDQESQK
PGHQRFEKLIGGLYLGEIVRHVLAKLAAEGALFSGTAAPALGTKDVFKTQQVLEIID------------
--DE--AGLAAARNTLQA-LGLAPSEQDCCHVQQVCRAVVTRAAGLCAAGLAAILSYMCRSR-ELERLKT
NVAVDGTVYRGH-PRFGEILHRITKLLAPECTVTLVPAMDGGGRGAAMVTAVALRLAAQRRELDQILAPL
RLSIADLQRVQALMRREMDLGLNKETNAQS-SVRMLPSYVRNTPDGT----------------------
---ERGDFLALDLGGTNFRVLVVHVEKE---GVRMASEIYAIPQAIMQGTGEA-------LFDHIIDCII
DFQQKQGLTGQ-VLPLGFTFSFPCRQLGLDKAVLLNWTKGFSASGCVGNDVVQMLREAARRKQ----HTG
LDIVA-----VVNDTVGTMMSCGYDDPSCEIGLIVGTGTNTCYMEEM---QNVGTVE------------
--GE------EG---------------------RMCINMEWGAFGENGC-LDDIFTDFDRMVDEKTINPG
KQR-FEKLISGMYLGEIVRYILLALVEKHILFRGKPAPKLQTRDIFQTKFLSTIEI-------------
--GADGPRCACSDGLALRQVRGVLHELEL-DASFEDSVLVREVCQTVSLRAAQLCAAGMAAVVEKMRESR
GQTQLAVTVGVDGTLYKLH-----------PRFSQHLQHTLRELAPKCFVTF---------LQSEDG---
----------SGKGAALVAAVQCRLAQ-AQGQ--Q-----------------
>XP_031454018|Phasianus colchicus|HK3
LLALTPTLEKLQEMKGRVLEALEQGL-SQQSGARST--MKMLPTFICSTPDGTENGDVLVAELC--QNHV
RTLWVTLTGD---GN-QS--PQLMYHVFEMPEDITQGSGEA---------------LFDFIAQCVKNFLE
EI-----GNP---QHRLPLGFIFPFSCRQTGLD-----------------------K-----------
-----------AELISWSKGFRCSDVEGKDVVQLLQAAISKQ----------E-----------------
--------------LCHVDVVALLNDTVGTMMTCSTREKPCEIALVV----------------------
-----G-----SGTNCCFMAESHLL-DMAEVTS--GRMCVNSEWGCLGDDGALDDVLTPYDHRVDQESSN
PGERRFEKMVGSLYLGEIVRHAMLMLASEQALFIGRNTSILATKAVLKTQQVLEIMD------------
--NE--DGMAKAQGILEA-LGLQPSERDCCRVQQLCRAVVSRAAALSAMGLAAVLSHMCRSR-QLERLVV
NVGVDGGLYHTS-ARFGEILRSVVGLLAPECAANLMPSADISGLGAAMVTAVALRTMAQRNKVERLLAPL
RLSRADLQRVQALMRQEMERGLDRERNAES-SLRMLPTYVCHTPDGT----------------------
---ERGKFLALDLGGTNFRVLVVHVAED---NIDIINEIYVIPTAIMQGTGEA-------LFNHIMECIM
DFQMKQELMDQ-VLPLGFTFSFPCQQLGLDKAVLLNWTKGFSASGCVGQDVVQMLREAAQRKQ----HTA
LKVVA-----VVNDTVGTMMACGYHDPKCEIGLIVGTGTNTCYMEER---AKVGTID------------
--GD------EG---------------------RMCINMEWGAFGDNGC-LDDFFTSFDRLVDEKSINAG
KQK-FEKLISGMYLGEIVRQILLALVEEQVLFCGKSSQKLQNRDIFQTKFLSTIET-------------
------------PGLALVQVQGILRNLEL-DASYEDSVLVREVCQTVSLRAAHLCAAGMAAVVEKMRESR
GLEQLTVTVGVDGTLYKMH-----------PRFSQNLQQMLQELAPLCNVTF---------LQSEDG---
----------SGKGAALVAAVACRGAD-PSSP--Q-----------------
>NXN92093|Rhinopomastus cyanomelas|HK3
LLALTIPLETLQAVKGRMLQAMHKGL-SRQTHAQAD--VRMLPTYVCSTPDGTEKGEFLVVEMC--QNHV
RTLWMALAGD---GN-QS--PQITYKTFDMPEDIMQGKGEA---------------LFDFIAQSLRQFLD
GI-----GRP---QHHLPLGFVFPFSCRQTQLD-----------------------K-----------
-----------AELISWSKGFSCSDVEGRDVVQLLQSAINKQ----------E-----------------
--------------LYHVEVVALLNDTVGTMMTCSLSGKPCEIALIV----------------------
-----D-----KGTNSCFMTEAHLV-EMVEDSS--GQMCVNTEWGYFGDDGALRDILTPYDHNVDKESSN
PGTKRFEKLIGSLYLGEIVRHVLITLAAEKALFIGRNIAILRKKGSIKTQQILEIID------------
--SE--KGMAEAKRTLEA-LGLQPSEQDCCRVQQVCRMVLSRAAALCATGLAAILSYMCRSR-ELEHLSV
NVAVDGDLYQGQ-SRFGEILQSVTGLLAPECSATLLPSVDGTGKGAAMVTAVALRLAAHRREVNELLAPL
RLSRADLEHVQALMRQEMELGLKQETNDTS-SLRMLPTYVCGTPDGT----------------------
---EQGDFLALDLGGTNFRVLVVRIAED---GIRMASEIYIIPINIMQGTGEA-------LFDHIVNCIA
DFQLKHELMGQ-VLPLGFTFSFPCQQLGLDKAVLLSWTKGFSASGCVGQDVVQLLRQAAQRKQ----YSG
LKVVA-----VVNDTVGTMMSCGHEDPKCEIGLIVGTGTNACYMEEM---QNVGTVE------------
--GD------EG---------------------RMCINMEWGAFGDNGC-LNDFFTDFDRLVDEKTINPG
RQR-FEKLISGMYLGEIVRHILLTLVEKQLLFQGRPCPKLHTKDIFQTKFLSEI----------------
------------DGLAVQHVQTILQNLEL-KASFEDSALVREVCQTVSLRAAQLCAAGLAAVAEKMRQSR
GLPHLAVTVGVDGTLYKMH-----------PSFSKHIEQTLKYLAPHCAVTF---------LRSEDG---
----------SGKGAALVAAVACRGAE------------------------
>XP_047936329|Anser cygnoides|HK3

LMLLTPSLEKLQAVKCRMLEAMDKGL-SRQTHAQAT--VRMLPTFVCSTPDGTEKGDFMVVELC--QSHV
RTLFVTLLGD---GN-QS--PQLMYKIFDMPADMLQGTGEA---------------LFDFIAQCVHKFLE
EI-----GSL---QHRLPLGFVFPFSCRQTGLG------------------------K------------
-----------AELISWSKGFQCSDVEGKDVVQLLQAAINKL----------E-----------------
--------------LYHVQVIALLNDTVGTMMASSWGGKPCEIALVT-----------------------
-----G-----TGTNSCFMAEAELV-EVVEESN--GRMCVNSEWGCFGDDGALGDVLTPYDHRVDQESSN
PGERRFEKMVGGLYLGEIVRHAVLMLAAEQALFVGASTSVLGAKDAFKTQQVLEIMD------------
--NE--TGMAKARSALEA-LGLQPSERDCCRVQQICRAVVSRAAALSATGLAAVLSLMCRSR-ELERLVV
NVGVDGELYRAH-ARFGEILQSVAGLLAPECTATLVPSADTNGLGVAMVTAVAQRLLTQRHEVEQLLAPL
RLSRADLERVQALMRQEMERGLGRDSNANA-SVRMLPTYVCNTPDGT-----------------------
---ERGDFLALDLGGTNFRVLVVRVMQE---GIHMASEIYVVPTSVMQGTGEA-------LFDHIIECIM
DFQTKQNLMEQ-VLPLGFTFSFPCQQLGLDKAVLLTWTKGFSASGCVGQDVVQLLRDAAQRKQ----HVA
LKVVA-----VVNDTVGTMMACGYDDPKCEIGLIVGTGTNACYMEER---KNVGTVE------------
--GD------EG---------------------RMCINMEWGAFGDNGC-LDDIFTTFDRMVDEKTINAG
KQR-FEKLISGMYLGEIVRHILLAMVDKQLLFRGKPSLKLQTRDIFQTKFLSTIET-------------
------------DGLALQQVQGILRDLEL-DASFEDSMLVREVCQTVSLRAAQLCAAGLAAVVEKMRENR
GLAQLAITVGVDGTLYKMH-----------PHFSQNLQQMLQDLAPNCAVTF---------LQSEDG---
----------SGKGAALVAAVACRETS----P--------------------
>NXS59539|Brachypteracias leptosomus|HK3
LLELTIPLETLQAVKGHMLQAMRKGL-SRQTHAQAN--VQMLPTYICSTPNGTEKGDLLVVELC--QSRV
RTLWVTLLGD---GN-QS--PKVMYKIFDMPEDIKQGKGEA---------------LFDFIAQCVRQFLA
GI-----GSP---QHRLPLGFVFPFNCKQTQLD------------------------K------------
-----------AEFICWSKDFTCSDMEGKDVVQLLQSAINKQ----------E-----------------
--------------LYHVDVVALVNDTVGTMMTCSTRRKPCEVALIV-----------------------
-----D-----KGTNCCFMAEAQEV-ETVDETS--GQMCVNTEWGYFGDDGILGNIVTPYDQHVDQESSN
PGEKRFEKLVGSLYLGEIVRHMLITLATEKALFVGSSVAALMTKDVLKTHQILEITN------------
--NE--EGMDEAKRTLEA-LGLQPSERDCCRVQQVCRVVVNRAAALCAAGLAAILSHMCQSR-ELERLVV
NVAVDGELCQSHTSRFGEILQSVTGLLAPECTFTLVPSVDGTGQGTAIVTAVALRLAAKRCEVNKLLAPL
WLSRADLERVQALMRREMELGLGRETSTSA-SVRMLPTYVRGTPDGT-----------------------
---ERGEFLALDLGGTNFRVLVVHVEDD---GIRVANEIYVIPTTITQGTGEA-------LFDHIVECIM
DFQLKQNLMDQ-VLPLAFTFSFPCQQLGLDKAVLLSWTKGFSASGCEGEDVVQLLREAAQRKQ----HSG
LNVVA-----VVNDTVGTMMSCGFDDPKCEIGLIVGTGTNACYMEEM---QNVGTVE------------
--GD------QG---------------------RMCINMEWGAFGDNGC-LDDFFTSFDRMVDEKTINPG
KQR-FEKLISGMYLGEIVRHILLALVEKQLLFCGKSSPKLQTKDIFQTKFLSTIEG-------------
------------DGLALQQVRSILQDLEL-EASIEDSVLVREVCQTVSLRAANLCAAGLAAVVEKMRENR
GLAQLAVTVGVDGTLYKLH-----------PHFTQNLQQMLQDLAPNCTVTF---------LQSQDG---
----------SGKGAALVAAVACRGTE----P--------------------
>NXK17988|Arenaria interpres|HK3
LLALTIPLDLLQMVKARMMLAMNKGL-NRQTHAQAT--VKMLPTYICSTPNGTENGDVLVVELR--QSHV
RILLMTLLGD---GN-CS--PQLTDKIFDVPEDLTQGTGEA---------------LFDFIAQCLRQFLG
DM-----CNS---KERLPLGFVFPFSCRQTRLD------------------------K-----------
-----------AELISWSKGFSCSDVEGKDVVQLLQLAINKQ----------E-----------------
--------------LSQVEVVALMNDTVGTMMTCSTREIPCEVALIV-----------------------
-----D-----KGTNSCFMADAQHV-QTAEETS--GRMCVNTEWGCFGDDGALNDILTPYDQHVDEESSN
PGVKRFEKLVGSLYLGDIVRHALIALAAEKVLFIGSNVTVLGTKGVLTTQQVVDIID------------
--YE--EGMTKARMTLEA-LGLRPSERDCCRVQQVCRAVVSRAAALCAAGLAAILSLISQSR-GLERLMI
NVAVDGELYRPQ-GRFKEILQSVTGLLAPECLISLKLSVDGTGRGAAMVTAVALRLADQRREVNQLLAPL
RLTRADLERVQALMRQEMDLGLGQETNANA-SVRMLPTYVRGTPDGT-----------------------
---ERGEFLALDLGGTNFRVLAVRVAEK---GIRIASQIYAIPTNIMQGTGEA-------LFDHIVDCIM
DFQMKQDLKGQ-VLPLGFTFSFPCKQLGLDKAVLLNWTKGFSASDCVGKDVVQLLREAAQRKQ----HLA
LKVVA-----VVNDTVGTMMSCGYDDPECEIGLIVGTGTNACYMEEM---KNVGTVE------------
--GE------QG---------------------RMCINMEWGAFGDNGC-LDHLFTEFDRMVDEKTINPG

```
KQR-FEKLISGMYLGEIVRHILLALVEKEILFQDKACPTLQNRDIFKTKFLSTIEM-------------
-VVLTMSLVRGSDGLALTKVQAILRDLGL-QANFEECSLVREVCQTVSLRAAQLCAAGLAAVVEKMRENR
GLGQLAVTVGVDGTLYKMH-----------PCFSQHLQQTLQDLAPNCTVTF---------LLSEDG---
----------SGKGAALVAAVAVREAE-------------------------
>NXJ66512|Rostratula benghalensis|HK3
LLALTLPLEVLQMVKDRMKEAMNKGL-SRQTHAQAN--VQMLPTYICSTPNGTENGNVLVVELC--QSHV
RILLTTLLGN---GN-NS--PQLTDKIFDMPGDITQGTGEA---------------LFDFIAQCLRQFLE
SN-----DVS---NDYLPLGFVFPFSCKQTRLD-----------------------K-----------
-----------AELISWSKGFSCSDVEGKDVVQLLQSAINKQ----------E-----------------
--------------LSQVKVVALLNDTVGTMMTCSTREKPCEVALIV----------------------
-----D-----KGTNSCFMAEAQYV-QIVEETS--GRMCVNTEWGCFGDDGALNDILTPYDQHVDQESSN
PGLKRFEKLVGSLYLGDIVRHALIALATEKALFIGSNVTALETKGVLKTQQVLEIIN------------
--NE--EGMTKARMTLEA-LGLRPSERDCCRVQQVCRAVVSRSAALCAAGLAAILSLIRQSR-GLERLMI
NVAVDGELYRAH-ARFREILQSVAGLLAPECMASFVPSMDGSGWGAAMVTAVALRMAQQRCEVDKLLAPL
RLSRADLERVQALMRQEMDLGLDQETNANS-SVRMLPTYVRGTPDGS----------------------
---ERGEFLALDLGGTNFRVLVVRLGEN---GIHMASEIYVIPTNIMQGTGEA-------LFDHIIDCIM
DFQLKQDLMEQ-VLPLGFTFSFPCKQLGLDKAVLLSWTKGFSASECVGKDVVQLLREAAQRKQ----HLA
LKVVA-----LVNDTVGTMMSCGYDDPKCEIGLIVGTGTNACYMEEM---QNVGTVE------------
--GE------QG--------------------RMCINMEWGAFGDNGC-LDHIFTEFDRMVDEKTINPG
KQR-FEKLISGMYLGEIVRHILLALVEKRVLFHDMPCFTLQSKDIFKTKFLSTIEI-------------
------------DGLALKQVQSILRDLDL-HATFDDSVLVREVCQTVSLRAAQLCAAGLAAVVEKMRENR
GLSQLAVTVGVDGTLYKMH-----------PCFSQHLQQTLQDLAPNCTVTF---------LQSEDG---
----------SGKGAALVAAVAVR----------------------------
>NXW42121|Nyctiprogne leucopyga|HK3
LLALTPPLETLKVVKGRMLQAMRKGL-SRQTHAQAS--VRMLPTYICSTPNGTEKGDYLVVELC--QSHV
RTLWVTLLGD---GN-QS--PQVMHEIFIMPADITQGKGET---------------LFDFIAQCVHQFLS
VL-----GNP---QRRLPLGFVFPFSCRQTGLD-----------------------K-----------
-----------AELISWAKGFSCSDVEGQDVVQLLQSAINKQ----------E-----------------
--------------LHHVDVIALMNDTVGTMMTCSMGGKPCEVALIM----------------------
-----D-----KGTNACFMAEAHQV-EMAEESS--GRMCVNTEWGCFGDDGTLSDIMTPYDQRVDQESSN
PGQKRFEKLVGSLYLGEIVRHALTALAVEKALFVGSNITVLRSKDVLKTQQVLEIID------------
--KE--EGTAKARKVLMA-LGLQPSEQDCSRVQQICRAVVSRAAALSAAGLAAILSHMRQSR-ELKQLVV
NVGVDGELYRSH-ARFGEIVQSVTGLLAPECTVTLVPSVDSTGQGAAIVMAVAVRLAAQRREVDQLLAPL
RLSRTDLQRVQALMRREMELGLGRETNANA-SVRMLPTYVCGTPDGT----------------------
---ERGEFLALDLGGTNFRVLVVHVERD---GIRMASEIYVIPTTIMQGTGEA-------LFNHIVECIM
DFQLKQDLMEQ-VLPLGFTFSFPCQQLGLDKAVLLTWTKGFNASDCVGQDVVQLLRDAAQRKQ----HFG
LKVVA-----VVNDTVGTMMSCGYDDPKCEIGLIVGTGTNACYMEEM---RNVGTVE------------
--GD------QG--------------------RMCINMEWGAFGDNGC-LDDIFTIFDQLVDEKTINAG
KQR-FEKLISGMYLGEIVRHILLSLVEKQVLFCGKPCPKLQTRDIFKTKFLSTIEMVMPPPSRGMGTVIG
MVVLMVPLVWGSDGLALRQVRAILQELEL-QASFEDSMLVREVCQTVSLRAAQLCAAGLAAVVEKMRENR
GLAQLAVTVGVDGTLYKMH-----------PRFSQHLQQTLKDLAPNCSVTF---------LQSEDG---
----------SGKGAALVAAVACRGAE----P--------------------
>NWQ87670|Burhinus bistriatus|HK3
LLALTVPLEMLQAVKGRMLQAMRKGL-SRQTHAQAN--VRMLPTYICSTPDGTEKGDLLVVELC--QSHV
RTLWMTLLGD---GN-QR--PQVTNKIFDMPGDIMQGKGEA---------------LFDFIAQCVRQFLA
GI-----SSP---QHRLPLGFVFPFSCRQTRLD-----------------------K-----------
-----------AELISWSKGFSCSDVERKDVVQLLQSAINKQ----------E-----------------
--------------LSHVDVVALMNDTVGTMMTCSIGEKPCEVALVVGEQRASHVPGWWLLVGGLHWCSS
PLLPAD-----KGTNSCFMAEAQQV-EMAEETS--GRMCVNTEWGCFGDDGTLSDILTPYDQRVDQESSN
PGEKRFEKLVGSLYLGEIVRHALTALAAEKALFIGSSVAVLGTKGVLKTQQVLEIID------------
--NE--EGMAKARKALED-LGLRPSERDCCRVQQICRAVVSRAAALCAAGLAAILSHMCQSR-ELDRLVV
NVGVDGELYRGH-SRFREILQSVTGLLAPECMATLLPSVDGTGRGAAMVTAVALRLAAHRREVDRLLAPL
```

RLSRTDLERIQALMRREMELGLGQESNANA-SIHMLPTYVRGTPDGT----------------------
---ERGEFLALDLGGTNFRVLVVRVAQD---GIRMASEIYVIPTAVMQGTGEA-------LFDHIIECIM
DFQLKQDLMER-VLPLGFTFSFPCQQLGLDKAVLLSWTKGFSASGCVGQDVVQLLREAAQRKQ----HLA
LKVVA-----VVNDTVGTMMSCGYDDPKCEIGLIVGTGTNACYMEEM---QNVGTVD-------------
--GE------QG---------------------RMCINMEWGAFGDNGC-LDDIFTDFDRLVDEKTINAG
KQR-FEKLISGMYLGEIVRHILLALVEKQLLFRSKPCPKLQTRDIFKTEFLSTIEI--------------
------------DGLALRQVCAILQDLEL-QASFEDSVLVREVCQTVSLRAAQLCAAGLAAVVEKMRENR
GLARLAVTVGVDGTLYKMH-----------PRFSQHLQQMLQDLAPNCTVTF---------LQSEDG---
----------SGKGAALVAAVACCGAE----P--------------------
>NXV43473|Uria aalge|HK3
LLALTIPLEMLQVVKGRMMQAMSNGL-SRQTHAQAN--VQMLPTYICSTPNGTEKGNVLVAELC--HSHV
RILLMTLLGD---GN-HN--PHVTDKIFDIPGDSMQGPGEA---------------LFDFIAQSVRQFLS
GI-----TSA---EERLPLGFVFPFSCKQTQLN------------------------K------------
-----------AELLSWSKGFNCSDVEGKDVVQLLQSAINKQ----------E-----------------
--------------LFHVEVVALMNDTVGTMMTCSTRERPCEVALVV-----------------------
-----D-----KGTNSCFMAEAQYV-DMVEETS--GRMCINTEWGSFGDDDALSDILTPYDQRVDQESSN
PGKKRFEKLVGSLYLGDIVRHTLMALSAEKALFIGSNVTVLGTKGALETQQVLEIIN-------------
--NE--EGMAKARAALEA-LGLRPSERDCCRVQQICRAVVSRSAALCAAGLAAILSYIRHSR-GLERLVI
NVAVDGELYRAH-SRFGEILQSVTGLLAPECMAALVPSVDGTGRGAAIVTAVALRLANQRREMDQLLAPL
RLSRADLEHVQALMRREMELGLGQETNANS-SVRMLPTYVRGTPDGTGEGWGPGEGGPGEAVAPLMPPCW
LPAERGEFLALDLGGTNFRVLVVRVAED---GIRMASEIYLIPTDIMQGTGEA-------LFDHIIDCIM
DFQLKQDLMEQ-VLPLGFTFSFPCQQLGLDKAVLLTWTKGFSASGCVGEDVVQLLREAAQRKQ----HLA
LKVVA-----VVNDTVGTMMSCGYDDPKCEIGLIVGTGTNACYMEET---QNVGTVE-------------
--GE------QG---------------------RMCINMEWGAFGDNGC-LDHIFTDFDRLVDEKTINAG
KQR-FEKLISGMYLGEIVRYILLALVEKQVLFRGKPCPRLQTRDIFKTKFLSTIEI--------------
------------DGLALQQVQAILQDLEL-DASFEDSVLVREVCQTVSLRAAQLCAAGLAAVVEKMRENR
GLSQLAVTVGVDGTLYKMH-----------PCFSQHLQKMLRELAPNCTVTF---------LQSEDG---
----------SGKGAALVAAVACRGAE-------------------------

**S2.** Phylogenetic files for the reconciled 100 kDa HK tree.

Reconciled tree for branch support:


```
(((((((Erpetoichthys calabaricus|HK1|uGyuBPeKnL:0.042827,(((Thalassoph
ryne amazonica|HK1|JsEOEDBkRa:0.048126,(Doryrhamphus excisus|HK1|cALFu
wyBrJ:0.017496,((Pelmatolapia mariae|HK1|kIEIvXJyGq:0.031923,Acanthoch
romis polyacanthus|HK1|BTvAaCUYBV:0.02435)74:0.003269,((Centropristis
striata|HK1|MsaUHPpxrS:0.008969,Pseudoliparis swirei|HK1|qSMnnuMCBW:0.
067126)100:0.026092,Micropterus salmoides|HK1|mczuyQEJNy:0.023986)81:0
.008495)65:0.0061)72:0.009603)95:0.017616,Lampris incognitus|HK1|PuwQU
XYfgX:0.030171)83:1e-06,Gadus macrocephalus|HK1|LWxXzurkMl:0.057181)10
0:0.088159)100:0.030805,((((((((Opisthocomus hoazin|HK1|WeUlkjVgVj:0.
001456,(Athene cunicularia|HK1|XfWuHJYvcU:0.001456,((Myiozetetes cayan
ensis|HK1|emzIUjIaqN:1e-06,(Corvus cornix cornix|HK1|rKHLvnvBYe:0.0029
24,Malurus melanocephalus|HK1|MlNzhWNIZC:1e-06)91:0.00146)100:0.006877
,Melopsittacus undulatus|HK1|PCYyNVKAZi:0.002476)100:0.003827)100:1e-0
6)100:0.010489,Dromaius novaehollandiae|HK1|RhrdaRtdfb:0.011616)100:0.
010016,Alligator sinensis|HK1|NuiileKrUF:0.027384)86:0.008391,Pelodisc
us sinensis|HK1|tqUNgzXpKh:0.023037)91:0.008235,Pogona vitticeps|HK1|J
BzVERGLeG:0.037744)98:0.009889,(Tachyglossus aculeatus|HK1|EipBvNDojf:
0.030522,((Monodelphis domestica|HK1|jKrGuSGcUc:0.126861,Dromiciops gl
iroides|HK1|kAxRfmrNyH:0.045883)100:0.134336,(((Homo sapiens|HK1|HikRH
BmLYx:0.022674,Phodopus roborovskii|HK1|iXEhcskcgQ:0.067077)80:0.01386
7,(Myotis myotis|HK1|ArELCCLoEK:0.064895,Lagenorhynchus albirostris|HK
1|wLZZSHxlti:0.044937)80:0.007598)80:0.016086,Orycteropus afer afer|HK
1|ARXWtKFged:0.047294)100:0.05847)100:0.056927)99:0.038663)98:0.023838
,Rhinatrema bivittatum|HK1|ADrBoRHQCz:0.051733)73:0.015923,Protopterus
 annectens|HK1|OSnuUfJPUx:0.111823)54:0.00772,Latimeria chalumnae|HK1|
OgOKnwCKyc:0.106276)68:0.014831)100:0.050126,Leucoraja erinacea|HK1|gF
eVmhUjNL:0.160266)98:0.048348,(((Polyodon spathula|HKDC1|FblhIVtssc:0.
106835,((Amia calva|HKDC1|uqqMvJNjEt:0.089454,Atractosteus spatula|HKD
C1|WZsmwTEtMw:0.09822)92:0.015618,(((Scleropages formosus|HKDC1|TPLznO
DygH:0.134745,Brienomyrus brachyistius|HKDC1|bZDMaqdqSA:0.178449)97:0.
051788,(((((Ictalurus punctatus|HKDC1|TGRsMCfawE:0.069373,Electrophoru
s electricus|HKDC1|vtZUynMZlo:0.092751)100:0.026504,Paramisgurnus dabr
yanus|HKDC1|vrcEEbpAUS:0.181241)82:0.013955,Chanos chanos|HKDC1|YHNqFM
cxNZ:0.138704)69:0.023831,(Clupea harengus|HKDC1|bFsdyTfXGl:0.113984,D
enticeps clupeoides|HKDC1|QIqYcckl UB:0.104353)54:0.019469)77:0.01788,(
Hypomesus transpacificus|HKDC1|aLetOARBie:0.160867,(((Myripristis murd
jan|HKDC1|BsaruMxntV:0.13315,(Thunnus albacares|HKDC1|ZmGyNEPRji:0.079
9,(((Centropristis striata|HKDC1|RSdDyCyTBC:0.081794,Epinephelus fusco
guttatus|HKDC1|AeFKeZcVXW:0.095916)76:0.013063,((Pseudoliparis swirei|
HKDC1|iCcmbGeuop:0.125383,Anoplopoma fimbria|HKDC1|VaFZQoiiBx:0.088181
)99:0.024085,((Cottoperca gobio|HKDC1|SdfTRkeJir:0.056577,Eleginops ma
clovinus|HKDC1|GQwKbHfHXa:0.145751)97:0.023319,Sebastes umbrosus|HKDC1
|ElnqstjQmQ:0.079493)88:1e-06)90:0.008725)92:0.027025,((Salarias fasci
atus|HKDC1|oPtZfrHMEn:0.21552,Gouania willdenowi|HKDC1|scCjzbmRDo:0.17
2378)94:0.042332,((Oryzias latipes|HKDC1|APKqDVmZwa:0.164753,Cololabis
 saira|HKDC1|dSBRkeWooW:0.140333)82:0.026185,(Nothobranchius furzeri|H
KDC1|vthLsNmGae:0.147621,((Fundulus heteroclitus|HKDC1|XFXAGhmEdt:0.05
```

6429,Cyprinodon tularosa|HKDC1|BxkPqfOcsU:0.107857)97:0.019443,Poecilia formosa|HKDC1|gXFNGdeqfM:0.103393)100:0.065108)85:0.031436)97:0.041641)89:0.028514)67:0.009116)100:0.129655)71:0.022445,Lampris incognitus|HKDC1|hNHyWARdjX:0.162052)73:0.024504,Gadus morhua|HKDC1|ygnXzsWQlY:0.2318)100:0.160587)91:0.075133)43:0.019401)43:0.0101,(Megalops cyprinoides|HKDC1|JiTDiqJWqq:0.063872,(Anguilla anguilla|HKDC1|hrkEPumxwE:0.105985,Conger conger|HKDC1|KyPRaGQWJn:0.144742)100:0.107401)83:0.02612)99:0.04707)100:0.076352)100:0.086505,(Latimeria chalumnae|HKDC1|kAoIdxkVCy:0.133948,((((Dromiciops gliroides|HKDC1|oKbmBUVIIJ:0.104678,(((Echinops telfairi|HKDC1|CTvbLSGChX:0.158559,Chrysochloris asiatica|HKDC1|TPMNSTrvzb:0.112338)99:0.025103,Orycteropus afer afer|HKDC1|IjoVSXORKf:0.041286)99:0.016377,(((Meles meles|HKDC1|cmKXcqnZVo:0.061083,Myotis brandtii|HKDC1|gNtcFCJNRn:0.053214)70:0.006724,(Suncus etruscus|HKDC1|SYiclaDRrZ:0.10866,Erinaceus europaeus|HKDC1|dAnfsJuzzw:0.051941)81:0.015025)68:0.006833,Homo sapiens|HKDC1|SJSMcTzbZx:0.054838)87:0.009068)100:0.049098)89:0.021471,Ornithorhynchus anatinus|HKDC1|YgLNdLnanC:0.104949)100:0.07321,(((((((Gavia stellata|HKDC1|XGlDGGvmcy:0.011982,((Neopsephotus bourkii|HKDC1|cfQkjMGkNp:0.018884,(Acanthisitta chloris|HKDC1|yZBvKdwupL:0.041304,(Zonotrichia albicollis|HKDC1|uymusyCedr:0.100731,Malurus melanocephalus|HKDC1|FzUHEqEplI:0.015182)94:0.014198)93:0.012751)80:0.00658,Athene cunicularia|HKDC1|umPqkFBiYK:0.010564)49:1e-06)55:1e-06,Grus americana|HKDC1|eAvPfRizJy:0.014941)61:0.004363,Calypte anna|HKDC1|utbBrXfyrJ:0.055557)71:0.011448,Coturnix japonica|HKDC1|luLQhYohCx:0.040949)100:0.088984,Alligator mississippiensis|HKDC1|wTCBdToqMz:0.048152)87:0.010923,Emys orbicularis|HKDC1|iqhvTiaZIT:0.063257)90:0.006857,((Podarcis muralis|HKDC1|IVIgXHPWxQ:0.072952,Python bivittatus|HKDC1|nahIDFGyme:0.095862)98:0.011449,Euleptes europaea|HKDC1|xIoXkCYtzw:0.069898)100:0.033993)100:0.028546)98:0.021759,(((Bufo bufo|HKDC1|EijMExSFZR:0.065147,Rana temporaria|HKDC1|PNiWpIvXnO:0.125618)100:0.037841,Xenopus tropicalis|HKDC1|cKpdtvXrTj:0.090193)100:0.117427,Rhinatrema bivittatum|HKDC1|DYIUurqojh:0.16561)92:0.027073)100:0.072769)64:0.017236)100:0.069519,(Stegostoma fasciatum|HKDC1|PxpsLTmAmN:0.350636,(Pristis pectinata|HKDC1|FrKkoLgmfd:0.060099,Dasyatis sabina|HKDC1|oOYqDdZzSb:0.19203)100:0.174183)100:0.206247)100:0.100981)94:0.067913,((((Protopterus annectens|HK2|EUsHUvJhlS:0.139272,((Spea bombifrons|HK2|kuBDQaekeh:0.080905,Rhinatrema bivittatum|HK2|rrWWLZdCcr:0.105941)54:0.006914,((Tachyglossus aculeatus|HK2|PsAKGFFUJM:0.172896,(Vombatus ursinus|HK2|VPnyoqRgzk:0.041309,((Homo sapiens|HK2|pkVOClMpeg:0.035019,((Hyaena hyaena|HK2|yOGOMOiisv:0.017576,Budorcas taxicolor|HK2|MwgMQvftWF:0.023783)92:0.007814,Galemys pyrenaicus|HK2|vgIzhLDEor:0.044853)82:0.005585)97:0.009985,Echinops telfairi|HK2|xzlgnvJvEl:0.043699)100:0.02739)100:0.036854)100:0.051232,((Terrapene carolina triunguis|HK2|TlpAKSJfbS:0.050967,(Alligator sinensis|HK2|KcAcsyJVLt:0.051181,((((Accipiter gentilis|HK2|ONiFiGZKDZ:0.010426,((Corapipo altera|HK2|gUAxZacVZH:0.025217,((Ptilonorhynchus violaceus|HK2|omwJgFJSDB:0.008377,(Orthonyx spaldingii|HK2|bjdmRLkbSC:0.008402,(Drymodes brunneopygia|HK2|hCroEBgKFM:0.000623,Chaetops frenatus|HK2|pNSXxqafgA:0.027434)98:0.006169)80:0.003031)78:0.002062,Atrichornis clamosus|HK2|JvNvvnoBoc:0.003184)72:0.003932)100:0.018456,Lathamus discolor|HK2|FJwoHpOswW:0.012488)84:0.007066)47:1e-06,((Gavia stellata|HK2|wKENSYxIvv:0.002969,Calonectris borealis|HK2|aWMhpJyWGa:0.002976)90:0.001506,Chroicocephalus ridibundus|HK2|VMOkiKMwtj:0.011957)61:0.00154)83:1e-06,Columba livia|HK2|bXWVFYgEU

U:0.008911)97:0.009515,Dromaius novaehollandiae|HK2|GAdXLAPtGH:0.04538
7)97:0.020237)62:1e-06)66:0.007021,(Eublepharis macularius|HK2|vGXLdoq
ggE:0.052086,(Pseudonaja textilis|HK2|DkEEDvenrP:0.182319,Elgaria mult
icarinata webbii|HK2|ZMtHoKJean:0.076352)100:0.033583)100:0.046569)97:
0.018503)85:0.017823)76:0.024773)39:0.008293,Latimeria chalumnae|HK2|I
KmASFgAXD:0.131939)49:0.006494,(Polypterus senegalus|HK2|RXwXFroLbj:0.
087665,(Polyodon spathula|HK2|PcUmlCXAsz:0.124769,((((((((((Gasteroste
us aculeatus aculeatus|HK2|twGyJDlrlX:0.08488,Eleginops maclovinus|HK2
|HmjDiAqQqY:0.080872)94:0.013561,Centropristis striata|HK2|XcZnpOSbXX:
0.064787)82:0.01489,((Sparus aurata|HK2|pXmfIKjGzQ:0.052502,Takifugu f
lavidus|HK2|DewUMDFyVe:0.197991)96:0.027372,(Xyrichtys novacula|HK2|CD
wEEimyOz:0.073151,Labrus mixtus|HK2|uyzlfzIdCF:0.074346)100:0.043741)9
1:0.020714)70:0.011446,((((Gouania willdenowi|HK2|mvMbYoycnf:0.155049,
Salarias fasciatus|HK2|buwEgcGvFp:0.092607)85:0.037202,Parambassis ran
ga|HK2|WCTACmqHwb:0.074222)38:0.001866,(Maylandia zebra|HK2|mwzhgngitf
:0.113708,((Cololabis saira|HK2|rKDgEdBXSe:0.126143,Oryzias melastigma
|HK2|tChcjADeIz:0.125624)77:0.018733,(Nothobranchius furzeri|HK2|mTMdG
JVSyy:0.078747,Kryptolebias marmoratus|HK2|ejpzKAiXzB:0.078091)96:0.04
0241)71:0.024425)45:0.008679)95:0.030966,(Echeneis naucrates|HK2|feLDl
jjCjp:0.118394,Seriola dumerili|HK2|fLDqICFZTw:0.035219)91:0.019877)82
:0.008409)55:0.003451,((Synchiropus splendidus|HK2|YUaKrzfYXJ:0.222826
,(Entelurus aequoreus|HK2|HJiPJMaXCk:0.122121,Phycodurus eques|HK2|pGb
tdXQYlk:0.100637)100:0.075363)94:0.020728,Thunnus maccoyii|HK2|oTODkFQ
bVt:0.052924)97:0.023575)98:0.040768,Thalassophryne amazonica|HK2|FViz
izcrOp:0.241586)100:0.061396,Lampris incognitus|HK2|SvQzutreJy:0.10933
)100:0.034378,Merluccius polli|HK2|fcOSUWMZAk:0.13425)100:0.023958,Sal
velinus namaycush|HK2|YbxquaaXVr:0.111815)70:0.005977,((Denticeps clup
eoides|HK2|qjFPaapWkZ:0.098493,(Engraulis encrasicolus|HK2|SiPVyFcTKH:
0.136163,Alosa sapidissima|HK2|FQIQwDCBql:0.077091)100:0.054204)64:0.0
1208,((Astyanax mexicanus|HK2|UCSURQaOGS:0.06453,(Trichomycterus rosab
lanca|HK2|PeNBFeARln:0.096706,Silurus meridionalis|HK2|XsMCJQiJBO:0.07
0129)100:0.046303)98:0.033694,Xyrauchen texanus|HK2|YSBDvLgDiW:0.10385
1)87:0.009968)78:0.025043)100:0.147162)74:0.023422)99:0.036841)100:0.0
57029,(Callorhinchus milii|HK2|HxvSQZRczB:0.105874,((Scyliorhinus cani
cula|HK2|KartHAlQEB:0.078369,Hemiscyllium ocellatum|HK2|jrOmbuyPcp:0.1
54145)68:0.014444,(Amblyraja radiata|HK2|WUGYLAukGn:0.184299,Dasyatis
sabina|HK2|AJKTmVRJQz:0.070922)80:0.033089)100:0.080567)100:0.050241)1
00:0.081811)44:0.053869,(Protopterus annectens|HK3|NvqwsvPsyX:0.320144
,(((Tachyglossus aculeatus|HK3|AvxvjOUqzE:0.298989,(((Trichechus manat
us latirostris|HK3|nxdYWiMVaR:0.156081,(((Chrysochloris asiatica|HK3|K
CyjjKuQTm:0.079569,Echinops telfairi|HK3|flIbqulDzJ:0.112912)69:0.0134
66,Elephantulus edwardii|HK3|HxnlTjCZOb:0.122379)65:0.007543,Orycterop
us afer afer|HK3|kuGXdlwnDq:0.097764)88:0.016081)99:0.041367,(((Erinac
eus europaeus|HK3|SbEBIJiXGc:0.172974,(Suncus etruscus|HK3|WIGLlbztRS:
0.269263,Talpa occidentalis|HK3|MbnVjEhKCF:0.083482)73:0.021716)64:0.0
21448,(Suricata suricatta|HK3|czgRgyZtFJ:0.072076,(Miniopterus natalen
sis|HK3|jBHUmwqCXu:0.083853,Desmodus rotundus|HK3|dXRmcPxDid:0.105319)
99:0.032772)38:0.007435)27:0.003043,(((Ochotona curzoniae|HK3|UpCmrXEd
ks:0.194381,Oryctolagus cuniculus|HK3|JhPnTeDAUk:0.058749)98:0.042522,
(((Fukomys damarensis|HK3|zgJUZbGElA:0.051128,Cavia porcellus|HK3|fcUj
fYTTiq:0.149711)100:0.050124,(Dipodomys spectabilis|HK3|pfTUUCWeHp:0.1
30777,(Jaculus jaculus|HK3|TWvySMGveq:0.110541,(Nannospalax galili|HK3

|CrmROQqfyG:0.062087,((Grammomys surdaster|HK3|BSLllRRzgE:0.069997,Psa
mmomys obesus|HK3|iPRQzzZeJH:0.10101)95:1e-06,Chionomys nivalis|HK3|oM
WYbiCulg:0.169145)100:0.046204)100:0.029695)94:0.011875)80:0.014882)47
:1e-06,Marmota marmota marmota|HK3|KJwThyPynx:0.074303)76:0.01064)83:0
.015955,((Microcebus murinus|HK3|EMaweTluNF:0.103193,Otolemur garnetti
i|HK3|ZxsMJahiAq:0.080177)28:0.015939,((Callithrix jacchus|HK3|AeeGloz
WKL:0.174635,Homo sapiens|HK3|BFYESbMjCp:0.029952)100:0.020526,Carlito
 syrichta|HK3|PTXqKIbVOb:0.116466)24:0.006289)61:0.012477)82:0.01765)6
9:0.0262)100:0.183963,((Dromiciops gliroides|HK3|fIRXultwTn:0.06181,(A
ntechinus flavipes|HK3|vtWrfmQvsB:0.143088,Trichosurus vulpecula|HK3|w
fGnpkHXaz:0.085759)94:0.01919)90:0.025469,Monodelphis domestica|HK3|wf
OMlvxmMu:0.092493)100:0.154968)100:0.125745)100:0.238306,((Protobothro
ps mucrosquamatus|HK3|YLYYaWThsj:0.371042,Elgaria multicarinata webbii
|HK3|MdNSXAHZft:0.255297)100:0.55215,((((Phasianus colchicus|HK3|vZvlm
TAEnR:0.218987,Anser cygnoides|HK3|RHUMEJUeYm:0.105836)100:0.08187,(Ny
ctiprogne leucopyga|HK3|kZwwNgzPCt:0.13009,((Burhinus bistriatus|HK3|U
lCPfPqcEA:0.05145,((Turnix velox|HK3|rqPQKInsSW:0.295639,Uria aalge|HK
3|IkbPSCMcHs:0.063222)61:0.02397,(Rostratula benghalensis|HK3|ApEQAGwb
CF:0.081485,Arenaria interpres|HK3|ZRJDvVqaYr:0.120128)100:0.065248)10
0:0.06193)95:0.032522,((((((Passer domesticus|HK3|OorMPQmfvK:0.105724,
Regulus satrapa|HK3|pghRAHgkQE:0.090448)73:0.019386,(Acrocephalus arun
dinaceus|HK3|ZdrTnDWctd:0.062305,Sylvia atricapilla|HK3|XgJeOwDcHb:0.0
90871)98:0.031509)98:0.039793,Drymodes brunneopygia|HK3|XiigvmLSSv:0.0
92388)100:0.048404,Climacteris rufus|HK3|DQXSMRkhuo:0.086005)100:0.095
566,(((Xiphorhynchus elegans|HK3|EJMVMuomip:0.076914,Scytalopus superc
iliaris|HK3|IdEaORUUBk:0.096661)75:0.016855,Grallaria varia|HK3|XpgMnW
JnrP:0.113128)100:0.056754,(Pitta sordida|HK3|pxizwQDLmo:0.13872,(Smit
hornis capensis|HK3|CYlBlIZKVA:0.085856,Calyptomena viridis|HK3|RPoOpE
Knrc:0.080459)100:0.050806)100:0.063332)100:0.025179)100:0.147054,(Rhi
nopomastus cyanomelas|HK3|PbQqymDllh:0.19345,Brachypteracias leptosomu
s|HK3|ajpcOmWbRH:0.137607)84:0.018573)80:0.027205)81:0.008561)91:0.040
854)100:0.130129,(Dromaius novaehollandiae|HK3|jOWGeTwolc:0.070103,Cry
pturellus undulatus|HK3|tLTLumxuGH:0.13036)93:0.056697)100:0.345781,De
rmochelys coriacea|HK3|KaFweMEudV:0.315832)92:0.097151)100:0.14141)100
:0.261851,((Pseudophryne corroboree|HK3|qSucPCjdGI:0.246067,(Rana temp
oraria|HK3|zOzfeCXiPf:0.099756,Nanorana parkeri|HK3|JEOPzKCIiz:0.05835
7)100:0.113625)100:0.141915,Bombina bombina|HK3|ExGhVFjtQF:0.175774)10
0:0.235897)83:0.077389)100:0.273291)1:0.402186,Lethenteron reissneri|H
K1|dGLCSoWnUb:0.402186);

Reconciled tree for duplication and speciation events:

((((((Erpetoichthys calabaricus|HK1|uGyuBPeKnL:0.042827,(((Thalassoph
ryne amazonica|HK1|JsEOEDBkRa:0.048126,(Doryrhamphus excisus|HK1|cALFu
wyBrJ:0.017496,((Pelmatolapia mariae|HK1|kIEIvXJyGq:0.031923,Acanthoch
romis polyacanthus|HK1|BTvAaCUYBV:0.02435)S:0.003269,((Centropristis s
triata|HK1|MsaUHPpxrS:0.008969,Pseudoliparis swirei|HK1|qSMnnuMCBW:0.0
67126)S:0.026092,Micropterus salmoides|HK1|mczuyQEJNy:0.023986)S:0.008
495)S:0.0061)S:0.009603)S:0.017616,Lampris incognitus|HK1|PuwQUXYfgX:0
.030171)S:1e-06,Gadus macrocephalus|HK1|LWxXzurkMl:0.057181)S:0.088159
)S:0.030805,((((((((Opisthocomus hoazin|HK1|WeUlkjVgVj:0.001456,(Athe

ne cunicularia|HK1|XfWuHJYvcU:0.001456,((Myiozetetes cayanensis|HK1|em
zIUjIaqN:1e-06,(Corvus cornix cornix|HK1|rKHLvnvBYe:0.002924,Malurus m
elanocephalus|HK1|MlNzhWNIZC:1e-06)S:0.00146)S:0.006877,Melopsittacus
undulatus|HK1|PCYyNVKAZi:0.002476)S:0.003827)S:1e-06)S:0.010489,Dromai
us novaehollandiae|HK1|RhrdaRtdfb:0.011616)S:0.010016,Alligator sinens
is|HK1|NuiileKrUF:0.027384)S:0.008391,Pelodiscus sinensis|HK1|tqUNgzXp
Kh:0.023037)S:0.008235,Pogona vitticeps|HK1|JBzVERGLeG:0.037744)S:0.00
9889,(Tachyglossus aculeatus|HK1|EipBvNDojf:0.030522,((Monodelphis dom
estica|HK1|jKrGuSGcUc:0.126861,Dromiciops gliroides|HK1|kAxRfmrNyH:0.0
45883)S:0.134336,(((Homo sapiens|HK1|HikRHBmLYx:0.022674,Phodopus robo
rovskii|HK1|iXEhcskcgQ:0.067077)S:0.013867,(Myotis myotis|HK1|ArELCCLo
EK:0.064895,Lagenorhynchus albirostris|HK1|wLZZSHxlti:0.044937)S:0.007
598)S:0.016086,Orycteropus afer afer|HK1|ARXWtKFged:0.047294)S:0.05847
)S:0.056927)S:0.038663)S:0.023838,Rhinatrema bivittatum|HK1|ADrBoRHQCz
:0.051733)S:0.015923,Protopterus annectens|HK1|OSnuUfJPUx:0.111823)S:0
.00772,Latimeria chalumnae|HK1|OgOKnwCKyc:0.106276)S:0.014831)S:0.0501
26,Leucoraja erinacea|HK1|gFeVmhUjNL:0.160266)S:0.048348,(((Polyodon s
pathula|HKDC1|FblhIVtssc:0.106835,((Amia calva|HKDC1|uqqMvJNjEt:0.0894
54,Atractosteus spatula|HKDC1|WZsmwTEtMw:0.09822)S:0.015618,(((Sclerop
ages formosus|HKDC1|TPLznODygH:0.134745,Brienomyrus brachyistius|HKDC1
|bZDMaqdqSA:0.178449)S:0.051788,(((((Ictalurus punctatus|HKDC1|TGRsMCf
awE:0.069373,Electrophorus electricus|HKDC1|vtZUynMZlo:0.092751)S:0.02
6504,Paramisgurnus dabryanus|HKDC1|vrcEEbpAUS:0.181241)S:0.013955,Chan
os chanos|HKDC1|YHNqFMcxNZ:0.138704)S:0.023831,(Clupea harengus|HKDC1|
bFsdyTfXGl:0.113984,Denticeps clupeoides|HKDC1|QIqYccklUB:0.104353)S:0
.019469)S:0.01788,(Hypomesus transpacificus|HKDC1|aLetOARBie:0.160867,
(((Myripristis murdjan|HKDC1|BsaruMxntV:0.13315,(Thunnus albacares|HKD
C1|ZmGyNEPRji:0.0799,(((Centropristis striata|HKDC1|RSdDyCyTBC:0.08179
4,Epinephelus fuscoguttatus|HKDC1|AeFKeZcVXW:0.095916)S:0.013063,((Pse
udoliparis swirei|HKDC1|iCcmbGeuop:0.125383,Anoplopoma fimbria|HKDC1|V
aFZQoiiBx:0.088181)S:0.024085,((Cottoperca gobio|HKDC1|SdfTRkeJir:0.05
6577,Eleginops maclovinus|HKDC1|GQwKbHfHXa:0.145751)S:0.023319,Sebaste
s umbrosus|HKDC1|ElnqstjQmQ:0.079493)S:1e-06)S:0.008725)S:0.027025,((S
alarias fasciatus|HKDC1|oPtZfrHMEn:0.21552,Gouania willdenowi|HKDC1|sc
CjzbmRDo:0.172378)S:0.042332,((Oryzias latipes|HKDC1|APKqDVmZwa:0.1647
53,Cololabis saira|HKDC1|dSBRkeWooW:0.140333)S:0.026185,(Nothobranchiu
s furzeri|HKDC1|vthLsNmGae:0.147621,((Fundulus heteroclitus|HKDC1|XFXA
GhmEdt:0.056429,Cyprinodon tularosa|HKDC1|BxkPqfOcsU:0.107857)S:0.0194
43,Poecilia formosa|HKDC1|gXFNGdeqfM:0.103393)S:0.065108)S:0.031436)S:
0.041641)S:0.028514)S:0.009116)S:0.129655)S:0.022445,Lampris incognitu
s|HKDC1|hNHyWARdjX:0.162052)S:0.024504,Gadus morhua|HKDC1|ygnXzsWQlY:0
.2318)S:0.160587)S:0.075133)S:0.019401)S:0.0101,(Megalops cyprinoides|
HKDC1|JiTDiqJWqq:0.063872,(Anguilla anguilla|HKDC1|hrkEPumxwE:0.105985
,Conger conger|HKDC1|KyPRaGQWJn:0.144742)S:0.107401)S:0.02612)S:0.0470
7)S:0.076352)S:0.086505,(Latimeria chalumnae|HKDC1|kAoIdxkVCy:0.133948
,((((Dromiciops gliroides|HKDC1|oKbmBUVIIJ:0.104678,(((Echinops telfai
ri|HKDC1|CTvbLSGChX:0.158559,Chrysochloris asiatica|HKDC1|TPMNSTrvzb:0
.112338)S:0.025103,Orycteropus afer afer|HKDC1|IjoVSXORKf:0.041286)S:0
.016377,(((Meles meles|HKDC1|cmKXcqnZVo:0.061083,Myotis brandtii|HKDC1
|gNtcFCJNRn:0.053214)S:0.006724,(Suncus etruscus|HKDC1|SYiclaDRrZ:0.10
866,Erinaceus europaeus|HKDC1|dAnfsJuzzw:0.051941)S:0.015025)S:0.00683
3,Homo sapiens|HKDC1|SJSMcTzbZx:0.054838)S:0.009068)S:0.049098)S:0.021

471,Ornithorhynchus anatinus|HKDC1|YgLNdLnanC:0.104949)S:0.07321,(((((
((Gavia stellata|HKDC1|XGlDGGvmcy:0.011982,((Neopsephotus bourkii|HKDC
1|cfQkjMGkNp:0.018884,(Acanthisitta chloris|HKDC1|yZBvKdwupL:0.041304,
(Zonotrichia albicollis|HKDC1|uymusyCedr:0.100731,Malurus melanocephal
us|HKDC1|FzUHEqEplI:0.015182)S:0.014198)S:0.012751)S:0.00658,Athene cu
nicularia|HKDC1|umPqkFBiYK:0.010564)S:1e-06)S:1e-06,Grus americana|HKD
C1|eAvPfRizJy:0.014941)S:0.004363,Calypte anna|HKDC1|utbBrXfyrJ:0.0555
57)S:0.011448,Coturnix japonica|HKDC1|luLQhYohCx:0.040949)S:0.088984,A
lligator mississippiensis|HKDC1|wTCBdToqMz:0.048152)S:0.010923,Emys or
bicularis|HKDC1|iqhvTiaZIT:0.063257)S:0.006857,((Podarcis muralis|HKDC
1|IVIgXHPWxQ:0.072952,Python bivittatus|HKDC1|nahIDFGyme:0.095862)S:0.
011449,Euleptes europaea|HKDC1|xIoXkCYtzw:0.069898)S:0.033993)S:0.0285
46)S:0.021759,(((Bufo bufo|HKDC1|EijMExSFZR:0.065147,Rana temporaria|H
KDC1|PNiWpIvXnO:0.125618)S:0.037841,Xenopus tropicalis|HKDC1|cKpdtvXrT
j:0.090193)S:0.117427,Rhinatrema bivittatum|HKDC1|DYIUurqojh:0.16561)S
:0.027073)S:0.072769)S:0.017236)S:0.069519,(Stegostoma fasciatum|HKDC1
|PxpsLTmAmN:0.350636,(Pristis pectinata|HKDC1|FrKkoLgmfd:0.060099,Dasy
atis sabina|HKDC1|oOYqDdZzSb:0.19203)S:0.174183)S:0.206247)S:0.100981)
D:0.067913,((((Protopterus annectens|HK2|EUsHUvJhlS:0.139272,((Spea bo
mbifrons|HK2|kuBDQaekeh:0.080905,Rhinatrema bivittatum|HK2|rrWWLZdCcr:
0.105941)S:0.006914,((Tachyglossus aculeatus|HK2|PsAKGFFUJM:0.172896,(
Vombatus ursinus|HK2|VPnyoqRgzk:0.041309,((Homo sapiens|HK2|pkVOClMpeg
:0.035019,((Hyaena hyaena|HK2|yOGOMOiisv:0.017576,Budorcas taxicolor|H
K2|MwgMQvftWF:0.023783)S:0.007814,Galemys pyrenaicus|HK2|vgIzhLDEor:0.
044853)S:0.005585)S:0.009985,Echinops telfairi|HK2|xzlgnvJvEl:0.043699
)S:0.02739)S:0.036854)S:0.051232,((Terrapene carolina triunguis|HK2|Tl
pAKSJfbS:0.050967,(Alligator sinensis|HK2|KcAcsyJVLt:0.051181,((((Acci
piter gentilis|HK2|ONiFiGZKDZ:0.010426,((Corapipo altera|HK2|gUAxZacVZ
H:0.025217,((Ptilonorhynchus violaceus|HK2|omwJgFJSDB:0.008377,(Orthon
yx spaldingii|HK2|bjdmRLkbSC:0.008402,(Drymodes brunneopygia|HK2|hCroE
BgKFM:0.000623,Chaetops frenatus|HK2|pNSXxqafgA:0.027434)S:0.006169)S:
0.003031)S:0.002062,Atrichornis clamosus|HK2|JvNvvnoBoc:0.003184)S:0.0
03932)S:0.018456,Lathamus discolor|HK2|FJwoHpOswW:0.012488)S:0.007066)
S:1e-06,((Gavia stellata|HK2|wKENSYxIvv:0.002969,Calonectris borealis|
HK2|aWMhpJyWGa:0.002976)S:0.001506,Chroicocephalus ridibundus|HK2|VMOk
iKMwtj:0.011957)S:0.00154)S:1e-06,Columba livia|HK2|bXWVFYgEUU:0.00891
1)S:0.009515,Dromaius novaehollandiae|HK2|GAdXLAPtGH:0.045387)S:0.0202
37)S:1e-06)S:0.007021,(Eublepharis macularius|HK2|vGXLdoqggE:0.052086,
(Pseudonaja textilis|HK2|DkEEDvenrP:0.182319,Elgaria multicarinata web
bii|HK2|ZMtHoKJean:0.076352)S:0.033583)S:0.046569)S:0.018503)S:0.01782
3)S:0.024773)S:0.008293,Latimeria chalumnae|HK2|IKmASFgAXD:0.131939)S:
0.006494,(Polypterus senegalus|HK2|RXwXFroLbj:0.087665,(Polyodon spath
ula|HK2|PcUmlCXAsz:0.124769,(((((((((Gasterosteus aculeatus aculeatus
|HK2|twGyJDlrlX:0.08488,Eleginops maclovinus|HK2|HmjDiAqQqY:0.080872)S
:0.013561,Centropristis striata|HK2|XcZnpOSbXX:0.064787)S:0.01489,((Sp
arus aurata|HK2|pXmfIKjGzQ:0.052502,Takifugu flavidus|HK2|DewUMDFyVe:0
.197991)S:0.027372,(Xyrichtys novacula|HK2|CDwEEimyOz:0.073151,Labrus
mixtus|HK2|uyzlfzIdCF:0.074346)S:0.043741)S:0.020714)S:0.011446,((((Go
uania willdenowi|HK2|mvMbYoycnf:0.155049,Salarias fasciatus|HK2|buwEgc
GvFp:0.092607)S:0.037202,Parambassis ranga|HK2|WCTACmqHwb:0.074222)S:0
.001866,(Maylandia zebra|HK2|mwzhgngitf:0.113708,((Cololabis saira|HK2
|rKDgEdBXSe:0.126143,Oryzias melastigma|HK2|tChcjADeIz:0.125624)S:0.01

8733,(Nothobranchius furzeri|HK2|mTMdGJVSyy:0.078747,Kryptolebias marm
oratus|HK2|ejpzKAiXzB:0.078091)S:0.040241)S:0.024425)S:0.008679)S:0.03
0966,(Echeneis naucrates|HK2|feLDljjCjp:0.118394,Seriola dumerili|HK2|
fLDqICFZTw:0.035219)S:0.019877)S:0.008409)S:0.003451,((Synchiropus spl
endidus|HK2|YUaKrzfYXJ:0.222826,(Entelurus aequoreus|HK2|HJiPJMaXCk:0.
122121,Phycodurus eques|HK2|pGbtdXQYlk:0.100637)S:0.075363)S:0.020728,
Thunnus maccoyii|HK2|oTODkFQbVt:0.052924)S:0.023575)S:0.040768,Thalass
ophryne amazonica|HK2|FVizizcrOp:0.241586)S:0.061396,Lampris incognitu
s|HK2|SvQzutreJy:0.10933)S:0.034378,Merluccius polli|HK2|fcOSUWMZAk:0.
13425)S:0.023958,Salvelinus namaycush|HK2|YbxquaaXVr:0.111815)S:0.0059
77,((Denticeps clupeoides|HK2|qjFPaapWkZ:0.098493,(Engraulis encrasico
lus|HK2|SiPVyFcTKH:0.136163,Alosa sapidissima|HK2|FQIQwDCBql:0.077091)
S:0.054204)S:0.01208,((Astyanax mexicanus|HK2|UCSURQaOGS:0.06453,(Tric
homycterus rosablanca|HK2|PeNBFeARln:0.096706,Silurus meridionalis|HK2
|XsMCJQiJBO:0.070129)S:0.046303)S:0.033694,Xyrauchen texanus|HK2|YSBDv
LgDiW:0.103851)S:0.009968)S:0.025043)S:0.147162)S:0.023422)S:0.036841)
S:0.057029,(Callorhinchus milii|HK2|HxvSQZRczB:0.105874,((Scyliorhinus
canicula|HK2|KartHAlQEB:0.078369,Hemiscyllium ocellatum|HK2|jrOmbuyPc
p:0.154145)S:0.014444,(Amblyraja radiata|HK2|WUGYLAukGn:0.184299,Dasya
tis sabina|HK2|AJKTmVRJQz:0.070922)S:0.033089)S:0.080567)S:0.050241)S:
0.081811)D:0.053869,(Protopterus annectens|HK3|NvqwsvPsyX:0.320144,(((
Tachyglossus aculeatus|HK3|AvxvjOUqzE:0.298989,(((Trichechus manatus l
atirostris|HK3|nxdYWiMVaR:0.156081,(((Chrysochloris asiatica|HK3|KCyjj
KuQTm:0.079569,Echinops telfairi|HK3|flIbqulDzJ:0.112912)S:0.013466,El
ephantulus edwardii|HK3|HxnlTjCZOb:0.122379)S:0.007543,Orycteropus afe
r afer|HK3|kuGXdlwnDq:0.097764)S:0.016081)S:0.041367,(((Erinaceus euro
paeus|HK3|SbEBIJiXGc:0.172974,(Suncus etruscus|HK3|WIGLlbztRS:0.269263
,Talpa occidentalis|HK3|MbnVjEhKCF:0.083482)S:0.021716)S:0.021448,(Sur
icata suricatta|HK3|czgRgyZtFJ:0.072076,(Miniopterus natalensis|HK3|jB
HUmwqCXu:0.083853,Desmodus rotundus|HK3|dXRmcPxDid:0.105319)S:0.032772
)S:0.007435)S:0.003043,(((Ochotona curzoniae|HK3|UpCmrXEdks:0.194381,O
ryctolagus cuniculus|HK3|JhPnTeDAUk:0.058749)S:0.042522,(((Fukomys dam
arensis|HK3|zgJUZbGElA:0.051128,Cavia porcellus|HK3|fcUjfYTTiq:0.14971
1)S:0.050124,(Dipodomys spectabilis|HK3|pfTUUCWeHp:0.130777,(Jaculus j
aculus|HK3|TWvySMGveq:0.110541,(Nannospalax galili|HK3|CrmROQqfyG:0.06
2087,((Grammomys surdaster|HK3|BSLllRRzgE:0.069997,Psammomys obesus|HK
3|iPRQzzZeJH:0.10101)S:1e-06,Chionomys nivalis|HK3|oMWYbiCulg:0.169145
)S:0.046204)S:0.029695)S:0.011875)S:0.014882)S:1e-06,Marmota marmota m
armota|HK3|KJwThyPynx:0.074303)S:0.01064)S:0.015955,((Microcebus murin
us|HK3|EMaweTluNF:0.103193,Otolemur garnettii|HK3|ZxsMJahiAq:0.080177)
S:0.015939,((Callithrix jacchus|HK3|AeeGlozWKL:0.174635,Homo sapiens|H
K3|BFYESbMjCp:0.029952)S:0.020526,Carlito syrichta|HK3|PTXqKIbVOb:0.11
6466)S:0.006289)S:0.012477)S:0.01765)S:0.0262)S:0.183963,((Dromiciops
gliroides|HK3|fIRXultwTn:0.06181,(Antechinus flavipes|HK3|vtWrfmQvsB:0
.143088,Trichosurus vulpecula|HK3|wfGnpkHXaz:0.085759)S:0.01919)S:0.02
5469,Monodelphis domestica|HK3|wfOMlvxmMu:0.092493)S:0.154968)S:0.1257
45)S:0.238306,((Protobothrops mucrosquamatus|HK3|YLYYaWThsj:0.371042,E
lgaria multicarinata webbii|HK3|MdNSXAHZft:0.255297)S:0.55215,((((Phas
ianus colchicus|HK3|vZvlmTAEnR:0.218987,Anser cygnoides|HK3|RHUMEJUeYm
:0.105836)S:0.08187,(Nyctiprogne leucopyga|HK3|kZwwNgzPCt:0.13009,((Bu
rhinus bistriatus|HK3|UlCPfPqcEA:0.05145,((Turnix velox|HK3|rqPQKInsSW
:0.295639,Uria aalge|HK3|IkbPSCMcHs:0.063222)S:0.02397,(Rostratula ben

```
ghalensis|HK3|ApEQAGwbCF:0.081485,Arenaria interpres|HK3|ZRJDvVqaYr:0.
120128)S:0.065248)S:0.06193)S:0.032522,((((((Passer domesticus|HK3|Oor
MPQmfvK:0.105724,Regulus satrapa|HK3|pghRAHgkQE:0.090448)S:0.019386,(A
crocephalus arundinaceus|HK3|ZdrTnDWctd:0.062305,Sylvia atricapilla|HK
3|XgJeOwDcHb:0.090871)S:0.031509)S:0.039793,Drymodes brunneopygia|HK3|
XiigvmLSSv:0.092388)S:0.048404,Climacteris rufus|HK3|DQXSMRkhuo:0.0860
05)S:0.095566,(((Xiphorhynchus elegans|HK3|EJMVMuomip:0.076914,Scytalo
pus superciliaris|HK3|IdEaORUUBk:0.096661)S:0.016855,Grallaria varia|H
K3|XpgMnWJnrP:0.113128)S:0.056754,(Pitta sordida|HK3|pxizwQDLmo:0.1387
2,(Smithornis capensis|HK3|CYlBlIZKVA:0.085856,Calyptomena viridis|HK3
|RPoOpEKnrc:0.080459)S:0.050806)S:0.063332)S:0.025179)S:0.147054,(Rhin
opomastus cyanomelas|HK3|PbQqymDllh:0.19345,Brachypteracias leptosomus
|HK3|ajpcOmWbRH:0.137607)S:0.018573)S:0.027205)S:0.008561)S:0.040854)S
:0.130129,(Dromaius novaehollandiae|HK3|jOWGeTwolc:0.070103,Crypturell
us undulatus|HK3|tLTLumxuGH:0.13036)S:0.056697)S:0.345781,Dermochelys
coriacea|HK3|KaFweMEudV:0.315832)S:0.097151)S:0.14141)S:0.261851,((Pse
udophryne corroboree|HK3|qSucPCjdGI:0.246067,(Rana temporaria|HK3|zOzf
eCXiPf:0.099756,Nanorana parkeri|HK3|JEOPzKCIiz:0.058357)S:0.113625)S:
0.141915,Bombina bombina|HK3|ExGhVFjtQF:0.175774)S:0.235897)S:0.077389
)S:0.273291)D:0.402186,Lethenteron reissneri|HK1|dGLCSoWnUb:0.402186);
```

Reconciled tree for ancestor identities:

```
(((((((Erpetoichthys calabaricus|HK1|uGyuBPeKnL:0.042827,(((Thalassoph
ryne amazonica|HK1|JsEOEDBkRa:0.048126,(Doryrhamphus excisus|HK1|cALFu
wyBrJ:0.017496,((Pelmatolapia mariae|HK1|kIEIvXJyGq:0.031923,Acanthoch
romis polyacanthus|HK1|BTvAaCUYBV:0.02435)a1:0.003269,((Centropristis
striata|HK1|MsaUHPpxrS:0.008969,Pseudoliparis swirei|HK1|qSMnnuMCBW:0.
067126)a2:0.026092,Micropterus salmoides|HK1|mczuyQEJNy:0.023986)a3:0.
008495)a4:0.0061)a5:0.009603)a6:0.017616,Lampris incognitus|HK1|PuwQUX
YfgX:0.030171)a7:1e-06,Gadus macrocephalus|HK1|LWxXzurkMl:0.057181)a8:
0.088159)a9:0.030805,((((((((Opisthocomus hoazin|HK1|WeUlkjVgVj:0.001
456,(Athene cunicularia|HK1|XfWuHJYvcU:0.001456,((Myiozetetes cayanens
is|HK1|emzIUjIaqN:1e-06,(Corvus cornix cornix|HK1|rKHLvnvBYe:0.002924,
Malurus melanocephalus|HK1|MlNzhWNIZC:1e-06)a10:0.00146)a11:0.006877,M
elopsittacus undulatus|HK1|PCYyNVKAZi:0.002476)a12:0.003827)a13:1e-06)
a14:0.010489,Dromaius novaehollandiae|HK1|RhrdaRtdfb:0.011616)a15:0.01
0016,Alligator sinensis|HK1|NuiileKrUF:0.027384)a16:0.008391,Pelodiscu
s sinensis|HK1|tqUNgzXpKh:0.023037)a17:0.008235,Pogona vitticeps|HK1|J
BzVERGLeG:0.037744)a18:0.009889,(Tachyglossus aculeatus|HK1|EipBvNDojf
:0.030522,((Monodelphis domestica|HK1|jKrGuSGcUc:0.126861,Dromiciops g
liroides|HK1|kAxRfmrNyH:0.045883)a19:0.134336,(((Homo sapiens|HK1|HikR
HBmLYx:0.022674,Phodopus roborovskii|HK1|iXEhcskcgQ:0.067077)a20:0.013
867,(Myotis myotis|HK1|ArELCCLoEK:0.064895,Lagenorhynchus albirostris|
HK1|wLZZSHxlti:0.044937)a21:0.007598)a22:0.016086,Orycteropus afer afe
r|HK1|ARXWtKFged:0.047294)a23:0.05847)a24:0.056927)a25:0.038663)a26:0.
023838,Rhinatrema bivittatum|HK1|ADrBoRHQCz:0.051733)a27:0.015923,Prot
opterus annectens|HK1|OSnuUfJPUx:0.111823)a28:0.00772,Latimeria chalum
nae|HK1|OgOKnwCKyc:0.106276)a29:0.014831)a30:0.050126,Leucoraja erinac
```

ea|HK1|gFeVmhUjNL:0.160266)a31:0.048348,(((Polyodon spathula|HKDC1|Fbl
hIVtssc:0.106835,((Amia calva|HKDC1|uqqMvJNjEt:0.089454,Atractosteus s
patula|HKDC1|WZsmwTEtMw:0.09822)a32:0.015618,(((Scleropages formosus|H
KDC1|TPLznODygH:0.134745,Brienomyrus brachyistius|HKDC1|bZDMaqdqSA:0.1
78449)a33:0.051788,(((((Ictalurus punctatus|HKDC1|TGRsMCfawE:0.069373,
Electrophorus electricus|HKDC1|vtZUynMZlo:0.092751)a34:0.026504,Parami
sgurnus dabryanus|HKDC1|vrcEEbpAUS:0.181241)a35:0.013955,Chanos chanos
|HKDC1|YHNqFMcxNZ:0.138704)a36:0.023831,(Clupea harengus|HKDC1|bFsdyTf
XGl:0.113984,Denticeps clupeoides|HKDC1|QIqYccklUB:0.104353)a37:0.0194
69)a38:0.01788,(Hypomesus transpacificus|HKDC1|aLetOARBie:0.160867,(((
Myripristis murdjan|HKDC1|BsaruMxntV:0.13315,(Thunnus albacares|HKDC1|
ZmGyNEPRji:0.0799,(((Centropristis striata|HKDC1|RSdDyCyTBC:0.081794,E
pinephelus fuscoguttatus|HKDC1|AeFKeZcVXW:0.095916)a39:0.013063,((Pseu
doliparis swirei|HKDC1|iCcmbGeuop:0.125383,Anoplopoma fimbria|HKDC1|Va
FZQoiiBx:0.088181)a40:0.024085,((Cottoperca gobio|HKDC1|SdfTRkeJir:0.0
56577,Eleginops maclovinus|HKDC1|GQwKbHfHXa:0.145751)a41:0.023319,Seba
stes umbrosus|HKDC1|ElnqstjQmQ:0.079493)a42:1e-06)a43:0.008725)a44:0.0
27025,((Salarias fasciatus|HKDC1|oPtZfrHMEn:0.21552,Gouania willdenowi
|HKDC1|scCjzbmRDo:0.172378)a45:0.042332,((Oryzias latipes|HKDC1|APKqDV
mZwa:0.164753,Cololabis saira|HKDC1|dSBRkeWooW:0.140333)a46:0.026185,(
Nothobranchius furzeri|HKDC1|vthLsNmGae:0.147621,((Fundulus heteroclit
us|HKDC1|XFXAGhmEdt:0.056429,Cyprinodon tularosa|HKDC1|BxkPqfOcsU:0.10
7857)a47:0.019443,Poecilia formosa|HKDC1|gXFNGdeqfM:0.103393)a48:0.065
108)a49:0.031436)a50:0.041641)a51:0.028514)a52:0.009116)a53:0.129655)a
54:0.022445,Lampris incognitus|HKDC1|hNHyWARdjX:0.162052)a55:0.024504,
Gadus morhua|HKDC1|ygnXzsWQlY:0.2318)a56:0.160587)a57:0.075133)a58:0.0
19401)a59:0.0101,(Megalops cyprinoides|HKDC1|JiTDiqJWqq:0.063872,(Angu
illa anguilla|HKDC1|hrkEPumxwE:0.105985,Conger conger|HKDC1|KyPRaGQWJn
:0.144742)a60:0.107401)a61:0.02612)a62:0.04707)a63:0.076352)a64:0.0865
05,(Latimeria chalumnae|HKDC1|kAoIdxkVCy:0.133948,((((Dromiciops gliro
ides|HKDC1|oKbmBUVIIJ:0.104678,(((Echinops telfairi|HKDC1|CTvbLSGChX:0
.158559,Chrysochloris asiatica|HKDC1|TPMNSTrvzb:0.112338)a65:0.025103,
Orycteropus afer afer|HKDC1|IjoVSXORKf:0.041286)a66:0.016377,(((Meles
meles|HKDC1|cmKXcqnZVo:0.061083,Myotis brandtii|HKDC1|gNtcFCJNRn:0.053
214)a67:0.006724,(Suncus etruscus|HKDC1|SYiclaDRrZ:0.10866,Erinaceus e
uropaeus|HKDC1|dAnfsJuzzw:0.051941)a68:0.015025)a69:0.006833,Homo sapi
ens|HKDC1|SJSMcTzbZx:0.054838)a70:0.009068)a71:0.049098)a72:0.021471,O
rnithorhynchus anatinus|HKDC1|YgLNdLnanC:0.104949)a73:0.07321,((((((G
avia stellata|HKDC1|XGlDGGvmcy:0.011982,((Neopsephotus bourkii|HKDC1|c
fQkjMGkNp:0.018884,(Acanthisitta chloris|HKDC1|yZBvKdwupL:0.041304,(Zo
notrichia albicollis|HKDC1|uymusyCedr:0.100731,Malurus melanocephalus|
HKDC1|FzUHEqEplI:0.015182)a74:0.014198)a75:0.012751)a76:0.00658,Athene
cunicularia|HKDC1|umPqkFBiYK:0.010564)a77:1e-06)a78:1e-06,Grus americ
ana|HKDC1|eAvPfRizJy:0.014941)a79:0.004363,Calypte anna|HKDC1|utbBrXfy
rJ:0.055557)a80:0.011448,Coturnix japonica|HKDC1|luLQhYohCx:0.040949)a
81:0.088984,Alligator mississippiensis|HKDC1|wTCBdToqMz:0.048152)a82:0
.010923,Emys orbicularis|HKDC1|iqhvTiaZIT:0.063257)a83:0.006857,((Poda
rcis muralis|HKDC1|IVIgXHPWxQ:0.072952,Python bivittatus|HKDC1|nahIDFG
yme:0.095862)a84:0.011449,Euleptes europaea|HKDC1|xIoXkCYtzw:0.069898)
a85:0.033993)a86:0.028546)a87:0.021759,(((Bufo bufo|HKDC1|EijMExSFZR:0
.065147,Rana temporaria|HKDC1|PNiWpIvXnO:0.125618)a88:0.037841,Xenopus
tropicalis|HKDC1|cKpdtvXrTj:0.090193)a89:0.117427,Rhinatrema bivittat

um|HKDC1|DYIUurqojh:0.16561)a90:0.027073)a91:0.072769)a92:0.017236)a93:0.069519,(Stegostoma fasciatum|HKDC1|PxpsLTmAmN:0.350636,(Pristis pectinata|HKDC1|FrKkoLgmfd:0.060099,Dasyatis sabina|HKDC1|oOYqDdZzSb:0.19203)a94:0.174183)a95:0.206247)a96:0.100981)a97:0.067913,((((Protopterus annectens|HK2|EUsHUvJhlS:0.139272,((Spea bombifrons|HK2|kuBDQaekeh:0.080905,Rhinatrema bivittatum|HK2|rrWWLZdCcr:0.105941)a98:0.006914,((Tachyglossus aculeatus|HK2|PsAKGFFUJM:0.172896,(Vombatus ursinus|HK2|VPnyoqRgzk:0.041309,((Homo sapiens|HK2|pkVOClMpeg:0.035019,((Hyaena hyaena|HK2|yOGOMOiisv:0.017576,Budorcas taxicolor|HK2|MwgMQvftWF:0.023783)a99:0.007814,Galemys pyrenaicus|HK2|vgIzhLDEor:0.044853)a100:0.005585)a101:0.009985,Echinops telfairi|HK2|xzlgnvJvEl:0.043699)a102:0.02739)a103:0.036854)a104:0.051232,((Terrapene carolina triunguis|HK2|TlpAKSJfbS:0.050967,(Alligator sinensis|HK2|KcAcsyJVLt:0.051181,((((Accipiter gentilis|HK2|ONiFiGZKDZ:0.010426,((Corapipo altera|HK2|gUAxZacVZH:0.025217,((Ptilonorhynchus violaceus|HK2|omwJgFJSDB:0.008377,(Orthonyx spaldingii|HK2|bjdmRLkbSC:0.008402,(Drymodes brunneopygia|HK2|hCroEBgKFM:0.000623,Chaetops frenatus|HK2|pNSXxqafgA:0.027434)a105:0.006169)a106:0.003031)a107:0.002062,Atrichornis clamosus|HK2|JvNvvnoBoc:0.003184)a108:0.003932)a109:0.018456,Lathamus discolor|HK2|FJwoHpOswW:0.012488)a110:0.007066)a111:1e-06,((Gavia stellata|HK2|wKENSYxIvv:0.002969,Calonectris borealis|HK2|aWMhpJyWGa:0.002976)a112:0.001506,Chroicocephalus ridibundus|HK2|VMOkiKMwtj:0.011957)a113:0.00154)a114:1e-06,Columba livia|HK2|bXWVFYgEUU:0.008911)a115:0.009515,Dromaius novaehollandiae|HK2|GAdXLAPtGH:0.045387)a116:0.020237)a117:1e-06)a118:0.007021,(Eublepharis macularius|HK2|vGXLdoqggE:0.052086,(Pseudonaja textilis|HK2|DkEEDvenrP:0.182319,Elgaria multicarinata webbii|HK2|ZMtHoKJean:0.076352)a119:0.033583)a120:0.046569)a121:0.018503)a122:0.017823)a123:0.024773)a124:0.008293,Latimeria chalumnae|HK2|IKmASFgAXD:0.131939)a125:0.006494,(Polypterus senegalus|HK2|RXwXFroLbj:0.087665,(Polyodon spathula|HK2|PcUmlCXAsz:0.124769,(((((((((Gasterosteus aculeatus aculeatus|HK2|twGyJDlrlX:0.08488,Eleginops maclovinus|HK2|HmjDiAqQqY:0.080872)a126:0.013561,Centropristis striata|HK2|XcZnpOSbXX:0.064787)a127:0.01489,((Sparus aurata|HK2|pXmfIKjGzQ:0.052502,Takifugu flavidus|HK2|DewUMDFyVe:0.197991)a128:0.027372,(Xyrichtys novacula|HK2|CDwEEimyOz:0.073151,Labrus mixtus|HK2|uyzlfzIdCF:0.074346)a129:0.043741)a130:0.020714)a131:0.011446,((((Gouania willdenowi|HK2|mvMbYoycnf:0.155049,Salarias fasciatus|HK2|buwEgcGvFp:0.092607)a132:0.037202,Parambassis ranga|HK2|WCTACmqHwb:0.074222)a133:0.001866,(Maylandia zebra|HK2|mwzhgngitf:0.113708,((Cololabis saira|HK2|rKDgEdBXSe:0.126143,Oryzias melastigma|HK2|tChcjADeIz:0.125624)a134:0.018733,(Nothobranchius furzeri|HK2|mTMdGJVSyy:0.078747,Kryptolebias marmoratus|HK2|ejpzKAiXzB:0.078091)a135:0.040241)a136:0.024425)a137:0.008679)a138:0.030966,(Echeneis naucrates|HK2|feLDljjCjp:0.118394,Seriola dumerili|HK2|fLDqICFZTw:0.035219)a139:0.019877)a140:0.008409)a141:0.003451,((Synchiropus splendidus|HK2|YUaKrzfYXJ:0.222826,(Entelurus aequoreus|HK2|HJiPJMaXCk:0.122121,Phycodurus eques|HK2|pGbtdXQYlk:0.100637)a142:0.075363)a143:0.020728,Thunnus maccoyii|HK2|oTODkFQbVt:0.052924)a144:0.023575)a145:0.040768,Thalassophryne amazonica|HK2|FVizizcrOp:0.241586)a146:0.061396,Lampris incognitus|HK2|SvQzutreJy:0.10933)a147:0.034378,Merluccius polli|HK2|fcOSUWMZAk:0.13425)a148:0.023958,Salvelinus namaycush|HK2|YbxquaaXVr:0.111815)a149:0.005977,((Denticeps clupeoides|HK2|qjFPaapWkZ:0.098493,(Engraulis encrasicolus|HK2|SiPVyFcTKH:0.136163,Alosa sapidissima|HK2|FQIQwDCBql:0.077091)a150:0.0542

04)a151:0.01208,((Astyanax mexicanus|HK2|UCSURQaOGS:0.06453,(Trichomyc
terus rosablanca|HK2|PeNBFeARln:0.096706,Silurus meridionalis|HK2|XsMC
JQiJBO:0.070129)a152:0.046303)a153:0.033694,Xyrauchen texanus|HK2|YSBD
vLgDiW:0.103851)a154:0.009968)a155:0.025043)a156:0.147162)a157:0.02342
2)a158:0.036841)a159:0.057029,(Callorhinchus milii|HK2|HxvSQZRczB:0.10
5874,((Scyliorhinus canicula|HK2|KartHAlQEB:0.078369,Hemiscyllium ocel
latum|HK2|jrOmbuyPcp:0.154145)a160:0.014444,(Amblyraja radiata|HK2|WUG
YLAukGn:0.184299,Dasyatis sabina|HK2|AJKTmVRJQz:0.070922)a161:0.033089
)a162:0.080567)a163:0.050241)a164:0.081811)a165:0.053869,(Protopterus
annectens|HK3|NvqwsvPsyX:0.320144,(((Tachyglossus aculeatus|HK3|AvxvjO
UqzE:0.298989,(((Trichechus manatus latirostris|HK3|nxdYWiMVaR:0.15608
1,(((Chrysochloris asiatica|HK3|KCyjjKuQTm:0.079569,Echinops telfairi|
HK3|flIbqulDzJ:0.112912)a166:0.013466,Elephantulus edwardii|HK3|HxnlTj
CZOb:0.122379)a167:0.007543,Orycteropus afer afer|HK3|kuGXdlwnDq:0.097
764)a168:0.016081)a169:0.041367,(((Erinaceus europaeus|HK3|SbEBIJiXGc:
0.172974,(Suncus etruscus|HK3|WIGLlbztRS:0.269263,Talpa occidentalis|H
K3|MbnVjEhKCF:0.083482)a170:0.021716)a171:0.021448,(Suricata suricatta
|HK3|czgRgyZtFJ:0.072076,(Miniopterus natalensis|HK3|jBHUmwqCXu:0.0838
53,Desmodus rotundus|HK3|dXRmcPxDid:0.105319)a172:0.032772)a173:0.0074
35)a174:0.003043,(((Ochotona curzoniae|HK3|UpCmrXEdks:0.194381,Oryctol
agus cuniculus|HK3|JhPnTeDAUk:0.058749)a175:0.042522,(((Fukomys damare
nsis|HK3|zgJUZbGElA:0.051128,Cavia porcellus|HK3|fcUjfYTTiq:0.149711)a
176:0.050124,(Dipodomys spectabilis|HK3|pfTUUCWeHp:0.130777,(Jaculus j
aculus|HK3|TWvySMGveq:0.110541,(Nannospalax galili|HK3|CrmROQqfyG:0.06
2087,((Grammomys surdaster|HK3|BSLllRRzgE:0.069997,Psammomys obesus|HK
3|iPRQzzZeJH:0.10101)a177:1e-06,Chionomys nivalis|HK3|oMWYbiCulg:0.169
145)a178:0.046204)a179:0.029695)a180:0.011875)a181:0.014882)a182:1e-06
,Marmota marmota marmota|HK3|KJwThyPynx:0.074303)a183:0.01064)a184:0.0
15955,((Microcebus murinus|HK3|EMaweTluNF:0.103193,Otolemur garnettii|
HK3|ZxsMJahiAq:0.080177)a185:0.015939,((Callithrix jacchus|HK3|AeeGloz
WKL:0.174635,Homo sapiens|HK3|BFYESbMjCp:0.029952)a186:0.020526,Carlit
o syrichta|HK3|PTXqKIbVOb:0.116466)a187:0.006289)a188:0.012477)a189:0.
01765)a190:0.0262)a191:0.183963,((Dromiciops gliroides|HK3|fIRXultwTn:
0.06181,(Antechinus flavipes|HK3|vtWrfmQvsB:0.143088,Trichosurus vulpe
cula|HK3|wfGnpkHXaz:0.085759)a192:0.01919)a193:0.025469,Monodelphis do
mestica|HK3|wfOMlvxmMu:0.092493)a194:0.154968)a195:0.125745)a196:0.238
306,((Protobothrops mucrosquamatus|HK3|YLYYaWThsj:0.371042,Elgaria mul
ticarinata webbii|HK3|MdNSXAHZft:0.255297)a197:0.55215,((((Phasianus c
olchicus|HK3|vZvlmTAEnR:0.218987,Anser cygnoides|HK3|RHUMEJUeYm:0.1058
36)a198:0.08187,(Nyctiprogne leucopyga|HK3|kZwwNgzPCt:0.13009,((Burhin
us bistriatus|HK3|UlCPfPqcEA:0.05145,((Turnix velox|HK3|rqPQKInsSW:0.2
95639,Uria aalge|HK3|IkbPSCMcHs:0.063222)a199:0.02397,(Rostratula beng
halensis|HK3|ApEQAGwbCF:0.081485,Arenaria interpres|HK3|ZRJDvVqaYr:0.1
20128)a200:0.065248)a201:0.06193)a202:0.032522,(((((Passer domesticus
|HK3|OorMPQmfvK:0.105724,Regulus satrapa|HK3|pghRAHgkQE:0.090448)a203:
0.019386,(Acrocephalus arundinaceus|HK3|ZdrTnDWctd:0.062305,Sylvia atr
icapilla|HK3|XgJeOwDcHb:0.090871)a204:0.031509)a205:0.039793,Drymodes
brunneopygia|HK3|XiigvmLSSv:0.092388)a206:0.048404,Climacteris rufus|H
K3|DQXSMRkhuo:0.086005)a207:0.095566,(((Xiphorhynchus elegans|HK3|EJMV
Muomip:0.076914,Scytalopus superciliaris|HK3|IdEaORUUBk:0.096661)a208:
0.016855,Grallaria varia|HK3|XpgMnWJnrP:0.113128)a209:0.056754,(Pitta
sordida|HK3|pxizwQDLmo:0.13872,(Smithornis capensis|HK3|CYlBlIZKVA:0.0

```
85856,Calyptomena viridis|HK3|RPoOpEKnrc:0.080459)a210:0.050806)a211:0
.063332)a212:0.025179)a213:0.147054,(Rhinopomastus cyanomelas|HK3|PbQq
ymDllh:0.19345,Brachypteracias leptosomus|HK3|ajpcOmWbRH:0.137607)a214
:0.018573)a215:0.027205)a216:0.008561)a217:0.040854)a218:0.130129,(Dro
maius novaehollandiae|HK3|jOWGeTwolc:0.070103,Crypturellus undulatus|H
K3|tLTLumxuGH:0.13036)a219:0.056697)a220:0.345781,Dermochelys coriacea
|HK3|KaFweMEudV:0.315832)a221:0.097151)a222:0.14141)a223:0.261851,((Ps
eudophryne corroboree|HK3|qSucPCjdGI:0.246067,(Rana temporaria|HK3|zOz
feCXiPf:0.099756,Nanorana parkeri|HK3|JEOPzKCIiz:0.058357)a224:0.11362
5)a225:0.141915,Bombina bombina|HK3|ExGhVFjtQF:0.175774)a226:0.235897)
a227:0.077389)a228:0.273291)a229:0.402186,Lethenteron reissneri|HK1|dG
LCSoWnUb:0.402186);
```

Reconciled tree for posterior probabilities:

```
((((((Erpetoichthys calabaricus|HK1|uGyuBPeKnL:0.042827,(((Thalassoph
ryne amazonica|HK1|JsEOEDBkRa:0.048126,(Doryrhamphus excisus|HK1|cALFu
wyBrJ:0.017496,((Pelmatolapia mariae|HK1|kIEIvXJyGq:0.031923,Acanthoch
romis polyacanthus|HK1|BTvAaCUYBV:0.02435)0.998044:0.003269,((Centropr
istis striata|HK1|MsaUHPpxrS:0.008969,Pseudoliparis swirei|HK1|qSMnnuM
CBW:0.067126)0.998054:0.026092,Micropterus salmoides|HK1|mczuyQEJNy:0.
023986)0.998497:0.008495)0.998969:0.0061)0.998605:0.009603)0.998065:0.
017616,Lampris incognitus|HK1|PuwQUXYfgX:0.030171)0.997257:1e-06,Gadus
 macrocephalus|HK1|LWxXzurkMl:0.057181)0.997257:0.088159)0.99319:0.030
805,((((((((Opisthocomus hoazin|HK1|WeUlkjVgVj:0.001456,(Athene cunic
ularia|HK1|XfWuHJYvcU:0.001456,((Myiozetetes cayanensis|HK1|emzIUjIaqN
:1e-06,(Corvus cornix cornix|HK1|rKHLvnvBYe:0.002924,Malurus melanocep
halus|HK1|MlNzhWNIZC:1e-06)1:0.00146)1:0.006877,Melopsittacus undulatu
s|HK1|PCYyNVKAZi:0.002476)0.999489:0.003827)1:1e-06)1:0.010489,Dromaiu
s novaehollandiae|HK1|RhrdaRtdfb:0.011616)0.999322:0.010016,Alligator
sinensis|HK1|NuiileKrUF:0.027384)0.998126:0.008391,Pelodiscus sinensis
|HK1|tqUNgzXpKh:0.023037)0.99854:0.008235,Pogona vitticeps|HK1|JBzVERG
LeG:0.037744)0.998341:0.009889,(Tachyglossus aculeatus|HK1|EipBvNDojf:
0.030522,((Monodelphis domestica|HK1|jKrGuSGcUc:0.126861,Dromiciops gl
iroides|HK1|kAxRfmrNyH:0.045883)0.989664:0.134336,(((Homo sapiens|HK1|
HikRHBmLYx:0.022674,Phodopus roborovskii|HK1|iXEhcskcgQ:0.067077)0.995
167:0.013867,(Myotis myotis|HK1|ArELCCLoEK:0.064895,Lagenorhynchus alb
irostris|HK1|wLZZSHxlti:0.044937)0.996335:0.007598)0.994332:0.016086,O
rycteropus afer afer|HK1|ARXWtKFged:0.047294)0.99336:0.05847)0.986444:
0.056927)0.99403:0.038663)0.99701:0.023838,Rhinatrema bivittatum|HK1|A
DrBoRHQCz:0.051733)0.995839:0.015923,Protopterus annectens|HK1|OSnuUfJ
PUx:0.111823)0.994322:0.00772,Latimeria chalumnae|HK1|OgOKnwCKyc:0.106
276)0.994158:0.014831)0.992585:0.050126,Leucoraja erinacea|HK1|gFeVmhU
jNL:0.160266)0.985455:0.048348,(((Polyodon spathula|HKDC1|FblhIVtssc:0
.106835,((Amia calva|HKDC1|uqqMvJNjEt:0.089454,Atractosteus spatula|HK
DC1|WZsmwTEtMw:0.09822)0.989715:0.015618,(((Scleropages formosus|HKDC1
|TPLznODygH:0.134745,Brienomyrus brachyistius|HKDC1|bZDMaqdqSA:0.17844
9)0.984796:0.051788,(((((Ictalurus punctatus|HKDC1|TGRsMCfawE:0.069373
,Electrophorus electricus|HKDC1|vtZUynMZlo:0.092751)0.991721:0.026504,
Paramisgurnus dabryanus|HKDC1|vrcEEbpAUS:0.181241)0.990407:0.013955,Ch
```

anos chanos|HKDC1|YHNqFMcxNZ:0.138704)0.990705:0.023831,(Clupea hareng
us|HKDC1|bFsdyTfXGl:0.113984,Denticeps clupeoides|HKDC1|QIqYccklUB:0.1
04353)0.98841:0.019469)0.990693:0.01788,(Hypomesus transpacificus|HKDC
1|aLetOARBie:0.160867,(((Myripristis murdjan|HKDC1|BsaruMxntV:0.13315,
(Thunnus albacares|HKDC1|ZmGyNEPRji:0.0799,(((Centropristis striata|HK
DC1|RSdDyCyTBC:0.081794,Epinephelus fuscoguttatus|HKDC1|AeFKeZcVXW:0.0
95916)0.994453:0.013063,((Pseudoliparis swirei|HKDC1|iCcmbGeuop:0.1253
83,Anoplopoma fimbria|HKDC1|VaFZQoiiBx:0.088181)0.993588:0.024085,((Co
ttoperca gobio|HKDC1|SdfTRkeJir:0.056577,Eleginops maclovinus|HKDC1|GQ
wKbHfHXa:0.145751)0.994627:0.023319,Sebastes umbrosus|HKDC1|ElnqstjQmQ
:0.079493)0.998038:1e-06)0.998038:0.008725)0.996063:0.027025,((Salaria
s fasciatus|HKDC1|oPtZfrHMEn:0.21552,Gouania willdenowi|HKDC1|scCjzbmR
Do:0.172378)0.97542:0.042332,((Oryzias latipes|HKDC1|APKqDVmZwa:0.1647
53,Cololabis saira|HKDC1|dSBRkeWooW:0.140333)0.979545:0.026185,(Nothob
ranchius furzeri|HKDC1|vthLsNmGae:0.147621,((Fundulus heteroclitus|HKD
C1|XFXAGhmEdt:0.056429,Cyprinodon tularosa|HKDC1|BxkPqfOcsU:0.107857)0
.988962:0.019443,Poecilia formosa|HKDC1|gXFNGdeqfM:0.103393)0.987066:0
.065108)0.985099:0.031436)0.985133:0.041641)0.985122:0.028514)0.992435
:0.009116)0.989024:0.129655)0.977775:0.022445,Lampris incognitus|HKDC1
|hNHyWARdjX:0.162052)0.97415:0.024504,Gadus morhua|HKDC1|ygnXzsWQlY:0.
2318)0.969761:0.160587)0.975085:0.075133)0.988742:0.019401)0.992164:0.
0101,(Megalops cyprinoides|HKDC1|JiTDiqJWqq:0.063872,(Anguilla anguill
a|HKDC1|hrkEPumxwE:0.105985,Conger conger|HKDC1|KyPRaGQWJn:0.144742)0.
984795:0.107401)0.989042:0.02612)0.99128:0.04707)0.989616:0.076352)0.9
84428:0.086505,(Latimeria chalumnae|HKDC1|kAoIdxkVCy:0.133948,((((Drom
iciops gliroides|HKDC1|oKbmBUVIIJ:0.104678,(((Echinops telfairi|HKDC1|
CTvbLSGChX:0.158559,Chrysochloris asiatica|HKDC1|TPMNSTrvzb:0.112338)0
.991301:0.025103,Orycteropus afer afer|HKDC1|IjoVSXORKf:0.041286)0.996
276:0.016377,(((Meles meles|HKDC1|cmKXcqnZVo:0.061083,Myotis brandtii|
HKDC1|gNtcFCJNRn:0.053214)0.997138:0.006724,(Suncus etruscus|HKDC1|SYi
claDRrZ:0.10866,Erinaceus europaeus|HKDC1|dAnfsJuzzw:0.051941)0.996515
:0.015025)0.997455:0.006833,Homo sapiens|HKDC1|SJSMcTzbZx:0.054838)0.9
97434:0.009068)0.997252:0.049098)0.989674:0.021471,Ornithorhynchus ana
tinus|HKDC1|YgLNdLnanC:0.104949)0.989179:0.07321,((((((Gavia stellata
|HKDC1|XGlDGGvmcy:0.011982,((Neopsephotus bourkii|HKDC1|cfQkjMGkNp:0.0
18884,(Acanthisitta chloris|HKDC1|yZBvKdwupL:0.041304,(Zonotrichia alb
icollis|HKDC1|uymusyCedr:0.100731,Malurus melanocephalus|HKDC1|FzUHEqE
plI:0.015182)0.997253:0.014198)0.997948:0.012751)0.999549:0.00658,Athe
ne cunicularia|HKDC1|umPqkFBiYK:0.010564)0.999933:1e-06)0.999933:1e-06
,Grus americana|HKDC1|eAvPfRizJy:0.014941)0.999933:0.004363,Calypte an
na|HKDC1|utbBrXfyrJ:0.055557)0.999207:0.011448,Coturnix japonica|HKDC1
|luLQhYohCx:0.040949)0.997545:0.088984,Alligator mississippiensis|HKDC
1|wTCBdToqMz:0.048152)0.995612:0.010923,Emys orbicularis|HKDC1|iqhvTia
ZIT:0.063257)0.995139:0.006857,((Podarcis muralis|HKDC1|IVIgXHPWxQ:0.0
72952,Python bivittatus|HKDC1|nahIDFGyme:0.095862)0.994499:0.011449,Eu
leptes europaea|HKDC1|xIoXkCYtzw:0.069898)0.995226:0.033993)0.995982:0
.028546)0.991918:0.021759,(((Bufo bufo|HKDC1|EijMExSFZR:0.065147,Rana
temporaria|HKDC1|PNiWpIvXnO:0.125618)0.990563:0.037841,Xenopus tropica
lis|HKDC1|cKpdtvXrTj:0.090193)0.986191:0.117427,Rhinatrema bivittatum|
HKDC1|DYIUurqojh:0.16561)0.984156:0.027073)0.988129:0.072769)0.98503:0
.017236)0.984258:0.069519,(Stegostoma fasciatum|HKDC1|PxpsLTmAmN:0.350
636,(Pristis pectinata|HKDC1|FrKkoLgmfd:0.060099,Dasyatis sabina|HKDC1

|oOYqDdZzSb:0.19203)0.982202:0.174183)0.95217:0.206247)0.973411:0.1009
81)0.981298:0.067913,((((Protopterus annectens|HK2|EUsHUvJhlS:0.139272
,((Spea bombifrons|HK2|kuBDQaekeh:0.080905,Rhinatrema bivittatum|HK2|r
rWWLZdCcr:0.105941)0.992875:0.006914,((Tachyglossus aculeatus|HK2|PsAK
GFFUJM:0.172896,(Vombatus ursinus|HK2|VPnyoqRgzk:0.041309,((Homo sapie
ns|HK2|pkVOClMpeg:0.035019,((Hyaena hyaena|HK2|yOGOMOiisv:0.017576,Bud
orcas taxicolor|HK2|MwgMQvftWF:0.023783)0.999219:0.007814,Galemys pyre
naicus|HK2|vgIzhLDEor:0.044853)0.998469:0.005585)0.997787:0.009985,Ech
inops telfairi|HK2|xzlgnvJvEl:0.043699)0.996549:0.02739)0.994813:0.036
854)0.991561:0.051232,((Terrapene carolina triunguis|HK2|TlpAKSJfbS:0.
050967,(Alligator sinensis|HK2|KcAcsyJVLt:0.051181,((((Accipiter genti
lis|HK2|ONiFiGZKDZ:0.010426,((Corapipo altera|HK2|gUAxZacVZH:0.025217,
((Ptilonorhynchus violaceus|HK2|omwJgFJSDB:0.008377,(Orthonyx spalding
ii|HK2|bjdmRLkbSC:0.008402,(Drymodes brunneopygia|HK2|hCroEBgKFM:0.000
623,Chaetops frenatus|HK2|pNSXxqafgA:0.027434)0.999665:0.006169)0.9995
59:0.003031)0.999577:0.002062,Atrichornis clamosus|HK2|JvNvvnoBoc:0.00
3184)0.999478:0.003932)0.998883:0.018456,Lathamus discolor|HK2|FJwoHpO
swW:0.012488)0.999657:0.007066)0.99996:1e-06,((Gavia stellata|HK2|wKEN
SYxIvv:0.002969,Calonectris borealis|HK2|aWMhpJyWGa:0.002976)0.999997:
0.001506,Chroicocephalus ridibundus|HK2|VMOkiKMwtj:0.011957)0.999986:0
.00154)0.99996:1e-06,Columba livia|HK2|bXWVFYgEUU:0.008911)0.99996:0.0
09515,Dromaius novaehollandiae|HK2|GAdXLAPtGH:0.045387)0.998758:0.0202
37)0.998177:1e-06)0.998177:0.007021,(Eublepharis macularius|HK2|vGXLdo
qggE:0.052086,(Pseudonaja textilis|HK2|DkEEDvenrP:0.182319,Elgaria mul
ticarinata webbii|HK2|ZMtHoKJean:0.076352)0.988219:0.033583)0.993344:0
.046569)0.997336:0.018503)0.995473:0.017823)0.99435:0.024773)0.99352:0
.008293,Latimeria chalumnae|HK2|IKmASFgAXD:0.131939)0.99342:0.006494,(
Polypterus senegalus|HK2|RXwXFroLbj:0.087665,(Polyodon spathula|HK2|Pc
UmlCXAsz:0.124769,(((((((((Gasterosteus aculeatus aculeatus|HK2|twGyJ
DlrlX:0.08488,Eleginops maclovinus|HK2|HmjDiAqQqY:0.080872)0.995598:0.
013561,Centropristis striata|HK2|XcZnpOSbXX:0.064787)0.996092:0.01489,
((Sparus aurata|HK2|pXmfIKjGzQ:0.052502,Takifugu flavidus|HK2|DewUMDFy
Ve:0.197991)0.992627:0.027372,(Xyrichtys novacula|HK2|CDwEEimyOz:0.073
151,Labrus mixtus|HK2|uyzlfzIdCF:0.074346)0.993227:0.043741)0.995481:0
.020714)0.996692:0.011446,((((Gouania willdenowi|HK2|mvMbYoycnf:0.1550
49,Salarias fasciatus|HK2|buwEgcGvFp:0.092607)0.990033:0.037202,Paramb
assis ranga|HK2|WCTACmqHwb:0.074222)0.996513:0.001866,(Maylandia zebra
|HK2|mwzhgngitf:0.113708,((Cololabis saira|HK2|rKDgEdBXSe:0.126143,Ory
zias melastigma|HK2|tChcjADeIz:0.125624)0.991375:0.018733,(Nothobranch
ius furzeri|HK2|mTMdGJVSyy:0.078747,Kryptolebias marmoratus|HK2|ejpzKA
iXzB:0.078091)0.994722:0.040241)0.994346:0.024425)0.996446:0.008679)0.
996817:0.030966,(Echeneis naucrates|HK2|feLDljjCjp:0.118394,Seriola du
merili|HK2|fLDqICFZTw:0.035219)0.995595:0.019877)0.997548:0.008409)0.9
98109:0.003451,((Synchiropus splendidus|HK2|YUaKrzfYXJ:0.222826,(Entel
urus aequoreus|HK2|HJiPJMaXCk:0.122121,Phycodurus eques|HK2|pGbtdXQYlk
:0.100637)0.984951:0.075363)0.990017:0.020728,Thunnus maccoyii|HK2|oTO
DkFQbVt:0.052924)0.996312:0.023575)0.997633:0.040768,Thalassophryne am
azonica|HK2|FVizizcrOp:0.241586)0.988885:0.061396,Lampris incognitus|H
K2|SvQzutreJy:0.10933)0.990561:0.034378,Merluccius polli|HK2|fcOSUWMZA
k:0.13425)0.99262:0.023958,Salvelinus namaycush|HK2|YbxquaaXVr:0.11181
5)0.99347:0.005977,((Denticeps clupeoides|HK2|qjFPaapWkZ:0.098493,(Eng
raulis encrasicolus|HK2|SiPVyFcTKH:0.136163,Alosa sapidissima|HK2|FQIQ

wDCBql:0.077091)0.987033:0.054204)0.991263:0.01208,((Astyanax mexicanu
s|HK2|UCSURQaOGS:0.06453,(Trichomycterus rosablanca|HK2|PeNBFeARln:0.0
96706,Silurus meridionalis|HK2|XsMCJQiJBO:0.070129)0.991133:0.046303)0
.991598:0.033694,Xyrauchen texanus|HK2|YSBDvLgDiW:0.103851)0.992467:0.
009968)0.992678:0.025043)0.992731:0.147162)0.986191:0.023422)0.989517:
0.036841)0.993988:0.057029,(Callorhinchus milii|HK2|HxvSQZRczB:0.10587
4,((Scyliorhinus canicula|HK2|KartHAlQEB:0.078369,Hemiscyllium ocellat
um|HK2|jrOmbuyPcp:0.154145)0.988851:0.014444,(Amblyraja radiata|HK2|WU
GYLAukGn:0.184299,Dasyatis sabina|HK2|AJKTmVRJQz:0.070922)0.987655:0.0
33089)0.987717:0.080567)0.98527:0.050241)0.985928:0.081811)0.974525:0.
053869,(Protopterus annectens|HK3|NvqwsvPsyX:0.320144,(((Tachyglossus
aculeatus|HK3|AvxvjOUqzE:0.298989,(((Trichechus manatus latirostris|HK
3|nxdYWiMVaR:0.156081,(((Chrysochloris asiatica|HK3|KCyjjKuQTm:0.07956
9,Echinops telfairi|HK3|flIbqulDzJ:0.112912)0.993554:0.013466,Elephant
ulus edwardii|HK3|HxnlTjCZOb:0.122379)0.995066:0.007543,Orycteropus af
er afer|HK3|kuGXdlwnDq:0.097764)0.995459:0.016081)0.993994:0.041367,((
(Erinaceus europaeus|HK3|SbEBIJiXGc:0.172974,(Suncus etruscus|HK3|WIGL
lbztRS:0.269263,Talpa occidentalis|HK3|MbnVjEhKCF:0.083482)0.989001:0.
021716)0.992748:0.021448,(Suricata suricatta|HK3|czgRgyZtFJ:0.072076,(
Miniopterus natalensis|HK3|jBHUmwqCXu:0.083853,Desmodus rotundus|HK3|d
XRmcPxDid:0.105319)0.993016:0.032772)0.996275:0.007435)0.997567:0.0030
43,(((Ochotona curzoniae|HK3|UpCmrXEdks:0.194381,Oryctolagus cuniculus
|HK3|JhPnTeDAUk:0.058749)0.991253:0.042522,(((Fukomys damarensis|HK3|z
gJUZbGElA:0.051128,Cavia porcellus|HK3|fcUjfYTTiq:0.149711)0.993504:0.
050124,(Dipodomys spectabilis|HK3|pfTUUCWeHp:0.130777,(Jaculus jaculus
|HK3|TWvySMGveq:0.110541,(Nannospalax galili|HK3|CrmROQqfyG:0.062087,(
(Grammomys surdaster|HK3|BSLllRRzgE:0.069997,Psammomys obesus|HK3|iPRQ
zzZeJH:0.10101)0.993502:1e-06,Chionomys nivalis|HK3|oMWYbiCulg:0.16914
5)0.993502:0.046204)0.994361:0.029695)0.995089:0.011875)0.995001:0.014
882)0.997052:1e-06,Marmota marmota marmota|HK3|KJwThyPynx:0.074303)0.9
97052:0.01064)0.996916:0.015955,((Microcebus murinus|HK3|EMaweTluNF:0.
103193,Otolemur garnettii|HK3|ZxsMJahiAq:0.080177)0.994989:0.015939,((
Callithrix jacchus|HK3|AeeGlozWKL:0.174635,Homo sapiens|HK3|BFYESbMjCp
:0.029952)0.995284:0.020526,Carlito syrichta|HK3|PTXqKIbVOb:0.116466)0
.996832:0.006289)0.996627:0.012477)0.996953:0.01765)0.997583:0.0262)0.
990547:0.183963,((Dromiciops gliroides|HK3|fIRXultwTn:0.06181,(Antechi
nus flavipes|HK3|vtWrfmQvsB:0.143088,Trichosurus vulpecula|HK3|wfGnpkH
Xaz:0.085759)0.991958:0.01919)0.992289:0.025469,Monodelphis domestica|
HK3|wfOMlvxmMu:0.092493)0.988664:0.154968)0.957556:0.125745)0.940813:0
.238306,((Protobothrops mucrosquamatus|HK3|YLYYaWThsj:0.371042,Elgaria
multicarinata webbii|HK3|MdNSXAHZft:0.255297)0.929637:0.55215,((((Pha
sianus colchicus|HK3|vZvlmTAEnR:0.218987,Anser cygnoides|HK3|RHUMEJUeY
m:0.105836)0.981573:0.08187,(Nyctiprogne leucopyga|HK3|kZwwNgzPCt:0.13
009,((Burhinus bistriatus|HK3|UlCPfPqcEA:0.05145,((Turnix velox|HK3|rq
PQKInsSW:0.295639,Uria aalge|HK3|IkbPSCMcHs:0.063222)0.987971:0.02397,
(Rostratula benghalensis|HK3|ApEQAGwbCF:0.081485,Arenaria interpres|HK
3|ZRJDvVqaYr:0.120128)0.987721:0.065248)0.987723:0.06193)0.99396:0.032
522,((((((Passer domesticus|HK3|OorMPQmfvK:0.105724,Regulus satrapa|HK
3|pghRAHgkQE:0.090448)0.990147:0.019386,(Acrocephalus arundinaceus|HK3
|ZdrTnDWctd:0.062305,Sylvia atricapilla|HK3|XgJeOwDcHb:0.090871)0.9944
44:0.031509)0.988798:0.039793,Drymodes brunneopygia|HK3|XiigvmLSSv:0.0
92388)0.988639:0.048404,Climacteris rufus|HK3|DQXSMRkhuo:0.086005)0.98

5147:0.095566,(((Xiphorhynchus elegans|HK3|EJMVMuomip:0.076914,Scytalo
pus superciliaris|HK3|IdEaORUUBk:0.096661)0.993012:0.016855,Grallaria
varia|HK3|XpgMnWJnrP:0.113128)0.991537:0.056754,(Pitta sordida|HK3|pxi
zwQDLmo:0.13872,(Smithornis capensis|HK3|CYlBlIZKVA:0.085856,Calyptome
na viridis|HK3|RPoOpEKnrc:0.080459)0.990017:0.050806)0.98651:0.063332)
0.988487:0.025179)0.985629:0.147054,(Rhinopomastus cyanomelas|HK3|PbQq
ymDllh:0.19345,Brachypteracias leptosomus|HK3|ajpcOmWbRH:0.137607)0.98
7939:0.018573)0.989946:0.027205)0.992929:0.008561)0.991792:0.040854)0.
984763:0.130129,(Dromaius novaehollandiae|HK3|jOWGeTwolc:0.070103,Cryp
turellus undulatus|HK3|tLTLumxuGH:0.13036)0.983482:0.056697)0.972947:0
.345781,Dermochelys coriacea|HK3|KaFweMEudV:0.315832)0.927799:0.097151
)0.917076:0.14141)0.919104:0.261851,((Pseudophryne corroboree|HK3|qSuc
PCjdGI:0.246067,(Rana temporaria|HK3|zOzfeCXiPf:0.099756,Nanorana park
eri|HK3|JEOPzKCIiz:0.058357)0.986221:0.113625)0.963745:0.141915,Bombin
a bombina|HK3|ExGhVFjtQF:0.175774)0.952228:0.235897)0.928434:0.077389)
0.932867:0.273291)0.954919:0.402186,Lethenteron reissneri|HK1|dGLCSoWn
Ub:0.402186);

**S3.** Reconstructed maximum-likelihood and altAll sequences of HK ancestors.

```
>anc1|avgPP:0.998|lnPP:-3.142|num_ambig:4|num_ambig_gaps:0
LYSMRFSDETLKDIMNRFRREMENGLGRDTNPTATVKMLPTFVRSIPDGSEKGDFIALDLGGSNFRILRV
KVTQDKKQPVQMESQVYETPDDIIHGSGTQLFDHVADCLGDFMEKQKIKDKKLPVGFTFSFPCAQTKLDE
AVLVTWTKKFKASGVEGMDVVKLLNKAIKKRGDYEADIMAVVNDTVGTMMTCGFDDQRCEVGIIIGTGTN
ACYMEELRHIDLVEGDEGRMCINTEWGAFGDDGSLEDIRTEFDREIDRGSINPGKQLFEKMASGMYMGEL
VRLILVKMAKEGLLFEGRITPELLTKGKIETKHVSAIEKTKEGLKKCMEILTRLGVEPSDEDCLAVQHVC
TIVSFRSANLISATLGAILSRLKENKGVARLRTTVGIDGSLYKMHPQYARRLHKTVRRLVPDSDVRFLLS
ESGSAKGAAMVTAVAYRLTEQARQIQQTLAEFRLSKAQLLEVKKRMRVEIERGLKKDSHEEATVKMLPTF
VRSTPDGTENGDFLALDLGGTNFRVLLVKIRSGKRRSVEMHNKIYAIPIEVMQGTGEELFDHIVHCISDF
LDYMGMKSARLPLGFTFSFPCHQTSLDAGILVTWTKGFKATDCEGEDVVELLREAIKRKEEFELDVVAIV
NDTVGTMMTCAYEEPTCEVGLIAGTGSNACYMEEMKNIEIVEGNEGRMCVNMEWGAFGDNGCLDDIRTTY
DQAVDENSLNEGKQRYEKMCSGMYLGEIVRQILIDLTKRGFLFRGQISETLKTRGIFETKFLSQIESDRL
ALLQVRAILQQLGLDSTCDDSIIVKEVCGTVSRRAAQICGAGMAAVVDKIRENRGLDHLDVTVGVDGTLY
KLHPHFSRIFQQTVKELAPKCDVNFLLSEDGSGKGAALITAVGCRQRELEAQQH
>anc1_altAll|avgPP:0.998|lnPP:-4.581|num_ambig:4|num_ambig_gaps:0
LYSMRFSDETLKDIMNRFRREMENGLGRDTNPTATVKMLPTFVRSIPDGSEKGDFIALDLGGSNFRILRV
KVTQDKKQPVQMESQVYETPDDIIHGSGTQLFDHVADCLGDFMEKQKIKDKKLPVGFTFSFPCAQTKLDE
AVLVTWTKKFKASGVEGMDVVKLLNKAIKKRGDYEADIMAVVNDTVGTMMTCGFDDQRCEVGIIIGTGTN
ACYMEELRHIDLVEGDEGRMCINTEWGAFGDDGSLEDIRTEFDREIDRGSINPGKQLFEKMASGMYMGEL
VRLILVKMAKEGLLFEGRITPELLTKGKIETKHVSAIEKTKEGLKKCMEILTRLGVEPSDEDCLAVQHVC
TIVSFRSANLISATLGAILSRLKENKGVARLRTTVGIDGSLYKMHPQYARRLHKTVRRLVPDSDVRFLLS
ESGSAKGAAMVTAVAYRLTEQARQIQQTLAEFRLSKTQLLEVKKRMRVEIERGLKKDSHKEATVKMLPTF
VRSTPDGTENGDFLALDLGGTNFRVLLVKIRSGKRRSVEMHNKIYAIPIEVMQGTGEELFDHIVHCISDF
LDYMGMKSARLPLGFTFSFPCHQTSLDAGILVTWTKGFKATDCEGEDVVELLREAIKRKEEFELDVVAIV
NDTVGTMMTCAYEEPTCEVGLIAGTGSNACYMEEMKNIEIVEGNEGRMCVNMEWGAFGDNGCLDDIRTKY
DQAVDENSLNEGKQRYEKMCSGMYLGEIVRQILIDLTKRGFLFRGQISETLKTRGIFETKFLSQIESDRL
ALLQVRAILQQLGLDSTCDDSIIVKEVCGTVSRRAAQICGAGMAAVVDKIRENRGLDHLDVTVGVDGTLY
KLHPHFSRIFQQTVKELAPKCDVTFLLSEDGSGKGAALITAVGCRQRELEAQQH
>anc2|avgPP:0.998|lnPP:-2.746|num_ambig:4|num_ambig_gaps:0
LYSMRFSDETLKDIMNRFRREMENGLGRDTSPTATVKMLPTFVRSIPDGSEKGDFIALDLGGSNFRILRV
KVTQDKKQPVQMESQVYETPDDIIHGSGTRLFGHVADCLGDFMEKQKIKDKKLPVGFTFSFPCAQTKLDE
AVLVTWTKKFKASGVEGMDVVSLLNRAIKKRGDYEADIMAVVNDTVGTMMTCGFDDQRCEVGIIIGTGTN
ACYMEELRHIDQVEGDEGRMCINTEWGAFGDDGSLEDIRTEFDREIDRGSINPGKQLFEKMASGMYMGEL
VRLILVKMAKEGLLFEGRITPELLTKGKIETKHVSAIEKSKEGLKKCMEILTRLGVEPSDEDCLAVQHVC
TIVSFRSANLIASTLGGILSRLKENKGVARLRTTVGIDGSLYKMHPQYARRLHKTVRRLVPDSDVRFLLS
ESGSAKGAAMVTAVAYRLTEQARQIQQTMAEFRLSKAQLLEVKKRMRVEIERGLKKTSHEEATVKMLPTF
VRTTPDGTENGDFLALDLGGTNFRVLLVKIRSGKKRSVEMHNKIYAIPIEVMQGTGEELFDHIVHCISDF
LDYMGMKSARLPLGFTFSFPCHQTSLDAGILVTWTKGFKATDCEGEDVVELLREAIKRKEEFELDVVAIV
NDTVGTMMTCAYEEPSCEVGLIAGTGSNACYMEEAKNIEIVEGSEGRMCVNMEWGAFGDNGCLDDIRTKY
DQAVDENSLNEGKQRYEKMCSGMYLGEIVRQILIDLTRRGFLFRGQISETLKTRGIFETKFLSQIESDRL
ALLQVRAILQQLGLDSTCDDSIIVKEVCGTVSRRAAQICGAGMAAVVDKIRENRGLDHLDITVGVDGTLY
KLHPHFSRIFQQTVKELAPKCDVNFLLSEDGSGKGAALITAVGCRQRELEAQQH
>anc2_altAll|avgPP:0.997|lnPP:-5.949|num_ambig:4|num_ambig_gaps:0
LYSMRFSDETLKDIMNRFRREMENGLGRDTSPTATVKMLPTFVRSIPDGSEKGDFIALDLGGSNFRILRV
KVTQDKKQPVQMESQVYETPDDIIHGSGTRLFGHVADCLGDFMEKQKIKDKKLPVGFTFSFPCAQTKLDE
AVLVTWTKKFKASGVEGMDVVSLLNRAIKKRGDYEADIMAVVNDTVGTMMTCGFDDQRCEVGIIIGTGTN
ACYMEELRHIDQVEGDEGRMCINTEWGAFGDDGSLEDIRTEFDREIDRGSINPGKQLFEKMASGMYMGEL
VRLILVKMAKEGLLFEGRITPELLTKGKIETKHVSAIEKSKEGLKKCMEILTRLGVEPSDEDCLAVQHVC
TIVSFRSANLIAATLGGILSRLKENKGVARLRTTVGIDGSLYKMHPQYARRLHKTVRRLVPDSDVRFLLS
```

ESGSAKGAAMVTAVAYRLTEQARQIQQTMAEFRLSKAQLLEVKKRMRVEIERGLKKTSHEDATVKMLPTF
VRSTPDGTENGDFLALDLGGTNFRVLLVKIRSGKKRSVEMHNKIYAIPIEVMQGTGEELFDHIVHCISDF
LDYMGMKSARLPLGFTFSFPCHQTSLDAGILVTWTKGFKATDCEGEDVVELLREAIKRKEEFELDVVAIV
NDTVGTMMTCAYEEPSCEVGLIAGTGSNACYMEEAKNIEIVEGSEGRMCVNMEWGAFGDNGCLDDIRTKY
DQAVDENSLNEGKQRYEKMCSGMYLGEIVRQILIDLTRRGFLFRGQISETLKTRGIFETKFLSQIESDRL
ALLQVRAILQQLGLDSTCDDSIIVKEVCGTVSRRAAQICGAGMAAVVDKIRENRGLDHLDITVGVDGTLY
KLHPHFSRIFQQTVKELAPKCDVNFLLSEDGSGKGAALITAVGCRQRELEAHQH
>anc3|avgPP:0.998|lnPP:-2.229|num_ambig:3|num_ambig_gaps:0
LYSMRFSDETLKDIMNRFRREMENGLGRDTNPTATVKMLPTFVRSIPDGSEKGDFIALDLGGSNFRILRV
KVTQDKKQPVQMESQVYETPDDIIHGSGTQLFDHVADCLGDFMEKQKIKDKKLPVGFTFSFPCAQTKLDE
AVLVTWTKKFKASGVEGMDVVKLLNKAIKKRGDYEADIMAVVNDTVGTMMTCGFDDQRCEVGIIIGTGTN
ACYMEELRHIDLVEGDEGRMCINTEWGAFGDDGSLEDIRTEFDREIDRGSINPGKQLFEKMASGMYMGEL
VRLILVKMAKEGLLFEGRITPELLTKGKIETKHVSAIEKTKEGLKKCMEILTRLGVEPSDEDCLAVQHVC
TIVSFRSANLIASTLGGILSRLKENKGVARLRTTVGIDGSLYKMHPQYARRLHKTVRRLVPDSDVRFLLS
ESGSAKGAAMVTAVAYRLTEQARQIQQTMAEFRLSKAQLLEVKKRMRVEIERGLKKDSHEEATVKMLPTF
VRSTPDGTENGDFLALDLGGTNFRVLLVKIRSGKRRSVEMHNKIYAIPIEVMQGTGEELFDHIVHCISDF
LDYMGMKSARLPLGFTFSFPCHQTSLDAGILVTWTKGFKATDCEGEDVVELLREAIKRKEEFELDVVAIV
NDTVGTMMTCAYEEPTCEVGLIAGTGSNACYMEETKNIEIVEGNEGRMCVNMEWGAFGDNGCLDDIRTKY
DQAVDENSLNEGKQRYEKMCSGMYLGEIVRQILIDLTRRGFLFRGQISETLKTRGIFETKFLSQIESDRL
ALLQVRAILQQLGLDSTCDDSIIVKEVCGTVSRRAAQICGAGMAAVVDKIRENRGLDHLDITVGVDGTLY
KLHPHFSRIFQQTVKELAPKCDVNFLLSEDGSGKGAALITAVGCRQRELEAQQH
>anc3_altAll|avgPP:0.998|lnPP:-4.092|num_ambig:3|num_ambig_gaps:0
LYSMRFSDETLKDIMNRFRREMENGLGRDTNPTATVKMLPTFVRSIPDGSEKGDFIALDLGGSNFRILRV
KVTQDKKQPVQMESQVYETPDDIIHGSGTQLFDHVADCLGDFMEKQKIKDKKLPVGFTFSFPCAQTKLDE
AVLVTWTKKFKASGVEGMDVVKLLNKAIKKRGDYEADIMAVVNDTVGTMMTCGFDDQRCEVGIIIGTGTN
ACYMEELRHIDLVEGDEGRMCINTEWGAFGDDGSLEDIRTEFDREIDRGSINPGKQLFEKMASGMYMGEL
VRLILVKMAKEGLLFEGRITPELLTKGKIETKHVSAIEKTKEGLKKCMEILTRLGVEPSDEDCLAVQHVC
TIVSFRSANLIAATLGGILSRLKENKGVARLRTTVGIDGSLYKMHPQYARRLHKTVRRLVPDSDVRFLLS
ESGSAKGAAMVTAVAYRLTEQARQIQQTMAEFRLSKAQLLEVKKRMRVEIERGLKKDSHKEATVKMLPTF
VRSTPDGTENGDFLALDLGGTNFRVLLVKIRSGKRRSVEMHNKIYAIPIEVMQGTGEELFDHIVHCISDF
LDYMGMKSARLPLGFTFSFPCHQTSLDAGILVTWTKGFKATDCEGEDVVELLREAIKRKEEFELDVVAIV
NDTVGTMMTCAYEEPTCEVGLIAGTGSNACYMEETKNIEIVEGNEGRMCVNMEWGAFGDNGCLDDIRTKY
DQAVDENSLNEGKQRYEKMCSGMYLGEIVRQILIDLTRRGFLFRGQISETLKTRGIFETKFLSQIESDRL
ALLQVRAILQQLGLDSTCDDSIIVKEVCGTVSRRAAQICGAGMAAVVDKIRENRGLDHLDITVGVDGTLY
KLHPHFSRIFQQTVKELAPKCDVTFLLSEDGSGKGAALITAVGCRQRELEAQQH
>anc4|avgPP:0.999|lnPP:-1.520|num_ambig:2|num_ambig_gaps:0
LYSMRFSDETLKDIMNRFRREMENGLGRDTNPTATVKMLPTFVRSIPDGSEKGDFIALDLGGSNFRILRV
KVTQDKKQPVQMESQVYETPDDIIHGSGTQLFDHVADCLGDFMEKQKIKDKKLPVGFTFSFPCAQTKLDE
AVLVTWTKKFKASGVEGMDVVKLLNKAIKKRGDYEADIMAVVNDTVGTMMTCGFDDQRCEVGIIIGTGTN
ACYMEELRHIDLVEGDEGRMCINTEWGAFGDDGSLEDIRTEFDREIDRGSINPGKQLFEKMASGMYMGEL
VRLILVKMAKEGLLFEGRITPELLTKGKIETKHVSAIEKTKEGLKKCMEILTRLGVEPSDEDCLAVQHVC
TIVSFRSANLISATLGGILSRLKENKGVARLRTTVGIDGSLYKMHPQYARRLHKTVRRLVPDSDVRFLLS
ESGSAKGAAMVTAVAYRLTEQARQIQQTLAEFRLSKAQLLEVKKRMRVEIERGLKKDSHEEATVKMLPTF
VRSTPDGTENGDFLALDLGGTNFRVLLVKIRSGKRRSVEMHNKIYAIPIEVMQGTGEELFDHIVHCISDF
LDYMGMKSARLPLGFTFSFPCHQTSLDAGILVTWTKGFKATDCEGEDVVELLREAIKRKEEFELDVVAIV
NDTVGTMMTCAYEEPTCEVGLIAGTGSNACYMEEMKNIEIVEGNEGRMCVNMEWGAFGDNGCLDDIRTKY
DQAVDENSLNEGKQRYEKMCSGMYLGEIVRQILIDLTKRGFLFRGQISETLKTRGIFETKFLSQIESDRL
ALLQVRAILQQLGLDSTCDDSIIVKEVCGTVSRRAAQICGAGMAAVVDKIRENRGLDHLDVTVGVDGTLY
KLHPHFSRIFQQTVKELAPKCDVNFLLSEDGSGKGAALITAVGCRQRELEAQQH
>anc4_altAll|avgPP:0.999|lnPP:-2.567|num_ambig:2|num_ambig_gaps:0
LYSMRFSDETLKDIMNRFRREMENGLGRDTNPTATVKMLPTFVRSIPDGSEKGDFIALDLGGSNFRILRV
KVTQDKKQPVQMESQVYETPDDIIHGSGTQLFDHVADCLGDFMEKQKIKDKKLPVGFTFSFPCAQTKLDE

AVLVTWTKKFKASGVEGMDVVKLLNKAIKKRGDYEADIMAVVNDTVGTMMTCGFDDQRCEVGIIIGTGTN
ACYMEELRHIDLVEGDEGRMCINTEWGAFGDDGSLEDIRTEFDREIDRGSINPGKQLFEKMASGMYMGEL
VRLILVKMAKEGLLFEGRITPELLTKGKIETKHVSAIEKTKEGLKKCMEILTRLGVEPSDEDCLAVQHVC
TIVSFRSANLISATLGGILSRLKENKGVARLRTTVGIDGSLYKMHPQYARRLHKTVRRLVPDSDVRFLLS
ESGSAKGAAMVTAVAYRLTEQARQIQQTLAEFRLSKAQLLEVKKRMRVEIERGLKKDSHKEATVKMLPTF
VRSTPDGTENGDFLALDLGGTNFRVLLVKIRSGKRRSVEMHNKIYAIPIEVMQGTGEELFDHIVHCISDF
LDYMGMKSARLPLGFTFSFPCHQTSLDAGILVTWTKGFKATDCEGEDVVELLREAIKRKEEFELDVVAIV
NDTVGTMMTCAYEEPTCEVGLIAGTGSNACYMEEMKNIEIVEGNEGRMCVNMEWGAFGDNGCLDDIRTKY
DQAVDENSLNEGKQRYEKMCSGMYLGEIVRQILIDLTKRGFLFRGQISETLKTRGIFETKFLSQIESDRL
ALLQVRAILQQLGLDSTCDDSIIVKEVCGTVSRRAAQICGAGMAAVVDKIRENRGLDHLDVTVGVDGTLY
KLHPHFSRIFQQTVKELAPKCDVTFLLSEDGSGKGAALITAVGCRQRELEAQQH
>anc5|avgPP:0.999|lnPP:-1.997|num_ambig:2|num_ambig_gaps:0
LYSMRFSDETLKDIMNRFRREMENGLGRDTNPTATVKMLPTFVRSIPDGSEKGDFIALDLGGSNFRILRV
KVTQDKKQPVQMESQVYETPDDIIHGSGTQLFDHVADCLGDFMEKQKIKDKKLPVGFTFSFPCAQTKLDE
AVLVTWTKKFKASGVEGMDVVKLLNKAIKKRGDYEADIMAVVNDTVGTMMTCGFDDQRCEVGIIIGTGTN
ACYMEELRHIDLVEGDEGRMCINTEWGAFGDDGSLEDIRTEFDREIDRGSINPGKQLFEKMASGMYMGEL
VRLILVKMAKEGLLFEGRITPELLTKGKIETKHVSAIEKTKEGLKKCMEILTRLGVEPSDEDCLAVQHVC
TIVSFRSANLISATLGGILSRLKENKGVARLRTTVGIDGSLYKMHPQYARRLHKTVRRLVPDSDVRFLLS
ESGSAKGAAMVTAVAYRLTEQARQIQQTLAEFRLSKAQLLEVKKRMRVEIERGLKKDTHEEATVKMLPTF
VRSTPDGTENGDFLALDLGGTNFRVLLVKIRSGKRRSVEMHNKIYAIPIEVMQGTGEELFDHIVHCISDF
LDYMGMKSARLPLGFTFSFPCHQTSLDAGILVTWTKGFKATDCEGEDVVELLREAIKRKEEFELDVVAIV
NDTVGTMMTCAYEEPTCEVGLIAGTGSNACYMEEMRNIEIVDGNEGRMCVNMEWGAFGDNGCLDDIRTKY
DQAVDENSLNEGKQRYEKMCSGMYLGEIVRQILIDLTKRGFLFRGQISETLKTRGIFETKFLSQIESDRL
ALLQVRAILQQLGLDSTCDDSIIVKEVCGTVSRRAAQICGAGMAAVVDKIRENRGLDHLDVTVGVDGTLY
KLHPHFSRIFHQTVKELAPKCDVNFLLSEDGSGKGAALITAVGCRQRELEAQQH
>anc5_altAll|avgPP:0.998|lnPP:-3.389|num_ambig:2|num_ambig_gaps:0
LYSMRFSDETLKDIMNRFRREMENGLGRDTNPTATVKMLPTFVRSIPDGSEKGDFIALDLGGSNFRILRV
KVTQDKKQPVQMESQVYETPDDIIHGSGTQLFDHVAECLGDFMEKQKIKDKKLPVGFTFSFPCAQTKLDE
AVLVTWTKKFKASGVEGMDVVKLLNKAIKKRGDYEADIMAVVNDTVGTMMTCGFDDQRCEVGIIIGTGTN
ACYMEELRHIDLVEGDEGRMCINTEWGAFGDDGSLEDIRTEFDREIDRGSINPGKQLFEKMASGMYMGEL
VRLILVKMAKEGLLFEGRITPELLTKGKIETKHVSAIEKTKEGLKKCMEILTRLGVEPSDEDCLAVQHVC
TIVSFRSANLISATLGGILSRLKENKGVARLRTTVGIDGSLYKMHPQYARRLHKTVRRLVPDSDVRFLLS
ESGSAKGAAMVTAVAYRLTEQARQIQQTLAEFRLSKAQLLEVKKRMRVEIERGLKKDTHEEATVKMLPTF
VRSTPDGTENGDFLALDLGGTNFRVLLVKIRSGKRRSVEMHNKIYAIPIEVMQGTGEELFDHIVHCISDF
LDYMGMKSARLPLGFTFSFPCHQTSLDAGILVTWTKGFKATDCEGEDVVELLREAIKRKEEFELDVVAIV
NDTVGTMMTCAYEEPTCEVGLIAGTGSNACYMEEMRNIEIVDGNEGRMCVNMEWGAFGDNGCLDDIRTKY
DQAVDENSLNEGKQRYEKMCSGMYLGEIVRQILIDLTKRGFLFRGQISETLKTRGIFETKFLSQIESDRL
ALLQVRAILQQLGLDSTCDDSIIVKEVCGTVSRRAAQICGAGMAAVVDKIRENRGLDHLDVTVGVDGTLY
KLHPHFSRIFHQTVKELAPKCDVTFLLSEDGSGKGAALITAVGCRQRELEAQQH
>anc6|avgPP:0.998|lnPP:-2.986|num_ambig:3|num_ambig_gaps:0
LYSMRFSDETLKDIMNRFRREMENGLGRDTNPTATVKMLPTFVRSIPDGSEKGDFIALDLGGSNFRILRV
KVTQDKKQPVQMESQVYETPDDIIHGSGTQLFDHVADCLGDFMEKQKIKDKKLPVGFTFSFPCAQTKLNE
AVLVTWTKKFKASGVEGMDVVKLLNKAIKKRGDYEADIMAVVNDTVGTMMTCGFDDQRCEVGIIIGTGTN
ACYMEELRHIDLVEGDEGRMCINTEWGAFGDDGSLEDIRTEFDREIDRGSINPGKQLFEKMASGMYMGEL
VRLILVKMAKEGLLFEGRITPELLTKGKIETKHVSAIEKTKEGLKKCMEILTRLGVEPSDEDCLAVQHVC
TIVSFRSANLISATLGGILSRLKENKGVARLRTTVGIDGSLYKMHPQYARRLHKTVRRLVPDSDVRFLLS
ESGSAKGAAMVTAVAYRLTEQARQIQQTLAEFRLSKAQLLEVKKRMRVEIERGLKKDTHEDATVKMLPTF
VRSTPDGTENGDFLALDLGGTNFRVLLVKIRSGKRRTVEMHNKIYAIPIEVMQGTGEELFDHIVHCISDF
LDYMGMKSARLPLGFTFSFPCHQTSLDAGILVTWTKGFKATDCEGEDVVELLREAIKRKEEFELDVVAIV
NDTVGTMMTCAYEEPTCEVGLIAGTGSNACYMEEMRNIEMVDGNEGRMCVNMEWGAFGDNGCLDDIRTKY
DRAVDENSLNEGKQRYEKMCSGMYLGEIVRNILIDLTKRGFLFRGQISETLKTRGIFETKFLSQIESDRL
ALLQVRAILQQLGLDSTCDDSIIVKEVCGTVSRRAAQICGAGMAAVVDKIRENRGLDHLDVTVGVDGTLY

KLHPHFSRIFHQTVKELAPKCDVNFLLSEDGSGKGAALITAVGCRQREQEAQQH
>anc6_altAll|avgPP:0.998|lnPP:-4.112|num_ambig:3|num_ambig_gaps:0
LYSMRFSDETLKDIMNRFRREMENGLGRDTNPTATVKMLPTFVRSIPDGSEKGDFIALDLGGSNFRILRV
KVTQDKKQPVQMESQVYETPDDIIHGSGTQLFDHVAECLGDFMEKQKIKDKKLPVGFTFSFPCAQTKLNE
AVLVTWTKKFKASGVEGMDVVKLLNKAIKKRGDYEADIMAVVNDTVGTMMTCGFDDQRCEVGIIIGTGTN
ACYMEELRHIDLVEGDEGRMCINTEWGAFGDDGSLEDIRTEFDREIDRGSINPGKQLFEKMASGMYMGEL
VRLILVKMAKEGLLFEGRITPELLTKGKIETKHVSAIEKTKEGLKKCMEILTRLGVEPSDEDCLAVQHVC
TIVSFRSANLISATLGGILSRLKENKGVARLRTTVGIDGSLYKMHPQYARRLHKTVRRLVPDSDVRFLLS
ESGSAKGAAMVTAVAYRLTEQARQIQQTLAEFRLSKAQLLEVKKRMRVEIERGLKKDTHEEATVKMLPTF
VRSTPDGTENGDFLALDLGGTNFRVLLVKIRSGKRRTVEMHNKIYAIPIEVMQGTGEELFDHIVHCISDF
LDYMGMKSARLPLGFTFSFPCHQTSLDAGILVTWTKGFKATDCEGEDVVELLREAIKRKEEFELDVVAIV
NDTVGTMMTCAYEEPTCEVGLIAGTGSNACYMEEMRNIEMVDGNEGRMCVNMEWGAFGDNGCLDDIRTKY
DQAVDENSLNEGKQRYEKMCSGMYLGEIVRNILIDLTKRGFLFRGQISETLKTRGIFETKFLSQIESDRL
ALLQVRAILQQLGLDSTCDDSIIVKEVCGTVSRRAAQICGAGMAAVVDKIRENRGLDHLDVTVGVDGTLY
KLHPHFSRIFHQTVKELAPKCDVNFLLSEDGSGKGAALITAVGCRQREQEAQQH
>anc7|avgPP:0.997|lnPP:-4.370|num_ambig:5|num_ambig_gaps:2
LYSMRFSDETLKDIMTRFRREMENGLGRDTNPTATVKMLPTFVRSIPDGSEKGDFIALDLGGSNFRILRV
KVTQDKKQPVQMESQVYETPDDIIHGSGTQLFDHVADCLGDFMEKQKIKDKKLPVGFTFSFPCAQTKLNE
AVLVTWTKKFKASGVEGMDVVKLLNKAIKKRGDYEADIMAVVNDTVGTMMTCGFDDQRCEVGIIIGTGTN
ACYMEELRHIDLVEGDEGRMCINTEWGAFGDDGSLEDIRTEFDREIDRGSLNPGKQLFEKMASGMYMGEL
VRLILVKMAKEGLLFEGRITPELLTKGKIETKHVSAIEKSKEGLKKCMEILTRLGVEPSDEDCLAVQHVC
TIVSFRSANLIAATLGGILSRLKENKGVARLRTTVGIDGSLYKMHPQYARRLHKTVRRLVPDSDVRFLLS
ESGSAKGAAMVTAVAYRLADQARQIQQTLAEFRLSKAQLLEVKKRMRTEIERGLKKDTHEDATVKMLPTF
VRSTPDGTENGDFLALDLGGTNFRVLLVKIRSGKRRTVEMHNKIYAIPIEVMQGTGEELFDHIVHCISDF
LDYMGMKSARLPLGFTFSFPCHQTSLDAGILVNWTKGFKATDCEGEDVVELLREGIKRKEEFELDVVAIV
NDTVGTMMTCAYEEPTCEVGLIAGTGSNACYMEEMRNIEMVDGNEGRMCVNMEWGAFGDNGCLDDIRTQY
DRAVDENSLNEGKQRYEKMCSGMYLGEIVRNILIDLTKRGFLFRGQISETLKTRGIFETKFLSQIESDRL
ALLQVRAILQQLGLDSTCDDSIIVKEVCGTVSRRAAQICGAGMAAVVDKIRENRGLDHLDITVGVDGTLY
KLHPHFSRIFHQTVKELAPKCDVNFLLSEDGSGKGAALITAVGCRQREQEAQQH
>anc7_altAll|avgPP:0.996|lnPP:-6.574|num_ambig:5|num_ambig_gaps:2
LYSMRFSDETLKDIMNRFRREMENGLGRDTNPTATVKMLPTFVRSIPDGSEKGDFIALDLGGSNFRILRV
KVTQDKKQPVQMESQVYETPDDIIHGSGTQLFDHVADCLGDFMEKQKIKDKKLPVGFTFSFPCAQTKLNE
AVLVTWTKKFKASGVEGMDVVKLLNKAIKKRGDYDADIMAVVNDTVGTMMTCGFDDQRCEVGIIIGTGTN
ACYMEELRHIDLVEGDEGRMCINTEWGAFGDDGSLEDIRTEFDREIDRGSLNPGKQLFEKMASGMYMGEL
VRLILVKMAKEGLLFEGRITPELLTKGKIETKHVSAIEKSKEGLKKCMEILTRLGVEPSDEDCLAVQHVC
TIVSFRSANLIAATLGGILSRLKENKGVARLRTTVGIDGSLYKMHPQYARRLHKTVRRLVPDSDVRFLLS
ESGSAKGAAMVTAVAYRLAEQARQIQQTLAEFRLSKAQLLEVKKRMRTEIERGLKKDTHEEATVKMLPTF
VRSTPDGTENGDFLALDLGGTNFRVLLVKIRSGKRRTVEMHNKIYAIPIEVMQGTGEELFDHIVHCISDF
LDYMGMKSARLPLGFTFSFPCHQTSLDAGILVNWTKGFKATDCEGEDVVELLREGIKRKEEFELDVVAIV
NDTVGTMMTCAYEEPTCEVGLIAGTGSNACYMEEMRNIEMVDGNEGRMCVNMEWGAFGDNGCLDDIRTQY
DRAVDENSLNEGKQRYEKMCSGMYLGEIVRNILIDLTKRGFLFRGQISETLKTRGIFETKFLSQIESDRL
ALLQVRAILQQLGLDSTCDDSIIVKEVCGTVSRRAAQICGAGMAAVVDKIRENRGLDHLDITVGVDGTLY
KLHPHFSRIFHQTVKELAPKCDVNFLLSEDGSGKGAALITAVGCRQREQEAEQH
>anc8|avgPP:0.997|lnPP:-4.370|num_ambig:5|num_ambig_gaps:2
LYSMRFSDETLKDIMTRFRREMENGLGRDTNPTATVKMLPTFVRSIPDGSEKGDFIALDLGGSNFRILRV
KVTQDKKQPVQMESQVYETPDDIIHGSGTQLFDHVADCLGDFMEKQKIKDKKLPVGFTFSFPCAQTKLNE
AVLVTWTKKFKASGVEGMDVVKLLNKAIKKRGDYEADIMAVVNDTVGTMMTCGFDDQRCEVGIIIGTGTN
ACYMEELRHIDLVEGDEGRMCINTEWGAFGDDGSLEDIRTEFDREIDRGSLNPGKQLFEKMASGMYMGEL
VRLILVKMAKEGLLFEGRITPELLTKGKIETKHVSAIEKSKEGLKKCMEILTRLGVEPSDEDCLAVQHVC
TIVSFRSANLIAATLGGILSRLKENKGVARLRTTVGIDGSLYKMHPQYARRLHKTVRRLVPDSDVRFLLS
ESGSAKGAAMVTAVAYRLADQARQIQQTLAEFRLSKAQLLEVKKRMRTEIERGLKKDTHEDATVKMLPTF
VRSTPDGTENGDFLALDLGGTNFRVLLVKIRSGKRRTVEMHNKIYAIPIEVMQGTGEELFDHIVHCISDF

```
LDYMGMKSARLPLGFTFSFPCHQTSLDAGILVNWTKGFKATDCEGEDVVELLREGIKRKEEFELDVVAIV
NDTVGTMMTCAYEEPTCEVGLIAGTGSNACYMEEMRNIEMVDGNEGRMCVNMEWGAFGDNGCLDDIRTQY
DRAVDENSLNEGKQRYEKMCSGMYLGEIVRNILIDLTKRGFLFRGQISETLKTRGIFETKFLSQIESDRL
ALLQVRAILQQLGLDSTCDDSIIVKEVCGTVSRRAAQICGAGMAAVVDKIRENRGLDHLDITVGVDGTLY
KLHPHFSRIFHQTVKELAPKCDVNFLLSEDGSGKGAALITAVGCRQREQEAQQH
>anc8_altAll|avgPP:0.996|lnPP:-6.574|num_ambig:5|num_ambig_gaps:2
LYSMRFSDETLKDIMNRFRREMENGLGRDTNPTATVKMLPTFVRSIPDGSEKGDFIALDLGGSNFRILRV
KVTQDKKQPVQMESQVYETPDDIIHGSGTQLFDHVADCLGDFMEKQKIKDKKLPVGFTFSFPCAQTKLNE
AVLVTWTKKFKASGVEGMDVVKLLNKAIKKRGDYDADIMAVVNDTVGTMMTCGFDDQRCEVGIIIGTGTN
ACYMEELRHIDLVEGDEGRMCINTEWGAFGDDGSLEDIRTEFDREIDRGSLNPGKQLFEKMASGMYMGEL
VRLILVKMAKEGLLFEGRITPELLTKGKIETKHVSAIEKSKEGLKKCMEILTRLGVEPSDEDCLAVQHVC
TIVSFRSANLIAATLGGILSRLKENKGVARLRTTVGIDGSLYKMHPQYARRLHKTVRRLVPDSDVRFLLS
ESGSAKGAAMVTAVAYRLAEQARQIQQTLAEFRLSKAQLLEVKKRMRTEIERGLKKDTHEEATVKMLPTF
VRSTPDGTENGDFLALDLGGTNFRVLLVKIRSGKRRTVEMHNKIYAIPIEVMQGTGEELFDHIVHCISDF
LDYMGMKSARLPLGFTFSFPCHQTSLDAGILVNWTKGFKATDCEGEDVVELLREGIKRKEEFELDVVAIV
NDTVGTMMTCAYEEPTCEVGLIAGTGSNACYMEEMRNIEMVDGNEGRMCVNMEWGAFGDNGCLDDIRTQY
DRAVDENSLNEGKQRYEKMCSGMYLGEIVRNILIDLTKRGFLFRGQISETLKTRGIFETKFLSQIESDRL
ALLQVRAILQQLGLDSTCDDSIIVKEVCGTVSRRAAQICGAGMAAVVDKIRENRGLDHLDITVGVDGTLY
KLHPHFSRIFHQTVKELAPKCDVNFLLSEDGSGKGAALITAVGCRQREQEAEQH
>anc9|avgPP:0.993|lnPP:-10.786|num_ambig:10|num_ambig_gaps:0
LYSMRFSDETLKDIMTRFRREMENGLGRDTNPTATVKMLPTFVRSIPDGSEKGDFIALDLGGSNFRILRV
KVSHEKKQTVQMESQIYETPEDIIHGSGTQLFDHVAECLGDFMEKQKIKDKKLPVGFTFSFPCRQTKLDE
GILITWTKRFKASGVEGADVVKLLNKAIKKRGDYDADIMAVVNDTVGTMMTCGFDDQRCEVGIIIGTGTN
ACYMEELRHIDLVEGDEGRMCINTEWGAFGDDGSLEDIRTEFDREIDRGSLNPGKQLFEKMVSGMYMGEL
VRLILVKMAKEGLLFEGRITPELLTKGKFETKHVSAIEKSKEGLNKAKEILTRLGVEPSHEDCIAVQHVC
TIVSFRSANLIAATLGGILSRLKDNKGVPRLRTTVGVDGSLYKMHPQYARRLHKTVRRLVPDSDVRFLLS
ESGSGKGAAMVTAVAYRLAEQHRQIEETLAEFKLSHAQLLEVKKRMRTEMERGLRKKTHESATVKMLPTF
VRSTPDGTENGDFLALDLGGTNFRVLLVKIRSGKRRTVEMHNKIYAIPIEVMQGTGEELFDHIVHCISDF
LDYMGMKSARLPLGFTFSFPCKQTSLDAGILINWTKGFKATDCEGEDVVNLLREGIKRREEFELDVVAIV
NDTVGTMMTCAYEEPTCEVGLIAGTGSNACYMEEMRNIEMVEGDEGRMCVNMEWGAFGDNGCLDDIRTQY
DRAVDELSLNPGKQRYEKMCSGMYLGEIVRNILIDLTKRGFLFRGQISETLKTRGIFETKFLSQIESDRL
ALLQVRAILQQLGLDSTCDDSIIVKEVCGAVSRRAAQLCGAGMAAVVDKIRENRGLDHLEITVGVDGTLY
KLHPHFSKIMHQTVKELAPKCDVTFLLSEDGSGKGAALITAVGCRLREAEQQ
>anc9_altAll|avgPP:0.992|lnPP:-14.534|num_ambig:10|num_ambig_gaps:0
LYSMRFSDETLTDIMTRFRREMENGLGRDTNPTATVKMLPTFVRSIPDGSEKGDFIALDLGGSNFRILRV
KVSHEKKQTVQMESQIYETPEDIIHGSGTQLFDHVAECLGDFMEKQKIKDKKLPVGFTFSFPCRQTKLDE
GVLITWTKRFKASGVEGADVVKLLNKAIKKRGDYDADIMAVVNDTVGTMMTCGFDDQRCEVGIIIGTGTN
ACYMEELRHIDLVEGDEGRMCINTEWGAFGDDGSLEDIRTEFDREIDRGSLNPGKQLFEKMVSGMYMGEL
VRLILVKMAKEGLLFEGRITPELLTKGKFETKHVSAIEKSKEGLNKAKEILTRLGVEPSHEDCIAVQHVC
TIVSFRSANLIAATLGGILSRLKDNKGVPRLRTTVGIDGSLYKMHPQYARRLHKTVRRLVPDSDVRFLLS
ESGSGKGAAMVTAVAYRLAEQHRQIEETLAEFKLSHVQLLEVKKRMRTEMERGLRKKTHESATVKMLPTF
VRSTPDGTENGDFLALDLGGTNFRVLLVKIRSGKRRSVEMHNKIYAIPIEVMQGTGEELFDHIVHCISDF
LDYMGMKNARLPLGFTFSFPCKQTSLDAGILVNWTKGFKATDCEGEDVVNLLREGIKRREEFELDVVAIV
NDTVGTMMTCAYEEPTCEVGLIAGTGSNACYMEEMRNIEMVDGDEGRMCVNMEWGAFGDNGCLDDIRTQY
DRAVDELSLNPGKQRYEKMCSGMYLGEIVRNILIDLTKRGFLFRGQISEALKTRGIFETKFLSQIESDRL
ALLQVRAILQQLGLDSTCDDSIIVKEVCGAVSRRAAQLCGAGMAAVVDKIRENRGLDHLEITVGVDGTLY
KLHPHFSKIMHQTVKELAPKCDVTFLLSEDGSGKGAALITAVGCRLREAEQH
>anc10|avgPP:1.000|lnPP:0.000|num_ambig:0|num_ambig_gaps:0
LYAMRLSDETLLDVMARFRREMKNGLSRDFNPTAAVKMLPTFVRSIPDGSEKGDFIALDLGGSYFRILRV
KVSHEKKQTVQMESEIYNTPEDIMHGSGTRLFDHVAECLGDFMEKQQIKDKKLPVGFTFSFPCRQSKLDE
GILITWTKRFKASGVEGADVVRLLNKAIKKRGDYDADIMAVVNDTVGTMMTCGFDDQRCEVGLIIGTGTN
ACYMEEMRHIDLVEGDEGRMCINTEWGAFGDDGSLEDIRTEFDREIDRGSLNPGKQLFEKMVSGLYMGEL
```

VRLILVKMAKEGLLFEGRITPELLTKGKFETKHVSAIEKSKEGLNKAKEILTRLGVEPSHEDCIAVQHVC
TIVSFRSANLVASTLGAILNQLRDNKGVGRLRTTVGVDGSLYKMHPQYARRLHKTTRRLVPDSEVRFLLS
ESGSGKGAAMVTAVAYRLSEQHRLIDETLAEFKLTHEQLLQVKKRMRAEMEAGLKKKTHETAKVKMLPTF
VRSTPDGTENGDFLALDLGGTNFRVLLVKIRSGKRRTVEMHNKIYAIPIEVMQGTGEELFDHIVTCISDF
LDYMGIKGARLPLGFTFSFPCKQTSLDAGILLNWTKGFKATDCEGEDVVYLLREGIKRREEFDLDVVAVV
NDTVGTMMTCAYEDPNCEIGLIVGTGSNACYMEEMRNIEMVDGEQGRMCVNTEWGAFGDNGCLDDIRTIY
DKAVDDFSLNAGKQRYEKMISGMYLGEIVRNILIDFTKRGFLFRGQISETLKTRHIFETKFLSQIESDRL
ALLQVRTILQQLGLNSTCDDSIIVKTVCGAVSRRAAQLCGAGMAAIVDKIRENRGLERLEITVGVDGTLY
KLHPHFSRIMHQTVKDLAPNCDVTFLLSEDGSGKGAALITAVGCRLREAEQN
>anc10_altAll|avgPP:1.000|lnPP:0.000|num_ambig:0|num_ambig_gaps:0
LYAMRLSDETLLDVMARFRREMKNGLSRDFNPTAAVKMLPTFVRSIPDGSEKGDFIALDLGGSYFRILRV
KVSHEKKQTVQMESEIYNTPEDIMHGSGTRLFDHVAECLGDFMEKQQIKDKKLPVGFTFSFPCRQSKLDE
GILITWTKRFKASGVEGADVVRLLNKAIKKRGDYDADIMAVVNDTVGTMMTCGFDDQRCEVGLIIGTGTN
ACYMEEMRHIDLVEGDEGRMCINTEWGAFGDDGSLEDIRTEFDREIDRGSLNPGKQLFEKMVSGLYMGEL
VRLILVKMAKEGLLFEGRITPELLTKGKFETKHVSAIEKSKEGLNKAKEILTRLGVEPSHEDCIAVQHVC
TIVSFRSANLVASTLGAILNQLRDNKGVGRLRTTVGVDGSLYKMHPQYARRLHKTTRRLVPDSEVRFLLS
ESGSGKGAAMVTAVAYRLSEQHRLIDETLAEFKLTHEQLLQVKKRMRAEMEAGLKKKTHETAKVKMLPTF
VRSTPDGTENGDFLALDLGGTNFRVLLVKIRSGKRRTVEMHNKIYAIPIEVMQGTGEELFDHIVTCISDF
LDYMGIKGARLPLGFTFSFPCKQTSLDAGILLNWTKGFKATDCEGEDVVYLLREGIKRREEFDLDVVAVV
NDTVGTMMTCAYEDPNCEIGLIVGTGSNACYMEEMRNIEMVDGEQGRMCVNTEWGAFGDNGCLDDIRTIY
DKAVDDFSLNAGKQRYEKMISGMYLGEIVRNILIDFTKRGFLFRGQISETLKTRHIFETKFLSQIESDRL
ALLQVRTILQQLGLNSTCDDSIIVKTVCGAVSRRAAQLCGAGMAAIVDKIRENRGLERLEITVGVDGTLY
KLHPHFSRIMHQTVKDLAPNCDVTFLLSEDGSGKGAALITAVGCRLREAEQN
>anc11|avgPP:1.000|lnPP:0.000|num_ambig:0|num_ambig_gaps:0
LYAMRLSDETLLDVMARFRREMKNGLSRDFNPTAAVKMLPTFVRSIPDGSEKGDFIALDLGGSYFRILRV
KVSHEKKQTVQMESEIYNTPEDIMHGSGTRLFDHVAECLGDFMEKQQIKDKKLPVGFTFSFPCRQSKLDE
GILITWTKRFKASGVEGADVVRLLNKAIKKRGDYDADIMAVVNDTVGTMMTCGFDDQRCEVGLIIGTGTN
ACYMEEMRHIDLVEGDEGRMCINTEWGAFGDDGSLEDIRTEFDREIDRGSLNPGKQLFEKMVSGLYMGEL
VRLILVKMAKEGLLFEGRITPELLTKGKFETKHVSAIEKSKEGLNKAKEILTRLGVEPSHEDCIAVQHVC
TIVSFRSANLVASTLGAILNQLRDNKGVGRLRTTVGVDGSLYKMHPQYARRLHKTTRRLVPDSEVRFLLS
ESGSGKGAAMVTAVAYRLSEQHRLIDETLAEFKLTHEQLLQVKKRMRAEMEAGLKKKTHETAKVKMLPTF
VRSTPDGTENGDFLALDLGGTNFRVLLVKIRSGKRRTVEMHNKIYAIPIEVMQGTGEELFDHIVTCISDF
LDYMGIKGARLPLGFTFSFPCKQTSLDAGILLNWTKGFKATDCEGEDVVYLLREGIKRREEFDLDVVAVV
NDTVGTMMTCAYEDPNCEIGLIVGTGSNACYMEEMRNIEMVDGEQGRMCVNTEWGAFGDNGCLDDIRTIY
DKAVDDYSLNAGKQRYEKMISGMYLGEIVRNILIDFTKRGFLFRGQISETLKTRHIFETKFLSQIESDRL
ALLQVRTILQQLGLNSTCDDSIIVKTVCGAVSRRAAQLCGAGMAAIVDKIRENRGLERLEITVGVDGTLY
KLHPHFSRIMHQTVKDLAPNCDVTFLLSEDGSGKGAALITAVGCRLREAEQN
>anc11_altAll|avgPP:1.000|lnPP:0.000|num_ambig:0|num_ambig_gaps:0
LYAMRLSDETLLDVMARFRREMKNGLSRDFNPTAAVKMLPTFVRSIPDGSEKGDFIALDLGGSYFRILRV
KVSHEKKQTVQMESEIYNTPEDIMHGSGTRLFDHVAECLGDFMEKQQIKDKKLPVGFTFSFPCRQSKLDE
GILITWTKRFKASGVEGADVVRLLNKAIKKRGDYDADIMAVVNDTVGTMMTCGFDDQRCEVGLIIGTGTN
ACYMEEMRHIDLVEGDEGRMCINTEWGAFGDDGSLEDIRTEFDREIDRGSLNPGKQLFEKMVSGLYMGEL
VRLILVKMAKEGLLFEGRITPELLTKGKFETKHVSAIEKSKEGLNKAKEILTRLGVEPSHEDCIAVQHVC
TIVSFRSANLVASTLGAILNQLRDNKGVGRLRTTVGVDGSLYKMHPQYARRLHKTTRRLVPDSEVRFLLS
ESGSGKGAAMVTAVAYRLSEQHRLIDETLAEFKLTHEQLLQVKKRMRAEMEAGLKKKTHETAKVKMLPTF
VRSTPDGTENGDFLALDLGGTNFRVLLVKIRSGKRRTVEMHNKIYAIPIEVMQGTGEELFDHIVTCISDF
LDYMGIKGARLPLGFTFSFPCKQTSLDAGILLNWTKGFKATDCEGEDVVYLLREGIKRREEFDLDVVAVV
NDTVGTMMTCAYEDPNCEIGLIVGTGSNACYMEEMRNIEMVDGEQGRMCVNTEWGAFGDNGCLDDIRTIY
DKAVDDYSLNAGKQRYEKMISGMYLGEIVRNILIDFTKRGFLFRGQISETLKTRHIFETKFLSQIESDRL
ALLQVRTILQQLGLNSTCDDSIIVKTVCGAVSRRAAQLCGAGMAAIVDKIRENRGLERLEITVGVDGTLY
KLHPHFSRIMHQTVKDLAPNCDVTFLLSEDGSGKGAALITAVGCRLREAEQN
>anc12|avgPP:0.999|lnPP:-0.974|num_ambig:1|num_ambig_gaps:0

LYAMRLSDETLIDVMARFRREMKNGLSRDFNPTAAVKMLPTFVRSIPDGSEKGDFIALDLGGSYFRILRV
KVSHEKKQTVQMESEIYNTPEDIMHGSGTRLFDHVAECLGDFMEKQQIKDKKLPVGFTFSFPCRQSKLDE
GILITWTKRFKASGVEGADVVRLLNKAIKKRGDYDADIMAVVNDTVGTMMTCGFDDQRCEVGLIIGTGTN
ACYMEEMRHIDLVEGDEGRMCINTEWGAFGDDGSLEDIRTEFDREIDRGSLNPGKQLFEKMVSGLYMGEL
VRLILVKMAKEGLLFEGRITPELLTKGKFETKHVSAIEKSKEGLNKAKEILTRLGVEPSHEDCIAVQHVC
TIVSFRSANLVASTLGAILNQLRDNKGVGRLRTTVGVDGSLYKMHPQYARRLHKTTRRLVPDSEVRFLLS
ESGSGKGAAMVTAVAYRLSEQHRLIDETLAEFKLTHEQLLQVKKRMRAEMEAGLKKKTHETAKVKMLPTF
VRSTPDGTENGDFLALDLGGTNFRVLLVKIRSGKRRTVEMHNKIYAIPIEVMQGTGEELFDHIVTCISDF
LDYMGIKGARLPLGFTFSFPCKQTSLDAGILLNWTKGFKATDCEGEDVVYLLREGIKRREEFDLDVVAVV
NDTVGTMMTCAYEDPNCEIGLIVGTGSNACYMEEMRNIEMVDGEQGRMCVNTEWGAFGDNGCLDDIRTIY
DKAVDDYSLNAGKQRYEKMISGMYLGEIVRNILIDFTKRGFLFRGQISETLKTRHIFETKFLSQIESDRL
ALLQVRAILQQLGLNSTCDDSIIVKTVCGAVSRRAAQLCGAGMAAVVDKIRENRGLEHLEITVGVDGTLY
KLHPHFSRIMHQTVKDLAPNCDVTFLLSEDGSGKGAALITAVGCRLRDAEQN
>anc12_altAll|avgPP:0.999|lnPP:-1.167|num_ambig:1|num_ambig_gaps:0
LYAMRLSDETLIDVMARFRREMKNGLSRDFNPTAAVKMLPTFVRSIPDGSEKGDFIALDLGGSYFRILRV
KVSHEKKQTVQMESEIYNTPEDIMHGSGTRLFDHVAECLGDFMEKQQIKDKKLPVGFTFSFPCRQSKLDE
GILITWTKRFKASGVEGADVVRLLNKAIKKRGDYDADIMAVVNDTVGTMMTCGFDDQRCEVGLIIGTGTN
ACYMEEMRHIDLVEGDEGRMCINTEWGAFGDDGSLEDIRTEFDREIDRGSLNPGKQLFEKMVSGLYMGEL
VRLILVKMAKEGLLFEGRITPELLTKGKFETKHVSAIEKSKEGLNKAKEILTRLGVEPSHEDCIAVQHVC
TIVSFRSANLVASTLGAILNQLRDNKGVGRLRTTVGVDGSLYKMHPQYARRLHKTTRRLVPDSEVRFLLS
ESGSGKGAAMVTAVAYRLSEQHRLIDETLAEFKLTHEQLLQVKKRMRAEMEAGLKKKTHETAKVKMLPTF
VRSTPDGTENGDFLALDLGGTNFRVLLVKIRSGKRRTVEMHNKIYAIPIEVMQGTGEELFDHIVTCISDF
LDYMGIKGARLPLGFTFSFPCKQTSLDAGILLNWTKGFKATDCEGEDVVYLLREGIKRREEFDLDVVAVV
NDTVGTMMTCAYEDPNCEIGLIVGTGSNACYMEEMRNIEMVDGEQGRMCVNTEWGAFGDNGCLDDIRTIY
DKAVDDYSLNAGKQRYEKMISGMYLGEIVRNILIDFTKRGFLFRGQISETLKTRHIFETKFLSQIESDRL
ALLQVRAILQQLGLNSTCDDSIIVKTVCGAVSRRAAQLCGAGMAAIVDKIRENRGLEHLEITVGVDGTLY
KLHPHFSRIMHQTVKDLAPNCDVTFLLSEDGSGKGAALITAVGCRLRDAEQN
>anc13|avgPP:1.000|lnPP:-0.000|num_ambig:0|num_ambig_gaps:0
LYAMRLSDETLIDIMARFRREMKNGLSRDFNPTATVKMLPTFVRSIPDGSEKGDFIALDLGGSYFRILRV
KVSHEKKQTVQMESEIYNTPEDIMHGSGTRLFDHVAECLGDFMEKQQIKDKKLPVGFTFSFPCRQSKLDE
GILITWTKRFKASGVEGADVVRLLNKAIKKRGDYDADIMAVVNDTVGTMMTCGFDDQRCEVGLIIGTGTN
ACYMEEMRHIDLVEGDEGRMCINTEWGAFGDDGSLEDIRTEFDREIDRGSLNPGKQLFEKMVSGLYMGEL
VRLILVKMAKEGLLFEGRITPELLTKGKFETKHVSAIEKSKEGLNKAKEILTRLGVEPSHEDCIAVQHVC
TIVSFRSANLVASTLGAILNQLRDNKGVGRLRTTVGVDGSLYKMHPQYARRLHKTTRRLVPDSEVRFLLS
ESGSGKGAAMVTAVAYRLSEQHRLIDETLAEFKLTHEQLLQVKKRMRAEMEAGLKKKTHETAKVKMLPTF
VRSTPDGTENGDFLALDLGGTNFRVLLVKIRSGKRRTVEMHNKIYAIPIEVMQGTGEELFDHIVTCISDF
LDYMGIKGARLPLGFTFSFPCKQTSLDAGILLNWTKGFKATDCEGEDVVYLLREGIKRREEFDLDVVAVV
NDTVGTMMTCAYEDPNCEIGLIVGTGSNACYMEEMRNIEMVDGEQGRMCVNTEWGAFGDNGCLDDIRTIY
DKAVDDYSLNAGKQRYEKMISGMYLGEIVRNILIDFTKRGFLFRGQISETLKTRHIFETKFLSQIESDRL
ALLQVRAILQQLGLNSTCDDSIIVKTVCGAVSRRAAQLCGAGMAAVVDKIRENRGLEHLEITVGVDGTLY
KLHPHFSRIMHQTVKDLAPNCDVTFLLSEDGSGKGAALITAVGCRLRDAEQN
>anc13_altAll|avgPP:1.000|lnPP:-0.000|num_ambig:0|num_ambig_gaps:0
LYAMRLSDETLIDIMARFRREMKNGLSRDFNPTATVKMLPTFVRSIPDGSEKGDFIALDLGGSYFRILRV
KVSHEKKQTVQMESEIYNTPEDIMHGSGTRLFDHVAECLGDFMEKQQIKDKKLPVGFTFSFPCRQSKLDE
GILITWTKRFKASGVEGADVVRLLNKAIKKRGDYDADIMAVVNDTVGTMMTCGFDDQRCEVGLIIGTGTN
ACYMEEMRHIDLVEGDEGRMCINTEWGAFGDDGSLEDIRTEFDREIDRGSLNPGKQLFEKMVSGLYMGEL
VRLILVKMAKEGLLFEGRITPELLTKGKFETKHVSAIEKSKEGLNKAKEILTRLGVEPSHEDCIAVQHVC
TIVSFRSANLVASTLGAILNQLRDNKGVGRLRTTVGVDGSLYKMHPQYARRLHKTTRRLVPDSEVRFLLS
ESGSGKGAAMVTAVAYRLSEQHRLIDETLAEFKLTHEQLLQVKKRMRAEMEAGLKKKTHETAKVKMLPTF
VRSTPDGTENGDFLALDLGGTNFRVLLVKIRSGKRRTVEMHNKIYAIPIEVMQGTGEELFDHIVTCISDF
LDYMGIKGARLPLGFTFSFPCKQTSLDAGILLNWTKGFKATDCEGEDVVYLLREGIKRREEFDLDVVAVV
NDTVGTMMTCAYEDPNCEIGLIVGTGSNACYMEEMRNIEMVDGEQGRMCVNTEWGAFGDNGCLDDIRTIY

DKAVDDYSLNAGKQRYEKMISGMYLGEIVRNILIDFTKRGFLFRGQISETLKTRHIFETKFLSQIESDRL
ALLQVRAILQQLGLNSTCDDSIIVKTVCGAVSRRAAQLCGAGMAAVVDKIRENRGLEHLEITVGVDGTLY
KLHPHFSRIMHQTVKDLAPNCDVTFLLSEDGSGKGAALITAVGCRLRDAEQN
>anc14|avgPP:1.000|lnPP:-0.000|num_ambig:0|num_ambig_gaps:0
LYAMRLSDETLIDIMARFRREMKNGLSRDFNPTATVKMLPTFVRSIPDGSEKGDFIALDLGGSYFRILRV
KVSHEKKQTVQMESEIYNTPEDIMHGSGTRLFDHVAECLGDFMEKQQIKDKKLPVGFTFSFPCRQSKLDE
GILITWTKRFKASGVEGADVVRLLNKAIKKRGDYDADIMAVVNDTVGTMMTCGFDDQRCEVGLIIGTGTN
ACYMEEMRHIDLVEGDEGRMCINTEWGAFGDDGSLEDIRTEFDREIDRGSLNPGKQLFEKMVSGLYMGEL
VRLILVKMAKEGLLFEGRITPELLTKGKFETKHVSAIEKSKEGLNKAKEILTRLGVEPSHEDCIAVQHVC
TIVSFRSANLVASTLGAILNQLRDNKGVGRLRTTVGVDGSLYKMHPQYARRLHKTTRRLVPDSEVRFLLS
ESGSGKGAAMVTAVAYRLSEQHRLIDETLAEFKLTHEQLLQVKKRMRAEMEAGLKKKTHETAKVKMLPTF
VRSTPDGTENGDFLALDLGGTNFRVLLVKIRSGKRRTVEMHNKIYAIPIEVMQGTGEELFDHIVTCISDF
LDYMGIKGARLPLGFTFSFPCKQTSLDAGILLNWTKGFKATDCEGEDVVYLLREGIKRREEFDLDVVAVV
NDTVGTMMTCAYEDPNCEIGLIVGTGSNACYMEEMRNIEMVDGEQGRMCVNTEWGAFGDNGCLDDIRTIY
DKAVDDYSLNAGKQRYEKMISGMYLGEIVRNILIDFTKRGFLFRGQISETLKTRHIFETKFLSQIESDRL
ALLQVRAILQQLGLNSTCDDSIIVKTVCGAVSRRAAQLCGAGMAAVVDKIRENRGLEHLEITVGVDGTLY
KLHPHFSRIMHQTVKDLAPNCDVTFLLSEDGSGKGAALITAVGCRLRDAEQN
>anc14_altAll|avgPP:1.000|lnPP:-0.000|num_ambig:0|num_ambig_gaps:0
LYAMRLSDETLIDIMARFRREMKNGLSRDFNPTATVKMLPTFVRSIPDGSEKGDFIALDLGGSYFRILRV
KVSHEKKQTVQMESEIYNTPEDIMHGSGTRLFDHVAECLGDFMEKQQIKDKKLPVGFTFSFPCRQSKLDE
GILITWTKRFKASGVEGADVVRLLNKAIKKRGDYDADIMAVVNDTVGTMMTCGFDDQRCEVGLIIGTGTN
ACYMEEMRHIDLVEGDEGRMCINTEWGAFGDDGSLEDIRTEFDREIDRGSLNPGKQLFEKMVSGLYMGEL
VRLILVKMAKEGLLFEGRITPELLTKGKFETKHVSAIEKSKEGLNKAKEILTRLGVEPSHEDCIAVQHVC
TIVSFRSANLVASTLGAILNQLRDNKGVGRLRTTVGVDGSLYKMHPQYARRLHKTTRRLVPDSEVRFLLS
ESGSGKGAAMVTAVAYRLSEQHRLIDETLAEFKLTHEQLLQVKKRMRAEMEAGLKKKTHETAKVKMLPTF
VRSTPDGTENGDFLALDLGGTNFRVLLVKIRSGKRRTVEMHNKIYAIPIEVMQGTGEELFDHIVTCISDF
LDYMGIKGARLPLGFTFSFPCKQTSLDAGILLNWTKGFKATDCEGEDVVYLLREGIKRREEFDLDVVAVV
NDTVGTMMTCAYEDPNCEIGLIVGTGSNACYMEEMRNIEMVDGEQGRMCVNTEWGAFGDNGCLDDIRTIY
DKAVDDYSLNAGKQRYEKMISGMYLGEIVRNILIDFTKRGFLFRGQISETLKTRHIFETKFLSQIESDRL
ALLQVRAILQQLGLNSTCDDSIIVKTVCGAVSRRAAQLCGAGMAAVVDKIRENRGLEHLEITVGVDGTLY
KLHPHFSRIMHQTVKDLAPNCDVTFLLSEDGSGKGAALITAVGCRLRDAEQN
>anc15|avgPP:0.999|lnPP:-1.024|num_ambig:2|num_ambig_gaps:0
LYAMRLSDETLIDIMARFRREMKNGLSRDFNPTATVKMLPTFVRSIPDGSEKGDFIALDLGGSYFRILRV
KVSHEKKQTVQMETEIYNTPEDIMHGSGTRLFDHVAECLGDFMEKQQIKDKKLPVGFTFSFPCRQSKLDE
GILITWTKRFKASGVEGADVVKLLNKAIKKRGDYDADIMAVVNDTVGTMMTCGFDDQRCEVGLIIGTGTN
ACYMEEMRHIDLVEGDEGRMCINTEWGAFGDDGSLEDIRTEFDREIDRGSLNPGKQLFEKMVSGLYMGEL
VRLILVKMAKEGLLFEGRITPELLTKGKFETKHVSAIEKSKEGLNKAKEILTRLGVEPSHEDCIAVQHVC
TIVSFRSANLVASTLGAILNQLRDNKGVGRLRTTVGVDGSLYKMHPQYARRLHKTTRRLVPDSDVRFLLS
ESGSGKGAAMVTAVAYRLSEQHRLIDETLAEFKLTHEQLLQVKKRMRTEMEAGLRKKTHETAKVKMLPTF
VRSTPDGTENGDFLALDLGGTNFRVLLVKIRSGKRRTVEMHNKIYAIPIEVMQGTGEELFDHIVTCISDF
LDYMGIKGARLPLGFTFSFPCKQTSLDAGILLNWTKGFKATDCEGEDVVYLLREGIKRREEFDLDVVAVV
NDTVGTMMTCAYEDPNCEIGLIVGTGSNACYMEEMRNIEMVDGEQGRMCVNMEWGAFGDNGCLDDIRTIY
DKAVDDYSLNAGKQRYEKMISGMYLGEIVRNILIDFTKRGFLFRGQISETLKTRHIFETKFLSQIESDRL
ALLQVRAILQQLGLNSTCDDSIIVKTVCGAVSRRAAQLCGAGMAAVVDKIRENRGLEHLEITVGVDGTLY
KLHPHFSRIMHQTVKDLAPKCDVTFLLSEDGSGKGAALITAVGCRLRDAEQN
>anc15_altAll|avgPP:0.999|lnPP:-2.192|num_ambig:2|num_ambig_gaps:0
LYAMRLSDETLIDIMARFRREMKNGLSRDFNPTATVKMLPTFVRSIPDGSEKGDFIALDLGGSYFRILRV
KVSHEKKQTVQMETEIYNTPEDIMHGSGTRLFDHVAECLGDFMEKQQIKDKKLPVGFTFSFPCRQSKLDE
GILITWTKRFKASGVEGADVVKLLNKAIKKRGDYDADIMAVVNDTVGTMMTCGFDDQRCEVGLIIGTGTN
ACYMEEMRHIDLVEGDEGRMCINTEWGAFGDDGSLEDIRTEFDREIDRGSLNPGKQLFEKMVSGLYMGEL
VRLILVKMAKEGLLFEGRITPELLTKGKFETKHVSAIEKSKEGLNKAKEILTRLGVEPSHEDCIAVQHVC
TIVSFRSANLVAATLGAILNQLRDNKGVGRLRTTVGVDGSLYKMHPQYARRLHKTTRRLVPDSDVRFLLS

ESGSGKGAAMVTAVAYRLSEQHRLIDETLAEFKLTHEQLLQVKKRMRTEMEAGLKKKTHETAKVKMLPTF
VRSTPDGTENGDFLALDLGGTNFRVLLVKIRSGKRRTVEMHNKIYAIPIEVMQGTGEELFDHIVTCISDF
LDYMGIKGARLPLGFTFSFPCKQTSLDAGILLNWTKGFKATDCEGEDVVYLLREGIKRREEFDLDVVAVV
NDTVGTMMTCAYEDPNCEIGLIVGTGSNACYMEEMRNIEMVDGEQGRMCVNMEWGAFGDNGCLDDIRTIY
DKAVDDYSLNAGKQRYEKMISGMYLGEIVRNILIDFTKRGFLFRGQISETLKTRHIFETKFLSQIESDRL
ALLQVRAILQQLGLNSTCDDSIIVKTVCGAVSRRAAQLCGAGMAAVVDKIRENRGLEHLEITVGVDGTLY
KLHPHFSRIMHQTVKDLAPKCDVTFLLSEDGSGKGAALITAVGCRLRDAEQN
>anc16|avgPP:0.998|lnPP:-2.903|num_ambig:5|num_ambig_gaps:0
LYAMRLSDETLLDIMARFRREMKNGLSRDFNPTATVKMLPTFVRSIPDGSEKGDFIALDLGGSYFRILRV
KVSHEKKQTVQMETEIYNTPEDIMHGSGTRLFDHVAECLGDFMEKQQIKDKKLPVGFTFSFPCRQSKLDE
GILITWTKRFKASGVEGADVVKLLNKAIKKRGDYDADIMAVVNDTVGTMMTCGFDDQRCEVGLIIGTGTN
ACYMEEMRHIDLVEGDEGRMCINTEWGAFGDDGSLEDIRTEFDREIDRGSLNPGKQLFEKMVSGMYMGEL
VRLILVKMAKEGLLFEGRITPELLTKGKFETKHVSAIEKSKEGLNKAKEILTRLGVEPSHEDCIAVQHVC
TIVSFRSANLVASTLGAILNQLRDNKGVPRLRTTVGVDGSLYKMHPQYARRLHKTTRRLVPDSDVRFLLS
ESGSGKGAAMVTAVAYRLAEQHRLIDETLAEFKLTHEQLLQVKKRMRTEMEAGLRKKTHETAKVKMLPTF
VRSTPDGTENGDFLALDLGGTNFRVLLVKIRSGKRRTVEMHNKIYAIPIEVMQGTGEELFDHIVTCISDF
LDYMGIKGARLPLGFTFSFPCKQTSLDAGILLNWTKGFKATDCEGEDVVYLLREGIKRREEFDLDVVAVV
NDTVGTMMTCAYEDPNCEIGLIVGTGSNACYMEEMRNIEMVEGDQGRMCVNMEWGAFGDNGCLDDIRTIY
DKAVDDYSLNAGKQRYEKMISGMYLGEIVRNILIDFTKRGFLFRGQISETLKTRSIFETKFLSQIESDRL
ALLQVRAILQQLGLNSTCDDSIIVKTVCGAVSRRAAQLCGAGMAAVVDKIRENRGLEHLEITVGVDGTLY
KLHPHFSRIMHQTVKDLAPKCDVTFLLSEDGSGKGAALITAVGCRLRDAEQN
>anc16_altAll|avgPP:0.997|lnPP:-5.497|num_ambig:5|num_ambig_gaps:0
LYAMRLSDETLLDIMARFRREMKNGLSRDFNPTATVKMLPTFVRSIPDGSEKGDFIALDLGGSYFRILRV
KVSHEKKQTVQMETEIYNTPEDIMHGSGTRLFDHVAECLGDFMEKQQIKDKKLPVGFTFSFPCRQSKLDE
GILITWTKRFKASGVEGADVVKLLNKAIKKRGDYDADIMAVVNDTVGTMMTCGFDDQRCEVGLIIGTGTN
ACYMEEMRHIDLVEGDEGRMCINTEWGAFGDDGSLEDIRTEFDREIDRGSLNPGKQLFEKMVSGMYMGEL
VRLILVKMAKEGLLFEGRITPELLTKGKFETKHVSAIEKSKEGLNKAKEILTRLGVEPSHEDCIAVQHVC
TIVSFRSANLVAATLGAILNQLRDNKGVPRLRTTVGVDGSLYKMHPQYARRLHKTTRRLVPDSDVRFLLS
ESGSGKGAAMVTAVAYRLSEQHRLIDETLAEFKLTHEQLLQVKKRMRTEMEAGLKKKTHESAKVKMLPTF
VRSTPDGTENGDFLALDLGGTNFRVLLVKIRSGKRRTVEMHNKIYAIPIEVMQGTGEELFDHIVTCISDF
LDYMGIKGARLPLGFTFSFPCKQTSLDAGILLNWTKGFKATDCEGEDVVYLLREGIKRREEFDLDVVAVV
NDTVGTMMTCAYEDPNCEIGLIVGTGSNACYMEEMRNIEMVEGDQGRMCVNMEWGAFGDNGCLDDIRTIY
DKAVDDYSLNAGKQRYEKMISGMYLGEIVRNILIDFTKRGFLFRGQISETLKTRGIFETKFLSQIESDRL
ALLQVRAILQQLGLNSTCDDSIIVKTVCGAVSRRAAQLCGAGMAAVVDKIRENRGLEHLEITVGVDGTLY
KLHPHFSRIMHQTVKDLAPKCDVTFLLSEDGSGKGAALITAVGCRLRDAEQN
>anc17|avgPP:0.999|lnPP:-2.190|num_ambig:3|num_ambig_gaps:0
LYAMRLSDETLLDIMARFRREMKNGLSRDFNPTATVKMLPTFVRSIPDGSEKGDFIALDLGGSYFRILRV
KVSHEKKQTVQMETEIYDTPEDIMHGSGTRLFDHVAECLGDFMEKQQIKDKKLPVGFTFSFPCRQSKLDE
GILITWTKRFKASGVEGADVVKLLNKAIKKRGDYDADIMAVVNDTVGTMMTCGFDDQRCEVGLIIGTGTN
ACYMEEMRHIDLVEGDEGRMCINTEWGAFGDDGSLEDIRTEFDREIDRGSLNPGKQLFEKMVSGMYMGEL
VRLILVKMAKEGLLFEGRITPELLTKGKFETKHVSAIEKSKEGLNKAKEILTRLGVEPSHEDCIAVQHVC
TIVSFRSANLVASTLGAILNQLRDNKGVPRLRTTVGVDGSLYKMHPQYSRRLHKTTRRLVPDSDVRFLLS
ESGSGKGAAMVTAVAYRLAEQHRLIDETLAEFKLTHEQLLQVKKRMRTEIEAGLRKKTHETAKVKMLPTF
VRSTPDGTENGDFLALDLGGTNFRVLLVKIRSGKRRTVEMHNKIYAIPIEVMQGTGEELFDHIVTCISDF
LDYMGIKGARLPLGFTFSFPCKQTSLDAGILLNWTKGFKATDCEGEDVVYLLREGIKRREEFDLDVVAVV
NDTVGTMMTCAYEDPNCEIGLIVGTGSNACYMEEMRNIEMVEGDQGRMCVNMEWGAFGDNGCLDDIRTIY
DKAVDDYSLNAGKQRYEKMISGMYLGEIVRNILIDFTKRGFLFRGQISETLKTRGIFETKFLSQIESDRL
ALLQVRAILQQLGLNGTCDDSIIVKTVCGAVSRRAAQLCGAGMAAVVDKIRENRGLDHLEITVGVDGTLY
KLHPHFSRIMHQTVKDLAPKCDVTFLLSEDGSGKGAALITAVGCRLRDAEQN
>anc17_altAll|avgPP:0.998|lnPP:-3.497|num_ambig:3|num_ambig_gaps:0
LYAMRLSDETLLDIMARFRREMKNGLSRDFNPTATVKMLPTFVRSIPDGSEKGDFIALDLGGSYFRILRV
KVSHEKKQTVQMETEIYDTPEDIMHGSGTRLFDHVAECLGDFMEKQQIKDKKLPVGFTFSFPCRQSKLDE

GILITWTKRFKASGVEGADVVKLLNKAIKKRGDYDADIMAVVNDTVGTMMTCGFDDQRCEVGLIIGTGTN
ACYMEEMRHIDLVEGDEGRMCINTEWGAFGDDGSLEDIRTEFDREIDRGSLNPGKQLFEKMVSGMYMGEL
VRLILVKMAKEGLLFEGRITPELLTKGKFETKHVSAIEKSKEGLNKAKEILTRLGVEPSHEDCIAVQHVC
TIVSFRSANLVAATLGAILNQLRDNKGVPRLRTTVGVDGSLYKMHPQYSRRLHKTTRRLVPDSDVRFLLS
ESGSGKGAAMVTAVAYRLSEQHRLIDETLAEFKLTHEQLLQVKKRMRTEIEAGLRKKTHESAKVKMLPTF
VRSTPDGTENGDFLALDLGGTNFRVLLVKIRSGKRRTVEMHNKIYAIPIEVMQGTGEELFDHIVTCISDF
LDYMGIKGARLPLGFTFSFPCKQTSLDAGILLNWTKGFKATDCEGEDVVYLLREGIKRREEFDLDVVAVV
NDTVGTMMTCAYEDPNCEIGLIVGTGSNACYMEEMRNIEMVEGDQGRMCVNMEWGAFGDNGCLDDIRTIY
DKAVDDYSLNAGKQRYEKMISGMYLGEIVRNILIDFTKRGFLFRGQISETLKTRGIFETKFLSQIESDRL
ALLQVRAILQQLGLNGTCDDSIIVKTVCGAVSRRAAQLCGAGMAAVVDKIRENRGLDHLEITVGVDGTLY
KLHPHFSRIMHQTVKDLAPKCDVTFLLSEDGSGKGAALITAVGCRLRDAEQN
>anc18|avgPP:0.998|lnPP:-2.368|num_ambig:2|num_ambig_gaps:0
LYAMRLSDETLLDIMARFRREMKNGLSRDFNPTATVKMLPTFVRSIPDGSEKGDFIALDLGGSYFRILRV
KVSHEKKQTVQMETEIYDTPEDIMHGSGTRLFDHVAECLGDFMEKQQIKDKKLPVGFTFSFPCRQSKLDE
GILITWTKRFKASGVEGADVVKLLNKAIKKRGDYDADIMAVVNDTVGTMMTCGFDDQRCEVGLIIGTGTN
ACYMEEMRHIDLVEGDEGRMCINTEWGAFGDDGSLEDIRTEFDREIDRGSLNPGKQLFEKMVSGMYMGEL
VRLILVKMAKEGLLFEGRITPELLTKGKFETKHVSAIEKSKEGLNKAKEILTRLGVEPSHEDCIAVQHVC
TIVSFRSANLVAATLGAILNQLRDNKGVPRLRTTVGVDGSLYKMHPQYSRRLHKTVRRLVPDSDVRFLLS
ESGSGKGAAMVTAVAYRLAEQHRLIDETLAEFKLTHEQLLQVKKRMRTEIEAGLRKKTHESAKVKMLPTF
VRSTPDGTENGDFLALDLGGTNFRVLLVKIRSGKRRTVEMHNKIYAIPIEVMQGTGEELFDHIVTCISDF
LDYMGIKGARLPLGFTFSFPCKQTSLDAGILLNWTKGFKATDCEGEDVVNLLREGIKRREEFDLDVVAVV
NDTVGTMMTCAYEDPNCEIGLIVGTGSNACYMEEMRNIEMVEGDQGRMCVNMEWGAFGDNGCLDDIRTIY
DKAVDEYSLNAGKQRYEKMISGMYLGEIVRNILIDFTKRGFLFRGQISETLKTRGIFETKFLSQIESDRL
ALLQVRAILQQLGLNGTCDDSIIVKTVCGAVSRRAAQLCGAGMAAVVDKIRENRGLDHLEITVGVDGTLY
KLHPHFSRIMHQTVKDLAPKCDVTFLLSEDGSGKGAALITAVGCRLRDAEQN
>anc18_altAll|avgPP:0.998|lnPP:-3.691|num_ambig:2|num_ambig_gaps:0
LYAMRLSDETLLDIMARFRREMKNGLSRDFNPTATVKMLPTFVRSIPDGSEKGDFIALDLGGSYFRILRV
KVSHEKKQTVQMETEIYDTPEDIMHGSGTRLFDHVAECLGDFMEKQQIKDKKLPVGFTFSFPCRQSKLDE
GILITWTKRFKASGVEGADVVKLLNKAIKKRGDYDADIMAVVNDTVGTMMTCGFDDQRCEVGLIIGTGTN
ACYMEEMRHIDLVEGDEGRMCINTEWGAFGDDGSLEDIRTEFDREIDRGSLNPGKQLFEKMVSGMYMGEL
VRLILVKMAKEGLLFEGRITPELLTKGKFETKHVSAIEKSKEGLNKAKEILTRLGVEPSHEDCIAVQHVC
TIVSFRSANLVAATLGAILNQLRDNKGVPRLRTTVGVDGSLYKMHPQYSRRLHKTVRRLVPDSDVRFLLS
ESGSGKGAAMVTAVAYRLAEQHRLIDETLAEFKLTHEQLLQVKKRMRTEIEAGLRKKTHETAKVKMLPTF
VRSTPDGTENGDFLALDLGGTNFRVLLVKIRSGKRRTVEMHNKIYAIPIEVMQGTGEELFDHIVSCISDF
LDYMGIKGARLPLGFTFSFPCKQTSLDAGILLNWTKGFKATDCEGEDVVNLLREGIKRREEFDLDVVAVV
NDTVGTMMTCAYEDPNCEIGLIVGTGSNACYMEEMRNIEMVEGDQGRMCVNMEWGAFGDNGCLDDIRTIY
DKAVDEYSLNAGKQRYEKMISGMYLGEIVRNILIDFTKRGFLFRGQISETLKTRGIFETKFLSQIESDRL
ALLQVRAILQQLGLNGTCDDSIIVKTVCGAVSRRAAQLCGAGMAAVVDKIRENRGLDHLEITVGVDGTLY
KLHPHFSRIMHQTVKDLAPKCDVTFLLSEDGSGKGAALITAVGCRLRDAEQN
>anc19|avgPP:0.990|lnPP:-15.496|num_ambig:14|num_ambig_gaps:0
LYAMRLSDEMLHEVTNRFKKAMQNGLSRNYNVTASVKMLPTFVRSIPDGSEKGDFIALDLGGSSFRILRV
QVSHEKNQTVQMESEVYETPEGIMHGSGSQLFDHVAECLGDFMEKKKIKDKNLPVGFTFSFPCRQTKIDE
AVLITWTKRFKASGVEGSDVVQLLNKAIKKRGDYDANIVAVVNDTVGTMMTCGFDDQECEVGLIIGTGTN
ACYMEELRHIDMVEGDEGRMCINTEWGAFGDDGCIQEIRTEFDKEIDRGSLNPGKQLFEKMVSGMYMGEL
VRLIIVKMAKGGLLFEGRITPELLTKGKFETKNVSAIEKTKEGLQRAKEILTNLGLEPSPEDCISVQHVC
AIVSHRSANLVAAMLAGILHRLKENKGVPRLRTTVGVDGTLYKFHPQYSRRFHKTLRRLVPDSDVRFILS
ESGSGKGAAMVTAVAYRLAEQHKQIEEMLAPFKLTTEQLMQVKNRMRVEMDLGLQKKTHESAKVKMLPSF
VCGTPDGTENGDFLALDLGGTNFRVLLVKIRSGKKRSVEMHNKIYAIPEEVMQGTGEELFDHIVSCISDF
LDYMGIKGARLPLGFTFSFPCRQTSLDGGILLTWTKGFNASNCEGQDVPTLLREAIKRREEFDLDVVALV
NDTVGTMMTCAYEEPTCEVGLIVGTGSNACYMEEMKNIEMVEGNEGRMCINMEWGAFGDNGCLEDIRTEY
DRIVDDYSLNAGKQRYEKMISGMYLGEIVRNILIDFTKKGFLFRGQISETLKTRGIFKTKFLSQIESDRL
ALLQVRTILQELGLNSTCDDSILVKMVCGVVSKRAAQLCGAGMAAVVDKIRENRGLETLNVTVGVDGTLY

KLHPHFSRIMHQTVQELAPKCNVKFLLSEDGSGKGAALISAVGVRLRDAKAKAS
>anc19_altAll|avgPP:0.987|lnPP:-23.511|num_ambig:14|num_ambig_gaps:0
LYAMRLSDEMLNEVTNRFKKAMQNGLSRNYNVTASVKMLPTFVRSIPDGSEKGDFIALDLGGSSFRILRV
QVSHEKNQTVQMESEVYETPEGIMNGSGSELFDHVAECLGDFMEKKNIKDKNLPVGFTFSFPCRQTKIDE
AVLITWTKRFKASGVEGSDVVQLLNKAIKKRGDYDANIVAVVNDTVGTMMTCGFDDQECEVGLIIGTGTN
ACYMEELRHIDMVEGDEGRMCINTEWGAFGDDGCIQEIRTEFDKEIDRGSLNPGKQLFEKMVSGMYMGEL
VRLIIVKMAKGGLLFEGRITPELLTKGKFETKNVSAIEKTKEGLQRAKEILTNLGLEPSPEDCISVQHVC
AIVSHRSANLVAAMLAGILHRLKENKGVPRLRTTVGVDGTLYKFHPQYSRRFHKTLRRLVPDSDVRFILS
ESGSGKGAAMVTAVAYRLAEQHKQIEEMLAPFKLTTEQLMQVKNRMRVEMDLGLKKKTHDSAKVKMLPSF
VRGTPDGTENGDFLALDLGGTNFRVLLVKIRSGKKRSVEMHNKIYAIPEEVMQGTGEELFDHIVSCISDF
LDYMGIKGARLPLGFTFSFPCRQTSLDGGILLTWTKGFNASNCEGQDVPTLLRDAIKRREEFDLDVVALV
NDTVGTMMTCAYEEPTCEVGLIVGTGSNACYMEEMKNIEMVEGDEGRMCINMEWGAFGDNGCLEDIRTEY
DRIVDDYSLNAGKQRYEKMISGMYLGEIVRNILIDFTKKGFLFRGQISETLKTRGIFKTKFLSQIESDRL
ALLQVRAILQELGLNSTCDDSILVKMVCGIVSKRAAQLCGAGMAAVVDKIRENRGLETLNVTVGVDGTLY
KLHPHFSRIMHQTVKDLAPKCNVKFLLSEDGSGKGAALISAVGVRLRDGKAKAS
>anc20|avgPP:0.995|lnPP:-7.697|num_ambig:8|num_ambig_gaps:1
LYAMRLSDETLIDIMTRFKKEMKNGLSRDFNPTATVKMLPTFVRSIPDGSEKGDFIALDLGGSSFRILRV
QVNHEKNQNVHMESEVYDTPENIVHGSGSQLFDHVAECLGDFMEKRKIKDKKLPVGFTFSFPCRQSKIDE
AILITWTKRFKASGVEGADVVKLLNKAIKKRGDYDANIVAVVNDTVGTMMTCGYDDQHCEVGLIIGTGTN
ACYMEELRHIDLVEGDEGRMCINTEWGAFGDDGSLEDIRTEFDREIDRGSLNPGKQLFEKMVSGMYMGEL
VRLILVKMAKEGLLFEGRITPELLTRGKFNTSDVSAIEKNKEGLHNAKEILTRLGVEPSDDDCVSVQHVC
TIVSFRSANLVAATLGAILNRLRDNKGTPRLRTTVGVDGSLYKTHPQYSRRFHKTLRRLVPDSDVRFLLS
ESGSGKGAAMVTAVAYRLAEQHRQIEETLAHFRLTKEMLLEVKKRMRAEMELGLKKQTHNKAVVKMLPSF
VRSTPDGTENGDFLALDLGGTNFRVLLVKIRSGKKRTVEMHNKIYAIPIEIMQGTGEELFDHIVSCISDF
LDYMGIKGPRMPLGFTFSFPCKQTSLDAGILITWTKGFKATDCVGHDVVTLLRDAIKRREEFDLDVVAVV
NDTVGTMMTCAYEEPTCEVGLIVGTGSNACYMEEMKNVEMVEGNQGQMCINMEWGAFGDNGCLDDIRTDY
DRLVDEYSLNAGKQRYEKMISGMYLGEIVRNILIDFTKKGFLFRGQISEPLKTRGIFETKFLSQIESDRL
ALLQVRAILQQLGLNSTCDDSILVKTVCGVVSKRAAQLCGAGMAAVVDKIRENRGLDHLNVTVGVDGTLY
KLHPHFSRIMHQTVKELSPKCNVSFLLSEDGSGKGAALITAVGVRLRNEASSS
>anc20_altAll|avgPP:0.994|lnPP:-12.022|num_ambig:8|num_ambig_gaps:1
LYAMRLSDETLIDIMTRFRKEMKNGLSRDFNPTATVKMLPTFVRSIPDGSEKGDFIALDLGGSSFRILRV
QVNHEKNQNVHMESEVYDTPENIVHGSGSQLFDHVAECLGDFMEKRKIKDKKLPVGFTFSFPCRQSKIDE
AILITWTKRFKASGVEGADVVKLLNKAIKKRGDYDANIVAVVNDTVGTMMTCGYDDQHCEVGLIIGTGTN
ACYMEELRHIDLVEGDEGRMCINTEWGAFGDDGALEDIRTEFDREIDRGSLNPGKQLFEKMVSGMYMGEL
VRLILVKMAKEGLLFEGRITPELLTRGKFNTSDVSAIEKNKEGLHNAKEILTRLGVEPSDDDCVSVQHVC
TIVSFRSANLVAATLGAILNRLRDNKGTPRLRTTVGVDGSLYKTHPQYSRRFHKTLRRLVPDSDVRFLLS
ESGSGKGAAMVTAVAYRLAEQHRQIEETLAHFRLTKDMLLEVKKRMRTEMELGLKKQTHNKAVVKMLPSF
VRSTPDGTENGDFLALDLGGTNFRVLLVKIRSGKKRTVEMHNKIYAIPIEIMQGTGEELFDHIVSCISDF
LDYMGIKGPRMPLGFTFSFPCKQTSLDAGILITWTKGFKATDCVGHDVVTLLRDAVKRREEFDLDVVAVV
NDTVGTMMTCAYEEPTCEVGLIVGTGSNACYMEEMKNVEMVEGNQGQMCINMEWGAFGDNGCLDDIRTNY
DRLVDEYSLNAGKQRYEKMISGMYLGEIVRNILIDFTKKGFLFRGQISETLKTRGIFETKFLSQIESDRL
ALLQVRAILQQLGLNSTCDDSILVKTVCGVVSKRAAQLCGAGMAAVVDKIRENRGLDHLNVTVGVDGTLY
KLHPHFSRIMHQTVKELSPKCNVSFLLSEDGSGKGAALITAVGVRLRNEASSA
>anc21|avgPP:0.996|lnPP:-5.754|num_ambig:6|num_ambig_gaps:0
LYAMRLSDETLIDIMTRFKKEMKNGLSRDFNPTATVKMLPTFVRSIPDGSEKGDFIALDLGGSSFRILRV
QVNHEKNQNVHMESEVYDTPENIMHGSGSQLFDHVAECLGDFMEKKKIKDKKLPVGFTFSFPCRQSKIDE
AILITWTKRFKASGVEGADVVKLLNKAIKKRGDYDANIVAVVNDTVGTMMTCGYDDQHCEVGLIIGTGTN
ACYMEELRHIDLVEGDEGRMCINTEWGAFGDDGALEDIRTEFDREIDRGSLNPGKQLFEKMVSGMYMGEL
VRLILVKMAKEGLLFEGRITPELLTRGKFNTSDVSAIEKNKEGLHNAKEILTRLGVEPSDDDCISVQHVC
TIVSFRSANLVAATLGAILNRLRDNKGTPRLRTTVGVDGSLYKTHPQYSRRFHKTLRRLVPDCDVRFLLS
ESGSGKGAAMVTAVAYRLAEQHRQIEETLAHFRLTKEMLLEVKKRMRAEMELGLKKQTHDKAVVKMLPSF
VRSTPDGTENGDFLALDLGGTNFRVLLVKIRSGKKRTVEMHNKIYAIPIEIMQGTGEELFDHIVSCISDF

LDYMGIKGPRMPLGFTFSFPCKQTSLDAGILITWTKGFKATDCVGHDVATLLRDAVKRREEFDLDVVAVV
NDTVGTMMTCAYEEPTCEVGLIVGTGSNACYMEEMKNVEMLEGNEGQMCINMEWGAFGDNGCLDDIRTDY
DKLVDEYSLNAGKQRYEKMISGMYLGEIVRNILIDFTKKGFLFRGQISEPLKTRGIFQTKYLSQIESDRL
ALLQVRAILQQLGLNSTCDDSILVKTVCGVVSKRAAQLCGAGMAAVVDKIRENRGLDHLNVTVGVDGTLY
KLHPHFSRIMHQTVKELSPKCNVSFLLSEDGSGKGAALITAVGVRLRNEASS
>anc21_altAll|avgPP:0.995|lnPP:-8.873|num_ambig:6|num_ambig_gaps:0
LYAMRLSDETLLDIMARFKKEMKNGLSRDFNPTATVKMLPTFVRSIPDGSEKGDFIALDLGGSSFRILRV
QVNHEKNQNVHMESEVYDTPENIMHGSGSQLFDHVAECLGDFMEKKKIKDKKLPVGFTFSFPCRQSKIDE
AILITWTKRFKASGVEGADVVKLLNKAIKKRGDYDANIVAVVNDTVGTMMTCGYDDQHCEVGLIIGTGTN
ACYMEELRHIDLVEGDEGRMCINTEWGAFGDDGALEDIRTEFDREIDRGSLNPGKQLFEKMVSGMYMGEL
VRLILVKMAKEGLLFEGRITPELLTRGKFNTSDVSAIEKNKEGLHNAKEILTRLGVEPSDDDCVSVQHVC
TIVSFRSANLVAATLGAILNRLRDNKGTPRLRTTVGVDGSLYKTHPQYSRRFHKTLRRLVPDCDVRFLLS
ESGSGKGAAMVTAVAYRLAEQHRQIEETLAHFRLTKDMLLEVKKRMRTEMELGLKKQTHDKAVVKMLPSF
VRSTPDGTENGDFLALDLGGTNFRVLLVKIRSGKKRTVEMHNKIYAIPIEIMQGTGEELFDHIVSCISDF
LDYMGIKGPRMPLGFTFSFPCKQTSLDAGILITWTKGFKATDCVGHDVATLLRDAVKRREEFDLDVVAVV
NDTVGTMMTCAYEEPTCEVGLIVGTGSNACYMEEMKNVEMLEGNEGQMCINMEWGAFGDNGCLDDIRTNY
DKLVDEYSLNAGKQRYEKMISGMYLGEIVRNILIDFTKKGFLFRGQISEPLKTRGIFQTKYLSQIESDRL
ALLQVRAILQQLGLNSTCDDSILVKTVCGVVSKRAAQLCGAGMAAVVDKIRENRGLDHLNVTVGVDGTLY
KLHPHFSRIMHQTVKELSPKCNVSFLLSEDGSGKGAALITAVGVRLRNEASS
>anc22|avgPP:0.994|lnPP:-8.866|num_ambig:11|num_ambig_gaps:1
LYAMRLSDETLIDIMTRFKKEMKNGLSRDFNPTATVKMLPTFVRSIPDGSEKGDFIALDLGGSSFRILRV
QVNHEKNQNVHMESEVYDTPENIMHGSGSQLFDHVAECLGDFMEKKKIKDKKLPVGFTFSFPCRQSKIDE
AILITWTKRFKASGVEGADVVKLLNKAIKKRGDYDANIVAVVNDTVGTMMTCGYDDQHCEVGLIIGTGTN
ACYMEELRHIDLVEGDEGRMCINTEWGAFGDDGSLEDIRTEFDREIDRGSLNPGKQLFEKMVSGMYMGEL
VRLILVKMAKEGLLFEGRITPELLTRGKFNTSDVSAIEKNKEGLHNAKEILTRLGVEPSDDDCISVQHVC
TIVSFRSANLVAATLGAILNRLRDNKGTPRLRTTVGVDGSLYKTHPQYSRRFHKTLRRLVPDSDVRFLLS
ESGSGKGAAMVTAVAYRLAEQHRQIEETLAHFRLTKEMLLEVKKRMRAEMELGLKKQTHDKAVVKMLPSF
VRSTPDGTENGDFLALDLGGTNFRVLLVKIRSGKKRTVEMHNKIYAIPIEIMQGTGEELFDHIVSCISDF
LDYMGIKGPRMPLGFTFSFPCKQTSLDAGILITWTKGFKATDCVGHDVATLLRDAIKRREEFDLDVVAVV
NDTVGTMMTCAYEEPTCEVGLIVGTGSNACYMEEMKNVEMLEGNEGQMCINMEWGAFGDNGCLDDIRTDY
DKLVDEYSLNAGKQRYEKMISGMYLGEIVRNILIDFTKKGFLFRGQISEPLKTRGIFETKYLSQIESDRL
ALLQVRAILQQLGLNSTCDDSILVKTVCGVVSKRAAQLCGAGMAAVVDKIRENRGLDHLNVTVGVDGTLY
KLHPHFSRIMHQTVKELSPKCNVSFLLSEDGSGKGAALITAVGVRLRNEASSS
>anc22_altAll|avgPP:0.992|lnPP:-13.916|num_ambig:11|num_ambig_gaps:1
LYAMRLSDETLLDIMARFRKEMKNGLSRDFNPTATVKMLPTFVRSIPDGSEKGDFIALDLGGSSFRILRV
QVNHEKNQNVHMESEVYDTPENIMHGSGSQLFDHVAECLGDFMEKKKIKDKKLPVGFTFSFPCRQSKIDE
AILITWTKRFKASGVEGADVVKLLNKAIKKRGDYDANIVAVVNDTVGTMMTCGYDDQHCEVGLIIGTGTN
ACYMEELRHIDLVEGDEGRMCINTEWGAFGDDGALEDIRTEFDREIDRGSLNPGKQLFEKMVSGMYMGEL
VRLILVKMAKEGLLFEGRITPELLTRGKFNTSDVSAIEKNKEGLHNAKEILTRLGVEPSDDDCVSVQHVC
TIVSFRSANLVAATLGAILNRLRDNKGTPRLRTTVGVDGSLYKTHPQYSRRFHKTLRRLVPDSDVRFLLS
ESGSGKGAAMVTAVAYRLAEQHRQIEETLAHFRLTKEMLLEVKKRMRTEMELGLKKQTHDKAVVKMLPSF
VRSTPDGTENGDFLALDLGGTNFRVLLVKIRSGKKRTVEMHNKIYAIPIEIMQGTGEELFDHIVSCISDF
LDYMGIKGPRMPLGFTFSFPCKQTSLDAGILITWTKGFKATDCVGHDVATLLRDAVKRREEFDLDVVAVV
NDTVGTMMTCAYEEPTCEVGLIVGTGSNACYMEEMKNVEMVEGNEGQMCINMEWGAFGDNGCLDDIRTNY
DKLVDEYSLNAGKQRYEKMISGMYLGEIVRNILIDFTKKGFLFRGQISETLKTRGIFETKYLSQIESDRL
ALLQVRAILQQLGLNSTCDDSILVKTVCGVVSKRAAQLCGAGMAAVVDKIRENRGLDHLNVTVGVDGTLY
KLHPHFSRIMHQTVKELSPKCNVSFLLSEDGSGKGAALITAVGVRLREEASSS
>anc23|avgPP:0.993|lnPP:-10.498|num_ambig:13|num_ambig_gaps:1
LYAMRLSDETLLDIMARFRKEMKNGLSRDFNPTATVKMLPTFVRSIPDGSEKGDFIALDLGGSSFRILRV
QVNHEKNQNVHMESEVYDTPENIMHGSGSQLFDHVAECLGDFMEKKKIKDKKLPVGFTFSFPCRQSKIDE
AVLITWTKRFKASGVEGADVVKLLNKAIKKRGDYDANIVAVVNDTVGTMMTCGYDDQHCEVGLIIGTGTN
ACYMEELRHIDLVEGDEGRMCINTEWGAFGDDGSLEDIRTEFDREIDRGSLNPGKQLFEKMVSGMYMGEL

VRLILVKMAKEGLLFEGRITPELLTKGKFNTSDVSAIEKNKEGLHNAKEILTRLGVEPSHDDCISVQHVC
TIVSFRSANLVAATLGAILNRLRDNKGTPRLRTTVGVDGSLYKTHPQYSRRFHKTLRRLVPDSDVRFLLS
ESGSGKGAAMVTAVAYRLAEQHRQIEETLAHFRLTKEMLLEVKKRMRAEMELGLKKQTHDSATVKMLPSF
VRSTPDGTENGDFLALDLGGTNFRVLLVKIRSGKKRTVEMHNKIYAIPIEIMQGTGEELFDHIVSCISDF
LDYMGIKGPRMPLGFTFSFPCKQTSLDAGILITWTKGFKATDCVGHDVATLLRDAIKRREEFDLDVVAVV
NDTVGTMMTCAYEEPTCEVGLIVGTGSNACYMEEMKNVEMLEGNEGQMCINMEWGAFGDNGCLDDIRTNY
DKLVDEYSLNAGKQRYEKMISGMYLGEIVRNILIDFTKKGFLFRGQISETLKTRGIFETKYLSQIESDRL
ALLQVRAILQQLGLNSTCDDSILVKTVCGVVSKRAAQLCGAGMAAVVDKIRENRGLDHLNVTVGVDGTLY
KLHPHFSRIMHQTVKELAPKCNVSFLLSEDGSGKGAALITAVGVRLRDEASSS
>anc23_altAll|avgPP:0.991|lnPP:-16.564|num_ambig:13|num_ambig_gaps:1
LYAMRLSDETLLDIMTRFKKEMKNGLSRDFNPTATVKMLPTFVRSIPDGSEKGDFIALDLGGSSFRILRV
QVNHEKNQNVHMESEVYDTPENIMHGSGSQLFDHVAECLGDFMEKKKIKDKKLPVGFTFSFPCRQSKIDE
AILITWTKRFKASGVEGADVVKLLNKAIKKRGDYDANIVAVVNDTVGTMMTCGYDDQHCEVGLIIGTGTN
ACYMEELRHIDLVEGDEGRMCINTEWGAFGDDGSLEDIRTEFDREIDRGSLNPGKQLFEKMVSGMYMGEL
VRLILVKMAKEGLLFEGRITPELLTKGKFNTSDVSAIEKNKEGLHNAKEILTRLGVEPSHDDCISVQHVC
TIVSFRSANLVAATLGAILNRLRDNKGTPRLRTTVGVDGSLYKTHPQYSRRFHKTLRRLVPDSDVRFLLS
ESGSGKGAAMVTAVAYRLAEQHRQIEETLAHFRLTKEMLMEVKKRMRTEMELGLKKKTHDAATVKMLPSF
VRSTPDGTENGDFLALDLGGTNFRVLLVKIRSGKKRTVEMHNKIYAIPIEIMQGTGEELFDHIVSCISDF
LDYMGIKGPRMPLGFTFSFPCKQTSLDAGILITWTKGFKATDCVGHDVATLLRDAIKRREEFDLDVVAVV
NDTVGTMMTCAYEEPTCEVGLIVGTGSNACYMEEMKNVEMVEGNEGQMCINMEWGAFGDNGCLDDIRTDY
DKLVDEYSLNAGKQRYEKMISGMYLGEIVRNILIDFTKKGFLFRGQISETLKTRGIFETKYLSQIESDRL
ALLQVRAILQQLGLNSTCDDSILVKTVCGVVSKRAAQLCGAGMAAVVDKIRENRGLDHLNVTVGVDGTLY
KLHPHFSRIMHQTVKELAPKCNVSFLLSEDGSGKGAALITAVGVRLREESPSS
>anc24|avgPP:0.986|lnPP:-22.081|num_ambig:20|num_ambig_gaps:1
LYAMRLSDETLLDIMARFRKEMKNGLSRDFNPTATVKMLPTFVRSIPDGSEKGDFIALDLGGSSFRILRV
QVSHEKNQTVQMESEVYDTPEDIMHGSGSQLFDHVAECLGDFMEKKKIKDKKLPVGFTFSFPCRQSKIDE
AVLITWTKRFKASGVEGADVVKLLNKAIKKRGDYDANIVAVVNDTVGTMMTCGFDDQRCEVGLIIGTGTN
ACYMEELRHIDLVEGDEGRMCINTEWGAFGDDGSLEDIRTEFDREIDRGSLNPGKQLFEKMVSGMYMGEL
VRLILVKMAKEGLLFEGRITPELLTKGKFETKNVSAIEKSKEGLHKAKEILTRLGVEPSHEDCISVQHVC
TIVSFRSANLVAATLGAILNRLRDNKGVPRLRTTVGVDGSLYKMHPQYSRRFHKTLRRLVPDSDVRFLLS
ESGSGKGAAMVTAVAYRLAEQHRQIEETLAHFKLTKEQLLQVKKRMRVEMELGLKKKTHESAKVKMLPSF
VRSTPDGTENGDFLALDLGGTNFRVLLVKIRSGKKRTVEMHNKIYAIPIEVMQGTGEELFDHIVSCISDF
LDYMGIKGARLPLGFTFSFPCKQTSLDAGILLTWTKGFKATDCEGQDVATLLREAIKRREEFDLDVVAVV
NDTVGTMMTCAYEEPTCEVGLIVGTGSNACYMEEMKNIEMVEGNEGRMCINMEWGAFGDNGCLDDIRTDY
DKLVDEYSLNAGKQRYEKMISGMYLGEIVRNILIDFTKKGFLFRGQISETLKTRGIFETKFLSQIESDRL
ALLQVRAILQQLGLNSTCDDSILVKTVCGVVSKRAAQLCGAGMAAVVDKIRENRGLDHLNVTVGVDGTLY
KLHPHFSRIMHQTVKELAPKCNVTFLLSEDGSGKGAALITAVGVRLRDAGSSN
>anc24_altAll|avgPP:0.983|lnPP:-32.222|num_ambig:20|num_ambig_gaps:1
LYAMRLSDETLLDVMARFKKEMKNGLSRDFNPTATVKMLPTFVRSIPDGSEKGDFIALDLGGSSFRILRV
QVSHEKNQTVQMESEVYDTPENIMHGSGSQLFDHVAECLGDFMEKKKIKDKKLPVGFTFSFPCRQSKIDE
AVLITWTKRFKASGVEGADVVKLLNKAIKKRGDYDANIVAVVNDTVGTMMTCGFDDQHCEVGLIIGTGTN
ACYMEELRHIDLVEGDEGRMCINTEWGAFGDDGSLEDIRTEFDREIDRGSLNPGKQLFEKMVSGMYMGEL
VRLILVKMAKEGLLFEGRITPELLTKGKFETKDVSAIEKNKEGLNKAKEILTRLGVEPSHEDCISVQHVC
AIVSFRSANLVAATLGAILNRLRDNKGVPRLRTTVGVDGSLYKMHPQYSRRFHKTLRRLVPDSDVRFILS
ESGSGKGAAMVTAVAYRLAEQHRQIEETLAHFKLTKEQLMQVKKRMRTEMELGLQKKTHDSAKVKMLPSF
VRSTPDGTENGDFLALDLGGTNFRVLLVKIRSGKKRTVEMHNKIYAIPIEVMQGTGEELFDHIVSCISDF
LDYMGIKGARLPLGFTFSFPCKQTSLDAGILLTWTKGFKATDCEGQDVATLLRDAIKRREEFDLDVVAVV
NDTVGTMMTCAYEEPTCEVGLIVGTGSNACYMEEMKNIEMVEGDEGRMCINMEWGAFGDNGCLDDIRTNY
DKLVDEYSLNAGKQRYEKMISGMYLGEIVRNILIDFTKKGFLFRGQISETLKTRGIFETKFLSQIESDRL
ALLQVRAILQQLGLNSTCDDSILVKTVCGVVSKRAAQLCGAGMAAVVDKIRENRGLDHLNVTVGVDGTLY
KLHPHFSRIMHQTVKDLAPKCNVSFLLSEDGSGKGAALITAVGVRLRDGGSSN
>anc25|avgPP:0.994|lnPP:-9.102|num_ambig:11|num_ambig_gaps:0

LYAMRLSDETLLDIMARFRKEMKNGLSRDFNPTATVKMLPTFVRSIPDGSEKGDFIALDLGGSCFRILRV
KVSHEKKQTVQMETEIYDTPEDIMHGSGSQLFDHVAECLGDFMEKKQIKDKKLPVGFTFSFPCRQSKIDE
GVLITWTKRFKASGVEGADVVKLLNKAIKKRGDYDADIVAVVNDTVGTMMTCGFDDQRCEVGLIIGTGTN
ACYMEELRHIDLVEGDEGRMCINTEWGAFGDDGSLEDIRTEFDREIDRGSLNPGKQLFEKMVSGMYMGEL
VRLILVKMAKEGLLFEGRITPELLTKGKFETKHVSAIEKSKEGLNKAKEILTRLGVEPSHEDCIAVQHVC
TIVSFRSANLVAATLGAILNRLRDNKGVPRLRTTVGVDGSLYKMHPQYSRRFHKTLRRLVPDSDVRFLLS
ESGSGKGAAMVTAVAYRLAEQHRLIEETLAEFKLTHEQLLQVKKRMRVEIEAGLQKKTHESAKVKMLPTF
VRSTPDGTENGDFLALDLGGTNFRVLLVKIRSGKRRTVEMHNKIYAIPIEVMQGTGEELFDHIVHCISDF
LDYMGIKGARLPLGFTFSFPCKQTSLDAGILLNWTKGFKATDCEGQDVANLLREGIKRREEFDLDVVAVV
NDTVGTMMTCAYEEPTCEVGLIVGTGSNACYMEEMKNIEMVEGNEGRMCVNMEWGAFGDNGCLDDIRTIY
DKAVDEYSLNAGKQRYEKMISGMYLGEIVRNILIDFTKRGFLFRGQISETLKTRGIFETKFLSQIESDRL
ALLQVRAILQQLGLNGTCDDSILVKTVCGAVSRRAAQLCGAGMAAVVDKIRENRGLDHLEVTVGVDGTLY
KLHPHFSRIMHQTVKDLAPKCNVTFLLSEDGSGKGAALITAVGVRLRDAGQS
>anc25_altAll|avgPP:0.992|lnPP:-14.348|num_ambig:11|num_ambig_gaps:0
LYAMRLSDETLLDVMARFRKEMKNGLSRDFNPTATVKMLPTFVRSIPDGSEKGDFIALDLGGSCFRILRV
KVSHEKKQTVQMETEIYDTPEDIMHGSGSQLFDHVAECLGDFMEKKQIKDKKLPVGFTFSFPCRQSKIDE
GVLITWTKRFKASGVEGADVVKLLNKAIKKRGDYDADIVAVVNDTVGTMMTCGFDDQRCEVGLIIGTGTN
ACYMEELRHIDLVEGDEGRMCINTEWGAFGDDGSLEDIRTEFDREIDRGSLNPGKQLFEKMVSGMYMGEL
VRLILVKMAKEGLLFEGRITPELLTKGKFETKHVSAIEKSKEGLNKAKEILTRLGVEPSHEDCIAVQHVC
AIVSFRSANLVAATLGAILTRLRDNKGVPRLRTTVGVDGSLYKMHPQYSRRFHKTLRRLVPDSDVRFILS
ESGSGKGAAMVTAVAYRLAEQHRQIEETLAEFKLTHEQLLQVKKRMRTEIEAGLRKKTHESAKVKMLPTF
VRSTPDGTENGDFLALDLGGTNFRVLLVKIRSGKRRTVEMHNKIYAIPIEVMQGTGEELFDHIVHCISDF
LDYMGIKGARLPLGFTFSFPCKQTSLDAGILLNWTKGFKATDCEGQDVANLLREGIKRREEFDLDVVAVV
NDTVGTMMTCAYEEPTCEVGLIVGTGSNACYMEEMKNIEMVEGDEGRMCVNMEWGAFGDNGCLDDIRTIY
DKAVDEYSLNAGKQRYEKMISGMYLGEIVRNILIDFTKRGFLFRGQISETLKTRGIFETKFLSQIESDRL
ALLQVRAILQQLGLNGTCDDSILVKTVCGTVSRRAAQLCGAGMAAVVDKIRENRGLDHLEVTVGVDGTLY
KLHPHFSRIMHQTVKDLAPKCNVTFLLSEDGSGKGAALITAVGVRLRDGEQS
>anc26|avgPP:0.997|lnPP:-4.389|num_ambig:4|num_ambig_gaps:0
LYAMRLSDETLLDIMARFRREMKNGLSRDFNPTATVKMLPTFVRSIPDGSEKGDFIALDLGGSYFRILRV
KVSHEKKQTVQMETEIYDTPEDIMHGSGTRLFDHVAECLGDFMEKQQIKDKKLPVGFTFSFPCRQSKLDE
GILITWTKRFKASGVEGADVVKLLNKAIKKRGDYDADIMAVVNDTVGTMMTCGFDDQRCEVGLIIGTGTN
ACYMEELRHIDLVEGDEGRMCINTEWGAFGDDGSLEDIRTEFDREIDRGSLNPGKQLFEKMVSGMYMGEL
VRLILVKMAKEGLLFEGRITPELLTKGKFETKHVSAIEKSKEGLNKAKEILTRLGVEPSHEDCIAVQHVC
TIVSFRSANLVAATLGAILNRLRDNKGVPRLRTTVGVDGSLYKMHPQYSRRLHKTVRRLVPDSDVRFLLS
ESGSGKGAAMVTAVAYRLAEQHRLIDETLAEFKLTHEQLLQVKKRMRTEIEAGLRKKTHESAKVKMLPTF
VRSTPDGTENGDFLALDLGGTNFRVLLVKIRSGKRRTVEMHNKIYAIPIEVMQGTGEELFDHIVHCISDF
LDYMGIKGARLPLGFTFSFPCKQTSLDAGILLNWTKGFKATDCEGEDVVNLLREGIKRREEFDLDVVAVV
NDTVGTMMTCAYEDPNCEIGLIVGTGSNACYMEEMRNIEMVEGDEGRMCVNMEWGAFGDNGCLDDIRTIY
DKAVDEYSLNAGKQRYEKMISGMYLGEIVRNILIDFTKRGFLFRGQISETLKTRGIFETKFLSQIESDRL
ALLQVRAILQQLGLNGTCDDSIIVKTVCGAVSRRAAQLCGAGMAAVVDKIRENRGLDHLEITVGVDGTLY
KLHPHFSRIMHQTVKDLAPKCDVTFLLSEDGSGKGAALITAVGCRLRDAEQS
>anc26_altAll|avgPP:0.996|lnPP:-6.158|num_ambig:4|num_ambig_gaps:0
LYAMRLSDETLLDIMARFRREMKNGLSRDFNPTATVKMLPTFVRSIPDGSEKGDFIALDLGGSYFRILRV
KVSHEKKQTVQMETEIYDTPEDIMHGSGTRLFDHVAECLGDFMEKQQIKDKKLPVGFTFSFPCRQSKLDE
GVLITWTKRFKASGVEGADVVKLLNKAIKKRGDYDADIMAVVNDTVGTMMTCGFDDQRCEVGLIIGTGTN
ACYMEELRHIDLVEGDEGRMCINTEWGAFGDDGSLEDIRTEFDREIDRGSLNPGKQLFEKMVSGMYMGEL
VRLILVKMAKEGLLFEGRITPELLTKGKFETKHVSAIEKSKEGLNKAKEILTRLGVEPSHEDCIAVQHVC
TIVSFRSANLVAATLGAILTRLRDNKGVPRLRTTVGVDGSLYKMHPQYSRRLHKTVRRLVPDSDVRFLLS
ESGSGKGAAMVTAVAYRLAEQHRQIDETLAEFKLTHEQLLQVKKRMRTEIEAGLRKKTHESAKVKMLPTF
VRSTPDGTENGDFLALDLGGTNFRVLLVKIRSGKRRTVEMHNKIYAIPIEVMQGTGEELFDHIVHCISDF
LDYMGIKGARLPLGFTFSFPCKQTSLDAGILLNWTKGFKATDCEGEDVVNLLREGIKRREEFDLDVVAVV
NDTVGTMMTCAYEDPNCEIGLIVGTGSNACYMEEMRNIEMVEGDEGRMCVNMEWGAFGDNGCLDDIRTIY

DKAVDEYSLNAGKQRYEKMISGMYLGEIVRNILIDFTKRGFLFRGQISETLKTRGIFETKFLSQIESDRL
ALLQVRAILQQLGLNGTCDDSIIVKTVCGAVSRRAAQLCGAGMAAVVDKIRENRGLDHLEITVGVDGTLY
KLHPHFSRIMHQTVKDLAPKCDVTFLLSEDGSGKGAALITAVGCRLRDAEQN
>anc27|avgPP:0.996|lnPP:-6.125|num_ambig:6|num_ambig_gaps:0
LYSMRLSDETLLDIMARFRREMENGLSRDFNPTATVKMLPTFVRSIPDGSEKGDFIALDLGGSYFRILRV
KVSHEKKQTVQMESEIYDTPEDIMHGSGTRLFDHVAECLGDFMEKQQIKDKKLPVGFTFSFPCRQSKLDE
GILITWTKRFKASGVEGADVVKLLNKAIKKRGDYDADIMAVVNDTVGTMMTCGFDDQRCEVGIIIGTGTN
ACYMEELRHIDLVEGDEGRMCINTEWGAFGDDGSLEDIRTEFDREIDRGSLNPGKQLFEKMVSGMYMGEL
VRLILVKMAKEGLLFEGRITPELLTKGKFETKHVSAIEKSKEGLNKAKEILTRLGVEPSHEDCIAVQHVC
TIVSFRSANLIAATLGAILTRLRDNKGVPRLRTTVGVDGSLYKMHPQYARRLHKTVRRLVPDSDVRFLLS
ESGSGKGAAMVTAVAYRLAEQHRQIDETLAEFKLSHEQLLEVKKRMRTEIEAGLRKKTHESAKVKMLPTF
VRSTPDGTENGDFLALDLGGTNFRVLLVKIRSGKRRTVEMHNKIYAIPIEVMQGTGEELFDHIVHCISDF
LDYMGIKGARLPLGFTFSFPCKQTSLDAGILLNWTKGFKATDCEGEDVVNLLREGIKRREEFELDVVAVV
NDTVGTMMTCAYEDPNCEIGLIVGTGSNACYMEEMRNIEMVEGDEGRMCVNMEWGAFGDNGCLDDIRTIY
DKAVDELSLNSGKQRYEKMISGMYLGEIVRNILIDFTKRGFLFRGQISETLKTRGIFETKFLSQIESDRL
ALLQVRAILQQLGLDGTCDDSIIVKEVCGAVSRRAAQLCGAGMAAVVDKIRENRGLDHLEITVGVDGTLY
KLHPHFSRIMHQTVKELAPKCDVTFLLSEDGSGKGAALITAVGCRLREAEQS
>anc27_altAll|avgPP:0.995|lnPP:-9.038|num_ambig:6|num_ambig_gaps:0
LYAMRLSDETLLDIMARFRREMENGLSRDFNPTATVKMLPTFVRSIPDGSEKGDFIALDLGGSYFRILRV
KVSHEKKQTVQMESEIYDTPEDIMHGSGTRLFDHVAECLGDFMEKQQIKDKKLPVGFTFSFPCRQSKLDE
GVLITWTKRFKASGVEGADVVKLLNKAIKKRGDYDADIMAVVNDTVGTMMTCGFDDQRCEVGIIIGTGTN
ACYMEELRHIDLVEGDEGRMCINTEWGAFGDDGSLEDIRTEFDREIDRGSLNPGKQLFEKMVSGMYMGEL
VRLILVKMAKEGLLFEGRITPELLTKGKFETKHVSAIEKSKEGLNKAKEILTRLGVEPSHEDCIAVQHVC
TIVSFRSANLIAATLGAILTRLRDNKGVPRLRTTVGVDGSLYKMHPQYARRLHKTVRRLVPDSDVRFLLS
ESGSGKGAAMVTAVAYRLAEQHRQIDETLAEFKLSHEQLLEVKKRMRTEIEAGLRKKTHESAKVKMLPTF
VRSTPDGTENGDFLALDLGGTNFRVLLVKIRSGKRRTVEMHNKIYAIPIEVMQGTGEELFDHIVHCISDF
LDYMGIKGARLPLGFTFSFPCKQTSLDAGILLNWTKGFKATDCEGEDVVNLLREGIKRREEFELDVVAVV
NDTVGTMMTCAYEDPNCEIGLIVGTGSNACYMEEMRNIEMVEGDEGRMCVNMEWGAFGDNGCLDDIRTIY
DKAVDELSLNSGKQRYEKMISGMYLGEIVRNILIDFTKRGFLFRGQISEALKTRGIFETKFLSQIESDRL
ALLQVRAILQQLGLDGTCDDSIIVKEVCGAVSRRAAQLCGAGMAAVVDKIRENRGLDHLDITVGVDGTLY
KLHPHFSRIMHQTVKDLAPKCDVTFLLSEDGSGKGAALITAVGCRLREAEQN
>anc28|avgPP:0.994|lnPP:-8.540|num_ambig:8|num_ambig_gaps:0
LYSMRLSDETLLDIMARFRREMENGLSRDFNPTATVKMLPTFVRSIPDGSEKGDFIALDLGGSYFRILRV
KVSHEKKQTVQMESQIYDTPEDIMHGSGTRLFDHVAECLGDFMEKQQIKDKKLPVGFTFSFPCRQTKLDE
GILITWTKRFKASGVEGADVVKLLNKAIKKRGDYDADIMAVVNDTVGTMMTCGFDDQRCEVGIIIGTGTN
ACYMEELRHIDLVEGDEGRMCINTEWGAFGDDGSLEDIRTEFDREIDRGSLNPGKQLFEKMVSGMYMGEL
VRLILVKMAKEGLLFEGRITPELLTKGKFETKHVSAIEKSKEGLNKAKEILTRLGVEPSHEDCIAVQHVC
TIVSFRSANLIAATLGAILTRLRDNKGVPRLRTTVGVDGSLYKMHPQYARRLHKTVRRLVPDCDVRFLLS
ESGSGKGAAMVTAVAYRLAEQHRQIDETLAEFKLSHEQLLEVKKRMRTEIEAGLRKKTHESATVKMLPTF
VRSTPDGTENGDFLALDLGGTNFRVLLVKIRSGKRRTVEMHNKIYAIPIEVMQGTGEELFDHIVHCISDF
LDYMGMKGARLPLGFTFSFPCKQTSLDAGILLNWTKGFKATDCEGEDVVNLLREAIKRREEFELDVVAVV
NDTVGTMMTCAYEDPNCEIGLIVGTGSNACYMEEMRNIEMVEGDEGRMCVNMEWGAFGDNGCLDDIRTIY
DKAVDELSLNSGKQRYEKMISGMYLGEIVRNILIDFTKRGFLFRGQISETLKTRGIFETKFLSQIESDRL
ALLQVRAILQQLGLDSTCDDSIIVKEVCGAVSRRAAQLCGAGMAAVVDKIRENRGLDHLEITVGVDGTLY
KLHPHFSKIMHQTVKELAPKCDVTFLLSEDGSGKGAALITAVGCRLREAEQQ
>anc28_altAll|avgPP:0.993|lnPP:-12.710|num_ambig:8|num_ambig_gaps:0
LYAMRLSDETLLDIMTRFRREMENGLSRDFNPTATVKMLPTFVRSIPDGSEKGDFIALDLGGSFFRILRV
KVSHEKKQTVQMESQIYDTPEDIIHGSGTRLFDHVAECLGDFMEKQQIKDKKLPVGFTFSFPCRQTKLDE
GVLITWTKRFKASGVEGADVVKLLNKAIKKRGDYDADIMAVVNDTVGTMMTCGFDDQRCEVGIIIGTGTN
ACYMEELRHIDLVEGDEGRMCINTEWGAFGDDGSLEDIRTEFDREIDRGSLNPGKQLFEKMVSGMYMGEL
VRLILVKMAKEGLLFEGRITPELLTKGKFETKHVSAIEKSKEGLNKAKEILTRLGVEPSHEDCIAVQHVC
TIVSFRSANLIAATLGAILTRLRDNKGVPRLRTTVGVDGSLYKMHPQYARRLHKTVRRLVPDCDVRFLLS

ESGSGKGAAMVTAVAYRLAEQHRQIDETLAEFKLSHEQLLEVKKRMRTEIEAGLRKKTHESAKVKMLPTF
VRSTPDGTENGDFLALDLGGTNFRVLLVKIRSGKRRTVEMHNKIYAIPIEVMQGTGEELFDHIVHCISDF
LDYMGMKGARLPLGFTFSFPCKQTSLDAGILLNWTKGFKATDCEGEDVVNLLREAIKRREEFELDVVAVV
NDTVGTMMTCAYEDPNCEIGLIVGTGSNACYMEEMRNIEMVEGDEGRMCVNMEWGAFGDNGCLDDIRTIY
DKAVDELSLNSGKQRYEKMISGMYLGEIVRNILIDFTKRGFLFRGQISEALKTRGIFETKFLSQIESDRL
ALLQVRAILQQLGLDSTCDDSIIVKEVCGAVSRRAAQLCGAGMAAVVDKIRENRGLDHLDITVGVDGTLY
KLHPHFSKIMHQTVKELAPKCDVTFLLSEDGSGKGAALITAVGCRLREAEQQ
>anc29|avgPP:0.994|lnPP:-8.781|num_ambig:9|num_ambig_gaps:0
LYSMRLSDETLLDIMTRFRREMENGLSRDFNPTATVKMLPTFVRSIPDGSEKGDFIALDLGGSFFRILRV
KVSHEKKQTVQMESQIYDTPEDIMHGSGTRLFDHVAECLGDFMEKQQIKDKKLPVGFTFSFPCRQTKLDE
GILITWTKRFKASGVEGADVVKLLNKAIKKRGDYDADIMAVVNDTVGTMMTCGFDDQRCEVGIIIGTGTN
ACYMEELRHIDLVEGDEGRMCINTEWGAFGDDGSLEDIRTEFDREIDRGSLNPGKQLFEKMVSGMYMGEL
VRLILVKMAKEGLLFEGRITPELLTKGKFETKHVSAIEKSKEGLNKAKEILTRLGVEPSHEDCIAVQHVC
TIVSFRSANLIAATLGAILTRLKDNKGVPRLRTTVGVDGSLYKMHPQYARRLHKTVRRLVPDCDVRFLLS
ESGSGKGAAMVTAVAYRLAEQHRQIDETLAEFKLSHEQLLEVKKRMRTEMEAGLRKKTHESATVKMLPTF
VRSTPDGTENGDFLALDLGGTNFRVLLVKIRSGKRRTVEMHNKIYAIPIEVMQGTGEELFDHIVHCISDF
LDYMGMKGARLPLGFTFSFPCKQTSLDAGILLNWTKGFKATDCEGEDVVNLLREAIKRREEFELDVVAIV
NDTVGTMMTCAYEDPNCEIGLIVGTGSNACYMEEMRNIEMVEGDEGRMCVNMEWGAFGDNGCLDDIRTIY
DKAVDELSLNPGKQRYEKMISGMYLGEIVRNILIDFTKRGFLFRGQISETLKTRGIFETKFLSQIESDRL
ALLQVRAILQQLGLDSTCDDSIIVKEVCGAVSRRAAQLCGAGMAAVVDKIRENRGLDHLEITVGVDGTLY
KLHPHFSKIMHQTVKELAPKCDVTFLLSEDGSGKGAALITAVGCRLREAEQQ
>anc29_altAll|avgPP:0.993|lnPP:-12.220|num_ambig:9|num_ambig_gaps:0
LYSMRLSDETLLDIMARFRREMENGLSRDFNPTATVKMLPTFVRSIPDGSEKGDFIALDLGGSYFRILRV
KVSHEKKQTVQMESQIYDTPEDIIHGSGTRLFDHVAECLGDFMEKQQIKDKKLPVGFTFSFPCRQTKLDE
GVLITWTKRFKASGVEGADVVKLLNKAIKKRGDYDADIMAVVNDTVGTMMTCGFDDQRCEVGIIIGTGTN
ACYMEELRHIDLVEGDEGRMCINTEWGAFGDDGSLEDIRTEFDREIDRGSLNPGKQLFEKMVSGMYMGEL
VRLILVKMAKEGLLFEGRITPELLTKGKFETKHVSAIEKSKEGLNKAKEILTRLGVEPSHEDCIAVQHVC
TIVSFRSANLIAATLGAILTRLRDNKGVPRLRTTVGVDGSLYKMHPQYARRLHKTVRRLVPDCDVRFLLS
ESGSGKGAAMVTAVAYRLAEQHRQIDETLAEFKLSHEQLLEVKKRMRTEMEAGLRKKTHESAKVKMLPTF
VRSTPDGTENGDFLALDLGGTNFRVLLVKIRSGKRRTVEMHNKIYAIPIEVMQGTGEELFDHIVHCISDF
LDYMGMKGARLPLGFTFSFPCKQTSLDAGILLNWTKGFKATDCEGEDVVNLLREAIKRREEFELDVVAIV
NDTVGTMMTCAYEDPNCEIGLIVGTGSNACYMEEMRNIEMVEGDEGRMCVNMEWGAFGDNGCLDDIRTVY
DKAVDELSLNPGKQRYEKMISGMYLGEIVRNILIDFTKRGFLFRGQISEALKTRGIFETKFLSQIESDRL
ALLQVRAILQQLGLDSTCDDSIIVKEVCGAVSRRAAQLCGAGMAAVVDKIRENRGLDHLEITVGVDGTLY
KLHPHFSKIMHQTVKELAPKCDVTFLLSEDGSGKGAALITAVGCRLREAEQH
>anc30|avgPP:0.993|lnPP:-11.267|num_ambig:13|num_ambig_gaps:0
LYSMRLSDETLQDIMTRFRREMENGLGRDTNPTATVKMLPTFVRSIPDGSEKGDFIALDLGGSNFRILRV
KVSHEKKQTVQMESQIYDTPEDIIHGSGTQLFDHVAECLGDFMEKQQIKDKKLPVGFTFSFPCRQTKLDE
GILITWTKRFKASGVEGADVVKLLNKAIKKRGDYDADIMAVVNDTVGTMMTCGFDDQRCEVGIIIGTGTN
ACYMEELRHIDLVEGDEGRMCINTEWGAFGDDGSLEDIRTEFDREIDRGSLNPGKQLFEKMVSGMYMGEL
VRLILVKMAKEGLLFEGRITPELLTKGKFETKHVSAIEKSKEGLNKAKEILTRLGVEPSHEDCIAVQHVC
TIVSFRSANLIAATLGAILTRLKDNKGVPRLRTTVGVDGSLYKMHPQYARRLHKTVRRLVPDCDVRFLLS
ESGSGKGAAMVTAVAYRLAEQHRQIDETLAEFKLSHEQLLEVKKRMRTEMERGLRKKTHESATVKMLPTF
VRSTPDGTENGDFLALDLGGTNFRVLLVKIRSGKRRTVEMHNKIYAIPIEVMQGTGEELFDHIVHCISDF
LDYMGMKGARLPLGFTFSFPCKQTSLDAGILLNWTKGFKATDCEGEDVVNLLREAIKRREEFELDVVAIV
NDTVGTMMTCAYEDPNCEIGLIVGTGSNACYMEEMRNIEMVEGDEGRMCVNMEWGAFGDNGCLDDIRTQY
DKAVDELSLNPGKQRYEKMISGMYLGEIVRNILIDFTKRGFLFRGQISETLKTRGIFETKFLSQIESDRL
ALLQVRAILQQLGLDSTCDDSIIVKEVCGAVSRRAAQLCGAGMAAVVDKIRENRGLDHLEITVGVDGTLY
KLHPHFSKIMHQTVKELAPKCDVTFLLSEDGSGKGAALITAVGCRLREAEQQ
>anc30_altAll|avgPP:0.990|lnPP:-17.816|num_ambig:13|num_ambig_gaps:0
LYSMRLSDETLLDIMTRFRREMENGLGRDTNPTATVKMLPTFVRSIPDGSEKGDFIALDLGGSNFRILRV
KVSHEKKQTVQMESQIYETPEDIMHGSGTRLFDHVAECLGDFMEKQKIKDKKLPVGFTFSFPCRQTKLDE

GVLITWTKRFKASGVEGADVVKLLNKAIKKRGDYDADIMAVVNDTVGTMMTCGFDDQRCEVGIIIGTGTN
ACYMEELRHIDLVEGDEGRMCINTEWGAFGDDGSLEDIRTEFDREIDRGSLNPGKQLFEKMVSGMYMGEL
VRLILVKMAKEGLLFEGRITPELLTKGKFETKHVSAIEKSKEGLNKAKEILTRLGVEPSHEDCIAVQHVC
TIVSFRSANLIAATLGAILTRLRDNKGVPRLRTTVGVDGSLYKMHPQYARRLHKTVRRLVPDCDVRFLLS
ESGSGKGAAMVTAVAYRLAEQHRQIDETLAEFKLSHEQLLEVKKRMRTEMERGLRKKTHESATVKMLPTF
VRSTPDGTENGDFLALDLGGTNFRVLLVKIRSGKRRSVEMHNKIYAIPIEVMQGTGEELFDHIVHCISDF
LDYMGMKGARLPLGFTFSFPCKQTSLDAGILLNWTKGFKATDCEGEDVVNLLREAIKRREEFELDVVAIV
NDTVGTMMTCAYEDPNCEVGLIVGTGSNACYMEEMRNIEMVEGDEGRMCVNMEWGAFGDNGCLDDIRTEY
DRAVDELSLNPGKQRYEKMISGMYLGEIVRNILIDFTKRGFLFRGQISEALKTRGIFETKFLSQIESDRL
ALLQVRAILQQLGLDSTCDDSIIVKEVCGAVSRRAAQLCGAGMAAVVDKIRENRGLDHLEITVGVDGTLY
KLHPHFSKIMHQTVKELAPKCDVTFLLSEDGSGKGAALITAVGCRLREAEQH
>anc31|avgPP:0.985|lnPP:-22.415|num_ambig:20|num_ambig_gaps:0
LYSMRLSDETLQDIMTRFRTEMEKGLGRDTNPTATVKMLPTFVRSTPDGSEKGDFLALDLGGSNFRILRV
KVSHERKQKVQMESQIYATPEDIIHGSGTQLFDHVAECLGDFMEKQQIKDKKLPLGFTFSFPCRQTKLDE
GILITWTKGFKASGVEGADVVKLLRKAIKKRGDYDVDIMAVVNDTVGTMMTCGFDDQRCEVGLIIGTGTN
ACYMEELRHIDLVEGDEGRMCINTEWGAFGDDGSLEDIRTEFDREIDRGSLNPGKQLFEKMISGMYMGEL
VRLILVKMAKEGLLFEGKITPELLTKGKFETKHVSAIEKSKEGLSKAKEILTRLGLEPSEEDCIAVQHVC
TIVSFRSANLCAATLGAILTRLKDNKGVPRLRTTVGVDGSLYKMHPQYARRLHKTVRRLVPDCDVRFLLS
ESGSGKGAAMVTAVAYRLASQRKQIDETLAPFKLSHEQLLEVKKRMRTEMERGLRKKTHESATVKMLPTF
VRSTPDGTENGDFLALDLGGTNFRVLLVKIRSGKRRSVEMHNKIYAIPLEVMQGTGEELFDHIVHCISDF
LDYMGMKGARLPLGFTFSFPCKQTSLDEGILLNWTKGFKATDCEGEDVVNLLREAIKRREEFELDVVAIV
NDTVGTMMTCAYEDPNCEIGLIVGTGSNACYMEEMRNIEMVEGDEGRMCVNMEWGAFGDNGCLDDIRTEY
DRAVDELSLNPGKQRYEKMISGMYLGEIVRNILIDFTKRGFLFRGQISERLKTRGIFETKFLSQIESDRL
ALLQVRAILQQLGLDSTCDDSIIVKEVCGAVSRRAAQLCGAGMAAVVDKIRENRGLDHLEITVGVDGTLY
KLHPHFSKIMHETVKELAPKCDVTFLLSEDGSGKGAALITAVACRLREAGQH
>anc31_altAll|avgPP:0.982|lnPP:-31.829|num_ambig:20|num_ambig_gaps:0
LYSMRLSDETLQDIMTRFRTEMEKGLGRDTNPTATVKMLPTFVRSTPDGSEKGDFLALDLGGSNFRILRV
KVSHEKKQKVEMESQIYSTPEDIMHGSGTRLFDHVAECLGDFMEKQQIKDKKLPLGFTFSFPCRQTKLDE
SVLITWTKGFKASGVEGADVVKLLRKAIKKRGDYDVDIMAVVNDTVGTMMTCGFDDQRCEVGLIIGTGTN
ACYMEELRHIDLVEGDEGRMCINTEWGAFGDDGSLEDIRTEFDREIDRGSLNPGKQLFEKMISGMYMGEL
VRLILVKMAKEGLLFEGRITPELLTKGKFETKHVSAIEKSKEGLNKAKEILTRLGLEPSEEDCIAVQHVC
TIVSFRSANLSAAALGAILTRLRENKGVPRLRTTVGVDGSLYKMHPQYARRLHKTVRRLVPDCDVRFLLS
ESGSGKGAAMVTAVAYRLASQRKQIDETLAPFKLSHEQLLEVKKRMRTEMERGLKKKTHESATVKMLPTF
VRSTPDGTENGDFLALDLGGTNFRVLLVKIRSGKRRSVEMHNKIYAIPLEVMQGTGEELFDHIVHCISDF
LDYMGMKGARLPLGFTFSFPCKQTSLDQGILLNWTKGFKATDCEGEDVVNLLREAIKRREEFELDVVAIV
NDTVGTMMTCAYEDPNCEVGLIVGTGSNACYMEEMRNIEMVEGDEGRMCVNMEWGAFGDNGCLDDIRTQY
DKAVDELSLNPGKQRYEKMISGMYLGEIVRNILIDFTKRGFLFRGQISERLKTRGIFETKFLSQIESDRL
ALLQVRAILQQLGLDSTCDDSIIVKEVCGAVSRRAAQLCGAGMAAVVDKIRENRGLDHLEITVGVDGTLY
KLHPHFSKIMHETVKELAPKCDVTFLLSEDGSGKGAALITAVACRLREAEQQ
>anc32|avgPP:0.990|lnPP:-16.577|num_ambig:15|num_ambig_gaps:0
LYAMRLSDDQLLDIAARFRAEMEKGLSAESNPTAAVKMLPTHVRSTPDGSEKGEFLALDLGGSKFKVLQV
KVSEDGKRKVEMESQTYPTPEEILSGKGTDLFDHVAECLKDFMQKKHINQKKQPLGFTFSFPCRQSKLDE
GVLLSWSKNFKARGVQGTDVVELLRKAIDRAGGIDVDVLALVNDTVGTMMTCGYDDQHCEVGVIIGTGTN
ACYMEEMRHIDLVEGDEGRMCINTEWGAFGDDGSLDDFITDFDREIDAASINPGKQLFEKMISGLYMGEL
VRLILLKMTKKGLLFKGRASDALRTKGKFETKHVAAIEKYKEGLSNTKEILTDLGLNPSEDDCIAVQHVC
TIVSFRSANLCAAALAAILTRIRENKKLKRLRTTVGVDGTVYRTHPQYAKRLHKVVRRLVPDCHVRFVLS
ESGSGKGAAMVTAVAQRLVSQRKQIDETLAPFRLSRQQLQQVRDKMRVELERGLKKDTHHTASVKMLPSY
VYRTPDGTERGKFLALDLGGTNFRVLVVKIRSGLRRSVRMYNKIYAIPLEIMQGTGEELFDHIVQCISDF
LDYMGMKNTRLPLGFTFSFPCRQTGIDKGTLVSWTKGFKATDCEGNDVVDMLREAIKRRNEFDLDIVAVV
NDTVGTMMTCAYEDPNCEIGLIAGTGSNVCYMEEMRNIETVEGDEGRMCVNTEWGGFGENGCIEDIRTKF
DREVDEGSLNVGKQRFEKMTSGMYLGEIVRQILIDLTRRGLLFRGRITERLKTRGIFETKFLSQIESDRL
ALLQVRAILQQLGLDSTCDDSIIVKEVCGAVSRRAAQLCGAGMAAVVDKIRENRGLDKLDITVGVDGTLY

KLHPHFSSILKETVQELAPQCDVTFVLSEDGSGKGAALITAVARKLHSPAT
>anc32_altAll|avgPP:0.987|lnPP:-22.847|num_ambig:15|num_ambig_gaps:0
LYAMRLSDDQLLDIAARFRAEMEKGLSAESNPTAAVKMLPTHVRSTPDGSEKGEFLALDLGGSKFKVLQV
KVSEDGKRRVEMESKTYPTPEEILSGKGTDLFDHVAESLKDFMQRKHINQKKQPLGFTFSFPCRQSKLDE
GVLLSWSKNFKARGVQGTDVVELLRKAIDRAGGIDVDVLALVNDTVGTMMTCGYDDQHCEVGVIVGTGTN
ACYMEEMRHIDLVEGDEGRMCINTEWGAFGDDGSLDDFITDFDREIDAASINPGKQLFEKMISGLYMGEL
VRLILLKMTKKGLLFKGRASDALRTKGKFETKHVSAIEKYKEGLNNTKEILTALGLDPSEDDCIAVQHVC
TIVSFRSANLCAAALAAILTRIRENKKLKRLRTTVGVDGTVYRTHPQYAKRLHKVVRRLVPDCHVRFLLS
ESGSGKGAAMVTAVAQRLVSQRRQIDETLAPFRLSRQQLQQVRDKMRVELERGLKKDTHHTASVKMLPSY
VYRTPDGTERGKFLALDLGGTNFRVLVVKIRSGLRRSVRMYNKIYAIPLEIMQGTGEELFDHIVQCISDF
LDYMGMKNTRLPLGFTFSFPCRQTGIDKGTLVSWTKGFKATDCEGNDVVDMLREAIKRRNEFDLDIVAVV
NDTVGTMMTCAYEDPNCEIGLIAGTGSNVCYMEEMRNIETVEGDEGRMCVNTEWGGFGENGCIEDIRTKF
DREVDEGSLNVGKQRFEKMTSGMYLGEIVRQILIDLTRRGLLFRGRITERLKTRGIFETKFLSQIESDRL
ALLQVRAILQQLGLDSTCDDSIIVKEVCGAVSRRAAQLCGAGMAAVVDKIRENRGLDKLDITVGVDGTLY
KLHPHFSRILKETVQELAPQCDVTFVLSEDGSGKGAALITAVARRLHSQAR
>anc33|avgPP:0.985|lnPP:-23.182|num_ambig:12|num_ambig_gaps:1
LYAMRLSDDQLLDISARFRAEMEKGLSAESNATATVRMLPTHVRSTPDGSEKGEFLALDLGGSKFKVLQV
KVSEDGKRKVEMESQTYPIPEEVLNGRGTDLFDHVAESLRDFMQKNNINQKKKPLGFTFSFPCRQSKLDE
GVLLSWSKNFKARGVQGTDVVESLRQAIDRVGDIDVNVLALVNDTVGTMMTCGFDDQHCEVGVIVGTGTN
ACYMEEMRHIDLVEGDEGRMCINTEWGAFGDDGALDDFITDFDREIDAASINPGKQLFEKMVSGMYMGEL
VRIILLKMARKGLLFGGRVSDALCTKGKLQTKHVALMEKYKEGLKNTKNILTDLGLNPSEDDCVAVQHVC
TIVSFRSANLCAAALAAILTRIRENRKLKTLRTTVGVDGTVYRTHPQYAKRLHKVVRRLVPECHVRFVLS
ESGSGKGAAMVTAVAQRLLSQRRQVDQTLAPFRLSAEQLQQVRDRMRVELERGLRKDTHSTASVKMLPSY
VYRTPSGTERGKYLALDLGGTNFRVLVVKIRSGLRRSVRMYNKIYAIPLEIMQGTGEELFDHIVQCISDF
LDYMGMKNTRLPLGFTFSFPCRQTGIDKGTLVSWTKGFKATDCEGNDVVDMLREAIKRRNEFDLDIVAVV
NDTVGTMMTCAYEDPKCEIGLIAGTGSNVCYMEEMRNIETVEGDEGRMCVNTEWGGFGENGALEDIRTKF
DREVDEGSLNVGKQRFEKMTSGMYLGEIVRQILIDLTRRGLLFRGRITERLKTPGIFETKFLSQIESDRL
ALLQVRSILQHLGLDSTCDDSIIVKEVCGAVSRRAAQLCGAGMAAVVDKIRENRGLDHLDITVGVDGTLY
KLHPHFSSILKETVKELAPKCDVTFVLSEDGSGKGAALITAVARSLCGGEV
>anc33_altAll|avgPP:0.983|lnPP:-28.925|num_ambig:12|num_ambig_gaps:1
LYAMRLSDDQLLDISARFRAEMEKGLSAESNATAAVRMLPTHVRSTPDGSEKGEFLALDLGGSKFKVLQV
KVSEDGKRKVEMESETYPIPEEVLNGRGTDLFDHVAESLRDFMQKNNINQKKKPLGFTFSFPCRQSKLDE
GVLLSWSKNFKARGVQGTDVVESLRQAIDRVGDIDVNVLALVNDTVGTMMTCGFDDQHCEVGVIIGTGTN
ACYMEEMRHIDLVEGDEGRMCINTEWGAFGDDGALDDFITDFDREIDAASINPGKQLFEKMVSGMYMGEL
VRIILLKMARRGLLFGGRVSDALCTKGKLQTRHVALMEKYKEGLKNTRNILTDLGLNPSEDDCIAVQHVC
TIVSFRSANLCAAALAAILTRIRENRKLKTLRTTVGVDGTVYRTHPQYAKRLHKVVRRLVPECHVRFVLS
ESGSGKGAAMVTAVAQRLVSQRRQVDQTLAPFSLSAEQLQQVRDRMRVELERGLRKDTHSTASVKMLPSY
VYNTPSGTERGKYLALDLGGTNFRVLVVKIRSGLRRSVRMYNKIYAIPLEIMQGTGEELFDHIVQCISDF
LDYMGMKNTRLPLGFTFSFPCRQTGIDKGTLVSWTKGFKATDCEGNDVVDMLREAIKRRNEFDLDIVAVV
NDTVGTMMTCAYEDPKCEIGLIAGTGSNVCYMEEMRNIETVEGDEGRMCVNTEWGGFGENGALEDIRTKF
DREVDEGSLNVGKQRFEKMTSGMYLGEIVRQILIDLTRRGLLFRGRITERLKTPGIFETKFLSQIESDRL
ALLQVRSILQHLGLDSTCDDSIIVKEVCGAVSRRAAQLCGAGMAAVVDKIRENRGLDHLDITVGVDGTLY
KLHPHFSRILKETVNELAPKCDVTFVLSEDGSGKGAALITAVARSLCGGEV
>anc34|avgPP:0.992|lnPP:-12.376|num_ambig:11|num_ambig_gaps:0
LYAMRLSDEQLLDISSRFRAEMEKGLSGESNATATVKMLPTHVRSTPDGSEKGEFLALDLGGSKFKVLQV
KVSEDGKRKVQMESETFPVPEEILNGRGTDLFDHVAESLKAFMQKNNINQKKKPLGFTFSFPCLQSKLDE
GVLLSWSKNYKARGVQGTNVVQLLRQAINNVGGIDVDVLALVNDTVGTMMTCGYDDQHCEVGVIIGTGTN
ACYMEEMRHIDLVEGDEGRMCINTEWGAFGDDGALDDFITSFDREIDAASINPGKQLFEKMVSGMYMGEL
VRLILLKMAKKGLLFKGRVSDALRTKGKFQTKHVCLIEEYKDGLKNTRDILTELGLSPSEDDCIAVQHVC
TIVSFRSANLCAATLAAILTRIRENKKLKTLRTTVGVDGTVYRTHPQYAKRLHKVVRRLVPECHVRFVLS
ESGSGKGAAMVTAVAQRLASQRKEVDDTLAGFALSHTQLQQVKDKMRVELERGLRKDSHGSASVKMLPSY
VYRTPDGTERGKFLALDLGGTNFRVLVVKIRSGIRKSVRMYNKIYAIPLEIMQGTGEELFDHIVQCISDF

LDYMGMKNTCLPLGFTFSFPCRQTGIDKGILVSWTKGFKATDCEGNDVVDMLREAIKRRNEFDLDIVAVV
NDTVGTMMTCAYEDPKCEIGLIAGTGSNVCYMEEMKNIETVEGDEGQMCVNTEWGGFGDNDNIEDIRTRF
DREVDEGSLNVGKQRFEKMTSGMYLGEIVRQILIDLTKRGLLFRGRITEPLKTRGIFETKFLSQIESDRL
ALLQVRSILQHLGLDSTCDDSIIVKEVCGAVSRRAAQLCGAGMAAVVDKIRENRGLDHLDITVGVDGTLY
KLHPHFSRILKDTVKELAPKCDVDFVLSEDGSGKGAALITAVARQYCRKT
>anc34_altAll|avgPP:0.990|lnPP:-18.551|num_ambig:11|num_ambig_gaps:0
LYAMRLSDEQLLDISSRFRAEMEKGLSSESNATATVKMLPTHVRSTPDGSEKGEFLALDLGGSKFKVLQV
KVSEDGKRKVQMESETFPVPEEILNGRGTDLFDHVAESLKAFMQKNNINQKKKPLGFTFSFPCLQSKLDE
GVLLSWSKNYKARGVQGTNVVQLLRQAINNVGGIDVDVLALVNDTVGTMMTCGYDDQHCEVGVIIGTGTN
ACYMEEMRHIDLVEGDEGRMCINTEWGAFGDDGALDDFITSFDREIDAASINPGKQLFEKMVSGMYMGEL
VRLILLKMAKRGLLFKGRVSDALRTKGKFQTKHVCLIEEYKDGLKNTKDILTELGLSPSEDDCIAVQHVC
TIVSFRSANLCAATLAAILTRIRENKKLKTLRTTVGVDGTVYRTHPQYAKRLHKVVRRLVPECHVRFVLS
ESGSGKGAAMVTAVAQRLASQRKEIDDTLAAFALSHAQLQQVKDKMRVELERGLRKDSHGSASVKMLPSY
VYRTPDGTERGKFLALDLGGTNFRVLVVKIRSGIRKSVRMYNKIYAIPLEIMQGTGEELFDHIVQCISDF
LDYMGMKNTCLPLGFTFSFPCRQTGIDKGILVSWTKGFKATDCEGNDVVDMLREAIKRRNEFDLDIVAVV
NDTVGTMMTCAYEDPKCEIGLIAGTGSNVCYMEEMRNIETVEGDEGHMCVNTEWGGFGDNDNIEDIRTRF
DREVDEGSLNVGKQRFEKMTSGMYLGEIVRQILIDLTKRGLLFRGRITEPLKTRGIFETKFLSQIESDRL
ALLQVRSILQHLGLDSTCDDSIIVKEVCGAVSRRAAQLCGAGMAAVVDKIRENRGLDHLDITVGVDGTLY
KLHPHFSRILKETVKELAPKCNVDFVLSEDGSGKGAALITAVARQHCRKT
>anc35|avgPP:0.990|lnPP:-14.539|num_ambig:15|num_ambig_gaps:1
LYAMRLSDDQLLDISARFRAEMEKGLSGESNATATVKMLPTHVRSTPDGSEKGEFLALDLGGSKFKVLQV
KVSEDGKRKVQMESETYPIPEEILNGRGTDLFDHVAESLKAFMQKNNINQKKKPLGFTFSFPCAQSKLDE
GVLLSWSKNYKARGVQGTNVVQSLRQAINKVGGIDVDVLALVNDTVGTMMTCGYDDQHCEVGVIIGTGTN
ACYMEEMRHIDLVEGDEGRMCINTEWGAFGDDGALDDFITDFDREIDAASINPGKQLFEKMVSGMYMGEL
VRLILLKMAKKGLLFKGRVSDALRTKGKFQTKHVCLIEEYKDGLKNTRDILTELGLNPSEDDCIAVQHVC
TIVSFRSANLCAAALAAILTRIRENKKLKTLRTTVGVDGTVYRTHPQYAKRLHKVVRRLVPDCHVRFVLS
ESGSGKGAAMVTAVAQRLASQRKEVDDTLAAFALSHTQLQQVKDKMRVELERGLRKDSHGSASVKMLPSY
VYRTPDGTERGKFLALDLGGTNFRVLVVKIRSGIRKSVRMYNKIYAIPLEIMQGTGEELFDHIVQCISDF
LDYMGMKNTCLPLGFTFSFPCRQTGIDKGTLVSWTKGFKATDCEGNDVVDMLREAIKRRNEFDLDIVAVV
NDTVGTMMTCAYEDPKCEIGLIAGTGSNVCYMEEMRNIETVEGDEGQMCVNTEWGGFGENDNIEDIRTRF
DREVDEGSLNVGKQRFEKMTSGMYLGEIVRQILIDLTKRGLLFRGRITEPLKTRGIFETKFLSQIESDRL
ALLQVRSILQHLGLDSTCDDSIIVKEVCGAVSRRAAQLCGAGMAAVVDKIRENRGLDHLDITVGVDGTLY
KLHPHFSRILKDTVKELAPKCDVDFVLSEDGSGKGAALITAVARQYCGGKT
>anc35_altAll|avgPP:0.987|lnPP:-22.513|num_ambig:15|num_ambig_gaps:1
LYAMRLSDDQLLDISARFRAEMEKGLSSESNATATVKMLPTHVRSTPDGSEKGEFLALDLGGSKFKVLQV
KVSEDGKRKVQMESETYPIPEEILNGRGTDLFDHVAESLKAFMQKNNINQKKKPLGFTFSFPCAQSKLDE
GVLLSWSKNFKARGVQGTNVVQSLRQAINNVGGIDVDVLALVNDTVGTMMTCGYDDQHCEVGVIIGTGTN
ACYMEEMRHIDLVEGDEGRMCINTEWGAFGDDGALDDFITDFDREIDAASINPGKQLFEKMVSGMYMGEL
VRLILLKMAKKGLLFKGRVSDALRTKGKFQTKHVCLIEEYKDGLKNTKDILTELGLSPSEDDCIAVQHVC
TIVSFRSANLCAAALAAILTRIRENKKLKTLRTTVGVDGTVYRTHPQYAKRLHKVVRRLVPDCHVRFVLS
ESGSGKGAAMVTAVAQRLASQRKEIDDTLASFALSHAQLQQVKDKMRVELERGLRKDSHSSASVKMLPSY
VYRTPDGTERGKFLALDLGGTNFRVLVVKIRSGIRKSVRMYNKIYAIPLEIMQGTGEELFDHIVQCISDF
LDYMGMKNTCLPLGFTFSFPCRQTGIDKGSLVSWTKGFKATDCEGNDVVDMLREAIKRRNEFDLDIVAVV
NDTVGTMMTCAYEDPKCEIGLIAGTGSNVCYMEEMKNIETVEGDEGQMCVNTEWGGFGENDNIEDIRTKF
DREVDEGSLNVGKQRFEKMTSGMYLGEIVRQILIDLTKRGLLFRGRITEPLKTRGIFETKFLSQIESDRL
ALLQVRSILQHLGLDSTCDDSIIVKEVCGAVSRRAAQLCGAGMAAVVDKIRENRGLDHLDITVGVDGTLY
KLHPHFSRILKETVKELAPKCNVDFVLSEDGSGKGAALITAVARQHCGGKT
>anc36|avgPP:0.991|lnPP:-14.261|num_ambig:17|num_ambig_gaps:1
LYAMRLSDDQLLDISARFRAEMEKGLSGESNATAAVKMLPTHVRSTPDGSEKGEFLALDLGGSKFKVLQV
KVSEDGKRKVQMESETYPIPEEILNGRGTDLFDHVAESLKAFMQKNNINQKKKPLGFTFSFPCAQSKLDE
GVLLSWSKNYKARGVQGTNVVQSLRQAINKVGGIDVDVLALVNDTVGTMMTCGYDDQHCEVGVIIGTGTN
ACYMEEMRHIDLVEGDEGRMCINTEWGAFGDDGALDDFITDFDREIDAASINPGKQLFEKMVSGMYMGEL

VRLILLKMAKKGLLFKGRVSDALRTKGKFQTKHVCLIEEYKDGLKNTRDILTELGLNPSEDDCIAVQHVC
TIVSFRSANLCAAALAAILTRIRENKKLKTLRTTVGVDGTVYRTHPQYAKRLHKVVRRLVPDCHVRFVLS
ESGSGKGAAMVTAVAQRLASQRKEVDDTLASFRLSHEQLQQVRDKMRVELERGLRKDSHGSASVKMLPSY
VYRTPDGTERGKYLALDLGGTNFRVLVVKIRSGLRKSVRMYNKIYAIPLEIMQGTGEELFDHIVQCISDF
LDYMGMKNTCLPLGFTFSFPCRQTGIDKGTLVSWTKGFKATDCEGNDVVDMLREAIKRRNEFDLDIVAVV
NDTVGTMMTCAYEDPKCEIGLIAGTGSNVCYMEEMRNIETVEGDEGQMCVNTEWGGFGENDNIEDIRTKF
DREVDEGSLNVGKQRFEKMTSGMYLGEIVRQILIDLTKRGLLFRGRITEPLKTRGIFETKFLSQIESDRL
ALLQVRSILQHLGLDSTCDDSIIVKEVCGAVSRRAAQLCGAGMAAVVDKIRENRGLDHLDITVGVDGTLY
KLHPHFSRILKDTVKELAPKCDVDFVLSEDGSGKGAALITAVARQYCGGKT
>anc36_altAll|avgPP:0.987|lnPP:-23.692|num_ambig:17|num_ambig_gaps:1
LYAMRLSDDQLKDISARFRAEMEKGLSSESNATATVKMLPTHVRSTPDGSEKGEFLALDLGGSKFKVLQV
KVSEDGKRKVQMESETYPIPEEILNGRGTDLFDHVAESLKAFMQKNNINQKKKPLGFTFSFPCAQSKLDE
GVLLSWSKNFKARGVQGTNVVQSLRQAINRVGDIDVDVLALVNDTVGTMMTCGYDDQHCEVGVIIGTGTN
ACYMEEMRHIDLVEGDEGRMCINTEWGAFGDDGALDDFITDFDREIDAASINPGKQLFEKMVSGMYMGEL
VRLILLKMAKKGLLFKGRVSDALRTKGKFQTKHVCLIEEYKDGLKNTKDILTDLGLNPSEDDCIAVQHVC
TIVSFRSANLCAAALAAILTRIRENKKLKTLRTTVGVDGTVYRTHPQYAKRLHKVVRRLVPDCHVRFVLS
ESGSGKGAAMVTAVAQRLASQRKEIDDTLAAFRLSREQLQQVRDKMRVELERGLRKDSHSSASVKMLPSY
VYRTPDGTERGKYLALDLGGTNFRVLVVKIRSGIRKSVRMYNKIYAIPLEIMQGTGEELFDHIVQCISDF
LDYMGMKNTCLPLGFTFSFPCRQTGIDKGSLVSWTKGFKATDCEGNDVVDMLREAIKRRNEFDLDIVAVV
NDTVGTMMTCAYEDPKCEIGLIAGTGSNVCYMEEMRNIETVEGDEGQMCVNTEWGGFGENDNIEDIRTKF
DREVDEGSLNVGKQRFEKMTSGMYLGEIVRQILIDLTKRGLLFRGRITEPLKTRGIFETKFLSQIESDRL
ALLQVRSILQHLGLDSTCDDSIIVKEVCGAVSRRAAQLCGAGMAAVVDKIRENRGLDHLDITVGVDGTLY
KLHPHFSRILKETVKELAPKCNVDFVLSEDGSGKGAALITAVARQHCGGKT
>anc37|avgPP:0.988|lnPP:-18.607|num_ambig:19|num_ambig_gaps:1
LYAMRLSDDQLKDISARFRAEMEKGLSGESNATAAVKMLPTHVRSTPDGSEKGEFLALDLGGSRFKVLQV
KVSENGKRKVEMESETYPIPEEILNGRGTDLFDHVAESLKDFMQKNNINQKKKPLGFTFSFPCAQSKLDE
GVLLSWSKNFKARGVQGTNVVQSLRQAINRVGDIDVDVLALVNDTVGTMMTCGYDDQHCEVGVIVGTGTN
ACYMEEMRHIDLVEGDEGRMCINTEWGAFGDDGALDDFITDFDREIDAASINPGKQLFEKMVSGMYMGEL
VRLILLKMAKKGLLFGGMVSDALRTKGKFQTKHVCLIEEYKDGLKNTKDILTELGLNPSEDDCIAVQHVC
TIVSFRSANLCAAALAAILTRIRENRKLKTLRTTVGVDGTVYRTHPQYAKRLHKVVRRLVPDCHVRFVLS
ESGSGKGAAMVTAVAQRLMSQRKQVDDTLAPFRLNREQLQQVRDKMRVELERGLRKDSHSSASVKMLPSY
VYRTPDGTERGKYLALDLGGTNFRVLVVKIRSGLRKSVRMYNKIYAIPQEIMQGTGEELFDHIVQCISDF
LDYMGMKNTRLPLGFTFSFPCRQTGIDKGTLVSWTKGFKATDCEGNDVVDMLREAIKRRNEFDLDIVAVV
NDTVGTMMTCAYEDPKCEIGLIAGTGSNVCYMEEMRNIETVEGDEGQMCVNTEWGGFGENGCIEDIRTKF
DREVDEGSLNSGKQRFEKMTSGMYLGEIVRQILIDLTKRGLLFRGRITEPLKTRGIFETKFLSQIESDRL
ALLQVRCILQQLGLDSTCDDSIIVKEVCGAVSHRAAQLCGAGMAAVVDKIRENRGLDHLDITVGVDGTLY
KLHPHFSRILKDTVKELAPKCDVDFVLSEDGSGKGAALITAVARQYCGGET
>anc37_altAll|avgPP:0.985|lnPP:-28.481|num_ambig:19|num_ambig_gaps:1
LYAMRLSDDQLLDISARFRAEMEKGLSSESNATAAVKMLPTHVRSTPDGSEKGEFLALDLGGSKFKVLQV
KVSENGKRKVQMESETYPIPEEILNGRGTDLFDHVAESLKAFMQKNNINQKKKPLGFTFSFPCAQSKLDE
GVLLSWSKNFKARGVQGTNVVQSLRQAINRVGDIDVDVLALVNDTVGTMMTCGYDDQHCEVGVIVGTGTN
ACYMEEMRHIDLVEGDEGRMCINTEWGAFGDDGALDDFITGFDREIDAASINPGKQLFEKMVSGMYMGEL
VRLILLKMAKKGLLFGGMVSDALRTKGKFQTKHVCLIEEYKDGLKNTKEILTELGLNPSDDDCIAVQHVC
TIVSFRSANLCAAALAAILTRIRENRKLKTLRTTVGVDGTVYRTHPQYAKRLHKVVRRLVPDCHVRFVLS
ESGSGKGAAMVTAVAQRLLSQRKEIDDTLASFRLSQEQLQQVRDKMRVELERGLKKDSHSTASVKMLPSY
VYRTPDGTERGKYLALDLGGTNFRVLVVKIRSGIRKSVRMYNKIYAIPHEIMQGTGEELFDHIVQCISDF
LDYMGMKNTRLPLGFTFSFPCRQTGIDKGTLVSWTKGFKATDCEGNDVVDMLREAIKRRNEFDLDIVAVV
NDTVGTMMTCAYEDPKCEIGLIAGTGSNVCYMEEMRNIETVEGDEGQMCVNTEWGGFGENGCIEDIRTKF
DREVDEGSLNSGKQRFEKMTSGMYLGEIVRQILIDLTKRGLLFRGRITEPLKTRGIFETKFLSQIESDRL
ALLQVRCILQQLGLDSTCDDSIIVKEVCGAVSHRAAQLCGAGMAAVVDKIRENRGLDHLDITVGVDGTLY
KLHPHFSRILKDTVKELAPKCDVDFVLSEDGSGKGAALITAVARRYCGGET
>anc38|avgPP:0.991|lnPP:-14.248|num_ambig:18|num_ambig_gaps:1

LYAMRLSDDQLLDISARFRAEMEKGLSGESNATAAVKMLPTHVRSTPDGSEKGEFLALDLGGSKFKVLQV
KVSEDGKRKVEMESETYPIPEEILNGRGTDLFDHVAESLKDFMQKNNINQKKKPLGFTFSFPCAQSKLDE
GVLLSWSKNFKARGVQGTNVVQSLRQAINRVGGIDVDVLALVNDTVGTMMTCGYDDQHCEVGVIVGTGTN
ACYMEEMRHIDLVEGDEGRMCINTEWGAFGDDGALDDFITDFDREIDAASINPGKQLFEKMVSGMYMGEL
VRLILLKMAKKGLLFGGRVSDALRTKGKFQTKHVCLIEEYKDGLKNTKDILTELGLNPSEDDCIAVQHVC
TIVSFRSANLCAAALAAILTRIRENRKLKTLRTTVGVDGTVYRTHPQYAKRLHKVVRRLVPDCHVRFVLS
ESGSGKGAAMVTAVAQRLASQRKQVDDTLAPFRLSREQLQQVRDKMRVELERGLRKDSHSSASVKMLPSY
VYRTPDGTERGKYLALDLGGTNFRVLVVKIRSGLRKSVRMYNKIYAIPLEIMQGTGEELFDHIVQCISDF
LDYMGMKNTRLPLGFTFSFPCRQTGIDKGTLVSWTKGFKATDCEGNDVVDMLREAIKRRNEFDLDIVAVV
NDTVGTMMTCAYEDPKCEIGLIAGTGSNVCYMEEMRNIETVEGDEGQMCVNTEWGGFGENGCIEDIRTKF
DREVDEGSLNVGKQRFEKMTSGMYLGEIVRQILIDLTKRGLLFRGRITEPLKTRGIFETKFLSQIESDRL
ALLQVRSILQHLGLDSTCDDSIIVKEVCGAVSRRAAQLCGAGMAAVVDKIRENRGLDHLDITVGVDGTLY
KLHPHFSRILKDTVKELAPKCDVDFVLSEDGSGKGAALITAVARQYCGGET
>anc38_altAll|avgPP:0.987|lnPP:-23.713|num_ambig:18|num_ambig_gaps:1
LYAMRLSDDQLKDISARFRAEMEKGLSSESNATATVKMLPTHVRSTPDGSEKGEFLALDLGGSKFKVLQV
KVSEDGKRKVQMESETYPIPEEILNGRGTDLFDHVAESLKAFMQKNNINQKKKPLGFTFSFPCAQSKLDE
GVLLSWSKNFKARGVQGTNVVQSLRQAINRVGDIDVDVLALVNDTVGTMMTCGYDDQHCEVGVIIGTGTN
ACYMEEMRHIDLVEGDEGRMCINTEWGAFGDDGALDDFITDFDREIDAASINPGKQLFEKMVSGMYMGEL
VRLILLKMAKKGLLFGGRVSDALRTKGKFQTKHVCLIEEYKDGLKNTREILTDLGLNPSEDDCIAVQHVC
TIVSFRSANLCAAALAAILTRIRENRKLKTLRTTVGVDGTVYRTHPQYAKRLHKVVRRLVPDCHVRFVLS
ESGSGKGAAMVTAVAQRLASQRKEIDDTLASFRLSREQLQQVRDKMRVELERGLKKDSHSTASVKMLPSY
VYRTPDGTERGKYLALDLGGTNFRVLVVKIRSGIRKSVRMYNKIYAIPLEIMQGTGEELFDHIVQCISDF
LDYMGMKNTRLPLGFTFSFPCRQTGIDKGTLVSWTKGFKATDCEGNDVVDMLREAIKRRNEFDLDIVAVV
NDTVGTMMTCAYEDPKCEIGLIAGTGSNVCYMEEMRNIETVEGDEGQMCVNTEWGGFGENGCIEDIRTKF
DREVDEGSLNVGKQRFEKMTSGMYLGEIVRQILIDLTKRGLLFRGRITEPLKTRGIFETKFLSQIESDRL
ALLQVRSILQHLGLDSTCDDSIIVKEVCGAVSHRAAQLCGAGMAAVVDKIRENRGLDHLDITVGVDGTLY
KLHPHFSRILKETVKELAPKCDVDFVLSEDGSGKGAALITAVARQYCGGET
>anc39|avgPP:0.994|lnPP:-8.556|num_ambig:5|num_ambig_gaps:0
LYAMRLHDDQLSDISARFQAEMKKGLSAESNAAAAVKMLPTHVRSTPDGSEKGQFLALDLGGSRFKVLQV
KVREGMGIRRGGVEMEEKTYPIPKELLNGRGTELFDHVSESLKDFLHEKNISLEKKHPLAFTFSFPCEQS
TLNQGLLLNWSKNYRARGLQGKDVVQALRGAIDRTGGMDVEVLAMVNDTVATMMTCGFDDQYCEVGLIIG
TGTNACYMEELRHVDLVEGDEGRMCVNTEWGAFGDDGALNDYITEFDRDVDAASNNPGKQIFEKMVSGMY
LGELVRLVVLKMAKLGLLFDGHVSDALKTKGKITTAHVAAMEEYKNGLKNTRNILMDLDLTPTSDDCVAV
QHVSTIVSFRSSNLVAAGLAAILTRIKQNRNLRTFRITVGVDGTVYKTHPQYPKRLHKVVRRLLPECHVR
FVLSDSGSSKGAALVTAVAQRLASRRRQVDETLSPFRLSQEQLQLVKTRMRAGLEAGLKSKGPSAIKMLP
SFVYRTPDGTEHGKYLALDLGGTNFRALLVNFKKGLQQNTRLYHKIYTIPLEIMQGSGEELFDHIAQCVS
DFLDYMGMKNARLPAGFTFSFPCEQTAIDTGTLVSWTKGFKATDCEGHDVVSMLREAIKRRNEFDLDIVA
VVNDTVGTMMSCAYEDPQCEIGLIAGTGTNVCYMEELKNIEKTEQKTGQVEREEGTPEGEEGKQDDGTER
ERKEGDAEIPKMCINTEWGGLGDDGSLDDIITPYDIEVDQNSVNPGKQRFEKLTSGMYLGEVVRQVLLDL
TRGGLLFRGHVTEALKTPGIFQTKYLSQIESDRLALLQVRSILQQLGLDSTCDDSIIVKEVCGAVSRRAA
QLCGAGMAAVVDKIRENRGLDHLNITVGVDGALYKLHPHFSGVLQETARVLAPQCNVTFLPSEEGSGKGA
ALITAVARQKPEM
>anc39_altAll|avgPP:0.993|lnPP:-11.761|num_ambig:5|num_ambig_gaps:0
LYAMRLHDDQLSDISARFQAEMKKGLSAESNAAAAVKMLPTHVRSTPDGSEKGQFLALDLGGSRFKVLQV
KVREGMGIRRGGVEMEEKTYPIPKELLNGRGTELFDHVSESLKDFLHEKNISLEKKHPLAFTFSFPCEQS
TLNQGLLLNWSKNYRARGLQGKDVVQALRGAIDRTGGMDVEVLAMVNDTVATMMTCGFDDQYCEVGLIIG
TGTNACYMEELRHVDLVEGDEGRMCVNTEWGAFGDDGALNDYITEFDRDVDAASNNPGKQIFEKMVSGMY
LGELVRLVVLKMAKLGLLFDGHVSDALKTKGKITTAHVAAMEEYKNGLKNTRNILMDLDLTPTSDDCVAV
QHVSTIVSFRSSNLVAAGLAAILTRIKQNRNLRTFRITVGVDGTVYKTHPQYPKRLHKVVRRLLPECHVR
FVLSDSGSSKGAALVTAVAQRLASRRRQVDETLSPFRLSQEQLQLVKTRMRAGLEAGLKSKGPSAIKMLP
SFVYRTPDGTEHGKYLALDLGGTNFRALLVNFKKGLQQNTQLYHKIYTIPLEIMQGTGEELFDHIAQCVS
DFLDYMGMKNARLPAGFTFSFPCEQTAIDTGTLVSWTKGFKATDCEGHDVVSMLREAIKRRNEFDLDIVA

VVNDTVGTMMSCAYEDPQCEIGLIAGTGTNVCYMEELKNIEKTEQKKGQVEREEGTPEGEEGKQDDGTER
EGKEGDAEIPKMCINTEWGGLGDDGSLDDIITPYDIEVDQNSINPGKQRFEKLTSGMYLGEVVRQVLLDL
TRGGLLFRGHVTEALKTPGIFQTKYLSQIESDRLALLQVRSILQQLGLDSTCDDSIIVKEVCGAVSRRAA
QLCGAGMAAVVDKIRENRGLDHLNITVGVDGALYKLHPHFSGVLQETARVLAPQCNVTFLPSEEGSGKGA
ALITAVARQKPEM
>anc40|avgPP:0.994|lnPP:-9.347|num_ambig:4|num_ambig_gaps:3
LYAMRLHDDQLSDISARFQAEMKKGLSAESNAAAAVKMLPTHVRSTPDGSEKGQFLALDLGGSKFKVLQV
KVREGMGIRRGGVEMEEKTYPIPKELHIGRGTELFDHVSESLKDFLHEKNISLEKKHPLAFTFSFPCEQS
TLNQGLLLNWSKNFRARGLQGKDVVQALRGAIDRTGGMDVEVLAMVNDTVATMMTCGFDDQYCEVGLIIG
TGTNACYMEELRHVDLVEGDEGRMCVNTEWGAFGDDGALNDYITEFDRDVDAASNNPGKQILEKMVSGMY
LGELVRLVVLKMAKLRLLFDGHVSDALRTKGTITTEHVAAMEEYKNGLKNTRNILMDLDLTPTSDDCVAV
QHVSTIVSFRSSNLVAAGLAAILTRIKQNRNLRTFRITVGVDGSVYKTHPQYPKRLHKVVRRLLPECHVR
FVLSDSGSSKGAALVTAVAQRLASRRRQVDETLSPFRLSQEQLQLVKTRMRAGLEAGLKSKGPSAIKMLP
SFVYHTPDGTEHGKYLALDLGGTNFRALLVNFKQGLQQNSQLYHKIYTIPLEIMQGSGEELFDHIAQCVS
DFLDYMGMKNARLPAGFIFSFPCEQTAIDTGTLVSWTKGFKATDCEGHDVVSMLREAIKRRNEFDLDIAA
VVNDTVATMMSCAYEDPQCEIGLIAGTGTNVCYMEELKNIEKTEQKTGQKEREEETEEGEEGKQDDGRER
ERKEGDAEIPKMCINTEWGGLGDDGSLDDIITPYDIEVDQNSLNPGKQRFEKLTSGMYLGEVVRQVLLDL
TRGGLLFRGHVTEALKTPGMFQTKYLSQIESDRLALLQVRSILQQLGLDSTCDDSIIVKEVCGAVSRRAA
QLCGAGMAAVVDKIRENRGLDHLNITVGVDGTLYKLHPHFSGILQETARVLAPQCNVTFLPSEEGSGKGA
ALITAVARQKLEM
>anc40_altAll|avgPP:0.993|lnPP:-10.774|num_ambig:4|num_ambig_gaps:3
LYAMRLHDDQLSDISARFQAEMKKGLSAESNAAAAVKMLPTHVRSTPDGSEKGQFLALDLGGSKFKVLQV
KVREGMGIRRGGVEMEEKTYPIPKELHIGRGTELFDHVSESLKDFLHEKNISLEKKHPLAFTFSFPCEQS
TLNQGLLLNWSKNFRARGLQGKDVVQALRGAIDRTGGMDVEVLAMVNDTVATMMTCGFDDQYCEVGLIIG
TGTNACYMEELRHVDLVEGDEGRMCVNTEWGAFGDDGALNDYITEFDRDVDAASNNPGKQILEKMVSGMY
LGELVRLVVLKMAKLRLLFNGHVSDALRTKGTITTEHVAAMEEYKNGLKNTRNILMDLDLTPTSDDCVAV
QHVSTIVSFRSSNLVAAGLAAILTRIKQNRNLRTFRITVGVDGSVYKTHPQYPKRLHKVVRRLLPECHVR
FVLSDSGSSKGAALVTAVAQRLASRRRQVDETLSPFRLSQEQLQLVKTRMRAGLEAGLKSKGPSAIKMLP
SFVYHTPDGTEHGKYLALDLGGTNFRALLVNFKQGLQQNSRLYHKIYTIPLEIMQGSGEELFDHIAQCVS
DFLDYMGMKNARLPASFIFSFPCEQTAIDTGTLVSWTKGFKATDCEGHDVVSMLREAIKRRNEFDLDIAA
VVNDTVATMMSCAYEDPQCEIGLIAGTGTNVCYMEELKNIEKTEQKKGQKEREEETEEGEEGKQDDGRER
ERKEGDAEIPKMCINTEWGGLGDDGSLDDIITPYDIEVDQNSLNPGKQRFEKLTSGMYLGEVVRQVLLDL
TRGGLLFRGHVTEALKTPGMFQTKYLSQIESDRLALLQVRSILQQLGLDSTCDDSIIVKEVCGAVSRRAA
QLCGAGMAAVVDKIRENRGLDHLNITVGVDGTLYKLHPHFSGILQETARVLAPQCNVTFLPSEEGSGKGA
ALITAVARQKLEM
>anc41|avgPP:0.995|lnPP:-7.948|num_ambig:5|num_ambig_gaps:3
LYAMRLHDDQLSDISACFQAEMKKGLSAESNAAAAVKMLPTHVRSTPDGSEKGQFLALDLGGSKFKVLQV
KVREGTGIRRGGVEMEEKTYPIPEELLTGRGTELFDHVSESLKDFLHEKNISLEKKHPLAFTFSFPCEQS
TLNQGLLLNWSKNYRARGLQGKDVVQALRGAIDRTGGMDVEVLAMVNDTVATMMTCGFDDQYCEVGLIVG
TGTNACYMEELRHVDLVEGDEGRMCVNTEWGAFGDDGALNDYITEFDRDVDAASNNPGKQILEKMVSGMY
LGELVRLVVLKMAKLGLLFDGHVSDALRTKGKITTAHVAAMEEYKNGLKNTRNILVDLDLTPTSDDCVAV
QHVSTIVSFRSSNLVAAGLAAILTRIKQNRNLRTLRITVGVDGTVYKTHPRYPNRLHKVVRRLLPECHVR
FVLSDSGSSKGAALVTAVAQRLASRRRQVDETLSPFRLSQEQLQLVKTRMRAGLEAGLKSKGPSAIKMLP
SFVYHTPDGTEHGKYLALGLGGTNFRALLVNFKKGLQQNSRLHHKIYTIPLEIMQGSGEELFDHIAQCVS
DFLDYMGMKNARLPAGFTFSFPCEQTAIDTGTLVSWTKGFKATDCEGHDVVSMLREAIKRRNDFDLDIVA
VVNDTVGTMMSCAYEDPQCEIGLIAGTGTNVCYMEELKNIEKTEQKKGKKEREEETEEGEEGKQDDGTER
ERKEGDAEIPKMCINTEWGGLGDDGSLDDIITPYDIEVDQNSLNPGKQRFEKLTSGMYLGEVVRQVLLDL
TRGGLLFRGHVTEALKTPGIFQTKYLSQIESDRLALLQVRSILQQLGLDSTCDDSIIVKEVCGAVSHRAA
QLCGAGMAAVVDKIRENRGLDHLNITVGVDGTLYKLHPHFSGVLQETAKVLAPQCNVTFLPSEGGSGKGA
ALITAVARQKLEM
>anc41_altAll|avgPP:0.994|lnPP:-9.973|num_ambig:5|num_ambig_gaps:3
LYAMRLHDDQLSDISACFQAEMKKGLSAESNAAAAVKMLPTHVRSTPDGSEKGQFLALDLGGSKFKVLQV

KVREGTGIRRGGVEMEEKTYPIPEELLTGRGTELFDHVSESLKDFLHEKNISLEKKHPLAFTFSFPCEQS
TLNQGLLLNWSKNYRARGLQGKDVVQALRGAIDRTGGMDVEVLAMVNDTVATMMTCGFDDQYCEVGLIVG
TGTNACYMEELRHVDLVEGDEGRMCVNTEWGAFGDDGALNDYITEFDRDVDAASNNPGKQILEKMVSGMY
LGELVRLVVLKMAKLGLLFDGHVSDALRTKGKITTAHVAAMEEYKNGLKNTRNILVDLDLTPTSDDCVAV
QHVSTIVSFRSSNLVAAGLAAILTRIKQNRNLRTLRITVGVDGTVYKTHPRYPNRLHKVVRRLLPECHVR
FVLSDSGSSKGAALVTAVAQRLASRRRQVDETLSPFRLSQEQLQLVKTRMRAGLEAGLKSKGPSAIKMLP
SFVYHTPDGTEHGKYLALGLGGTNFRALLVNFKKGLQQNSQLYHKIYTIPLEIMQGSGEELFDHIAQCVS
DFLDYMGMKNARLPAGFTFSFPCEQTAIDTGTLVSWTKGFKATDCEGHDVVSMLREAIKRRNDFDLDIVA
VVNDTVGTMMSCAYEDPQCEIGLIAGTGTNVCYMEELKNIEKTEQKTGKKEREEEMEEGEEGKQDDGTER
ERKEGDAEIPKMCINTEWGGLGDDGSLDDIITPYDIEVDQNSLNPGKQRFEKLTSGMYLGEVVRQVLLDL
TRGGLLFRGHVTEALKTPGIFQTKYLSQIESDRLALLQVRSILQQLGLDSTCDDSIIVKEVCGAVSHRAA
QLCGAGMAAVVDKIRENRGLDHLNITVGVDGTLYKLHPHFSGVLQETAKVLAPQCNVTFLPSEGGSGKGA
ALITAVARRKLEM
>anc42|avgPP:0.998|lnPP:-2.831|num_ambig:2|num_ambig_gaps:3
LYAMRLHDDQLSDISARFQAEMKKGLSAESNAAAAVKMLPTHVRSTPDGSEKGQFLALDLGGSKFKVLQV
KVREGMGIRRGGVEMEEKTYPIPKELLIGRGTELFDHVSESLKDFLHEKNISLEKKHPLAFTFSFPCEQS
TLNQGLLLNWSKNYRARGLQGKDVVQALRGAIDRTGGMDVEVLAMVNDTVATMMTCGFDDQYCEVGLIIG
TGTNACYMEELRHVDLVEGDEGRMCVNTEWGAFGDDGALNDYITEFDRDVDAASNNPGKQILEKMVSGMY
LGELVRLVVLKMAKLGLLFDGHVSDALRTKGKITTAHVAAMEEYKNGLKNTRNILMDLDLTPTSDDCVAV
QHVSTIVSFRSSNLVAAGLAAILTRIKQNRNLRTFRITVGVDGTVYKTHPQYPKRLHKVVRRLLPECHVR
FVLSDSGSSKGAALVTAVAQRLASRRRQVDETLSPFRLSQEQLQLVKTRMRAGLEAGLKSKGPSAIKMLP
SFVYHTPDGTEHGKYLALDLGGTNFRALLVNFKKGLQQNSRLYHKIYTIPLEIMQGSGEELFDHIAQCVS
DFLDYMGMKNARLPAGFTFSFPCEQTAIDTGTLVSWTKGFKATDCEGHDVVSMLREAIKRRNEFDLDIVA
VVNDTVGTMMSCAYEDPQCEIGLIAGTGTNVCYMEELKNIEKTEQKTGQKEREEETEEGEEGKQDDGTER
ERKEGDAEIPKMCINTEWGGLGDDGSLDDIITPYDIEVDQNSLNPGKQRFEKLTSGMYLGEVVRQVLLDL
TRGGLLFRGHVTEALKTPGIFQTKYLSQIESDRLALLQVRSILQQLGLDSTCDDSIIVKEVCGAVSRRAA
QLCGAGMAAVVDKIRENRGLDHLNITVGVDGTLYKLHPHFSGVLQETARVLAPQCNVTFLPSEEGSGKGA
ALITAVARQKLEM
>anc42_altAll|avgPP:0.998|lnPP:-3.556|num_ambig:2|num_ambig_gaps:3
LYAMRLHDDQLSDISARFQAEMKKGLSAESNAAAAVKMLPTHVRSTPDGSEKGQFLALDLGGSKFKVLQV
KVREGMGIRRGGVEMEEKTYPIPKELLIGRGTELFDHVSESLKDFLHEKNISLEKKHPLAFTFSFPCEQS
TLNQGLLLNWSKNYRARGLQGKDVVQALRGAIDRTGGMDVEVLAMVNDTVATMMTCGFDDQYCEVGLIIG
TGTNACYMEELRHVDLVEGDEGRMCVNTEWGAFGDDGALNDYITEFDRDVDAASNNPGKQILEKMVSGMY
LGELVRLVVLKMAKLGLLFDGHVSDALRTKGKITTAHVAAMEEYKNGLKNTRNILMDLDLTPTSDDCVAV
QHVSTIVSFRSSNLVAAGLAAILTRIKQNRNLRTFRITVGVDGTVYKTHPQYPKRLHKVVRRLLPECHVR
FVLSDSGSSKGAALVTAVAQRLASRRRQVDETLSPFRLSQEQLQLVKTRMRAGLEAGLKSKGPSAIKMLP
SFVYHTPDGTEHGKYLALDLGGTNFRALLVNFKKGLQQNSQLYHKIYTIPLEIMQGSGEELFDHIAQCVS
DFLDYMGMKNARLPAGFTFSFPCEQTAIDTGTLVSWTKGFKATDCEGHDVVSMLREAIKRRNEFDLDIVA
VVNDTVGTMMSCAYEDPQCEIGLIAGTGTNVCYMEELKNIEKTEQKKGQKEREEETEEGEEGKQDDGTER
ERKEGDAEIPKMCINTEWGGLGDDGSLDDIITPYDIEVDQNSLNPGKQRFEKLTSGMYLGEVVRQVLLDL
TRGGLLFRGHVTEALKTPGIFQTKYLSQIESDRLALLQVRSILQQLGLDSTCDDSIIVKEVCGAVSRRAA
QLCGAGMAAVVDKIRENRGLDHLNITVGVDGTLYKLHPHFSGVLQETARVLAPQCNVTFLPSEEGSGKGA
ALITAVARQKLEM
>anc43|avgPP:0.998|lnPP:-2.831|num_ambig:2|num_ambig_gaps:3
LYAMRLHDDQLSDISARFQAEMKKGLSAESNAAAAVKMLPTHVRSTPDGSEKGQFLALDLGGSKFKVLQV
KVREGMGIRRGGVEMEEKTYPIPKELLIGRGTELFDHVSESLKDFLHEKNISLEKKHPLAFTFSFPCEQS
TLNQGLLLNWSKNYRARGLQGKDVVQALRGAIDRTGGMDVEVLAMVNDTVATMMTCGFDDQYCEVGLIIG
TGTNACYMEELRHVDLVEGDEGRMCVNTEWGAFGDDGALNDYITEFDRDVDAASNNPGKQILEKMVSGMY
LGELVRLVVLKMAKLGLLFDGHVSDALRTKGKITTAHVAAMEEYKNGLKNTRNILMDLDLTPTSDDCVAV
QHVSTIVSFRSSNLVAAGLAAILTRIKQNRNLRTFRITVGVDGTVYKTHPQYPKRLHKVVRRLLPECHVR
FVLSDSGSSKGAALVTAVAQRLASRRRQVDETLSPFRLSQEQLQLVKTRMRAGLEAGLKSKGPSAIKMLP
SFVYHTPDGTEHGKYLALDLGGTNFRALLVNFKKGLQQNSRLYHKIYTIPLEIMQGSGEELFDHIAQCVS

DFLDYMGMKNARLPAGFTFSFPCEQTAIDTGTLVSWTKGFKATDCEGHDVVSMLREAIKRRNEFDLDIVA
VVNDTVGTMMSCAYEDPQCEIGLIAGTGTNVCYMEELKNIEKTEQKTGQKEREEETEEGEEGKQDDGTER
ERKEGDAEIPKMCINTEWGGLGDDGSLDDIITPYDIEVDQNSLNPGKQRFEKLTSGMYLGEVVRQVLLDL
TRGGLLFRGHVTEALKTPGIFQTKYLSQIESDRLALLQVRSILQQLGLDSTCDDSIIVKEVCGAVSRRAA
QLCGAGMAAVVDKIRENRGLDHLNITVGVDGTLYKLHPHFSGVLQETARVLAPQCNVTFLPSEEGSGKGA
ALITAVARQKLEM
>anc43_altAll|avgPP:0.998|lnPP:-3.556|num_ambig:2|num_ambig_gaps:3
LYAMRLHDDQLSDISARFQAEMKKGLSAESNAAAAVKMLPTHVRSTPDGSEKGQFLALDLGGSKFKVLQV
KVREGMGIRRGGVEMEEKTYPIPKELLIGRGTELFDHVSESLKDFLHEKNISLEKKHPLAFTFSFPCEQS
TLNQGLLLNWSKNYRARGLQGKDVVQALRGAIDRTGGMDVEVLAMVNDTVATMMTCGFDDQYCEVGLIIG
TGTNACYMEELRHVDLVEGDEGRMCVNTEWGAFGDDGALNDYITEFDRDVDAASNNPGKQILEKMVSGMY
LGELVRLVVLKMAKLGLLFDGHVSDALRTKGKITTAHVAAMEEYKNGLKNTRNILMDLDLTPTSDDCVAV
QHVSTIVSFRSSNLVAAGLAAILTRIKQNRNLRTFRITVGVDGTVYKTHPQYPKRLHKVVRRLLPECHVR
FVLSDSGSSKGAALVTAVAQRLASRRRQVDETLSPFRLSQEQLQLVKTRMRAGLEAGLKSKGPSAIKMLP
SFVYHTPDGTEHGKYLALDLGGTNFRALLVNFKKGLQQNSQLYHKIYTIPLEIMQGSGEELFDHIAQCVS
DFLDYMGMKNARLPAGFTFSFPCEQTAIDTGTLVSWTKGFKATDCEGHDVVSMLREAIKRRNEFDLDIVA
VVNDTVGTMMSCAYEDPQCEIGLIAGTGTNVCYMEELKNIEKTEQKKGQKEREEETEEGEEGKQDDGTER
ERKEGDAEIPKMCINTEWGGLGDDGSLDDIITPYDIEVDQNSLNPGKQRFEKLTSGMYLGEVVRQVLLDL
TRGGLLFRGHVTEALKTPGIFQTKYLSQIESDRLALLQVRSILQQLGLDSTCDDSIIVKEVCGAVSRRAA
QLCGAGMAAVVDKIRENRGLDHLNITVGVDGTLYKLHPHFSGVLQETARVLAPQCNVTFLPSEEGSGKGA
ALITAVARQKLEM
>anc44|avgPP:0.996|lnPP:-5.831|num_ambig:6|num_ambig_gaps:3
LYAMRLHDDQLSDISARFQAEMKKGLSAESNAAAAVKMLPTHVRSTPDGSEKGQFLALDLGGSKFKVLQV
KVREGMGIRRGGVEMEEKTYPIPKELLIGRGTELFDHVSESLKDFLHEKNISLEKKHPLAFTFSFPCEQS
TLNQGLLLNWSKNYRARGLQGKDVVQALRGAIDRTGGMDVEVLAMVNDTVATMMTCGFDDQYCEVGLIIG
TGTNACYMEELRHVDLVEGDEGRMCVNTEWGAFGDDGALNDYITEFDRDVDAASNNPGKQIFEKMVSGMY
LGELVRLVVLKMAKLGLLFDGHVSDALRTKGKITTAHVAAMEEYKNGLKNTRNILMDLDLTPTSDDCVAV
QHVSTIVSFRSSNLVAAGLAAILTRIKQNRNLRTFRITVGVDGTVYKTHPQYPKRLHKVVRRLLPECHVR
FVLSDSGSSKGAALVTAVAQRLASRRRQVDETLSPFRLSQEQLQLVKTRMRAGLEAGLKSKGPSAIKMLP
SFVYHTPDGTEHGKYLALDLGGTNFRALLVNFKKGLQQNSRLYHKIYTIPLEIMQGSGEELFDHIAQCVS
DFLDYMGMKNARLPAGFTFSFPCEQTAIDTGTLVSWTKGFKATDCEGHDVVSMLREAIKRRNEFDLDIVA
VVNDTVGTMMSCAYEDPQCEIGLIAGTGTNVCYMEELKNIEKTEQKTGQKEREEETEEGEEGKQDDGTER
ERKEGDAEIPKMCINTEWGGLGDDGSLDDIITPYDIEVDQNSVNPGKQRFEKLTSGMYLGEVVRQVLLDL
TRGGLLFRGHVTEALKTPGIFQTKYLSQIESDRLALLQVRSILQQLGLDSTCDDSIIVKEVCGAVSRRAA
QLCGAGMAAVVDKIRENRGLDHLNITVGVDGALYKLHPHFSGVLQETARVLAPQCNVTFLPSEEGSGKGA
ALITAVARQKPEM
>anc44_altAll|avgPP:0.995|lnPP:-9.552|num_ambig:6|num_ambig_gaps:3
LYAMRLHDDQLSDISARFQAEMKKGLSAESNAAAAVKMLPTHVRSTPDGSEKGQFLALDLGGSKFKVLQV
KVREGMGIRRGGVEMEEKTYPIPKELLIGRGTELFDHVSESLKDFLHEKNISLEKKHPLAFTFSFPCEQS
TLNQGLLLNWSKNYRARGLQGKDVVQALRGAIDRTGGMDVEVLAMVNDTVATMMTCGFDDQYCEVGLIIG
TGTNACYMEELRHVDLVEGDEGRMCVNTEWGAFGDDGALNDYITEFDRDVDAASNNPGKQIFEKMVSGMY
LGELVRLVVLKMAKLGLLFDGHVSDALRTKGKITTAHVAAMEEYKNGLKNTRNILMDLDLTPTSDDCVAV
QHVSTIVSFRSSNLVAAGLAAILTRIKQNRNLRTFRITVGVDGTVYKTHPQYPKRLHKVVRRLLPECHVR
FVLSDSGSSKGAALVTAVAQRLASRRRQVDETLSPFRLSQEQLQLVKTRMRAGLEAGLKSKGPSAIKMLP
SFVYHTPDGTEHGKYLALDLGGTNFRALLVNFKKGLQQNSQLYHKIYTIPLEIMQGTGEELFDHIAQCVS
DFLDYMGMKNARLPAGFTFSFPCEQTAIDTGTLVSWTKGFKATDCEGHDVVSMLREAIKRRNEFDLDIVA
VVNDTVGTMMSCAYEDPQCEIGLIAGTGTNVCYMEELKNIEKTEQKKGQKEREEETEEGEEGKQDDGTER
EGKEGDAEIPKMCINTEWGGLGDDGSLDDIITPYDIEVDQNSINPGKQRFEKLTSGMYLGEVVRQVLLDL
TRGGLLFRGHVTEALKTPGIFQTKYLSQIESDRLALLQVRSILQQLGLDSTCDDSIIVKEVCGAVSRRAA
QLCGAGMAAVVDKIRENRGLDHLNITVGVDGALYKLHPHFSGVLQETARVLAPQCNVTFLPSEEGSGKGA
ALITAVARQKLEM
>anc45|avgPP:0.975|lnPP:-39.496|num_ambig:33|num_ambig_gaps:3

LYAMRLNDDQLSDISARFQAEMKKGLSAESNAAAAVKMLPTHVRSTPDGSEKGQFLALDLGGSKFKVLQV
KVREGRGIRKGGVEMEEKTYPIPKELLTGRGTELFDHVSESLKDFLHEKNISLEKKHPLAFTFSFPCEQP
TLDQGFLLSWSKNYRVRGLQGKDVVQALRQAIDRTGGMDVEVLAMVNDTVATMMTCGFDDQYCEVGLIIG
TGTNACYMEELRHVDLVEGDEGRMCVNTEWGAFGDDGALNDFITEFDRDVDAASINPGKQIFEKMVSGMY
LGELVRLVVLKMAKLGLLFDGHVSDALRTKGKITTAHVAAMEEYKNGLKNTRDILMDLDLSPSPDDCIAV
QHVSTIVSFRSSNLVAAGLAAILTRIKKNRNVRGLRITIGVDGTVYKTHPQYPKRLHKVVRRLLPECHVR
FVLSESGSSKGAALVAAVAQRLASRKRQVDETLSPFRLSQEQLQLVKARMRAGLEAGLKSKGPSAVKMLP
SFVYHTPDGTEHGKYLALDLGGTNFRALLVNFKKGLQQNSRLYHKIYTIPLEIMQGTGEELFDYLAQCVG
DFLDYMGMKNARLPAGFTFSFPCEQTAIDTGTLVSWTKGFKATDCEGHDVVNMLREAIKRRNEFDLDIVA
LVNDTVGTMMSCAYEDPQCEVALIAGTGSNVCYMEELKNIEKIKQKSGQKERDEDTEEGEEGKEDDGERE
GKEGDADISKMCINTEWGGLGDDGSLDDIITSYDTEVDQNSINPGKQRFEKLTSGMYLGEIVRQVLLDLT
RRGLLFRGHVTEALKTPGIFETKYLSQIESDRLALLQVRSILQQLGLDSTCDDSIIVKEVCGAISRRAAQ
LCGAGMAAVVDKIRENRGLDRLNITVGVDGALYKLHPHFSQVLQETTRVLAPQCDVTFLPSEEGSGKGAA
LITAVAKQKQE
>anc45_altAll|avgPP:0.969|lnPP:-56.149|num_ambig:33|num_ambig_gaps:3
LHAMRLNDDQLSDISARFQAEMKKGLSAESNAAAAVKMLPTHVRSTPDGSEKGQFLALDLGGSKFKVLQV
KVREGRGIRKGGVEMEEKIYPIPKELLSGRGAELFDHVSESLKDFLHEKNISLEKKHPLAFTFSFPCEQP
SLDQGFLLSWSKNYRVRGLQGKDVVHALREAIDRTGGMDVEVLAMVNDTVATMMTCGFDDQYCEVGLIIG
TGTNACYMEELRHVDLVEGDEGRMCVNTEWGAFGDDGALDDYITEFDRDVDAASVNPGKQIFEKMVSGMY
LGELVRLVVLKMAKLGLLFDGHVSDALRTKGKITTAHVAAMEEYKNGLKNTREILMGLDLTPSPDDCIAV
QHVSTIVSFRSSNLVAAGLAAILTRIKKNRNVRGLRITIGVDGTVYKTHPQYPKRLHKVVRRLLPECHVR
FVLSESGSSKGAALVTAVAQRLASRKRQVDETLSPFRLSQEQLQLVKARMRAGLEAGLKSKGPSAIKMLP
SFVYHTPDGTEHGKYLALDLGGTNFRALLVNFKKGLQQNSRLYHKIYTIPLEIMQGTGEELFDYLAQCVG
DFLDYMGMKNARLPAGFTFSFPCEQTAIDTGTLVSWTKGFKATDCEGHDVVSMLREAIKRRNEFDLDIVA
LVNDTVGTMMSCAYEDPQCEIALIAGTGSNVCYMEELKNIEKIKQQTGQKEGDEETEEGEEGKEDDGERK
GTNGDAEIPKMCINTEWGGLGDDGSLDDIITPYDTEVDQNSINPGKQRFEKLTSGMYLGEVVRQVLLDLT
RRGLLFRGHVTETLKTPGIFETKYLSQIESDRLALLQVRSILQQLGLDSTCDDSIIVKEVCGAISRRAAQ
LCGAGMAAVVDKIRENRGLDHLNITVGVDGALYKLHPHFSRVLQETTRVLAPQCDVTFLPSEEGSGKGAA
LIAAVARQKQE
>anc46|avgPP:0.980|lnPP:-32.933|num_ambig:29|num_ambig_gaps:4
LHAMRLNDDQLSDISARFQAEMKKGLSAESNAAAAVKMLPTHVRSTPDGSEKGQFLALDLGGSKFKVLQV
KVREGVGIRKGEVEMEEKIYPIPKELQTGRGAELFDHVSESLKDFLHEKNISLEKKHPLAFTFSFPCEQP
SLDQGLLLSWCKNYRARGLQGKDVVQALRDAISRAGDMDMEVLAMVNDTVATMMTCGFDDQYCEVGLIIG
TGTNACYMEELRHVDLVEGDEGRMCVNTEWGAFGDDGALNDFITEFDRDVDAASSNPGKQIFEKMVSGMY
LGELVRLVVLKMAKLGLLFDGHVSDALRTKGKITTAHVAAMEEYRNGLKNTKDILMQLDLTPSPDDCIAV
QHVSTIVSFRSSNLVAAGLAAILTRIKQNRNVRGLRITVGVDGTVYKTHPQYPKRLHKVVRRLLPECHVR
FVLSDSGSSKGAALVTAVAQRLTSQSRKVDETLSPFRLSQEQLQLVKARMRAGLEAGLKSQGPFTIKMLP
SFVYHTPDGTEHGKYLALDLGGTNFRALLVRFKKGKQKSRLYHKIYTIPLEIMQGTGEELFDHLAQCVCD
FLDYMGMKNARLPAGFTFSFPCEQTAIDTGTLVSWTKGFKATDCEGHDVVSMLREAIKRRNEFDLDIVAL
VNDTVGTMMSCAYEDPQCEIGLIAGTGSNLCYMEELKNIEKIEQKSGTTETDEETPEGEEGKEDDGTEAE
KTEEDAEIPKMCINTEWGGLGDDGSLDDIITAFDTEVDQNSVNPGKQRFEKLTSGMYLGEVVRQVLLDLT
RRGLLFRGHVTETLKKPGIFETKYLSQIESDHLALLQVRSILQKLGLDSTCDDSIIVKEVCGAVSRRAAQ
LCGAGMAAVVDKIRENRGLDHLNITVGVDGALYKLHPHFSQILQETVKVMAPQCDVTFLPSEEGSGKGAA
LIAAVARQKQEVTM
>anc46_altAll|avgPP:0.975|lnPP:-44.575|num_ambig:29|num_ambig_gaps:4
LHAMRLNDDQLSDISARFQAEMKKGLSAESNAAAAVKMLPTHVRSTPDGSEKGQFLALDLGGSKFKVLQV
KVREGVGIKKGEVEMEEKIYPIPKELLTGRGTELFDHVSESLKDFLHEKNISLEKKHPLAFTFSFPCEQP
SLDQGLLLSWCKNYRVRGLQGKDVVQALREAISRAGDMDMEVLAMVNDTVATMMTCGFDDQYCEVGLIIG
TGTNACYMEELRHVDLVEGDEGRMCVNTEWGAFGDDGALDDFITEFDRDVDAASSNPGKQIFEKMVSGMY
LGELVRLVVLKMAKLGLLFDGHVSDALRTKGKITTADVAAMEEYRNGLKNTKDILMHLDLSPSPDDCIAV
QHVSTIVSFRSSNLVAAGLAAILTRIRQNRNIRALRITVGVDGTVYKTHPQYPKRLHKVVRRLLPECQVR
FVLSDSGSSKGAALVAAVAQRLASQSRKVDETLSAFRLSQEQLQLVKARMRAGLEAGLKSQGPFTIKMLP

SFVYHTPDGTEHGKYLALDLGGTNFRALLVRFKGGKQKSRLYHKIYTIPLEIMQGTGEELFDHLAQCVCD
FLDYMGMKNARLPAGFTFSFPCEQTAIDTGTLVSWTKGFKATDCEGHDVVSMLREAIKRRNEFDLDIVAL
VNDTVGTMMSCAYEDPQCEIGLIAGTGSNLCYMEELKNIEKIKQRSGTTETDEETQEGEEGKEDDGTEAE
KNEGDAEISKMCINTEWGGLGDDGSLDDIITPFDTEVDQNSVNPGKQRFEKLTSGMYLGEIVRQVLLDLT
RRGLLFRGHVTETLKRPGIFETKYLSQIESDRLALLQVRSILQKLGLDSTCDDSIIVKEVCGAVSRRAAQ
LCGAGMAAVVDKIRENRGLDHLNITVGVDGALYKLHPHFSQVLQETVRVMAPQCDVTFLPSEEGSGKGAA
LIAAVARQKQEVTM
>anc47|avgPP:0.989|lnPP:-17.345|num_ambig:17|num_ambig_gaps:0
LHAMRLQDDQLSDISKRFLAEMKKGLSAESTAAAAVKMLPTHVRSTPDGSEKGQFLALDLGGSKFKVLRV
KVREDVGIKKGGVEMEEKIYPIPKELHAGRAAELFDHVSESLKDFLHEKNISPEKKHPLAFTFSFPCEHP
SLDQGLLVSWCKNYKVRGLQGKDVVQALRQAIGRTGDMDVEVLAMVNDTVATMMTCGFDDQCCEVGLIIG
TGTNACYMEELRHIDLVEGDEGRMCVNTEWGAFGDDGALDDFITEFDRDVDAASLNPGKQIFEKMVSGMY
LGELVRLVVLKMAKLGLLFDGHVSDALRTKGKITTADVAAMEEYRNGLKNTKDILTRLHLTPSPDDCIAV
QHVSTIVSFRSSNLVAAGLAAILTRIRQNRNLRGLRITVGVDGTVYKTHPQYPKRLHKVVRRLLPECQVR
FVLSESGSSKGAALVAAVAQRLTSQRRKVDEALSLFTLNQQQLQLVKSRMREGLEAGLKSKGCTVKMLPS
FVYHTPDGTEHGKYLALDLGGTNFRAMLVRFKRQKSRLYHKIYTIPLEIMQGTGEELFDHLAQCVCDFLD
YMGLKKARLPAGFTFSFPCEQSGIDTGTLVSWTKGFKATDCEGQDVVSMLREAIKRRNEFDLDIVALVND
TVGTMMSCAYEDPQCEIGMIAGTGSNLCYMEELKNIEKIKQGNGITKTDKETLEGEEDDDATEAEKSAVE
ADKSKMCINTEWGGLGDDRCLDDIITPYDAEVDIHSVNPGKQRFEKLTSGMYLGEIVRQVLLDLTRRGLL
FKGHVTETLKTPGIFETKYLSQIESDRLALLQVRSILQQLGLRGTCDDSIIVKEVCGAVSRRAAQLCGAG
MAAVVDKIREDRRLERLSITVGVDGALYKLHPHFSQVLKETVRVMAPQCDVTFLPSEEGSGKGAALIAAV
VRQKQEM
>anc47_altAll|avgPP:0.986|lnPP:-26.622|num_ambig:17|num_ambig_gaps:0
LHAMRLQDDQLSDISKRFLAEMKKGLSAESTAAAAVKMLPTHVRSTPDGSEKGQFLALDLGGSKFKVLRV
KVREDVGIRKGGVEMEEKTYPIPKELHAGRAAELFDHVSESLKDFLHEKNISLEKKHPLAFTFSFPCEHP
SLDQGLLVSWCKNYKVRGLQGKDVVQALREAIGRTGDMDVEVLAMVNDTVATMMTCGFDDQCCEVGLIIG
TGTNACYMEELRHIDLVEGDEGRMCVNTEWGAFGDDGALDDFITEFDRDVDAASLNPGKQIFEKMVSGMY
LGELVRLVVLKMAKLGLLFDGRVSDALRTKGKITTADVAAMEEYRNGLKNTKDILTHLHLSPSPDDCIAV
QHVSTIVSFRSSNLVAAGLAAILTRIRQNRNVRGLRITVGVDGTVYKTHPQYPKRLHKVVRRLLPDCQVR
FVLSESGSSKGAALVAAVAQRLTSQRRKVDEALSLFRLNQQQLQLVKSRMREGLEAGLKSKGCTVKMLPS
FVYHTPDGTEHGKYLALDLGGTNFRAMLVRFKRQKSRLYHKIYTIPLEIMQGTGEELFDHLAQCVCDFLD
YMGLKKARLPAGFTFSFPCEQSGIDTGTLVSWTKGFKATDCEGQDVVSMLREAIKRRNEFDLDIVALVND
TVGTMMSCAYEDPQCEIGMIAGTGSNLCYMEELKNIEKIKQGNGTTKTDKEALEGEEDDDVTEAEKSAVE
AGESKMCINTEWGGLGDDRCLDDIITPYDAEVDIHSVNPGKQRFEKLTSGMYLGEIVRQVLLDLTRRGLL
FKGHVTETLKTPGIFETKYLSQIESDRLALLQVRSILQHLGLRGTCDDSIIVKEVCGAVSRRAAQLCGAG
MAAVVDKIREDRRLERLSITVGVDGALYKLHPHFSQVLKETVRVMAPQCDVTFLPSEEGSGKGAALIAAV
VQQKQEM
>anc48|avgPP:0.987|lnPP:-20.569|num_ambig:17|num_ambig_gaps:2
LHAMRLQDDQLSDISARFLAEMKKGLSAESTAAAAVKMLPTHVRSTPDGSEKGQFLALDLGGSKFKVLRV
KVREDVGIKKGGVEMEEKIYPIPKELHAGRAAELFDHVSESLKDFLHEKNISPEKKHPLAFTFSFPCEHP
SLDQGLLLSWCKNYKVRGLQGKDVVQALRQAIDRTGDMDVEVLAMVNDTVATMMTCGFDDQYCEVGLIIG
TGTNACYMEELRHIDLVEGDEGRMCVNTEWGAFGDDGALDDFITEFDRDVDAASLNPGKQIFEKMVSGMY
LGELVRLVVLKMAKLGLLFDGHVSDALRTKGKITTADVAAMEEYRNGLKNTKDILTRLHLTPSPDDCIAV
QHVSTIVSFRSSNLVAAGLAAILTRIRQNRNLRGLRITVGVDGTVYKTHPQYPKRLHKVVRRLLPECQVR
FVLSESGSSKGAALVAAVAQRLTSQRRKVDEALSLFTLNQEQLQLVKSRMREGLEAGLKSKGCTIKMLPS
FVYHTPDGTEHGKYLALDLGGTNFRAMLVRFKKQKSRLYHKIYTIPLEIMQGTGEELFDHLAQCVCDFLD
YMGLKKARLPAGFTFSFPCEQSGIDTGTLVSWTKGFKATDCEGQDVVSMLREAIKRRNEFDLDIVALVND
TVGTMMSCAYEDPQCEIGMIAGTGSNLCYMEELKNIEKIEQGNGITKTDKETLEGEEGDDATEAEKSAAE
ADKSKMCINTEWGGLGDDRCLDDIITPYDAEVDIHSVNPGKQRFEKLTSGMYLGEIVRQVLLDLTRRGLL
FKGHVTETLKTPGIFETKYLSQIESDRLALLQVRSILQQLGLHGTCDDSIIVKEVCGAVSRRAAQLCGAG
MAAVVDKIREDRGLDRLSITVGVDGALYKLHPHFSQVLKETVRVMAPQCDVTFLPSEEGSGKGAALIAAV
VRQKQEM

```
>anc48_altAll|avgPP:0.984|lnPP:-28.913|num_ambig:17|num_ambig_gaps:2
LHAMRLQDDQLSDISSRFLAEMKKGLSAESTAAAAVKMLPTHVRSTPDGSEKGQFLALDLGGSKFKVLRV
KVREDVGIRKGGVEMEEKTYPIPKELHAGRAAELFDHVSESLKDFLHEKNISLEKKHPLAFTFSFPCEHP
SLDQGLLLSWCKNYKVRGLQGKDVVQALREAIDRTGDMDVEVLAMVNDTVATMMTCGFDDQCCEVGLIIG
TGTNACYMEELRHIDLVEGDEGRMCVNTEWGAFGDDGALDDFITEFDRDVDAASLNPGKQIFEKMVSGMY
LGELVRLVVLKMAKLGLLFDGHVSDALRTKGKITTADVAAMEEYRNGLKNTKDILTHLHLSPSPDDCIAV
QHVSTIVSFRSSNLVAAGLAAILTRIRQNRNVRGLRITVGVDGTVYKTHPQYPKRLHKVVRRLLPDCQVR
FVLSESGSSKGAALVAAVAQRLTSQRRKVDEALSLFRLNQEQLQLVKSRMREGLEAGLKSKGCTIKMLPS
FVYHTPDGTEHGKYLALDLGGTNFRAMLVRFKKQKSRLYHKIYTIPLEIMQGTGEELFDHLAQCVCDFLD
YMGLKKARLPAGFTFSFPCEQSGIDTGTLVSWTKGFKATDCEGQDVVSMLREAIKRRNEFDLDIVALVND
TVGTMMSCAYEDPQCEIGMIAGTGSNLCYMEELKNIEKIKQGNGTTKTDKETLEGEEGDDATEAEKSAVE
AEESKMCINTEWGGLGDDRCLDDIITPYDAEVDIHSVNPGKQRFEKLTSGMYLGEIVRQVLLDLTRRGLL
FKGHVTETLKTPGIFETKYLSQIESDRLALLQVRSILQQLGLHGTCDDSIIVKEVCGAVSRRAAQLCGAG
MAAVVDKIREDRGLDRLSITVGVDGALYKLHPHFSQVLKETVRVMAPQCDVTFLPSEEGSGKGAALIAAV
VQQKQEM
>anc49|avgPP:0.985|lnPP:-23.747|num_ambig:19|num_ambig_gaps:3
LHAMRLNDDQLSDISARFQAEMKKGLSAESNAAAAVKMLPTHVRSTPDGSEKGQFLALDLGGSKFKVLQV
KVREGVGIKKGGVEMEEKIYPIPKELLTGRAAELFDHVSESLKDFLHEKNISLEKKHPLAFTFSFPCEQP
SLDQGLLLSWCKNYKARGLQGKDVVQALREAIDRTGDMDVEVLAMVNDTVATMMTCGFDDQYCEVGLIIG
TGTNACYMEELRHIDLVEGDEGRMCVNTEWGAFGDDGALDDFITEFDRDVDAASSNPGKQIFEKMVSGMY
LGELVRLVVLKMAKLGLLFDGHVSDALRTKGKITTADVAAMEEYRNGLKNTKDILTHLDLTPSPDDCIAV
QHVSTIVSFRSSNLVAAGLAAILTRIRQNRNLRGLRITVGVDGTVYKTHPQYPKRLHKVVRRLLPECQVR
FVLSESGSSKGAALVAAVAQRLTSQRRKVDETLSLFRLNQEQLQLVKARMRAGLEAGLKSKGSSTIKMLP
SFVYHTPDGTEHGKYLALDLGGTNFRALLVRFKKQKSRLYHKIYTIPLEIMQGTGEELFDHLAQCVCDFL
DYMGLKNARLPAGFTFSFPCEQSAIDTGTLVSWTKGFKATDCEGHDVVSMLREAIKRRNEFDLDIVALVN
DTVGTMMSCAYEDPQCEIGMIAGTGSNLCYMEELKNIEKIEQKNGTTKTDEETQEGEEGDDGTEAEKNEG
DAEIPKMCINTEWGGLGDDGCLDDIITPYDTEVDQNSVNPGKQRFEKLTSGMYLGEVVRQVLLDLTRRGL
LFKGHVTETLKTPGIFETKYLSQIESDRLALLQVRSILQQLGLHGTCDDSIIVKEVCGAVSRRAAQLCGA
GMAAVVDKIRENRGLDHLNITVGVDGALYKLHPHFSQVLKETVRVMAPQCDVTFLPSEEGSGKGAALIAA
VARQKQEM
>anc49_altAll|avgPP:0.982|lnPP:-31.673|num_ambig:19|num_ambig_gaps:3
LHAMRLNDDQLSDISARFQAEMKKGLSAESNAAAAVKMLPTHVRSTPDGSEKGQFLALDLGGSKFKVLQV
KVREGVGIRKGGVEMEEKIYPIPKELLTGRATELFDHVSESLKDFLHEKNISLEKKHPLAFTFSFPCEQP
SLDQGLLLSWCKNYKVRGLQGKDVVQALREAIDRTGGMDVEVLAMVNDTVATMMTCGFDDQYCEVGLIIG
TGTNACYMEELRHIDLVEGDEGRMCVNTEWGAFGDDGALNDFITEFDRDVDAASSNPGKQIFEKMVSGMY
LGELVRLVVLKMAKLGLLFDGHVSDALRTKGKITTADVAAMEEYRNGLKNTKDILMQLDLSPSPDDCIAV
QHVSTIVSFRSSNLVAAGLAAILTRIRQNRNVRALRITVGVDGTVYKTHPQYPKRLHKVVRRLLPECQVR
FVLSESGSSKGAALVAAVAQRLTSQRRQVDETLSLFRLSQEQLQLVKARMRAGLEAGLKSKGSSTIKMLP
SFVYHTPDGTEHGKYLALDLGGTNFRALLVRFKKQKSRLYHKIYTIPLEIMQGTGEELFDHLAQCVCDFL
DYMGLKNARLPAGFTFSFPCEQSAIDTGTLVSWTKGFKATDCEGHDVVSMLREAIKRRNEFDLDIVALVN
DTVGTMMSCAYEDPQCEIGMIAGTGSNLCYMEELKNIEKIKQRNGTTKTDEETPEGEEGDDGTEAEKSEG
DAEMSKMCINTEWGGLGDDGCLDDIITPYDTEVDQNSVNPGKQRFEKLTSGMYLGEIVRQVLLDLTRRGL
LFKGHVTETLKTPGIFETKYLSQIESDRLALLQVRSILQQLGLHGTCDDSIIVKEVCGAVSRRAAQLCGA
GMAAVVDKIRENRGLDHLNITVGVDGALYKLHPHFSQVLKETVRVMAPQCDVTFLPSEEGSGKGAALIAA
VARQKQEM
>anc50|avgPP:0.985|lnPP:-23.867|num_ambig:23|num_ambig_gaps:6
LHAMRLNDDQLSDISARFQAEMKKGLSAESNAAAAVKMLPTHVRSTPDGSEKGQFLALDLGGSKFKVLQV
KVREGVGIRKGGVEMEEKIYPIPKELLTGRGAELFDHVSESLKDFLHEKNISLEKKHPLAFTFSFPCEQP
SLDQGLLLSWCKNYRARGLQGKDVVQALREAIDRTGDMDVEVLAMVNDTVATMMTCGFDDQYCEVGLIIG
TGTNACYMEELRHVDLVEGDEGRMCVNTEWGAFGDDGALNDFITEFDRDVDAASSNPGKQIFEKMVSGMY
LGELVRLVVLKMAKLGLLFDGHVSDALRTKGKITTAHVAAMEEYRNGLKNTKDILMHLDLTPSPDDCIAV
QHVSTIVSFRSSNLVAAGLAAILTRIKQNRNLRGLRITVGVDGTVYKTHPQYPKRLHKVVRRLLPECHVR
```

FVLSDSGSSKGAALVAAVAQRLTSQRRKVDETLSPFRLSQEQLQLVKARMRAGLEAGLKSKGPSTIKMLP
SFVYHTPDGTEHGKYLALDLGGTNFRALLVRFKKGKQKSRLYHKIYTIPLEIMQGTGEELFDHLAQCVCD
FLDYMGMKNARLPAGFTFSFPCEQTAIDTGTLVSWTKGFKATDCEGHDVVSMLREAIKRRNEFDLDIVAL
VNDTVGTMMSCAYEDPQCEIGLIAGTGSNLCYMEELKNIEKIEQKSGTTETDEETQEGEEGKEDDGTEAE
KNEGDAEIPKMCINTEWGGLGDDGSLDDIITPYDTEVDQNSVNPGKQRFEKLTSGMYLGEVVRQVLLDLT
RRGLLFRGHVTETLKTPGIFETKYLSQIESDRLALLQVRSILQQLGLDSTCDDSIIVKEVCGAVSRRAAQ
LCGAGMAAVVDKIRENRGLDHLNITVGVDGALYKLHPHFSQVLQETVRVMAPQCDVTFLPSEEGSGKGAA
LIAAVARQKQEM
>anc50_altAll|avgPP:0.982|lnPP:-31.313|num_ambig:23|num_ambig_gaps:6
LHAMRLNDDQLSDISARFQAEMKKGLSAESNAAAAVKMLPTHVRSTPDGSEKGQFLALDLGGSKFKVLQV
KVREGVGIKKGGVEMEEKIYPIPKELLTGRGTELFDHVSESLKDFLHEKNISLEKKHPLAFTFSFPCEQP
SLDQGLLLSWCKNYRVRGLQGKDVVQALREAIDRTGGMDVEVLAMVNDTVATMMTCGFDDQYCEVGLIIG
TGTNACYMEELRHVDLVEGDEGRMCVNTEWGAFGDDGALDDFITEFDRDVDAASSNPGKQIFEKMVSGMY
LGELVRLVVLKMAKLGLLFDGHVSDALRTKGKITTADVAAMEEYRNGLKNTKDILMQLDLSPSPDDCIAV
QHVSTIVSFRSSNLVAAGLAAILTRIRQNRNVRALRITVGVDGTVYKTHPQYPKRLHKVVRRLLPECQVR
FVLSESGSSKGAALVTAVAQRLASQRRQVDETLSPFRLSQEQLQLVKARMRAGLEAGLKSKGPSTIKMLP
SFVYHTPDGTEHGKYLALDLGGTNFRALLVRFKKGKQKSRLYHKIYTIPLEIMQGTGEELFDHLAQCVCD
FLDYMGMKNARLPAGFTFSFPCEQTAIDTGTLVSWTKGFKATDCEGHDVVSMLREAIKRRNEFDLDIVAL
VNDTVGTMMSCAYEDPQCEIGLIAGTGSNLCYMEELKNIEKIKQRNGTTETDEETPEGEEGKEDDGTEAE
KTEGDAEISKMCINTEWGGLGDDGSLDDIITPYDTEVDQNSVNPGKQRFEKLTSGMYLGEIVRQVLLDLT
RRGLLFRGHVTETLKTPGIFETKYLSQIESDRLALLQVRSILQQLGLDSTCDDSIIVKEVCGAVSRRAAQ
LCGAGMAAVVDKIRENRGLDHLNITVGVDGALYKLHPHFSQVLQETVRVMAPQCDVTFLPSEEGSGKGAA
LIAAVARQKQEM
>anc51|avgPP:0.985|lnPP:-23.318|num_ambig:23|num_ambig_gaps:4
LYAMRLNDDQLSDISARFQAEMKKGLSAESNAAAAVKMLPTHVRSTPDGSEKGQFLALDLGGSKFKVLQV
KVREGMGIRKGGVEMEEKTYPIPKELLTGRGTELFDHVSESLKDFLHEKNISLEKKHPLAFTFSFPCEQP
TLDQGLLLSWSKNYRARGLQGKDVVQALREAIDRTGGMDVEVLAMVNDTVATMMTCGFDDQYCEVGLIIG
TGTNACYMEELRHVDLVEGDEGRMCVNTEWGAFGDDGALNDFITEFDRDVDAASNNPGKQIFEKMVSGMY
LGELVRLVVLKMAKLGLLFDGHVSDALRTKGKITTAHVAAMEEYKNGLKNTRDILMDLDLTPSPDDCIAV
QHVSTIVSFRSSNLVAAGLAAILTRIKQNRNLRGLRITVGVDGTVYKTHPQYPKRLHKVVRRLLPECHVR
FVLSDSGSSKGAALVAAVAQRLASQRRQVDETLSPFRLSQEQLQLVKARMRAGLEAGLKSKGPSAIKMLP
SFVYHTPDGTEHGKYLALDLGGTNFRALLVNFKKGLQQNSRLYHKIYTIPLEIMQGTGEELFDHLAQCVS
DFLDYMGMKNARLPAGFTFSFPCEQTAIDTGTLVSWTKGFKATDCEGHDVVSMLREAIKRRNEFDLDIVA
LVNDTVGTMMSCAYEDPQCEIGLIAGTGSNVCYMEELKNIEKIEQKTGQKERDEETEEGEEGKEDDGTER
EGKEGDAEIPKMCINTEWGGLGDDGSLDDIITPYDTEVDQNSVNPGKQRFEKLTSGMYLGEVVRQVLLDL
TRRGLLFRGHVTEALKTPGIFETKYLSQIESDRLALLQVRSILQQLGLDSTCDDSIIVKEVCGAVSRRAA
QLCGAGMAAVVDKIRENRGLDHLNITVGVDGALYKLHPHFSQVLQETTRVLAPQCDVTFLPSEEGSGKGA
ALITAVARQKQEM
>anc51_altAll|avgPP:0.981|lnPP:-34.068|num_ambig:23|num_ambig_gaps:4
LHAMRLNDDQLSDISARFQAEMKKGLSAESNAAAAVKMLPTHVRSTPDGSEKGQFLALDLGGSKFKVLQV
KVREGMGIRKGGVEMEEKIYPIPKELLTGRGAELFDHVSESLKDFLHEKNISLEKKHPLAFTFSFPCEQP
SLDQGLLLSWSKNYRVRGLQGKDVVQALREAIDRTGGMDVEVLAMVNDTVATMMTCGFDDQYCEVGLIIG
TGTNACYMEELRHVDLVEGDEGRMCVNTEWGAFGDDGALDDYITEFDRDVDAASSNPGKQIFEKMVSGMY
LGELVRLVVLKMAKLGLLFDGHVSDALRTKGKITTAHVAAMEEYKNGLKNTRDILMDLDLSPSPDDCIAV
QHVSTIVSFRSSNLVAAGLAAILTRIKQNRNVRALRITVGVDGTVYKTHPQYPKRLHKVVRRLLPECHVR
FVLSESGSSKGAALVTAVAQRLASRRRQVDETLSPFRLSQEQLQLVKARMRAGLEAGLKSKGPSAIKMLP
SFVYHTPDGTEHGKYLALDLGGTNFRALLVNFKKGLQQNSRLYHKIYTIPLEIMQGTGEELFDHLAQCVS
DFLDYMGMKNARLPAGFTFSFPCEQTAIDTGTLVSWTKGFKATDCEGHDVVSMLREAIKRRNEFDLDIVA
LVNDTVGTMMSCAYEDPQCEIGLIAGTGSNVCYMEELKNIEKIKQKTGQKERDEETEEGEEGKEDDGTEA
EGTEGDAEISKMCINTEWGGLGDDGSLDDIITPYDTEVDQNSINPGKQRFEKLTSGMYLGEIVRQVLLDL
TRRGLLFRGHVTETLKTPGIFETKYLSQIESDRLALLQVRSILQQLGLDSTCDDSIIVKEVCGAVSRRAA
QLCGAGMAAVVDKIRENRGLDHLNITVGVDGALYKLHPHFSRVLQETTRVLAPQCDVTFLPSEEGSGKGA

ALIAAVARQKQEM
>anc52|avgPP:0.992|lnPP:-11.480|num_ambig:13|num_ambig_gaps:4
LYAMRLHDDQLSDISARFQAEMKKGLSAESNAAAAVKMLPTHVRSTPDGSEKGQFLALDLGGSKFKVLQV
KVREGMGIRRGGVEMEEKTYPIPKELLIGRGTELFDHVSESLKDFLHEKNISLEKKHPLAFTFSFPCEQS
TLNQGLLLNWSKNYRARGLQGKDVVQALREAIDRTGGMDVEVLAMVNDTVATMMTCGFDDQYCEVGLIIG
TGTNACYMEELRHVDLVEGDEGRMCVNTEWGAFGDDGALNDYITEFDRDVDAASNNPGKQIFEKMVSGMY
LGELVRLVVLKMAKLGLLFDGHVSDALRTKGKITTAHVAAMEEYKNGLKNTRDILMDLDLTPSPDDCIAV
QHVSTIVSFRSSNLVAAGLAAILTRIKQNRNLRTLRITVGVDGTVYKTHPQYPKRLHKVVRRLLPECHVR
FVLSDSGSSKGAALVTAVAQRLASQRRQVDETLSPFRLSQEQLQLVKARMRAGLEAGLKSKGPSAIKMLP
SFVYHTPDGTEHGKYLALDLGGTNFRALLVNFKKGLQQNSRLYHKIYTIPLEIMQGTGEELFDHIAQCVS
DFLDYMGMKNARLPAGFTFSFPCEQTAIDTGTLVSWTKGFKATDCEGHDVVSMLREAIKRRNEFDLDIVA
VVNDTVGTMMSCAYEDPQCEIGLIAGTGSNVCYMEELKNIEKIEQKTGQKEREEETEEGEEGKEDDGTER
EGKEGDAEIPKMCINTEWGGLGDDGSLDDIITPYDIEVDQNSVNPGKQRFEKLTSGMYLGEVVRQVLLDL
TRGGLLFRGHVTEALKTPGIFQTKYLSQIESDRLALLQVRSILQQLGLDSTCDDSIIVKEVCGAVSRRAA
QLCGAGMAAVVDKIRENRGLDHLNITVGVDGALYKLHPHFSRVLQETTRVLAPQCDVTFLPSEEGSGKGA
ALITAVARQKQEM
>anc52_altAll|avgPP:0.990|lnPP:-18.836|num_ambig:13|num_ambig_gaps:4
LYAMRLHDDQLSDISARFQAEMKKGLSAESNAAAAVKMLPTHVRSTPDGSEKGQFLALDLGGSKFKVLQV
KVREGMGIRRGGVEMEEKTYPIPKELLIGRGTELFDHVSESLKDFLHEKNISLEKKHPLAFTFSFPCEQS
TLNQGLLLNWSKNYRARGLQGKDVVQALREAIDRTGGMDVEVLAMVNDTVATMMTCGFDDQYCEVGLIIG
TGTNACYMEELRHVDLVEGDEGRMCVNTEWGAFGDDGALDDFITEFDRDVDAASNNPGKQIFEKMVSGMY
LGELVRLVVLKMAKLGLLFDGHVSDALRTKGKITTAHVAAMEEYKNGLKNTRDILMDLDLTPSPDDCIAV
QHVSTIVSFRSSNLVAAGLAAILTRIKQNRNLRTLRITVGVDGTVYKTHPQYPKRLHKVVRRLLPECHVR
FVLSDSGSSKGAALVAAVAQRLASRRRQVDETLSPFRLSQEQLQLVKARMRAGLEAGLKSKGPSAIKMLP
SFVYHTPDGTEHGKYLALDLGGTNFRALLVNFKKGLQQNSRLYHKIYTIPLEIMQGTGEELFDHIAQCVS
DFLDYMGMKNARLPAGFTFSFPCEQTAIDTGTLVSWTKGFKATDCEGHDVVSMLREAIKRRNEFDLDIVA
VVNDTVGTMMSCAYEDPQCEIGLIAGTGSNVCYMEELKNIEKIKQKKGQKERDEETEEGEEGKEDDGTER
ERKEGDAEIPKMCINTEWGGLGDDGSLDDIITPYDTEVDQNSINPGKQRFEKLTSGMYLGEVVRQVLLDL
TRRGLLFRGHVTEALKTPGIFETKYLSQIESDRLALLQVRSILQQLGLDSTCDDSIIVKEVCGAVSRRAA
QLCGAGMAAVVDKIRENRGLDHLNITVGVDGALYKLHPHFSRVLQETTRVLAPQCNVTFLPSEEGSGKGA
ALITAVARQKQEM
>anc53|avgPP:0.989|lnPP:-17.041|num_ambig:16|num_ambig_gaps:7
LYAMRLHDDQLSDISARFQAEMKKGLSAESNAAAAVKMLPTHVRSTPDGSEKGQFLALDLGGSKFKVLQV
KVREGMGIRRGGVEMEEKTYPIPKELLIGRGTELFDHVSESLKDFLHEKNISLEKKHPLAFTFSFPCEQS
TLNQGLLLNWSKNYRARGLQGKDVVQALREAIDRTGGMDVEVLAMVNDTVATMMTCGFDDQYCEVGLIIG
TGTNACYMEELRHVDLVEGDEGRMCVNTEWGAFGDDGALNDYITEFDRDVDAASNNPGKQLFEKMVSGMY
LGELVRLVVLKMAKLGLLFDGHVSDALRTKGKITTAHVAAMEEYKNGLKNTRDILMDLDLTPSPDDCIAV
QHVSTIVSFRSSNLVAAGLAAILTRIKQNRNLRTLRITVGVDGTVYKTHPQYPKRLHKVVRRLLPECHVR
FVLSDSGSSKGAALVTAVAQRLASQRRQVDETLSPFRLSQEQLQLVKARMRAGLEAGLKSKGPSAIKMLP
SFVYHTPDGTEHGKYLALDLGGTNFRALLVNFKKGLQQNSRLYHKIYTIPLEIMQGTGEELFDHIVQCIS
DFLDYMGMKNARLPAGFTFSFPCEQTAIDTGTLVSWTKGFKATDCEGHDVVSMLREAIKRRNEFDLDIVA
VVNDTVGTMMSCAYEDPQCEIGLIAGTGSNVCYMEELKNIEKIKQKTGQKEREEETEEGEEGAMEKEDDG
TEREGKEGDAEIPRMCINTEWGGLGDDGSLDDIITPYDIEVDQNSVNPGKQRFEKLTSGMYLGEVVRQVL
LDLTRGGLLFRGHVTEALKTPGIFQTKYLSQIESDRLALLQVRSILQQLGLDSTCDDSIIVKEVCGAVSR
RAAQLCGAGMAAVVDKIRENRGLDHLNITVGVDGALYKLHPHFSRVLQETTRVLAPQCDVTFLPSEEGSG
KGAALITAVARQKQEM
>anc53_altAll|avgPP:0.986|lnPP:-25.495|num_ambig:16|num_ambig_gaps:7
LYAMRLHDDQLSDISARFQAEMKKGLSAESNAAAAVKMLPTHVRSTPDGSEKGQFLALDLGGSKFKVLQV
KVREGMGIRRGGVEMEEKTYPIPKELLIGRGTELFDHVSESLKDFLHEKNISLEKKHPLAFTFSFPCEQS
TLNQGLLLNWSKNYRARGLQGKDVVQALREAIDRTGGMDVEVLAMVNDTVATMMTCGFDDQYCEVGLIIG
TGTNACYMEELRHVDLVEGDEGRMCVNTEWGAFGDDGALDDFITEFDRDVDAASNNPGKQLFEKMVSGMY
LGELVRLVVLKMAKLGLLFDGHVSDALRTKGKITTAHVAAMEEYKNGLKNTRDILMDLDLTPSPDDCIAV

QHVSTIVSFRSSNLVAAGLAAILTRIKQNRNLRTLRITVGVDGTVYKTHPQYPKRLHKVVRRLLPECHVR
FVLSDSGSSKGAALVAAVAQRLASQRRQVDETLSPFRLSQEQLQLVKARMRAGLEAGLKSKGPSAIKMLP
SFVYHTPDGTEHGKYLALDLGGTNFRALLVNFKKGLQQNSRLYHKIYTIPLEIMQGTGEELFDHIVQCIS
DFLDYMGMKNARLPAGFTFSFPCEQTAIDTGTLVSWTKGFKATDCEGHDVVSMLREAIKRRNEFDLDIVA
VVNDTVGTMMSCAYEDPQCEIGLIAGTGSNVCYMEELKNIEKIEQKKGQKERDEETEEEEEGAQEKEDDG
TEREGKEGDAEIRKMCINTEWGGLGDDGSLDDIITPYDTEVDQNSINPGKQRFEKLTSGMYLGEVVRQVL
LDLTRRGLLFRGHVTEALKTPGIFETKYLSQIESDRLALLQVRSILQQLGLDSTCDDSIIVKEVCGAVSR
RAAQLCGAGMAAVVDKIRENRGLDHLNITVGVDGALYKLHPHFSKVLQETTRVLAPQCNVTFLPSEEGSG
KGAALITAVARQKQEM
>anc54|avgPP:0.978|lnPP:-35.985|num_ambig:23|num_ambig_gaps:7
LYAMRLHDDQLRDISARFQAEMKKGLSAESNAAATVKMLPTHVRSTPDGSEKGEFLALDLGGSKFKVLQV
KVREEAGRVRRRVEMEEQTYPIPKELLIGRGTELFDHVSESLKDFLHQKNISLEKKHPLGFTFSFPCQQS
TLNQGLLLNWSKNYRARGLQGKDVVQALRQAINRTGGMDVDVLALVNDTVGTMMTCGFDDQHCEVGLIIG
TGTNACYMEELRHIDLVEGDEGRMCVNTEWGAFGDDGALDDYITEFDRDIDAASINPGKQLFEKMVSGLY
LGELVRLVVLKMAKLGLLFDGHVSDALRTKGKITTAHVAAMEEYKDGLKNTRDILLDLGLTPSPDDCIAV
QHVSTVVSFRSSNLVAAGLAAILTRIRQNRKLKTLRITVGVDGTVYKTHPQYAKRLHKVVRRLVPDCHVR
FVLSESGSSKGAAMVTAVAQRLASQRRQVDETLSPFRLSQEQLRLVKARMRAGLEAGLKSNSPSPVKMLP
SYVYRTPDGTEHGKYLALDLGGTNFRVLLVKIRSGLRRSVRLYNKIYAIPLEIMQGTGEELFDHIVQCIS
DFLDYMGMKNARLPLGFTFSFPCQQTAIDTGTLVSWTKGFKATDCEGNDVVDMLREAIKRRNEFDLDIVA
IVNDTVGTMMSCAYEDPQCEIGLIAGTGSNVCYMEELRNIEKIKQENGEEERKEGSEEEEEGVEEKEDDG
MEREGKEGNGEMQRMCINTEWGGLGEDGCLDDIITPYDIEVDQDSINHGKQRFEKLTSGLYLGEVVRQVL
LDLTRRGLLFRGHVTEALKTPGIFETKYLSQIESDRLALLQVRSILQQLGLDSTCDDSIIVKEVCGAVSR
RAAQLCGAGMAAVVDKIRENRGLDHLNITVGVDGTLYKLHPHFSRVLQETVRALAPQCDVTFLLSEEGSG
KGAALITAVARQQQEVEM
>anc54_altAll|avgPP:0.974|lnPP:-47.395|num_ambig:23|num_ambig_gaps:7
LYAMRLHDDQLRDISARFQAEMKKGLSAESNAAATVKMLPTHVRSTPDGSEKGEFLALDLGGSKFKVLQV
KVREEAGKVRRRVEMEEQTYPIPKELLIGRGVELFDHVSECLKDFLHQKNISLEKKHPLGFTFSFPCQQT
TLNQGLLLNWSKNYRARGLQGKDVVQALRQAINRTGGMDVDVLALVNDTVGTMMTCGFDDQHCEVGLIIG
TGTNACYMEELRHIDLVEGDEGRMCVNTEWGAFGDDGALDDYITEFDRDIDAASTNPGKQLFEKMVSGLY
LGELVRLVVLKMAKLGLLFDGHVSDALRTKGKITTAHVAAMEEYKDGLKNTRDILMDLGLTPSPDDCIAV
QHVCTVVSFRSSNLVAAGLAAILTRIRQNRKLKTLRITVGVDGTVYKTHPQYAKRLHKVVRRLVPDCHVR
FVLSESGSSKGAAMVTAVAQRLASQRRQVDETLSPFRLSQEQLGLVKARMRAGLEAGLKSKSPSPVKMLP
SYVYRTPDGTEHGKYLALDLGGTNFRVLLVKIRSGLRRSVRLYNKIYAIPLEIMQGTGEELFDHIVQCIS
DFLDYMGMKNARLPLGFTFSFPCQQTAIDTGTLVSWTKGFKATDCEGNDVVDMLREAIKRRNEFDLDIVA
IVNDTVGTMMSCAYEDPQCEIGLIAGTGSNVCYMEELRNIEKVEQQNGEEERNEDSDEEEEGVEEKEDDG
MEREGKEGNGEIQRMCINTEWGGLGEDGCLDDIITPYDTEVDQESLNHGKQRFEKLTSGLYLGEVVRQVL
LDLTRRGLLFRGHVTEALKTPGIFETKYLSQIESDRLALLQVRSILQHLGLDSTCDDSIIVKEVCGAVSR
RAAQLCGAGMAAVVDKIRENRGLDHLDITVGVDGTLYKLHPHFSRVLQETVRELAPQCNVTFLLSEEGSG
KGAALITAVARQQQEVEM
>anc55|avgPP:0.974|lnPP:-42.154|num_ambig:22|num_ambig_gaps:13
LYAMRLHDDQLRDISARFLAEMKKGLSSESNAAATVKMLPTHVRSTPDGSEKGEFLALDLGGSKFKVLQV
KVREEAGRVRRRVEMEEQTYPIPKELLIGRGTELFDHVSESLKDFLHQKNISLEKKHPLGFTFSFPCQQS
TLNQGVLLNWSKNYRARGLQGKDVVQALRQAINRTGGMDVDVLALVNDTVGTMMTCGFDDQHCEVGLIIG
TGTNACYMEELRHIDLVEGDEGRMCVNTEWGAFGDDGALDDYITEFDRDIDAASINPGKQLFEKMVSGLY
LGELVRLVVLKMAKLGLLFDGHVSDALRTKGKITTAHVAAMEEYKDGLKNTRDILLDLGLTPSPDDCIAV
QHVSTVVSFRSSNLVAAGLAAILTRIRQNRKLKTLRITVGVDGTVYRTHPQYAKRLHKVVRRLVPDCHVR
FVLSESGSSKGAAMVTAVAQRLASQRRQVDETLSPFRLSQEQLGLVKARMRAGLEAGLKSNSPSPVKMLP
SYVYRTPDGTEHGKYLALDLGGTNFRVLLVKIRSGLRRSVRLYNKIYAIPLEIMQGTGEELFDHIVQCIS
DFLDYMGMKNARLPLGFTFSFPCQQTAIDTGTLVSWTKGFKATDCEGNDVVDMLREAIKRRNEFDLDIVA
IVNDTVGTMMSCAYEDPQCEIGLIAGTGSNVCYMEELRNIEKIEQQNGEEGRKEGSEEEEQEEGRAEGVE
EKEDDGMEREGKEGNGEMQRMCINTEWGGLGEDGCLDDIITPYDIEVDQDSLNHGKQRFEKLTSGLYLGE
IVRQVLLDLTRRGLLFRGHVTEALKTPGIFETKYLSQIESDRLALLQVRSILQQLGLDSTCDDSIIVKEV

CGAVSHRAAQLCGAGMAAVVDKIRENRGLDHLDITVGVDGTLYKLHPHFSRVLQETVRELAPQCDVTFLL
SEEGSGKGAALITAVARQQQEVEM
>anc55_altAll|avgPP:0.970|lnPP:-52.470|num_ambig:22|num_ambig_gaps:13
LYAMRLHDDQLRDISARFLAEMKKGLSAESNAAATVKMLPTHVRSTPDGSEKGEFLALDLGGSKFKVLQV
KVREEAGKVRRKVEMEEQTYPIPKELLIGRGVELFDHVSECLKDFLHQKNISLEKKHPLGFTFSFPCQQT
TLNQGILLNWSKNYRARGLQGKDVVQALRQAINRTGGMDVDVLALVNDTVGTMMTCGFDDQHCEVGLIIG
TGTNACYMEELRHIDLVEGDEGRMCVNTEWGAFGDDGALDDYITEFDRDIDAASTNPGKQLFEKMVSGLY
LGELVRLIVLKMAKLGLLFDGHVSDALRTKGKITTAHVAAMEEYKDGLRNTRDILMDLGLTPSPDDCIAV
QHVCTVVSFRSSNLVAAGLAAILTRIRQNRKLKTLRITVGVDGTVYRTHPQYAKRLHKVVRRLVPDCHVR
FVLSESGSSKGAAMVTAVAQRLASQRRQVDETLSPFRLSQEQLGLVKARMRAGLEAGLKSNSPSPVKMLP
SYVYRTPDGTEHGKYLALDLGGTNFRVLLVKIRSGLRRSVRLYNKIYAIPLEIMQGTGEELFDHIVQCIS
DFLDYMGMKNARLPLGFTFSFPCKQTAIDTGTLVSWTKGFKATDCEGNDVVDMLREAIKRRNEFDLDIVA
IVNDTVGTMMSCAYEDPQCEIGLIAGTGSNVCYMEELRNIEKVKQENGEEGRNEGSEEEEQEEGRAEGVE
EKEDDGMEREGKEGNGEMQRMCINTEWGGLGEDGCLDDIITPYDTEVDQDSLNHGKQRFEKLTSGLYLGE
IVRQVLLDLTRRGLLFRGHVTEALKTPGIFETKYLSQIESDRLALLQVRSILQHLGLDSTCDDSIIVKEV
CGAVSHRAAQLCGAGMAAVVDKIRENRGLDHLDITVGVDGTLYKLHPHFSRVLQETVRELAPQCNVTFVL
SEEGSGKGAALITAVARQQREVEM
>anc56|avgPP:0.970|lnPP:-49.515|num_ambig:31|num_ambig_gaps:12
LYAMRLHDDQLRDISARFLAEMKKGLSSESNAAATVKMLPTHVRSTPDGSEKGEFLALDLGGSKFKVLQV
KVREEVRRKVEMEEQTYPIPKELLNGRGTELFDHVSESLKDFLHQKNISLEKKHPLGFTFSFPCQQSTLN
QGVLLNWSKNYRARGVQGKDVVQALRQAINRTGGMDVDVLALVNDTVGTMMTCGFDDQHCEVGLIIGTGT
NACYMEELRHIDLVEGDEGRMCVNTEWGAFGDDGALDDYITEFDRDIDAASINPGKQLFEKMVSGLYLGE
LVRLVVLKMAKQGLLFDGHVSDALRTKGKITTAHVAAMEEYKDGLKNTRDILLDLGLTPSDDDCIAVQHV
STVVSFRSSNLVAAGLAAILTRIRQNRKLKTLRITVGVDGTVYRTHPQYAKRLHKVVRRLVPDCHVRFVL
SESGSSKGAAMVTAVAQRLASQRRQVDETLSPFRLSQEQLGLVKARMRAGLEAGLKSNSPSPVKMLPSYV
YRTPDGTEHGKYLALDLGGTNFRVLLVKIRSGLRRSVRLYNKIYAIPLEIMQGTGEELFDHIVQCISDFL
DYMGMKNTRLPLGFTFSFPCQQTAIDTGTLVSWTKGFKATDCEGNDVVDMLREAIKRRNEFDLDIVAIVN
DTVGTMMSCAYEDPQCEIGLIAGTGSNVCYMEELRNIEKIEQQNGEEGRKEGSEEEEQEEGRAEGVEEKE
DDGMEREGKEGNGEMQRMCINTEWGGLGEDGCLDDIITPYDSEVDQDSLNHGKQRFEKMTSGLYLGEIVR
QVLLDLTRRGLLFRGHVTEALKTPGIFETKYLSQIESDRLALLQVRSILQQLGLDSTCDDSIIVKEVCGA
VSHRAAQLCGAGMAAVVDKIRENRGLDHLDITVGVDGTLYKLHPHFSRVLQETVRELAPQCDVTFLLSED
GSGKGAALITAVARQQREVEM
>anc56_altAll|avgPP:0.964|lnPP:-66.996|num_ambig:31|num_ambig_gaps:12
LYAMRLNDDQLRDISARFLAEMKKGLSAESNAAATVKMLPTHVRSTPDGSEKGEFLALDLGGSKFKVLQV
KVREEGRRRVEMEEQTYPIPKELLNGRGTELFDHVSESLKDFLHQKNINLEKKHPLGFTFSFPCQQTTLN
QGVLLNWSKNYRARGVQGKDVVQALRQAINRTGGMDVDVLALVNDTVGTMMTCGFDDQHCEVGLIIGTGT
NACYMEELRHIDLVEGDEGRMCVNTEWGAFGDDGALDDYITDFDRDIDAASTNPGKQLFEKMVSGLYLGE
LVRLIVLKMAKLGLLFDGHVSDALRTKGKITTAHVAAMEEYKDGLRNTRDILMDLGLTPSDDDCIAVQHV
CTVVSFRSSNLVAAGLAAILTRIRQNRKLKTLRITVGVDGTVYRTHPQYAKRLHKVVRRLVPDCHVRFVL
SESGSSKGAAMVTAVAQRLASQRRQVDETLSPFRLSQEQLGLVKARMRAGLEAGLKSNSPSPVKMLPSYV
YRTPDGTEHGKYLALDLGGTNFRVLLVKIRSGLRRSVRLYNKIYAIPLEIMQGTGEELFDHIVQCISDFL
DYMGMKNARLPLGFTFSFPCQQTAIDTGTLVSWTKGFKATDCEGNDVVDMLREAIKRRNEFDLDIVAIVN
DTVGTMMSCAYEDPQCEIGLIAGTGSNVCYMEELRNIEKVKPENGEEGRNEGSEEEECTEGRAEGVEEKE
DDGMEREGKEGSGEMQRMCINTEWGGLGEDGCLDDIITPYDSEVDRDSLNHGKQRFEKMTSGLYLGEIVR
QVLLDLTRRGLLFRGHVTEALKTPGIFETKYLSQIESDRLALLQVRSILQHLGLDSTCDDSIIVKEVCGA
VSHRAAQLCGAGMAAVVDKIRENRGLDHLDITVGVDGTLYKLHPHFSRVLQETVKELAPQCNVNFVLSED
GSGKGAALITAVARQLQEVET
>anc57|avgPP:0.975|lnPP:-39.367|num_ambig:38|num_ambig_gaps:1
LYAMRLSDDQLKDISARFMAEMEKGLSSESNATATVKMLPTHVRSTPDGSEKGEFLALDLGGSKFKVLQV
KVSEDGRRKVEMESETYPIPEELLNGRGTELFDHVAESLKDFMQKKNINQKKQPLGFTFSFPCRQSTLDQ
GVLLSWSKNFKARGVQGTNVVESLRQAINRAGGIDVDVLALVNDTVGTMMTCGYDDQHCEVGVIVGTGTN
ACYMEELRHIDLVEGDEGRMCINTEWGAFGDDGALDDFITDFDREIDAASINPGKQLFEKMVSGLYLGEL

VRLILLKMAKKGLLFGGRVSDALRTKGKLQTKHVCLMEEYKDGLKNTRDILVDLGLSPSDDDCIAVQHVC
TIVSFRSSNLCAAALAAILTRIRENRKLKTLRITVGVDGTVYRTHPQYAKRLHKVVRRLVPDCHVRFVLS
ESGSGKGAAMVTAVAQRLASQRRQVDETLAPFRLSQEQLGLVRDRMRAGLERGLKKDSHSSAPVKMLPSY
VYRTPDGTERGKYLALDLGGTNFRVLVVKIRSGLRRSVRMYNKIYAIPLEIMQGTGEELFDHIVQCISDF
LDYMGMKNTRLPLGFTFSFPCQQTGIDKGTLVSWTKGFKATDCEGNDVVDMLREAIKRRNEFDLDIVAVV
NDTVGTMMTCAYEDPKCEIGLIAGTGSNVCYMEEMRNIETVEGDEGQMCINTEWGGFGEDGCIDDIRTKF
DREVDKGSLNVGKQRFEKMTSGMYLGEIVRQILIDLTRRGLLFRGHVTEALKTRGIFETKFLSQIESDRL
ALLQVRSILQHLGLDSTCDDSIIVKEVCGAVSHRAAQLCGAGMAAVVDKIRENRGLDHLDITVGVDGTLY
KLHPHFSRVLQETVKELAPQCDVTFVLSEDGSGKGAALITAVARQLCGGET
>anc57_altAll|avgPP:0.969|lnPP:-56.298|num_ambig:38|num_ambig_gaps:1
LYAMRLSDDQLRDISARFLAEMEKGLSAESNATATVKMLPTHVRSTPDGSEKGEFLALDLGGSKFKVLQV
KVCEDGRRKVEMESQTYPIPEELLNGRGTELFDHVAESLKDFMQKNNINQKKQPLGFTFSFPCQQSTLDQ
GVLLSWSKNFKARGVQGTDVVQSLRQAINRAGGIDVDVLALVNDTVGTMMTCGYDDQHCEVGVIIGTGTN
ACYMEELRHIDLVEGDEGRMCINTEWGAFGDDGALDDFITEFDREIDAASINPGKQLFEKMVSGLYLGEL
VRLILLKMAKKGLLFGGRVSDALRTKGKFETRHVAMMEEYKDGLRNTREILTDLGLNPSEDDCIAVQHVC
TIVSFRSSNLCAAALAAILTRIRQNRKLKTLRITVGVDGTVYRTHPQYAKRLHKVVRRLVPDCHVRFVLS
ESGSGKGAAMVTAVAQRLASQRRQVDDTLAPFRLSREQLGLVRDRMRAGLEQGLKRESHSTVPVKMLPSY
VYRTPDGTERGKYLALDLGGTNFRVLVVKIRSGLRRSVRMYNKIYAIPLEIMQGTGEELFDHIVQCISDF
LDYMGMKNTRLPLGFTFSFPCRQTGIDKGTLVSWTKGFKATDCEGNDVVDMLREAIKRRNEFDLDIVAVV
NDTVGTMMTCAYEDPRCEIGLIAGTGSNVCYMEEMRNIETVEGDEGRMCINTEWGGFGEDGCIDDIRTRF
DGEVDEGSLNIGKQRFEKMTSGMYLGEIVRQILIDLTRRGLLFRGHVTETLKTRGIFETKFLSQIESDRL
ALLQVRSILQHLGLDSTCDDSIIVKEVCGAVSHRAAQLCGAGMAAVVDKIRENRGLDHLDITVGVDGTLY
KLHPHFSRVLQETVKELAPQCDVNFVLSEDGSGKGAALITAVARQLCGGET
>anc58|avgPP:0.989|lnPP:-17.337|num_ambig:21|num_ambig_gaps:1
LYAMRLSDDQLLDISARFRAEMEKGLSAESNATAAVKMLPTHVRSTPDGSEKGEFLALDLGGSKFKVLQV
KVSEDGKRKVEMESETYPIPEEILNGRGTDLFDHVAESLKDFMQKNNINQKKKPLGFTFSFPCRQSKLDE
GVLLSWSKNFKARGVQGTNVVESLRQAINRVGGIDVDVLALVNDTVGTMMTCGYDDQHCEVGVIVGTGTN
ACYMEEMRHIDLVEGDEGRMCINTEWGAFGDDGALDDFITDFDREIDAASINPGKQLFEKMVSGMYMGEL
VRLILLKMAKKGLLFGGRVSDALRTKGKFQTKHVCLIEEYKDGLKNTKDILTDLGLNPSEDDCIAVQHVC
TIVSFRSANLCAAALAAILTRIRENRKLKTLRTTVGVDGTVYRTHPQYAKRLHKVVRRLVPDCHVRFVLS
ESGSGKGAAMVTAVAQRLASQRRQVDETLAPFRLSREQLQQVRDKMRVELERGLKKDSHSSASVKMLPSY
VYRTPDGTERGKYLALDLGGTNFRVLVVKIRSGLRKSVRMYNKIYAIPLEIMQGTGEELFDHIVQCISDF
LDYMGMKNTRLPLGFTFSFPCRQTGIDKGTLVSWTKGFKATDCEGNDVVDMLREAIKRRNEFDLDIVAVV
NDTVGTMMTCAYEDPKCEIGLIAGTGSNVCYMEEMRNIETVEGDEGQMCVNTEWGGFGENGCIEDIRTKF
DREVDEGSLNVGKQRFEKMTSGMYLGEIVRQILIDLTRRGLLFRGRITEPLKTRGIFETKFLSQIESDRL
ALLQVRSILQHLGLDSTCDDSIIVKEVCGAVSRRAAQLCGAGMAAVVDKIRENRGLDHLDITVGVDGTLY
KLHPHFSRILKETVKELAPKCDVTFVLSEDGSGKGAALITAVARQLCGGET
>anc58_altAll|avgPP:0.985|lnPP:-26.591|num_ambig:21|num_ambig_gaps:1
LYAMRLSDDQLKDISARFRAEMEKGLSSESNATATVKMLPTHVRSTPDGSEKGEFLALDLGGSKFKVLQV
KVSEDGKRKVEMESQTYPIPEEILNGRGTDLFDHVAESLKDFMQKNNINQKKKPLGFTFSFPCRQSKLDE
GVLLSWSKNFKARGVQGTDVVQSLRQAINRAGGIDVDVLALVNDTVGTMMTCGYDDQHCEVGVIIGTGTN
ACYMEEMRHIDLVEGDEGRMCINTEWGAFGDDGALDDFITDFDREIDAASINPGKQLFEKMVSGMYMGEL
VRLILLKMAKKGLLFGGRVSDALRTKGKFQTKHVALIEEYKDGLKNTREILTDLGLNPSEDDCIAVQHVC
TIVSFRSANLCAAALAAILTRIRENRKLKTLRTTVGVDGTVYRTHPQYAKRLHKVVRRLVPDCHVRFVLS
ESGSGKGAAMVTAVAQRLASQRKQIDDTLAPFRLSREQLQQVRDRMRVELERGLRKDSHSTASVKMLPSY
VYRTPDGTERGKYLALDLGGTNFRVLVVKIRSGLRRSVRMYNKIYAIPLEIMQGTGEELFDHIVQCISDF
LDYMGMKNTRLPLGFTFSFPCRQTGIDKGTLVSWTKGFKATDCEGNDVVDMLREAIKRRNEFDLDIVAVV
NDTVGTMMTCAYEDPKCEIGLIAGTGSNVCYMEEMRNIETVEGDEGRMCVNTEWGGFGENGCIEDIRTKF
DREVDEGSLNVGKQRFEKMTSGMYLGEIVRQILIDLTKRGLLFRGRITEPLKTRGIFETKFLSQIESDRL
ALLQVRSILQHLGLDSTCDDSIIVKEVCGAVSHRAAQLCGAGMAAVVDKIRENRGLDHLDITVGVDGTLY
KLHPHFSRILKETVKELAPKCDVTFVLSEDGSGKGAALITAVARQLCGGET
>anc59|avgPP:0.992|lnPP:-11.601|num_ambig:16|num_ambig_gaps:1

LYAMRLSDDQLLDISARFRAEMEKGLSAESNATAAVKMLPTHVRSTPDGSEKGEFLALDLGGSKFKVLQV
KVSEDGKRKVEMESQTYPIPEEILNGRGTDLFDHVAESLKDFMQKNNINQKKKPLGFTFSFPCRQSKLDE
GVLLSWSKNFKARGVQGTDVVESLRQAIDRVGGIDVDVLALVNDTVGTMMTCGYDDQHCEVGVIVGTGTN
ACYMEEMRHIDLVEGDEGRMCINTEWGAFGDDGALDDFITDFDREIDAASINPGKQLFEKMVSGMYMGEL
VRLILLKMAKKGLLFGGRVSDALRTKGKFQTKHVALIEKYKEGLKNTKDILTDLGLNPSEDDCIAVQHVC
TIVSFRSANLCAAALAAILTRIRENRKLKTLRTTVGVDGTVYRTHPQYAKRLHKVVRRLVPDCHVRFVLS
ESGSGKGAAMVTAVAQRLVSQRRQVDETLAPFRLSREQLQQVRDKMRVELERGLKKDTHSTASVKMLPSY
VYRTPDGTERGKYLALDLGGTNFRVLVVKIRSGLRKSVRMYNKIYAIPLEIMQGTGEELFDHIVQCISDF
LDYMGMKNTRLPLGFTFSFPCRQTGIDKGTLVSWTKGFKATDCEGNDVVDMLREAIKRRNEFDLDIVAVV
NDTVGTMMTCAYEDPKCEIGLIAGTGSNVCYMEEMRNIETVEGDEGRMCVNTEWGGFGENGCIEDIRTKF
DREVDEGSLNVGKQRFEKMTSGMYLGEIVRQILIDLTRRGLLFRGRITERLKTRGIFETKFLSQIESDRL
ALLQVRSILQHLGLDSTCDDSIIVKEVCGAVSRRAAQLCGAGMAAVVDKIRENRGLDHLDITVGVDGTLY
KLHPHFSRILKETVKELAPKCDVTFVLSEDGSGKGAALITAVARQLCGGET
>anc59_altAll|avgPP:0.988|lnPP:-21.345|num_ambig:16|num_ambig_gaps:1
LYAMRLSDDQLLDISARFRAEMEKGLSAESNATATVKMLPTHVRSTPDGSEKGEFLALDLGGSKFKVLQV
KVSEDGKRKVEMESETYPIPEEILNGRGTDLFDHVAESLKDFMQKNHINQKKKPLGFTFSFPCRQSKLDE
GVLLSWSKNFKARGVQGTDVVESLRQAINRAGGIDVDVLALVNDTVGTMMTCGYDDQHCEVGVIIGTGTN
ACYMEEMRHIDLVEGDEGRMCINTEWGAFGDDGALDDFITDFDREIDAASINPGKQLFEKMVSGMYMGEL
VRLILLKMAKKGLLFGGRVSDALRTKGKFQTKHVALIEKYKEGLKNTREILTDLGLNPSEDDCIAVQHVC
TIVSFRSANLCAAALAAILTRIRENRKLKTLRTTVGVDGTVYRTHPQYAKRLHKVVRRLVPDCHVRFVLS
ESGSGKGAAMVTAVAQRLVSQRKQIDDTLAPFRLSRQQLQQVRDRMRVELERGLRKDTHSTASVKMLPSY
VYRTPDGTERGKYLALDLGGTNFRVLVVKIRSGLRRSVRMYNKIYAIPLEIMQGTGEELFDHIVQCISDF
LDYMGMKNTRLPLGFTFSFPCRQTGIDKGTLVSWTKGFKATDCEGNDVVDMLREAIKRRNEFDLDIVAVV
NDTVGTMMTCAYEDPKCEIGLIAGTGSNVCYMEEMRNIETVEGDEGRMCVNTEWGGFGENGCIEDIRTKF
DREVDEGSLNVGKQRFEKMTSGMYLGEIVRQILIDLTRRGLLFRGRITERLKTRGIFETKFLSQIESDRL
ALLQVRSILQHLGLDSTCDDSIIVKEVCGAVSRRAAQLCGAGMAAVVDKIRENRGLDHLDITVGVDGTLY
KLHPHFSRILKETVKELAPKCDVTFVLSEDGSGKGAALITAVARRLCGGET
>anc60|avgPP:0.985|lnPP:-23.202|num_ambig:19|num_ambig_gaps:0
LYAMRLSDEQLQDISARFRAEMEKGLSAKSNATATVKMLPTHVLSTPDGSEKGEFLALDLGGSSLKVLQV
KVSEDGKRKVEMKSHTYPIPDGIINGRGTDLFNHVAESLKDFMEKNHINNKKTPLGFTFSFPCRQSKLDE
GVLLSWSKNYKARGVQGMDVVESLRQAINRTGGMDVEVLALVNDTVGTMMSCGYDDQHCEVGVTVGTGTN
ACYMEELRHIDLVEGDEGRMCISTEWGAFGDDGVLNDFITDFDREIDAASSNPGKQLFEKMVSGLYMGEL
VRIILLKMTKKGLLFEGRASDALRTKGKFQTKHIALIEENKEGLKNTKEILVDLDLNPSEEDCIAVQHVC
TIVSFRSANLCAAALAAILTRIRQNRKLKTLRTTVGVDGTVYRTHPQYPKRLHEVVRRLVPDCHVRFMLS
ESGSGKGVAMVTAVAQRLVIQRQQISETLAAFKLSPEQLQEVRDKMRIEMERGLRKDTHSTASVKMLPTY
VYSIPDGTETGKYLALDLGGTNFRVLVVKIRSGLRKSVHMDNKIYAIPLEIMQGTREELFDHIVQCISDF
LDYMGMKNTCLPLGFTFSFPCRQTGIDKGSLVCWTKGFKATDCEGNDVVDMLREAIKRRNEFDLDIVAVM
NDTVGTMMTCAYEDPKCEIGLIAGTGSNVCYMEDMRNIETVEGDEGRMCVNTEWGGFGENGCIDDIRTKF
DREVDEGSLNVGKQRFEKMMSGMYLGEIVRQILIDLTRRGLLFRGHITERLKTRGIFETKFLSQIESDRL
ALLQVRSILQHLGLDSTCDDSIIVKEVCGTVSRRAAQLCGAGMAAVVDKIRENRGLDHLDITVGVDGTLY
KLHPHFSRILKETVRELAPKCDVTFVLSEDGSGKGAALITAVAQQRYSEM
>anc60_altAll|avgPP:0.982|lnPP:-31.583|num_ambig:19|num_ambig_gaps:0
LYAMRLSDEQLQDISARFRAEMEKGLSAKSNATATVKMLPTHVLSTPDGSEKGEFLALDLGGSSLKVLQV
KVSEDGKRKVEMNSHTYPIPDAVINGRGTDLFNHVAESLKDFMEKNHINNRKTPLGFTFSFPCRQSKLDE
GVLLSWSKNYKARGVQGVDVVESLRQAINRTGGMDVEVLALVNDTVGTMMSCGYDDQHCEVGVTVGTGTN
ACYMEELRHIDLVEGDEGRMCISTEWGAFGDDGVLNDFITDFDREIDAASSNPGKQLFEKMVSGLYMGEL
VRIILLKMTKKGLLFEGRASDALRTKGKFQTKHIALIEEDKEGLKNTKEILVDLDLNPSEEDCIAVQHVC
TIVSFRSANLCAAALAAILTRIRQNRKLKTLRTTVGVDGTVYRTHPQYPKRLHEVVRRLVPDCHVRFMLS
ESGSGKGVAMVTAVAQRLVMQRQQISETLAAFKLSLEQLQEVRDKMRIEMERGLRKDTHSTASVKMLPTY
VYNMPDGTETGKYLALDLGGTNFRVLVVKIRSGLHKSVHMDNKIYTVPLEIMQGTREELFDHIVQCISDF
LDYMGMKNTCLPLGFTFSFPCRQTGIDKGSLVCWTKGFKATDCEGNDVVDMLREAIKRRNEFELDIVAVM
NDTVGTMMTCAYEDSECEIGLIAGTGSNVCYMEDMRNIETVEGDEGRMCVNTEWGGFGENGCIDDIRTKF

DREVDEGSLNVGKQRFEKMISGMYLGEIVRQILIDLTRRGLLFRGHITERLKTRGIFETKFLSQIESDRL
ALLQVRSILQHLGLDSTCDDSIIVKEVCGTVSRRAAQLCGAGMAAVVDKIRENRGLDHLDITVGVDGTLY
KLHPHFSRILKETVRELAPKCDVTFVLSEDGSGKGAALITTVAQRRYSEM
>anc61|avgPP:0.989|lnPP:-16.990|num_ambig:21|num_ambig_gaps:1
LYAMRLSDDQLLDISARFRAEMEKGLSAESNPTAAVKMLPTHVLSTPDGSEKGEFLALDLGGSKFKVLQV
KVSEDGKRKVEMESQTYPIPEEIVNGRGTDLFNHVAESLKDFMQKNHINQKKKPLGFTFSFPCRQSKLDE
GVLLSWSKNYKARGVQGTDVVESLRQAIDRAGGMDVEVLALVNDTVGTMMTCGYDDQHCEVGVIVGTGTN
ACYMEELRHIDLVEGDEGRMCINTEWGAFGDDGSLDDFITDFDREIDAASINPGKQLFEKMVSGMYMGEL
VRLILLKMAKKGLLFEGRASDALRTKGKFQTKHVALIEKYKEGLKNTKDILTDLGLNPSEDDCIAVQHVC
TIVSFRSANLCAAALAAILTRIRENRKLKTLRTTVGVDGTVYRTHPQYPKRLHKVVRRLVPDCHVRFVLS
ESGSGKGAAMVTAVAQRLVSQRRQVDETLAPFRLSRQQLQQVRDKMRVELERGLKKDTHSTASVKMLPSY
VYRTPDGTETGKYLALDLGGTNFRVLVVKIRSGLRKSVRMYNKIYAIPLEIMQGTGEELFDHIVQCISDF
LDYMGMKNTCLPLGFTFSFPCRQTGIDKGSLVSWTKGFKATDCEGNDVVDMLREAIKRRNEFDLDIVAVV
NDTVGTMMTCAYEDPKCEIGLIAGTGSNVCYMEEMRNIETVEGDEGRMCVNTEWGGFGENGCIEDIRTKF
DREVDEGSLNVGKQRFEKMTSGMYLGEIVRQILIDLTRRGLLFRGHITERLKTRGIFETKFLSQIESDRL
ALLQVRSILQHLGLDSTCDDSIIVKEVCGAVSRRAAQLCGAGMAAVVDKIRENRGLDHLDITVGVDGTLY
KLHPHFSRILKETVRELAPKCDVTFVLSEDGSGKGAALITAVARQLYSGET
>anc61_altAll|avgPP:0.985|lnPP:-26.874|num_ambig:21|num_ambig_gaps:1
LYAMRLSDDQLLDISARFRAEMEKGLSAESNATATVKMLPTHVLSTPDGSEKGEFLALDLGGSRFKVLQV
KVSEDGKRKVEMESQTYPIPEEVINGRGTDLFNHVAESLKDFMQKNHINQKKKPLGFTFSFPCRQSKLDE
GVLLSWSKNYKARGVQGTDVVESLRQAINRAGGMDVEVLALVNDTVGTMMTCGYDDQHCEVGVIVGTGTN
ACYMEELRHIDLVEGDEGRMCINTEWGAFGDDGALDDFITDFDREIDAASINPGKQLFEKMVSGMYMGEL
VRLILLKMTKKGLLFEGRASDALRTKGKFQTKHVALIEKYKEGLKNTREILTDLGLNPSEDDCIAVQHVC
TIVSFRSANLCAAALAAILTRIRENRKLKTLRTTVGVDGTVYRTHPQYPKRLHKVVRRLVPDCHVRFVLS
ESGSGKGAAMVTAVAQRLVAQRRQIDDTLAPFRLSREQLQQVRDRMRVELERGLRKDTHSTASVKMLPSY
VYRTPDGTETGKYLALDLGGTNFRVLVVKIRSGLRKSVRMYNKIYAIPLEIMQGTGEELFDHIVQCISDF
LDYMGMKNTCLPLGFTFSFPCRQTGIDKGSLVSWTKGFKATDCEGNDVVDMLREAIKRRNEFELDIVAVV
NDTVGTMMTCAYEDPECEIGLIAGTGSNVCYMEEMRNIETVEGDEGRMCVNTEWGGFGENGCIEDIRTKF
DREVDEGSLNVGKQRFEKMTSGMYLGEIVRQILIDLTKRGLLFRGHITERLKTRGIFETKFLSQIESDRL
ALLQVRSILQHLGLDSTCDDSIIVKEVCGAVSRRAAQLCGAGMAAVVDKIRENRGLDHLDITVGVDGTLY
KLHPHFSRILKETVRELAPKCDVTFVLSEDGSGKGAALITAVARRLCSGET
>anc62|avgPP:0.991|lnPP:-13.353|num_ambig:19|num_ambig_gaps:1
LYAMRLSDDQLLDISARFRAEMEKGLSAESNATAAVKMLPTHVRSTPDGSEKGEFLALDLGGSKFKVLQV
KVSEDGKRKVEMESQTYPIPEEILNGRGTDLFDHVAESLKDFMQKNHINQKKKPLGFTFSFPCRQSKLDE
GVLLSWSKNFKARGVQGTDVVESLRQAIDRAGGIDVDVLALVNDTVGTMMTCGYDDQHCEVGVIVGTGTN
ACYMEEMRHIDLVEGDEGRMCINTEWGAFGDDGSLDDFITDFDREIDAASINPGKQLFEKMVSGMYMGEL
VRLILLKMAKKGLLFEGRASDALRTKGKFQTKHVALIEKYKEGLKNTKDILTDLGLNPSEDDCIAVQHVC
TIVSFRSANLCAAALAAILTRIRENRKLKTLRTTVGVDGTVYRTHPQYAKRLHKVVRRLVPDCHVRFVLS
ESGSGKGAAMVTAVAQRLVSQRRQVDETLAPFRLSRQQLQQVRDKMRVELERGLKKDTHSTASVKMLPSY
VYRTPDGTERGKYLALDLGGTNFRVLVVKIRSGLRKSVRMYNKIYAIPLEIMQGTGEELFDHIVQCISDF
LDYMGMKNTRLPLGFTFSFPCRQTGIDKGTLVSWTKGFKATDCEGNDVVDMLREAIKRRNEFDLDIVAVV
NDTVGTMMTCAYEDPKCEIGLIAGTGSNVCYMEEMRNIETVEGDEGRMCVNTEWGGFGENGCIEDIRTKF
DREVDEGSLNVGKQRFEKMTSGMYLGEIVRQILIDLTRRGLLFRGRITERLKTRGIFETKFLSQIESDRL
ALLQVRSILQHLGLDSTCDDSIIVKEVCGAVSRRAAQLCGAGMAAVVDKIRENRGLDHLDITVGVDGTLY
KLHPHFSRILKETVKELAPKCDVTFVLSEDGSGKGAALITAVARQLCSGET
>anc62_altAll|avgPP:0.988|lnPP:-22.891|num_ambig:19|num_ambig_gaps:1
LYAMRLSDDQLLDISARFRAEMEKGLSAESNPTATVKMLPTHVRSTPDGSEKGEFLALDLGGSKFKVLQV
KVSEDGKRKVEMESQTYPIPEEILNGRGTDLFDHVAESLKDFMQKNHINQKKKPLGFTFSFPCRQSKLDE
GVLLSWSKNFKARGVQGTDVVESLRQAINRAGGIDVDVLALVNDTVGTMMTCGYDDQHCEVGVIIGTGTN
ACYMEEMRHIDLVEGDEGRMCINTEWGAFGDDGALDDFITDFDREIDAASINPGKQLFEKMVSGMYMGEL
VRLILLKMAKKGLLFGGRVSDALRTKGKFQTKHVALIEKYKEGLKNTREILTDLGLNPSEDDCIAVQHVC
TIVSFRSANLCAAALAAILTRIRENRKLKTLRTTVGVDGTVYRTHPQYAKRLHKVVRRLVPDCHVRFVLS

```
ESGSGKGAAMVTAVAQRLVSQRKQIDDTLAPFRLSREQLQQVRDRMRVELERGLRKDTHSTASVKMLPSY
VYRTPDGTERGKYLALDLGGTNFRVLVVKIRSGLRRSVRMYNKIYAIPLEIMQGTGEELFDHIVQCISDF
LDYMGMKNTRLPLGFTFSFPCRQTGIDKGTLVSWTKGFKATDCEGNDVVDMLREAIKRRNEFDLDIVAVV
NDTVGTMMTCAYEDPKCEIGLIAGTGSNVCYMEEMRNIETVEGDEGRMCVNTEWGGFGENGCIEDIRTKF
DREVDEGSLNVGKQRFEKMTSGMYLGEIVRQILIDLTRRGLLFRGRITERLKTRGIFETKFLSQIESDRL
ALLQVRSILQHLGLDSTCDDSIIVKEVCGAVSRRAAQLCGAGMAAVVDKIRENRGLDHLDITVGVDGTLY
KLHPHFSRILKETVKELAPKCDVTFVLSEDGSGKGAALITAVARRLYGGET
>anc63|avgPP:0.990|lnPP:-15.978|num_ambig:13|num_ambig_gaps:1
LYAMRLSDDQLLDIAARFRAEMEKGLSAESNPTAAVKMLPTHVRSTPDGSEKGEFLALDLGGSKFKVLQV
KVSEDGKRKVEMESQTYPTPEEILSGKGTDLFDHVAECLKDFMQKKHINQKKQPLGFTFSFPCRQSKLDE
GVLLSWSKNFKARGVQGTDVVESLRKAIDRAGGIDVDVLALVNDTVGTMMTCGYDDQHCEVGVIIGTGTN
ACYMEEMRHIDLVEGDEGRMCINTEWGAFGDDGSLDDFITDFDREIDAASINPGKQLFEKMISGMYMGEL
VRLILLKMAKKGLLFKGRASDALRTKGKFETKHVAAIEKYKEGLSNTKEILTDLGLNPSEDDCIAVQHVC
TIVSFRSANLCAAALAAILTRIRENKKLKRLRTTVGVDGTVYRTHPQYAKRLHKVVRRLVPDCHVRFVLS
ESGSGKGAAMVTAVAQRLVSQRKQIDETLAPFRLSRQQLQQVRDKMRVELERGLKKDTHHTASVKMLPSY
VYRTPDGTERGKFLALDLGGTNFRVLVVKIRSGLRRSVRMYNKIYAIPLEIMQGTGEELFDHIVQCISDF
LDYMGMKNTRLPLGFTFSFPCRQTGIDKGTLVSWTKGFKATDCEGNDVVDMLREAIKRRNEFDLDIVAVV
NDTVGTMMTCAYEDPNCEIGLIAGTGSNVCYMEEMRNIETVEGDEGRMCVNTEWGGFGENGCIEDIRTKF
DREVDEGSLNVGKQRFEKMTSGMYLGEIVRQILIDLTRRGLLFRGRITERLKTRGIFETKFLSQIESDRL
ALLQVRSILQQLGLDSTCDDSIIVKEVCGAVSRRAAQLCGAGMAAVVDKIRENRGLDHLDITVGVDGTLY
KLHPHFSRILKETVKELAPQCDVTFVLSEDGSGKGAALITAVARKLHSGET
>anc63_altAll|avgPP:0.987|lnPP:-23.460|num_ambig:13|num_ambig_gaps:1
LYAMRLSDDQLLDIAARFRAEMEKGLSAESNPTAAVKMLPTHVRSTPDGSEKGEFLALDLGGSKFKVLQV
KVSEDGKRKVEMESQTYPTPEEILSGKGTDLFDHVAESLKDFMQRNHINQKKQPLGFTFSFPCRQSKLDE
GVLLSWSKNFKARGVQGTDVVESLRKAIDRAGGIDVDVLALVNDTVGTMMTCGYDDQHCEVGVIVGTGTN
ACYMEEMRHIDLVEGDEGRMCINTEWGAFGDDGSLDDFITDFDREIDAASINPGKQLFEKMISGMYMGEL
VRLILLKMAKKGLLFKGRVSDALRTKGKFETKHVAAIEKYKEGLNNTKEILTDLGLDPSEDDCIAVQHVC
TIVSFRSANLCAAALAAILTRIRENKKLKRLRTTVGVDGTVYRTHPQYAKRLHKVVRRLVPDCHVRFLLS
ESGSGKGAAMVTAVAQRLVSQRRQIDETLAPFRLSRQQLQQVRDKMRVELERGLKKDTHHTASVKMLPSY
VYRTPDGTERGKFLALDLGGTNFRVLVVKIRSGLRKSVRMYNKIYAIPLEIMQGTGEELFDHIVQCISDF
LDYMGMKNTRLPLGFTFSFPCRQTGIDKGTLVSWTKGFKATDCEGNDVVDMLREAIKRRNEFDLDIVAVV
NDTVGTMMTCAYEDPNCEIGLIAGTGSNVCYMEEMRNIETVEGDEGRMCVNTEWGGFGENGCIEDIRTKF
DREVDEGSLNVGKQRFEKMTSGMYLGEIVRQILIDLTRRGLLFRGRITERLKTRGIFETKFLSQIESDRL
ALLQVRAILQQLGLDSTCDDSIIVKEVCGAVSRRAAQLCGAGMAAVVDKIRENRGLDHLDITVGVDGTLY
KLHPHFSRILKETVKELAPQCDVTFVLSEDGSGKGAALITAVARRLHSGAT
>anc64|avgPP:0.984|lnPP:-25.142|num_ambig:19|num_ambig_gaps:1
LYHMRLSDDQLLDIMTRFRAEMEKGLSRETNPTAAVKMLPTHVRSTPDGSEKGEFLALDLGGSKFKVLRV
KVSEDGKQKVEMESQTYPTPEEIISGKGTDLFDHVAECLKDFMEKKHINQKKQPLGFTFSFPCRQTKLDE
GVLLSWSKNFKARGVQGTDVVDSLRKAIDKTGGIDVDVLALVNDTVGTMMTCGYDDQYCEVGVIIGTGTN
ACYMEEMRHIDLVEGDEGRMCINTEWGAFGDDGSLEDIRTEFDREIDLGSINPGKQLFEKMISGMYMGEL
VRLILVKMAKKGLLFKGRISDALRTKGKFETKHVAAIEKYKEGLSNAKEILTDLGLNPSEDDCIAVQHVC
TIVSFRSANLCAAALAAILTRLRENKKLKRLRTTVGMDGTVYRTHPQYAKRLHKVVRRLVPDCNVRFLLS
ESGSGKGAAMVTAVAYRLVSQRKQIDETLAPFKLSRQQLQEVKDKMRVELERGLKKDTHHTASVKMLPSY
VYGTPDGTERGKFLALDLGGTNFRVLLVKIRSGLRRSVRMYNKIYAIPLEIMQGTGEELFDHIVQCISDF
LDYMGMKNARLPLGFTFSFPCRQTGIDKGTLVSWTKGFKATDCEGQDVVDMLREAIKRRNEFDLDIVAVV
NDTVGTMMTCAYEDPNCEIGLIAGTGSNVCYMEEMRNIETVEGDEGRMCVNTEWGGFGDNGCIEDIRTKF
DREVDEGSLNPGKQRYEKMTSGMYLGEIVRQILIDLTKRGLLFRGQISERLKTRGIFETKFLSQIESDRL
ALLQVRSILQQLGLDSTCDDSIIVKEVCGAVSRRAAQLCGAGMAAVVDKIRENRGLDHLDITVGVDGTLY
KLHPHFSRILKETVKELAPQCDVTFVLSEDGSGKGAALITAVARKLHSAGS
>anc64_altAll|avgPP:0.981|lnPP:-34.199|num_ambig:19|num_ambig_gaps:1
LYHMRLSDDQLLDIMARFRAEMEKGLSRDTNPTAAVKMLPTHVRSTPDGSEKGEFLALDLGGSKFKVLRV
KVSEDGKQKVEMESQTYPTPEEIISGKGTELFDHVAECLKDFMERNHINQKKHPLGFTFSFPCRQTKLDE
```

GVLLSWSKNFKARGVQGTDVVESLRKAIDKAGGIDVDVLALVNDTVGTMMTCGYDDQHCEVGVIIGTGTN
ACYMEEMRHIDLVEGDEGRMCINTEWGAFGDDGSLEDIRTEFDREIDLGSINPGKQLFEKMISGMYMGEL
VRLILVKMAKKGLLFKGRMSDALRTKGKFETKHVAAIEKYKEGLSNAKEILTDLGLDPSEDDCIAVQHVC
TIVSFRSANLCAAALAAILTRLRENKKLKRLRTTVGMDGTVYRTHPQYAKRLHKVVRRLVPDCNVRFLLS
ESGSGKGAAMVTAVAYRLVSQRKQIDETLAPFKLSRQQLQDVKDKMRVELERGLKKDTHHTASVKMLPSY
VYGTPDGTERGKFLALDLGGTNFRVLLVKIRSGLRKSVRMYNKIYAIPLEIMQGTGEELFDHIVQCISDF
LDYMGMKNARLPLGFTFSFPCRQAGIDKGTLVSWTKGFKATDCEGEDVVDMLREAIKRRNEFDLDIVAVV
NDTVGTMMTCAYEDPNCEIGLIAGTGSNVCYMEEMRNIETVEGDEGRMCVNTEWGGFGDNGCIEDIRTKF
DREVDEGSLNQGKQRYEKMTSGMYLGEIVRQILIDLTKRGLLFRGQISERLKTRGIFETKFLSQIESDRL
ALLQVRAILQQLGLDSTCDDSIIVKEVCGAVSRRAAQLCGAGMAAVVDKIRENRGLDHLDITVGVDGTLY
KLHPHFSKILKETVKELAPQCDVTFVLSEDGSGKGAALITAVARRLHSAGS
>anc65|avgPP:0.991|lnPP:-13.163|num_ambig:13|num_ambig_gaps:0
LYHMRLSDETLLDIMARFQAEMEKGLGKDTNPTASVKMLPTFVRAIPDGSENGEFLSLDLGGSKFRVLKV
QVSEEGKRNVHMESQFYPTTNEITRGNGSELFEYVADCLADFMKAKDLKHKKLPLGFTFSFPCRQTKLDE
GVLLSWTKKFKARGVQDTDVVSHLTKAMKKHPDMDVDILALVNDTVGTMMTCAYDDPYCEVGVIIGTGSN
ACYMEDMSNIDLVEGDEGRMCINTEWGAFGDDGALEDIRTEFDRELDLGSLNPGKQLFEKMISGLYLGEL
VRLILLKMARAGLLFGGKTSSALHTKDKIETRHVAAMEKYKEGLANTKEILTDLGLEPSEADCIAVQHVC
TIVSFRSANLCAAALAAILTRLRENKKLVRLRTTVGVDGTLYKIHPQYPKRLHKVVRKLVPNCDVRFLLS
ESGSTKGAAMVTAVASRVQAQRKQIDEVLALFRLTHEHLVDVQDKMRAELEYGLKKETHPMATVKMLPTY
VCGMPDGTEKGKFLALDLGGTNFRVLLVKIRRGRKSVRMYNKIFAIPLEIMQGTGEELFDHIVQCIADFL
DYMGLKGAQLPLGFTFSFPCRQTSIDKGILIEWTKGFKATDCEGEDVVDMLREAIKRRNEFDLDIVALVN
DTVGTMMTCAYEDPNCEIGLIAGTGSNVCYMEEMRNIELVEEDKGKMCINTEWGGFGDNGCIDDIQTQYD
KEVDEGSLNPGKQRYEKMTSGMYLGEIVRQILIDLTKQGLLFRGQISERLRTRGIFETKFLSQIESDRLA
LLQVRGILQQLGLDSTCEDSIVVKEVCGAVSRRAAQLCGAGLAAIVEKRREDQGLEHLKITVGVDGTLYK
LHPHFSRILQETVKELAPQCDVTFMLSEDGSGKGAALITAVAKRLQQAQKEN
>anc65_altAll|avgPP:0.989|lnPP:-19.767|num_ambig:13|num_ambig_gaps:0
LYHMRLSDETLLDIMARFQAEMEKGLGKDTNPTASVKMLPTFVRAIPDGSENGEFLSLDLGGSKFRVLKV
QVSEEGKRNVHMESQFYPMTNEITRGNGSELFEYVADCLADFMKTKDLKHKKLPLGFTFSFPCRQTKLDE
GVLLSWTKKFKARGVQDTDVVSRLTKAMKKHPDMDVDILALVNDTVGTMMTCAYDDPYCEVGVIIGTGSN
ACYMEDMSNIDLVEGDEGRMCINTEWGAFGDDGALEDIRTEFDRELDLGSLNPGKQLFEKMISGLYLGEL
VRLILLKMARAGLLFGGKKSSALHTKDKIETRHVAAMEKYKEGLANTREILTDLGLEPSEADCIAVQHVC
TIVSFRSANLCAAALAAILTRLRENKKLVRLRTTVGVDGTLYKIHPQYPKRLHKVVRKLVPNCDVRFLLS
ESGSAKGAAMVTAVASRVQAQRKQIDEVLALFRLTREHLVDVRDKMRAELEYGLKKETHPMATVKMLPTY
VYGMPDGTEKGKFLALDLGGTNFRVLLVKIRRGRKSVRMYNKIFAIPLEIMQGTGEELFDHIVQCIADFL
DYMGLKGAQLPLGFTFSFPCRQTSIDKGILIEWTKGFKATDCEGEDVVDMLREAIKRRNEFDLDIVALVN
DTVGTMMTCAYEDPNCEIGLIAGTGSNVCYMEEMRNIELVEEDKGKMCINTEWGGFGDNGCIDDIRTQYD
REVDEGSLNPGKQRYEKMTSGMYLGEIVRQILIDLTKQGLLFRGQVSERLRTRGIFETKFLSQIESDRLA
LLQVRGILQQLGLDSTCEDSIVVKEVCGAVSRRAAQLCGAGLAAIVEKRREDQGLEHLKITVGVDGTLYK
LHPHFSRILQETVKELAPQCDVTFMLSEDGSGKGAALITAVAKRLQQPQKEN
>anc66|avgPP:0.996|lnPP:-5.669|num_ambig:7|num_ambig_gaps:0
LYHMRLSDETLLDIMARFQAEMEKGLGKDTNPTASVKMLPTFVRAIPDGSENGEFLSLDLGGSKFRVLKV
QVSEEGKRNVQMESQFYPTPNEIIRGNGSELFEYVADCLADFMKTKDLKHKKLPLGLTFSFPCRQTKLEE
GVLLSWTKKFKARGVQDTDVVSHLTKAMKKHPDMDVDILALVNDTVGTMMTCAYDDPYCEVGVIIGTGTN
ACYMEDMSNIDLVEGDEGRMCINTEWGAFGDDGALEDIRTEFDRELDLGSLNPGKQLFEKMISGLYLGEL
VRLILLKMARAGLLFGGKKSSALHTKGKIETRHVAAMEKYKEGLANTREILTDLGLEPSEADCIAVQHVC
TIVSFRSANLCAAALAAILTRLRENKKLVRLRTTVGMDGTLYKIHPQYPKRLHKVVRKLVPNCDVRFLLS
ESGSTKGAAMVTAVASRVQAQRKQIDEVLALFRLTHEHLVDVQDKMRAELEYGLKKETHPMATVKMLPTY
VCGMPDGTEKGKFLALDLGGTNFRVLLVKIRRGRKSVRMYNKIFAIPLEIMQGTGEELFDHIVQCIADFL
DYMGLKGAQLPLGFTFSFPCRQTSIDKGILIEWTKGFKATDCEGEDVVDMLREAIKRRNEFDLDIVAVVN
DTVGTMMTCAYEDPNCEIGLIAGTGSNVCYMEEMRNIELVEEDEGKMCINTEWGGFGDNGCIDDIRTQYD
KEVDEGSLNPGKQRYEKMTSGMYLGEIVRQILIDLTKQGLLFRGQISERLRTRGIFETKFLSQIESDRLA
LLQVRGILQQLGLDSTCEDSIVVKEVCGAVSRRAAQLCGAGLAAIVEKRREDQGLEHLKITVGVDGTLYK

LHPHFSRILQETVKELAPQCDVTFMLSEDGSGKGAALITAVAKRLQQAQKEN
>anc66_altAll|avgPP:0.995|lnPP:-8.368|num_ambig:7|num_ambig_gaps:0
LYHMRLSDETLLDIMARFQAEMEKGLGKDTNPTASVKMLPTFVRAIPDGSENGEFLSLDLGGSKFRVLKV
QVSEEGKRNVQMESQFYPTPNEIIRGNGSELFDYVADCLADFMKTKDLKHKKLPLGLTFSFPCRQTKLEE
GVLLSWTKKFKARGVQDTDVVSRLTKAMKKHPDMDVDILALVNDTVGTMMTCAYDDPYCEVGVIIGTGTN
ACYMEDMSNIDLVEGDEGRMCINTEWGAFGDDGALEDIRTEFDRELDLGSLNPGKQLFEKMISGLYLGEL
VRLILLKMARAGLLFGGKKSSALHTKGKIETRHVAAMEKYKEGLANTREILTDLGLEPSEADCIAVQHVC
TIVSFRSANLCAAALAAILTRLRENKKLVRLRTTVGMDGTLYKIHPQYPKRLHKVVRKLVPNCDVRFLLS
ESGSAKGAAMVTAVASRVQAQRKQIDEVLALFRLTREHLVDVQDKMRAELEYGLKKETHPMATVKMLPTY
VYGMPDGTEKGKFLALDLGGTNFRVLLVKIRRGRKSVRMYNKIFAIPLEIMQGTGEELFDHIVQCIADFL
DYMGLKGAQLPLGFTFSFPCRQTSIDKGILIEWTKGFKATDCEGEDVVDMLREAIKRRNEFDLDIVAVVN
DTVGTMMTCAYEDPNCEIGLIAGTGSNVCYMEEMRNIELVEEDEGKMCINTEWGGFGDNGCIDDIRTQYD
KEVDEGSLNPGKQRYEKMTSGMYLGEIVRQILIDLTKQGLLFRGQVSERLRTRGIFETKFLSQIESDRLA
LLQVRGILQQLGLDSTCEDSIVVKEVCGAVSRRAAQLCGAGLAAIVEKRREDQGLEHLKITVGVDGTLYK
LHPHFSRILQETVKELAPQCDVTFMLSEDGSGKGAALITAVAKRLQQPQKEN
>anc67|avgPP:0.997|lnPP:-4.408|num_ambig:5|num_ambig_gaps:0
LYHMRLSDETLLDIMARFQAEMQKGLGKDTNPTASVKMLPTFVRAIPDGSENGEFLSLDLGGSKFRVLKV
QVSEEGKRKVQMESQFYPTPNEIIRGNGSELFEYVADCLADFMKTKGLKHKKLPLGLTFSFPCRQTKLEE
GVLLSWTKKFKARGVQDTDVVSRLTKAMKKHKDIDVDILALVNDTVGTMMTCAYDDPYCEVGVIIGTGTN
ACYMEDMSNIDLVEGDEGRMCINTEWGAFGDDGALEDIRTEFDRELDLGSLNPGKQLFEKMISGLYLGEL
VRLILLKMAKAGLLFGGKKSSALHTKGKIETRHVAAMEKYKEGLANTREILTDLGLEPSEADCIAVQHVC
TIVSFRSANLCAAALAAILTRLRENKKLVRLRTTVGVDGTLYKIHPQYPKRLHKVVRRLVPNCDVRFLLS
ESGSTKGAAMVTAVASRVQAQRKQIDQVLALFRLTREQLVDVQDKMRAELEYGLKKDTHLLATVKMLPTY
VCGMPDGTEKGKFLALDLGGTNFRVLLVKIRSGRRSVRMYNKIFAIPLEIMQGTGEELFDHIVQCIADFL
DYMGLKGAQLPLGFTFSFPCRQTSIDKGILIEWTKGFKATDCEGEDMVDMLREAIKRRNEFDLDIVAVVN
DTVGTMMTCGYEDPNCEIGLIAGTGSNMCYMEEMRNIEMVEGDEGKMCINTEWGGFGDNGCIDDIRTQYD
KEVDEGSLNPGKQRYEKMTSGMYLGEIVRQILIDLTKQGLLFRGQISERLRTRGIFETKFLSQIESDRLA
LLQVRRILQQLGLDSTCEDSIVVKEVCGAVSRRAAQLCGAGLAAIVEKRREDQGLEHLKITVGVDGTLYK
LHPHFSRILQETVKELAPQCDVTFMLSEDGSGKGAALITAVAKRLQQAQKEN
>anc67_altAll|avgPP:0.996|lnPP:-6.786|num_ambig:5|num_ambig_gaps:0
LYHMRLSDETLLDIMARFQAEMQKGLGKDTNPTASVKMLPTFVRAIPDGSENGEFLSLDLGGSKFRVLKV
QVSEEGKRKVQMESQFYPTPNEIIRGNGSELFEYVADCLADFMKTKGLKHKKLPLGLTFSFPCRQTKLEE
GVLLSWTKKFKARGVQDTDVVSRLTKAMKKHKDIDVDILALVNDTVGTMMTCAYDDPYCEVGVIIGTGTN
ACYMEDMSNIDLVEGDEGRMCINTEWGAFGDDGALEDIRTEFDRELDLGSLNPGKQLFEKMISGLYLGEL
VRLILLKMAKAGLLFGGKKSSALHTKGKIETRHVAAMEKYKEGLANTREILTDLGLEPSEADCIAVQHVC
TIVSFRSANLCAAALAAILTRLRENKKLARLRTTVGVDGTLYKIHPQYPKRLHKVVRRLVPNCDVRFLLS
ESGSTKGAAMVTAVASRVQAQRKQIDQVLALFRLTREQLVGVQDKMRAELEYGLKRDTHLLATVKMLPTY
VCGMPDGTEKGKFLALDLGGTNFRVLLVKIRSGRRSVRMYNKIFAIPLEIMQGTGEELFDHIVQCIADFL
DYMGLKGAQLPLGFTFSFPCRQTSIDKGILIEWTKGFKATDCEGEDMVDMLREAIKRRNEFDLDIVAVVN
DTVGTMMTCGYEDPNCEIGLIAGTGSNMCYMEEMRNIELVEGDEGKMCINTEWGGFGDNGCIDDIRTQYD
KEVDEGSLNPGKQRYEKMTSGMYLGEIVRQILIDLTKQGLLFRGQISERLRTRGIFETKFLSQIESDRLA
LLQVRRILQQLGLDSTCEDSIVVKEVCGAVSRRAAQLCGAGLAAIVEKRREDQGLEHLKITVGVDGTLYK
LHPHFSRILQETVKELAPRCDVTFMLSEDGSGKGAALITAVAKRLQQAQKEN
>anc68|avgPP:0.997|lnPP:-5.185|num_ambig:3|num_ambig_gaps:0
LYHMRLSDETLLDIMARFQAEMQKGLDKDTNPTASVKMLPTFVRAIPDGSENGEFLSLDLGGSKFRVLKV
QVSEEGKRNVQMESQFYPTPNEIIRGNGSELFEYVADCLADFMKTKGLKHKKLPLGLTFSFPCRQTKLEE
GVLLSWTKKFKARGVQDTDVVSRLKKAMKKHKQDIDVDILALVNDTVGTMMTCAYDDPYCEVGVIIGTGT
NACYMEDMSNIDLVEGDEGRMCINTEWGAFGDDGALEDIRTEFDRELDLGSLNPGKQLFEKMISGLYLGE
LVRLILLKMAKAGLLFGGKKSSALHTKGKIETRHVAAMEKYKEGLANTREILTDLGLEPSEADCIAVQHV
CTIVSFRSANLCAAALAAILTRLQENKKLARLRTTVGMDGTLYKIHPQYPKRLHKVVRRLVPNCDVRFLL
SESGSTKGAAMVTAVACRLQAQRKQIDQVLALFRLTREQLVGVQDKMRAELEYGLKRDTHPLATVKMLPT
YVYGMPDGTEKGKFLALDLGGTNFRVLLVKIRSGRRSVRMYNKIFAIPLEIMQGTGEELFDHIVQCIADF

LDYMGLKGAQLPLGFTFSFPCRQTSIDKGILIEWTKGFKATDCEGEDVVDMLREAIKRRNEFDLDIVAVV
NDTVGTMMTCGYEDPNCEIGLIAGTGSNMCYMEEMRNIEMVEGDEGKMCINTEWGGFGDNGCIDDIRTQY
DKEVDEGSLNPGKQRYEKMTSGMYLGEIVRQILIDLTKQGLLFRGQISERLRTRGIFETKFLSQIESDRL
ALLQVRSILQQLGLDSTCEDSIVVKEVCGAVSRRAAQLCGAGLAAIVEKRREDQGLEHLKITVGVDGTLY
KLHPHFSRILQETVKELAPQCDVTFMLSEDGSGKGAALITAVAKRLQQAQKEN
>anc68_altAll|avgPP:0.996|lnPP:-6.187|num_ambig:3|num_ambig_gaps:0
LYHMRLSDETLLDIMARFQAEMQKGLDKDTNPTASVKMLPTFVRAIPDGSENGEFLSLDLGGSKFRVLKV
QVSEEGKRNVQMESQFYPTPNEIIRGNGSELFEYVADCLADFMKTKGLKHKKLPLGLTFSFPCRQTKLEE
GVLLSWTKKFKARGVQDTDVVSSLKKAMKKHKQDIDVDILALVNDTVGTMMTCAYDDPYCEVGVIIGTGT
NACYMEDMSNIDLVEGDEGRMCINTEWGAFGDDGALEDIRTEFDRELDLGSLNPGKQLFEKMISGLYLGE
LVRLILLKMAKAGLLFGGKKSSALHTKGKIETRHVAAMEKYKEGLANTREILTDLGLEPSEADCIAVQHV
CTIVSFRSANLCAAALAAILTRLQENKKLVRLRTTVGMDGTLYKIHPQYPKRLHKVVRKLVPNCDVRFLL
SESGSTKGAAMVTAVACRLQAQRKQIDQVLALFRLTREQLVGVQDKMRAELEYGLKRDTHPLATVKMLPT
YVYGMPDGTEKGKFLALDLGGTNFRVLLVKIRSGRRSVRMYNKIFAIPLEIMQGTGEELFDHIVQCIADF
LDYMGLKGAQLPLGFTFSFPCRQTSIDKGILIEWTKGFKATDCEGEDVVDMLREAIKRRNEFDLDIVAVV
NDTVGTMMTCGYEDPNCEIGLIAGTGSNMCYMEEMRNIEMVEGDEGKMCINTEWGGFGDNGCIDDIRTQY
DKEVDEGSLNPGKQRYEKMTSGMYLGEIVRQILIDLTKQGLLFRGQISERLRTRGIFETKFLSQIESDRL
ALLQVRSILQQLGLDSTCEDSIVVKEVCGAVSRRAAQLCGAGLAAIVEKRREDQGLEHLKITVGVDGTLY
KLHPHFSRILQETVKELAPQCDVTFMLSEDGSGKGAALITAVAKRLQQAQKEN
>anc69|avgPP:0.997|lnPP:-3.815|num_ambig:6|num_ambig_gaps:0
LYHMRLSDETLLDIMARFQAEMQKGLGKDTNPTASVKMLPTFVRAIPDGSENGEFLSLDLGGSKFRVLKV
QVSEEGKRNVQMESQFYPTPNEIIRGNGSELFEYVADCLADFMKTKGLKHKKLPLGLTFSFPCRQTKLEE
GVLLSWTKKFKARGVQDTDVVSRLTKAMKKHKDIDVDILALVNDTVGTMMTCAYDDPYCEVGVIIGTGTN
ACYMEDMSNIDLVEGDEGRMCINTEWGAFGDDGALEDIRTEFDRELDLGSLNPGKQLFEKMISGLYLGEL
VRLILLKMAKAGLLFGGKKSSALHTKGKIETRHVAAMEKYKEGLANTREILTDLGLEPSEADCIAVQHVC
TIVSFRSANLCAAALAAILTRLRENKKLVRLRTTVGMDGTLYKIHPQYPKRLHKVVRRLVPNCDVRFLLS
ESGSTKGAAMVTAVASRVQAQRKQIDQVLALFRLTREQLVDVQDKMRAELEYGLKKDTHPLATVKMLPTY
VCGMPDGTEKGKFLALDLGGTNFRVLLVKIRSGRRSVRMYNKIFAIPLEIMQGTGEELFDHIVQCIADFL
DYMGLKGAQLPLGFTFSFPCRQTSIDKGILIEWTKGFKATDCEGEDVVDMLREAIKRRNEFDLDIVAVVN
DTVGTMMTCGYEDPNCEIGLIAGTGSNMCYMEEMRNIEMVEGDEGKMCINTEWGGFGDNGCIDDIRTQYD
KEVDEGSLNPGKQRYEKMTSGMYLGEIVRQILIDLTKQGLLFRGQISERLRTRGIFETKFLSQIESDRLA
LLQVRRILQQLGLDSTCEDSIVVKEVCGAVSRRAAQLCGAGLAAIVEKRREDQGLEHLKITVGVDGTLYK
LHPHFSRILQETVKELAPQCDVTFMLSEDGSGKGAALITAVAKRLQQAQKEN
>anc69_altAll|avgPP:0.996|lnPP:-7.699|num_ambig:6|num_ambig_gaps:0
LYHMRLSDETLLDIMARFQAEMQKGLGKDTNPTASVKMLPTFVRAIPDGSENGEFLSLDLGGSKFRVLKV
QVSEEGKRNVQMESQFYPTPNEIIRGNGSELFEYVADCLADFMKTKGLKHKKLPLGLTFSFPCRQTKLEE
GVLLSWTKKFKARGVQDTDVVSRLTKAMKKHKDIDVDILALVNDTVGTMMTCAYDDPYCEVGVIIGTGTN
ACYMEDMSNIDLVEGDEGRMCINTEWGAFGDDGALEDIRTEFDRELDLGSLNPGKQLFEKMISGLYLGEL
VRLILLKMAKAGLLFGGKKSSALHTKGKIETRHVAAMEKYKEGLANTREILTDLGLEPSEADCIAVQHVC
TIVSFRSANLCAAALAAILTRLRENKKLARLRTTVGMDGTLYKIHPQYPKRLHKVVRKLVPNCDVRFLLS
ESGSTKGAAMVTAVASRVQAQRKQIDQVLALFRLTREQLVGVQDKMRAELEYGLKRDTHPLATVKMLPTY
VCGMPDGTEKGKFLALDLGGTNFRVLLVKIRSGRRSVRMYNKIFAIPLEIMQGTGEELFDHIVQCIADFL
DYMGLKGAQLPLGFTFSFPCRQTSIDKGILIEWTKGFKATDCEGEDVVDMLREAIKRRNEFDLDIVAVVN
DTVGTMMTCGYEDPNCEIGLIAGTGSNMCYMEEMRNIELVEGDEGKMCINTEWGGFGDNGCIDDIRTQYD
KEVDEGSLNPGKQRYEKMTSGMYLGEIVRQILIDLTKQGLLFRGQISERLRTRGIFETKFLSQIESDRLA
LLQVRRILQQLGLDSTCEDSIVVKEVCGAVSRRAAQLCGAGLAAIVEKRREDQGLEHLKITVGVDGTLYK
LHPHFSRILQETVKELAPRCDVTFMLSEDGSGKGAALITAVAKRLQQAQKEN
>anc70|avgPP:0.997|lnPP:-3.759|num_ambig:4|num_ambig_gaps:0
LYHMRLSDETLLDIMARFQAEMEKGLGKDTNPTASVKMLPTFVRAIPDGSENGEFLSLDLGGSKFRVLKV
QVSEEGKRNVQMESQFYPTPNEIIRGNGSELFEYVADCLADFMKTKDLKHKKLPLGLTFSFPCRQTKLEE
GVLLSWTKKFKARGVQDTDVVSRLTKAMKKHKDIDVDILALVNDTVGTMMTCAYDDPYCEVGVIIGTGTN
ACYMEDMSNIDLVEGDEGRMCINTEWGAFGDDGALEDIRTEFDRELDLGSLNPGKQLFEKMISGLYLGEL

VRLILLKMAKAGLLFGGKKSSALHTKGKIETRHVAAMEKYKEGLANTREILTDLGLEPSEADCIAVQHVC
TIVSFRSANLCAAALAAILTRLRENKKLVRLRTTVGMDGTLYKIHPQYPKRLHKVVRKLVPNCDVRFLLS
ESGSTKGAAMVTAVASRVQAQRKQIDQVLALFRLTREQLVDVQDKMRAELEYGLKKETHPLATVKMLPTY
VCGMPDGTEKGKFLALDLGGTNFRVLLVKIRSGRRSVRMYNKIFAIPLEIMQGTGEELFDHIVQCIADFL
DYMGLKGAQLPLGFTFSFPCRQTSIDKGILIEWTKGFKATDCEGEDVVDMLREAIKRRNEFDLDIVAVVN
DTVGTMMTCGYEDPNCEIGLIAGTGSNMCYMEEMRNIEMVEGDEGKMCINTEWGGFGDNGCIDDIRTQYD
KEVDEGSLNPGKQRYEKMTSGMYLGEIVRQILIDLTKQGLLFRGQISERLRTRGIFETKFLSQIESDRLA
LLQVRRILQQLGLDSTCEDSIVVKEVCGAVSRRAAQLCGAGLAAIVEKRREDQGLEHLKITVGVDGTLYK
LHPHFSRILQETVKELAPQCDVTFMLSEDGSGKGAALITAVAKRLQQAQKEN
>anc70_altAll|avgPP:0.996|lnPP:-6.459|num_ambig:4|num_ambig_gaps:0
LYHMRLSDETLLDIMARFQAEMEKGLGKDTNPTASVKMLPTFVRAIPDGSENGEFLSLDLGGSKFRVLKV
QVSEEGKRNVQMESQFYPTPNEIIRGNGSELFEYVADCLADFMKTKDLKHKKLPLGLTFSFPCRQTKLEE
GVLLSWTKKFKARGVQDTDVVSRLTKAMKKHKDMDVDILALVNDTVGTMMTCAYDDPYCEVGVIIGTGTN
ACYMEDMSNIDLVEGDEGRMCINTEWGAFGDDGALEDIRTEFDRELDLGSLNPGKQLFEKMISGLYLGEL
VRLILLKMAKAGLLFGGKKSSALHTKGKIETRHVAAMEKYKEGLANTREILTDLGLEPSEADCIAVQHVC
TIVSFRSANLCAAALAAILTRLRENKKLARLRTTVGMDGTLYKIHPQYPKRLHKVVRKLVPNCDVRFLLS
ESGSTKGAAMVTAVASRVQAQRKQIDQVLALFRLTREQLVDVQDKMRAELEYGLKKETHPLATVKMLPTY
VCGMPDGTEKGKFLALDLGGTNFRVLLVKIRSGRRSVRMYNKIFAIPLEIMQGTGEELFDHIVQCIADFL
DYMGLKGAQLPLGFTFSFPCRQTSIDKGILIEWTKGFKATDCEGEDVVDMLREAIKRRNEFDLDIVAVVN
DTVGTMMTCGYEDPNCEIGLIAGTGSNMCYMEEMRNIELVEGDEGKMCINTEWGGFGDNGCIDDIRTQYD
KEVDEGSLNPGKQRYEKMTSGMYLGEIVRQILIDLTKQGLLFRGQISERLRTRGIFETKFLSQIESDRLA
LLQVRRILQQLGLDSTCEDSIVVKEVCGAVSRRAAQLCGAGLAAIVEKRREDQGLEHLKITVGVDGTLYK
LHPHFSRILQETVKELAPRCDVTFMLSEDGSGKGAALITAVAKRLQQAQKEN
>anc71|avgPP:0.997|lnPP:-3.934|num_ambig:4|num_ambig_gaps:0
LYHMRLSDETLLDIMARFQAEMEKGLGKDTNPTASVKMLPTFVRAIPDGSENGEFLSLDLGGSKFRVLKV
QVSEEGKRNVQMESQFYPTPNEIIRGNGSELFEYVADCLADFMKTKDLKHKKLPLGLTFSFPCRQTKLEE
GVLLSWTKKFKARGVQDTDVVSRLTKAMKKHKDIDVDILALVNDTVGTMMTCAYDDPYCEVGVIIGTGTN
ACYMEDMSNIDLVEGDEGRMCINTEWGAFGDDGALEDIRTEFDRELDLGSLNPGKQLFEKMISGLYLGEL
VRLILLKMAKAGLLFGGKKSSALHTKGKIETRHVAAMEKYKEGLANTREILTDLGLEPSEADCIAVQHVC
TIVSFRSANLCAAALAAILTRLRENKKLVRLRTTVGMDGTLYKIHPQYPKRLHKVVRKLVPNCDVRFLLS
ESGSTKGAAMVTAVASRVQAQRKQIDEVLALFRLTREHLVDVQDKMRAELEYGLKKETHPLATVKMLPTY
VCGMPDGTEKGKFLALDLGGTNFRVLLVKIRSGRKSVRMYNKIFAIPLEIMQGTGEELFDHIVQCIADFL
DYMGLKGAQLPLGFTFSFPCRQTSIDKGILIEWTKGFKATDCEGEDVVDMLREAIKRRNEFDLDIVAVVN
DTVGTMMTCGYEDPNCEIGLIAGTGSNVCYMEEMRNIELVEGDEGKMCINTEWGGFGDNGCIDDIRTQYD
KEVDEGSLNPGKQRYEKMTSGMYLGEIVRQILIDLTKQGLLFRGQISERLRTRGIFETKFLSQIESDRLA
LLQVRRILQQLGLDSTCEDSIVVKEVCGAVSRRAAQLCGAGLAAIVEKRREDQGLEHLKITVGVDGTLYK
LHPHFSRILQETVKELAPQCDVTFMLSEDGSGKGAALITAVAKRLQQAQKEN
>anc71_altAll|avgPP:0.996|lnPP:-6.203|num_ambig:4|num_ambig_gaps:0
LYHMRLSDETLLDIMARFQAEMEKGLGKDTNPTASVKMLPTFVRAIPDGSENGEFLSLDLGGSKFRVLKV
QVSEEGKRNVQMESQFYPTPNEIIRGNGSELFDYVADCLADFMKTKDLKHKKLPLGLTFSFPCRQTKLEE
GVLLSWTKKFKARGVQDTDVVSRLTKAMKKHKDMDVDILALVNDTVGTMMTCAYDDPYCEVGVIIGTGTN
ACYMEDMSNIDLVEGDEGRMCINTEWGAFGDDGALEDIRTEFDRELDLGSLNPGKQLFEKMISGLYLGEL
VRLILLKMAKAGLLFGGKKSSALHTKGKIETRHVAAMEKYKEGLANTREILTDLGLEPSEADCIAVQHVC
TIVSFRSANLCAAALAAILTRLRENKKLVRLRTTVGMDGTLYKIHPQYPKRLHKVVRKLVPNCDVRFLLS
ESGSAKGAAMVTAVASRVQAQRKQIDEVLALFRLTREHLVDVQDKMRAELEYGLKKETHPLATVKMLPTY
VCGMPDGTEKGKFLALDLGGTNFRVLLVKIRSGRRSVRMYNKIFAIPLEIMQGTGEELFDHIVQCIADFL
DYMGLKGAQLPLGFTFSFPCRQTSIDKGILIEWTKGFKATDCEGEDVVDMLREAIKRRNEFDLDIVAVVN
DTVGTMMTCGYEDPNCEIGLIAGTGSNVCYMEEMRNIELVEGDEGKMCINTEWGGFGDNGCIDDIRTQYD
KEVDEGSLNPGKQRYEKMTSGMYLGEIVRQILIDLTKQGLLFRGQISERLRTRGIFETKFLSQIESDRLA
LLQVRRILQQLGLDSTCEDSIVVKEVCGAVSRRAAQLCGAGLAAIVEKRREDQGLEHLKITVGVDGTLYK
LHPHFSRILQETVKELAPQCDVTFMLSEDGSGKGAALITAVAKRLQQAQKEN
>anc72|avgPP:0.990|lnPP:-15.401|num_ambig:20|num_ambig_gaps:0

LYHMRLSDDTLLDIMARFQAEMEKGLGKDTNPTASVKMLPTFVRAIPDGSEKGEFLALDLGGSKFRVLKV
QVSEEGKQNVQMESQFYPTPKEIIRGNGSELFDYVADCLADFMKTKDIKHKKLPLGLTFSFPCKQTKLEE
GVLLSWTKKFKVRGVQDTDVVSSLTKALKKHKDIDVDVLALVNDTVGTMMTCAYDDPHCEVGVIIGTGTN
ACYMEDMSNIDLVEGDEGRMCINTEWGAFGDDGSLEDIRTEFDREIDLGSINPGKQLFEKMISGLYLGEL
VRLILLKMAKAGLLFGGKKSTALHTKGKIETQHVAAMEKYKEGLANTREILMELGLEPSEADCIAVQHVC
TIVSFRSANLCAAALAAILTRLRENKKLVRLRTTVGMDGTLYKTHPQYPKRLHKVVRKLVPNCDVRFLLS
ESGSAKGAAMVTAVASRVQAQRKQIDEVLAMFKLTREHLVDVKNKMRAELEYGLKKETHPLATVKMLPTY
VCGMPDGTEKGKFLALDLGGTNFRVLLVKIRSGRKSVRMYNKIFAIPLEIMQGTGEELFDHIVQCIADFL
DYMGIKGARLPLGFTFSFPCRQTSIDKGILIEWTKGFKATDCEGEDVVDMLREAIKRRNEFDLDIVAVVN
DTVGTMMTCGYEDPNCEIGLIAGTGSNVCYMEEMRNIELVEGDEGKMCINTEWGGFGDNGCIDDIRTQYD
KEVDEGSLNPGKQRYEKMTSGMYLGEIVRQILIDLTKQGLLFRGQISERLRTRGIFETKFLSQIESDRLA
LLQVRRILQQLGLDSTCEDSIVVKEVCGAVSRRAAQLCGAGLAAIVEKRREDRGLEHLKITVGVDGTLYK
LHPHFSRILQETVKELAPQCDVTFMLSEDGSGKGAALITAVAKRLHQAKEN
>anc72_altAll|avgPP:0.985|lnPP:-27.943|num_ambig:20|num_ambig_gaps:0
LYHMRLSDDTLLDIMTRFQAEMEKGLGKDTNPTATVKMLPTFVRAIPDGSEKGEFLALDLGGSKFRVLKV
QVSEEGKQNVQMESQFYPTPNEIIRGNGSELFEYVADCLADFMKTKNIKHKKLPLGLTFSFPCRQTKLEE
GVLLSWTKKFKVRGVQDTDVVSSLTKALKKHKDIDVDVLALVNDTVGTMMTCAYDDPHCEVGVIIGTGTN
ACYMEDMSNIDLVEGDEGRMCINTEWGAFGDDGSLEDIRTEFDRELDLGSINPGKQLFEKMISGLYLGEL
VRLILLKMAKAGLLFGGKKSSALHTKGKIETQHVAAMEKYKEGLSNTREILTDLGLEPSEADCIAVQHVC
TIVSFRSANLCAAALAAILTRLRENKKLVRLRTTVGMDGTLYKTHPQYPKRLHKVVRKLVPNCDVRFLLS
ESGSAKGAAMVTAVASRVQAQRKQIDEVLALFRLTREHLVSVKNKMRAELEYGLKKETHPLATVKMLPTY
VCGMPDGTEKGKYLALDLGGTNFRVLLVKIRSGRRSVRMYNKIFAIPLEIMQGTGEELFDHIVQCIADFL
DYMGLKGARLPLGFTFSFPCRQTSIDKGILVEWTKGFKATDCEGEDVVDMLREAIKRRNEFDLDIVAVVN
DTVGTMMTCGYEDPNCEIGLIAGTGSNVCYMEEMRNIELVEGDEGKMCINTEWGGFGDNGCLDDIRTQYD
KEVDEGSLNPGKQRYEKMTSGMYLGEIVRQILIDLTKQGLLFRGQISERLRTRGIFETKFLSQIESDRLA
LLQVRRILQQLGLDSTCEDSIVVKEVCGAVSRRAAQLCGAGLAAIVEKRREDRGLEHLKITVGVDGTLYK
LHPHFSRILQETVKELAPQCDVTFMLSEDGSGKGAALITAVAKRLQQAKEN
>anc73|avgPP:0.989|lnPP:-16.256|num_ambig:19|num_ambig_gaps:0
LYHMRLSDDTLLDIMARFQAEMEKGLGKDTNPTATVKMLPTFVRAIPDGSEKGEFLALDLGGSKFRVLKV
KVSEDGKQNVQMESQFYPTPKEIIRGNGSELFDYVADCLADFMKTKDIKHKKLPLGFTFSFPCKQTKLEE
GVLLSWTKKFKVRGVQGTDVVSSLTKALKKHKDIDVDVLALVNDTVGTMMTCAYDDPHCEVGVIIGTGTN
ACYMEDMSNIDLVEGDEGRMCINTEWGAFGDDGSLEDIRTEFDREIDLGSINPGKQLFEKMISGLYLGEL
VRLILLKMAKEGLLFGGKTSTALHTKGKIETQHVAAMEKYKEGLSNTKEILMELGLEPSEADCIAVQHVC
TIVSFRSANLCAAALAAILTRLRENKKLVRLRTTVGMDGTLYKTHPQYPKRLHKVVRRLVPNCDVRFLLS
ESGSAKGAAMVTAVASRVQAQRKQIDEVLAMFKLSREHLVDVKNKMRAELEYGLKKETHPLATVKMLPTY
VCGMPDGTEKGKFLALDLGGTNFRVLLVKIRSGRKSVRMYNKIFAIPLEIMQGTGEELFDHIVQCIADFL
DYMGIKGARLPLGFTFSFPCRQTSIDKGILIGWTKGFKATDCEGEDVVDMLREAIKRRNEFDLDIVAVVN
DTVGTMMTCGYEDPNCEIGLIAGTGSNVCYMEEMRNIELVEGDEGKMCINTEWGGFGDNGCIDDIRTQYD
KEVDEGSLNPGKQRYEKMTSGMYLGEIVRQILIDLTKQGLLFRGQISERLRTRGIFETKFLSQIESDRLA
LLQVRRILQQLGLDSTCEDSIVVKEVCGAVSRRAAQLCGAGLAAIVEKRREDRGLEHLKITVGVDGTLYK
LHPHFSRILQETVKELAPQCDVTFMLSEDGSGKGAALITAVAKRLHQAKEN
>anc73_altAll|avgPP:0.985|lnPP:-27.441|num_ambig:19|num_ambig_gaps:0
LYHMRLSDDTLLDIMTRFQAEMEKGLGRDTNPTATVKMLPTFVRAIPDGSEKGEFLALDLGGSKFRVLKV
KVSEDGKQNVQMESQFYPTPKEIIHGNGSELFEYVADCLADFMKTKNIKHKKLPLGFTFSFPCKQTKLEE
GVLLSWTKKFKVRGVQGTDVVSSLTKALKKHKDIDVDVLALVNDTVGTMMTCAYDDPHCEVGVIIGTGTN
ACYMEDMSNIDLVEGDEGRMCINTEWGAFGDDGSLEDIRTEFDRELDLGSINPGKQLFEKMISGLYLGEL
VRLILLKMAKEGLLFGGKKSTALHTKGKIETQHVAAMEKYKEGLANTKEILTELGLEPSEADCIAVQHVC
TIVSFRSANLCAAALAAILTRLRENKKLARLRTTVGMDGTLYKTHPQYPKRLHKVVRRLVPNCDVRFLLS
ESGSAKGAAMVTAVASRVQAQRKQIDEVLALFKLTREHLVSVKNKMRAELEYGLKKETHPLATVKMLPTY
VCGMPDGTEKGKYLALDLGGTNFRVLLVKIRSGRKSVRMYNKIFAIPLEIMQGTGEELFDHIVQCIADFL
DYMGLKGARLPLGFTFSFPCRQTSIDKGILVGWTKGFKATDCEGEDVVDMLREAIKRRNEFDLDIVAVVN
DTVGTMMTCGYEDPNCEIGLIAGTGSNVCYMEEMRNIELVEGDEGKMCINTEWGGFGDNGCLDDIRTQYD

```
KEVDEGSLNPGKQRYEKMTSGMYLGEIVRQILIDLTKQGLLFRGQISERLRTRGIFETKFLSQIESDRLA
LLQVRRILQQLGLDSTCEDSIVVKEVCGAVSRRAAQLCGAGLAAVVEKRREDRGLEHLKITVGVDGTLYK
LHPHFSRILQETVKELAPQCDVTFMLSEDGSGKGAALITAVAKRLQQAKEN
>anc74|avgPP:0.997|lnPP:-4.040|num_ambig:5|num_ambig_gaps:0
LYHLRLSDDVLLDVMARFQAEMVKGLGRDTNPTATVKMLPSFVRSLPDGSEKGDFLAVDLGGSQFRAHQV
KVFDDGKQSSQLESKFYPTPKEVTQGNGAELFDYIADCLLDFMETKNLKHKKLPLGFTFSFPCKQTKLEE
GVLLAWTKHFKVRGVQDTDVVSSLRKALQKHKDIDVDVLALVNDTVGTMMTCGYDDQRCEVGLIIGTGTN
ACYMEEMRHIDLVEGDEGRMCINTEWGAFGDDGALDDLRTEFDRELDLGSLNPGKQLFEKMISSLYLGEL
VRLILLKMTKEGLLFNGKVSTALLTKGKIEMKHVSAMEKYKEGLSNTKEILTELNLFPSEEDCVAVQHVC
TIVSFRSANLCAAALAAILTRLRENKKLLRMRTTVGIDGGLYKTHPQYAKRLHKVVRRLVPTCDVRFLLS
VSGSGRGAAMVTAVAHRLAAQRKQIDATLAPFLLPLGTLREVKDKMRAELEYGLKRETQGNATVKMLPTY
VCGTPDGTEKGKFLALDLGGTNFRVLLVKIKSGRRRSVQMYNKIFAIPLEIMQGTGEELFDHIVQCIADF
LEYMGIKGARLPLGFTFSFPCRQASIDKGTLVGWTKGFKATDCEGEDVVDMLREAIRRRNEFDLDIVAVV
NDTVGTMMTCGYEDPNCEIGLIAGTGSNVCYMEDMKNIEIVEGNEGKMCINTEWGAFGDNGCIDNIRTKY
DKEVDEGSLNPGKQRYEKMTSGMYLGEIVRQILIDLTKQGLLFRGHISESLRKRGIFETKFLSQIESDRL
ALLQVRRILQQLGLDSTCDDSIIVKEVCGVVSRRAAQLCGAGLAAIVEKKRENRGVERLQITVGVDGTLY
KLHPHFSRVLQETVKELAPQCDVTFMLSEDGSGKGAALITAVAKRLHNVGQK
>anc74_altAll|avgPP:0.996|lnPP:-7.332|num_ambig:5|num_ambig_gaps:0
LYHLRLSDDVLLDVMARFQAEMVKGLGRDTNPTATVKMLPSFVRSLPDGSEKGDFLAVDLGGSQFRAHQV
KVFDDGKQSSQLESKFYPTPKEVTQGNGAELFDYIADCLLDFMETKNLKHKKLPLGFTFSFPCKQTKLEE
GVLLAWTKHFKVRGVQDTDVVSSLRKALQKHKDIDVDVLALVNDTVGTMMTCGYDDQRCEVGLIIGTGTN
ACYMEEMRHIDLVEGDEGRMCINTEWGAFGDDGALDDLRTEFDRELDLGSLNPGKQLFEKMISSLYLGEL
VRLILLKMTKEGLLFNGKVSTALLTKGKIEMKHVSAMEKYKEGLSNTKEILTELNLFPSEEDCVAVQHVC
TIVSFRSANLCAAALAAILTRLRENKKLLRMRTTVGIDGGLYKTHPQYAKRLHKVVRRLVPTCDVRFLLS
VSGSGRGAAMVTAVARRLAAQRQQIDAALAPFLLPLGTLREVKDKMRAELEYGLKRETQGNATVKMLPTY
VCGTPDGTEKGKFLALDLGGTNFRVLLVKIKSGRRRSVQMYNKIFAIPLEIMQGTGEELFDHIVQCIADF
LEYMGIKGARLPLGFTFSFPCRQASIDKGTLVGWTKGFKATDCEGEDVVDMLREAIRRRNEFDLDIVAVV
NDTVGTMMTCGYEDPNCEIGLIAGTGSNVCYMEDMKNIEIVEGNEGKMCINTEWGAFGDNGCIDNIRTKY
DKEVDEGSLNPGKQRYEKMTSGMYLGEIVRQILIDLTKQGLLFRGHISESLRKRGIFETKFLSQIESDRL
ALLQVRRILQQLGLDSTCEDSIIVKEVCGAVSRRAAQLCGAGLAAIVEKKRENRGVERLQITVGVDGTLY
KLHPHFSRVLQETVKELAPQCDVTFMLSEDGSGKGAALITAVAKRLHNVGQK
>anc75|avgPP:0.998|lnPP:-2.903|num_ambig:4|num_ambig_gaps:0
LYHLRLSDDVLLDVMARFQAEMVKGLGRDTNPTATVKMLPSFVRSLPDGSEKGDFLAVDLGGSQFRAHQV
KVFDDGKQSSQLESKFYPTPKEVTQGNGAELFDYIADCLLDFMETKNLKHKKLPLGFTFSFPCKQTKLEE
GVLLAWTKHFKVRGVQDTDVVSSLRKALQKHKDIDVDVLALVNDTVGTMMTCGYDDQRCEVGLIIGTGTN
ACYMEEMRHIDLVEGDEGRMCINTEWGAFGDDGALDDLRTEFDRELDLGSLNPGKQLFEKMISSLYLGEL
VRLILLKMTKEGLLFNGKVSTALLTKGKIEMKHVSAMEKYKEGLSNTKEILTELNLFPSEEDCIAVQHVC
TIVSFRSANLCAAALAAILTRLRENKKLLRMRTTVGIDGGLYKTHPQYAKRLHKVVRRLVPNCDVRFLLS
VSGSGKGAAMVTAVAHRLAAQRKQIDATLAPFLLSLETLREVKDKMRTELEYGLKRETQANATVKMLPTY
VCGTPDGTEKGKFLALDLGGTNFRVLLVKIRSGRRRSVQMYNKIFAIPLEIMQGTGEELFDHIVQCIADF
LEYMGIKGARLPLGFTFSFPCRQASIDKGTLVGWTKGFKATDCEGEDVVDMLREAIRRRNEFDLDIVAVV
NDTVGTMMTCGYEDPNCEIGLIAGTGSNVCYMEDMKNIEIVEGNEGKMCINTEWGAFGDNGCIDNIRTKY
DKEVDEGSLNPGKQRYEKMTSGMYLGEIVRQILIDLTKQGLLFRGHISESLRKRGIFETKFLSQIESDRL
ALLQVRRILQQLGLDSTCDDSIIVKEVCGVVSRRAAQLCGAGLAAIVEKKRENRGVEHLQITVGVDGTLY
KLHPHFSRVLQETVKELAPQCDVTFMLSEDGSGKGAALITAVAKRLHNVGQK
>anc75_altAll|avgPP:0.997|lnPP:-5.786|num_ambig:4|num_ambig_gaps:0
LYHLRLSDDVLLDVMARFQAEMVKGLGRDTNPTATVKMLPSFVRSLPDGSEKGDFLAVDLGGSQFRAHQV
KVFDDGKQSSQLESKFYPTPKEVTQGNGAELFDYIADCLLDFMETKNLKHKKLPLGFTFSFPCKQTKLEE
GVLLAWTKHFKVRGVQDTDVVSSLRKALQKHKDIDVDVLALVNDTVGTMMTCGYDDQRCEVGLIIGTGTN
ACYMEEMRHIDLVEGDEGRMCINTEWGAFGDDGALDDLRTEFDRELDLGSLNPGKQLFEKMISSLYLGEL
VRLILLKMTKEGLLFNGKVSTALLTKGKIEMKHVSAMEKYKEGLSNTKEILTELNLFPSEEDCIAVQHVC
TIVSFRSANLCAAALAAILTRLRENKKLLRMRTTVGIDGGLYKTHPQYAKRLHKVVRRLVPNCDVRFLLS
```

VSGSGKGAAMVTAVAHRLAAQRKQIDAALAPFLLSLETLREVKDKMRTELEYGLKRETQANATVKMLPTY
VCGTPDGTEKGKFLALDLGGTNFRVLLVKIRSGRRRSVQMYNKIFAIPLEIMQGTGEELFDHIVQCIADF
LEYMGIKGARLPLGFTFSFPCRQASIDKGTLVGWTKGFKATDCEGEDVVDMLREAIRRRNEFDLDIVAVV
NDTVGTMMTCGYEDPNCEIGLIAGTGSNVCYMEDMKNIEIVEGNEGKMCINTEWGAFGDNGCIDNIRTKY
DKEVDEGSLNPGKQRYEKMTSGMYLGEIVRQILIDLTKQGLLFRGHISESLRKRGIFETKFLSQIESDRL
ALLQVRRILQQLGLDSTCEDSIIVKEVCGAVSRRAAQLCGAGLAAIVEKKRENRGVERLQITVGVDGTLY
KLHPHFSRVLQETVKELAPQCDVTFMLSEDGSGKGAALITAVAKRLHNVGQK
>anc76|avgPP:1.000|lnPP:-0.647|num_ambig:1|num_ambig_gaps:0
LYHMRLSDDVLLDVMARFQAEMVKGLGRDTNPTATVKMLPSFVRSLPDGSEKGDFLAVDLGGSQFRAHQV
KVFDDGKQSSQLESKFYPTPKEVTQGNGAELFDYIADCLLDFMETKNLKHKKLPLGFTFSFPCKQTKLEE
GVLLAWTKHFKVRGVQDTDVVSSLRKALQKHKDIDVDVLALVNDTVGTMMTCGYDDQRCEVGLIIGTGTN
ACYMEEMRHIDLVEGDEGRMCINTEWGAFGDDGALDDLRTEFDRELDLGSLNPGKQLFEKMISSLYLGEL
VRLILLKMTKEGLLFNGKVSTALLTKGKIEMKHVSAMEKYKEGLSNTKEILTELNLFPSEEDCIAVQHVC
TIVSFRSANLCAAALAAILTRLRENKKLLRMRTTVGIDGGLYKTHPQYAKRLHKVVRRLVPNCDVRFLLS
VSGSGKGAAMVTAVAYRLAAQRKQIDAALAPFLLSLETLREVKNKMRTELEYGLKRETQASATVKMLPTY
VCGTPDGTEKGKFLALDLGGTNFRVLLVKIRSGRRRSVQMYNKIFAIPLEIMQGTGEELFDHIVQCIADF
LEYMGIKGARLPLGFTFSFPCRQASIDKGTLVGWTKGFKATDCEGEDVVDMLREAIRRRNEFDLDIVAVV
NDTVGTMMTCGYEDPNCEIGLIAGTGSNVCYMEDMKNIEIVEGNEGKMCINTEWGAFGDNGCIDNIRTKY
DKEVDEGSLNPGKQRYEKMTSGMYLGEIVRQILIDLTKQGLLFRGQISESLRKRGIFETKFLSQIESDRL
ALLQVRRILQQLGLDSTCEDSIIVKEVCGAVSKRAAQLCGAGLAAIVEKKRENRGVEHLQITVGVDGTLY
KLHPHFSRVLQETVKELAPQCDVTFMLSEDGSGKGAALITAVAKRLHNVGQK
>anc76_altAll|avgPP:0.999|lnPP:-1.280|num_ambig:1|num_ambig_gaps:0
LYHMRLSDDVLLDVMARFQAEMVKGLGRDTNPTATVKMLPSFVRSLPDGSEKGDFLAVDLGGSQFRAHQV
KVFDDGKQSSQLESKFYPTPKEVTQGNGAELFDYIADCLLDFMETKNLKHKKLPLGFTFSFPCKQTKLEE
GVLLAWTKHFKVRGVQDTDVVSSLRKALQKHKDIDVDVLALVNDTVGTMMTCGYDDQRCEVGLIIGTGTN
ACYMEEMRHIDLVEGDEGRMCINTEWGAFGDDGALDDLRTEFDRELDLGSLNPGKQLFEKMISSLYLGEL
VRLILLKMTKEGLLFNGKVSTALLTKGKIEMKHVSAMEKYKEGLSNTKEILTELNLFPSEEDCIAVQHVC
TIVSFRSANLCAAALAAILTRLRENKKLLRMRTTVGIDGGLYKTHPQYAKRLHKVVRRLVPNCDVRFLLS
VSGSGKGAAMVTAVAYRLAAQRKQIDAALAPFLLSLETLREVKNKMRTELEYGLKRETQASATVKMLPTY
VCGTPDGTEKGKFLALDLGGTNFRVLLVKIRSGRRRSVQMYNKIFAIPLEIMQGTGEELFDHIVQCIADF
LEYMGIKGARLPLGFTFSFPCRQASIDKGTLVGWTKGFKATDCEGEDVVDMLREAIRRRNEFDLDIVAVV
NDTVGTMMTCGYEDPNCEIGLIAGTGSNVCYMEDMKNIEIVEGNEGKMCINTEWGAFGDNGCIDNIRTKY
DKEVDEGSLNPGKQRYEKMTSGMYLGEIVRQILIDLTKQGLLFRGQISESLRKRGIFETKFLSQIESDRL
ALLQVRRILQQLGLDSTCEDSIIVKEVCGAVSKRAAQLCGAGLAAIVEKKRENRGVERLQITVGVDGTLY
KLHPHFSRVLQETVKELAPQCDVTFMLSEDGSGKGAALITAVAKRLHNVGQK
>anc77|avgPP:1.000|lnPP:-0.084|num_ambig:0|num_ambig_gaps:0
LYHMRLSDDVLLDVMARFQAEMVKGLGRDTNPTATVKMLPSFVRSLPDGSEKGDFLAVDLGGSQFRAHQV
KVFDDGKQSSQLESKFYPTPKEVIQGNGAELFDYIADCLLDFMETKNLKHKKLPLGFTFSFPCKQTKLDE
GVLLAWTKHFKVRGVQDTDVVSSLRKALQKHKDIDVDVLALVNDTVGTMMTCGYDDQRCEVGLIIGTGTN
ACYMEEMRHIDLVEGDEGRMCINTEWGAFGDDGALDDLRTEFDRELDLGSLNPGKQLFEKMISSLYLGEL
VRLILLKMTKEGLLFNGKVSTALLTKGKIEMKHVSAMEKYKEGLSNTKEILTELNLFPSEEDCIAVQHVC
TIVSFRSANLCAAALAAILTRLRENKKLLRMRTTVGIDGGLYKTHPQYAKRLHKVVRRLVPNCDVRFLLS
VSGSGKGAAMVTAVAYRLAAQRKQIDAALAPFLLSLETLREVKNKMRTELEYGLKRETQASATVKMLPTY
VCGTPDGTEKGKFLALDLGGTNFRVLLVKIRSGRRRSVQMYNKIFAIPLEIMQGTGEELFDHIVQCIADF
LEYMGIKGARLPLGFTFSFPCRQASIDKGTLVGWTKGFKATDCEGEDVVDMLREAIRRRNEFDLDIVAVV
NDTVGTMMTCGYEDPNCEIGLIAGTGSNVCYMEDMKNIEIVEGNEGKMCINTEWGGFGDNGCIDNIRTKY
DKEVDEGSLNPGKQRYEKMTSGMYLGEIVRQILIDLTKQGLLFRGQISESLRKRGIFETKFLSQIESDRL
ALLQVRRILQQLGLDSTCEDSIIVKEVCGAVSKRAAQLCGAGLAAIVEKKRENRGVEHLQITVGVDGTLY
KLHPHFSRVLRETVKELAPQCDVTFMLSEDGSGKGAALITAVAKRLHNVGQK
>anc77_altAll|avgPP:1.000|lnPP:-0.084|num_ambig:0|num_ambig_gaps:0
LYHMRLSDDVLLDVMARFQAEMVKGLGRDTNPTATVKMLPSFVRSLPDGSEKGDFLAVDLGGSQFRAHQV
KVFDDGKQSSQLESKFYPTPKEVIQGNGAELFDYIADCLLDFMETKNLKHKKLPLGFTFSFPCKQTKLDE

GVLLAWTKHFKVRGVQDTDVVSSLRKALQKHKDIDVDVLALVNDTVGTMMTCGYDDQRCEVGLIIGTGTN
ACYMEEMRHIDLVEGDEGRMCINTEWGAFGDDGALDDLRTEFDRELDLGSLNPGKQLFEKMISSLYLGEL
VRLILLKMTKEGLLFNGKVSTALLTKGKIEMKHVSAMEKYKEGLSNTKEILTELNLFPSEEDCIAVQHVC
TIVSFRSANLCAAALAAILTRLRENKKLLRMRTTVGIDGGLYKTHPQYAKRLHKVVRRLVPNCDVRFLLS
VSGSGKGAAMVTAVAYRLAAQRKQIDAALAPFLLSLETLREVKNKMRTELEYGLKRETQASATVKMLPTY
VCGTPDGTEKGKFLALDLGGTNFRVLLVKIRSGRRRSVQMYNKIFAIPLEIMQGTGEELFDHIVQCIADF
LEYMGIKGARLPLGFTFSFPCRQASIDKGTLVGWTKGFKATDCEGEDVVDMLREAIRRRNEFDLDIVAVV
NDTVGTMMTCGYEDPNCEIGLIAGTGSNVCYMEDMKNIEIVEGNEGKMCINTEWGGFGDNGCIDNIRTKY
DKEVDEGSLNPGKQRYEKMTSGMYLGEIVRQILIDLTKQGLLFRGQISESLRKRGIFETKFLSQIESDRL
ALLQVRRILQQLGLDSTCEDSIIVKEVCGAVSKRAAQLCGAGLAAIVEKKRENRGVEHLQITVGVDGTLY
KLHPHFSRVLRETVKELAPQCDVTFMLSEDGSGKGAALITAVAKRLHNVGQK
>anc78|avgPP:1.000|lnPP:-0.084|num_ambig:0|num_ambig_gaps:0
LYHMRLSDDVLLDVMARFQAEMVKGLGRDTNPTATVKMLPSFVRSLPDGSEKGDFLAVDLGGSQFRAHQV
KVFDDGKQSSQLESKFYPTPKEVIQGNGAELFDYIADCLLDFMETKNLKHKKLPLGFTFSFPCKQTKLDE
GVLLAWTKHFKVRGVQDTDVVSSLRKALQKHKDIDVDVLALVNDTVGTMMTCGYDDQRCEVGLIIGTGTN
ACYMEEMRHIDLVEGDEGRMCINTEWGAFGDDGALDDLRTEFDRELDLGSLNPGKQLFEKMISSLYLGEL
VRLILLKMTKEGLLFNGKVSTALLTKGKIEMKHVSAMEKYKEGLSNTKEILTELNLFPSEEDCIAVQHVC
TIVSFRSANLCAAALAAILTRLRENKKLLRMRTTVGIDGGLYKTHPQYAKRLHKVVRRLVPNCDVRFLLS
VSGSGKGAAMVTAVAYRLAAQRKQIDAALAPFLLSLETLREVKNKMRTELEYGLKRETQASATVKMLPTY
VCGTPDGTEKGKFLALDLGGTNFRVLLVKIRSGRRRSVQMYNKIFAIPLEIMQGTGEELFDHIVQCIADF
LEYMGIKGARLPLGFTFSFPCRQASIDKGTLVGWTKGFKATDCEGEDVVDMLREAIRRRNEFDLDIVAVV
NDTVGTMMTCGYEDPNCEIGLIAGTGSNVCYMEDMKNIEIVEGNEGKMCINTEWGGFGDNGCIDNIRTKY
DKEVDEGSLNPGKQRYEKMTSGMYLGEIVRQILIDLTKQGLLFRGQISESLRKRGIFETKFLSQIESDRL
ALLQVRRILQQLGLDSTCEDSIIVKEVCGAVSKRAAQLCGAGLAAIVEKKRENRGVEHLQITVGVDGTLY
KLHPHFSRVLRETVKELAPQCDVTFMLSEDGSGKGAALITAVAKRLHNVGQK
>anc78_altAll|avgPP:1.000|lnPP:-0.084|num_ambig:0|num_ambig_gaps:0
LYHMRLSDDVLLDVMARFQAEMVKGLGRDTNPTATVKMLPSFVRSLPDGSEKGDFLAVDLGGSQFRAHQV
KVFDDGKQSSQLESKFYPTPKEVIQGNGAELFDYIADCLLDFMETKNLKHKKLPLGFTFSFPCKQTKLDE
GVLLAWTKHFKVRGVQDTDVVSSLRKALQKHKDIDVDVLALVNDTVGTMMTCGYDDQRCEVGLIIGTGTN
ACYMEEMRHIDLVEGDEGRMCINTEWGAFGDDGALDDLRTEFDRELDLGSLNPGKQLFEKMISSLYLGEL
VRLILLKMTKEGLLFNGKVSTALLTKGKIEMKHVSAMEKYKEGLSNTKEILTELNLFPSEEDCIAVQHVC
TIVSFRSANLCAAALAAILTRLRENKKLLRMRTTVGIDGGLYKTHPQYAKRLHKVVRRLVPNCDVRFLLS
VSGSGKGAAMVTAVAYRLAAQRKQIDAALAPFLLSLETLREVKNKMRTELEYGLKRETQASATVKMLPTY
VCGTPDGTEKGKFLALDLGGTNFRVLLVKIRSGRRRSVQMYNKIFAIPLEIMQGTGEELFDHIVQCIADF
LEYMGIKGARLPLGFTFSFPCRQASIDKGTLVGWTKGFKATDCEGEDVVDMLREAIRRRNEFDLDIVAVV
NDTVGTMMTCGYEDPNCEIGLIAGTGSNVCYMEDMKNIEIVEGNEGKMCINTEWGGFGDNGCIDNIRTKY
DKEVDEGSLNPGKQRYEKMTSGMYLGEIVRQILIDLTKQGLLFRGQISESLRKRGIFETKFLSQIESDRL
ALLQVRRILQQLGLDSTCEDSIIVKEVCGAVSKRAAQLCGAGLAAIVEKKRENRGVEHLQITVGVDGTLY
KLHPHFSRVLRETVKELAPQCDVTFMLSEDGSGKGAALITAVAKRLHNVGQK
>anc79|avgPP:1.000|lnPP:-0.084|num_ambig:0|num_ambig_gaps:0
LYHMRLSDDVLLDVMARFQAEMVKGLGRDTNPTATVKMLPSFVRSLPDGSEKGDFLAVDLGGSQFRAHQV
KVFDDGKQSSQLESKFYPTPKEVIQGNGAELFDYIADCLLDFMETKNLKHKKLPLGFTFSFPCKQTKLDE
GVLLAWTKHFKVRGVQDTDVVSSLRKALQKHKDIDVDVLALVNDTVGTMMTCGYDDQRCEVGLIIGTGTN
ACYMEEMRHIDLVEGDEGRMCINTEWGAFGDDGALDDLRTEFDRELDLGSLNPGKQLFEKMISSLYLGEL
VRLILLKMTKEGLLFNGKVSTALLTKGKIEMKHVSAMEKYKEGLSNTKEILTELNLFPSEEDCIAVQHVC
TIVSFRSANLCAAALAAILTRLRENKKLLRMRTTVGIDGGLYKTHPQYAKRLHKVVRRLVPNCDVRFLLS
VSGSGKGAAMVTAVAYRLAAQRKQIDAALAPFLLSLETLREVKNKMRTELEYGLKRETQASATVKMLPTY
VCGTPDGTEKGKFLALDLGGTNFRVLLVKIRSGRRRSVQMYNKIFAIPLEIMQGTGEELFDHIVQCIADF
LEYMGIKGARLPLGFTFSFPCRQASIDKGTLVGWTKGFKATDCEGEDVVDMLREAIRRRNEFDLDIVAVV
NDTVGTMMTCGYEDPNCEIGLIAGTGSNVCYMEDMKNIEIVEGNEGKMCINTEWGGFGDNGCIDNIRTKY
DKEVDEGSLNPGKQRYEKMTSGMYLGEIVRQILIDLTKQGLLFRGQISESLRKRGIFETKFLSQIESDRL
ALLQVRRILQQLGLDSTCEDSIIVKEVCGAVSKRAAQLCGAGLAAIVEKKRENRGVEHLQITVGVDGTLY

KLHPHFSRVLRETVKELAPQCDVTFMLSEDGSGKGAALITAVAKRLHNVGQK
>anc79_altAll|avgPP:1.000|lnPP:-0.084|num_ambig:0|num_ambig_gaps:0
LYHMRLSDDVLLDVMARFQAEMVKGLGRDTNPTATVKMLPSFVRSLPDGSEKGDFLAVDLGGSQFRAHQV
KVFDDGKQSSQLESKFYPTPKEVIQGNGAELFDYIADCLLDFMETKNLKHKKLPLGFTFSFPCKQTKLDE
GVLLAWTKHFKVRGVQDTDVVSSLRKALQKHKDIDVDVLALVNDTVGTMMTCGYDDQRCEVGLIIGTGTN
ACYMEEMRHIDLVEGDEGRMCINTEWGAFGDDGALDDLRTEFDRELDLGSLNPGKQLFEKMISSLYLGEL
VRLILLKMTKEGLLFNGKVSTALLTKGKIEMKHVSAMEKYKEGLSNTKEILTELNLFPSEEDCIAVQHVC
TIVSFRSANLCAAALAAILTRLRENKKLLRMRTTVGIDGGLYKTHPQYAKRLHKVVRRLVPNCDVRFLLS
VSGSGKGAAMVTAVAYRLAAQRKQIDAALAPFLLSLETLREVKNKMRTELEYGLKRETQASATVKMLPTY
VCGTPDGTEKGKFLALDLGGTNFRVLLVKIRSGRRRSVQMYNKIFAIPLEIMQGTGEELFDHIVQCIADF
LEYMGIKGARLPLGFTFSFPCRQASIDKGTLVGWTKGFKATDCEGEDVVDMLREAIRRRNEFDLDIVAVV
NDTVGTMMTCGYEDPNCEIGLIAGTGSNVCYMEDMKNIEIVEGNEGKMCINTEWGGFGDNGCIDNIRTKY
DKEVDEGSLNPGKQRYEKMTSGMYLGEIVRQILIDLTKQGLLFRGQISESLRKRGIFETKFLSQIESDRL
ALLQVRRILQQLGLDSTCEDSIIVKEVCGAVSKRAAQLCGAGLAAIVEKKRENRGVEHLQITVGVDGTLY
KLHPHFSRVLRETVKELAPQCDVTFMLSEDGSGKGAALITAVAKRLHNVGQK
>anc80|avgPP:0.999|lnPP:-1.154|num_ambig:1|num_ambig_gaps:0
LYHMRLSDDVLLDVMARFQAEMVKGLGRDTNPTATVKMLPSFVRSLPDGSEKGDFLAVDLGGSQFRAHQV
KVFDDGKQSSQLESKFYPTPKEVIQGNGAELFDYIADCLSDFMETKNLKHKKLPLGFTFSFPCKQTKLDE
GVLLSWTKHFKVRGVQDTDVVSSLRKALQKHKDIDVDVLALVNDTVGTMMTCGYDDQRCEVGLIIGTGTN
ACYMEEMRHIDLVEGDEGRMCINTEWGAFGDDGALDDLRTEFDRELDLGSLNPGKQLFEKMISSLYLGEL
VRLILLKMTKEGLLFNGKVSTALLTKGKIEMKHVSAMEKYKEGLSNTKEILTELNLFPSEEDCIAVQHVC
TIVSFRSANLCAAALAAILTRLRENKKLLRMRTTVGIDGGLYKTHPQYAKRLHKVVRRLVPNCDVRFLLS
VSGSGKGAAMVTAVAYRLAAQRKQIDAALAPFLLSLETLREVKNKMRTELEYGLKRETQASATVKMLPTY
VCGTPDGTEKGKFLALDLGGTNFRVLLVKIRSGRRRSVQMYNKIFAIPLEIMQGTGEELFDHIVQCIADF
LEYMGIKGARLPLGFTFSFPCRQASIDKGTLVGWTKGFKATDCEGEDVVDMLREAIRRRNEFDLDIVAVV
NDTVGTMMTCGYEDPNCEIGLIAGTGSNVCYMEDMKNIEIVEGNEGKMCINTEWGGFGDNGCIDNIRTKY
DKEVDEGSLNPGKQRYEKMTSGMYLGEIVRQILIDLTKQGLLFRGQISESLRKRGIFETKFLSQIESDRL
ALLQVRRILQQLGLDSTCEDSIIVKEVCGAVSKRAAQLCGAGLAAIVEKKRENRGVEHLQITVGVDGTLY
KLHPHFSRVLQETVKELAPQCDVTFMLSEDGSGKGAALITAVAKRLHNVGQK
>anc80_altAll|avgPP:0.999|lnPP:-1.366|num_ambig:1|num_ambig_gaps:0
LYHMRLSDDVLLDVMARFQAEMVKGLGRDTNPTATVKMLPSFVRSLPDGSEKGDFLAVDLGGSQFRAHQV
KVFDDGKQSSQLESKFYPTPKEVIQGNGAELFDYIADCLSDFMETKNLKHKKLPLGFTFSFPCKQTKLDE
GVLLSWTKHFKVRGVQDTDVVSSLRKALQKHKDIDVDVLALVNDTVGTMMTCGYDDQRCEVGLIIGTGTN
ACYMEEMRHIDLVEGDEGRMCINTEWGAFGDDGALDDLRTEFDRELDLGSLNPGKQLFEKMISSLYLGEL
VRLILLKMTKEGLLFNGKVSTALLTKGKIEMKHVSAMEKYKEGLSNTKEILTELNLFPSEEDCIAVQHVC
TIVSFRSANLCAAALAAILTRLRENKKLLRMRTTVGIDGGLYKTHPQYAKRLHKVVRRLVPNCDVRFLLS
VSGSGKGAAMVTAVAYRLAAQRKQIDAALAPFLLSLETLREVKNKMRTELEYGLKRETQASATVKMLPTY
VCGTPDGTEKGKFLALDLGGTNFRVLLVKIRSGRRRSVQMYNKIFAIPLEIMQGTGEELFDHIVQCIADF
LEYMGIKGARLPLGFTFSFPCRQASIDKGTLVGWTKGFKATDCEGEDVVDMLREAIRRRNEFDLDIVAVV
NDTVGTMMTCGYEDPNCEIGLIAGTGSNVCYMEDMKNIEIVEGNEGKMCINTEWGGFGDNGCIDNIRTKY
DKEVDEGSLNPGKQRYEKMTSGMYLGEIVRQILIDLTKQGLLFRGQISESLRKRGIFETKFLSQIESDRL
ALLQVRRILQQLGLDSTCEDSIIVKEVCGAVSKRAAQLCGAGLAAIVEKKRENRGVEHLQITVGVDGTLY
KLHPHFSRVLRETVKELAPQCDVTFMLSEDGSGKGAALITAVAKRLHNVGQK
>anc81|avgPP:0.998|lnPP:-3.583|num_ambig:3|num_ambig_gaps:0
LYHMRLSDDVLLDVMARFQAEMVKGLGRDTNPTATVKMLPSFVRSLPDGSEKGDFLAVDLGGSQFRALQV
KVFDDGKQSSQLESKFYPTPKEVIQGNGTELFDYVADCLSDFMETKNLKHKKLPLGFTFSFPCKQTKLDE
GVLLSWTKHFKVRGVQDTDVVSSLRKALQKHKDIDVDVLALVNDTVGTMMTCGYDDQRCEVGLIIGTGTN
ACYMEEMRHIDLVEGDEGRMCINTEWGAFGDDGALDDLRTEFDRELDLGSLNPGKQLFEKMISSLYLGEL
VRLILLKMTKEGLLFNGKVSTALLTKGKIEMKHVSAMEKYKEGLSNTKEILTELNLFPSEEDCIAVQHVC
TIVSFRSANLCAAALAAILTRLRENKKLLRLRTTVGIDGGLYKTHPQYAKRLHKVVRRLVPNCDVRFLLS
QSGSGKGAAMVTAVAYRLAAQRKQIDAVLAPFLLSLETLREVKNKMRTELEYGLKRETQASATVKMLPTY
VCGTPDGTEKGKFLALDLGGTNFRVLLVKIRSGRRRSVRMYNKIFAIPLEIMQGTGEELFDHIVQCIADF

LEYMGIKGARLPLGFTFSFPCRQASIDKGTLVGWTKGFKATDCEGEDVVDMLREAIRRRNEFDLDIVAVV
NDTVGTMMTCGYEDPNCEIGLIAGTGSNVCYMEDMKNIEIVEGNEGKMCINTEWGGFGDNGCIDNIRTKY
DKEVDEGSLNPGKQRYEKMTSGMYLGEIVRQILIDLTKQGLLFRGQISESLRKRGIFETKFLSQIESDRL
ALLQVRRILQQLGLDSTCEDSIIVKEVCGAVSKRAAQLCGAGLAAIVEKKRENRGVEHLQITVGVDGTLY
KLHPHFSRVLQETVKELAPQCDVTFMLSEDGSGKGAALITAVAKRLHNVGQK
>anc81_altAll|avgPP:0.997|lnPP:-5.209|num_ambig:3|num_ambig_gaps:0
LYHMRLSDDVLLDVMARFQAEMVKGLGRDTNPTATVKMLPSFVRSLPDGSEKGDFLAVDLGGSQFRALQV
KVFDDGKQSSQLESKFYPTPKEVIQGNGTELFDYVADCLSDFMETKNLKHKKLPLGFTFSFPCKQTKLDE
GVLLSWTKHFKARGVQDTDVVSSLRKALQKHKDIDVDVLALVNDTVGTMMTCGYDDQRCEVGLIIGTGTN
ACYMEEMRHIDLVEGDEGRMCINTEWGAFGDDGALDDLRTEFDRELDLGSLNPGKQLFEKMISSLYLGEL
VRLILLKMTKEGLLFNGKVSTALLTKGKIEMKHVSAMEKYKEGLSNTKEILTELNLFPSEEDCIAVQHVC
TIVSFRSANLCAAALAAILTRLRENKKLLRLRTTVGIDGGLYKTHPQYAKRLHKVVRRLVPNCDVRFLLS
ESGSGKGAAMVTAVAYRLAAQRKQIDAVLAPFLLSLETLREVKNKMRTELEYGLKRETQASATVKMLPTY
VCGTPDGTEKGKFLALDLGGTNFRVLLVKIRSGRRRSVRMYNKIFAIPLEIMQGTGEELFDHIVQCIADF
LEYMGIKGARLPLGFTFSFPCRQASIDKGTLVGWTKGFKATDCEGEDVVDMLREAIRRRNEFDLDIVAVV
NDTVGTMMTCGYEDPNCEIGLIAGTGSNVCYMEDMKNIEIVEGNEGKMCINTEWGGFGDNGCIDNIRTKY
DKEVDEGSLNPGKQRYEKMTSGMYLGEIVRQILIDLTKQGLLFRGQISESLRKRGIFETKFLSQIESDRL
ALLQVRRILQQLGLDSTCEDSIIVKEVCGAVSKRAAQLCGAGLAAIVEKKRENRGVEHLQITVGVDGTLY
KLHPHFSRVLRETVKELAPQCDVTFMLSEDGSGKGAALITAVAKRLHNVGQK
>anc82|avgPP:0.996|lnPP:-6.637|num_ambig:9|num_ambig_gaps:0
LYHMRLSDDVLLDVMTRFQAEMVKGLGRDTNPTATVKMLPTFVRSIPDGSEKGDFLALDLGGSKFRVLKV
KVSEDGKQNVQMESQFYPTPKEIIHGNGTELFDYVADCLADFMETKNIKHKKLPLGFTFSFPCKQTKLDE
GVLLSWTKHFKVRGVQETDVVSSLRKALRKHKDIDVDVLALVNDTVGTMMTCGYDDQRCEVGVIIGTGTN
ACYMEEMSHIDLVEGDEGRMCINTEWGAFGDDGSLEDIRTEFDREIDLGSINPGKQLFEKMISGLYLGEL
VRLILLKMAKEGLLFNGKISTALRTKGKIETKHVAAMEKYKEGLSNTKEILMELGLFPSEEDCIAVQHVC
TIVSFRSANLCAAALAAILTRLRENKKLIRLRTTVGMDGTLYKTHPQYAKRLHKVVRRLVPNCDVRFLLS
ESGSGKGAAMVTAVAYRLASQRKQIDEILAPFKLSKENLREVKNKMRAELEYGLKKETQPTATVKMLPTY
VCGTPDGTEKGKFLALDLGGTNFRVLLVKIRSGRRKSVRMYNKIFAIPLEIMQGTGEELFDHIVQCIADF
LEYMGIKGARLPLGFTFSFPCRQASIDKGTLVGWTKGFKATDCEGEDVVDMLREAIKRRNEFDLDIVAVV
NDTVGTMMTCGYEDPNCEIGLIAGTGSNVCYMEDMKNIEIVEGNEGKMCINTEWGGFGDNGCIDNIRTKY
DKEVDEGSLNPGKQRYEKMTSGMYLGEIVRQILIDLTKQGLLFRGQISESLRTRGIFETKFLSQIESDRL
ALLQVRRILQQLGLDSTCDDSIIVKEVCGAVSKRAAQLCGAGLAAIVEKKRENRGVEHLKITVGVDGTLY
KLHPHFSRILRETVKELAPQCDVTFMLSEDGSGKGAALITAVAKRLHNVGEN
>anc82_altAll|avgPP:0.994|lnPP:-11.998|num_ambig:9|num_ambig_gaps:0
LYHMRLSDDVLLDVMTRFQAEMVKGLGRDTNPTATVKMLPTFVRSIPDGSEKGDFLALDLGGSKFRVLKV
KVSEDGKQNVQMESQFYPTPKEIIHGNGTELFDYVADCLADFMETKNIKHKKLPLGFTFSFPCKQTKLDE
GVLLSWTKHFKARGVQETDVVSSLRKALRKHKDIDVDVLALVNDTVGTMMTCGYDDQRCEVGVIIGTGTN
ACYMEEMSHIDLVEGDEGRMCINTEWGAFGDDGSLDDIRTEFDREIDLGSINPGKQLFEKMISGLYLGEL
VRLILLKMAKEGLLFNGKISTALRTKGKIETKHVAAMEKYKEGLSNTKEILMELGLFPSEEDCIAVQHVC
TIVSFRSANLCAAALAAILTRLRENKKLMRLRTTVGMDGTLYKTHPQYAKRLHKVVRRLVPNCDVRFLLS
ESGSGKGAAMVTAVAYRLASQRKQIDEVLAPFKLSKENLREVKNKMRTELEYGLKKETQPTATVKMLPTY
VCGTPDGTEKGKFLALDLGGTNFRVLLVKIRSGRRKSVRMYNKIFAIPLEIMQGTGEELFDHIVQCIADF
LEYMGIKGARLPLGFTFSFPCRQASIDKGTLVGWTKGFKATDCEGEDVVDMLREAIKRRNEFDLDIVAVV
NDTVGTMMTCGYEDPNCEIGLIAGTGSNVCYMEDMKNIEIVEGNEGKMCINTEWGGFGDNGCIDNIRTKY
DKEVDEGSLNPGKQRYEKMTSGMYLGEIVRQILIDLTKHGLLFRGQISESLRTRGIFETKFLSQIESDRL
ALLQVRRILQQLGLDSTCDDSIIVKEVCGAVSRRAAQLCGAGLAAVVEKKRENRGVEHLKITVGVDGTLY
KLHPHFSRILQETVKELAPQCDVTFMLSEDGSGKGAALITAVAKRLHNVGEN
>anc83|avgPP:0.995|lnPP:-7.297|num_ambig:9|num_ambig_gaps:0
LYHMRLSDDVLLDVMTRFQAEMVKGLGRDTNPTATVKMLPTFVRSIPDGSEKGDFLALDLGGSKFRVLKV
KVSEDGKQNVQMESQFYPTPKEIIHGNGTELFDYVADCLADFMETKNIKHKKLPLGFTFSFPCKQTKLDE
GVLLSWTKHFKVRGVQETDVVSSLRKALRKHKDIDVDVLALVNDTVGTMMTCGYDDQRCEVGVIIGTGTN
ACYMEEMSNIDLVEGDEGRMCINTEWGAFGDDGSLEDIRTEFDREIDLGSINPGKQLFEKMISGLYLGEL

VRLILLKMAKEGLLFNGKISTALRTKGKIETKHVAAMEKYKEGLSNTKEILMELGLAPSEEDCIAVQHVC
TIVSFRSANLCAAALAAILTRLRENKKLIRLRTTVGMDGTLYKTHPQYAKRLHKVVRRLVPNCDVRFLLS
ESGSGKGAAMVTAVAYRLVSQRKQIDEILAPFKLSKENLIEVKNKMRAELEYGLKKETHPTATVKMLPTY
VCGTPDGTEKGKFLALDLGGTNFRVLLVKIRSGRRKSVRMYNKIFAIPLEIMQGTGEELFDHIVQCIADF
LEYMGIKGARLPLGFTFSFPCRQASIDKGTLVGWTKGFKATDCEGEDVVDMLREAIKRRNEFDLDIVAVV
NDTVGTMMTCGYEDPNCEIGLIAGTGSNVCYMEDMKNIEIVEGNEGKMCINTEWGGFGDNGCIDNIRTKY
DKEVDEGSLNPGKQRYEKMTSGMYLGEIVRQILIDLTKQGLLFRGQISESLRTRGIFETKFLSQIESDRL
ALLQVRRILQQLGLDSTCDDSIIVKEVCGAVSKRAAQLCGAGLAAIVEKKRENRGLEHLKITVGVDGTLY
KLHPHFSRILRETVKELAPQCDVTFMLSEDGSGKGAALITAVAKRLHNVGEN
>anc83_altAll|avgPP:0.993|lnPP:-13.421|num_ambig:9|num_ambig_gaps:0
LYHMRLSDDVLLDVMTRFQAEMVKGLGRDTNPTATVKMLPTFVRSIPDGSEKGDFLALDLGGSKFRVLKV
KVSEDGKQNVQMESQFYPTPKEIIHGNGTELFDYVADCLADFMETKNIKHKKLPLGFTFSFPCKQTKLDE
GVLLSWTKHFKVRGVQETDVVSSLRKALRKHKDIDVDVLALVNDTVGTMMTCGYDDQRCEVGVIIGTGTN
ACYMEEMSNIDLVEGDEGRMCINTEWGAFGDDGSLDDIRTEFDREIDLGSINPGKQLFEKMISGLYLGEL
VRLILLKMAKEGLLFNGKISTALRTKGKIETKHVAAMEKYKEGLSNTKEILMELGLAPSEEDCIAVQHVC
TIVSFRSANLCAAALAAILTRLRENKKLIRLRTTVGMDGTLYKTHPQYAKRLHKVVRRLVPNCDVRFLLS
ESGSGKGAAMVTAVAYRLASQRKQIDEVLAPFKLSKENLIEVKNKMRTELEYGLKKETHPTATVKMLPTY
VCGTPDGTEKGKFLALDLGGTNFRVLLVKIRSGRRKSVRMYNKIFAIPLEIMQGTGEELFDHIVQCIADF
LEYMGIKGARLPLGFTFSFPCRQASIDKGTLVGWTKGFKATDCEGEDVVDMLREAIKRRNEFDLDIVAVV
NDTVGTMMTCGYEDPNCEIGLIAGTGSNVCYMEEMKNIEIVEGDEGKMCINTEWGGFGDNGCIDNIRTKY
DKEVDEGSLNPGKQRYEKMTSGMYLGEIVRQILIDLTKHGLLFRGQISESLRTRGIFETKFLSQIESDRL
ALLQVRRILQQLGLDSTCDDSIIVKEVCGAVSKRAAQLCGAGLAAVVEKKRENRGLEHLKITVGVDGTLY
KLHPHFSRILQETVKELAPQCDVTFMLSEDGSGKGAALITAVAKRLHNVGEN
>anc84|avgPP:0.994|lnPP:-7.973|num_ambig:9|num_ambig_gaps:0
LYHMRLSDDVLLDVMTRFQAEMVKGLGRDTNPTATVKMLPTFVRSIPDGSEKGEFLALDLGGSKFRVLKV
KVSEDGKQNVQMESQFYPTPKEIIHGSGTDLFEYVADCLADFMETKNIKHKKLPLGFTFSFPCRQNKLEE
GVLLSWTKHFKVRGVQQTDVVSSLRKALRKHQDIDVDVLALVNDTVGTMMTCGYDDQRCEVGVIIGTGTN
ACYMEEMSNIDLVEGDEGRMCINTEWGAFGDDASLEDIRTEFDKEIDLGSINPGKQLFEKMISGLYLGEL
VRLILLKMTREGLLFNGKLSTALCTKGKIETKHVAAMEKYKEGLSNTKEILMELGLAPSEEDCIAVQHVC
TIVSFRSANLCAAALAAILTRLRENKKVARLRTTVGMDGTLYKTHPQYAKRLHKVVRRLVPNCDVRFLLS
ESGSGKGAAMVTAVAYRLLSQRKQIDEVLAPFKLTKENLIEVKNKMRAELEYGLKKETHPTATVKMLPTF
VYGTPDGTEKGKFLALDLGGTNFRVLLVKIRSGRSKSVRMYNKIFAIPLEIMQGTGEELFDHIVQCIADF
LEYMGIKGARLPLGFTFSFPCRQASIDKGTLVGWTKGFKATDCEGEDVVDMLREAIKRRNEFDLDIVAVV
NDTVGTMMTCGYEDPNCEIGLIAGTGSNVCYMEDMKNIEIVEGDEGKMCINTEWGGFGDNGCIDNLRTKY
DKEVDEGSLNPGKQRYEKMTSGMYLGEIVRQILIDLTKQGLLFRGQISESLRTRGIFETKFLSQIESDRL
ALLQVRRILQQLGLDSTCDDSIIVKEVCGCVSKRAAQLCGAGLAAIVEKIRDNRKLEHLKTTVGVDGTLY
KLHPHFSRILQETVKELAPQCDVTFMLSEDGSGKGAALITAVAKRMHNVAEN
>anc84_altAll|avgPP:0.992|lnPP:-13.908|num_ambig:9|num_ambig_gaps:0
LYHMRLSDDVLLDVMTRFQAEMVKGLGRDTNPTATLKMLPTFVRSIPDGSEKGEFLALDLGGSRFRVLKV
KVSEDGKQNVQMESQFYPTPKEIIHGNGTDLFEYVADCLADFMETKNIKHKKLPLGFTFSFPCRQNKLEE
GVLLSWTKHFKVRGVQQTDVVSSLRKALRKHQDIDVDVLALVNDTVGTMMTCGYDDQRCEVGVIIGTGTN
ACYMEEMSNIDLVEGDEGRMCINTEWGAFGDDASLEDIRTEFDREIDLGSINPGKQLFEKMISGLYLGEL
VRLILLKMTREGLLFNGKLSTALCTKGKIETKHVAAMEKYKEGLSNTKEILMELGLAPSEEDCIAVQHVC
TIVSFRSANLCAAALAAILTRLRENKKVARLRTTVGIDGTLYKTHPQYAKRLHKVVRRLVPNCDVRFLLS
ESGSGKGAAMVTAVAYRLLSQRKQIDEVLAPFKLTKENLIEVKNKMRTELEYGLKKETHPTATVKMLPTF
VCGTPDGTEKGKFLALDLGGTNFRVLLVKIRSGRSKSVRMYNKIFAIPLEIMQGTGEELFDHIVQCIADF
LEYMGIKGARLPLGFTFSFPCRQASIDKGALVGWTKGFKATDCEGEDVVDMLREAIKRRNEFDLDIVAVV
NDTVGTMMTCGYEDPNCEIGLIAGTGSNVCYMEDMKNIEIVEGDEGKMCINTEWGGFGDNGCIDNLRTKY
DKEVDEGSLNPGKQRYEKMTSGMYLGEIVRQILIDLTKQGLLFRGQISESLRTRGIFETKFLSQIESDRL
ALLQVRRILQQLGLDSTCDDSIIVKEVCGCVSKRAAQLCGAGLAAVVEKIRDNRKLEHLKTTVGVDGTLY
KLHPHFSRILQETVKELAPQCDVTFMLSEDGSGKGAALITAVAKRMHNVAEN
>anc85|avgPP:0.995|lnPP:-6.959|num_ambig:6|num_ambig_gaps:0

LYHMRLSDDVLLDVMTRFQAEMVKGLGRDTNPTATVKMLPTFVRSIPDGSEKGDFLALDLGGSKFRVLKV
KVSEDGKQNVQMESQFYPTPKEIIHGNGTDLFEYVADCLADFMETKNIKHKKLPLGFTFSFPCRQNKLEE
GVLLSWTKHFKVRGVQQTDVVSSLRKALRKHQDIDVDVLALVNDTVGTMMTCGYDDQRCEVGVIIGTGTN
ACYMEEMSNIDLVEGDEGRMCINTEWGAFGDDGSLEDIRTEFDREIDLGSINPGKQLFEKMISGLYLGEL
VRLILLKMTREGLLFNGKLSTALCTKGKIETKHVAAMEKYKEGLSNTKEILMELGLAPSEEDCIAVQHVC
TIVSFRSANLCAAALAAILTRLRENKKLIRLRTTVGMDGTLYKTHPQYAKRLHKVVRRLVPNCDVRFLLS
ESGSGKGAAMVTAVAYRLLSQRKQIDEVLAPFKLTKENLIEVKNKMRAELEYGLKKETHPTATVKMLPTF
VCGTPDGTEKGKFLALDLGGTNFRVLLVKIRSGRSKSVRMYNKIFAIPLEIMQGTGEELFDHIVQCIADF
LEYMGIKGARLPLGFTFSFPCRQASIDKGTLVGWTKGFKATDCEGEDVVDMLREAIKRRNEFDLDIVAVV
NDTVGTMMTCGYEDPNCEIGLIAGTGSNVCYMEDMKNIEIVEGDEGKMCINTEWGGFGDNGCIDNLRTKY
DKEVDEGSLNPGKQRYEKMTSGMYLGEIVRQILIDLTKQGLLFRGQISESLRTRGIFETKFLSQIESDRL
ALLQVRRILQQLGLDSTCDDSIIVKEVCGCVSKRAAQLCGAGLAAIVEKIRENRKLEHLKTTVGVDGTLY
KLHPHFSRILQETVKELAPQCDVTFMLSEDGSGKGAALITAVAKRMHNVAEN
>anc85_altAll|avgPP:0.994|lnPP:-10.185|num_ambig:6|num_ambig_gaps:0
LYHMRLSDDVLLDVMTRFQAEMVKGLGRDTNPTATLKMLPTFVRSIPDGSEKGDFLALDLGGSRFRVLKV
KVSEDGKQNVQMESQFYPTPKEIIHGNGTDLFEYVADCLADFMETKNIKHKKLPLGFTFSFPCRQNKLEE
GVLLSWTKHFKVRGVQQTDVVSSLRKALRKHQDIDVDVLALVNDTVGTMMTCGYDDQRCEVGVIIGTGTN
ACYMEEMSNIDLVEGDEGRMCINTEWGAFGDDGSLEDIRTEFDREIDLGSINPGKQLFEKMISGLYLGEL
VRLILLKMTREGLLFNGKLSTALCTKGKIETKHVAAMEKYKEGLSNTKEILMELGLAPSEEDCIAVQHVC
TIVSFRSANLCAAALAAILTRLRENKKLIRLRTTVGMDGTLYKTHPQYAKRLHKVVRRLVPNCDVRFLLS
ESGSGKGAAMVTAVAYRLLSQRKQIDEVLAPFKLTKENLIEVKNKMRTELEYGLKKETHPTATVKMLPTF
VCGTPDGTEKGKFLALDLGGTNFRVLLVKIRSGRSKSVRMYNKIFAIPLEIMQGTGEELFDHIVQCIADF
LEYMGIKGARLPLGFTFSFPCRQASIDKGALVGWTKGFKATDCEGEDVVDMLREAIKRRNEFDLDIVAVV
NDTVGTMMTCGYEDPNCEIGLIAGTGSNVCYMEDMKNIEIVEGDEGKMCINTEWGGFGDNGCIDNLRTKY
DKEVDEGSLNPGKQRYEKMTSGMYLGEIVRQILIDLTKQGLLFRGQISESLRTRGIFETKFLSQIESDRL
ALLQVRRILQQLGLDSTCDDSIIVKEVCGCVSKRAAQLCGAGLAAVVEKIRENRNLEHLKTTVGVDGTLY
KLHPHFSRILQETVKELAPQCDVTFMLSEDGSGKGAALITAVAKRMHNVAEN
>anc86|avgPP:0.996|lnPP:-5.743|num_ambig:5|num_ambig_gaps:0
LYHMRLSDDVLLDVMTRFQAEMVKGLGRDTNPTATVKMLPTFVRSIPDGSEKGDFLALDLGGSKFRVLKV
KVSEDGKQNVQMESQFYPTPKEIIHGNGTELFDYVADCLADFMETKNIKHKKLPLGFTFSFPCKQTKLDE
GVLLSWTKHFKVRGVQETDVVSSLRKALRKHKDIDVDVLALVNDTVGTMMTCGYDDQRCEVGVIIGTGTN
ACYMEEMSNIDLVEGDEGRMCINTEWGAFGDDGSLEDIRTEFDREIDLGSINPGKQLFEKMISGLYLGEL
VRLILLKMAKEGLLFNGKISTALRTKGKIETKHVAAMEKYKEGLSNTKEILMELGLAPSEEDCIAVQHVC
TIVSFRSANLCAAALAAILTRLRENKKLIRLRTTVGMDGTLYKTHPQYAKRLHKVVRRLVPNCDVRFLLS
ESGSGKGAAMVTAVAYRLLSQRKQIDEVLAPFKLSKENLIEVKNKMRAELEYGLKKETHPTATVKMLPTY
VCGTPDGTEKGKFLALDLGGTNFRVLLVKIRSGRRKSVRMYNKIFAIPLEIMQGTGEELFDHIVQCIADF
LEYMGIKGARLPLGFTFSFPCRQASIDKGTLVGWTKGFKATDCEGEDVVDMLREAIKRRNEFDLDIVAVV
NDTVGTMMTCGYEDPNCEIGLIAGTGSNVCYMEDMKNIEIVEGDEGKMCINTEWGGFGDNGCIDNIRTKY
DKEVDEGSLNPGKQRYEKMTSGMYLGEIVRQILIDLTKQGLLFRGQISESLRTRGIFETKFLSQIESDRL
ALLQVRRILQQLGLDSTCDDSIIVKEVCGAVSKRAAQLCGAGLAAIVEKIRENRGLEHLKITVGVDGTLY
KLHPHFSRILQETVKELAPQCDVTFMLSEDGSGKGAALITAVAKRLHNVGEN
>anc86_altAll|avgPP:0.994|lnPP:-9.751|num_ambig:5|num_ambig_gaps:0
LYHMRLSDDVLLDVMTRFQAEMVKGLGRDTNPTATVKMLPTFVRSIPDGSEKGDFLALDLGGSKFRVLKV
KVSEDGKQNVQMESQFYPTPKEIIHGNGTELFDYVADCLADFMETKNIKHKKLPLGFTFSFPCKQTKLDE
GVLLSWTKHFKVRGVQETDVVSSLRKALRKHKDIDVDVLALVNDTVGTMMTCGYDDQRCEVGVIIGTGTN
ACYMEEMSNIDLVEGDEGRMCINTEWGAFGDDGSLDDIRTEFDREIDLGSINPGKQLFEKMISGLYLGEL
VRLILLKMAKEGLLFNGKISTALRTKGKIETKHVAAMEKYKEGLSNTKEILMELGLAPSEEDCIAVQHVC
TIVSFRSANLCAAALAAILTRLRENKKLIRLRTTVGMDGTLYKTHPQYAKRLHKVVRRLVPNCDVRFLLS
ESGSGKGAAMVTAVAYRLLSQRKQIDEILAPFKLSKENLIEVKNKMRTELEYGLKKETHPTATVKMLPTY
VCGTPDGTEKGKFLALDLGGTNFRVLLVKIRSGRRKSVRMYNKIFAIPLEIMQGTGEELFDHIVQCIADF
LEYMGIKGARLPLGFTFSFPCRQASIDKGTLVGWTKGFKATDCEGEDVVDMLREAIKRRNEFDLDIVAVV
NDTVGTMMTCGYEDPNCEIGLIAGTGSNVCYMEEMKNIEIVEGDEGKMCINTEWGGFGDNGCIDNIRTKY

DKEVDEGSLNPGKQRYEKMTSGMYLGEIVRQILIDLTKQGLLFRGQISESLRTRGIFETKFLSQIESDRL
ALLQVRRILQQLGLDSTCDDSIIVKEVCGAVSKRAAQLCGAGLAAVVEKIRENRGLEHLKITVGVDGTLY
KLHPHFSRILQETVKELAPQCDVTFMLSEDGSGKGAALITAVAKRLHNVGEN
>anc87|avgPP:0.992|lnPP:-12.119|num_ambig:11|num_ambig_gaps:0
LYHMRLSDDTLLDIMTRFQAEMEKGLGKDTNPTATVKMLPTFVRSIPDGSEKGDFLALDLGGSKFRVLKV
KVSEDGKQNVQMESQFYPTPKEIIHGNGTELFDYVADCLADFMETKNIKHKKLPLGFTFSFPCKQTKLDE
GVLLSWTKNFKVRGVQGTDVVSSLRKALKKHKDIDVDVLALVNDTVGTMMTCGYDDQHCEVGVIIGTGTN
ACYMEEMSNIDLVEGDEGRMCINTEWGAFGDDGSLEDIRTEFDREIDLGSINPGKQLFEKMISGLYLGEL
VRLILLKMAKEGLLFNGKISTALRTKGKIETKHVAAMEKYKEGLSNTKEILMELGLEPSEEDCIAVQHVC
TIVSFRSANLCAAALAAILTRLRENKKLIRLRTTVGMDGTLYKTHPQYAKRLHKVVRRLVPNCDVRFLLS
ESGSGKGAAMVTAVAYRLLSQRKQIDEVLAPFKLSKEHLIEVKNKMRAELEYGLKKETHPTATVKMLPTY
VCGTPDGTEKGKFLALDLGGTNFRVLLVKIRSGRRKSVRMYNKIFAIPLEIMQGTGEELFDHIVQCIADF
LDYMGIKGARLPLGFTFSFPCRQTSIDKGTLVGWTKGFKATDCEGEDVVDMLREAIKRRNEFDLDIVAVV
NDTVGTMMTCGYEDPNCEIGLIAGTGSNVCYMEEMKNIEIVEGDEGKMCINTEWGGFGDNGCIDDIRTKY
DKEVDEGSLNPGKQRYEKMTSGMYLGEIVRQILIDLTKQGLLFRGQISERLRTRGIFETKFLSQIESDRL
ALLQVRRILQQLGLDSTCDDSIIVKEVCGAVSRRAAQLCGAGLAAIVEKIRENRGLEHLKITVGVDGTLY
KLHPHFSRILQETVKELAPQCDVTFMLSEDGSGKGAALITAVAKRLHNAGEN
>anc87_altAll|avgPP:0.989|lnPP:-19.528|num_ambig:11|num_ambig_gaps:0
LYHMRLSDDTLLDIMTRFQAEMEKGLGRDTNPTATVKMLPTFVRSIPDGSEKGDFLALDLGGSKFRVLKV
KVSEDGKQNVQMESQFYPTPKEIIHGNGTELFDYVADCLADFMETKNIKHKKLPLGFTFSFPCKQTKLDE
GVLLSWTKNFKARGVQGTDVVSSLRKALKKHKDIDVDVLALVNDTVGTMMTCGYDDQHCEVGVIIGTGTN
ACYMEEMSNIDLVEGDEGRMCINTEWGAFGDDGSLEDIRTEFDREIDLGSINPGKQLFEKMISGLYLGEL
VRLILLKMAKEGLLFNGKISTALRTKGKIETKHVAAMEKYKEGLSNTKEILTDLGLEPSEEDCIAVQHVC
TIVSFRSANLCAAALAAILTRLRENKKLVRLRTTVGMDGTLYKTHPQYAKRLHKVVRRLVPNCDVRFLLS
ESGSGKGAAMVTAVAYRLLSQRKQIDEILAPFKLSKEHLIDVKNKMRTELEYGLKKETHPTATVKMLPTY
VCGTPDGTEKGKFLALDLGGTNFRVLLVKIRSGRRKSVRMYNKIFAIPLEIMQGTGEELFDHIVQCIADF
LDYMGIKGARLPLGFTFSFPCRQASIDKGTLVGWTKGFKATDCEGEDVVDMLREAIKRRNEFDLDIVAVV
NDTVGTMMTCGYEDPNCEIGLIAGTGSNVCYMEEMKNIEIVEGDEGKMCINTEWGGFGDNGCIDDIRTQY
DKEVDEGSLNPGKQRYEKMTSGMYLGEIVRQILIDLTKQGLLFRGQISERLRTRGIFETKFLSQIESDRL
ALLQVRRILQQLGLDSTCDDSIIVKEVCGAVSRRAAQLCGAGLAAVVEKIRENRGLEHLKITVGVDGTLY
KLHPHFSRILQETVKELAPQCDVTFMLSEDGSGKGAALITAVAKRLHNAGEN
>anc88|avgPP:0.991|lnPP:-14.075|num_ambig:10|num_ambig_gaps:0
LYHMRLSDETLLDIMARFQAEMEKGLKKDTNSTASVKMLPTYVRSTPDGSEKGDFLALDLGGSKFRVLRV
KVSEDGKQKVQMDSQFYPTPSEIITGSKAELFDYVADCLGDFMKKRQITEKLPLGFTFSFPCKQTKLDEG
VLISWTKEFKVRGVQGTDVVNTLRKTLKKKKYTNVDVLALVNDTVGTMMTCGYDDQLCEVGVIVGTGTNA
CYMENLSNIDLVEGDEGRMCINTEWGAFGDDGSLDDIRTEFDKEIDLGSVNPGKQLFEKMISGLYMGELV
RLILLKMAKKGLLFGGKISNALRTKGKIETSHVASMEKYKEGLYNTKQILLDLGLEPSEDDCIAVQHVAT
IVSFRSANLCAAALAAILTRLRENKKLDRMRTTVGMDGTVYKTHPQYAKRLHKVVRRLVPNCDVRFLLSE
SGSGKGAAMVTAVACRLLSQHKQIDEVLDLFRLSDEKLADIKQRMMTELKYGLKKETHDSATVKMLPTYV
YGTPDGTEHGKFLALDLGGTNFRVLLVKIRSGRRRYVRMYNKIFAIPLEIMQGTGEELFDHIVECIAEFL
DYMGIKGAPLPLGFTFSFPCKQDRIDKGTLVGWTKGFKATDCEGQDVVDMLREAIKRRNEFDLDIVAIVN
DTVGTMMTCGYEDPNCEIGLIAGTGSNVCYMEEMKNIEFVDGDEGKMCINTEWGGFGDNDCLADIQTQYD
IHVDEKSLNPGKQRYEKMTSGMYLGEIVRQILIDLTKRGLLFRGQISERLRTKGIFETKFLSQIESDRLA
LLQVRKILQQLGLDSTCDDSIIVKEVCGAVSTRAAKLCGAGLAAVVDKIRENRGLDHLKITVGVDGTLYK
LHPHFSRILQETVKKLSPQCDVTFMLSEDGSGKGAALITAVAKRFHSQGEEGKN
>anc88_altAll|avgPP:0.989|lnPP:-17.722|num_ambig:10|num_ambig_gaps:0
LYHMRLSDETLLDIMARFQAEMEKGLKKDTNSTASLKMLPTYVRSTPDGSEKGDFLALDLGGSKFRVLRV
KVSEDGKQNVQMDSQFYPTPSEIITGNKAELFDYVADCLGDFIKKRQITEKLPLGFTFSFPCKQTKLDEG
VLISWTKDFKVRGVQGTDVVNTLRKTLKKKKYTNVDVLALVNDTVGTMMTCGYDDQLCEVGVIVGTGTNA
CYMENLSNIDLVEGDEGRMCINTEWGAFGDDGSLDDIRTEFDKEIDLGSVNPGKQLFEKMISGLYMGELV
RLILLKMAKKGLLFGGKISNALRTKGKIETSHVASMEKYKEGLYNTKQILLDLGLEPSEDDCIAVQHVAT
IVSFRSANLCAAALAAILTRLRENKKLDRMRTTVGMDGTVYKTHPQYAKRLHKVVRRLVPNCDVRFLLSE

SGSGKGAAMVTAVACRLLSQHKQIDEVLDLLQISDEKLADIKQRMMTELKYGLKKETHDSATVKMLPTYV
YGTPDGTEHGKFLALDLGGTNFRVLLVKIRSGRRRYVRMYNKIFAIPLEIMQGTGEELFDHIVECIAEFL
DYMGIKGAPLPLGFTFSFPCKQDRIDKGTLVGWTKGFKATDCEGQDVVDMLREAIKRRNEFDLDIVAIVN
DTVGTMMTCGYEDPNCEIGLIAGTGSNVCYMEEMKNIEFIDGDEGKMCINTEWGGFGDNDCLADIQTQYD
IRVDEKSLNPGKQRYEKMTSGMYLGEIVRQILIDLTKRGLLFRGQISERLRTKGIFETKFLSQIESDRLA
LLQVRKILQQLGLDSTCDDSIIVKEVCGAVSTRAAKLCGAGLAAVVDKIRENRGLDHLKITVGVDGTLYK
LHPHFSRILQETVKKLSPQCDVTFMLSEDGSGKGAALITAVAKRFHSQGEEGKN
>anc89|avgPP:0.986|lnPP:-21.465|num_ambig:22|num_ambig_gaps:0
LYHMRLSDETLLDIMARFQAEMEKGLKKDTNPTASVKMLPTHVRSTPDGSEKGDFLALDLGGSKFRVLRV
KVSEDGKQKVQMDSQFYPTPNEIITGSGAELFDYVADCLGDFMKKRQIKEKLPLGFTFSFPCKQTKLDEG
VLISWTKEFKARGVQGTDVVNTLRKTLKKKKYTNVDVLALVNDTVGTMMTCGYDDQLCEVGVIVGTGTNA
CYMENLSNIDLVEGDEGRMCINTEWGAFGDDGSLDDIRTEFDKEIDLGSVNPGKQLFEKMISGLYMGELV
RLILLKMAKKGLLFGGKISNALRTKGKIETSHVAAMEKYKEGLYNTKQILTDLGLEPSEEDCIAVQHVCT
IVSFRSANLCAAALAAILTRLRENKKLERMRTTVGMDGTVYKTHPQYAKRLHKVVRRLVPNCDVRFLLSE
SGSGKGAAMVTAVACRLLSQHKQIDEVLALFQLSNENLADIKQRMMTELKYGLKKETHDSATVKMLPTYV
YGTPDGTEHGKFLALDLGGTNFRVLLVKIRSGRRKYVRMYNKIFAIPLEIMQGTGEELFDHIVECIAEFL
DYMGIKGAPLPLGFTFSFPCKQDRIDKGTLVGWTKGFKATDCEGQDVVDMLREAIKRRNEFDLDIVAIVN
DTVGTMMTCGYEDPNCEIGLIAGTGSNVCYMEEMKNIEIVDGDEGKMCINTEWGGFGDNGCLADIQTKYD
IDVDEKSLNPGKQRYEKMTSGMYLGEIVRQILIDLTKRGLLFRGQISERLRTKGIFETKFLSQIESDRLA
LLQVRKILQQLGLDSTCDDSIIVKEVCGAVSTRAAKLCGAGLAAVVDKIRENRGLDHLKITVGVDGTLYK
LHPHFSRILQETVKKLSPQCDVTFMLSEDGSGKGAALITAVAKRFHSQGEEGTN
>anc89_altAll|avgPP:0.983|lnPP:-31.413|num_ambig:22|num_ambig_gaps:0
LYHMRLSDETLLDIMARFQAEMEKGLKKDTNPTASLKMLPTYVRSTPDGSEKGDFLALDLGGSKFRVLRV
KVSEDGKQNVQMDSQFYPTPSEIITGNGAELFDYVADCLGDFIKKRQIKEKLPLGFTFSFPCKQTKLDEG
VLISWTKDFKARGVQGTDVVNTLRKTLKKKKYANVDVLALVNDTVGTMMTCGYDDQLCEVGVIVGTGTNA
CYMENLSNIDLVEGDEGRMCINTEWGAFGDDGSLDDIRTEFDKEIDLGSINPGKQLFEKMISGLYMGELV
RLILLKMAKKGLLFGGKISNALRTKGKIETSHVAAMEKYKEGLYNTKQILMDLGLEPSEEDCIAVQHVCT
IVSFRSANLCAAALAAILTRLRENKKLDRMRTTVGMDGTVYKTHPQYAKRLHKVVRRLVPNCDVRFLLSE
SGSGKGAAMVTAVACRLLSQHKQIDEILALLRLSDENLVDIKERMMTELKYGLKKETHDSATVKMLPTYV
YGTPDGTERGKFLALDLGGTNFRVLLVKIRSGRRKYVRMYNKIFAIPLEIMQGTGEELFDHIVECIAEFL
DYMGIKGAPLPLGFTFSFPCKQERIDKGTLVGWTKGFKATDCEGQDVVDMLREAIKRRNEFDLDIVAIVN
DTVGTMMTCGYEDPNCEIGLIAGTGSNVCYMEEMKNIEIVDGDEGKMCINTEWGGFGDNGCLADIQTQYD
IEVDEKSLNPGKQRYEKMTSGMYLGEIVRQILIDLTKRGLLFRGQISERLRTKGIFETKFLSQIESDRLA
LLQVRKILQQLGLDSTCDDSIIVKEVCGAVSTRAAKLCGAGLAAVVDKIRENRGLDHLKITVGVDGTLYK
LHPHFSRILQETVKKLSPQCDVTFMLSEDGSGKGAALITAVAKRFHSQGEEGKN
>anc90|avgPP:0.984|lnPP:-24.812|num_ambig:25|num_ambig_gaps:0
LYHMRLSDDTLLDIMTRFQAEMEKGLKKDTNPTATVKMLPTHVRSTPDGSEKGDFLALDLGGSKFRVLKV
KVSEDGKQNVQMESQFYPTPKEIIHGNGTELFDYVADCLADFMETKNIKHKKLPLGFTFSFPCKQTKLDE
GVLLSWTKNFKARGVQGTDVVSSLRKALKKHKDIDVDVLALVNDTVGTMMTCGYDDQHCEVGVIIGTGTN
ACYMEEMSNIDLVEGDEGRMCINTEWGAFGDDGSLDDIRTEFDKEIDLGSINPGKQLFEKMISGLYMGEL
VRLILLKMAKKGLLFKGKISTALRTKGKIETKHVAAMEKYKEGLSNTKQILMDLGLEPSEEDCIAVQHVC
TIVSFRSANLCAAALAAILTRLRENKKLRRLRTTVGMDGTVYKTHPQYAKRLHKVVRRLVPNCDVRFLLS
ESGSGKGAAMVTAVAYRLLSQRKQIDEILAPFKLSKEHLIDVKNKMMTELKYGLKKETHDSATVKMLPTY
VYGTPDGTEHGKFLALDLGGTNFRVLLVKIRSGRRKSVRMYNKIFAIPLEIMQGTGEELFDHIVECIAEF
LDYMGIKGAPLPLGFTFSFPCRQTSIDKGTLVGWTKGFKATDCEGEDVVDMLREAIKRRNEFDLDIVAVV
NDTVGTMMTCGYEDPNCEIGLIAGTGSNVCYMEEMKNIEIVEGDEGKMCINTEWGGFGDNGCLDDIRTKY
DKEVDEGSLNPGKQRYEKMTSGMYLGEIVRQILIDLTKQGLLFRGQISERLRTKGIFETKFLSQIESDRL
ALLQVRRILQQLGLDSTCDDSIIVKEVCGAVSTRAAQLCGAGLAAIVEKIRENRGLDHLKITVGVDGTLY
KLHPHFSRILQETVKELAPQCDVTFMLSEDGSGKGAALITAVAKRLHNNGENGTN
>anc90_altAll|avgPP:0.979|lnPP:-39.556|num_ambig:25|num_ambig_gaps:0
LYHMRLSDDTLLDIMTRFQAEMEKGLKKDTNPTATVKMLPTYVRSTPDGSEKGDFLALDLGGSKFRVLRV
KVSEDGKQKVQMESQFYPTPKEIIHGNGAELFDYVADCLADFMEKKNIKHKKLPLGFTFSFPCKQTKLDE

GVLISWTKNFKARGVQGTDVVSSLRKALKKHKDIDVDVLALVNDTVGTMMTCGYDDQHCEVGVIIGTGTN
ACYMEEMSNIDLVEGDEGRMCINTEWGAFGDDGSLEDIRTEFDREIDLGSINPGKQLFEKMISGLYLGEL
VRLILLKMAKKGLLFKGKISTALRTKGKIETKHVAAMEKYKEGLSNTKEILTELGLDPSEEDCIAVQHVC
TIVSFRSANLCAAALAAILTRLRENKKLKRLRTTVGMDGTVYKTHPQYAKRLHKVVRRLVPNCDVRFLLS
ESGSGKGAAMVTAVAYRLLSQRKQIDEVLAPFKLSKEHLIDVKNKMMAELKYGLKKETHETATVKMLPTY
VYGTPDGTERGKFLALDLGGTNFRVLLVKIRSGRRKSVRMYNKIFAIPLEIMQGTGEELFDHIVECIAEF
LDYMGIKGAPLPLGFTFSFPCRQASIDKGTLVGWTKGFKATDCEGEDVVDMLREAIKRRNEFDLDIVAVV
NDTVGTMMTCGYEDPNCEIGLIAGTGSNVCYMEEMKNIEIVDGDEGKMCINTEWGGFGDNGCLDDIRTQY
DKEVDEGSLNPGKQRYEKMTSGMYLGEIVRQILIDLTKQGLLFRGQISERLRTKGIFETKFLSQIESDRL
ALLQVRRILQQLGLDSTCDDSIIVKEVCGAVSTRAAQLCGAGLAAVVDKIRENRGLDHLKITVGVDGTLY
KLHPHFSRILQETVKELAPQCDVTFMLSEDGSGKGAALITAVAKRLHSNGENGTN
>anc91|avgPP:0.988|lnPP:-18.214|num_ambig:23|num_ambig_gaps:0
LYHMRLSDDTLLDIMTRFQAEMEKGLGKDTNPTATVKMLPTFVRSTPDGSEKGDFLALDLGGSKFRVLKV
KVSEDGKQNVQMESQFYPTPKEIIHGNGTELFDYVADCLADFMETKNIKHKKLPLGFTFSFPCKQTKLDE
GVLLSWTKNFKARGVQGTDVVSSLRKALKKHKDIDVDVLALVNDTVGTMMTCGYDDQHCEVGVIIGTGTN
ACYMEEMSNIDLVEGDEGRMCINTEWGAFGDDGSLEDIRTEFDREIDLGSINPGKQLFEKMISGLYMGEL
VRLILLKMAKKGLLFKGKISTALRTKGKIETKHVAAMEKYKEGLSNTKEILMDLGLEPSEEDCIAVQHVC
TIVSFRSANLCAAALAAILTRLRENKKLKRLRTTVGMDGTVYKTHPQYAKRLHKVVRRLVPNCDVRFLLS
ESGSGKGAAMVTAVAYRLLSQRKQIDEVLAPFKLSKEHLIEVKNKMRTELEYGLKKETHPTATVKMLPTY
VYGTPDGTERGKFLALDLGGTNFRVLLVKIRSGRRKSVRMYNKIFAIPLEIMQGTGEELFDHIVQCIADF
LDYMGIKGARLPLGFTFSFPCRQTSIDKGTLVGWTKGFKATDCEGEDVVDMLREAIKRRNEFDLDIVAVV
NDTVGTMMTCGYEDPNCEIGLIAGTGSNVCYMEEMKNIEIVEGDEGKMCINTEWGGFGDNGCIDDIRTKY
DKEVDEGSLNPGKQRYEKMTSGMYLGEIVRQILIDLTKQGLLFRGQISERLRTRGIFETKFLSQIESDRL
ALLQVRRILQQLGLDSTCDDSIIVKEVCGAVSRRAAQLCGAGLAAIVEKIRENRGLDHLKITVGVDGTLY
KLHPHFSRILQETVKELAPQCDVTFMLSEDGSGKGAALITAVAKRLHNNGEN
>anc91_altAll|avgPP:0.983|lnPP:-31.016|num_ambig:23|num_ambig_gaps:0
LYHMRLSDDTLLDIMTRFQAEMEKGLGRDTNPTATVKMLPTFVRSTPDGSEKGDFLALDLGGSKFRVLRV
KVSEDGKQKVQMESQFYPTPKEIIHGNGTELFDYVADCLADFMEKKNIKHKKLPLGFTFSFPCKQTKLDE
GVLISWTKNFKARGVQGTDVVSSLRKALKKHKDIDVDVLALVNDTVGTMMTCGYDDQHCEVGVIIGTGTN
ACYMEEMSNIDLVEGDEGRMCINTEWGAFGDDGSLDDIRTEFDREIDLGSINPGKQLFEKMISGLYLGEL
VRLILLKMAKKGLLFKGKISTALRTKGKIETKHVAAMEKYKEGLSNTKEILTELGLDPSEEDCIAVQHVC
TIVSFRSANLCAAALAAILTRLRENKKLRRLRTTVGMDGTVYKTHPQYAKRLHKVVRRLVPNCDVRFLLS
ESGSGKGAAMVTAVAYRLLSQRKQIDEILAPFKLSKEHLIDVKKKMRAELEYGLKKETHHTATVKMLPTY
VYGTPDGTEKGKFLALDLGGTNFRVLLVKIRSGRRKSVRMYNKIFAIPLEIMQGTGEELFDHIVQCIADF
LDYMGIKGARLPLGFTFSFPCRQASIDKGTLVGWTKGFKATDCEGEDVVDMLREAIKRRNEFDLDIVAVV
NDTVGTMMTCGYEDPNCEIGLIAGTGSNVCYMEEMKNIEIVEGDEGKMCINTEWGGFGDNGCIDDIRTQY
DKEVDEGSLNPGKQRYEKMTSGMYLGEIVRQILIDLTKQGLLFRGQISERLRTRGIFETKFLSQIESDRL
ALLQVRRILQQLGLDSTCDDSIIVKEVCGAVSRRAAQLCGAGLAAVVDKIRENRGLDHLKITVGVDGTLY
KLHPHFSRILQETVKELAPQCDVTFMLSEDGSGKGAALITAVAKRLHSAGEN
>anc92|avgPP:0.985|lnPP:-22.766|num_ambig:22|num_ambig_gaps:0
LYHMRLSDDTLLDIMTRFRAEMEKGLGKDTNPTATVKMLPTFVRSTPDGSEKGDFLALDLGGSKFRVLRV
KVSEDGKQKVEMESQFYPTPEEIIHGNGTELFDYVAECLADFMEKKNIKDKKLPLGFTFSFPCKQTKLDE
GVLISWTKNFKARGVQGTDVVSSLRKAIKKHKDIDVDVLALVNDTVGTMMTCGYDDQYCEVGVIIGTGTN
ACYMEEMRHIDLVEGDEGRMCINTEWGAFGDDGSLEDIRTEFDREIDLGSINPGKQLFEKMISGLYMGEL
VRLILVKMAKKGLLFKGKISAALRTKGKFETKHVAAIEKYKEGLSNAKEILTDLGLDPSEEDCIAVQHVC
TIVSFRSANLCAAALAAILTRLRENKKLKRLRTTVGMDGTVYKTHPQYAKRLHKVVRRLVPDCDVRFLLS
ESGSGKGAAMVTAVAYRLVSQRKQIDETLAPFKLSKEQLLEVKKKMRTELEHGLKKETHHTATVKMLPTY
VYGTPDGTERGKFLALDLGGTNFRVLLVKIRSGLRRSVRMYNKIYAIPLEIMQGTGEELFDHIVQCISDF
LDYMGMKGARLPLGFTFSFPCRQTGIDKGTLVGWTKGFKATDCEGEDVVDMLREAIKRRNEFDLDIVAVV
NDTVGTMMTCGYEDPNCEIGLIAGTGSNVCYMEEMRNIEIVEGDEGRMCINTEWGGFGDNGCIDDIRTKF
DREVDEGSLNPGKQRYEKMTSGMYLGEIVRQILIDLTKQGLLFRGQISERLKTRGIFETKFLSQIESDRL
ALLQVRSILQQLGLDSTCDDSIIVKEVCGAVSRRAAQLCGAGLAAIVDKIRENRGLDHLEITVGVDGTLY

```
KLHPHFSRILQETVKELAPQCDVTFMLSEDGSGKGAALITAVARRLHSNGQH
>anc92_altAll|avgPP:0.980|lnPP:-35.231|num_ambig:22|num_ambig_gaps:0
LYHMRLSDDTLLDIMTRFRAEMEKGLGRDTNPTATVKMLPTFVRSTPDGSEKGDFLALDLGGSKFRVLRV
KVSEDGKQKVEMESQFYPTPEEIIHGNGTQLFDYVAECLADFMEKKNIKDKKLPLGFTFSFPCKQTKLDE
GILLSWTKNFKARGVQGTDVVSSLRKAIKKHKDIDVDVLALVNDTVGTMMTCGYDDQHCEVGVIIGTGTN
ACYMEEMRHIDLVEGDEGRMCINTEWGAFGDDGSLEDIRTEFDREIDLGSINPGKQLFEKMISGMYMGEL
VRLILVKMAKKGLLFKGKISTALRTKGKFETKHVAAIEKYKEGLSNAKEILTDLGLEPSEEDCIAVQHVC
TIVSFRSANLCAATLAAILTRLRENKKLRRLRTTVGMDGTVYKTHPQYAKRLHKVVRRLVPDCDVRFLLS
ESGSGKGAAMVTAVAYRLVSQRKQIDETLAPFKLSREQLLEVKNKMRTELEHGLKKETHHTASVKMLPTY
VYGTPDGTERGKFLALDLGGTNFRVLLVKIRSGLRKSVRMYNKIYAIPLEIMQGTGEELFDHIVQCISDF
LDYMGMKGARLPLGFTFSFPCRQASIDKGTLVGWTKGFKATDCEGEDVVDMLREAIKRRNEFDLDIVAVV
NDTVGTMMTCGYEDPNCEIGLIAGTGSNVCYMEEMRNIEIVEGDEGRMCINTEWGAFGDNGCIDDIRTQF
DREVDEGSLNPGKQRYEKMTSGMYLGEIVRQILIDLTKQGLLFRGQISERLKTRGIFETKFLSQIESDRL
ALLQVRAILQQLGLDSTCDDSIIVKEVCGAVSRRAAQLCGAGLAAVVDKIRENRGLDHLEITVGVDGTLY
KLHPHFSRILQETVKELAPQCDVTFMLSEDGSGKGAALITAVARRLHNAGQH
>anc93|avgPP:0.984|lnPP:-24.123|num_ambig:27|num_ambig_gaps:0
LYHMRLSDDTLLDIMTRFRAEMEKGLGRDTNPTATVKMLPTFVRSTPDGSEKGDFLALDLGGSKFRVLRV
KVSEDGKQKVEMESQFYPTPEEIIHGNGTELFDYVAECLADFMEKKNIKDKKLPLGFTFSFPCRQTKLDE
GVLISWTKNFKARGVQGTDVVSSLRKAIKKHGDIDVDVLALVNDTVGTMMTCGYDDQYCEVGVIIGTGTN
ACYMEEMRHIDLVEGDEGRMCINTEWGAFGDDGSLEDIRTEFDREIDLGSINPGKQLFEKMISGMYMGEL
VRLILVKMAKKGLLFKGKISAALRTKGKFETKHVAAIEKYKEGLSNAKEILTDLGLDPSEEDCIAVQHVC
TIVSFRSANLCAAALAAILTRLRENKKLKRLRTTVGMDGTVYKTHPQYAKRLHKVVRRLVPDCDVRFLLS
ESGSGKGAAMVTAVAYRLVSQRKQIDETLAPFKLSREQLLEVKKKMRTELERGLKKETHHTATVKMLPTY
VYGTPDGTERGKFLALDLGGTNFRVLLVKIRSGLRRSVRMYNKIYAIPLEIMQGTGEELFDHIVQCISDF
LDYMGMKGARLPLGFTFSFPCRQTGIDKGTLVSWTKGFKATDCEGEDVVDMLREAIKRRNEFDLDIVAVV
NDTVGTMMTCAYEDPNCEIGLIAGTGSNVCYMEEMRNIEIVEGDEGRMCINTEWGGFGDNGCIDDIRTKF
DREVDEGSLNPGKQRYEKMTSGMYLGEIVRQILIDLTKRGLLFRGQISERLKTRGIFETKFLSQIESDRL
ALLQVRSILQQLGLDSTCDDSIIVKEVCGAVSRRAAQLCGAGMAAVVDKIRENRGLDHLEITVGVDGTLY
KLHPHFSRILQETVKELAPQCDVTFMLSEDGSGKGAALITAVARRLHSNGQH
>anc93_altAll|avgPP:0.979|lnPP:-39.557|num_ambig:27|num_ambig_gaps:0
LYHMRLSDDTLLDIMTRFRAEMEKGLGKDTNPTATVKMLPTFVRSTPDGSEKGDFLALDLGGSKFRVLRV
KVSEDGKQKVEMESQIYPTPEEIIHGNGTQLFDYVAECLGDFMEKKHIKDKKLPLGFTFSFPCKQTKLDE
GILLSWTKNFKARGVQGTDVVSSLRKAIKKHGDIDVDVLALVNDTVGTMMTCGYDDQHCEVGVIIGTGTN
ACYMEEMRHIDLVEGDEGRMCINTEWGAFGDDGSLEDIRTEFDREIDLGSINPGKQLFEKMISGMYMGEL
VRLILVKMAKKGLLFKGKISTALRTKGKFETKHVAAIEKYKEGLSNAKEILTDLGLEPSEEDCIAVQHVC
TIVSFRSANLCAATLAAILTRLRENKKLRRLRTTVGMDGTVYKTHPQYAKRLHKVVRRLVPDCDVRFLLS
ESGSGKGAAMVTAVAYRLVSQRKQIDETLAPFKLSKEQLLEVKNKMRTELEHGLKKETHHTASVKMLPTY
VYGTPDGTERGKFLALDLGGTNFRVLLVKIRSGLRKSVRMYNKIYAIPLEIMQGTGEELFDHIVQCISDF
LDYMGMKGARLPLGFTFSFPCRQASIDKGTLVGWTKGFKATDCEGEDVVDMLREAIKRRNEFDLDIVAVV
NDTVGTMMTCAYEDPNCEIGLIAGTGSNVCYMEEMRNIEIVEGDEGRMCINTEWGAFGDNGCIDDIRTQF
DREVDEGSLNPGKQRYEKMTSGMYLGEIVRQILIDLTKQGLLFRGQISERLKTRGIFETKFLSQIESDRL
ALLQVRAILQQLGLDSTCDDSIIVKEVCGAVSRRAAQLCGAGMAAVVDKIRENRGLDHLEITVGVDGTLY
KLHPHFSRILQETVKELAPQCDVTFMLSEDGSGKGAALITAVARRLHNAGQH
>anc94|avgPP:0.982|lnPP:-26.896|num_ambig:22|num_ambig_gaps:0
LYNMQLNSEQLQDIMARFDNEMRKGLSTNETSTVKMLPTFVRSTPDGTEKGDFLALDLGGSRFEVLRVKV
SEDGRQVIHTERQIYPMPDEMKQGNAPEFFNYVAECLGDFMEKQEIKDKKLPLGFTFSFPCRQTKLDEGF
LISWGKGFKIKGTKGANVISLLRKGIEQQGDFDVDVLALVNDTVGTMMSGGFDDPYCEVGVIIGTGTNAC
YMEELSHIESVEGDIGRMCINTEWGAFGDDGSLEDIRTEFDRELDSGSLNPGKQLFEKMISAMYLGELVR
LILVKIAKNGMLFKGNTSAKLLTKGSFGIQHVSAIMNHKSGLENAMQILTDLDLNPSEEDCLAVQHICTI
VLSRSVSLCAATLATILIRLKRNKKVKRLRTTVGIDGTVYRTVPQYSKNLHRMVRRLAPECDVRFLLVET
GTGKGAAMVTAVAYRLVHQRKYQDKTLAPLKLSREQLLKVMKMMRSEMEQGLKKETHSTATVKMLPTFIQ
KFPDGTERGKFLALDLGGTNFRVLLLVLQSRPRRSTQVYQKSYSIPMEIMQGTGEELFDYIVECIVEFLD
```

YMGMKCAQLPLGFTFSFPCVQHRLDQGILVNWTKGFKATDVEGDDVVTLLKEAIKRHGELDLDIVALLND
TVGTMMTCAYENPNCEVGMIAGTGSNACYMEEMKNIEIVEGDEGRMCINTEWGAFGDNGCLDDIRTTFDR
LVDINSLNPGKQKYEKMISGMYLGEIVRNILINFTKHGLLFRGLVSPVLKTSGIFETSYISQIESDRLAL
LQVRTILQNLGLDSTCDDSIIVKEVCGVVSKRAAQLCGAGMAAVVDKIRENRGLDHLEITVGVDGTLYKL
HPHFSTILQETVKDLAPNCDVTFMLSEDGSGKGAALVTAAVKRPD
>anc94_altAll|avgPP:0.978|lnPP:-38.950|num_ambig:22|num_ambig_gaps:0
LYNMHLNSEQLQDIMARFGNEMRKGLSTNETSTVKMLPTFVRSTPDGTEKGDFLALDLGGSRFEVLRVKV
SEDGRQVIHTERQTYPMPDEMKQGNASEFFNYVAECLGDFMEKQEIKGKKLPLGFTFSFPCRQTKLDEGF
LISWGKGFKIKGTKGANVISLLRKGIEQQGDFDVDVLALVNDTVGTMMSGGFDDPYCEVGVIIGTGTNAC
YMEEMSHIESVEGDIGRMCINTEWGAFGDDGSLEDIRTEFDRELDSGSLNPGKQLFEKMISAMYLGELVR
LILVKIAKNGMLFKGNTSAKLLTKGSFGIQHVSAIVNHKSGLENAMQILTDLDLNPSEEDCLAVQHICTI
VLSRSVSLCAATLATILIHLKKNKKIKRLRTTVGIDGTVYRTVPQYSKNLHRMVRRLAPECDVRFILVET
GTGKGAAMVTAVAYRLVHQRKYQDKTLAPLKLTREQLLKVMKMMRSEMEQGLKKETHSTATVKMLPTFIQ
KFPDGTERGKFLALDLGGTNLRVLLLVLRNRPRRSTQMYQKSYRIPMEIMQGTGEELFDYIVECIVEFLD
YMGMKCAQLPLGFTFSFPCVQHQLDQGVLVNWTKGFKATDVEGNDVVTLLKEAIKRHGELDLDIVALLND
TVGTMMTCAYENPNCEVGMIAGTGSNACYMEEMKNIEIVEGDEGRMCINTEWGAFGDNGCLDDIRTTFDR
LVDINSLNPGKQKYEKMISGMYLGEIVRNILINFTKHGLLFRGLVSPVLKTSGIFETSYMSQIESDRLAL
LQVRTILQNLGLDSTCDDSIIVKEVCGVVSKRAAQLCGAGMAAVVDKIRENRGLDHLEITVGVDGTLYKL
HPHFSTILKETVKDLAPNCDVTFMLSEDGSGKGAALVTAAVKRPD
>anc95|avgPP:0.952|lnPP:-78.468|num_ambig:57|num_ambig_gaps:0
LYNMQLNNEQLQDIMARFRAEMRKGLDTNPTSTVKMLPTFVRSTPDGSEKGDFLALDLGGSRFRVLRVKV
SEDGRQMVHMESQIYPMPEEMKHGNATEFFDYVAECLGDFMEKQQIKDKKLPLGFTFSFPCRQTKLDEGI
LISWSKGFKVKGVQGADVVSLLRKAIEKRGDFDVDVLALVNDTVGTMMTCGFDDPYCEVGVIIGTGTNAC
YMEELRHIELVEGDEGRMCINTEWGAFGDDGSLEDIRTEFDREIDSGSLNPGKQLFEKMISAMYMGELVR
LILVKIAKNGMLFKGKTSAKLLTKGSFEIQHVSAIENYKSGLENAMQILTDLDLEPSEEDCIAVQHICTI
VSFRSASLCAATLAAILTRLKKNKKVKRLRTTVGIDGTVYRTHPQYAKNLHKMVRRLVPDCDVRFLLAES
GSGKGAAMVTAVAYRLVHQRKYIDKTLAPLKLSREQLLEVMKKMRSEMEQGLKKETHNTATVKMLPTFIR
KFPDGTERGKFLALDLGGTNFRVLLVELQSGLRRSVRMYNKIYAIPLEIMQGTGEELFDHIVECIADFLD
YMGMKGARLPLGFTFSFPCVQTRLDQGILVNWTKGFKATDVEGEDVVSLLKEAIKRHGELDLDIVALLND
TVGTMMTCAYENPNCEIGMIAGTGSNACYMEEMRNIEIVEGDEGRMCINTEWGAFGDNGCLDDIRTTFDR
LVDINSLNPGKQKYEKMISGMYLGEIVRNILINFTKQGLLFRGLISQSLKTSGIFETKYLSQIESDRLAL
LQVRTILQKLGLDSTCDDSIIVKEVCGVVSKRAAQLCGAGMAAVVDKIRENRGLDHLEITVGVDGTLYKL
HPHFSTILQETVKDLAPNCDVTFMLSEDGSGKGAALVTAAAKRPDKDGQH
>anc95_altAll|avgPP:0.943|lnPP:-104.821|num_ambig:57|num_ambig_gaps:0
LYHMQLNSDQLQDIMTRFKAEMRKGLDTNSTSTVKMLPTFVRSTPDGSEKGDFLALDLGGSKFRVLRVKV
SEDGRQVVHIESQVYPMPEDMKHGNSSQFFDYVAECLGDFMEKQEIKDKKLPLGFTFSFPCRQTKLDEGV
LISWSKGFKIKGVQGADVVSLLRKAIERRGDYDVDVLALVNDTVGTMMTCGFDDPYCEVGVIIGTGTNAC
YMEEMRHIELVEGDEGRMCINTEWGAFGDDGSLEDIRTEFDREVDSGSINPGKQLFEKMISAMYMGELVR
LILVKIAKNGLLFKGKTSSKLLTKGSFDIKHVSAIQNYKTGLENAMEILTDLELDPSEEDCIAVQHICTI
VSSRSANLCAATLATILTHLKKNKKIKRLRTTVGIDGTVYRTNPQYAKHLHKMVRRLVPDCDVRFLLAET
GSGKGAAMVTAVAYRLVHQRKYIDKTLAPLKLSREQLLEVMKKMRSEMEQGLKKETHSTATVKMLPTFIR
KFPDGTERGKFLALDLGGTNLRVLLVKIRSGLRKSVQMYNKIYSIPMEIMQGTGEELFDHIVECIADFLD
YMGMKGARLPLGFTFSFPCVQTRLDQGILVNWTKGFKATDVEGEDVVTLLKEAIKRHGELDLDIVALLND
TVGTMMTCAYENPNCEIGMIAGTGSNACYMEEMKNIEIVEGDEGRMCINTEWGAFGDNGCLDDIRTTFDK
LVDINSLNPGKQKYEKMISGMYLGEIVRNILINFTKHGLLFRGLVSQSLKTSGIFETKYLSQIESDRLAL
LQVRTILQQLGLDSTCDDSIIVKEVCGVVSRRAAQLCGAGMAAVVDKIRENRGLDHLEITVGVDGTLYKL
HPHFSTILQETVKELAPNCDVTFMLSEDGSGKGAALVTAAARRPDKDGQH
>anc96|avgPP:0.973|lnPP:-41.981|num_ambig:29|num_ambig_gaps:0
LYHMRLSDDTLQDIMTRFRAEMEKGLGRDTNPTATVKMLPTFVRSTPDGSEKGDFLALDLGGSKFRVLRV
KVSEDGKQKVEMESQIYPTPEEIIHGNGTQLFDYVAECLGDFMEKQQIKDKKLPLGFTFSFPCRQTKLDE
GILISWTKGFKARGVQGADVVSLLRKAIKKRGDFDVDVLALVNDTVGTMMTCGYDDQYCEVGVIIGTGTN
ACYMEEMRHIDLVEGDEGRMCINTEWGAFGDDGSLEDIRTEFDREIDRGSLNPGKQLFEKMISGMYMGEL

VRLILVKMAKKGLLFKGKISAELLTKGKFETKHVSAIEKYKEGLSNAKEILTDLGLEPSEEDCIAVQHVC
TIVSFRSANLCAATLAAILTRLRENKKVKRLRTTVGMDGTVYKTHPQYAKRLHKVVRRLVPDCDVRFLLS
ESGSGKGAAMVTAVAYRLVSQRKQIDETLAPFKLSREQLLEVKKKMRTEMERGLKKETHHTATVKMLPTY
VRGTPDGTERGKFLALDLGGTNFRVLLVKIRSGLRRSVRMYNKIYAIPLEIMQGTGEELFDHIVQCISDF
LDYMGMKGARLPLGFTFSFPCRQTSLDQGILVNWTKGFKATDCEGEDVVNLLREAIKRRNEFDLDIVAVV
NDTVGTMMTCAYEDPNCEIGLIAGTGSNACYMEEMRNIEIVEGDEGRMCINTEWGAFGDNGCLDDIRTKF
DRAVDEGSLNPGKQRYEKMISGMYLGEIVRNILIDFTKRGLLFRGQISERLKTRGIFETKFLSQIESDRL
ALLQVRAILQQLGLDSTCDDSIIVKEVCGAVSRRAAQLCGAGMAAVVDKIRENRGLDHLEITVGVDGTLY
KLHPHFSRILQETVKELAPKCDVTFMLSEDGSGKGAALITAVARRLHEAGQH
>anc96_altAll|avgPP:0.970|lnPP:-52.879|num_ambig:29|num_ambig_gaps:0
LYHMRLSDETLQDIMTRFRAEMEKGLGRDTNPTATVKMLPTFVRSTPDGSEKGDFLALDLGGSKFRVLRV
KVSEDGKQKVQMESQIYPTPEDIIHGNGTELFDYVAECLGDFMEKQQIKDKKLPLGFTFSFPCRQTKLDE
GVLISWTKGFKARGVQGTDVVSLLRKAIKKRGDYDVDVLALVNDTVGTMMTCGFDDQHCEVGVIIGTGTN
ACYMEEMRHIDLVEGDEGRMCINTEWGAFGDDGSLEDIRTEFDREIDLGSINPGKQLFEKMISGMYMGEL
VRLILVKMAKKGLLFKGKISPELLTKGKFETKHVSAIEKYKEGLSNAKEILTDLGLDPSEEDCIAVQHVC
TIVSFRSANLCAAALAAILTRLKENKKVKRLRTTVGVDGTVYKTHPQYAKRLHKMVRRLVPDCDVRFLLS
ESGSGKGAAMVTAVAYRLVSQRKQIDETLAPFKLSREQLLEVKKKMRTEMERGLKKETHNTATVKMLPTY
VRSTPDGTERGKFLALDLGGTNFRVLLVKIRSGLRRSVRMYNKIYAIPLEIMQGTGEELFDHIVQCISDF
LDYMGMKGARLPLGFTFSFPCRQTGLDQGILVNWTKGFKATDCEGEDVVDLLREAIKRREEFDLDIVAVV
NDTVGTMMTCAYEDPNCEIGLIAGTGSNACYMEEMRNIEMVEGDEGRMCINTEWGAFGDNGCLDDIRTQF
DREVDEGSLNPGKQRYEKMISGMYLGEIVRNILIDFTKQGLLFRGQISERLKTRGIFETKFLSQIESDRL
ALLQVRSILQQLGLDSTCDDSIIVKEVCGAVSRRAAQLCGAGMAAVVDKIRENRGLDHLEITVGVDGTLY
KLHPHFSKILQETVKELAPQCDVTFMLSEDGSGKGAALITAVARRLHEAGQH
>anc97|avgPP:0.981|lnPP:-28.484|num_ambig:24|num_ambig_gaps:0
LYHMRLSDETLQDIMTRFRTEMEKGLGRDTNPTATVKMLPTFVRSTPDGSEKGDFLALDLGGSNFRVLRV
KVSDDGKQKVEMESQIYATPEDIIHGSGTQLFDHVAECLGDFMEKQQIKDKKLPLGFTFSFPCRQTKLDE
GILITWTKGFKASGVEGADVVKLLRKAIKKRGDYDVDIMAVVNDTVGTMMTCGYDDQRCEVGLIIGTGTN
ACYMEEMRHIDLVEGDEGRMCINTEWGAFGDDGSLEDIRTEFDREIDRGSLNPGKQLFEKMISGMYMGEL
VRLILVKMAKEGLLFEGKITPELLTKGKFETKHVSAIEKSKEGLSKAKEILTKLGLEPSEEDCIAVQHVC
TIVSFRSANLCAATLAAILTRLKENKGVKRLRTTVGVDGTVYKTHPQYARRLHKTVRRLVPDCDVRFLLS
ESGSGKGAAMVTAVAYRLASQRKQIDETLAPFKLSHEQLLEVKKRMRTEMERGLKKETHNSATVKMLPTY
VRSTPDGTEKGDFLALDLGGTNFRVLLVKIRSGKRRSVEMHNKIYAIPLEVMQGTGEELFDHIVHCISDF
LDYMGMKGARLPLGFTFSFPCKQTSLDQGILLNWTKGFKATDCEGEDVVNLLREAIKRREEFDLDVVAVV
NDTVGTMMTCAYEDPNCEIGLIVGTGSNACYMEEMRNIEMVEGDEGRMCINMEWGAFGDNGCLDDIRTEF
DRAVDELSLNPGKQRYEKMISGMYLGEIVRNILIDFTKRGLLFRGQISERLKTRGIFETKFLSQIESDRL
ALLQVRAILQQLGLDSTCDDSIIVKEVCGAVSRRAAQLCGAGMAAVVDKIRENRGLDHLEITVGVDGTLY
KLHPHFSKIMHETVKELAPKCDVTFLLSEDGSGKGAALITAVACRLREAGQH
>anc97_altAll|avgPP:0.976|lnPP:-44.247|num_ambig:24|num_ambig_gaps:0
LYHMRLSDETLQDIMTRFRAEMEKGLGRDTNPTATVKMLPTFVRSTPDGSEKGDFLALDLGGSNFRVLRV
KVSDDGKQKVQMESQIYATPEDIMHGSGTQLFDHVAECLGDFMEKQQIKDKKLPLGFTFSFPCRQTKLDE
SVLITWTKGFKASGVEGADVVSLLRKAIKKRGDFDVDIMAVVNDTVGTMMTCGFDDQHCEVGLIIGTGTN
ACYMEEMRHIDLVEGDEGRMCINTEWGAFGDDGSLEDIRTEFDREIDRGSLNPGKQLFEKMISGMYMGEL
VRLILVKMAKEGLLFEGRITPELLTKGKFETKHVSAIEKSKEGLNKAKEILTRLGLEPSEEDCIAVQHVC
TIVSFRSANLCAAALAAILTRLRENKGVQRLRTTVGVDGTVYKTHPQYARRLHKTVRRLVPDCDVRFLLS
ESGSGKGAAMVTAVAYRLASQRKQIDETLAPFKLSHEQLLEVKKRMRTEMERGLKKETHNTATVKMLPTY
VRSTPDGTEKGDFLALDLGGTNFRVLLVKIRSGKRRSVEMHNKIYAIPLEVMQGTGEELFDHIVHCISDF
LDYMGMKGARLPLGFTFSFPCRQTSLDQGILLNWTKGFKATDCEGEDVVNLLREAIKRREEFELDVVAIV
NDTVGTMMTCAYEDPNCEVGLIVGTGSNACYMEEMRNIEMVEGDEGRMCVNMEWGAFGDNGCLDDIRTQF
DKAVDELSLNPGKQRYEKMISGMYLGEIVRNILIDFTKRGLLFRGQISERLKTRGIFETKFLSQIESDRL
ALLQVRAILQQLGLDSTCDDSIIVKEVCGAVSRRAAQLCGAGMAAVVDKIRENRGLDHLEITVGVDGTLY
KLHPHFSRIMHETVKELAPKCDVTFLLSEDGSGKGAALITAVACRLREAGQH
>anc98|avgPP:0.993|lnPP:-10.767|num_ambig:11|num_ambig_gaps:0

LYHMRLSDETLQEVSNRFRKEMEKGLGADTNPTASVKMLPTFVRSTPDGTEDGDFLALDLGGTNFRVLRV
KVSDNGLKKVEMENQIYAIPEDLMRGSGTQLFDHIAECLANFMEKLHIKDKKLPLGFTFSFPCHQTKLDE
SILVTWTKGFKASGVEGKDVVSLLRKAIKKRGDFDIDIVAVVNDTVGTMMTCGYDDHNCEIGLIVGTGTN
ACYMEEMRHIDLVEGDEGRMCINMEWGAFGDDGVLDDIRTEFDREIDMGSLNPGKQLFEKMISGMYMGEL
VRLILVKMAKEDLLFGGRITPDLLTTGHFETRFVSAIEKDKEGIQKAHEILTKLGLEPSQEDCVATHRIC
QIVSTRSANLCAATLAAVLRRIKENKGVERLRSTVGVDGSVYKKHPHFARRLHKTVRRLVPDCDIRFVRS
EDGSGKGAAMVTAVAYRLAAQQKARQKTLEALKLSHEQLLEVKKRMRIEMERGLGKETHNSATVKMLPTY
VRSTPDGTEKGDFLALDLGGTNFRVLLVRVRNGMRRGVEMHNKIYSIPQDIMQGTGEELFDHIVHCIADF
LEYMGMKGVSLPLGFTFSFPCHQNNLDEGILLKWTKGFKATGCEGEDVVNLLKEAIHRREEFDLDVVAVV
NDTVGTMMTCGYEDPYCEVGLIVGTGSNACYMEEMRNVELVEGEEGRMCINMEWGAFGDNGCLEDFRTEF
DRAVDELSLNPGKQRYEKMISGMYLGEIVRNILIDFTKRGLLFRGRISERLKTRGIFETKFLSQIESDCL
ALLQVRSILQHLGLESTCDDSIIVKEVCTVVARRAAQLCGAGMAAVVDKIRENRGLDFLKVTVGVDGTLY
KLHPHFSTIMHETVKELSPKCEVTFLQSEDGSGKGAALITAVACRIREAGQR
>anc98_altAll|avgPP:0.991|lnPP:-14.784|num_ambig:11|num_ambig_gaps:0
LYHMRLSDETLQDVSNRFRKEMEKGLGADTNPTASVKMLPTFVRSTPDGTENGDFLALDLGGTNFRVLRV
KVSDNGLQKVEMENQIYAIPEDLMRGSGTQLFDHIAECLANFMEKLHIKDKKLPLGFTFSFPCHQTKLDE
SILVTWTKGFKASGVEGKDVVSLLRKAIKKRGDFDIDIVAVVNDTVGTMMTCGYDDHNCEIGLIVGTGTN
ACYMEEMRHIDLVEGDEGRMCINMEWGAFGDDGVLNDIRTEFDREIDMGSLNPGKQLFEKMISGMYMGEL
VRLILVKMAKEDLLFGGRITPDLLTTGHFETRFVSAIEKDKEGIQKAHEILTKLGLEPSQEDCVATHRIC
QIVSTRSANLCAATLAAVLRRIKENKGVERLRSTVGVDGSVYKKHPHFARRLHKTVRRLVPDCDIRFVRS
EDGSGKGAAMVTAVAYRLAAQQKARQKTLEALKLSHEQLLEVKKRMRIEMERGLGKETHSSATVKMLPTY
VRSTPDGTEKGDFLALDLGGTNFRVLLVRVRNGMRRGVEMHNKIYSIPQEIMQGTGEELFDHIVHCIADF
LEYMGMKGVSLPLGFTFSFPCHQNNLDEGILLKWTKGFKATGCEGEDVVNLLKEAIHRREEFDLDVVAVV
NDTVGTMMTCGYEDPYCEVGLIVGTGSNACYMEEMRNVELVEGEEGRMCVNMEWGAFGDNGCLDDFRTEF
DRAVDELSLNPGKQRFEKMISGMYLGEIVRNILIDFTKRGLLFRGRISERLKTRGIFETKFLSQIESDRL
ALLQVRSILQHLGLESTCDDSIIVKEVCTVVARRAAQLCGAGMAAVVDKIRENRGLDFLKVTVGVDGTLY
KLHPHFSTVMHETVKELSPKCEVTFLQSEDGSGKGAALITAVACRIREAGQR
>anc99|avgPP:0.999|lnPP:-1.081|num_ambig:0|num_ambig_gaps:0
LYHMRLSDETLLEISKRFRKEMEKGLGATTHPTASVKMLPTFVRSTPDGTEHGEFLALDLGGTNFRVLRV
RVTDNGLQKVEMENQIYAIPEDIMRGSGTQLFDHIAECLANFMDKLQIKDKKLPLGFTFSFPCLQTKLDE
SFLVSWTKGFKSSGVEGKDVVTLIRKAIQRRGDFDIDIVAVVNDTVGTMMTCGYDDQNCEIGLIVGTGSN
ACYMEEMRHIDTVEGDEGRMCINMEWGAFGDDGALDDIRTEFDQEIDMGSLNPGKQLFEKMISGMYMGEL
VRLILVKMAKEELLFGGKLSPELLATGHFETKDVSDIEGEKDGIRKAREVLVRLGLDPTQEDCVATHRVC
QIVSTRSASLCAATLAAVLRRIKENKGEERLRSTIGVDGSVYKKHPHFAKRLHKTVRRLVPDCDVRFLRS
EDGSGKGAAMVTAVAYRLADQHRARQKTLESLKLSREQLLEVKRRMKVEMERGLSKETHAIAPVKMLPTY
VCATPDGTEKGDFLALDLGGTNFRVLLVRVRNGKRRGVEMHNKIYSIPQEVMHGTGDELFDHIVQCIADF
LEYMGMKGVSLPLGFTFSFPCQQNSLDESILLKWTKGFKASGCEGEDVVTLLKEAIHRREEFDLDVVAVV
NDTVGTMMTCGYEDPHCEVGLIVGTGSNACYMEEMRNVELVEGEEGRMCVNMEWGAFGDNGCLDDFRTEF
DVAVDELSLNPGKQRFEKMISGMYLGEIVRNILIDFTKRGLLFRGRISERLKTRGIFETKFLSQIESDCL
ALLQVRAILHHLGLESTCDDSIIVKEVCTVVARRAAQLCGAGMAAVVDKIRENRGLDTLKVTVGVDGTLY
KLHPHFAKVMHETVKDLAPKCDVSFLESEDGSGKGAALITAVACRIREAGQR
>anc99_altAll|avgPP:0.999|lnPP:-1.081|num_ambig:0|num_ambig_gaps:0
LYHMRLSDETLLEISKRFRKEMEKGLGATTHPTASVKMLPTFVRSTPDGTEHGEFLALDLGGTNFRVLRV
RVTDNGLQKVEMENQIYAIPEDIMRGSGTQLFDHIAECLANFMDKLQIKDKKLPLGFTFSFPCLQTKLDE
SFLVSWTKGFKSSGVEGKDVVTLIRKAIQRRGDFDIDIVAVVNDTVGTMMTCGYDDQNCEIGLIVGTGSN
ACYMEEMRHIDTVEGDEGRMCINMEWGAFGDDGALDDIRTEFDQEIDMGSLNPGKQLFEKMISGMYMGEL
VRLILVKMAKEELLFGGKLSPELLATGHFETKDVSDIEGEKDGIRKAREVLVRLGLDPTQEDCVATHRVC
QIVSTRSASLCAATLAAVLRRIKENKGEERLRSTIGVDGSVYKKHPHFAKRLHKTVRRLVPDCDVRFLRS
EDGSGKGAAMVTAVAYRLADQHRARQKTLESLKLSREQLLEVKRRMKVEMERGLSKETHAIAPVKMLPTY
VCATPDGTEKGDFLALDLGGTNFRVLLVRVRNGKRRGVEMHNKIYSIPQEVMHGTGDELFDHIVQCIADF
LEYMGMKGVSLPLGFTFSFPCQQNSLDESILLKWTKGFKASGCEGEDVVTLLKEAIHRREEFDLDVVAVV
NDTVGTMMTCGYEDPHCEVGLIVGTGSNACYMEEMRNVELVEGEEGRMCVNMEWGAFGDNGCLDDFRTEF

DVAVDELSLNPGKQRFEKMISGMYLGEIVRNILIDFTKRGLLFRGRISERLKTRGIFETKFLSQIESDCL
ALLQVRAILHHLGLESTCDDSIIVKEVCTVVARRAAQLCGAGMAAVVDKIRENRGLDTLKVTVGVDGTLY
KLHPHFAKVMHETVKDLAPKCDVSFLESEDGSGKGAALITAVACRIREAGQR
>anc100|avgPP:0.998|lnPP:-2.359|num_ambig:2|num_ambig_gaps:0
LYHMRLSDETLQEISKRFRKEMEKGLGATTHPTASVKMLPTFVRSTPDGTEHGEFLALDLGGTNFRVLRV
RVTDNGLQKVEMENQIYAIPEDIMRGSGTQLFDHIAECLANFMDKLQIKDKKLPLGFTFSFPCHQTKLDE
SFLVSWTKGFKSSGVEGRDVVTLIRKAIQRRGDFDIDIVAVVNDTVGTMMTCGYDDQNCEIGLIVGTGSN
ACYMEEMRHIDTVEGDEGRMCINMEWGAFGDDGALDDIRTEFDQEIDMGSLNPGKQLFEKMISGMYMGEL
VRLILVKMAKEELLFGGKLSPELLATGHFETKDVSDIEGEKDGIRKAREVLVRLGLDPTQEDCVATHRIC
QIVSTRSASLCAATLAAVLRRIKENKGEERLRSTIGVDGSVYKKHPHFAKRLHKTVRRLVPDCDVRFLRS
EDGSGKGAAMVTAVAYRLADQHRARQKTLESLKLSHEQLLEVKRRMKVEMERGLSKETHAIAPVKMLPTY
VCATPDGTEKGDFLALDLGGTNFRVLLVRVRNGKRRGVEMHNKIYSIPQEVMHGTGDELFDHIVQCIADF
LEYMGMKGVSLPLGFTFSFPCQQNSLDESILLKWTKGFKASGCEGEDVVTLLKEAIHRREEFDLDVVAVV
NDTVGTMMTCGYEDPHCEVGLIVGTGSNACYMEEMRNVELVEGEEGRMCVNMEWGAFGDNGCLDDFRTEF
DVAVDELSLNPGKQRFEKMISGMYLGEIVRNILIDFTKRGLLFRGRISERLKTRGIFETKFLSQIESDCL
ALLQVRAILHHLGLESTCDDSIIVKEVCTVVARRAAQLCGAGMAAVVDKIRENRGLDTLKVTVGVDGTLY
KLHPHFAKVMHETVKDLAPKCDVSFLESEDGSGKGAALITAVACRIREAGQR
>anc100_altAll|avgPP:0.998|lnPP:-2.419|num_ambig:2|num_ambig_gaps:0
LYHMRLSDETLLEISKRFRKEMEKGLGATTHPTASVKMLPTFVRSTPDGTEHGEFLALDLGGTNFRVLRV
RVTDNGLQKVEMENQIYAIPEDIMRGSGTQLFDHIAECLANFMDKLQIKDKKLPLGFTFSFPCHQTKLDE
SFLVSWTKGFKSSGVEGRDVVTLIRKAIQRRGDFDIDIVAVVNDTVGTMMTCGYDDQNCEIGLIVGTGSN
ACYMEEMRHIDTVEGDEGRMCINMEWGAFGDDGALDDIRTEFDQEIDMGSLNPGKQLFEKMISGMYMGEL
VRLILVKMAKEELLFGGKLSPELLATGHFETKDVSDIEGEKDGIRKAREVLMRLGLDPTQEDCVATHRIC
QIVSTRSASLCAATLAAVLRRIKENKGEERLRSTIGVDGSVYKKHPHFAKRLHKTVRRLVPDCDVRFLRS
EDGSGKGAAMVTAVAYRLADQHRARQKTLESLKLSHEQLLEVKRRMKVEMERGLSKETHAIAPVKMLPTY
VCATPDGTEKGDFLALDLGGTNFRVLLVRVRNGKRRGVEMHNKIYSIPQEVMHGTGDELFDHIVQCIADF
LEYMGMKGVSLPLGFTFSFPCQQNSLDESILLKWTKGFKASGCEGEDVVTLLKEAIHRREEFDLDVVAVV
NDTVGTMMTCGYEDPHCEVGLIVGTGSNACYMEEMRNVELVEGEEGRMCVNMEWGAFGDNGCLDDFRTEF
DVAVDELSLNPGKQRFEKMISGMYLGEIVRNILIDFTKRGLLFRGRISERLKTRGIFETKFLSQIESDCL
ALLQVRAILHHLGLESTCDDSIIVKEVCTVVARRAAQLCGAGMAAVVDKIRENRGLDTLKVTVGVDGTLY
KLHPHFAKVMHETVKDLAPKCDVSFLESEDGSGKGAALITAVACRIREAGQR
>anc101|avgPP:0.998|lnPP:-3.433|num_ambig:4|num_ambig_gaps:0
LYHMRLSDETLQEISKRFRKEMEKGLGATTHPTASVKMLPTFVRSTPDGTEHGEFLALDLGGTNFRVLRV
RVTDNGLQKVEMENQIYAIPEDIMRGSGTQLFDHIAECLANFMDKLQIKDKKLPLGFTFSFPCHQTKLDE
SFLVSWTKGFKSSGVEGRDVVTLIRKAIQRRGDFDIDIVAVVNDTVGTMMTCGYDDQNCEIGLIVGTGSN
ACYMEEMRHIDTVEGDEGRMCINMEWGAFGDDGALDDIRTEFDQEIDMGSLNPGKQLFEKMISGMYMGEL
VRLILVKMAKEELLFGGKLSPELLTTGHFETKDVSDIEGEKDGIRKAREVLMRLGLDPTQEDCVATHRIC
QIVSTRSASLCAATLAAVLRRIKENKGEERLRSTIGVDGSVYKKHPHFAKRLHKTVRRLVPDCDVRFLRS
EDGSGKGAAMVTAVAYRLADQHRARQKTLESLKLSHEQLLEVKRRMKVEMERGLSKETHASAPVKMLPTY
VCATPDGTEKGDFLALDLGGTNFRVLLVRVRNGKRRGVEMHNKIYSIPQEVMHGTGDELFDHIVQCIADF
LEYMGMKGVSLPLGFTFSFPCQQNSLDESILLKWTKGFKASGCEGEDVVTLLKEAIHRREEFDLDVVAVV
NDTVGTMMTCGYEDPHCEVGLIVGTGSNACYMEEMRNVELVEGEEGRMCVNMEWGAFGDNGCLDDFRTEF
DVAVDELSLNPGKQRFEKMISGMYLGEIVRNILIDFTKRGLLFRGRISERLKTRGIFETKFLSQIESDCL
ALLQVRAILQHLGLESTCDDSIIVKEVCTVVARRAAQLCGAGMAAVVDKIRENRGLDTLKVTVGVDGTLY
KLHPHFAKVMHETVKDLAPKCDVSFLESEDGSGKGAALITAVACRIREAGQR
>anc101_altAll|avgPP:0.997|lnPP:-4.513|num_ambig:4|num_ambig_gaps:0
LYHMRLSDETLLEISKRFRKEMEKGLGATTHPTASVKMLPTFVRSTPDGTEHGEFLALDLGGTNFRVLRV
RVTDNGLQKVEMENQIYAIPEDIMRGSGTQLFDHIAECLANFMDKLQIKDKKLPLGFTFSFPCHQTKLDE
SFLVSWTKGFKSSGVEGRDVVTLIRKAIQRRGDFDIDIVAVVNDTVGTMMTCGYDDQNCEIGLIVGTGSN
ACYMEEMRHIDTVEGDEGRMCINMEWGAFGDDGALNDIRTEFDQEIDMGSLNPGKQLFEKMISGMYMGEL
VRLILVKMAKEELLFGGKLSPELLATGHFETKDVSDIEGEKDGIRKAREVLVRLGLDPTQEDCVATHRIC
QIVSTRSASLCAATLAAVLRRIKENKGEERLRSTIGVDGSVYKKHPHFAKRLHKTVRRLVPDCDVRFLRS

EDGSGKGAAMVTAVAYRLADQHRARQKTLESLKLSHEQLLEVKRRMKVEMERGLSKETHASAPVKMLPTY
VCATPDGTEKGDFLALDLGGTNFRVLLVRVRNGKRRGVEMHNKIYSIPQEVMHGTGDELFDHIVQCIADF
LEYMGMKGVSLPLGFTFSFPCQQNSLDESILLKWTKGFKASGCEGEDVVTLLKEAIHRREEFDLDVVAVV
NDTVGTMMTCGYEDPHCEVGLIVGTGSNACYMEEMRNVELVEGEEGRMCVNMEWGAFGDNGCLDDFRTEF
DVAVDELSLNPGKQRFEKMISGMYLGEIVRNILIDFTKRGLLFRGRISERLKTRGIFETKFLSQIESDCL
ALLQVRAILQHLGLESTCDDSIIVKEVCTVVARRAAQLCGAGMAAVVDKIRENRGLDTLKVTVGVDGTLY
KLHPHFAKVMHETVKDLAPKCDVSFLESEDGSGKGAALITAVACRIREAGQR
>anc102|avgPP:0.997|lnPP:-5.384|num_ambig:4|num_ambig_gaps:0
LYHMRLSDETLQEISKRFRKEMEKGLGATTHPTASVKMLPTFVRSTPDGTEHGEFLALDLGGTNFRVLRV
RVTDNGLQKVEMENQIYAIPEDIMRGSGTQLFDHIAECLANFMDKLQIKDKKLPLGFTFSFPCHQTKLDE
SFLVSWTKGFKSSGVEGRDVVTLIRKAIQRRGDFDIDIVAVVNDTVGTMMTCGYDDQNCEIGLIVGTGSN
ACYMEEMRHIDTVEGDEGRMCINMEWGAFGDDGALDDIRTEFDQEIDMGSLNPGKQLFEKMISGMYMGEL
VRLILVKMAKEELLFGGKLSPELLTTGHFETKDVSDIEGEKDGIRKAHEVLVRLGLQPTQEDCVATHRIC
QIVSTRSANLCAATLAAVLRRLKENKGEERLRSTIGVDGSVYKKHPHFAKRLHKTVRRLVPDCDVRFLRS
EDGSGKGAAMVTAVAYRLADQHRARQKTLESLKLSHEQLLEVKRRMKVEMERGLSKETHASAPVKMLPTY
VCATPDGTEKGDFLALDLGGTNFRVLLVRVRNGKRRGVEMHNKIYSIPQEVMHGTGDELFDHIVQCIADF
LEYMGMKGVSLPLGFTFSFPCQQNSLDESILLKWTKGFKASGCEGEDVVNLLKEAIHRREEFDLDVVAVV
NDTVGTMMTCGYEDPHCEVGLIVGTGSNACYMEEMRNVELVEGEEGRMCVNMEWGAFGDNGCLDDFRTEF
DVAVDELSLNPGKQRFEKMISGMYLGEIVRNILIDFTKRGLLFRGRISERLKTRGIFETKFLSQIESDCL
ALLQVRAILQHLGLESTCDDSIIVKEVCTVVARRAAQLCGAGMAAVVDKIRENRGLDTLKVTVGVDGTLY
KLHPHFAKVMHETVKDLAPKCDVSFLESEDGSGKGAALITAVACRIREAGQR
>anc102_altAll|avgPP:0.996|lnPP:-6.293|num_ambig:4|num_ambig_gaps:0
LYHMRLSDETLQEISKRFRKEMEKGLGATTHPTASVKMLPTFVRSTPDGTEHGEFLALDLGGTNFRVLRV
RVTDNGLQKVEMENQIYAIPEDIMRGSGTQLFDHIAECLANFMDKLQIKDKKLPLGFTFSFPCHQTKLDE
SFLVSWTKGFKSSGVEGRDVVSLIRKAIQRRGDFDIDIVAVVNDTVGTMMTCGYDDQNCEIGLIVGTGSN
ACYMEEMRHIDTVEGDEGRMCINMEWGAFGDDGALNDIRTEFDQEIDMGSLNPGKQLFEKMISGMYMGEL
VRLILVKMAKEELLFGGKLSPELLTTGHFETKDVSDIEGEKDGIRKAHEVLMRLGLQPTQEDCVATHRIC
QIVSTRSANLCAATLAAVLRRLKENKGEERLRSTIGVDGSVYKKHPHFAKRLHKTVRRLVPDCDVRFLRS
EDGSGKGAAMVTAVAYRLADQHRARQKTLESLKLSHEQLLEVKRRMKVEMERGLSKETHASAPVKMLPTY
VCATPDGTEKGDFLALDLGGTNFRVLLVRVRNGKRRGVEMHNKIYSIPQEVMHGTGDELFDHIVQCIADF
LEYMGMKGVSLPLGFTFSFPCQQNSLDESILLKWTKGFKASGCEGEDVVTLLKEAIHRREEFDLDVVAVV
NDTVGTMMTCGYEDPHCEVGLIVGTGSNACYMEEMRNVELVEGEEGRMCVNMEWGAFGDNGCLDDFRTEF
DVAVDELSLNPGKQRFEKMISGMYLGEIVRNILIDFTKRGLLFRGRISERLKTRGIFETKFLSQIESDCL
ALLQVRAILQHLGLESTCDDSIIVKEVCTVVARRAAQLCGAGMAAVVDKIRENRGLDTLKVTVGVDGTLY
KLHPHFAKVMHETVKDLAPKCDVSFLESEDGSGKGAALITAVACRIREAGQR
>anc103|avgPP:0.995|lnPP:-7.625|num_ambig:9|num_ambig_gaps:0
LYHMRLSDETLQEVSKRFRKEMEKGLGADTHPTASVKMLPTFVRSTPDGTEDGDFLALDLGGTNFRVLRV
RVTDNGLQKVEMENQIYAIPEDIMRGSGTQLFDHIAECLANFMDKLQIKDKKLPLGFTFSFPCHQTKLDE
SFLVSWTKGFKSSGVEGRDVVSLIRKAIQRRGDFDIDIVAVVNDTVGTMMTCGYDDQNCEIGLIVGTGTN
ACYMEEMRHIDTVEGDEGRMCINMEWGAFGDDGVLDDIRTEFDREIDMGSLNPGKQLFEKMISGMYMGEL
VRLILVKMAKEELLFGGKLSPELLTTGHFETKDVSDIEGEKDGIRKAHEVLTRLGLQPSQEDCVATHRIC
QIVSTRSANLCAATLAAVLRRLKENKGEERLRSTIGVDGSVYKKHPHFAKRLHKTVRRLVPDCDVRFLRS
EDGSGKGAAMVTAVAYRLADQHRARQKTLEALKLSHEQLLEVKKRMKVEMERGLGKETHASASVKMLPTY
VCATPDGTEKGDFLALDLGGTNFRVLLVRVRNGMRRGVEMHNKIYSIPQEVMHGTGDELFDHIVQCIADF
LEYMGMKGVSLPLGFTFSFPCQQNSLDESILLKWTKGFKATGCEGEDVVNLLKEAIHRREEFDLDVVAVV
NDTVGTMMTCGYEDPHCEVGLIVGTGSNACYMEEMRNVELVEGEEGRMCINMEWGAFGDNGCLDDFRTEF
DVAVDELSLNPGKQRFEKMISGMYLGEIVRNILIDFTKRGLLFRGRISERLKTRGIFETKFLSQIESDCL
ALLQVRAILQHLGLESTCDDSIIVKEVCTVVARRAAQLCGAGMAAVVDKIRENRGLDSLKVTVGVDGTLY
KLHPHFAKVMHETVKELAPKCDVSFLESEDGSGKGAALITAVACRIREAGQR
>anc103_altAll|avgPP:0.993|lnPP:-12.869|num_ambig:9|num_ambig_gaps:0
LYHMRLSDETLQEISKRFRKEMEKGLGADTHPTASVKMLPTFVRSTPDGTEDGDFLALDLGGTNFRVLRV
RVTDNGLQKVEMENQIYAIPEDIMRGSGTQLFDHIAECLANFMDKLQIKDKKLPLGFTFSFPCHQTKLDE

SFLVSWTKGFKSSGVEGRDVVSLIRKAIQRRGDFDIDIVAVVNDTVGTMMTCGYDDQNCEIGLIVGTGSN
ACYMEEMRHIDTVEGDEGRMCINMEWGAFGDDGVLNDIRTEFDREIDMGSLNPGKQLFEKMISGMYMGEL
VRLILVKMAKEELLFGGKLSPELLTTGHFETKDVSDIEGEKDGIRKAHEVLTRLGLEPTQEDCVATHRIC
QIVSTRSANLCAATLAAVLRRLKENKGEERLRSTIGVDGSVYKKHPHFAKRLHKTVRRLVPDCDVRFLRS
EDGSGKGAAMVTAVAYRLADQHRARQKTLEALKLSHEQLLEVKRRMKVEMERGLGKETHASASVKMLPTY
VCATPDGTEKGDFLALDLGGTNFRVLLVRVRNGMRRGVEMHNKIYSIPQEVMHGTGDELFDHIVQCIADF
LEYMGMKGVSLPLGFTFSFPCQQNSLDESILLKWTKGFKATGCEGEDVVSLLKEAIHRREEFDLDVVAVV
NDTVGTMMTCGYEDPHCEVGLIVGTGSNACYMEEMRNVELVEGDEGRMCVNMEWGAFGDNGCLDDFRTEF
DVAVDELSLNPGKQRFEKMISGMYLGEIVRNILIDFTKRGLLFRGRISERLKTRGIFETKFLSQIESDCL
ALLQVRAILQHLGLESTCDDSIIVKEVCTVVARRAAQLCGAGMAAVVDKIRENRGLDSLKVTVGVDGTLY
KLHPHFAKVMHETVKELAPKCDVSFLESEDGSGKGAALITAVACRIREAGQR
>anc104|avgPP:0.992|lnPP:-12.261|num_ambig:10|num_ambig_gaps:0
LYHMRLSDETLQEVSKRFRKEMEKGLGADTHPTASVKMLPTFVRSTPDGTEDGDFLALDLGGTNFRVLRV
RVTDNGLQKVEMENQIYAIPEDLMRGSGTQLFDHIAECLANFMEKLQIKDKKLPLGFTFSFPCHQTKLDE
SFLVTWTKGFKSSGVEGRDVVSLLRKAIKKRGDFDIDIVAVVNDTVGTMMTCGYDDQNCEIGLIVGTGTN
ACYMEEMRHIDLVEGDEGRMCINMEWGAFGDDGVLDDIRTEFDREIDMGSLNPGKQLFEKMISGMYMGEL
VRLILVKMAKEELLFGGKLTPELLTTGHFETRDVSAIEGEKDGIRKAHEVLTRLGLEPSQEDCVATHRIC
QIVSTRSANLCAATLAAVLRRLKENKGEERLRSTIGVDGSVYKKHPHFARRLHKTVRRLVPDCDVRFLRS
EDGSGKGAAMVTAVAYRLADQHRARQKTLEALKLSHEQLLEVKKRMRVEMERGLGKETHASASVKMLPTY
VCATPDGTEKGDFLALDLGGTNFRVLLVRVRNGMRRGVEMHNKIYSIPQEIMQGTGDELFDHIVHCIADF
LEYMGMKGVSLPLGFTFSFPCHQNNLDEGILLKWTKGFKATGCEGEDVVNLLKEAIHRREEFDLDVVAVV
NDTVGTMMTCGYEDPHCEVGLIVGTGSNACYMEEMRNVELVEGEEGRMCINMEWGAFGDNGCLDDFRTEF
DVAVDELSLNPGKQRFEKMISGMYLGEIVRNILIDFTKRGLLFRGRISERLKTRGIFETKFLSQIESDCL
ALLQVRAILQHLGLESTCDDSIIVKEVCTVVARRAAQLCGAGMAAVVDKIRENRGLDSLKVTVGVDGTLY
KLHPHFSTIMHETVKELAPKCEVSFLQSEDGSGKGAALITAVACRIREAGQR
>anc104_altAll|avgPP:0.990|lnPP:-17.006|num_ambig:10|num_ambig_gaps:0
LYHMRLSDETLQEISKRFRKEMEKGLGADTHPTASVKMLPTFVRSTPDGTEDGDFLALDLGGTNFRVLRV
RVTDNGLQKVEMENQIYAIPEDLMRGSGTQLFDHIAECLANFMEKLQIKDKKLPLGFTFSFPCHQTKLDE
SFLVTWTKGFKSSGVEGRDVVSLLRKAIQKRGDFDIDIVAVVNDTVGTMMTCGYDDQNCEIGLIVGTGSN
ACYMEEMRHIDLVEGDEGRMCINMEWGAFGDDGVLNDIRTEFDREIDMGSLNPGKQLFEKMISGMYMGEL
VRLILVKMAKEELLFGGKLTPELLTTGHFETRDVSAIEGEKDGIRKAHEVLTRLGLEPTQEDCVATHRIC
QIVSTRSANLCAATLAAVLRRLKENKGEERLRSTIGVDGSVYKKHPHFARRLHKTVRRLVPDCDVRFLRS
EDGSGKGAAMVTAVAYRLADQHRARQKTLEALKLSHEQLLEVKRRMRIEMERGLGKETHASASVKMLPTY
VCATPDGTEKGDFLALDLGGTNFRVLLVRVRNGMRRGVEMHNKIYSIPQEIMQGTGDELFDHIVHCIADF
LEYMGMKGVSLPLGFTFSFPCHQNNLDEGILLKWTKGFKATGCEGEDVVSLLKEAIHRREEFDLDVVAVV
NDTVGTMMTCGYEDPHCEVGLIVGTGSNACYMEEMRNVELVEGEEGRMCVNMEWGAFGDNGCLDDFRTEF
DVAVDELSLNPGKQRFEKMISGMYLGEIVRNILIDFTKRGLLFRGRISERLKTRGIFETKFLSQIESDCL
ALLQVRAILQHLGLESTCDDSIIVKEVCTVVARRAAQLCGAGMAAVVDKIRENRGLDSLKVTVGVDGTLY
KLHPHFSTVMHETVKELAPKCEVSFLQSEDGSGKGAALITAVACRIREAGQR
>anc105|avgPP:1.000|lnPP:-0.505|num_ambig:1|num_ambig_gaps:0
LYHMRLSDETLQEVSQRFEKEMEKGLGAETNPTASVKMLPTFVRSTPDGTEDGDFLALDLGGTNFRVLRV
KVSDNGLQKVEMENQIYAIPEELMRGSGVQLFDHIAECLANFMDKLQIKDKKLPLGFTFSFPCHQTKLDE
SILVNWTKGFKCSSVEGRDVVSLLRKSIKKRGDFDIDIVAVVNDTVGTMMTCGYDDHNCEVGLIVGTGTN
ACYMEEMRHIDLVEGDEGRMCINMEWGAFGDDGVLNDIRTEFDREIDMGSLNPGKQLFEKMISGMYMGEL
VRLILVKMTKEGLLFGGRLTPDLLTTGHFETRFVSAIEKEKEGLQKAHEILSKLGLEPSHEDCLATLRIC
QIVSTRSASLCGATLAAVLRRIKDNKGVDRLRSTVGVDGSVYKKHPHFARRLHKTVRKLLPDCEIRFIRS
EDGSGKGAAMVTAVAYRLAAQHKARQKILEPLKLSHEQLLEVKERMRIEMEKGLGKETHAEATVKMLPTY
VCSTPDGTEKGDFLALDLGGTNFRVLLVRVRNGMRRGVEMHNKIYSIPVEIMQGTGEELFDHIVHCISDF
LEYMGMKGVSLPLGFTFSFPCQQNNLDEGILLKWTKGFKATGCEGEDVVGLLKEAIHRREEFDLDVVAVV
NDTVGTMMTCGYEDPYCEVGLIVGTGSNACYMEEMRNVELVEGDEGRMCVNMEWGAFGDSGCLDDIRTEF
DLAVDELSLNPGKQRFEKMISGMYLGEIVRNILMDFTKRGLLFRGRISERLKTRGIFETKFLSQIESDCL
ALLQVRSILQHLGLESTCDDSIIVKEVCTVVARRAAQLCGAGMAAVVDKIRENRGLDFLKITVGVDGTLY

```
KLHPHFSTVMHETVKQLSPKCEVTFLQSEDGSGKGAALITAVACRIREAGQR
>anc105_altAll|avgPP:0.999|lnPP:-1.083|num_ambig:1|num_ambig_gaps:0
LYHMRLSDETLQEVSQRFEKEMEKGLGAETNPTASVKMLPTFVRSTPDGTEDGDFLALDLGGTNFRVLRV
KVSDNGLQKVEMENQIYAIPEELMRGSGVQLFDHIAECLANFMDKLQIKDKKLPLGFTFSFPCHQTKLDE
SILVNWTKGFKCSSVEGRDVVSLLRKSIKKRGDFDIDIVAVVNDTVGTMMTCGYDDHNCEVGLIVGTGTN
ACYMEEMRHIDLVEGDEGRMCINMEWGAFGDDGVLNDIRTEFDREIDMGSLNPGKQLFEKMISGMYMGEL
VRLILVKMTKEGLLFGGRLTPDLLTTGHFETRFVSAIEKEKEGLQKAHEILSKLGLEPSHEDCLATLRIC
QIVSTRSASLCGATLAAVLRRIKDNKGVERLRSTVGVDGSVYKKHPHFARRLHKTVRKLLPDCEIRFIRS
EDGSGKGAAMVTAVAYRLAAQHKARQKILEPLKLSHEQLLEVKERMRIEMEKGLGKETHAEATVKMLPTY
VCSTPDGTEKGDFLALDLGGTNFRVLLVRVRNGMRRGVEMHNKIYSIPVEIMQGTGEELFDHIVHCISDF
LEYMGMKGVSLPLGFTFSFPCQQNNLDEGILLKWTKGFKATGCEGEDVVGLLKEAIHRREEFDLDVVAVV
NDTVGTMMTCGYEDPYCEVGLIVGTGSNACYMEEMRNVELVEGDEGRMCVNMEWGAFGDSGCLDDIRTEF
DLAVDELSLNPGKQRFEKMISGMYLGEIVRNILMDFTKRGLLFRGRISERLKTRGIFETKFLSQIESDCL
ALLQVRSILQHLGLESTCDDSIIVKEVCTVVARRAAQLCGAGMAAVVDKIRENRGLDFLKITVGVDGTLY
KLHPHFSTVMHETVKQLSPKCEVTFLQSEDGSGKGAALITAVACRIREAGQR
>anc106|avgPP:1.000|lnPP:-0.714|num_ambig:1|num_ambig_gaps:0
LYHMRLSDETLLEVSQRFEKEMEKGLGAETNPTASVKMLPTFVRSTPDGTEDGDFLALDLGGTNFRVLRV
KVSDNGLQKVEMENQIYAIPEELMRGSGVQLFDHIAECLANFMDKLQIKDKKLPLGFTFSFPCHQTKLDE
SILVNWTKGFKCSSVEGRDVVSMLRKSIKKRGDFDIDIVAVVNDTVGTMMTCGYDDHNCEVGLIVGTGTN
ACYMEEMRHIDLVEGDEGRMCINMEWGAFGDDGVLNDIRTEFDREIDMGSLNPGKQLFEKMISGMYMGEL
VRLILVKMAKEGLLFGGRLTPDLLTTGHFETRFVSAIEKEKEGLQKAHEILSKLGLEPSHEDCLATLRIC
QIVSTRSASLCGATLAAVLRRIKDNKGVERLRSTVGVDGSVYKKHPHFARRLHKTVRKLLPDCEIRFIRS
EDGSGKGAAMVTAVAYRLAAQHKARQKILEPLKLSHEQLLEVKERMRIEMEKGLGKETHAEATVKMLPTY
VCSTPDGTEKGDFLALDLGGTNFRVLLVRVRNGMRRGVEMHNKIYSIPVEIMQGTGEELFDHIVHCISDF
LEYMGMKGVSLPLGFTFSFPCQQNNLDEGILLKWTKGFKATGCEGEDVVGLLKEAIHRREEFDLDVVAVV
NDTVGTMMTCGYEDPYCEVGLIVGTGSNACYMEEMRNVELVEGDEGRMCVNMEWGAFGDSGCLDDIRTEF
DLAVDELSLNPGKQRFEKMISGMYLGEIVRNILMDFTKRGLLFRGRISERLKTRGIFETKFLSQIESDCL
ALLQVRSILQHLGLESTCDDSIIVKEVCTVVARRAAQLCGAGMAAVVDKIRENRGLDFLKITVGVDGTLY
KLHPHFSTVMHETVKQLSPKCEVTFLQSEDGSGKGAALITAVACRIREAGQR
>anc106_altAll|avgPP:1.000|lnPP:-0.837|num_ambig:1|num_ambig_gaps:0
LYHMRLSDETLLEVSQRFEKEMEKGLGADTNPTASVKMLPTFVRSTPDGTEDGDFLALDLGGTNFRVLRV
KVSDNGLQKVEMENQIYAIPEELMRGSGVQLFDHIAECLANFMDKLQIKDKKLPLGFTFSFPCHQTKLDE
SILVNWTKGFKCSSVEGRDVVSMLRKSIKKRGDFDIDIVAVVNDTVGTMMTCGYDDHNCEVGLIVGTGTN
ACYMEEMRHIDLVEGDEGRMCINMEWGAFGDDGVLNDIRTEFDREIDMGSLNPGKQLFEKMISGMYMGEL
VRLILVKMAKEGLLFGGRLTPDLLTTGHFETRFVSAIEKEKEGLQKAHEILSKLGLEPSHEDCLATLRIC
QIVSTRSASLCGATLAAVLRRIKDNKGVERLRSTVGVDGSVYKKHPHFARRLHKTVRKLLPDCEIRFIRS
EDGSGKGAAMVTAVAYRLAAQHKARQKILEPLKLSHEQLLEVKERMRIEMEKGLGKETHAEATVKMLPTY
VCSTPDGTEKGDFLALDLGGTNFRVLLVRVRNGMRRGVEMHNKIYSIPVEIMQGTGEELFDHIVHCISDF
LEYMGMKGVSLPLGFTFSFPCQQNNLDEGILLKWTKGFKATGCEGEDVVGLLKEAIHRREEFDLDVVAVV
NDTVGTMMTCGYEDPYCEVGLIVGTGSNACYMEEMRNVELVEGDEGRMCVNMEWGAFGDSGCLDDIRTEF
DLAVDELSLNPGKQRFEKMISGMYLGEIVRNILMDFTKRGLLFRGRISERLKTRGIFETKFLSQIESDCL
ALLQVRSILQHLGLESTCDDSIIVKEVCTVVARRAAQLCGAGMAAVVDKIRENRGLDFLKITVGVDGTLY
KLHPHFSTVMHETVKQLSPKCEVTFLQSEDGSGKGAALITAVACRIREAGQR
>anc107|avgPP:1.000|lnPP:-0.690|num_ambig:1|num_ambig_gaps:0
LYHMRLSDETLLEVSQRFEKEMEKGLGAETNPTASVKMLPTFVRSTPDGTEDGDFLALDLGGTNFRVLRV
KVSDNGLQKVEMENQIYAIPEELMRGSGVQLFDHIAECLANFMDKLQIKDKKLPLGFTFSFPCHQTKLDE
SILVTWTKGFKCSSVEGRDVVSMLRKSIKKRGDFDIDIVAVVNDTVGTMMTCGYDDHNCEVGLIVGTGTN
ACYMEEMRHIDLVEGDEGRMCINMEWGAFGDDGVLNDIRTEFDREIDMGSLNPGKQLFEKMISGMYMGEL
VRLILVKMAKEGLLFGGRLTPDLLTTGHFETRFVSAIEKEKEGLQKAHEILSKLGLEPSHEDCLATLRIC
QIVSTRSASLCGATLAAVLRRIKDNKGVERLRSTVGVDGSVYKKHPHFARRLHKTVRKLLPDCEIRFIRS
EDGSGKGAAMVTAVAYRLAAQHKARQKILEPLKLSHEQLLEVKERMRIEMEKGLGKETHAEATVKMLPTY
VCSTPDGTEKGDFLALDLGGTNFRVLLVRVRNGMRRGVEMHNKIYSIPVEIMQGTGEELFDHIVHCISDF
```

LEYMGMKGVSLPLGFTFSFPCQQNNLDEGILLKWTKGFKATGCEGEDVVGLLKEAIHRREEFDLDVVAVV
NDTVGTMMTCGYEDPYCEVGLIVGTGSNACYMEEMRNVELVEGDEGRMCVNMEWGAFGDSGCLDDIRTEF
DLAVDELSLNPGKQRFEKMISGMYLGEIVRNILMDFTKRGLLFRGRISERLKTRGIFETKFLSQIESDCL
ALLQVRSILQHLGLESTCDDSIIVKEVCTVVARRAAQLCGAGMAAVVDKIRENRGLDFLKVTVGVDGTLY
KLHPHFSTVMHETVKQLSPKCEVTFLQSEDGSGKGAALITAVACRIREAGQR
>anc107_altAll|avgPP:1.000|lnPP:-0.812|num_ambig:1|num_ambig_gaps:0
LYHMRLSDETLLEVSQRFEKEMEKGLGADTNPTASVKMLPTFVRSTPDGTEDGDFLALDLGGTNFRVLRV
KVSDNGLQKVEMENQIYAIPEELMRGSGVQLFDHIAECLANFMDKLQIKDKKLPLGFTFSFPCHQTKLDE
SILVTWTKGFKCSSVEGRDVVSMLRKSIKKRGDFDIDIVAVVNDTVGTMMTCGYDDHNCEVGLIVGTGTN
ACYMEEMRHIDLVEGDEGRMCINMEWGAFGDDGVLNDIRTEFDREIDMGSLNPGKQLFEKMISGMYMGEL
VRLILVKMAKEGLLFGGRLTPDLLTTGHFETRFVSAIEKEKEGLQKAHEILSKLGLEPSHEDCLATLRIC
QIVSTRSASLCGATLAAVLRRIKDNKGVERLRSTVGVDGSVYKKHPHFARRLHKTVRKLLPDCEIRFIRS
EDGSGKGAAMVTAVAYRLAAQHKARQKILEPLKLSHEQLLEVKERMRIEMEKGLGKETHAEATVKMLPTY
VCSTPDGTEKGDFLALDLGGTNFRVLLVRVRNGMRRGVEMHNKIYSIPVEIMQGTGEELFDHIVHCISDF
LEYMGMKGVSLPLGFTFSFPCQQNNLDEGILLKWTKGFKATGCEGEDVVGLLKEAIHRREEFDLDVVAVV
NDTVGTMMTCGYEDPYCEVGLIVGTGSNACYMEEMRNVELVEGDEGRMCVNMEWGAFGDSGCLDDIRTEF
DLAVDELSLNPGKQRFEKMISGMYLGEIVRNILMDFTKRGLLFRGRISERLKTRGIFETKFLSQIESDCL
ALLQVRSILQHLGLESTCDDSIIVKEVCTVVARRAAQLCGAGMAAVVDKIRENRGLDFLKVTVGVDGTLY
KLHPHFSTVMHETVKQLSPKCEVTFLQSEDGSGKGAALITAVACRIREAGQR
>anc108|avgPP:0.999|lnPP:-0.766|num_ambig:2|num_ambig_gaps:0
LYHMRLSDETLLEVSQRFEKEMEKGLGAETNPTASVKMLPTFVRSTPDGTEDGDFLALDLGGTNFRVLRV
KVSDNGLQKVEMENQIYAIPEELMRGSGVQLFDHIAECLANFMDKLQIKDKKLPLGFTFSFPCHQTKLDE
SILVTWTKGFKCSSVEGKDVVSMLRKSIKKRGDFDIDIVAVVNDTVGTMMTCGYDDHNCEVGLIVGTGTN
ACYMEEMRHIDLVEGDEGRMCINMEWGAFGDDGVLNDIRTEFDREIDMGSLNPGKQLFEKMISGMYMGEL
VRLILVKMAKEGLLFGGRLTPDLLTTGHFETRFVSAIEKEKEGLQKAHEILSKLGLEPSHEDCLATLRIC
QIVSTRSASLCGATLAAVLRRIKDNKGVERLRSTVGVDGSVYKKHPHFARRLHKTVRKLLPDCEIRFIRS
EDGSGKGAAMVTAVAYRLAAQHKARQKILEPLKLSHEQLLEVKERMRIEMEKGLGKETHAEATVKMLPTY
VCSTPDGTEKGDFLALDLGGTNFRVLLVRVRNGMRRGVEMHNKIYSIPVEIMQGTGEELFDHIVHCISDF
LEYMGMKGVSLPLGFTFSFPCQQNNLDEGILLKWTKGFKATGCEGEDVVGLLKEAIHRREEFDLDVVAVV
NDTVGTMMTCGYEDPYCEVGLIVGTGSNACYMEEMRNVELVEGDEGRMCVNMEWGAFGDSGCLDDIRTEF
DLAVDELSLNPGKQRFEKMISGMYLGEIVRNILMDFTKRGLLFRGRISERLKTRGIFETKFLSQIESDCL
ALLQVRSILQHLGLESTCDDSIIVKEVCTVVARRAAQLCGAGMAAVVDKIRENRGLDFLKVTVGVDGTLY
KLHPHFSTVMHETVKQLSPKCEVTFLQSEDGSGKGAALITAVACRIREAGQR
>anc108_altAll|avgPP:0.999|lnPP:-2.484|num_ambig:2|num_ambig_gaps:0
LYHMRLSDETLLEVSQRFEKEMEKGLGADTNPTASVKMLPTFVRSTPDGTEDGDFLALDLGGTNFRVLRV
KVSDNGLQKVEMENQIYAIPEELMRGSGVQLFDHIAECLANFMDKLQIKDKKLPLGFTFSFPCHQTKLDE
SILVTWTKGFKCSSVEGKDVVSMLRKSIKKRGDFDIDIVAVVNDTVGTMMTCGYDDHNCEVGLIVGTGTN
ACYMEEMRHIDLVEGDEGRMCINMEWGAFGDDGVLNDIRTEFDREIDMGSLNPGKQLFEKMISGMYMGEL
VRLILVKMAKEGLLFGGRLTPDLLTTGHFETRFVSAIEKEKEGLQKAHEILSKLGLEPSHEDCLATLRIC
QIVSTRSASLCGATLAAVLRRIKDNKGVERLRSTVGVDGSVYKKHPHFARRLHKTVRKLLPDCEIRFIRS
EDGSGKGAAMVTAVAYRLAAQHKARQKILEALKLSHEQLLEVKERMRIEMEKGLGKETHAEATVKMLPTY
VCSTPDGTEKGDFLALDLGGTNFRVLLVRVRNGMRRGVEMHNKIYSIPVEIMQGTGEELFDHIVHCISDF
LEYMGMKGVSLPLGFTFSFPCQQNNLDEGILLKWTKGFKATGCEGEDVVGLLKEAIHRREEFDLDVVAVV
NDTVGTMMTCGYEDPYCEVGLIVGTGSNACYMEEMRNVELVEGDEGRMCVNMEWGAFGDSGCLDDIRTEF
DLAVDELSLNPGKQRFEKMISGMYLGEIVRNILMDFTKRGLLFRGRISERLKTRGIFETKFLSQIESDCL
ALLQVRSILQHLGLESTCDDSIIVKEVCTVVARRAAQLCGAGMAAVVDKIRENRGLDFLKVTVGVDGTLY
KLHPHFSTVMHETVKQLSPKCEVTFLQSEDGSGKGAALITAVACRIREAGQR
>anc109|avgPP:0.999|lnPP:-1.674|num_ambig:4|num_ambig_gaps:0
LYHMRLSDETLLEVSQRFEKEMEKGLGAETNPTASVKMLPTFVRSTPDGTEDGDFLALDLGGTNFRVLRV
KVSDNGLQKVEMENQIYAIPEELMRGSGVQLFDHIAECLANFMDKLQIKDKKLPLGFTFSFPCHQTKLDE
SILVTWTKGFKCSSVEGKDVVSMLRKSIKKRGDFDIDIVAVVNDTVGTMMTCGYDDHNCEVGLIVGTGTN
ACYMEEMRHIDLVEGDEGRMCINMEWGAFGDDGVLNDIRTEFDREIDMGSLNPGKQLFEKMISGMYMGEL

VRLILVKMAKEGLLFGGRLTPDLLTTGHFETRYVSAIEKEKEGLQKAHEILSKLGLEPSHEDCVATLRIC
QIVSTRSASLCGATLAAVLRRIKDNKGVERLRSTVGVDGSVYKKHPHFARRLHKTVRKLLPDCEIRFIRS
EDGSGKGAAMVTAVAYRLAAQHKARQKILEPLKLSHEQLLEVKERMRIEMEKGLGKETHAEATVKMLPTY
VCSTPDGTEKGDFLALDLGGTNFRVLLVRVRNGMRRGVEMHNKIYSIPVEIMQGTGEELFDHIVHCISDF
LEYMGMKGVSLPLGFTFSFPCQQNNLDEGILLKWTKGFKATGCEGEDVVGLLKEAIHRREEFDLDVVAVV
NDTVGTMMTCGYEDPYCEVGLIVGTGSNACYMEEMRNVELVEGDEGRMCVNMEWGAFGDSGCLDDIRTEF
DLAVDELSLNPGKQRFEKMISGMYLGEIVRNILMDFTKRGLLFRGRISERLKTRGIFETKFLSQIESDCL
ALLQVRSILQHLGLESTCDDSIIVKEVCTVVARRAAQLCGAGMAAVVDKIRENRGLDFLKVTVGVDGTLY
KLHPHFSTVMHETVKQLSPKCEVTFLQSEDGSGKGAALITAVACRIREAGQR
>anc109_altAll|avgPP:0.998|lnPP:-4.952|num_ambig:4|num_ambig_gaps:0
LYHMRLSDETLLEVSQRFEKEMEKGLGADTNPTASVKMLPTFVRSTPDGTEDGDFLALDLGGTNFRVLRV
KVSDNGLQKVEMENQIYAIPEELMRGSGVQLFDHIAECLANFMDKLKIKDKKLPLGFTFSFPCHQTKLDE
SILVTWTKGFKCSSVEGKDVVSMLRKSIKKRGDFDIDIVAVVNDTVGTMMTCGYDDHNCEVGLIVGTGTN
ACYMEEMRHIDLVEGDEGRMCINMEWGAFGDDGVLNDIRTEFDREIDMGSLNPGKQLFEKMISGMYMGEL
VRLILVKMAKEGLLFGGRLTPDLLTTGHFETRYVSAIEKEKEGLQKAHEILSKLGLEPSHEDCVATLRIC
QIVSTRSASLCGATLAAVLRRIKDNKGVDRLRSTVGVDGSVYKKHPHFARRLHKTVRKLLPDCEIRFIRS
EDGSGKGAAMVTAVAYRLAAQHKARQKILEALKLSHEQLLEVKERMRIEMEKGLGKETHAEATVKMLPTY
VCSTPDGTEKGDFLALDLGGTNFRVLLVRVRNGMRRGVEMHNKIYSIPVEIMQGTGEELFDHIVHCISDF
LEYMGMKGVSLPLGFTFSFPCQQNNLDEGILLKWTKGFKATGCEGEDVVGLLKEAIHRREEFDLDVVAVV
NDTVGTMMTCGYEDPYCEVGLIVGTGSNACYMEEMRNVELVEGDEGRMCVNMEWGAFGDSGCLDDIRTEF
DLAVDELSLNPGKQRFEKMISGMYLGEIVRNILMDFTKRGLLFRGRISERLKTRGIFETKFLSQIESDCL
ALLQVRSILQHLGLESTCDDSIIVKEVCTVVARRAAQLCGAGMAAVVDKIRENRGLDFLKVTVGVDGTLY
KLHPHFSTVMHETVKQLSPKCEVTFLQSEDGSGKGAALITAVACRIREAGQR
>anc110|avgPP:1.000|lnPP:-0.480|num_ambig:1|num_ambig_gaps:0
LYHMRLSEETLQEVSQRFGKEMEKGLGADTNPTASVKMLPTFVRSTPDGTEDGDFLALDLGGTNFRVLRV
KVSDNGLQKVEMENQIYAIPEELMRGSGVQLFDHIAECLANFMDKLKIKDKKLPLGFTFSFPCHQTKLDE
SILVTWTKGFKCSSVEGKDVVSMLRKSIKKRGDFDIDIVAVVNDTVGTMMTCGYDDHNCEVGLIVGTGTN
ACYMEEMRHIDLVEGDEGRMCINMEWGAFGDDGVLNDIRTEFDREIDMGSLNPGKQLFEKMISGMYMGEL
VRLILVKMAKEGLLFGGRLTPDLLTTGHFETRYVSAIEKEKEGLQKAHEILSKLGLEPSHEDCVATLRIC
QIVSTRSANLCGATLAAVLRRIKDNKGVERLRSTVGVDGSVYKKHPHFARRLHKTVRKLLPDCEIRFVRS
EDGSGKGAAMVTAVAYRLAAQHKARQKILEALKLSHEQLLEVKQRMRIEMEKGLGKETHAEATVKMLPTY
VCSTPDGTEKGDFLALDLGGTNFRVLLVRVRNGMRRGVEMHNKIYSIPVEIMQGTGEELFDHIVHCISDF
LEYMGMKGVSLPLGFTFSFPCQQTNLDEGILLKWTKGFKATGCEGEDVVNLLKEAIHRREEFDLDVVAVV
NDTVGTMMTCGYEDPYCEVGLIVGTGSNACYMEEMRNVELVEGEEGRMCVNMEWGAFGDNGCLDDIRTEF
DLAVDELSLNPGKQRFEKMISGMYLGEIVRNILMDFTKRGLLFRGRISERLKTRGIFETKFLSQIESDCL
ALLQVRSILQHLGLESTCDDSIIVKEVCTVVARRAAQLCGAGMAAVVDKIRENRGLDFLKVTVGVDGTLY
KLHPHFSTVMHETVKQLSPKCEVTFLQSEDGSGKGAALITAVACRIREAGQR
>anc110_altAll|avgPP:0.999|lnPP:-1.381|num_ambig:1|num_ambig_gaps:0
LYHMRLSEETLQEVSQRFGKEMEKGLGADTNPTASVKMLPTFVRSTPDGTEDGDFLALDLGGTNFRVLRV
KVSDNGLQKVEMENQIYAIPEELMRGSGVQLFDHIAECLANFMDKLKIKDKKLPLGFTFSFPCHQTKLDE
SILVTWTKGFKCSSVEGKDVVSMLRKSIKKRGDFDIDIVAVVNDTVGTMMTCGYDDHNCEVGLIVGTGTN
ACYMEEMRHIDLVEGDEGRMCINMEWGAFGDDGVLNDIRTEFDREIDMGSLNPGKQLFEKMISGMYMGEL
VRLILVKMAKEGLLFGGRLTPDLLTTGHFETRYVSAIEKEKEGLQKAHEILSKLGLEPSHEDCVATLRIC
QIVSTRSANLCGATLAAVLRRIKDNKGVDRLRSTVGVDGSVYKKHPHFARRLHKTVRKLLPDCEIRFVRS
EDGSGKGAAMVTAVAYRLAAQHKARQKILEALKLSHEQLLEVKQRMRIEMEKGLGKETHAEATVKMLPTY
VCSTPDGTEKGDFLALDLGGTNFRVLLVRVRNGMRRGVEMHNKIYSIPVEIMQGTGEELFDHIVHCISDF
LEYMGMKGVSLPLGFTFSFPCQQTNLDEGILLKWTKGFKATGCEGEDVVNLLKEAIHRREEFDLDVVAVV
NDTVGTMMTCGYEDPYCEVGLIVGTGSNACYMEEMRNVELVEGEEGRMCVNMEWGAFGDNGCLDDIRTEF
DLAVDELSLNPGKQRFEKMISGMYLGEIVRNILMDFTKRGLLFRGRISERLKTRGIFETKFLSQIESDCL
ALLQVRSILQHLGLESTCDDSIIVKEVCTVVARRAAQLCGAGMAAVVDKIRENRGLDFLKVTVGVDGTLY
KLHPHFSTVMHETVKQLSPKCEVTFLQSEDGSGKGAALITAVACRIREAGQR
>anc111|avgPP:1.000|lnPP:-0.051|num_ambig:0|num_ambig_gaps:0

LYHMRLSEETLQEVSQRFGKEMEKGLGADTNPTASVKMLPTFVRSTPDGTEDGDFLALDLGGTNFRVLRV
KVSDNGLQKVEMENQIYAIPEELMRGSGVQLFDHIAECLANFMEKLKIKDKKLPLGFTFSFPCHQTKLDE
SILVTWTKGFKCSSVEGKDVVSMLRKSIKKRGDFDIDIVAVVNDTVGTMMTCGYDDHNCEVGLIVGTGTN
ACYMEEMRHIDLVEGDEGRMCINMEWGAFGDDGVLNDIRTEFDREIDMGSLNPGKQLFEKMISGMYMGEL
VRLILVKMAKEGLLFGGRLTPDLLTTGHFETRYVSAIEKEKEGLQKAHEILTKLGLEPSHEDCVATHRIC
QIVSTRSANLCGATLAAVLRRIKENKGVDRLRSTVGVDGSVYKKHPHFARRLHKTVRKLLPDCEIRFVRS
EDGSGKGAAMVTAVAYRLAAQHKARQKILEALKLSHEQLLEVKQRMRIEMEKGLGKETHAEATVKMLPTY
VCSTPDGTEKGDFLALDLGGTNFRVLLVRVRNGMRRGVEMHNKIYSIPVEIMQGTGEELFDHIVHCISDF
LEYMGMKGVSLPLGFTFSFPCQQTNLDEGILLKWTKGFKATGCEGEDVVNLLKEAIHRREEFDLDVVAVV
NDTVGTMMTCGYEDPYCEVGLIVGTGSNACYMEEMRNVELVEGEEGRMCVNMEWGAFGDNGCLDDIRTEF
DLAVDELSLNPGKQRFEKMISGMYLGEIVRNILMDFTKRGLLFRGRISERLKTRGIFETKFLSQIESDCL
ALLQVRSILQHLGLESTCDDSIIVKEVCTVVARRAAQLCGAGMAAVVDKIRENRGLDFLKVTVGVDGTLY
KLHPHFSTVMHETVKQLSPKCEVTFLQSEDGSGKGAALITAVACRIREAGQR
>anc111_altAll|avgPP:1.000|lnPP:-0.051|num_ambig:0|num_ambig_gaps:0
LYHMRLSEETLQEVSQRFGKEMEKGLGADTNPTASVKMLPTFVRSTPDGTEDGDFLALDLGGTNFRVLRV
KVSDNGLQKVEMENQIYAIPEELMRGSGVQLFDHIAECLANFMEKLKIKDKKLPLGFTFSFPCHQTKLDE
SILVTWTKGFKCSSVEGKDVVSMLRKSIKKRGDFDIDIVAVVNDTVGTMMTCGYDDHNCEVGLIVGTGTN
ACYMEEMRHIDLVEGDEGRMCINMEWGAFGDDGVLNDIRTEFDREIDMGSLNPGKQLFEKMISGMYMGEL
VRLILVKMAKEGLLFGGRLTPDLLTTGHFETRYVSAIEKEKEGLQKAHEILTKLGLEPSHEDCVATHRIC
QIVSTRSANLCGATLAAVLRRIKENKGVDRLRSTVGVDGSVYKKHPHFARRLHKTVRKLLPDCEIRFVRS
EDGSGKGAAMVTAVAYRLAAQHKARQKILEALKLSHEQLLEVKQRMRIEMEKGLGKETHAEATVKMLPTY
VCSTPDGTEKGDFLALDLGGTNFRVLLVRVRNGMRRGVEMHNKIYSIPVEIMQGTGEELFDHIVHCISDF
LEYMGMKGVSLPLGFTFSFPCQQTNLDEGILLKWTKGFKATGCEGEDVVNLLKEAIHRREEFDLDVVAVV
NDTVGTMMTCGYEDPYCEVGLIVGTGSNACYMEEMRNVELVEGEEGRMCVNMEWGAFGDNGCLDDIRTEF
DLAVDELSLNPGKQRFEKMISGMYLGEIVRNILMDFTKRGLLFRGRISERLKTRGIFETKFLSQIESDCL
ALLQVRSILQHLGLESTCDDSIIVKEVCTVVARRAAQLCGAGMAAVVDKIRENRGLDFLKVTVGVDGTLY
KLHPHFSTVMHETVKQLSPKCEVTFLQSEDGSGKGAALITAVACRIREAGQR
>anc112|avgPP:1.000|lnPP:-0.003|num_ambig:0|num_ambig_gaps:0
LYHMRLSEETLQEVSQRFGKEMEKGLGADTNPTASVKMLPTFVRSTPDGTEDGDFLALDLGGTNFRVLRV
KVSDNGLQKVEMENQIYAIPEELMRGSGVQLFDHIAECLANFMEKLKIKDKKLPLGFTFSFPCHQTKLDE
SILVTWTKGFKCSSVEGKDVVSMLRKSIKKRGDFDIDIVAVVNDTVGTMMTCGYDDHNCEVGLIVGTGTN
ACYMEEMRHIDLVEGDEGRMCINMEWGAFGDDGVLNDIRTEFDREIDMGSLNPGKQLFEKMISGMYMGEL
VRLILVKMAKEGLLFGGKLTPDLLTTGHFETRYVSAIEKEKEGLQKAHEILTKLGLEPSHEDCVATHRIC
QIVSTRSANLCGATLAAVLRRIKENKGVDRLRSTVGVDGSVYKKHPHFARRLHKTVRKLLPDCEIRFVRS
EDGSGKGAAMVTAVAYRLAAQHKARQKILEALKLSHEQLLEVKQRMRIEMEKGLGKETHAEATVKMLPTY
VCSTPDGTEKGDFLALDLGGTNFRVLLVRVRNGMRRGVEMHNKIYSIPVEIMQGTGEELFDHIVHCISDF
LEYMGMKGVSLPLGFTFSFPCQQTNLDEGILLKWTKGFKATGCEGEDVVNLLKEAIHRREEFDLDVVAVV
NDTVGTMMTCGYEDPYCEVGLIVGTGSNACYMEEMRNVELVEGEEGRMCVNMEWGAFGDNGCLDDIRTEF
DLAVDELSLNPGKQRFEKMISGMYLGEIVRNILMDFTKRGLLFRGRISERLKTRGIFETKFLSQIESDCL
ALLQVRSILQHLGLETTCDDSIIVKEVCTVVARRAAQLCGAGMAAVVDKIRENRGLDFLKVTVGVDGTLY
KLHPHFSTVMHETVKQLSPKCEVTFLQSEDGSGKGAALITAVACRIREAGQR
>anc112_altAll|avgPP:1.000|lnPP:-0.003|num_ambig:0|num_ambig_gaps:0
LYHMRLSEETLQEVSQRFGKEMEKGLGADTNPTASVKMLPTFVRSTPDGTEDGDFLALDLGGTNFRVLRV
KVSDNGLQKVEMENQIYAIPEELMRGSGVQLFDHIAECLANFMEKLKIKDKKLPLGFTFSFPCHQTKLDE
SILVTWTKGFKCSSVEGKDVVSMLRKSIKKRGDFDIDIVAVVNDTVGTMMTCGYDDHNCEVGLIVGTGTN
ACYMEEMRHIDLVEGDEGRMCINMEWGAFGDDGVLNDIRTEFDREIDMGSLNPGKQLFEKMISGMYMGEL
VRLILVKMAKEGLLFGGKLTPDLLTTGHFETRYVSAIEKEKEGLQKAHEILTKLGLEPSHEDCVATHRIC
QIVSTRSANLCGATLAAVLRRIKENKGVDRLRSTVGVDGSVYKKHPHFARRLHKTVRKLLPDCEIRFVRS
EDGSGKGAAMVTAVAYRLAAQHKARQKILEALKLSHEQLLEVKQRMRIEMEKGLGKETHAEATVKMLPTY
VCSTPDGTEKGDFLALDLGGTNFRVLLVRVRNGMRRGVEMHNKIYSIPVEIMQGTGEELFDHIVHCISDF
LEYMGMKGVSLPLGFTFSFPCQQTNLDEGILLKWTKGFKATGCEGEDVVNLLKEAIHRREEFDLDVVAVV
NDTVGTMMTCGYEDPYCEVGLIVGTGSNACYMEEMRNVELVEGEEGRMCVNMEWGAFGDNGCLDDIRTEF

DLAVDELSLNPGKQRFEKMISGMYLGEIVRNILMDFTKRGLLFRGRISERLKTRGIFETKFLSQIESDCL
ALLQVRSILQHLGLETTCDDSIIVKEVCTVVARRAAQLCGAGMAAVVDKIRENRGLDFLKVTVGVDGTLY
KLHPHFSTVMHETVKQLSPKCEVTFLQSEDGSGKGAALITAVACRIREAGQR
>anc113|avgPP:1.000|lnPP:-0.018|num_ambig:0|num_ambig_gaps:0
LYHMRLSEETLQEVSQRFGKEMEKGLGADTNPTASVKMLPTFVRSTPDGTEDGDFLALDLGGTNFRVLRV
KVSDNGLQKVEMENQIYAIPEELMRGSGVQLFDHIAECLANFMEKLKIKDKKLPLGFTFSFPCHQTKLDE
SILVTWTKGFKCSSVEGKDVVSMLRKSIKKRGDFDIDIVAVVNDTVGTMMTCGYDDHNCEVGLIVGTGTN
ACYMEEMRHIDLVEGDEGRMCINMEWGAFGDDGVLNDIRTEFDREIDMGSLNPGKQLFEKMISGMYMGEL
VRLILVKMAKEGLLFGGKLTPDLLTTGHFETRYVSAIEKEKEGLQKAHEILTKLGLEPSHEDCVATHRIC
QIVSTRSANLCGATLAAVLRRIKENKGVDRLRSTVGVDGSVYKKHPHFARRLHKTVRKLLPDCEIRFVRS
EDGSGKGAAMVTAVAYRLAAQHKARQKILEALKLSHEQLLEVKQRMRIEMEKGLGKETHAEATVKMLPTY
VCSTPDGTEKGDFLALDLGGTNFRVLLVRVRNGMRRGVEMHNKIYSIPVEIMQGTGEELFDHIVHCISDF
LEYMGMKGVSLPLGFTFSFPCQQTNLDEGILLKWTKGFKATGCEGEDVVNLLKEAIHRREEFDLDVVAVV
NDTVGTMMTCGYEDPYCEVGLIVGTGSNACYMEEMRNVELVEGEEGRMCVNMEWGAFGDNGCLDDIRTEF
DLAVDELSLNPGKQRFEKMISGMYLGEIVRNILMDFTKRGLLFRGRISERLKTRGIFETKFLSQIESDCL
ALLQVRSILQHLGLESTCDDSIIVKEVCTVVARRAAQLCGAGMAAVVDKIRENRGLDFLKVTVGVDGTLY
KLHPHFSTVMHETVKQLSPKCEVTFLQSEDGSGKGAALITAVACRIREAGQR
>anc113_altAll|avgPP:1.000|lnPP:-0.018|num_ambig:0|num_ambig_gaps:0
LYHMRLSEETLQEVSQRFGKEMEKGLGADTNPTASVKMLPTFVRSTPDGTEDGDFLALDLGGTNFRVLRV
KVSDNGLQKVEMENQIYAIPEELMRGSGVQLFDHIAECLANFMEKLKIKDKKLPLGFTFSFPCHQTKLDE
SILVTWTKGFKCSSVEGKDVVSMLRKSIKKRGDFDIDIVAVVNDTVGTMMTCGYDDHNCEVGLIVGTGTN
ACYMEEMRHIDLVEGDEGRMCINMEWGAFGDDGVLNDIRTEFDREIDMGSLNPGKQLFEKMISGMYMGEL
VRLILVKMAKEGLLFGGKLTPDLLTTGHFETRYVSAIEKEKEGLQKAHEILTKLGLEPSHEDCVATHRIC
QIVSTRSANLCGATLAAVLRRIKENKGVDRLRSTVGVDGSVYKKHPHFARRLHKTVRKLLPDCEIRFVRS
EDGSGKGAAMVTAVAYRLAAQHKARQKILEALKLSHEQLLEVKQRMRIEMEKGLGKETHAEATVKMLPTY
VCSTPDGTEKGDFLALDLGGTNFRVLLVRVRNGMRRGVEMHNKIYSIPVEIMQGTGEELFDHIVHCISDF
LEYMGMKGVSLPLGFTFSFPCQQTNLDEGILLKWTKGFKATGCEGEDVVNLLKEAIHRREEFDLDVVAVV
NDTVGTMMTCGYEDPYCEVGLIVGTGSNACYMEEMRNVELVEGEEGRMCVNMEWGAFGDNGCLDDIRTEF
DLAVDELSLNPGKQRFEKMISGMYLGEIVRNILMDFTKRGLLFRGRISERLKTRGIFETKFLSQIESDCL
ALLQVRSILQHLGLESTCDDSIIVKEVCTVVARRAAQLCGAGMAAVVDKIRENRGLDFLKVTVGVDGTLY
KLHPHFSTVMHETVKQLSPKCEVTFLQSEDGSGKGAALITAVACRIREAGQR
>anc114|avgPP:1.000|lnPP:-0.050|num_ambig:0|num_ambig_gaps:0
LYHMRLSEETLQEVSQRFGKEMEKGLGADTNPTASVKMLPTFVRSTPDGTEDGDFLALDLGGTNFRVLRV
KVSDNGLQKVEMENQIYAIPEELMRGSGVQLFDHIAECLANFMEKLKIKDKKLPLGFTFSFPCHQTKLDE
SILVTWTKGFKCSSVEGKDVVSMLRKSIKKRGDFDIDIVAVVNDTVGTMMTCGYDDHNCEVGLIVGTGTN
ACYMEEMRHIDLVEGDEGRMCINMEWGAFGDDGVLNDIRTEFDREIDMGSLNPGKQLFEKMISGMYMGEL
VRLILVKMAKEGLLFGGRLTPDLLTTGHFETRYVSAIEKEKEGLQKAHEILTKLGLEPSHEDCVATHRIC
QIVSTRSANLCGATLAAVLRRIKENKGVDRLRSTVGVDGSVYKKHPHFARRLHKTVRKLLPDCEIRFVRS
EDGSGKGAAMVTAVAYRLAAQHKARQKILEALKLSHEQLLEVKQRMRIEMEKGLGKETHAEATVKMLPTY
VCSTPDGTEKGDFLALDLGGTNFRVLLVRVRNGMRRGVEMHNKIYSIPVEIMQGTGEELFDHIVHCISDF
LEYMGMKGVSLPLGFTFSFPCQQTNLDEGILLKWTKGFKATGCEGEDVVNLLKEAIHRREEFDLDVVAVV
NDTVGTMMTCGYEDPYCEVGLIVGTGSNACYMEEMRNVELVEGEEGRMCVNMEWGAFGDNGCLDDIRTEF
DLAVDELSLNPGKQRFEKMISGMYLGEIVRNILMDFTKRGLLFRGRISERLKTRGIFETKFLSQIESDCL
ALLQVRSILQHLGLESTCDDSIIVKEVCTVVARRAAQLCGAGMAAVVDKIRENRGLDFLKVTVGVDGTLY
KLHPHFSTVMHETVKQLSPKCEVTFLQSEDGSGKGAALITAVACRIREAGQR
>anc114_altAll|avgPP:1.000|lnPP:-0.050|num_ambig:0|num_ambig_gaps:0
LYHMRLSEETLQEVSQRFGKEMEKGLGADTNPTASVKMLPTFVRSTPDGTEDGDFLALDLGGTNFRVLRV
KVSDNGLQKVEMENQIYAIPEELMRGSGVQLFDHIAECLANFMEKLKIKDKKLPLGFTFSFPCHQTKLDE
SILVTWTKGFKCSSVEGKDVVSMLRKSIKKRGDFDIDIVAVVNDTVGTMMTCGYDDHNCEVGLIVGTGTN
ACYMEEMRHIDLVEGDEGRMCINMEWGAFGDDGVLNDIRTEFDREIDMGSLNPGKQLFEKMISGMYMGEL
VRLILVKMAKEGLLFGGRLTPDLLTTGHFETRYVSAIEKEKEGLQKAHEILTKLGLEPSHEDCVATHRIC
QIVSTRSANLCGATLAAVLRRIKENKGVDRLRSTVGVDGSVYKKHPHFARRLHKTVRKLLPDCEIRFVRS

EDGSGKGAAMVTAVAYRLAAQHKARQKILEALKLSHEQLLEVKQRMRIEMEKGLGKETHAEATVKMLPTY
VCSTPDGTEKGDFLALDLGGTNFRVLLVRVRNGMRRGVEMHNKIYSIPVEIMQGTGEELFDHIVHCISDF
LEYMGMKGVSLPLGFTFSFPCQQTNLDEGILLKWTKGFKATGCEGEDVVNLLKEAIHRREEFDLDVVAVV
NDTVGTMMTCGYEDPYCEVGLIVGTGSNACYMEEMRNVELVEGEEGRMCVNMEWGAFGDNGCLDDIRTEF
DLAVDELSLNPGKQRFEKMISGMYLGEIVRNILMDFTKRGLLFRGRISERLKTRGIFETKFLSQIESDCL
ALLQVRSILQHLGLESTCDDSIIVKEVCTVVARRAAQLCGAGMAAVVDKIRENRGLDFLKVTVGVDGTLY
KLHPHFSTVMHETVKQLSPKCEVTFLQSEDGSGKGAALITAVACRIREAGQR
>anc115|avgPP:1.000|lnPP:-0.050|num_ambig:0|num_ambig_gaps:0
LYHMRLSEETLQEVSQRFGKEMEKGLGADTNPTASVKMLPTFVRSTPDGTEDGDFLALDLGGTNFRVLRV
KVSDNGLQKVEMENQIYAIPEELMRGSGVQLFDHIAECLANFMEKLKIKDKKLPLGFTFSFPCHQTKLDE
SILVTWTKGFKCSSVEGKDVVSMLRKSIKKRGDFDIDIVAVVNDTVGTMMTCGYDDHNCEVGLIVGTGTN
ACYMEEMRHIDLVEGDEGRMCINMEWGAFGDDGVLNDIRTEFDREIDMGSLNPGKQLFEKMISGMYMGEL
VRLILVKMAKEGLLFGGRLTPDLLTTGHFETRYVSAIEKEKEGLQKAHEILTKLGLEPSHEDCVATHRIC
QIVSTRSANLCGATLAAVLRRIKENKGVDRLRSTVGVDGSVYKKHPHFARRLHKTVRKLLPDCEIRFVRS
EDGSGKGAAMVTAVAYRLAAQHKARQKILEALKLSHEQLLEVKQRMRIEMEKGLGKETHAEATVKMLPTY
VCSTPDGTEKGDFLALDLGGTNFRVLLVRVRNGMRRGVEMHNKIYSIPVEIMQGTGEELFDHIVHCISDF
LEYMGMKGVSLPLGFTFSFPCQQTNLDEGILLKWTKGFKATGCEGEDVVNLLKEAIHRREEFDLDVVAVV
NDTVGTMMTCGYEDPYCEVGLIVGTGSNACYMEEMRNVELVEGEEGRMCVNMEWGAFGDNGCLDDIRTEF
DLAVDELSLNPGKQRFEKMISGMYLGEIVRNILMDFTKRGLLFRGRISERLKTRGIFETKFLSQIESDCL
ALLQVRSILQHLGLESTCDDSIIVKEVCTVVARRAAQLCGAGMAAVVDKIRENRGLDFLKVTVGVDGTLY
KLHPHFSTVMHETVKQLSPKCEVTFLQSEDGSGKGAALITAVACRIREAGQR
>anc115_altAll|avgPP:1.000|lnPP:-0.050|num_ambig:0|num_ambig_gaps:0
LYHMRLSEETLQEVSQRFGKEMEKGLGADTNPTASVKMLPTFVRSTPDGTEDGDFLALDLGGTNFRVLRV
KVSDNGLQKVEMENQIYAIPEELMRGSGVQLFDHIAECLANFMEKLKIKDKKLPLGFTFSFPCHQTKLDE
SILVTWTKGFKCSSVEGKDVVSMLRKSIKKRGDFDIDIVAVVNDTVGTMMTCGYDDHNCEVGLIVGTGTN
ACYMEEMRHIDLVEGDEGRMCINMEWGAFGDDGVLNDIRTEFDREIDMGSLNPGKQLFEKMISGMYMGEL
VRLILVKMAKEGLLFGGRLTPDLLTTGHFETRYVSAIEKEKEGLQKAHEILTKLGLEPSHEDCVATHRIC
QIVSTRSANLCGATLAAVLRRIKENKGVDRLRSTVGVDGSVYKKHPHFARRLHKTVRKLLPDCEIRFVRS
EDGSGKGAAMVTAVAYRLAAQHKARQKILEALKLSHEQLLEVKQRMRIEMEKGLGKETHAEATVKMLPTY
VCSTPDGTEKGDFLALDLGGTNFRVLLVRVRNGMRRGVEMHNKIYSIPVEIMQGTGEELFDHIVHCISDF
LEYMGMKGVSLPLGFTFSFPCQQTNLDEGILLKWTKGFKATGCEGEDVVNLLKEAIHRREEFDLDVVAVV
NDTVGTMMTCGYEDPYCEVGLIVGTGSNACYMEEMRNVELVEGEEGRMCVNMEWGAFGDNGCLDDIRTEF
DLAVDELSLNPGKQRFEKMISGMYLGEIVRNILMDFTKRGLLFRGRISERLKTRGIFETKFLSQIESDCL
ALLQVRSILQHLGLESTCDDSIIVKEVCTVVARRAAQLCGAGMAAVVDKIRENRGLDFLKVTVGVDGTLY
KLHPHFSTVMHETVKQLSPKCEVTFLQSEDGSGKGAALITAVACRIREAGQR
>anc116|avgPP:0.999|lnPP:-1.998|num_ambig:2|num_ambig_gaps:0
LYHMRLSEETLQEVSQRFRKEMEKGLGADTNPTASVKMLPTFVRSTPDGTEDGDFLALDLGGTNFRVLRV
KVSDNGLQKVEMENQIYAIPEELMRGSGVQLFDHIAECLANFMEKLNIKDKKLPLGFTFSFPCHQTKLDE
SILVTWTKGFKCSGVEGKDVVSMLRKSIKKRGDFDIDIVAVVNDTVGTMMTCGYDDHNCEVGLIVGTGTN
ACYMEEMRHIDLVEGDEGRMCINMEWGAFGDDGVLNDIRTEFDREIDMGSLNPGKQLFEKMISGMYMGEL
VRLILVKMAKEGLLFGGRLTPDLLTTGHFETRYVSAIEKEKEGLQKAHEILTKLGLEPSHEDCVATHRIC
QIVSTRSANLCGATLAAVLRRIKENKGVDRLRSTVGVDGSVYKKHPHFARRLHKTVRKLLPDCDIRFVRS
EDGSGKGAAMVTAVAYRLAAQHKARQKTLEALKLSHEQLLEVKQRMRIEMEKGLGKETHAEATVKMLPTY
VCSTPDGTEKGDFLALDLGGTNFRVLLVRVRNGMRRGVEMHNKIYSIPVEIMQGTGEELFDHIVHCISDF
LEYMGMKGVSLPLGFTFSFPCQQTNLDEGILLKWTKGFKATGCEGEDVVNLLKEAIHRREEFDLDVVAVV
NDTVGTMMTCGYEDPYCEVGLIVGTGSNACYMEEMRNVELVEGEEGRMCVNMEWGAFGDNGCLDDIRTEF
DLAVDELSLNPGKQRFEKMISGMYLGEIVRNILIDFTKRGLLFRGRISERLKTRGIFETKFLSQIESDCL
ALLQVRSILQHLGLESTCDDSIIVKEVCTVVARRAAQLCGAGMAAVVDKIRENRGLDFLKVTVGVDGTLY
KLHPHFSTVMHETVKQLSPKCEVTFLQSEDGSGKGAALITAVACRIREAGQR
>anc116_altAll|avgPP:0.999|lnPP:-2.440|num_ambig:2|num_ambig_gaps:0
LYHMRLSEETLQEVSQRFRKEMEKGLGADTNPTASVKMLPTFVRSTPDGTEDGDFLALDLGGTNFRVLRV
KVSDNGLQKVEMENQIYAIPEELMRGSGVQLFDHIAECLANFMEKLKIKDKKLPLGFTFSFPCHQTKLDE

SILVTWTKGFKCSGVEGKDVVSMLRKSIKKRGDFDIDIVAVVNDTVGTMMTCGYDDHNCEVGLIVGTGTN
ACYMEEMRHIDLVEGDEGRMCINMEWGAFGDDGVLNDIRTEFDREIDMGSLNPGKQLFEKMISGMYMGEL
VRLILVKMAKEGLLFGGRLTPDLLTTGHFETRYVSAIEKEKEGLQKAHEILTKLGLEPSHEDCVATHRIC
QIVSTRSANLCGATLAAVLRRIKENKGVDRLRSTVGVDGSVYKKHPHFARRLHKTVRKLLPDCDIRFVRS
EDGSGKGAAMVTAVAYRLAAQHKARQKTLEALKLSHEQLLEVKQRMRIEMEKGLGKETHAEATVKMLPTY
VCSTPDGTEKGDFLALDLGGTNFRVLLVRVRNGMRRGVEMHNKIYSIPLEIMQGTGEELFDHIVHCISDF
LEYMGMKGVSLPLGFTFSFPCQQTNLDEGILLKWTKGFKATGCEGEDVVNLLKEAIHRREEFDLDVVAVV
NDTVGTMMTCGYEDPYCEVGLIVGTGSNACYMEEMRNVELVEGEEGRMCVNMEWGAFGDNGCLDDIRTEF
DLAVDELSLNPGKQRFEKMISGMYLGEIVRNILIDFTKRGLLFRGRISERLKTRGIFETKFLSQIESDCL
ALLQVRSILQHLGLESTCDDSIIVKEVCTVVARRAAQLCGAGMAAVVDKIRENRGLDFLKVTVGVDGTLY
KLHPHFSTVMHETVKQLSPKCEVTFLQSEDGSGKGAALITAVACRIREAGQR
>anc117|avgPP:0.998|lnPP:-2.605|num_ambig:1|num_ambig_gaps:0
LYHMRLSDETLQEVSERFRKEMEKGLGADTNPTASVKMLPTFVRSTPDGTEDGDFLALDLGGTNFRVLRV
KVSDNGLQKVEMENQIYAIPEDLMRGSGVQLFDHIAECLANFMEKLNIKDKKLPLGFTFSFPCHQTKLDE
SILVTWTKGFKSSGVEGKDVVSMLRKAIKKRGDFDIDIVAVVNDTVGTMMTCGYDDHNCEVGLIVGTGTN
ACYMEEMRHIDLVEGDEGRMCINMEWGAFGDDGVLNDIRTEFDREIDMGSLNPGKQLFEKMISGMYMGEL
VRLILVKMAKEELLFGGRLTPDLLTTGHFETRYVSAIEKEKEGLQKAHEILTKLGLEPSHEDCVATHRIC
QIVSTRSANLCGATLAAVLRRIKENKGVDRLRSTVGVDGSVYKKHPHFARRLHKTVRKLLPDCDIRFVRS
EDGSGKGAAMVTAVAYRLAAQHKARQKTLEALKLSHEQLLEVKKRMRIEMERGLGKETHAEATVKMLPTY
VCSTPDGTEKGDFLALDLGGTNFRVLLVRVRNGMRRGVEMHNKIYSIPQEIMQGTGEELFDHIVHCISDF
LEYMGMKGVSLPLGFTFSFPCQQNNLDEGILLKWTKGFKATGCEGEDVVNLLKEAIHRREEFDLDVVAVV
NDTVGTMMTCGYEDPYCEVGLIVGTGSNACYMEEMRNVELVEGEEGRMCVNMEWGAFGDNGCLDDFRTEF
DVAVDELSLNPGKQRFEKMISGMYLGEIVRNILIDFTKRGLLFRGRISERLKTRGIFETKFLSQIESDCL
ALLQVRSILQHLGLESTCDDSIIVKEVCTVVARRAAQLCGAGMAAVVDKIRENRGLDFLKVTVGVDGTLY
KLHPHFSTIMHETVKQLSPKCEVTFLQSEDGSGKGAALITAVACRIREAGQR
>anc117_altAll|avgPP:0.998|lnPP:-3.076|num_ambig:1|num_ambig_gaps:0
LYHMRLSDETLQEVSERFRKEMEKGLGADTNPTASVKMLPTFVRSTPDGTEDGDFLALDLGGTNFRVLRV
KVSDNGLQKVEMENQIYAIPEDLMRGSGVQLFDHIAECLANFMEKLNIKDKKLPLGFTFSFPCHQTKLDE
SILVTWTKGFKSSGVEGKDVVSMLRKAIKKRGDFDIDIVAVVNDTVGTMMTCGYDDHNCEVGLIVGTGTN
ACYMEEMRHIDLVEGDEGRMCINMEWGAFGDDGVLNDIRTEFDREIDMGSLNPGKQLFEKMISGMYMGEL
VRLILVKMAKEELLFGGRLTPDLLTTGHFETRYVSAIEKEKEGLQKAHEILTKLGLEPSHEDCVATHRIC
QIVSTRSANLCGATLAAVLRRIKENKGVDRLRSTVGVDGSVYKKHPHFARRLHKTVRKLLPDCDIRFVRS
EDGSGKGAAMVTAVAYRLAAQHKARQKTLEALKLSHEQLLEVKKRMRIEMERGLGKETHAEATVKMLPTY
VCSTPDGTEKGDFLALDLGGTNFRVLLVRVRNGMRRGVEMHNKIYSIPQEIMQGTGEELFDHIVHCISDF
LEYMGMKGVSLPLGFTFSFPCQQNNLDEGILLKWTKGFKATGCEGEDVVNLLKEAIHRREEFDLDVVAVV
NDTVGTMMTCGYEDPYCEVGLIVGTGSNACYMEEMRNVELVEGEEGRMCVNMEWGAFGDNGCLDDFRTEF
DVAVDELSLNPGKQRFEKMISGMYLGEIVRNILIDFTKRGLLFRGRISERLKTRGIFETKFLSQIESDCL
ALLQVRSILQHLGLESTCDDSIIVKEVCTVVARRAAQLCGAGMAAVVDKIRENRGLDFLKVTVGVDGTLY
KLHPHFSTVMHETVKQLSPKCEVTFLQSEDGSGKGAALITAVACRIREAGQR
>anc118|avgPP:0.998|lnPP:-2.605|num_ambig:1|num_ambig_gaps:0
LYHMRLSDETLQEVSERFRKEMEKGLGADTNPTASVKMLPTFVRSTPDGTEDGDFLALDLGGTNFRVLRV
KVSDNGLQKVEMENQIYAIPEDLMRGSGVQLFDHIAECLANFMEKLNIKDKKLPLGFTFSFPCHQTKLDE
SILVTWTKGFKSSGVEGKDVVSMLRKAIKKRGDFDIDIVAVVNDTVGTMMTCGYDDHNCEVGLIVGTGTN
ACYMEEMRHIDLVEGDEGRMCINMEWGAFGDDGVLNDIRTEFDREIDMGSLNPGKQLFEKMISGMYMGEL
VRLILVKMAKEELLFGGRLTPDLLTTGHFETRYVSAIEKEKEGLQKAHEILTKLGLEPSHEDCVATHRIC
QIVSTRSANLCGATLAAVLRRIKENKGVDRLRSTVGVDGSVYKKHPHFARRLHKTVRKLLPDCDIRFVRS
EDGSGKGAAMVTAVAYRLAAQHKARQKTLEALKLSHEQLLEVKKRMRIEMERGLGKETHAEATVKMLPTY
VCSTPDGTEKGDFLALDLGGTNFRVLLVRVRNGMRRGVEMHNKIYSIPQEIMQGTGEELFDHIVHCISDF
LEYMGMKGVSLPLGFTFSFPCQQNNLDEGILLKWTKGFKATGCEGEDVVNLLKEAIHRREEFDLDVVAVV
NDTVGTMMTCGYEDPYCEVGLIVGTGSNACYMEEMRNVELVEGEEGRMCVNMEWGAFGDNGCLDDFRTEF
DVAVDELSLNPGKQRFEKMISGMYLGEIVRNILIDFTKRGLLFRGRISERLKTRGIFETKFLSQIESDCL
ALLQVRSILQHLGLESTCDDSIIVKEVCTVVARRAAQLCGAGMAAVVDKIRENRGLDFLKVTVGVDGTLY

```
KLHPHFSTIMHETVKQLSPKCEVTFLQSEDGSGKGAALITAVACRIREAGQR
>anc118_altAll|avgPP:0.998|lnPP:-3.076|num_ambig:1|num_ambig_gaps:0
LYHMRLSDETLQEVSERFRKEMEKGLGADTNPTASVKMLPTFVRSTPDGTEDGDFLALDLGGTNFRVLRV
KVSDNGLQKVEMENQIYAIPEDLMRGSGVQLFDHIAECLANFMEKLNIKDKKLPLGFTFSFPCHQTKLDE
SILVTWTKGFKSSGVEGKDVVSMLRKAIKKRGDFDIDIVAVVNDTVGTMMTCGYDDHNCEVGLIVGTGTN
ACYMEEMRHIDLVEGDEGRMCINMEWGAFGDDGVLNDIRTEFDREIDMGSLNPGKQLFEKMISGMYMGEL
VRLILVKMAKEELLFGGRLTPDLLTTGHFETRYVSAIEKEKEGLQKAHEILTKLGLEPSHEDCVATHRIC
QIVSTRSANLCGATLAAVLRRIKENKGVDRLRSTVGVDGSVYKKHPHFARRLHKTVRKLLPDCDIRFVRS
EDGSGKGAAMVTAVAYRLAAQHKARQKTLEALKLSHEQLLEVKKRMRIEMERGLGKETHAEATVKMLPTY
VCSTPDGTEKGDFLALDLGGTNFRVLLVRVRNGMRRGVEMHNKIYSIPQEIMQGTGEELFDHIVHCISDF
LEYMGMKGVSLPLGFTFSFPCQQNNLDEGILLKWTKGFKATGCEGEDVVNLLKEAIHRREEFDLDVVAVV
NDTVGTMMTCGYEDPYCEVGLIVGTGSNACYMEEMRNVELVEGEEGRMCVNMEWGAFGDNGCLDDFRTEF
DVAVDELSLNPGKQRFEKMISGMYLGEIVRNILIDFTKRGLLFRGRISERLKTRGIFETKFLSQIESDCL
ALLQVRSILQHLGLESTCDDSIIVKEVCTVVARRAAQLCGAGMAAVVDKIRENRGLDFLKVTVGVDGTLY
KLHPHFSTVMHETVKQLSPKCEVTFLQSEDGSGKGAALITAVACRIREAGQR
>anc119|avgPP:0.988|lnPP:-17.872|num_ambig:15|num_ambig_gaps:0
LYHMRLSDDTLQEISNRFRKEMEKGLGADTNPTASVKMLPTFVRSTPDGTEEGDFLALDLGGTNFRVLRV
KVADNGSKKVEMENQIYAIPEDLMRGSGVQLFDHIAECLANFMEKLQMKDKKLPLGFTFSFPCYQTKLDE
SILVTWTKGFKSSGVEGKDVVSLLRKAIKKRGDFDIDVVAVVNDTVGTMMTCGYDDHNCEVGLIVGTGSN
ACYMEEMRHIELVEGDEGRMCINMEWGAFGDDGALNDIRTEFDRDIDMGSLNPGKQLFEKMISGMYMGEL
VRLILVKMAKEELLFGGRLTPDLLTTGCFETRYVSAIEKEKEGLQKAHEILTKLGLEPSHEDCVATRRVC
EIVSTRSANLCGAALAAVLRRIKENKGVDRLRSTVGVDGTVYKKHPHFARRLHKTVRRLLPDCDIRFVRS
EDGSGKGAAMVTAVAYRLAAQHKARQKILEPLRLSHEQLLEVKKRMRQEMARGLARETHAEATVKMLPTY
VCSTPDGTEKGDFLALDLGGTNFRVLLVRIRNGMRRSVEMHNKIYSIPQDIMQGTGEELFDHIVHCISDF
LEYMGMKGVSLPLGFTFSFPCHQKNLDEGILLKWTKGFSATGCEGEDVVGLLKDAIHRREEFDLDVVALV
NDTVGTMMTCGYEDPLCEVGLIVGTGSNACYMEEMRNVELVEGDQGRMCVNMEWGAFGDNGCLDDFRTEF
DAAVDENSLNPGKQRFEKMISGMYLGEIVRNILIDFTKRGLLFRGRISERLKTRGIFETKFLSQIESDRL
ALLQVRSILQHLGLESTCDDSIIVKEVCTVIARRAAQLCGAGIAAVVDKIRENRGLDFLKVTVGVDGTLY
KLHPHFSTVMHETVKQLSPKCEVTFLQSEDGSGKGAALITAVACRIREAGQH
>anc119_altAll|avgPP:0.985|lnPP:-27.280|num_ambig:15|num_ambig_gaps:0
LYHMRLSDDTLQEISNRFRKEMEKGLGADTNPTASVKMLPTFVRSTPDGTEEGDFLALDLGGTNFRVLRV
KVADNGSKKVEMENQIYAIPEDLMRSSGVQLFDHIAECLANFMEQLQMKDKKLPLGFTFSFPCYQTKLDE
SVLVTWTKGFKSSGVEGKDVVSLLRQAIKKRGDFDIDVVAVVNDTVGTMMTCGYDDHNCEVGLIVGTGSN
ACYMEEMRHIELVEGDEGRMCINMEWGAFGDDGALNDIRTEFDREIDMGSLNPGKQLFEKMISGMYMGEL
VRLILVKMAKEELLFGGKLTPDLLTTGCFETRYVSAIEKEKEGLLKAHEILTKLGLEPSHEDCVATRRVC
EIVSTRSANLCGAALAAVLRRIKENKGVDRLRSTVGVDGTVYKKHPHFARRLHKTVRRLLPDCDIRFVRS
EDGSGKGAAMVTAVAYRLAAQHKARQEILEPLRLSHEQLLEVKKRMRQEMARGLARETHAQATVKMLPTY
VCATPDGTEKGDFLALDLGGTNFRVLLVRVRNGMRRSVEMHNKIYSIPQDIMQGTGEELFDHIVHCISDF
LEYMGMKGVSLPLGFTFSFPCHQKNLDEGILLKWTKGFSATGCEGEDVVGLLKEAIHRREEFDLDVVALV
NDTVGTMMTCGYEDPLCEVGLIVGTGSNACYMEEMRNVELVEGDQGRMCVNMEWGAFGDNGCLDDFRTEF
DAAVDENSLNPGKQRFEKMISGMYLGEIVRNILIDFTKRGLLFRGRISERLKTRGIFETKFLSQIESDCL
ALLQVRSILQHLGLESTCDDSIIVKEVCTVIARRAAQLCGAGMAAVVDKIRENRGLDFLKVTVGVDGTLY
KLHPHFSTVMDETVKQLSPKCEVTFLQSEDGSGKGAALITAVACRIREAGQH
>anc120|avgPP:0.993|lnPP:-9.911|num_ambig:10|num_ambig_gaps:0
LYHMRLSDETLQEVSNRFRKEMEKGLGADTNPTASVKMLPTFVRSTPDGTEEGDFLALDLGGTNFRVLRV
KVSDNGSKKVEMENQIYAIPEDLMRGSGVQLFDHIAECLANFMEKLQMKDKKLPLGFTFSFPCYQTKLDE
SILVTWTKGFKSSGVEGKDVVSLLRKAIKKRGDFDIDIVAVVNDTVGTMMTCGYDDHNCEVGLIVGTGSN
ACYMEEMRHIDLVEGDEGRMCINMEWGAFGDDGVLNDIRTEFDRDIDMGSLNPGKQLFEKMISGMYMGEL
VRLILVKMAKEELLFGGRLTPDLLTTGHFETRYVSAIEKEKEGLQKAHEILTKLGLEPSHEDCVATRRIC
EIVSTRSANLCGAALAAVLRRIKENKGVDRLRSTVGVDGTVYKKHPHFARRLHKTVRRLLPDCDIRFVRS
EDGSGKGAAMVTAVAYRLAAQHKARQKILEPLRLSHEQLLEVKKRMREEMERGLARETHAEATVKMLPTY
VCSTPDGTEKGDFLALDLGGTNFRVLLVRIRNGMRRSVEMHNKIYSIPQDIMQGTGEELFDHIVHCISDF
```

LEYMGMKGVSLPLGFTFSFPCHQNNLDEGILLKWTKGFKATGCEGEDVVGLLKDAIHRREEFDLDVVAVV
NDTVGTMMTCGYEDPYCEVGLIVGTGSNACYMEEMRNVELVEGDEGRMCVNMEWGAFGDNGCLDDFRTEF
DVAVDEHSLNPGKQRFEKMISGMYLGEIVRNILIDFTKRGLLFRGRISERLKTRGIFETKFLSQIESDRL
ALLQVRSILQHLGLESTCDDSIIVKEVCTVIARRAAQLCGAGMAAVVDKIRENRGLDFLKVTVGVDGTLY
KLHPHFSTVMHETVKQLSPKCEVTFLQSEDGSGKGAALITAVACRIREAGQH
>anc120_altAll|avgPP:0.991|lnPP:-16.099|num_ambig:10|num_ambig_gaps:0
LYHMRLSDETLQEVSNRFRKEMEKGLGADTNPTASVKMLPTFVRSTPDGTEEGDFLALDLGGTNFRVLRV
KVSDNGSKKVEMENQIYAIPEDLMRGSGVQLFDHIAECLANFMEQLQMKDKKLPLGFTFSFPCYQTKLDE
SILVTWTKGFKSSGVEGKDVVSLLRKAIKKRGDFDIDIVAVVNDTVGTMMTCGYDDHNCEVGLIVGTGSN
ACYMEEMRHIDLVEGDEGRMCINMEWGAFGDDGVLNDIRTEFDREIDMGSLNPGKQLFEKMISGMYMGEL
VRLILVKMAKEELLFGGKLTPDLLTTGHFETRYVSAIEKEKEGLQKAHEILSKLGLEPSHEDCVATRRIC
EIVSTRSANLCGAALAAVLRRIKENKGVDRLRSTVGVDGTVYKKHPHFARRLHKTVRRLLPDCDIRFVRS
EDGSGKGAAMVTAVAYRLAAQHKARQKILEPLRLSHEQLLEVKKRMRQEMERGLARETHAEATVKMLPTY
VCATPDGTEKGDFLALDLGGTNFRVLLVRVRNGMRRSVEMHNKIYSIPQDIMQGTGEELFDHIVHCISDF
LEYMGMKGVSLPLGFTFSFPCHQTNLDEGILLKWTKGFKATGCEGEDVVGLLKEAIHRREEFDLDVVAVV
NDTVGTMMTCGYEDPYCEVGLIVGTGSNACYMEEMRNVELVEGDEGRMCVNMEWGAFGDNGCLDDFRTEF
DVAVDEHSLNPGKQRFEKMISGMYLGEIVRNILIDFTKRGLLFRGRISERLKTRGIFETKFLSQIESDCL
ALLQVRSILQHLGLESTCDDSIIVKEVCTVIARRAAQLCGAGMAAVVDKIRENRGLDFLKVTVGVDGTLY
KLHPHFSTVMHETVKQLSPKCEVTFLQSEDGSGKGAALITAVACRIREAGQH
>anc121|avgPP:0.997|lnPP:-3.826|num_ambig:2|num_ambig_gaps:0
LYHMRLSDETLQEVSNRFRKEMEKGLGADTNPTASVKMLPTFVRSTPDGTEDGDFLALDLGGTNFRVLRV
KVSDNGLQKVEMENQIYAIPEDLMRGSGVQLFDHIAECLANFMEKLQIKDKKLPLGFTFSFPCHQTKLDE
SILVTWTKGFKSSGVEGKDVVSLLRKAIKKRGDFDIDIVAVVNDTVGTMMTCGYDDHNCEVGLIVGTGTN
ACYMEEMRHIDLVEGDEGRMCINMEWGAFGDDGVLNDIRTEFDREIDMGSLNPGKQLFEKMISGMYMGEL
VRLILVKMAKEELLFGGRLTPDLLTTGHFETRYVSAIEKEKEGLQKAHEILTKLGLEPSHEDCVATHRIC
QIVSTRSANLCGATLAAVLRRIKENKGVDRLRSTVGVDGSVYKKHPHFARRLHKTVRRLLPDCDIRFVRS
EDGSGKGAAMVTAVAYRLAAQHKARQKTLEALKLSHEQLLEVKKRMRIEMERGLGKETHAEATVKMLPTY
VCSTPDGTEKGDFLALDLGGTNFRVLLVRVRNGMRRGVEMHNKIYSIPQEIMQGTGEELFDHIVHCISDF
LEYMGMKGVSLPLGFTFSFPCHQNNLDEGILLKWTKGFKATGCEGEDVVNLLKEAIHRREEFDLDVVAVV
NDTVGTMMTCGYEDPYCEVGLIVGTGSNACYMEEMRNVELVEGEEGRMCVNMEWGAFGDNGCLDDFRTEF
DVAVDELSLNPGKQRFEKMISGMYLGEIVRNILIDFTKRGLLFRGRISERLKTRGIFETKFLSQIESDCL
ALLQVRSILQHLGLESTCDDSIIVKEVCTVVARRAAQLCGAGMAAVVDKIRENRGLDFLKVTVGVDGTLY
KLHPHFSTIMHETVKQLSPKCEVTFLQSEDGSGKGAALITAVACRIREAGQR
>anc121_altAll|avgPP:0.997|lnPP:-4.729|num_ambig:2|num_ambig_gaps:0
LYHMRLSDETLQEVSNRFRKEMEKGLGADTNPTASVKMLPTFVRSTPDGTEDGDFLALDLGGTNFRVLRV
KVSDNGLQKVEMENQIYAIPEDLMRGSGVQLFDHIAECLANFMEKLNIKDKKLPLGFTFSFPCHQTKLDE
SILVTWTKGFKSSGVEGKDVVSLLRKAIKKRGDFDIDIVAVVNDTVGTMMTCGYDDHNCEVGLIVGTGTN
ACYMEEMRHIDLVEGDEGRMCINMEWGAFGDDGVLNDIRTEFDREIDMGSLNPGKQLFEKMISGMYMGEL
VRLILVKMAKEELLFGGRLTPDLLTTGHFETRYVSAIEKEKEGLQKAHEILTKLGLEPSHEDCVATHRIC
QIVSTRSANLCGATLAAVLRRIKENKGVDRLRSTVGVDGSVYKKHPHFARRLHKTVRRLLPDCDIRFVRS
EDGSGKGAAMVTAVAYRLAAQHKARQKTLEALKLSHEQLLEVKKRMRIEMERGLGKETHAEATVKMLPTY
VCSTPDGTEKGDFLALDLGGTNFRVLLVRVRNGMRRGVEMHNKIYSIPQEIMQGTGEELFDHIVHCISDF
LEYMGMKGVSLPLGFTFSFPCHQNNLDEGILLKWTKGFKATGCEGEDVVNLLKEAIHRREEFDLDVVAVV
NDTVGTMMTCGYEDPYCEVGLIVGTGSNACYMEEMRNVELVEGEEGRMCVNMEWGAFGDNGCLDDFRTEF
DVAVDELSLNPGKQRFEKMISGMYLGEIVRNILIDFTKRGLLFRGRISERLKTRGIFETKFLSQIESDCL
ALLQVRSILQHLGLESTCDDSIIVKEVCTVVARRAAQLCGAGMAAVVDKIRENRGLDFLKVTVGVDGTLY
KLHPHFSTVMHETVKQLSPKCEVTFLQSEDGSGKGAALITAVACRIREAGQR
>anc122|avgPP:0.995|lnPP:-6.581|num_ambig:5|num_ambig_gaps:0
LYHMRLSDETLQEVSNRFRKEMEKGLGADTNPTASVKMLPTFVRSTPDGTEDGDFLALDLGGTNFRVLRV
KVSDNGLQKVEMENQIYAIPEDLMRGSGTQLFDHIAECLANFMEKLQIKDKKLPLGFTFSFPCHQTKLDE
SILVTWTKGFKSSGVEGKDVVSLLRKAIKKRGDFDIDIVAVVNDTVGTMMTCGYDDHNCEIGLIVGTGTN
ACYMEEMRHIDLVEGDEGRMCINMEWGAFGDDGVLNDIRTEFDREIDMGSLNPGKQLFEKMISGMYMGEL

VRLILVKMAKEELLFGGRLTPDLLTTGHFETRYVSAIEKEKEGIQKAHEILTKLGLEPSQEDCVATHRIC
QIVSTRSANLCAATLAAVLRRIKENKGVERLRSTVGVDGSVYKKHPHFARRLHKTVRRLVPDCDIRFVRS
EDGSGKGAAMVTAVAYRLAAQHKARQKTLEALKLSHEQLLEVKKRMRIEMERGLGKETHASATVKMLPTY
VCSTPDGTEKGDFLALDLGGTNFRVLLVRVRNGMRRGVEMHNKIYSIPQEIMQGTGEELFDHIVHCIADF
LEYMGMKGVSLPLGFTFSFPCHQNNLDEGILLKWTKGFKATGCEGEDVVNLLKEAIHRREEFDLDVVAVV
NDTVGTMMTCGYEDPYCEVGLIVGTGSNACYMEEMRNVELVEGEEGRMCINMEWGAFGDNGCLDDFRTEF
DVAVDELSLNPGKQRFEKMISGMYLGEIVRNILIDFTKRGLLFRGRISERLKTRGIFETKFLSQIESDCL
ALLQVRSILQHLGLESTCDDSIIVKEVCTVVARRAAQLCGAGMAAVVDKIRENRGLDFLKVTVGVDGTLY
KLHPHFSTIMHETVKELSPKCEVTFLQSEDGSGKGAALITAVACRIREAGQR
>anc122_altAll|avgPP:0.995|lnPP:-9.015|num_ambig:5|num_ambig_gaps:0
LYHMRLSDETLQEVSNRFRKEMEKGLGADTNPTASVKMLPTFVRSTPDGTEDGDFLALDLGGTNFRVLRV
KVSDNGLQKVEMENQIYAIPEDLMRGSGTQLFDHIAECLANFMEKLNIKDKKLPLGFTFSFPCHQTKLDE
SILVTWTKGFKSSGVEGKDVVSLLRKAIKKRGDFDIDIVAVVNDTVGTMMTCGYDDHNCEIGLIVGTGTN
ACYMEEMRHIDLVEGDEGRMCINMEWGAFGDDGVLDDIRTEFDREIDMGSLNPGKQLFEKMISGMYMGEL
VRLILVKMAKEELLFGGRLTPDLLTTGHFETRYVSAIEKEKEGIQKAHEILTKLGLEPSQEDCVATHRIC
QIVSTRSANLCAATLAAVLRRIKENKGVERLRSTVGVDGSVYKKHPHFARRLHKTVRRLVPDCDIRFVRS
EDGSGKGAAMVTAVAYRLAAQHKARQKTLEALKLSHEQLLEVKKRMRVEMERGLGKETHASATVKMLPTY
VCSTPDGTEKGDFLALDLGGTNFRVLLVRVRNGMRRGVEMHNKIYSIPQEIMQGTGEELFDHIVHCIADF
LEYMGMKGVSLPLGFTFSFPCHQNNLDEGILLKWTKGFKATGCEGEDVVNLLKEAIHRREEFDLDVVAVV
NDTVGTMMTCGYEDPYCEVGLIVGTGSNACYMEEMRNVELVEGEEGRMCVNMEWGAFGDNGCLDDFRTEF
DVAVDELSLNPGKQRFEKMISGMYLGEIVRNILIDFTKRGLLFRGRISERLKTRGIFETKFLSQIESDCL
ALLQVRSILQHLGLESTCDDSIIVKEVCTVVARRAAQLCGAGMAAVVDKIRENRGLDFLKVTVGVDGTLY
KLHPHFSTVMHETVKELSPKCEVTFLQSEDGSGKGAALITAVACRIREAGQR
>anc123|avgPP:0.994|lnPP:-8.686|num_ambig:11|num_ambig_gaps:0
LYHMRLSDETLQEVSNRFRKEMEKGLGADTNPTASVKMLPTFVRSTPDGTEDGDFLALDLGGTNFRVLRV
KVSDNGLQKVEMENQIYAIPEDLMRGSGTQLFDHIAECLANFMEKLHIKDKKLPLGFTFSFPCHQTKLDE
SILVTWTKGFKASGVEGKDVVSLLRKAIKKRGDFDIDIVAVVNDTVGTMMTCGYDDHNCEIGLIVGTGTN
ACYMEEMRHIDLVEGDEGRMCINMEWGAFGDDGVLDDIRTEFDREIDMGSLNPGKQLFEKMISGMYMGEL
VRLILVKMAKEDLLFGGRITPDLLTTGHFETRFVSAIEKDKEGIQKAHEILTKLGLEPSQEDCVATHRIC
QIVSTRSANLCAATLAAVLRRIKENKGVERLRSTVGVDGSVYKKHPHFARRLHKTVRRLVPDCDIRFVRS
EDGSGKGAAMVTAVAYRLAAQQKARQKTLEALKLSHEQLLEVKKRMRIEMERGLGKETHNSATVKMLPTY
VRSTPDGTEKGDFLALDLGGTNFRVLLVRVRNGMRRGVEMHNKIYSIPQEIMQGTGEELFDHIVHCIADF
LEYMGMKGVSLPLGFTFSFPCHQNNLDEGILLKWTKGFKATGCEGEDVVNLLKEAIHRREEFDLDVVAVV
NDTVGTMMTCGYEDPYCEVGLIVGTGSNACYMEEMRNVELVEGEEGRMCINMEWGAFGDNGCLDDFRTEF
DRAVDELSLNPGKQRYEKMISGMYLGEIVRNILIDFTKRGLLFRGRISERLKTRGIFETKFLSQIESDCL
ALLQVRSILQHLGLESTCDDSIIVKEVCTVVARRAAQLCGAGMAAVVDKIRENRGLDFLKVTVGVDGTLY
KLHPHFSTIMHETVKELSPKCEVTFLQSEDGSGKGAALITAVACRIREAGQR
>anc123_altAll|avgPP:0.993|lnPP:-13.211|num_ambig:11|num_ambig_gaps:0
LYHMRLSDETLQDVSNRFRKEMEKGLGADTNPTASVKMLPTFVRSTPDGTENGDFLALDLGGTNFRVLRV
KVSDNGLQKVEMENQIYAIPEDLMRGSGTQLFDHIAECLANFMEKLNIKDKKLPLGFTFSFPCHQTKLDE
SILVTWTKGFKASGVEGKDVVSLLRKAIKKRGDFDIDIVAVVNDTVGTMMTCGYDDHNCEIGLIVGTGTN
ACYMEEMRHIDLVEGDEGRMCINMEWGAFGDDGVLNDIRTEFDREIDMGSLNPGKQLFEKMISGMYMGEL
VRLILVKMAKEELLFGGRITPDLLTTGHFETRFVSAIEKDKEGIQKAHEILTKLGLEPSQEDCVATHRIC
QIVSTRSANLCAATLAAVLRRIKENKGVERLRSTVGVDGSVYKKHPHFARRLHKTVRRLVPDCDIRFVRS
EDGSGKGAAMVTAVAYRLAAQQKARQKTLEALKLSHEQLLEVKKRMRVEMERGLGKETHSSATVKMLPTY
VRSTPDGTEKGDFLALDLGGTNFRVLLVRVRNGMRRGVEMHNKIYSIPQEIMQGTGEELFDHIVHCIADF
LEYMGMKGVSLPLGFTFSFPCHQNNLDEGILLKWTKGFKATGCEGEDVVNLLKEAIHRREEFDLDVVAVV
NDTVGTMMTCGYEDPYCEVGLIVGTGSNACYMEEMRNVELVEGEEGRMCVNMEWGAFGDNGCLDDFRTEF
DRAVDELSLNPGKQRFEKMISGMYLGEIVRNILIDFTKRGLLFRGRISERLKTRGIFETKFLSQIESDRL
ALLQVRSILQHLGLESTCDDSIIVKEVCTVVARRAAQLCGAGMAAVVDKIRENRGLDFLKVTVGVDGTLY
KLHPHFSTVMHETVKELSPKCEVTFLQSEDGSGKGAALITAVACRIREAGQR
>anc124|avgPP:0.994|lnPP:-9.636|num_ambig:14|num_ambig_gaps:0

LYHMRLSDETLQDISNRFRKEMEKGLGRDTNPTASVKMLPTFVRSTPDGTENGDFLALDLGGTNFRVLRV
KVSDNGKQKVEMENQIYAIPEDLMRGSGTQLFDHIAECLANFMEKLNIKDKKLPLGFTFSFPCQQTKLDE
SILVTWTKGFKASGVEGRDVVSLLRKAIKKRGDFDIDIVAVVNDTVGTMMTCGYDDHNCEIGLIVGTGTN
ACYMEEMRHIDLVEGDEGRMCINMEWGAFGDDGVLDDIRTEFDREIDMGSLNPGKQLFEKMISGMYMGEL
VRLILVKMAKEGLLFGGRITPDLLTTGHFETRFVSAIEKDKEGIQKAHEILTKLGLEPSQEDCVATQRIC
QIVSTRSANLCAATLAAVLRRIKENKGVERLRTTVGVDGTVYKKHPQFARRLHKAVRRLVPDCDVRFVRS
EDGSGKGAAMVTAVAYRLASQQKARQKTLEALKLSHEQLLEVKKRMRIEMERGLKKETHNSATVKMLPTY
VRSTPDGTEKGDFLALDLGGTNFRVLLVRVRNGMRRGVEMHNKIYSIPQEVMQGTGEELFDHIVHCIADF
LEYMGMKGVSLPLGFTFSFPCHQNNLDEGILLKWTKGFKATGCEGEDVVNLLKEAIHRREEFDLDVVAVV
NDTVGTMMTCGYEDPYCEVGLIVGTGSNACYMEEMRNVELVEGEEGRMCINMEWGAFGDNGCLDDFRTEF
DRAVDELSLNPGKQRYEKMISGMYLGEIVRNILIDFTKRGLLFRGRISERLKTRGIFETKFLSQIESDRL
ALLQVRSILQHLGLESTCDDSIIVKEVCTVVARRAAQLCGAGMAAVVDKIRENRGLDFLKVTVGVDGTLY
KLHPHFSTIMHETVKELSPKCEVTFLQSEDGSGKGAALITAVACRIREAGQR
>anc124_altAll|avgPP:0.990|lnPP:-17.832|num_ambig:14|num_ambig_gaps:0
LYHMRLSDETLQDISNRFRKEMEKGLGADTNPTASVKMLPTFVRSTPDGTENGDFLALDLGGTNFRVLRV
KVSDNGRQKVEMENQIYAIPEDLMRGSGTQLFDHIAECLANFMEKLNIKDKKLPLGFTFSFPCHQTKLDE
SILVTWTKGFKASGVEGKDVVSLLRKAIKKRGDFDIDIVAVVNDTVGTMMTCGYDDHNCEIGLIVGTGTN
ACYMEEMRHIDLVEGDEGRMCINMEWGAFGDDGVLNDIRTEFDREIDMGSLNPGKQLFEKMISGMYMGEL
VRLILVKMAKEGLLFGGKITPDLLTTGHFETRFVSAIEKDKEGIQKAHEILTKLGLEPSQEDCVATHRIC
QIVSTRSANLCAATLAAVLRRIKENKGVERLRSTVGVDGTVYKKHPQFARRLHKTVRRLVPDCDVRFVRS
EDGSGKGAAMVTAVAYRLASQQKARQKTLEALKLSHEQLLEVKKRMRVEMERGLKKETHSSATVKMLPTY
VRSTPDGTEKGDFLALDLGGTNFRVLLVRVRNGMRRGVEMHNKIYSIPQEVMQGTGEELFDHIVHCIADF
LEYMGMKGVSLPLGFTFSFPCHQNNLDEGILLKWTKGFKATGCEGEDVVNLLKEAIHRREEFDLDVVAVV
NDTVGTMMTCGYEDPYCEVGLIVGTGSNACYMEEMRNVELVEGEEGRMCVNMEWGAFGDNGCLDDFRTEF
DRAVDELSLNPGKQRYEKMISGMYLGEIVRNILIDFTKRGLLFRGRISERLKTRGIFETKFLSQIESDRL
ALLQVRSILQHLGLESTCDDSIIVKEVCTVVARRAAQLCGAGMAAVVDKIRENRGLDFLKVTVGVDGTLY
KLHPHFSTVMHETVKELSPKCEVTFLQSEDGSGKGAALITAVACRIREAGQR
>anc125|avgPP:0.993|lnPP:-9.724|num_ambig:9|num_ambig_gaps:0
LYHMRLSDETLQDISNRFRKEMEKGLGRDTNPTASVKMLPTFVRSTPDGTENGDFLALDLGGTNFRVLRV
KVSDNGKQKVEMENQIYAIPEDLMRGSGTQLFDHIAECLANFMEKLNIKDKKLPLGFTFSFPCQQTKLDE
SILVTWTKGFKASGVEGRDVVSLLRKAIKKRGDFDIDIVAVVNDTVGTMMTCGYDDHNCEIGLIVGTGTN
ACYMEEMRHIDLVEGDEGRMCINMEWGAFGDDGVLDDIRTEFDREIDMGSLNPGKQLFEKMISGMYMGEL
VRLILVKMAKEGLLFGGKITPDLLTTGHFETRFVSAIEKDKEGIQKAHEILTKLGLEPSQEDCVATQRIC
QIVSTRSANLCAATLAAVLRRIKENKGVERLRTTVGVDGTVYKKHPQFARRLHKAVRRLVPDCDVRFVRS
EDGSGKGAAMVTAVAYRLASQQKARQKTLDAFKLSHEQLLEVKKRMRVEMERGLKKETHNSATVKMLPTY
VRSTPDGTEKGDFLALDLGGTNFRVLLVRVRNGMRRGVEMHNKIYSIPQEVMQGTGEELFDHIVHCIADF
LEYMGMKGVSLPLGFTFSFPCHQNNLDEGILLKWTKGFKATGCEGEDVVNLLKEAIHRREEFDLDVVAVV
NDTVGTMMTCGYEDPYCEVGLIVGTGSNACYMEEMRNVELVEGEEGRMCINMEWGAFGDNGCLDDFRTEF
DRAVDELSLNPGKQRYEKMISGMYLGEIVRNILIDFTKRGLLFRGRISERLKTRGIFETKFLSQIESDRL
ALLQVRSILQHLGLESTCDDSIIVKEVCTVVARRAAQLCGAGMAAVVDKIRENRGLDFLKVTVGVDGTLY
KLHPHFSTIMHETVKELAPKCEVTFLQSEDGSGKGAALITAVACRIREAGQR
>anc125_altAll|avgPP:0.992|lnPP:-13.411|num_ambig:9|num_ambig_gaps:0
LYHMRLSDETLQDISNRFRKEMEKGLGRDTNPTASVKMLPTFVRSTPDGTENGDFLALDLGGTNFRVLRV
KVSDNGKQKVEMENQIYAIPEDLMRGSGTQLFDHIAECLANFMEKLNIKDKKLPLGFTFSFPCHQTKLDE
SILVTWTKGFKASGVEGKDVVSLLRKAIKKRGDFDIDIVAVVNDTVGTMMTCGYDDHNCEIGLIVGTGTN
ACYMEEMRHIDLVEGDEGRMCINMEWGAFGDDGVLDDIRTEFDREIDMGSLNPGKQLFEKMISGMYMGEL
VRLILVKMAKEGLLFGGRITPDLLTTGHFETRFVSAIEKDKEGIQKAHEILTKLGLEPSQEDCVATHRIC
QIVSTRSANLCAATLAAVLRRIKENKGVERLRTTVGVDGTVYKKHPQFARRLHKAVRRLVPDCDVRFVRS
EDGSGKGAAMVTAVAYRLASQQKARQKTLDAFKLSHEQLLEVKKRMRIEMERGLKKETHSSATVKMLPTY
VRSTPDGTEKGDFLALDLGGTNFRVLLVRVRNGMRRGVEMHNKIYSIPQEVMQGTGEELFDHIVHCIADF
LEYMGMKGVSLPLGFTFSFPCHQNNLDEGILLKWTKGFKATGCEGEDVVNLLKEAIHRREEFDLDVVAVV
NDTVGTMMTCGYEDPYCEVGLIVGTGSNACYMEEMRNVELVEGEEGRMCVNMEWGAFGDNGCLDDFRTEF

DRAVDELSLNPGKQRYEKMISGMYLGEIVRNILIDFTKRGLLFRGRISERLKTRGIFETKFLSQIESDRL
ALLQVRSILQHLGLESTCDDSIIVKEVCTVVARRAAQLCGAGMAAVVDKIRENRGLDFLKVTVGVDGTLY
KLHPHFSTVMHETVKELAPKCEVTFLQSEDGSGKGAALITAVACRIREAGQR
>anc126|avgPP:0.996|lnPP:-6.451|num_ambig:7|num_ambig_gaps:0
LHHFQLSDKTLMDLSIRFRREMDKGLCRDTNPTAAVKMLPTFVRSTPDGTEQGEFLALDLGGSNFRVLLV
KVMANGKQEVEMENQIYDIPEHLMRGSGSELFDHIADCLANFLEKLGIKDKKLPLGFTFSFPCQQTKLDE
SVLVSWTKSFKSSGVEGKDVVSLLRKSIKKRGDFDVDILAVINDTVGTMMTCGYDDHHCEIGLIVGTGTN
ACYMEQMRNLDLLDGDEGRMCVNTEWGAFGDDGALEDLRTDIDREIDAGSLNPGKQLFEKMISGLYMGEL
VRLILVRMTKEQLLFQGKTTAQLLTTGSFNTSYIYAIENDKDEEGLASAEKVLRGLGLDPSVEDCIATQR
VCQIVSTRAAHLCAATLVAVLRQIRDNKAAEKLRTTIGVDGSVYKNHPEFSRRLHKMVRRLVPDCDVRFL
QSQCGSGKGAAMVTAVAYRLAAQHAERQRILDTLRLSREQLLEVKKRMSEEMVRGLSKQTHEQASVKMLP
TYVRSTPDGTEHGDFLALDLGGSSFRVLLVRLRSGKGHNVDMHHKIYSIPQETMQGTGEELFSHIVDCIA
DFLEYMGMRGASLPLGFTFSFPCHQSKLDQGILLKWTKGFKASGCEGQDVITLLKDAVRRRQEFDLNFVA
VVNDTVGTMMTCSYEDPQCEVGLIVGTGTNACYMEEMQNIELVEGDDGRMCVNMEWGAFGDNGELDDFCT
QFDRIVDDSSNNPGKQRYEKMISGMYLGEIVRNVLIDFTAKGLLFRGKLSERLKTRGIFETKFLAQIESD
RLAMRQVRSILQHLGLTTSTCDDSVLVKEVCSVVSRRAAQLCGAGLAAVVDKIRQNRNLNQLSITVGVDG
TLYKTHPHFSSIMQETLQDLAPQCQVTFHKSEDGSGKGAALITAVACRVKSEGQH
>anc126_altAll|avgPP:0.994|lnPP:-10.904|num_ambig:7|num_ambig_gaps:0
LHHFQLSDKTLMDLSIRFRREMDKGLCRDTNPTAAVKMLPTFVRSTPDGTEQGEFLALDLGGSNFRVLLV
KVMANGKQEVEMENQIYDIPEHLMRGCGSELFDHIADCLANFLEKLGIKDKKLPLGFTFSFPCQQTKLDE
SVLVSWTKSFKSSGVEGKDVVSLLRKSIKKRGDFDVDILAVINDTVGTMMTCGYDDHHCEIGLIVGTGTN
ACYMEQMRNLDLLDGDEGRMCVNTEWGAFGDDGALEDLRTDIDREIDAGSLNPGKQLFEKMISGLYMGEL
VRLILVRMTKEKLLFQGKTTAQLLTTGSFNTSYIYAIENDKDEEGLASAEKVLRGLGLDPSVEDCIATQR
VCQIVSTRAAHLCAATLVAVLRQIRDNKAAEKLRTTIGVDGSVYKNHPEFSRRLHKMVRRLVPDCDVRFL
QSQCGSGKGAAMVTAVAYRLAAQHAERQRILDTLRLSREQLLEVKKRMSEEMARGLSKQTHEQASVKMLP
TYVRSTPDGTEHGDFLALDLGGSSFRVLLVRLRSGKGHNVDMHHKIYSIPQETMQGTGEELFSHIVDCIA
DFLEYMGMRGASLPLGFTFSFPCHQSKLDQGILLKWTKGFKASGCEGQDVITLLKDAVRRRQEFDLNFVA
VVNDTVGTMMTCSYEDPQCEVGLIVGTGTNACYMEEMQNIELVEGDDGRMCVNMEWGAFGDNGELDDFCT
QFDHVVDDSSNNPGKQRYEKMISGMYLGEIVRNVLIDFTAKGLLFRGKLSERLKTRGIFETKFLAQIESD
RLAMRQIRSILQHLGLTTSTCDDSVLVKEVCSVVSRRAAQLCGAGLAAVVDKIRQNRNLNQLSITVGVDG
TLYKTHPHFSSIMQETLQDLAPQCQVTFHKSEDGSGKGAALITAVACRIKSEGQH
>anc127|avgPP:0.996|lnPP:-5.865|num_ambig:8|num_ambig_gaps:0
LHHFQLSDKTLMDLSIRFRREMDKGLCRDTNPTAAVKMLPTFVRSTPDGTEQGEFLALDLGGSNFRVLLV
KVMANGKQEVEMENQIYDIPEHLMRGSGSELFDHIADCLANFLEKLGIKDKKLPLGFTFSFPCQQTKLDE
SVLVSWTKGFKSSGVEGKDVVSLLRKSIKKRGDFDIDILAVINDTVGTMMTCGYDDHHCEIGLIVGTGTN
ACYMEQMRNLELLDGDEGRMCVNTEWGAFGDDGALEDLRTDIDREIDAGSLNPGKQLFEKMISGLYMGEL
VRLILVRMTKEQLLFQGKTTAQLLTTGSFNTSYIYAIENDKDEEGLASAEKVLRGLGLDPSVEDCIATQR
VCQIVSTRAAHLCAATLVAVLRQIRDNKAAEKLRTTIGVDGSVYKNHPEFSRRLHKMVRRLVPDCDVRFL
QSQCGSGKGAAMVTAVAYRLAAQHAERQRILDTLRLSREQLLEVKKRMSEEMVRGLSKQTHEQASVKMLP
TYVRSTPDGTEHGDFLALDLGGSSFRVLLVRVRSGKRHNVDMHHKIYSIPQETMQGTGEELFSHIVDCIA
DFLEYMGMRGASLPLGFTFSFPCHQSKLDQGILLKWTKGFKASGCEGQDVITLLKDAVRRRQEFDLNFVA
VVNDTVGTMMTCSYEDPQCEVGLIVGTGTNACYMEEMQNIELVEGDDGRMCVNMEWGAFGDNGELDDFCT
QFDRIVDDCSNNPGKQRYEKMISGMYLGEIVRNVLIDFTAKGLLFRGKLSERLKTRGIFETKFLSQIESD
RLAMRQVRSILQHLGLTTSTCDDSVLVKEVCSVVARRAAQLCGAGLAAVVDKIRQNRNLNQLSITVGVDG
TLYKTHPHFSSIMQETLQDLAPQCQVTFHKSEDGSGKGAALITAVACRIKSEGQH
>anc127_altAll|avgPP:0.994|lnPP:-10.131|num_ambig:8|num_ambig_gaps:0
LHHFQLSDKTLMDLSIRFRREMDKGLCRDTNPTAAVKMLPTFVRSTPDGTEQGEFLALDLGGSNFRVLLV
KVMANGKQEVEMENQIYDIPEHLMRGCGSELFDHIADCLANFLEKLGIKDKKLPLGFTFSFPCQQTKLDE
SVLVSWTKGFKSSGVEGKDVVSLLRKSIKKRGDFDIDIVAVINDTVGTMMTCGYDDHHCEIGLIVGTGTN
ACYMEQMRNLELLDGDEGRMCVNTEWGAFGDDGALEDLRTDIDREIDAGSLNPGKQLFEKMISGLYMGEL
VRLILVRMTKEKLLFQGKTTAQLLTTGSFNTSYIYAIENDKDEEGLASAEKVLRGLGLDPSVEDCIATQR
VCQIVSTRAAHLCAATLVAVLRQIRDNKAAEKLRTTIGVDGSVYKNHPEFSRRLHKMVRRLVPDCDVRFL

QSQCGSGKGAAMVTAVAYRLAAQHAERQRILDTLRLSREQLLEVKKRMSEEMVRGLSKQTHEQASVKMLP
TYVRSTPDGTEHGDFLALDLGGSSFRVLLVRVRSGKKHNVDMHHKIYSIPQETMQGTGEELFSHIVDCIA
DFLEYMGMRGASLPLGFTFSFPCHQSKLDQGILLKWTKGFKASGCEGQDVITLLKDAVRRRQEFDLNFVA
VVNDTVGTMMTCSYEDPQCEVGLIVGTGTNACYMEEMQNIELVEGDDGRMCVNMEWGAFGDNGELDDFCT
QFDHIVDDSSNNPGKQRYEKMISGMYLGEIVRNVLIDFTAKGLLFRGKLSERLKTRGIFETKFLSQIESD
RLAMRQIRSILQHLGLTTSTCDDSVLVKEVCSVVARRAAQLCGAGLAAVVDKIRQNRNLNQLSITVGVDG
TLYKTHPHFSSIMQETLQDLAPQCQVTFHKSEDGSGKGAALITAVACRVKSEGQH
>anc128|avgPP:0.993|lnPP:-11.253|num_ambig:12|num_ambig_gaps:0
LQHFQLSDETLMDVSIRFRREMDKGLCRDTNPTAAVKMLPTFVLSTPDGTEQGEFLALDLGGSNFRVLLV
KVMADGEQKVEMENQIYDIPEHLMRGSGTELFDHIADCLANFLEKLGIKDKKLPLGFTFSFPCQQTKLDE
SVLLSWTKAFKSSGVEGKDVVSLLRKSIKKRGDFDVDILAVINDTVGTMMTCGYDDHLCEIGLIVGTGTN
ACYMEQMRNIELLDGDEGRMCVNTEWGAFGDDGTLEDLRTDIDREIDAGSLNPGKQLFEKMISGMYMGEL
VRLILVRMTKEQLLFQGKTTAELLTTGSFKTSYIYAIDNDNDEEGLASAEKVLRGLNLDPSVEDCIATQR
VCQIVSTRAAHLCAATLVAVLRQIRDNKAAEKLRTTIGVDGSVYKNHPEFSRRLHKMVRRLVPDCDVRFL
QSQCGSGKGAAMVTAVAYRLAAQNAERQRILDTLRLSREQLLEVKRRMSEEMVRGLSKQTHEQASVKMLP
TYVRSTPDGTEHGDFLALDLGGSSFRVLLVQVQSGKRHNADMHQKIYSIPQETMQGTGEELFDHIVDCIA
DFLEYMGMRGASLPLGFTFSFPCHQSRLDQGILLKWTKGFKASGCEGQDVIVLLKDAVRRRQEFDLNFVA
VVNDTVGTMMTCGYEDPKCEVGLIVGTGTNACYMEEMQNIELVEGDEGRMCVNMEWGAFGDNGELDDFCT
RFDRIVDDSSNNPGKQRYEKMISGMYLGEIVRNVLIDFTAKGLLFRGKLSERLKTRGIFATKFLSQIESD
RLAMRQVRSILQHLGLTSSTCDDSVLVKEVCSVVARRAAQLCGAGLAAVVDKMRENRNLNQLSVTVGVDG
TLYKTHPHFSSIMQETLQDLAPQCQVTFHKSEDGSGKGAALITAVACRIKSEGQH
>anc128_altAll|avgPP:0.990|lnPP:-17.524|num_ambig:12|num_ambig_gaps:0
LQHLQLSDETLMDVSIRFRREMDKGLCRDTNPTAAVKMLPTFVLSTPDGTEQGEFLALDLGGSNFRVLLV
KVMADGEQKVEMENQIYDIPEHLMRGSGSELFDHIADCLANFMEKLGIKDKKLPLGFTFSFPCQQTKLDE
SVLLSWTKAFKSSGVEGKDVVSLLRKSIKKRGDFDVDILAVINDTVGTMMTCGYDDHLCEIGLIVGTGTN
ACYMEQMRNIELLDGDEGRMCVNTEWGAFGDDGTLEDLRTDIDREIDAGSLNPGKQLFEKMISGMYMGEL
VRLILVRMTKEQLLFQGKTTAQLLTTGSFKTSYIYTIDNDNDEEGLASAEKVLRGLNLDPSVEDCIATQR
VCQIVSTRAAHLCAATLVAVLRQIRDNKAAEKLRTTIGVDGSVYKNHPEFSRRLHKMVRRLVPDCDVRFL
QSQCGSGKGAAMVTAVAYRLAAQNAERQRILDTLRLSREQLLEVKKRMSEEMVRGLSKQTHEQASVKMLP
TYVRSTPDGTEHGDFLALDLGGSSFRVLLVQVQSGKKHNVKMHQKIYSIPQETMQGTGEELFDHIVDCIA
DFLEYMGMKGASLPLGFTFSFPCHQSRLDQGILLKWTKGFKASGCEGQDVIVLLKDAVRRRQEFDLNFVA
VVNDTVGTMMTCGYEDPKCEVGLIVGTGTNACYMEEMQNIELVEGDEGRMCVNMEWGAFGDNGELDDFCT
RFDHIVDDSSNNPGKQRYEKMISGMYLGEIVRNVLIDFTSKGLLFRGKLSERLKTRGIFATKFLSQIESD
RLAMRQVRSILQHLGLTSSTCDDSVLVKEVCSVVARRAAQLCGAGLAAVVDKMRENRNLNQLSVTVGVDG
TLYKTHPHFSSIMQETLQDLAPQCQVTFHKSEDGSGKGAALITAVACRIKSEGQH
>anc129|avgPP:0.993|lnPP:-10.880|num_ambig:9|num_ambig_gaps:0
LPHFQLSDETIMDVSVRFRREMDKGLCRDTNPAAAVKMLPTFVLSTPDGTEQGEFLALDLGGSNFRVLLV
KVRANGKQEVEMENQIYDIPENLMRGSGSELFDHIADCLADFMEKLGIKDKKLPLGFTFSFPCQQTKLDE
SVLLSWTKAFKASGVEGKDVVSLLRKSIKKRGDFDIDILAVINDTVGTMMTCGYDDHLCEIGLIVGTGTN
ACYMEQMRNLELLDGDEGRMCVNTEWGAFGDDGALEDLRTDIDREIDAGSLNPGKQLFEKMISGMYMGEM
VRLILVRMTKEKLLFQGKTTAQLLTTGSFKTSYIYTIDTDKDEEGLQSAEEVLRGLGLDPSVEDCIATQR
VCQIVSTRAAHLCAASLVAVLRQIRDNKAAEKLRATIGVDGSVYKNHAEFSRRLHKMVRRLVPDCDVRFL
QSQCGSGKGAAMVTAVAYRLAAQNAERQRILDTLRLSREQLLEVKKRMSEEMVRGLSRQTHEQASVKMLP
TYVRSTPDGTEHGDFLALDLGGSSFRVLLVQLRSGKKHTVDMHQKIYSIPQETMQSTGEELFDHIVDCIA
DFLEYMGMRGASLPLGFTFSFPCHQSKLDQGILLKWTKGFKASGCEGQDVIMLLKDAVRRRQEFDLNFVA
VVNDTVGTMMTCGYEDPKCEVGLIVGTGTNACYMEEMRNIELVEGDDGRMCVNMEWGAFGENGELDDFCT
QFDRIVDESSNNPGKQRYEKMISGMYLGEIVRNVLIDYTAKGLLFRGKLSERLKTRGIFETKFLSQIERD
RLAMRQVRSILQHLGLTSSTCDDSVLVKEVCSVVARRAAQLCGAGLAAVVDKIRENRNLNQLSITVGVDG
TLYKTHPHFSSIMQETLQDLAPQCQVTFHKSEDGSGKGAALITAVACRIQNEGQH
>anc129_altAll|avgPP:0.992|lnPP:-14.420|num_ambig:9|num_ambig_gaps:0
LPHLQLSDETIMDVSVRFRREMDKGLCRDTNPAAAVKMLPTFVLSTPDGTEQGEFLALDLGGSNFRVLLV
KVSANGKQEVEMENQIYDIPENLMRGSGSELFDHIADCLADFMEKLGIKDKKLPLGFTFSFPCQQTKLDE

SVLLSWTKAFKASGVEGKDVVSLLRKSIKKRGDFDIDILAVINDTVGTMMTCGYDDHLCEIGLIVGTGTN
ACYMEQMRNLELLDGDEGRMCVNTEWGAFGDDGALEDLRTDIDREIDAGSLNPGKQLFEKMISGMYMGEM
VRLILVRMTKEKLLFQGKTTAQLLTTGSFKTSYIYAIDTDKDEEGLASAEKVLRGLGLDPSVEDCIATQR
VCQIVSTRAAHLCAASLVAVLRQIRDNKAAEKLRATIGVDGSVYKNHTEFSRRLHKMVRRLVPDCDVRFL
QSQCGSGKGAAMVTAVAYRLAAQNAERQRILDTLRLSREQLLEVKKRMSEEMVRGLSRQTHEQTSVKMLP
TYVRSTPDGTEHGDFLALDLGGSSFRVLLVQLRSGKKHTVDMHQKIYSIPQETMQSTGEELFDHIVDCIA
DFLEYMGMRGASLPLGFTFSFPCHQSKLDQGILLKWTKGFKASGCEGQDVIVLLKDAVRRRQEFDLNFVA
VVNDTVGTMMTCGYEDPKCEVGLIVGTGTNACYMEEMRNIELVEGDDGRMCVNMEWGAFGENGELDDFCT
QFDRIVDESSNNPGKQRYEKMISGMYLGEIVRNVLIDYTAKGLLFRGKLSERLKTRGIFETKFLSQIERD
RLAMRQVRSILQHLGLTSSTCDDSVLVKEVCSVVARRAAQLCGAGLAAVVDKIRENRNLNQLSVTVGVDG
TLYKTHPHFSSIMQETLQDLAPQCQVTFHKSEDGSGKGAALITAVACRIQNEGQH
>anc130|avgPP:0.995|lnPP:-6.813|num_ambig:8|num_ambig_gaps:0
LHHFQLSDETLMDVSIRFRREMDKGLCRDTNPTAAVKMLPTFVLSTPDGTEQGEFLALDLGGSNFRVLLV
KVMANGKQKVEMENQIYDIPEHLMRGSGSELFDHIADCLANFLEKLGIKDKKLPLGFTFSFPCQQTKLDE
SVLLSWTKAFKSSGVEGKDVVSLLRKSIKKRGDFDIDILAVINDTVGTMMTCGYDDHLCEIGLIVGTGTN
ACYMEQMRNLELLDGDEGRMCVNTEWGAFGDDGALEDLRTDIDREIDAGSLNPGKQLFEKMISGMYMGEL
VRLILVRMTKEQLLFQGKTTAQLLTTGSFKTSYIYAIDNDKDEEGLASAEKVLRGLGLDPSVEDCIATQR
VCQIVSTRAAHLCAATLVAVLRQIRDNKAAEKLRTTIGVDGSVYKNHPEFSRRLHKMVRRLVPDCDVRFL
QSQCGSGKGAAMVTAVAYRLAAQNAERQRILDTLRLSREQLLEVKKRMSEEMVRGLSKQTHEQASVKMLP
TYVRSTPDGTEHGDFLALDLGGSSFRVLLVQVRSGKRHNVDMHQKIYSIPQETMQGTGEELFDHIVDCIA
DFLEYMGMRGASLPLGFTFSFPCHQSKLDQGILLKWTKGFKASGCEGQDVIVLLKDAVRRRQEFDLNFVA
VVNDTVGTMMTCGYEDPKCEVGLIVGTGTNACYMEEMQNIELVEGDDGRMCVNMEWGAFGDNGELDDFCT
QFDRIVDDSSNNPGKQRYEKMISGMYLGEIVRNVLIDFTAKGLLFRGKLSERLKTRGIFETKFLSQIESD
RLAMRQVRSILQHLGLTSSTCDDSVLVKEVCSVVARRAAQLCGAGLAAVVDKIRENRNLNQLSITVGVDG
TLYKTHPHFSSIMQETLQDLAPQCQVTFHKSEDGSGKGAALITAVACRIKSEGQH
>anc130_altAll|avgPP:0.994|lnPP:-11.013|num_ambig:8|num_ambig_gaps:0
LQHLQLSDETLMDVSIRFRREMDKGLCRDTNPTAAVKMLPTFVLSTPDGTEQGEFLALDLGGSNFRVLLV
KVMANGKQEVEMENQIYDIPEHLMRGSGSELFDHIADCLANFMEKLGIKDKKLPLGFTFSFPCQQTKLDE
SVLLSWTKAFKSSGVEGKDVVSLLRKSIKKRGDFDIDILAVINDTVGTMMTCGYDDHLCEIGLIVGTGTN
ACYMEQMRNLELLDGDEGRMCVNTEWGAFGDDGALEDLRTDIDREIDAGSLNPGKQLFEKMISGMYMGEL
VRLILVRMTKEQLLFQGKTTAQLLTTGSFKTSYIYTIDNDKDEEGLASAEKVLRGLGLDPSVEDCIATQR
VCQIVSTRAAHLCAATLVAVLRQIRDNKAAEKLRTTIGVDGSVYKNHPEFSRRLHKMVRRLVPDCDVRFL
QSQCGSGKGAAMVTAVAYRLAAQNAERQRILDTLRLSREQLLEVKKRMSEEMVRGLSKQTHEQASVKMLP
TYVRSTPDGTEHGDFLALDLGGSSFRVLLVQVRSGKKHNVDMHQKIYSIPQETMQGTGEELFDHIVDCIA
DFLEYMGMRGASLPLGFTFSFPCHQSKLDQGILLKWTKGFKASGCEGQDVIMLLKDAVRRRQEFDLNFVA
VVNDTVGTMMTCGYEDPKCEVGLIVGTGTNACYMEEMQNIELVEGDDGRMCVNMEWGAFGDNGELDDFCT
QFDRIVDDSSNNPGKQRYEKMISGMYLGEIVRNVLIDFTAKGLLFRGKLSERLKTRGIFETKFLSQIESD
RLAMRQVRSILQHLGLTSSTCDDSVLVKEVCSVVARRAAQLCGAGLAAVVDKIRENRNLNQLSVTVGVDG
TLYKTHPHFSSIMQETLQDLAPQCQVTFHKSEDGSGKGAALITAVACRIKSEGQH
>anc131|avgPP:0.997|lnPP:-4.931|num_ambig:6|num_ambig_gaps:0
LHHFQLSDETLMDVSIRFRREMDKGLCRDTNPTAAVKMLPTFVRSTPDGTEQGEFLALDLGGSNFRVLLV
KVMANGKQKVEMENQIYDIPEHLMRGSGSELFDHIADCLANFLEKLGIKDKKLPLGFTFSFPCQQTKLDE
SVLVSWTKGFKSSGVEGKDVVSLLRKSIKKRGDFDIDILAVINDTVGTMMTCGYDDHHCEIGLIVGTGTN
ACYMEQMRNLELLDGDEGRMCVNTEWGAFGDDGALEDLRTDIDREIDAGSLNPGKQLFEKMISGMYMGEL
VRLILVRMTKEQLLFQGKTTAQLLTTGSFNTSYIYAIENDKDEEGLASAEKVLRGLGLDPSVEDCIATQR
VCQIVSTRAAHLCAATLVAVLRQIRDNKAAEKLRTTIGVDGSVYKNHPEFSRRLHKMVRRLVPDCDVRFL
QSQCGSGKGAAMVTAVAYRLAAQHAERQRILDTLRLSREQLLEVKKRMSEEMVRGLSKQTHEQASVKMLP
TYVRSTPDGTEHGDFLALDLGGSSFRVLLVRVRSGKRHNVDMHHKIYSIPQETMQGTGEELFDHIVDCIA
DFLEYMGMRGASLPLGFTFSFPCHQSKLDQGILLKWTKGFKASGCEGQDVITLLKDAVRRRQEFDLNFVA
VVNDTVGTMMTCGYEDPKCEVGLIVGTGTNACYMEEMQNIELVEGDDGRMCVNMEWGAFGDNGELDDFCT
QFDRIVDDCSNNPGKQRYEKMISGMYLGEIVRNVLIDFTAKGLLFRGKLSERLKTRGIFETKFLSQIESD
RLAMRQVRSILQHLGLTSSTCDDSVLVKEVCSVVARRAAQLCGAGLAAVVDKIRQNRNLNQLSITVGVDG

TLYKTHPHFSSIMQETLQDLAPQCQVTFHKSEDGSGKGAALITAVACRIKSEGQH
>anc131_altAll|avgPP:0.995|lnPP:-8.112|num_ambig:6|num_ambig_gaps:0
LHHLQLSDETLMDVSIRFRREMDKGLCRDTNPTAAVKMLPTFVRSTPDGTEQGEFLALDLGGSNFRVLLV
KVMANGKQEVEMENQIYDIPEHLMRGSGSELFDHIADCLANFLEKLGIKDKKLPLGFTFSFPCQQTKLDE
SVLVSWTKGFKSSGVEGKDVVSLLRKSIKKRGDFDIDIVAVINDTVGTMMTCGYDDHHCEIGLIVGTGTN
ACYMEQMRNLELLDGDEGRMCVNTEWGAFGDDGALEDLRTDIDREIDAGSLNPGKQLFEKMISGMYMGEL
VRLILVRMTKEQLLFQGKTTAQLLTTGSFNTSYIYAIENDKDEEGLASAEKVLRGLGLDPSVEDCIATQR
VCQIVSTRAAHLCAATLVAVLRQIRDNKAAEKLRTTIGVDGSVYKNHPEFSRRLHKMVRRLVPDCDVRFL
QSQCGSGKGAAMVTAVAYRLAAQHAERQRILDTLRLSREQLLEVKKRMSEEMVRGLSKQTHEQASVKMLP
TYVRSTPDGTEHGDFLALDLGGSSFRVLLVRVRSGKKHNVDMHHKIYSIPQETMQGTGEELFNHIVDCIA
DFLEYMGMRGASLPLGFTFSFPCHQSKLDQGILLKWTKGFKASGCEGQDVITLLKDAVRRRQEFDLNFVA
VVNDTVGTMMTCGYEDPKCEVGLIVGTGTNACYMEEMQNIELVEGDDGRMCVNMEWGAFGDNGELDDFCT
QFDRIVDDSSNNPGKQRYEKMISGMYLGEIVRNVLIDFTAKGLLFRGKLSERLKTRGIFETKFLSQIESD
RLAMRQVRSILQHLGLTSSTCDDSVLVKEVCSVVARRAAQLCGAGLAAVVDKIRQNRNLNQLSITVGVDG
TLYKTHPHFSSIMQETLQDLAPQCQVTFHKSEDGSGKGAALITAVACRIKSEGQH
>anc132|avgPP:0.990|lnPP:-15.294|num_ambig:12|num_ambig_gaps:0
LHHFQLSDQTVMDVSMRFRREMDKGLCRDTNPTAAVKMLPTFVRSTPDGTEQGEFLALDLGGSNFRVLLV
KVMGNGEQKVEMENQIYEIPEHIMRGSGSELFDHVADCLANFLEKMGIKDKKLPLGFTFSFPCQQTKLDE
SVLLSWTKGFRSSGVEGKDVVSLLRKSIKKRGDFDIDIVAVINDTVGTMMTCGFDDHHCEIGLIVGTGTN
ACYMEHMRNLELLDGDEGRMCVNTEWGAFGDDGALEDLRTDIDREIDAGSLNPGKQLFEKMISGMYMGEL
VRLILVKMAREQLLFQGKTTPELLTTGRFSTSYIYSIDNDKEEEGLASAEKVLRGLGLDPSAEDCIATRR
VCQIVSTRAAHLCAATLAALLRQIRDNKAAEKLRTTIGVDGSVYKNHPEFSRRLNKMVRRLVPDCDVRFL
LSQDGSSKGAAMVTAVAFRLAAQQAERQRILDTLRLSREQLLEVKRRMSEEMLRGLSKQTHEQASVKMLP
TYVRSTPDGTEHGDFLALDLGGSSFRVLLVRVRSGKRHNVDMHHKIYSIPQETMQGTGEELFSHIVNCIA
DFLEYMGMRGASLPLGFTFSFPCHQSKLDQGILLKWTKGFKASGCEGQDVVTLLKEAVRRKQEFDLNFVA
VVNDTVGTMMTCGYEDPKCEVGLIVGTGTNACYMEEMHNIELVEGDDGRMCVNVEWGAFGDNGELDDFCT
QFDRAVDDCSNYPGKQRYEKMISGMYLGEIVRNVLMDFTAKGLLFRGKVSERLKTRGIFETKFLSQIESD
RLAMRQVRSILQHLGLTGSTCDDSVLVKEVCAVVARRAAQLCGAGLAAVVDKIRQNRNLNQLSVTVGVDG
TLYKTHPHFSSIMQETLQDLAPQCQVTFHKSEDGSGKGAALITAVACRMKSEGQH
>anc132_altAll|avgPP:0.988|lnPP:-21.609|num_ambig:12|num_ambig_gaps:0
LHNFQLSDQTIMDVSMRFRREMDKGLCRDTNPTAAVKMLPTFVRSTPDGTEQGEFLALDLGGSNFRVLLV
KVMGNGEQKVEMENQIYDIPEHIMRGSGSELFDHVADCLANFLEKMGIKDKKLPLGFTFSFPCQQTKLDE
SVLLSWTKGFRSSGVEGKDVVSLLRKSIKKRGDFDIDIVAVINDTVGTMMTCGFDDHHCEIGLIVGTGTN
ACYMEHMRNLELLDGDEGRMCVNTEWGAFGDDGALEDLRTDIDREIDAGSLNPGKQLFEKMISGMYMGEL
VRLILVKMAREQLLFQGKTTPALLTTGRFSTSYIYAIDNDKDEEGLASAEKVLRGLGLDPSAEDCIATRR
VCQIVSTRAAHLCAATLAALLRQIRDNKAAEKLRTTIGVDGSVYKNHAEFSRRLNKMVRRLVPDCDVRFL
LSQDGSSKGAAMVTAVAFRLAAQQAERQRILDTLRLSREQLLEVKRRMSEEMLRGLSKQTHEQTSVKMLP
TYVRSTPDGTEHGDFLALDLGGSSFRVLLVRVRSGKRHNVDMHHKIYSIPQETMQGTGEELFSHIVDCIA
DFLEYMGMRGASLPLGFTFSFPCHQSKLDQGILLKWTKGFKASGCEGQDVVTLLKEAVRRKQEFDLNFVA
VVNDTVGTMMTCGYEDPRCEVGLIVGTGTNACYMEEMHNIELVEGDDGRMCVNVEWGAFGDNGELDDFCT
QFDRAVDDCSNYPGKQRYEKMISGMYLGEIVRNVLLDFTAKGLLFRGKVSERLKTRGIFETKFLSQIESD
RLAMRQVRSILQHLGLTGSTCDDSVLVKEVCAVVARRAAQLCGAGLAAVVDKIRQNRNLNQLSVTVGVDG
TLYKTHPHFSSIMQETLQDLAPQCQVTFHKSEDGSGKGAALITAVACRMKSEGQR
>anc133|avgPP:0.997|lnPP:-5.031|num_ambig:4|num_ambig_gaps:0
LHHLQLSDETLMDVSMRFRREMDKGLCRDTNPTAAVKMLPTFVRSTPDGTEQGEFLALDLGGSNFRVLLV
KVMANGEQKVEMENQIYAIPEHIMRGSGSELFDHVADCLANFLEKMGIKDKKLPLGFTFSFPCQQTKLDE
SVLLSWTKGFRSSGVEGKDVVSLLRKSIKKRGDFDIDIVAVVNDTVGTMMTCGYDDHHCEIGLIVGTGTN
ACYMEQMRNLELLDGDEGRMCVNTEWGAFGDDGALEDLRTDIDREIDAGSLNPGKQLFEKMISGMYMGEL
VRLILVKMAREQLLFQGKTTPELLTTGRFSTSYIYAIENDKDEEGLASAEKVLRGLGLDPSVEDCIATRR
VCQIVSTRAAHLCAATLAAVLRQIRDNKAAEKLRTTIGVDGSVYKNHPEFSRRLNKMVRRLVPDCDVRFL
LSQDGSGKGAAMVTAVAYRLAAQHAERQRILDTLRLSREQLLEVKRRMSEEMVRGLSKQTHEQASVKMLP
TYVRSTPDGTEHGDFLALDLGGSSFRVLLVRVRSGKRHNVDMHHKIYSIPQETMQGTGEELFNHIVDCIA

DFLEYMGMRGASLPLGFTFSFPCHQSKLDQGILLKWTKGFKASGCEGQDVVTLLKEAVRRKQEFDLNFVA
VVNDTVGTMMTCGYEDPKCEVGLIVGTGTNACYMEEMHNIELVEGDDGRMCVNVEWGAFGDNGELDDFCT
QFDRIVDDCSNYPGKQRYEKMISGMYLGEIVRNVLLDFTAKGLLFRGKVSERLKTRGIFETKFLSQIESD
RLAMRQVRSILQHLGLTGSTCDDSVLVKEVCSVVARRAAQLCGAGLAAVVDKIRQNRNLNQLSITVGVDG
TLYKTHPHFSSIMQETLQDLAPQCQVTFHKSEDGSGKGAALITAVACRIKSEGQH
>anc133_altAll|avgPP:0.996|lnPP:-7.711|num_ambig:4|num_ambig_gaps:0
LHHLQLSDETLMDVSMRFRREMDKGLCRDTNPTAAVKMLPTFVRSTPDGTEQGEFLALDLGGSNFRVLLV
KVMGNGEQKVEMENQIYAIPEHIMRGSGSELFDHIADCLANFLEKMGIKDKKLPLGFTFSFPCQQTKLDE
SVLLSWTKGFRSSGVEGKDVVSLLRKSIKKRGDFDIDIVAVVNDTVGTMMTCGYDDHHCEIGLIVGTGTN
ACYMEQMRNLELLDGDEGRMCVNTEWGAFGDDGALEDLRTDIDREIDAGSLNPGKQLFEKMISGMYMGEL
VRLILVKMAREQLLFQGKTTPELLTTGRFSTSYIYSIENDKDEEGLASAEKVLRGLGLDPSVEDCIATRR
VCQIVSTRAAHLCAATLAAVLRQIRDNKAAEKLRTTIGVDGSVYKNHPEFSRRLNKMVRRLVPDCDVRFL
LSQDGSGKGAAMVTAVAYRLAAQHAERQRILDTLRLSREQLLEVKRRMSEEMVRGLSKQTHEQASVKMLP
TYVRSTPDGTEHGDFLALDLGGSSFRVLLVRVRSGKRHNVDMHHKIYSIPQETMQGTGEELFNHIVDCIA
DFLEYMGMRGASLPLGFTFSFPCHQSKLDQGILLKWTKGFKASGCEGQDVVTLLKEAVRRKQEFDLNFVA
VVNDTVGTMMTCGYEDPKCEVGLIVGTGTNACYMEEMHNIELVEGDDGRMCVNVEWGAFGDNGELDDFCT
QFDRVVDDCSNYPGKQRYEKMISGMYLGEIVRNVLLDFTAKGLLFRGKVSERLKTRGIFETKFLSQIESD
RLAMRQVRSILQHLGLTGSTCDDSVLVKEVCSVVARRAAQLCGAGLAAVVDKIRQNRNLNQLSITVGVDG
TLYKTHPHFSSIMQETLQDLAPQCQVTFHKSEDGSGKGAALITAVACRIKSEGQH
>anc134|avgPP:0.991|lnPP:-12.967|num_ambig:9|num_ambig_gaps:0
LHQFKLSDQTLMDLSMRFRREMDKGLCRDTNPTAAVKMLPTFVRSTPDGTEQGEFLALDLGGSNFRVLLV
KVKANGEQKVEMENQIYAIPEHIMRGSGSELFDHIVDCLANFMEMMGIKDKKLPLGFTFSFPCQQTKLDE
SVLLSWTKGFKLSGVEGKDVVSLLRKSIKKRGDFDIDVVAVVNDTVGTMMTCGYDDHHCEIGLIVGTGTN
ACYMEQMRNLDLLDGDEGRMCVNTEWGAFGDDGALEDLRTDIDREIDAGSLNPGKQLFEKMISGMYMGEL
VRLILVKMAREQLLFQGKTTPELLTTGRFSTSYIYTIENDKDEEGLASAEKVLRGLGLDPSVEDCIATRR
VCQIVSTRAAHLCAATLAAVLRQIRDNKAAEKLRTTIGVDGSVYKNHPEFSRRLNKMVRRLVPDCDVRFL
QSQDGSGKGAAMVTAVAYRLAAQHAERQRILDTLRLSREQLLEVKRRMTEEMVRGLSKQTHEQTSVKMLP
TYVRSTPDGTEHGDFLALDLGGSSFRVLLVRLRSGKRHKVDMHQKIYSIPQETMQGTGEELFDHIVDCIA
DFLEYMGMRGASLPLGFTFSFPCHQSKLDEGILLKWTKGFKASGCEGRDVVTLLKEAVRRKQEFDLNFVA
VVNDTVGTMMTCGYEDPKCEVGLIVGTGTNACYMEEMHNIELVEGNDGQMCVNVEWGAFGDNGELDDFCT
QFDRIVDDCSNYPGKQRYEKMISGMYLGEIVRNVLLDFTAKGLLFRGKMSERLKTRGIFETKFLSQIESD
RLAMRQVRSILQHLGLTGSTCDDSVLVKEVCSVVARRAAQLCGAGLAAVVDKIRQNRNLNQLSITVGVDG
TLYKTHPHFSSIMQETLQDLAPQCQVTFLKSEDGSGKGAALITAVACRINSEGQ
>anc134_altAll|avgPP:0.990|lnPP:-17.145|num_ambig:9|num_ambig_gaps:0
LHQFKLSDETLMDLSMRFRREMDKGLCRDTNPTAAVKMLPTFVRSTPDGTEQGEFLALDLGGSNFRVLLV
KVKANGEQKVEMENQIYAIPEHVMRGSGSELFDHIVDCLANFMEMMGIKDKKLPLGFTFSFPCQQTKLDE
SVLLSWTKGFKLSGVEGKDVVSLLRKSIKKRGDFDIDVVAVVNDTVGTMMTCGYDDHHCEIGLIVGTGTN
ACYMEQMRNLDLLDGDEGRMCVNTEWGAFGDDGALEDLRTDIDREIDAGSLNPGKQLFEKMISGMYLGEL
VRLILVKMAREQLLFQGKTTPELLTTGRFSTSYIYTIENDKDEEGLASAEKVLRGLGLEPSVEDCIATRR
VCQIVSTRAAHLCAATLAAVLRQIRDNKAAEKLRTTIGVDGSVYKNHPEFSRRLNKMVRRLVPDCDVRFL
QSQDGSGKGAAMVTAVAYRLAAQHAERQRILDTLRLSREQLLEVKRRMTEGMVRGLSKQTHEQTSVKMLP
TYVRSTPDGTEHGDFLALDLGGSSFRVLLVRLRSGKRHKVDMHQKIYSIPQETMQGTGEELFDHIVDCIA
DFLEYMGMRGASLPLGFTFSFPCHQSKLDEGILLKWTKGFKASGCEGKDVVTLLKEAVRRKQEFDLNFVA
VVNDTVGTMMTCGYEDPKCEVGLIVGTGTNACYMEEMHNIEVVEGNDGQMCVNVEWGAFGDNGELDDFCT
QFDRAVDDCSNYPGKQRYEKMISGMYLGEIVRNVLLDFTSKGLLFRGKMSERLKTRGIFETKFLSQIESD
RLAMRQVRSILQHLGLTGSTCDDSVLVKEVCSVVARRAAQLCGAGLAAVVDKIRQNRNLNQLSITVGVDG
TLYKTHPHFSSIMQETLQDLAPQCQVTFLKSEDGSGKGAALITAVACRINSEGQ
>anc135|avgPP:0.995|lnPP:-7.468|num_ambig:3|num_ambig_gaps:0
LQKLKLSDETLMDLSVRFRREMDKGLCRDTNPTAAVKMLPTFVRSTPDGTEQGEFLALDLGGSNFRVLLV
KVKGNGEQKVEMENQIYDIPEHVMRGSGSELFDHIADCLANFLEKMGIKDKKLPLGFTFSFPCQQTKLDE
SVLIYWTKGFRLSGVEGKEVVSLLRKSIKKRGDFDVDILAVINDTVGTMMTCGYDDHHCEIGLIVGTGTN
ACYMEQMRNLDLLDGDEGRMCVNTEWGAFGDDGALEDLRTDIDREIDAGSLNPGKQLFEKMISGMYMGEL

VRLILVKMARDQLLFQGKTTPELLTTGRFSTSYIYTIETDKDEEGLANAEKVLRGLGLDPSAEDCIATRR
VCQIVSTRAAHLCAATLAAVLRQIRDNKAAEKLRTTIGVDGSVYKNHPEFSRRLHKMVRRLVPDCDVRFL
QSQCGSGKGAAMVTAVAYRLAAQQAERQRILDTLRLSREQLLEVKRRMTEGMVLGLSKQTHEQTSVKMLP
TYVRSTPDGTEHGDFLALDLGGSSFRVLLVRLRSGKRHKVDMHQKIYSIPQETMQGTGEELFDHIVHCIA
DFLEYMGMRGASLPLGFTFSFPCHQSKLDEGILLKWTKGFKASDCEGKDVVMLLKEAVRRKQEFDLNFVA
VVNDTVGTMMTCGYEDPKCEVGLIVGTGTNACYMEEMHNIELVEGNEGRMCVNVEWGAFGDNGELDDFCT
QFDRAVDDCSNYPGKQRYEKMISGMYLGEIVRNVLLDFTSKGLLFRGKMSERLKTRGIFETKFLSQIESD
RLAMRQVRSILQHLGLTGSTCDDSVLVKEVCSVVARRAAQLCGAGLAAVVDRIRQNRNLNQLSITVGVDG
TLYKTHPHFSSIMQETLQDLAPQCQVTFLKSEDGSGKGAALITAVACRVNSEGQH
>anc135_altAll|avgPP:0.994|lnPP:-9.400|num_ambig:3|num_ambig_gaps:0
LQKLKLSDETLMDLSVRFRREMDKGLCRDTNPTAAVKMLPTFVRSTPDGTEQGEFLALDLGGSNFRVLLV
KVKGNGEQKVEMENQIYDIPEHVMRGSGSELFDHIADCLANFLEKMGIKDKKLPLGFTFSFPCQQTKLDE
SVLIYWTKGFRLSGVEGKEVVSLLRKSIKKRGDFDVDILAVINDTVGTMMTCGYDDHHCEIGLIVGTGTN
ACYMEQMRNLELLDGDEGRMCVNTEWGAFGDDGALEDLRTDIDREIDAGSLNPGKQLFEKMISGMYMGEL
VRLILVKMARDQLLFQGKTTPELLTTGRFSTSYIYTIETDKDEEGLANAEKVLRGLGLDPSAEDCIATRR
VCQIVSTRAAHLCAATLAAVLRQIRDNKAAEKLRTTIGVDGSVYKNHPEFSRRLHKMVRRLVPDCDVRFL
QSQCGSGKGAAMVTAVAYRLAAQQAERQRILDTLRLSREQLLEVKRRMTEGMVLGLSKQTHEQTSVKMLP
TYVRSTPDGTEHGDFLALDLGGSSFRVLLVRLRSGKRHKVDMHQKIYSIPQETMQGTGEELFDHIVQCIA
DFLEYMGMRGASLPLGFTFSFPCHQSKLDEGILLKWTKGFKASDCEGKDVVMLLKEAVRRKQEFDLNFVA
VVNDTVGTMMTCGYEDPKCEVGLIVGTGTNACYMEEMHNIELVEGNEGRMCVNVEWGAFGDNGELDDFCT
QFDRAVDDCSNYPGKQRYEKMISGMYLGEIVRNVLLDFTSKGLLFRGKMSERLKTRGIFETKFLSQIESD
RLAMRQVRSILQHLGLTGSTCDDSVLVKEVCSVVARRAAQLCGAGLAAVVDRIRQNRNLNQLSITVGVDG
TLYKTHPHFSSIMQETLQDLAPQCQVTFLKSEDGSGKGAALITAVACRINSEGQH
>anc136|avgPP:0.994|lnPP:-8.351|num_ambig:6|num_ambig_gaps:0
LHQLKLSDETLMDLSMRFRREMDKGLCRDTNPTAAVKMLPTFVRSTPDGTEQGEFLALDLGGSNFRVLLV
KVKANGEQKVEMENQIYAIPEHIMRGSGSELFDHIADCLANFLEKMGIKDKKLPLGFTFSFPCQQTKLDE
SVLLSWTKGFRLSGVEGKDVVSLLRKSIKKRGDFDIDIVAVVNDTVGTMMTCGYDDHHCEIGLIVGTGTN
ACYMEQMRNLDLLDGDEGRMCVNTEWGAFGDDGALEDLRTDIDREIDAGSLNPGKQLFEKMISGMYMGEL
VRLILVKMAREQLLFQGKTTPELLTTGRFSTSYIYTIENDKDEEGLASAEKVLRGLGLDPSVEDCIATRR
VCQIVSTRAAHLCAATLAAVLRQIRDNKAAEKLRTTIGVDGSVYKNHPEFSRRLNKMVRRLVPDCDVRFL
QSQDGSGKGAAMVTAVAYRLAAQHAERQRILDTLRLSREQLLEVKRRMTEEMVRGLSKQTHEQTSVKMLP
TYVRSTPDGTEHGDFLALDLGGSSFRVLLVRLRSGKRHKVDMHQKIYSIPQETMQGTGEELFDHIVDCIA
DFLEYMGMRGASLPLGFTFSFPCHQSKLDEGILLKWTKGFKASGCEGKDVVTLLKEAVRRKQEFDLNFVA
VVNDTVGTMMTCGYEDPKCEVGLIVGTGTNACYMEEMHNIELVEGNDGRMCVNVEWGAFGDNGELDDFCT
QFDRIVDDCSNYPGKQRYEKMISGMYLGEIVRNVLLDFTAKGLLFRGKMSERLKTRGIFETKFLSQIESD
RLAMRQVRSILQHLGLTGSTCDDSVLVKEVCSVVARRAAQLCGAGLAAVVDKIRQNRNLNQLSITVGVDG
TLYKTHPHFSSIMQETLQDLAPQCQVTFLKSEDGSGKGAALITAVACRINSEGQH
>anc136_altAll|avgPP:0.993|lnPP:-11.353|num_ambig:6|num_ambig_gaps:0
LHQLKLSDETLMDLSMRFRREMDKGLCRDTNPTAAVKMLPTFVRSTPDGTEQGEFLALDLGGSNFRVLLV
KVKGNGEQKVEMENQIYAIPEHVMRGSGSELFDHIADCLANFLEKMGIKDKKLPLGFTFSFPCQQTKLDE
SVLLSWTKGFRLSGVEGKDVVSLLRKSIKKRGDFDIDIVAVVNDTVGTMMTCGYDDHHCEIGLIVGTGTN
ACYMEQMRNLELLDGDEGRMCVNTEWGAFGDDGALEDLRTDIDREIDAGSLNPGKQLFEKMISGMYMGEL
VRLILVKMAREQLLFQGKTTPELLTTGRFSTSYIYTIENDKDEEGLASAEKVLRGLGLDPSVEDCIATRR
VCQIVSTRAAHLCAATLAAVLRQIRDNKAAEKLRTTIGVDGSVYKNHPEFSRRLNKMVRRLVPDCDVRFL
QSQDGSGKGAAMVTAVAYRLAAQHAERQRILDTLRLSREQLLEVKRRMTEGMVRGLSKQTHEQTSVKMLP
TYVRSTPDGTEHGDFLALDLGGSSFRVLLVRLRSGKRHKVDMHQKIYSIPQETMQGTGEELFDHIVDCIA
DFLEYMGMRGASLPLGFTFSFPCHQSKLDEGILLKWTKGFKASGCEGKDVVTLLKEAVRRKQEFDLNFVA
VVNDTVGTMMTCGYEDPKCEVGLIVGTGTNACYMEEMHNIELVEGNDGRMCVNVEWGAFGDNGELDDFCT
QFDRAVDDCSNYPGKQRYEKMISGMYLGEIVRNVLLDFTSKGLLFRGKMSERLKTRGIFETKFLSQIESD
RLAMRQVRSILQHLGLTGSTCDDSVLVKEVCSVVARRAAQLCGAGLAAVVDKIRQNRNLNQLSITVGVDG
TLYKTHPHFSSIMQETLQDLAPQCQVTFLKSEDGSGKGAALITAVACRINSEGQH
>anc137|avgPP:0.996|lnPP:-5.071|num_ambig:2|num_ambig_gaps:0

LHHLQLSDETLMDVSMRFRREMDKGLCRDTNPTAAVKMLPTFVRSTPDGTEQGEFLALDLGGSNFRVLLV
KVMANGEQKVEMENQIYAIPEHIMRGSGSELFDHIADCLANFLEKMGIKDKKLPLGFTFSFPCQQTKLDE
SVLLSWTKGFRSSGVEGKDVVSLLRKSIKKRGDFDIDIVAVVNDTVGTMMTCGYDDHHCEIGLIVGTGTN
ACYMEQMRNLELLDGDEGRMCVNTEWGAFGDDGALEDLRTDIDREIDAGSLNPGKQLFEKMISGMYMGEL
VRLILVKMAREQLLFQGKTTPELLTTGRFSTSYIYAIENDKDEEGLASAEKVLRGLGLDPSVEDCIATRR
VCQIVSTRAAHLCAATLAAVLRQIRDNKAAEKLRTTIGVDGSVYKNHPEFSRRLNKMVRRLVPDCDVRFL
QSQDGSGKGAAMVTAVAYRLAAQHAERQRILDTLRLSREQLLEVKRRMSEEMVRGLSKQTHEQASVKMLP
TYVRSTPDGTEHGDFLALDLGGSSFRVLLVRVRSGKRHNVDMHQKIYSIPQETMQGTGEELFNHIVDCIA
DFLEYMGMRGASLPLGFTFSFPCHQSKLDEGILLKWTKGFKASGCEGKDVVTLLKEAVRRKQEFDLNFVA
VVNDTVGTMMTCGYEDPKCEVGLIVGTGTNACYMEEMHNIELVEGDDGRMCVNVEWGAFGDNGELDDFCT
QFDRIVDDCSNYPGKQRYEKMISGMYLGEIVRNVLLDFTAKGLLFRGKVSERLKTRGIFETKFLSQIESD
RLAMRQVRSILQHLGLTGSTCDDSVLVKEVCSVVARRAAQLCGAGLAAVVDKIRQNRNLNQLSITVGVDG
TLYKTHPHFSSIMQETLQDLAPQCQVTFLKSEDGSGKGAALITAVACRIKSEGQH
>anc137_altAll|avgPP:0.996|lnPP:-6.452|num_ambig:2|num_ambig_gaps:0
LHHLQLSDETLMDVSMRFRREMDKGLCRDTNPTAAVKMLPTFVRSTPDGTEQGEFLALDLGGSNFRVLLV
KVMGNGEQKVEMENQIYAIPEHIMRGSGSELFDHIADCLANFLEKMGIKDKKLPLGFTFSFPCQQTKLDE
SVLLSWTKGFRSSGVEGKDVVSLLRKSIKKRGDFDIDIVAVVNDTVGTMMTCGYDDHHCEIGLIVGTGTN
ACYMEQMRNLELLDGDEGRMCVNTEWGAFGDDGALEDLRTDIDREIDAGSLNPGKQLFEKMISGMYMGEL
VRLILVKMAREQLLFQGKTTPELLTTGRFSTSYIYAIENDKDEEGLASAEKVLRGLGLDPSVEDCIATRR
VCQIVSTRAAHLCAATLAAVLRQIRDNKAAEKLRTTIGVDGSVYKNHPEFSRRLNKMVRRLVPDCDVRFL
QSQDGSGKGAAMVTAVAYRLAAQHAERQRILDTLRLSREQLLEVKRRMSEEMVRGLSKQTHEQASVKMLP
TYVRSTPDGTEHGDFLALDLGGSSFRVLLVRVRSGKRHNVDMHQKIYSIPQETMQGTGEELFNHIVDCIA
DFLEYMGMRGASLPLGFTFSFPCHQSKLDEGILLKWTKGFKASGCEGKDVVTLLKEAVRRKQEFDLNFVA
VVNDTVGTMMTCGYEDPKCEVGLIVGTGTNACYMEEMHNIELVEGDDGRMCVNVEWGAFGDNGELDDFCT
QFDRVVDDCSNYPGKQRYEKMISGMYLGEIVRNVLLDFTAKGLLFRGKVSERLKTRGIFETKFLSQIESD
RLAMRQVRSILQHLGLTGSTCDDSVLVKEVCSVVARRAAQLCGAGLAAVVDKIRQNRNLNQLSITVGVDG
TLYKTHPHFSSIMQETLQDLAPQCQVTFLKSEDGSGKGAALITAVACRIKSEGQH
>anc138|avgPP:0.997|lnPP:-4.485|num_ambig:2|num_ambig_gaps:0
LHHLQLSDETLMDVSMRFRREMDKGLCRDTNPTAAVKMLPTFVRSTPDGTEQGEFLALDLGGSNFRVLLV
KVMANGEQKVEMENQIYAIPEHIMRGSGSELFDHIADCLANFLEKMGIKDKKLPLGFTFSFPCQQTKLDE
SVLLSWTKGFRSSGVEGKDVVSLLRKSIKKRGDFDIDIVAVVNDTVGTMMTCGYDDHHCEIGLIVGTGTN
ACYMEQMRNLELLDGDEGRMCVNTEWGAFGDDGALEDLRTDIDREIDAGSLNPGKQLFEKMISGMYMGEL
VRLILVKMAREQLLFQGKTTPELLTTGRFSTSYIYAIENDKDEEGLASAEKVLRGLGLDPSVEDCIATRR
VCQIVSTRAAHLCAATLAAVLRQIRDNKAAEKLRTTIGVDGSVYKNHPEFSRRLNKMVRRLVPDCDVRFL
LSQDGSGKGAAMVTAVAYRLAAQHAERQRILDTLRLSREQLLEVKRRMSEEMVRGLSKQTHEQASVKMLP
TYVRSTPDGTEHGDFLALDLGGSSFRVLLVRVRSGKRHNVDMHHKIYSIPQETMQGTGEELFNHIVDCIA
DFLEYMGMRGASLPLGFTFSFPCHQSKLDQGILLKWTKGFKASGCEGQDVVTLLKEAVRRKQEFDLNFVA
VVNDTVGTMMTCGYEDPKCEVGLIVGTGTNACYMEEMHNIELVEGDDGRMCVNVEWGAFGDNGELDDFCT
QFDRIVDDCSNYPGKQRYEKMISGMYLGEIVRNVLLDFTAKGLLFRGKVSERLKTRGIFETKFLSQIESD
RLAMRQVRSILQHLGLTGSTCDDSVLVKEVCSVVARRAAQLCGAGLAAVVDKIRQNRNLNQLSITVGVDG
TLYKTHPHFSSIMQETLQDLAPQCQVTFHKSEDGSGKGAALITAVACRIKSEGQH
>anc138_altAll|avgPP:0.996|lnPP:-6.117|num_ambig:2|num_ambig_gaps:0
LHHLQLSDETLMDVSMRFRREMDKGLCRDTNPTAAVKMLPTFVRSTPDGTEQGEFLALDLGGSNFRVLLV
KVMGNGEQKVEMENQIYAIPEHIMRGSGSELFDHIADCLANFLEKMGIKDKKLPLGFTFSFPCQQTKLDE
SVLLSWTKGFRSSGVEGKDVVSLLRKSIKKRGDFDIDIVAVVNDTVGTMMTCGYDDHHCEIGLIVGTGTN
ACYMEQMRNLELLDGDEGRMCVNTEWGAFGDDGALEDLRTDIDREIDAGSLNPGKQLFEKMISGMYMGEL
VRLILVKMAREQLLFQGKTTPELLTTGRFSTSYIYAIENDKDEEGLASAEKVLRGLGLDPSVEDCIATRR
VCQIVSTRAAHLCAATLAAVLRQIRDNKAAEKLRTTIGVDGSVYKNHPEFSRRLNKMVRRLVPDCDVRFL
LSQDGSGKGAAMVTAVAYRLAAQHAERQRILDTLRLSREQLLEVKRRMSEEMVRGLSKQTHEQASVKMLP
TYVRSTPDGTEHGDFLALDLGGSSFRVLLVRVRSGKRHNVDMHHKIYSIPQETMQGTGEELFNHIVDCIA
DFLEYMGMRGASLPLGFTFSFPCHQSKLDQGILLKWTKGFKASGCEGQDVVTLLKEAVRRKQEFDLNFVA
VVNDTVGTMMTCGYEDPKCEVGLIVGTGTNACYMEEMHNIELVEGDDGRMCVNVEWGAFGDNGELDDFCT

QFDRVVDDCSNYPGKQRYEKMISGMYLGEIVRNVLLDFTAKGLLFRGKVSERLKTRGIFETKFLSQIESD
RLAMRQVRSILQHLGLTGSTCDDSVLVKEVCSVVARRAAQLCGAGLAAVVDKIRQNRNLNQLSITVGVDG
TLYKTHPHFSSIMQETLQDLAPQCQVTFHKSEDGSGKGAALITAVACRIKSEGQH
>anc139|avgPP:0.996|lnPP:-6.596|num_ambig:7|num_ambig_gaps:0
LHRLQLSDETLMDLSIRFRREMDKGLCRDTNPTAAVKMLPTFVRSTPDGTEQGEFLALDLGGSNFRVLLV
KVMADGEQKVEMENQIYAIPEHLMRGSGSELFDHIADCLSNFLEKMGIKDKKLPLGFTFSFPCQQTKLDE
SVLVSWTKGFKSSGVEGKDVVSLLRKSIKKRGDFDIDIVAVVNDTVGTMMTCGYDDQHCEIGLIVGTGTN
ACYMEQMRNLELLDGDEGRMCVNTEWGAFGDDGALEDLRTDIDRELDAGSLNPGKQLFEKMISGMYMGEL
VRLILVRMTKEQLLFQGKTTAQLLTTGSFSTSYIYAIENDKGEEGLASAEKVLRGLGLEPSVEDCIATQR
VCQIVSTRAAHLCAATLVAVLRQIRDNKAAEKLRTTIGVDGSVYKNHPEFSRRLHKMVRRLVPDCDVRFL
LSQDGSGKGSAMVTAVAYRLAAQHAERQRILDTLRLSREQLLEVKKRMSEEMVRGLSKQTHEQASVKMLP
TYVRSTPDGTEHGDFLALDLGGSSFRVLLVRVRSGKRHNVDMHHKIYSIPQETMQGTGEELFNHIVDCIA
DFLEYMGMRGASLPLGFTFSFPCHQSKLDQGILLKWTKGFKASGCEGQDVVTLLKDAVRRRQEFDLNFVA
VVNDTVGTMMTCGYEDPKCEVGLIVGTGTNACYMEEMHNIELVEGDDGRMCVNMEWGAFGDNGELDDFCT
QFDRIVDDCSNNPGKQRYEKMISGMYLGEIVRNVLLDFTAKGLLFRGKLSERLKTHGIFETKFLSQIEKD
RLAMRQIRSILQHLGLTSSTCDDSVLVKEVCSVVARRAAQLCGAGLAAVVDKIRQNRNLNQLSITVGVDG
TLYKTHPHFSSIMQETLQDLAPQCQVTFHKSEDGSGKGAALITAVACRIKSEGQH
>anc139_altAll|avgPP:0.994|lnPP:-10.706|num_ambig:7|num_ambig_gaps:0
LHRLQLSDETLMDLSIRFRREMDKGLCRDTNPTAAVKMLPTFVRSTPDGTEQGEFLALDLGGSNFRVLLV
KVMADGEQKVEMENQIYAIPEHLMRGSGSELFDHIADCLSNFLEKMGIKDKKLPLGFTFSFPCQQTKLDE
SVLVSWTKGFKSSGVEGKDVVSLLRKSIKKRGDFDIDIVAVVNDTVGTMMTCGYDDQHCEIGLIVGTGTN
ACYMEQMRNLELLDGDEGRMCVNTEWGAFGDDGALEDLRTDIDRELDAGSLNPGKQLFEKMISGMYMGEL
VRLILVRMTKEQLLFQGKTTAQLLTTGSFNTSYIYAIENDKDEEGLASAEKVLRGLGLEPSVEDCIATQR
VCQIVSTRAAHLCAATLVAVLRQIRDNKAAEKLRTTIGVDGSVYKNHPEFSRRLHKMVRRLVPDCDVRFL
LSQDGSGKGSAMVTAVAYRLAAQHAERQRILDTLRLSREQLLEVKKRMSEEMARGLSKQTHEQASVKMLP
TYVRSTPDGTEHGDFLALDLGGSSFRVLLVRVRSGKRHNVDMHHKIYSIPQETMQGTGEELFDHIVDCIA
DFLEYMGMRGASLPLGFTFSFPCHQSKLDQGILLKWTKGFKASGCEGQDVVTLLKDAVRRRQEFDLNFVA
VVNDTVGTMMTCGYEDPKCEVGLIVGTGTNACYMEEMHNIELVEGDDGCMCVNMEWGAFGDNGELDDFCT
QFDRVVDDCSNNPGKQRYEKMISGMYLGEIVRNVLMDFTAKGLLFRGKLSERLKTHGIFETKFLSQIEKD
RLAMRQIRSILQHLGLTSSTCDDSVLVKEVCSVVARRAAQLCGAGLAAVVDKIRQNRNLNQLSITVGVDG
TLYKTHPHFSSIMQETLQDLAPQCQVTFHKSEDGSGKGAALITAVACRIKSEGQH
>anc140|avgPP:0.998|lnPP:-3.507|num_ambig:4|num_ambig_gaps:0
LHHLQLSDETLMDVSIRFRREMDKGLCRDTNPTAAVKMLPTFVRSTPDGTEQGEFLALDLGGSNFRVLLV
KVMANGEQKVEMENQIYAIPEHLMRGSGSELFDHIADCLANFLEKMGIKDKKLPLGFTFSFPCQQTKLDE
SVLVSWTKGFKSSGVEGKDVVSLLRKSIKKRGDFDIDIVAVVNDTVGTMMTCGYDDHHCEIGLIVGTGTN
ACYMEQMRNLELLDGDEGRMCVNTEWGAFGDDGALEDLRTDIDREIDAGSLNPGKQLFEKMISGMYMGEL
VRLILVRMTKEQLLFQGKTTAQLLTTGSFSTSYIYAIENDKDEEGLASAEKVLRGLGLDPSVEDCIATQR
VCQIVSTRAAHLCAATLVAVLRQIRDNKAAEKLRTTIGVDGSVYKNHPEFSRRLHKMVRRLVPDCDVRFL
LSQDGSGKGAAMVTAVAYRLAAQHAERQRILDTLRLSREQLLEVKKRMSEEMVRGLSKQTHEQASVKMLP
TYVRSTPDGTEHGDFLALDLGGSSFRVLLVRVRSGKRHNVDMHHKIYSIPQETMQGTGEELFNHIVDCIA
DFLEYMGMRGASLPLGFTFSFPCHQSKLDQGILLKWTKGFKASGCEGQDVVTLLKDAVRRRQEFDLNFVA
VVNDTVGTMMTCGYEDPKCEVGLIVGTGTNACYMEEMHNIELVEGDDGRMCVNMEWGAFGDNGELDDFCT
QFDRIVDDCSNNPGKQRYEKMISGMYLGEIVRNVLLDFTAKGLLFRGKLSERLKTRGIFETKFLSQIESD
RLAMRQVRSILQHLGLTSSTCDDSVLVKEVCSVVARRAAQLCGAGLAAVVDKIRQNRNLNQLSITVGVDG
TLYKTHPHFSSIMQETLQDLAPQCQVTFHKSEDGSGKGAALITAVACRIKSEGQH
>anc140_altAll|avgPP:0.997|lnPP:-6.149|num_ambig:4|num_ambig_gaps:0
LHHLQLSDETLMDVSIRFRREMDKGLCRDTNPTAAVKMLPTFVRSTPDGTEQGEFLALDLGGSNFRVLLV
KVMANGEQKVEMENQIYAIPEHLMRGSGSELFDHIADCLANFLEKMGIKDKKLPLGFTFSFPCQQTKLDE
SVLVSWTKGFKSSGVEGKDVVSLLRKSIKKRGDFDIDIVAVVNDTVGTMMTCGYDDHHCEIGLIVGTGTN
ACYMEQMRNLELLDGDEGRMCVNTEWGAFGDDGALEDLRTDIDREIDAGSLNPGKQLFEKMISGMYMGEL
VRLILVRMTKEQLLFQGKTTAQLLTTGSFNTSYIYAIENDKDEEGLASAEKVLRGLGLDPSVEDCIATQR
VCQIVSTRAAHLCAATLVAVLRQIRDNKAAEKLRTTIGVDGSVYKNHPEFSRRLHKMVRRLVPDCDVRFL

LSQDGSGKGAAMVTAVAYRLAAQHAERQRILDTLRLSREQLLEVKKRMSEEMVRGLSKQTHEQASVKMLP
TYVRSTPDGTEHGDFLALDLGGSSFRVLLVRVRSGKRHNVDMHHKIYSIPQETMQGTGEELFDHIVDCIA
DFLEYMGMRGASLPLGFTFSFPCHQSKLDQGILLKWTKGFKASGCEGQDVVTLLKDAVRRRQEFDLNFVA
VVNDTVGTMMTCGYEDPKCEVGLIVGTGTNACYMEEMHNIELVEGDDGRMCVNMEWGAFGDNGELDDFCT
QFDRVVDDCSNNPGKQRYEKMISGMYLGEIVRNVLMDFTAKGLLFRGKLSERLKTRGIFETKFLSQIESD
RLAMRQVRSILQHLGLTSSTCDDSVLVKEVCSVVARRAAQLCGAGLAAVVDKIRQNRNLNQLSITVGVDG
TLYKTHPHFSSIMQETLQDLAPQCQVTFHKSEDGSGKGAALITAVACRIKSEGQH
>anc141|avgPP:0.998|lnPP:-2.850|num_ambig:3|num_ambig_gaps:0
LHHLQLSDETLMDVSIRFRREMDKGLCRDTNPTAAVKMLPTFVRSTPDGTEQGEFLALDLGGSNFRVLLV
KVMANGKQKVEMENQIYAIPEHLMRGSGSELFDHIADCLANFLEKLGIKDKKLPLGFTFSFPCQQTKLDE
SVLVSWTKGFKSSGVEGKDVVSLLRKSIKKRGDFDIDIVAVINDTVGTMMTCGYDDHHCEIGLIVGTGTN
ACYMEQMRNLELLDGDEGRMCVNTEWGAFGDDGALEDLRTDIDREIDAGSLNPGKQLFEKMISGMYMGEL
VRLILVRMTKEQLLFQGKTTAQLLTTGSFNTSYIYAIENDKDEEGLASAEKVLRGLGLDPSVEDCIATQR
VCQIVSTRAAHLCAATLVAVLRQIRDNKAAEKLRTTIGVDGSVYKNHPEFSRRLHKMVRRLVPDCDVRFL
LSQDGSGKGAAMVTAVAYRLAAQHAERQRILDTLRLSREQLLEVKKRMSEEMVRGLSKQTHEQASVKMLP
TYVRSTPDGTEHGDFLALDLGGSSFRVLLVRVRSGKRHNVDMHHKIYSIPQETMQGTGEELFDHIVDCIA
DFLEYMGMRGASLPLGFTFSFPCHQSKLDQGILLKWTKGFKASGCEGQDVVTLLKDAVRRRQEFDLNFVA
VVNDTVGTMMTCGYEDPKCEVGLIVGTGTNACYMEEMQNIELVEGDDGRMCVNMEWGAFGDNGELDDFCT
QFDRIVDDCSNNPGKQRYEKMISGMYLGEIVRNVLLDFTAKGLLFRGKLSERLKTRGIFETKFLSQIESD
RLAMRQVRSILQHLGLTSSTCDDSVLVKEVCSVVARRAAQLCGAGLAAVVDKIRQNRNLNQLSITVGVDG
TLYKTHPHFSSIMQETLQDLAPQCQVTFHKSEDGSGKGAALITAVACRIKSEGQH
>anc141_altAll|avgPP:0.998|lnPP:-4.500|num_ambig:3|num_ambig_gaps:0
LHHLQLSDETLMDVSIRFRREMDKGLCRDTNPTAAVKMLPTFVRSTPDGTEQGEFLALDLGGSNFRVLLV
KVMANGKQKVEMENQIYAIPEHLMRGSGSELFDHIADCLANFLEKLGIKDKKLPLGFTFSFPCQQTKLDE
SVLVSWTKGFKSSGVEGKDVVSLLRKSIKKRGDFDIDIVAVINDTVGTMMTCGYDDHHCEIGLIVGTGTN
ACYMEQMRNLELLDGDEGRMCVNTEWGAFGDDGALEDLRTDIDREIDAGSLNPGKQLFEKMISGMYMGEL
VRLILVRMTKEQLLFQGKTTAQLLTTGSFNTSYIYAIENDKDEEGLASAEKVLRGLGLDPSVEDCIATQR
VCQIVSTRAAHLCAATLVAVLRQIRDNKAAEKLRTTIGVDGSVYKNHPEFSRRLHKMVRRLVPDCDVRFL
LSQDGSGKGAAMVTAVAYRLAAQHAERQRILDTLRLSREQLLEVKKRMSEEMVRGLSKQTHEQASVKMLP
TYVRSTPDGTEHGDFLALDLGGSSFRVLLVRVRSGKRHNVDMHHKIYSIPQETMQGTGEELFNHIVDCIA
DFLEYMGMRGASLPLGFTFSFPCHQSKLDQGILLKWTKGFKASGCEGQDVVTLLKDAVRRRQEFDLNFVA
VVNDTVGTMMTCGYEDPKCEVGLIVGTGTNACYMEEMQNIELVEGDDGRMCVNMEWGAFGDNGELDDFCT
QFDRVVDDCSNNPGKQRYEKMISGMYLGEIVRNVLMDFTAKGLLFRGKLSERLKTRGIFETKFLSQIESD
RLAMRQVRSILQHLGLTSSTCDDSVLVKEVCSVVARRAAQLCGAGLAAVVDKIRQNRNLNQLSITVGVDG
TLYKTHPHFSSIMQETLQDLAPQCQVTFHKSEDGSGKGAALITAVACRIKSEGQH
>anc142|avgPP:0.985|lnPP:-24.512|num_ambig:17|num_ambig_gaps:0
LHHLQLSDETLMDISMRFRREMDKGLCRDTNPTAAVKMLPTYVLSTPDGTEQGEFLALDLGGSTFRVLLV
TVMANGEQKVDMENQIYAIPEHLMRGSGSELFDHIAECLANFLEKLGIKDKKLPLGFTFSFPCQQSKLDE
SVLISWTKAFRASGVEGKDVVSLLQRSIKKRGDINVDIMAVINDTVGAMMTCGYDDHHCEIGLIVGTGTN
ACYMEQMRNLEVLDGDEGQMCVNTEWGAFGDDGALEDLRTDFDREIDAGSLNPGKQLFEKMISGMYMGEL
VRLILVRMTKDQLLFQGKTTSQLLTTGKFTTNYIYSIESDNDEEGLASAQNVLRSLGLDPTLEDCIATQR
VCQIVSTRAAHLCAATLVAVLRQIRDNKAAEKLRTTIGVDGSVYKNHPEFSRRLHKMVRRLVPDCDVRFL
RSEVGSGKGAAMVTAVALRLAAQNAERQRILDTLRLNREQLLEVKKLISEEMVRGLSKQAHDQASIKMLP
TFVGSTPDGTEHGDFLALDLGGSSFRVLLVRVRSGKRHNVDMHHKIYSIPQETMQGTGEELFSHIVDCIA
NFLENMGMRGASLPLGFTFSFPCHQSKLDQGILLKWTKGFKASGCEGQDVVTLLKDAVRRRGEFDLSFVA
VVNDTVGTMMTCGYEDPKCEVGLIVGTGTNACYMEEMQNIEVVDGDDGRMCVNMEWGAFGENGELDDFLT
QFDRVVDASSNNPGKQRYEKMISGMYLGEIVRHVLMEFTAKGLLFRGKLSERLKTRGIFETKFLSQIESD
RLAMRQVRSILQHLGLTSSTCDDSVLVKEVCSVVSRRAAQLCGAGLAAVVDKIRQNRNMQQLSITVGVDG
TLYKMHPHFSRIMQETLQDLAPQCKVTFHKSEDGSGKGAALITAVACRIKSNGQNNSP
>anc142_altAll|avgPP:0.983|lnPP:-30.583|num_ambig:17|num_ambig_gaps:0
LHHLQLSDETLMDISMRFRREMDKGLCRDTNPTAAVKMLPTYVLSTPDGTEQGEFLALDLGGSTFRVLLV
TVMANGEQKVDMENQIYAIPEHLMRGSGSELFDHIAECLANFLEKLGIKDKKLPLGFTFSFPCQQSKLDE

```
SVLISWTKAFRASGVEGKDVVSLLQRSIKKRGDINVDIMAVINDTVGAMMTCGYDDHHCEIGLIVGTGTN
ACYMEQMRNLELLDGDEGQMCVNTEWGAFGDDGALEDLRTDFDREIDAGSLNPGKQLFEKMISGMYMGEL
VRLILVRMTKDQLLFQGKTTSQLLTTGKFTTKYIYSIESDNDEEGLMSAQNVLRALGLDPTLEDCIATQR
VCQIVSTRAAHLCAATLVAVLRQIRDNKAAEKLRTTIGVDGSVYKNHPEFPRRLHKMVRRLVPDCDVRFL
RSEVGSGKGAAMVTAVALRLAAQNAERQRILDTLRLNREQLLEVKRLVSEDMVRGLSKQTHDQASIKMLP
TFVGSTPDGTEHGDFLALDLGGSSFRVLLVRVRSGKRHNVDMHHKIYSIPQETMQGTGEELFSHIVDCIA
NFLENMGMRGASLPLGFTFSFPCHQSKLDQGILLKWTKGFKASGCEGQDVVTLLKDAVRRRGEFDLNFVA
VVNDTVGTMMTCGYEDPKCEVGLIVGTGTNACYMEEMQNIEVVDGDDGRMCVNMEWGAFGENGELDDFFT
EFDRVVDASSNNPGKQRYEKMISGMYLGEIVRHVLMEFTAKGLLFRGKLSERLKTRGIFETKFLSQIESD
RLAMRQVRCILQHLGLTSSTCDDSVLVKEVCSVVSRRAAQLCGAGLAAVVDKIRQNRNMQQLSITVGVDG
TLYKMHPHFSGIMQETLQDLAPQCKVTFHKSEDGSGKGAALITAVACRIKSNGQHHSG
>anc143|avgPP:0.990|lnPP:-14.846|num_ambig:12|num_ambig_gaps:0
LHHLQLSDETLMDISIRFRREMDKGLCRDTNPTAAVKMLPTFVLSTPDGTEQGEFLALDLGGSNFRVLLV
KVMANGKQKVEMENQIYAIPEHLMRGSGSELFDHIADCLANFLEKLGIKDKKLPLGFTFSFPCQQTKLDE
SVLVSWTKGFRSSGVEGKDVVSLLRKSIKKRGDFDVDIMAVINDTVGAMMTCGYDDHHCEIGLIVGTGTN
ACYMEQMRNLELLDGDEGRMCVNTEWGAFGDDGALEDLRTDFDREIDAGSLNPGKQLFEKMISGMYMGEL
VRLILVRMTKEQLLFQGKTTAQLLTTGSFNTKYIYAIESDKDEEGLASAEKVLRGLGLDPSLEDCIATQR
VCQIVSTRAAHLCAATLVAVLRQIRDNKAAEKLRTTIGVDGSVYKNHPEFSRRLHKMVRRLVPDCDVRFL
RSEVGSGKGAAMVTAVAYRLAAQNAERQRILDTLRLSREQLLEVKKRMSEEMVRGLSKQTHEQASVKMLP
TFVRSTPDGTEHGDFLALDLGGSSFRVLLVRVRSGKRHNVDMHHKIYSIPQETMQGTGEELFSHIVDCIA
NFLENMGMRGASLPLGFTFSFPCHQSKLDQGILLKWTKGFKASGCEGQDVVTLLKDAVRRRGEFDLNFVA
VVNDTVGTMMTCGYEDPKCEVGLIVGTGTNACYMEEMQNIEVVEGDDGRMCVNMEWGAFGDNGELDDFFT
QFDRVVDDSSNNPGKQRYEKMISGMYLGEIVRNVLMDFTAKGLLFRGKLSERLKTRGIFETKFLSQIESD
RLAMRQVRSILQHLGLTSSTCDDSVLVKEVCSVVSRRAAQLCGAGLAAVVDKIRQNRNLQQLSITVGVDG
TLYKTHPHFSSIMQETLQDLAPQCKVTFHKSEDGSGKGAALITAVACRIKSEGQH
>anc143_altAll|avgPP:0.988|lnPP:-20.920|num_ambig:12|num_ambig_gaps:0
LHHLQLSDETLMDVSMRFRREMDKGLCRDTNPTAAVKMLPTFVLSTPDGTEQGEFLALDLGGSNFRVLLV
KVMANGKQKVEMENQIYAIPEHLMRGSGSELFDHIADCLANFLEKLGIKDKKLPLGFTFSFPCQQSKLDE
SVLVSWTKGFRTSGVEGKDVVSLLRKSIKKRGDFDVDIMAVINDTVGAMMTCGYDDHHCEIGLIVGTGTN
ACYMEQMRNLEVLDGDEGRMCVNTEWGAFGDDGALEDLRTDFDREIDAGSLNPGKQLFEKMISGMYMGEL
VRLILVRMTKEQLLFQGKTTAQLLTTGSFNTSYIYAIENDKDEEGLASAEKVLRGLGLDPSVEDCIATQR
VCQIVSTRAAHLCAATLVAVLRQIRDNKAAEKLRTTIGVDGSVYKNHPEFPRRLHKMVRRLVPDCDVRFL
RSEVGSGKGAAMVTAVAYRLAAQNAERQRILDTLRLSREQLLEVKKRMSEEMVRGLSKQTHEQASVKMLP
TFVRSTPDGTEHGDFLALDLGGSSFRVLLVRVRSGKRHNVDMHHKIYSIPQETMQGTGEELFSHIVDCIA
NFLENMGMRGASLPLGFTFSFPCHQSKLDQGILLKWTKGFKASGCEGQDVVTLLKDAVRRRGEFDLNFVA
VVNDTVGTMMTCGYEDPKCEVGLIVGTGTNACYMEEMQNIEVVEGDDGRMCVNMEWGAFGDNGELDDFFT
QFDRIVDDCSNNPGKQRYEKMISGMYLGEIVRNVLMDFTAKGLLFRGKLSERLKTRGIFETKFLSQIESD
RLAMRQVRSILQHLGLTSSTCDDSVLVKEVCSVVSRRAAQLCGAGLAAVVDKIRQNRNLQQLSITVGVDG
TLYKTHPHFSRIMQETLQDLAPQCKVTFHKSEDGSGKGAALITAVACRIKSEGQH
>anc144|avgPP:0.996|lnPP:-5.276|num_ambig:4|num_ambig_gaps:0
LHHLQLSDETLMDVSIRFRREMDKGLCRDTNPTAAVKMLPTFVLSTPDGTEQGEFLALDLGGSNFRVLLV
KVMANGKQKVEMENQIYAIPEHLMRGSGSELFDHIADCLANFLEKLGIKDKKLPLGFTFSFPCQQTKLDE
SVLVSWTKGFRSSGVEGKDVVSLLRKSIKKRGDFDVDIMAVINDTVGAMMTCGYDDHHCEIGLIVGTGTN
ACYMEQMRNLELLDGDEGRMCVNTEWGAFGDDGALEDLRTDIDREIDAGSLNPGKQLFEKMISGMYMGEL
VRLILVRMTKEQLLFQGKTTAQLLTTGSFNTSYIYAIENDKDEEGLASAEKVLRGLGLDPSLEDCIATQR
VCQIVSTRAAHLCAATLVAVLRQIRDNKAAEKLRTTIGVDGSVYKNHPEFSRRLHKMVRRLVPDCDVRFL
RSEVGSGKGAAMVTAVAYRLAAQHAERQRILDTLRLSREQLLEVKKRMSEEMVRGLSKQTHEQASVKMLP
TYVRSTPDGTEHGDFLALDLGGSSFRVLLVRVRSGKRHNVDMHHKIYSIPQETMQGTGEELFSHIVDCIA
DFLEYMGMRGASLPLGFTFSFPCHQSKLDQGILLKWTKGFKASGCEGQDVVTLLKDAVRRRGEFDLNFVA
VVNDTVGTMMTCGYEDPKCEVGLIVGTGTNACYMEEMQNIEVVEGDDGRMCVNMEWGAFGDNGELDDFFT
QFDRIVDDCSNNPGKQRYEKMISGMYLGEIVRNVLMDFTAKGLLFRGKLSERLKTRGIFETKFLSQIESD
RLAMRQVRSILQHLGLTSSTCDDSVLVKEVCSVVSRRAAQLCGAGLAAVVDKIRQNRNLQQLSITVGVDG
```

```
TLYKTHPHFSSIMQETLQDLAPQCKVTFHKSEDGSGKGAALITAVACRIKSEGQH
>anc144_altAll|avgPP:0.995|lnPP:-8.080|num_ambig:4|num_ambig_gaps:0
LHHLQLSDETLMDVSIRFRREMDKGLCRDTNPTAAVKMLPTFVLSTPDGTEQGEFLALDLGGSNFRVLLV
KVMANGKQKVEMENQIYAIPEHLMRGSGSELFDHIADCLANFLEKLGIKDKKLPLGFTFSFPCQQTKLDE
SVLVSWTKGFRTSGVEGKDVVSLLRKSIKKRGDFDVDIMAVINDTVGAMMTCGYDDHHCEIGLIVGTGTN
ACYMEQMRNLELLDGDEGRMCVNTEWGAFGDDGALEDLRTDIDREIDAGSLNPGKQLFEKMISGMYMGEL
VRLILVRMTKEQLLFQGKTTAQLLTTGSFNTSYIYAIENDKDEEGLASAEKVLRGLGLDPSVEDCIATQR
VCQIVSTRAAHLCAATLVAVLRQIRDNKAAEKLRTTIGVDGSVYKNHPEFPRRLHKMVRRLVPDCDVRFL
RSEVGSGKGAAMVTAVAYRLAAQHAERQRILDTLRLSREQLLEVKKRMSEEMVRGLSKQTHEQASVKMLP
TYVRSTPDGTEHGDFLALDLGGSSFRVLLVRVRSGKRHNVDMHHKIYSIPQETMQGTGEELFSHIVDCIA
DFLEYMGMRGASLPLGFTFSFPCHQSKLDQGILLKWTKGFKASGCEGQDVVTLLKDAVRRRGEFDLNFVA
VVNDTVGTMMTCGYEDPKCEVGLIVGTGTNACYMEEMQNIEVVEGDDGRMCVNMEWGAFGDNGELDDFFT
QFDRVVDDCSNNPGKQRYEKMISGMYLGEIVRNVLMDFTAKGLLFRGKLSERLKTRGIFETKFLSQIESD
RLAMRQVRSILQHLGLTSSTCDDSVLVKEVCSVVSRRAAQLCGAGLAAVVDKIRQNRNLQQLSITVGVDG
TLYKTHPHFSSIMQETLQDLAPQCKVTFHKSEDGSGKGAALITAVACRIKSEGQH
>anc145|avgPP:0.998|lnPP:-3.486|num_ambig:3|num_ambig_gaps:0
LHHLQLSDETLMDVSIRFRREMDKGLCRDTNPTAAVKMLPTFVRSTPDGTEQGEFLALDLGGSNFRVLLV
KVMANGKQKVEMENQIYAIPEHLMRGSGSELFDHIADCLANFLEKLGIKDKKLPLGFTFSFPCQQTKLDE
SVLVSWTKGFKSSGVEGKDVVSLLRKSIKKRGDFDIDIVAVINDTVGTMMTCGYDDHHCEIGLIVGTGTN
ACYMEQMRNLELLDGDEGRMCVNTEWGAFGDDGALEDLRTDIDREIDAGSLNPGKQLFEKMISGMYMGEL
VRLILVRMTKEQLLFQGKTTAQLLTTGSFNTSYIYAIENDKDEEGLASAEKVLRGLGLDPSVEDCIATQR
VCQIVSTRAAHLCAATLVAVLRQIRDNKAAEKLRTTIGVDGSVYKNHPEFSRRLHKMVRRLVPDCDVRFL
LSEDGSGKGAAMVTAVAYRLAAQHAERQRILDTLRLSREQLLEVKKRMSEEMVRGLSKQTHEQASVKMLP
TYVRSTPDGTEHGDFLALDLGGSSFRVLLVRVRSGKRHNVDMHHKIYSIPQETMQGTGEELFDHIVDCIA
DFLEYMGMRGASLPLGFTFSFPCHQSKLDQGILLKWTKGFKASGCEGQDVVTLLKDAVRRRQEFDLNFVA
VVNDTVGTMMTCGYEDPKCEVGLIVGTGTNACYMEEMQNIELVEGDDGRMCVNMEWGAFGDNGELDDFCT
QFDRIVDDCSNNPGKQRYEKMISGMYLGEIVRNVLLDFTAKGLLFRGKLSERLKTRGIFETKFLSQIESD
RLAMRQVRSILQHLGLTSSTCDDSVLVKEVCSVVARRAAQLCGAGLAAVVDKIRQNRNLQQLSITVGVDG
TLYKTHPHFSSIMQETLQDLAPQCQVTFHKSEDGSGKGAALITAVACRIKSEGQH
>anc145_altAll|avgPP:0.997|lnPP:-5.119|num_ambig:3|num_ambig_gaps:0
LHHLQLSDETLMDVSIRFRREMDKGLCRDTNPTAAVKMLPTFVRSTPDGTEQGEFLALDLGGSNFRVLLV
KVMANGKQKVEMENQIYAIPEHLMRGSGSELFDHIADCLANFLEKLGIKDKKLPLGFTFSFPCQQTKLDE
SVLVSWTKGFKSSGVEGKDVVSLLRKSIKKRGDFDIDIVAVINDTVGTMMTCGYDDHHCEIGLIVGTGTN
ACYMEQMRNLELLDGDEGRMCVNTEWGAFGDDGALEDLRTDIDREIDAGSLNPGKQLFEKMISGMYMGEL
VRLILVRMTKEQLLFQGKTTAQLLTTGSFNTSYIYAIENDKDEEGLASAEKVLRGLGLDPSVEDCIATQR
VCQIVSTRAAHLCAATLVAVLRQIRDNKAAEKLRTTIGVDGSVYKNHPEFSRRLHKMVRRLVPDCDVRFL
LSEDGSGKGAAMVTAVAYRLAAQHAERQRILDTLRLSREQLLEVKKRMSEEMVRGLSKQTHEQASVKMLP
TYVRSTPDGTEHGDFLALDLGGSSFRVLLVRVRSGKRHNVDMHHKIYSIPQETMQGTGEELFNHIVDCIA
DFLEYMGMRGASLPLGFTFSFPCHQSKLDQGILLKWTKGFKASGCEGQDVVTLLKDAVRRRQEFDLNFVA
VVNDTVGTMMTCGYEDPKCEVGLIVGTGTNACYMEEMQNIELVEGDDGRMCVNMEWGAFGDNGELDDFCT
QFDRVVDDCSNNPGKQRYEKMISGMYLGEIVRNVLMDFTAKGLLFRGKLSERLKTRGIFETKFLSQIESD
RLAMRQVRSILQHLGLTSSTCDDSVLVKEVCSVVARRAAQLCGAGLAAVVDKIRQNRNLQQLSITVGVDG
TLYKTHPHFSSIMQETLQDLAPQCQVTFHKSEDGSGKGAALITAVACRIKSEGQH
>anc146|avgPP:0.989|lnPP:-17.016|num_ambig:12|num_ambig_gaps:0
LHHLQLSDENLMDVSIRFRREMDKGLCRDTNPTAAVKMLPTFVRSTPDGTEKGEFLALDLGGSNFRVLLV
KVMANGKQKVEMENQIYAIPEHLMRGSGSELFDHIADCLANFLEKLGIKDKKLPLGFTFSFPCQQTKLDE
SILVSWTKGFKSSGVEGKDVVSLLRKAIKKRGDFDIDIVAVINDTVGTMMTCGYDDHHCEIGLIVGTGTN
ACYMEEMRNLELVEGDEGRMCVNMEWGAFGDDGALEDLRTDIDREIDAGSLNPGKQLFEKMISGMYMGEL
VRLILVRMTKEQLLFQGKTTAQLLTTGHFHTSYIYAIENDKDEEGLLSAEKVLRGLGLDPSVEDCIATQR
VCQIVSTRAAHLCAATLVAVLRQIRDNKAAEKLRTTIGVDGSVYKNHPEFSRRLHKMVRRLVPDCDVRFL
LSEDGSGKGAAMVTAVAYRLAAQHAERQRILDTLRLSREQLLEVKKRMSEEMVRGLSKQTHEQASVKMLP
TYVRSTPDGTEHGDFLALDLGGSSFRVLLVRVRSGKRRNVEMHNKIYSIPQETMQGTGEELFDHIVDCIA
```

DFLEYMGMKGASLPLGFTFSFPCHQSKLDQGILLKWTKGFKASGCEGQDVVTLLKDAVHRRQEFDLNFVA
VVNDTVGTMMTCGYEDPKCEVGLIVGTGTNACYMEEMQNIELVEGDEGRMCVNMEWGAFGDHGELDDFCT
QFDRTVDDCSNNPGKQRYEKMISGMYLGEIVRNVLLDFTAKGLLFRGKLSERLKTRGIFETKFLSQIESD
RLALRQVRSILQHLGLTSSTCDDSVLVKEVCSVVARRAAQLCGAGLAAVVDKIRQNRNLQQLCITVGVDG
TLYKTHPHFSSIMQETLQDLAPQCQVTFLKSEDGSGKGAALITAVACRIKSEGQH
>anc146_altAll|avgPP:0.987|lnPP:-23.516|num_ambig:12|num_ambig_gaps:0
LHHLQLSDETLMDVSVRFRREMDKGLCRDTNPTAAVKMLPTFVRSTPDGTEKGEFLALDLGGSNFRVLLV
KVMANGKQKVEMENQIYAIPEHLMRGSGSELFDHIADCLANFLEKLGIKDKKLPLGFTFSFPCQQTKLDE
SILVSWTKGFKSSGVEGKDVVSLLRKAIKKRGDFDIDIVAVINDTVGTMMTCGYDDHHCEIGLIVGTGTN
ACYMEEMRNLELVEGDEGRMCVNTEWGAFGDDGALEDLRTDIDREIDAGSLNPGKQLFEKMISGMYMGEL
VRLILVRMTKEQLLFQGKTTAQLLTTGHFNTSYIYAIENDKDNEGLLSAEKVLRGLGLDPSVEDCIATQR
VCQIVSTRAAHLCAATLVAVLRQIRDNKAAEKLRTTIGVDGSVYKNHPEFARRLHKMVRRLVPDCDVRFL
LSEDGSGKGAAMVTAVAYRLAAQHAERQRILDTLRLSREQLLEVKRRMSEEMVRGLSKQTHEQASVKMLP
TYVRSTPDGTEHGDFLALDLGGSSFRVLLVRVRSGKRRRVEMHNKIYSIPQETMQGTGEELFDHIVDCIA
DFLEYMGMKGASLPLGFTFSFPCHQSKLDQGILLKWTKGFKASGCEGEDVVTLLKDAVHRRQEFDLNFVA
VVNDTVGTMMTCGYEDPKCEVGLIVGTGTNACYMEEMQNIELVEGDDGRMCVNMEWGAFGDNGELDDFCT
QFDRTVDDCSNNPGKQRYEKMISGMYLGEIVRNVLLDFTAKGLLFRGKLSERLKTRGIFETKFLSQIESD
RLALRQVRSILQHLGLTSSTCDDSVLVKEVCSVVARRAAQLCGAGLAAVVDKIRQNRNLQQLCITVGVDG
TLYKMHPHFSSIMQETLQDLAPQCQVTFLKSEDGSGKGAALITAVACRIKSEGQH
>anc147|avgPP:0.991|lnPP:-14.093|num_ambig:8|num_ambig_gaps:0
LYHLRLSDENLMDVSVRFRREMDKGLCRDTNPTAAVKMLPTFVRSTPDGTEKGDFLALDLGGTNFRVLLV
KVSANGKQKVEMENQIYAIPEHLMRGSGSELFDHIADCLANFLEKLGIKDKKLPLGFTFSFPCQQTKLDE
SILVSWTKGFKASGVEGKDVVSLLRKAIKKRGDFDIDIVAVINDTVGTMMTCGYDDHHCEIGLIVGTGTN
ACYMEEMRNLELVEGDEGRMCVNMEWGAFGDDGALEDLRTDFDREIDAGSLNPGKQLFEKMISGMYMGEL
VRLILVKMTKEQLLFQGRTTPDLLTTGHFHTSYIYAIENDKDNEGLLSAEKVLRGLGLDPSAEDCIATQR
VCQIVSTRAAHLCAATLVAVLRQIRDNKAAERLRTTIGVDGSVYKNHPQFARRLHKMVRRLVPDCDVRFL
LSEDGSGKGAAMVTAVAYRLATQHAERQRILDTLRLSREQLLEVKRRMAEEMVRGLAKQTHEQASVKMLP
TYVRSTPDGTEHGDFLALDLGGSNFRVLLVRVRSGKRRSVEMHNKIYSIPQETMQGTGEELFDHIVHCIA
DFLEYMGMKGASLPLGFTFSFPCHQSKLDQGILLKWTKGFKASGCEGEDVVTLLKDAVHRRQEFDLDVVA
VVNDTVGTMMTCGYEDPLCEVGLIVGTGTNACYMEEMQNVELVEGDEGRMCVNMEWGAFGDHGELDDFCT
EFDRTVDDHSNNPGKQRYEKMISGMYLGEIVRNVLLDFTAKGLLFRGKLSERLKTRGIFETKFLSQIESD
RLALRQVRSILQHLGLTSSTCDDSILVKEVCSVVARRAAQLCGAGLAAVVDKIRQNRNLEQLCITVGVDG
TLYKMHPHFSSIMRETVQDLAPQCEVTFLQSEDGSGKGAALITAVACRIRDAGQH
>anc147_altAll|avgPP:0.989|lnPP:-18.093|num_ambig:8|num_ambig_gaps:0
LYHLRLSDENLMDVSIRFRREMDKGLCRDTNPTAAVKMLPTFVRSTPDGTEKGDFLALDLGGTNFRVLLV
KVSANGKQKVEMENQIYAIPEHLMRGSGSELFDHIADCLANFLEKLGIKDKKLPLGFTFSFPCQQTKLDE
SILVSWTKGFKVSGVEGKDVVSLLRKAIKKRGDFDIDIVAVINDTVGTMMTCGYDDHHCEIGLIVGTGTN
ACYMEEMRNLELVEGDEGRMCVNMEWGAFGDDGALEDLRTDFDREIDAGSLNPGKQLFEKMISGMYMGEL
VRLILVKMTKEQLLFQGRTTPDLLTTGHFQTSYIFAIENDKDNEGLLSAEKVLRGLGLDPSAEDCIATQR
VCQIVSTRAAHLCAATLVAVLRQIRDNKAAEKLRTTIGVDGSVYKNHPQFARRLHKMVRRLVPDCDVRFL
LSEDGSGKGAAMVTAVAYRLATQHAERQRILDTLRLSREQLLEVKRRMAEEMVRGLSKQTHEQASVKMLP
TYVRSTPDGTEHGDFLALDLGGSNFRVLLVRVRSGKRRRVEMHNKIYSIPQETMQGTGEELFDHIVHCIA
DFLEYMGMKGASLPLGFTFSFPCHQSKLDQGILLKWTKGFKASGCEGEDVVTLLKDAVHRRQEFDLDVVA
VVNDTVGTMMTCGYEDPLCEVGLIVGTGTNACYMEEMQNVELVEGDEGRMCVNMEWGAFGDHGELDDFCT
EFDRTVDDHSNNPGKQRYEKMISGMYLGEIVRNVLLDFTAKGLLFRGKLSERLKTRGIFETKFLSQIESD
RLALRQVRSILQHLGLTSSTCDDSILVKEVCSVVARRAAQLCGAGLAAVVDKIRQNRNLEQLCITVGVDG
TLYKLHPHFSSIMRETVQDLAPQCEVTFLQSEDGSGKGAALITAVACRIRDAGQH
>anc148|avgPP:0.993|lnPP:-11.203|num_ambig:7|num_ambig_gaps:0
LYHLRLSDENLMDVSIRFRREMDKGLGRDTNPTAAVKMLPTFVRSTPDGTEKGDFLALDLGGTNFRVLLV
KVSANGKQKVEMENQIYAIPENLMRGSGSELFDHIADCLANFLEKLGIKDKKLPLGFTFSFPCQQTKLDE
SILVSWTKGFKASGVEGKDVVSLLRKAIKKRGDFDIDIVAVINDTVGTMMTCGYDDHHCEIGLIVGTGTN
ACYMEEMRNLELVEGDEGRMCVNMEWGAFGDDGALDDLRTDFDREIDAGSLNPGKQLFEKMISGMYMGEL

```
VRLILVKMAKEQLLFQGHTTPDLLTTGHFQTSYVSAIENDKDNEGLLSAEKVLRGLGLDPSAEDCVATQR
VCQIVSTRAAHLCAATLVAVLRQIRDNKAAERLRTTIGVDGSVYKNHPQFARRLHKMVRRLVPDCDVRFL
RSEDGSGKGAAMVTAVAYRLATQHAERQRILDTLRLSREQLLEVKRRMAEEMERGLAKETHDQASVKMLP
TYVRSTPDGTEHGDFLALDLGGTNFRVLLVRVRSGKRRSVEMHNKIYSIPQETMQGTGEELFDHIVHCIA
DFLEYMGMKGASLPLGFTFSFPCHQSKLDQGILLKWTKGFKATGCEGEDVVTLLKDAIHRREEFDLDVVA
VVNDTVGTMMTCGYEDPLCEVGLIVGTGTNACYMEEMQNVELVEGDEGRMCVNMEWGAFGDHGELDDFCT
EFDRAVDEHSNNPGKQRYEKMISGMYLGEIVRNVLLDFTAKGLLFRGKLSERLKTRGIFETKFLSQIESD
RLALRQVRSILQHLGLTSSTCDDSILVKEVCSVVARRAAQLCGAGLAAVVDKIRQNRNLEQLRITVGVDG
TLYKLHPHFSSIMRETVQDLAPQCEVTFLQSEDGSGKGAALITAVACRIRDAGQH
>anc148_altAll|avgPP:0.991|lnPP:-14.783|num_ambig:7|num_ambig_gaps:0
LYHLRLSDENLMDVSVRFRREMDKGLGRDTNPTAAVKMLPTFVRSTPDGTEKGDFLALDLGGTNFRVLLV
KVSANGKQKVEMENQIYAIPENLMRGSGSELFDHIADCLANFLEKLGIKDKKLPLGFTFSFPCQQTKLDE
SILVSWTKGFKVSGVEGKDVVSLLRKAIKKRGDFDIDIVAVINDTVGTMMTCGYDDHHCEIGLIVGTGTN
ACYMEEMRNLELVEGDEGRMCVNMEWGAFGDDGALDDLRTDFDREIDAGSLNPGKQLFEKMISGMYMGEL
VRLILVKMAKEQLLFQGHTTPDLLTTGHFQTSYVSAIENDKDNEGLLSAEKVLRGLGLDPSAEDCVATQR
VCQIVSTRAAHLCAATLVAVLRQIRDNKAAEKLRTTIGVDGSVYKNHPQFARRLHKMVRRLVPDCDVRFL
QSEDGSGKGAAMVTAVAYRLATQHAERQRILDTLRLSREQLLEVKRRMAEEMERGLAKETHEQASVKMLP
TYVRSTPDGTEHGDFLALDLGGTNFRVLLVRVRSGKRRSVEMHNKIYSIPQETMQGTGEELFDHIVHCIA
DFLEYMGMKGASLPLGFTFSFPCHQSKLDQGILLKWTKGFKATGCEGEDVVTLLKDAIHRREEFDLDVVA
VVNDTVGTMMTCGYEDPLCEVGLIVGTGTNACYMEEMQNVELVEGDEGRMCVNMEWGAFGDHGELDDFCT
EFDRTVDERSNNPGKQRYEKMISGMYLGEIVRNVLLDFTAKGLLFRGKLSERLKTRGIFETKFLSQIESD
RLALRQVRSILQHLGLTSSTCDDSILVKEVCSVVARRAAQLCGAGLAAVVDKIRQNRNLEQLRITVGVDG
TLYKLHPHFSSIMRETVQDLAPQCEVTFLQSEDGSGKGAALITAVACRIRDAGQH
>anc149|avgPP:0.993|lnPP:-9.470|num_ambig:8|num_ambig_gaps:0
LYHLRLSDENLMDVSKRFRREMDKGLGRDTNPTAAVKMLPTFVRSTPDGTEKGDFLALDLGGTNFRVLLV
KVSANGKQKVEMENQIYAIPENLMRGSGTELFDHIAECLANFLEKLGIKDKKLPLGFTFSFPCQQTKLDE
SILVSWTKGFKASGVEGRDVVSLLRKAIKKRGDFDIDIVAVINDTVGTMMTCGYDDHHCEIGLIVGTGTN
ACYMEEMRHLELVEGDEGRMCVNMEWGAFGDDGALDDLRTDFDREIDAGSLNPGKQLFEKMISGMYMGEL
VRLILVKMAKEGLLFQGHTTPDLLTTGHFQTSFVSAIENDKDNEGLLSAEKVLRGLGLDPSAEDCVATQR
VCQIVSTRAAHLCAATLAAVLRQIRDNKAAERLRTTIGVDGSVYKNHPQFARRLHKMVRRLVPDCDVRFL
RSEDGSGKGAAMVTAVAYRLATQHAERQRILDTLRLSREQLLEVKRRMAEEMERGLAKETHDQATVKMLP
TYVRSTPDGTERGDFLALDLGGTNFRVLLVRVRSGKRRSVEMHNKIYSIPQETMQGTGEELFDHIVHCIA
DFLEYMGMKGASLPLGFTFSFPCHQSKLDQGILLKWTKGFKATGCEGEDVVTLLKDAIHRREEFDLDVVA
VVNDTVGTMMTCGYEDPLCEVGLIVGTGTNACYMEEMQNVELVEGDEGRMCVNMEWGAFGDHGELDDFCT
EFDRAVDEHSANPGKQRYEKMISGMYLGEIVRNVLLDFTAKGLLFRGKLSERLKTRGIFETKFLSQIESD
RLALRQVRSILQHLGLTSSTCDDSILVKEVCSVVARRAAQLCGAGLAAVVDKIRQNRNLEQLRITVGVDG
TLYKLHPHFSTIMRETVRDLAPQCEVTFLQSEDGSGKGAALITAVACRIRDAGQH
>anc149_altAll|avgPP:0.992|lnPP:-14.413|num_ambig:8|num_ambig_gaps:0
LYHLRLSDENLMDVSKRFRREMDKGLGRDTNPTAAVKMLPTFVRSTPDGTEKGDFLALDLGGTNFRVLLV
KVSANGKQKVEMENQIYAIPENLMRGSGSELFDHIAECLANFLEKLGIKDKKLPLGFTFSFPCQQTKLDE
SILVSWTKGFKASGVEGKDVVSLLRKAIKKRGDFDIDIVAVINDTVGTMMTCGYDDHHCEIGLIVGTGTN
ACYMEEMRHLELVEGDEGRMCVNMEWGAFGDDGALDDLRTDFDREIDAGSLNPGKQLFEKMISGMYMGEL
VRLILVKMAKEGLLFQGHTTPDLLTTGHFQTSFVSAIENDKDNEGLLSAEKVLRGLGLDPSAEDCVATQR
VCQIVSTRAAHLCAATLAAVLRQIRDNKAAERLRTTIGVDGSVYKNHPQFARRLHKMVRRLVPDCDVRFL
RSEDGSGKGAAMVTAVAYRLATQHAERQRILDTLRLSREQLLEVKRRMAEEMERGLAKETHDQASVKMLP
TYVRSTPDGTERGDFLALDLGGTNFRVLLVRVRSGKRRSVEMHNKIYSIPQETMQGTGEELFDHIVHCIA
DFLEYMGMKGASLPLGFTFSFPCHQSKLDQGILLKWTKGFKATGCEGEDVVTLLKDAIHRREEFDLDVVA
VVNDTVGTMMTCGYEDPLCEVGLIVGTGTNACYMEEMQNVELVEGDEGRMCVNMEWGAFGDHGELDDFCT
EFDRAVDERSANPGKQRYEKMISGMYLGEIVRNVLLDFTAKGLLFRGKLSERLKTRGIFETKFLSQIESD
RLALRQVRSILQHLGLTSSTCDDSILVKEVCSVVARRAAQLCGAGLAAVVDKIRQNRNLDQLKITVGVDG
TLYKLHPHFSSIMRETVRDLAPQCEVTFLQSEDGSGKGAALITAVACRIRDAGQR
>anc150|avgPP:0.987|lnPP:-20.827|num_ambig:20|num_ambig_gaps:0
```

LYHLRLSEENLMDVSLRFRREMDRGLGRDTNPTAAVKMLPTFVRSTPDGTEQGDFLALDLGGTNFRVLLV
KVSANGKQKVEMENQIYAIPEDLMRGSGSELFDHIAECLANFLEKLGIKDKKLPLGFTFSFPCQQTKLDE
SILVSWTKGFKASGVEGKDVVSMLRAAIKKRGDFDIDIVAVINDTVGTMMTCGYDDHHCEIGLIVGTGTN
ACYMEEMRNLETVEGDEGRMCVNMEWGAFGDDGALDDLRTSFDREIDAGSLNPGKQLFEKMISGMYMGEL
VRLILVKMAKEELLFQGKTTPELLTTGHFQTSFVSAIENEKDNEGLLSAEQVLQGLGLNPSAEDCVATQR
VCQIVSTRAAHLCAATLAAVLRQIRDNKAAERLRTTVGVDGSVYKNHPQFARRLHKMVRRLVPDCDVRFL
RSEDGSGKGAAMVTAVAYRLANQHAERQRILDTLRLSREQLLEVKRRMAEAMERGLARDTHGNATVKMLP
TYVRSTPNGTERGDFLALDLGGTNFRVLLVRVRSGKKRSVEMHNKIYAIPQDLMQGTGEELFDHIVHCIA
DFLEYMGMKGASLPLGFTFSFPCQQNRLDQGILLKWTKGFRATGCEGQDVVTLLKDAIHRHEEFDLDVVA
VVNDTVGTMMTCGYEDPHCEVGLIVGTGTNACYMEEMENVELVEGNEGRMCVNMEWGAFGDHGELDDFCT
TFDHAVDERSANPGKQRYEKMISGMYLGEIVRNVLLDFTAKGLLFRGRLSERLKTRGIFETKFLSQIESD
RLALRQVRSILQHLGLTSSTCDDSILVKEVCSVVSRRAAQLCGAGLAAVVDKIRKNRHLEQLRVTVGVDG
TLYKLHPHFSTIMRETLRELAPQCEVTLMQSEDGSGKGAALITAVACRLRDAGK
>anc150_altAll|avgPP:0.983|lnPP:-31.117|num_ambig:20|num_ambig_gaps:0
LYHLRLSEENLMDVSLRFRREMDRGLGRDTNPTAAVKMLPTFVRSTPDGTEQGDFLALDLGGTNFRVLLV
KVSSNGKQKVEMENQIYAIPEDLMRGSGSELFDHIAECLANFLEKLGIKDKKLPLGFTFSFPCQQTKLDE
SILVSWTKGFKASGVEGKDVVSMLRAAIKKRGDFDIDIVAVINDTVGTMMTCGYDDHHCEIGLIVGTGTN
ACYMEEMRNLETVEGDEGRMCVNMEWGAFGDDGALDDLRTPFDQEIDAGSLNPGKQLFEKMISGMYMGEL
VRLILVKMAKEELLFQGHTTPELLTTGHFQTSFVSAIENEKDNEGLLNAEQILRGLGLNPSAEDCVATQR
VCQIVSTRAAHLCAATLAAVLRQIRDNKAAERLRTTVGVDGSVYKNHPQFARRLHKMVRRLVPDCDVRFL
RSEDGSGKGAAMVTAVAYRLANQHAERQRILDTLRLSRDQLLEVKRRMAEEMERGLARETHGNATVKMLP
TYVRSTPNGTERGDFLALDLGGTNFRVLLVRVRSGKKRSVEMHNKIYAIPQDIMQGTGEELFDHIVHCIA
DFLEYMGMKGASLPLGFTFSFPCQQNRLDQGILLKWTKGFRATGCEGQDVVTLLKDAIHRHEEFDLDVVA
VVNDTVGTMMTCGYEDPHCEVGLIVGTGTNACYMEEMENVELVEGNEGRMCINMEWGAFGDHGELDDFCT
TFDHAVDERSANPGKQRYEKMISGMYLGEIVRNVLLDFTAKGLLFRGRLSERLKTRGIFETKFLSQIESD
RLALRQVRSILQHLGLTSSTCDDSILVKEVCTVVSRRAAQLCGAGLAAVVDKIRQNRHLEHLRITVGVDG
TLYKLHPHFSTIMRETLRELAPQCEVSLLQSEDGSGKGAALITAVACRLRDAAQ
>anc151|avgPP:0.991|lnPP:-13.374|num_ambig:14|num_ambig_gaps:0
LYHLRLSEENLMDVSKRFRREMDRGLGRDTNPTAAVKMLPTFVRSTPDGTEKGDFLALDLGGTNFRVLLV
KVSANGKQKVEMENQIYAIPENLMRGSGTELFDHIAECLANFLEKLGIKDKKLPLGFTFSFPCQQTKLDE
SILVSWTKGFKASGVEGKDVVSLLRKAIKKRGDFDIDIVAVINDTVGTMMTCGYDDHHCEIGLIVGTGTN
ACYMEEMRHLELVEGDEGRMCVNMEWGAFGDDGALDDLRTDFDREIDAGSLNPGKQLFEKMISGMYMGEL
VRLILVKMAKEGLLFQGHTTPDLLTTGHFQTSFVSAIENEKDNEGLLSAEQVLRGLGLDPSAEDCVATQR
VCQIVSTRAAHLCAATLAAVLRQIRDNKAAERLRTTIGVDGSVYKNHPQFARRLHKMVRRLVPDCDVRFL
RSEDGSGKGAAMVTAVAYRLAKQHAERQRILDTLRLSREQLLEVKRRMAEEMERGLARETHDSATVKMLP
TYVRSTPDGTERGDFLALDLGGTNFRVLLVRVRSGKRRSVEMHNKIYAIPQDLMQGTGEELFDHIVHCIA
DFLEYMGMKGASLPLGFTFSFPCHQTRLDQGILLKWTKGFRATGCEGEDVVTLLKDAIHRHEEFDLDVVA
VVNDTVGTMMTCGYEDPQCEVGLIVGTGTNACYMEEMQNVELVEGNEGRMCVNMEWGAFGDHGELDDFCT
EFDRAVDERSANPGKQRYEKMISGMYLGEIVRNVLLDFTAKGLLFRGKLSERLKTRGIFETKFLSQIESD
RLALRQVRSILQHLGLTSSTCDDSILVKEVCSVVSRRAAQLCGAGLAAVVDKIRQNRNLEQLRVTVGVDG
TLYKLHPHFSTIMRETLRDLAPQCEVTLVQSEDGSGKGAALITAVACRLRDAGK
>anc151_altAll|avgPP:0.989|lnPP:-19.501|num_ambig:14|num_ambig_gaps:0
LYHLRLSEENLMDVSKRFRREMDRGLGRDTNPTAAVKMLPTFVRSTPDGTEKGDFLALDLGGTNFRVLLV
KVSSNGKQKVEMENQIYAIPENLMRGSGTELFDHIAECLANFLEKLGIKDKKLPLGFTFSFPCQQTKLDE
SILVSWTKGFKASGVEGRDVVSLLRKAIKKRGDFDIDIVAVINDTVGTMMTCGYDDHHCEIGLIVGTGTN
ACYMEEMRHLELVEGDEGRMCVNMEWGAFGDDGALDDLRTDFDREIDAGSLNPGKQLFEKMISGMYMGEL
VRLILVKMAKEGLLFQGHTTPDLLTTGHFQTSFVSAIENEKDNEGLLSAEQILRGLGLNPSAEDCVATQR
VCQIVSTRAAHLCAATLAAVLRQIRDNKAAERLRTTIGVDGSVYKNHPQFARRLHKMVRRLVPDCDVRFL
RSEDGSGKGAAMVTAVAYRLAKQHAERQRILDTLRLSREQLLEVKRRMAEEMERGLARETHSSASVKMLP
TYVRSTPDGTERGDFLALDLGGTNFRVLLVRVRSGKRRSVEMHNKIYAIPQDIMQGTGEELFDHIVHCIA
DFLEYMGMKGASLPLGFTFSFPCHQTRLDQGILLKWTKGFRATGCEGEDVVTLLKDAIHRHEEFDLDVVA
VVNDTVGTMMTCGYEDPHCEVGLIVGTGTNACYMEEMQNVELVEGDEGRMCVNMEWGAFGDHGELDDFCT

```
EFDRAVDERSANPGKQRYEKMISGMYLGEIVRNVLLDFTAKGLLFRGKLSERLKTRGIFETKFLSQIESD
RLALRQVRSILQHLGLTSSTCDDSILVKEVCSVVSRRAAQLCGAGLAAVVDKIRQNRNLDHLKITVGVDG
TLYKLHPHFSTIMRETLRDLAPQCEVTLVQSEDGSGKGAALITAVACRLRDAGQ
>anc152|avgPP:0.991|lnPP:-13.908|num_ambig:13|num_ambig_gaps:0
LYHLRLSEENLMDVSKRFRREMDKGLGRDTNLTASVKMLPTFVRSTPDGTEKGDFLALDLGGTNFRVLLV
KVSSNGKQTVEMENQIYAIPETLMRGSGTELFDHIAECLANFLEKLGIKEKKLPLGFTFSFPCQQTKLDE
SILVSWTKGFKASGVEGRDVVGLLRKAIKKRGDFDIDIVAVINDTVGTMMTCGFDDHHCEIGLIVGTGTN
ACYMEEMRNLELVEGDEGRMCVNMEWGAFGDDGALDDIRTDFDREIDAGSLNPGKQLFEKMISGMYMGEL
VRIILVKMTKEGLLFQGHTTPDLLTTGHFKTSYVSAIENDKSKEGLVCVEQVLRGLGLEPSAEDCVAVQR
VCQIVSTRAAHLCAATLAAVLRQIRDNKAAERLRTTIGVDGSVYKNHPQFARRLHKMVRRLVPDCDVRFL
RSEDGSGKGAAMVTAVAYRLAKQHAERQRVLDTLRLSREQLMEVKRRMEEQMKRGLAKDTHDSASVKMLP
TYVRATPDGTERGDFLALDLGGTNFRVLLVRVRSGKRRSVEMHNKIYTIPQDIMQGTGEELFDHIVHCIA
DFLEYMGMKGASLPLGFTFSFPCHQTGLDQGILLKWTKGFKATGCEGEDVAMLLKDAIHRMEEFDLDVVA
VVNDTVGTMMSCGYEDPQCEVGLIVGTGTNACYMEEMRNVELVEGDEGRMCVNMEWGAFGDDGELDDICT
PFDRAVDENSTNPGKQRFEKMISGMYLGEIVRNVLLDFTAKGLLFRGKLSERLKTRGIFETKFLSQIESD
RLALRQVRSILQHLGLTSSTCDDSILVKEVCSVVSQRAAQLCGAALAAVVDKIRQNRNLNSLSVTVGVDG
TLYKLHPHFSSIMRETLRDLAPQCDVTLVQSEDGSGKGAALITAVACRIRDAKK
>anc152_altAll|avgPP:0.989|lnPP:-19.628|num_ambig:13|num_ambig_gaps:0
LYHLRLSDENLMDVSKRFRREMDKGLGRDTNLTASVKMLPTFVRSTPDGTEKGDFLALDLGGTNFRVLLV
KVSSNGKQTVEMENQIYAIPETLMRGSGTELFDHIAECLANFLDKLGIKEKKLPLGFTFSFPCQQTKLDE
SILVSWTKGFKASGVEGRDVVSLLRKAIKKRGDFDIDIVAVINDTVGTMMTCGFDDHHCEIGLIVGTGTN
ACYMEEMRNLELVEGDEGRMCVNMEWGAFGDDGALDDIRTDFDREIDAGSLNPGKQLFEKMISGMYMGEL
VRIILVKMTKEGLLFQGHTTPDLLTTGHFKTSFVSAIENDKSKEGLVSVEQVLRGLGLEPSQEDCVAVQR
VCQIVSTRAAHLCAASLAAVLRQIRDNKAAERLRTTIGVDGSVYKNHPQFARRLHKMVRRLVPDCDVRFL
RSEDGSGKGAAMVTAVAYRLAKQHAERQKVLDTLRLSREQLMEVKRRMEEQMKRGLAKDTHDSASVKMLP
TYVRATPDGTERGDFLALDLGGTNFRVLLVRVRSGKRRSVEMHNKIYTIPQDIMQGTGEELFDHIVHCIA
DFLEYMGMKGASLPLGFTFSFPCHQTRLDQGILLKWTKGFKATGCEGEDVAVLLKDAIHRMEEFDLDVVA
VVNDTVGTMMSCGYEDPQCEVGLIVGTGTNACYMEEMRNVELVEGDEGRMCVNMEWGAFGDDGELDDVCT
PFDRAVDENSTNPGKQRFEKMISGMYLGEIVRNVLLDFTAKGLLFRGKLSERLKTRGIFETKFLSQIESD
RLALRQVRSILQHLGLTSSTCDDSILVKEVCSVVSQRAAQLCGAALAAVVDKIRQNRNLNTLSVTVGVDG
TLYKLHPHFSTIMRETLRDLAPQCDVTLVQSEDGSGKGAALITAVACRIRDAKK
>anc153|avgPP:0.992|lnPP:-12.501|num_ambig:13|num_ambig_gaps:0
LYHLRLSEENLMDVSKRFRREMDKGLGRDTNLTASVKMLPTFVRSTPDGTEKGDFLALDLGGTNFRVLLV
KVSANGKQKVEMENQIYAIPETLMRGSGTELFDHIAECLANFLEKLGIKDKKLPLGFTFSFPCQQTKLDE
SILVSWTKGFKASGVEGRDVVGLLRKAIKKRGDFDIDIVAVINDTVGTMMTCGYDDHHCEIGLIVGTGTN
ACYMEEMRNLELVEGDEGRMCVNMEWGAFGDDGALDDLRTDFDREIDAGSLNPGKQLFEKMISGMYMGEL
VRIILVKMAKEGLLFQGHTTPDLLTTGHFKTSYVSAIENEKDKEGLLSAEQVLRGLGLDPSAEDCVAVQR
VCQIVSTRAAHLCAATLAAVLRQIRDNKAAERLRTTIGVDGSVYKNHPQFARRLHKMVRRLVPDCDVRFL
RSEDGSGKGAAMVTAVAYRLAKQHAERQRVLDTLRLSREQLMEVKRRMEEEMKRGLAKDTHDSASVKMLP
TYVRSTPDGTERGDFLALDLGGTNFRVLLVRVRSGKRRSVEMHNKIYTIPQDIMQGTGEELFDHIVHCIA
DFLEYMGMKGASLPLGFTFSFPCHQTRLDQGILLKWTKGFKATGCEGEDVATLLKDAIHRMEEFDLDVVA
VVNDTVGTMMTCGYEDPQCEVGLIVGTGTNACYMEEMQNVELVEGDEGRMCVNMEWGAFGDHGELDDFCT
QFDRAVDERSANPGKQRFEKMISGMYLGEIVRNVLLDFTAKGLLFRGKLSERLKTRGIFETKFLSQIESD
RLALRQVRSILQHLGLTSSTCDDSILVKEVCSVVSRRAAQLCGAGLAAVVDKIRQNRNLEQLSVTVGVDG
TLYKLHPHFSTIMRETLRDLAPQCDVTLVQSEDGSGKGAALITAVACRLRDAKK
>anc153_altAll|avgPP:0.989|lnPP:-20.056|num_ambig:13|num_ambig_gaps:0
LYHLRLSEENLMDVSKRFRREMDKGLGRDTNLTASVKMLPTFVRSTPDGTEKGDFLALDLGGTNFRVLLV
KVSSNGKQKVEMENQIYAIPETLMRGSGTELFDHIAECLANFLDKLGIKDKKLPLGFTFSFPCQQTKLDE
SILVSWTKGFKASGVEGKDVVSLLRKAIKKRGDFDIDIVAVINDTVGTMMTCGYDDHHCEIGLIVGTGTN
ACYMEEMRNLELVEGDEGRMCVNMEWGAFGDDGALDDLRTDFDREIDAGSLNPGKQLFEKMISGMYMGEL
VRIILVKMAKEGLLFQGHTTPDLLTTGHFKTSFVSAIENDKDKEGLVCAEQVLRGLGLDPSAEDCVAVQR
VCQIVSTRAAHLCAATLAAVLRQIRDNKAAERLRTTIGVDGSVYKNHPQFARRLHKMVRRLVPDCDVRFL
```

RSEDGSGKGAAMVTAVAYRLAKQHAERQRVLDTLRLSREQLMEVKRRMEEEMKRGLAKDTHDSASVKMLP
TYVRSTPDGTERGDFLALDLGGTNFRVLLVRVRSGKRRSVEMHNKIYTIPQDIMQGTGEELFDHIVHCIA
DFLEYMGMKGASLPLGFTFSFPCHQTRLDQGILLKWTKGFKATGCEGEDVAALLKDAIHRMEEFDLDVVA
VVNDTVGTMMTCGYEDPQCEVGLIVGTGTNACYMEEMRNVELVEGDEGRMCVNMEWGAFGDHGELDDFCT
QFDRAVDERSANPGKQRFEKMISGMYLGEIVRNVLLDFTAKGLLFRGKLSERLKTRGIFETKFLSQIESD
RLALRQVRSILQHLGLTSSTCDDSILVKEVCSVVSRRAAQLCGAGLAAVVDKIRQNRNLDQLTVTVGVDG
TLYKLHPHFSTIMRETLKDLAPQCDVTLVQSEDGSGKGAALITAVACRLRDAKK
>anc154|avgPP:0.992|lnPP:-11.263|num_ambig:9|num_ambig_gaps:0
LYHLRLSEENLMDVSKRFRREMDKGLGRDTNLTAAVKMLPTFVRSTPDGTEKGDFLALDLGGTNFRVLLV
KVSANGKQKVEMENQIYAIPENLMRGSGTELFDHIAECLANFLEKLGIKDKKLPLGFTFSFPCQQTKLDE
SILVSWTKGFKASGVEGRDVVSLLRKAIKKRGDFDIDIVAVINDTVGTMMTCGYDDHHCEIGLIVGTGTN
ACYMEEMRHLELVEGDEGRMCVNMEWGAFGDDGALDDLRTDFDREIDAGSLNPGKQLFEKMISGMYMGEL
VRLILVKMAKEGLLFQGHTTPDLLTTGHFQTSFVSAIENEKDNEGLLSAEQVLRGLGLDPSAEDCVATQR
VCQIVSTRAAHLCAATLAAVLRQIRDNKAAERLRTTIGVDGSVYKNHPQFARRLHKMVRRLVPDCDVRFL
RSEDGSGKGAAMVTAVAYRLAKQHAERQRILDTLRLSREQLLEVKRRMAEEMKRGLAKETHDSASVKMLP
TYVRSTPDGTERGDFLALDLGGTNFRVLLVRVRSGKRRSVEMHNKIYTIPQDIMQGTGEELFDHIVHCIA
DFLEYMGMKGASLPLGFTFSFPCHQTRLDQGILLKWTKGFKATGCEGEDVATLLKDAIHRHEEFDLDVVA
VVNDTVGTMMTCGYEDPQCEVGLIVGTGTNACYMEEMQNVELVEGDEGRMCVNMEWGAFGDHGELDDFCT
EFDRAVDERSANPGKQRYEKMISGMYLGEIVRNVLLDFTAKGLLFRGKLSERLKTRGIFETKFLSQIESD
RLALRQVRSILQHLGLTSSTCDDSILVKEVCSVVSRRAAQLCGAGLAAVVDKIRQNRNLEQLKVTVGVDG
TLYKLHPHFSTIMRETLRDLAPQCEVTLVQSEDGSGKGAALITAVACRLRDAGK
>anc154_altAll|avgPP:0.991|lnPP:-15.072|num_ambig:9|num_ambig_gaps:0
LYHLRLSEENLMDVSKRFRREMDKGLGRDTNLTAAVKMLPTFVRSTPDGTEKGDFLALDLGGTNFRVLLV
KVSSNGKQKVEMENQIYAIPENLMRGSGTELFDHIAECLANFLEKLGIKDKKLPLGFTFSFPCQQTKLDE
SILVSWTKGFKASGVEGKDVVSLLRKAIKKRGDFDIDIVAVINDTVGTMMTCGYDDHHCEIGLIVGTGTN
ACYMEEMRHLELVEGDEGRMCVNMEWGAFGDDGALDDLRTDFDREIDAGSLNPGKQLFEKMISGMYMGEL
VRLILVKMAKEGLLFQGHTTPDLLTTGHFQTSFVSAIENEKDNEGLVSAEQVLRGLGLDPSAEDCVATQR
VCQIVSTRAAHLCAATLAAVLRQIRDNKAAERLRTTIGVDGSVYKNHPQFARRLHKMVRRLVPDCDVRFL
RSEDGSGKGAAMVTAVAYRLAKQHAERQRILDTLRLSREQLLEVKRRMAEEMERGLAKETHSSASVKMLP
TYVRSTPDGTERGDFLALDLGGTNFRVLLVRVRSGKRRSVEMHNKIYTIPQDIMQGTGEELFDHIVHCIA
DFLEYMGMKGASLPLGFTFSFPCHQTRLDQGILLKWTKGFKATGCEGEDVATLLKDAIHRREEFDLDVVA
VVNDTVGTMMTCGYEDPQCEVGLIVGTGTNACYMEEMQNVELVEGDEGRMCVNMEWGAFGDHGELDDFCT
EFDRAVDERSANPGKQRYEKMISGMYLGEIVRNVLLDFTAKGLLFRGKLSERLKTRGIFETKFLSQIESD
RLALRQVRSILQHLGLTSSTCDDSILVKEVCSVVSRRAAQLCGAGLAAVVDKIRQNRNLDQLRITVGVDG
TLYKLHPHFSTIMRETLRDLAPQCEVTLVQSEDGSGKGAALITAVACRLRDAGK
>anc155|avgPP:0.993|lnPP:-11.068|num_ambig:11|num_ambig_gaps:0
LYHLRLSEENLMDVSKRFRREMDKGLGRDTNPTAAVKMLPTFVRSTPDGTEKGDFLALDLGGTNFRVLLV
KVSANGKQKVEMENQIYAIPENLMRGSGTELFDHIAECLANFLEKLGIKDKKLPLGFTFSFPCQQTKLDE
SILVSWTKGFKASGVEGRDVVSLLRKAIKKRGDFDIDIVAVINDTVGTMMTCGYDDHHCEIGLIVGTGTN
ACYMEEMRHLELVEGDEGRMCVNMEWGAFGDDGALDDLRTDFDREIDAGSLNPGKQLFEKMISGMYMGEL
VRLILVKMAKEGLLFQGHTTPDLLTTGHFQTSFVSAIENEKDNEGLLSAEQVLRGLGLDPSAEDCVATQR
VCQIVSTRAAHLCAATLAAVLRQIRDNKAAERLRTTIGVDGSVYKNHPQFARRLHKMVRRLVPDCDVRFL
RSEDGSGKGAAMVTAVAYRLAKQHAERQRILDTLRLSREQLLEVKRRMAEEMERGLAKETHDSATVKMLP
TYVRSTPDGTERGDFLALDLGGTNFRVLLVRVRSGKRRSVEMHNKIYTIPQDIMQGTGEELFDHIVHCIA
DFLEYMGMKGASLPLGFTFSFPCHQTRLDQGILLKWTKGFKATGCEGEDVVTLLKDAIHRHEEFDLDVVA
VVNDTVGTMMTCGYEDPQCEVGLIVGTGTNACYMEEMQNVELVEGDEGRMCVNMEWGAFGDHGELDDFCT
EFDRAVDERSANPGKQRYEKMISGMYLGEIVRNVLLDFTAKGLLFRGKLSERLKTRGIFETKFLSQIESD
RLALRQVRSILQHLGLTSSTCDDSILVKEVCSVVSRRAAQLCGAGLAAVVDKIRQNRNLEQLRVTVGVDG
TLYKLHPHFSTIMRETLRDLAPQCEVTLVQSEDGSGKGAALITAVACRLRDAGK
>anc155_altAll|avgPP:0.991|lnPP:-15.659|num_ambig:11|num_ambig_gaps:0
LYHLRLSEENLMDVSKRFRREMDKGLGRDTNPTAAVKMLPTFVRSTPDGTEKGDFLALDLGGTNFRVLLV
KVSSNGKQKVEMENQIYAIPENLMRGSGTELFDHIAECLANFLEKLGIKDKKLPLGFTFSFPCQQTKLDE

SILVSWTKGFKASGVEGKDVVSLLRKAIKKRGDFDIDIVAVINDTVGTMMTCGYDDHHCEIGLIVGTGTN
ACYMEEMRHLELVEGDEGRMCVNMEWGAFGDDGALDDLRTDFDREIDAGSLNPGKQLFEKMISGMYMGEL
VRLILVKMAKEGLLFQGHTTPDLLTTGHFQTSFVSAIENEKDNEGLLSAEQVLRGLGLDPSAEDCVATQR
VCQIVSTRAAHLCAATLAAVLRQIRDNKAAERLRTTIGVDGSVYKNHPQFARRLHKMVRRLVPDCDVRFL
RSEDGSGKGAAMVTAVAYRLAKQHAERQRILDTLRLSREQLLEVKRRMAEEMERGLAKETHSSASVKMLP
TYVRSTPDGTERGDFLALDLGGTNFRVLLVRVRSGKRRSVEMHNKIYAIPQDIMQGTGEELFDHIVHCIA
DFLEYMGMKGASLPLGFTFSFPCHQTRLDQGILLKWTKGFKATGCEGEDVVTLLKDAIHRREEFDLDVVA
VVNDTVGTMMTCGYEDPHCEVGLIVGTGTNACYMEEMQNVELVEGDEGRMCVNMEWGAFGDHGELDDFCT
EFDRAVDERSANPGKQRYEKMISGMYLGEIVRNVLLDFTAKGLLFRGKLSERLKTRGIFETKFLSQIESD
RLALRQVRSILQHLGLTSSTCDDSILVKEVCSVVSRRAAQLCGAGLAAVVDKIRQNRNLDQLKITVGVDG
TLYKLHPHFSTIMRETLRDLAPQCEVTLVQSEDGSGKGAALITAVACRLRDAGQ
>anc156|avgPP:0.993|lnPP:-10.841|num_ambig:10|num_ambig_gaps:0
LYHLRLSDENLMDVSKRFRREMDKGLGRDTNPTAAVKMLPTFVRSTPDGTEKGDFLALDLGGTNFRVLLV
KVSANGKQKVEMENQIYAIPENLMRGSGTELFDHIAECLANFLEKLGIKDKKLPLGFTFSFPCQQTKLDE
SILVSWTKGFKASGVEGRDVVSLLRKAIKKRGDFDIDIVAVINDTVGTMMTCGYDDHHCEIGLIVGTGTN
ACYMEEMRHLELVEGDEGRMCVNMEWGAFGDDGALDDLRTDFDREIDAGSLNPGKQLFEKMISGMYMGEL
VRLILVKMAKEGLLFQGHTTPDLLTTGHFQTSFVSAIENDKDNEGLLSAEKVLRGLGLDPSAEDCVATQR
VCQIVSTRAAHLCAATLAAVLRQIRDNKAAERLRTTIGVDGSVYKNHPQFARRLHKMVRRLVPDCDVRFL
RSEDGSGKGAAMVTAVAYRLATQHAERQRILDTLRLSREQLLEVKRRMAEEMERGLAKETHDSATVKMLP
TYVRSTPDGTERGDFLALDLGGTNFRVLLVRVRSGKRRSVEMHNKIYSIPQEIMQGTGEELFDHIVHCIA
DFLEYMGMKGASLPLGFTFSFPCHQSKLDQGILLKWTKGFKATGCEGEDVVTLLKDAIHRREEFDLDVVA
VVNDTVGTMMTCGYEDPLCEVGLIVGTGTNACYMEEMQNVELVEGDEGRMCVNMEWGAFGDHGELDDFCT
EFDRAVDEHSANPGKQRYEKMISGMYLGEIVRNVLLDFTAKGLLFRGKLSERLKTRGIFETKFLSQIESD
RLALRQVRSILQHLGLTSSTCDDSILVKEVCSVVARRAAQLCGAGLAAVVDKIRQNRNLEQLRITVGVDG
TLYKLHPHFSTIMRETVRDLAPQCEVTFLQSEDGSGKGAALITAVACRIRDAGQH
>anc156_altAll|avgPP:0.991|lnPP:-16.366|num_ambig:10|num_ambig_gaps:0
LYHLRLSDENLMDVSKRFRREMDKGLGRDTNPTAAVKMLPTFVRSTPDGTEKGDFLALDLGGTNFRVLLV
KVSANGKQKVEMENQIYAIPEDLMRGSGTELFDHIAECLANFLEKLGIKDKKLPLGFTFSFPCQQTKLDE
SILVSWTKGFKASGVEGKDVVSLLRKAIKKRGDFDIDIVAVINDTVGTMMTCGYDDHHCEIGLIVGTGTN
ACYMEEMRHLELVEGDEGRMCVNMEWGAFGDDGALDDLRTDFDREIDAGSLNPGKQLFEKMISGMYMGEL
VRLILVKMAKEGLLFQGHTTPDLLTTGHFQTSFVSAIENDKDNEGLLSAEQVLRGLGLDPSAEDCVATQR
VCQIVSTRAAHLCAATLAAVLRQIRDNKAAERLRTTIGVDGSVYKNHPQFARRLHKMVRRLVPDCDVRFL
RSEDGSGKGAAMVTAVAYRLATQHAERQRILDTLRLSREQLLEVKRRMAEEMERGLAKETHDSASVKMLP
TYVRSTPDGTERGDFLALDLGGTNFRVLLVRVRSGKRRSVEMHNKIYSIPQETMQGTGEELFDHIVHCIA
DFLEYMGMKGASLPLGFTFSFPCHQSKLDQGILLKWTKGFKATGCEGEDVVTLLKDAIHRREEFDLDVVA
VVNDTVGTMMTCGYEDPLCEVGLIVGTGTNACYMEEMQNVELVEGDEGRMCVNMEWGAFGDHGELDDFCT
EFDRAVDERSANPGKQRYEKMISGMYLGEIVRNVLLDFTAKGLLFRGKLSERLKTRGIFETKFLSQIESD
RLALRQVRSILQHLGLTSSTCDDSILVKEVCSVVARRAAQLCGAGLAAVVDKIRQNRNLDQLKVTVGVDG
TLYKLHPHFSTIMRETVRDLAPQCEVTFLQSEDGSGKGAALITAVACRIRDAGQR
>anc157|avgPP:0.986|lnPP:-21.164|num_ambig:19|num_ambig_gaps:0
LYHLRLSDETLLDVSKRFRKEMEKGLGRDTNPTASVKMLPTFVRSTPDGTENGDFLALDLGGTNFRVLWV
KVSDNGKQKVEMENQIYAIPEDLMRGSGTQLFDHIAECLANFLEKLNIKDKKLPLGFTFSFPCQQTKLDE
SILVSWTKGFKASGVEGRDVVSLLRKAIKKRGDFDIDIVAVINDTVGTMMTCGYDDHNCEIGLIVGTGTN
ACYMEEMRHIDLVEGDEGRMCVNMEWGAFGDDGVLDDIRTEFDREIDMGSLNPGKQLFEKMISGMYMGEL
VRLILVKMAKEGLLFGGRITPDLLTTGHFETRFVSAIEKDKEGLQKAEEILSKLGLEPSQEDCVATQRIC
QIVSTRAANLCAATLAAVLRQIKENKGVERLRTTVGVDGSVYKKHPQFARRLHKMVRRLVPDCDVRFLRS
EDGSGKGAAMVTAVAYRLASQQAERQKTLDAFRLSREQLLEVKRRMREEMERGLKKETHDSATVKMLPTY
VRSTPDGTEKGDFLALDLGGTNFRVLLVRVRNGMRRSVEMHNKIYSIPQEVMQGTGEELFDHIVHCIADF
LEYMGMKGVSLPMGFTFSFPCHQNNLDEGILLKWTKGFKATGCEGEDVVTLLKDAIHRREEFDLDVVAVV
NDTVGTMMTCGYEDPYCEVGLIVGTGSNACYMEEMRNVELVEGDEGRMCVNMEWGAFGDNGCLDDFRTPF
DRAVDELSLNPGKQRYEKMISGMYLGEIVRNILIDFTKRGLLFRGRISERLKTRGIFETKFLSQIESDRL
ALRQVRSILQHLGLESTCDDSIIVKEVCTVVARRAAQLCGAGLAAVVDKIRENRGLDSLKVTVGVDGTLY

KLHPHFSTVMHETVRDLAPKCEVTFLQSEDGSGKGAALITAVACRIREAGQH
>anc157_altAll|avgPP:0.983|lnPP:-30.681|num_ambig:19|num_ambig_gaps:0
LYHLRLSDETLLDVSNRFRKEMEKGLGRDTNPTASVKMLPTFVRSTPDGTEKGDFLALDLGGTNFRVLRV
KVSDNGKQKVEMENQIYAIPEDLMRGSGTQLFDHIAECLANFLEKLEIKDKKLPLGFTFSFPCQQTKLDE
SILVSWTKGFKASGVEGRDVVSLLRKAIKKRGDFDIDIVAVINDTVGTMMTCGYDDHNCEIGLIVGTGTN
ACYMEEMRHIDLVEGDEGRMCVNMEWGAFGDDGVLDDIRTEFDREIDMGSLNPGKQLFEKMISGMYMGEL
VRLILVKMAKEGLLFGGKITPDLLTTGHFETSFVSAIEKDKEGLQKAEEILSKLGLDPSQEDCVATQRIC
QIVSERAANLCAATLAAVLRQIKENKGVERLRTTVGVDGSVYKKHPQFARRLHKMVRRLVPDCDVRFLRS
EDGSGKGAAMVTAVAYRLASQQAERQKTLDAFRLSREQLLEVKRRMREEMERGLKKETHDSATVKMLPTY
VRSTPDGTEKGDFLALDLGGTNFRVLLVRVRNGMRRGVEMHNKIYSIPQEAMQGTGEELFDHIVHCIADF
LEYMGMKGVSLPLGFTFSFPCHQNNLDEGILLKWTKGFKATGCEGEDVVTLLKDAIHRREEFDLDVVAVV
NDTVGTMMTCGYEDPYCEVGLIVGTGSNACYMEEMRNVELVEGEEGRMCVNMEWGAFGDNGCLDDFRTEF
DRAVDELSLNPGKQRYEKMISGMYLGEIVRNILIDFTKRGLLFRGRISERLKTRGIFETKFLSQIESDRL
ALRQVRSILQHLGLESTCDDSIIVKEVCTAVARRAAQLCGAGLAAVVDKIRENRGLDFLRITVGVDGTLY
KLHPHFSTIMHETVRDLAPKCEVTFLQSEDGSGKGAALITAVACRIREAGQR
>anc158|avgPP:0.990|lnPP:-16.304|num_ambig:18|num_ambig_gaps:0
LYHMRLSDETLLDVSNRFRKEMEKGLGRDTNPTASVKMLPTFVRSTPDGTENGDFLALDLGGTNFRVLRV
KVSDNGKQKVEMENQIYAIPEDLMRGSGTQLFDHIAECLANFLEKLNIKDKKLPLGFTFSFPCQQTKLDE
SILVTWTKGFKASGVEGRDVVSLLRKAIKKRGDFDIDIVAVVNDTVGTMMTCGYDDHNCEIGLIVGTGTN
ACYMEEMRHIDLVEGDEGRMCINMEWGAFGDDGVLDDIRTEFDREIDMGSLNPGKQLFEKMISGMYMGEL
VRLILVKMAKEGLLFGGKITPDLLTTGHFETRFVSAIEKDKEGLQKAHEILSKLGLEPSQEDCVATQRIC
QIVSTRAANLCAATLAAVLRQIKENKGVERLRTTVGVDGSVYKKHPQFARRLHKMVRRLVPDCDVRFLRS
EDGSGKGAAMVTAVAYRLASQQAERQKTLDAFRLSREQLLEVKRRMRVEMERGLKKETHNSATVKMLPTY
VRSTPDGTEKGDFLALDLGGTNFRVLLVRVRNGMRRGVEMHNKIYSIPQEVMQGTGEELFDHIVHCIADF
LEYMGMKGVSLPMGFTFSFPCHQNNLDEGILLKWTKGFKATGCEGEDVVTLLKEAIHRREEFDLDVVAVV
NDTVGTMMTCGYEDPYCEVGLIVGTGSNACYMEEMRNVELVEGEEGRMCINMEWGAFGDNGCLDDFRTPF
DRAVDELSLNPGKQRYEKMISGMYLGEIVRNILIDFTKRGLLFRGRISERLKTRGIFETKFLSQIESDRL
ALRQVRSILQHLGLESTCDDSIIVKEVCTVVARRAAQLCGAGLAAVVDKIRENRGLDFLKVTVGVDGTLY
KLHPHFSTVMHETVKDLAPKCEVTFLQSEDGSGKGAALITAVACRIREAGQR
>anc158_altAll|avgPP:0.987|lnPP:-23.179|num_ambig:18|num_ambig_gaps:0
LYHMRLSDETLLDVSNRFRKEMEKGLGRDTNPTASVKMLPTFVRSTPDGTEKGDFLALDLGGTNFRVLWV
KVSDNGKQKVEMENQIYAIPEDLMRGSGTQLFDHIAECLANFLEKLEIKDKKLPLGFTFSFPCQQTKLDE
SILVTWTKGFKASGVEGRDVVSLLRKAIKKRGDFDIDIVAVVNDTVGTMMTCGYDDHNCEIGLIVGTGTN
ACYMEEMRHIDLVEGDEGRMCINMEWGAFGDDGVLDDIRTEFDREIDMGSLNPGKQLFEKMISGMYMGEL
VRLILVKMAKEGLLFGGRITPDLLTTGHFETRFVSAIEKDKEGLQKANEILSKLGLDPSQEDCVATQRIC
QIVSERAANLCAATLAAVLRQIKENKGVERLRTTVGVDGSVYKKHPQFARRLHKMVRRLVPDCDVRFLRS
EDGSGKGAAMVTAVAYRLASQQAERQTTLDAFRLSREQLLEVKRRMRVEMERGLKKETHSSATVKMLPTY
VRSTPDGTEKGDFLALDLGGTNFRVLLVRVRNGMRRGVEMHNKIYSIPQEVMQGTGEELFDHIVHCIADF
LEYMGMKGVSLPLGFTFSFPCHQNNLDEGILLKWTKGFKATGCEGEDVVTLLKEAIHRREEFDLDVVAVV
NDTVGTMMTCGYEDPYCEVGLIVGTGSNACYMEEMRNVELVEGDEGRMCVNMEWGAFGDNGCLDDFRTEF
DRAVDELSLNPGKQRYEKMISGMYLGEIVRNILIDFTKRGLLFRGRISERLKTRGIFETKFLSQIESDRL
ALRQVRSILQHLGLESTCDDSIIVKEVCTAVARRAAQLCGAGLAAVVDKIRENRGLDFLKITVGVDGTLY
KLHPHFSTIMHETVRDLAPKCEVTFLQSEDGSGKGAALITAVACRIREAGQH
>anc159|avgPP:0.994|lnPP:-8.638|num_ambig:8|num_ambig_gaps:0
LYHMRLSDETLQDISNRFRKEMEKGLGRDTNPTASVKMLPTFVRSTPDGTENGDFLALDLGGTNFRVLRV
KVSDNGKQKVEMENQIYAIPEDLMRGSGTQLFDHIAECLANFMEKLNIKDKKLPLGFTFSFPCQQTKLDE
SILVTWTKGFKASGVEGRDVVSLLRKAIKKRGDFDIDIVAVVNDTVGTMMTCGYDDHNCEIGLIVGTGTN
ACYMEEMRHIDLVEGDEGRMCINMEWGAFGDDGVLDDIRTEFDREIDMGSLNPGKQLFEKMISGMYMGEL
VRLILVKMAKEGLLFGGKITPDLLTTGHFETRFVSAIEKDKEGLQKAHEILTKLGLEPSQEDCVATQRIC
QIVSTRSANLCAATLAAVLRRIKENKGVERLRTTVGVDGTVYKKHPQFARRLHKAVRRLVPDCDVRFLRS
EDGSGKGAAMVTAVAYRLASQQKARQKTLDAFKLSHEQLLEVKRRMRVEMERGLKKETHNSATVKMLPTY
VRSTPDGTEKGDFLALDLGGTNFRVLLVRVRNGMRRGVEMHNKIYSIPQEVMQGTGEELFDHIVHCIADF

LEYMGMKGVSLPLGFTFSFPCHQNNLDEGILLKWTKGFKATGCEGEDVVNLLKEAIHRREEFDLDVVAVV
NDTVGTMMTCGYEDPYCEVGLIVGTGSNACYMEEMRNVELVEGEEGRMCINMEWGAFGDNGCLDDFRTEF
DRAVDELSLNPGKQRYEKMISGMYLGEIVRNILIDFTKRGLLFRGRISERLKTRGIFETKFLSQIESDRL
ALLQVRSILQHLGLESTCDDSIIVKEVCTVVARRAAQLCGAGMAAVVDKIRENRGLDFLKVTVGVDGTLY
KLHPHFSTIMHETVKELAPKCEVTFLQSEDGSGKGAALITAVACRIREAGQR
>anc159_altAll|avgPP:0.992|lnPP:-13.108|num_ambig:8|num_ambig_gaps:0
LYHMRLSDETLQDISNRFRKEMEKGLGRDTNPTASVKMLPTFVRSTPDGTENGDFLALDLGGTNFRVLRV
KVSDNGKQKVEMENQIYAIPEDLMRGSGTQLFDHIAECLANFMEKLNIKDKKLPLGFTFSFPCQQTKLDE
SILVTWTKGFKASGVEGRDVVSLLRKAIKKRGDFDIDIVAVVNDTVGTMMTCGYDDHNCEIGLIVGTGTN
ACYMEEMRHIDLVEGDEGRMCINMEWGAFGDDGVLDDIRTEFDREIDMGSLNPGKQLFEKMISGMYMGEL
VRLILVKMAKEGLLFGGRITPDLLTTGHFETRFVSAIEKDKEGLQKAHEILTKLGLEPSQEDCVATQRIC
QIVSTRSANLCAATLAAVLRRIKENKGVERLRTTVGVDGTVYKKHPQFARRLHKAVRRLVPDCDVRFLRS
EDGSGKGAAMVTAVAYRLASQQKARQKTLDAFKLSHEQLLEVKKRMRIEMERGLKKETHSSATVKMLPTY
VRSTPDGTEKGDFLALDLGGTNFRVLLVRVRNGMRRGVEMHNKIYSIPQEVMQGTGEELFDHIVHCIADF
LEYMGMKGVSLPLGFTFSFPCHQNNLDEGILLKWTKGFKATGCEGEDVVNLLKEAIHRREEFDLDVVAVV
NDTVGTMMTCGYEDPYCEVGLIVGTGSNACYMEEMRNVELVEGDEGRMCVNMEWGAFGDNGCLDDFRTEF
DRAVDELSLNPGKQRYEKMISGMYLGEIVRNILIDFTKRGLLFRGRISERLKTRGIFETKFLSQIESDRL
ALLQVRSILQHLGLESTCDDSIIVKEVCTVVARRAAQLCGAGMAAVVDKIRENRGLDFLKVTVGVDGTLY
KLHPHFSTVMHETVKELAPKCEVTFLQSEDGSGKGAALITAVACRIREAGQH
>anc160|avgPP:0.989|lnPP:-16.871|num_ambig:13|num_ambig_gaps:0
LYHMRLSDEALHDISNRFLIEMKKGLRKDTNPTSSVKMLPTFVRSTPDGTESGDFLALDLGGTNFRVLRV
KVSDNGKQKVEIENRICAIPEDIMHGTGTQLFDHIANCLGDVMEELDIKDKKLPLGFTFSFPCQQSKLDE
SILVHWTKGFKASGVEGRDVVTLLRQAIQRRGDFDIDVVAVVNDTVGTMMSCGYDDHNCEIGLIVGTGTN
ACYMEEMQHIDLVEGDEGRMCINMEWGAFGDLGELDDIRTEFDREIDRGSINPGKQLFEKMISGMYMGEL
VRIILVKMAKDGLLFGGKISPELLIKGHFETKYVSAIEKDSDGLQKAHEILTKLGLEPTADDCAATQRVC
TIVSTRSSNLCAATLAAVLRRIKENKGVQRLRTTIGVDGTVYKKHPQFARRLHKAVRRLVPNCEVRFLPS
EDGSGKGAAMVTAVAYRLAKQQKVREETLAALKLSDEQLLEVKRRMRVEMERGLKKETHSTATVKMLATY
VRSTPDGTEKGDFLALDLGGTNFRVLLVKVRSGKRRAVEMHNKIYTIPMDAMEGTGEELFDHIVHCISDF
LEYMGMKGCVLPLGFTFSFPCEQRNLDQGILLKWTKGFKATGCEGEDVVDLLREAIKRREEFDLDVVAIV
NDTVGTMMSCGYDDPFCEVGLIVGTGCNACYMEEMRNVEMVEGDEGRMCVNMEWGAFGDNGCLDDIRTEF
DREVDKLSMNPGKQRYEKLISGMYLGEIVRNILIDFTKRGLLFRGRISERLKTRGIFQTKFLSQIESDRL
ALLQVRAILQQLGLQSTCDDSIIVKEVCGVVSRRAAQLCGAGMAAVVDKIRENRNLDHLKVTVGVDGTLY
KLHPHFSTIMHETVARLAPKCKVTFVQSEDGSGKGAALITAVACRIREAGQH
>anc160_altAll|avgPP:0.987|lnPP:-22.314|num_ambig:13|num_ambig_gaps:0
LYHMRLSDEVLHDISNRFLIEMEKGLRKDTNPTSSVKMLPTFVRSTPDGTESGDFLALDLGGTNFRVLRV
KVSDNGKQKVEIENRICAIPEDIMHGTGTQLFDHIANCLGDVMEELDIKDEKLPLGFTFSFPCQQSKLDE
SILVHWTKGFKASGVEGRDVVSLLRQAIQRRGDFDIDVVAVVNDTVGTMMSCGYDDHNCEIGLIVGTGTN
ACYMEEMQHIDLVEGDEGRMCINMEWGAFGDLGELDDIRTEFDREIDRGSINPGKQLFEKMISGMYMGEL
VRIILVKMAKDGLLFGGKISPELLIKGHFETKYVSAIEKDSDGLQKAHEILTKLGLEPTADDCAATQRVC
TIVSTRSANLCAATLAAVLRRIKENKGVERLRTTIGVDGTVYKKHPQFARRLHKAVRRLVPNCEVRFLPS
EDGSGKGAAMVTAVAYRLAKQQKMREEILAALKLSDEQLLEVKRRMRAEMERGLKKETHSSATVKMLATY
VRSTPDGTEKGDFLALDLGGTNFRVLLVKVRSGKRRAVEMHNKIYTIPMDAMEGTGEELFDHIVHCIADF
LEYMGMKGCVLPLGFTFSFPCEQRNLDQGILLKWTKGFKATGCEGEDVVDLLREAIKRREEFDLDVVAIV
NDTVGTMMSCGYDDPFCEVGLIVGTGCNACYMEEMRNVEMVEGDEGRMCVNMEWGAFGDNGCLDDIRTEF
DREVDKLSMNPGKQRYEKLISGMYLGEIVRNILINFTKRGLLFRGRISERLKTRGIFQTKFLSQIESDRL
ALLQVRAILQQLGLQSTCDDSIIVKEVCGVVSRRAAQLCGAGLAAVVDKIRENRNLDHLKVTVGVDGTLY
KLHPHFSTIMHETVARLAPKCKVTFVQSEDGSGKGAALITAVACRIREAGQH
>anc161|avgPP:0.988|lnPP:-19.354|num_ambig:14|num_ambig_gaps:0
LYHMRLSDEALHDISKRFLTEMKKGLRKDTNPTSSVKMLPSFVRSTPDGTESGDFLALDLGGTNFRVLRV
KVSDNGKQKVEIENRICAIPEDIMHGTGTQLFDHIANCLGDVMEELDIKDKKLPLGFTFSFPCQQSKLDE
SILVNWTKGFKASGVEGRDVVGLLRQAIQRRGDFDIDVVAVVNDTVGTMMSCGYDDHNCEIGLIVGTGTN
ACYMEEMQHIDLVEGDEGRMCINMEWGAFGDLGELDDIRTEFDREIDRGSINPGKQLFEKMISGMYMGEL

VRLILVKMAKDGLLFRGKITPELLIKGHFETKYVSAIEKDSDGLQKAHEILTKLGLEPTADDCAATQRVC
SIVSTRSSNLCAATLAAVLRRIKENKGVQRLRTTIGVDGTVYKKHPQFARRLHKAVRRLVPNCEVRFLPS
EDGSGKGAAMVTAVAYRLAKQQKMREEILSAFKLSDEQLLEVKRRMRVEMERGLKKETHSTATVKMLATY
VRATPDGTEKGDFLALDLGGTNFRVLLVKVRGGKRRAVEMHNKIYTIPMDAMEGTGEELFDHIVHCIADF
LEYMGMKGCRLPLGFTFSFPCEQRNLDQGILLKWTKGFKATGCEGEDVVDLLREAIKRREEFDLDVVAIV
NDTVGTMMSCGYDDPLCEVGLIVGTGCNACYMEEMRNVEMVEGDEGRMCVNMEWGAFGDNGCLDDIRTEF
DMEVDKLSLNPGKQRYEKMISGMYLGEIVRNILIKFTKQGLLFRGRISERLKTRGIFQTKFLSQIESDRL
ALLQVRAILQQLGLQSTCDDSIIVKEVCGVVSRRAAQLCGAGLAAIVDKIRENRNLDHLKVTVGVDGTLY
KLHPHFSTIMHETVERLSPKCKVTFVQSEDGSGKGAALITAVACRIREAGQH
>anc161_altAll|avgPP:0.986|lnPP:-22.866|num_ambig:14|num_ambig_gaps:0
LYHMRLSDEVLHDISNRFLTEMKKGLRKDTNPTSSVKMLPSFVRSTPDGTESGDFLALDLGGTNFRVLRV
KVSDNGKQKVEIENRICAIPEDIMHGTGTQLFDHIANCLGDVMEELDIKDEKLPLGFTFSFPCQQSKLDE
SILVNWTKGFKASGVEGRDVVGLLRKAIQKRGDFDIDVVAVVNDTVGTMMSCGYDDHNCEIGLIVGTGTN
ACYMEEMQHIDLVEGDEGRMCINMEWGAFGDLGELDDIRTEFDKEIDRGSINPGKQLFEKMISGMYMGEL
VRIILVKMAKDGLLFRGKITPELLIKGHFETKYVSAIEKDSDGLQKAHEILTKLGLEPTADDCAATKRVC
SIVSTRSSNLCAATLAAVLRRIKENKGVERLRTTIGVDGTVYKKHPQFARRLHKAVRRLVPNCEVRFLPS
EDGSGKGAAMVTAVAYRLAKQQKMREETLSAFKLSDEQLLEVKRRMRVEMERGLKKETHSTATVKMLATY
VRATPDGTEKGDFLALDLGGTNFRVLLVKVRSGKRRAVEMHNKIYTIPMDAMEGTGEELFDHIVHCISDF
LEYMGMKGCRLPLGFTFSFPCEQRNLDQGILLKWTKGFKATGCEGEDVVDLLREAIKRREEFDLDVVAIV
NDTVGTMMSCGYDDPLCEVGLIVGTGCNACYMEEMRNVEMVEGDEGRMCVNMEWGAFGDNGCLDDIRTEF
DMEVDKLSLNPGKQRYEKMISGMYLGEIVRNILIKFTKRGLLFRGRISERLKTRGIFQTKFLSQIESDRL
ALLQVRAILQQLGLQSTCDDSIIVKEVCGVVSRRAAQLCGAGLAAIVDKIRENRNLDNLKVTVGVDGTLY
KLHPHFSTIMHETVERLSPKCKVTFVQSEDGSGKGAALITAVACRIREAGQH
>anc162|avgPP:0.988|lnPP:-18.838|num_ambig:18|num_ambig_gaps:0
LYHMRLSDEALHDISNRFLIEMKKGLRKDTNPTSSVKMLPTFVRSTPDGTESGDFLALDLGGTNFRVLRV
KVSDNGKQKVEIENRICAIPEDIMHGTGTQLFDHIANCLGDVMEELDIKDKKLPLGFTFSFPCQQSKLDE
SILVNWTKGFKASGVEGRDVVTLLRQAIQRRGDFDIDVVAVVNDTVGTMMSCGYDDHNCEIGLIVGTGTN
ACYMEEMQHIDLVEGDEGRMCINMEWGAFGDLGELDDIRTEFDREIDRGSINPGKQLFEKMISGMYMGEL
VRIILVKMAKDGLLFGGKITPELLIKGHFETKYVSAIEKDSDGLQKAHEILTKLGLEPTADDCAATQRVC
TIVSTRSSNLCAATLAAVLRRIKENKGVQRLRTTIGVDGTVYKKHPQFARRLHKAVRRLVPNCEVRFLPS
EDGSGKGAAMVTAVAYRLAKQQKMREETLAAFKLSDEQLLEVKRRMRVEMERGLKKETHSTATVKMLATY
VRSTPDGTEKGDFLALDLGGTNFRVLLVKVRSGKRRAVEMHNKIYTIPMDAMEGTGEELFDHIVHCISDF
LEYMGMKGCRLPLGFTFSFPCEQRNLDQGILLKWTKGFKATGCEGEDVVDLLREAIKRREEFDLDVVAIV
NDTVGTMMSCGYDDPFCEVGLIVGTGCNACYMEEMRNVEMVEGDEGRMCVNMEWGAFGDNGCLDDIRTEF
DREVDKLSMNPGKQRYEKMISGMYLGEIVRNILIDFTKRGLLFRGRISERLKTRGIFQTKFLSQIESDRL
ALLQVRAILQQLGLQSTCDDSIIVKEVCGVVSRRAAQLCGAGMAAVVDKIRENRNLDHLKVTVGVDGTLY
KLHPHFSTIMHETVARLAPKCKVTFVQSEDGSGKGAALITAVACRIREAGQH
>anc162_altAll|avgPP:0.985|lnPP:-25.445|num_ambig:18|num_ambig_gaps:0
LYHMRLSDEVLHDISKRFLTEMEKGLRKDTNPTSSVKMLPTFVRSTPDGTESGDFLALDLGGTNFRVLRV
KVSDNGKQKVEIENRICAIPEDIMHGTGTQLFDHIANCLGDVMEELDIKDEKLPLGFTFSFPCQQSKLDE
SILVNWTKGFKASGVEGRDVVSLLRQAIQKRGDFDIDVVAVVNDTVGTMMSCGYDDHNCEIGLIVGTGTN
ACYMEEMQHIDLVEGDEGRMCINMEWGAFGDLGELDDIRTEFDREIDRGSINPGKQLFEKMISGMYMGEL
VRLILVKMAKDGLLFGGKISPELLIKGHFETKYVSAIEKDSDGLQKAHEILTKLGLEPTADDCAATQRVC
TIVSTRSANLCAATLAAVLRRIKENKGVERLRTTIGVDGTVYKKHPQFARRLHKAVRRLVPNCEVRFLPS
EDGSGKGAAMVTAVAYRLAKQQKVREEILAAFKLSDEQLLEVKRRMRVEMERGLKKETHSSATVKMLATY
VRSTPDGTEKGDFLALDLGGTNFRVLLVKVRSGKRRAVEMHNKIYTIPMDAMEGTGEELFDHIVHCIADF
LEYMGMKGCRLPLGFTFSFPCEQRNLDQGILLKWTKGFKATGCEGEDVVDLLREAIKRREEFDLDVVAIV
NDTVGTMMSCGYDDPFCEVGLIVGTGCNACYMEEMRNVEMVEGDEGRMCVNMEWGAFGDNGCLDDIRTEF
DREVDKLSLNPGKQRYEKMISGMYLGEIVRNILINFTKRGLLFRGRISERLKTRGIFQTKFLSQIESDRL
ALLQVRAILQQLGLQSTCDDSIIVKEVCGVVSRRAAQLCGAGLAAVVDKIRENRNLDHLKVTVGVDGTLY
KLHPHFSTIMHETVARLAPKCKVTFVQSEDGSGKGAALITAVACRIREAGQH
>anc163|avgPP:0.985|lnPP:-22.631|num_ambig:22|num_ambig_gaps:0

LYHMRLSDETLQDISNRFLTEMEKGLGRDTNPTSSVKMLPTFVRSTPDGTESGDFLALDLGGTNFRVLRV
KVSDNGKQKVEMENQIYAIPEDIMRGSGTQLFDHIADCLGNFMEKLDIKDKKLPLGFTFSFPCQQTKLDE
SILVNWTKGFKASGVEGRDVVSLLRQAIQKRGDFDIDIVAVVNDTVGTMMTCGYDDHNCEIGLIVGTGTN
ACYMEEMRHIDLVEGDEGRMCINMEWGAFGDQGALDDIRTEFDREIDRGSLNPGKQLFEKMISGMYMGEL
VRLILVKMAKEGLLFGGKITPELLIKGHFETKYVSAIEKDSDGLQKAHEILTKLGLEPSQEDCVATQRIC
TIVSTRSANLCAATLAAVLRRIKENKGVQRLRTTIGVDGTVYKKHPQFARRLHKAVRRLVPDCDVRFLRS
EDGSGKGAAMVTAVAYRLASQQKAREETLAAFKLSHEQLLEVKRRMRVEMERGLKKETHSSATVKMLATY
VRSTPDGTEKGDFLALDLGGTNFRVLLVKVRNGMRRGVEMHNKIYAIPMEAMEGTGEELFDHIVHCISDF
LEYMGMKGVRLPLGFTFSFPCEQRNLDQGILLKWTKGFKATGCEGEDVVDLLREAIKRREEFDLDVVAVV
NDTVGTMMSCGYDDPYCEVGLIVGTGSNACYMEEMRNVEMVEGDEGRMCINMEWGAFGDNGCLDDIRTEF
DREVDKLSMNPGKQRYEKMISGMYLGEIVRNILIDFTKRGLLFRGRISERLKTRGIFETKFLSQIESDRL
ALLQVRAILQQLGLESTCDDSIIVKEVCGVVARRAAQLCGAGMAAVVDKIRENRNLDHLKVTVGVDGTLY
KLHPHFSTIMHETVAKLAPKCKVTFVQSEDGSGKGAALITAVACRIREAGQH
>anc163_altAll|avgPP:0.981|lnPP:-33.545|num_ambig:22|num_ambig_gaps:0
LYHMRLSDETLQDISKRFLVEMEKGLGRDTNPTSSVKMLPTFVRSTPDGTESGDFLALDLGGTNFRVLRV
KVSDNGKQKVEMENQIYAIPEDLMRGSGTQLFDHIADCLGDFMEKLNIKDKKLPLGFTFSFPCQQTKLDE
SILVNWTKGFKASGVEGRDVVTLLRKAIQKRGDFDIDIVAVVNDTVGTMMTCGYDDHNCEIGLIVGTGTN
ACYMEEMRHIDLVEGDEGRMCINMEWGAFGDQGALDDIRTEFDREIDRGSLNPGKQLFEKMISGMYMGEL
VRLILVKMAKEGLLFGGKISPELLIKGHFETKYVSAIEKDSDGLQKAHEILTKLGLEPSQEDCIATQRIC
TIVSTRSANLCAATLAAVLRRIKENKGVERLRTTIGVDGTVYKKHPQFARRLHKAVRRLVPDCDVRFLRS
EDGSGKGAAMVTAVAYRLAAQRKAREETLAAFKLSHEQLLEVKRRMRVEMERGLKKETHSSATVKMLATY
VRSTPDGTEKGDFLALDLGGTNFRVLLVKVRSGKRRGVEMHNKIYAIPTEAMEGTGEELFDHIVHCIADF
LEYMGMKGVRLPLGFTFSFPCEQRNLDQGILLKWTKGFKATGCEGEDVVDLLREAIRRREEFDLDVVAVV
NDTVGTMMSCGYDDPHCEVGLIVGTGSNACYMEEMRNVEMVEGDEGRMCVNMEWGAFGDNGCLDDIRTEF
DREVDKLSLNPGKQRYEKMISGMYLGEIVRNILIDFTKRGLLFRGRISERLKTRGIFETKFLSQIESDRL
ALLQVRAILQQLGLESTCDDSIIVKEVCGVVSRRAAQLCGAGMAAVVDKIRENRNLDHLKITVGVDGTLY
KLHPHFSTIMHETVAKLAPKCKVTFVQSEDGSGKGAALITAVACRIREAGQH
>anc164|avgPP:0.986|lnPP:-22.099|num_ambig:20|num_ambig_gaps:0
LYHMRLSDETLQDISNRFRTEMEKGLGRDTNPTASVKMLPTFVRSTPDGTENGDFLALDLGGTNFRVLRV
KVSDNGKQKVEMENQIYAIPEDIMRGSGTQLFDHIAECLGNFMEKLNIKDKKLPLGFTFSFPCQQTKLDE
SILVTWTKGFKASGVEGRDVVSLLRKAIQKRGDFDIDIVAVVNDTVGTMMTCGYDDHNCEIGLIVGTGTN
ACYMEEMRHIDLVEGDEGRMCINMEWGAFGDDGALDDIRTEFDREIDRGSLNPGKQLFEKMISGMYMGEL
VRLILVKMAKEGLLFGGKITPELLTKGHFETKYVSAIEKDKEGLQKAHEILTKLGLEPSQEDCVATQRIC
TIVSTRSANLCAATLAAVLRRIKENKGVQRLRTTVGVDGTVYKKHPQFARRLHKAVRRLVPDCDVRFLRS
EDGSGKGAAMVTAVAYRLASQQKAREETLAAFKLSHEQLLEVKRRMRVEMERGLKKETHSSATVKMLPTY
VRSTPDGTEKGDFLALDLGGTNFRVLLVKVRNGMRRGVEMHNKIYAIPQEVMQGTGEELFDHIVHCISDF
LEYMGMKGVRLPLGFTFSFPCHQNNLDQGILLKWTKGFKATGCEGEDVVNLLREAIKRREEFDLDVVAVV
NDTVGTMMTCGYEDPYCEVGLIVGTGSNACYMEEMRNVEMVEGDEGRMCINMEWGAFGDNGCLDDIRTEF
DRAVDELSLNPGKQRYEKMISGMYLGEIVRNILIDFTKRGLLFRGRISERLKTRGIFETKFLSQIESDRL
ALLQVRAILQQLGLESTCDDSIIVKEVCGVVARRAAQLCGAGMAAVVDKIRENRGLDHLKVTVGVDGTLY
KLHPHFSTIMHETVKELAPKCEVTFLQSEDGSGKGAALITAVACRIREAGQH
>anc164_altAll|avgPP:0.982|lnPP:-31.610|num_ambig:20|num_ambig_gaps:0
LYHMRLSDETLQDISNRFRKEMEKGLGRDTNPTASVKMLPTFVRSTPDGTENGDFLALDLGGTNFRVLRV
KVSDNGKQKVEMENQIYAIPEDLMRGSGTQLFDHIAECLGDFMEKLNIKDKKLPLGFTFSFPCQQTKLDE
SILVTWTKGFKASGVEGRDVVSLLRKAIKKRGDFDIDIVAVVNDTVGTMMTCGYDDHNCEIGLIVGTGTN
ACYMEEMRHIDLVEGDEGRMCINMEWGAFGDDGALDDIRTEFDREIDRGSLNPGKQLFEKMISGMYMGEL
VRLILVKMAKEGLLFGGKITPELLTKGHFETKYVSAIEKDKEGLQKAHEILTKLGLEPSQEDCIATQRIC
TIVSTRSANLCAATLAAVLRRIKENKGVERLRTTVGVDGTVYKKHPQFARRLHKAVRRLVPDCDVRFLRS
EDGSGKGAAMVTAVAYRLASQRKAREETLAAFKLSHEQLLEVKKRMRVEMERGLKKETHNSATVKMLPTY
VRSTPDGTEKGDFLALDLGGTNFRVLLVKVRSGKRRGVEMHNKIYSIPMEVMQGTGEELFDHIVHCIADF
LEYMGMKGVRLPLGFTFSFPCHQTNLDQGILLKWTKGFKATGCEGEDVVNLLREAIRRREEFDLDVVAVV
NDTVGTMMTCGYEDPHCEVGLIVGTGSNACYMEEMRNVEMVEGDEGRMCVNMEWGAFGDNGCLDDIRTEF

DRAVDELSLNPGKQRYEKMISGMYLGEIVRNILIDFTKRGLLFRGRISERLKTRGIFETKFLSQIESDRL
ALLQVRAILQQLGLESTCDDSIIVKEVCGVVSRRAAQLCGAGMAAVVDKIRENRGLDHLKITVGVDGTLY
KLHPHFSTIMHETVKELAPKCEVTFLQSEDGSGKGAALITAVACRIREAGQH
>anc165|avgPP:0.975|lnPP:-39.769|num_ambig:40|num_ambig_gaps:0
LYHMRLSDETLQDIMNRFRTEMEKGLGRDTNPTATVKMLPTFVRSTPDGTEKGDFLALDLGGSNFRVLRV
KVSDDGKQKVEMESQIYAIPEDIMRGSGTQLFDHVAECLGNFMEKLQIKDKKLPLGFTFSFPCRQTKLDE
SILITWTKGFKASGVEGRDVVKLLRKAIKKRGDFDIDIVAVVNDTVGTMMTCGYDDHRCEIGLIVGTGTN
ACYMEEMRHIDLVEGDEGRMCINMEWGAFGDDGSLDDIRTEFDREIDRGSLNPGKQLFEKMISGMYMGEL
VRLILVKMAKEGLLFGGKITPELLTKGHFETKYVSAIEKDKEGLSKAKEILTKLGLEPSEEDCIAVQRIC
TIVSTRSANLCAATLAAVLTRIKENKGVKRLRTTVGVDGTVYKTHPQFARRLHKTVRRLVPDCDVRFLLS
EDGSGKGAAMVTAVAYRLASQRKQIDETLAPFKLSHEQLLEVKKRMRTEMERGLKKETHSSATVKMLPTY
VRSTPDGTEKGDFLALDLGGTNFRVLLVKIRSGKRRGVEMHNKIYAIPQEVMQGTGEELFDHIVHCISDF
LDYMGMKGARLPLGFTFSFPCRQTSLDQGILLNWTKGFKATGCEGEDVVNLLREAIKRREEFDLDVVAVV
NDTVGTMMTCAYEDPNCEIGLIVGTGSNACYMEEMRNVEMVEGDEGRMCINMEWGAFGDNGCLDDIRTEF
DRAVDELSLNPGKQRYEKMISGMYLGEIVRNILIDFTKRGLLFRGRISERLKTRGIFETKFLSQIESDRL
ALLQVRAILQQLGLESTCDDSIIVKEVCGVVSRRAAQLCGAGMAAVVDKIRENRGLDHLKITVGVDGTLY
KLHPHFSKIMHETVKELAPKCDVTFLQSEDGSGKGAALITAVACRLREAGQH
>anc165_altAll|avgPP:0.967|lnPP:-61.155|num_ambig:40|num_ambig_gaps:0
LYHMRLSDETLQDIMNRFRTEMEKGLGRDTNPTATVKMLPTFVRSTPDGSEKGDFLALDLGGTNFRVLRV
KVSDDGKQKVEMENQIYAIPEDIIHGSGTQLFDHVAECLGDFMEKQQIKDKKLPLGFTFSFPCRQTKLDE
SVLITWTKGFKASGVEGRDVVSLLRKAIQKRGDYDVDIVAVVNDTVGTMMTCGYDDHNCEIGLIVGTGTN
ACYMEEMRHIDLVEGDEGRMCINMEWGAFGDDGALDDIRTEFDREIDRGSLNPGKQLFEKMISGMYMGEL
VRLILVKMAKEGLLFGGKITPELLTKGHFETKYVSAIEKDKEGLSKAKEILTKLGLEPSEEDCVAVQHIC
TIVSTRSANLCAAALAAVLTRLRENKGVQRLRTTVGVDGTVYKKHPQYARRLHKTVRRLVPDCDVRFLLS
EDGSGKGAAMVTAVAYRLASQRKQIDETLAPFKLSHEQLLEVKRRMRTEMERGLKKETHNTATVKMLPTY
VRSTPDGTEKGDFLALDLGGTNFRVLLVKVRSGKRRSVEMHNKIYSIPQEVMQGTGEELFDHIVHCISDF
LEYMGMKGVRLPLGFTFSFPCKQTSLDQGILLNWTKGFKATGCEGEDVVNLLREAIKRREEFELDVVAIV
NDTVGTMMTCAYEDPNCEVGLIVGTGSNACYMEEMRNVEMVEGDEGRMCINMEWGAFGDNGCLDDIRTEF
DRAVDELSLNPGKQRYEKMISGMYLGEIVRNILIDFTKRGLLFRGQISERLKTRGIFETKFLSQIESDRL
ALLQVRAILQQLGLDSTCDDSIIVKEVCGVVSRRAAQLCGAGMAAVVDKIRENRGLDHLKVTVGVDGTLY
KLHPHFSRIMHETVKELAPKCEVTFLQSEDGSGKGAALITAVACRIREAGQH
>anc166|avgPP:0.994|lnPP:-9.758|num_ambig:5|num_ambig_gaps:0
LQQFMVTEAQLQQIQASLLGSMEQALMGQSSPAPAVRMLPTYVGSTPHGTEQGDFVVLELGATGASLRVL
WVTLTGTEGHRVEPRSREFVIPPEVMLGPGQQLFDFAARCLAEFLEALPVGNRCLPLGFNFSFPCHQTGL
DRSTLISWTKGFRCSGVEGQDVVQLLREAIERQGASSLDVVAVVNDTVGTMMGCDPGLGPCEVGLVVDTG
TNACYMEEARHVAALDEDRGRVCVSVEWGSFGDDKALGPVLTPFDHALDQESLNPGAQRFEKMIGGLYLG
ELVRLVLAHLAQRGLLFGGCTSPALLSQGSVGLEHVAEMEDPSAGAARVLAVLQNLGLSPGASDVELVQH
VCEAVSTRAARLCAAALAAVLGRLQHSREQHALQIAVATGGRVFERHPRFHSLLQETVMLLAPECDVSFI
PSVDGGGRGVAVVTAVAARLAAHRRLLEETLAPFRLDREQLAAVQAQMREAMAKGLRGEASSLRMLPTYV
RATPDGSERGDFLALDLGGTNFRVLLVRVAAGGVQITSQVYAIPEQVAQGSGQQLFDHIVDCIVDFQQRQ
GLSGQSLPLGFTFSFPCRQLGLDQGILLNWTKGFNASDCEGQDVVALLREAIGRRQAVELNVVAIVNDTV
GTMMSCGYEDHRCEVGLIVGTGTNACYMEELQNVAGVPGDSGHMCINMEWGAFGDDGTLSVPSTCFDASV
DRASINPGKQRFEKLISGMYLGEIVRHILLHLNSLGVLFRGQQTQCLQTRDIFKTKFLSEIESDSLALRQ
VRAILEDLGLPLTSDDALVVREVCQAVSRRAAQLCGAGVAAVVEKIRENRGLEELTVTVGVDGTLYKLHP
HFSSLVAATVRELAPRCAVTFLQSEDGSGKGAALVTAVACRLAQTTRV
>anc166_altAll|avgPP:0.993|lnPP:-12.333|num_ambig:5|num_ambig_gaps:0
LQQFMVTEAQLQQIQASLLGSMEQALMGQNGPAPAVRMLPTYVGSTPHGTEQGDFVVLELGATGASLRVL
WVTLTGTEGHRVEPRSREFVIPPEVMLGPGQQLFDFAARCLAEFLEALPVGNRCLPLGFNFSFPCHQTGL
DRSTLISWTKGFRCSGVEGQDVVQLLREAIERQGASSLAVVAVVNDTVGTMMGCDPGLGPCEVGLVVDTG
TNACYMEEARHVAALDEDRGRVCVSVEWGSFGDDKALGPVLTPFDHALDQESLNPGAQRFEKMIGGLYLG
ELVRLVLAHLAQRGLLFGGCTSPALLSQGSVGLEHVAEMEDPSAGAARVLAVLQNLGLSPGASDVELVQH
VCEAVSTRAARLCAAALAAVLGRLQHSREQHALQIAVATGGRVFERHPRFHSLLQETVMLLAPECDVSFI

PSVDGGGRGVAVVTAVAARLAAHRRLLEETLAPFRLDREQLAAVQVQMREAMAKGLRGEASSLRMLPTYV
RATPDGSERGDFLALDLGGTNFRVLLVRVAAGGVQITSQVYAIPERVAQGSGQQLFDHIVDCIVDFQQRQ
GLSGQSLPLGFTFSFPCRQLGLDQGILLNWTKGFNASDCEGQDVVALLREAIGRRQAVELNVVAIVNDTV
GTMMSCGYEDHRCEVGLIVGTGTNACYMEELQNVAGVPGDSGHMCINMEWGAFGDDGTLSVPSTCFDASV
DRASINPGKQRFEKLISGMYLGEIVRHILLHLNSLGVLFRGQQTQCLQTRDIFKTKFLSEIESDSLALRQ
VRAILEDLGLPLTSDDALVVREVCQAVSRRAAQLCGAGVAAVVEKIRENRGLEELTVTVGVDGTLYKLHP
HFSSLVAATVRELAPRCAVTFLQSEDGSGKGAALVTAVACRLAQTTRV
>anc167|avgPP:0.995|lnPP:-7.159|num_ambig:5|num_ambig_gaps:0
LQQFMVTEAQLQQIQASLLGSMEQALMGQSSPAPAVRMLPTYVGSTPHGTEQGDFVVLELGATGASLRVL
WVTLTGTEGHRVEPRSREFVIPPEVMLGPGQQLFDFAARCLAEFLEALPVGNRCLPLGFNFSFPCHQTGL
DRSTLISWTKGFRCSGVEGQDVVQLLREAIERQGASSLDVVAVVNDTVGTMMGCDPGLGPCEVGLVVDTG
TNACYMEEARHVAALDEDRGRVCVSVEWGSFGDDEALGPVLTPFDHALDQESLNPGAQRFEKMIGGLYLG
ELVRLVLAHLAQRGLLFGGCTSPALLSQGSVTLEHVAEMEDPSAGAARVLAVLQNLGLSPGASDVELVQH
VCEAVSTRAARLCAAALAAVLGRLQHSREQHTLQIAVATGGRVFERHPRFHSILQETVMLLAPECDVSFI
PSVDGGGRGVAMVTAVAARLAAHRRLLEETLAPFRLDREQLAAVQAQMREAMAKGLRGEASSLRMLPTYV
RATPDGSERGDFLALDLGGTNFRVLLVRVAAGGVQITSQVYAIPEQVAQGSGQQLFDHIVDCIVDFQQRQ
GLSGQSLPLGFTFSFPCRQLGLDQGILLNWTKGFNASDCEGQDVVALLREAIGRRQAVELNVVAIVNDTV
GTMMSCGYEDHRCEVGLIVGTGTNACYMEELQNVAGVPGDSGHMCINMEWGAFGDDGTLDVPSTCFDASV
DRASINPGKQRFEKLISGMYLGEIVRHILLHLNSLGVLFRGQQTQCLQTRDIFKTKFLSEIESDSLALRQ
VRAILEDLGLPLTSDDALVVREVCQAVSRRAAQLCGAGVAAVVEKIRENRGLEELTVTVGVDGTLYKLHP
HFSSLVAATVRELAPRCAVTFLQSEDGSGKGAALVTAVACRLAQTTRV
>anc167_altAll|avgPP:0.994|lnPP:-9.544|num_ambig:5|num_ambig_gaps:0
LQQFMVTEAQLQQIQASLLGSMEQALMGQSGPAPAVRMLPTYVGSTPHGTEQGDFVVLELGATGASLRVL
WVTLTGTEGHRVEPRSREFVIPPEVMLGPGQQLFDFAARCLAEFLEALPVGNRCLPLGFNFSFPCHQTGL
DRSTLISWTKGFRCSGVEGQDVVQLLREAIERQGASNLDVVAVVNDTVGTMMGCDPGLGPCEVGLVVDTG
TNACYMEEARHVAVLDEDRGRVCVSVEWGSFGDDEALGPVLTPFDHALDQESLNPGAQRFEKMIGGLYLG
ELVRLVLAHLAQRGLLFGGCTSPALLSQGSVTLEHVAEMEDPSAGAARVLAVLQNLGLSPGASDVELVQH
VCEAVSTRAARLCAAALAAVLGRLQHSREQHTLQIAVATGGRVFERHPRFHSILQETVMLLAPECDVSFI
PSVDGGGRGVAMVTAVAARLAAHRRLLEETLAPFRLDREQLAAVQVQMREAMAKGLRGEASSLRMLPTYV
RATPDGSERGDFLALDLGGTNFRVLLVRVAAGGVQITSQVYAIPERVAQGSGQQLFDHIVDCIVDFQQRQ
GLSGQSLPLGFTFSFPCRQLGLDQGILLNWTKGFNASDCEGQDVVALLREAIGRRQAVELNVVAIVNDTV
GTMMSCGYEDHRCEVGLIVGTGTNACYMEELQNVAGVPGDSGHMCINMEWGAFGDDGTLDVPSTCFDASV
DRASINPGKQRFEKLISGMYLGEIVRHILLHLNSLGVLFRGQQTQCLQTRDIFKTKFLSEIESDSLALRQ
VRAILEDLGLPLTSDDALVVREVCQAVSRRAAQLCGAGVAAVVEKIRENRGLEELTVTVGVDGTLYKLHP
HFSSLVAATVRELAPRCAVTFLQSEDGSGKGAALVTAVACRLAQTTRV
>anc168|avgPP:0.995|lnPP:-6.511|num_ambig:5|num_ambig_gaps:0
LQQFMVTEAQLQQIQASLLGSMEQALMGQSSPAPAVRMLPTYVGSTPHGTEQGDFVVLELGATGASLRVL
WVTLTGTEGHRVEPRSQEFVIPPEVMLGPGQQLFDFAARCLAEFLEALPVGNQCLPLGFNFSFPCHQTGL
DRSTLISWTKGFRCSGVEGQDVVQLLREAIERQGASSLDVVAVVNDTVGTMMGCDPGLGPCEVGLVVDTG
TNACYMEEARHVAALDEDRGRVCVSVEWGSFGDDEALGPVLTPFDHALDQESLNPGAQRFEKMIGGLYLG
ELVRLVLAHLAQRGLLFGGCTSPALLSQGSVTLEHVAEMEDPSAGAARVLAVLQNLGLSPGASDVELVQH
VCAAVSTRAARLCAAALAAVLGRLQHSREQHTLQIAVATGGRVFERHPRFHSILQETVMLLAPECDVSFI
PSVDGGGRGVAMVTAVAARLAAHRRLLEETLAPFRLDREQLAAVQAQMREAMAKGLRGEASSLRMLPTYV
RATPDGSERGDFLALDLGGTNFRVLLVRVAAGGVQITSQVYAIPECVAQGSGQQLFDHIVDCIVDFQQRQ
GLSGQSLPLGFTFSFPCRQLGLDQGILLNWTKGFNASDCEGQDVVALLREAIGRRQAVELNVVAIVNDTV
GTMMSCGYEDHRCEVGLIVGTGTNACYMEELQNVAGVPGDSGHMCINMEWGAFGDDGTLDVPSTCFDASV
DRASINPGKQRFEKLISGMYLGEIVRHILLHLNSLGVLFRGQQTQCLQTRDIFKTKFLSEIESDSLALRQ
VRAILEDLGLPLTSDDALVVREVCQAVSRRAAQLCGAGVAAVVEKIRENRGLEELTVTVGVDGTLYKLHP
HFSSLVAATVRELAPRCAVTFLQSEDGSGKGAALVTAVACRLAQTTRV
>anc168_altAll|avgPP:0.994|lnPP:-9.286|num_ambig:5|num_ambig_gaps:0
LQQFMVTEAQLQQIQASLLGSMEQALMGQSGPAPAVRMLPTYVGSTPHGTEQGDFVVLELGATGASLRVL
WVTLTGTEGHRVEPRSQEFVIPPEVMLGPGQQLFDFAARCLAEFLEALPVGNQCLPLGFNFSFPCHQTGL

DRSTLISWTKGFRCSGVEGQDVVQLLREAIERQGASNLDVVAVVNDTVGTMMGCDPGLGPCEVGLVVDTG
TNACYMEEARHVAVLDEDRGRVCVSVEWGSFGDDEALGPVLTPFDRALDQESLNPGAQRFEKMIGGLYLG
ELVRLVLAHLAQRGLLFGGCTSPALLSQGSVTLEHVAEMEDPSAGAARVLAVLQNLGLSPGASDVELVQH
VCAAVSTRAARLCAAALAAVLGRLQHSREQHTLQIAVATGGRVFERHPRFHSILQETVMLLAPECDVSFI
PSVDGGGRGVAMVTAVAARLAAHRRLLEETLAPFRLDREQLAAVQVQMREAMAKGLRGEASSLRMLPTYV
RATPDGSERGDFLALDLGGTNFRVLLVRVAAGGVQITSQVYAIPECVAQGSGQQLFDHIVDCIVDFQQRQ
GLSGQSLPLGFTFSFPCRQLGLDQGILLNWTKGFNASDCEGQDVVALLREAIGRRQAVELNVVAIVNDTV
GTMMSCGYEDHRCEVGLIVGTGTNACYMEELQNVAGVPGDSGHMCINMEWGAFGDDGTLDVPSTCFDASV
DRASINPGKQRFEKLISGMYLGEIVRHILLHLNSLGVLFRGQQTQCLQTRDIFKTKFLSEIESDSLALRQ
VRAILEDLGLPLTSDDALVVREVCQAVSRRAAQLCGAGVAAVVEKIRENRGLEELTVTVGVDGTLYKLHP
HFSSLVAATVRELAPRCAVTFLQSEDGSGKGAALVTAVACRLAQTTRV
>anc169|avgPP:0.994|lnPP:-8.519|num_ambig:4|num_ambig_gaps:0
LQQFMVTEAQLQQIQASLLGSMEQALMGQSSPAPAVRMLPTYVGSTPHGTEQGDFVVLELGATGASLRVL
WVTLTGTEGHRVEPRSQEFVIPPEVMLGPGQQLFDFAARCLAEFLEALPVGNQCLPLGFSFSFPCHQTGL
DRSTLISWTKGFRCSGVEGQDVVQLLRDAIERQGASSLDVVAVVNDTVGTMMGCDPGLGPCEVGLVVDTG
TNACYMEEARHVAVLDEDRGRVCVSVEWGSFGDDEALGPVLTPFDHALDQESLNPGAQRFEKMIGGLYLG
ELVRLVLAHLAQRGLLFGGCTSPALLSQGSVTLEHVAEMEDPSAGAARVLAVLQNLGLSPGASDVELVQH
VCAAVSTRAARLCAAALAAVLGRLQHSREQHTLQIAVATGGRVFERHPRFHSILQETVMLLAPECDVSFI
PSVDGGGRGVAMVTAVAARLAAHRRLLEETLAPFRLDREQLAAVQAQMREAMAKGLRGEASSLRMLPTYV
RATPDGSERGDFLALDLGGTNFRVLLVRVAAGGVQITSQVYAIPECVAQGSGQQLFDHIVDCIVDFQQKQ
GLSGQSLPLGFTFSFPCRQLGLDQGILLNWTKGFNASDCEGQDVVALLREAIRRRQAVELNVVAIVNDTV
GTMMSCGYEDPRCEVGLIVGTGTNACYMEELQNVAGVPGDSGHMCINMEWGAFGDDGTLDVLSTCFDASV
DRASINPGKQRFEKLISGMYLGEIVRHILLHLTSLGVLFRGQQTQRLQTRDIFKTKFLSEIESDSLALRQ
VRAILEDLGLPLTSDDALVVREVCQAVSRRAAQLCGAGVAAVVEKIRENRGLEELTVTVGVDGTLYKLHP
HFSSLVAATVRELAPRCAVTFLQSEDGSGKGAALVTAVACRLAQTTRV
>anc169_altAll|avgPP:0.993|lnPP:-11.075|num_ambig:4|num_ambig_gaps:0
LQQFMVTEAQLQQIQASLLGSMEQALMGQSSPAPAVRMLPTYVGSTPHGTEQGDFVVLELGATGASLRVL
WVTLTGTEGHRVEPRSQEFVIPPEVMLGPGQQLFDFAARCLAEFLEALPVGNQCLPLGFSFSFPCHQTGL
DRSTLISWTKGFRCSGVEGQDVVQLLREAIERQGASNLDVVAVVNDTVGTMMGCDPGLGPCEVGLVVDTG
TNACYMEEARHVAVLDEDRGRVCVSVEWGSFGDDEALGPVLTPFDHALDQESLNPGAQRFEKMIGGLYLG
ELVRLVLAHLAQRGLLFGGCTSPALLSQGSVTLEHVAEMEDPSAGAARVLAVLQNLGLSPGASDVELVQH
VCAAVSTRAARLCAAALAAVLGRLQHSREQHTLQIAVATGGRVFERHPRFLSILQETVMLLAPECDVSFI
PSVDGGGRGVAMVTAVAARLAAHRRLLEETLAPFRLDREQLAAVQVQMREAMAKGLRGEASSLRMLPTYV
RATPDGSERGDFLALDLGGTNFRVLLVRVAAGGVQITSQVYAIPECVAQGSGQQLFDHIVDCIVDFQQKQ
GLSGQSLPLGFTFSFPCRQLGLDQGILLNWTKGFNASDCEGQDVVALLREAIRRRQAVELNVVAIVNDTV
GTMMSCGYEDPRCEVGLIVGTGTNACYMEELQNVAGVPGDSGHMCINMEWGAFGDDGTLDVLSTCFDASV
DRASINPGKQRFEKLISGMYLGEIVRHILLHLTSLGVLFRGQQTQRLQTRDIFKTKFLSEIESDSLALRQ
VRAILEDLGLPLTSDDALVVREVCQAVSRRAAQLCGAGVAAVVEKIRENRGLEELTVTVGVDGTLYKLHP
HFSSLVAATVRELAPRCAVTFLQSEDGSGKGAALVTAVACRLAQTTRV
>anc170|avgPP:0.989|lnPP:-16.428|num_ambig:17|num_ambig_gaps:0
LQQFKVTEAQLRQIQASLLASMEQALKGQASPAPAVRMLPTYVGSTPHGTEQGDFVVLELGATGASLRVL
WVTLMGTEGPRMEPRSQEFVIPQEVMLGPGQQLFDFAARCLSEFLDALPLGKQSLQLGFSFSFPCHQTGL
DKSTLISWTKGFRCSGVEGQDVVQLLRDAIQRQGAYSIDVVAVVNDTVGTMMGCEPGVGPCEIGLVVDTG
TNACYMEEARHVAVLDEDRGRVCVSVEWGSFGDDGALGPVLTTFDHALDHESLNPGAQRFEKMIGGLYLG
ELVRLVLVHLAQRGVLFGGCTSSALLSQGSILLEHVAEMEDPSAGAARVHAVLQDLGLSVGASDAKLVQH
VCVAVCTRAARLCAAALAAVLSRLQHSREQQTLQIAVATGGRVFERHPRFLSILQETVMLLAPECDVSFI
PSVDGGGRGVAMVTAVAARLAAHRRLLEETLAPFRLSREQLAAVQAQMREAMAKGLRGEASSLRMLPTYV
RATPDGSERGDFLALDLGGTNFRVLLVRVAAGGVQITNQVYSIPESVAQGSGQQLFDHIVDCIVDFQQKQ
GLSGQSLPLGFTFSFPCRQLGLDQGILLNWTKGFKASDCEGQDVVCLLREAIRRKQGVELNVVAIVNDTV
GTMMSCGYEDPRCEVGLIVGTGTNACYMEELQNVAGVAGDSGHMCINMEWGAFGDDGSLDMLSTCFDASV
DQASINPGKQRFEKMISGMYLGEIVRHVLLHLTSLGVLFRGQQTQRLQTRDIFKTKFLSEIESDSLALRQ
VRAILEDLGLPLTSDDALMVLEVCQAVSQRAAQLCGAGVAAVVEKIRENRGLEELTVSVGVDGTLYKLHP

HFSSLVAATVRELAPRCAVTFLQSEDGSGKGAALVTAVACRLAQTTRV
>anc170_altAll|avgPP:0.986|lnPP:-25.562|num_ambig:17|num_ambig_gaps:0
LQQFKVTEAQLQQIQASLLASMEQALKGQASPAPAVRMLPTYVGSTPHGTEQGDFVVLELGATGASLRVL
WVTLMGTEGPRMEPRSQEFVIPQEVMLGPGQQLFDFAARCLSEFLDALPVGNQSLQLGFSFSFPCHQTGL
DKSTLISWTKGFRCSGVEGQDVVQMLREAIQRQGAYNIDVVAVVNDTVGTMMGCEPGVGPCEVGLVVDTG
TNACYMEEARHVAVLDEDRGRVCVSVEWGSFGDDGALGPVLTTFDHALDHESLNPGAQRFEKMIGGLYLG
ELVRLVLVHLAERGVLFGGCTSPALLSQGSILLEHVAEMESPSAGAARVHAILQDLGLSVGASDAKLVQH
VCVAVCTRAARLCAAALAAVLSRLQHSREQQTLQIAVATGGRVFERHPRFLSILQETVMLLAPECDVSFI
PSVDGGGRGVAMVTAVAARLAAHRRLLEETLAPFRLSREQLAAVQAQMREAMAKGLRGEASSLRMLPTYV
RATPDGSERGDFLALDLGGTNFRVLLVRVAAGGVQITNQVYSIPECVAQGSGQQLFDHIVDCIVDFQQKQ
GLSGQSLPLGFTFSFPCRQLGLDQGILLNWTKGFKASDCEGQDVVCLLREAIGRKQGVELNVVAIVNDTV
GTMMSCGYEDPRCEVGLIVGTGTNACYMEELRNVAGVPGDSGHMCINMEWGAFGDDGSLDMLSTCFDASV
DQASINPGKQRFEKMISGMYLGEIVRHVLLHLTSLGVLFRGQQTQRLQTRDIFKTKFLSEIESDSLALRQ
VRAILEDLGLPLTSDDALMVLEVCQAVSQRAAQLCGAGVAAVVEKIRENRGLEELTVSVGVDGTLYKLHP
HFSSLVAATVQELAPRCAVTFLQSEDGSGKGAALVTAVACRLAQTTHV
>anc171|avgPP:0.993|lnPP:-10.854|num_ambig:11|num_ambig_gaps:0
LQQFKVTEAQLQQIQASLLASMEQALRGQASPAPAVRMLPTYVGSTPHGTEQGDFVVLELGATGASLRVL
WVTLTGTEGPRMEPRSQEFVIPQEVMLGPGQQLFDFAARCLSEFLDALPLGNQSLQLGFSFSFPCHQTGL
DKSTLISWTKGFRCSGVEGQDVVQLLRDAIQRQGAYSIDVVAVVNDTVGTMMGCEPGVGPCEVGLVVDTG
TNACYMEEARHVAVLDEDRGRVCVSVEWGSFGDDGALGPVLTTFDHALDHESLNPGAQRFEKMIGGLYLG
ELVRLVLAHLAQRGVLFGGCTSPALLSQGSILLEHVAEMEDPSAGAARVHAILQDLGLSPGASDAKLVQH
VCVAVCTRAARLCAAALAAVLSRLQHSREQQTLQIAVATGGRVFERHPRFLSILQETVMLLAPECDVSFI
PSVDGGGRGVAMVTAVAARLAAHRRLLEETLAPFRLSREQLAAVQAQMREAMAKGLRGEASSLRMLPTYV
RATPDGSERGDFLALDLGGTNFRVLLVRVAAGGVQITNQVYSIPECVAQGSGQQLFDHIVDCIVDFQQKQ
GLSGQSLPLGFTFSFPCRQLGLDQGILLNWTKGFKASDCEGQDVVCLLREAIRRRQAVELNVVAIVNDTV
GTMMSCGYEDPRCEVGLIVGTGTNACYMEELRNVAGVAGDSGHMCINMEWGAFGDDGSLDMLSTCFDASV
DQASINPGKQRFEKMISGMYLGEIVRHILLHLTSLGVLFRGQQTQRLQTRDIFKTKFLSEIESDSLALRQ
VRAILEDLGLPLTSDDALMVLEVCQAVSQRAAQLCGAGVAAVVEKIRENRGLEELTVSVGVDGTLYKLHP
HFSSLVAATVRELAPRCAVTFLQSEDGSGKGAALVTAVACRLAQTTRV
>anc171_altAll|avgPP:0.991|lnPP:-15.584|num_ambig:11|num_ambig_gaps:0
LQQFKVTEAQLQQIQASLLASMEQALKGQASPAPAVRMLPTYVGSTPHGTEQGDFVVLELGATGASLRVL
WVTLTGTEGPRMEPRSQEFVIPQEVMLGPGQQLFDFAARCLSEFLDALPVGKQSLQLGFSFSFPCHQTGL
DKSTLISWTKGFRCSGVEGQDVVQMLRDAIQRQGAYNIDVVAVVNDTVGTMMGCEPGVGPCEVGLVVDTG
TNACYMEEARHVAVLDEDRGRVCVSVEWGSFGDDGALGPVLTTFDHALDHESLNPGAQRFEKMIGGLYLG
ELVRLVLAHLAERGVLFGGCTSPALLSQGSILLEHVAEMEDPSAGAARVHAVLQDLGLSVGASDAKLVQH
VCAAVCTRAARLCAAALAAVLSRLQHSREQQTLQIAVATGGRVFERHPRFLSILQETVMLLAPECDVSFI
PSVDGGGRGVAMVTAVAARLAAHRRLLEETLAPFRLNREQLAAVQAQMREAMAKGLRGEASSLRMLPTYV
RATPDGSERGDFLALDLGGTNFRVLLVRVAAGGVQITNQVYSIPESVAQGSGQQLFDHIVDCIVDFQQKQ
GLSGQSLPLGFTFSFPCRQLGLDQGILLNWTKGFKASDCEGQDVVCLLREAIRRRQAVELNVVAIVNDTV
GTMMSCGYEDPRCEVGLIVGTGTNACYMEELRNVAGVAGDSGHMCINMEWGAFGDDGSLDMLSTCFDASV
DQASINPGKQRFEKMISGMYLGEIVRHILLHLTSLGVLFRGQQTQRLQTRDIFKTKFLSEIESDSLALRQ
VRAILEDLGLPLTSDDALMVLEVCQAVSQRAAQLCGAGVAAVVEKIRENRGLEELTVSVGVDGTLYKLHP
HFSSLVAATVRELAPRCAVTFLQSEDGSGKGAALVTAVACRLAQTTRV
>anc172|avgPP:0.993|lnPP:-10.887|num_ambig:7|num_ambig_gaps:0
LNQFKVTEAQLRQIQASLLGSMEQALRGQASPAPAVRMLPTYVGSIPHGTEQGDFLVLELGATGASLRVL
WVTLTGTEGHRIEPRSQEFLIPQEVMLGPGQQLFDFAARCLSEFLDALPVGNQGLQLGFNFSFPCHQTGL
DRSTLISWTKGFRCSGVEGQDVVQLLRDAIKRQGAYSINVVAVVNDTVGTMMGCEPGTGPCEVGLVVDTG
TNACYMEEARHVAVLDEDRGRVCVSIEWGSFGDDGALGPVLTTFDHILDQESLNPGAQRFEKMIGGLYLG
ELVRLVLAHLAQRGVLFGGCTSPALLSQGSILLEHVAEMEDPSAGTARVHAILQDLGLSPGASDAELVQH
VCAAVCTRAARLCAAALAAVLTRLQHSREQQTLQIAVATGGRVFERHPRFLSILQETVKLLAPECDVSFI
PSVDGGGRGVAMVTAVAARLAAHRRLLEETLAPFRLNREQLAAVQAQMREAMAKGLRGEASSLRMLPTYV
RATPDGSERGDFLALDLGGTNFRVLLVRVAAGGVQITSQIYSIPECVAQGSGQQLFDHIVDCILDFQQKQ

GLSGQSLPLGFTFSFPCRQLGLDQGILLNWTKGFNASDCEGQDVVCLLREAIGRRQAVELNVVAIVNDTV
GTMMSCGYEDPRCEVGLIVGTGTNACYMEELRNVAGVAGDTGHMCINMEWGAFGDDGSLDMLSTCFDASV
DQASINPGKQRFEKMISGMYLGEIVRHILLHLTSLGVLFRGQQTQRLQTRDIFKTKFLSEIESDSLALRQ
VRAILEDLGLPLTSDDALMVLEVCQAVSQRAAQLCGAGVAAVVEKIRENRGLEELTVSVGVDGTLYKLHP
HFSGLVAATVRELAPRCAVTFLQSEDGSGKGAALVTAVACRLAQTAHV
>anc172_altAll|avgPP:0.992|lnPP:-13.399|num_ambig:7|num_ambig_gaps:0
LDQFKVTEAQLRQIQASLLGSMEQALRGQASPAPAVRMLPTYVGSIPHGTEQGDFLVLELGATGASLRVL
WVTLTGTEGHRIEPRSQEFLIPQEVMLGPGQQLFDFAARCLSEFLDALPVGNKSLQLGFNFSFPCHQTGL
DRSTLISWTKGFRCSGVEGQDVVQLLRDAIQRQGAYSINVVAVVNDTVGTMMGCEPGTGPCEVGLVVDTG
TNACYMEEARHVAVLDEDRGRVCVSIEWGSFGDDGALGPVLTTFDHILDQESLNPGAQRFEKMIGGLYLG
ELVRLVLAHLAQQGVLFGGCTSPALLSQGSILLEHVAEMEDPSAGTARVHAILQDLGLSPGASDAELVQH
VCAAVCTRAARLCAAALAAVLTRLQHSREQQTLQIAVATGGRVFERHPRFLSILQETVKLLAPECDVSFI
PSVDGGGRGVAMVTAVAARLAAHRRLLEETLAPFRLNREQLAAVQAQMREAMAKGLRGEASSLRMLPTYV
RATPDGSERGDFLALDLGGTNFRVLLVRVAAGGVQITSQIYSIPECVAQGSGQQLFDHIVDCILDFQQKQ
GLSGQSLPLGFTFSFPCRQLGLDQGILLNWTKGFNASDCEGQDVVRLLREAIGRRQAVELNVVAIVNDTV
GTMMSCGYEDPRCEVGLIVGTGTNACYMEELRNVAGVAGDTGHMCINMEWGAFGDDGSLDMLSTCFDASV
DQASINPGKQRFEKMISGMYLGEIVRHILLHLTSLGVLFRGQQTQRLQTRDIFKTKFLSEIESDSLALRQ
VRAILEDLGLPLTSDDALMVLEVCQAVSQRAAQLCGAGVAAVVEKIRENRGLEELTVSVGVDGTLYKLHP
HFSGLVAATVRELAPRCAVTFLQSEDGSGKGAALVTAVACRLAQPAHV
>anc173|avgPP:0.996|lnPP:-5.464|num_ambig:6|num_ambig_gaps:0
LQQFKVTEAQLQQIQASLLGSMEQALRGQASPAPAVRMLPTYVGSTPHGTEQGDFLVLELGATGASLRVL
WVTLTGTEGHRMEPRSQEFVIPQEVMLGPGQQLFDFAARCLSEFLDALPVGNQGLQLGFSFSFPCHQTGL
DRSTLISWTKGFRCSGVEGQDVVQLLRDAIKRQGAYSIDVVAVVNDTVGTMMGCEPGVGPCEVGLVVDTG
TNACYMEEARHVAVLDEDRGRVCVSVEWGSFGDDGALGPVLTTFDHALDQESLNPGAQRFEKMIGGLYLG
ELVRLVLAHLAQRGVLFGGCTSPALLSQGSILLEHVAEMEDPSAGAARVHAILQDLGLSPGASDAELVQH
VCAAVCTRAARLCAAALAAVLSRLQHSREQQTLQIAVATGGRVFERHPRFLSILQETVKLLAPECDVSFI
PSVDGGGRGVAMVTAVAARLAAHRRLLEETLAPFRLNREQLAAVQAQMREAMAKGLRGEASSLRMLPTYV
RATPDGSERGDFLALDLGGTNFRVLLVRVAAGGVQITNQVYSIPECVAQGSGQQLFDHIVDCIVDFQQKQ
GLSGQSLPLGFTFSFPCRQLGLDQGILLNWTKGFNASDCEGQDVVCLLREAIRRRQAVELNVVAIVNDTV
GTMMSCGYEDPRCEVGLIVGTGTNACYMEELRNVAGVAGDSGHMCINMEWGAFGDDGSLDMLSTCFDASV
DQASINPGKQRFEKMISGMYLGEIVRHILLHLTSLGVLFRGQQTQRLQTRDIFKTKFLSEIESDSLALRQ
VRAILEDLGLPLTSDDALMVLEVCQAVSQRAAQLCGAGVAAVVEKIRENRGLEELTVSVGVDGTLYKLHP
HFSGLVAATVRELAPRCAVTFLQSEDGSGKGAALVTAVACRLAQTTHV
>anc173_altAll|avgPP:0.995|lnPP:-8.672|num_ambig:6|num_ambig_gaps:0
LQQFKVTEAQLQQIQASLLGSMEQALRGQASPAPAVRMLPTYVGSTPHGTEQGDFLVLELGATGASLRVL
WVTLTGTEGHRMEPRSQEFVIPQEVMLGPGQQLFDFAARCLSEFLDALPVGNQSLQLGFSFSFPCHQTGL
DRSTLISWTKGFRCSGVEGQDVVQLLRDAIQRQGAYNIDVVAVVNDTVGTMMGCEPGVGPCEVGLVVDTG
TNACYMEEARHVAVLDEDRGRVCVSVEWGSFGDDGALGPVLTTFDHALDQESLNPGAQRFEKMIGGLYLG
ELVRLVLAHLAQRGVLFGGCTSPALLSQGSILLEHVAEMEDPSAGAARVHAILQDLGLSPGASDAELVQH
VCAAVCTRAARLCAAALAAVLSRLQHSREQQTLQIAVATGGRVFERHPRFLSILQETVKLLAPECDVSFI
PSVDGGGRGVAMVTAVAARLAAHRRLLEETLAPFRLNREQLAAVQAQMREAMAKGLRGEASSLRMLPTYV
RATPDGSERGDFLALDLGGTNFRVLLVRVAAGGVQITNQVYSIPECVAQGSGQQLFDHIVDCIVDFQQKQ
GLSGQSLPLGFTFSFPCRQLGLDQGILLNWTKGFNASDCEGQDVVCLLREAIRRRQAVELNVVAIVNDTV
GTMMSCGYEDPRCEVGLIVGTGTNACYMEELRNVAGVAGDSGHMCINMEWGAFGDDGSLDMLSTCFDASV
DQASINPGKQRFEKMISGMYLGEIVRHILLHLTSLGVLFRGQQTQRLQTRDIFKTKFLSEIESDSLALRQ
VRAILEDLGLPLTSDDALMVLEVCQAVSQRAAQLCGAGVAAVVEKIRENRGLEELTVSVGVDGTLYKLHP
HFSSLVAATVRELAPRCAVTFLQSEDGSGKGAALVTAVACRLAQTARV
>anc174|avgPP:0.998|lnPP:-3.587|num_ambig:4|num_ambig_gaps:0
LQQFKVTEAQLQQIQASLLGSMEQALRGQASPAPAVRMLPTYVGSTPHGTEQGDFLVLELGATGASLRVL
WVTLTGTEGHRMEPRSQEFVIPQEVMLGPGQQLFDFAARCLSEFLDALPVGNQSLQLGFSFSFPCHQTGL
DRSTLISWTKGFRCSGVEGQDVVQLLRDAIQRQGAYSIDVVAVVNDTVGTMMGCEPGVGPCEVGLVVDTG
TNACYMEEARHVAVLDEDRGRVCVSVEWGSFGDDGALGPVLTTFDHALDQESLNPGAQRFEKMIGGLYLG

ELVRLVLAHLAQRGVLFGGCTSPALLSQGSILLEHVAEMEDPSAGAARVHAILQDLGLSPGASDAELVQH
VCAAVCTRAARLCAAALAAVLSRLQHSREQQTLQIAVATGGRVFERHPRFLSILQETVMLLAPECDVSFI
PSVDGGGRGVAMVTAVAARLAAHRRLLEETLAPFRLNREQLAAVQAQMREAMAKGLRGEASSLRMLPTYV
RATPDGSERGDFLALDLGGTNFRVLLVRVAAGGVQITNQVYSIPECVAQGSGQQLFDHIVDCIVDFQQKQ
GLSGQSLPLGFTFSFPCRQLGLDQGILLNWTKGFNASDCEGQDVVCLLREAIRRRQAVELNVVAIVNDTV
GTMMSCGYEDPRCEVGLIVGTGTNACYMEELRNVAGVAGDSGHMCINMEWGAFGDDGSLDMLSTCFDASV
DQASINPGKQRFEKMISGMYLGEIVRHILLHLTSLGVLFRGQQTQRLQTRDIFKTKFLSEIESDSLALRQ
VRAILEDLGLPLTSDDALMVLEVCQAVSQRAAQLCGAGVAAVVEKIRENRGLEELTVSVGVDGTLYKLHP
HFSSLVAATVRELAPRCAVTFLQSEDGSGKGAALVTAVACRLAQTTRV
>anc174_altAll|avgPP:0.997|lnPP:-5.399|num_ambig:4|num_ambig_gaps:0
LQQFKVTEAQLQQIQASLLGSMEQALRGQASPAPAVRMLPTYVGSTPHGTEQGDFVVLELGATGASLRVL
WVTLTGTEGHRMEPRSQEFVIPQEVMLGPGQQLFDFAARCLSEFLDALPVGNQGLQLGFSFSFPCHQTGL
DRSTLISWTKGFRCSGVEGQDVVQLLRDAIQRQGAYNIDVVAVVNDTVGTMMGCEPGVGPCEVGLVVDTG
TNACYMEEARHVAVLDEDRGRVCVSVEWGSFGDDGALGPVLTTFDHALDHESLNPGAQRFEKMIGGLYLG
ELVRLVLAHLAQRGVLFGGCTSPALLSQGSILLEHVAEMEDPSAGAARVHAILQDLGLSPGASDAELVQH
VCAAVCTRAARLCAAALAAVLSRLQHSREQQTLQIAVATGGRVFERHPRFLSILQETVMLLAPECDVSFI
PSVDGGGRGVAMVTAVAARLAAHRRLLEETLAPFRLNREQLAAVQAQMREAMAKGLRGEASSLRMLPTYV
RATPDGSERGDFLALDLGGTNFRVLLVRVAAGGVQITNQVYSIPECVAQGSGQQLFDHIVDCIVDFQQKQ
GLSGQSLPLGFTFSFPCRQLGLDQGILLNWTKGFNASDCEGQDVVCLLREAIRRRQAVELNVVAIVNDTV
GTMMSCGYEDPRCEVGLIVGTGTNACYMEELRNVAGVAGDSGHMCINMEWGAFGDDGSLDMLSTCFDASV
DQASINPGKQRFEKMISGMYLGEIVRHILLHLTSLGVLFRGQQTQRLQTRDIFKTKFLSEIESDSLALRQ
VRAILEDLGLPLTSDDALMVLEVCQAVSQRAAQLCGAGVAAVVEKIRENRGLEELTVSVGVDGTLYKLHP
HFSSLVAATVRELAPRCAVTFLQSEDGSGKGAALVTAVACRLAQTTRV
>anc175|avgPP:0.991|lnPP:-13.506|num_ambig:11|num_ambig_gaps:0
LQQFQVTRAQLQQIQASLLGSMEQALRGQASPAPAVRMLPTYVGSTPHGTEQGDFLVLELGATGASLRVL
WVTLTGIDGHRVEPRSQEFVIPQEVMLGTGQQLFDFAARCLSEFLDAHPVGNQGLQLGFSFSFPCHQTGL
DRSTLISWTKGFRCSGVEGQDVVQLLRDAIQRQGAYNIDVIAVVNDTVGTMMGCEPGVRPCEVGLVVDTG
TNACYMEEARHVAVLDEDRGRVCVSVEWGSFNDDGALGPVLTTFDHALDHESLNPGAQRFEKMIGGLYLG
ELVRLVLAHLAQRGVLFGGCTSPALQSQGSILPEHVAEMEDPSTGAARVLTILQDLGLSPEASDAELVQS
VCAAVSTRAARLCAAALAAVLSRLQRSREQQTLQVTVATGGRVFEQHPRFHSILQETVVLLVPECDVSFI
PSVDGGGRGVAVVTAVAARLAAHRRLLEETLMPFRLNREQLAAVQAQMREAMAKGLRGEASSLRMLPTYV
RATPDGSERGDFLALDLGGTNFRVLLVRVTEGGVQITNQVYSVPEHVAQGSGQQLFDHIVDCIVDFQRKQ
GLSGQSLPLGFTFSFPCKQLGLDQGILLNWTKGFNASDCEGRDVVCLLREAIRRRQAVELNVVAIVNDTV
GTMMSCGYEDPRCEMGLIVGTGTNACYMEELRNVAGVAGDSGHMCINMEWGAFGDDGSLGSLSTCFDASV
DQASINPGKQRFEKMISGMYLGEIVRHILLHLTSLGVLFRGQQTQRLQTRDIFKTKFLSDIESDSLALRQ
VRAILEDLGLPVTSDDALMVLEVCQAVSRRAAQLCGAGVAAVVEKIRENRGLEELTVSVGVDGTLYKLHP
HFSSLVAATVRQLAPRCTVTFLESEDGSGKGAALVTAVACRLAQTTRV
>anc175_altAll|avgPP:0.989|lnPP:-18.655|num_ambig:11|num_ambig_gaps:0
LQQFQVTRAQLQQIQASLLGSMEQALRGQASPAPAVRMLPTYVGSTPHGTEQGDFLVLELGATGASLRVL
WVTLTGVDGHRVEPRSQEFVIPQEVMLGTGQQLFDFAARCLSEFLDAHPVGNQGLQLGFSFSFPCHQTGL
DRSTLISWTKGFRCSGVEGQDVVQLLRDAIQRQGAYNIDVIAVVNDTVGTMMGCEPGVRPCEVGLVVDTG
TNACYMEEARHVAVLDEDRGRVCVSVEWGSFNDDGALGPVLTTFDHALDQESLNPGAQRFEKMVGGLYLG
ELVRLVLAHLAQRGVLFGGCTSPALQSQGSILPEHVAEMEDPSAGAARVLAILQDLGLSPEASDAELVQS
VCAAVSTRAARLCAAALAAVLSRLQRSREQQTLQVTVATGGRVFERHPRFHSILQETVMLLVPECDVSFI
PSVDGGGRGVAVVTAVAARLAAHRRLLEETLMPFRLNREQLAGVQAQMREAMAKGLRGEASSLRMLPTYV
RATPDGSERGDFLALDLGGTNFRVLLVRVTEGGVQITNQVYSVPEHVAQGSGQQLFDHIVDCIVDFQRKQ
GLSGQSLPLGFTFSFPCKQLGLDQGILLNWTKGFNASDCEGQDVVCLLREAIRRRQAVELNVVAIVNDTV
GTMMSCGYEDPRCEMGLIVGTGTNACYMEELRNVAGVAGDSGQMCINMEWGAFGDDGSLGSLSTCFDDSV
DQASINPGKQRFEKMISGMYLGEIVRHILLHLTSLGVLFRGQQTQRLQTRDIFKTKFLSDIESDSLALRQ
VRAILEDLGLPVTSDDALMVLEVCQAVSRRAAQLCGAGVAAVVEKIRENRGLEELTVSVGVDGTLYKLHP
HFSSLVAATVRQLAPRCTVTFLESEDGSGKGAALVTAVACRLAQTTRV
>anc176|avgPP:0.994|lnPP:-10.161|num_ambig:9|num_ambig_gaps:0

```
LQQFKVTRTQLQQIQASLLGSMEQALTGKASPAPGVRMLPTYVGSTPHGNEQGNFVVLELGASGASLRVL
WVTLTGIKGQRVEPKSQEFVIPQEVVLGTGQQLFDFAARCLSEFLDAHSVGNQGLQLGFSFSFPCHQTGL
DRSTLISWTKGFRCSGVEGQDVVQLLRDAIQRQGAYNIDVVAVVNDTVGTMMGCEPGVRPCEVGLIVDTG
TNACYMEEARHVAVLDEDRGRTCVSVEWGSFNDDGALGPVLTTFDHALDHASLNPGAQRFEKMIGGLYLG
ELVRLVLAHLAQHRVLFGGYTSPALLSQGSILLEHVAEMEDPSTGAARVHAILQDLGLSPEASDAELVQR
VCAAVCSRAAQLCAAALAAILSRLQHSREQQTLQVAVATGGRVLEQHPRFHSILQETVMLLASECDVSFI
PSVDGGGRGVAMVTAVAARLAAHRRLLEETLAPFRLSREQLAAVQTQMREAMAKGLRGEASSLRMLPTYV
RATPDGSERGDFLALDLGGTNFRVLLVRVTEGGVQIVNQVYSIPECVAQGSGQQLFDHIVDCIVDFQEKQ
GLRGQSLPLGFTFSFPCKQLGLDQGILLNWTKGFNASGCEGQDVVYLLREAIRRKQAVELNVVAIVNDTV
GTMMSCGYEDPRCEMGLIVGTGTNACYMEELRNVAGVAGDSGHMCINMEWGAFGDDGSLGMLSTCFDARV
DQASINPGKQRFEKMISGMYLGEIVRHILLHLTSLGVLFRGQQTQRLQTRDIFKTKFLSEIESDSLALRQ
VRAILEDLGLPLTSDDALIVLEVCQAVSRRAAQLCGAGVAAVVEKIRENRGLEELTVSVGVDGTLYKLHP
HFSSLVAATVRELAPRCTVTFLQSEDGSGKGAALVTAVACRLAQTTRV
>anc176_altAll|avgPP:0.993|lnPP:-12.748|num_ambig:9|num_ambig_gaps:0
LQQFKVTRTQLQQIQASLLGSMEQALTGQASPAPGVRMLPTYVGSTPHGNEQGNFVVLELGASGASLRVL
WVTLTGIKGQRVEPKSQEFVIPQEVVLGTGQQLFDFAARCLSEFLDAHSVGNQGLQLGFSFSFPCHQTGL
DRSTLISWTKGFRCSGVEGQDVVQLLRDAIQRQGAYNIDVVAVVNDTVGTMMGCEPGVRPCEVGLIVDTG
TNACYMEEARHVAVLDEDRGRTCVSVEWGSFNDDGALGPVLTTFDHALDHASLNPGAQRFEKMIGGLYLG
ELVRLVLAHLAQHRVLFGGYTSPALLSQGSILLEHVAEMEDPSTGAARVHAILQDLGLSPEASDAELVQR
VCAAVCSRAAQLCAAALAAILSRLQHSREQQTLQVAVATGGRVLEQHPRFHSILQETVTLLASECDVSFI
PSVDGGGRGVAMVTAVAARLAAHRRLLEETLAPFRLNREQLAAVQTQMREAMAKGLRGEASSLRMLPTYV
RATPDGSERGDFLALDLGGTNFRVLLVRVTEGGVQIINQVYSIPECVAQGSGQQLFDHIVDCIVDFQEKQ
GLRGQSLPLGFTFSFPCKQLGLDQGILLNWTKGFNASGCEGQDVVYLLREAIRRKQAVELNVVAIVNDTV
GTMMSCGYEDPRCEMGLIVGTGTNACYMEELRNVAGVAGDSGHMCINMEWGAFGDDGSLGMLSTCFDESV
DQASINPGKQRFEKMISGMYLGEIVRHILLHLTSLGVLFRGQQTQRLQTRDIFKTKFLSEIESDSLALRQ
VRAILEDLGLPLTSDDALIVLEVCQAVSRRAAQLCGAGVAAVVEKIRENRGLEELTVSVGVDGTLYKLHP
HFSSLVAATVRELAPRCAVTFLQSEDGSGKGAALVTAVACRLAQTAHV
>anc177|avgPP:0.994|lnPP:-9.958|num_ambig:11|num_ambig_gaps:0
LQQFKVTRTQLQQIQASLLCSMEQALKGQDSPAPSVRMLPTYVGSTPHGTEQGDFLVLELGATGASLRVL
WVTLTGTKEHRVEPRSQEFVIPQEVMLGAGQQLFDFAARCLSEFLDAHPVKNQDLKLGFNFSFPCHQTGL
DRSTLISWTKGFRCSGVEGQDVVQLLRDAIQRQGTYSIDVVAMVNDTVGTMMGCELGARPCEVGLIVDTG
TNACYMEEARHVAALDEDRGRTCVSIEWGSFCDEEALGPVLTTFDHALDHESLTPGAQRFEKMIGGLYLG
ELVRLVLVHLAQHGVLFGGCTSPALLSQGSILLEHVAEMEDPATGTAHVHTILQDLGLSPQASDAELVQR
VCAAVCTRAAQLCAAALAAVLSRLQHSREQQTLRVAVATGGRVFEQHPRFHSILKETVMLLAPECDVSFI
PSVDGGGRGVAMVTAVAARLAAHRHILEETLAPFQLTLEQLAVVQAQMREAMIRGLQGEASSLRMLPTYV
RATPDGSERGDFLALDLGGTNFRVLLVRVAEGSVQITNQVYSIPECVAQGSGQQLFDHIVDCIVDFQQRQ
GLSGQSLPLGFTFSFPCKQLGLDQGILLNWTKGFNASGCEGQDVVYLLREAIRRRQAVELNVVAIVNDTV
GTMMSCGYDDPRCEMGLIVGTGTNACYMEELRNVASVAGDSGRMCINMEWGAFGDDGSLGMLSTCFDASV
DQASINPGKQRFEKMISGMYLGEIVRHILLHLTSLGVLFRGQKTQCLQTRDIFKTKFLSEIESDSLALRQ
VRAILEDLGLTLTSDDALMVLEVCQAVSRRAAQLCGAGVAAVVEKIRENRGLQELTVSVGVDGTLYKLHP
HFSRLVSATVRKLAPHCTVTFLQSEDGSGKGAALVTAVACRLTQMARV
>anc177_altAll|avgPP:0.991|lnPP:-15.517|num_ambig:11|num_ambig_gaps:0
LQQFKVTRTQLQQIQASLLCSMEQALKGQDSPAPSVRMLPTYVGSTPHGTEQGDFLVLELGATGASLRVL
WVTLTGTKEHRVEPRSQEFVIPQEVMLGTGQQLFDFAACCLSEFLDAHPVKNQDLKLGFNFSFPCHQTGL
DRSTLISWTKGFRCSGVEGQDVVQLLRDAIQRQGTYSIDVVAMVNDTVGTMMGCELGARPCEVGLIVDTG
TNACYMEEARHVAALDEDRGRTCVSIEWGSFCDEEALGPVLTTFDHALDHESLTPGAQRFEKMIGGLYLG
ELVRLVLVHLAQHGVLFGGCTSPALLSQGSILLEHVAEMEDPATGTAHVHAILQDLGLSPQASDAELVQH
VCAAVCTRAAQLCAAALAAVLSRLQHSREQQTLRVAVATGGRVFERHPRFHRILKETVMLLAPECDVSFI
PSVDGGGRGVAMVTAVAARLAAHRRILEETLAPFQLTLEQLTVVQAQMREAMIRGLQGEASSLRMLPTYV
RATPDGSERGDFLALDLGGTNFRVLLVRVAEGSVQITNQVYSIPECVAQGSGQQLFDHIVDCIVDFQQRQ
GLSGQSLPLGFTFSFPCKQLGLDQGILLNWTKGFNASGCEGQDVVHLLREAIRRRQAVELNVVAIVNDTV
GTMMSCGYDDPRCEMGLIVGTGTNACYMEELRNVASVAGDSGRMCINMEWGAFGDDGSLGMLSTCFDASV
```

DQASINPGKQRFEKMISGMYLGEIVRHILLHLTSLGVLFRGQKTQCLQTRDIFKTKFLSEIESDSLALRQ
VRAILEDLGLTLTSDDALMVLEVCQAVSRRAAQLCGAGVAAVVEKIRENRGLQELTVSVGVDGTLYKLHP
HFSRLVSATVRKLAPHCTVTFLQSEDGSGKGAALVTAVACRLAQMAHV
>anc178|avgPP:0.994|lnPP:-9.958|num_ambig:11|num_ambig_gaps:0
LQQFKVTRTQLQQIQASLLCSMEQALKGQDSPAPSVRMLPTYVGSTPHGTEQGDFLVLELGATGASLRVL
WVTLTGTKEHRVEPRSQEFVIPQEVMLGAGQQLFDFAARCLSEFLDAHPVKNQDLKLGFNFSFPCHQTGL
DRSTLISWTKGFRCSGVEGQDVVQLLRDAIQRQGTYSIDVVAMVNDTVGTMMGCELGARPCEVGLIVDTG
TNACYMEEARHVAALDEDRGRTCVSIEWGSFCDEEALGPVLTTFDHALDHESLTPGAQRFEKMIGGLYLG
ELVRLVLVHLAQHGVLFGGCTSPALLSQGSILLEHVAEMEDPATGTAHVHTILQDLGLSPQASDAELVQR
VCAAVCTRAAQLCAAALAAVLSRLQHSREQQTLRVAVATGGRVFEQHPRFHSILKETVMLLAPECDVSFI
PSVDGGGRGVAMVTAVAARLAAHRHILEETLAPFQLTLEQLAVVQAQMREAMIRGLQGEASSLRMLPTYV
RATPDGSERGDFLALDLGGTNFRVLLVRVAEGSVQITNQVYSIPECVAQGSGQQLFDHIVDCIVDFQQRQ
GLSGQSLPLGFTFSFPCKQLGLDQGILLNWTKGFNASGCEGQDVVYLLREAIRRRQAVELNVVAIVNDTV
GTMMSCGYDDPRCEMGLIVGTGTNACYMEELRNVASVAGDSGRMCINMEWGAFGDDGSLGMLSTCFDASV
DQASINPGKQRFEKMISGMYLGEIVRHILLHLTSLGVLFRGQKTQCLQTRDIFKTKFLSEIESDSLALRQ
VRAILEDLGLTLTSDDALMVLEVCQAVSRRAAQLCGAGVAAVVEKIRENRGLQELTVSVGVDGTLYKLHP
HFSRLVSATVRKLAPHCTVTFLQSEDGSGKGAALVTAVACRLTQMARV
>anc178_altAll|avgPP:0.991|lnPP:-15.516|num_ambig:11|num_ambig_gaps:0
LQQFKVTRTQLQQIQASLLCSMEQALKGQDSPAPSVRMLPTYVGSTPHGTEQGDFLVLELGATGASLRVL
WVTLTGTKEHRVEPRSQEFVIPQEVMLGTGQQLFDFAACCLSEFLDAHPVKNQDLKLGFNFSFPCHQTGL
DRSTLISWTKGFRCSGVEGQDVVQLLRDAIQRQGTYSIDVVAMVNDTVGTMMGCELGARPCEVGLIVDTG
TNACYMEEARHVAALDEDRGRTCVSIEWGSFCDEEALGPVLTTFDHALDHESLTPGAQRFEKMIGGLYLG
ELVRLVLVHLAQHGVLFGGCTSPALLSQGSILLEHVAEMEDPATGTAHVHAILQDLGLSPQASDAELVQH
VCAAVCTRAAQLCAAALAAVLSRLQHSREQQTLRVAVATGGRVFERHPRFHRILKETVMLLAPECDVSFI
PSVDGGGRGVAMVTAVAARLAAHRRILEETLAPFQLTLEQLTVVQAQMREAMIRGLQGEASSLRMLPTYV
RATPDGSERGDFLALDLGGTNFRVLLVRVAEGSVQITNQVYSIPECVAQGSGQQLFDHIVDCIVDFQQRQ
GLSGQSLPLGFTFSFPCKQLGLDQGILLNWTKGFNASGCEGQDVVHLLREAIRRRQAVELNVVAIVNDTV
GTMMSCGYDDPRCEMGLIVGTGTNACYMEELRNVASVAGDSGRMCINMEWGAFGDDGSLGMLSTCFDASV
DQASINPGKQRFEKMISGMYLGEIVRHILLHLTSLGVLFRGQKTQCLQTRDIFKTKFLSEIESDSLALRQ
VRAILEDLGLTLTSDDALMVLEVCQAVSRRAAQLCGAGVAAVVEKIRENRGLQELTVSVGVDGTLYKLHP
HFSRLVSATVRKLAPHCTVTFLQSEDGSGKGAALVTAVACRLAQMAHV
>anc179|avgPP:0.994|lnPP:-8.590|num_ambig:11|num_ambig_gaps:0
LQQFKVTRTQLQQIQASLLGSMEQALKGQDSPAPAVRMLPTYVGSTPHGTEQGDFLVLELGATGASLRVL
WVTLTGTEEHRVEPRSQEFVIPQEVVLGTGQQLFDFAARCLSEFLDAHPVGNQDLKLGFSFSFPCHQTGL
DRSTLISWTKGFRCSGVEGQDVVQLLRDAIQRQGTYNIDVVAVVNDTVGTMMGCELGARPCEVGLIVDTG
TNACYMEEARHVAALDEDRGRTCVSIEWGSFNDDGALGPVLTTFDHALDHESLNPGAQRFEKMIGGLYLG
ELVRLVLAHLAQHGVLFGGCTSPALLSQGSILLEHVAEMEDPSTGAAHVHAILQDLGLSPQASDAELVQR
VCAAVCTRAAQLCAAALAAVLSRLQHSREQQTLQVAVATGGRVFEQHPRFHSILKETVMLLAPECDVSFI
PSVDGGGRGVAMVTAVAARLAAHRHLLEETLAPFQLNREQLAVVQAQMREAMIRGLQGEASSLRMLPTYV
RATPDGSERGDFLALDLGGTNFRVLLVRVAEGSVQITNQVYSIPECVAQGSGQQLFDHIVDCIVDFQQRQ
GLSGQSLPLGFTFSFPCKQLGLDQGILLNWTKGFNASGCEGQDVVCLLREAIRRRQAVELNVVAIVNDTV
GTMMSCGYDDPRCEMGLIVGTGTNACYMEELRNVASVAGDSGHMCINMEWGAFGDDGSLGMLSTCFDASV
DQASINPGKQRFEKMISGMYLGEIVRHILLHLTSLGVLFRGQQTQCLQTRDIFKTKFLSEIESDSLALRQ
VRAILEDLGLTLTSDDALMVLEVCQAVSRRAAQLCGAGVAAVVEKIRENRGLEELTVSVGVDGTLYKLHP
HFSSLVSATVRKLAPHCTVTFLQSEDGSGKGAALVTAVACRLAQTARV
>anc179_altAll|avgPP:0.992|lnPP:-15.186|num_ambig:11|num_ambig_gaps:0
LQQFKVTRTQLQQIQASLLRSMEQALKGQNSPAPAVRMLPTYVGSTPHGTEQGDFLVLELGATGASLRVL
WVTLTGTEEHRVEPRSQEFVIPQEVVLGAGQQLFDFAACCLSEFLDAHPVGNQDLKLGFSFSFPCHQTGL
DRSTLISWTKGFRCSGVEGQDVVQLLRDAIQRQGTYNIDVVAVVNDTVGTMMGCELGARPCEVGLIVDTG
TNACYMEEARHVAALDEDRGRTCVSIEWGSFNDDGALGPVLTTFDHALDHESLNPGAQRFEKMIGGLYLG
ELVRLVLAHLAQHGVLFGGCTSPALLSQGSILLEHVAEMEDPSTGAAHVHTILQDLGLSPQASDAELVQH
VCAAVCTRAAQLCAAALAAVLSRLQHSREQQTLQVAVATGGRVFERHPRFHSILKETVMLLAPECDVSFI

PSVDGGGRGVAMVTAVAARLAAHRRLLEETLAPFQLNREQLTVVQAQMREAMIRGLQGEASSLRMLPTYV
RATPDGSERGDFLALDLGGTNFRVLLVRVAEGSVQITNQVYSIPECVAQGSGQQLFDHIVDCIVDFQQRQ
GLSGQSLPLGFTFSFPCKQLGLDQGILLNWTKGFNASGCEGQDVVCLLREAIRRRQAVELNVVAIVNDTV
GTMMSCGYDDPRCEMGLIVGTGTNACYMEELRNVASVAGDSGHMCINMEWGAFGDDGSLGMLSTCFDASV
DQASINPGKQRFEKMISGMYLGEIVRHILLHLTSLGVLFRGQQTQCLQTRDIFKTKFLSEIESDSLALRQ
VRAILEDLGLTLTSDDALMVLEVCQAVSRRAAQLCGAGVAAVVEKIRENRGLEELTVSVGVDGTLYKLHP
HFSSLVSATVRKLAPRCTVTFLQSEDGSGKGAALVTAVACRLAQTTRV
>anc180|avgPP:0.995|lnPP:-7.237|num_ambig:7|num_ambig_gaps:0
LQQFKVTRTQLQQIQASLLGSMEQALKGQASPAPAVRMLPTYVGSTPHGTEQGDFLVLELGATGASLRVL
WVTLTGTEGHRVEPRSQEFVIPQEVVLGTGQQLFDFAARCLSEFLDAHPVGNQGLQLGFSFSFPCHQTGL
DRSTLISWTKGFRCSGVEGQDVVQLLRDAIQRQGTYNIDVVAVVNDTVGTMMGCELGARPCEVGLIVDTG
TNACYMEEARHVAALDEDRGRTCVSVEWGSFNDDGALGPVLTTFDHALDHESLNPGAQRFEKMIGGLYLG
ELVRLVLAHLAQHGVLFGGCTSPALLSQGSILLEHVAEMEDPSTGAAHVHAILQDLGLSPEASDAELVQR
VCAAVCTRAAQLCAAALAAVLSRLQHSREQQTLQVAVATGGRVFEQHPRFHSILKETVMLLAPECDVSFI
PSVDGGGRGVAMVTAVAARLAAHRHLLEETLAPFQLNREQLAAVQAQMREAMVRGLRGEASSLRMLPTYV
RATPDGSERGDFLALDLGGTNFRVLLVRVTEGGVQITNQVYSIPECVAQGSGQQLFDHIVDCIVDFQQRQ
GLSGQSLPLGFTFSFPCKQLGLDQGILLNWTKGFNASDCEGQDVVCLLREAIRRRQAVELNVVAIVNDTV
GTMMSCGYEDPRCEMGLIVGTGTNACYMEELRNVASVAGDSGHMCINMEWGAFGDDGSLGMLSTCFDASV
DQASINPGKQRFEKMISGMYLGEIVRHILLHLTSLGVLFRGQQTQCLQTRDIFKTKFLSEIESDSLALRQ
VRAILEDLGLPLTSDDALMVLEVCQAVSRRAAQLCGAGVAAVVEKIRENRGLEELTVSVGVDGTLYKLHP
HFSSLVAATVRELAPRCAVTFLQSEDGSGKGAALVTAVACRLAQTARV
>anc180_altAll|avgPP:0.994|lnPP:-10.712|num_ambig:7|num_ambig_gaps:0
LQQFKVTRTQLQQIQASLLGSMEQALKGQASPAPAVRMLPTYVGSTPHGTEQGDFLVLELGATGASLRVL
WVTLTGTEGHRVEPRSQEFVIPQEVVLGAGQQLFDFAARCLSEFLDAHPVGNQGLQLGFSFSFPCHQTGL
DRSTLISWTKGFRCSGVEGQDVVQLLRDAIQRQGTYNIDVVAVVNDTVGTMMGCELGARPCEVGLIVDTG
TNACYMEEARHVAALDEDRGRTCVSVEWGSFNDDGALGPVLTTFDHALDHESLNPGAQRFEKMIGGLYLG
ELVRLVLAHLAQHGVLFGGCTSPALLSQGSILLEHVAEMEDPSTGAAHVHTILQDLGLSPQASDAELVQR
VCAAVCTRAAQLCAAALAAVLSRLQHSREQQTLQVAVATGGRVFERHPRFHSILKETVMLLAPECDVSFI
PSVDGGGRGVAMVTAVAARLAAHRRLLEETLAPFQLNREQLAAVQAQMREAMVRGLRGEASSLRMLPTYV
RATPDGSERGDFLALDLGGTNFRVLLVRVTEGGVQITNQVYSIPECVAQGSGQQLFDHIVDCIVDFQQRQ
GLSGQSLPLGFTFSFPCKQLGLDQGILLNWTKGFNASDCEGQDVVCLLREAIRRRQAVELNVVAIVNDTV
GTMMSCGYEDPRCEMGLIVGTGTNACYMEELRNVASVAGDSGHMCINMEWGAFGDDGSLGMLSTCFDASV
DQASINPGKQRFEKMISGMYLGEIVRHILLHLTSLGVLFRGQQTQCLQTRDIFKTKFLSEIESDSLALRQ
VRAILEDLGLPLTSDDALMVLEVCQAVSRRAAQLCGAGVAAVVEKIRENRGLEELTVSVGVDGTLYKLHP
HFSSLVAATVRELAPRCTVTFLQSEDGSGKGAALVTAVACRLAQTTRV
>anc181|avgPP:0.995|lnPP:-7.398|num_ambig:10|num_ambig_gaps:0
LQQFKVTRTQLQQIQASLLGSMEQALKGQASPAPAVRMLPTYVGSTPHGTEQGDFLVLELGATGASLRVL
WVTLTGTEGHRVEPRSQEFVIPQEVVLGTGQQLFDFAARCLSEFLDAHPVGNQGLQLGFSFSFPCHQTGL
DRSTLISWTKGFRCSGVEGQDVVQLLRDAIQRQGTYNIDVVAVVNDTVGTMMGCELGARPCEVGLIVDTG
TNACYMEEARHVAVLDEDRGRTCVSVEWGSFNDDGALGPVLTTFDHALDHESLNPGAQRFEKMIGGLYLG
ELVRLVLAHLAQRGVLFGGCTSPALLSQGSILLEHVAEMEDPSTGAARVHAILQDLGLSPEASDAELVQR
VCAAVCTRAAQLCAAALAAVLSRLQHSREQQTLQVAVATGGRVFEQHPRFHSILKETVMLLAPECDVSFI
PSVDGGGRGVAMVTAVAARLAAHRHLLEETLAPFRLNREQLAAVQAQMREAMARGLRGEASSLRMLPTYV
RATPDGSERGDFLALDLGGTNFRVLLVRVTEGGVQITNQVYSIPECVAQGSGQQLFDHIVDCIVDFQQRQ
GLSGQSLPLGFTFSFPCKQLGLDQGILLNWTKGFNASDCEGQDVVCLLREAIRRRQAVELNVVAIVNDTV
GTMMSCGYEDPRCEMGLIVGTGTNACYMEELRNVAGVAGDSGHMCINMEWGAFGDDGSLGMLSTCFDASV
DQASINPGKQRFEKMISGMYLGEIVRHILLHLTSLGVLFRGQQTQCLQTRDIFKTKFLSEIESDSLALRQ
VRAILEDLGLPLTSDDALMVLEVCQAVSRRAAQLCGAGVAAVVEKIRENRGLEELTVSVGVDGTLYKLHP
HFSSLVAATVRELAPRCAVTFLQSEDGSGKGAALVTAVACRLAQTARV
>anc181_altAll|avgPP:0.993|lnPP:-12.962|num_ambig:10|num_ambig_gaps:0
LQQFKVTRTQLQQIQASLLGSMEQALRGQASPAPAVRMLPTYVGSTPHGTEQGDFLVLELGATGASLRVL
WVTLTGTEGHRVEPRSQEFVIPQEVVLGAGQQLFDFAARCLSEFLDAHPVGNQGLQLGFSFSFPCHQTGL

DRSTLISWTKGFRCSGVEGQDVVQLLRDAIQRQGTYNIDVVAVVNDTVGTMMGCELGARPCEVGLIVDTG
TNACYMEEARHVAVLDEDRGRTCVSVEWGSFNDDGALGPVLTTFDHALDHESLNPGAQRFEKMIGGLYLG
ELVRLVLAHLAQRGVLFGGCTSPALLSQGSILLEHVAEMEDPSTGAAHVHTILQDLGLSPEASDAELVQR
VCAAVCTRAAQLCAAALAAVLSRLQHSREQQTLQVAVATGGRVFEQHPRFHSILQETVMLLAPECDVSFI
PSVDGGGRGVAMVTAVAARLAAHRRLLEETLAPFRLNREQLAAVQAQMREAMARGLRGEASSLRMLPTYV
RATPDGSERGDFLALDLGGTNFRVLLVRVTEGGVQITNQVYSIPECVAQGSGQQLFDHIVDCIVDFQQKQ
GLSGQSLPLGFTFSFPCKQLGLDQGILLNWTKGFNASDCEGQDVVCLLREAIRRRQAVELNVVAIVNDTV
GTMMSCGYEDPRCEMGLIVGTGTNACYMEELRNVAGVAGDSGHMCINMEWGAFGDDGSLGMLSTCFDESV
DQASINPGKQRFEKMISGMYLGEIVRHILLHLTSLGVLFRGQQTQCLQTRDIFKTKFLSEIESDSLALRQ
VRAILEDLGLPLTSDDALMVLEVCQAVSRRAAQLCGAGVAAVVEKIRENRGLEELTVSVGVDGTLYKLHP
HFSSLVAATVRELAPRCTVTFLQSEDGSGKGAALVTAVACRLAQTTRV
>anc182|avgPP:0.997|lnPP:-4.382|num_ambig:5|num_ambig_gaps:0
LQQFKVTRTQLQQIQASLLGSMEQALKGQASPAPAVRMLPTYVGSTPHGTEQGDFLVLELGATGASLRVL
WVTLTGIEGHRVEPRSQEFVIPQEVVLGTGQQLFDFAARCLSEFLDAHPVGNQGLQLGFSFSFPCHQTGL
DRSTLISWTKGFRCSGVEGQDVVQLLRDAIQRQGAYNIDVVAVVNDTVGTMMGCEPGVRPCEVGLIVDTG
TNACYMEEARHVAVLDEDRGRTCVSVEWGSFNDDGALGPVLTTFDHALDHESLNPGAQRFEKMIGGLYLG
ELVRLVLAHLAQRGVLFGGCTSPALLSQGSILLEHVAEMEDPSTGAARVHAILQDLGLSPEASDAELVQR
VCAAVCTRAAQLCAAALAAVLSRLQHSREQQTLQVAVATGGRVFEQHPRFHSILQETVMLLAPECDVSFI
PSVDGGGRGVAMVTAVAARLAAHRRLLEETLAPFRLNREQLAAVQAQMREAMAKGLRGEASSLRMLPTYV
RATPDGSERGDFLALDLGGTNFRVLLVRVTEGGVQITNQVYSIPECVAQGSGQQLFDHIVDCIVDFQQKQ
GLSGQSLPLGFTFSFPCKQLGLDQGILLNWTKGFNASDCEGQDVVCLLREAIRRRQAVELNVVAIVNDTV
GTMMSCGYEDPRCEMGLIVGTGTNACYMEELRNVAGVAGDSGHMCINMEWGAFGDDGSLGMLSTCFDASV
DQASINPGKQRFEKMISGMYLGEIVRHILLHLTSLGVLFRGQQTQRLQTRDIFKTKFLSEIESDSLALRQ
VRAILEDLGLPLTSDDALMVLEVCQAVSRRAAQLCGAGVAAVVEKIRENRGLEELTVSVGVDGTLYKLHP
HFSSLVAATVRELAPRCAVTFLQSEDGSGKGAALVTAVACRLAQTTRV
>anc182_altAll|avgPP:0.996|lnPP:-6.493|num_ambig:5|num_ambig_gaps:0
LQQFKVTRTQLQQIQASLLGSMEQALRGQASPAPAVRMLPTYVGSTPHGTEQGDFLVLELGATGASLRVL
WVTLTGIEGHRVEPRSQEFVIPQEVVLGTGQQLFDFAARCLSEFLDAHPVGNQGLQLGFSFSFPCHQTGL
DRSTLISWTKGFRCSGVEGQDVVQLLRDAIQRQGAYNIDVVAVVNDTVGTMMGCEPGVRPCEVGLIVDTG
TNACYMEEARHVAVLDEDRGRTCVSVEWGSFNDDGALGPVLTTFDHALDHESLNPGAQRFEKMIGGLYLG
ELVRLVLAHLAQRGVLFGGCTSPALLSQGSILLEHVAEMEDPSTGAARVHTILQDLGLSPEASDAELVQR
VCAAVCTRAAQLCAAALAAVLSRLQHSREQQTLQVAVATGGRVFEQHPRFHSILQETVMLLAPECDVSFI
PSVDGGGRGVAMVTAVAARLAAHRRLLEETLAPFRLNREQLAAVQAQMREAMAKGLRGEASSLRMLPTYV
RATPDGSERGDFLALDLGGTNFRVLLVRVTEGGVQITNQVYSIPECVAQGSGQQLFDHIVDCIVDFQQKQ
GLSGQSLPLGFTFSFPCKQLGLDQGILLNWTKGFNASDCEGQDVVCLLREAIRRRQAVELNVVAIVNDTV
GTMMSCGYEDPRCEMGLIVGTGTNACYMEELRNVAGVAGDSGHMCINMEWGAFGDDGSLGMLSTCFDESV
DQASINPGKQRFEKMISGMYLGEIVRHILLHLTSLGVLFRGQQTQCLQTRDIFKTKFLSEIESDSLALRQ
VRAILEDLGLPLTSDDALMVLEVCQAVSRRAAQLCGAGVAAVVEKIRENRGLEELTVSVGVDGTLYKLHP
HFSSLVAATVRELAPRCTVTFLQSEDGSGKGAALVTAVACRLAQTTRV
>anc183|avgPP:0.997|lnPP:-4.381|num_ambig:5|num_ambig_gaps:0
LQQFKVTRTQLQQIQASLLGSMEQALKGQASPAPAVRMLPTYVGSTPHGTEQGDFLVLELGATGASLRVL
WVTLTGIEGHRVEPRSQEFVIPQEVVLGTGQQLFDFAARCLSEFLDAHPVGNQGLQLGFSFSFPCHQTGL
DRSTLISWTKGFRCSGVEGQDVVQLLRDAIQRQGAYNIDVVAVVNDTVGTMMGCEPGVRPCEVGLIVDTG
TNACYMEEARHVAVLDEDRGRTCVSVEWGSFNDDGALGPVLTTFDHALDHESLNPGAQRFEKMIGGLYLG
ELVRLVLAHLAQRGVLFGGCTSPALLSQGSILLEHVAEMEDPSTGAARVHAILQDLGLSPEASDAELVQR
VCAAVCTRAAQLCAAALAAVLSRLQHSREQQTLQVAVATGGRVFEQHPRFHSILQETVMLLAPECDVSFI
PSVDGGGRGVAMVTAVAARLAAHRRLLEETLAPFRLNREQLAAVQAQMREAMAKGLRGEASSLRMLPTYV
RATPDGSERGDFLALDLGGTNFRVLLVRVTEGGVQITNQVYSIPECVAQGSGQQLFDHIVDCIVDFQQKQ
GLSGQSLPLGFTFSFPCKQLGLDQGILLNWTKGFNASDCEGQDVVCLLREAIRRRQAVELNVVAIVNDTV
GTMMSCGYEDPRCEMGLIVGTGTNACYMEELRNVAGVAGDSGHMCINMEWGAFGDDGSLGMLSTCFDASV
DQASINPGKQRFEKMISGMYLGEIVRHILLHLTSLGVLFRGQQTQRLQTRDIFKTKFLSEIESDSLALRQ
VRAILEDLGLPLTSDDALMVLEVCQAVSRRAAQLCGAGVAAVVEKIRENRGLEELTVSVGVDGTLYKLHP

HFSSLVAATVRELAPRCAVTFLQSEDGSGKGAALVTAVACRLAQTTRV
>anc183_altAll|avgPP:0.996|lnPP:-6.493|num_ambig:5|num_ambig_gaps:0
LQQFKVTRTQLQQIQASLLGSMEQALRGQASPAPAVRMLPTYVGSTPHGTEQGDFLVLELGATGASLRVL
WVTLTGIEGHRVEPRSQEFVIPQEVVLGTGQQLFDFAARCLSEFLDAHPVGNQGLQLGFSFSFPCHQTGL
DRSTLISWTKGFRCSGVEGQDVVQLLRDAIQRQGAYNIDVVAVVNDTVGTMMGCEPGVRPCEVGLIVDTG
TNACYMEEARHVAVLDEDRGRTCVSVEWGSFNDDGALGPVLTTFDHALDHESLNPGAQRFEKMIGGLYLG
ELVRLVLAHLAQRGVLFGGCTSPALLSQGSILLEHVAEMEDPSTGAARVHTILQDLGLSPEASDAELVQR
VCAAVCTRAAQLCAAALAAVLSRLQHSREQQTLQVAVATGGRVFEQHPRFHSILQETVMLLAPECDVSFI
PSVDGGGRGVAMVTAVAARLAAHRRLLEETLAPFRLNREQLAAVQAQMREAMAKGLRGEASSLRMLPTYV
RATPDGSERGDFLALDLGGTNFRVLLVRVTEGGVQITNQVYSIPECVAQGSGQQLFDHIVDCIVDFQQKQ
GLSGQSLPLGFTFSFPCKQLGLDQGILLNWTKGFNASDCEGQDVVCLLREAIRRRQAVELNVVAIVNDTV
GTMMSCGYEDPRCEMGLIVGTGTNACYMEELRNVAGVAGDSGHMCINMEWGAFGDDGSLGMLSTCFDESV
DQASINPGKQRFEKMISGMYLGEIVRHILLHLTSLGVLFRGQQTQCLQTRDIFKTKFLSEIESDSLALRQ
VRAILEDLGLPLTSDDALMVLEVCQAVSRRAAQLCGAGVAAVVEKIRENRGLEELTVSVGVDGTLYKLHP
HFSSLVAATVRELAPRCTVTFLQSEDGSGKGAALVTAVACRLAQTTRV
>anc184|avgPP:0.997|lnPP:-4.513|num_ambig:6|num_ambig_gaps:0
LQQFKVTRAQLQQIQASLLGSMEQALRGQASPAPAVRMLPTYVGSTPHGTEQGDFLVLELGATGASLRVL
WVTLTGIEGHRVEPRSQEFVIPQEVMLGTGQQLFDFAARCLSEFLDAHPVGNQGLQLGFSFSFPCHQTGL
DRSTLISWTKGFRCSGVEGQDVVQLLRDAIQRQGAYNIDVVAVVNDTVGTMMGCEPGVRPCEVGLVVDTG
TNACYMEEARHVAVLDEDRGRVCVSVEWGSFNDDGALGPVLTTFDHALDHESLNPGAQRFEKMIGGLYLG
ELVRLVLAHLAQRGVLFGGCTSPALLSQGSILLEHVAEMEDPSTGAARVHAILQDLGLSPEASDAELVQR
VCAAVCTRAAQLCAAALAAVLSRLQHSREQQTLQVAVATGGRVFEQHPRFHSILQETVMLLAPECDVSFI
PSVDGGGRGVAMVTAVAARLAAHRRLLEETLAPFRLNREQLAAVQAQMREAMAKGLRGEASSLRMLPTYV
RATPDGSERGDFLALDLGGTNFRVLLVRVTEGGVQITNQVYSIPECVAQGSGQQLFDHIVDCIVDFQQKQ
GLSGQSLPLGFTFSFPCKQLGLDQGILLNWTKGFNASDCEGQDVVCLLREAIRRRQAVELNVVAIVNDTV
GTMMSCGYEDPRCEMGLIVGTGTNACYMEELRNVAGVAGDSGHMCINMEWGAFGDDGSLGMLSTCFDASV
DQASINPGKQRFEKMISGMYLGEIVRHILLHLTSLGVLFRGQQTQRLQTRDIFKTKFLSEIESDSLALRQ
VRAILEDLGLPLTSDDALMVLEVCQAVSRRAAQLCGAGVAAVVEKIRENRGLEELTVSVGVDGTLYKLHP
HFSSLVAATVRELAPRCAVTFLQSEDGSGKGAALVTAVACRLAQTTRV
>anc184_altAll|avgPP:0.996|lnPP:-8.060|num_ambig:6|num_ambig_gaps:0
LQQFKVTRAQLQQIQASLLGSMEQALRGQASPAPAVRMLPTYVGSTPHGTEQGDFLVLELGATGASLRVL
WVTLTGIEGHRVEPRSQEFVIPQEVMLGTGQQLFDFAARCLSEFLDAHPVGNQGLQLGFSFSFPCHQTGL
DRSTLISWTKGFRCSGVEGQDVVQLLRDAIQRQGAYNIDVVAVVNDTVGTMMGCEPGVRPCEVGLVVDTG
TNACYMEEARHVAVLDEDRGRVCVSVEWGSFNDDGALGPVLTTFDHALDHESLNPGAQRFEKMIGGLYLG
ELVRLVLAHLAQRGVLFGGCTSPALLSQGSILLEHVAEMEDPSAGAARVHTILQDLGLSPEASDAELVQH
VCAAVCTRAAQLCAAALAAVLSRLQHSREQQTLQVAVATGGRVFERHPRFHSILQETVMLLAPECDVSFI
PSVDGGGRGVAMVTAVAARLAAHRRLLEETLAPFRLNREQLAAVQAQMREAMAKGLRGEASSLRMLPTYV
RATPDGSERGDFLALDLGGTNFRVLLVRVTEGGVQITNQVYSIPECVAQGSGQQLFDHIVDCIVDFQQKQ
GLSGQSLPLGFTFSFPCKQLGLDQGILLNWTKGFNASDCEGQDVVCLLREAIRRRQAVELNVVAIVNDTV
GTMMSCGYEDPRCEMGLIVGTGTNACYMEELRNVAGVAGDSGHMCINMEWGAFGDDGSLGMLSTCFDESV
DQASINPGKQRFEKMISGMYLGEIVRHILLHLTSLGVLFRGQQTQRLQTRDIFKTKFLSEIESDSLALRQ
VRAILEDLGLPLTSDDALMVLEVCQAVSRRAAQLCGAGVAAVVEKIRENRGLEELTVSVGVDGTLYKLHP
HFSSLVAATVRELAPRCTVTFLQSEDGSGKGAALVTAVACRLAQTTRV
>anc185|avgPP:0.995|lnPP:-7.650|num_ambig:8|num_ambig_gaps:0
LQQFKVTEAQLQQIQASLLASMEQALRGQASPAPAVRMLPTYVGSTPHGTEQGDFLVLELGAKGASLRVL
WVTLTGIEGHRVEPRSQEFVIPQEVMLGAGQQLFDFAAHCLSEFLDAHPVSNQGLQLGFSFSFPCHQTGL
DRSTLISWTKGFRCSGVEGQDVVQLLRDAIQRQGAYNIDVVAVVNDTVGTMMGCEPGVRPCEVGLVVDTG
TNACYMEEARHVAVLDDDRGRMCVSVEWGSFSDDGALGPVLTTFDHALDHESLNPGAQRFEKMIGGLYLG
ELVRLVLVHLARRGVLFGGCTSPALLSQGSILLEHVAEMEDPSAGAARVHAILQDLGLSPEASDAELVQH
VCAAVCTRAAQLCAAALAAVLSRLQRSREHQTLQVAVATGGRVCEQHPRFCSILQETVMLLAPECDVSFV
PSVDGGGRGVAMVTAVAARLAAHRRLLEETLAPFRLNREQLAAVQAQMREAMAKGLRGEASSLRMLPTYV
RATPDGSERGDFLALDLGGTNFRVLLVRVAAGGVQITSQVYSIPECVAQGSGQQLFDHIVDCIVDFQQKQ

GLSGQSLPLGFTFSFPCRQLGLDQGILLNWTKGFNASDCEGQDVVCLLREAIRRRQAVELNVVAIVNDTV
GTMMSCGYEDPRCEIGLIVGTGTNACYMEELRNVAGVAGDSGHMCINMEWGAFGDDGSLGMLSTRFDASV
DQASINPGKQRFEKMISGMYLGEIVRHILLHLTSLGVLFRGQQTQRLQTRDIFKTKFLSEIESDSLALRQ
VRAILEDLGLPLTSDDALMVLEVCQAVSQRAAQLCGAGVAAVVERIRENRGLEELTVSVGVDGTLYKLHP
HFSSLVAATVRELAPRCAVTFLQSEDGSGKGAALVTAVACRLAQKTRV
>anc185_altAll|avgPP:0.994|lnPP:-10.550|num_ambig:8|num_ambig_gaps:0
LQQFKVTEAQLQQIQASLLGSMEQALRGQASPAPAVRMLPTYVGSTPHGTEQGDFLVLELGAKGASLRVL
WVTLTGIEGHRVEPRSQEFVIPQEVMLGAGQQLFDFAAHCLSEFLDAHPVSSQGLQLGFSFSFPCHQTGL
DRSTLISWTKGFRCSGVEGQDVVQLLRDAIQRQGAYNIDVVAVVNDTVGTMMGCEPGVRPCEVGLVVDTG
TNACYMEEARHVAVLDEDRGRMCVSVEWGSFSDDGALGPVLTTFDHALDHESLNPGAQRFEKMIGGLYLG
ELVRLVLVHLARRGVLFGGCTSPALLSQGSILLEHVAEMEDPSAGAARVHAILQDLGLSPEASDAELVQH
VCAAVCTRAAQLCAAALAAVLSRLQRSREHQTLQVAVATGGRVFEQHPRFCSILQETVMLLAPECDVSFI
PSVDGGGRGVAMVTAVAARLAAHRRLLEETLAPFRLNREQLAAVQAQMREAMAKGLRGEASSLRMLPTYV
RATPDGSERGDFLALDLGGTNFRVLLVRVAAGGVQITSQVYSIPECVAQGSGQQLFDHIVDCIVDFQQKQ
GLSGQSLPLGFTFSFPCRQLGLDQGILLNWTKGFNASDCEGQDVVSLLREAIRRRQAVELNVVAIVNDTV
GTMMSCGYEDPRCEIGLIVGTGTNACYMEELRNVAGVAGDSGHMCINMEWGAFGDDGSLGMLSTCFDASV
DQASINPGKQRFEKMISGMYLGEIVRHILLHLTSLGVLFRGQQIQRLQTRDIFKTKFLSEIESDSLALRQ
VRAILEDLGLPLTSDDALMVLEVCQAVSQRAAQLCGAGVAAVVERIRENRGLEELTVSVGVDGTLYKLHP
HFSSLVAATVRELAPRCAVTFLQSEDGSGKGAALVTAVACRLAQKTRV
>anc186|avgPP:0.995|lnPP:-7.244|num_ambig:6|num_ambig_gaps:0
LQQFKVTEAQLQQIQASLLGSMEQALRGQASPAPAVRMLPTYVGSTPHGTEQGDFVVLELGATGASLRVL
WVTLTGIEGHRVEPRSQEFVIPQEVMLGAGQQLFDFAAHCLSEFLDAQPVSKQGLQLGFSFSFPCHQTGL
DRSTLISWTKGFRCSGVEGQDVVQLLRDAIQRQGAYNIDVVAVVNDTVGTMMGCEPGVRPCEVGLVVDTG
TNACYMEEARHVAVLDEDRGRVCVSVEWGSFSDDGALGPVLTTFDHTLDHESLNPGAQRFEKMIGGLYLG
ELVRLVLAHLARRGVLFGGCTSPALLSQGSILLEHVAEMEDPSAGAARVHAILQDLGLSPEASDVELVQH
VCAAVCTRAAQLCAAALAAVLSRLQHSREQQTLQVAVATGGRVCERHPRFCSILQGTVMLLAPECDVSFI
PSVDGGGRGVAMVTAVAARLAAHRRLLEETLAPFRLNHDQLAAVQAQMREAMAKGLRGEASSLRMLPTYV
RATPDGSERGDFLALDLGGTNFRVLLVRVTTGGVQITSQIYSIPETVAQGSGQQLFDHIVDCIVDFQQKQ
GLSGQSLPLGFTFSFPCRQLGLDQGILLNWTKGFNASDCEGQDVVSLLREAITRRQAVELNVVAIVNDTV
GTMMSCGYEDPRCEIGLIVGTGTNACYMEELRNVAGVAGDSGHMCINMEWGAFGDDGSLGMLSTRFDASV
DQASINPGKQRFEKMISGMYLGEIVRHILLHLTSLGVLFRGQQTQRLQTRDIFKTKFLSEIESDSLALRQ
VRAILEDLGLPLTSDDALMVLEVCQAVSQRAAQLCGAGVAAVVEKIRENRGLEELAVSVGVDGTLYKLHP
HFSSLVAATVRELAPRCAVTFLQSEDGSGKGAALVTAVACRLAQTTRV
>anc186_altAll|avgPP:0.994|lnPP:-10.022|num_ambig:6|num_ambig_gaps:0
LQQFKVTEAQLQQIQASLLGSMEQALRGQASPAPAVRMLPTYVGSTPHGTEQGDFVVLELGATGASLRVL
WVTLTGIEGHRVEPRSQEFVIPQEVMLGAGQQLFDFAAHCLSEFLDAQPVSKQGLQLGFSFSFPCHQTGL
DRSTLISWTKGFRCSGVEGQDVVQLLRDAIRRQGAYNIDVVAVVNDTVGTMMGCEPGVRPCEVGLVVDTG
TNACYMEEARHVAVLDEDRGRVCVSVEWGSFSDDGALGPVLTTFDHTLDHESLNPGAQRFEKMIGGLYLG
ELVRLVLAHLARRGVLFGGCTSPALLSQGSILLEHVAEMEDPSTGAARVHAILQDLGLSPEASDVELVQH
VCAAVCTRAAQLCAAALAAVLSRLQHSREQQTLQVAVATGGRVCERHPRFCSILQGTVMLLAPECDVSFI
PSVDGGGRGVAMVTAVAARLAAHRRLLEETLAPFRLNHDQLAAVQAQMREAMAKGLRGEASSLRMLPTYV
RATPDGSERGDFLALDLGGTNFRVLLVRVTTGGVQITSQIYSIPESVAQGSGQQLFDHIVDCIVDFQQKQ
GLSGQSLPLGFTFSFPCRQLGLDQGILLNWTKGFNASDCEGQDVVSLLREAITRRQAVELNVVAIVNDTV
GTMMSCGYEDPRCEIGLIVGTGTNACYMEELRNVAGVPGDSGRMCINMEWGAFGDDGSLGMLSTRFDASV
DQASINPGKQRFEKMISGMYLGEIVRHILLHLTSLGVLFRGQQTQRLQTRDIFKTKFLSEIESDSLALRQ
VRAILEDLGLPLTSDDALMVLEVCQAVSQRAAQLCGAGVAAVVEKIRENRGLEELAVSVGVDGTLYKLHP
HFSSLVAATVRELAPRCAVTFLQSEDGSGKGAALVTAVACRLAQSTRV
>anc187|avgPP:0.997|lnPP:-4.586|num_ambig:3|num_ambig_gaps:0
LQQFKVTEAQLQQIQASLLGSMEQALRGQASPAPAVRMLPTYVGSTPHGTEQGDFVVLELGATGASLRVL
WVTLTGIEGHRVEPRSQEFVIPQEVMLGAGQQLFDFAAHCLSEFLDAHPVSNQGLQLGFSFSFPCHQTGL
DRSTLISWTKGFRCSGVEGQDVVQLLRDAIQRQGAYNIDVVAVVNDTVGTMMGCEPGVRPCEVGLVVDTG
TNACYMEEARHVAVLDEDRGRVCVSVEWGSFSDDGALGPVLTTFDHALDHESLNPGAQRFEKMIGGLYLG

ELVRLVLAHLARRGVLFGGCTSPALLSQGSILLEHVAEMEDPSAGAARVHAILQDLGLSPEASDAELVQH
VCAAVCTRAAQLCAAALAAVLSRLQHSREQQTLQVAVATGGRVCERHPRFCSILQETVMLLAPECDVSFI
PSVDGGGRGVAMVTAVAARLAAHRRLLEETLAPFRLNREQLAAVQAQMREAMAKGLRGEASSLRMLPTYV
RATPDGSERGDFLALDLGGTNFRVLLVRVAAGGVQITSQVYSIPECVAQGSGQQLFDHIVDCIVDFQQKQ
GLSGQSLPLGFTFSFPCRQLGLDQGILLNWTKGFNASDCEGQDVVSLLREAITRRQAVELNVVAIVNDTV
GTMMSCGYEDPRCEIGLIVGTGTNACYMEELRNVAGVAGDSGHMCINMEWGAFGDDGSLGMLSTRFDASV
DQASINPGKQRFEKMISGMYLGEIVRHILLHLTSLGVLFRGQQTQRLQTRDIFKTKFLSEIESDSLALRQ
VRAILEDLGLPLTSDDALMVLEVCQAVSQRAAQLCGAGVAAVVEKIRENRGLEELTVSVGVDGTLYKLHP
HFSSLVAATVRELAPRCAVTFLQSEDGSGKGAALVTAVACRLAQTTRV
>anc187_altAll|avgPP:0.996|lnPP:-6.011|num_ambig:3|num_ambig_gaps:0
LQQFKVTEAQLQQIQASLLGSMEQALRGQASPAPAVRMLPTYVGSTPHGTEQGDFVVLELGATGASLRVL
WVTLTGIEGHRVEPRSQEFVIPQEVMLGAGQQLFDFAAHCLSEFLDAHPVSSQGLQLGFSFSFPCHQTGL
DRSTLISWTKGFRCSGVEGQDVVQLLRDAIQRQGAYNIDVVAVVNDTVGTMMGCEPGVRPCEVGLVVDTG
TNACYMEEARHVAVLDEDRGRVCVSVEWGSFSDDGALGPVLTTFDHALDHESLNPGAQRFEKMIGGLYLG
ELVRLVLAHLARRGVLFGGCTSPALLSQGSILLEHVAEMEDPSTGAARVHAILQDLGLSPEASDAELVQH
VCAAVCTRAAQLCAAALAAVLSRLQHSREQQTLQVAVATGGRVCERHPRFCSILQETVMLLAPECDVSFI
PSVDGGGRGVAMVTAVAARLAAHRRLLEETLAPFRLNREQLAAVQAQMREAMAKGLRGEASSLRMLPTYV
RATPDGSERGDFLALDLGGTNFRVLLVRVAAGGVQITSQVYSIPECVAQGSGQQLFDHIVDCIVDFQQKQ
GLSGQSLPLGFTFSFPCRQLGLDQGILLNWTKGFNASDCEGQDVVSLLREAIRRRQAVELNVVAIVNDTV
GTMMSCGYEDPRCEIGLIVGTGTNACYMEELRNVAGVAGDSGHMCINMEWGAFGDDGSLGMLSTRFDASV
DQASINPGKQRFEKMISGMYLGEIVRHILLHLTSLGVLFRGQQTQRLQTRDIFKTKFLSEIESDSLALRQ
VRAILEDLGLPLTSDDALMVLEVCQAVSQRAAQLCGAGVAAVVEKIRENRGLEELTVSVGVDGTLYKLHP
HFSSLVAATVRELAPRCAVTFLQSEDGSGKGAALVTAVACRLAQTTRV
>anc188|avgPP:0.997|lnPP:-4.887|num_ambig:6|num_ambig_gaps:0
LQQFKVTEAQLQQIQASLLGSMEQALRGQASPAPAVRMLPTYVGSTPHGTEQGDFLVLELGATGASLRVL
WVTLTGIEGHRVEPRSQEFVIPQEVMLGAGQQLFDFAAHCLSEFLDAHPVSNQGLQLGFSFSFPCHQTGL
DRSTLISWTKGFRCSGVEGQDVVQLLRDAIQRQGAYNIDVVAVVNDTVGTMMGCEPGVRPCEVGLVVDTG
TNACYMEEARHVAVLDEDRGRVCVSVEWGSFSDDGALGPVLTTFDHALDHESLNPGAQRFEKMIGGLYLG
ELVRLVLAHLARRGVLFGGCTSPALLSQGSILLEHVAEMEDPSAGAARVHAILQDLGLSPEASDAELVQH
VCAAVCTRAAQLCAAALAAVLSRLQHSREQQTLQVAVATGGRVCERHPRFCSILQETVMLLAPECDVSFI
PSVDGGGRGVAMVTAVAARLAAHRRLLEETLAPFRLNREQLAAVQAQMREAMAKGLRGEASSLRMLPTYV
RATPDGSERGDFLALDLGGTNFRVLLVRVAAGGVQITSQVYSIPECVAQGSGQQLFDHIVDCIVDFQQKQ
GLSGQSLPLGFTFSFPCRQLGLDQGILLNWTKGFNASDCEGQDVVCLLREAIRRRQAVELNVVAIVNDTV
GTMMSCGYEDPRCEIGLIVGTGTNACYMEELRNVAGVAGDSGHMCINMEWGAFGDDGSLGMLSTRFDASV
DQASINPGKQRFEKMISGMYLGEIVRHILLHLTSLGVLFRGQQTQRLQTRDIFKTKFLSEIESDSLALRQ
VRAILEDLGLPLTSDDALMVLEVCQAVSQRAAQLCGAGVAAVVEKIRENRGLEELTVSVGVDGTLYKLHP
HFSSLVAATVRELAPRCAVTFLQSEDGSGKGAALVTAVACRLAQTTRV
>anc188_altAll|avgPP:0.995|lnPP:-8.671|num_ambig:6|num_ambig_gaps:0
LQQFKVTEAQLQQIQASLLGSMEQALRGQASPAPAVRMLPTYVGSTPHGTEQGDFVVLELGATGASLRVL
WVTLTGIEGHRVEPRSQEFVIPQEVMLGAGQQLFDFAAHCLSEFLDAHPVSSQGLQLGFSFSFPCHQTGL
DRSTLISWTKGFRCSGVEGQDVVQLLRDAIQRQGAYNIDVVAVVNDTVGTMMGCEPGVRPCEVGLVVDTG
TNACYMEEARHVAVLDEDRGRVCVSVEWGSFSDDGALGPVLTTFDHALDHESLNPGAQRFEKMIGGLYLG
ELVRLVLAHLARRGVLFGGCTSPALLSQGSILLEHVAEMEDPSTGAARVHAILQDLGLSPEASDAELVQH
VCAAVCTRAAQLCAAALAAVLSRLQHSREQQTLQVAVATGGRVFERHPRFCSILQETVMLLAPECDVSFI
PSVDGGGRGVAMVTAVAARLAAHRRLLEETLAPFRLNREQLAAVQAQMREAMAKGLRGEASSLRMLPTYV
RATPDGSERGDFLALDLGGTNFRVLLVRVAAGGVQITSQVYSIPECVAQGSGQQLFDHIVDCIVDFQQKQ
GLSGQSLPLGFTFSFPCRQLGLDQGILLNWTKGFNASDCEGQDVVSLLREAIRRRQAVELNVVAIVNDTV
GTMMSCGYEDPRCEIGLIVGTGTNACYMEELRNVAGVAGDSGHMCINMEWGAFGDDGSLGMLSTCFDASV
DQASINPGKQRFEKMISGMYLGEIVRHILLHLTSLGVLFRGQQTQRLQTRDIFKTKFLSEIESDSLALRQ
VRAILEDLGLPLTSDDALMVLEVCQAVSQRAAQLCGAGVAAVVEKIRENRGLEELTVSVGVDGTLYKLHP
HFSSLVAATVRELAPRCAVTFLQSEDGSGKGAALVTAVACRLAQTTRV
>anc189|avgPP:0.997|lnPP:-4.493|num_ambig:4|num_ambig_gaps:0

LQQFKVTEAQLQQIQASLLGSMEQALRGQASPAPAVRMLPTYVGSTPHGTEQGDFLVLELGATGASLRVL
WVTLTGIEGHRVEPRSQEFVIPQEVMLGAGQQLFDFAARCLSEFLDAHPVGNQGLQLGFSFSFPCHQTGL
DRSTLISWTKGFRCSGVEGQDVVQLLRDAIQRQGAYNIDVVAVVNDTVGTMMGCEPGVRPCEVGLVVDTG
TNACYMEEARHVAVLDEDRGRVCVSVEWGSFSDDGALGPVLTTFDHALDHESLNPGAQRFEKMIGGLYLG
ELVRLVLAHLAQRGVLFGGCTSPALLSQGSILLEHVAEMEDPSAGAARVHAILQDLGLSPEASDAELVQH
VCAAVCTRAAQLCAAALAAVLSRLQHSREQQTLQVAVATGGRVFERHPRFHSILQETVMLLAPECDVSFI
PSVDGGGRGVAMVTAVAARLAAHRRLLEETLAPFRLNREQLAAVQAQMREAMAKGLRGEASSLRMLPTYV
RATPDGSERGDFLALDLGGTNFRVLLVRVAAGGVQITNQVYSIPECVAQGSGQQLFDHIVDCIVDFQQKQ
GLSGQSLPLGFTFSFPCRQLGLDQGILLNWTKGFNASDCEGQDVVCLLREAIRRRQAVELNVVAIVNDTV
GTMMSCGYEDPRCEIGLIVGTGTNACYMEELRNVAGVAGDSGHMCINMEWGAFGDDGSLGMLSTCFDASV
DQASINPGKQRFEKMISGMYLGEIVRHILLHLTSLGVLFRGQQTQRLQTRDIFKTKFLSEIESDSLALRQ
VRAILEDLGLPLTSDDALMVLEVCQAVSQRAAQLCGAGVAAVVEKIRENRGLEELTVSVGVDGTLYKLHP
HFSSLVAATVRELAPRCAVTFLQSEDGSGKGAALVTAVACRLAQTTRV
>anc189_altAll|avgPP:0.996|lnPP:-7.752|num_ambig:4|num_ambig_gaps:0
LQQFKVTEAQLQQIQASLLGSMEQALRGQASPAPAVRMLPTYVGSTPHGTEQGDFVVLELGATGASLRVL
WVTLTGIEGHRVEPRSQEFVIPQEVMLGPGQQLFDFAARCLSEFLDAHPVGNQGLQLGFSFSFPCHQTGL
DRSTLISWTKGFRCSGVEGQDVVQLLRDAIQRQGAYNIDVVAVVNDTVGTMMGCEPGVRPCEVGLVVDTG
TNACYMEEARHVAVLDEDRGRVCVSVEWGSFSDDGALGPVLTTFDHALDHESLNPGAQRFEKMIGGLYLG
ELVRLVLAHLAQRGVLFGGCTSPALLSQGSILLEHVAEMEDPSTGAARVHAILQDLGLSPEASDAELVQH
VCAAVCTRAAQLCAAALAAVLSRLQHSREQQTLQVAVATGGRVFERHPRFHSILQETVMLLAPECDVSFI
PSVDGGGRGVAMVTAVAARLAAHRRLLEETLAPFRLNREQLAAVQAQMREAMAKGLRGEASSLRMLPTYV
RATPDGSERGDFLALDLGGTNFRVLLVRVAAGGVQITNQVYSIPECVAQGSGQQLFDHIVDCIVDFQQKQ
GLSGQSLPLGFTFSFPCRQLGLDQGILLNWTKGFNASDCEGQDVVCLLREAIRRRQAVELNVVAIVNDTV
GTMMSCGYEDPRCEMGLIVGTGTNACYMEELRNVAGVAGDSGHMCINMEWGAFGDDGSLGMLSTCFDASV
DQASINPGKQRFEKMISGMYLGEIVRHILLHLTSLGVLFRGQQTQRLQTRDIFKTKFLSEIESDSLALRQ
VRAILEDLGLPLTSDDALMVLEVCQAVSQRAAQLCGAGVAAVVEKIRENRGLEELTVSVGVDGTLYKLHP
HFSSLVAATVRELAPRCAVTFLQSEDGSGKGAALVTAVACRLAQTTRV
>anc190|avgPP:0.998|lnPP:-3.596|num_ambig:5|num_ambig_gaps:0
LQQFKVTEAQLQQIQASLLGSMEQALRGQASPAPAVRMLPTYVGSTPHGTEQGDFLVLELGATGASLRVL
WVTLTGTEGHRVEPRSQEFVIPQEVMLGPGQQLFDFAARCLSEFLDALPVGNQSLQLGFSFSFPCHQTGL
DRSTLISWTKGFRCSGVEGQDVVQLLRDAIQRQGAYSIDVVAVVNDTVGTMMGCEPGVGPCEVGLVVDTG
TNACYMEEARHVAVLDEDRGRVCVSVEWGSFGDDGALGPVLTTFDHALDQESLNPGAQRFEKMIGGLYLG
ELVRLVLAHLAQRGVLFGGCTSPALLSQGSILLEHVAEMEDPSAGAARVHAILQDLGLSPGASDAELVQH
VCAAVCTRAARLCAAALAAVLSRLQHSREQQTLQIAVATGGRVFERHPRFHSILQETVMLLAPECDVSFI
PSVDGGGRGVAMVTAVAARLAAHRRLLEETLAPFRLNREQLAAVQAQMREAMAKGLRGEASSLRMLPTYV
RATPDGSERGDFLALDLGGTNFRVLLVRVAAGGVQITNQVYSIPECVAQGSGQQLFDHIVDCIVDFQQKQ
GLSGQSLPLGFTFSFPCRQLGLDQGILLNWTKGFNASDCEGQDVVCLLREAIRRRQAVELNVVAIVNDTV
GTMMSCGYEDPRCEVGLIVGTGTNACYMEELRNVAGVAGDSGHMCINMEWGAFGDDGSLDMLSTCFDASV
DQASINPGKQRFEKMISGMYLGEIVRHILLHLTSLGVLFRGQQTQRLQTRDIFKTKFLSEIESDSLALRQ
VRAILEDLGLPLTSDDALMVLEVCQAVSQRAAQLCGAGVAAVVEKIRENRGLEELTVSVGVDGTLYKLHP
HFSSLVAATVRELAPRCAVTFLQSEDGSGKGAALVTAVACRLAQTTRV
>anc190_altAll|avgPP:0.997|lnPP:-6.208|num_ambig:5|num_ambig_gaps:0
LQQFKVTEAQLQQIQASLLGSMEQALRGQASPAPAVRMLPTYVGSTPHGTEQGDFVVLELGATGASLRVL
WVTLTGTEGHRVEPRSQEFVIPQEVMLGPGQQLFDFAARCLSEFLDALPVGNQGLQLGFSFSFPCHQTGL
DRSTLISWTKGFRCSGVEGQDVVQLLRDAIQRQGAYNIDVVAVVNDTVGTMMGCEPGVGPCEVGLVVDTG
TNACYMEEARHVAVLDEDRGRVCVSVEWGSFGDDGALGPVLTTFDHALDHESLNPGAQRFEKMIGGLYLG
ELVRLVLAHLAQRGVLFGGCTSPALLSQGSILLEHVAEMEDPSAGAARVHAILQDLGLSPGASDAELVQH
VCAAVCTRAARLCAAALAAVLSRLQHSREQQTLQIAVATGGRVFERHPRFLSILQETVMLLAPECDVSFI
PSVDGGGRGVAMVTAVAARLAAHRRLLEETLAPFRLNREQLAAVQAQMREAMAKGLRGEASSLRMLPTYV
RATPDGSERGDFLALDLGGTNFRVLLVRVAAGGVQITNQVYSIPECVAQGSGQQLFDHIVDCIVDFQQKQ
GLSGQSLPLGFTFSFPCRQLGLDQGILLNWTKGFNASDCEGQDVVCLLREAIRRRQAVELNVVAIVNDTV
GTMMSCGYEDPRCEVGLIVGTGTNACYMEELRNVAGVAGDSGHMCINMEWGAFGDDGSLDMLSTCFDASV

DQASINPGKQRFEKMISGMYLGEIVRHILLHLTSLGVLFRGQQTQRLQTRDIFKTKFLSEIESDSLALRQ
VRAILEDLGLPLTSDDALMVLEVCQAVSQRAAQLCGAGVAAVVEKIRENRGLEELTVSVGVDGTLYKLHP
HFSSLVAATVRELAPRCAVTFLQSEDGSGKGAALVTAVACRLAQTTRV
>anc191|avgPP:0.991|lnPP:-14.377|num_ambig:17|num_ambig_gaps:0
LQQFKVTEAQLQQIQASLLGSMEQALRGQASPAPAVRMLPTYVGSTPHGTEQGDFLVLELGATGASLRVL
WVTLTGTEGHRVEPRSQEFVIPPEVMLGPGQQLFDFAARCLSEFLDALPVGNQSLQLGFSFSFPCHQTGL
DRSTLISWTKGFRCSGVEGQDVVQLLRDAIQRQGAYSIDVVAVVNDTVGTMMGCDPGVGPCEVGLVVDTG
TNACYMEEARHVAVLDEDRGRVCVSVEWGSFGDDGALGPVLTPFDHALDQESLNPGAQRFEKMIGGLYLG
ELVRLVLAHLAQRGVLFGGCTSPALLSQGSILLEHVAEMEDPSAGAARVHAVLQDLGLSPGASDAELVQH
VCAAVSTRAARLCAAALAAVLSRLQHSREQQTLQIAVATGGRVFERHPRFHSILQETVMLLAPECDVSFI
PSVDGGGRGVAMVTAVAARLAAHRRLLEETLAPFRLNREQLAAVQAQMREAMAKGLRGEASSLRMLPTYV
RATPDGSERGDFLALDLGGTNFRVLLVRVAAGGVQITNQVYSIPECVAQGSGQQLFDHIVDCIVDFQQKQ
GLSGQSLPLGFTFSFPCRQLGLDQGILLNWTKGFNASDCEGQDVVALLREAIRRRQAVELNVVAIVNDTV
GTMMSCGYEDPRCEVGLIVGTGTNACYMEELRNVAGVPGDSGHMCINMEWGAFGDDGSLDVLSTCFDASV
DRASINPGKQRFEKLISGMYLGEIVRHILLHLTSLGVLFRGQQTQRLQTRDIFKTKFLSEIESDSLALRQ
VRAILEDLGLPLTSDDALMVREVCQAVSQRAAQLCGAGVAAVVEKIRENRGLEELTVTVGVDGTLYKLHP
HFSSLVAATVRELAPRCAVTFLQSEDGSGKGAALVTAVACRLAQTTRV
>anc191_altAll|avgPP:0.988|lnPP:-22.387|num_ambig:17|num_ambig_gaps:0
LQQFKVTEAQLQQIQASLLGSMEQALRGQSSPAPAVRMLPTYVGSTPHGTEQGDFVVLELGATGASLRVL
WVTLTGTEGHRVEPRSQEFVIPPEVMLGPGQQLFDFAARCLAEFLDALPVGNQGLQLGFSFSFPCHQTGL
DRSTLISWTKGFRCSGVEGQDVVQLLRDAIQRQGAYNIDVVAVVNDTVGTMMGCDPGLGPCEVGLVVDTG
TNACYMEEARHVAVLDEDRGRVCVSVEWGSFGDDGALGPVLTPFDHALDQESLNPGAQRFEKMIGGLYLG
ELVRLVLAHLAQRGVLFGGCTSPALLSQGSVLLEHVAEMEDPSAGAARVHAILQDLGLSPGASDVELVQH
VCAAVCTRAARLCAAALAAVLSRLQHSREQHTLQIAVATGGRVFERHPRFLSILQETVMLLAPECDVSFI
PSVDGGGRGVAMVTAVAARLAAHRRLLEETLAPFRLNREQLAAVQAQMREAMAKGLRGEASSLRMLPTYV
RATPDGSERGDFLALDLGGTNFRVLLVRVAAGGVQITSQVYSIPECVAQGSGQQLFDHIVDCIVDFQQKQ
GLSGQSLPLGFTFSFPCRQLGLDQGILLNWTKGFNASDCEGQDVVCLLREAIRRRQAVELNVVAIVNDTV
GTMMSCGYEDPRCEVGLIVGTGTNACYMEELRNVAGVPGDSGHMCINMEWGAFGDDGSLDMLSTCFDASV
DQASINPGKQRFEKLISGMYLGEIVRHILLHLTSLGVLFRGQQTQRLQTRDIFKTKFLSEIESDSLALRQ
VRAILEDLGLPLTSDDALMVREVCQAVSRRAAQLCGAGVAAVVEKIRENRGLEELTVTVGVDGTLYKLHP
HFSSLVAATVRELAPRCAVTFLQSEDGSGKGAALVTAVACRLAQTTRV
>anc192|avgPP:0.992|lnPP:-11.885|num_ambig:9|num_ambig_gaps:0
LKQFDVTEEQLQQIQAGILQDMEQALSRQTDPSPTVRMLPTYVRSTPHGTEQGDFLVLELGASGATLRVL
WVTLTGITGPRKEPISREFTIPTEIMLGTGQQLFDFAAGCLVEFLDQLKVGSQPIQLGFNFSFPCHQTGL
DQSTLISWTKGFRCSGVEGQDVVQLLREAIQRQGAYSIEVVAVVNDTVGTMMSCDPRPQACEIGLVVDTG
TNACYMEEARHVAELGEDQGRICVNVQWGSFGDSGALEPVQTPFDLQLDQESLNPGTQRFEKMVGGLYLG
ELVRLVLIHLAGQRALFGGVTSPALLTHGSVHIEHVMEIEDPSHGRARAQAVLQGLGLNPSTQDCEQVQQ
VCGAVSARAARLCAAALAAILARVKSNQERSQPGIAVATGGHVYEQHPRFSTILQQTVRLLAPDSDITFV
PSEDGGGRGVAVVTAVASRLALHRRQLEETLAPFRLNSEQLKAVQAQMREAMVRGLRGEASSLRMLPTYV
CATPDGTERGDFLALDLGGTNFRVLWVKVGSRSEGGVKIINQVFSLPESVTQGTGQQLFDHIVDCIVDFQ
KEHDLSGQTLPLGFTFSFPCKQLGLDQGILLNWTKGFKASDCEGQDVVSLLRAAIRRRQGVELNVVAIVN
DTVGTMMSCGYEDPHCEVGLIVGTGTNACYMEELCNVSAVPGDQGRMCINMEWGAFGDDGLLKEFCTPFD
ISVDKSSINPGKQRFEKLISGMYLGEIVRQILLHLTRLGVLFGGQVFQCLQTKDIFKTKFLSEIESDNLE
RWQVRAILEKLGLPVSCDDALMVREVCQAVSQRAAKLCGAGVAAVVEKIRENRGLEELSVTVGVDGTLYK
LHPHFSDQVKDTVRHLAPRCSVRFLQSEDGSGKGAALVTAVACRLAESTQA
>anc192_altAll|avgPP:0.990|lnPP:-15.988|num_ambig:9|num_ambig_gaps:0
LKQFDVTEKQLQQIQAGILQDMEQALSRQTDPSPTVRMLPTYVRSTPHGTEQGDFLVLELGASGATLRVL
WVTLTGITGPRKEPISREFTIPTEIMLGTGQQLFDFAAGCLVEFLDQLKVGSQPIQLGFNFSFPCHQTGL
DQSTLISWTKGFRCSGVEGQDVVQLLREAIQRQGAYSIEVVAVVNDTVGTMMSCDPRPQACEIGLVVDTG
TNACYMEEARHVAELGEDQGRICVNVQWGSFGDSGALEPVQTPFDLQLDQESLNPGTQRFEKMVGGLYLG
ELVRLVLIHLAGQRALFGGVTSPALLTRGSVHIEHVMEIEDPSHGRARAQAVLQGLGLNPSTQDCEQVQQ
VCGAVCARAARLCAAALAAILARVKSNQERSQPGIAVATGGRVYEQHPRFSTILQQTVRLLAPDSDVTFV

```
PSEDGGGRGVAVVTAVASRLALHRRQLEETLAPFRLNSEQLKAVQAQMREAMVRGLRGEASSLRMLPTYV
CATPDGTERGDFLALDLGGTNFRVLWVKVGSRSEGGVKIINQVFSLPESVTQGTGQQLFDHIVDCIVDFQ
KQHGLSGQTLPLGFTFSFPCKQLGLDQGILLNWTKGFKASDCEGQDVVSLLRAAIRRRQGVELNVVAIVN
DTVGTMMSCGYEDPHCEVGLIVGTGTNACYMEELCNVSAVPGDQGRMCINMEWGAFGDDGLLKEFCTPFD
INVDKSSINPGKQRFEKLISGMYLGEIVRQILLHLTRLGVLFGGQVFQCLQTKDIFKTKFLSEIESDNLE
RWQVRAILEKLGLSVSCDDALMVREVCQAVSQRAAKLCGAGVAAVVEKIRENRGLEELSVTVGVDGTLYK
LHPHFSDQVKDTVRHLAPRCSVRFLQSEDGSGKGAALVTAVACRLAESTQA
>anc193|avgPP:0.992|lnPP:-11.428|num_ambig:13|num_ambig_gaps:0
LKQFDVTEEQLQQIQAGILQDMEQALSRQTDPSPTVRMLPTYVRSTPHGTEQGDFLVLELGASGATLRVL
WVTLTGITGPRKEPISREFTIPTEIMLGTGQQLFDFAAGCLVEFLDQLKVGSQPIQLGFNFSFPCHQTGL
DQSTLISWTKGFRCSGVEGQDVVQLLREAIQRQGAYSIDVVAVVNDTVGTMMSCDPRPQACEIGLVVDTG
TNACYMEEARHVAELGEDQGRVCVNVQWGSFGDSGALEPVQTPFDLQLDQESLNPGTQRFEKMVGGLYLG
ELVRLVLIHLAGQRALFGGVTSPALLTHGSVHIEHVMEIEDPSHGRARAQAVLQGLGLNPSTQDCELVQQ
VCGAVSARAARLCAAALAAILARVKSNQERSQPGIAVATGGHVYEQHPRFSTILQQTVRLLAPDSDITFV
PSEDGGGRGVAVVTAVASRLALHRRQLEETLAPFRLNSEQLKAVQAQMREAMVRGLRGEASSLPMLPTYV
CATPDGTERGDFLALDLGGTNFRVLWVKVGSRSEGGVKITNQVFSIPESVTQGTGQQLFDHIVDCIVDFQ
KEHGLSGQTLPLGFTFSFPCKQLGLDQGILLNWTKGFKASDCEGQDVVSLLRAAIRRRQGVELNVVAIVN
DTVGTMMSCGYEDPHCEVGLIVGTGTNACYMEELCNVSAVPGDQGRMCINMEWGAFGDDGLLKELCTPFD
ISVDKSSINPGKQRFEKLISGMYLGEIVRQILLHLTRLGVLFGGQVFQCLQTKDIFKTKFLSEIESDNLE
RRQVRAILEELGLPVSCDDALMVREVCQAVSQRAAKLCGAGVAAVVEKIRENRGLEELSVTVGVDGTLYK
LHPHFSDQVKDTVRHLAPRCSVRFLQSEDGSGKGAALVTAVACRLAESTQA
>anc193_altAll|avgPP:0.990|lnPP:-18.382|num_ambig:13|num_ambig_gaps:0
LKQFNVTEKQLQQIQAGILQDMEQALSRQTDPSPTVRMLPTYVRSTPHGTEQGDFLVLELGASGATLRVL
WVTLTGITGPRKEPISREFTIPTEIMLGTGQQLFDFAAGCLVEFLDQLKVGSQPIQLGFNFSFPCHQTGL
DQSTLISWTKGFRCSGVEGQDVVQLLREAIQRQGAYSIDVVAVVNDTVGTMMSCDPRPQACEIGLVVDTG
TNACYMEEARHVAELGEDQGRVCVNVQWGSFGDSGALEPVQTPFDLQLDQESLNPGTQRFEKMVGGLYLG
ELVRLVLIHLAGQRALFGGVTSPALLTRGSVHIEHVMEIEDPSHGRARAQAVLQGLGLNPSTQDCELVQQ
VCGAVCARAARLCAAALAAILARVKSNRERSQPGIAVATGGRVYEQHPRFSSILQQTVRLLAPDSDVTFV
PSEDGGGRGVAVVTAVASRLALHRRQLEETLAPFRLNSEQLKAVQAQMREAMVRGLRGEASSLRMLPTYV
CATPDGTERGDFLALDLGGTNFRVLWVKVGSRSEGGVKITNQVFSIPESVTQGTGQQLFDHIVDCIVDFQ
KEHGLSGQTLPLGFTFSFPCKQLGLDQGILLNWTKGFKASDCEGQDVVSLLRAAIRRRQGVELNVVAIVN
DTVGTMMSCGYEDPQCEVGLIVGTGTNACYMEELCNVSAVPGDQGRMCINMEWGAFGDDGLLKELCTPFD
INVDKSSINPGKQRFEKLISGMYLGEIVRQILLHLTRLGVLFGGQVFQCLQTKDIFKTKFLSEIESDDLE
RRQVRAILEELGLPVSCDDALMVREVCQAVSQRAAKLCGAGVAAVVEKIRENRGLEELSVAVGVDGTLYK
LHPHFSDQVKDTVRHLAPRCSVRFLQSEDGSGKGAALVTAVACRLAESTQA
>anc194|avgPP:0.989|lnPP:-16.791|num_ambig:16|num_ambig_gaps:0
LKQFDVTEEQLQQIQAGILQDMEQALSRQTDPSPTVRMLPTYVRSTPHGTEQGDFLVLELGASGATLRVL
WVTLTGTTGPRKEPISREFTIPPEIMLGTGQQLFDFAAGCLVEFLDQLKVGSQPIQLGFNFSFPCHQTGL
DRSTLISWTKGFRCSGVEGQDVVQLLREAIQRQGAYSIDVVAVVNDTVGTMMSCDPRPQACEIGLVVDTG
TNACYMEEARHVAELGEDQGRVCVNVQWGSFGDSGALEPVQTPFDLQLDQESLNPGTQRFEKMVGGLYLG
ELVRLVLIHLAGQRALFGGVTSPALLTHGSVHIEHVMEIEDPSHGRARAQAVLQGLGLNPSTQDCELVQQ
VCGAVSARAARLCAAALAAILARVKHNQERSQPGIAVATGGRVYEQHPRFSTILQQTVKLLAPDSDVTFV
PSEDGGGRGVAVVTAVASRLALHRRQLEETLAPFRLNSEQLKAVQAQMREAMVRGLRGEASSLPMLPTYV
CATPDGTERGDFLALDLGGTNFRVLLVKVGSRSEGGVKITNQVFSIPESVTQGTGQQLFDHIVDCIVDFQ
KEHGLSGQTLPLGFTFSFPCKQLGLDQGILLNWTKGFKASDCEGQDVVSLLRAAIRRRQGVELNVVAIVN
DTVGTMMSCGYEDPRCEVGLIVGTGTNACYMEELCNVSAVPGDQGRMCINMEWGAFGDDGLLKELCTPFD
ISVDKSSINPGKQRFEKLISGMYLGEIVRQILLHLTRLGVLFGGQVSQRLQTKDIFKTKFLSEIESDNLE
RRQVRAILEELGLPVSCDDALMVREVCQAVSQRAAKLCGAGVAAVVEKIRENRGLEELSVTVGVDGTLYK
LHPHFSDQVKDTVRHLAPRCSVTFLQSEDGSGKGAALVTAVACRLAESTQA
>anc194_altAll|avgPP:0.985|lnPP:-26.690|num_ambig:16|num_ambig_gaps:0
LKQFNVTEEQLQQIQAGILQDMEQALSRQTDPSPTVRMLPTYVRSTPHGTEQGDFLVLELGASGATLRVL
WVTLTGTSGPRKEPISREFTIPPEIMLGTGQQLFDFAAGCLVEFLDQLKVGSQPIQLGFNFSFPCHQTGL
```

```
DRSTLISWTKGFRCSGVEGQDVVQLLREAIQRQGAYSIDVVAVVNDTVGTMMSCDPRPQACEIGLVVDTG
TNACYMEEARHVAELGEDQGRVCVNVQWGSFGDSGALEPVQTPFDLQLDQESLNPGTQRFEKMVGGLYLG
ELVRLVLIHLAGQRALFGGATSPALLTHGSVPIEHVMEIEDPSQGRARAQAVLQGLGLNPSTQDCELVQQ
VCGAVCARAARLCAAALAAILARVKHNRERSQPGIAVATGGRVYEQHPRFSSILQQTVRLLAPDSDVTFV
PSEDGGGRGVAVVTAVASRLALHRRQLEETLAPFRLNSEQLKAVQAQMREAMVRGLRGEASSLRMLPTYV
CATPDGTERGDFLALDLGGTNFRVLLVKVGSRSEGGVKITNQVFSIPESVTQGTGQQLFDHIVDCIVDFQ
KEHGLSGQTLPLGFTFSFPCKQLGLDQGILLNWTKGFKASDCEGQDVVALLRAAIRRRQGVELNVVAIVN
DTVGTMMSCGYEDPRCEVGLIVGTGTNACYMEELCNVSAVPGDQGRMCINMEWGAFGDDGLLKELCTPFD
INVDKSSINPGKQRFEKLISGMYLGEIVRQILLHLTRLGVLFGGQVSQRLQTKDIFKTKFLSEIESDNLE
RRQVRAILEELGLPVNCDDALMVREVCQAVSQRAAKLCGAGVAAVVEKIRENRGLEELSVAVGVDGTLYK
LHPHFSEQVKDTVRHLAPRCSVTFLQSEDGSGKGAALVTAVACRLAESAQA
>anc195|avgPP:0.958|lnPP:-69.857|num_ambig:58|num_ambig_gaps:0
LQQFDVTEEQLQQIQASILEDMEQALRRQTNPTPTVRMLPTYVRSTPHGTEQGDFLVLELGATGATLRVL
WVTLTGTAGHRVEPKSQEFTIPPEIMLGTGQQLFDFAARCLAEFLDQLQVGNQPIQLGFSFSFPCHQTGL
DRSTLISWTKGFRCSGVEGQDVVQLLRDAIQRQGAYSIDVVAVVNDTVGTMMSCDPGPQPCEIGLVVDTG
TNACYMEEARHVAELEEDQGRVCVNVEWGSFGDDGALEPVQTPFDLQLDQESLNPGTQRFEKMIGGLYLG
ELVRLVLVHLAEQGALFGGATSPALLTHGSVLIEHVMEIEDPSAGRARAQAVLQDLGLSPSAQDCELVQQ
VCAAVSTRAARLCAAALAAILSRLQHNREQSQLQIAVATGGRVYEQHPRFSSILQQTVKLLAPECDVTFV
PSEDGGGRGVAMVTAVASRLAAHRRQLEETLAPFRLNSEQLKAVQAQMREAMARGLRGEASSLRMLPTYV
RATPDGTERGDFLALDLGGTNFRVLLVKVGSRSEGGVQITNQVYSIPESVTQGTGQQLFDHIVDCIVDFQ
KKQGLSGQTLPLGFTFSFPCKQLGLDQGILLNWTKGFNASDCEGQDVVSLLREAIRRRQGVELNVVAIVN
DTVGTMMSCGYEDPRCEVGLIVGTGTNACYMEELRNVAAVPGDQGRMCINMEWGAFGDDGSLDELSTPFD
ASVDKSSINPGKQRFEKLISGMYLGEIVRHILLHLTRLGVLFRGQQSQRLQTKDIFKTKFLSEIESDSLA
LRQVRAILEDLGLPVTCDDALMVREVCQAVSQRAAQLCGAGVAAVVEKIRENRGLEELSVTVGVDGTLYK
LHPHFSSQVQDTVRELAPRCSVTFLQSEDGSGKGAALVTAVACRLAETTQA
>anc195_altAll|avgPP:0.950|lnPP:-92.596|num_ambig:58|num_ambig_gaps:0
LQQFNVTEEQLQQIQASILEDMEQALRRQTDPSPTVRMLPTYVRSTPHGTEQGDFLVLELGATGATLRVL
WVTLTGTTGHRVEPRSQEFTIPPEVMLGPGQQLFDFAARCLSEFLDQLKVGNQPLQLGFSFSFPCHQTGL
DRSTLISWTKGFRCSGVEGQDVVQLLREAIQRQGAYNIDVVAVVNDTVGTMMSCDPRPQPCEIGLVVDTG
TNACYMEEARHVAVLDEDQGRVCVNVQWGSFGDDGALEPVQTPFDLQLDQESLNPGTQRFEKMVGGLYLG
ELVRLVLVHLAEQGALFGGATSPALLTQGSILIEHVVEMEDPSAGLARAQAILQDLGLNPSAQDCELVQQ
VCGAVCTRAARLCAAALAAILSRLQHNRERSQLQIAVATGGRVYEQHPRFSTILQETVKLLAPDCDVSFV
PSEDGGGRGVAVVTAVASRLAAHRRLLEETLAPFRLNREQLKAVQAQMREAMVRGLRGEASSLRMLPTYV
RATPDGSERGDFLALDLGGTNFRVLLVRVGARSEGGVQITSQVYSIPESVTQGTGQQLFDHIVDCIVDFQ
KKHGLSGQTLPLGFTFSFPCRQLGLDQGILLNWTKGFNASDCEGQDVVALLREAIRRRQGVELNVVAIVN
DTVGTMMSCGYEDPRCEVGLIVGTGTNACYMEELRNVSAVPGDEGRMCINMEWGAFGDDGSLDDLCTPFD
ASVDKASINPGKQRFEKLISGMYLGEIVRHILLHLTKLGVLFRGQQSQRLQTRDIFKTKFLSEIESDSLA
LRQVRAILEDLGLPLTCDDALMVREVCQAVSRRAAQLCGAGVAAVVEKIRENRGLEELSVTVGVDGTLYK
LHPHFSSQVKETVRELAPRCAVTFLQSEDGSGKGAALVTAVACRLAQSAQV
>anc196|avgPP:0.941|lnPP:-99.942|num_ambig:72|num_ambig_gaps:0
LQQFDLSEEQLQQIQASILEDMERALRRQTNPTSTVRMLPTYVRSTPHGTEKGDFLVLELGATGATLRVL
WVTLTGDAGHRVEPKSQVFSIPPDIMLGTGQQLFDFVAKCLSEFLDQLQVGKQPIQLGFSFSFPCHQTGL
DRSTLISWTKGFRCSGVEGKDVVQLLRDAIQRQGKYSIDVVAVVNDTVGTMMSCGSEEQPCEIGLVVDTG
TNACYMEEARHVAGLEEDQGRVCVNTEWGSFGDDGALKPVQTPFDLQLDQESLNPGKQRFEKMISGMYLG
ELVRLVLVHLAEQGALFGGSTSPALLTHGSILIQHVVEIEDPSAGLARAQAVLQELGLSPSEQDCELVQQ
VCSAVSTRAARLCAAALAAILSRLQHNRELSQLQIAVATGGAVYQQHPRFSSILQQTVKLLAPECDVTFV
PSEDGGGRGVAMVTAVASRLAAHRRQLEETLAPFRLNSEQLQAVQAQMREAMERGLRGEASSLRMLPTYV
RATPDGTERGDFLALDLGGTNFRVLLVQVGARAEGGVQITSQVYSIPEDVTQGTGEQLFDHIVDCIVDFQ
KKQGLSGQTLPLGFTFSFPCQQLGLDQGILLSWTKGFNASDCEGQDVVTLLREAIRRRQDVELNVVAIVN
DTVGTMMSCGYDDPRCEVGLIVGTGTNACYMEEMRNVATVPGDEGRMCINMEWGAFGDDGSLDDLITPFD
CSVDKSSINPGKQRFEKLISGMYLGEIVRHILLHLTKRGVLFRGRQSQRLQTKDIFKTKFLSEIESDSLA
LRQVRAILEDLGLQVTCDDALVVREVCQAVSRRAAQLCGAGVAAVVEKIRENRGLEQLSVTVGVDGTLYK
```

LHPHFSSQVQETVRELAPRCTVTFLQSEDGSGKGAALITAVACRLAETGPA
>anc196_altAll|avgPP:0.929|lnPP:-132.688|num_ambig:72|num_ambig_gaps:0
LQPFNLSEEQLQQIQASILEDMERALRRQTNPTSTVRMLPTYVRSTPHGTEQGDFLVLELGGTGATLRVL
WVTLTGDAGHRVEPKSQVFTIPSEVMLGSGQQLFDFVARCLANFLDQLQVGNQPLQLGFSFSFPCHQTGL
DRSTLISWTKGFKCSGVEGQDVVQLLRDAIKRQEKYSIDVVAVVNDTVGTMMSCGPEEQPCEIGLIVDTG
TNSCYMEEARHVAGLEEDQGRICVNTQWGSFGDDGALEDVQTPFDLQLDQESLNPGKQRFEKMVSGLYLG
ELVRLILVHLAEQGALFGGSTSPALLTHGSVLIQHVMEMEDPSAGLARAQAILKDLGLSPSEQDCELVQQ
VCSAVCTRAARLCAAALAAILSRLQHNRELSRLRIAVATGGAVYEQHPRFSRILQETVKQLAPDCDVSFV
PSEDGGGRGVAVVTAVASRLAAHRRLLEETLAPFRLNSEQLKAVQAQMREAMERGLRGEASSLRMLPTYV
RATPDGSERGDFLALDLGGTNFRVLLVQVGSRAEGGVQITNQVYSIPQDVTQGTGEQLFDHIVDCIVDFQ
KKHGLAGQVLPLGFTFSFPCKQLGLDQGILLNWTKGFNASDCEGQDVVSLLREAIRRRQDVELNVVAIVN
DTVGTMMSCGYDDPRCEVGLIVGTGTNACYMEEMKNVSTVPGDEGRMCINMEWGAFGDDGSLDELVTPFD
CSVDKSSINPGKQRFEKLISGMYLGEIVRHILLHLTRRGVLFRGQESQRLQTRDIFKTKFLSDIESDSLA
LRQVRAILEDLGLPLTCDDALVVREVCQAVSRRAAQLCGAGVAAVVEKIRENRGLDQLSVTVGVDGTLYK
LHPHFSSHVQETVRELAPRCSVTFLQSEDGSGKGAALITAVACRVAETGQA
>anc197|avgPP:0.930|lnPP:-116.197|num_ambig:81|num_ambig_gaps:0
LQPFYLSLEQLQQVKNSMLAAMERGLKQKVKATSSLPMLPTYVCTLPDGTERGDFLGLELSQTQVKVLLL
QLTDDVDEPVVVKRRTFPLPEKILIGSGQEVFEFMAQCLSTFLGGLNLSKKYFPLGFSFPFPCEHTSLNQ
CKLIRWTKDFRWPGVEGRDVTQLLQLAINQHCKNYQIEVLAVVNDTVGTLMSCSSNKQPCEIGLMIDSGT
NCCYMEETQRIAGVEESEGHMCVNTEWGSFGNNGELNDMLTEFDLLVDKESMDPGKCRFEKMISSMYLRE
TVRVILANLAEKAELFSGVLTPPLLTKGKLEIQDILEITDDKVGLAKAKSFLCRLGLVASNQDCFHVQQI
CQAVFTRSANLCAAGLAAILTHIRKTQELSELKTCVAVDGEMYRSHTQYSEILQQTLKELVPECTITFVP
SEDGSTLGAAMVTAVAVRLRNQRQGVSQALAPFRLSMVDLERLRNLMRQEMEKGLSKETNCKSSVRMLPT
YVCNLPDGTERGDFLALDLGGTNFRVLMVQVQSKEEGGVHMISETYTIPPEITQSTGEQLFDHIVNCIVD
FQTKHGMLGHMLPLGFTFSFPCNQIGLDKGILLRWTKGFSASGCVGQDVVNLLREAVQRQQHVDVDVVAL
VNDTVGTMMSCAYIDPKCEVGLIVGTGTNACYMEEMRNVGTVEGDEGRMCINMEWGAFGDDGCLDHYITP
FDTIVDKNSLNPGQQRFEKLISGMYLGEIVRHILLDLTSRKILFRGRESKGLQTRNIFPTKFLSDIENDS
QSRRHVCDILEDLGLSVTPEDAEVVKEVCHAVSSRAAKMCAAGVAAVVEKIRENKRMTHLEVTVGVDGTL
YKMHPYFSRKLQDTVPLLAPGCSVKFLLSEDGSGKGAALIAAVACRTKIS
>anc197_altAll|avgPP:0.918|lnPP:-150.268|num_ambig:81|num_ambig_gaps:0
LQPFYLSLEQLQQVKNSMLAAMEKGLKQKVKATSSLPMLPTYVCALPDGTEKGDFLGLELSQTQVKVLLL
QLTDDIDEPMVVKKKTFPLPEKILIGSGQQVFEFMAQCLATFLSGLNLNKKYFPLGFSFPFPCDHSALNQ
CKLIRWTKDFRWPGVEGKDVTQLLQLAINQQCKNYQIEVLAVVNDTVGILMSCSSTKQPCEIGLLIDAGT
NCCYMEETQRIGGVEETEGHMCVNTEWGSFGNDGELDDLLTEFDLLVDKESMDPGKCRFEKMISSMYFCE
TVRIILASLAEKAELFNGVLTPTLLTKGKLEIQDILEIVDEKVGLAKAKNFLFRLGLVASNQDCFHVQQI
CQAVYTRSANLCAAGLAAILTHIHRNQELPELKTFVAVDGDLYKSHSQYGEILQQTLKQLIPECSITFIP
SEDGSTLGAAIVTAVAVRLRNQRQGVSQALAPLKLSMVDLERLRNLMRQEMEKGLSKETNCKSSVRMLPT
YVCNLPDGTEKGDFLALDLGGTNFRVLVVQVKSKEEGGVHIISESYTIPPEIAQSTGEQLFDHIVSCIVD
FQTKHGMLGHILPLGFTFSFPCNQIGLDKGILLRWTKGFTASDCVGQDVVNLLREAVQRQQHIDVDVVAL
VNDTVGTMMSCAYLDPNCEVGLIVGTGTNACYMEEMRNVGTVEGDEGRMCINMEWGAFGDDGCLDHYITP
FDTIVDKNSLNPGQQRFEKLISGMYLGEIVRHILLELTSRKILFRGRKSKGLQTKDIFPTKFLSEIENDS
QGNRHVCDILEDLGLPVTPEDAEVVKEVCHTVSSRAAKVCAAGIAAVVEKIRENKRMTQLEVTVGVDGTL
YKMHPYFSKKLQDTVPLLAPNCNVKFLLSEDGSGKGAALIAAVACRTKIS
>anc198|avgPP:0.982|lnPP:-28.921|num_ambig:26|num_ambig_gaps:3
LLALTPSLEKLQAVKGRMLEAMEKGLSRQTHAQATVRMLPTFICSTPDGTEKGDFLVVELCQSHVRTLWV
TLLGDGNQSPQLMYKIFDMPEDIMQGTGEALFDFIAQCVRKFLEEIGSPQHRLPLGFVFPFSCRQTGLDK
AELISWSKGFRCSDVEGKDVVQLLQAAINKQELYHVDVVALLNDTVGTMMTCSTGGKPCEIALVVGTGTN
SCFMAEAHLVEMVEETSGRMCVNSEWGCFGDDGALGDVLTPYDHRVDQESSNPGERRFEKMVGSLYLGEI
VRHALLMLAAEQALFVGSNTSVLGTKDVFKTQQVLEIMDNEEGMAKARSTLEALGLQPSERDCCRVQQIC
RAVVSRAAALSATGLAAVLSHMCRSRELERLVVNVGVDGELYRAHARFGEILQSVAGLLAPECTATLVPS
ADTSGLGAAMVTAVALRLLAQRHEVEQLLAPLRLSRADLERVQALMRQEMERGLGRESNANASVRMLPTY
VCNTPDGTERGDFLALDLGGTNFRVLVVRVAQDGIHMASEIYVIPTAIMQGTGEALFDHIIECIMDFQMK

QDLMEQVLPLGFTFSFPCQQLGLDKAVLLTWTKGFSASGCVGQDVVQLLREAAQRKQHVALKVVAVVNDT
VGTMMACGYDDPKCEIGLIVGTGTNACYMEERQNVGTVEGDEGRMCINMEWGAFGDNGCLDDIFTTFDRL
VDEKTINAGKQRFEKLISGMYLGEIVRHILLALVEKQLLFRGKPSPKLQTRDIFQTKFLSTIETDGLALR
QVQGILRDLELDASFEDSVLVREVCQTVSLRAAQLCAAGLAAVVEKMRENRGLAQLAVTVGVDGTLYKMH
PRFSQNLQQMLQDLAPNCTVTFLQSEDGSGKGAALVAAVACRGAESGPP
>anc198_altAll|avgPP:0.978|lnPP:-38.589|num_ambig:26|num_ambig_gaps:3
LLALTPSLEKLQAVKGRMLEAMDKGLSRQTHAQATVRMLPTFICSTPDGTEKGDFLVVELCQSHVRTLWV
TLLGDGNQSPQLMYKIFDMPEDIMQGTGEALFDFIAQCVHQFLEEIGSPQHRLPLGFVFPFSCRQTGLDK
AELISWSKGFRCSDVEGKDVVQLLQAAINKQELYHVDVVALLNDTVGTMMTCSTGGKPCEIALVVGTGTN
SCFMAEAHLVEMVEETSGRMCVNSEWGCFGDDGALSDVLTPYDHRVDQESSNPGERRFEKMVGGLYLGEI
VRHAMLMLAAEQALFIGASTSVLGTKDVLKTQQVLEIMDNEDGMAKARSALEALGLQPSERDCCRVQQIC
RAVVSRAAALSATGLAAVLSHMCRSRELERLVVNVGVDGELYRAHARFGEILQSVTGLLAPECTATLVPS
ADTSGLGAAMVTAVALRLMAQRHEVEQLLAPLRLSRADLQRVQALMRQEMERGLGRETNANASVRMLPTY
VCNTPDGTERGDFLALDLGGTNFRVLVVRVAEDGIHMASEIYVIPTAIMQGTGEALFDHIIECIMDFQMK
QDLMEQVLPLGFTFSFPCQQLGLDKAVLLSWTKGFSASGCVGQDVVQLLREAAQRKQHVALKVVAVVNDT
VGTMMACGYDDPKCEIGLIVGTGTNACYMEERKNVGTVEGDEGRMCINMEWGAFGDNGCLDDIFTSFDRM
VDEKTINAGKQRFEKLISGMYLGEIVRHILLALVEKQVLFRGKPSPKLQTRDIFQTKFLSTIETDGLALQ
QVQGILRDLELDASFEDSVLVREVCQTVSLRAAQLCAAGLAAVVEKMRENRGLAQLAVTVGVDGTLYKMH
PRFSQNLQQMLQDLAPNCAVTFLQSEDGSGKGAALVAAVACRGADQGPP
>anc199|avgPP:0.988|lnPP:-18.136|num_ambig:18|num_ambig_gaps:1
LLALTIPLEMLQVVKGRMMQAMSKGLSRQTHAQANVQMLPTYICSTPNGTEKGNVLVVELCQSHVRILLM
TLLGDGNHSPHVTDKIFDMPGDSMQGTGEALFDFIAQCVRQFLSGISSAQERLPLGFVFPFSCRQTQLDK
AELISWSKGFSCSDVEGKDVVQLLQSAINKQELSHVEVVALMNDTVGTMMTCSTREKPCEVALVVDKGTN
SCFMAEAQYVEMVEETSGRMCVNTEWGSFGDDDALSDILTPYDQRVDQESSNPGKKRFEKLVGSLYLGDI
VRHTLMALAAEKALFIGSNVTVLGTKGALKTQQVLEIIDNEEGMAKARAALEALGLRPSERDCCRVQQIC
RAVVSRSAALCAAGLAAILSHIRQSRGLERLVINVAVDGELYRAHSRFGEILQSVTGLLAPECMASLVPS
VDGTGRGAAIVTAVALRLADQRREVDQLLAPLRLSRADLERVQALMRREMELGLGQETNANSSVRMLPTY
VRGTPDGTERGEFLALDLGGTNFRVLVVRVAEDGIRMASEIYLIPTNIMQGTGEALFDHIIDCIMDFQLK
QDLMEQVLPLGFTFSFPCQQLGLDKAVLLTWTKGFSASGCVGEDVVQLLREAAQRKQHLALKVVAVVNDT
VGTMMSCGYDDPKCEIGLIVGTGTNACYMEEMQNVGTVEGEQGRMCINMEWGAFGDNGCLDHIFTDFDRL
VDEKTINPGKQRFEKLISGMYLGEIVRHILLALVEKQVLFRGKPCPTLQTRDIFKTKFLSTIEIDGLALQ
QVQAILQDLELDASFEDSVLVREVCQTVSLRAAQLCAAGLAAVVEKMRENRGLSQLAVTVGVDGTLYKMH
PCFSQHLQQMLQDLAPNCTVTFLQSEDGSGKGAALVAAVACRGAEP
>anc199_altAll|avgPP:0.984|lnPP:-28.982|num_ambig:18|num_ambig_gaps:1
LLALTIPLEMLQVVKGRMMQAMNKGLSRQTHAQANVQMLPTYICSTPNGTEKGDVLVVELCQSHVRILLM
TLLGDGNQSPQVTDKIFDMPGDIMQGTGEALFDFIAQCVRQFLAGITSSQERLPLGFVFPFSCKQTQLDK
AELISWSKGFSCSDVEGKDVVQLLQSAINKQELSHVEVVALMNDTVGTMMTCSTRERPCEVALVVDKGTN
SCFMAEAQYVEMVEETSGRMCVNTEWGSFGDDDALNDILTPYDQHVDQESSNPGKKRFEKLVGSLYLGDI
VRHTLMALAAEKALFIGSNVTVLGTKGALKTQQVLEIIDNEEGMAKARAALEALGLRPSERDCCRVQQIC
RAVVSRSAALCAAGLAAILSHIRHSRGLERLVINVAVDGELYRAHSRFGEILQSVTGLLAPECMATLVPS
VDGTGRGAAIVTAVALRLANQRREVDQLLAPLRLSRADLERVQALMRREMELGLGQETNANSSVRMLPTY
VRGTPDGTERGEFLALDLGGTNFRVLVVRVAEDGIRMASEIYLIPTNIMQGTGEALFDHIIDCIMDFQLK
QDLMEQVLPLGFTFSFPCQQLGLDKAVLLTWTKGFSASGCVGEDVVQLLREAAQRKQHLALKVVAVVNDT
VGTMMSCGYDDPKCEIGLIVGTGTNACYMEEMQNVGTVEGEQGRMCINMEWGAFGDNGCLDHIFTDFDRM
VDEKTINPGKQRFEKLISGMYLGEIVRHILLALVEKQVLFRGKPCPTLQTRDIFKTKFLSTIEIDGLALR
QVQAILQDLELDASFEDSVLVREVCQTVSLRAAQLCAAGLAAVVEKMRENRGLAQLAVTVGVDGTLYKMH
PCFSQHLQQMLQDLAPNCTVTFLQSEDGSGKGAALVAAVACRGAEP
>anc200|avgPP:0.988|lnPP:-19.376|num_ambig:15|num_ambig_gaps:0
LLALTIPLEMLQMVKGRMMQAMNKGLSRQTHAQANVQMLPTYICSTPNGTENGNVLVVELCQSHVRILLM
TLLGDGNHSPQLTDKIFDMPGDITQGTGEALFDFIAQCLRQFLGGISSSKERLPLGFVFPFSCRQTRLDK
AELISWSKGFSCSDVEGKDVVQLLQSAINKQELSQVEVVALMNDTVGTMMTCSTREKPCEVALIVDKGTN
SCFMAEAQYVQMVEETSGRMCVNTEWGCFGDDGALNDILTPYDQHVDQESSNPGVKRFEKLVGSLYLGDI

VRHALIALAAEKALFIGSNVTVLGTKGVLKTQQVLEIIDNEEGMTKARMTLEALGLRPSERDCCRVQQVC
RAVVSRSAALCAAGLAAILSLIRQSRGLERLMINVAVDGELYRAHSRFREILQSVTGLLAPECMASLVPS
VDGTGRGAAMVTAVALRLADQRREVDQLLAPLRLSRADLERVQALMRQEMDLGLGQETNANSSVRMLPTY
VRGTPDGTERGEFLALDLGGTNFRVLVVRVAENGIRMASEIYVIPTNIMQGTGEALFDHIIDCIMDFQLK
QDLMEQVLPLGFTFSFPCKQLGLDKAVLLSWTKGFSASECVGKDVVQLLREAAQRKQHLALKVVAVVNDT
VGTMMSCGYDDPKCEIGLIVGTGTNACYMEEMQNVGTVEGEQGRMCINMEWGAFGDNGCLDHIFTEFDRM
VDEKTINPGKQRFEKLISGMYLGEIVRHILLALVEKQVLFHDKPCPTLQSRDIFKTKFLSTIEIDGLALK
QVQAILRDLELQASFEDSVLVREVCQTVSLRAAQLCAAGLAAVVEKMRENRGLSQLAVTVGVDGTLYKMH
PCFSQHLQQTLQDLAPNCTVTFLQSEDGSGKGAALVAAVAVRGAE
>anc200_altAll|avgPP:0.986|lnPP:-25.613|num_ambig:15|num_ambig_gaps:0
LLALTIPLEMLQMVKGRMMQAMNKGLSRQTHAQANVQMLPTYICSTPNGTENGDVLVVELCQSHVRILLM
TLLGDGNNSPQLTDKIFDMPGDITQGTGEALFDFIAQCLRQFLAGMSSSQERLPLGFVFPFSCKQTRLDK
AELISWSKGFSCSDVEGKDVVQLLQSAINKQELSQVEVVALMNDTVGTMMTCSTREKPCEVALIVDKGTN
SCFMAEAQYVQIVEETSGRMCVNTEWGCFGDDGALNDILTPYDQHVDQESSNPGLKRFEKLVGSLYLGDI
VRHALIALAAEKALFIGSNVTVLGTKGVLKTQQVLEIIDNEEGMTKARMTLEALGLRPSERDCCRVQQVC
RAVVSRSAALCAAGLAAILSLIRQSRGLERLMINVAVDGELYRAHARFREILQSVTGLLAPECMASLVPS
VDGTGRGAAMVTAVALRLADQRREVDQLLAPLRLSRADLERVQALMRQEMDLGLGQETNANASVRMLPTY
VRGTPDGTERGEFLALDLGGTNFRVLVVRVAENGIRMASEIYVIPTNIMQGTGEALFDHIIDCIMDFQLK
QDLMEQVLPLGFTFSFPCKQLGLDKAVLLSWTKGFSASDCVGKDVVQLLREAAQRKQHLALKVVAVVNDT
VGTMMSCGYDDPKCEIGLIVGTGTNACYMEEMQNVGTVEGEQGRMCINMEWGAFGDNGCLDHIFTEFDRM
VDEKTINPGKQRFEKLISGMYLGEIVRHILLALVEKQVLFHDKPCPTLQTRDIFKTKFLSTIEIDGLALK
QVQAILRDLDLHASFEDSVLVREVCQTVSLRAAQLCAAGLAAVVEKMRENRGLSQLAVTVGVDGTLYKMH
PCFSQHLQQTLQDLAPNCTVTFLQSEDGSGKGAALVAAVAVREAE
>anc201|avgPP:0.988|lnPP:-18.905|num_ambig:22|num_ambig_gaps:1
LLALTIPLEMLQVVKGRMMQAMSKGLSRQTHAQANVQMLPTYICSTPNGTEKGNVLVVELCQSHVRILLM
TLLGDGNHSPQVTDKIFDMPGDIMQGTGEALFDFIAQCVRQFLAGISSSQERLPLGFVFPFSCRQTRLDK
AELISWSKGFSCSDVEGKDVVQLLQSAINKQELSHVEVVALMNDTVGTMMTCSTREKPCEVALVVDKGTN
SCFMAEAQYVEMVEETSGRMCVNTEWGCFGDDGALSDILTPYDQRVDQESSNPGKKRFEKLVGSLYLGDI
VRHALIALAAEKALFIGSNVTVLGTKGVLKTQQVLEIIDNEEGMAKARAALEALGLRPSERDCCRVQQIC
RAVVSRSAALCAAGLAAILSHIRQSRGLERLVINVAVDGELYRAHSRFGEILQSVTGLLAPECMASLVPS
VDGTGRGAAMVTAVALRLADQRREVDQLLAPLRLSRADLERVQALMRREMELGLGQETNANSSVRMLPTY
VRGTPDGTERGEFLALDLGGTNFRVLVVRVAEDGIRMASEIYVIPTNIMQGTGEALFDHIIDCIMDFQLK
QDLMEQVLPLGFTFSFPCQQLGLDKAVLLSWTKGFSASGCVGEDVVQLLREAAQRKQHLALKVVAVVNDT
VGTMMSCGYDDPKCEIGLIVGTGTNACYMEEMQNVGTVEGEQGRMCINMEWGAFGDNGCLDHIFTDFDRL
VDEKTINPGKQRFEKLISGMYLGEIVRHILLALVEKQVLFRGKPCPTLQTRDIFKTKFLSTIEIDGLALR
QVQAILQDLELDASFEDSVLVREVCQTVSLRAAQLCAAGLAAVVEKMRENRGLSQLAVTVGVDGTLYKMH
PCFSQHLQQMLQDLAPNCTVTFLQSEDGSGKGAALVAAVACRGAEP
>anc201_altAll|avgPP:0.983|lnPP:-33.073|num_ambig:22|num_ambig_gaps:1
LLALTIPLEMLQAVKGRMMQAMNKGLSRQTHAQANVQMLPTYICSTPNGTEKGDVLVVELCQSHVRILLM
TLLGDGNQSPQVTDKIFDMPGDIMQGTGEALFDFIAQCVRQFLSGISSAQERLPLGFVFPFSCKQTRLDK
AELISWSKGFSCSDVEGKDVVQLLQSAINKQELSHVEVVALMNDTVGTMMTCSTREKPCEVALVVDKGTN
SCFMAEAQYVEMVEETSGRMCVNTEWGCFGDDGALNDILTPYDQHVDQESSNPGEKRFEKLVGSLYLGDI
VRHALMALAAEKALFIGSNVTVLGTKGVLKTQQVLEIIDNEEGMAKARATLEALGLRPSERDCCRVQQIC
RAVVSRSAALCAAGLAAILSHIRQSRGLERLVINVAVDGELYRAHSRFGEILQSVTGLLAPECMATLVPS
VDGTGRGAAMVTAVALRLADQRREVDQLLAPLRLSRADLERVQALMRREMELGLGQETNANASVRMLPTY
VRGTPDGTERGEFLALDLGGTNFRVLVVRVAEDGIRMASEIYVIPTNIMQGTGEALFDHIIDCIMDFQLK
QDLMEQVLPLGFTFSFPCQQLGLDKAVLLTWTKGFSASGCVGQDVVQLLREAAQRKQHLALKVVAVVNDT
VGTMMSCGYDDPKCEIGLIVGTGTNACYMEEMQNVGTVEGEQGRMCINMEWGAFGDNGCLDHIFTDFDRM
VDEKTINPGKQRFEKLISGMYLGEIVRHILLALVEKQVLFRGKPCPKLQTRDIFKTKFLSTIEIDGLALQ
QVQAILQDLELQASFEDSVLVREVCQTVSLRAAQLCAAGLAAVVEKMRENRGLAQLAVTVGVDGTLYKMH
PCFSQHLQQTLQDLAPNCTVTFLQSEDGSGKGAALVAAVACRGAEP
>anc202|avgPP:0.994|lnPP:-8.822|num_ambig:11|num_ambig_gaps:1

LLALTIPLEMLQAVKGRMLQAMRKGLSRQTHAQANVRMLPTYICSTPDGTEKGDLLVVELCQSHVRTLWM
TLLGDGNQSPQVTNKIFDMPGDIMQGKGEALFDFIAQCVRQFLAGISSPQHRLPLGFVFPFSCRQTRLDK
AELISWSKGFSCSDVEGKDVVQLLQSAINKQELSHVDVVALMNDTVGTMMTCSTGEKPCEVALVVDKGTN
SCFMAEAQQVEMVEETSGRMCVNTEWGCFGDDGTLSDILTPYDQRVDQESSNPGEKRFEKLVGSLYLGEI
VRHALIALAAEKALFIGSNVAVLGTKGVLKTQQVLEIIDNEEGMAKARKALEALGLRPSERDCCRVQQIC
RAVVSRAAALCAAGLAAILSHMCQSRELERLVVNVGVDGELYRGHSRFGEILQSVTGLLAPECMATLVPS
VDGTGRGAAMVTAVALRLAAQRREVDQLLAPLRLSRADLERVQALMRREMELGLGQETNANASVRMLPTY
VRGTPDGTERGEFLALDLGGTNFRVLVVRVAQDGIRMASEIYVIPTAIMQGTGEALFDHIIECIMDFQLK
QDLMEQVLPLGFTFSFPCQQLGLDKAVLLSWTKGFSASGCVGQDVVQLLREAAQRKQHLALKVVAVVNDT
VGTMMSCGYDDPKCEIGLIVGTGTNACYMEEMQNVGTVEGEQGRMCINMEWGAFGDNGCLDDIFTDFDRL
VDEKTINPGKQRFEKLISGMYLGEIVRHILLALVEKQLLFRGKPCPKLQTRDIFKTKFLSTIEIDGLALR
QVRAILQDLELDASFEDSVLVREVCQTVSLRAAQLCAAGLAAVVEKMRENRGLAQLAVTVGVDGTLYKMH
PRFSQHLQQMLQDLAPNCTVTFLQSEDGSGKGAALVAAVACRGAEP
>anc202_altAll|avgPP:0.991|lnPP:-15.722|num_ambig:11|num_ambig_gaps:1
LLALTIPLEMLQAVKGRMLQAMRKGLSRQTHAQANVRMLPTYICSTPDGTEKGDFLVVELCQSHVRTLWM
TLLGDGNQSPQVTDKIFDMPGDIMQGKGEALFDFIAQCVRQFLAGISSPQHRLPLGFVFPFSCRQTRLDK
AELISWSKGFSCSDVEGKDVVQLLQSAINKQELSHVDVVALMNDTVGTMMTCSTGEKPCEVALVVDKGTN
SCFMAEAQQVEMVEETSGRMCVNTEWGCFGDDGALSDILTPYDQRVDQESSNPGEKRFEKLVGSLYLGEI
VRHALTALAAEKALFIGSNVAVLGTKGVLKTQQVLEIIDNEEGMAKARKTLEALGLRPSERDCCRVQQIC
RAVVSRAAALCAAGLAAILSHMCQSRELERLVVNVGVDGELYRGHSRFGEILQSVTGLLAPECMATLVPS
VDGTGRGAAMVTAVALRLAAQRREVDQLLAPLRLSRADLERVQALMRREMELGLGQETNANASVRMLPTY
VRGTPDGTERGEFLALDLGGTNFRVLVVRVAEDGIRMASEIYVIPTAIMQGTGEALFDHIIDCIMDFQLK
QDLMEQVLPLGFTFSFPCQQLGLDKAVLLSWTKGFSASGCVGQDVVQLLREAAQRKQHLALKVVAVVNDT
VGTMMSCGYDDPKCEIGLIVGTGTNACYMEEMQNVGTVEGEQGRMCINMEWGAFGDNGCLDDIFTDFDRM
VDEKTINPGKQRFEKLISGMYLGEIVRHILLALVEKQVLFRGKPCPKLQTRDIFKTKFLSTIEIDGLALR
QVRAILQDLELQASFEDSVLVREVCQTVSLRAAQLCAAGLAAVVEKMRENRGLAQLAVTVGVDGTLYKMH
PRFSQHLQQTLQDLAPNCTVTFLQSEDGSGKGAALVAAVACRGAEP
>anc203|avgPP:0.990|lnPP:-15.249|num_ambig:15|num_ambig_gaps:7
LRSLSIPLERLHIMKGHMIQDMCKGLSRQTHAQAKVRMLPTYICSTPNGTEKGNFLVVELCQNQVRTLLV
TLYGDGNMSPQMMYKIFDMPEGIMQGEGEALFDFIAQCVSQFLAETISPDTSTSEEHLPLGFVFPFTCRQ
TQLDKAELLSWSKGFSCSGVVGKDVVQLLQSAIDKQEAGGGGGGGNWLSSWKGRKPSQSAPGQLCHVDVV
ALMNDTVGTMMTCSMEGRPCEVAIVADKGSNCCFMAEAYLVETTEETSGRMCVNTEWGCFGDDGTLNDIF
TPYDECVDVESSNPGEKRFEKMVGTLYLGEIVRHALVALTAEKAVFTGTNAAVLKEKGVFTMQHVLDIIN
NEEGITEVRRILEVLGLQPSERDCGRVQQICRAVVGRAATLHATGLAAILSYMCQTRDLETLMVNVGVEG
ELYTGYPRFEEILLSVSRLLSPECMATLLPSRDGSGRGAAMVTAVALRLAAQRREVNEVLAPLRLSRADL
EKVQALMRQEMDQGLGKETNPTASVRMLPTYVSHTPDGTERGDFLALDLGGTNFRVLVVRVTEEGISMAS
EIYVIPAAIMQGTGEGLFDHIIDCIIDFQTKQNLVTQMLPLGFTFSFPCKQVGLDKALLLTWTKGFSASG
CVGQDVVQLLRDAAQRKQHLGMQVVALVNDTVGTMMACGYDDPKCEIGLIVGTGTNACYMEEMKNVGTVE
GDQGRMCINMEWGAFGDNGCLDHIFTHFDKVVDETTINPGKQRFEKLISGMYLGEIVRQILLVMTEKQLL
FQGKPCPKLQTKDIFKTKFLSTIELSNGLALRQIRAILNELEIDASFEDSVLMREVCQTVSLRAAQLCAA
GLAAVVEKMRENRGVDQLSVTVGVDGTLYKLHPCFSSNLQKTLKDLAPNCNVTFLLSEDGSGKGAALVAA
VANRAANAME
>anc203_altAll|avgPP:0.987|lnPP:-23.491|num_ambig:15|num_ambig_gaps:7
LRSLSIPLERLHIMKGHMIQDMCKGLSRQTHAQAKVRMLPTYICSTPNGTEKGNFLVVELCQNQVRTLLV
TLYGDGNMSPQMMYKIFDLPEGIMQGEGEALFDFIAQCVSQFLAETISPDTNSSEEHLPLGFVFPFSCRQ
TQLDKAELLSWSKGFSCSGVVGKDVVQLLQSAIDKQEDGEGNGSGRWLSSWKGRKPSQSAPGQLCHVDVV
ALMNDTVGTMMTCSMEGRPCEVAIVADKGSNCCFMAEAYLVETTEETSGRMCVNTEWGCFGDDGTLNDIF
TPYDECVDEESSNPGEKRFEKMVGTLYLGEIVRHALVALTAEKAVFTGTNAAVLKEKGVFTMQHVLDIIN
NEEGITEVKRILEVLGLQPSERDCGRVQQICRAVVGRAATLHATGLAAILSYMCQTRDLETLMVNVGVEG
ELYTGYPRFEEILLSVSRLLSPECMATLLPSRDGSGRGAAMVTAVALRLAAQRREVNEVLAPLRLSRADL
EKVQALMRQEMEQGLGKETNPTASVRMLPTYVSHTPDGTERGDFLALDLGGTNFRVLVVRVTEEGISMAS
EIYVIPAAIMQGTGEALFDHIIDCIIDFQTKQNLVTQMLPLGFTFSFPCKQVGLDKALLLTWTKGFSASG

CVGQDVVQLLRDAAQRKQHLGMQVVALVNDTVGTMMACGYDDPKCEIGLIVGTGTNACYMEEMKNVGTVE
GDQGRMCINMEWGAFGDNGCLDHIFTHFDKVVDESTINPGKQRFEKLISGMYLGEIVRQILLVMTEKQLL
FQGKPSPKLQTKDIFKTKFLSTIELSNGLALRQIRAILNELEIDASFEDSVLMREVCQTVSLRAAQLCAA
GLAAVVEKMRENRGVDQLSVTVGVDGTLYKLHPCFSSNLQKTLKDLAPNCNVTFLLSEDGSGKGAALVAA
VANRAANAME
>anc204|avgPP:0.994|lnPP:-8.079|num_ambig:4|num_ambig_gaps:0
LRSLSIPLERLHIMKGHMIQDMCKGLSRQTHAQAKVRMLPTYICSTPNGTEKGNFLVVELCQSQVRTLLV
TLYGDGNMSPHMVYKIFDLPEGITQGEGEALFDFIAQCVSQFLAETISSDTSKSEERLPLGFVFPFTCRQ
TQLDKAELLSWSKGFSCSGVVGKDVVQLLQSAINKQELCHVDVVALMNDTVGTMMTCSTEGRPCEVAIVA
DKGSNCCFMAEAYLVETTDETSGRMCVNTEWGCFGDDGTLNDIFTPYDECVDVESSNPGEKRFEKMVGTL
YLGEIVRHALIALTAEKAIFTGTNIAVLKEKGVFTMQHILDIINNEEGITEVKRILEVLGLQPSERDCGR
VQQICRAVVGRAATLHATGLAAILSYMCQTRDMETLMVNVGVEGELYTGYPRFEEILLSVSRLLSPECMA
TLLPSRDGSGRGAAMVTAVALRLAALRREVNEVLAPLRLSRADLEKVQALMRQEMDQGLGKETNATASVR
MLPTYVSHTPDGTERGDFLALDLGGTNFRVLVVHVTEEGISMASEIYVIPAAIMQGTGEALFDHIIDCII
DFQTKQNLVTDMLPLGFTFSFPCKQMGLDKALLLTWTKGFSASGCVGQDVVQLLREAARRKQHLGMQVVA
LVNDTVGTMMACGYDDPKCEIGLIVGTGTNACYMEEMKNVGTVEGDQGRMCINMEWGAFGDNGCLDHIFT
HFDKVVDETTINPGKQRFEKLISGMYLGEIVRQILLVMTEKQLLFQGKLSPKLQTKDIFKTKFLSTIEVS
NGLALRQIRAILNELEIDASFEDSVLLREVCQTVSLRAAQLCAAGLAAVVEKMRENRGVDQLSVTVGVDG
TLYKLHPCFSDNLQKTLKDLAPNCNVSFLLSEDGSGKGAALVAAVASRAANAME
>anc204_altAll|avgPP:0.994|lnPP:-10.312|num_ambig:4|num_ambig_gaps:0
LRSLSIPLERLHIMKGHMIQDMCKGLSRQTHAQAKVRMLPTYICSTPNGTEKGNFLVVELCQSQVRTLLV
TLYGDGNMSPHMVYKIFDLPEGITQGEGEALFDFIAQCVSQFLAETISSDTSKSEERLPLGFVFPFTCRQ
TQLDKAELLSWSKGFSCSGVVGKDVVQLLQSAINKQELCHVDVVALMNDTVGTMMTCSTEGRPCEVAIVA
DKGSNCCFMAEAYLVETTDETSGRMCVNTEWGCFGDDGTLNDIFTPYDECVDVESSNPGEKRFEKMVGTL
YLGEIVRHALIALTAEKAIFTGTNIAVLKEKGVFTMQHILDIINNEEGITEVKRILEVLGLQPSERDCGR
VQQICRAVVGRAATLHATGLAAILSYMCQTRDLETLMVNVGVEGELYTGYPRFEEILLSVSRLLSPECMA
TLLPSRDGSGRGAAMVTAVALRLAALRREVNEVLAPLRLSRADLEKVQALMRQEMDQGLGKETNATASVR
MLPTYVSHTPDGTERGDFLALDLGGTNFRVLVVHVTEEGISMASEIYVIPAAIMQGTGEALFDHIIDCII
DFQTKQNLVTDMLPLGFTFSFPCKQVGLDKALLLTWTKGFSASGCVGQDVVQLLREAARRKQHLGMQVVA
LVNDTVGTMMACGYDDPKCEIGLIVGTGTNACYMEEMKNVGTVEGDQGRMCINMEWGAFGDNGCLDHIFT
HFDKVVDESTINPGKQRFEKLISGMYLGEIVRQILLVMTEKQLLFQGKLSPKLQTKDIFQTKFLSTIEVS
NGLALRQIRAILNELEIDASFEDSVLLREVCQTVSLRAAQLCAAGLAAVVEKMRENRGVDQLSVTVGVDG
TLYKLHPCFSDNLQKTLKDLAPNCNVSFLLSEDGSGKGAALVAAVASRAANAME
>anc205|avgPP:0.989|lnPP:-16.957|num_ambig:18|num_ambig_gaps:7
LRSLSIPLERLHIMKGHMIQDMCKGLSRQTHAQAKVRMLPTYICSTPNGTEKGNFLVVELCQNQVRTLLV
TLYGDGNMSPQMMYKIFDMPEGIMQGEGEALFDFIAQCVSQFLAETISPDTSTSEERLPLGFVFPFTCRQ
TQLDKAELLSWSKGFSCSGVVGKDVVQLLQSAINKQEAGGGGGGGNWLSSWKGRKPSQSAPGQLCHVDVV
ALMNDTVGTMMTCSMEGRPCEVAIVADKGSNCCFMAEAYLVETTEETSGRMCVNTEWGCFGDDGTLNDIF
TPYDECVDVESSNPGEKRFEKMVGTLYLGEIVRHALIALTAEKAIFTGTNVAVLKEKGVFTMQHVLDIIN
NEEGITEVKRILEVLGLQPSERDCGRVQQICRAVVGRAATLHATGLAAILSYMCQTRDMETLMVNVGVEG
ELYTGYPRFEEILLSVSRLLSPECMATLLPSRDGSGRGAAMVTAVALRLAAQRREVNEVLAPLRLSRADL
EKVQALMRQEMDQGLGKETNATASVRMLPTYVSHTPDGTERGDFLALDLGGTNFRVLVVRVTEEGISMAS
EIYVIPAAIMQGTGEALFDHIIDCIIDFQTKQNLVTDMLPLGFTFSFPCKQVGLDKALLLTWTKGFSASG
CVGQDVVQLLREAAQRKQHLGMQVVALVNDTVGTMMACGYDDPKCEIGLIVGTGTNACYMEEMKNVGTVE
GDQGRMCINMEWGAFGDNGCLDHIFTHFDKVVDETTINPGKQRFEKLISGMYLGEIVRQILLVMTEKQLL
FQGKPCPKLQTKDIFKTKFLSTIELSNGLALRQIRAILNELEIDASFEDSVLMREVCQTVSLRAAQLCAA
GLAAVVEKMRENRGVDQLSVTVGVDGTLYKLHPCFSDNLQKTLKDLAPNCNVTFLLSEDGSGKGAALVAA
VANRAANAME
>anc205_altAll|avgPP:0.985|lnPP:-26.679|num_ambig:18|num_ambig_gaps:7
LRSLSIPLERLHIMKGHMIQDMCKGLSRQTHAQAKVRMLPTYICSTPNGTEKGNFLVVELCQSQVRTLLV
TLYGDGNMSPHMMYKIFDLPEGIMQGEGEALFDFIAQCVSQFLAETISPDTSSSEERLPLGFVFPFSCRQ
TQLDKAELLSWSKGFSCSGVVGKDVVQLLQSAINKQEAGEGGGGGRWLSSWRGRKPSQSAPGQLCHVDVV

```
ALMNDTVGTMMTCSTEGRPCEVAIVADKGSNCCFMAEAYLVETTDETSGRMCVNTEWGCFGDDGTLNDIF
TPYDECVDEESSNPGEKRFEKMVGTLYLGEIVRHALIALTAEKAIFTGTNVAVLKEKGVFTMQHVLDIIN
NEEGITEVKRILEVLGLQPSERDCGRVQQICRAVVGRAATLHATGLAAILSYMCQTRDLETLMVNVGVEG
ELYTGYPRFEEILLSVSRLLSPECMATLLPSRDGSGRGAAMVTAVALRLAAQRREVNEVLAPLRLSRADL
EKVQALMRQEMEQGLGKETNATASVRMLPTYVSHTPDGTERGDFLALDLGGTNFRVLVVRVTEEGISMAS
EIYVIPAAIMQGTGEGLFDHIIDCIIDFQTKQNLVTDMLPLGFTFSFPCKQVGLDKALLLTWTKGFSASG
CVGQDVVQLLREAAQRKQHLGMQVVALVNDTVGTMMACGYDDPKCEIGLIVGTGTNACYMEEMKNVGTVE
GDQGRMCINMEWGAFGDNGCLDHIFTHFDKVVDESTINPGKQRFEKLISGMYLGEIVRQILLVMTEKQLL
FQGKPSPKLQTKDIFQTKFLSTIELSNGLALRQIRAILNELEIDASFEDSVLMREVCQTVSLRAAQLCAA
GLAAVVEKMRENRGVDQLSVTVGVDGTLYKLHPCFSDNLQKTLKDLAPNCNVTFLLSEDGSGKGAALVAA
VASRAANAME
>anc206|avgPP:0.989|lnPP:-17.002|num_ambig:17|num_ambig_gaps:8
LRSLSIPLERLHIMKGHMMQDMCKGLSRQTHAQAKVRMLPTYICSTPNGTEKGNFLVVELCQNQVRTLLV
TLYGDGNMSPQMMYKIFDMPEGIMQGEGEALFDFIAQCVSQFLAETISPDTSTSEERLPLGFVFPFSCRQ
TQLDKAELLSWSKGFSCSGVVGKDVVQLLQSAINKQEAGGGGGGGNWLSSWRGRKPSQSAPGQLCHVDVV
ALMNDTVGTMMTCSMEGRPCEVAVVADKGSNCCFMAEAYLVETTEETSGRMCVNTEWGCFGDDGTLNDIF
TPYDECVDEESSNPGEKRFEKLVGSLYLGEIVRHALIALTAEKAIFTGTDVAVLKQKGVFTMQHVLEIIN
NEEGTTEVKRILEVLGLQPSERDCGRVQQICRAVVGRAATLYATGLAAILSYMCQTREMETLMVNVGVEG
ELYTGYPRFEEILQSVSRLLAPECMATLLPSRDGSGRGAAMVTAVALRLAAQRREVNEVLAPLRLSRADL
EKVQALMRQEMEQGLGKETNATASVRMLPTYVSHTPDGTERGDFLALDLGGTNFRVLVVRVTEEGISMAS
EIYVIPAAIMQGTGEALFDHIIDCIIDFQTKQNLMTDVLPLGFTFSFPCKQVGLDKALLLTWTKGFSASG
CVGQDVVQLLREAAQRKQHSGLQVVALVNDTVGTMMSCGYDDPKCEIGLIVGTGTNACYMEEMKNVGTVE
GDQGRMCINMEWGAFGDNGCLDHIFTHFDKVVDETTINPGKQRFEKLISGMYLGEIVRQILLVMTEKQLL
FQGKPCPKLQTKDIFQTKFLSTIELSNGLALRQIRAILNELELDASFEDSVLMREVCQTVSLRAAQLCAA
GLAAVVEKMRENRGLDQLSVTVGVDGTLYKLHPCFSDNLQKTLKDLAPNCNVTFLLSEDGSGKGAALVAA
VANRAANAGQ
>anc206_altAll|avgPP:0.985|lnPP:-25.342|num_ambig:17|num_ambig_gaps:8
LRSLSIPLERLHIMKGHMMQDMCKGLSRQTHTQAKVRMLPTYICSTPNGTEKGNFLVVELCQSQVRTLLV
TLYGDGNMSPHMMYKIFDMPEGIMQGEGEALFDFIAQCVSQFLAETISPDTSSSEERLPLGFVFPFSCRQ
TQLDKAELLSWSKGFSCSGVVGKDVVQLLQSAINKQEAGGGGGGGRWLSSWRGRKPSQPAPGQLCHVDVV
ALMNDTVGTMMTCSTEGRPCEVAVVADKGSNCCFMAEAYLVETTDETSGRMCVNTEWGCFGDDGTLNDIF
TPYDECVDEESSNPGEKRFEKLVGSLYLGEIVRHALIALTAEKAIFTGTNVAVLKEKGVFTMQHVLEIIN
NEEGTTEVKRILEALGLQPSERDCGRVQQICRAVVGRAATLYATGLAAILSYMCQTREMETLMVNVGVEG
ELYTGYPRFEEILQSVSRLLAPECMATLLPSRDGSGRGAAMVTAVALRLAAQRREVNEVLAPLRLSRADL
EKVQALMRQEMERGLGKETNATASVRMLPTYVSHTPDGTERGDFLALDLGGTNFRVLVVRVTEEGISMAS
EIYVIPAAIMQGTGEGLFDHIIDCIMDFQTKQNLMTDVLPLGFTFSFPCKQVGLDKALLLTWTKGFSASG
CVGQDVVQLLREAAQRKQHSGLQVVALVNDTVGTMMSCGYDDPKCEIGLIVGTGTNACYMEEMKNVGTVE
GDQGRMCINMEWGAFGDNGCLDHIFTHFDKVVDESTINPGKQRFEKLISGMYLGEIVRQILLVMTEKQLL
FQGRPCPKLQTKDIFQTKFLSTIELSNGLALRQIRAILNELELDASFEDSVLMREVCQTVSLRAAQLCAA
GLAAVVEKMRENRGLDQLSVTVGVDGTLYKLHPCFSDNLQKTLKDLAPNCNVTFLLSEDGSGKGAALVAA
VASRAANAGQ
>anc207|avgPP:0.985|lnPP:-22.705|num_ambig:25|num_ambig_gaps:9
LRSLSIPLERLHIMKGHMMQDMCKGLSRQTHAQAKVRMLPTYICSTPNGTEKGNFLVVELCQSQVRTLLV
TLYGDGNLSPQMMYKIFDMPEGIMQGEGEALFDFIAQCVSQFLAETISPDTSSSEERLPLGFVFPFSCRQ
TQLDKAELISWSKGFSCSGVVGKDVVQLLQSAINKQEVGGGGGGGRWLSSWRGRKPSQSAPGQLSHVDVV
ALMNDTVGTMMTCSTEGRPCEVAVVADKGSNCCFMAEAHLVETTEETSGRMCVNTEWGCFGDDGTLNDIL
TPYDECVDQESSNPGEKRFEKLVGSLYLGEIVRHALIALTAEKAIFTGTDVAVLKQKGVFTTQQVLEIIN
NEEGTTEVKRILEVLGLQPSERDCGRVQQICRAVVGRAATLYATGLAAILSYMCQSREMEQLMVNVGVEG
ELYQGYTRFEEILQSVSRLLAPECMATLLPSRDGSGRGAAMVTAVALRLEAQRREVNEVLAPLRLSRADL
EKVQALMRQEMELGLGRETNATASVRMLPTYVCGTPDGTERGDFLALDLGGTNFRVLVVRVTEEGISMAS
EIYVIPTAIMQGTGEALFDHIIDCIMDFQMKQNLMTDVLPLGFTFSFPCQQVGLDKALLLTWTKGFSASG
CVGQDVVQLLREAAQRKQHSGLQVVALVNDTVGTMMSCGYDDPKCEIGLIVGTGTNACYMEEMKNVGTVE
```

GDQGRMCINMEWGAFGDNGCLDHIFTHFDKVVDESTINPGKQRFEKLISGMYLGEIVRHILLAMTERQLL
FQGKPCPKLQTKDIFQTKFLSTIELSNGLALRQIRAILQELELDASFEDSVLMREVCQTVSLRAAQLCAA
GLAAVVEKMRENRGLDQLSVTVGVDGTLYKLHPCFSHNLQKTLKDLAPNCNVTFLLSEDGSGKGAALVAA
VASRAANQ
>anc207_altAll|avgPP:0.981|lnPP:-33.335|num_ambig:25|num_ambig_gaps:9
LRSLSIPLERLHIMKGHMMQDMCKGLSRQTHTQAKVRMLPTYICSTPNGTEKGNFLVVELCQSQVRTLLV
TLYGDGNLSPQMMYKIFDMPEGIMQGEGEALFDFIAQCVSQFLAETISSDSSSSEERLPLGFVFPFSCRQ
TQLDKAELISWSKGFSCSGVVGKDVVQLLQSAINKQEAGSGGAGGNWLSPWRSWKLSPPAPGQLCHVDVV
ALMNDTVGTMMTCSTEGRPCEVAVVADKGSNCCFMAEAHLVETTEETSGRMCVNTEWGCFGDDGTLNDIF
TPYDECVDQESSNPGEKRFEKLVGSLYLGEIVRHALIALTAEKAIFTGTNVAVLKEKGVFTTQQVLEIIN
NEEGTTEVKRILEALGLQPSERDCGRVQQICRAVVGRAATLYATGLAAILSYMCQSREMEQLMVNVGVEG
ELYQGYTRFEEILQSVSRLLAPECMATLLPSRDGSGRGAAMVTAVALRLEAQRREVNEVLAPLRLSRADL
EKVQALMRQEMELGLGKETNATASVRMLPTYVCGTPDGTERGDFLALDLGGTNFRVLVVRVTEEGISMAS
EIYVIPTAIMQGTGEALFDHIIDCIMDFQMKQNLMTDVLPLGFTFSFPCQQVGLDKALLLTWTKGFSASG
CVGQDVVQLLREAAQRKQHSGLQVVALVNDTVGTMMSCGYDDPKCEIGLIVGTGTNACYMEEMKNVGTVE
GDQGRMCINMEWGAFGDNGCLDHIFTHFDRVVDETTINPGKQRFEKLISGMYLGEIVRHILLVMTEKQLL
FQGRPCPKLQTKDIFQTKFLSTIELSNGLALRQIRAILQELELDASFEDSVLMREVCQTVSLRAAQLCAA
GLAAVVEKMRENRGLDQLSVTVGVDGTLYKLHPCFSHNLQKTLKDLAPNCNVTFLLSEDGSGKGAALVAA
VASRAASQ
>anc208|avgPP:0.993|lnPP:-10.535|num_ambig:11|num_ambig_gaps:0
LRALTIPLERLQVMKNHMMQAMRKGLSRQTHAQATVQMLPTYICSTPDGTEKGDFLVVELCQSQVRTLLV
TLLGDGNLSPQMMYKTFDMPEEIMKGEGEALFDFIAQCVSQFLADTGSSQERLPLGFVFPFSCRQTQLDK
AELISWSKGFSCSDVEGKDVVQLLQSAINKQELSHVEVVALMNDTVGTMMTCSTEGRPCEVALVMDKGSN
CCFMAEAHLVEIAEETSGRMCVNTEWGCFGDNGSLNDIVTPYDECVDNESSSPGEKRFEKMVGSLYLGEI
VRHVLITLAAEKVLFTGSNVAVLKEKGVLKTQQVLEIINHEEGMAATKRILEALGLQPNERDCSRVQQIC
LAVVSRAASLYAAGLAAVLTYMCQTRDMDQLLVNVGVDGELYHGHTRFEEILQSVTRLLAPECVATFLPS
TDGSGRGAAMVTAVALRLEAQRREVNEVLAPLRLSRADLEQVQALMRQEMELGLGQETNANASVRMLPTY
VCNTPDGTEQGEFLALDLGGTNFRVLVVRVTEEGIRMASEIYVIPTAIMQGTGEALFDHIIDCIIDFQMK
QNLMNHVLPLGFTFSFPCKQVGLDKALLLTWTKGFSASDCVGQDVVQLLREAAQRKHHSGLNVVALVNDT
VGTMMSCGYDDPKCEIGLIVGTGTNACYMEEMRNVGTVEGDEGRMCINMEWGAFGDNGCLDHIFTSFDKV
VDDQTINPGKQRFEKLISGMYLGEIVRHILLAMADRQLLFHGKPSPKLYCKDIFKTKFLSTIELNGLALR
QIRAILQDLELNASYEDSVLMREVCETVSLRAAQLCAAGLAAVVEKMRENRGLDQLSVTVGVDGTLYKLH
PNFSHNLQKTLQDLAPNCTVTFMPSEDGSGKGAALVAAVACRITS
>anc208_altAll|avgPP:0.991|lnPP:-15.539|num_ambig:11|num_ambig_gaps:0
LRALTIPLERLQVMKDHMMQAMRKGLSRQTHAQATVQMLPTYICSTPDGTEKGDFLVVELCQSQVRTLLV
TLLGDGNLSPQMMYKTFDMPEEIMKGEGEALFDFIAQCVSQFLADTSSSQERLPLGFVFPFSCRQTQLDK
AELISWSKGFSCSDVEGKDVGQLLQSAINKQELSHVEVVALMNDTVGTMMTCSTEGRPCEVALVMDKGSN
CCFMAEAHLVEMAKETSGRMCVNTEWGCFGDNGSLNDIVTPYDECVDNESSSPGEKRFEKMVGSLYLGEI
VRHVLITLAAEKVLFTGSNVAVLKEKGVLKTQQVLEIINHEEGMAVTKRILEALGLQPNERDCSRVQQIC
LAVVSRAASLYAAGLAAVLTYMCQTREMDQLLVNVGVDGELYHGHTRFEEILQSVTRLLAPECVATFLPS
TDGSGRGAAMVTAVALRLEAQRREVNEVLAPLRLSRADLEQVQALMRQEMELGLGRETNANASVRMLPTY
VCNTPDGTEQGEFLALDLGGTNFRVLVVRVTEEGIRMASEIYVIPTAIMQGTGEALFDHIIDCIIDFQMK
QDLMNHVLPLGFTFSFPCKQVGLDKALLLTWTKGFSASDCVGQDVVQLLREAAQRKHHSGLNVVALVNDT
VGTMMSCGYDDPKCEIGLIVGTGTNACYMEEMRNVGTVEGDEGRMCINMEWGAFGDNGCLDHIFTSFDKV
VDDQTINPGKQRFEKLISGMYLGEIVRHILLAMADRQLLFQGKPSPKLYCKDIFKTKFLSTIELNGLALR
QIRAILQDLELNASYEDSVLMREVCETVSLRAAQLCAAGLAAVVEKMRENRGLDQLSVTVGVDGTLYKLH
PNFSQNLQKTLQDLAPNCTVTFMPSEDGSGKGAALVAAVACRITS
>anc209|avgPP:0.992|lnPP:-12.996|num_ambig:14|num_ambig_gaps:0
LRALTIPLERLQVMKNHMMQAMRKGLSRQTHAQATVQMLPTYICSTPDGTEKGDFLVVELCQSQVRTLLV
TLFGDGNLSPQMMYKTFDMPEEIMQGEGEALFDFIAQCVSQFLADTGSSQERLPLGFVFPFSCRQTQLDK
AELISWSKGFSCSDVEGKDVVQLLQSAINKQELSHVEVVALMNDTVGTMMTCSTEGRPCEVALVMDKGSN
CCFMAEAHLVEIAEETSGRMCVNTEWGCFGDNGTLNDIVTPYDECVDNESSSPGEKRFEKMVGSLYLGEI

VRHVLITLAAEKALFTGSNVAVLKEKGVLKTQQVLEIINNEEGMAATKRILEALGLQPNERDCSRVQQIC
RAVVSRAATLYAAGLAAVLSYMCQTRDMDQLLVNVGVDGELYHGHTRFEEILQSVTRLLAPECVATFLPS
TDGSGRGAAMVTAVALRLEAQRREVNEVLAPLRLSRADLEQVQALMRQEMELGLGQETNANASVRMLPTY
VCNTPDGTEQGEFLALDLGGTNFRVLVVRVTEEGIRMASEIYVIPTAIMQGTGEALFDHIIDCIIDFQMK
QNLMNHVLPLGFTFSFPCQQVGLDKALLLTWTKGFSASDCVGQDVVQLLREAAQRKHHSGLNVVALVNDT
VGTMMSCGYDDPKCEIGLIVGTGTNACYMEEMRNVGTVEGDQGRMCINMEWGAFGDNGCLDHIFTSFDKV
VDDQTINPGKQRFEKLISGMYLGEIVRHILLAMADRQLLFHGKPSPKLYCKDIFKTKFLSTIELNGLALR
QIRAILQDLELNASYEDSVLMREVCETVSLRAAQLCAAGLAAVVEKMRENRGLDQLSVTVGVDGTLYKLH
PNFSHNLQKTLQDLAPNCTVTFMPSEDGSGKGAALVAAVACRITS
>anc209_altAll|avgPP:0.989|lnPP:-19.108|num_ambig:14|num_ambig_gaps:0
LRALTIPLERLQVMKGHMMQAMRKGLSRQTHAQATVQMLPTYICSTPDGTEKGDFLVVELCQSQVRTLLV
TLFGDGNLSPQMMYKTFDMPEEIMQGEGEALFDFIAQCVSQFLADTSSSQERLPLGFVFPFSCRQTQLDK
AELISWSKGFSCSDVEGKDVGQLLQSAINKQELSHVEVVALMNDTVGTMMTCSTEGRPCEVALVMDKGSN
CCFMAEAHLVEMAKETSGRMCVNTEWGCFGDNGSLNDIVTPYDECVDNESSSPGEKRFEKMVGSLYLGEI
VRHVLITLAAEKALFTGSNVAVLKEKGVLKTQQVLEIINHEEGMAVTKRILEALGLQPNERDCSRVQQIC
RAVVSRAATLYAAGLAAVLSYMCQTREMDQLLVNVGVDGELYHGHTRFEEILQSVTRLLAPECVATFLPS
TDGSGRGAAMVTAVALRLEAQRREVNEVLAPLRLSRADLEQVQALMRQEMELGLGRETNANASVRMLPTY
VCNTPDGTEQGEFLALDLGGTNFRVLVVRVTEEGIRMASEIYVIPTAIMQGTGEALFDHIIDCIIDFQMK
QDLMEHVLPLGFTFSFPCQQVGLDKALLLTWTKGFSASDCVGQDVVQLLREAAQRKHHSGLNVVALVNDT
VGTMMSCGYDDPKCEIGLIVGTGTNACYMEEMRNVGTVEGDQGRMCINMEWGAFGDNGCLDHIFTSFDRV
VDDQTINPGKQRFEKLISGMYLGEIVRHILLAMADRQLLFQGKPSPKLYCKDIFKTKFLSTIELNGLALR
QIRAILQDLELNASYEDSVLMREVCETVSLRAAQLCAAGLAAVVEKMRENRGLDQLSVTVGVDGTLYKLH
PNFSHNLQKTLQDLAPNCTVTFMPSEDGSGKGAALVAAVACRITS
>anc210|avgPP:0.990|lnPP:-15.242|num_ambig:17|num_ambig_gaps:1
LRALTIPLERLQMMKGHMMQAMRKGLSRQTHAQATVQMLPTYICSTPDGTEKGDFLVVELCQSQVRILLV
TLFGDGNLSPQMVYKIFDMPEDIMHGEGEALFDFIAQRLSEFLGDTSSSQERIPLGFVFPFSCRQTKLDK
AELISWSKGFSCSDVEGKDVVQLLQWAIYKQELSHVDVVALMNDTVGTMMTCSTEGRPCEIALIADKGSN
CCFMAEAHLVEIAEETSGRMCVNTEWGCFGDDGMLSDILTPYDESMDDESSNPGLKRFEKLVGTLYLGEI
VRHVLITLAAQKALFTGSNVAVLKEKGVLKTQQVLEIINNGEGMAAAKRILEALGLESSERDCSRVQQIC
QAVVTRAATLYAAGLAAVLSYMCQSRDMDQLLVNVGVDGELYHGCTRFKEILENVVKLLAPECTATLLPS
TDGSGRGAAMVTAVAVRLEAQRREVDEVLAPLRLSRADLERVQALMRQEMELGLGQETNATACVRMLPTY
VCGTPDGTEQGDFLALDLGGTNFRVLVVHVSEEGIRMASEIYVIPTAIMQGTGEQLFDHIVDCIMDFQLK
QNLMEQVLPLGFTFSFPCQQVGLDKAFLLTWTKGFSASGCVGQDVVQLLREAAQRKNVGLQVVALVNDTV
GTMMSCGYDDPKCEIGLIVGTGTNACYMEEMQNVGTVEGEQGRMCINMEWGAFGDNGCLDHIFTSFDTVV
DKDTINPGKQRFEKLISGMYLGEIVRHILLAMVDRQLLFKGNPSPKLQTKNIFQTKFLSTIELNGLALRQ
IRAILQDLELNASFEDSVLMREVCHTVSQRAAKLCAAGLAAVVEKMRENRGLDQLSVTVGVDGTLYKLHP
TFSHILQETLQDLAPKCTVTFMPSEDGSGKGAALVAAVACRISSP
>anc210_altAll|avgPP:0.987|lnPP:-22.721|num_ambig:17|num_ambig_gaps:1
LRALTIPLERLQMMKGHMMQAMRKGLSRQTHAQATVQMLPTYICSTPDGTEKGDFLVVELCQSQVRILLV
TLFGDGNLSPQMVYKIFDMPEDIMHGEGEALFDFIAQRLSEFLGDTSSSQERIPLGFVFPFSCHQTKLDK
AELISWSKGFSCSDVEGKDVVQLLQWAIYKQELCHVDVVALMNDTVGTMMTCSTEGRPCEIALIADKGSN
CCFMAEAHLVEIAEETSGRMCVNTEWGCFGDDGMLSDILTPYDESMDDESSNPGLKRFEKLVGTLYLGEI
VRHVLITLAAQKALFTGSDVAVLKQKGVLKTQQVLEIINNGEGTAAAKRILEALGLESSEQDCSRVQQIC
QAVVTRAATLYAAGLAAVLSYMCQSRDMDQLLVNVGVDGELYHGCTRFKEILENVVKLLAPECTATLLPS
TDGSGRGAAMVTAVAVRLEAQRREVDEVLAPLRLSRADLERVQALMRQEMELGLGRETNATASVRMLPTY
VCGTPDGTEQGDFLALDLGGTNFRVLVVRVSQEGIRMASEIYVIPTGIMQGTGEQLFDHIVDCIMDFQLK
QNLMEQVLPLGFTFSFPCQQVGLDKALLLTWTKGFSASGCVGQDVVQLLREAAQRKNVGLEVVALVNDTV
GTMMSCGYDDPKCEIGLIVGTGTNACYMEEMQNVGTVEGEQGRMCINMEWGAFGDNGCLDHIFTSFDTVV
DKDTINPGKQRFEKLISGMYLGEIVRHILLAMVDRQLLFKGNPSPKLQTKNIFQTKFLSTIELNGLALRQ
IRAILQDLELNASFEDSVLMREVCHTVSQRAAKLCAAGLAAVVEKMRENRGLDQLSVTVGVDGTLYKLHP
TFSHVVQETLQDLAPKCTVTFMPSEDGSGKGAALVAAVACNITSP
>anc211|avgPP:0.987|lnPP:-20.701|num_ambig:25|num_ambig_gaps:1

LRALTIPLERLQMMKGHMMQAMRKGLSRQTHAQATVQMLPTYICSTPDGTEKGDFLVVELCQSQVRTLLV
TLFGDGNLSPQMVYKIFDMPEDIMHGEGEALFDFIAQCLSQFLADTSSSQERLPLGFVFPFSCRQTKLDK
AELISWSKGFSCSDVEGKDVVQLLQSAIYKQELSHVDVVALMNDTVGTMMTCSTEGRPCEIALIADKGSN
CCFMAEAHLVEMAEETSGRMCVNTEWGCFGDDGMLSDILTPYDESMDNESSNPGVKRFEKLVGTLYLGEI
VRHVLITLAAQKALFTGSNVAVLKEKGVLKTQQVLEIINNEEGMAAAKRILEALGLEPSERDCSRVQQIC
QAVVTRAATLYAAGLAAVLSYMCQSRDMDQLLVNVGVDGELYHGHTRFKEILQNVVKLLAPECTATLLPS
TDGSGRGAAMVTAVAVRLEAQRREVNEVLAPLRLSRADLERVQALMRQEMELGLGRETNATACVRMLPTY
VCGTPDGTEQGDFLALDLGGTNFRVLVVHVSEEGIRMASEIYVIPTAIMQGTGEQLFDHIIDCIMDFQLK
QNLMEQVLPLGFTFSFPCQQVGLDKAFLLTWTKGFSASGCVGQDVVQLLREAAQRKNHSGLNVVALVNDT
VGTMMSCGYDDPKCEIGLIVGTGTNACYMEEMQNVGTVEGEQGRMCINMEWGAFGDNGCLDHIFTSFDTV
VDEQTINPGKQRFEKLISGMYLGEIVRHILLAMTDRQLLFKGNPSPKLQTKDIFQTKFLSTIELNGLALR
QIRAILQDLELNASFEDSVLMREVCQTVSLRAAQLCAAGLAAVVEKMRENRGLDQLSVTVGVDGTLYKLH
PNFSHNLQKTLQDLAPNCTVTFMPSEDGSGKGAALVAAVACRITSP
>anc211_altAll|avgPP:0.982|lnPP:-31.466|num_ambig:25|num_ambig_gaps:1
LRALTIPLERLQMMKGHMMQAMRKGLSRQTHAQATVQMLPTYICSTPDGTEKGDFLVVELCQSQVRTLLV
TLFGDGNLSPQMVYKIFDMPEDIMHGEGEALFDFIAQCLSEFLADTSSSQERLPLGFVFPFSCHQTKLDK
AELISWSKGFSCSDVEGKDVVQLLQSAIYKQELCHVDVVALMNDTVGTMMTCSTEGRPCEIALIADKGSN
CCFMAEAHLVEIAEETSGRMCVNTEWGCFGDDGMLSDILTPYDESMDNESSNPGLKRFEKLVGTLYLGEI
VRHVLITLAAQKALFTGSDVAVLKQKGVLKTQQVLEIINNEEGTAAAKRILEALGLEPSEQDCSRVQQIC
QAVVTRAATLYAAGLAAVLSYMCQSREMEQLLVNVGVDGELYHGHTRFKEILQNVIKLLAPECTATLLPS
TDGSGRGAAMVTAVAVRLEAQRREVNEVLAPLRLSRADLEKVQALMRQEMELGLGQETNATASVRMLPTY
VCGTPDGTERGDFLALDLGGTNFRVLVVRVSQEGIRMASEIYVIPTAIMQGTGEQLFDHIVDCIIDFQLK
QNLMEQVLPLGFTFSFPCQQVGLDKALLLTWTKGFSASGCVGQDVVQLLREAAQRKNHSGLNVVALVNDT
VGTMMSCGYDDPKCEIGLIVGTGTNACYMEEMQNVGTVEGEQGRMCINMEWGAFGDNGCLDHIFTSFDTV
VDEQTINPGKQRFEKLISGMYLGEIVRHILLAMVERQLLFKGNPSPKLQTKDIFQTKFLSTIELNGLALR
QIRAILQDLELNASFEDSVLMREVCQTVSLRAAQLCAAGLAAVVEKMRENRGLDQLSVTVGVDGTLYKLH
PNFSHNVQKTLQDLAPNCTVTFMPSEDGSGKGAALVAAVACRTTSP
>anc212|avgPP:0.988|lnPP:-17.297|num_ambig:19|num_ambig_gaps:0
LRALTIPLERLQVMKGHMMQAMRKGLSRQTHAQATVQMLPTYICSTPDGTEKGDFLVVELCQSQVRTLLV
TLFGDGNLSPQMMYKIFDMPEDIMQGEGEALFDFIAQCVSQFLADTSSSQERLPLGFVFPFSCRQTQLDK
AELISWSKGFSCSDVEGKDVVQLLQSAINKQELSHVDVVALMNDTVGTMMTCSTEGRPCEVALVADKGSN
CCFMAEAHLVEMAEETSGRMCVNTEWGCFGDDGTLNDILTPYDECVDNESSNPGEKRFEKLVGSLYLGEI
VRHVLITLAAEKALFTGSNVAVLKEKGVLKTQQVLEIINNEEGMAAAKRILEALGLQPSERDCSRVQQIC
RAVVSRAATLYAAGLAAVLSYMCQSREMDQLLVNVGVDGELYHGHTRFEEILQSVTRLLAPECTATLLPS
TDGSGRGAAMVTAVALRLEAQRREVNEVLAPLRLSRADLERVQALMRQEMELGLGRETNATASVRMLPTY
VCGTPDGTEQGDFLALDLGGTNFRVLVVRVTEEGIRMASEIYVIPTAIMQGTGEALFDHIIDCIMDFQMK
QNLMEQVLPLGFTFSFPCQQVGLDKALLLTWTKGFSASGCVGQDVVQLLREAAQRKHHSGLNVVALVNDT
VGTMMSCGYDDPKCEIGLIVGTGTNACYMEEMQNVGTVEGDQGRMCINMEWGAFGDNGCLDHIFTSFDKV
VDEQTINPGKQRFEKLISGMYLGEIVRHILLAMTDRQLLFQGKPSPKLQTKDIFQTKFLSTIELNGLALR
QIRAILQDLELNASFEDSVLMREVCQTVSLRAAQLCAAGLAAVVEKMRENRGLDQLSVTVGVDGTLYKLH
PNFSHNLQKTLQDLAPNCTVTFMPSEDGSGKGAALVAAVACRITS
>anc212_altAll|avgPP:0.985|lnPP:-27.489|num_ambig:19|num_ambig_gaps:0
LRALTIPLERLQVMKGHMMQAMRKGLSRQTHAQATVQMLPTYICSTPDGTEKGDFLVVELCQSQVRTLLV
TLFGDGNLSPQMMYKIFDMPEDIMQGEGEALFDFIAQCVSQFLADTGSSQERLPLGFVFPFSCRQTQLDK
AELISWSKGFSCSDVEGKDVVQLLQSAINKQELCHVDVVALMNDTVGTMMTCSTEGRPCEVALVADKGSN
CCFMAEAHLVEIAEETSGRMCVNTEWGCFGDDGTLNDILTPYDECVDNESSNPGEKRFEKLVGSLYLGEI
VRHVLITLAAEKALFTGSNVAVLKQKGVLKTQQVLEIINNEEGMAAAKRILEALGLQPSERDCSRVQQIC
RAVVSRAATLYAAGLAAVLSYMCQSRDMEQLLVNVGVDGELYHGHTRFEEILQSVTRLLAPECTATLLPS
TDGSGRGAAMVTAVALRLEAQRREVNEVLAPLRLSRADLEKVQALMRQEMELGLGQETNANASVRMLPTY
VCGTPDGTERGEFLALDLGGTNFRVLVVRVTEEGIRMASEIYVIPTAIMQGTGEALFDHIIDCIIDFQLK
QNLMEQVLPLGFTFSFPCQQVGLDKALLLTWTKGFSASGCVGQDVVQLLREAAQRKQHSGLNVVALVNDT
VGTMMSCGYDDPKCEIGLIVGTGTNACYMEEMRNVGTVEGDQGRMCINMEWGAFGDNGCLDHIFTSFDRV

VDEQTINPGKQRFEKLISGMYLGEIVRHILLAMVERQLLFQGKPSPKLQTKDIFQTKFLSTIELNGLALR
QIRAILQDLELDASFEDSVLMREVCQTVSLRAAQLCAAGLAAVVEKMRENRGLDQLSVTVGVDGTLYKLH
PNFSHNLQKTLQDLAPNCTVTFMPSEDGSGKGAALVAAVACRITS
>anc213|avgPP:0.986|lnPP:-21.557|num_ambig:20|num_ambig_gaps:1
LRALTIPLERLQVMKGHMMQAMRKGLSRQTHAQATVRMLPTYICSTPDGTEKGDFLVVELCQSQVRTLLV
TLFGDGNLSPQMMYKIFDMPEDIMQGEGEALFDFIAQCVSQFLADTSSSQERLPLGFVFPFSCRQTQLDK
AELISWSKGFSCSDVEGKDVVQLLQSAINKQELSHVDVVALMNDTVGTMMTCSTEGRPCEVALVADKGSN
CCFMAEAHLVEMAEETSGRMCVNTEWGCFGDDGTLNDILTPYDECVDQESSNPGEKRFEKLVGSLYLGEI
VRHVLITLAAEKALFTGSNVAVLKEKGVLKTQQVLEIINNEEGMAEAKRILEALGLQPSERDCSRVQQIC
RAVVSRAATLYAAGLAAILSYMCQSREMEQLLVNVGVDGELYQGHTRFEEILQSVTRLLAPECTATLLPS
TDGSGRGAAMVTAVALRLEAQRREVNEVLAPLRLSRADLERVQALMRQEMELGLGRETNATASVRMLPTY
VCGTPDGTERGDFLALDLGGTNFRVLVVRVTEEGIRMASEIYVIPTAIMQGTGEALFDHIIDCIMDFQMK
QNLMEQVLPLGFTFSFPCQQVGLDKALLLTWTKGFSASGCVGQDVVQLLREAAQRKQHSGLNVVALVNDT
VGTMMSCGYDDPKCEIGLIVGTGTNACYMEEMQNVGTVEGDQGRMCINMEWGAFGDNGCLDHIFTSFDKV
VDEQTINPGKQRFEKLISGMYLGEIVRHILLAMTERQLLFQGKPCPKLQTKDIFQTKFLSTIELNGLALR
QIRAILQDLELDASFEDSVLMREVCQTVSLRAAQLCAAGLAAVVEKMRENRGLDQLSVTVGVDGTLYKLH
PNFSHNLQKTLQDLAPNCTVTFLPSEDGSGKGAALVAAVACRAASQ
>anc213_altAll|avgPP:0.981|lnPP:-33.007|num_ambig:20|num_ambig_gaps:1
LRALTIPLERLQVMKGHMMQAMRKGLSRQTHAQATVRMLPTYICSTPDGTEKGDFLVVELCQSQVRTLLV
TLFGDGNLSPQMMYKIFDMPEDIMQGEGEALFDFIAQCVSQFLADTGSSQERLPLGFVFPFSCRQTQLDK
AELISWSKGFSCSDVEGKDVVQLLQSAINKQELCHVDVVALMNDTVGTMMTCSTEGRPCEVALVADKGSN
CCFMAEAHLVEIAEETSGRMCVNTEWGCFGDDGTLNDILTPYDECVDQESSNPGEKRFEKLVGSLYLGEI
VRHALITLAAEKALFTGSNVAVLKQKGVLKTQQVLEIINNEEGMAEAKRILEALGLQPSERDCSRVQQIC
RAVVSRAATLYAAGLAAILSYMCQSREMDQLMVNVGVDGELYHGHTRFEEILQSVTRLLAPECTATLLPS
TDGSGRGAAMVTAVALRLEAQRREVNEVLAPLRLSRADLEKVQALMRQEMELGLGQETNANASVRMLPTY
VCGTPDGTERGEFLALDLGGTNFRVLVVRVTEEGIRMASEIYVIPTAIMQGTGEALFDHIIDCIMDFQLK
QNLMEQVLPLGFTFSFPCQQVGLDKALLLTWTKGFSASGCVGQDVVQLLREAAQRKQHSGLQVVALVNDT
VGTMMSCGYDDPKCEIGLIVGTGTNACYMEEMRNVGTVEGDQGRMCINMEWGAFGDNGCLDHIFTNFDRV
VDEQTINPGKQRFEKLISGMYLGEIVRHILLAMVEKQLLFQGKPCPKLQTKDIFQTKFLSTIELNGLALR
QIRAILQDLELNASFEDSVLMREVCQTVSLRAAQLCAAGLAAVVEKMRENRGLDQLSVTVGVDGTLYKLH
PNFSHNLQKTLQDLAPNCTVTFLPSEDGSGKGAALVAAVACRAASQ
>anc214|avgPP:0.988|lnPP:-18.453|num_ambig:21|num_ambig_gaps:1
LLALTIPLETLQAVKGRMLQAMRKGLSRQTHAQANVRMLPTYICSTPDGTEKGDFLVVELCQSHVRTLWV
TLLGDGNQSPQVMYKIFDMPEDIMQGKGEALFDFIAQCVRQFLAGIGSPQHRLPLGFVFPFSCRQTQLDK
AELISWSKGFSCSDVEGKDVVQLLQSAINKQELYHVDVVALMNDTVGTMMTCSTGGKPCEVALIVDKGTN
SCFMAEAHQVEMVEETSGQMCVNTEWGYFGDDGTLSDILTPYDQHVDQESSNPGEKRFEKLVGSLYLGEI
VRHVLITLAAEKALFIGSNVAVLRTKGVLKTQQILEIIDNEEGMAEAKRTLEALGLQPSERDCCRVQQVC
RAVVSRAAALCAAGLAAILSHMCQSRELERLVVNVAVDGELYQGHSRFGEILQSVTGLLAPECTATLVPS
VDGTGRGAAMVTAVALRLAAQRREVNELLAPLRLSRADLERVQALMRREMELGLGRETNANASVRMLPTY
VCGTPDGTERGEFLALDLGGTNFRVLVVRVAEDGIRMASEIYVIPTAIMQGTGEALFDHIVECIMDFQLK
QDLMEQVLPLGFTFSFPCQQLGLDKAVLLSWTKGFSASGCVGQDVVQLLREAAQRKQHSGLKVVAVVNDT
VGTMMSCGYDDPKCEIGLIVGTGTNACYMEEMQNVGTVEGDQGRMCINMEWGAFGDNGCLDDFFTDFDRL
VDEKTINPGKQRFEKLISGMYLGEIVRHILLALVEKQLLFRGKPCPKLQTKDIFQTKFLSTIEMDGLALQ
QVRAILQDLELEASFEDSVLVREVCQTVSLRAAQLCAAGLAAVVEKMRENRGLAQLAVTVGVDGTLYKMH
PRFSQHLQQTLQDLAPNCTVTFLQSEDGSGKGAALVAAVACRGAEP
>anc214_altAll|avgPP:0.984|lnPP:-29.404|num_ambig:21|num_ambig_gaps:1
LLALTIPLETLQAVKGHMLQAMRKGLSRQTHAQANVRMLPTYICSTPDGTEKGDFLVVELCQSHVRTLWV
TLLGDGNQSPQVMYKIFDMPEDIMQGKGEALFDFIAQCVRQFLAGIGSPQHRLPLGFVFPFSCRQTQLDK
AELISWSKGFSCSDVEGKDVVQLLQSAINKQELYHVDVVALMNDTVGTMMTCSTGGKPCEVALIVDKGTN
CCFMAEAHLVEMVEETSGQMCVNTEWGYFGDDGALSDILTPYDQRVDQESSNPGEKRFEKLVGSLYLGEI
VRHVLITLAAEKALFVGSNVAVLRTKGVLKTQQVLEIIDNEEGMAEAKRTLEALGLQPSERDCCRVQQIC
RVVVSRAAALCAAGLAAILSHMCQSRELERLVVNVAVDGELYQGHSRFGEILQSVTGLLAPECTATLVPS

VDGTGRGAAMVTAVALRLAAQRREVNQLLAPLRLSRADLERVQALMRREMELGLGQETNATASVRMLPTY
VCGTPDGTERGDFLALDLGGTNFRVLVVRVAEDGIRMASEIYVIPTTIMQGTGEALFDHIVDCIMDFQLK
QNLMEQVLPLGFTFSFPCQQLGLDKAVLLSWTKGFSASGCVGQDVVQLLREAAQRKQHSGLKVVAVVNDT
VGTMMSCGYDDPKCEIGLIVGTGTNACYMEEMQNVGTVEGDQGRMCINMEWGAFGDNGCLDDFFTDFDRM
VDEKTINPGKQRFEKLISGMYLGEIVRHILLALVEKQLLFQGKPCPKLQTKDIFQTKFLSTIEIDGLALQ
QVRAILQDLELDASFEDSVLVREVCQTVSLRAAQLCAAGLAAVVEKMRENRGLAQLAVTVGVDGTLYKMH
PHFSQHLQQTLQDLAPNCTVTFLQSEDGSGKGAALVAAVACRGAEP
>anc215|avgPP:0.990|lnPP:-15.086|num_ambig:17|num_ambig_gaps:1
LLALTIPLETLQAVKGRMLQAMRKGLSRQTHAQANVRMLPTYICSTPDGTEKGDFLVVELCQSHVRTLWV
TLLGDGNQSPQVMYKIFDMPEDIMQGKGEALFDFIAQCVRQFLAGIGSPQHRLPLGFVFPFSCRQTQLDK
AELISWSKGFSCSDVEGKDVVQLLQSAINKQELYHVDVVALMNDTVGTMMTCSTGGKPCEVALVVDKGTN
SCFMAEAHQVEMVEETSGRMCVNTEWGCFGDDGTLSDILTPYDQRVDQESSNPGEKRFEKLVGSLYLGEI
VRHVLITLAAEKALFIGSNVAVLRTKGVLKTQQVLEIIDNEEGMAEAKRTLEALGLQPSERDCCRVQQIC
RAVVSRAAALCAAGLAAILSHMCQSRELERLVVNVGVDGELYQGHSRFGEILQSVTGLLAPECTATLVPS
VDGTGRGAAMVTAVALRLAAQRREVNELLAPLRLSRADLERVQALMRREMELGLGRETNANASVRMLPTY
VCGTPDGTERGEFLALDLGGTNFRVLVVRVAEDGIRMASEIYVIPTAIMQGTGEALFDHIIECIMDFQLK
QDLMEQVLPLGFTFSFPCQQLGLDKAVLLSWTKGFSASGCVGQDVVQLLREAAQRKQHSGLKVVAVVNDT
VGTMMSCGYDDPKCEIGLIVGTGTNACYMEEMQNVGTVEGDQGRMCINMEWGAFGDNGCLDDIFTDFDRL
VDEKTINPGKQRFEKLISGMYLGEIVRHILLALVEKQLLFRGKPCPKLQTKDIFQTKFLSTIEMDGLALR
QVRAILQDLELDASFEDSVLVREVCQTVSLRAAQLCAAGLAAVVEKMRENRGLAQLAVTVGVDGTLYKMH
PRFSQHLQQTLQDLAPNCTVTFLQSEDGSGKGAALVAAVACRGAEP
>anc215_altAll|avgPP:0.986|lnPP:-24.862|num_ambig:17|num_ambig_gaps:1
LLALTIPLETLQAVKGHMLQAMRKGLSRQTHAQANVRMLPTYICSTPDGTEKGDFLVVELCQSHVRTLWV
TLLGDGNQSPQVMYKIFDMPEDIMQGKGEALFDFIAQCVRQFLAGIGSPQHRLPLGFVFPFSCRQTQLDK
AELISWSKGFSCSDVEGKDVVQLLQSAINKQELYHVDVVALMNDTVGTMMTCSTGGKPCEVALVVDKGTN
CCFMAEAHLVEMVEETSGRMCVNTEWGCFGDDGALSDILTPYDQHVDQESSNPGEKRFEKLVGSLYLGEI
VRHALITLAAEKALFVGSNVAVLRTKGVLKTQQVLEIIDNEEGMAEAKRTLEALGLQPSERDCCRVQQIC
RAVVSRAAALCAAGLAAILSHMCQSRELERLVVNVGVDGELYQGHSRFGEILQSVTGLLAPECTATLVPS
VDGTGRGAAMVTAVALRLAAQRREVNQLLAPLRLSRADLERVQALMRREMELGLGQETNATASVRMLPTY
VCGTPDGTERGDFLALDLGGTNFRVLVVRVAEDGIRMASEIYVIPTAIMQGTGEALFDHIIDCIMDFQLK
QNLMEQVLPLGFTFSFPCQQLGLDKAVLLSWTKGFSASGCVGQDVVQLLREAAQRKQHSGLKVVAVVNDT
VGTMMSCGYDDPKCEIGLIVGTGTNACYMEEMQNVGTVEGDQGRMCINMEWGAFGDNGCLDDIFTDFDRM
VDEKTINPGKQRFEKLISGMYLGEIVRHILLALVEKQLLFQGKPCPKLQTKDIFQTKFLSTIEIDGLALR
QVRAILQDLELDASFEDSVLVREVCQTVSLRAAQLCAAGLAAVVEKMRENRGLAQLAVTVGVDGTLYKMH
PHFSQHLQQTLQDLAPNCTVTFLQSEDGSGKGAALVAAVACRGAEP
>anc216|avgPP:0.993|lnPP:-10.367|num_ambig:13|num_ambig_gaps:1
LLALTIPLETLQAVKGRMLQAMRKGLSRQTHAQANVRMLPTYICSTPDGTEKGDFLVVELCQSHVRTLWV
TLLGDGNQSPQVMYKIFDMPEDIMQGKGEALFDFIAQCVRQFLAGIGSPQHRLPLGFVFPFSCRQTRLDK
AELISWSKGFSCSDVEGKDVVQLLQSAINKQELYHVDVVALMNDTVGTMMTCSTGGKPCEVALVVDKGTN
SCFMAEAHQVEMVEETSGRMCVNTEWGCFGDDGTLSDILTPYDQRVDQESSNPGEKRFEKLVGSLYLGEI
VRHALIALAAEKALFIGSNVAVLGTKGVLKTQQVLEIIDNEEGMAKARKTLEALGLQPSERDCCRVQQIC
RAVVSRAAALCAAGLAAILSHMCQSRELERLVVNVGVDGELYRGHSRFGEILQSVTGLLAPECTATLVPS
VDGTGRGAAMVTAVALRLAAQRREVDQLLAPLRLSRADLERVQALMRREMELGLGRETNANASVRMLPTY
VCGTPDGTERGEFLALDLGGTNFRVLVVRVAQDGIRMASEIYVIPTAIMQGTGEALFDHIIECIMDFQLK
QDLMEQVLPLGFTFSFPCQQLGLDKAVLLSWTKGFSASGCVGQDVVQLLREAAQRKQHFGLKVVAVVNDT
VGTMMSCGYDDPKCEIGLIVGTGTNACYMEEMQNVGTVEGDQGRMCINMEWGAFGDNGCLDDIFTDFDRL
VDEKTINPGKQRFEKLISGMYLGEIVRHILLALVEKQLLFRGKPCPKLQTRDIFQTKFLSTIEMDGLALR
QVRAILQDLELDASFEDSVLVREVCQTVSLRAAQLCAAGLAAVVEKMRENRGLAQLAVTVGVDGTLYKMH
PRFSQHLQQTLQDLAPNCTVTFLQSEDGSGKGAALVAAVACRGAEP
>anc216_altAll|avgPP:0.990|lnPP:-18.515|num_ambig:13|num_ambig_gaps:1
LLALTIPLETLQAVKGRMLQAMRKGLSRQTHAQANVRMLPTYICSTPDGTEKGDFLVVELCQSHVRTLWV
TLLGDGNQSPQVMYKIFDMPEDIMQGKGEALFDFIAQCVRQFLAGIGSPQHRLPLGFVFPFSCRQTRLDK

AELISWSKGFSCSDVEGKDVVQLLQSAINKQELYHVDVVALMNDTVGTMMTCSTGGKPCEVALVVDKGTN
SCFMAEAHQVEMVEETSGRMCVNTEWGCFGDDGALSDILTPYDQRVDQESSNPGEKRFEKLVGSLYLGEI
VRHALTALAAEKALFVGSNVAVLGTKGVLKTQQVLEIIDNEEGMAKARRALEALGLQPSERDCCRVQQIC
RAVVSRAAALCAAGLAAILSHMCQSRELERLVVNVGVDGELYRGHSRFGEILQSVTGLLAPECTATLVPS
VDGTGRGAAMVTAVALRLAAQRREVDQLLAPLRLSRADLERVQALMRREMELGLGQETNANASVRMLPTY
VCGTPDGTERGEFLALDLGGTNFRVLVVRVAEDGIRMASEIYVIPTAIMQGTGEALFDHIIDCIMDFQLK
QDLMEQVLPLGFTFSFPCQQLGLDKAVLLTWTKGFSASGCVGQDVVQLLREAAQRKQHFGLKVVAVVNDT
VGTMMSCGYDDPKCEIGLIVGTGTNACYMEEMQNVGTVEGDQGRMCINMEWGAFGDNGCLDDIFTDFDRM
VDEKTINPGKQRFEKLISGMYLGEIVRHILLALVEKQVLFRGKPCPKLQTRDIFQTKFLSTIEIDGLALR
QVRAILQDLELQASFEDSVLVREVCQTVSLRAAQLCAAGLAAVVEKMRENRGLAQLAVTVGVDGTLYKMH
PRFSQHLQQTLQDLAPNCTVTFLQSEDGSGKGAALVAAVACRGAEP
>anc217|avgPP:0.992|lnPP:-11.954|num_ambig:12|num_ambig_gaps:0
LLALTPPLETLQAVKGRMLQAMRKGLSRQTHAQANVRMLPTYICSTPDGTEKGDFLVVELCQSHVRTLWV
TLLGDGNQSPQVMYKIFDMPEDIMQGKGEALFDFIAQCVRQFLAGIGSPQHRLPLGFVFPFSCRQTRLDK
AELISWSKGFSCSDVEGKDVVQLLQSAINKQELYHVDVVALMNDTVGTMMTCSTGGKPCEVALVVDKGTN
SCFMAEAHQVEMVEETSGRMCVNTEWGCFGDDGTLSDILTPYDQRVDQESSNPGEKRFEKLVGSLYLGEI
VRHALIALAAEKALFIGSNVAVLGTKDVLKTQQVLEIIDNEEGMAKARKTLEALGLQPSERDCCRVQQIC
RAVVSRAAALCAAGLAAILSHMCQSRELERLVVNVGVDGELYRGHARFGEILQSVTGLLAPECTATLVPS
VDGTGRGAAMVTAVALRLAAQRREVDQLLAPLRLSRADLERVQALMRREMELGLGRETNANASVRMLPTY
VCGTPDGTERGEFLALDLGGTNFRVLVVRVAQDGIRMASEIYVIPTAIMQGTGEALFDHIIECIMDFQLK
QDLMEQVLPLGFTFSFPCQQLGLDKAVLLSWTKGFSASGCVGQDVVQLLREAAQRKQHFGLKVVAVVNDT
VGTMMSCGYDDPKCEIGLIVGTGTNACYMEEMQNVGTVEGDQGRMCINMEWGAFGDNGCLDDIFTDFDRL
VDEKTINPGKQRFEKLISGMYLGEIVRHILLALVEKQLLFRGKPCPKLQTRDIFQTKFLSTIEMDGLALR
QVRAILQDLELDASFEDSVLVREVCQTVSLRAAQLCAAGLAAVVEKMRENRGLAQLAVTVGVDGTLYKMH
PRFSQHLQQTLQDLAPNCTVTFLQSEDGSGKGAALVAAVACRGAEP
>anc217_altAll|avgPP:0.989|lnPP:-18.735|num_ambig:12|num_ambig_gaps:0
LLALTPPLETLQAVKGRMLQAMRKGLSRQTHAQANVRMLPTYICSTPDGTEKGDFLVVELCQSHVRTLWV
TLLGDGNQSPQVMYKIFDMPEDIMQGKGEALFDFIAQCVRQFLAGIGSPQHRLPLGFVFPFSCRQTGLDK
AELISWSKGFSCSDVEGKDVVQLLQSAINKQELYHVDVVALMNDTVGTMMTCSTGGKPCEVALVVDKGTN
SCFMAEAHQVEMVEETSGRMCVNTEWGCFGDDGALSDILTPYDQRVDQESSNPGEKRFEKLVGSLYLGEI
VRHALTALAAEKALFVGSNVAVLGTKDVLKTQQVLEIIDNEEGMAKARRALEALGLQPSERDCCRVQQIC
RAVVSRAAALCAAGLAAILSHMCQSRELERLVVNVGVDGELYRGHARFGEILQSVTGLLAPECTATLVPS
VDGTGRGAAMVTAVALRLAAQRREVDQLLAPLRLSRADLERVQALMRREMELGLGRETNANASVRMLPTY
VCGTPDGTERGEFLALDLGGTNFRVLVVRVAEDGIRMASEIYVIPTAIMQGTGEALFDHIIECIMDFQLK
QDLMEQVLPLGFTFSFPCQQLGLDKAVLLTWTKGFSASGCVGQDVVQLLREAAQRKQHFGLKVVAVVNDT
VGTMMSCGYDDPKCEIGLIVGTGTNACYMEEMQNVGTVEGDQGRMCINMEWGAFGDNGCLDDIFTDFDRM
VDEKTINPGKQRFEKLISGMYLGEIVRHILLALVEKQVLFRGKPCPKLQTRDIFQTKFLSTIEIDGLALR
QVRAILQDLELQASFEDSVLVREVCQTVSLRAAQLCAAGLAAVVEKMRENRGLAQLAVTVGVDGTLYKMH
PRFSQHLQQTLQDLAPNCTVTFLQSEDGSGKGAALVAAVACRGAEP
>anc218|avgPP:0.985|lnPP:-23.100|num_ambig:22|num_ambig_gaps:3
LLALTPSLETLQAVKGRMLEAMEKGLSRQTHAQATVRMLPTYICSTPDGTEKGDFLVVELCQSHVRTLWV
TLLGDGNQSPQLMYKIFDMPEDIMQGKGEALFDFIAQCVRQFLEGIGSPQHRLPLGFVFPFSCRQTRLDK
AELISWSKGFRCSDVEGKDVVQLLQAAINKQELYHVDVVALMNDTVGTMMTCSTGGKPCEVALVVDTGTN
SCFMAEAHQVEMVEETSGRMCVNTEWGCFGDDGALSDVLTPYDQRVDQESSNPGEKRFEKLVGSLYLGEI
VRHALIALAAEKALFIGSNTAVLGTKDVFKTQQVLEIIDNEEGMAKARSTLEALGLQPSERDCCRVQQIC
RAVVSRAAALCAAGLAAILSHMCQSRELERLVVNVGVDGELYRGHARFGEILQSVTGLLAPECTATLVPS
ADGSGRGAAMVTAVALRLAAQRREVDQLLAPLRLSRADLERVQALMRREMELGLGRETNANASVRMLPTY
VCNTPDGTERGDFLALDLGGTNFRVLVVRVAQDGIHMASEIYVIPTAIMQGTGEALFDHIIECIMDFQLK
QDLMEQVLPLGFTFSFPCQQLGLDKAVLLSWTKGFSASGCVGQDVVQLLREAAQRKQHFGLKVVAVVNDT
VGTMMSCGYDDPKCEIGLIVGTGTNACYMEEMQNVGTVEGDEGRMCINMEWGAFGDNGCLDDIFTDFDRL
VDEKTINPGKQRFEKLISGMYLGEIVRHILLALVEKQLLFRGKPSPKLQTRDIFQTKFLSTIEMDGLALR
QVRGILQDLELDASFEDSVLVREVCQTVSLRAAQLCAAGLAAVVEKMRENRGLAQLAVTVGVDGTLYKMH

PRFSQHLQQTLQDLAPNCTVTFLQSEDGSGKGAALVAAVACRGAEQGPP
>anc218_altAll|avgPP:0.980|lnPP:-36.016|num_ambig:22|num_ambig_gaps:3
LLALTPSLETLQAVKGRMLEAMDKGLSRQTHAQATVRMLPTYICSTPDGTEKGDFLVVELCQSHVRTLWV
TLLGDGNQSPQVMYKIFDMPEDIMQGKGEALFDFIAQCVRQFLEGIGSPQHRLPLGFVFPFSCRQTGLDK
AELISWSKGFRCSDVEGKDVVQLLQSAINKQELYHVDVVALMNDTVGTMMTCSTGGKPCEVALVVDTGTN
SCFMAEAHQVEMVEETSGRMCVNTEWGCFGDDGALSDILTPYDQRVDQESSNPGEKRFEKLVGGLYLGEI
VRHALTMLAAEQALFVGSNTAVLGTKDVLKTQQVLEIIDNEEGMAKARSALEALGLQPSERDCCRVQQIC
RAVVSRAAALCAAGLAAILSHMCRSRELERLVVNVGVDGELYRGHARFGEILQSVTGLLAPECTATLVPS
ADGSGRGAAMVTAVALRLAAQRREVDQLLAPLRLSRADLQRVQALMRREMELGLGRETNANASVRMLPTY
VCNTPDGTERGEFLALDLGGTNFRVLVVRVAEDGIRMASEIYVIPTAIMQGTGEALFDHIIECIMDFQLK
QDLMEQVLPLGFTFSFPCQQLGLDKAVLLTWTKGFSASGCVGQDVVQLLREAAQRKQHFGLKVVAVVNDT
VGTMMSCGYDDPKCEIGLIVGTGTNACYMEEMQNVGTVEGDQGRMCINMEWGAFGDNGCLDDIFTDFDRM
VDEKTINPGKQRFEKLISGMYLGEIVRHILLALVEKQVLFRGKPSPKLQTRDIFQTKFLSTIEIDGLALR
QVRGILQDLELDASFEDSVLVREVCQTVSLRAAQLCAAGLAAVVEKMRENRGLAQLAVTVGVDGTLYKMH
PRFSQHLQQTLQDLAPNCTVTFLQSEDGSGKGAALVAAVACRGAEQGPP
>anc219|avgPP:0.983|lnPP:-26.153|num_ambig:25|num_ambig_gaps:0
LLELTLSLETLQAVKGHMLEAMAKGLSRQTHAQATVRMLPTYICSTPDGTEKGDFLVVELRRRHVRTLWV
TLPGDGNQSPQLKDQLFDMPEDIMQGKGEALFDFIAQCLCQFLEGVGLSEPQQRLPLGFVFPFSCRQTRL
DQGELITWSKGFRCSDVEGHDVVQLLQAAINKQERYQVDVVALMNDTVGTMMSCSTEGMPCEVALVVDTG
TNSCFMAEGCHVEMVEETSGRMCVNTEWGSFGDDGALSNVLTPYDQCVDQESAKPGEKRFEKLIGGLYLG
EIVRHVLAKLAAEGALFGGTSTPVLGTKDVFKTQQVLEIIDDEAGLAMARSTLQALGLAPSEQDCCHVQQ
VCRAVVTRAAGLCAAGLAAILSHMCQSRELERLKTNVGVDGAVYRGHARFGEILQRITKLLAPECTATLV
PSADGGGRGAAMVTAVALRLAAQRREVDQLLAPLRLSPADLQRVQALMRREMELGLNKETNAQSSVRMLP
TYVRSTPDGTERGDFLALDLGGTNFRVLVVRVGQDGVHMASEIYAIPQAIMQGTGEALFDHIIDCIVDFQ
QKQGLTGQVLPLGFTFSFPCQQLGLDKAVLLSWTKGFSASGCVGKDVVQMLREAAQRKQHFGLDVVAVVN
DTVGTMMSCGYDDPSCEIGLIVGTGTNACYMEEMQNVGTVEGEEGRMCINMEWGAFGDNGCLDDIFTDFD
RLVDEKTINPGKQRFEKLISGMYLGEIVRYILLALVEKHILFRGKPSPKLQTRDIFQTKFLSTIEMGPRC
AGSDGLALRQVRGVLHDLELDASFEDSVLVREVCQTVSLRAAQLCAAGLAAVVEKMRESRGLAQLAVTVG
VDGTLYKMHPRFSQHLQRTLQDLAPNCTVTFLQSEDGSGKGAALVAAVQCRVAQAQGPP
>anc219_altAll|avgPP:0.980|lnPP:-37.151|num_ambig:25|num_ambig_gaps:0
LLELTLSLETLQAVKGHMLEAMAKGLSRQTHAQATVRMLPTYICSTPDGTEKGDFLVVELRRRHVRTLWV
TLPGDGNQSPQLKERLFDMPEDIMQGKGEALFDFIAQCLCQFLDGVGPSEPQQRLPLGFVFPFSCRQTRL
DQGELISWSKGFRCSDVEGHDVVQLLQAAINKQERYQVDVVALMNDTVGTMMSCSTEGMPCEVGLVVDTG
TNSCFMAEGWHVEIVEETSGRMCVNTEWGSFGDDGALSNVLTPYDQSVDQESAKPGEKRFEKLIGGLYLG
EIVRHVLAKLAAEGALFSGTSTAALGTKDVFKTQQVLEIIDDEAGLAKARSTLQALGLAPSEQDCCHVQQ
VCRAVVTRAAGLCAAGLAAILSHMCRSRELEQLKANVGVDGAVYRGHARFGEILQRITKLLAPECTATLV
PSADGGGRGAAMVTAVALRLAAQRREVDQLLAPLRLSPADLQRVQALMRREMELGLNKETNAQSSVRMLP
TYVRNTPDGTERGDFLALDLGGTNFRVLVVRVGQDGVRMASEIYAIPLAIMQGTGEALFDHIIDCIIDFQ
QKQGLTGQVLPLGFTFSFPCQQLGLDKAVLLSWTKGFSASGCVGKDVVQMLREAAQRKQHFGLDVVAVVN
DTVGTMMSCGYDDPSCEIGLIVGTGTNACYMEEMQNVGTVEGEQGRMCINMEWGAFGDNGCLDDIFTDFD
RMVDEKTINPGKQRFEKLISGMYLGEIVRYILLALVEKHILFRGKPSPKLQTRDIFQTKFLSTIEIGPRC
AGSDGLALRQVRGVLHDLELDASFEDSVLVREVCQTVSLRAAQLCAAGLAAVVEKMRESRGLAQLAVTVG
VDGTLYKMHPRFSQHLQQTLQDLAPNCTVTFLQSEDGSGKGAALVAAVHCRVAQAQGPP
>anc220|avgPP:0.973|lnPP:-41.913|num_ambig:35|num_ambig_gaps:0
LLALTLSLETLQAVKGHMLEAMEKGLSRQTHAQATVRMLPTYICSTPDGTEKGDFLVVELRQSHVRTLWV
TLPGDGNQSPQLKDQLFDMPEDIMQGKGEALFDFIAQCLCQFLEGVGLPQQRLPLGFVFPFSCRQTRLDQ
GELISWSKGFRCSDVEGKDVVQLLQAAINKQERYQVDVVALMNDTVGTMMSCSTEGKPCEVALVVDTGTN
SCFMAEARHVEMVEETSGRMCVNTEWGSFGDDGALSDVLTPYDQRVDQESSNPGEKRFEKLIGGLYLGEI
VRHVLAKLAAEGALFGGSSTPVLGTKDVFKTQQVLEIIDDEAGLAKARSTLQALGLAPSEQDCCHVQQVC
RAVVTRAAGLCAAGLAAILSHMCRSRELERLKTNVGVDGAVYRGHARFGEILQRITKLLAPECTATLVPS
ADGSGRGAAMVTAVALRLAAQRREVDQLLAPLRLSPADLQRVQALMRREMELGLNKETNAQSSVRMLPTY
VCSTPDGTERGDFLALDLGGTNFRVLVVRVGQDGVHMASEIYAIPPAIMQGTGEALFDHIIDCIMDFQQK

QGLMGQVLPLGFTFSFPCQQLGLDKAVLLSWTKGFSASGCVGQDVVQLLREAAQRKQHFGLDVVAVVNDT
VGTMMSCGYDDPSCEIGLIVGTGTNACYMEEMQNVGTVEGDEGRMCINMEWGAFGDNGCLDDIFTDFDRL
VDEKTINPGKQRFEKLISGMYLGEIVRHILLALVEKHILFRGKPSPKLQTRDIFQTKFLSTIEMDGLALR
QVRGILQDLELDASFEDSVLVREVCQTVSLRAAQLCAAGLAAVVEKMRENRGLAQLAVTVGVDGTLYKMH
PRFSQHLQQTLQDLAPNCTVTFLQSEDGSGKGAALVAAVACRVAQQGPP
>anc220_altAll|avgPP:0.967|lnPP:-59.885|num_ambig:35|num_ambig_gaps:0
LLELTLSLETLQAVKGRMLEAMAKGLSRQTHAQATVRMLPTYICSTPDGTEKGDFLVVELRRSHVRTLWV
TLPGDGNQSPQLKEQIFDMPEDIMQGKGEALFDFIAQCLCQFLDGIGLSQQRLPLGFVFPFSCRQTRLDQ
AELISWSKGFRCSDVEGHDVVQLLQAAINKQERYQVDVVALMNDTVGTMMSCSTGGKPCEVGLVVDTGTN
SCFMAEAHHVEMVEETSGRMCVNTEWGSFGDDGALSDVLTPYDQRVDQESANPGEKRFEKLIGGLYLGEI
VRHVLAKLAAEGALFSGTSTAALGTKDVFKTQQVLEIIDDEAGLAKARSTLQALGLQPSEQDCCHVQQVC
RAVVTRAAALCAAGLAAILSHMCQSRELERLKANVGVDGAVYRGHARFGEILQRITKLLAPECTATLVPS
ADGSGRGAAMVTAVALRLAAQRREVDQLLAPLRLSPADLERVQALMRREMELGLNKETNAQSSVRMLPTY
VCNTPDGTERGDFLALDLGGTNFRVLVVRVGQDGVRMASEIYAIPPAIMQGTGEALFDHIIDCIVDFQQK
QGLTGQVLPLGFTFSFPCQQLGLDKAVLLSWTKGFSASGCVGQDVVQLLREAAQRKQHFGLDVVAVVNDT
VGTMMSCGYDDPSCEIGLIVGTGTNACYMEEMQNVGTVEGEQGRMCINMEWGAFGDNGCLDDIFTDFDRM
VDEKTINPGKQRFEKLISGMYLGEIVRHILLALVEKQILFRGKPSPKLQTRDIFQTKFLSTIEIDGLALR
QVRGILHDLELDASFEDSVLVREVCQTVSLRAAQLCAAGLAAVVEKMRENRGLAQLAVTVGVDGTLYKMH
PRFSQHLQQTLQDLAPNCTVTFLQSEDGSGKGAALVAAVACRVAEQGPP
>anc221|avgPP:0.928|lnPP:-123.384|num_ambig:91|num_ambig_gaps:0
LQPLCLSLEQLQQVKDSMLEAMERGLSRQTHAQATVRMLPTYIRSTPDGTERGDFLVLELRQTHLRVLWV
TLSGDTNQSVEVKQQLFPIPEDIMQGPGEELFDFMAQCLCEFLDGLRLPKQRFPLGFSFPFPCRQTRLDQ
SELISWTKGFRCPGVEGRDVVQLLQNAINKQERYQIDVVAVVNDTVGTMMSCSSEAQPCEVGLVVDTGTN
SCFMAEARHVEMVEESEGRMCVNTEWGSFGDDGALNDVLTPYDQLVDQESLNPGKSRFEKMISGLYLGEI
VRHILAKLAAQGALFGGTTTPALLTKGSFQTQDVLEITDDQAGLAKARSVLRQLGLEPSEQDCFHVQQVC
SAVCTRAASLCAAGLAAILSHMCRNRELTHLKTSVAVDGAVYRGHAKFSEILQQIVKQLAPECDVTLVPS
EDGSGRGAAMVTAVASRLAAQRRQVNQTLAPLRLSPDDLEKVQALMRREMELGLKKETHAKSSVRMLPTY
VCATPDGTERGDFLALDLGGTNFRVLLVQVGSREEGGVQMVSEIYSIPPDIMQGTGEQLFDHIVDCIMDF
QKKQGMMGQTLPLGFTFSFPCQQLGLDQGVLLSWTKGFNASGCVGQDVVKLLREAVKRKQHFGLDVVAIV
NDTVGTMMSCGYDDPSCEVGLIVGTGTNACYMEEMRNVGTVEGDEGRMCINMEWGAFGDNGCLDDFVTPF
DRLVDEASINPGKQRFEKLISGMYLGEIVRHILLELTEKGILFRGRPSPKLQTRDIFQTKFLSSIESDGL
ALRQVRAILEDLGLDVTCEDSVLVREVCQAVSRRAAQLCGAGVAAVVEKIRENRGLEQLSVTVGVDGTLY
KLHPHFSQQVQETVQELAPRCSVTFLQSEDGSGKGAALIAAVACRVAQHGPA
>anc221_altAll|avgPP:0.915|lnPP:-159.713|num_ambig:91|num_ambig_gaps:0
LQPLCLSLEKLQRVKDSMLEAMDKGLSRQTHAQATVRMLPTYICSTPDGTEKGDFLVLELRGTHVRVLWV
TLTGDANQSVQVKQQIFPVPEDIMQGSGEQLFDFMAQCLCQFLDRLRLSRRALPLGFSFPFPCRQTRLDQ
SELISWTKGFKCSGVEGKDVVQLLQDAINKQDRYQIDVIAVVNDTVGTMMSCGTEAQPCEVGLVVDTGTN
SCFMAEARHVEMVEETEGRMCVNTEWGSFGDDGALNDMLTPYDRLVDQESLNPGKQRFEKMISGLYLGEI
VRHILVKLATQGALFSGTTTPALLTKGSFQTQHILEITDEKVGLAKAQSILQQLGLEPSDQDCFHVQQVC
RAVCTRAANLCAAGLAAILSHMCRNRELTHLKTSVAVDGAMYRGHARFSEILQQTVKQLAPECDVTLVPS
EDGSGKGAAMVTAVASRLAAQRQQVDQTLAPLRLSPEDLQRVQALMRREMELGLKKETNAKSSVRMLPTY
VCATPDGTERGDFLALDLGGTNFRVLLVQVGSREQGGVHIVSEVYSIPPEVTQGTGEQLFDHIVDCIVDF
QKKQGMMGQVLPLGFTFSFPCQQTGLDQGILLSWTKGFNASDCVGQDVVKLLREAVQRKQHFDLDVVAIV
NDTVGTMMSCGYDDPSCEVGLIVGTGTNACYMEEMRNVGTVEGDEGRMCINMEWGAFGDNGCLDDIITPF
DRLVDEASINPGKQRFEKLISGMYLGEIVRHILLDLTAKGILFRGKPSPKLQTRDIFQTKFLSTIESDGL
ALRQVCAILEDLGLHVTCEDSVLVREVCQTVSRRAAQLCGAGVAAVVEKIRENRGLDQLAVTVGVDGTLY
KMHPHFSQHVQKTVQDLAPRCTVTFLQSEDGSGKGAALIAAVACRVAQQGQP
>anc222|avgPP:0.917|lnPP:-143.416|num_ambig:93|num_ambig_gaps:0
LQPFCLSLEQLQQVKDSMLEAMERGLRRQTHATSTVRMLPTYVRSTPDGTERGDFLVLELGQTQLRVLWV
TLTGDANQPVEVKQQIFPIPEDIMQGSGQQLFDFMAQCLSEFLDGLRLSKKRFPLGFSFPFPCRQTRLDQ
STLISWTKGFRCSGVEGRDVVQLLQDAINKQERYQIDVVAVVNDTVGTMMSCSSEAQPCEIGLVVDTGTN
SCYMEEARHVEMVEESEGRMCVNTEWGSFGDDGALNDVLTPFDLLVDQESLNPGKQRFEKMISGMYLGEI

VRLILAKLAEQGALFGGATTPALLTKGSFQTQDVLEITDDKAGLAKAKSVLRQLGLEPSEQDCFHVQQVC
SAVCTRAANLCAAGLAAILSHIRRNRELTHLKTSVAVDGAVYRSHAKFSEILQQTVKQLAPECDVTFVPS
EDGSGRGAAMVTAVASRLAAQRRQVNQTLAPFRLSPEDLEKVQALMRREMERGLKKETHAKSSVRMLPTY
VCATPDGTERGDFLALDLGGTNFRVLLVQVGSREEGGVQMVSEIYSIPPEITQGTGEQLFDHIVDCIVDF
QKKQGMMGQTLPLGFTFSFPCQQLGLDQGILLSWTKGFNASGCVGQDVVKLLREAVKRKQHFDLDVVAIV
NDTVGTMMSCGYDDPSCEVGLIVGTGTNACYMEEMRNVGTVEGDEGRMCINMEWGAFGDNGCLDDFITPF
DRLVDKASINPGKQRFEKLISGMYLGEIVRHILLDLTKKGILFRGRPSQRLQTRDIFQTKFLSDIESDSL
ALRQVRAILEDLGLDVTCEDAVVVREVCQAVSRRAAQLCGAGVAAVVEKIRENRGLEQLSVTVGVDGTLY
KLHPHFSQQVQETVQELAPRCSVTFLQSEDGSGKGAALIAAVACRVAETGPA
>anc222_altAll|avgPP:0.904|lnPP:-180.266|num_ambig:93|num_ambig_gaps:0
LQPFCLSLEQLQQVKDSMLEAMEKGLKRQTHATSTVRMLPTYVCSTPDGTEKGDFLVLELRGTQVRVLWV
TLSGDTGQSVEVKSQVFPIPEDIMQGTGEELFDFMAQCLCEFLDRLRLPKQRLPLGFSFPFPCRQTRLDQ
STLISWTKGFKCPGVEGKDVVQLLQNAINRQERYQIDVIAVVNDTVGTMMSCGSEAQPCEVGLVVDTGTN
SCYMEEARHVEMVEETEGRMCVNTEWGSFGDDGALNDILTPFDLQVDQESLNPGKSRFEKMISGLYLGEI
VRLILVKLAAQGALFSGATTPALLTKGSFETQHILEITDEKVGLAKAQSILQQLGLEPSDQDCFHVQQVC
SAVSTRAANLCAAGLAAILTHMQRNRELTQLKTSVAVDGAMYRSHPRFSEILQQTVKQLAPECDVTFVPS
EDGSGKGAAMVTAVASRLAAQRQQVDQTLAPLRLSPDDLQRVQALMRREMERGLRKETNAKSSVRMLPTY
VCATPDGTERGDFLALDLGGTNFRVLLVQVGSREEGGVHIVSEVYSIPPDVMQGTGEQLFDHIVDCIVDF
QKKQGMMGQVLPLGFTFSFPCQQTGLDQGILLSWTKGFNASDCVGQDVVKLLREAVQRKQHVELDVVAIV
NDTVGTMMSCGYDDPSCEVGLIVGTGTNACYMEEMRNVGTVEGDEGRMCINMEWGAFGDNGCLDDIVTPF
DRLVDEASINPGKQRFEKLISGMYLGEIVRHILLELTKRGILFRGKPSPKLQTRDIFQTKFLSEIESDSL
ALRQVCAILEDLGLDVTCEDAVVVREVCQTVSRRAAQLCGAGVAAVVEKIRENRGLDQLAVTVGVDGTLY
KMHPHFSQHVQETVQELAPRCSVTFLQSEDGSGKGAALIAAVACRVAQTGQP
>anc223|avgPP:0.919|lnPP:-143.890|num_ambig:72|num_ambig_gaps:0
LQPFCLSEEQLQQIKDSMLEDMERGLRRQTHPTSTVRMLPTYVRSTPDGTERGDFLVLELGGTKLRVLWV
TLTGDAGQPVEVKSQIFAIPEDIMLGTGQQLFDFVAQCLSEFLDQLQLSKKPLPLGFSFSFPCRQTRLDR
STLISWTKGFRCSGVEGKDVVQLLRDAIQRQGKYQIDVVAVVNDTVGTMMSCGSEDQPCEIGLVVDTGTN
ACYMEEARHVEAVEEDEGRMCVNTEWGSFGDDGALNDIQTPFDLQVDQESLNPGKQRFEKMISGMYLGEI
VRLILVKLAEQGALFGGATTPALLTKGSFQTQHVLEIEDDKAGLAKAKAILQELGLEPSEQDCFHVQQVC
SAVSTRAANLCAAGLAAILSRIQRNRELSHLRTSVAVDGAVYKQHPKFSEILQQTVKQLAPECDVTFVPS
EDGSGRGAAMVTAVASRLAAQRRQVNETLAPFRLSPEQLQEVQAQMREEMERGLRKETHAKSSVRMLPTY
VRATPDGTERGDFLALDLGGTNFRVLLVQVGSREEGGVQMTSQIYSIPEEVTQGTGEQLFDHIVDCIVDF
QKKQGMMGQTLPLGFTFSFPCQQLGLDQGILLSWTKGFNASGCEGQDVVKLLREAIKRKQDVELDVVAIV
NDTVGTMMSCGYDDPNCEVGLIVGTGTNACYMEEMRNVGTVEGDEGRMCINMEWGAFGDNGCLDDFITPF
DRMVDKSSINPGKQRFEKLISGMYLGEIVRHILLDLTKRGILFRGRQSQRLQTRDIFQTKFLSEIESDSL
ALRQVRAILEDLGLDVTCDDAVVVREVCQAVSRRAAQLCGAGVAAVVEKIRENRGLDQLSVTVGVDGTLY
KLHPHFSQQVQETVRELAPRCSVTFLQSEDGSGKGAALITAVACRVAETGPA
>anc223_altAll|avgPP:0.908|lnPP:-176.493|num_ambig:72|num_ambig_gaps:0
LQPFCLSEEQLQQVKDSMLENMEKGLRRQTNPTSTVRMLPTYVRSTPDGTEKGDFLVLELGGAKLRVLWV
KLTGDAGQSVEVKSQVFAIPEDIMLGSGQQLFDFVAQCLSEFLDRLQLSKQPLPLGFSFSFPCRQTGLDK
STLISWTKGFKCSGVEGKDVVQLLRDAIQRQEKYNIDVIAVVNDTVGTMMSCGSEEQPCEVGLIVDTGTN
SCYMEEARHVEAVEEDEGRMCVNTEWGSFGDDGALNDVQTPFDLQVDQESLNPGKQRFEKMISGMYLGEI
VRLVLVKLAEQGALFGGATTPALLTKGSFETQHVLEIEDEKVGLAKAQAVLQELGLEPSDQDCFHVQQVC
SAVCTRAANLCAAGLAAILTRIQRNRELSQLKTTVAVDGAVYRQHPRFSEILQQTVKQLAPDCDVTFVPS
EDGSGKGAAMVTAVASRLAAQRQQIDETLAPFRLSPEQLQEVQARMREEMERGLKKETNAKSSVRMLPTY
VRATPDGTERGDFLALDLGGTNFRVLLVQVGSREEGGVQIVSEVYSIPQDVMQGTGEQLFDHIVDCIVDF
QKKQGLMGQTLPLGFTFSFPCKQTGLDQGILLSWTKGFNASDCEGQDVVKLLREAIRRKQDFDLDVVAIV
NDTVGTMMSCGYDDPNCEVGLIVGTGTNACYMEEMKNVGTVEGDEGRMCINMEWGAFGDNGCLDDLVTPF
DRMVDKSSINPGKQRFEKLISGMYLGEIVRHILLDLTKKGILFRGKQSQRLQTRDIFQTKFLSDIESDSL
ALRQVRAILEDLGLDVTCDDAVIVREVCQAVSRRAAQLCGAGIAAVVEKIRENRGLEQLSVTVGVDGTLY
KLHPHFSQHVQETVKELAPRCSVTFLQSEDGSGKGAALITAVACRLAETGQA
>anc224|avgPP:0.986|lnPP:-21.801|num_ambig:21|num_ambig_gaps:0

LQPFLLSIGQLEKVRDLMLDNMERGLKAKPRKQSTLKMLPTYVRSTPDGTERGDFLVLDLGGAKFRILKV
KLSRASGVEMDSEVFDISEDIMTGTGEQLFDHVAECLGRFLQHRNLSEEVFPLGFTFSFPCEQSQLDKSV
LISWTKGFRCSGVEGKDVVQLLKDAIQRRGDYKVDVIAIVNDTVGTMMSCGYKDHSCEVGFIVGTGTNVC
YMEEMGNVEAVEGDEGTMCINMEWGSLGDDGALNDILTEYDLQVDQTSRVPGKQRFDKMISGMYMGEIVR
HVLVKLTNQGLLFGGVSTQQLLNKGTFGTKDVSDIENDNFPFEKTKNVLTALGLKPSEKDCVQVKCVCSA
VSMRSARLCAAGLAAVANRIKNNRRHQLGRITVGVDGSVYKTNPKFGEHLNQALKELAPGCQIKFLVSED
GSGKGTAIVTAVAQRLASQRKHINEILAPFLMSHEKLKEVQSRMRNEMEIGLHKQTQPEATVKMLPTYVR
ATPDGTEEGEFIALDLGGTNFRVLCVNVGLKNEGGVQMKSKTFTLPTEVIQGTGEALFDHIVDCITEFQK
ENGLQGKKLPLGFTFSFPCKQTSLDQGILISWTKGFSASGCVGEDVVRLLREAANRKNNHDFRVVALVND
TVGTMMSCGYDDTACEIGLIVGTGTNACYMEEMRNVEVLEGDEGKMCINMEWGAFGDNGCLDDITTSFDS
NVDKFSINPGKQRYEKMISGMYLGEIVRQVLIELTKDRILFRGKISEQLNTRDLFPTKYLSQIESDSLGL
LQVRSILTKLGLNSTCDDTVIVKEVCTIVSKRAAQLCAAGIAAVVEKMRENRGLDHLKITVGVDGTLYKL
HPHFAGVLQKTVKTLAPQCDVSFHQSEDGSGKGAGLITAVACRVADAEPRAA
>anc224_altAll|avgPP:0.982|lnPP:-32.747|num_ambig:21|num_ambig_gaps:0
LQPFLLSIGQLEKVRDLMLDNMERGLKAKPKRQSTLKMLPTYVRSTPDGTECGDFLVLDLGGAKFRILKV
KLSRASGVEMDSEVFDISEDIMTGTGEQLFDHVAECLGHFLQHRNLSEEVFPLGFTFSFPCEQSQLDKSV
LISWTKGFRCSGVEGKDVVQLLKDAIQRRGDYKVDVVAIVNDTVGTMMSCGYKDHSCEVGFIVGTGTNVC
YMEEMGNVEAVEGDEGTMCINLEWGALGDDGALNDIVTEYDSQVDQTSRVPGKQRFDKMISGMYMGEIVR
HVLVKLTNRGLLFGGVSTLQLLNKGTFGTKDVSDIENDNFPFEKTKSVLTALGLKPSEQDCVQVKCVCSA
VSMRSARLCAAGLAAVANRIKNNRRHHLGRITVGVDGSVYKTNPKFGEHLNQALKELAPGCQIKFLVSED
GSGKGTAIVTAVAQRLATQRKHINEILAPFLMSHEKLKEVQSRMRNEMEIGLHKQTQPEATVKMLPTYVR
ATPDGTEEGEFIALDLGGTNFRVLCVNVGLKNEGGVQMKSKTFTLPTEVIQGTGEGLFDHIVDCITEFQK
ENGLQGKKLPLGFTFSFPCKQTSLDQGILISWTKGFSASGCVGEDVVRLLREAANRKNNHDFRVVALVND
TVGTMMSCGYDDTACEIGLIVGTGTNACYMEEMRNVEVLEGDEGRMCINMEWGAFGDNGCLDDITTSFDN
DVDKFSINPGKQRYEKMISGMYLGEIVRQVLIELTKDRILFRGKISEQLNTRDLFPTKYLSQIESDSLGL
LQVRSILTKLGLNSTCDDTVIVKEVCTVVSKRAAQLCAAGIAAVVEKMRENRGLDHLKITVGVDGTLYKL
HPHFAGVLQKTVKTLAPQCDVSFHLSEDGSGKGAGLITAVACRVADAEPRAA
>anc225|avgPP:0.964|lnPP:-58.559|num_ambig:39|num_ambig_gaps:2
LQPFLLSQAQLEKVRDLMLDNMERGLKAKPKRPSTLKMLPTYVRSTPDGTERGDFLVLDLGGAKFRILKV
KLDGASGRGVEMDSEVYDIPEDIMTGTGKQLFDHVAECLGRFLQQRNLKEEVLPLGFTFSFPCKQSQLDK
SVLISWTKGFRCSGVEGKDVVQLLKDAIQRRGDYKVDVIAIVNDTVGTMMSCGYKDHSCEVGFIVGTGTN
VCYMEEMGNVEAVEGDEGTMCINMEWGALGDDGALNDIQTEYDLQVDQTSRVPGKQRFDKMISGMYMGEI
VRHVLVKLTNQGLLFGGVKTSQLLTKGSFGTKDVSEIENDNFGLEKAKSVLTALGLKPSEQDCVQVKCVC
SAVSMRSAKLCAAGLAAVANRIKNNRQDQLGRITVGVDGSVYKTNPKFGEHLNQALKELAPGCQIKFLVS
EDGSGKGTAIVTAVAQRLAAQRQHINEILAPFIMSHEKLKEVQRRMRNEMEIGLHKETQPEATVKMLPTY
VRATPDGTERGEFLALDLGGTNFRVLYVNVGLKNEGGVQMKSKTFTLPTEVIQGTGEELFDHIVDCITEF
QKENGLKGKRLPLGFTFSFPCKQTSLDQGILISWTKGFSASGCVGEDVVRLLREAAIRKNNPDFHVVALV
NDTVGTMMSCGYDDTACEIGLIVGTGTNACYMEEMRNVEVLEGDEGQMCINMEWGAFGDNGCLDDITTSF
DNNVDKFSINPGKQRYEKMISGMYLGEIVRQVLIVLTKNGILFRGKISEQLNTRNLFPTKFLSDIESDSL
GLLQVRSILTELGLNSTCDDTVVVKEVCTIVSRRAAQLCAAGIAAVVEKIRENRGLDHLKITVGVDGTLY
KLHPHFAGVVQKTVKTLAPQCDVTFHQSEDGSGKGAALITAVACRVAGAEPRAA
>anc225_altAll|avgPP:0.957|lnPP:-76.726|num_ambig:39|num_ambig_gaps:2
LQPFLLSQGQLEQVRDLMLDNMERGLKAKPKRPSTLKMLPTYVRSTPDGTERGDFLVLDLGGAKFRILKV
KLDRASGRGVEMDSEVFDIPEDIMTGTGKQLFDHVAECLGRFLEQRNLKDEVLPLGFTFSFPCKQTQLDK
SVLISWTKGFRCSGVEGKDVVQLLKDAIQRRGDYKVDVIAIVNDTVGTMMSCGYEDHSCEVGFIVGTGTN
VCYMEEMGNVEAVEGDEGTMCINMEWGALGDDGALNDIQTEYDSQVDQTSRVPGKQRFEKMISGMYMGEI
VRHVLVKLTNQGLLFGGVKTSQLLTKGTFGTKDVSDIENDHFGLDKAKNILTSLGLKPSEQDCVQVKCVC
SAVSMRSARLCAAGLAAVANRIKNNRHDHLGRITVGVDGSVYKTNPKFGEHLNQALKELAPGCQIKFLVS
EDGSGKGTAIVTAVAQRLAAQRQHINEILAPFIMSHEKLKEVQKRMRNEMEIGLHKQTQPEATVKMLPTY
VRATPDGTERGEFLALDLGGTNFRVLYVHVGLKNEGGVQMKSKTFTLPTEVIQGTGEELFDHIVDCITEF
QKENGLKGKRLPLGFTFSFPCKQTSLDQGILISWTKGFSASGCVGEDVVRLLREAANRKNNHDFHVVALV
NDTVGTMMSCGYDDAACEIGLIVGTGTNACYMEEMRNVEVLEGDEGQMCINMEWGAFGDNGCLDDITTSF

DSDVDQFSINPGKQRYEKMISGMYLGEIVRQVLIELTKDGILFGGKISEQLNTRDLFPTKFLSDIESDTL
GLLQVRAILTELGLNSTCDDTVIVKEVCTTVSRRAAQLCAAGIAAVVEKIRENRGLDHLKITVGVDGTLY
KLHPHFAGVVQKTVKTLAPQCDVTFHQSEDGSGKGAALITAVACRVADAEARAA
>anc226|avgPP:0.952|lnPP:-78.945|num_ambig:59|num_ambig_gaps:2
LQPFCLSQAQLEQVRDLMLDNMERGLKKKPDRPSTLKMLPTYVRSTPDGTERGDFLVLDLGGAKFRVLQV
KLDGDSGRPVEMESEIYAIPEDIMTGTGRQLFDHVAECLGRFLQQRHLKEKALPLGFTFSFPCKQSQLDK
SVLISWTKGFRCSGVEGKDVVQLLRDAIHRRGDYNVDVIAIVNDTVGTMMSCGYEDHSCEVGFIVGTGTN
ACYMEEMGNVETVEGDEGKMCVNMEWGALGDDGALNDIQTEYDLQVDQNSLAPGKQRFDKMISGMYMGEI
VRLVLVKLASQGLLFGGAKTSALLTKGSFGTKHVSEIEDDHSGLAKAKQILTSLGLKPSEQDCIQVQRVC
SAVSTRSANLCAAGLAAVVTRIQRNRHDNHSRITVGVDGSVYKTHPKFSERLHQALKELAPECQIKFLVS
EDGSGKGTAIVTAVAQRLAAQRQHINEILAPFILSHEKLREVQKRMRVEMEMGLRKETQPKATVKMLPTY
VRATPDGTERGDFLALDLGGTNFRVLYVHVGLKEEGGVQMKSKTFSLPEEVIQGTGEELFDHIVDCITDF
QKENGLRGKRLPLGFTFSFPCKQTSLDQGILISWTKGFSASGCEGEDVVRLLREAAVRKNNSDLHVVALV
NDTVGTMMSCGYDDPSCEIGLIVGTGSNACYMEEMRNVEVLEGDEGQMCINMEWGAFGDNGCLDDITTSF
DHNVDQFSINPGKQRYEKMISGMYLGEIVRQILIVLTKNGILFRGKISERLQTRNIFPTKFLSDIESDSL
GLLQVRSILTDLGLDSTCDDTVVVKEVCTTVSRRAAQLCAAGIAAVVEKIRENRGLDHLDITVGVDGTLY
KLHPHFAGVVQETVKALAPQCDVTFLQSEDGSGKGAALITAVACRIAGAEPRAA
>anc226_altAll|avgPP:0.943|lnPP:-105.838|num_ambig:59|num_ambig_gaps:2
LQPFCLSQAQLEKIRDLMLDNMERGLKKKPNRPSTLKMLPTYVRSTPDGTERGDFLVLDLGGAKFRVLKV
KLDKDSGRSVEMESEVFDIPEDIMTGTGKQLFDHVAECLGRFLEQRNLKEKALPLGFTFSFPCKQTQLDK
SVLISWTKGFRCSGVEGEDVVQLLRDAIQRRGDYNVDVIAIVNDTVGTMMSCGYKDHSCEVGFIVGTGTN
ACYMEEMGNVEAVEGDEGRMCVNMEWGALGDDGALNDIQTDYDHQVDQNSLAPGKQRFEKMISGMYMGEI
VRLVLVKLANQGLLFGGVKTPALLTKGSFATKHVSEIEDDNFGLAKAKQILTSLGLKPSEQDCVQVQRVC
STVSTRSANLCAAGLAAVVTRIQRNRQDNHSRVTVGVDGSVYKTHPKFSERLHQALKQLAPECQIKFIVS
EDGSGKGTAIVTAVAQRLAAQRQQINEVLAPFILSNEKLKEVQRRMRVEMEMGLRKETQPKATVKMLPTY
VRATPDGTERGDFLALDLGGTNFRVLYVNVGLKEEGGVQMKSKTFSLPEEVIQGTGEELFDHIVDCITDF
QKENGLKGKRLPLGFTFSFPCKQTSLDQGILISWTKGFSASGCEGEDVVRLLREAAIRKNNSDLHVVALV
NDTVGTMMSCGYDDPACEIGLIVGTGTNACYMEEMKNVEVLEGDEGQMCINMEWGAFGDNGCLDDITTSF
DHNVDKFSINPGKQRYEKMISGMYLGEIVRQILIELTKNGILFGGKISERLNTRDIFPTKFLSDIESDTL
GLLQVRAILTDLGLDSTCDDAVIVKEVCTIVSRRAAQLCAAGIAAVVEKIRENRGLDHLDVTVGVDGTLY
KLHPHFAGVVQETVKALAPQCEVTFLQSEDGSGKGAALITAVACRVAGSEARAA
>anc227|avgPP:0.928|lnPP:-124.131|num_ambig:88|num_ambig_gaps:2
LQPFCLSEEQLQQIRDLMLENMERGLRKQTNPTSTLKMLPTYVRSTPDGTERGDFLVLDLGGTKFRVLRV
KLTGDSGQSVEMESEIFAIPEDIMTGTGQQLFDHVAECLGNFLDKLQLKDKKLPLGFTFSFPCRQTQLDK
SVLISWTKGFRCSGVEGKDVVQLLRDAIHRRGDYNVDVIAVVNDTVGTMMSCGYEDHSCEIGLIVGTGTN
ACYMEEMGNVETVEGDEGRMCVNMEWGAFGDDGALNDIQTEFDLQVDQNSLNPGKQRFEKMISGMYMGEI
VRLILVKLASQGLLFGGVTTPALLTKGSFETKHVSEIEDDKAGLAKAKEILTSLGLNPSEQDCVQVQRVC
SAVSTRSANLCAAGLAAVVTRIQRNRHLKHLRTTVGVDGTVYKTHPKFSERLQQTLKQLAPDCDVKFLLS
EDGSGKGTAMVTAVAQRLAAQRQQINETLAPFRLSHEQLREVQKRMRVEMERGLKKETQSTATVKMLPTY
VRATPDGTERGDFLALDLGGTNFRVLLVNVGSREEGGVQMVSKIYSIPEEVMQGTGEELFDHIVDCITDF
QKKQGMMGRRLPLGFTFSFPCKQTGLDQGILISWTKGFNASGCEGEDVVRLLREAIKRKQNVDLDVVAIV
NDTVGTMMSCGYDDPNCEIGLIVGTGSNACYMEEMRNVETVEGDEGRMCINMEWGAFGDNGCLDDIVTPF
DRVVDKSSINPGKQRYEKMISGMYLGEIVRHILIDLTKRGILFRGKISERLQTRDIFQTKFLSEIESDSL
ALLQVRAILQDLGLDSTCDDTVIVKEVCTVVSRRAAQLCGAGIAAVVEKIRENRGLDHLKITVGVDGTLY
KLHPHFSKIVQETVKELAPRCDVTFLQSEDGSGKGAALITAVACRLAEAGPH
>anc227_altAll|avgPP:0.916|lnPP:-160.360|num_ambig:88|num_ambig_gaps:2
LQPFCLSDEQLQKIKDLMLENMEKGLKRQTHPTSTVKMLPTYVRSTPDGTEKGDFLVLDLGGAKFRVLQV
KLSDDGGQPVEMESEVYAIPEDIMTGTGQQLFDHVAECLSNFLEQQQLKEKALPLGFTFSFPCKQTQLDK
SVLISWTKGFKCSGVEGKDVVQLLRDAIQRQGDYDIDVVAVVNDTVGTMMSCGYKDHPCEVGLIVGTGTN
ACYMEEMRNVETVEGDEGRMCVNMEWGAFGDDGALNDIQTEFDLQVDQNSLNPGKQRFEKMISGMYMGEI
VRLVLVKLANQGLLFGGATTPALLTKGTFETKHVSEIEDDKVGLAKAKEILTSLGLKPSEQDCIQVQQVC
STVSTRSANLCAAGLAAVVTRIQRNRHLKHLRTTVGVDGTVYKTHPKFSERLHQTVKQLAPECDVQFLLS

EDGSGKGAAMVTAVAHRLAAQRQQINETLAPFRLSHEQLREVQRRMRVEMERGLRKETHPTATVKMLPTY
VRATPDGTERGDFLALDLGGTNFRVLLVNVGSREEGGVQMTNKIYSIPEEVIQGTGEELFDHIVDCIIDF
QKKQGLKGKRLPLGFTFSFPCRQTSLDQGILISWTKGFNASGCEGEDVVRLLREAIKRKQDFELDVVAIV
NDTVGTMMSCGYDDPSCEIGLIVGTGTNACYMEEMKNVETVEGDEGQMCINMEWGAFGDNGCLDDIITPF
DQVVDKSSINPGKQRYEKMISGMYLGEIVRNILIDLTKRGILFRGRISERLNTRDIFETKFLSDIESDSL
ALLQVRAILQELGLDVTCDDAVVVKEVCTVVSRRAAQLCGAGIAAVVEKIRENRGLDHLKVTVGVDGTLY
KLHPHFSRVVQETVKELAPRCEVTFLQSEDGSGKGAALITAVACRIAETGQR
>anc228|avgPP:0.933|lnPP:-114.021|num_ambig:74|num_ambig_gaps:2
LQPFCLSDEQLQKIMDLMLENMEKGLRKQTNPTSTLKMLPTFVRSTPDGTERGDFLALDLGGTKFRVLRV
KLTDDGGQSVEMESEIFAIPEDIMTGTGTQLFDHVAECLGSFLDKQQLKDKKLPLGFTFSFPCRQTKLDK
SVLITWTKGFKCSGVEGKDVVQLLRDAIHRRGDYDVDVIAVVNDTVGTMMSCGYEDHSCEIGLIVGTGTN
ACYMEEMGNIEQVEGDEGRMCVNMEWGAFGDDGALNDIQTEFDLEVDRNSLNPGKQRFEKMISGMYMGEI
VRLILVKLASQGLLFGGVTTPALLTKGSFETKYVSEIEDDKVGLAKAKEILTSLGLNPSEQDCVAVQRIC
SAVSTRSANLCAAGLAAVVTRIQRNRHLKHLRTTVGVDGTVYKTHPKFSERLQQTLKQLAPDCDVQFLLS
EDGSGKGTAMVTAVAHRLAAQRQQINETLAPFRLSHEQLLEVQKRMRLEMERGLKKETQSTATVKMLPTY
VRATPDGTERGDFLALDLGGTNFRVLLVNVGSREEGGVQMVNKIYSIPEEVMQGTGEELFDHIVDCITDF
LKKQGMKGRRLPLGFTFSFPCRQTGLDQGILISWTKGFNATGCEGEDVVKLLREAIKRKQDVELDVVAIV
NDTVGTMMSCAYDDPNCEIGLIVGTGSNACYMEEMRNVETVEGDEGRMCINMEWGAFGDNGCLDDIVTQF
DQIVDKLSINPGKQRYEKMISGMYLGEIVRNILIDLTKRGILFRGQISERLNTRGIFETKFLSEIESDSL
ALLQVRAILQELGLDSTCDDTVIVKEVCTVVSRRAAQLCGAGIAAVVEKIRENRGLDHLKITVGVDGTLY
KLHPHFSKIVHETVKELAPRCDVTFLQSEDGSGKGAALITAVACRLRETGQH
>anc228_altAll|avgPP:0.921|lnPP:-147.058|num_ambig:74|num_ambig_gaps:2
LQPFCLSDEKLQQIMNLMLENMERGLRRQTHPTSTVKMLPTFVRSTPDGTEKGDFLALDLGGTKFRVLQV
KLSDDSGQSVEMESEVYEIPEDIMTGTGTQLFDHVAECLSNFLEKLQLKDKKLPLGFTFSFPCKQTQLDK
SVLISWTKGFKCSGVEGKDVVQLLRDAIQRQGDYDIDVVAVVNDTVGTMMSCGYKDHSCEIGLVVGTGTN
ACYMEEMRNIEQVEGDEGRMCVNMEWGAFGDDGSLNDIQTEFDLEVDQNSLNPGKQRFEKMISGMYMGEI
VRLILVKLASQGLLFGGVTTPALLTKGTFETKYVSEIEDDKAGLTKAKEILTSLGLNPSEQDCIAVQQVC
STVSTRSANLCAAGLAAVVTRIQRNRHLKHLQTTVGVDGTVYKKHPKFSGRLHQTVKQLAPDCDVKFMLS
EDGSGKGAAMVTAVAHRLAAQRQQINETLAPFRLSREQLLEVQRRMRLEMERGLRKETHSTATVKMLPTY
VRSTPDGTERGDFLALDLGGTNFRVLLVKVGSREEGGVQMVNKIYSIPEEVMQGTGEELFDHIVDCIIDF
LKKQGMMGRRLPLGFTFSFPCKQTSLDQGILINWTKGFNASGCEGEDVVKLLREAIKRKQGFDLDVVAIV
NDTVGTMMSCAYDDPNCEIGLIVGTGSNACYMEEMRNVETVEGDEGQMCINMEWGAFGDNGCLDDIVTQF
DRIVDKLSINPGKQRYEKMISGMYLGEIVRNILIDLTKRGILFRGKISERLQTRGIFETKFLSEIESDSL
ALLQVRAILQELGLEVTCDDAIIVKEVCTIVSRRAAQLCGAGIAAVVEKIRENRGLDHLKVTVGVDGTLY
KLHPHFSRIVHETVKELAPKCEVTFLQSEDGSGKGAALITAVACRLREAGQH
>anc229|avgPP:0.955|lnPP:-72.144|num_ambig:63|num_ambig_gaps:0
LYHMRLSDETLQDIMNRMRTEMEKGLGRDTNPTATVKMLPTFVRSTPDGTEKGDFLALDLGGSNFRVLRV
KVSDDGKQKVEMESEIYAIPEDIMRGSGTQLFDHVAECLGNFMEKLQIKDKKLPLGFTFSFPCRQTKLDE
SVLITWTKGFKASGVEGRDVVKLLRKAIQRRGDFDIDIVAVVNDTVGTMMTCGYDDHRCEIGLIVGTGTN
ACYMEEMRNIDLVEGDEGRMCINMEWGAFGDDGSLDDIRTEFDLEIDRGSLNPGKQLFEKMISGMYMGEL
VRLILVKMAKQGLLFGGKITPELLTKGHFETKYVSAIEKDKEGLSKAKEILTKLGLEPSEEDCVAVQRIC
TIVSTRSANLCAATLAAVVTRIKENKGVKRLRTTVGVDGTVYKTHPQFARRLHKTVRRLAPDCDVRFLLS
EDGSGKGAAMVTAVAYRLASQRKQIDETLAPFKLSHEQLLEVKKRMRTEMERGLKKETHSSATVKMLPTY
VRSTPDGTEKGDFLALDLGGTNFRVLLVKIRSGKRRGVEMHNKIYSIPQEVMQGTGEELFDHIVHCISDF
LDYMGMKGARLPLGFTFSFPCRQTSLDQGILLNWTKGFKATGCEGEDVVNLLREAIKRREEFDLDVVAVV
NDTVGTMMTCAYEDPNCEIGLIVGTGSNACYMEEMRNVEMVEGDEGRMCINMEWGAFGDNGCLDDIRTEF
DRAVDELSLNPGKQRYEKMISGMYLGEIVRNILIDFTKRGLLFRGRISERLKTRGIFETKFLSQIESDRL
ALLQVRAILQQLGLESTCDDSIIVKEVCGVVSRRAAQLCGAGMAAVVDKIRENRGLDHLKITVGVDGTLY
KLHPHFSKIMHETVKELAPKCDVTFLQSEDGSGKGAALITAVACRLREAGQH
>anc229_altAll|avgPP:0.944|lnPP:-101.475|num_ambig:63|num_ambig_gaps:0
LYHMRLSDETLQDIMNRMRTEMEKGLGKETNPTATVKMLPTFVRSTPDGSEKGDFLALDLGGTNFRVLRV
KVSDDGKQTVEMENQIYEIPEDIMHGTGTQLFDHVAECLGSFMEKQQIKDKKLPLGFTFSFPCRQTKLDE

SILITWTKGFKASGVEGQDVVKLLREAIKKRGDYDVDIVAVVNDTVGTMMTCGYDDHRCEIGLIVGTGTN
ACYMEEMRHIEQVEGDEGRMCINMEWGAFGDDGALDDIRTEFDREVDRGSLNPGKQLFEKMISGMYMGEL
VRLILVKMATEGLLFGGKTTPELLTKGHFETKYVSAIEKDKEGLSKAKEILTKLGLEPSEEDCIAVQQIC
TIVSTRSANLCAAALAAVVTRLRENKGVQRLRTTVGVDGTVYKKHPQYSKRLQQTVRRLVPDCDVQFLLS
EDGSGKGAAMVTAVAYRLAAQRKQIDETLAPFKLSHEQLLEVKRRMRTEMERGLKKETHSTATVKMLPTY
VRSTPDGTEKGDFLALDLGGTNFRVLLVKVRSGKKRSVEMHNKIYAIPQEVMQGTGEELFDHIVDCISDF
LEYMGMKGARLPLGFTFSFPCKQTSLDQGILVNWTKGFKASGCEGEDVVNLLREAIKRREEFELDVVAIV
NDTVGTMMTCAYEDPNCEIGLIVGTGSNACYMEEMRNVEMVEGDEGRMCINMEWGAFGDNGCLDDIRTEY
DRAVDELSLNPGKQRYEKMISGMYLGEIVRNILIDFTKRGLLFRGQISERLKTRGIFETKFLSQIESDRL
ALLQVRAILQELGLDSTCDDSIIVKEVCGVVSRRAAQLCGAGMAAVVDKIRENRGLDHLKVTVGVDGTLY
KLHPHFSRIMHETVKELAPKCEVTFLQSEDGSGKGAALITAVACRLREAGQH
